# MICROGLIAL CELL EXPRESSION OF THE TYPE 2 CANNABINOID RECEPTOR REGULATES IMMUNE-MEDIATED NEUROINFLAMMATION

**DOI:** 10.1101/2023.08.10.552854

**Authors:** Alison Moe, Aditya Rayasam, Garrett Sauber, Ravi K. Shah, Cheng-Yin Yuan, Aniko Szabo, Bob M. Moore, Marco Colonna, Weiguo Cui, Julian Romero, Anthony E. Zamora, Cecilia J. Hillard, William R. Drobyski

**Author notes:** A.M. and A.R. contributed equally to this study. Address correspondence to: William R. Drobyski, M.D. Hematopoietic Stem Cell Transplant and Cellular Therapy Program 9200 West Wisconsin Avenue Milwaukee, WI 53226.

## Abstract

Neuroinflammation is a recognized complication of immunotherapeutic approaches such as immune checkpoint inhibitor treatment, chimeric antigen receptor therapy, and graft versus host disease (GVHD) occurring after allogeneic hematopoietic stem cell transplantation. While T cells and inflammatory cytokines play a role in this process, the precise interplay between the adaptive and innate arms of the immune system that propagates inflammation in the central nervous system remains incompletely understood. Using a murine model of GVHD, we demonstrate that type 2 cannabinoid receptor (CB2R) signaling plays a critical role in the pathophysiology of neuroinflammation. In these studies, we identify that CB2R expression on microglial cells induces an activated inflammatory phenotype which potentiates the accumulation of donor-derived proinflammatory T cells, regulates chemokine gene regulatory networks, and promotes neuronal cell death. Pharmacological targeting of this receptor with a brain penetrant CB2R inverse agonist/antagonist selectively reduces neuroinflammation without deleteriously affecting systemic GVHD severity. Thus, these findings delineate a therapeutically targetable neuroinflammatory pathway and has implications for the attenuation of neurotoxicity after GVHD and potentially other T cell-based immunotherapeutic approaches.

## INTRODUCTION

Neurological inflammation and cognitive dysfunction are increasingly recognized complications of cancer immunotherapeutic approaches such as immune checkpoint inhibitor treatment (1,2), chimeric antigen receptor therapy (3,4), and graft versus host disease (GVHD) occurring after allogeneic hematopoietic stem cell transplantation (5-9). Adverse neurological events that have been described include encephalitis, vasculitis, demyelinating disorders, aphasia, and seizures, all of which can lead to severe cognitive alterations and, in some instances, death (6-8,10). The pathophysiology of this complication is thought to be due to the activation of T cells and subsequent release of inflammatory cytokines which promote pathological damage within the central nervous system (CNS) (9,11,12). Notably, these T cell populations do not reside in the brain but are able to enter from the periphery under inflammatory conditions when there is breakdown of the blood brain barrier (13). Resident immune cells in the brain, specifically microglial cells, have also been implicated in this pathological process (11,14), although the mechanistic pathways by which these cells contribute to neuroinflammation have not been completely delineated. Moreover, the mechanisms by which microglia, the primary innate immune cells of the brain, interact with non-resident T cells to promote this inflammatory milieu are not well understood.

Endocannabinoids are endogenously produced, bioactive lipids that are produced on demand and are present in all brain regions (15,16). The two primary endocannabinoids are anandamide (*N*-arachidonoylethanolamine) (AEA) (17) and 2-arachidonoyl glycerol (2-AG) (18). Endocannabinoid effects are mediated by two G protein coupled receptors; the type 1 cannabinoid receptor (CB1R) (19) that is highly expressed in the CNS (20), and the type 2 cannabinoid receptor (CB2R) (21) which is predominantly expressed on immune cells (22). The CB1R plays a role in the regulation of synaptic activity at glutamatergic and GABAergic receptors through presynaptic inhibition of neurotransmitter release (23), whereas the CB2R has been shown to exert immunoregulatory effects on T cells, B cells, macrophages, and microglial cells (24-27). Thus, endocannabinoids have both neurological and immunological effects and, importantly, the CB2R is expressed on cells that constitute the innate and adaptive arms of the immune system, both of which have pathophysiological roles in immunotherapy-mediated neuroinflammation.

In the current report, we employed a murine model to examine the pathophysiology of GVHD-induced neuroinflammation. These studies demonstrated that endocannabinoid signaling through the CB2R plays a critical role in the regulation of inflammation within the CNS during this disease. Mechanistically, host expression of the CB2R potentiated inflammation in the brain by promoting the accumulation of proinflammatory donor T cells, augmenting microglial cell numbers, and increasing neuronal cell death. Specifically, GVHD augmented CB2R expression on microglial cells resulting in an activated proinflammatory phenotype that was independent of interleukin 6 (IL-6) signaling blockade. Conversely, conditional deletion of the CB2R from microglia significantly reduced the accumulation of inflammatory T cells, decreased expression of genes associated with TNF signaling and chemotaxis and attenuated neuronal cell death. In addition, pharmacological inhibition of this pathway with a brain penetrant CB2R inverse agonist/antagonist mitigated neuroinflammation without exacerbating systemic GVHD, indicating that this pathway is amenable to focused pharmacological intervention. Thus, these results define microglial cell expression of the CB2R as a critical regulator of immune-mediated neuroinflammation.

## RESULTS

### Donor T cells accumulate in all regions of the brain during GVHD and induce neuronal cell death

The clinical manifestations of neuroinflammation which develop during GVHD are varied and include a broad spectrum of cognitive and functional neurological deficits (6-8), suggesting that multiple areas of the brain can be affected. Using beta actin-driven GFP-labeled T cells, both CD4^+^ and CD8^+^ donor T cells were present in significantly higher frequency and absolute numbers in GVHD animals in the prefrontal cortex (PFC), amygdala, brainstem, and cerebellum (**Figure 1A**) which represent regions of the brain that are critical for the processing of higher cortical function, emotional inputs, regulation of key involuntary actions, and balance, respectively (28-31). Gene expression of proinflammatory cytokines in whole brain tissue was significantly increased in GVHD animals (**Supplemental Figure 1A**). In addition, we observed that the frequency and absolute number of donor CD4^+^ and CD8^+^ T cells were augmented (**Supplemental Figure 1B**) and these cells produced multiple inflammatory cytokines (i.e., IFN-γ, TNF-α, IL-6 and GM-CSF) (**Supplemental Figures 1C and 1D**), indicative of generalized inflammation. Immunofluorescence staining of whole mounted brains confirmed the wide dissemination of T cells throughout white and gray matter in GVHD animals (**Figures 1B and 1C**). To account for the existing size disparity of these examined regions, we performed a density-based analysis which revealed similar numbers of T cells in equivalently demarcated areas of each region, except for the amygdala which had relatively reduced T cell accumulation (**Figure 1D**). We also observed an increased density of IBA-1^+^ macrophage/microglial cells in the PFC, amygdala, brain stem, and cerebellum in GVHD mice when compared to BM controls (**Figure 1E**). Most IBA-1^+^ cells in BM control animals displayed a ramified morphology, indicative of a resting phenotype (32), whereas the morphology in GVHD mice was hyper ramified and ameboid, indicative of a more activated phagocytic phenotype (33), particularly within the PFC (**Figure 1F**). Immunofluorescence staining demonstrated that there were adjacent GFP^+^ T cells and IBA-1^+^ cells in the brains of these mice (**Figure 1G**). To determine if this proinflammatory environment resulted in cellular damage, we performed western blot analysis which revealed increased cleaved spectrin and cleaved caspase 3 expression in the whole brains of GVHD compared to control animals (**Figure 1H**). The more significant increase in the 120kDa compared to the 150 kDa cleaved spectrin fragment in GVHD mice was indicative of a more prominent role for apoptotic as opposed to necrotic cell death (34). Immunofluorescence with NeuN also revealed that caspase positive cells were essentially all neuronal cells and depicted cleaved caspase 3 positive neurons in anatomic proximity to donor-derived T cells (**Figure 1I**). Collectively, these studies indicated that GVHD induced an inflammatory environment in the brain that was comprised of T cells, microglia/macrophages, and proinflammatory cytokines which resulted in neuronal cell death.

**Figure 1:**
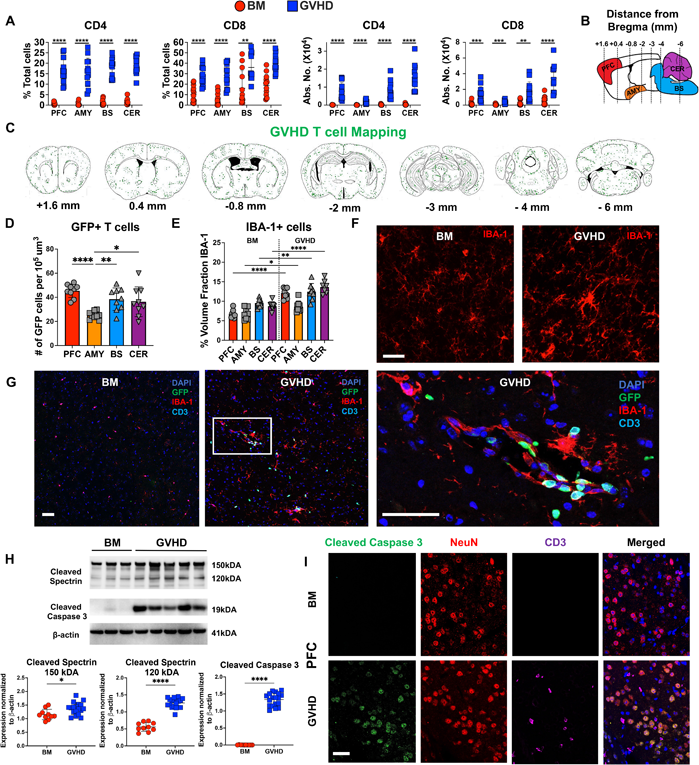
Donor T cells disseminate widely throughout the brain and induce neuronal cell death during GVHD. (A). Lethally irradiated (900 cGy) Balb/c mice were transplanted with B6^EGFP^ BM alone (5 x 10^6^) or B6^EGFP^ BM and B6^EGFP^ spleen cells (adjusted to yield αβ T cell dose of 0.75 x 10^6^). The percentage and absolute number of donor derived CD4^+^ and CD8^+^ T cells in the prefrontal cortex (PFC), amygdala (AMY), brainstem (BS), and cerebellum (CER) 14 days post transplantation is shown. Results are from three experiments. (B-G). Lethally irradiated Balb/c mice were transplanted with B6 Rag-1 BM (5 x 10^6^) alone or together with purified B6^EGFP^ splenic CD4^+^ (0.9 x 10^6^) and CD8^+^ (0.55 x 10^6^) T cells. (B). Sagittal graphical representation of the brain depicting the anatomic location of specified regions along with the distance (in millimeters) from the bregma that selected coronal slices were examined to assess the extent of donor T cell infiltration. (C). Distribution of donor GFP-labeled T cells in the specified coronal sections of animals transplanted with Rag-1 BM and purified GFP^+^ T cells 14 days post transplantation. (D). The number of GFP^+^ T cells in specified brain regions per 10^5^ um^3^. (E, F). Number of IBA-1^+^ cells in the specified brain regions of mice reconstituted with Rag-1 BM alone or together with purified GFP^+^ T cells depicted as the percent of the area fraction are shown in panel E and representative immunofluorescence images are depicted in panel F. White scale bar is 30 microns. Results from panels D and E are from two experiments. (G). Representative immunofluorescence images depicting co-localization of GFP^+^ T cells and IBA-1^+^ cells in the PFC. Scale bar is 30 microns. (H). Representative western blot images and scatterplots depicting normalized expression of cleaved spectrin (150 and 120 kDA) and cleaved caspase 3 in the PFC from Balb/c mice transplanted with B6 BM alone or together with B6 spleen cells (adjusted to yield an αβ T cell dose of 0.8 x 10^6^). Results are from two experiments. (I). Representative immunofluorescence images showing expression of cleaved caspase 3, NeuN (neurons), and CD3 (T cells) along with merged compilation. Statistics: *p<0.05, **p<0.01, ***p<0.001, ****p<0.0001.

### Microglial cells acquire an inflammatory transcriptional signature during GVHD

To further delineate the inflammatory environment in the brain, we performed single cell RNA sequencing (scRNAseq) analysis on immune cells isolated from the brains of BM and GVHD mice. This analysis revealed 9 transcriptionally distinct clusters that represented CD8^+^ T cells (2 clusters), CD4^+^ T cells (1 cluster), macrophages (*Lyz2*) (1 cluster), and microglia (*P2ry12/Tmem119*) (5 clusters) which constituted the largest set of clusters (**Figures 2A and 2B**). The majority of analyzed cells consisted of microglia and T cells, with only a small percentage (∼10%) consisting of macrophages (**Supplemental Figure 2A**). T cells represented 44% of all cells in GVHD mice, but only 5% in BM control animals. Transcriptional analysis of inflammatory cytokines in the brain revealed that CD4^+^ and CD8^+^ T cells primarily produced IFN-γ, TNF-α and GM-CSF (clusters 2, 4 and 5), macrophages produced IL-1β and IL-27 (cluster 3), and microglia produced IL-1α, TNF-α, and to a lesser extent IL-6 (clusters 0,1,6,7 and 8) (**Figure 2C**). There was no detectable expression of IL-10, IL-12, IL-17A, IL-22 or IL-23 in any of these cell types (data not shown). The inflammatory cytokine transcripts identified in T cells were consistent with cytokine protein expression that was observed by flow cytometry (**Supplemental Figures 1C and 1D**). T cells from GVHD mice had increased expression of *Icos*, *Stat1*, *Il12rb2*, *Ly6c*2, and *Ly6a*, indicative of an activated memory phenotype (**Supplemental Figure 2B and Supplemental Table 1**), whereas the one macrophage cluster from GVHD mice demonstrated increased expression of complement (*C1qa, C1qb*), chemokine (*Ccl5, Cxcl9, Cxcl10*), and S100a genes (*S100a8, S100a11*) which all are associated with inflammatory pathways (**Supplemental Figure 2C and Supplemental Table 2)**.

**Figure 2:**
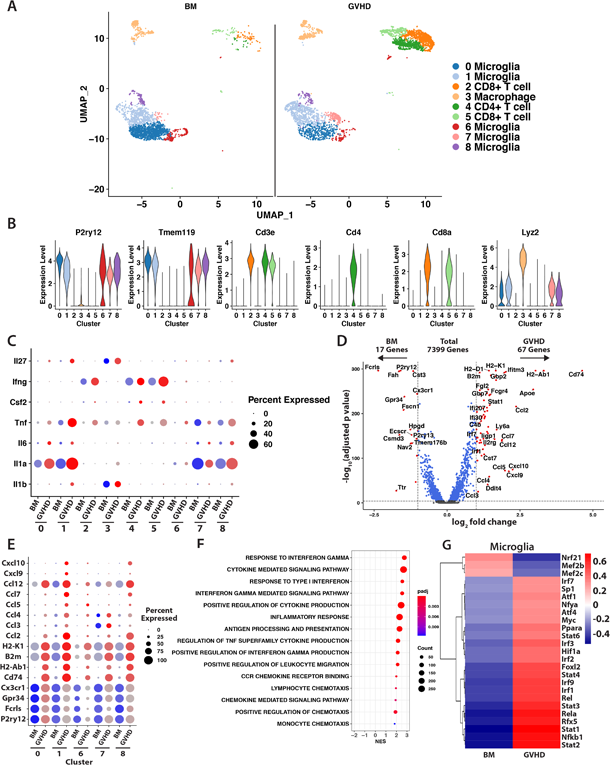
Microglial cells acquire an inflammatory transcriptional signature during GVHD. Lethally irradiated (1100 cGy) B6 mice were transplanted with B10.BR BM (5 x 10^6^) alone (BM) or together with B10.BR spleen cells (adjusted to yield an αβ T cell dose of 5 x 10^6^) (GVHD). Single live cells from pooled brains (n=5/group) were sorted 14 days post transplantation. (A). Uniform manifold approximation and projection (UMAP) dimensional reduction of scRNAseq data of flow-sorted live cells from pooled brains. Unsupervised clustering using Seurat revealed 9 transcriptionally distinct clusters using a resolution of 0.5. (B). Violin plots showing log normalized expression of indicated microglia (*P2ry12, Tmem119*), T cell (*CD3, CD4, CD8*) and macrophage markers (*Lyz2*). (C). Bubble plots depicting inflammatory cytokine profile in each cluster. (D). Volcano plot showing over/underrepresented genes in aggregated microglial clusters from BM versus GVHD mice. Cutoff parameters were |log2(fold change)| > 1.0 and p_adjusted_ <0.0001. (E). Bubble plots depicting microglia-specific, MHC class I and II, and chemokine gene expression in each microglial cell cluster. (F). Bubble plot demonstrating normalized enrichment score (NES) for pathways identified using the Gene Ontology database. (G). Heatmap showing binary regulon activity of the top 25 regulons that were differentially expressed in microglial cell clusters from BM versus GVHD animals.

Since microglia constituted the majority of total immune and identified clusters, we performed a more detailed transcriptome analysis focused on the five microglial cell subsets. This revealed that there were 84 differentially expressed genes with 17 over expressed in BM and 67 over expressed in GVHD mice based on defined cutoff criteria (log 2-fold change) >1.0 and p_adjusted_<0.0001, full list available in **Supplemental Table 3**), representing approximately 1.1% of the total sequenced transcriptome (**Figure 2D**). Microglial cells from BM control animals had increased expression of genes that are characteristic of a homeostatic phenotype (i.e., *Cx3cr1*, *Gpr34*, *Fcrls*, and *P2ry12*) (35) (**Figure 2E**). In contrast, microglia from GVHD mice exhibited increased expression of major histocompatibility complex class I and II MHC genes (i.e., H2-*K1, B2m, CD74*, and *H2-Ab1*), as well as chemokine genes (i.e., *Ccl2, Ccl3, Ccl4, Ccl5, Ccl7, Ccl12, Cxcl9, and Cxcl10*), indicative of an activated inflammatory phenotype (**Figure 2E**). This phenotype was most prominent in clusters 0, 1 and 8 which constituted the majority of microglial cells in the brain (**Supplemental Figure 2A**). Gene set enrichment analysis (GSEA) using the Gene Ontology (GO) dataset confirmed that microglia from GVHD mice had increased expression of genes associated with inflammatory pathways (e.g., cytokine mediated signaling pathway, response to interferon gamma, antigen processing and presentation, and TNF superfamily cytokine production), and chemotaxis (i.e., chemokine receptor binding, positive regulation of chemokine production, leukocyte chemotaxis) (**Figure 2F**). The activation and differentiation of microglia is driven by the expression of transcription factors and downstream target genes which constitute a gene regulatory network (i.e., regulons) and can be interrogated using DoRothEA (Discriminant Regulon Expression Analysis) which is a computational method for gene regulatory network construction of scRNAseq data (36). Using this approach, we observed that microglia from GVHD animals had enriched regulon activity for Stat (i.e., *Stat1*, *Stat2*, *Stat3*, *Stat4*, and *Stat6*), NF-κB (i.e., *Nfkb1*, *Rela*, and *Rel*), and interferon regulatory factor (i.e., *Irf1*, *Irf2*, *Irf3*, *Irf7*, and *Irf9*) family transcription factors (**Figure 2G**), which are all constituents of regulatory networks that mediate inflammation (37). Thus, GVHD induced a transcriptionally coordinated, inflammatory microglial phenotype that was characterized by prominent expression of chemokine genes.

### Microglial cells regulate neuronal cell death during GVHD

The acquisition of an inflammatory microglial transcriptional profile led us to examine the functional role of these cells in promoting neuroinflammation within the CNS. To address this question, we employed IL-34^−/−^ mice as recipients since the maintenance of microglial cell homeostasis is dependent upon interactions between the endogenous ligands CSF-1 and IL-34 with their cognate receptor, CSF-1R (38,39), and the genetic absence of IL-34 results in a significant reduction in microglial cell numbers (40). Consistent with these prior reports, the absence of IL-34 expression in recipient mice resulted in a significant reduction in the number of microglia when compared to wild type (WT) B6 GVHD animals (**Figure 3A**). This was accompanied by a reduction in the absolute number of microglia that expressed MHC class II (**Figure 3B**) and the costimulatory molecules CD80 and CD86 (**Figure 3C**). Immunofluorescence confirmed that there was a significant reduction in IBA-1^+^ microglia in the brains of IL-34^−/−^ versus WT GVHD animals (**Figures 3D and 3E**). Correspondingly, we observed that there was no decrease in the frequency of donor-derived CD4^+^ or CD8^+^ T cells, but there was a significant reduction in the absolute number of these T cell populations (**Figure 3F**) as well as the total number of CD4^+^ and CD8^+^ T cells that produced inflammatory cytokines (i.e., IFN-γ, TNF-α, IL-6 and GM-CSF) (**Figures 3G-3J**). Notably, there was also a decrease in neuronal cell death in IL-34^−/−^ GVHD recipients as evidenced by reduced cleaved caspase 3 protein expression (**Figure 3K**). Collectively, these results provide evidence that microglial cells promote the accumulation of proinflammatory T cells into the CNS and the induction of neuronal cell death during GVHD.

**Figure 3:**
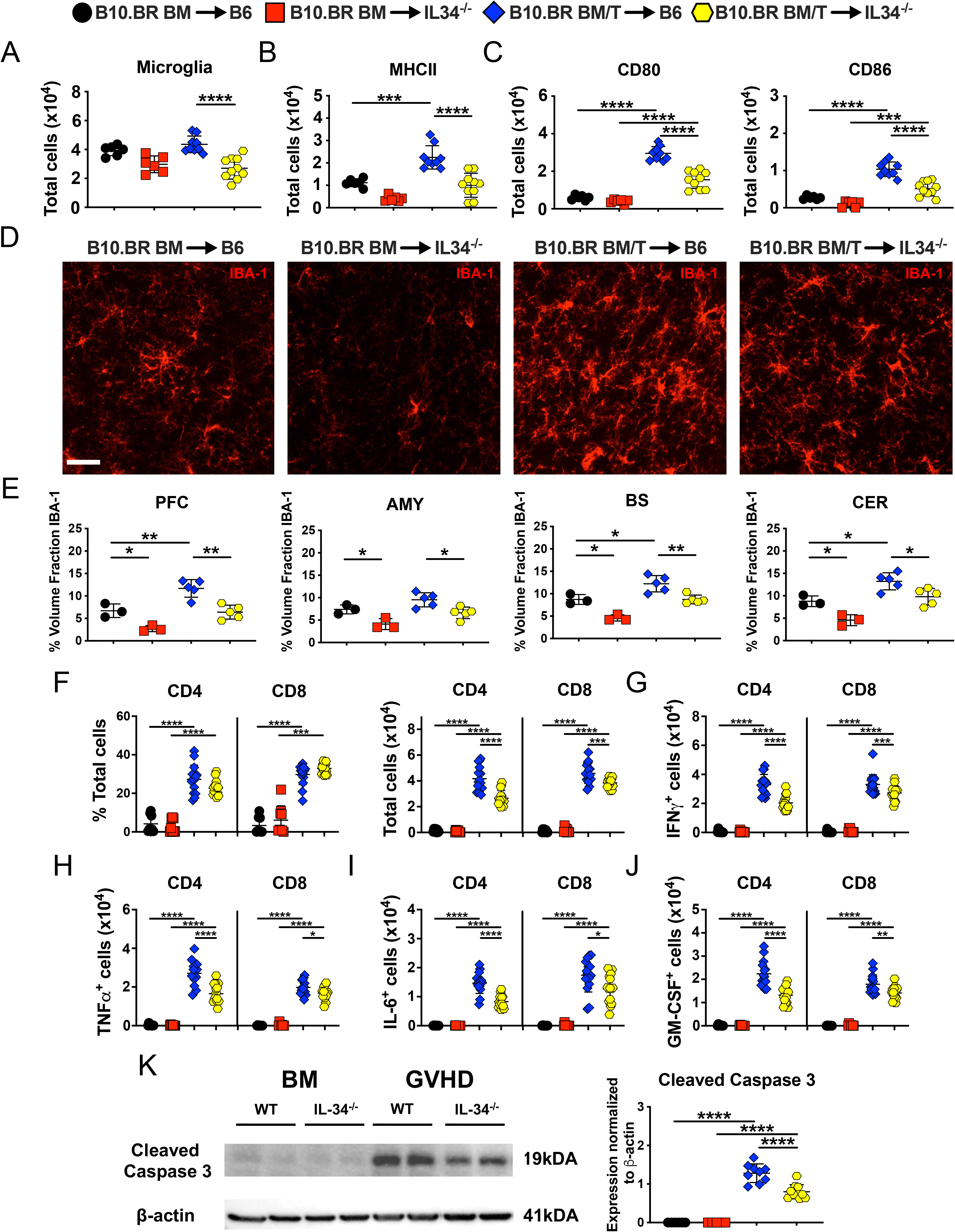
Microglial cells regulate neuronal cell death. (A-J). Lethally irradiated (1000 cGy) B6 or IL-34^−/−^ mice were transplanted with B10.BR BM alone or together with B10.BR spleen cells (adjusted to yield an αβ T cells dose of 4 x 10^6^ T cells). (A). Absolute number of CD45^+^ CD11b^+^ microglial cells. (B, C). Absolute number of MHC class II, CD80 and CD86 expressing microglial cells. Analysis of microglial cells was performed by flow cytometry. Results are from two experiments. (D). Representative immunofluorescence images of IBA-1^+^ cells in the PFC. The scale bar is 30 microns. (E). Quantification of IBA-1^+^ cells in the PFC, amygdala, brainstem, and cerebellum. (F-J). The frequency and absolute number of donor-derived CD4^+^ and CD8^+^ T cells and the absolute number of CD4^+^ and CD8^+^ T cells that produced IFN-γ, TNF-α, IL-6 or GM-CSF in the brains of mice 14 days post transplantation. Analysis of T cells was performed by flow cytometry. Data are from three experiments. (K). Representative western blot images and scatter plots depicting normalized expression of cleaved caspase 3 in the PFC from B6 or IL-34^−/−^ mice transplanted with B10.BR BM alone or together with B10.BR spleen cells. Results are from two experiments. Statistics: *p<0.05, **p<0.01, ***p<0.001, ****p<0.0001.

### Host expression of the CB2R drives neuroinflammation during GVHD

Given the inflammatory milieu and ensuing neuronal cell death driven by T cells and microglia, we sought to uncover mechanistic pathways that coordinately regulated these cell populations in the development of GVHD-induced neuroinflammation. We hypothesized that signaling through the CB2R expressed on immune cell populations might play an important pathophysiological role, since the CB2R has been shown to regulate both adaptive and innate immune responses during systemic GVHD (26), and our data indicate that both arms of the immune system are involved in neuroinflammation. To test this hypothesis, we first examined the role of CB2R expression on donor immune cells in CNS inflammation by transplanting recipients with marrow grafts from either WT or CB2R^−/−^ donors. We observed that there was no difference in expression of inflammatory cytokines in whole brains of animals reconstituted with grafts from WT versus CB2R^−/−^ mice **(Supplemental Figure 3A)**. Given the requirement for donor T cells in the induction of GVHD-induced neuroinflammation, we examined the role of CB2R expression on donor T cells and noted that absence of CB2R expression resulted in a significant decrease in the percentage and absolute number of donor-derived CD4^+^, but not CD8^+^, T cells in GVHD mice (**Supplemental Figure 3B**). There were also reduced numbers of CD4^+^ T cells that produced IFN-γ, TNF-α and GM-CSF, whereas the total number of CD8^+^ T cells that expressed these cytokines was not different between WT and CB2R^−/−^ groups (**Supplemental Figures 3C-3F**). No difference was observed in the absolute number of microglia (**Supplemental Figure 3G**) or microglia expressing class II, CD80 or CD86 (**Supplemental Figures 3H and 3I**). Notably, cleaved caspase 3 levels were not significantly different between animals in these two groups (**Supplemental Figure 3J**), indicating that absence of the CB2R on donor T cells reduced the number of proinflammatory CD4^+^ T cells, but had no effect on the accumulation of inflammatory CD8^+^ T cells and did not prevent neuronal cell death.

In contrast, when recipient mice lacked CB2R expression, we observed a significant reduction in gene expression of IFN-γ, IL-6, TNF-α, and GM-CSF in the brain **(Figures 4A)**. There was also a decrease in the absolute number of donor derived CD4^+^ and CD8^+^ T cells (**Figure 4B**) as well as CD4^+^ and CD8^+^ T cells that produced IFN-γ, IL-6 (CD8 only), TNF-α and GM-CSF in the brains of CB2R^−/−^ versus WT recipients (**Figures 4C-4F**). This was accompanied by a significant reduction in the total number of microglial cells (**Figure 4G**) as well as those expressing MHC class II, CD86 and CD80 (**Figures 4H and 4I**). Correspondingly, we noted decreased expression of cleaved caspase-3 in the brains of CB2R^−/−^ recipient mice (**Figures 4J**), indicating that host CB2R expression potentiated neuronal cell death. Of note, there was no increase in overall GVHD lethality in CB2R^−/−^ recipients, demonstrating that absence of CB2R in the host did not exacerbate survival (**Figure 4K**), unlike what has been reported after transplantation with donor CB2R^−/−^ immune cell populations (26). Thus, these studies revealed that absence of recipient CB2R expression resulted in decreased accumulation of proinflammatory donor T cells, reduced numbers of microglia with an activated phenotype, and significantly diminished neuronal cell death in the brain.

**Figure 4:**
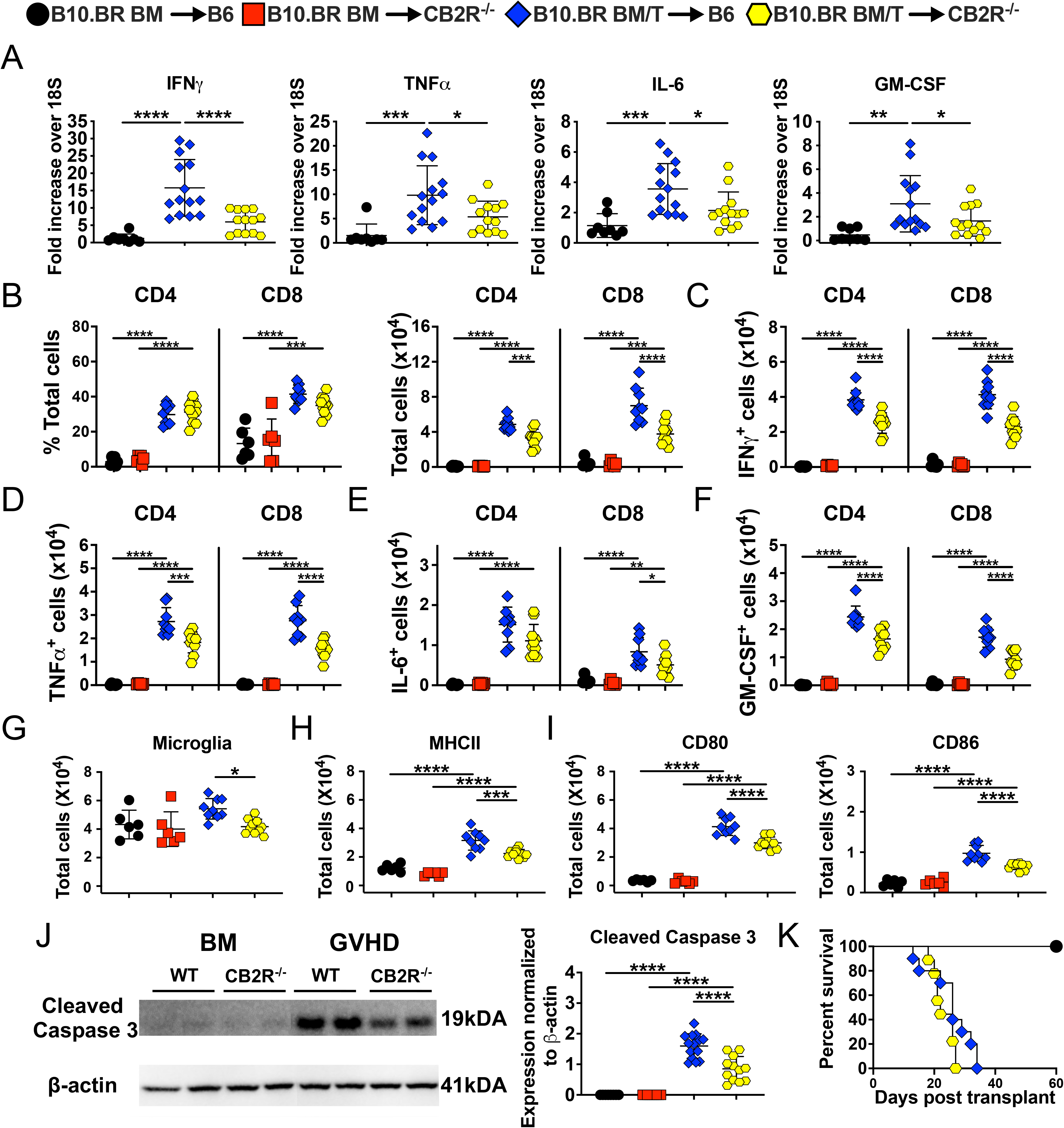
Host CB2R expression regulates neuroinflammation in the brain. Lethally irradiated (1100 cGy) B6 or CB2R^−/−^ mice were transplanted with B10.BR BM (5 x 10^6^) together with B10.BR spleen cells (adjusted to yield a αβ T cell dose of 4.5-5 x 10^6^ T cells). B6 animals transplanted with B10.BR BM served as controls. (A). IFN-γ, TNF-α, IL-6, and GM-CSF mRNA expression is depicted in whole brain. Results are from three experiments. (B-J). Lethally irradiated B6 or CB2R^−/−^ mice were transplanted with B10.BR BM (5 x 10^6^) alone or together with B10.BR spleen cells (adjusted to yield a T cell dose of 4.5-5 x 10^6^ T cells). (B). The percentage and absolute number of donor derived CD4^+^ and CD8^+^ T cells in the brains of mice 14 days post transplantation. (C-F). The absolute number of CD4^+^ and CD8^+^ T cells that produced IFN-γ, TNF-α, IL-6 or GM-CSF. (G). Absolute number of microglial cells. (H, I). Absolute number of MHC class II, CD80 and CD86 expressing microglial cells. Data are from two experiments. (J). Representative western blot images and scatter plots depicting normalized expression of cleaved caspase 3 in the brain from B6 or CB2R^−/−^ mice transplanted with B10.BR BM and spleen cells. (K). Lethally irradiated B6 (n=10) or CB2R^−/−^ (n=9) animals were transplanted with B10.BR BM and spleen cells. B6 mice transplanted with B10.BR BM alone (n=6) served as controls. Survival is shown. Results are from three experiments. Statistics: *p<0.05, **p<0.01, ***p<0.001, ****p<0.0001.

### A brain penetrant, but not peripherally restricted, CB2R inverse agonist/antagonist attenuates inflammation in the brain

Since GVHD is a systemic disease, inflammation is simultaneously induced in the periphery and in the CNS where host CB2R expressing cells both reside. Therefore, to define the location of the recipient CB2R^+^ cell population that was most critical for driving neuroinflammation, as well as determine if the CB2R signaling pathway could be therapeutically targeted, we pursued a pharmacological strategy in which mice were treated with either a peripherally restricted (SR144528) or brain penetrant (SMM-189) CB2R inverse agonist/antagonist. SRR144528 has been reported to not be distributed into the CNS (41) and to be an optimal tool for *in vivo* murine studies due to its high selectivity profile for the receptor (42). To corroborate that SR144528 did not enter the CNS, we treated mice with SR144528 and performed isotope-dilution mass spectrometric analysis to quantitate and compare the concentration of SR144528 in the blood versus the brain of transplant recipients (**Supplemental Figure 4)**. We observed that SR144528 **(Supplemental Figure 5A)** was measurable in the serum with mean concentrations of 150-250 pg/µl with no differences between naïve, BM controls, or GVHD recipients (**Supplemental Figure 5B**). In contrast, while SR144528 was detectable in the brain (mean 5-10 pg/mg), concentrations were significantly lower (15-50-fold) when compared to serum (**Supplemental Figure 5C**), indicating that very little SR144528 distributes to the CNS. To assess the functional effects of CB2R signaling blockade with this agent, we treated mice daily for 14 days and observed that there was a reduced frequency of CD4^+^ T cells, but no difference in the absolute number of CD4^+^ or CD8^+^ T cells in the brains of animals treated with SR144528 versus a vehicle control (**Supplemental Figure 5D**). Moreover, we noted no significant difference in the total number of CD4^+^ and CD8^+^ T cells that produced IFN-γ, TNF-α, or IL-6 in CB2R antagonist-treated animals (**Supplemental Figures 5E-5G**). Only CD4^+^ and CD8^+^ T cells that produced GM-CSF were found to be decreased in animals that were treated with SRR144528 (**Supplemental Figure 5H**). In addition, administration of SR144528 had no effect on cleaved caspase 3 expression levels (**Supplemental Figure 5I**) when compared to vehicle-treated mice. Thus, these studies indicated that pharmacological blockade with a peripherally restricted CB2R inverse agonist/antagonist had no substantive effect on mitigating neuroinflammation or preventing neuronal cell death in GVHD mice.

**Figure 5:**
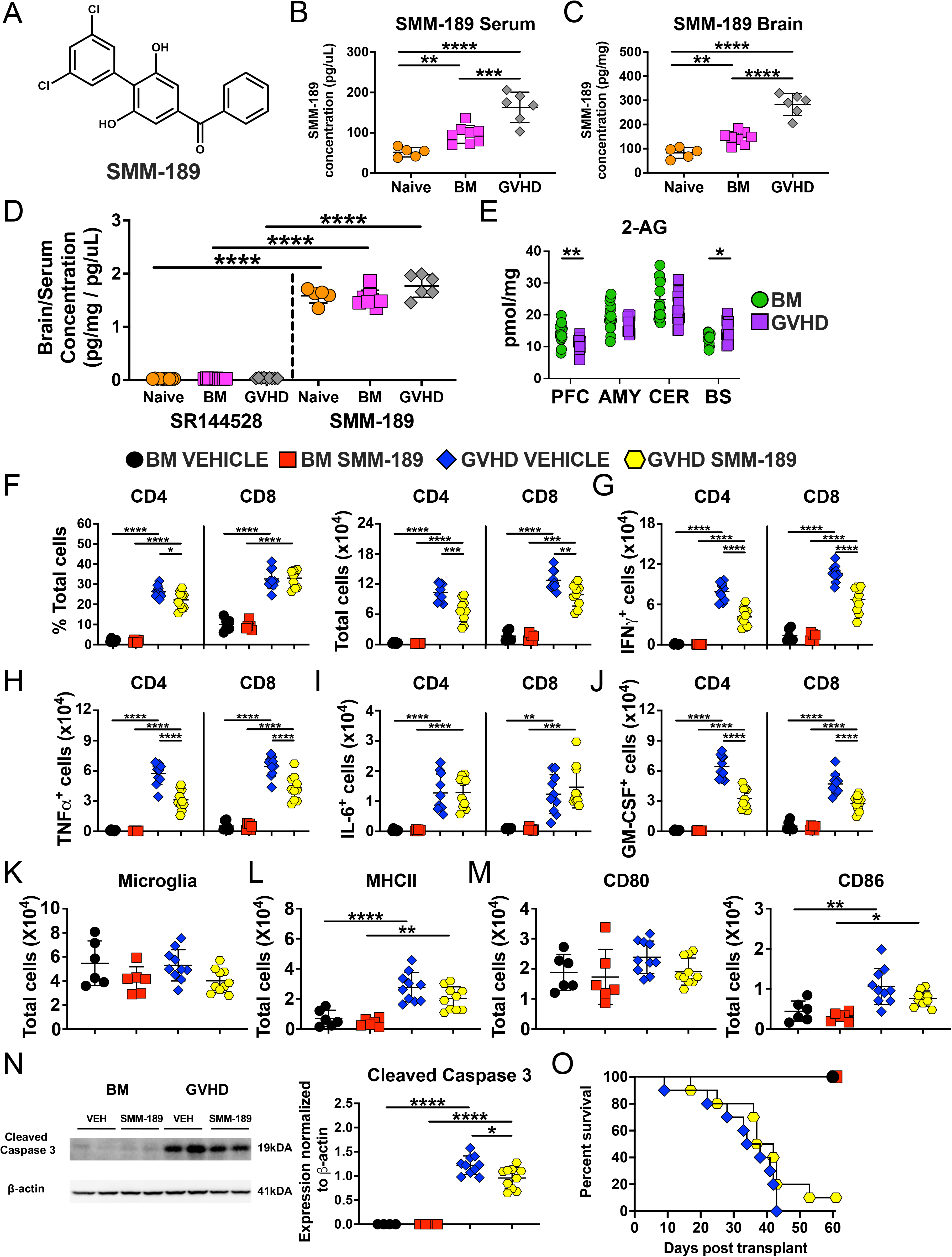
Pharmacological administration of a brain penetrant CB2R antagonist reduces inflammation in the CNS during GVHD. (A). Chemical structure of SMM-189. (B). Serum levels (pg/µl) of SMM-189 in naïve and lethally irradiated Balb/c mice transplanted with B6 BM alone or together with B6 spleen cells. (C). Concentration (pg/mg) of SMM-189 in the brain in naïve and lethally irradiated Balb/c mice transplanted with B6 BM alone or together with B6 spleen cells. (D). Ratio of brain to serum SR144528 and SMM-189 concentrations. Results are from two experiments. (E). Lethally irradiated Balb/c recipients were transplanted with B6 BM (5 x 10^6^) alone or together with B6 spleen cells (adjusted to yield an αβ T cell dose of 0.75 x 10^6^ cells). 2-AG levels in the amygdala, brainstem, cerebellum, and prefrontal cortex as determined by mass spectrometry are shown 14 days post transplantation. Results are from three experiments. (F-N). Lethally irradiated Balb/c recipients were transplanted with B6 BM and spleen cells (adjusted to yield an αβ T cell dose of 0.75 x 10^6^). Animals were then treated with a CB2R inverse agonist/antagonist, SMM-189 (6 mg/kg) or vehicle control for 14 days beginning on day 0. (F). The percentage and absolute number of donor derived CD4^+^ and CD8^+^ T cells in the brains of mice 14 days post transplantation. (G-J). The absolute number of CD4^+^ and CD8^+^ T cells that produced IFN-γ, TNF-α, IL-6 or GM-CSF. Data are from two experiments. (K). Absolute number of microglial cells. (L, M). Absolute number of MHC class II, CD80 and CD86 expressing microglial cells. (N). Representative western blot images and scatterplots depicting normalized expression of cleaved caspase 3 in the brain from mice treated with either SMM-189 or a vehicle control. Results are from two experiments. (O). Irradiated Balb/c recipients were transplanted with B6 BM and spleen cells. Animals were treated with SMM-189 (6 mg/kg) (n=10) or an equivalent amount of vehicle (n=10) for 14 days beginning on the day of transplant. Balb/c mice transplanted with B6 BM alone and then treated with either vehicle (n=6) or SMM-189 (n=6) served as controls. Survival is shown. Results are from two experiments. Statistics: *p<0.05, **p<0.01, ***p<0.001, ****p<0.0001.

To then determine whether host CB2R expressing cells within the brain played a critical pathophysiological role in promoting neuroinflammation, we examined the efficacy of SMM-189, which was synthesized in one of our labs (B.M. II). SMM-189 has a structurally unique tri-aryl core (43) (**Figure 5A**), is specific for the CB2R (43,44), and has been shown to reduce inflammation in murine models of traumatic brain injury (45), suggesting that this agent distributes into the brain. To confirm this premise, we performed isotope-dilution mass spectrometric analysis to measure SMM-189 concentrations in serum and the brain (**Supplemental Figure 6)**. We observed that GVHD mice had significantly higher concentrations of SMM-189 than BM control or naïve animals in both the serum (**Figure 5B**) and the brain (**Figure 5C**). In addition, these studies revealed that SMM-189 accumulates in the brain in all three groups with a brain/serum ratio of greater than one, whereas the ratio of brain to serum SR144528 concentration averaged only 0.02-0.03 (**Figure 5D**). Since SMM-189 competes with 2-arachidonoylglycerol (2-AG), the natural ligand for the CB2R (15), we measured 2-AG levels in specified regions of the brain to ascertain whether GVHD altered levels of this endocannabinoid. These studies demonstrated some discordant regional variation in the brainstem and PFC, but overall, no evidence that GVHD uniformly altered 2-AG levels (**Figure 5E**). Subsequent administration of SMM-189 resulted in a significant reduction in the frequency of CD4^+^ T cells, the absolute number of total CD4^+^ and CD8^+^ T cells (**Figure 5F**), and the total number of CD4^+^ and CD8^+^ T cells that produced IFN-γ, TNF-α, and GM-CSF (**Figures 5G-5J**). Treatment with SMM-189 had no effect on the absolute number of microglia **(Figure 5K)** or microglia with an activated phenotype (**Figure 5L-5M**) but did result in a significant decrease in the expression of cleaved caspase 3 (**Figure 5N**), indicative of reduced neuronal cell death. In addition, whereas administration of SR144258 exacerbated GVHD lethality (26), treatment with SMM-189 had no deleterious effect **(Figure 5O)**, demonstrating that CNS-directed blockade of CB2R signaling with SMM-189 selectively ameliorated neuroinflammation. Collectively, these results demonstrated that pharmacological blockade with a brain penetrant CB2R inverse agonist/antagonist attenuated GVHD-induced neuroinflammation and provided evidence that CB2R expression on a CNS resident population was critical for regulating inflammation in this tissue site.

### CB2R expression is increased on microglial cells during GVHD and is not regulated by IL-6

To define haematopoietically-derived recipient CB2R^+^ cells in the brain of GVHD mice, we gated on H-2K^b+^ CD45^+^ cells and observed that virtually all were CD45^lo^ CD11b^+^, indicative of a microglial cell phenotype (**Figure 6A**). Employing CB2R^EGFP^ reporter mice (46), we noted increased expression of the CB2R on microglia in the amygdala, brainstem, cerebellum, and prefrontal cortex in GVHD animals when compared to BM controls (**Figures 6B and 6C**), indicating that microglial cell expression of the CB2R was increased throughout the CNS under inflammatory conditions. This was further confirmed by immunofluorescence which demonstrated co-localization of GFP and TMEM119 expression in microglia from GVHD recipient CB2R^EGFP^ reporter mice (**Figure 6D**). Blockade of IL-6 signaling which has been shown to be ineffectual for the prevention and treatment of neuroinflammation occurring as a complication of immunotherapy in humans (47) had no effect on CB2R expression (**Figures 6E-6G**), demonstrating that microglial expression of CB2R was not regulated by IL-6. A more quantitative assessment revealed that the CB2R was expressed on only a small percentage (∼15%) of all microglia in GVHD animals (**Figure 6H**). Notably, the percentage of CB2R^+^ microglial cells that expressed MHC class II, CD80 and CD86 was significantly higher than CB2R non-expressing microglial cells (**Figures 6I and 6J**), indicating that CB2R expression was associated with an activated microglial phenotype.

**Figure 6:**
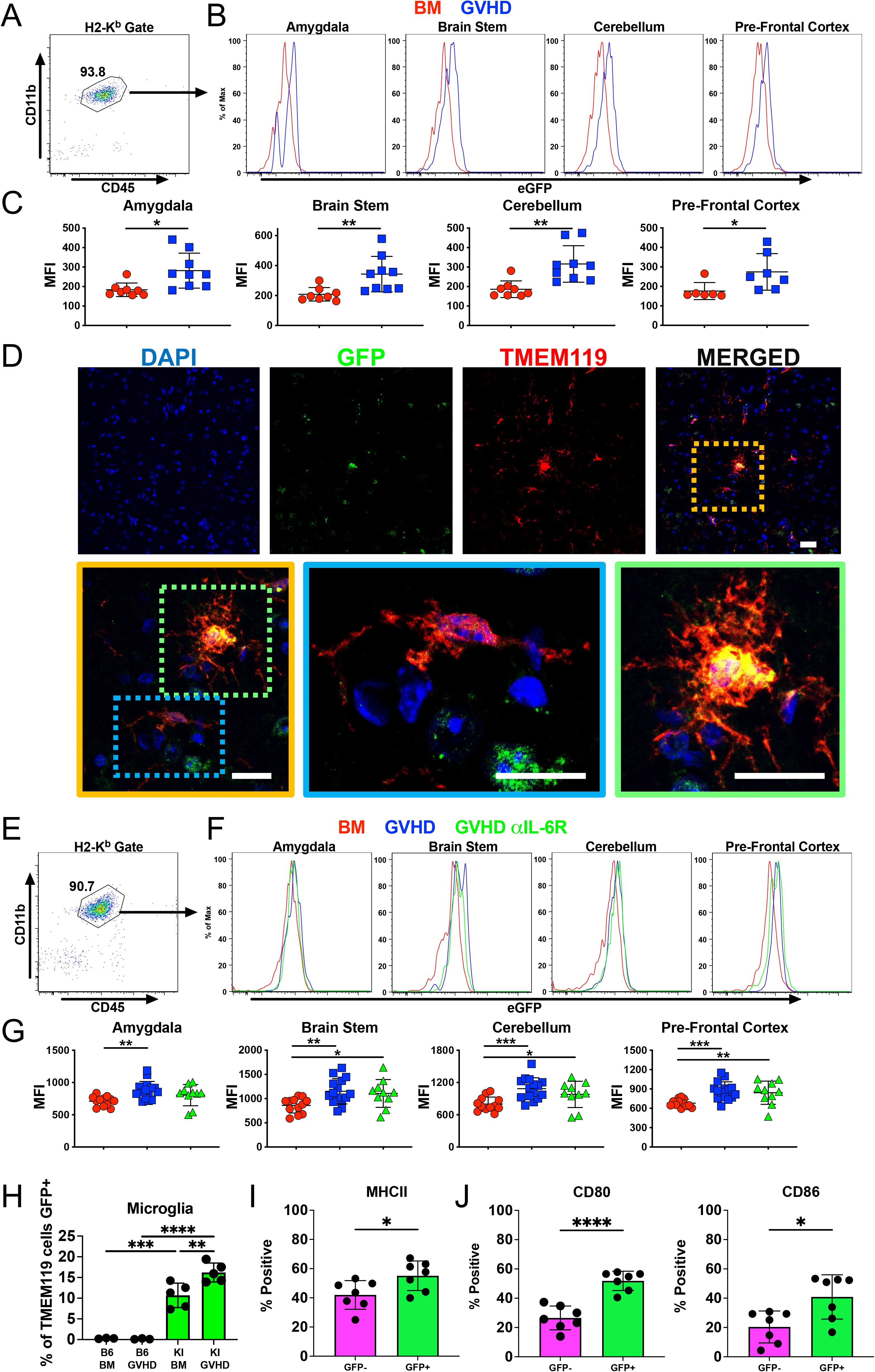
Microglial cell expression of the CB2R induces an activated phenotype and is not regulated by IL-6 signaling. (A-C). Lethally irradiated (1100c Gy) CB2R^EGFP+^ mice were transplanted with B10.BR BM alone or together with B10.BR spleen cells (adjusted to yield an αβ T cell dose of 5 x 10^6^). Animals were euthanized 14 days post transplantation. (A). Representative dot plot depicting recipient H-2K^b+^ CD45^+^ CD11b^+^ microglial cells. (B). Representative histograms depicting EGFP expression on recipient microglial cells obtained from the amygdala, brain stem, cerebellum, and prefrontal cortex of irradiated CB2R^EGFP^ animals reconstituted with B10.BR BM only (BM, red line) or CB2R^EGFP+^ mice transplanted with B10.BR BM and spleen cells (GVHD, blue line). (C). Scatterplot summary data showing cumulative median fluorescence intensity (MFI) shifts from four replicate experiments. Each data point represents results pooled from two mice. (D). Representative immunofluorescence staining showing CB2R (GFP), microglia (TMEM119), and merged images from the prefrontal cortex of lethally irradiated CB2R^EGFP+^ mice transplanted with B10.BR BM and spleen cells. Magnified yellow insert box depicts microglial cell that is CB2R^−^ (blue box) and one that expresses the CB2R (green box). White scale bar is 10 microns. (E-G). Lethally irradiated CB2R^EGFP+^ mice were transplanted with B10.BR BM alone or together with B10.BR spleen cells. Animals transplanted with adjunctive spleen cells were treated with either an anti-IL-6R or isotype control antibody on days 0 and 6 post transplantation. (E). Representative dot plot depicting recipient H-2K^b+^ CD45^+^ CD11b^+^ microglial cells. (F). Representative histograms depicting EGFP (CB2R) expression on recipient-derived microglial cells obtained from specified brain regions of CB2R^EGFP^ animals reconstituted with B10.BR BM only (BM, red line) or CB2R^EGFP+^ mice transplanted with B10.BR BM and spleen cells and then treated with isotype antibody (GVHD, blue line) or anti-IL-6R antibody (GVHD αIL-6R, green line) 14 days post transplantation. (G). Scatterplot summary data showing cumulative MFI shifts from five replicate experiments. Each data point represents results pooled from two mice. (H-J). Lethally irradiated CB2R^EGFP+^ (KI) or B6 mice were transplanted with B10.BR BM alone or together with B10.BR spleen cells. Percentage of microglial cells expressing CB2R (EGFP) is shown in panel H, and frequency of GFP^+^ (CB2R^+^) and GFP^−^ (CB2R^−^) microglial cells expressing MHC class II, CD80 and CD86 is depicted in panels I and J. Results are from two experiments. Statistics: *p<0.05, **p<0.01, ***p<0.001.

### CB2R expression on microglial cells regulates proinflammatory T cells and neuronal cell death in the brain

To further delineate the effect of CB2R expression on microglia and determine whether these cells had a more inflammatory signature, we performed scRNAseq analysis on sorted microglial cells obtained from the brains of WT versus CB2R^−/−^ GVHD recipient mice. This analysis revealed 6 transcriptionally distinct microglial clusters of which two were dominant (clusters 0 and 1) (**Figure 7A**). Two clusters that identified as T cells and macrophages, representing 1% of all cells, were deemed to be sort contaminants and were excluded from the analysis. Transcriptional analysis of these 6 clusters revealed 60 differentially expressed genes with 42 over expressed in microglia from WT and 18 over expressed in microglia from CB2R^−/−^ animals based on defined cutoff criteria (log 2-fold change) >0.2 and p_adjusted_<0.01, full list available in **Supplemental Table 4**), representing approximately 0.9% of the total sequenced transcriptome (**Figure 7B**). Microglial cells from WT mice had increased expression of inflammatory mediators such as *CCl3, CCl4* and *TNF,* whereas CB2R^−/−^ microglial cells had increased expression of genes associated with gamma interferon signaling such as the guanylate-binding proteins (*Gbp5* and *Gbp8)* (48) and *Iigp1* (49). To uncover biologically relevant pathways, we employed GSEA using the GO database which revealed increased expression of genes associated with TNF-alpha signaling, TGF-β responsiveness, leukocyte chemotaxis, and chemokine signaling in sorted WT microglia, whereas there was augmented expression of interferon gamma response genes in CB2R^−/−^ microglia **(Figure 7C)**. Similarly, GSEA using Hallmark gene annotation confirmed increased expression of genes associated with TNF-alpha signaling and interferon gamma response genes in WT and CB2R^−/−^ microglia, respectively (**Figure 7D)**. To determine whether CB2R expression on microglial cells was functionally significant and directly regulated neuroinflammation, we generated CX3CR1-Cre CB2R^fl/fl^ mice in which CB2R is deleted from microglial cells which express the fractalkine receptor, CX3CR1 (50), allowing microglia to be genetically targeted (51-53). Examination of CB2R expression on spleen cells from normal CX3CR1-Cre CB2R^fl/fl^ mice revealed that Cre-mediated deletion had no effect on B cells, T cells or macrophages (**Figure 7E**). In contrast, there was an 87% average reduction in CB2R expression on microglia from CX3CR1-Cre CB2R^fl/fl^ GVHD mice when compared to CB2R^fl/fl^ controls (**Figure 7F**), indicative of effective Cre-mediated recombination. To determine the functional significance of microglial CB2R expression, recipient CX3CR1-Cre, CB2R^fl/fl^, or CX3CR1-Cre CB2R^fl/fl^ animals were transplanted with MHC-mismatched BM and splenocytes from B10.BR mice. These studies revealed that there was a significant reduction in the absolute number of CD4^+^ and CD8^+^ T cells in the brains of recipient CX3CR1-Cre CB2R^fl/fl^ animals when compared to either CX3CR1-Cre or CB2R^fl/fl^ control mice (**Figure 7G**). In addition, we observed a significant reduction in the absolute number of CD4^+^ T cells that produced IFN-γ, TNF-α and GM-CSF, as well as CD8^+^ T cells that produced IFN-γ and TNF-α (**Figures 7H-7K**). There was also a corresponding decrease in cleaved caspase 3 expression in CX3CR1-Cre CB2R^fl/fl^ recipients (**Figure 7L**). Thus, microglial cell expression of the CB2R regulated TNF-α signaling and chemotaxis/chemokine signaling gene pathways, promoted the accumulation of proinflammatory CD4^+^ and CD8^+^ T cells, and augmented neuronal cell death in the brain during GVHD.

**Figure 7:**
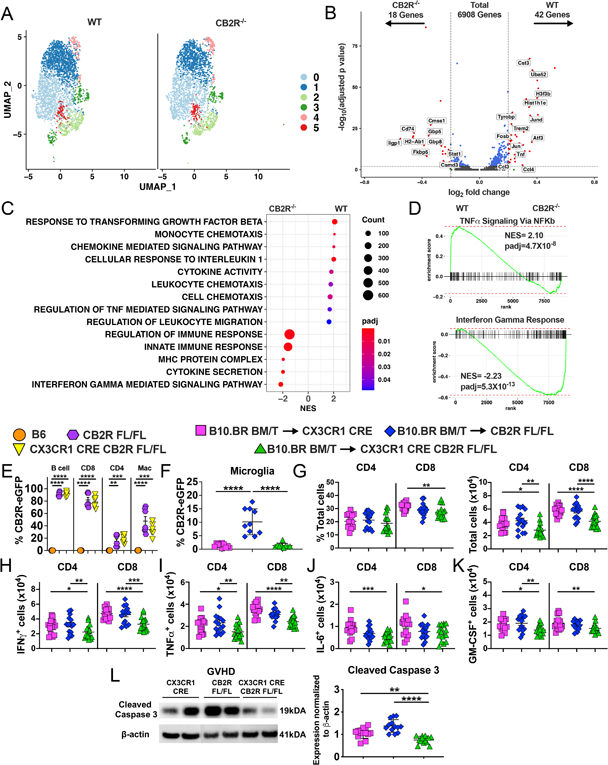
Conditional deletion of CB2R in microglial cells attenuates proinflammatory T cell accumulation and neuronal cell death. Lethally irradiated (1100 cGy) B6 or CB2R^−/−^ mice were transplanted with B10.BR BM and spleen cells (adjusted to yield an αβ T cell dose of 5 x 10^6^). Single live microglial cells from pooled brains (n=5/group) were sorted 14 days post transplantation. (A). UMAP dimensional reduction of scRNAseq data of flow-sorted live cells from pooled brains. Unsupervised clustering using Seurat revealed 6 transcriptionally distinct clusters using a resolution of 0.3. (B). Volcano plot showing over/underrepresented genes in aggregated microglial clusters from B6 versus CB2R^−/−^ mice. Cutoff parameters were |log2(fold change)| > 0.2 and p_adjusted_ <0.01. (C). Bubble plot demonstrating normalized enrichment score (NES) for pathways identified using the Gene Ontology database. (D). GSEA using Hallmark database comparing expression of TNF-α signaling and interferon response genes in microglial cells from B6 (WT) versus CB2R^−/−^ mice. (E). Percentage of CB2R expression on splenic B cells, CD8^+^ T cells, CD4^+^ T cells and macrophages in naïve, CB2R^EGFP^ fl/fl or CX3CR1-Cre CB2R^EGFP^ fl/fl mice (F-L). Lethally irradiated (1000 cGy) CX3CR1-Cre, CB2R^fl/fl^ or CX3CR1-Cre CB2R^fl/fl^ mice were transplanted with B10.BR BM and spleen cells (adjusted to yield an αβ T cell dose of 4 x 10^6^). Mice were euthanized in all cohorts 14 days post transplantation. (F). The percentage of microglial cells that expressed the CB2R based on eGFP expression. (G). The percentage and absolute number of donor derived CD4^+^ and CD8^+^ T cells in the brain. (H-K). The absolute number of CD4^+^ and CD8^+^ T cells that produced IFN-γ, TNF-α, IL-6 or GM-CSF. (L). Representative western blot images and bar graphs depicting normalized expression of cleaved caspase 3 in the brain. Data are from two-three experiments in panels E-L. Statistics: *p<0.05, **p<0.01, ***p<0.001, ****p<0.0001.

## DISCUSSION

Neuroinflammation in the brain is a recognized complication of immunotherapeutic approaches, such as allogeneic HSCT (5-9), immune checkpoint inhibitor treatment (1,2,54), and chimeric antigen receptor therapy (3,4) that are increasingly being used to treat patients with underlying malignant conditions. A distinguishing characteristic of this pathophysiological process is that the inciting immunological event, which is largely T cell driven, occurs in the periphery, but subsequently extends into the CNS due to T cells that cross the blood brain barrier and interact with resident immune cells in the brain (50). The mechanistic pathways by which cells of the adaptive and innate arms of the immune system intersect to promote a neuroinflammatory milieu in the CNS, however, have not been well delineated. Herein, using a murine model of GVHD, we have identified a critical role for CB2R signaling in the pathophysiology of immune-mediated CNS inflammation. Our studies indicate that host, but not donor, expression of the CB2R promotes the accumulation of proinflammatory T cells, increases microglial cell numbers, and induces neuronal cell death in the brain. Mechanistically, we show that microglial cell expression of the CB2R in the context of GVHD is associated with an activated phenotype characterized by increased expression of MHC class II and costimulatory molecules. Conversely, cell-specific deletion of the CB2R from microglia decreased the accumulation of inflammatory T cells, reduced expression of inflammatory and chemokine signaling gene pathways, and attenuated neuronal cell death. Furthermore, we demonstrate that targeting the CB2R with a brain penetrant, CB2R-specific inverse agonist/antagonist mitigated neuroinflammation without exacerbating systemic GVHD-induced lethality, indicating that this signaling pathway is amenable to CNS-directed pharmacological intervention.

Microglial cells have numerous functions that include phagocytosis, antigen presentation capabilities, and the production of inflammatory cytokines and chemokines (35,55), indicative of their role as a primary regulator of innate immunity in the CNS. During GVHD, we observed that these cells acquired an activated, inflammatory phenotype, characterized by a transcriptional profile that revealed increased expression of class I and II genes, along with a wide array of chemokine genes (i.e., *Ccl2*, *Ccl3*, *Ccl4*, *Ccl5*, *Ccl7*, *Ccl12*, *Cxcl9*, and *Cxcl10*) which function to recruit monocytes, T cells and other immune cells into the brain. This contrasted with the profile of microglia from BM control animals which maintained a more homeostatic phenotype (i.e., *P2ry12*, *Gpr34*, *Fscn1*, *Cx3CR1*, and *Fcrls*). Furthermore, applying DoRothEA analysis to the scRNAseq dataset, microglia from GVHD animals had increased expression of Stat, NF-KB and IRF family transcription factors that have been shown to be pathways by which microglia promote inflammation (56-58). The maintenance of microglial cell homeostasis is dependent upon interactions between the endogenous ligands CSF-1 and IL-34 with their cognate receptor, CSF-1R (38,39), and the genetic absence of IL-34 results in a profound reduction in microglial cell numbers (40). To formally define a role for microglia in mediating neuroinflammation, we employed IL-34^−/−^ mice as transplant recipients and demonstrated that these animals had a significant reduction in the absolute number of activated microglial cells, and that this was associated with a commensurate reduction in proinflammatory cytokine producing CD4^+^ and CD8^+^ T cells as well as a decrease in cleaved caspase 3 expression, indicative of reduced neuronal cell death. Thus, these data indicated that microglial cells play a critical role in mediating neuroinflammation during GVHD.

The critical roles that T cells and microglia have in mediating GVHD-associated neuroinflammation prompted us to examine the role of the type 2 cannabinoid receptor since this signaling pathway is known to regulate inflammatory cytokine production by T cells (25,27) and the receptor is also expressed on microglia (59). To address this question within the context of a murine GVHD model in which there are bidirectional immune responses, we first examined the role of donor CB2R expression and observed that absence of this receptor had very little effect on CNS inflammation. Whereas transplantation with CB2R^−/−^ marrow grafts resulted in a reduction in proinflammatory CD4^+^ T cells, there was no effect on CD8^+^ T cells nor did mice have any decrease in neuronal cell death as assessed by cleaved caspase 3 expression. Interestingly, prior studies in which animals were transplanted with donor CB2R^−/−^ marrow grafts (26) had demonstrated increased GVHD-induced lethality due to augmented systemic inflammation, indicative of discordant effects in the CNS versus periphery. In contrast, absence of host CB2R expression significantly reduced GVHD-induced neuroinflammation as evidenced by a decrease in the accumulation of proinflammatory, donor-derived CD4^+^ and CD8^+^ T cells, a reduced number of microglial cells that possessed an activated phenotype, and significantly less neuronal cell death. In addition, CB2R^−/−^ recipient mice had no increase in GVHD-induced mortality indicating that absence of recipient, in contrast to donor, CB2R expression did not result in discordant inflammatory responses in the periphery and the brain. Thus, recipient CB2R expression selectively regulated inflammation in the CNS during this disease.

Pharmacological targeting of the CB2R is complex and dependent upon several factors which include efficacy, ligand affinity, selectivity for intracellular signaling transduction pathways, and tissue distribution (42,60). Within the context of GVHD-induced neuroinflammation, tissue distribution is of particular importance since inflammation occurs concurrently within peripheral and central tissue sites, and our prior studies (26), coupled with current data, demonstrated that blockade of the CB2R signaling pathway resulted in discordant immune responses. Thus, additional questions which arise in this setting are whether effective pharmacological targeting is dependent upon penetration into the brain and if therapeutic blockade of the CB2R signaling pathway results in concordant or discordant effects in the periphery and CNS. To address these questions, we initially employed SR144528 which has been validated as the optimal CB2R inverse agonist/antagonist in murine studies due to its high selectivity profile (42). Mass spectrometry revealed that there was minimal accumulation of SR144528 in the brain under inflammatory conditions, demonstrating that this agent is essentially peripherally restricted. More importantly, treatment with this agent had only modest effects on the accumulation of proinflammatory T cells and did not mitigate neuronal cell death, indicating limited therapeutic utility in this context. In contrast, administration of a novel CB2R inverse agonist/antagonist, SMM-189, had excellent penetration into the brain which was further augmented under inflammatory conditions in GVHD recipients. Correspondingly, SMM-189 treatment of GVHD mice significantly reduced the accumulation of proinflammatory CD4^+^ and CD8^+^ T cells and decreased neuronal cell death. Furthermore, this agent had no adverse impact on overall survival indicating that, unlike SR144528 (26), there was no exacerbation of overall GVHD-related mortality. Thus, these results demonstrated the feasibility of therapeutically targeting the CB2R signaling pathway in the brain with an inverse agonist/antagonist that can cross the blood brain barrier and thereby ameliorate neuroinflammation. Presently, we do not know whether SMM-189 is acting primarily as a competitive antagonist (i.e., reducing endocannabinoid activation of CB2R signaling) or as an inverse agonist (i.e., reducing constitutive CB2R signaling). However, we did not find that GVHD produced consistent increases in brain 2-AG concentrations, which suggests that the inverse agonist efficacy of SMM-189 is of primary importance.

IL-6 has been shown to have a pivotal role in the pathophysiology of GVHD. Experimental studies in mice have demonstrated that blockade of this signaling pathway is effective for the prevention of GVHD target organ damage in the periphery (61,62). In addition, clinical studies have shown that inhibition of IL-6 by administration of tocilizumab, a humanized anti-IL-6R antibody, has efficacy for both the treatment and prevention of this disease (63-66). Within the context of GVHD-induced neuroinflammation, however, blockade of IL-6 signaling in pre-clinical murine models resulted in only partial protection and failed to correct the accumulation of neurotoxic kynurenine metabolites in the brain (11). Moreover, in other forms of immunotherapy such as CAR T cell administration, inhibition of IL-6 has proven efficacious for the abrogation of systemic side effects (67) but had little benefit for the prevention and/or treatment of neurotoxicity (47). Collectively, these data suggest that IL-6 independent pathways exist and contribute to immune-mediated neuroinflammation. In the current study, we observed that the increased expression of CB2R on microglial cells observed during GVHD was unaffected by administration of an anti-IL-6R antibody, and that this was evident in distinct regions of the brain that have unique functional roles. Thus, these results unveil the existence of an IL-6 independent pathway in the CNS that promotes inflammation, thereby providing a potential mechanistic explanation for the limited ability of IL-6-directed strategies to prevent and/or attenuate neurological complications that occur after GVHD and other forms of immunotherapy.

We observed that microglia were the only recipient hematopoietic population that expressed the CB2R in the brain, and that GVHD significantly increased expression of the receptor above that observed in BM control animals. This finding is consistent with prior studies which have shown that CB2R expression is very low in resting microglia but is upregulated by inflammatory signals, such as IFN-γ and granulocyte macrophage colony stimulating factor (68), that induce microglial progression to responsive and primed states (69). While microglial cells have been shown to express the CB2R in diverse models including Alzheimer’s, Huntington’s Disease (57), and traumatic brain injury, the role of receptor expression has been controversial. Whereas some studies have shown that CB2R activation inhibits cytokine release by activated microglia (70,71), others have demonstrated that *in vivo*-activated CB2R^−/−^ microglial cells release significantly lower amounts of proinflammatory cytokines than wild type microglia (72). A confounding factor has been that most studies have utilized global knockout CB2R^−/−^ animals which has precluded the ability to define the role of CB2R expression on microglial cells only (70). Furthermore, expression of this receptor on microglial cells has been difficult to quantitate accurately using alternative flow cytometric approaches due to antibody non-specificity and low receptor expression levels (73). To circumvent this obstacle, we created and utilized a CB2R reporter mouse (46) that faithfully identifies CB2R expressing immune cells (26). Using this reporter mouse, we observed somewhat surprisingly that only ∼10-15% of microglial cells expressed the CB2R during GVHD-induced inflammation. To address the functional role of this receptor on microglial cells, we selectively deleted CB2R from these cells and observed that recipients had reduced accumulation of proinflammatory T cells along with a reduction in neuronal cell death, indicating that a minor population of CB2R expressing microglia exerted potent immunomodulatory effects in the brain.

To delineate mechanistic pathways by which microglial expression of the CB2R modulated inflammatory responses in the brain, we performed GSEA on sorted microglial cells from WT or CB2R^−/−^ recipient mice. This analysis revealed higher expression of genes associated with chemotaxis, chemokine signaling and leukocyte migration in microglia from WT compared to CB2R^−/−^ recipients, supporting the premise that one of the main functions of CB2R signaling is to promote leukocyte recruitment into the brain during GVHD. These results are consistent with increased expression of chemokine pathway genes in microglia of GVHD mice when compared to BM controls (Figure 2). GSEA also revealed enrichment of genes associated with TNF-α signaling in recipients of WT compared to CB2R^−/−^ grafts, suggesting that activation of TNF-α mediated signaling may be another mechanistic pathway by which CB2R expression on microglia augments neuroinflammation. This observation is consistent with a prior report in which microglial cell production of TNF-α was shown to play a role in the pathogenesis of CNS inflammation during GVHD (14). Interestingly, we noted that microglia from recipient CX3CR1-Cre CB2R^fl/fl^ mice, that were protected from neuroinflammation, had increased expression of genes associated with the interferon gamma signaling pathway. Whereas IFN-γ has been shown to promote CNS inflammation, its role appears to be more nuanced as it can also exert anti-inflammatory effects in certain contexts and antagonize proinflammatory functions of TNF-α (74). Thus, another role of microglial CB2R signaling, in addition to the regulation of chemotaxis and leukocyte recruitment, may be to alter the balance between TNF-α and IFN-γ signaling pathways, and thereby regulate neuroinflammation during GVHD.

In summary, these studies have identified a critical role for the CB2R signaling pathway in the regulation of GVHD-induced neuroinflammation. Mechanistically, CB2R expression on microglia was associated with an activated inflammatory phenotype, potentiated the accumulation of donor-derived proinflammatory T cells, regulated chemokine and TNF-α gene regulatory networks, and promoted neuronal cell death. Conversely, administration of a brain penetrant CB2R inverse agonist/antagonist attenuated CNS inflammation without exacerbating systemic GVHD, indicating that this pathway can be therapeutically targeted which has clinical implications for the mitigation of GVHD-induced neuroinflammation and potentially other forms of immunotherapy that induce neurological dysfunction.

## EXPERIMENTAL MODEL AND SUBJECT DETAILS

### Mice

C57BL/6 (B6) (H-2^b^), Balb/c (H-2^d^), B10.BR (H-2^k^), B6 ACTb-EGFP, CX3CR1-Cre, CB2R^−/−^, IL-34^−/−^, CB2R^fl/fl^ eGFP, and CX3CR1-Cre CB2R^fl/fl^ eGFP mice were bred in the Biomedical Resource Center (BRC) at the Medical College of Wisconsin (MCW) or purchased from Jackson Laboratories (Bar Harbor, ME). CB2R^EGFP^ reporter mice were constructed by inserting an enhanced green fluorescent gene preceded by an internal ribosomal entry site (IRES) into the 3’ untranslated region of the *cnr2* mouse gene and have been previously described (46). In addition, the entire exon 3, including the 3′ UTR and knocked-in reporter, is flanked by *lox*P sites which allows for the conditional inactivation of the *cnr2* gene in cells expressing Cre recombinase. CB2R^-/-^ mice were generated by mating the CB2R^EGFP^ mice to Cre-deleter females and have been previously described (46). IL34^−/−^ mice that were created by using Cre recombinase to delete exons 3-5 and replacing them with an IRES-lacZ construct have been described (40). All animals were housed in the Association for Assessment and Accreditation of Laboratory Animal Care (AAALAC)-accredited Biomedical Resource Center of the Medical College of Wisconsin. Experiments were all carried out under protocols approved by the MCW Institutional Animal Care and Use Committee. Mice received standard 5L0D mouse chow and acidified tap water ad libitum.

### Bone Marrow Transplantation

Bone marrow (BM) was flushed from donor femurs and tibias with Dulbecco’s modified media (DMEM) (Thermo Fisher Scientific, Waltham, MA) and passed through sterile mesh filters to obtain single cell suspensions. Splenocytes were processed by mechanical disruption and red blood cells were lysed using Tris-buffered ammonium chloride (ACT) solution. Recipient mice were conditioned with total body irradiation (TBI) administered as a single exposure at a dose rate of 900-1100cGy using a Shepherd Mark I Cesium Irradiator (J.L. Shepherd and Associates, San Fernando, CA). Irradiated recipients received a single intravenous injection in the lateral tail vein of BM with or without added spleen cells. Mice were weighed two-three times per week and were euthanized when they attained pre-defined morbidity criteria.

## METHOD DETAILS

### Reagents

Anti-IL-6R antibody (MR-16-1) (Chugai Pharmaceuticals, Tokyo, Japan) is a rat IgG antibody that has been previously described (58). Animals received a loading dose of 2 mg intravenously on day 0, and then were treated with 0.5 mg on day 7 by intraperitoneal injection. Rat IgG (Jackson Immunoresearch Laboratories, West Grove, PA) was used as a control for MR-16-1. SR144528 is a CB2R inverse agonist/antagonist and was purchased from Tocris Bioscience (Bristol, UK). SR144528 was dissolved in 100% ethanol at a concentration 20 times greater than the final concentration, then an equal volume of Cremophor EL was added with vigorous vortexing, followed by the dropwise addition of sterile saline (18 times the volume of ethanol). Mice received a 3 mg/kg intraperitoneal injection of SR144528 daily. Control mice received vehicle (1:1:18 ratio of ethanol, Cremophor EL, and saline) only intraperitoneal injections. SMM-189, a CB2R inverse agonist/antagonist (43,44) was prepared using the same method as for SR144528 and administered at a dose of 6 mg/kg via daily intraperitoneal injection. Control mice received vehicle (1:1:18 ratio of ethanol, Cremophor EL, and saline) only daily intraperitoneal injections.

### Isolation of Cells

Cells were isolated from the brain by mechanical disruption followed by collagenase D digestion (Roche Pharmaceuticals, Mannheim, Germany). The resulting cell suspension was resuspended in 40% Percoll (GE Healthcare Biosciences, Uppsala, Sweden) in DMEM and layered on 70% Percoll in DMEM. The resultant gradients were centrifuged at 800g with no brake or acceleration at 4°C for 30 minutes. The interface was collected for further analysis. Cell counts were obtained from half of a brain for each experimental animal unless otherwise specified.

### Flow Cytometry

Isolated cells from the brain were labelled with LIVE/DEAD Fixable Aqua Dye (Thermo Fisher Scientific, Waltham, MA) according to manufacturer’s instructions. Cells were then stained with monoclonal antibodies conjugated to fluorescent molecules as listed in Supplemental Table 5 after treatment with Fc Block (BD Biosciences, Franklin Lakes, NJ). Cells were analyzed on either a BD LSR-II or BD LSRFortessa X-20 flow cytometer running BD FACSDiva software or a Cytek Aurora spectral cytometer running SpectroFlo software and analyzed using FlowJo software (TreeStar).

### Intracellular Cytokine Staining

Lymphocytes isolated from brain were stimulated with 50ng/ml PMA, 750ng/ml Ionomycin, and 2µM monensin (Thermo Fisher Scientific) for 3.5 hours. Cells were subsequently stained for viability and surface antigens and then intracellularly stained using the eBioscience Intracellular Fixation and Permeabilization Kit (Thermo Fisher Scientific) using the antibodies and reagents listed in Supplemental Table 5.

### cDNA Preparation

Total RNA was extracted from tissues by homogenization in Trizol (Thermo Fisher Scientific) followed by Phenol/Chloroform extraction, washing with isopropyl alcohol followed by 75% EtOH, drying pellets and resuspending in RNAse free water. cDNA was then made using the QuantiTect Reverse Transcription Kit (Qiagen, Valencia, CA).

### Real-time q-PCR

Real-time q-PCR was performed using QuantiTect SYBR Green PCR Kit (Qiagen) and run in a CFX C1000 Real-time Thermal Cycler (Bio-Rad, Hercules, CA). The 18S reference gene was amplified using QuantiTect Primer Assay Kit (Qiagen). The primers were purchased from Integrated DNA Technologies (Coralville, IA) and are listed in Supplemental Table 6. Primer specificity was verified by melt curve analysis. To calculate fold change in gene expression, the average C(t) value from triplicate wells was compared to the average 18S C(t) value from triplicate wells.

### Fluorescent Microscopy and Image Analyses

For immunofluorescence, mice were first perfused with cold PBS, followed by perfusion with 4% PFA/PBS. Harvested tissues were left in 25% sucrose/PBS overnight at 4°C. 10-40 μm-thick tissue cryosections were cut and stored at −20°C in glycerol/ethylene glycol-based antifreeze solution. Floating sections were incubated in PBS 2 times for 10 minutes min at room temperature before blocking with 5% normal goat serum in PBS for one hour. Tissue was then incubated with primary conjugated antibodies in staining buffer (PBS/ with 2% BSA/ and 0.1% Sodium Azide) with 0.1% Triton X-100 (1:1000) overnight at 37°C. Sections were then washed 2 times for 10 min each time with PBS and secondary antibodies were applied in PBS (1:500) for 2 hours if necessary. Lastly, sections were washed 3 times for 10 min each time with PBS and mounted with ProLong Gold antifade reagent containing DAPI (Invitrogen, Carlsbad, CA). All images were acquired with a confocal microscope (Leica SP8 Upright Confocal Microscope, Buffalo Grove, IL USA). The brightness/contrast of the acquired digital images was applied equally across the entire image and equally to control images and analyzed using Adobe Photoshop CS4 software (Adobe Systems Inc., San Jose, CA) and Image J/FIJI software (National Institute of Health, Bethesda, MD). Cells were quantified as a measure of number per volume or area/volume fraction per volume in 5-8 fields of view (FOV)/hemisphere/region per mouse. Automated protocols for signal intensity (SI) in a 1×10^6^/μm^3^ voxel and masks were created using ∼2SD SI threshold from the Mean Fluorescence Intensity (MFI) with background SI subtracted. Coronal/Sagittal brain maps were created utilizing Adobe Photoshop CS4 software. The following antibodies were purchased from AbCAM: Rat monoclonal anti-CD3 [CD3-12] (ab11089) and polyclonal anti-Cleaved Caspase-3 (ab4051). Polyclonal anti-IBA-1 was purchased from WAKO. Anti-NeuN [A60] (MAB377) was purchased from EDM Millipore. Polyclonal GFP Antibody (A10262) and all fluorescently conjugated secondary antibodies were purchased from Thermo Fisher.

### Western blot

Western brain lysates were homogenized in RIPA buffer with phosphatase/protease inhibitors (Thermo Fisher Scientific) and proteins were separated on SDS-polyacrylamide gels (Invitrogen Life Technologies) before being transferred to PVDF membranes. Transferred membranes were washed with TBST and incubated with anti-Spectrin [AA6] (EDM Millipore, Burlington, MA), anti-Cleaved Caspase 3 [ASP175], (Cell Signaling, Danvers, MA), or anti-β-actin (1:5000 Sigma Aldrich) in 5% milk in 0.2% Tween 20/TBS, 4°C, overnight. Membranes were washed and stained with appropriate HRP-labeled secondary antibodies and incubated with enhanced chemiluminescence (ECL) based substrates before being imaged utilizing ChemiDoc Bio-Rad imaging system (Hercules, CA). Band intensities were quantified using Image J/FIJI software (National Institute of Health, Bethesda, MD).

### Brain 2-AG Quantification

Brains were harvested from animals and immediately frozen in liquid nitrogen. Frozen brains were thawed enough for dissections; brain regions were quickly weighed then transferred to a liquid nitrogen cooled mortar and disrupted to a fine powder with the pestle. Pulverized, frozen samples were transferred to heavy-walled borosilicate glass tubes with a cell scraper. Two mL of cold acetonitrile and 9 µL of internal standard (deuterated 2-AG; 1800 pg/µl) were added to each tube. After brief homogenization with a glass rod, tubes were transferred to a bath sonicator at 4°C and sonicated for 30 minutes. Protein precipitation was carried out in each tube at −20°C for several hours. Samples were centrifuged at 2400rpm for 3 minutes and the supernatant was transferred to a fresh tube and dried down under nitrogen. Samples were resuspended in 400µL of methanol, and dried. Finally, samples were resuspended in 60µL of methanol. Samples were analyzed with an Agilent 6460 Triple Quadrupole Mass Spectrometer equipped with an HPLC system (Agilent Technologies, Santa Clara, CA), 5µL were used for injection following the methods reported previously (75).

### Mass Spectrometry and Assessment of Brain/Blood Distribution of SMM-189 and SR144528

SMM-189 and ^2^H-SMM-189 (SMM-189-D5) were synthesized as described previously (43). SR144528 was purchased from Abcam (Waltham, MA, catalog # ab146185) and ^2^H-SR144528 (SR144528-D7) was provided by the National Institute on Drug Abuse Drug Supply Program (Research Triangle Park, NC). HPLC-grade acetonitrile and water were obtained from Fisher Scientific (Waltham, MA) and reagent-grade formic acid from Sigma Aldrich. SMM-189 and SR144528 concentrations were measured in brain and whole blood using high performance liquid chromatography (HPLC) tandem mass spectrometry (MS/MS). HPLC separation was carried out on a Phenomenex Luna Omega PS C18 50×2.1 mm column (1.6 μm particle size) and MS was carried out on an Agilent 6460 mass spectrometer with Jetstream ionization source and equipped with an Agilent 1290 binary pump and autosampler.

For SMM-189, the aqueous mobile phase (A) was 0.1% formic acid in HPLC-grade water and the organic mobile phase (B) was 0.1% formic acid in acetonitrile. The flow rate was set to 0.2 mL/min. and the elution gradient was held at 10% B for 1.5 minutes, increased to 100% B over the next 6 minutes, held at 100% for 2.5 minutes, returned to 10% B over 2.5 minutes, and held at 10% B for an additional 2.5 minutes. The mass spectrometer was operated in the negative ion mode and the identified ion transitions for SMM-189 and SMM-189-D5 were 356.9 → 184.8 and 361.9 → 184.8, respectively. Retention times for both SMM-189 and SMM-189-D5 were between 6.5 and 7 min. Optimum ion signal was obtained with fragmentor voltage set to 160V and collision-induced dissociation at 25V. Quantification was carried out using a standard curve with 10, 50, 100, 500, 1000, 5000, 10000 ng/mL SMM-189, each also containing 1000 ng/mL SMM-189-D5 prepared in the initial mobile phase (90:10 0.1% formic acid in HPLC grade water: 0.1% formic acid in acetonitrile). A linear standard curve (concentration ratio versus peak area ratio) was generated from the prepared standards with 1/x^2^ weighting applied to the concentration ratio. To assess the brain and blood concentrations of SMM-189, the drug was prepared as described in “Reagents” and mice were injected intraperitoneally with SMM-189 (3 mg/kg) twice at 24 h intervals. Tissue collections were performed one-hour post-injection on the second day of treatment. Animals were anesthetized with isoflurane and then decapitated. Trunk blood was collected, and clotting was prevented by addition of 10% sample volume 0.5M K_2_EDTA in water. All samples were kept at - 80°C until analysis. Brains were weighed and homogenized and sonicated in phosphate buffered saline (2 μL PBS/1 mg tissue) until a homogenous solution was obtained. For extraction, 50 μL of brain homogenate or whole blood were added to 200 μL of ice-cold acetonitrile containing 50 ng of SMM-189-D5. Samples were vortexed for 1 minute and then sonicated in ice water bath for 10 minutes. After, they sat on ice for 20 minutes and were centrifuged at 3000 x g for 10 minutes. Supernatants were dried under N_2_ gas and resuspended in 50 μL acetonitrile for analysis. Tissue samples from untreated mice were extracted and no quantifiable data were obtained.

For SR144528, the aqueous mobile phase (A) was 0.1% formic acid in HPLC-grade water and the organic mobile phase (B) was 0.1% formic acid in acetonitrile. The flow rate was set to 0.2 mL/min. and the elution gradient was held at 30% B for 1.5 minutes, increased to 100% B over the next 6 minutes, held at 100% for 2.5 minutes, returned to 30% B over 2.5 minutes, and held at 10% B for an additional 2.5 minutes. The mass spectrometer was operated in the positive ion mode and the identified ion transitions for SR144528 and SR144528-D7 were 476.2 → 105.0 and 483.2 → 112.1 respectively. Retention times for both SR144528 and SR144528-D7 were between 5.5- and 6-min Optimum ion signal was obtained with fragmentor voltage set to 140V and collision-induced dissociation at 18V. Quantification was carried out using a standard curve with 1, 5, 10, 50, 100, 250, 500, ng/mL SR144528, each also containing 100 ng/mL SR144528-D7 prepared in the initial mobile phase (70:30 0.1% formic acid in HPLC grade water: 0.1% formic acid in acetonitrile). A linear standard curve (concentration ratio versus peak area ratio) was generated from the prepared standards with 1/x^2^ weighting applied to the concentration ratio. To assess the brain and blood concentrations of SR144528, the drug was prepared as noted in “Reagents” and mice were injected intraperitoneally with SR144528 (6 mg/kg) twice at 24 h intervals. Tissue collections were performed one-hour post-injection on the second day of treatment. Animals were anesthetized with isoflurane and then decapitated. Trunk blood was collected, and clotting was prevented by addition of 10% sample volume 0.5M K_2_EDTA in water. Blood was centrifuged at 1000 x g for 10 min and the resulting plasma was harvested. All samples were kept at −80°C until analysis. Brains were weighed and homogenized and sonicated in phosphate buffered saline (2 μL PBS/1 mg tissue) until a homogenous solution was obtained. For extraction, 100 μL of brain homogenate or plasma were added to 400 μL of ice-cold acetonitrile containing 40 ng of SR144528-D7. Samples were vortexed for 1 minute and then sonicated in ice water bath for 10 minutes. After, they sat on ice for 20 minutes and were centrifuged at 3000 x g for 10 minutes. Supernatants were dried under N_2_ gas and resuspended in 50 μL acetonitrile for analysis. Tissue samples from untreated mice were extracted and no quantifiable data were obtained.

### Single Cell RNA Sequencing

Cells were isolated from the brains of mice that were transplanted with BM alone or BM plus adjunctive spleen cells (GVHD) by collagenase D digestion and Percoll density centrifugation, and then stained with LIVE/DEAD Fixable Aqua Dye (Invitrogen) according to the manufacturer’s instructions. For the analysis of immune cells in BM versus GVHD mice, cells were sorted for live single cells, resuspended in PBS with 2% BSA at a concentration of 700-800 cells per microliter, and used as input for a target cell recovery of 5000 cells. For the analysis of microglial cells from WT versus CB2R^−/−^ animals, cells were sorted for live single recipient (H2K^b+^) microglia (CD45^low^, CD11b^+^), resuspended in PBS with 2% BSA at a concentration of 700-800 cells per microliter, and used as input for a target cell recovery of 5000 cells. Cells from five individual mice were pooled for each sample and were sorted on a BD FACSAria II cell sorter using a 100-micron nozzle. Cells were processed using the 10X Genomics Chromium controller and either the Chromium GEM Single Cell 3’ Reagent Kit v3 (10X Genomics, catalog # 1000092) or the Chromium Next GEM Single Cell 5’ Reagent Kits v2 (Dual Index) (catalog #1000265). Specifically, sequencing of immune cell populations in the brain from BM control versus GVHD animals was done with the Chromium Next GEM Single Cell 5’ Reagent Kit v2, while sequencing of microglia from WT and CB2R^−/−^ mice was done using the Chromium Single Cell 3’ Reagents Kit v3. 3’ or 5’ gene-expression (GEX) libraries were generated using 10X Genomics’ library preparation kits according to the manufacturer’s protocol. Following library preparation, 10X GEX libraries were converted and PCR amplification was carried out using Singular Genomics’ non-indexed S1-SP1 and S2-SP2 PCR primers at a final concentration of 0.3 μM each. PCR was performed with 2 ng of library input per reaction and 7 cycles of amplification using Q5 DNA polymerase (New England Biolabs, catalog # M0491S). Libraries were quantified using a Qubit 4 Fluorometer (Thermo Fisher Scientific) using the High Sensitivity dsDNA Assay kit (Thermo Fisher Scientific; catalog # Q32854). The quality of the libraries was assessed on a TapeStation 4200 (Agilent Technologies) using High Sensitivity D5000 ScreenTapes (Agilent Technologies; catalog # 5067-5592). Sample-specific GEX libraries were independently loaded into separate lanes of the G4 F2 flow cells and run on Singular Genomics’ G4 sequencer using G4 F2 Sequencing Kits (100 cycles) with the following conditions: 28 cycles for read 1 and 91 cycles for read 2.

### Single-Cell Gene Expression Data Analysis

Single-cell gene-expression data were processed using Cell Ranger (v.7.0.1, 10X Genomics) and aligned to the mouse (mm10) 2020-A reference. The resulting GEX matrices were analyzed using Seurat (76) (version 4.3.0) within the R statistical environment. For the initial QC step, cells were filtered out cells that expressed <200 or >6,000 genes for the BM versus GVHD analysis, and <200 or >7500 genes for the WT versus CB2R^−/−^ analysis. Cells with >10% mitochondrial transcripts were removed. scRNAseq datasets were integrated and then cell cycle scores were regressed when scaling gene expression values. Dimensionality reduction by principal component analysis was then performed on scaled and normalized data taking into consideration only the 2000 most variable genes in the dataset. Clustering was then performed on dimensionally reduced data using a shared nearest neighbor modularity optimization-based clustering algorithm based on the first 20 principal components and a clustering resolution of 0.5 for BM versus GVHD dataset and a clustering resolution of 0.3 for WT versus CB2R^−/−^ dataset. Clustered data was then visualized using the UMAP algorithm using Seurat’s default parameters.

### Gene Set Enrichment Analysis (GSEA)

To compare expression of certain gene sets between Seurat clusters, we used an R-based implementation of the GSEA method as previously published (77). Differentially expressed genes were first identified and were then used as an input for GSEA. Hallmark and GO gene sets were loaded from MSigDB and normalized expression scores and adjusted p values were obtained using fgsea.

### Dorothea Analysis

Regulon analysis was done using an R-based implementation of DoRothEA (36). Transcription factor-target interactions with confidence levels of A-C were used and the VIPER wrapper was run to determine the normalized enrichment scores for each transcription factor. Pheatmap was then used to graph the normalized enrichment scores.

## Data and Code Availability

The scRNA sequence data from this paper will be made available in the GEO database. All other raw data is available from the corresponding author upon request.

## Statistics

Data were analyzed using either a T test with Welch’s correction for two groups, a one-way ANOVA with Fisher’s LSD test for three groups, or a one-way ANOVA with Tukey’s test for four groups using GraphPad Prism software. Survival curves were compared using the log rank test. Results were considered significant at a p value of less than 0.05.

## AUTHOR CONTRIBUTIONS

A.M. and A.R. performed animal studies, flow cytometric analysis, single cell RNA sequencing, immunofluorescence staining, and helped write the manuscript. G.S. performed mass spectrophotometric analysis. C.C.Y. performed experimental research. B.M. provided pharmacological CB2R antagonists. M.C. and J.R. provided critical reagents. R.S., W.C., and A.Z. assisted with single cell RNA sequence analysis, and edited the manuscript. C.J.H. and W.R.D. developed the overall concept, designed experiments, supervised the study, and wrote the manuscript.

## ACKNOWLEDGMENTS

This research was supported by a grant from the National Institutes of Health (HL154579) to W.R.D. and C.J.H, and the Ministerio de Ciencia e Innovación / Agencia Estatal de Investigación (PID2019-108992RB-I00) to J.R. We thank Dr. Michael Thomas and the Department of Pharmacology Mass Spectroscopy Facility for assistance with the mass spectrometry analysis.

