## Supplemental Data for "MICROGLIAL CELL EXPRESSION OF THE TYPE 2 CANNABINOID RECEPTOR REGULATES IMMUNE-MEDIATED NEUROINFLAMMATION"

### Supplemental Figure 1

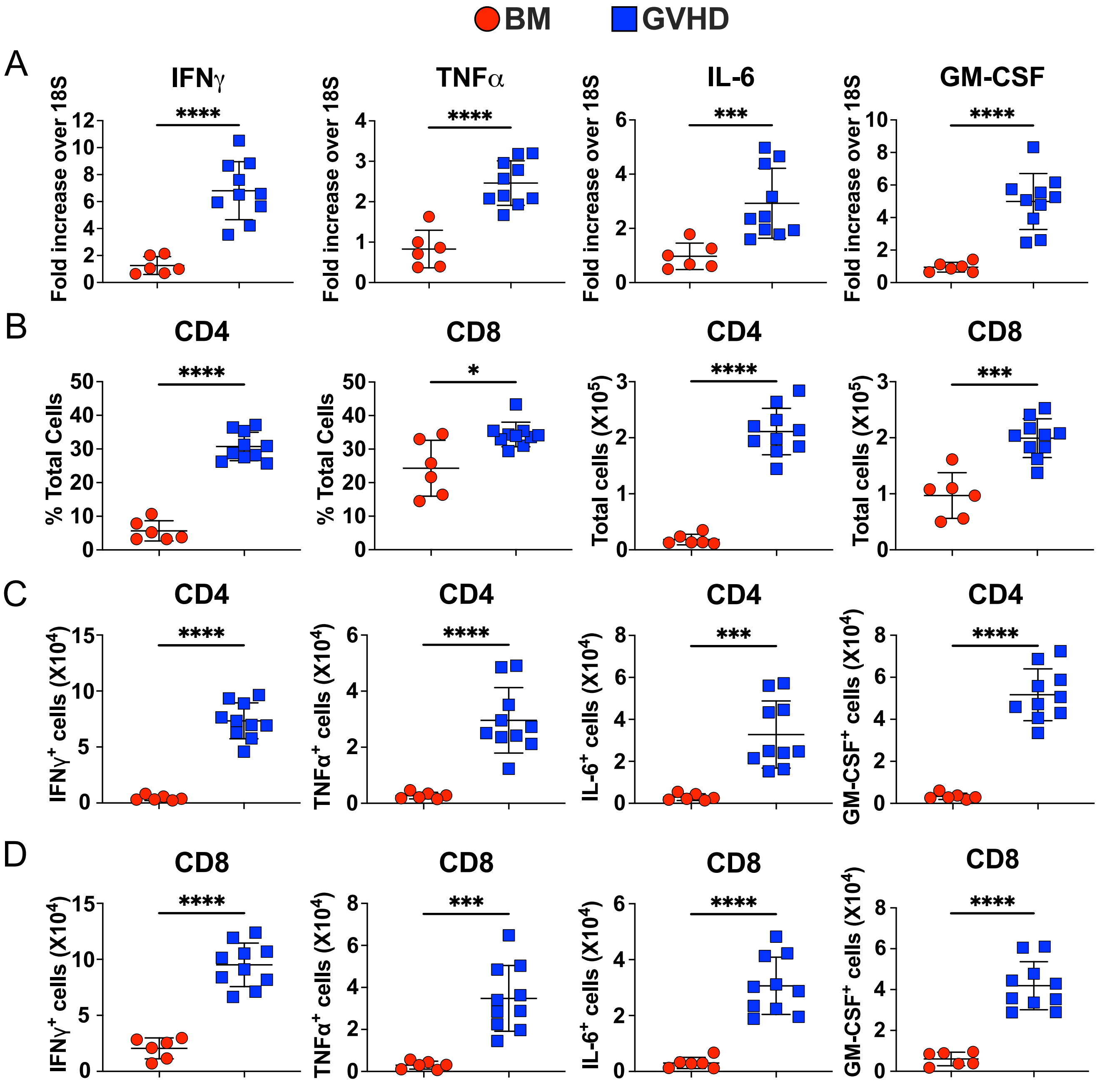

### Supplemental Figure 2

A

| Cluster | Cell Type | BM | GVHD | BM % | GVHD % |
|---|---|---|---|---|---|
| 0 | Microglia | 1046 | 404 | 42.7 | 11.2 |
| 1 | Microglia | 570 | 665 | 23.3 | 18.4 |
| 2 | CD8 T cell | 52 | 762 | 2.1 | 21.1 |
| 3 | Macrophage | 233 | 279 | 9.5 | 7.7 |
| 4 | CD4 T cell | 16 | 491 | 0.7 | 13.6 |
| 5 | CD8 T cell | 47 | 308 | 1.9 | 8.5 |
| 6 | Microglia | 185 | 107 | 7.6 | 3.0 |
| 7 | Microglia | 21 | 203 | 0.9 | 5.6 |
| 8 | Microglia | 113 | 83 | 4.6 | 2.3 |

B

#### T cells

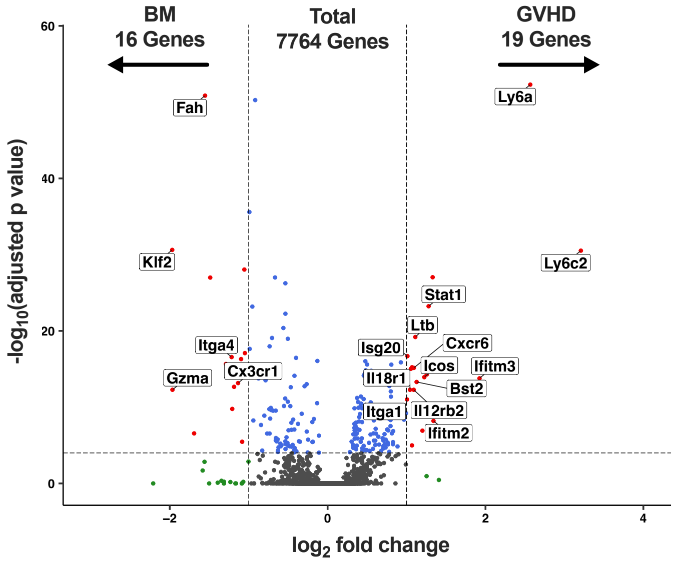


C

#### Macrophages

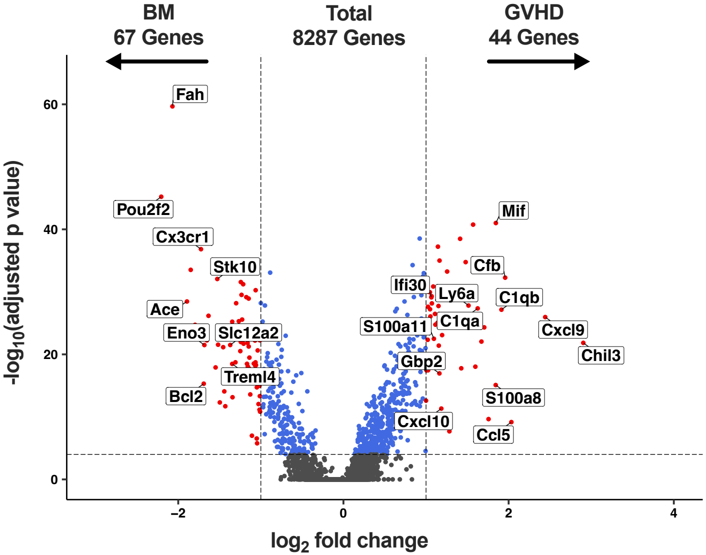

### Supplemental Figure 3

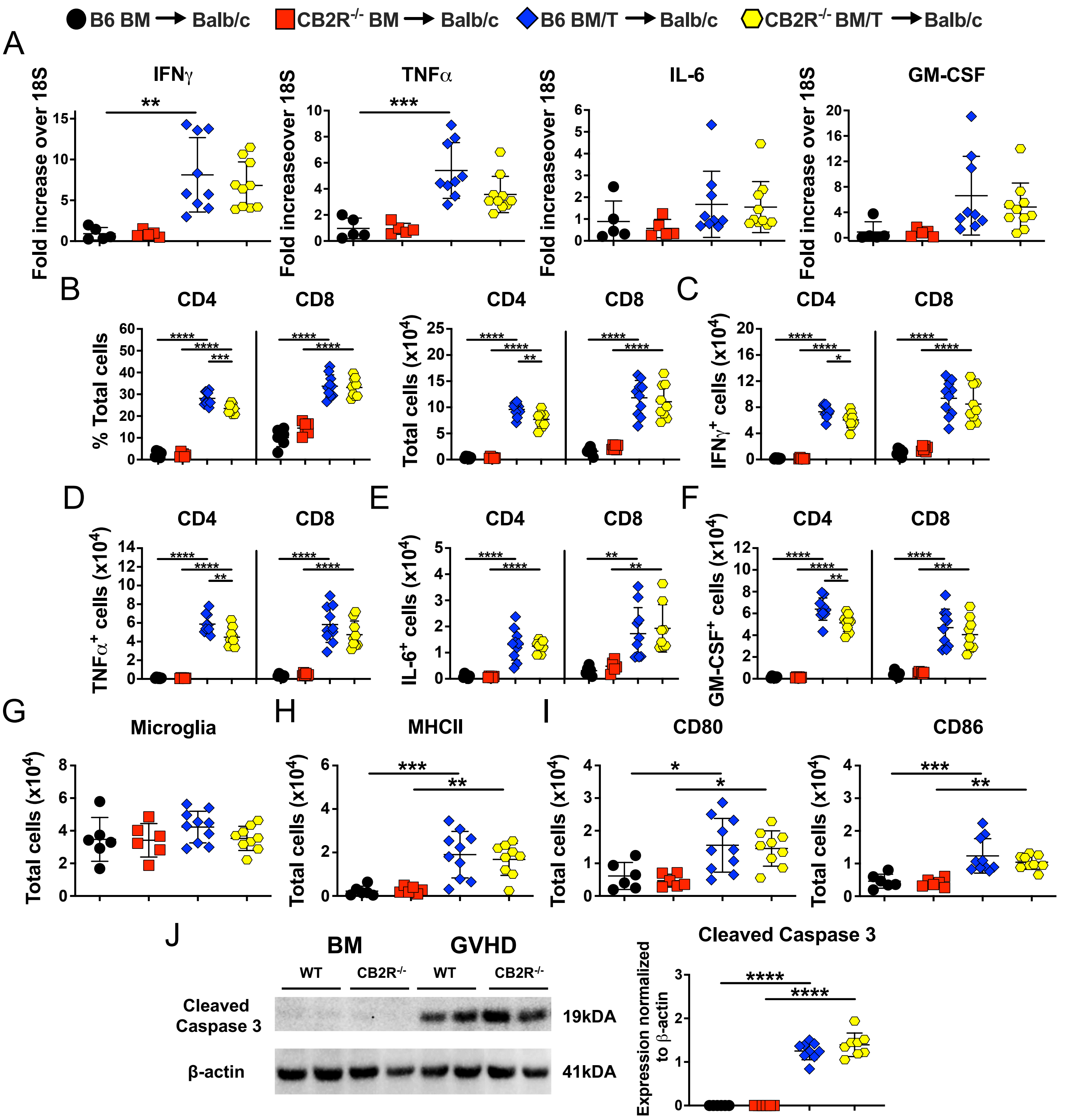

### Supplemental Figure 4

A

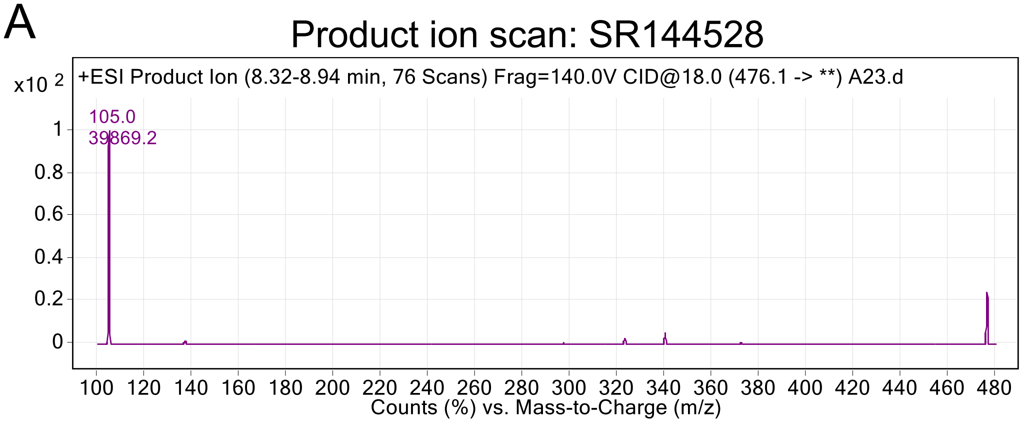


B

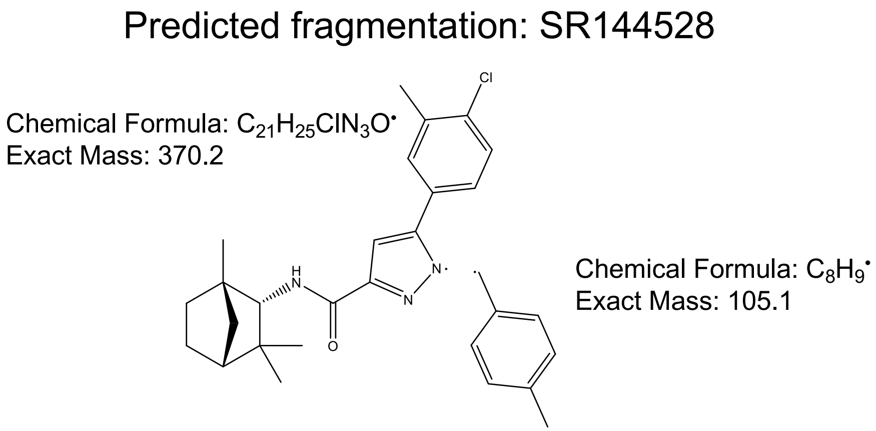


C

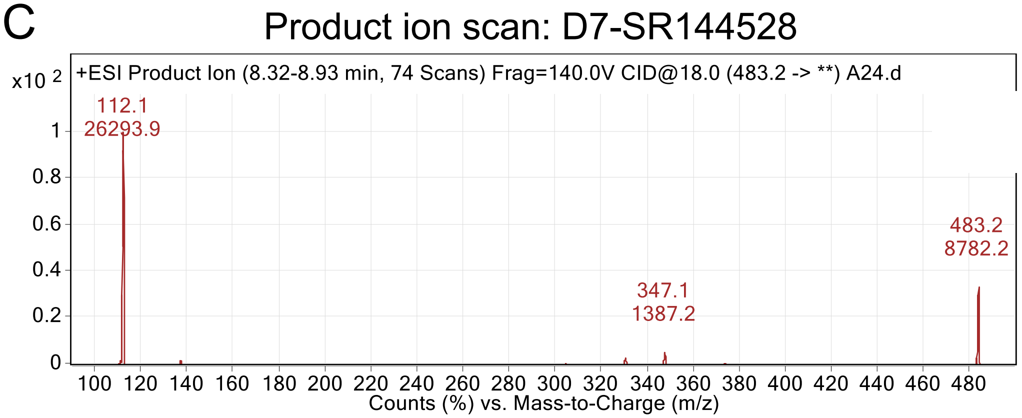


D

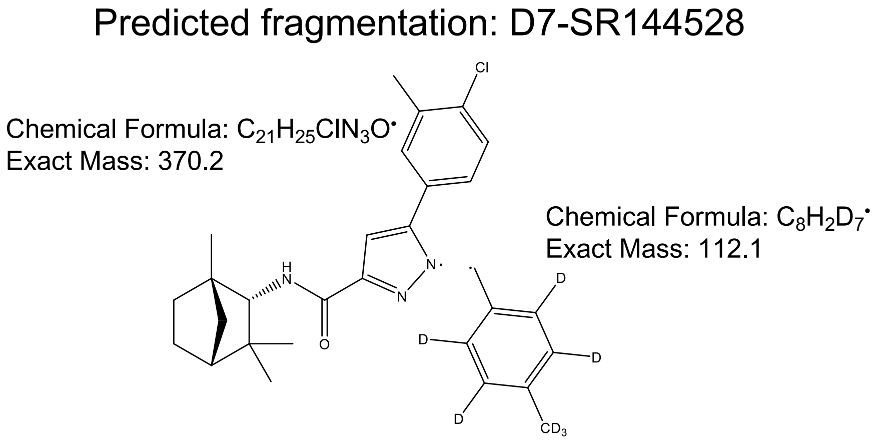


E

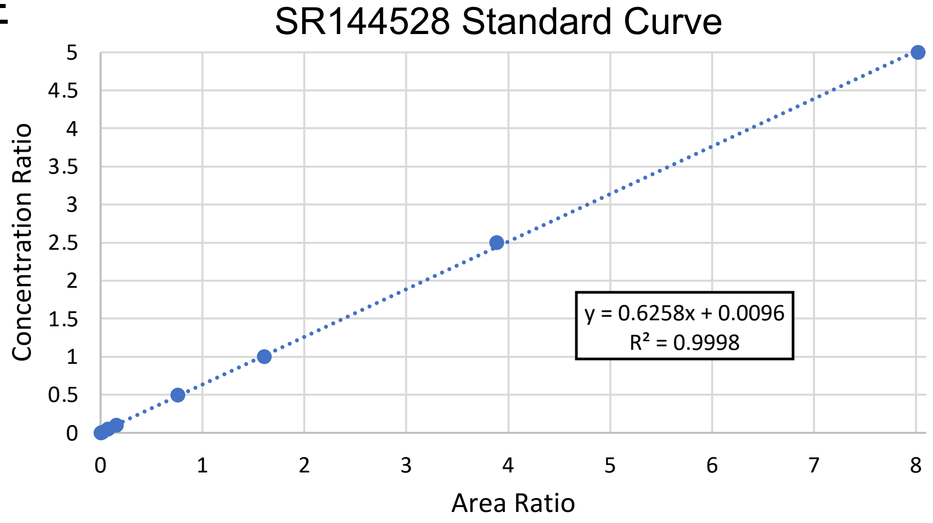

### Supplemental Figure 5

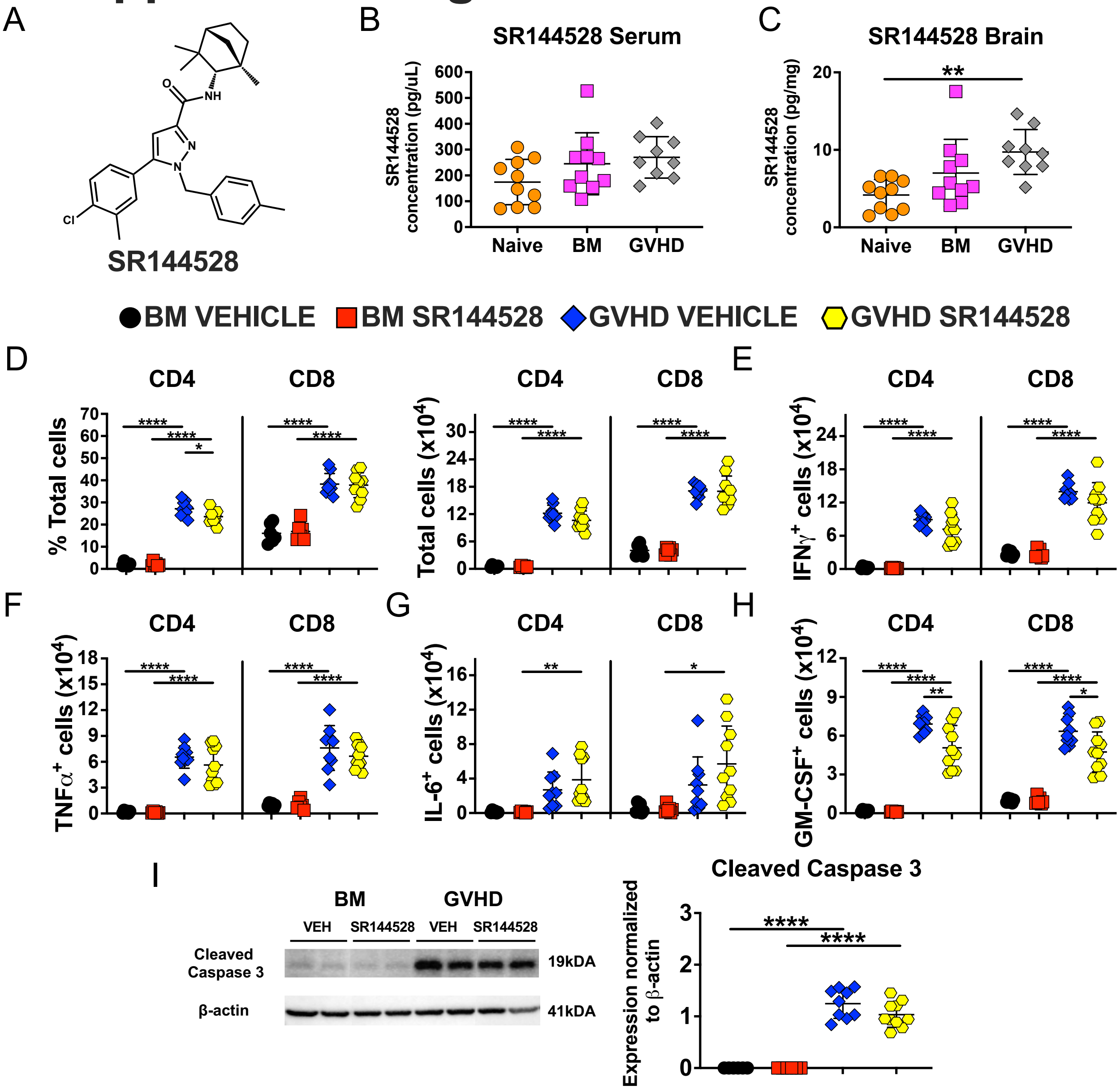

### Supplemental Figure 6

A

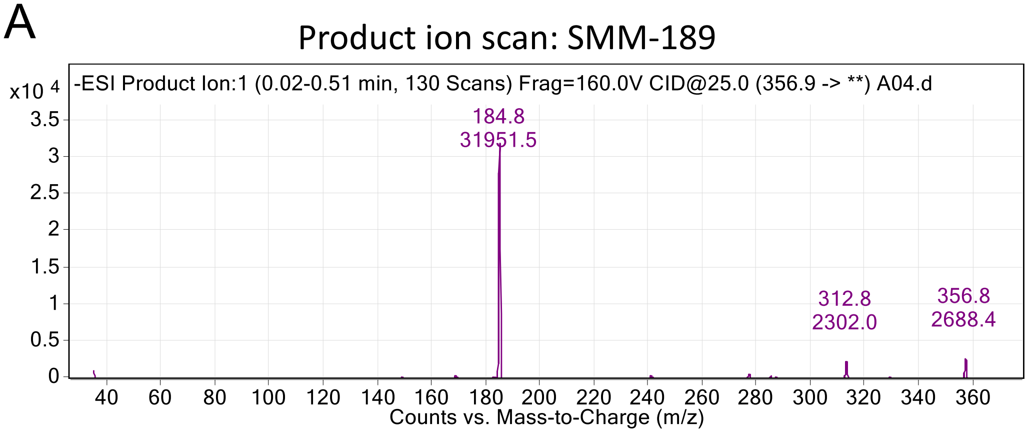


B

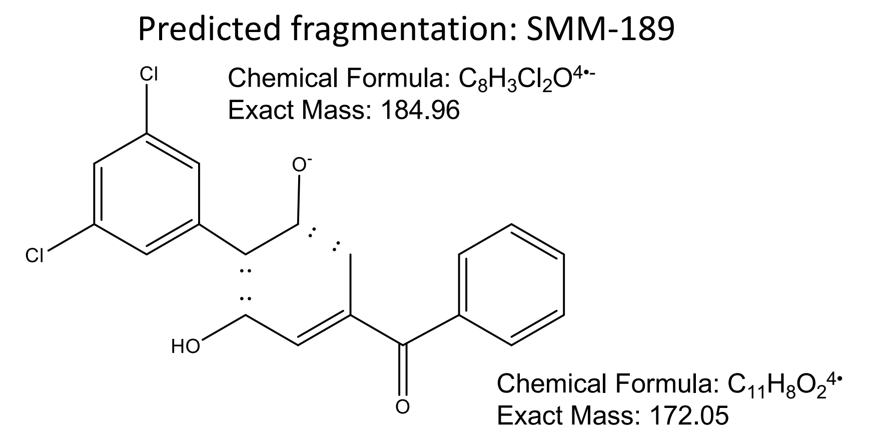


C

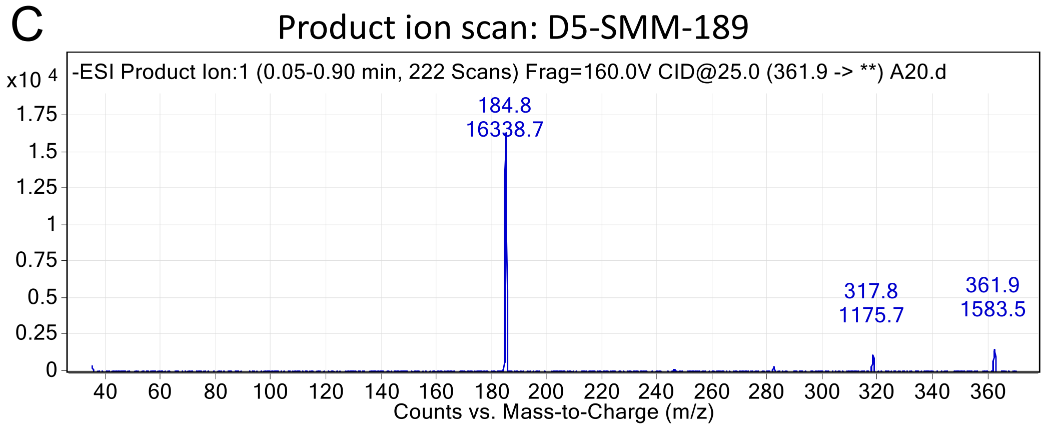


D

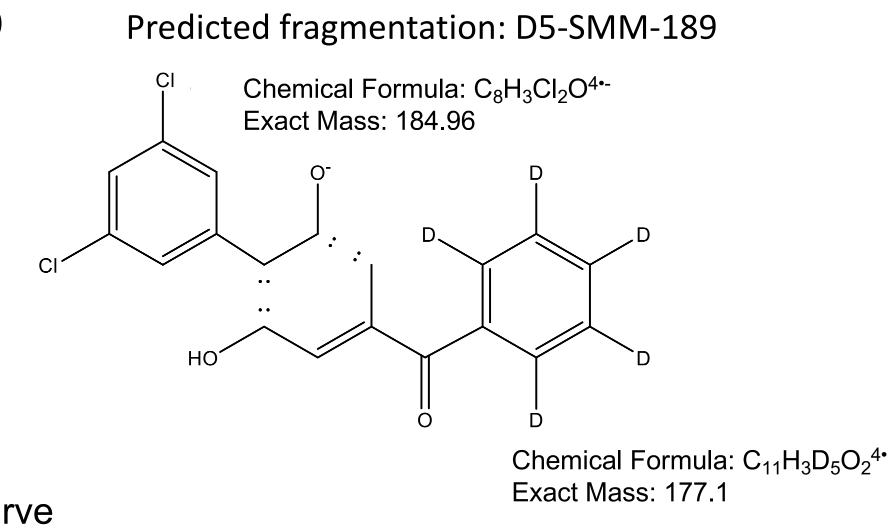


E

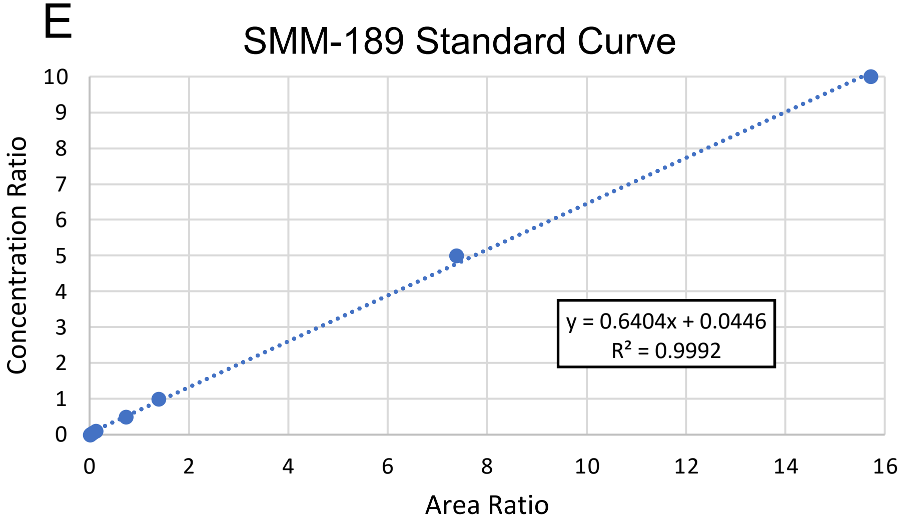

#### SUPPLEMENTAL FIGURE LEGENDS

Supplemental Figure 1**: Donor T cells from the brains of GVHD mice are polyfunctional and produce multiple inflammatory cytokines.** Lethally irradiated (900 cGy) Balb/c mice were transplanted with B6 BM alone (5 x $10^6$) or B6 BM and spleen cells (adjusted to yield αβ T cell dose of 0.75 x $10^6$). Animals were euthanized 14 days post transplantation. (A). IFN-γ, TNF-α, IL-6, and GM-CSF mRNA expression is depicted in the brain. (B). The percentage and absolute number of donor derived $CD4^+$ and $CD8^+$ T cells. (C, D). The absolute number of $CD4^+$ (panel C) and $CD8^+$ (panel D) T cells that produced IFN-γ, TNF-α, IL-6 or GM-CSF. Results are from two experiments. Statistics: *p<0.05; **p<0.01.

Supplemental Figure 2: **Transcriptional profile of T cells and macrophages in the brain of GVHD mice.** Lethally irradiated (1100 cGy) B6 mice were transplanted with B10.BR BM (5 x $10^6$) alone (BM) or together with B10.BR spleen cells (adjusted to yield an αβ T cell dose of 5 x $10^6$) (GVHD). Single live cells from pooled brains (n=5/group) were sorted 14 days post transplantation. (A). Absolute number and percentage of cells in each cluster from BM and GVHD mice that underwent single cell RNA sequence analysis. Six clusters representing 7% of cells in the BM scRNAseq dataset and 9.5% of cells in the GVHD dataset were excluded from the analysis. The transcriptional signature of these cells identified them as astrocytes, red blood cells, neurons, NK cells, an unclassifiable subset that had markers of T cells and microglia, and a minor macrophage population. (B). Volcano plot showing genes in aggregated T cell clusters that were over expressed in BM versus GVHD mice. Cutoff parameters were |log2(fold change)| > 1.0 and $p_{adjusted}$ <0.0001. (C). Volcano plot showing genes in macrophage cluster that were over expressed in BM versus GVHD mice. Cutoff parameters were |log2(fold change)| > 1.0 and $p_{adjusted}$ <0.0001.

Supplemental Figure 3: **Donor-derived CB2R expression has only modest effects on reducing inflammation in the brain during GVHD.** Lethally irradiated (900 cGy) Balb/c mice were transplanted with B6 or CB2R$^{-/-}$ BM alone or together with B6 or CB2R$^{-/-}$ spleen cells (adjusted to yield an αβ T cell dose of 0.75 x $10^6$ T cells). Animals were euthanized 14 days post transplantation. (A). IFN-γ, TNF-α, IL-6, and GM-CSF mRNA expression in the whole brain. (B). The percentage and absolute number of donor derived CD4$^+$ and CD8$^+$ T cells. (C-F). The absolute number of donor-derived CD4$^+$ and CD8$^+$ T cells that produced IFN-γ, TNF-α, IL-6 or GM-CSF 14 days post transplantation. (G). The absolute number of microglial cells. (H, I). The absolute number of microglia expressing MHC class II, CD80 and CD86. (J). Representative western blot images and scatterplots depicting normalized expression of cleaved caspase 3 in the brain. Results are from two-three experiments per panel. Statistics: *p<0.05, **p<0.01, ***p<0.001, ****p<0.0001.

Supplemental Figure 4: **Mass spectral characteristics of SR144528.** (A). Product ion scan of standard SR144528. The compound was injected directly into the mass spectrometer and analyzed using the parameters outlined in Methods. (B). The most abundant daughter ion had an m/z of 105, which is consistent with the fragmentation pattern shown in panel B. (C, D). The daughter ion with greatest abundance for the deuterium labeled SR144528 had an m/z ratio of 112.1 (panel C), consistent with a similar fragmentation pattern and with the presence of seven deuterium atoms on the benzoyl moiety (panel D). These daughter ions were monitored using selective ion monitoring. (E). A representative standard curve constructed using SR144528 concentrations of 1, 5, 10, 50, 100, 250, 500 ng/mL and 100 ng/ml SR144528-D7. SR144528 concentration/100 ng/ml is plotted on the y axis and the area of the peaks for SR144528/SR144528-D7 is plotted on the x axis.

Supplemental Figure 5: **Pharmacological administration of a brain impermeable CB2R antagonist has no significant effect on mitigating CNS inflammation.** (A). Chemical structure of SR144528. (B).

Serum levels (pg/µl) of SR144528 in naïve and lethally irradiated (900 cGy) Balb/c mice transplanted with B6 BM alone or together with B6 spleen cells (adjusted to yield an αβ T cell dose of 0.75 x $10^6$). (C). Concentration (pg/mg) of SR144528 in the brain. (D-I). Lethally irradiated Balb/c recipients were transplanted with B6 BM (5 x $10^6$) and spleen cells (adjusted to yield an αβ T cell dose of 0.75 x $10^6$ cells). Animals were then treated with SR 144528 (3 mg/kg) or a vehicle control for 14 days beginning on day 0. Balb/c mice transplanted with B6 BM alone and then treated with either vehicle or a SR144528 served as controls. Animals were euthanized 14 days post transplantation. (D). The percentage and absolute number of donor derived $CD4^+$ and $CD8^+$ T cells in the brain. (E-H). The absolute number of $CD4^+$ and $CD8^+$ T cells that produced IFN-γ, TNF-α, IL-6 or GM-CSF. (I). Representative western blot images and scatterplots depicting normalized expression of cleaved caspase 3 in the brain. Results are from two experiments. Statistics: *p<0.05, **p<0.01, ***p<0.001, ****p<0.0001.

Supplemental Figure 6: **Mass spectral characteristics of SMM-189.** (A). Product ion scan of standard SMM-189. The compound was injected directly into the mass spectrometer and analyzed using the parameters outlined in the methods. (B). The most abundant daughter ion had an m/z of 184.8 which is consistent with the fragmentation pattern shown in panel B. (C, D). The same daughter ion was present in greatest abundance for the deuterium labeled SMM-189 (panel C) consistent with a similar fragmentation pattern (panel D). This daughter ions were monitored using selective ion monitoring. (E). A representative standard curve constructed using SMM-189 concentrations of 10, 50, 100, 500, 1000, 5000, 10000 ng/mL and 1000 ng/ml SMM-189-D5. SMM-189 concentration/1000 ng/ml is plotted on the y axis and the area of the peaks for SMM-189/SMM-189-D5 is plotted on the x axis.

### SUPPLEMENTAL TABLE 1. Differentially Regulated Genes in GVHD versus BM T cells

| Gene | avg_log2FC | p_val | p_val_adj |
|---|---|---|---|
| Ly6c2 | 3.208077 | 9.28E-36 | 3E-31 |
| Ly6a | 2.567448 | 1.56E-57 | 5.03E-53 |
| Ifitm3 | 1.924385 | 5.08E-19 | 1.64E-14 |
| Trbv19 | 1.409484 | 1.08E-05 | 0.350053 |
| Ifitm2 | 1.341234 | 1.96E-13 | 6.33E-09 |
| AU020206 | 1.330486 | 2.84E-32 | 9.15E-28 |
| Stat1 | 1.279438 | 1.8E-28 | 5.81E-24 |
| Plac8 | 1.256192 | 1.59E-19 | 5.13E-15 |
| Trbv13-2 | 1.253727 | 3.43E-06 | 0.110689 |
| Icos | 1.224936 | 3.6E-19 | 1.16E-14 |
| Cd4 | 1.201073 | 3.75E-12 | 1.21E-07 |
| Bst2 | 1.127778 | 1.53E-18 | 4.96E-14 |
| Ltb | 1.111168 | 1.95E-24 | 6.31E-20 |
| Cxcr6 | 1.092178 | 2.18E-20 | 7.05E-16 |
| Il12rb2 | 1.090976 | 1.64E-17 | 5.3E-13 |
| Inpp4b | 1.069197 | 1.82E-20 | 5.88E-16 |
| Ifi27l2a | 1.068756 | 3.19E-10 | 1.03E-05 |
| Rora | 1.054915 | 2.85E-20 | 9.19E-16 |
| Il18r1 | 1.043753 | 1.62E-17 | 5.24E-13 |
| Isg20 | 1.011475 | 6.26E-22 | 2.02E-17 |
| Itga1 | 1.005883 | 3.02E-16 | 9.74E-12 |
| Jaml | 0.993767 | 1.93E-14 | 6.22E-10 |
| Tnfrsf4 | 0.990977 | 1.02E-07 | 0.003279 |
| Gbp6 | 0.979603 | 1.6E-19 | 5.16E-15 |
| Igtp | 0.970512 | 9.52E-19 | 3.07E-14 |
| Hif1a | 0.96805 | 1.34E-13 | 4.32E-09 |
| Zbp1 | 0.929497 | 2.93E-15 | 9.45E-11 |
| Efhd2 | 0.927916 | 4.1E-21 | 1.32E-16 |
| Slco3a1 | 0.918292 | 3.18E-15 | 1.03E-10 |
| Tnf | 0.898248 | 4.68E-09 | 0.000151 |
| Ms4a6b | 0.894583 | 1.81E-19 | 5.85E-15 |
| St6galnac3 | 0.885649 | 4.38E-10 | 1.41E-05 |
| Slfn1 | 0.880593 | 3.4E-15 | 1.1E-10 |
| AA467197 | 0.865882 | 3.36E-15 | 1.08E-10 |
| Mt1 | 0.861703 | 8.09E-09 | 0.000261 |
| Ifng | 0.859512 | 0.020298 | 1 |
| Socs1 | 0.830511 | 4.28E-12 | 1.38E-07 |
| Irf7 | 0.828664 | 1.18E-10 | 3.82E-06 |
| Ctla4 | 0.828294 | 2.22E-10 | 7.18E-06 |
| Il7r | 0.816251 | 2.12E-08 | 0.000685 |
| Rtp4 | 0.814762 | 8.43E-12 | 2.72E-07 |
| Junb | 0.812862 | 1.97E-08 | 0.000637 |
| AW112010 | 0.807195 | 8.17E-21 | 2.64E-16 |
| Srgn | 0.805845 | 1.06E-18 | 3.42E-14 |

| | | | |
|---|---|---|---|
| Satb1 | 0.802387 | 1.29E-16 | 4.18E-12 |
| mt-Atp8 | 0.800684 | 2.64E-17 | 8.52E-13 |
| Ifi47 | 0.797437 | 1.49E-13 | 4.82E-09 |
| Lilr4b | 0.797183 | 9.31E-13 | 3E-08 |
| Shisa5 | 0.782012 | 5.17E-18 | 1.67E-13 |
| Gstp3 | 0.781033 | 1.08E-13 | 3.48E-09 |
| Glrx | 0.779712 | 2.25E-09 | 7.26E-05 |
| Cd40lg | 0.777519 | 5.39E-09 | 0.000174 |
| Gramd3 | 0.768432 | 3.48E-12 | 1.12E-07 |
| Cd274 | 0.765578 | 2.36E-10 | 7.63E-06 |
| Aldoa | 0.761132 | 1.31E-14 | 4.23E-10 |
| Gm36723 | 0.76083 | 1.37E-10 | 4.44E-06 |
| Ifit3 | 0.759321 | 9.23E-09 | 0.000298 |
| Arhgap26 | 0.757268 | 2.54E-12 | 8.2E-08 |
| Bcl2a1b | 0.755196 | 2.72E-12 | 8.78E-08 |
| Lilrb4a | 0.735931 | 5.65E-11 | 1.82E-06 |
| Rnf213 | 0.733787 | 6.16E-10 | 1.99E-05 |
| Gbp9 | 0.723811 | 2.23E-10 | 7.19E-06 |
| Runx2 | 0.721161 | 1.07E-11 | 3.44E-07 |
| Rgs1 | 0.713811 | 6.49E-08 | 0.002096 |
| Cd69 | 0.712157 | 2.63E-06 | 0.084853 |
| Rgs10 | 0.706416 | 3.88E-10 | 1.25E-05 |
| Emb | 0.706167 | 7.31E-11 | 2.36E-06 |
| Irgm1 | 0.701917 | 8.53E-10 | 2.75E-05 |
| 2410006H16 | 0.697229 | 3.63E-11 | 1.17E-06 |
| Gzmb | 0.67888 | 0.278156 | 1 |
| Isg15 | 0.67732 | 1.72E-09 | 5.55E-05 |
| Tnfsf8 | 0.67702 | 4.09E-06 | 0.131913 |
| Ifi213 | 0.669539 | 9.01E-08 | 0.002908 |
| Cd226 | 0.666688 | 4.6E-09 | 0.000149 |
| Ms4a4c | 0.663378 | 7.55E-08 | 0.002438 |
| Apol7e | 0.651722 | 2.5E-10 | 8.07E-06 |
| Ly6e | 0.650538 | 4.64E-19 | 1.5E-14 |
| Gbp10 | 0.644742 | 3.38E-09 | 0.000109 |
| Stat3 | 0.643054 | 2.3E-11 | 7.43E-07 |
| Ddit4 | 0.641827 | 5.51E-08 | 0.00178 |
| Prkca | 0.635565 | 8.68E-08 | 0.002804 |
| Cd6 | 0.63289 | 5.57E-08 | 0.001798 |
| Trbv13-3 | 0.632491 | 0.010368 | 1 |
| Pde3b | 0.628523 | 1.46E-08 | 0.00047 |
| Xaf1 | 0.614219 | 2.65E-08 | 0.000857 |
| Csf1 | 0.613714 | 4.88E-05 | 1 |
| Nedd9 | 0.61152 | 9.01E-12 | 2.91E-07 |
| Ifit1 | 0.610595 | 1.32E-05 | 0.425389 |
| Prdx6 | 0.608777 | 2.12E-12 | 6.85E-08 |
| Id2 | 0.603822 | 7.5E-11 | 2.42E-06 |
| Parp9 | 0.603706 | 2.74E-10 | 8.86E-06 |
| Ccl4 | 0.598105 | 0.98797 | 1 |

| | | | |
|---|---|---|---|
| Sdf4 | 0.594783 | 1.92E-07 | 0.006211 |
| Cxcr3 | 0.593859 | 1.28E-08 | 0.000412 |
| Lpxn | 0.586472 | 7.49E-11 | 2.42E-06 |
| Gm10260 | 0.585503 | 1.7E-13 | 5.48E-09 |
| Nrgn | 0.585384 | 0.001744 | 1 |
| H2-Q1 | 0.580907 | 8.48E-12 | 2.74E-07 |
| Bcl2a1d | 0.579047 | 1.98E-08 | 0.000639 |
| Slfn2 | 0.576913 | 2.05E-08 | 0.00066 |
| Themis | 0.575268 | 2E-08 | 0.000646 |
| Dnajc15 | 0.574685 | 9.05E-10 | 2.92E-05 |
| Ifngr1 | 0.56603 | 1.01E-08 | 0.000326 |
| Gbp4 | 0.564895 | 4.75E-08 | 0.001534 |
| Rpl12 | 0.55992 | 1.1E-14 | 3.55E-10 |
| Rps12 | 0.553 | 5.22E-20 | 1.68E-15 |
| Zfp36l2 | 0.547805 | 1.12E-06 | 0.036233 |
| Clint1 | 0.547117 | 7.08E-08 | 0.002284 |
| Ccr8 | 0.546981 | 2E-05 | 0.645759 |
| Fth1 | 0.546322 | 7.24E-16 | 2.34E-11 |
| Rps2 | 0.546093 | 3.2E-15 | 1.03E-10 |
| Btg1 | 0.544147 | 1.19E-11 | 3.85E-07 |
| Peli1 | 0.541398 | 5.85E-08 | 0.001887 |
| Gm19585 | 0.540119 | 6.14E-08 | 0.001982 |
| Ehd1 | 0.539922 | 5.78E-07 | 0.018672 |
| Crlf2 | 0.537648 | 3.02E-08 | 0.000975 |
| Pag1 | 0.535883 | 2.07E-06 | 0.066693 |
| Krt83 | 0.535579 | 2.28E-06 | 0.073625 |
| Gas5 | 0.531706 | 1.65E-09 | 5.31E-05 |
| 4933406I18F | 0.52867 | 5.04E-08 | 0.001627 |
| H2-Q2 | 0.524532 | 1.48E-10 | 4.77E-06 |
| Nfkbia | 0.522275 | 0.000182 | 1 |
| Tgtp2 | 0.518373 | 2.85E-08 | 0.000922 |
| Gbp7 | 0.516668 | 7.2E-07 | 0.023259 |
| Pdcd1 | 0.512739 | 0.002166 | 1 |
| Rps14 | 0.512183 | 3.75E-23 | 1.21E-18 |
| Batf | 0.511283 | 2.36E-05 | 0.762662 |
| Gldc | 0.509709 | 2.13E-06 | 0.068712 |
| Rps15a | 0.503645 | 7.58E-20 | 2.45E-15 |
| Rpl27a | 0.500507 | 7.87E-21 | 2.54E-16 |
| Dusp2 | 0.499367 | 2.95E-07 | 0.009517 |
| Litaf | 0.498504 | 3.44E-07 | 0.011116 |
| Jun | 0.498424 | 1.4E-07 | 0.004534 |
| Gpr18 | 0.497137 | 1.2E-07 | 0.003884 |
| Tnfsf10 | 0.497081 | 1.17E-07 | 0.00379 |
| Pkm | 0.495373 | 1.33E-09 | 4.29E-05 |
| Ctsw | 0.494299 | 9.44E-08 | 0.003048 |
| Ifi208 | 0.493742 | 5.53E-07 | 0.017849 |
| Oas3 | 0.493509 | 9.58E-06 | 0.309314 |
| Smyd3 | 0.487761 | 3.99E-07 | 0.012872 |

| | | | |
|---|---|---|---|
| Mapkapk3 | 0.485675 | 4.74E-06 | 0.153192 |
| Rps23 | 0.483763 | 4.52E-20 | 1.46E-15 |
| Tab2 | 0.483045 | 2.23E-05 | 0.719359 |
| Epsti1 | 0.480123 | 3.9E-07 | 0.012602 |
| Gm15283 | 0.479388 | 3.22E-06 | 0.103967 |
| Rps24 | 0.478088 | 2.93E-21 | 9.46E-17 |
| Lrp10 | 0.477849 | 2.33E-07 | 0.007517 |
| Gbp2 | 0.477726 | 1.27E-09 | 4.09E-05 |
| Relb | 0.474035 | 1.92E-07 | 0.006193 |
| Rpl9 | 0.470242 | 1.28E-19 | 4.13E-15 |
| Traf1 | 0.469685 | 1.81E-06 | 0.058381 |
| Podnl1 | 0.469114 | 0.000371 | 1 |
| Pgk1 | 0.467341 | 4.22E-07 | 0.013614 |
| Gbp8 | 0.46325 | 7.05E-06 | 0.22774 |
| 1600014C10I | 0.462138 | 1.39E-05 | 0.447176 |
| Grina | 0.460714 | 8.64E-06 | 0.278995 |
| Tpi1 | 0.459356 | 0.000336 | 1 |
| Psme2 | 0.458475 | 9.2E-09 | 0.000297 |
| Rpl23 | 0.458293 | 2.23E-19 | 7.21E-15 |
| Anxa1 | 0.457188 | 3.68E-10 | 1.19E-05 |
| Rpl24 | 0.456554 | 2.65E-16 | 8.54E-12 |
| Chmp4b | 0.452948 | 2.49E-07 | 0.008052 |
| Serpina3f | 0.450512 | 0.000607 | 1 |
| H2afy | 0.446252 | 8.56E-07 | 0.02764 |
| Lef1 | 0.444917 | 8.62E-05 | 1 |
| Adam19 | 0.444002 | 6.61E-05 | 1 |
| Ddx24 | 0.4438 | 1.93E-06 | 0.062249 |
| Pdcd1lg2 | 0.441419 | 1.54E-08 | 0.000498 |
| Cd3g | 0.44106 | 7.12E-08 | 0.002297 |
| Npm1 | 0.440044 | 1.75E-09 | 5.65E-05 |
| Ubash3a | 0.438538 | 7.67E-10 | 2.48E-05 |
| Gm8909 | 0.437761 | 3.94E-08 | 0.00127 |
| Sipa1l1 | 0.437462 | 6.81E-06 | 0.219722 |
| Rpl7a | 0.436804 | 2.79E-13 | 8.99E-09 |
| Rplp1 | 0.435955 | 4.23E-15 | 1.37E-10 |
| Tapbp | 0.435011 | 5.25E-07 | 0.01695 |
| Rps20 | 0.433794 | 3.77E-11 | 1.22E-06 |
| Klk8 | 0.433719 | 4.01E-05 | 1 |
| Il2ra | 0.433488 | 3.65E-07 | 0.011779 |
| Mif | 0.432106 | 4.31E-06 | 0.13921 |
| Gramd1a | 0.429697 | 2.45E-06 | 0.079248 |
| Grcc10 | 0.429253 | 4.06E-08 | 0.00131 |
| Uhrf2 | 0.427688 | 7.82E-05 | 1 |
| Rpl17 | 0.427376 | 1.18E-15 | 3.8E-11 |
| Psmb9 | 0.426984 | 3.65E-09 | 0.000118 |
| Igflr1 | 0.425969 | 1.85E-06 | 0.059703 |
| Il2rg | 0.425083 | 1.5E-08 | 0.000485 |
| Cd5 | 0.424512 | 0.000127 | 1 |

| | | | |
|---|---|---|---|
| Oas1a | 0.423764 | 4.59E-05 | 1 |
| Hopx | 0.423646 | 0.000107 | 1 |
| Fam102a | 0.423202 | 4.26E-05 | 1 |
| Samhd1 | 0.422202 | 9.59E-05 | 1 |
| Rps9 | 0.421548 | 1.03E-15 | 3.31E-11 |
| Rpl13 | 0.421074 | 1.24E-16 | 4.01E-12 |
| Trac | 0.420427 | 2.89E-07 | 0.00934 |
| Rpl15 | 0.419772 | 1.09E-10 | 3.53E-06 |
| Lncpint | 0.418571 | 9.77E-05 | 1 |
| Oasl2 | 0.416691 | 4.24E-05 | 1 |
| Herc6 | 0.416022 | 0.000207 | 1 |
| Gm12216 | 0.415454 | 6.36E-06 | 0.205274 |
| Irgm2 | 0.414801 | 1.21E-06 | 0.038969 |
| Orai1 | 0.413016 | 8.17E-07 | 0.026392 |
| Rpl18 | 0.411463 | 2.89E-15 | 9.34E-11 |
| Crot | 0.411108 | 8.48E-06 | 0.273664 |
| Nek7 | 0.411012 | 0.000999 | 1 |
| Rpl29 | 0.409695 | 2.46E-14 | 7.94E-10 |
| Ipcef1 | 0.409146 | 3.64E-05 | 1 |
| H1f0 | 0.408731 | 9.23E-06 | 0.297984 |
| Tap1 | 0.408202 | 2.41E-06 | 0.077785 |
| Lag3 | 0.405971 | 0.085 | 1 |
| Stat2 | 0.403813 | 0.000762 | 1 |
| Rpl3 | 0.403622 | 2.98E-12 | 9.63E-08 |
| Tecpr1 | 0.40337 | 0.0005 | 1 |
| Irf1 | 0.401808 | 0.0002 | 1 |
| Lgals3bp | 0.401492 | 1.14E-05 | 0.367634 |
| Rpl11 | 0.400703 | 6.13E-14 | 1.98E-09 |
| Map2k2 | 0.397909 | 9.4E-07 | 0.030352 |
| Cd86 | 0.397713 | 0.000158 | 1 |
| Rpl26 | 0.397453 | 2.35E-13 | 7.59E-09 |
| Gpr55 | 0.39682 | 1.87E-06 | 0.060517 |
| Ppp3cc | 0.395929 | 0.000109 | 1 |
| Rbpj | 0.395547 | 0.00035 | 1 |
| Atf6 | 0.394983 | 0.00029 | 1 |
| Gpr183 | 0.394678 | 0.000384 | 1 |
| Gm7030 | 0.393947 | 4.45E-06 | 0.143668 |
| Slamf1 | 0.393242 | 2.53E-05 | 0.816591 |
| Ptpn18 | 0.391656 | 2.24E-08 | 0.000723 |
| Rpl18a | 0.390988 | 3.7E-15 | 1.19E-10 |
| Ly75 | 0.390632 | 4.54E-06 | 0.146695 |
| Mrpl52 | 0.388072 | 6.62E-07 | 0.021368 |
| Fgl2 | 0.386055 | 0.000234 | 1 |
| Rps13 | 0.385451 | 6.51E-16 | 2.1E-11 |
| Ogt | 0.384269 | 0.001286 | 1 |
| Rabac1 | 0.384093 | 8.99E-06 | 0.290225 |
| Ddx60 | 0.38349 | 1.5E-06 | 0.04835 |
| Gng2 | 0.383315 | 1.25E-05 | 0.403631 |

| | | | |
|---|---|---|---|
| Tigit | 0.382022 | 0.000133 | 1 |
| Celf2 | 0.381267 | 0.000104 | 1 |
| Tnfrsf18 | 0.380983 | 0.007871 | 1 |
| Rpl14 | 0.380512 | 4E-11 | 1.29E-06 |
| Arl15 | 0.379447 | 0.010097 | 1 |
| Arf5 | 0.378785 | 4.11E-07 | 0.013283 |
| Coro2a | 0.377987 | 6.43E-05 | 1 |
| Atxn1 | 0.377515 | 0.000328 | 1 |
| Vps54 | 0.377152 | 0.000196 | 1 |
| Trim30a | 0.3748 | 0.000792 | 1 |
| Itk | 0.374442 | 2.04E-06 | 0.065744 |
| Rps21 | 0.374011 | 6.27E-09 | 0.000202 |
| Ifit3b | 0.373748 | 9.34E-05 | 1 |
| Rpl30 | 0.373449 | 2.15E-12 | 6.93E-08 |
| Rps10 | 0.37338 | 1.92E-16 | 6.18E-12 |
| Itpr1 | 0.372986 | 0.000346 | 1 |
| Mdfic | 0.372528 | 0.000323 | 1 |
| Il18rap | 0.372275 | 4.21E-06 | 0.135841 |
| Nme2 | 0.371639 | 3.05E-06 | 0.098537 |
| Rpl6 | 0.371122 | 8.25E-12 | 2.66E-07 |
| Fndc3a | 0.370745 | 0.000137 | 1 |
| Rpl10 | 0.369459 | 3.42E-10 | 1.1E-05 |
| Ttr | 0.369371 | 1.24E-08 | 0.000399 |
| Marc2 | 0.369032 | 1.91E-05 | 0.616147 |
| Dtx3l | 0.36868 | 0.0004 | 1 |
| Jak3 | 0.368317 | 0.00024 | 1 |
| H2-Q4 | 0.367884 | 4.28E-08 | 0.001381 |
| Arid5a | 0.366534 | 0.00013 | 1 |
| Rplp2 | 0.36617 | 2.04E-12 | 6.58E-08 |
| Tap2 | 0.365908 | 1.18E-06 | 0.038252 |
| Nfkbil1 | 0.365464 | 1.33E-05 | 0.429749 |
| Rack1 | 0.360923 | 1.01E-09 | 3.25E-05 |
| Rpl34 | 0.360223 | 3.23E-13 | 1.04E-08 |
| Maf | 0.358293 | 0.009774 | 1 |
| Ypel3 | 0.357456 | 0.000113 | 1 |
| Psma5 | 0.357047 | 2.68E-05 | 0.865931 |
| Rpsa | 0.356363 | 1.19E-11 | 3.83E-07 |
| Rel | 0.354154 | 0.005332 | 1 |
| Rpl27 | 0.35371 | 9.64E-13 | 3.11E-08 |
| Kxd1 | 0.353654 | 0.00069 | 1 |
| Rgs16 | 0.352893 | 0.118375 | 1 |
| Npc2 | 0.352104 | 4.95E-05 | 1 |
| Itm2c | 0.351662 | 0.00027 | 1 |
| Sdcbp2 | 0.351092 | 3.56E-05 | 1 |
| Gpr171 | 0.349853 | 3.54E-05 | 1 |
| Ms4a4b | 0.348955 | 6.38E-05 | 1 |
| Flt3l | 0.348197 | 0.000723 | 1 |
| Nmi | 0.346877 | 0.001085 | 1 |

| | | | |
|---|---|---|---|
| Sdhaf1 | 0.345682 | 7.04E-05 | 1 |
| Rps5 | 0.345571 | 3.68E-12 | 1.19E-07 |
| Mrps34 | 0.345241 | 3.04E-05 | 0.982688 |
| Rpl32 | 0.343427 | 4.21E-11 | 1.36E-06 |
| Rps7 | 0.342839 | 3.4E-12 | 1.1E-07 |
| Gstp1 | 0.342792 | 0.000608 | 1 |
| Cish | 0.342781 | 0.001084 | 1 |
| Rpl8 | 0.342634 | 5.3E-10 | 1.71E-05 |
| Ifi35 | 0.341955 | 0.000176 | 1 |
| Cd96 | 0.340954 | 0.00016 | 1 |
| Npepl1 | 0.340939 | 6.4E-05 | 1 |
| Egr1 | 0.340747 | 0.001061 | 1 |
| Tpt1 | 0.340422 | 4.37E-15 | 1.41E-10 |
| H2-Q10 | 0.339813 | 5.18E-05 | 1 |
| Rpl19 | 0.339418 | 1.58E-12 | 5.11E-08 |
| Notch1 | 0.339393 | 0.001886 | 1 |
| B2m | 0.339328 | 4.78E-13 | 1.54E-08 |
| Plscr1 | 0.338506 | 0.014701 | 1 |
| Rpl5 | 0.337341 | 1.1E-06 | 0.035662 |
| Cd47 | 0.337144 | 9.96E-06 | 0.321584 |
| Ubac2 | 0.336715 | 7.5E-05 | 1 |
| Ndfip1 | 0.3361 | 0.001249 | 1 |
| Eef1b2 | 0.335657 | 1.54E-07 | 0.004957 |
| Pfdn5 | 0.335461 | 3.75E-07 | 0.01211 |
| Psmb8 | 0.334769 | 6.3E-09 | 0.000204 |
| Gimap3 | 0.334759 | 1.04E-05 | 0.336065 |
| Apobec3 | 0.334353 | 2.53E-05 | 0.815251 |
| Nrp1 | 0.333263 | 0.026466 | 1 |
| Cd8b1 | 0.331462 | 0.101471 | 1 |
| Cytip | 0.330319 | 0.000589 | 1 |
| Gls | 0.329876 | 0.001793 | 1 |
| Slfn8 | 0.328395 | 5.05E-06 | 0.162962 |
| Fxyd5 | 0.327838 | 2.53E-08 | 0.000818 |
| Glmn | 0.327575 | 0.00021 | 1 |
| Adgrg5 | 0.327341 | 2.55E-06 | 0.082268 |
| Rpl31 | 0.326294 | 3.26E-05 | 1 |
| Rpl21 | 0.326013 | 6.8E-10 | 2.19E-05 |
| Rps4x | 0.325929 | 1.11E-11 | 3.59E-07 |
| Tent5c | 0.325447 | 4.02E-05 | 1 |
| St6galnac4 | 0.325064 | 5.27E-05 | 1 |
| Eif3e | 0.324344 | 1.24E-05 | 0.401254 |
| B3gnt2 | 0.324113 | 0.000765 | 1 |
| Snhg12 | 0.324017 | 0.000177 | 1 |
| Rps25 | 0.323766 | 6.69E-06 | 0.216139 |
| Rps11 | 0.32298 | 2.77E-10 | 8.93E-06 |
| Abhd17a | 0.322921 | 0.000242 | 1 |
| Tmem176a | 0.322323 | 0.000567 | 1 |
| Gm8369 | 0.322301 | 6.25E-05 | 1 |

| | | | |
|---|---|---|---|
| Zup1 | 0.322106 | 0.003885 | 1 |
| Rps27a | 0.32112 | 3.11E-10 | 1E-05 |
| Csf2 | 0.320513 | 0.021528 | 1 |
| Thada | 0.319654 | 0.00262 | 1 |
| Cirbp | 0.319448 | 0.00324 | 1 |
| Rpl36 | 0.318477 | 1.2E-08 | 0.000388 |
| Cysltr2 | 0.318169 | 0.000165 | 1 |
| Nfkbiz | 0.317827 | 0.109599 | 1 |
| Susd6 | 0.316703 | 0.009793 | 1 |
| Gpr68 | 0.316505 | 0.000157 | 1 |
| Zfas1 | 0.315887 | 0.000915 | 1 |
| Pcca | 0.315587 | 3.6E-05 | 1 |
| H2-M3 | 0.315161 | 0.000143 | 1 |
| Uba7 | 0.314908 | 0.003284 | 1 |
| Eea1 | 0.314815 | 0.760457 | 1 |
| Ctla2a | 0.313393 | 0.015595 | 1 |
| Hcst | 0.312908 | 0.000382 | 1 |
| Dnaja1 | 0.310426 | 0.002423 | 1 |
| Pbxip1 | 0.310385 | 0.009532 | 1 |
| Ifi211 | 0.310327 | 3.17E-05 | 1 |
| Oxsr1 | 0.310178 | 0.026596 | 1 |
| Ptpn22 | 0.308644 | 0.00026 | 1 |
| Slc3a2 | 0.308487 | 0.000713 | 1 |
| Gpi1 | 0.307629 | 5.39E-05 | 1 |
| Atp8b4 | 0.30761 | 0.001567 | 1 |
| Rpl4 | 0.307001 | 8.61E-07 | 0.027794 |
| Pde4d | 0.305544 | 1.98E-05 | 0.638005 |
| Rab2a | 0.304245 | 7.4E-05 | 1 |
| Rps16 | 0.304027 | 1.01E-10 | 3.27E-06 |
| Klhl6 | 0.303705 | 0.000239 | 1 |
| Selenow | 0.30264 | 0.000511 | 1 |
| Ube2l6 | 0.302432 | 0.01585 | 1 |
| Armc7 | 0.301795 | 0.012328 | 1 |
| Psme1 | 0.301657 | 1.43E-05 | 0.461259 |
| Maml2 | 0.299927 | 0.018509 | 1 |
| Tex2 | 0.299175 | 0.00143 | 1 |
| Eef1g | 0.299057 | 0.000138 | 1 |
| Fkbp5 | 0.298413 | 0.004272 | 1 |
| Hnrnpll | 0.298361 | 0.000115 | 1 |
| Pfkp | 0.29769 | 0.0007 | 1 |
| Pfkl | 0.297316 | 0.00042 | 1 |
| Tmem106b | 0.296918 | 0.002771 | 1 |
| Birc2 | 0.296516 | 9.8E-05 | 1 |
| Eif4a2 | 0.295097 | 0.00231 | 1 |
| Pgam1 | 0.294797 | 0.012963 | 1 |
| Eef1d | 0.293661 | 0.000371 | 1 |
| Rbm3 | 0.293636 | 1.16E-05 | 0.375389 |
| Trafd1 | 0.29248 | 0.012932 | 1 |

| | | | |
|---|---|---|---|
| Fam78a | 0.292255 | 0.007956 | 1 |
| Hsp90ab1 | 0.292108 | 6.68E-05 | 1 |
| Tor1aip1 | 0.291942 | 0.000492 | 1 |
| Airn | 0.291632 | 0.00712 | 1 |
| Parp14 | 0.291211 | 0.009183 | 1 |
| Zeb1 | 0.291206 | 0.005829 | 1 |
| Endod1 | 0.290753 | 0.003082 | 1 |
| Havcr2 | 0.290059 | 0.214617 | 1 |
| Phf11b | 0.288336 | 0.026447 | 1 |
| Tspan31 | 0.288149 | 0.000527 | 1 |
| Atp5g2 | 0.287313 | 7.94E-06 | 0.256303 |
| Trim34a | 0.286985 | 0.008224 | 1 |
| Afg3l2 | 0.28674 | 0.000163 | 1 |
| Hspbp1 | 0.286286 | 0.001476 | 1 |
| Ccdc107 | 0.284389 | 0.000221 | 1 |
| Klrb1c | 0.283897 | 0.000229 | 1 |
| Ikbkb | 0.28376 | 0.003686 | 1 |
| Rpl36a | 0.282621 | 0.00012 | 1 |
| Fau | 0.282052 | 2.43E-09 | 7.85E-05 |
| Ccnl1 | 0.281939 | 0.002042 | 1 |
| Gmfg | 0.281884 | 0.002042 | 1 |
| Hpcal1 | 0.28163 | 0.001807 | 1 |
| Tmem176b | 0.28015 | 0.006479 | 1 |
| Nisch | 0.279717 | 0.004138 | 1 |
| Cd160 | 0.279222 | 0.017141 | 1 |
| Cebpb | 0.278818 | 2.89E-05 | 0.932519 |
| Stat4 | 0.278278 | 0.001033 | 1 |
| Ggh | 0.277965 | 0.001105 | 1 |
| Sit1 | 0.277771 | 0.014688 | 1 |
| 4930549P19I | 0.277706 | 0.000326 | 1 |
| Izumo1r | 0.277083 | 0.009163 | 1 |
| Pde4b | 0.27675 | 0.001267 | 1 |
| Crim1 | 0.275354 | 0.001646 | 1 |
| Pkp3 | 0.274869 | 0.08379 | 1 |
| Zbtb7b | 0.27419 | 4.53E-05 | 1 |
| Cd3d | 0.274076 | 0.000784 | 1 |
| Sik3 | 0.273903 | 0.003554 | 1 |
| Dhx58 | 0.27356 | 0.009928 | 1 |
| Rps8 | 0.27293 | 7.12E-06 | 0.229864 |
| Ybx3 | 0.272877 | 0.000935 | 1 |
| Dis3l2 | 0.272389 | 0.006232 | 1 |
| Skil | 0.271395 | 0.007267 | 1 |
| Tmem59 | 0.271119 | 0.000329 | 1 |
| Acbd6 | 0.271112 | 0.003949 | 1 |
| C1qbp | 0.270786 | 0.017 | 1 |
| Pdcd6 | 0.269536 | 0.001376 | 1 |
| Rplp0 | 0.26952 | 1.7E-06 | 0.054995 |
| Arap2 | 0.269002 | 0.097656 | 1 |

| | | | |
|---|---|---|---|
| Mrpl23 | 0.26838 | 0.001583 | 1 |
| Dcun1d5 | 0.267809 | 0.000652 | 1 |
| Pld3 | 0.266123 | 0.006047 | 1 |
| Rpl10a | 0.266021 | 1.12E-05 | 0.362193 |
| Eif3f | 0.265868 | 4.64E-06 | 0.149956 |
| Ppil4 | 0.26584 | 0.002859 | 1 |
| Limd2 | 0.265643 | 0.000968 | 1 |
| Mink1 | 0.265303 | 0.01061 | 1 |
| Rapgef6 | 0.265149 | 0.001009 | 1 |
| Mob2 | 0.264411 | 0.001688 | 1 |
| Tnfaip3 | 0.263802 | 0.026266 | 1 |
| Ric1 | 0.262129 | 0.000698 | 1 |
| Abcb1b | 0.261777 | 0.008766 | 1 |
| Sema7a | 0.261186 | 0.00014 | 1 |
| Frmd8 | 0.260917 | 0.007438 | 1 |
| Rpl13a | 0.26067 | 1.13E-05 | 0.363694 |
| Cycs | 0.260298 | 0.000469 | 1 |
| Lamtor4 | 0.259975 | 0.000687 | 1 |
| Sfi1 | 0.259738 | 0.002313 | 1 |
| Ifih1 | 0.259698 | 9.11E-06 | 0.293957 |
| Ubr4 | 0.258806 | 0.000328 | 1 |
| Eef1a1 | 0.258599 | 1.18E-07 | 0.003795 |
| Pofut2 | 0.257924 | 0.002704 | 1 |
| Ckb | 0.256916 | 0.00135 | 1 |
| D16Ertd472e | 0.256419 | 0.003654 | 1 |
| Btg2 | 0.256264 | 0.016397 | 1 |
| Serpinb6a | 0.256262 | 0.012877 | 1 |
| Rsad2 | 0.255964 | 0.01394 | 1 |
| Gbp5 | 0.255848 | 0.00127 | 1 |
| Gimap4 | 0.25486 | 0.00232 | 1 |
| Nfkbid | 0.254229 | 0.208389 | 1 |
| Lztfl1 | 0.254013 | 0.001969 | 1 |
| Mif4gd | 0.253556 | 0.002139 | 1 |
| mt-Co2 | 0.252388 | 5.95E-05 | 1 |
| Atp5g1 | 0.251965 | 0.002005 | 1 |
| Cln3 | 0.251961 | 0.005906 | 1 |
| Rnf181 | 0.250797 | 0.019874 | 1 |
| Rtf2 | 0.250423 | 0.005244 | 1 |
| Trmt1 | 0.250172 | 0.000736 | 1 |
| Sntb1 | 0.249555 | 0.095687 | 1 |
| Tmem50a | 0.249004 | 0.000374 | 1 |
| Dpm1 | 0.248956 | 0.012096 | 1 |
| Llph | 0.248819 | 0.006904 | 1 |
| Fam53a | 0.248809 | 0.00216 | 1 |
| Idnk | 0.248793 | 0.002502 | 1 |
| Fyco1 | 0.248397 | 0.053558 | 1 |
| Mndal | 0.248383 | 0.005322 | 1 |
| mt-Co1 | 0.248317 | 6.38E-06 | 0.206032 |

| | | | |
|---|---|---|---|
| Irak2 | 0.248266 | 0.020802 | 1 |
| Rbm17 | 0.247806 | 0.002439 | 1 |
| Cd8a | 0.247737 | 0.779781 | 1 |
| Ifi203 | 0.247591 | 0.017411 | 1 |
| Cyld | 0.246451 | 0.001673 | 1 |
| Ppp1r15a | 0.245747 | 0.000681 | 1 |
| Tmem173 | 0.245652 | 0.033656 | 1 |
| Phc3 | 0.245377 | 0.035213 | 1 |
| Ndufv3 | 0.245361 | 0.001711 | 1 |
| Psmb10 | 0.245267 | 0.001298 | 1 |
| Il12rb1 | 0.245195 | 0.003312 | 1 |
| Itgb3 | 0.245182 | 0.002479 | 1 |
| Ifi204 | 0.245133 | 0.00888 | 1 |
| Serpina3g | 0.245012 | 0.666432 | 1 |
| Lcn4 | 0.244382 | 0.001419 | 1 |
| Sptssa | 0.2442 | 0.015344 | 1 |
| Chd7 | 0.244152 | 0.031671 | 1 |
| Pink1 | 0.244106 | 0.000608 | 1 |
| Zfp292 | 0.243791 | 0.012093 | 1 |
| Malat1 | 0.24354 | 0.001764 | 1 |
| Trim12c | 0.242701 | 0.00095 | 1 |
| Rsrc1 | 0.242292 | 0.069594 | 1 |
| Pecam1 | 0.242168 | 0.000267 | 1 |
| Abr | 0.242007 | 0.005836 | 1 |
| Pde11a | 0.241842 | 0.000144 | 1 |
| Gimap6 | 0.241754 | 0.013828 | 1 |
| BC029722 | 0.240806 | 0.001308 | 1 |
| Smg1 | 0.24051 | 0.019366 | 1 |
| Tmbim4 | 0.240057 | 6.47E-05 | 1 |
| Mcrip1 | 0.239925 | 0.006444 | 1 |
| Zc3h7a | 0.239626 | 0.564199 | 1 |
| Rps3a1 | 0.239434 | 4.28E-06 | 0.138122 |
| Klrc1 | 0.239349 | 0.003054 | 1 |
| Rundc3b | 0.239322 | 0.000843 | 1 |
| Rps3 | 0.239233 | 1.42E-08 | 0.000459 |
| Rps19 | 0.239144 | 0.000329 | 1 |
| Rpl22 | 0.238941 | 1.15E-05 | 0.371452 |
| Cmpk2 | 0.238933 | 0.008854 | 1 |
| Supt4a | 0.238856 | 0.00908 | 1 |
| Clec2d | 0.237917 | 0.01103 | 1 |
| Hgsnat | 0.23777 | 0.001605 | 1 |
| Got2 | 0.237032 | 0.004039 | 1 |
| St14 | 0.236241 | 0.003128 | 1 |
| Trappc9 | 0.236226 | 0.003585 | 1 |
| Kdm2b | 0.236142 | 0.070082 | 1 |
| Rnf145 | 0.235406 | 0.012299 | 1 |
| Fam20a | 0.235317 | 3.58E-05 | 1 |
| Raly | 0.235269 | 0.005643 | 1 |

| | | | |
|---|---|---|---|
| Pitpna | 0.235242 | 0.012178 | 1 |
| Ppm1h | 0.234577 | 0.038327 | 1 |
| Rnf19a | 0.234523 | 0.124874 | 1 |
| Zap70 | 0.234413 | 0.037822 | 1 |
| Cst7 | 0.234107 | 0.015413 | 1 |
| Aars | 0.23362 | 0.025572 | 1 |
| Eif3k | 0.233071 | 0.001574 | 1 |
| Kbtbd11 | 0.232388 | 0.028682 | 1 |
| Hid1 | 0.232056 | 0.001132 | 1 |
| Casp4 | 0.231785 | 0.014887 | 1 |
| Cdk17 | 0.231699 | 0.018476 | 1 |
| Adcy7 | 0.231479 | 0.020301 | 1 |
| Tasor2 | 0.231471 | 0.029739 | 1 |
| Gbp3 | 0.230586 | 0.000472 | 1 |
| Ramp1 | 0.230478 | 0.006916 | 1 |
| Rpl23a | 0.230395 | 0.002065 | 1 |
| Dkkl1 | 0.230175 | 0.001686 | 1 |
| Ddit3 | 0.229616 | 0.004209 | 1 |
| Mitd1 | 0.229482 | 0.051582 | 1 |
| Nsun4 | 0.229424 | 0.000463 | 1 |
| Smyd1 | 0.229348 | 0.007634 | 1 |
| Ppp1r16b | 0.228872 | 0.066336 | 1 |
| Nfkb2 | 0.228367 | 0.007099 | 1 |
| Psma3 | 0.228189 | 0.000488 | 1 |
| Klrb1b | 0.228171 | 0.00163 | 1 |
| Mapre2 | 0.22791 | 0.022175 | 1 |
| Rpl37 | 0.2279 | 1.54E-06 | 0.049583 |
| Cflar | 0.227037 | 0.001006 | 1 |
| Rps27 | 0.226441 | 0.000328 | 1 |
| Tprg | 0.226225 | 0.011363 | 1 |
| Gstm5 | 0.225792 | 0.000291 | 1 |
| Srp72 | 0.22455 | 0.004348 | 1 |
| Cpd | 0.224513 | 0.000832 | 1 |
| Mt2 | 0.22445 | 0.000847 | 1 |
| Abhd18 | 0.224422 | 0.001097 | 1 |
| Malt1 | 0.223796 | 0.118629 | 1 |
| Sqstm1 | 0.22371 | 0.015189 | 1 |
| Tomm20 | 0.223436 | 0.009944 | 1 |
| Trbv12-2 | 0.223303 | 0.003842 | 1 |
| Rnf114 | 0.222855 | 0.020835 | 1 |
| Irf9 | 0.222833 | 0.013627 | 1 |
| Slfn5 | 0.222774 | 0.000933 | 1 |
| Cmip | 0.222761 | 0.04837 | 1 |
| Eno1b | 0.221684 | 0.018878 | 1 |
| Fdx2 | 0.221585 | 0.003769 | 1 |
| Trak2 | 0.221429 | 0.002791 | 1 |
| Cdkal1 | 0.221319 | 0.011833 | 1 |
| Usp18 | 0.221275 | 0.024246 | 1 |

| | | | |
|---|---|---|---|
| Fam174a | 0.221126 | 0.000982 | 1 |
| Ttc39c | 0.220959 | 0.004419 | 1 |
| Rilpl2 | 0.220861 | 0.078373 | 1 |
| Eprs | 0.220701 | 0.003501 | 1 |
| Nosip | 0.219666 | 0.008417 | 1 |
| Plcg1 | 0.21955 | 0.008801 | 1 |
| Ndufv1 | 0.219244 | 0.026079 | 1 |
| Sp110 | 0.219063 | 0.049799 | 1 |
| Plekhm3 | 0.218852 | 0.000955 | 1 |
| Cd247 | 0.218387 | 0.03022 | 1 |
| Gdpd5 | 0.218316 | 0.000759 | 1 |
| Rpl7 | 0.218134 | 7.12E-05 | 1 |
| Hspe1 | 0.217659 | 0.001068 | 1 |
| Scand1 | 0.21733 | 0.006375 | 1 |
| Gsto1 | 0.216987 | 0.003788 | 1 |
| H2-T24 | 0.21666 | 0.010219 | 1 |
| Hddc2 | 0.216348 | 0.006672 | 1 |
| Wdfy1 | 0.216287 | 0.03067 | 1 |
| Ext1 | 0.216108 | 0.027354 | 1 |
| Lcp2 | 0.215877 | 0.025904 | 1 |
| Gm12185 | 0.215473 | 0.065607 | 1 |
| Trim30d | 0.2151 | 0.041276 | 1 |
| Birc6 | 0.2151 | 0.028601 | 1 |
| Gm20400 | 0.214627 | 0.024255 | 1 |
| Camk1d | 0.214417 | 0.423069 | 1 |
| Bhlhe40 | 0.213349 | 0.024518 | 1 |
| Tuba4a | 0.213307 | 0.01389 | 1 |
| Ly6g5b | 0.213293 | 0.000603 | 1 |
| Asb2 | 0.212722 | 0.036134 | 1 |
| Uqcrh | 0.212591 | 0.002864 | 1 |
| Sbds | 0.212575 | 0.007889 | 1 |
| AC149090.1 | 0.212386 | 0.163848 | 1 |
| Ern1 | 0.212214 | 0.004734 | 1 |
| Rpl22l1 | 0.212111 | 0.001922 | 1 |
| Hspa5 | 0.211979 | 0.021169 | 1 |
| Ppp1r12a | 0.211939 | 0.081078 | 1 |
| Tespa1 | 0.211853 | 0.110519 | 1 |
| Eif2s2 | 0.211204 | 0.008351 | 1 |
| Prxl2a | 0.211107 | 0.0003 | 1 |
| Aebp2 | 0.211029 | 0.027017 | 1 |
| Rps6 | 0.210681 | 0.007307 | 1 |
| Srpk1 | 0.210233 | 0.018254 | 1 |
| Cmtm7 | 0.210201 | 0.085967 | 1 |
| Tmem241 | 0.209896 | 0.001419 | 1 |
| Max | 0.209869 | 0.039635 | 1 |
| Selenos | 0.209854 | 0.011433 | 1 |
| Aprt | 0.209422 | 0.017867 | 1 |
| Pcmtd1 | 0.209095 | 0.043914 | 1 |

| | | | |
|---|---|---|---|
| Alcam | 0.208741 | 0.129365 | 1 |
| Znrf1 | 0.208697 | 0.024451 | 1 |
| Naca | 0.20853 | 0.000365 | 1 |
| Rabgap1l | 0.208486 | 0.559109 | 1 |
| Atp2b4 | 0.208316 | 0.026363 | 1 |
| Txn2 | 0.207384 | 0.004726 | 1 |
| 2610035D17 | 0.207365 | 0.005455 | 1 |
| Slc39a11 | 0.206247 | 0.003868 | 1 |
| Tor4a | 0.206104 | 0.011102 | 1 |
| Sec22a | 0.206072 | 0.001283 | 1 |
| Ldha | 0.205776 | 0.01674 | 1 |
| Hhat | 0.205262 | 0.000108 | 1 |
| Rrp1b | 0.205259 | 0.005195 | 1 |
| Lta | 0.204378 | 0.072699 | 1 |
| Rasgrp1 | 0.204375 | 0.067462 | 1 |
| S100a6 | 0.204262 | 0.048174 | 1 |
| Cox5a | 0.204163 | 0.015009 | 1 |
| Mlxip | 0.204141 | 0.050366 | 1 |
| Mier1 | 0.20404 | 0.001669 | 1 |
| Akap13 | 0.20378 | 0.009767 | 1 |
| Clip1 | 0.203542 | 0.003075 | 1 |
| Psmb3 | 0.203346 | 0.009705 | 1 |
| Ccdc85b | 0.203184 | 0.055443 | 1 |
| Kdm4d | 0.202868 | 0.000479 | 1 |
| Mov10 | 0.202809 | 0.031433 | 1 |
| Iigp1 | 0.202669 | 5.07E-06 | 0.163694 |
| Spry1 | 0.202389 | 0.000662 | 1 |
| Kmt5b | 0.20226 | 0.042779 | 1 |
| 2810013P06I | 0.201906 | 0.028365 | 1 |
| Chrac1 | 0.201262 | 0.006688 | 1 |
| Ostf1 | 0.201208 | 0.022403 | 1 |
| Got1 | 0.201114 | 0.02051 | 1 |
| Smpd5 | 0.200946 | 0.013336 | 1 |
| Vmn2r97 | 0.200735 | 0.001352 | 1 |
| Gimap9 | 0.200526 | 0.029499 | 1 |
| Kdm5b | 0.200298 | 0.003843 | 1 |
| Bbx | 0.200253 | 0.026005 | 1 |
| Rpl37a | 0.200139 | 4.89E-05 | 1 |
| Tmub1 | 0.199737 | 0.002929 | 1 |
| Acot13 | 0.199547 | 0.005875 | 1 |
| Vax2 | 0.198933 | 0.001316 | 1 |
| Sla | 0.198859 | 0.003412 | 1 |
| Slc25a17 | 0.198636 | 0.058647 | 1 |
| Crtam | 0.198173 | 0.238985 | 1 |
| Ece1 | 0.197651 | 0.117862 | 1 |
| Capza2 | 0.197227 | 0.008177 | 1 |
| Tvp23b | 0.197097 | 0.007341 | 1 |
| Nenf | 0.196907 | 0.005138 | 1 |

| | | | |
|---|---|---|---|
| Eif3i | 0.196387 | 0.00726 | 1 |
| Mapkapk2 | 0.196282 | 0.022068 | 1 |
| Asb13 | 0.19628 | 0.003302 | 1 |
| Ccz1 | 0.196215 | 0.045051 | 1 |
| Dusp1 | 0.196156 | 0.907573 | 1 |
| Spats2 | 0.196112 | 0.000249 | 1 |
| Akr1a1 | 0.195702 | 0.014198 | 1 |
| Pim1 | 0.195675 | 0.143048 | 1 |
| Yy1 | 0.194649 | 0.03512 | 1 |
| Ifi206 | 0.19419 | 0.221348 | 1 |
| Cox7a2l | 0.193885 | 0.014342 | 1 |
| Zcchc7 | 0.19361 | 0.034867 | 1 |
| Hcfc1r1 | 0.193544 | 0.087469 | 1 |
| Actr10 | 0.193456 | 0.040194 | 1 |
| Pycr2 | 0.193314 | 0.014537 | 1 |
| Zc3h12a | 0.192836 | 0.02412 | 1 |
| Zfp53 | 0.192826 | 0.015821 | 1 |
| Saysd1 | 0.192726 | 0.086184 | 1 |
| Arhgap10 | 0.192689 | 0.088117 | 1 |
| Galm | 0.192577 | 0.005503 | 1 |
| Stk17b | 0.192525 | 0.020885 | 1 |
| B4galnt4 | 0.192412 | 0.000369 | 1 |
| Cd28 | 0.192234 | 0.036549 | 1 |
| Acp1 | 0.192025 | 0.027298 | 1 |
| Asb3 | 0.191861 | 0.058073 | 1 |
| Nfkbib | 0.191797 | 0.072774 | 1 |
| Aph1a | 0.191307 | 0.016228 | 1 |
| Gm19705 | 0.190397 | 0.000486 | 1 |
| Chd6 | 0.189907 | 0.043574 | 1 |
| Micu2 | 0.189831 | 0.00279 | 1 |
| Adam8 | 0.189498 | 0.040144 | 1 |
| Gmds | 0.189304 | 0.035408 | 1 |
| Parp10 | 0.189234 | 0.041474 | 1 |
| Aen | 0.189141 | 0.035087 | 1 |
| Agap2 | 0.188741 | 0.011858 | 1 |
| Gimap5 | 0.188687 | 0.072622 | 1 |
| Mast4 | 0.188317 | 0.000438 | 1 |
| Ola1 | 0.188232 | 0.023254 | 1 |
| Ubr1 | 0.187924 | 0.026612 | 1 |
| Gpr132 | 0.18787 | 0.053353 | 1 |
| Ptpn6 | 0.187669 | 5.13E-05 | 1 |
| Nop53 | 0.187388 | 0.038862 | 1 |
| Snx20 | 0.187367 | 0.008374 | 1 |
| Mbd2 | 0.187063 | 0.016008 | 1 |
| Smim10l1 | 0.187018 | 0.028509 | 1 |
| Fam189b | 0.186837 | 0.191222 | 1 |
| Xlr4b | 0.186747 | 0.024075 | 1 |
| Soat1 | 0.186584 | 0.006541 | 1 |

| | | | |
|---|---|---|---|
| Zfp1 | 0.186354 | 0.013597 | 1 |
| Hibadh | 0.186345 | 0.025893 | 1 |
| Rnf125 | 0.186326 | 0.061673 | 1 |
| Wwox | 0.186209 | 0.028073 | 1 |
| Zc3h6 | 0.18608 | 0.019948 | 1 |
| Ufc1 | 0.186021 | 0.026525 | 1 |
| Ifit2 | 0.185935 | 0.552854 | 1 |
| Sorl1 | 0.185133 | 0.036195 | 1 |
| Tmem192 | 0.185099 | 0.008468 | 1 |
| Gm26510 | 0.184972 | 0.014399 | 1 |
| Evi2a | 0.184962 | 0.043489 | 1 |
| Mrps6 | 0.18484 | 0.011694 | 1 |
| Trappc6a | 0.184528 | 0.012459 | 1 |
| Rab11fip1 | 0.184432 | 0.021621 | 1 |
| Mdn1 | 0.184418 | 0.02025 | 1 |
| Exph5 | 0.184415 | 0.000587 | 1 |
| Hdac2 | 0.184214 | 0.01909 | 1 |
| Vmp1 | 0.1842 | 0.008921 | 1 |
| Trnau1ap | 0.184098 | 0.015409 | 1 |
| Itgb7 | 0.18409 | 0.027466 | 1 |
| Hspa8 | 0.184029 | 0.002139 | 1 |
| Serbp1 | 0.18385 | 0.012482 | 1 |
| Gm20559 | 0.183842 | 0.017615 | 1 |
| Gstt2 | 0.183767 | 0.015417 | 1 |
| Zc3h13 | 0.183722 | 0.078281 | 1 |
| Gadd45b | 0.183462 | 0.008577 | 1 |
| Ncor2 | 0.183407 | 0.015661 | 1 |
| Fam169b | 0.183256 | 0.066849 | 1 |
| Gm11361 | 0.183169 | 0.000761 | 1 |
| Gm20275 | 0.182986 | 0.21823 | 1 |
| Rrp1 | 0.182661 | 0.003833 | 1 |
| Rsbn1 | 0.181891 | 0.039676 | 1 |
| Desi2 | 0.181205 | 0.037864 | 1 |
| Tmem189 | 0.181191 | 0.009516 | 1 |
| Ythdc1 | 0.180788 | 0.131312 | 1 |
| B230219D22 | 0.180668 | 0.014834 | 1 |
| Il3ra | 0.180578 | 0.023419 | 1 |
| Ssbp4 | 0.180129 | 0.011204 | 1 |
| Usf2 | 0.180051 | 0.001824 | 1 |
| Ptpn1 | 0.180044 | 0.011067 | 1 |
| Phb2 | 0.179816 | 0.004537 | 1 |
| Slain1 | 0.179787 | 0.051638 | 1 |
| Impdh2 | 0.179203 | 0.034117 | 1 |
| Ppm1k | 0.178909 | 0.007566 | 1 |
| Eva1b | 0.178592 | 0.097598 | 1 |
| Slc39a14 | 0.178371 | 0.001981 | 1 |
| Rabggtb | 0.178361 | 0.063201 | 1 |
| Asns | 0.17808 | 0.001931 | 1 |

| | | | |
|---|---|---|---|
| Setd3 | 0.177842 | 0.023194 | 1 |
| Ppp1cc | 0.177771 | 0.0098 | 1 |
| Mrpl9 | 0.177511 | 0.028913 | 1 |
| Bod1l | 0.177287 | 0.08007 | 1 |
| Aim2 | 0.176953 | 0.08239 | 1 |
| Sgms1 | 0.176908 | 0.010747 | 1 |
| Dctn2 | 0.176756 | 0.035866 | 1 |
| Tmed2 | 0.176497 | 0.048035 | 1 |
| Efr3a | 0.176484 | 0.082076 | 1 |
| Trim12a | 0.175971 | 0.063054 | 1 |
| Ifit1bl1 | 0.175584 | 9.3E-05 | 1 |
| Mrpl58 | 0.175549 | 0.097968 | 1 |
| Hectd1 | 0.17527 | 0.028492 | 1 |
| 1110038F14l | 0.174683 | 0.114112 | 1 |
| Etv6 | 0.174579 | 0.216029 | 1 |
| Esyt2 | 0.174076 | 0.058828 | 1 |
| Arntl | 0.174009 | 0.040101 | 1 |
| Arl6ip1 | 0.173922 | 0.012907 | 1 |
| Thy1 | 0.173367 | 0.038089 | 1 |
| Spata6 | 0.173244 | 0.021159 | 1 |
| Fto | 0.173238 | 0.020775 | 1 |
| Xbp1 | 0.173196 | 0.029119 | 1 |
| Sntb2 | 0.173101 | 0.00777 | 1 |
| Ptpn11 | 0.173027 | 0.151098 | 1 |
| Galnt10 | 0.173021 | 0.026004 | 1 |
| Zc3hav1 | 0.172922 | 0.036879 | 1 |
| Pip4k2a | 0.172827 | 0.050044 | 1 |
| Il27ra | 0.172737 | 0.03799 | 1 |
| Rpl28 | 0.172672 | 0.04616 | 1 |
| Rab5if | 0.172557 | 0.061521 | 1 |
| Rnf166 | 0.172306 | 0.017319 | 1 |
| Arid5b | 0.172279 | 0.435331 | 1 |
| Sigirr | 0.171962 | 0.042562 | 1 |
| Tmem251 | 0.171846 | 0.002181 | 1 |
| Smad7 | 0.171784 | 0.015345 | 1 |
| Tspan5 | 0.171663 | 0.025397 | 1 |
| Psma2 | 0.171352 | 0.030135 | 1 |
| Sytl1 | 0.171328 | 0.000171 | 1 |
| Ubxn2a | 0.170766 | 0.090211 | 1 |
| Denr | 0.170218 | 0.044884 | 1 |
| Phyh | 0.170109 | 0.087199 | 1 |
| Pgap2 | 0.170095 | 0.0743 | 1 |
| Apopt1 | 0.169937 | 0.040196 | 1 |
| Susd3 | 0.169913 | 0.015462 | 1 |
| Slc48a1 | 0.169762 | 0.019862 | 1 |
| Fam162a | 0.169553 | 0.030621 | 1 |
| Nubp2 | 0.169522 | 0.06597 | 1 |
| Dock8 | 0.169465 | 0.083331 | 1 |

| | | | |
|---|---|---|---|
| Rasal3 | 0.169454 | 0.026028 | 1 |
| Rest | 0.169302 | 0.050128 | 1 |
| Ech1 | 0.169183 | 0.060544 | 1 |
| Tmem38b | 0.169133 | 0.008951 | 1 |
| Rpl36al | 0.169014 | 0.011039 | 1 |
| Btbd19 | 0.168592 | 0.097404 | 1 |
| Trim14 | 0.168547 | 0.015013 | 1 |
| Psma7 | 0.168515 | 0.026379 | 1 |
| Hacd3 | 0.168491 | 0.136209 | 1 |
| Atp5l | 0.167878 | 0.005552 | 1 |
| Ssh2 | 0.167807 | 0.052768 | 1 |
| Serhl | 0.167764 | 0.031085 | 1 |
| Lsm14a | 0.167604 | 0.014627 | 1 |
| Trim21 | 0.167535 | 0.252308 | 1 |
| Pik3ca | 0.167483 | 0.026453 | 1 |
| Farp1 | 0.167455 | 0.139477 | 1 |
| Mpzl3 | 0.167441 | 0.00093 | 1 |
| Golga5 | 0.167215 | 0.035784 | 1 |
| Slk | 0.166899 | 0.157298 | 1 |
| Sdf2 | 0.166819 | 0.058258 | 1 |
| Nab1 | 0.166794 | 0.295301 | 1 |
| Tmem191c | 0.165958 | 0.006079 | 1 |
| Fam120c | 0.165933 | 0.018522 | 1 |
| Metrnl | 0.165586 | 0.0523 | 1 |
| Mpv17l2 | 0.165513 | 0.051005 | 1 |
| Rnf5 | 0.164935 | 0.058043 | 1 |
| Cry1 | 0.164478 | 0.02106 | 1 |
| Zcchc2 | 0.164443 | 0.027869 | 1 |
| Dock9 | 0.164383 | 0.16009 | 1 |
| Cox14 | 0.164366 | 0.078266 | 1 |
| Nars | 0.16426 | 0.277573 | 1 |
| Nup153 | 0.164195 | 0.055606 | 1 |
| Btbd11 | 0.163894 | 0.092874 | 1 |
| Dcaf8 | 0.163731 | 0.044148 | 1 |
| Sart1 | 0.16367 | 0.050858 | 1 |
| Epb41l4aos | 0.163484 | 0.006946 | 1 |
| Samd10 | 0.163108 | 0.0071 | 1 |
| Ndufs7 | 0.163103 | 0.002048 | 1 |
| Rpl35a | 0.163079 | 0.00037 | 1 |
| Fam32a | 0.162973 | 0.125953 | 1 |
| Mat2b | 0.162747 | 0.030724 | 1 |
| Mir155hg | 0.162639 | 0.248212 | 1 |
| Zfp945 | 0.16225 | 0.074478 | 1 |
| Ssu72 | 0.1619 | 0.05993 | 1 |
| Tet3 | 0.161896 | 0.058857 | 1 |
| Gm49417 | 0.161857 | 0.060625 | 1 |
| Grhpr | 0.16184 | 0.035979 | 1 |
| Dusp6 | 0.161831 | 0.078637 | 1 |

| | | | |
|---|---|---|---|
| Get4 | 0.161626 | 0.059635 | 1 |
| Znhit2 | 0.161583 | 0.002438 | 1 |
| Mfsd6 | 0.161539 | 0.010794 | 1 |
| Ndufa6 | 0.161535 | 0.046063 | 1 |
| Txnrd2 | 0.161515 | 0.020505 | 1 |
| Polr3k | 0.161171 | 0.087775 | 1 |
| Cotl1 | 0.161118 | 0.158151 | 1 |
| 4833438C02I | 0.160897 | 0.016998 | 1 |
| Ppp5c | 0.160547 | 0.159347 | 1 |
| Mcts1 | 0.160408 | 0.046177 | 1 |
| Ssr4 | 0.160273 | 0.020392 | 1 |
| Ddx6 | 0.159887 | 0.093825 | 1 |
| Xpot | 0.159878 | 0.007688 | 1 |
| Tut4 | 0.1597 | 0.150565 | 1 |
| Pithd1 | 0.159694 | 0.005103 | 1 |
| Cpne3 | 0.159486 | 0.005953 | 1 |
| Ubald1 | 0.159404 | 0.018096 | 1 |
| Parp8 | 0.159391 | 0.090376 | 1 |
| Odc1 | 0.159205 | 0.141465 | 1 |
| Fancc | 0.159136 | 0.046618 | 1 |
| Snd1 | 0.158792 | 0.04194 | 1 |
| Filip1 | 0.158743 | 0.006968 | 1 |
| Camk4 | 0.158615 | 0.112243 | 1 |
| Erich1 | 0.158584 | 0.197745 | 1 |
| Tmem131 | 0.158563 | 0.018725 | 1 |
| Helz2 | 0.158162 | 0.101641 | 1 |
| Gngt2 | 0.157991 | 0.019408 | 1 |
| Sdhd | 0.157966 | 0.062242 | 1 |
| Slc7a5 | 0.157685 | 0.024202 | 1 |
| Lrrc61 | 0.157556 | 0.008144 | 1 |
| Fyb | 0.157552 | 0.061356 | 1 |
| Card6 | 0.157291 | 0.073996 | 1 |
| Eya2 | 0.157224 | 0.008451 | 1 |
| Ctdnep1 | 0.157004 | 0.136524 | 1 |
| Mmadhc | 0.157 | 0.056888 | 1 |
| Foxj3 | 0.15686 | 0.023594 | 1 |
| Ahr | 0.156807 | 0.032915 | 1 |
| Gtpbp2 | 0.156698 | 0.048319 | 1 |
| Dynlt3 | 0.156662 | 0.007926 | 1 |
| Cdk5rap1 | 0.156421 | 0.01677 | 1 |
| Rmnd5b | 0.156067 | 0.055142 | 1 |
| Qrfp | 0.155965 | 0.101196 | 1 |
| Mapk14 | 0.155889 | 0.068467 | 1 |
| Crbn | 0.155694 | 0.19713 | 1 |
| Tnfsf11 | 0.155608 | 0.009126 | 1 |
| Tesc | 0.155595 | 0.015112 | 1 |
| Dennd4a | 0.155373 | 0.073311 | 1 |
| Las1l | 0.155098 | 0.142584 | 1 |

| | | | |
|---|---|---|---|
| Eif4e3 | 0.155095 | 0.053312 | 1 |
| 2310011J03F | 0.154881 | 0.087457 | 1 |
| Rala | 0.154737 | 0.043264 | 1 |
| Tpr | 0.154686 | 0.110123 | 1 |
| Chordc1 | 0.154615 | 0.158918 | 1 |
| Phf21a | 0.154517 | 0.082237 | 1 |
| Il15ra | 0.154419 | 0.071931 | 1 |
| Snhg1 | 0.154413 | 0.052857 | 1 |
| BC031181 | 0.154388 | 0.104651 | 1 |
| Rab5c | 0.154189 | 0.170993 | 1 |
| Hspa9 | 0.154122 | 0.325959 | 1 |
| Araf | 0.153956 | 0.124242 | 1 |
| Pcgf5 | 0.153954 | 0.056354 | 1 |
| Tacc1 | 0.153822 | 0.093321 | 1 |
| Fis1 | 0.153653 | 0.049946 | 1 |
| 2900041M22 | 0.153541 | 0.002962 | 1 |
| Gadd45gip1 | 0.15346 | 0.0329 | 1 |
| Foxp4 | 0.152976 | 0.11127 | 1 |
| Eif3m | 0.152537 | 0.00908 | 1 |
| Pdhb | 0.152533 | 0.092314 | 1 |
| Ppt2 | 0.151893 | 0.019738 | 1 |
| Focad | 0.151879 | 0.113167 | 1 |
| Tnfsf14 | 0.151838 | 0.032282 | 1 |
| Skap1 | 0.15172 | 0.09303 | 1 |
| Oxa1l | 0.151704 | 0.031016 | 1 |
| Apex1 | 0.151611 | 0.023825 | 1 |
| Grap2 | 0.151424 | 0.130658 | 1 |
| Mfng | 0.150774 | 0.100733 | 1 |
| Fbl | 0.150704 | 0.172962 | 1 |
| Pgam5 | 0.150556 | 0.03993 | 1 |
| Tmem39a | 0.150503 | 0.07702 | 1 |
| Crnkl1 | 0.150497 | 0.102522 | 1 |
| Trib2 | 0.15004 | 0.008123 | 1 |
| Gch1 | 0.150021 | 0.118176 | 1 |
| H2-Ke6 | 0.150019 | 0.070964 | 1 |
| Cox7b | 0.149982 | 0.023578 | 1 |
| Ttc39b | 0.149897 | 0.230808 | 1 |
| Rrs1 | 0.149625 | 0.024148 | 1 |
| Sun2 | 0.149269 | 0.133933 | 1 |
| Ms4a6c | 0.149207 | 0.210021 | 1 |
| Bnip3l | 0.14895 | 0.046682 | 1 |
| Cd3e | 0.148599 | 0.251752 | 1 |
| Oprm1 | 0.148565 | 0.013936 | 1 |
| mt-Nd1 | 0.148481 | 0.138163 | 1 |
| Gm11707 | 0.148475 | 0.039366 | 1 |
| Rhbdd1 | 0.148402 | 0.110892 | 1 |
| Dnajc5 | 0.14831 | 0.07146 | 1 |
| Sod1 | 0.147962 | 0.069994 | 1 |

| | | | |
|---|---|---|---|
| Ccdc28b | 0.147522 | 0.040672 | 1 |
| Gm49359 | 0.147179 | 0.034719 | 1 |
| Eci2 | 0.147146 | 0.076736 | 1 |
| Hilpda | 0.147136 | 0.007 | 1 |
| Izumo4 | 0.147116 | 0.167425 | 1 |
| Caprin1 | 0.14705 | 0.082793 | 1 |
| Phf11c | 0.147014 | 0.421344 | 1 |
| Nt5m | 0.146856 | 0.01452 | 1 |
| Tmem140 | 0.14672 | 0.158739 | 1 |
| Atpaf1 | 0.146633 | 0.043634 | 1 |
| Wdr83os | 0.14618 | 0.10365 | 1 |
| Ell2 | 0.14597 | 0.004294 | 1 |
| Chchd3 | 0.14593 | 0.013722 | 1 |
| Wdr7 | 0.145912 | 0.04847 | 1 |
| Cltb | 0.14591 | 0.1587 | 1 |
| Orc3 | 0.145801 | 0.097917 | 1 |
| Dgat1 | 0.145553 | 0.054659 | 1 |
| AI662270 | 0.145476 | 0.161684 | 1 |
| Rps15 | 0.145428 | 0.070146 | 1 |
| Leprotl1 | 0.14528 | 0.039535 | 1 |
| Dennd2d | 0.144985 | 0.04798 | 1 |
| Rtraf | 0.144751 | 0.092907 | 1 |
| Ifi214 | 0.144738 | 0.24934 | 1 |
| Pdhx | 0.144588 | 0.007503 | 1 |
| Ldah | 0.14443 | 0.151228 | 1 |
| Nucb1 | 0.144398 | 0.106583 | 1 |
| Phax | 0.144393 | 0.031551 | 1 |
| Matk | 0.144363 | 0.004797 | 1 |
| Capg | 0.144306 | 0.060877 | 1 |
| Rab43 | 0.144279 | 0.003819 | 1 |
| Prr13 | 0.144165 | 0.019815 | 1 |
| Nub1 | 0.144141 | 0.09455 | 1 |
| Acpp | 0.144071 | 0.008706 | 1 |
| Atg3 | 0.144037 | 0.131368 | 1 |
| Zdhhc18 | 0.144016 | 0.080708 | 1 |
| Vps13d | 0.143961 | 0.193518 | 1 |
| Uri1 | 0.143851 | 0.132836 | 1 |
| Foxo3 | 0.14384 | 0.055207 | 1 |
| Plaat3 | 0.14381 | 0.00879 | 1 |
| Usp48 | 0.143668 | 0.266139 | 1 |
| Pxn | 0.143578 | 0.061251 | 1 |
| Parl | 0.143558 | 0.036469 | 1 |
| Nfatc2 | 0.143546 | 0.323864 | 1 |
| Kiz | 0.143481 | 0.021189 | 1 |
| Ccdc92 | 0.143399 | 0.022811 | 1 |
| Dph6 | 0.14313 | 0.175715 | 1 |
| P4ha1 | 0.142901 | 0.06585 | 1 |
| Senp6 | 0.142851 | 0.094427 | 1 |

| | | | |
|---|---|---|---|
| Fbxo38 | 0.142848 | 0.022103 | 1 |
| Cep120 | 0.142842 | 0.227127 | 1 |
| Selenok | 0.1428 | 0.114676 | 1 |
| Ptprj | 0.142775 | 0.304731 | 1 |
| 1500011B03I | 0.142713 | 0.046493 | 1 |
| Clcc1 | 0.142536 | 0.076381 | 1 |
| Ndufaf4 | 0.142433 | 0.063772 | 1 |
| D5Ertd579e | 0.142376 | 0.094837 | 1 |
| Selenoo | 0.142255 | 0.043717 | 1 |
| Exosc4 | 0.142063 | 0.004435 | 1 |
| Adamts6 | 0.142041 | 0.331967 | 1 |
| Cir1 | 0.141656 | 0.072411 | 1 |
| Plcl1 | 0.141608 | 0.010404 | 1 |
| Plekho2 | 0.141343 | 0.131447 | 1 |
| Rinl | 0.141176 | 0.305404 | 1 |
| Tulp4 | 0.141092 | 0.18387 | 1 |
| Man2c1 | 0.14101 | 0.038482 | 1 |
| Eefsec | 0.140936 | 0.081529 | 1 |
| Yipf2 | 0.14084 | 0.002192 | 1 |
| Crtc2 | 0.140802 | 0.25184 | 1 |
| Fam53b | 0.140784 | 0.059928 | 1 |
| Ezh1 | 0.140638 | 0.095893 | 1 |
| Chchd4 | 0.140633 | 0.018206 | 1 |
| Bdh1 | 0.140371 | 0.052372 | 1 |
| Zfp397 | 0.140211 | 0.043468 | 1 |
| Pigx | 0.140105 | 0.03156 | 1 |
| Mtmr1 | 0.14009 | 0.044735 | 1 |
| Spint2 | 0.140024 | 0.227803 | 1 |
| Dnajc3 | 0.139923 | 0.097788 | 1 |
| Zfp672 | 0.139799 | 0.127022 | 1 |
| Csnk2b | 0.139646 | 0.108037 | 1 |
| Mtfr1l | 0.139525 | 0.100071 | 1 |
| Coq10b | 0.139109 | 0.116698 | 1 |
| Stk19 | 0.139004 | 0.054443 | 1 |
| Mbp | 0.138931 | 0.110951 | 1 |
| Wwp1 | 0.138916 | 0.025443 | 1 |
| Gpr65 | 0.138769 | 0.29532 | 1 |
| 0610012G03 | 0.138674 | 0.015579 | 1 |
| Mrpl37 | 0.138532 | 0.094681 | 1 |
| Med13l | 0.13853 | 0.085682 | 1 |
| Rassf5 | 0.138508 | 0.241675 | 1 |
| Agfg1 | 0.13843 | 0.19943 | 1 |
| Akr7a5 | 0.138413 | 0.034928 | 1 |
| Scamp3 | 0.138323 | 0.018595 | 1 |
| Sh2b1 | 0.138071 | 0.023521 | 1 |
| Snhg8 | 0.138012 | 0.046684 | 1 |
| Zfp683 | 0.137887 | 0.02755 | 1 |
| Aar2 | 0.137689 | 0.069789 | 1 |

| | | | |
|---|---|---|---|
| Ago2 | 0.137255 | 0.261589 | 1 |
| Nkiras1 | 0.137211 | 0.013302 | 1 |
| Utp4 | 0.137054 | 0.203815 | 1 |
| Abhd12 | 0.136914 | 0.046013 | 1 |
| Tk2 | 0.13691 | 0.062883 | 1 |
| Ginm1 | 0.136894 | 0.112771 | 1 |
| Asap1 | 0.136859 | 0.462596 | 1 |
| Ddx41 | 0.136838 | 0.254215 | 1 |
| Ubr5 | 0.136788 | 0.152778 | 1 |
| Dop1b | 0.13678 | 0.236065 | 1 |
| Aarsd1 | 0.136749 | 0.164862 | 1 |
| Tmem9 | 0.136682 | 0.041714 | 1 |
| Card19 | 0.136676 | 0.252931 | 1 |
| Slc25a30 | 0.136588 | 0.022578 | 1 |
| Uqcc3 | 0.136557 | 0.052073 | 1 |
| Ubn2 | 0.136513 | 0.044734 | 1 |
| Tiparp | 0.136254 | 0.024145 | 1 |
| Pebp1 | 0.136065 | 0.089181 | 1 |
| L3mbtl3 | 0.135972 | 0.177769 | 1 |
| Sars | 0.135866 | 0.314931 | 1 |
| Arf3 | 0.135689 | 0.051328 | 1 |
| Commd8 | 0.135588 | 0.338757 | 1 |
| Noc4l | 0.135503 | 0.067882 | 1 |
| Mdp1 | 0.135444 | 0.163488 | 1 |
| Cyp4f13 | 0.135414 | 0.030401 | 1 |
| Daxx | 0.135078 | 0.035928 | 1 |
| Sec24b | 0.135033 | 0.068072 | 1 |
| Mrps26 | 0.134784 | 0.110116 | 1 |
| 2410002F23I | 0.134752 | 0.112891 | 1 |
| Aig1 | 0.134672 | 0.044409 | 1 |
| Insig1 | 0.134602 | 0.011678 | 1 |
| Ocel1 | 0.134591 | 0.030116 | 1 |
| Crem | 0.134145 | 0.060887 | 1 |
| Asb8 | 0.134079 | 0.078576 | 1 |
| Uqcr10 | 0.134004 | 0.052059 | 1 |
| Fmnl3 | 0.133999 | 0.258273 | 1 |
| Pbx3 | 0.13384 | 0.033481 | 1 |
| mt-Nd5 | 0.133812 | 0.121001 | 1 |
| Pin4 | 0.13361 | 0.216532 | 1 |
| Tollip | 0.133605 | 0.06389 | 1 |
| Nprl3 | 0.133516 | 0.0871 | 1 |
| Xpo4 | 0.133485 | 0.192172 | 1 |
| Clk2 | 0.133365 | 0.019129 | 1 |
| Hbp1 | 0.133358 | 0.01166 | 1 |
| Sar1b | 0.133187 | 0.267423 | 1 |
| Snx29 | 0.133013 | 0.264769 | 1 |
| Emc7 | 0.132996 | 0.240106 | 1 |
| Elavl1 | 0.132927 | 0.063253 | 1 |

| | | | |
|---|---|---|---|
| Bclaf1 | 0.132916 | 0.118308 | 1 |
| Pde7a | 0.132828 | 0.710993 | 1 |
| Gpatch4 | 0.132791 | 0.026654 | 1 |
| Dnttip1 | 0.13261 | 0.011656 | 1 |
| Myo3b | 0.132568 | 0.237383 | 1 |
| Abi2 | 0.132506 | 0.071017 | 1 |
| Fut8 | 0.132192 | 0.157623 | 1 |
| Stxbp5 | 0.132181 | 0.129798 | 1 |
| Chka | 0.13217 | 0.057695 | 1 |
| Usp15 | 0.131536 | 0.311724 | 1 |
| Ssbp2 | 0.131341 | 0.005393 | 1 |
| Eif4ebp1 | 0.131243 | 0.025689 | 1 |
| Zmynd8 | 0.13121 | 0.139801 | 1 |
| Ifrd1 | 0.131174 | 0.166646 | 1 |
| Ccdc102a | 0.131113 | 0.025825 | 1 |
| Tbl1xr1 | 0.131027 | 0.244972 | 1 |
| Kdsr | 0.130905 | 0.115705 | 1 |
| Usp3 | 0.130804 | 0.198583 | 1 |
| Cox8a | 0.129935 | 0.026302 | 1 |
| Stx11 | 0.129821 | 0.153524 | 1 |
| Ube2v1 | 0.129648 | 0.227493 | 1 |
| Stx6 | 0.129566 | 0.270055 | 1 |
| Prdx5 | 0.129299 | 0.039432 | 1 |
| Bcas3 | 0.129291 | 0.33902 | 1 |
| Ntng2 | 0.129148 | 0.018316 | 1 |
| Eif3d | 0.128942 | 0.12011 | 1 |
| Trim44 | 0.12879 | 0.024063 | 1 |
| A430093F15I | 0.12877 | 0.532107 | 1 |
| Vps8 | 0.128698 | 0.100676 | 1 |
| Ms4a6d | 0.128683 | 0.089636 | 1 |
| Fkrp | 0.128565 | 0.011145 | 1 |
| Phf20l1 | 0.128471 | 0.059852 | 1 |
| Elmsan1 | 0.128462 | 0.040693 | 1 |
| Chmp1b | 0.128418 | 0.048362 | 1 |
| Uvssa | 0.128387 | 0.005864 | 1 |
| Ndufb9 | 0.128322 | 0.125107 | 1 |
| Npepps | 0.128189 | 0.361137 | 1 |
| Snhg20 | 0.128165 | 0.057803 | 1 |
| Taf3 | 0.128116 | 0.283468 | 1 |
| Rnf187 | 0.128101 | 0.085408 | 1 |
| Gabarapl2 | 0.127966 | 0.015186 | 1 |
| Poglut1 | 0.127841 | 0.01262 | 1 |
| Zfp414 | 0.127551 | 0.163799 | 1 |
| Ucp2 | 0.127415 | 0.12016 | 1 |
| Spr | 0.127288 | 0.257476 | 1 |
| Gfer | 0.127161 | 0.223616 | 1 |
| Lars | 0.126853 | 0.116466 | 1 |
| Mycbp2 | 0.126761 | 0.046378 | 1 |

| | | | |
|---|---|---|---|
| Map2k3 | 0.126705 | 0.474805 | 1 |
| Slc4a7 | 0.126207 | 0.111428 | 1 |
| Phtf1 | 0.126174 | 0.607037 | 1 |
| 2310015A10I | 0.126121 | 0.004699 | 1 |
| Cntrl | 0.126018 | 0.432187 | 1 |
| BC147527 | 0.125677 | 0.131105 | 1 |
| Rhbdd2 | 0.125631 | 0.03162 | 1 |
| Ubl7 | 0.125363 | 0.085181 | 1 |
| 9930021J03F | 0.125324 | 0.153498 | 1 |
| Mtln | 0.125187 | 0.09165 | 1 |
| Cln6 | 0.12505 | 0.011454 | 1 |
| Tmem164 | 0.124989 | 0.099678 | 1 |
| Zc3h12d | 0.124726 | 0.007147 | 1 |
| F2r | 0.124619 | 0.149823 | 1 |
| Fuom | 0.12458 | 0.017201 | 1 |
| Herc3 | 0.124576 | 0.143736 | 1 |
| Gm47283 | 0.124534 | 0.063652 | 1 |
| Dcaf6 | 0.124497 | 0.060055 | 1 |
| Ksr1 | 0.124497 | 0.34046 | 1 |
| Hsp90aa1 | 0.124495 | 0.204078 | 1 |
| Yars | 0.124481 | 0.304057 | 1 |
| Pus10 | 0.124396 | 0.01534 | 1 |
| Ino80 | 0.124376 | 0.110205 | 1 |
| Ccr2 | 0.124329 | 0.068337 | 1 |
| Rc3h2 | 0.124295 | 0.168498 | 1 |
| Ggct | 0.124116 | 0.159829 | 1 |
| Vapa | 0.123968 | 0.254423 | 1 |
| Fnbp1 | 0.123962 | 0.158702 | 1 |
| Dmtf1 | 0.123946 | 0.484979 | 1 |
| Retreg1 | 0.123853 | 0.090035 | 1 |
| Exosc7 | 0.123793 | 0.31658 | 1 |
| Rps28 | 0.123679 | 0.019021 | 1 |
| Fbxo34 | 0.123627 | 0.009478 | 1 |
| Exoc6b | 0.123579 | 0.213042 | 1 |
| Lyrm9 | 0.123377 | 0.001356 | 1 |
| Cluh | 0.123339 | 0.005334 | 1 |
| Nol7 | 0.1233 | 0.099229 | 1 |
| Psme4 | 0.123237 | 0.139259 | 1 |
| Pdcd2 | 0.123221 | 0.088909 | 1 |
| Rad23b | 0.123196 | 0.309991 | 1 |
| Gyg | 0.12313 | 0.123213 | 1 |
| Pip4k2b | 0.122936 | 0.16164 | 1 |
| Stx16 | 0.12293 | 0.13658 | 1 |
| Prpf18 | 0.122911 | 0.015484 | 1 |
| Dnajb13 | 0.122895 | 0.04074 | 1 |
| Eno1 | 0.122776 | 0.110682 | 1 |
| Fam49a | 0.122744 | 0.074592 | 1 |
| Arhgef12 | 0.122584 | 0.011698 | 1 |

| | | | |
|---|---|---|---|
| Map3k8 | 0.12249 | 0.243622 | 1 |
| Slc35c2 | 0.122396 | 0.336486 | 1 |
| Insl6 | 0.122279 | 0.146212 | 1 |
| Inpp4a | 0.122008 | 0.239985 | 1 |
| Suclg1 | 0.121958 | 0.194026 | 1 |
| Gypc | 0.121859 | 0.042238 | 1 |
| Timm50 | 0.121856 | 0.103075 | 1 |
| Gm26887 | 0.12184 | 0.087311 | 1 |
| 1700028E10I | 0.121793 | 0.036092 | 1 |
| 1700123O20 | 0.12174 | 0.139656 | 1 |
| Als2 | 0.12146 | 0.055233 | 1 |
| Rnf149 | 0.121456 | 0.039291 | 1 |
| Rab19 | 0.121318 | 0.150029 | 1 |
| Oma1 | 0.121256 | 0.226501 | 1 |
| Rabggta | 0.121201 | 0.253162 | 1 |
| Nxpe3 | 0.121134 | 0.09461 | 1 |
| Phgdh | 0.121083 | 0.179901 | 1 |
| Atf2 | 0.121039 | 0.057955 | 1 |
| Uhrf1bp1l | 0.120982 | 0.001025 | 1 |
| 1700021A07I | 0.1209 | 0.017766 | 1 |
| 2310039H08 | 0.120831 | 0.063343 | 1 |
| Slc19a1 | 0.1208 | 0.014436 | 1 |
| Fbxl17 | 0.120742 | 0.042256 | 1 |
| Esf1 | 0.120708 | 0.261686 | 1 |
| Axl | 0.120678 | 0.005884 | 1 |
| Arl14ep | 0.120674 | 0.021069 | 1 |
| Ndufaf2 | 0.120397 | 0.050073 | 1 |
| Parp11 | 0.120309 | 0.36895 | 1 |
| Lancl1 | 0.120293 | 0.12864 | 1 |
| Uck2 | 0.120054 | 0.251989 | 1 |
| Slc25a5 | 0.120029 | 0.09421 | 1 |
| Psmg4 | 0.119974 | 0.195818 | 1 |
| Tmem154 | 0.11991 | 0.39062 | 1 |
| Ahi1 | 0.119796 | 0.220915 | 1 |
| Trmt11 | 0.119699 | 0.100219 | 1 |
| Rfx7 | 0.119691 | 0.500798 | 1 |
| Hps3 | 0.119639 | 0.096532 | 1 |
| Epb41 | 0.119522 | 0.394708 | 1 |
| Tlk2 | 0.119235 | 0.222413 | 1 |
| Ppa1 | 0.118842 | 0.072006 | 1 |
| Itpr2 | 0.118589 | 0.18557 | 1 |
| Mrpl42 | 0.118552 | 0.204274 | 1 |
| mt-Co3 | 0.118547 | 0.084637 | 1 |
| Phactr2 | 0.118544 | 0.373141 | 1 |
| Gem | 0.118526 | 0.889314 | 1 |
| Rnft1 | 0.118366 | 0.056767 | 1 |
| Oser1 | 0.118318 | 0.132048 | 1 |
| Mrpl2 | 0.118272 | 0.254803 | 1 |

| | | | |
|---|---|---|---|
| Fam71b | 0.118201 | 0.061947 | 1 |
| Itgav | 0.118098 | 0.154045 | 1 |
| Nufip2 | 0.118047 | 0.318344 | 1 |
| Hipk1 | 0.117944 | 0.162368 | 1 |
| Cyb5a | 0.117909 | 0.03062 | 1 |
| Tmem134 | 0.117885 | 0.161712 | 1 |
| Gna15 | 0.117882 | 0.091556 | 1 |
| Irak4 | 0.117736 | 0.068092 | 1 |
| Pgm1 | 0.117669 | 0.078218 | 1 |
| Pomp | 0.11765 | 0.19147 | 1 |
| Slc9a9 | 0.117612 | 0.35097 | 1 |
| Vps13b | 0.117561 | 0.067688 | 1 |
| Prps1 | 0.117512 | 0.215263 | 1 |
| Taf8 | 0.117448 | 0.193104 | 1 |
| Abcf3 | 0.117103 | 0.079888 | 1 |
| Nop14 | 0.117001 | 0.175166 | 1 |
| Gpr160 | 0.116672 | 0.085162 | 1 |
| 5430405H02 | 0.116575 | 0.096502 | 1 |
| Acot9 | 0.116337 | 0.154092 | 1 |
| Car5b | 0.116262 | 0.161255 | 1 |
| Gas2 | 0.116101 | 0.019312 | 1 |
| Utp11 | 0.11606 | 0.257327 | 1 |
| Riox2 | 0.116052 | 0.186881 | 1 |
| Bag4 | 0.115898 | 0.075942 | 1 |
| Rai1 | 0.115653 | 0.123833 | 1 |
| Gnl3 | 0.115502 | 0.345904 | 1 |
| Tbc1d15 | 0.115498 | 0.224667 | 1 |
| Ralgapa2 | 0.115452 | 0.256743 | 1 |
| Pex16 | 0.115397 | 0.352538 | 1 |
| Ahcyl1 | 0.115296 | 0.112025 | 1 |
| Ttpal | 0.115252 | 0.386067 | 1 |
| N4bp1 | 0.115126 | 0.032929 | 1 |
| Invs | 0.115087 | 0.209188 | 1 |
| Atg10 | 0.114953 | 0.331127 | 1 |
| Ccdc58 | 0.114929 | 0.023488 | 1 |
| Nol12 | 0.114851 | 0.203753 | 1 |
| Phip | 0.11463 | 0.129714 | 1 |
| Nmrk1 | 0.114586 | 0.079739 | 1 |
| Scyl3 | 0.114576 | 0.245725 | 1 |
| Mgst2 | 0.114506 | 0.153207 | 1 |
| Mzt2 | 0.11448 | 0.011551 | 1 |
| Hip1 | 0.114385 | 0.240794 | 1 |
| Lck | 0.114341 | 0.094879 | 1 |
| Tusc2 | 0.114313 | 0.056321 | 1 |
| Ttc1 | 0.114187 | 0.288513 | 1 |
| Spata5 | 0.113963 | 0.061221 | 1 |
| Larp1 | 0.11383 | 0.396027 | 1 |
| F8a | 0.113811 | 0.364689 | 1 |

| | | | |
|---|---|---|---|
| Clock | 0.113769 | 0.085311 | 1 |
| Hagh | 0.113565 | 0.190033 | 1 |
| Fam204a | 0.113278 | 0.383797 | 1 |
| 1600002K03I | 0.113209 | 0.039606 | 1 |
| Pih1d1 | 0.113044 | 0.201734 | 1 |
| B4galnt1 | 0.113008 | 0.294663 | 1 |
| Nif3l1 | 0.112979 | 0.083586 | 1 |
| Rpap2 | 0.11284 | 0.049412 | 1 |
| Cdk7 | 0.11283 | 0.141589 | 1 |
| Lpin2 | 0.112826 | 0.240628 | 1 |
| Ints11 | 0.112693 | 0.187097 | 1 |
| Ip6k1 | 0.112641 | 0.10662 | 1 |
| Zrsr2 | 0.11259 | 0.185784 | 1 |
| 1190007I07F | 0.11241 | 0.167513 | 1 |
| Hmg20b | 0.112402 | 0.150319 | 1 |
| Trim24 | 0.112372 | 0.434101 | 1 |
| Gde1 | 0.112355 | 0.206598 | 1 |
| Hs3st3b1 | 0.112166 | 0.014648 | 1 |
| Parp4 | 0.112044 | 0.154498 | 1 |
| Mtx2 | 0.111593 | 0.302847 | 1 |
| Cracr2a | 0.111578 | 0.3701 | 1 |
| Mesd | 0.111373 | 0.086812 | 1 |
| Snip1 | 0.111301 | 0.171514 | 1 |
| Tor3a | 0.111129 | 0.231636 | 1 |
| Mrps28 | 0.110955 | 0.146068 | 1 |
| Gimap1 | 0.110917 | 0.149439 | 1 |
| L3mbtl2 | 0.110714 | 0.201981 | 1 |
| Csnk1g1 | 0.110507 | 0.174525 | 1 |
| Ino80b | 0.110444 | 0.150353 | 1 |
| Tox4 | 0.110364 | 0.288871 | 1 |
| Clta | 0.110227 | 0.05031 | 1 |
| Picalm | 0.110175 | 0.43153 | 1 |
| Abhd4 | 0.110097 | 0.014548 | 1 |
| Prelid1 | 0.109837 | 0.209271 | 1 |
| Rwdd4a | 0.109713 | 0.151863 | 1 |
| Ptpn3 | 0.109655 | 0.075092 | 1 |
| Marveld2 | 0.109545 | 0.098515 | 1 |
| Lrif1 | 0.109515 | 0.143171 | 1 |
| Tubgcp2 | 0.109476 | 0.279601 | 1 |
| Dpy19l3 | 0.109275 | 0.116092 | 1 |
| Slc30a9 | 0.109195 | 0.291556 | 1 |
| Atf4 | 0.109168 | 0.489311 | 1 |
| AY036118 | 0.109144 | 0.334454 | 1 |
| Surf1 | 0.109098 | 0.008329 | 1 |
| Klhl36 | 0.108927 | 0.137664 | 1 |
| Ncoa2 | 0.108873 | 0.25721 | 1 |
| Gnl2 | 0.108866 | 0.191859 | 1 |
| Tsn | 0.108663 | 0.363163 | 1 |

| | | | |
|---|---|---|---|
| Tbrg1 | 0.108592 | 0.00989 | 1 |
| Tmem160 | 0.108265 | 0.052895 | 1 |
| Zcchc17 | 0.108243 | 0.454263 | 1 |
| Pim3 | 0.10822 | 0.035497 | 1 |
| Fbxl20 | 0.108201 | 0.016766 | 1 |
| Chmp2a | 0.108096 | 0.230123 | 1 |
| Zer1 | 0.108071 | 0.07764 | 1 |
| Alg8 | 0.108041 | 0.050425 | 1 |
| Cnppd1 | 0.107769 | 0.053303 | 1 |
| H2-T22 | 0.107734 | 0.210732 | 1 |
| Ppp1r9b | 0.107673 | 0.170211 | 1 |
| Arfrp1 | 0.107638 | 0.172293 | 1 |
| Glrx5 | 0.107632 | 0.004883 | 1 |
| Mtfmt | 0.107533 | 0.071915 | 1 |
| Ripk2 | 0.107507 | 0.09942 | 1 |
| Rngtt | 0.107465 | 0.293507 | 1 |
| Golgb1 | 0.107348 | 0.083849 | 1 |
| Tax1bp1 | 0.10733 | 0.021377 | 1 |
| Stk26 | 0.107247 | 0.451689 | 1 |
| Rc3h1 | 0.107231 | 0.100258 | 1 |
| Galk2 | 0.107227 | 0.120443 | 1 |
| Aup1 | 0.107103 | 0.165475 | 1 |
| Gramd1b | 0.106876 | 0.347496 | 1 |
| 2310061I04F | 0.106809 | 0.206515 | 1 |
| Rictor | 0.106644 | 0.374203 | 1 |
| Bcl7b | 0.106288 | 0.401648 | 1 |
| Phf12 | 0.106285 | 0.221602 | 1 |
| Arfgap1 | 0.106147 | 0.129485 | 1 |
| Zmpste24 | 0.105965 | 0.199473 | 1 |
| Gm2a | 0.105871 | 0.230191 | 1 |
| Maip1 | 0.105824 | 0.083029 | 1 |
| Rmdn1 | 0.105762 | 0.032057 | 1 |
| Sptlc1 | 0.105739 | 0.285059 | 1 |
| Phkg2 | 0.105733 | 0.058828 | 1 |
| Dalrd3 | 0.1057 | 0.046005 | 1 |
| Chd1 | 0.105528 | 0.072054 | 1 |
| Trim27 | 0.105318 | 0.100255 | 1 |
| Coa7 | 0.105276 | 0.055956 | 1 |
| Eif1a | 0.105269 | 0.094227 | 1 |
| Cd52 | 0.105263 | 0.158527 | 1 |
| Pgls | 0.105226 | 0.168323 | 1 |
| Pnpla7 | 0.105158 | 0.112111 | 1 |
| Rbck1 | 0.105115 | 0.209131 | 1 |
| Rnf157 | 0.105068 | 0.108577 | 1 |
| Spsb3 | 0.104884 | 0.073067 | 1 |
| Amn1 | 0.104865 | 0.156314 | 1 |
| St3gal1 | 0.104782 | 0.255241 | 1 |
| Csnk1a1 | 0.104685 | 0.326105 | 1 |

| | | | |
|---|---|---|---|
| Lig3 | 0.10457 | 0.067494 | 1 |
| Sil1 | 0.104541 | 0.149678 | 1 |
| Serpinb1a | 0.104278 | 0.067516 | 1 |
| Rarg | 0.10422 | 0.056558 | 1 |
| Gm39323 | 0.104212 | 0.107698 | 1 |
| Tmem64 | 0.104067 | 0.04211 | 1 |
| Haghl | 0.104049 | 0.067398 | 1 |
| Tbcb | 0.10392 | 0.16373 | 1 |
| Phospho2 | 0.103904 | 0.100097 | 1 |
| Gtf3c2 | 0.103824 | 0.170534 | 1 |
| Zfp593 | 0.103808 | 0.215724 | 1 |
| Lrrc8d | 0.103806 | 0.197336 | 1 |
| Aurkaip1 | 0.1038 | 0.193524 | 1 |
| Tex264 | 0.1038 | 0.460321 | 1 |
| Wipi2 | 0.103598 | 0.164356 | 1 |
| Sinhcaf | 0.103595 | 0.177342 | 1 |
| Socs3 | 0.103446 | 0.266902 | 1 |
| A330040F15I | 0.103336 | 0.480408 | 1 |
| Dnajc19 | 0.103297 | 0.410111 | 1 |
| Tor1aip2 | 0.103188 | 0.036931 | 1 |
| Rnf14 | 0.102958 | 0.686404 | 1 |
| Pcnx | 0.102834 | 0.237188 | 1 |
| Iars | 0.102507 | 0.33495 | 1 |
| Clptm1l | 0.102498 | 0.076398 | 1 |
| Mrrf | 0.102341 | 0.065372 | 1 |
| Ccnq | 0.102313 | 0.254161 | 1 |
| Xpr1 | 0.102273 | 0.136192 | 1 |
| Magohb | 0.102259 | 0.228074 | 1 |
| Lrrc59 | 0.102218 | 0.141718 | 1 |
| Snx15 | 0.102176 | 0.094586 | 1 |
| Creb1 | 0.102155 | 0.305885 | 1 |
| Ptpn7 | 0.102087 | 0.296579 | 1 |
| B4galt5 | 0.102077 | 0.302635 | 1 |
| Meis3 | 0.102063 | 0.04685 | 1 |
| Asf1a | 0.102041 | 0.126552 | 1 |
| Rbms1 | 0.101818 | 0.295606 | 1 |
| E330009J07F | 0.101717 | 0.036827 | 1 |
| Ankzf1 | 0.101533 | 0.212124 | 1 |
| Cnot11 | 0.101506 | 0.121918 | 1 |
| Kidins220 | 0.101493 | 0.395917 | 1 |
| Tigd2 | 0.101391 | 0.473978 | 1 |
| Commd6 | 0.101379 | 0.119566 | 1 |
| Cdc123 | 0.101284 | 0.335172 | 1 |
| Rab27a | 0.101141 | 0.34473 | 1 |
| Sv2c | 0.101103 | 0.071607 | 1 |
| Celsr1 | 0.100918 | 0.178309 | 1 |
| Emp1 | 0.100872 | 0.133841 | 1 |
| Wdr45b | 0.100823 | 0.564921 | 1 |

| | | | |
|---|---|---|---|
| Rxylt1 | 0.100777 | 0.060628 | 1 |
| Ddx27 | 0.100547 | 0.280889 | 1 |
| Med30 | 0.100392 | 0.159455 | 1 |
| Nolc1 | 0.100376 | 0.450983 | 1 |
| Zfp64 | 0.100338 | 0.031896 | 1 |
| Cmss1 | 0.100328 | 0.248546 | 1 |
| B3gntl1 | 0.100293 | 0.174021 | 1 |
| Actr6 | 0.100208 | 0.021096 | 1 |
| Zxdc | 0.100107 | 0.113841 | 1 |
| Trps1 | 0.100033 | 0.407791 | 1 |
| Tfrc | 0.099987 | 0.109761 | 1 |
| A930037H05 | 0.099965 | 0.115587 | 1 |
| 2300009A05 | 0.099915 | 0.046087 | 1 |
| Poc5 | 0.099888 | 0.102666 | 1 |
| Wbp11 | 0.099839 | 0.452375 | 1 |
| Pank2 | 0.099631 | 0.134469 | 1 |
| Slc25a20 | 0.099577 | 0.087769 | 1 |
| Spg7 | 0.099493 | 0.063107 | 1 |
| Idh3b | 0.09946 | 0.404298 | 1 |
| Dnajb9 | 0.099392 | 0.34842 | 1 |
| Cdk13 | 0.099107 | 0.090591 | 1 |
| Mettl16 | 0.099036 | 0.726141 | 1 |
| Rnh1 | 0.098941 | 0.319822 | 1 |
| F730043M19 | 0.098868 | 0.778877 | 1 |
| Smyd5 | 0.098856 | 0.018861 | 1 |
| Fam210b | 0.09884 | 0.465006 | 1 |
| Phf6 | 0.09879 | 0.319896 | 1 |
| Nbdy | 0.098778 | 0.067682 | 1 |
| Plekha3 | 0.098673 | 0.173993 | 1 |
| Arl1 | 0.09867 | 0.440321 | 1 |
| Aak1 | 0.098657 | 0.377611 | 1 |
| F2rl2 | 0.098641 | 0.422105 | 1 |
| Dcaf5 | 0.09861 | 0.13812 | 1 |
| Ccni | 0.098607 | 0.07135 | 1 |
| Nr4a1 | 0.098587 | 0.039251 | 1 |
| Dnase1l1 | 0.098506 | 0.038326 | 1 |
| Polr3b | 0.098471 | 0.088524 | 1 |
| Sgsh | 0.09843 | 0.138558 | 1 |
| Thap7 | 0.098219 | 0.309691 | 1 |
| Furin | 0.098208 | 0.054773 | 1 |
| Stat5a | 0.098198 | 0.643651 | 1 |
| Atp10a | 0.098162 | 0.042968 | 1 |
| Itpkb | 0.098149 | 0.337491 | 1 |
| Ndufs3 | 0.09814 | 0.172903 | 1 |
| Spata5l1 | 0.098051 | 0.097281 | 1 |
| Dtd2 | 0.097887 | 0.118389 | 1 |
| Tpcn1 | 0.097871 | 0.072491 | 1 |
| Apba3 | 0.097844 | 0.116981 | 1 |

| | | | |
|---|---|---|---|
| Mia2 | 0.097828 | 0.412808 | 1 |
| N4bp2l2 | 0.097807 | 0.208916 | 1 |
| Nek6 | 0.097753 | 0.262955 | 1 |
| Tob2 | 0.097517 | 0.354372 | 1 |
| Ccnc | 0.097515 | 0.249863 | 1 |
| Tmem203 | 0.097468 | 0.068025 | 1 |
| Faf2 | 0.097434 | 0.13805 | 1 |
| Alkbh3 | 0.097407 | 0.28998 | 1 |
| Nanp | 0.09732 | 0.224092 | 1 |
| Zfp110 | 0.097267 | 0.146645 | 1 |
| Slc37a3 | 0.097067 | 0.211908 | 1 |
| Srek1ip1 | 0.097015 | 0.045563 | 1 |
| Mfap1b | 0.096928 | 0.41222 | 1 |
| Trerf1 | 0.096895 | 0.423473 | 1 |
| Api5 | 0.096892 | 0.520867 | 1 |
| Setd1a | 0.096869 | 0.220097 | 1 |
| Ctnna1 | 0.096842 | 0.170604 | 1 |
| Crybg1 | 0.09668 | 0.469578 | 1 |
| Kri1 | 0.096679 | 0.485733 | 1 |
| Angptl4 | 0.096548 | 0.114538 | 1 |
| Ears2 | 0.096547 | 0.088034 | 1 |
| Cdk9 | 0.096318 | 0.362135 | 1 |
| Runx2os1 | 0.096306 | 0.065454 | 1 |
| Gba | 0.096166 | 0.439497 | 1 |
| 2410022M11 | 0.095906 | 0.081311 | 1 |
| Tnip2 | 0.095709 | 0.010831 | 1 |
| Pdk1 | 0.095678 | 0.251779 | 1 |
| Ndufaf1 | 0.095608 | 0.104285 | 1 |
| Taz | 0.095568 | 0.105225 | 1 |
| Prkar1a | 0.095466 | 0.274933 | 1 |
| Rbm41 | 0.095374 | 0.332824 | 1 |
| Ptcd3 | 0.095307 | 0.195652 | 1 |
| Mrpl38 | 0.095244 | 0.236209 | 1 |
| Ccnh | 0.095166 | 0.157164 | 1 |
| 6030458C11 | 0.095149 | 0.347907 | 1 |
| Bicdl1 | 0.095064 | 0.047724 | 1 |
| Ddx18 | 0.095045 | 0.040783 | 1 |
| Bcl7c | 0.094998 | 0.585822 | 1 |
| Suco | 0.09497 | 0.52336 | 1 |
| Prxl2c | 0.094936 | 0.016866 | 1 |
| Stk25 | 0.094922 | 0.087455 | 1 |
| Snu13 | 0.094878 | 0.405025 | 1 |
| Cct3 | 0.094749 | 0.337793 | 1 |
| Rfxap | 0.0946 | 0.230421 | 1 |
| Plcxd2 | 0.094576 | 0.275434 | 1 |
| Ndufa5 | 0.094545 | 0.183392 | 1 |
| Ndufs6 | 0.094542 | 0.309277 | 1 |
| Zbtb2 | 0.094497 | 0.221481 | 1 |

| | | | |
|---|---|---|---|
| Uap1l1 | 0.094495 | 0.098444 | 1 |
| Rabepk | 0.094276 | 0.514753 | 1 |
| CAAA011183 | 0.094272 | 0.382637 | 1 |
| Fosb | 0.094249 | 0.501015 | 1 |
| Tbc1d9b | 0.09422 | 0.495927 | 1 |
| Aacs | 0.094206 | 0.107734 | 1 |
| Abi3 | 0.094127 | 0.103477 | 1 |
| Mrpl46 | 0.093998 | 0.330496 | 1 |
| Ccdc130 | 0.093964 | 0.07467 | 1 |
| Evl | 0.093889 | 0.188784 | 1 |
| Ercc6 | 0.093874 | 0.056906 | 1 |
| Zfp280c | 0.093851 | 0.230072 | 1 |
| Cfap36 | 0.093742 | 0.5086 | 1 |
| Nlrc5 | 0.093736 | 0.767867 | 1 |
| Lrrc45 | 0.093733 | 0.060568 | 1 |
| Unc50 | 0.093666 | 0.401882 | 1 |
| Zdhhc3 | 0.093599 | 0.028956 | 1 |
| Mettl23 | 0.093434 | 0.263732 | 1 |
| 1110032A03 | 0.093391 | 0.025224 | 1 |
| Ptpmt1 | 0.093345 | 0.222008 | 1 |
| Usp53 | 0.093269 | 0.210426 | 1 |
| Irf2bp2 | 0.093214 | 0.28578 | 1 |
| Ralgapb | 0.093171 | 0.136174 | 1 |
| Nol11 | 0.093164 | 0.188337 | 1 |
| Cyb5r1 | 0.093019 | 0.104679 | 1 |
| Stimate | 0.093009 | 0.008765 | 1 |
| Mrpl45 | 0.092992 | 0.023234 | 1 |
| Ypel5 | 0.092964 | 0.207728 | 1 |
| Phb | 0.092884 | 0.321988 | 1 |
| Eif2ak1 | 0.092759 | 0.256028 | 1 |
| Fbxl3 | 0.092732 | 0.194298 | 1 |
| Dgcr2 | 0.09266 | 0.188564 | 1 |
| Hsd17b12 | 0.092595 | 0.705857 | 1 |
| Ddx3x | 0.092547 | 0.139502 | 1 |
| Casc3 | 0.092518 | 0.320181 | 1 |
| Npm3 | 0.092355 | 0.295199 | 1 |
| Micu3 | 0.092178 | 0.264134 | 1 |
| Sclt1 | 0.092146 | 0.313113 | 1 |
| Rwdd1 | 0.092071 | 0.521686 | 1 |
| Ubac1 | 0.091975 | 0.152942 | 1 |
| Mtif2 | 0.091945 | 0.243473 | 1 |
| Sumf1 | 0.09189 | 0.140527 | 1 |
| Ndufb6 | 0.091879 | 0.431372 | 1 |
| Trmt112 | 0.091845 | 0.222747 | 1 |
| Guk1 | 0.091802 | 0.302387 | 1 |
| Tnrc18 | 0.091779 | 0.209402 | 1 |
| Rab11fip2 | 0.091737 | 0.130786 | 1 |
| Lrrc28 | 0.091719 | 0.089629 | 1 |

| | | | |
|---|---|---|---|
| Mbtps1 | 0.091714 | 0.196474 | 1 |
| Ccl3 | 0.091658 | 0.006486 | 1 |
| Mtg2 | 0.091607 | 0.38264 | 1 |
| Abcc1 | 0.091528 | 0.106 | 1 |
| Zfp664 | 0.09146 | 0.19394 | 1 |
| Stam2 | 0.091357 | 0.167007 | 1 |
| Arhgap27os3 | 0.091179 | 0.442215 | 1 |
| Lat | 0.091042 | 0.19595 | 1 |
| Nufip1 | 0.090944 | 0.273025 | 1 |
| Itgb3bp | 0.090912 | 0.22345 | 1 |
| Pmm2 | 0.090814 | 0.228468 | 1 |
| Polr3h | 0.090747 | 0.376646 | 1 |
| Parp12 | 0.09073 | 0.15108 | 1 |
| Pdha1 | 0.090673 | 0.228323 | 1 |
| Mboat7 | 0.090638 | 0.502515 | 1 |
| Ptpn9 | 0.090623 | 0.238708 | 1 |
| Tle5 | 0.090621 | 0.154892 | 1 |
| Ndufab1 | 0.090593 | 0.3402 | 1 |
| Rnf146 | 0.09042 | 0.174897 | 1 |
| Ndufa12 | 0.090416 | 0.410275 | 1 |
| Dph3 | 0.090345 | 0.115842 | 1 |
| Bcl6 | 0.090337 | 0.242886 | 1 |
| Bola3 | 0.090047 | 0.179742 | 1 |
| Ring1 | 0.090011 | 0.369175 | 1 |
| Mrpl14 | 0.090001 | 0.291744 | 1 |
| 5530601H04 | 0.089685 | 0.135545 | 1 |
| Dok1 | 0.089661 | 0.061009 | 1 |
| 2700062C07 | 0.089485 | 0.207133 | 1 |
| Noa1 | 0.089423 | 0.333126 | 1 |
| N4bp2l1 | 0.089412 | 0.566503 | 1 |
| Srsf5 | 0.08934 | 0.33735 | 1 |
| Fahd2a | 0.089333 | 0.188176 | 1 |
| Fam120b | 0.089316 | 0.338789 | 1 |
| Birc3 | 0.089304 | 0.022878 | 1 |
| Alg9 | 0.08929 | 0.059485 | 1 |
| Txndc9 | 0.089274 | 0.297431 | 1 |
| Psmf1 | 0.089193 | 0.254578 | 1 |
| Ankrd37 | 0.089125 | 0.321837 | 1 |
| Ints14 | 0.089082 | 0.275075 | 1 |
| Tbc1d14 | 0.089071 | 0.353905 | 1 |
| Klf13 | 0.088946 | 0.192267 | 1 |
| Tpgs2 | 0.088869 | 0.267696 | 1 |
| Spcs2 | 0.088823 | 0.216514 | 1 |
| Nfat5 | 0.088768 | 0.211476 | 1 |
| Psma4 | 0.088573 | 0.274401 | 1 |
| Taf10 | 0.088522 | 0.543394 | 1 |
| Tmem268 | 0.088483 | 0.162333 | 1 |
| Pold4 | 0.088404 | 0.004589 | 1 |

| | | | |
|---|---|---|---|
| Tmem42 | 0.088362 | 0.108811 | 1 |
| Mrpl24 | 0.088342 | 0.27294 | 1 |
| Raet1e | 0.088313 | 0.350402 | 1 |
| Slco4a1 | 0.088306 | 0.045537 | 1 |
| Dus3l | 0.088245 | 0.469743 | 1 |
| Yipf4 | 0.088233 | 0.31776 | 1 |
| Cars | 0.088231 | 0.227385 | 1 |
| Wdr45 | 0.088199 | 0.23491 | 1 |
| Dars | 0.088111 | 0.544914 | 1 |
| Socs4 | 0.088024 | 0.322877 | 1 |
| Stambpl1 | 0.087921 | 0.409842 | 1 |
| Gne | 0.087843 | 0.665768 | 1 |
| Gtf3c1 | 0.087692 | 0.195077 | 1 |
| Tmem14c | 0.087579 | 0.264981 | 1 |
| Poldip2 | 0.087558 | 0.092695 | 1 |
| Rlim | 0.087491 | 0.640442 | 1 |
| Kdm4b | 0.087473 | 0.094972 | 1 |
| Psma6 | 0.087459 | 0.108683 | 1 |
| Proser1 | 0.087453 | 0.15771 | 1 |
| Cyc1 | 0.087415 | 0.114358 | 1 |
| A430005L14I | 0.087399 | 0.170821 | 1 |
| Slc2a3 | 0.087219 | 0.417611 | 1 |
| Atxn7 | 0.087181 | 0.177948 | 1 |
| Ccdc86 | 0.087102 | 0.214054 | 1 |
| Rspry1 | 0.087085 | 0.399621 | 1 |
| Bfar | 0.086982 | 0.432022 | 1 |
| Hdac8 | 0.086948 | 0.327817 | 1 |
| A930024E05I | 0.086894 | 0.19224 | 1 |
| Hmgxb3 | 0.086877 | 0.111802 | 1 |
| Snx5 | 0.086825 | 0.360518 | 1 |
| Bet1l | 0.086812 | 0.135237 | 1 |
| Oxld1 | 0.086808 | 0.127686 | 1 |
| Rae1 | 0.0868 | 0.268558 | 1 |
| Abhd2 | 0.086737 | 0.330131 | 1 |
| Map2k4 | 0.086678 | 0.460228 | 1 |
| Rpp14 | 0.086509 | 0.093367 | 1 |
| Dpp4 | 0.086463 | 0.368177 | 1 |
| Lst1 | 0.086432 | 0.080773 | 1 |
| Rad18 | 0.086398 | 0.43844 | 1 |
| Ndst1 | 0.08631 | 0.01533 | 1 |
| Tgif1 | 0.086178 | 0.35626 | 1 |
| Acaca | 0.086081 | 0.257008 | 1 |
| Ktn1 | 0.086072 | 0.25484 | 1 |
| Zfp810 | 0.085978 | 0.535302 | 1 |
| Adam9 | 0.085975 | 0.289264 | 1 |
| Ccl25 | 0.08597 | 0.189652 | 1 |
| Carnmt1 | 0.085885 | 0.141811 | 1 |
| Mterf3 | 0.085844 | 0.016856 | 1 |

| | | | |
|---|---|---|---|
| Lrpprc | 0.085809 | 0.160297 | 1 |
| Cdc37l1 | 0.085703 | 0.164962 | 1 |
| Taf11 | 0.085662 | 0.204001 | 1 |
| Slc7a6 | 0.08566 | 0.348082 | 1 |
| Pmpcb | 0.085636 | 0.295165 | 1 |
| C87436 | 0.085616 | 0.195025 | 1 |
| Rab18 | 0.085542 | 0.57474 | 1 |
| Khnyn | 0.085421 | 0.720766 | 1 |
| Lap3 | 0.085381 | 0.088864 | 1 |
| Mtap | 0.085313 | 0.118799 | 1 |
| Pknox1 | 0.085262 | 0.240323 | 1 |
| Tor2a | 0.085137 | 0.072361 | 1 |
| Cdv3 | 0.085132 | 0.557449 | 1 |
| Wdr73 | 0.085113 | 0.258143 | 1 |
| Chd1l | 0.085061 | 0.084473 | 1 |
| Ydjc | 0.084898 | 0.0521 | 1 |
| Rcor3 | 0.084893 | 0.031681 | 1 |
| Ctss | 0.08483 | 0.004677 | 1 |
| Sec24c | 0.084793 | 0.571728 | 1 |
| Phf11a | 0.084707 | 0.933524 | 1 |
| Mrpl22 | 0.084707 | 0.402304 | 1 |
| Nabp1 | 0.084707 | 0.135822 | 1 |
| Ggps1 | 0.084618 | 0.498966 | 1 |
| Iscu | 0.084592 | 0.276923 | 1 |
| Cep164 | 0.084499 | 0.224667 | 1 |
| Gnl3l | 0.084481 | 0.260441 | 1 |
| Gm4316 | 0.084474 | 0.235041 | 1 |
| Mapk9 | 0.084458 | 0.130307 | 1 |
| Hsh2d | 0.084447 | 0.299727 | 1 |
| Il1rap | 0.084377 | 0.294707 | 1 |
| Smco4 | 0.084294 | 0.219668 | 1 |
| Galt | 0.084242 | 0.32755 | 1 |
| Creb3l2 | 0.084199 | 0.072032 | 1 |
| Zfp408 | 0.084158 | 0.197031 | 1 |
| Rnf7 | 0.084144 | 0.472703 | 1 |
| Mib1 | 0.0841 | 0.096896 | 1 |
| Arfgef1 | 0.084087 | 0.262661 | 1 |
| Zfp87 | 0.084041 | 0.1765 | 1 |
| Padi2 | 0.08401 | 0.019602 | 1 |
| Cabin1 | 0.083968 | 0.405944 | 1 |
| Nit2 | 0.083835 | 0.416377 | 1 |
| Emg1 | 0.083796 | 0.472541 | 1 |
| Uqcrq | 0.083748 | 0.108007 | 1 |
| Ubqln1 | 0.083732 | 0.265448 | 1 |
| Gtf2b | 0.083611 | 0.690072 | 1 |
| Zdhhc12 | 0.083601 | 0.113879 | 1 |
| Zdhhc17 | 0.083526 | 0.04902 | 1 |
| Ecd | 0.083446 | 0.555971 | 1 |

| | | | |
|---|---|---|---|
| 5430416N02 | 0.083427 | 0.246862 | 1 |
| Slc25a16 | 0.083399 | 0.17259 | 1 |
| Ipo4 | 0.083245 | 0.1245 | 1 |
| Aff4 | 0.083182 | 0.079191 | 1 |
| Clns1a | 0.083154 | 0.206129 | 1 |
| Prkdc | 0.083147 | 0.170188 | 1 |
| Gm5547 | 0.082853 | 0.153157 | 1 |
| Atp5f1 | 0.082754 | 0.403911 | 1 |
| Morf4l2 | 0.082743 | 0.306838 | 1 |
| Zfp36l1 | 0.08272 | 0.006114 | 1 |
| Eid1 | 0.082689 | 0.146905 | 1 |
| Trmt6 | 0.082625 | 0.178334 | 1 |
| Pfkfb3 | 0.082498 | 0.299762 | 1 |
| Ercc5 | 0.082488 | 0.262032 | 1 |
| Gm47015 | 0.082471 | 0.179047 | 1 |
| Gmfb | 0.082457 | 0.266021 | 1 |
| AW549877 | 0.082446 | 0.216585 | 1 |
| Itpk1 | 0.08243 | 0.216631 | 1 |
| Mettl22 | 0.082396 | 0.138153 | 1 |
| Sema4d | 0.082354 | 0.156139 | 1 |
| Zfp141 | 0.082274 | 0.147316 | 1 |
| Smpd4 | 0.082243 | 0.143643 | 1 |
| Usp45 | 0.082212 | 0.235974 | 1 |
| Rac1 | 0.082035 | 0.360148 | 1 |
| Inpp5k | 0.081957 | 0.108165 | 1 |
| Gmppa | 0.081941 | 0.351579 | 1 |
| Slc16a6 | 0.081811 | 0.446346 | 1 |
| Nsmce3 | 0.081755 | 0.306263 | 1 |
| Rusc1 | 0.081744 | 0.1297 | 1 |
| Med28 | 0.081693 | 0.482103 | 1 |
| Paox | 0.081675 | 0.133755 | 1 |
| Tfdp2 | 0.081596 | 0.279648 | 1 |
| Swt1 | 0.081545 | 0.183773 | 1 |
| Gpaa1 | 0.081531 | 0.345797 | 1 |
| Dpy19l4 | 0.081506 | 0.268392 | 1 |
| Maf1 | 0.08147 | 0.123276 | 1 |
| Tmem184b | 0.081432 | 0.317973 | 1 |
| Yars2 | 0.081318 | 0.050434 | 1 |
| Oas1c | 0.081313 | 0.474155 | 1 |
| Hltf | 0.08123 | 0.132279 | 1 |
| Mthfsl | 0.081189 | 0.409147 | 1 |
| Arhgef6 | 0.081159 | 0.447071 | 1 |
| Gon4l | 0.081142 | 0.792028 | 1 |
| Aldh18a1 | 0.081067 | 0.528509 | 1 |
| Gstcd | 0.081053 | 0.22976 | 1 |
| Anapc13 | 0.081047 | 0.487687 | 1 |
| Dnajc4 | 0.081015 | 0.272278 | 1 |
| Gimap8 | 0.081003 | 0.324012 | 1 |

| | | | |
|---|---|---|---|
| Kdm5d | 0.080989 | 0.323749 | 1 |
| Aptx | 0.080876 | 0.790503 | 1 |
| Imp4 | 0.08087 | 0.662005 | 1 |
| Vps39 | 0.08087 | 0.350102 | 1 |
| Sgf29 | 0.080837 | 0.331547 | 1 |
| Tpp2 | 0.080794 | 0.740195 | 1 |
| Zswim7 | 0.080776 | 0.216567 | 1 |
| Mrps5 | 0.080761 | 0.535684 | 1 |
| Dclre1b | 0.080649 | 0.33518 | 1 |
| Xab2 | 0.080637 | 0.740634 | 1 |
| Wsb1 | 0.080628 | 0.710089 | 1 |
| Ccnk | 0.080604 | 0.105764 | 1 |
| Med9 | 0.080603 | 0.525644 | 1 |
| Rnf139 | 0.080559 | 0.127598 | 1 |
| Cyb5r4 | 0.080536 | 0.262277 | 1 |
| Nme1 | 0.080459 | 0.203157 | 1 |
| Rmnd5a | 0.080423 | 0.100774 | 1 |
| Zfp644 | 0.080379 | 0.193033 | 1 |
| Fcf1 | 0.080363 | 0.207347 | 1 |
| Rnf113a2 | 0.080353 | 0.077468 | 1 |
| Gm36551 | 0.080328 | 0.124467 | 1 |
| Ndufb3 | 0.080228 | 0.53126 | 1 |
| Calhm6 | 0.080203 | 0.57629 | 1 |
| Cmtr1 | 0.080157 | 0.075269 | 1 |
| Mcur1 | 0.080119 | 0.019381 | 1 |
| Epas1 | 0.080081 | 0.190357 | 1 |
| Per1 | 0.07999 | 0.149938 | 1 |
| Kbtbd3 | 0.079886 | 0.617829 | 1 |
| Slc25a22 | 0.079833 | 0.477211 | 1 |
| Rrn3 | 0.0798 | 0.192234 | 1 |
| 1700017B05 | 0.079774 | 0.350242 | 1 |
| Rab28 | 0.079759 | 0.208037 | 1 |
| Nbas | 0.079627 | 0.299682 | 1 |
| Hlcs | 0.079533 | 0.060805 | 1 |
| Brms1 | 0.079405 | 0.268121 | 1 |
| Dtd1 | 0.079376 | 0.115919 | 1 |
| Hipk3 | 0.079367 | 0.281353 | 1 |
| Dhrs4 | 0.079362 | 0.19191 | 1 |
| Dad1 | 0.079334 | 0.203846 | 1 |
| Stat5b | 0.079325 | 0.642643 | 1 |
| Rgcc | 0.079144 | 0.262125 | 1 |
| AC113595.1 | 0.079108 | 0.148108 | 1 |
| Slc46a3 | 0.079056 | 0.371041 | 1 |
| Timm13 | 0.078932 | 0.542485 | 1 |
| Tmed3 | 0.078927 | 0.046133 | 1 |
| Herc4 | 0.07885 | 0.723434 | 1 |
| Nin | 0.078814 | 0.178516 | 1 |
| Vcpkmt | 0.078802 | 0.143431 | 1 |

| | | | |
|---|---|---|---|
| Trim11 | 0.078738 | 0.181147 | 1 |
| Micos13 | 0.078709 | 0.486747 | 1 |
| Dubr | 0.078637 | 0.383438 | 1 |
| A430088P11 | 0.07862 | 0.260809 | 1 |
| Setdb1 | 0.078555 | 0.257019 | 1 |
| Decr1 | 0.078536 | 0.303828 | 1 |
| Drosha | 0.078517 | 0.134731 | 1 |
| Pcid2 | 0.0785 | 0.53497 | 1 |
| Slc39a4 | 0.078471 | 0.117733 | 1 |
| Commd7 | 0.078308 | 0.30882 | 1 |
| Khdc4 | 0.078219 | 0.372217 | 1 |
| Eif4enif1 | 0.078179 | 0.291274 | 1 |
| Exosc9 | 0.078153 | 0.527597 | 1 |
| Ppfibp1 | 0.078134 | 0.104757 | 1 |
| Polr2m | 0.078025 | 0.046265 | 1 |
| Slc7a6os | 0.078012 | 0.406612 | 1 |
| Mecr | 0.07796 | 0.245852 | 1 |
| Eef2 | 0.07794 | 0.202282 | 1 |
| Impact | 0.077939 | 0.216076 | 1 |
| Nfe2l1 | 0.07788 | 0.287609 | 1 |
| Prrc2b | 0.077844 | 0.518744 | 1 |
| Ercc6l2 | 0.077831 | 0.704727 | 1 |
| Cdkn1b | 0.077828 | 0.676358 | 1 |
| Hsd17b10 | 0.077781 | 0.374325 | 1 |
| Fbxw11 | 0.077773 | 0.284535 | 1 |
| Rps19bp1 | 0.077729 | 0.359816 | 1 |
| Gna13 | 0.077689 | 0.333728 | 1 |
| Pex3 | 0.077673 | 0.792455 | 1 |
| Taf1c | 0.077632 | 0.317257 | 1 |
| Mrpl30 | 0.077628 | 0.398748 | 1 |
| Uqcrfs1 | 0.077547 | 0.38912 | 1 |
| Slc25a39 | 0.077543 | 0.274478 | 1 |
| Wars | 0.077497 | 0.144086 | 1 |
| Srp9 | 0.07749 | 0.154787 | 1 |
| Lamc1 | 0.077488 | 0.189913 | 1 |
| Aggf1 | 0.077464 | 0.611272 | 1 |
| Vwa5a | 0.077455 | 0.150781 | 1 |
| Mettl1 | 0.077337 | 0.056586 | 1 |
| Wapl | 0.07731 | 0.350505 | 1 |
| 2310001H17 | 0.077232 | 0.411884 | 1 |
| Nfatc1 | 0.077226 | 0.419441 | 1 |
| Map3k14 | 0.077202 | 0.298141 | 1 |
| Psmd4 | 0.077173 | 0.530149 | 1 |
| Odr4 | 0.077125 | 0.313624 | 1 |
| Gm6034 | 0.077093 | 0.089467 | 1 |
| Tcta | 0.077077 | 0.091124 | 1 |
| Klhl12 | 0.077068 | 0.246744 | 1 |
| Mpp7 | 0.077009 | 0.308702 | 1 |

| | | | |
|---|---|---|---|
| Dhodh | 0.076929 | 0.282204 | 1 |
| Eef1akmt4 | 0.076916 | 0.129309 | 1 |
| Chchd5 | 0.076902 | 0.16003 | 1 |
| Ppp1r12c | 0.076781 | 0.464026 | 1 |
| Deaf1 | 0.076704 | 0.306494 | 1 |
| Apool | 0.076521 | 0.550803 | 1 |
| Insyn2b | 0.076513 | 0.906967 | 1 |
| Ttc5 | 0.0765 | 0.411025 | 1 |
| Yif1b | 0.076496 | 0.11559 | 1 |
| CAAA0114731 | 0.076463 | 0.451875 | 1 |
| Gm44148 | 0.076418 | 0.378229 | 1 |
| Mrpl13 | 0.076288 | 0.178486 | 1 |
| Pdss2 | 0.076269 | 0.494538 | 1 |
| Mfsd8 | 0.076247 | 0.108828 | 1 |
| Ascc3 | 0.076233 | 0.554839 | 1 |
| Zfp995 | 0.076154 | 0.069595 | 1 |
| 9130401M01 | 0.07606 | 0.133457 | 1 |
| Cd3eap | 0.075899 | 0.24953 | 1 |
| Wdr75 | 0.075897 | 0.147927 | 1 |
| Kctd17 | 0.07589 | 0.318023 | 1 |
| Sirt6 | 0.075877 | 0.247952 | 1 |
| Ctcf | 0.075866 | 0.319693 | 1 |
| Mta3 | 0.075805 | 0.092527 | 1 |
| Ski | 0.075804 | 0.368126 | 1 |
| Atf6b | 0.075771 | 0.72664 | 1 |
| Cbr4 | 0.075683 | 0.161521 | 1 |
| Vezt | 0.075532 | 0.147907 | 1 |
| Elp5 | 0.075409 | 0.407171 | 1 |
| 2610507B11 | 0.075327 | 0.221556 | 1 |
| Slc35a1 | 0.075189 | 0.304135 | 1 |
| Mapkap1 | 0.075162 | 0.318325 | 1 |
| Aldh7a1 | 0.07509 | 0.289363 | 1 |
| Zbtb4 | 0.075007 | 0.365498 | 1 |
| Yae1d1 | 0.074987 | 0.26839 | 1 |
| Ammecr1 | 0.074861 | 0.318192 | 1 |
| B3galt6 | 0.074782 | 0.239503 | 1 |
| Ints12 | 0.074778 | 0.059706 | 1 |
| Zfp318 | 0.074678 | 0.078562 | 1 |
| Ctso | 0.074668 | 0.218849 | 1 |
| Eif2b3 | 0.074568 | 0.211332 | 1 |
| Osgin1 | 0.074467 | 0.013373 | 1 |
| 1110065P20 | 0.074452 | 0.13125 | 1 |
| Aimp2 | 0.074446 | 0.513731 | 1 |
| Psmg2 | 0.074444 | 0.523764 | 1 |
| Slc25a3 | 0.074396 | 0.527685 | 1 |
| Nsun3 | 0.074377 | 0.3109 | 1 |
| Nlk | 0.07436 | 0.469889 | 1 |
| Pgghg | 0.074312 | 0.144775 | 1 |

| | | | |
|---|---|---|---|
| Atp6ap2 | 0.074228 | 0.735367 | 1 |
| Blcap | 0.074225 | 0.247196 | 1 |
| Adss | 0.074155 | 0.379142 | 1 |
| Rab9 | 0.074143 | 0.19924 | 1 |
| Mrps10 | 0.074085 | 0.216867 | 1 |
| Cnih4 | 0.074055 | 0.305801 | 1 |
| Acat1 | 0.074032 | 0.303108 | 1 |
| Krcc1 | 0.073959 | 0.354015 | 1 |
| Rbm25 | 0.073933 | 0.692446 | 1 |
| Cinp | 0.073933 | 0.234933 | 1 |
| Psd4 | 0.073901 | 0.428147 | 1 |
| Casp6 | 0.073848 | 0.288111 | 1 |
| Malsu1 | 0.073808 | 0.206995 | 1 |
| 2610002M06 | 0.073765 | 0.256265 | 1 |
| Pmepa1 | 0.073751 | 0.30669 | 1 |
| Ttll12 | 0.073707 | 0.452621 | 1 |
| Ofd1 | 0.073606 | 0.183851 | 1 |
| Pikfyve | 0.073577 | 0.299318 | 1 |
| Srp14 | 0.07354 | 0.425618 | 1 |
| Txndc12 | 0.073539 | 0.250702 | 1 |
| Pnpo | 0.073453 | 0.072766 | 1 |
| Slc43a2 | 0.073433 | 0.080054 | 1 |
| Anks3 | 0.073424 | 0.092136 | 1 |
| Ascc1 | 0.073264 | 0.930886 | 1 |
| Tstd3 | 0.073102 | 0.125575 | 1 |
| Fbxw2 | 0.073059 | 0.788544 | 1 |
| 2900097C17 | 0.073014 | 0.255024 | 1 |
| Dedd2 | 0.073012 | 0.127661 | 1 |
| Gm17745 | 0.073008 | 0.277029 | 1 |
| Exosc1 | 0.072924 | 0.418306 | 1 |
| Tomm40 | 0.072904 | 0.745822 | 1 |
| Krtcap2 | 0.072833 | 0.361836 | 1 |
| Itfg1 | 0.072629 | 0.112657 | 1 |
| Rraga | 0.072607 | 0.341792 | 1 |
| Rbis | 0.072507 | 0.297736 | 1 |
| Riox1 | 0.072471 | 0.158522 | 1 |
| Notch2 | 0.072383 | 0.600642 | 1 |
| BC005537 | 0.072347 | 0.235538 | 1 |
| Dpm2 | 0.072273 | 0.298452 | 1 |
| Ap3d1 | 0.072231 | 0.28811 | 1 |
| Acp5 | 0.07222 | 0.615499 | 1 |
| Xylt2 | 0.072215 | 0.530565 | 1 |
| Prrc2c | 0.072023 | 0.831571 | 1 |
| Vapb | 0.072022 | 0.352664 | 1 |
| Polr2e | 0.071876 | 0.733652 | 1 |
| Ppme1 | 0.071872 | 0.135419 | 1 |
| Akr1b10 | 0.071844 | 0.265323 | 1 |
| Slc2a8 | 0.071782 | 0.024759 | 1 |

| | | | |
|---|---|---|---|
| Nr2c2 | 0.071775 | 0.577831 | 1 |
| Arhgef3 | 0.071737 | 0.493789 | 1 |
| Ubxn1 | 0.071684 | 0.478751 | 1 |
| Mthfsd | 0.071676 | 0.196904 | 1 |
| AU022252 | 0.071487 | 0.090384 | 1 |
| 2210016L21I | 0.071481 | 0.553331 | 1 |
| Pcf11 | 0.07145 | 0.157739 | 1 |
| Gin1 | 0.071378 | 0.294414 | 1 |
| Cstf1 | 0.071352 | 0.132724 | 1 |
| Ilf3 | 0.071341 | 0.641897 | 1 |
| Ndufs4 | 0.071331 | 0.619383 | 1 |
| Tfam | 0.071331 | 0.215781 | 1 |
| Psmd9 | 0.071188 | 0.761527 | 1 |
| Arid4b | 0.071099 | 0.929274 | 1 |
| Cplane1 | 0.070833 | 0.281269 | 1 |
| Cops7b | 0.070823 | 0.628309 | 1 |
| Pwp1 | 0.070823 | 0.358627 | 1 |
| Ndufaf3 | 0.070708 | 0.218015 | 1 |
| Mrps11 | 0.070409 | 0.336049 | 1 |
| Mettl2 | 0.0704 | 0.387568 | 1 |
| Rnf185 | 0.070277 | 0.184387 | 1 |
| Hsdl1 | 0.070257 | 0.113147 | 1 |
| Khdrbs1 | 0.070197 | 0.502323 | 1 |
| Eif3h | 0.070174 | 0.110692 | 1 |
| Arf4 | 0.07012 | 0.136345 | 1 |
| Usf1 | 0.070104 | 0.33623 | 1 |
| 2310009B15I | 0.070092 | 0.600019 | 1 |
| Tmem94 | 0.070055 | 0.104924 | 1 |
| Use1 | 0.070006 | 0.488247 | 1 |
| Dnajb6 | 0.069965 | 0.540345 | 1 |
| Zyg11b | 0.069899 | 0.126869 | 1 |
| Tprgl | 0.069881 | 0.419331 | 1 |
| Snrpb2 | 0.06984 | 0.697936 | 1 |
| Rsu1 | 0.069727 | 0.307201 | 1 |
| Surf2 | 0.069642 | 0.429835 | 1 |
| Gm43672 | 0.069579 | 0.164077 | 1 |
| Mboat1 | 0.069521 | 0.064941 | 1 |
| Atp9b | 0.069512 | 0.42395 | 1 |
| Aagab | 0.069469 | 0.508017 | 1 |
| Dennd4c | 0.069435 | 0.261485 | 1 |
| mt-Nd2 | 0.069375 | 0.480413 | 1 |
| Pprc1 | 0.069369 | 0.371181 | 1 |
| Pes1 | 0.069313 | 0.504329 | 1 |
| Idh3g | 0.069255 | 0.330677 | 1 |
| Supt20 | 0.069218 | 0.38227 | 1 |
| Gm44174 | 0.069217 | 0.588727 | 1 |
| Cnbp | 0.069137 | 0.06776 | 1 |
| 6530402F18I | 0.069108 | 0.057368 | 1 |

| | | | |
|---|---|---|---|
| Acads | 0.069082 | 0.189599 | 1 |
| Eif2ak2 | 0.068997 | 0.639385 | 1 |
| Cep44 | 0.068947 | 0.46861 | 1 |
| Tasp1 | 0.068874 | 0.363501 | 1 |
| Cfap97 | 0.06884 | 0.165632 | 1 |
| Wbp1 | 0.06879 | 0.129791 | 1 |
| Fam50a | 0.068788 | 0.758125 | 1 |
| Zfp512 | 0.068721 | 0.630884 | 1 |
| Txnip | 0.068711 | 0.1207 | 1 |
| Spag9 | 0.068707 | 0.515196 | 1 |
| Ar | 0.068674 | 0.924444 | 1 |
| Bzw2 | 0.068672 | 0.171868 | 1 |
| Psmc5 | 0.06866 | 0.20308 | 1 |
| Zfp560 | 0.068587 | 0.142703 | 1 |
| Sumo1 | 0.068554 | 0.338041 | 1 |
| Sco2 | 0.068469 | 0.441546 | 1 |
| Ift27 | 0.068269 | 0.389795 | 1 |
| Lias | 0.068232 | 0.168892 | 1 |
| Phlpp1 | 0.068194 | 0.520953 | 1 |
| Rab24 | 0.068073 | 0.353434 | 1 |
| Pex13 | 0.068002 | 0.35338 | 1 |
| Atp6v0e | 0.067925 | 0.565799 | 1 |
| Tsnax | 0.067907 | 0.541216 | 1 |
| Alad | 0.067882 | 0.691822 | 1 |
| Cyth4 | 0.067761 | 0.160875 | 1 |
| Fbxw17 | 0.067706 | 0.387732 | 1 |
| Rras | 0.067585 | 0.142692 | 1 |
| Zfp280b | 0.067575 | 0.329413 | 1 |
| Vps28 | 0.067495 | 0.584107 | 1 |
| Tmem120a | 0.067434 | 0.179268 | 1 |
| Ei24 | 0.067246 | 0.252666 | 1 |
| Trappc3 | 0.06717 | 0.965078 | 1 |
| Slc35b3 | 0.067112 | 0.411727 | 1 |
| Unc119b | 0.067062 | 0.137427 | 1 |
| Trp53 | 0.06693 | 0.278806 | 1 |
| Sugp1 | 0.066925 | 0.606688 | 1 |
| Ppan | 0.066862 | 0.45056 | 1 |
| Pcbp2 | 0.066838 | 0.487767 | 1 |
| Galnt12 | 0.06682 | 0.086498 | 1 |
| Zfp445 | 0.066818 | 0.971719 | 1 |
| Pam16 | 0.066807 | 0.99154 | 1 |
| Nudt18 | 0.066685 | 0.487121 | 1 |
| Nvl | 0.066671 | 0.843365 | 1 |
| Nudcd2 | 0.066659 | 0.067431 | 1 |
| Rbm39 | 0.066632 | 0.165896 | 1 |
| Zfp217 | 0.066627 | 0.352676 | 1 |
| Dmap1 | 0.066594 | 0.396411 | 1 |
| Zfp942 | 0.066521 | 0.484953 | 1 |

| | | | |
|---|---|---|---|
| Grap | 0.066501 | 0.557575 | 1 |
| Atp6v1h | 0.066483 | 0.382216 | 1 |
| Sik2 | 0.066478 | 0.341657 | 1 |
| Nabp2 | 0.066471 | 0.36666 | 1 |
| Ndufa7 | 0.066471 | 0.455202 | 1 |
| Pdcd11 | 0.066444 | 0.422488 | 1 |
| Fam122a | 0.066367 | 0.339203 | 1 |
| Zbtb17 | 0.066349 | 0.33177 | 1 |
| Vars | 0.066302 | 0.281391 | 1 |
| Slc25a32 | 0.066263 | 0.440717 | 1 |
| Rex1bd | 0.066213 | 0.144094 | 1 |
| Ak6 | 0.066153 | 0.629954 | 1 |
| Slc35d2 | 0.066129 | 0.184618 | 1 |
| Slfn9 | 0.066041 | 0.507976 | 1 |
| Tmem128 | 0.066027 | 0.099365 | 1 |
| Utp14a | 0.065998 | 0.211383 | 1 |
| Rnf123 | 0.065954 | 0.158468 | 1 |
| Dctn3 | 0.065924 | 0.357229 | 1 |
| Bet1 | 0.065766 | 0.445177 | 1 |
| Cops8 | 0.065701 | 0.73191 | 1 |
| Ttc19 | 0.065672 | 0.129738 | 1 |
| Thumpd3 | 0.065599 | 0.758717 | 1 |
| Ccpg1 | 0.065589 | 0.286393 | 1 |
| Huwe1 | 0.065573 | 0.835449 | 1 |
| BC003965 | 0.065558 | 0.65047 | 1 |
| Ftsj3 | 0.06545 | 0.552041 | 1 |
| Elovl5 | 0.065378 | 0.920941 | 1 |
| Tbc1d4 | 0.065102 | 0.65467 | 1 |
| Chic2 | 0.065087 | 0.420767 | 1 |
| Znhit3 | 0.065031 | 0.790636 | 1 |
| Nipsnap1 | 0.064883 | 0.390961 | 1 |
| Mpp6 | 0.064729 | 0.541888 | 1 |
| Tent4b | 0.064625 | 0.54883 | 1 |
| Cd44 | 0.064606 | 0.692285 | 1 |
| Snhg6 | 0.064481 | 0.598423 | 1 |
| Ncf4 | 0.064423 | 0.060004 | 1 |
| Taf7 | 0.06441 | 0.062278 | 1 |
| Gpx4 | 0.064357 | 0.123217 | 1 |
| Sos2 | 0.064317 | 0.973293 | 1 |
| Rnf44 | 0.064265 | 0.666831 | 1 |
| Nktr | 0.064224 | 0.298554 | 1 |
| Dcun1d3 | 0.064196 | 0.102786 | 1 |
| Amdhd2 | 0.064113 | 0.202926 | 1 |
| Rexo4 | 0.064005 | 0.76934 | 1 |
| Ctu2 | 0.063968 | 0.568412 | 1 |
| Vti1a | 0.063946 | 0.214905 | 1 |
| Ogfr | 0.06394 | 0.208082 | 1 |
| Spop | 0.063825 | 0.678508 | 1 |

| | | | |
|---|---|---|---|
| Alkbh6 | 0.063746 | 0.444135 | 1 |
| Rpl41 | 0.063726 | 0.118835 | 1 |
| Traf6 | 0.063699 | 0.311726 | 1 |
| Fnbp4 | 0.063676 | 0.560964 | 1 |
| Chmp1a | 0.063605 | 0.944318 | 1 |
| 4933434E20I | 0.063574 | 0.465514 | 1 |
| Zfp276 | 0.063448 | 0.108941 | 1 |
| Nop2 | 0.06341 | 0.29908 | 1 |
| Tex14 | 0.063382 | 0.422766 | 1 |
| Stam | 0.063364 | 0.427562 | 1 |
| Lyrm1 | 0.063344 | 0.455558 | 1 |
| 2010016I18F | 0.063333 | 0.776569 | 1 |
| Tg | 0.063275 | 0.547802 | 1 |
| Eaf1 | 0.06326 | 0.450058 | 1 |
| Sod2 | 0.063174 | 0.503799 | 1 |
| Ppil2 | 0.063146 | 0.199823 | 1 |
| Zfyve1 | 0.06314 | 0.108821 | 1 |
| Utp23 | 0.063117 | 0.537811 | 1 |
| Iah1 | 0.063074 | 0.734961 | 1 |
| Tmed4 | 0.063028 | 0.317997 | 1 |
| Polm | 0.063011 | 0.316554 | 1 |
| Jtb | 0.06298 | 0.471797 | 1 |
| Klhl24 | 0.062979 | 0.045452 | 1 |
| Cd200 | 0.062972 | 0.264989 | 1 |
| Dph7 | 0.062958 | 0.414459 | 1 |
| Plagl2 | 0.062909 | 0.283679 | 1 |
| Stx18 | 0.062828 | 0.326469 | 1 |
| Tbcel | 0.062732 | 0.176484 | 1 |
| Golph3 | 0.062713 | 0.490352 | 1 |
| Zfr | 0.06265 | 0.34091 | 1 |
| Slc33a1 | 0.062525 | 0.371515 | 1 |
| Arhgap9 | 0.062488 | 0.252262 | 1 |
| Ankrd28 | 0.062484 | 0.078441 | 1 |
| Myd88 | 0.06244 | 0.317795 | 1 |
| Nmt1 | 0.062387 | 0.332895 | 1 |
| Metap1 | 0.062281 | 0.550337 | 1 |
| Cd37 | 0.062254 | 0.008054 | 1 |
| Snai3 | 0.06224 | 0.408223 | 1 |
| Ifi209 | 0.062237 | 0.294164 | 1 |
| Optn | 0.062224 | 0.074249 | 1 |
| Gprin3 | 0.062199 | 0.960009 | 1 |
| Terf2ip | 0.062112 | 0.574273 | 1 |
| Git2 | 0.06208 | 0.977126 | 1 |
| Gpatch1 | 0.06208 | 0.510149 | 1 |
| Entpd5 | 0.061989 | 0.724407 | 1 |
| Cbr1 | 0.061871 | 0.31239 | 1 |
| Zfp429 | 0.06171 | 0.268321 | 1 |
| Flywch1 | 0.06164 | 0.488362 | 1 |

| | | | |
|---|---|---|---|
| Rnmt | 0.061616 | 0.553418 | 1 |
| Manba | 0.061577 | 0.78174 | 1 |
| Shld2 | 0.061556 | 0.298233 | 1 |
| Cript | 0.061532 | 0.413733 | 1 |
| Thoc1 | 0.061523 | 0.474625 | 1 |
| Pnpla8 | 0.061512 | 0.199605 | 1 |
| Maz | 0.061499 | 0.542295 | 1 |
| Utp15 | 0.061492 | 0.220228 | 1 |
| Tmem106a | 0.061492 | 0.177484 | 1 |
| Tnfsf9 | 0.06145 | 0.871749 | 1 |
| Tnrc6c | 0.061401 | 0.279006 | 1 |
| Chtf8 | 0.061332 | 0.390349 | 1 |
| Ssbp3 | 0.0613 | 0.511604 | 1 |
| Tep1 | 0.061202 | 0.201175 | 1 |
| Pick1 | 0.061171 | 0.513062 | 1 |
| Phf1 | 0.061147 | 0.827267 | 1 |
| Ctsa | 0.061107 | 0.406873 | 1 |
| Fech | 0.061104 | 0.119486 | 1 |
| Abcb1a | 0.061101 | 0.261692 | 1 |
| Pinx1 | 0.060996 | 0.469311 | 1 |
| Mdm4 | 0.060986 | 0.87268 | 1 |
| Dnajb4 | 0.060982 | 0.673244 | 1 |
| Nepro | 0.060975 | 0.243567 | 1 |
| Wdr74 | 0.060953 | 0.269659 | 1 |
| Srrm2 | 0.060877 | 0.452097 | 1 |
| Pdrg1 | 0.060877 | 0.276386 | 1 |
| Tprn | 0.06087 | 0.224787 | 1 |
| Tnrc6b | 0.060863 | 0.501845 | 1 |
| Galk1 | 0.060813 | 0.463059 | 1 |
| Sec63 | 0.060782 | 0.284586 | 1 |
| Tceanc2 | 0.060752 | 0.216467 | 1 |
| Mvp | 0.060743 | 0.262145 | 1 |
| Cpped1 | 0.060628 | 0.565212 | 1 |
| Art2b | 0.060527 | 0.723539 | 1 |
| Zdhhc15 | 0.060524 | 0.829548 | 1 |
| Gbe1 | 0.060418 | 0.954335 | 1 |
| Osbpl7 | 0.060397 | 0.894586 | 1 |
| Zfpm1 | 0.060365 | 0.361248 | 1 |
| Kras | 0.06035 | 0.881946 | 1 |
| Prpf31 | 0.06023 | 0.910341 | 1 |
| Ccdc115 | 0.060101 | 0.412053 | 1 |
| Psmd12 | 0.060081 | 0.244828 | 1 |
| Dnajc1 | 0.060013 | 0.573926 | 1 |
| Specc1l | 0.059988 | 0.406212 | 1 |
| Dnajc24 | 0.059956 | 0.421298 | 1 |
| Mau2 | 0.05991 | 0.428212 | 1 |
| Plk3 | 0.05989 | 0.127326 | 1 |
| Lyrm2 | 0.059853 | 0.467953 | 1 |

| | | | |
|---|---|---|---|
| Arhgap31 | 0.059835 | 0.963906 | 1 |
| Timm9 | 0.059816 | 0.066254 | 1 |
| Snapc2 | 0.059774 | 0.275598 | 1 |
| Pop7 | 0.059659 | 0.232874 | 1 |
| Dusp4 | 0.05963 | 0.8327 | 1 |
| Tmem248 | 0.05962 | 0.0734 | 1 |
| Faap20 | 0.059289 | 0.594028 | 1 |
| Hp1bp3 | 0.059148 | 0.754291 | 1 |
| Dhx38 | 0.059109 | 0.357505 | 1 |
| Polr1d | 0.058935 | 0.300605 | 1 |
| 2810001G20 | 0.058876 | 0.433351 | 1 |
| Itch | 0.058793 | 0.819247 | 1 |
| Strn4 | 0.058708 | 0.290962 | 1 |
| Gskip | 0.058691 | 0.432273 | 1 |
| Helz | 0.058631 | 0.545221 | 1 |
| Tmem87b | 0.05861 | 0.465364 | 1 |
| Dusp12 | 0.058581 | 0.448514 | 1 |
| Eif3b | 0.058569 | 0.356438 | 1 |
| Rsph3b | 0.058561 | 0.161372 | 1 |
| Slc37a4 | 0.058534 | 0.509278 | 1 |
| Kat6b | 0.058406 | 0.360817 | 1 |
| Slc38a6 | 0.058401 | 0.301401 | 1 |
| Eipr1 | 0.058369 | 0.621857 | 1 |
| 3830406C13 | 0.058289 | 0.512578 | 1 |
| Prorsd1 | 0.058278 | 0.412257 | 1 |
| Bcap29 | 0.05825 | 0.188652 | 1 |
| Ddx19b | 0.058238 | 0.385552 | 1 |
| Mctp2 | 0.058233 | 0.824389 | 1 |
| Fam172a | 0.058117 | 0.523889 | 1 |
| Pgrmc2 | 0.058088 | 0.648413 | 1 |
| Mrpl11 | 0.05808 | 0.348415 | 1 |
| Dcaf4 | 0.058072 | 0.467334 | 1 |
| Ube4a | 0.058037 | 0.650565 | 1 |
| Bmt2 | 0.05799 | 0.747477 | 1 |
| Akap8 | 0.057915 | 0.535048 | 1 |
| Gpalpp1 | 0.057911 | 0.611161 | 1 |
| Nap1l4 | 0.057886 | 0.683768 | 1 |
| Tmem209 | 0.057884 | 0.253368 | 1 |
| Gcc1 | 0.057879 | 0.468826 | 1 |
| Rap2c | 0.057854 | 0.27246 | 1 |
| Fkbp3 | 0.057758 | 0.599676 | 1 |
| Stap1 | 0.057653 | 0.336997 | 1 |
| Sufu | 0.057642 | 0.643435 | 1 |
| Spryd7 | 0.057585 | 0.541929 | 1 |
| Slc2a9 | 0.057582 | 0.415067 | 1 |
| Gnaq | 0.057566 | 0.235211 | 1 |
| Fkbp15 | 0.05755 | 0.163908 | 1 |
| Fbxl8 | 0.057548 | 0.448184 | 1 |

| | | | |
|---|---|---|---|
| Mtf2 | 0.057537 | 0.774373 | 1 |
| Dnajc17 | 0.057468 | 0.240608 | 1 |
| Dennd1b | 0.05745 | 0.766687 | 1 |
| Gsk3a | 0.057388 | 0.246414 | 1 |
| Cdc42se2 | 0.057277 | 0.511654 | 1 |
| Pex14 | 0.057246 | 0.768882 | 1 |
| Ift52 | 0.057103 | 0.736299 | 1 |
| Tes | 0.057071 | 0.390859 | 1 |
| Ccnt1 | 0.057058 | 0.567688 | 1 |
| Spg21 | 0.056986 | 0.465737 | 1 |
| Fundc1 | 0.056915 | 0.974758 | 1 |
| 9330175E14I | 0.056794 | 0.453514 | 1 |
| Blvra | 0.056695 | 0.841857 | 1 |
| Zfp617 | 0.056623 | 0.622318 | 1 |
| Dguok | 0.056601 | 0.184239 | 1 |
| Atg7 | 0.056487 | 0.534898 | 1 |
| Poli | 0.056391 | 0.602672 | 1 |
| Hivep2 | 0.056389 | 0.96203 | 1 |
| Mff | 0.056377 | 0.937709 | 1 |
| Heg1 | 0.056371 | 0.635502 | 1 |
| Acaa2 | 0.056352 | 0.350106 | 1 |
| Ppp2r2d | 0.056343 | 0.337859 | 1 |
| Ccdc25 | 0.05632 | 0.284887 | 1 |
| Coa3 | 0.056303 | 0.577787 | 1 |
| Rbks | 0.056225 | 0.204852 | 1 |
| Reep3 | 0.056214 | 0.257435 | 1 |
| Parp6 | 0.056168 | 0.195893 | 1 |
| Gsdmd | 0.056141 | 0.802561 | 1 |
| Rit1 | 0.056108 | 0.617631 | 1 |
| Tmbim1 | 0.056011 | 0.4978 | 1 |
| Ccrl2 | 0.056001 | 0.997869 | 1 |
| Rnf34 | 0.055975 | 0.326755 | 1 |
| Fgf13 | 0.055964 | 0.880646 | 1 |
| Ptges3 | 0.055941 | 0.301401 | 1 |
| Stk3 | 0.055886 | 0.344576 | 1 |
| Rps29 | 0.055783 | 0.27387 | 1 |
| Pigu | 0.055625 | 0.178181 | 1 |
| Rsf1 | 0.055537 | 0.268687 | 1 |
| Opa3 | 0.055528 | 0.113074 | 1 |
| Casp2 | 0.055511 | 0.713208 | 1 |
| Disp1 | 0.055479 | 0.564265 | 1 |
| Jmjd6 | 0.055473 | 0.787819 | 1 |
| Riok3 | 0.055385 | 0.813627 | 1 |
| Zkscan1 | 0.055376 | 0.950822 | 1 |
| Ube2j2 | 0.055367 | 0.604221 | 1 |
| Sra1 | 0.055339 | 0.480027 | 1 |
| Exoc3 | 0.055327 | 0.405294 | 1 |
| Zfpl1 | 0.055311 | 0.102783 | 1 |

| | | | |
|---|---|---|---|
| Crtc3 | 0.055166 | 0.361455 | 1 |
| Rhoh | 0.055162 | 0.369991 | 1 |
| March6 | 0.055076 | 0.542267 | 1 |
| Dcun1d1 | 0.055013 | 0.932571 | 1 |
| Atp5c1 | 0.055005 | 0.59907 | 1 |
| Lyar | 0.054913 | 0.997964 | 1 |
| Srpr | 0.054861 | 0.225116 | 1 |
| Ccdc125 | 0.054822 | 0.859296 | 1 |
| Ddx42 | 0.05478 | 0.535377 | 1 |
| Ambra1 | 0.054754 | 0.830764 | 1 |
| Mthfd2 | 0.054685 | 0.949036 | 1 |
| Ddx5 | 0.054625 | 0.280712 | 1 |
| Edc3 | 0.054591 | 0.356533 | 1 |
| Adar | 0.05458 | 0.66571 | 1 |
| Thoc3 | 0.054574 | 0.50712 | 1 |
| Tbc1d22a | 0.054543 | 0.291761 | 1 |
| Cep135 | 0.054469 | 0.754343 | 1 |
| Gcsh | 0.054436 | 0.257683 | 1 |
| Fas | 0.054417 | 0.094397 | 1 |
| Cic | 0.05422 | 0.92548 | 1 |
| Ube2z | 0.054189 | 0.409714 | 1 |
| C330018D20 | 0.054092 | 0.086383 | 1 |
| 1810013L24I | 0.054074 | 0.615647 | 1 |
| Zfp622 | 0.053985 | 0.705372 | 1 |
| Usp8 | 0.05395 | 0.423599 | 1 |
| Dzip3 | 0.053854 | 0.367614 | 1 |
| 1110038B12I | 0.053815 | 0.299141 | 1 |
| Trgv2 | 0.053814 | 0.196542 | 1 |
| Sirt5 | 0.053779 | 0.524818 | 1 |
| Cdk5rap3 | 0.053753 | 0.766518 | 1 |
| Prmt6 | 0.053693 | 0.048204 | 1 |
| Mrps31 | 0.053628 | 0.52255 | 1 |
| Gm14326 | 0.053617 | 0.121868 | 1 |
| Bnip1 | 0.05356 | 0.304854 | 1 |
| Chmp6 | 0.053548 | 0.625458 | 1 |
| Kif1bp | 0.053495 | 0.505375 | 1 |
| Adck1 | 0.053487 | 0.279673 | 1 |
| Tmed7 | 0.053404 | 0.406783 | 1 |
| Kdm6a | 0.0534 | 0.986961 | 1 |
| Atp6v1d | 0.0533 | 0.553509 | 1 |
| Bag3 | 0.053252 | 0.662686 | 1 |
| 4632427E13I | 0.05323 | 0.371537 | 1 |
| Tbl3 | 0.053204 | 0.600131 | 1 |
| Cdyl2 | 0.053177 | 0.238732 | 1 |
| Scap | 0.053164 | 0.773936 | 1 |
| Crk | 0.053021 | 0.64358 | 1 |
| Dhx57 | 0.052958 | 0.873815 | 1 |
| Mlkl | 0.052917 | 0.388026 | 1 |

| | | | |
|---|---|---|---|
| Trim65 | 0.052911 | 0.688455 | 1 |
| Mrps7 | 0.052897 | 0.780362 | 1 |
| Marcksl1 | 0.052803 | 0.988143 | 1 |
| Isoc2b | 0.052711 | 0.146392 | 1 |
| Vps26b | 0.052663 | 0.68099 | 1 |
| Kdm4c | 0.052643 | 0.663149 | 1 |
| Stk40 | 0.052623 | 0.291592 | 1 |
| Zfp213 | 0.052567 | 0.604554 | 1 |
| Rab2b | 0.052527 | 0.590062 | 1 |
| Ints3 | 0.052508 | 0.839231 | 1 |
| Rnf41 | 0.052444 | 0.963394 | 1 |
| Fam136a | 0.052364 | 0.467963 | 1 |
| Eya3 | 0.052292 | 0.369233 | 1 |
| Eif2ak3 | 0.052247 | 0.306561 | 1 |
| 4930523C07I | 0.052237 | 0.387268 | 1 |
| Rbm33 | 0.052229 | 0.479303 | 1 |
| Cmc4 | 0.05216 | 0.576478 | 1 |
| Smdt1 | 0.052004 | 0.440818 | 1 |
| G6pc3 | 0.051986 | 0.387147 | 1 |
| Vps9d1 | 0.051954 | 0.270558 | 1 |
| Prpf4 | 0.051927 | 0.879161 | 1 |
| Etfrf1 | 0.051867 | 0.461601 | 1 |
| Ptges2 | 0.051864 | 0.34382 | 1 |
| Ggact | 0.051739 | 0.380043 | 1 |
| Sar1a | 0.051711 | 0.884689 | 1 |
| Gtpbp4 | 0.051575 | 0.193234 | 1 |
| Taf13 | 0.051499 | 0.383216 | 1 |
| Coa4 | 0.051467 | 0.583824 | 1 |
| Dnajb12 | 0.051302 | 0.602814 | 1 |
| Mfn1 | 0.051297 | 0.744243 | 1 |
| Alg5 | 0.051292 | 0.03141 | 1 |
| Slc50a1 | 0.051247 | 0.50073 | 1 |
| Edem2 | 0.051235 | 0.352756 | 1 |
| Patl1 | 0.051186 | 0.847621 | 1 |
| Esm1 | 0.05118 | 0.7823 | 1 |
| Sec11a | 0.051173 | 0.490639 | 1 |
| Cox15 | 0.051118 | 0.678311 | 1 |
| Polr3e | 0.0511 | 0.279643 | 1 |
| Slc7a1 | 0.051068 | 0.715962 | 1 |
| Tmem123 | 0.051022 | 0.578532 | 1 |
| Btf3 | 0.050988 | 0.229632 | 1 |
| Actr8 | 0.050988 | 0.741711 | 1 |
| Mex3c | 0.050903 | 0.574519 | 1 |
| Anapc2 | 0.050791 | 0.362092 | 1 |
| Nt5dc1 | 0.050724 | 0.396403 | 1 |
| Chchd7 | 0.050687 | 0.506923 | 1 |
| Cpsf3 | 0.050545 | 0.47625 | 1 |
| Zfp507 | 0.050544 | 0.287285 | 1 |

| | | | |
|---|---|---|---|
| Top3a | 0.050467 | 0.398811 | 1 |
| Nudt13 | 0.050463 | 0.173547 | 1 |
| Hax1 | 0.050437 | 0.090956 | 1 |
| Bcl11b | 0.050361 | 0.092512 | 1 |
| Gm5617 | 0.050338 | 0.625269 | 1 |
| Med4 | 0.050245 | 0.929779 | 1 |
| 2610008E11I | 0.050211 | 0.267634 | 1 |
| Ecpas | 0.05019 | 0.911198 | 1 |
| Dhx40 | 0.050183 | 0.327871 | 1 |
| Zzz3 | 0.050175 | 0.283377 | 1 |
| Ube3a | 0.050145 | 0.593402 | 1 |
| Grsf1 | 0.050129 | 0.829272 | 1 |
| Rbm19 | 0.050121 | 0.353588 | 1 |
| Usp9x | 0.050073 | 0.405893 | 1 |
| Slc39a7 | 0.050055 | 0.825439 | 1 |
| Acbd3 | 0.050032 | 0.317482 | 1 |
| Mfsd1 | 0.049988 | 0.854614 | 1 |
| 1600020E01I | 0.049961 | 0.598092 | 1 |
| Tmem126b | 0.049874 | 0.081346 | 1 |
| Plbd2 | 0.049865 | 0.008997 | 1 |
| Chd2 | 0.049783 | 0.082447 | 1 |
| Pacsin1 | 0.049768 | 0.351639 | 1 |
| Mknk2 | 0.049669 | 0.823745 | 1 |
| Nsmce1 | 0.049645 | 0.466447 | 1 |
| Exosc2 | 0.049639 | 0.517923 | 1 |
| Tnfsf13b | 0.049625 | 0.069779 | 1 |
| Zdhhc6 | 0.049607 | 0.474812 | 1 |
| Il18bp | 0.049602 | 0.003297 | 1 |
| Csnk2a2 | 0.049594 | 0.169819 | 1 |
| Nudt2 | 0.049584 | 0.205256 | 1 |
| Zkscan14 | 0.049463 | 0.968908 | 1 |
| Abcd4 | 0.049461 | 0.400455 | 1 |
| Cited2 | 0.049413 | 0.792436 | 1 |
| Tmem106c | 0.049409 | 0.455756 | 1 |
| Stard7 | 0.049367 | 0.7778 | 1 |
| Eif4ebp2 | 0.049355 | 0.731643 | 1 |
| 4931406P16I | 0.049275 | 0.728473 | 1 |
| Adpgk | 0.049241 | 0.264163 | 1 |
| Rab3gap1 | 0.049211 | 0.911283 | 1 |
| Prcc | 0.04917 | 0.531899 | 1 |
| U2af1l4 | 0.049156 | 0.806444 | 1 |
| Kansl1l | 0.049093 | 0.379177 | 1 |
| Hacl1 | 0.049091 | 0.201385 | 1 |
| Snapc3 | 0.049077 | 0.896065 | 1 |
| Uggt1 | 0.049075 | 0.951153 | 1 |
| Mrpl55 | 0.049035 | 0.370949 | 1 |
| Polb | 0.049014 | 0.498586 | 1 |
| Secisbp2l | 0.048989 | 0.435535 | 1 |

| | | | |
|---|---|---|---|
| Ago1 | 0.04898 | 0.376826 | 1 |
| Sptlc2 | 0.048948 | 0.910332 | 1 |
| Mkrn1 | 0.048876 | 0.556742 | 1 |
| Cdk10 | 0.048863 | 0.519591 | 1 |
| Tmem230 | 0.048826 | 0.516847 | 1 |
| Rgs3 | 0.048757 | 0.459037 | 1 |
| Znhit1 | 0.048757 | 0.739834 | 1 |
| Pgs1 | 0.048584 | 0.38291 | 1 |
| Arglu1 | 0.048583 | 0.929871 | 1 |
| Usb1 | 0.048505 | 0.762858 | 1 |
| Dhcr7 | 0.048462 | 0.713009 | 1 |
| Gm9725 | 0.048433 | 0.493969 | 1 |
| Nop10 | 0.048422 | 0.576175 | 1 |
| Bach1 | 0.048402 | 0.559808 | 1 |
| Fam168a | 0.048331 | 0.841789 | 1 |
| Zfp984 | 0.048291 | 0.40252 | 1 |
| Mfn2 | 0.048284 | 0.304598 | 1 |
| Exoc7 | 0.048272 | 0.728611 | 1 |
| Yipf1 | 0.048227 | 0.375453 | 1 |
| Ergic1 | 0.048176 | 0.03373 | 1 |
| Kat2b | 0.048175 | 0.449453 | 1 |
| Lsg1 | 0.048173 | 0.882002 | 1 |
| Fra10ac1 | 0.04817 | 0.044117 | 1 |
| Acadl | 0.04816 | 0.691217 | 1 |
| Nrde2 | 0.048132 | 0.313619 | 1 |
| Cluap1 | 0.047954 | 0.904271 | 1 |
| Mindy3 | 0.047942 | 0.414793 | 1 |
| Tsr1 | 0.04794 | 0.601899 | 1 |
| Letm2 | 0.047928 | 0.478293 | 1 |
| Aatf | 0.047852 | 0.383536 | 1 |
| Stk16 | 0.047838 | 0.367856 | 1 |
| Eml2 | 0.047825 | 0.219143 | 1 |
| Lpgat1 | 0.047817 | 0.795544 | 1 |
| Zfp281 | 0.047805 | 0.551722 | 1 |
| Pex2 | 0.047793 | 0.342558 | 1 |
| Rufy2 | 0.04776 | 0.436231 | 1 |
| Kmt2b | 0.047724 | 0.416746 | 1 |
| Gclc | 0.0477 | 0.185524 | 1 |
| Phf3 | 0.047634 | 0.762768 | 1 |
| Mbd5 | 0.047626 | 0.463883 | 1 |
| Ndel1 | 0.04761 | 0.212707 | 1 |
| Lrrc40 | 0.047598 | 0.179716 | 1 |
| 5730455P16 | 0.047587 | 0.276895 | 1 |
| Bag1 | 0.047584 | 0.444333 | 1 |
| Arl4a | 0.047568 | 0.643329 | 1 |
| Mrpl21 | 0.047555 | 0.542432 | 1 |
| Wdr3 | 0.047304 | 0.837627 | 1 |
| Ppargc1b | 0.047291 | 0.18796 | 1 |

| | | | |
|---|---|---|---|
| Tmem104 | 0.047268 | 0.774593 | 1 |
| Gm26740 | 0.047204 | 0.815301 | 1 |
| Cenpb | 0.047106 | 0.617173 | 1 |
| Zfp950 | 0.047012 | 0.352474 | 1 |
| Washc4 | 0.04701 | 0.555644 | 1 |
| Thumpd2 | 0.046973 | 0.823901 | 1 |
| Tnrc6a | 0.046881 | 0.714263 | 1 |
| Trit1 | 0.046835 | 0.371038 | 1 |
| Ormdl2 | 0.04682 | 0.610134 | 1 |
| Vipas39 | 0.04671 | 0.601092 | 1 |
| Tcof1 | 0.046695 | 0.905524 | 1 |
| Tmem147 | 0.04668 | 0.460118 | 1 |
| Ndufa9 | 0.046673 | 0.521459 | 1 |
| Lpcat1 | 0.04665 | 0.521086 | 1 |
| Mark2 | 0.04654 | 0.797047 | 1 |
| Afg1l | 0.046499 | 0.449662 | 1 |
| Orc5 | 0.046417 | 0.613969 | 1 |
| Armt1 | 0.046283 | 0.535154 | 1 |
| Eftud2 | 0.046218 | 0.622929 | 1 |
| Plekhf1 | 0.046161 | 0.755943 | 1 |
| Trim39 | 0.046153 | 0.286339 | 1 |
| Arfgap3 | 0.046131 | 0.421552 | 1 |
| Polr2f | 0.04611 | 0.592526 | 1 |
| AI987944 | 0.046096 | 0.232241 | 1 |
| Tmem165 | 0.046053 | 0.734524 | 1 |
| Irak1 | 0.046045 | 0.480344 | 1 |
| Brd2 | 0.046037 | 0.671687 | 1 |
| Gtpbp1 | 0.045956 | 0.729552 | 1 |
| Mpg | 0.045891 | 0.51027 | 1 |
| Ccnl2 | 0.045863 | 0.906189 | 1 |
| Nsa2 | 0.045853 | 0.683868 | 1 |
| Zfand6 | 0.045798 | 0.793694 | 1 |
| Ahsa1 | 0.045748 | 0.953537 | 1 |
| Ube2h | 0.045739 | 0.885621 | 1 |
| Lrrc14 | 0.045713 | 0.33169 | 1 |
| Mrpl19 | 0.045711 | 0.532307 | 1 |
| Tmem138 | 0.04571 | 0.526548 | 1 |
| Prpf39 | 0.045613 | 0.79634 | 1 |
| Aldh3a2 | 0.045607 | 0.063459 | 1 |
| Wdr12 | 0.045515 | 0.829939 | 1 |
| Hint2 | 0.04548 | 0.004159 | 1 |
| Pmaip1 | 0.045477 | 0.745152 | 1 |
| Ralgapa1 | 0.045435 | 0.898158 | 1 |
| Psip1 | 0.045428 | 0.966808 | 1 |
| Kansl1 | 0.045423 | 0.701706 | 1 |
| Ndufb10 | 0.045359 | 0.51127 | 1 |
| Snupn | 0.045294 | 0.64805 | 1 |
| Rps26 | 0.045223 | 0.815844 | 1 |

| | | | |
|---|---|---|---|
| Tomm5 | 0.045218 | 0.694031 | 1 |
| Zfp609 | 0.04521 | 0.693968 | 1 |
| Pnpt1 | 0.045175 | 0.773585 | 1 |
| Nifk | 0.04516 | 0.779455 | 1 |
| Katnbl1 | 0.045088 | 0.750135 | 1 |
| Wdr77 | 0.045036 | 0.299296 | 1 |
| Ccdc9 | 0.044994 | 0.643015 | 1 |
| Smim3 | 0.044966 | 0.166039 | 1 |
| Trappc2l | 0.044951 | 0.771579 | 1 |
| Pop1 | 0.044929 | 0.173643 | 1 |
| Ero1l | 0.044884 | 0.035932 | 1 |
| Ddx21 | 0.044822 | 0.66202 | 1 |
| Resf1 | 0.044783 | 0.654619 | 1 |
| Kdm3a | 0.044782 | 0.93055 | 1 |
| Fbxo8 | 0.044674 | 0.28473 | 1 |
| Rnf121 | 0.044632 | 0.43668 | 1 |
| Ttc32 | 0.044575 | 0.287431 | 1 |
| Ap2a1 | 0.044574 | 0.74954 | 1 |
| Golga7 | 0.044558 | 0.741644 | 1 |
| Apoo | 0.044518 | 0.393329 | 1 |
| Ubald2 | 0.044499 | 0.460977 | 1 |
| Gm36975 | 0.044464 | 0.391875 | 1 |
| Mbtd1 | 0.044451 | 0.580108 | 1 |
| Dusp19 | 0.04445 | 0.833584 | 1 |
| Ube2e1 | 0.044415 | 0.512616 | 1 |
| D430042O09 | 0.044414 | 0.690927 | 1 |
| Ice2 | 0.044372 | 0.3912 | 1 |
| Dop1a | 0.04435 | 0.904116 | 1 |
| Ildr1 | 0.044349 | 0.994057 | 1 |
| Atxn7l3b | 0.044328 | 0.688559 | 1 |
| Tarbp2 | 0.04421 | 0.160025 | 1 |
| Sh3glb1 | 0.044019 | 0.671903 | 1 |
| Cox10 | 0.043924 | 0.890309 | 1 |
| Zfp738 | 0.043859 | 0.48256 | 1 |
| Trim35 | 0.043804 | 0.553795 | 1 |
| Snhg16 | 0.043799 | 0.472226 | 1 |
| Yaf2 | 0.043641 | 0.786121 | 1 |
| Cdadc1 | 0.043603 | 0.721986 | 1 |
| Tbc1d17 | 0.043565 | 0.45489 | 1 |
| Noc3l | 0.043563 | 0.794853 | 1 |
| Metap1d | 0.043549 | 0.394542 | 1 |
| Mapk8ip3 | 0.043541 | 0.744023 | 1 |
| Ngrn | 0.043533 | 0.702214 | 1 |
| Napa | 0.043511 | 0.36504 | 1 |
| Polr2d | 0.043467 | 0.597851 | 1 |
| Bora | 0.043446 | 0.387926 | 1 |
| Siva1 | 0.043417 | 0.927101 | 1 |
| Phlda1 | 0.043396 | 0.420302 | 1 |

| | | | |
|---|---|---|---|
| Lats1 | 0.043387 | 0.484092 | 1 |
| Gm39469 | 0.04338 | 0.94038 | 1 |
| Trappc11 | 0.043359 | 0.947349 | 1 |
| Rheb | 0.043331 | 0.733703 | 1 |
| Cd101 | 0.043314 | 0.417527 | 1 |
| Pcbd2 | 0.043276 | 0.879192 | 1 |
| Lxn | 0.043189 | 0.701066 | 1 |
| Ust | 0.043183 | 0.849581 | 1 |
| Gm16565 | 0.04316 | 0.343276 | 1 |
| Trappc8 | 0.043128 | 0.772308 | 1 |
| Hrh2 | 0.043077 | 0.469431 | 1 |
| Smyd2 | 0.043057 | 0.642385 | 1 |
| Gabarap | 0.043029 | 0.677725 | 1 |
| A530088E08I | 0.04296 | 0.295904 | 1 |
| Crebzf | 0.042943 | 0.691182 | 1 |
| Avl9 | 0.042936 | 0.555413 | 1 |
| Setd4 | 0.042909 | 0.591516 | 1 |
| Pyroxd1 | 0.042878 | 0.57257 | 1 |
| Znrf2 | 0.042872 | 0.527224 | 1 |
| Chmp3 | 0.04284 | 0.807039 | 1 |
| Elp2 | 0.042771 | 0.732396 | 1 |
| Dhx36 | 0.042728 | 0.915566 | 1 |
| Zmiz1 | 0.042693 | 0.742338 | 1 |
| Atp5d | 0.042611 | 0.61128 | 1 |
| Ptpn12 | 0.042579 | 0.946212 | 1 |
| Acsf2 | 0.042569 | 0.241032 | 1 |
| Cdip1 | 0.042498 | 0.329333 | 1 |
| Relch | 0.042477 | 0.757264 | 1 |
| Gcfc2 | 0.042458 | 0.58287 | 1 |
| Mtf1 | 0.042443 | 0.990045 | 1 |
| Eif2b2 | 0.042421 | 0.36812 | 1 |
| Mettl26 | 0.042379 | 0.712468 | 1 |
| Ankrd46 | 0.04234 | 0.280274 | 1 |
| Mrps18c | 0.042318 | 0.378703 | 1 |
| Pnisr | 0.042205 | 0.484964 | 1 |
| Lamb3 | 0.042199 | 0.556917 | 1 |
| Ivd | 0.042087 | 0.644126 | 1 |
| LTO1 | 0.042055 | 0.509142 | 1 |
| Wasl | 0.041979 | 0.770856 | 1 |
| Arel1 | 0.041938 | 0.426496 | 1 |
| Isg20l2 | 0.041876 | 0.615079 | 1 |
| Zfp831 | 0.041864 | 0.458694 | 1 |
| Pip4p2 | 0.041862 | 0.65364 | 1 |
| Ppp3cb | 0.041853 | 0.908941 | 1 |
| Timm10 | 0.041683 | 0.933112 | 1 |
| Enoph1 | 0.041682 | 0.468222 | 1 |
| Zc3hc1 | 0.041664 | 0.12269 | 1 |
| Chd4 | 0.041656 | 0.950351 | 1 |

| | | | |
|---|---|---|---|
| Eri3 | 0.041621 | 0.761472 | 1 |
| Ctla2b | 0.041515 | 0.782935 | 1 |
| Armc10 | 0.041441 | 0.662623 | 1 |
| Tomm40l | 0.041332 | 0.755039 | 1 |
| Nudt7 | 0.041282 | 0.549415 | 1 |
| Gmeb2 | 0.041261 | 0.844134 | 1 |
| Rtcb | 0.041207 | 0.158919 | 1 |
| Zfp451 | 0.0412 | 0.057086 | 1 |
| Zscan26 | 0.04117 | 0.726218 | 1 |
| Hsbp1 | 0.041167 | 0.533382 | 1 |
| Slc16a3 | 0.041126 | 0.862387 | 1 |
| Slc25a45 | 0.041035 | 0.879856 | 1 |
| Babam2 | 0.041009 | 0.673496 | 1 |
| Vcpip1 | 0.040924 | 0.979589 | 1 |
| Foxk1 | 0.040853 | 0.869304 | 1 |
| Mcm9 | 0.040844 | 0.912329 | 1 |
| Ap3m2 | 0.04083 | 0.526735 | 1 |
| Cul2 | 0.040807 | 0.982345 | 1 |
| Nras | 0.040707 | 0.935654 | 1 |
| Trim56 | 0.040702 | 0.467074 | 1 |
| Mthfs | 0.040694 | 0.972561 | 1 |
| Zfp472 | 0.040666 | 0.42844 | 1 |
| Zfp932 | 0.040647 | 0.466919 | 1 |
| AW554918 | 0.0406 | 0.817466 | 1 |
| Znhit6 | 0.040575 | 0.57842 | 1 |
| Bccip | 0.040573 | 0.656438 | 1 |
| Rab37 | 0.040567 | 0.494511 | 1 |
| E2f3 | 0.040553 | 0.992345 | 1 |
| Tubgcp5 | 0.040549 | 0.572787 | 1 |
| Nmnat3 | 0.040539 | 0.987772 | 1 |
| Napg | 0.040495 | 0.619529 | 1 |
| Lin52 | 0.04049 | 0.769495 | 1 |
| 1810058I24R | 0.040336 | 0.493074 | 1 |
| Arfip2 | 0.040291 | 0.9234 | 1 |
| Elac2 | 0.040288 | 0.492617 | 1 |
| Nfyc | 0.04022 | 0.665596 | 1 |
| Clp1 | 0.04021 | 0.622034 | 1 |
| Sec62 | 0.040197 | 0.991296 | 1 |
| Map3k5 | 0.040181 | 0.67162 | 1 |
| Shprh | 0.040101 | 0.92527 | 1 |
| Xlr4a | 0.040015 | 0.293012 | 1 |
| Hps1 | 0.04 | 0.752593 | 1 |
| Dus1l | 0.039987 | 0.918121 | 1 |
| Cd99l2 | 0.039957 | 0.311226 | 1 |
| Rbm45 | 0.039913 | 0.822475 | 1 |
| Ebag9 | 0.039905 | 0.412838 | 1 |
| Jund | 0.039877 | 0.36692 | 1 |
| Itpripl2 | 0.039876 | 0.326596 | 1 |

| | | | |
|---|---|---|---|
| Gm16124 | 0.039862 | 0.287622 | 1 |
| Klhl26 | 0.039848 | 0.406591 | 1 |
| Edem1 | 0.039846 | 0.611589 | 1 |
| Manbal | 0.039814 | 0.723323 | 1 |
| Nmt2 | 0.039759 | 0.817623 | 1 |
| Msi2 | 0.039747 | 0.891626 | 1 |
| Tcirg1 | 0.039685 | 0.937081 | 1 |
| Apobr | 0.039657 | 0.073527 | 1 |
| Gatd3a | 0.039625 | 0.21149 | 1 |
| Crlf3 | 0.039571 | 0.746181 | 1 |
| Slc29a1 | 0.039541 | 0.533564 | 1 |
| Mt3 | 0.039537 | 0.180354 | 1 |
| Timm23 | 0.039523 | 0.543876 | 1 |
| Rab22a | 0.039513 | 0.918699 | 1 |
| Lcorl | 0.039509 | 0.888865 | 1 |
| Plekhb2 | 0.039479 | 0.661641 | 1 |
| Caap1 | 0.039472 | 0.250288 | 1 |
| Ice1 | 0.039468 | 0.40491 | 1 |
| Dusp11 | 0.039466 | 0.724677 | 1 |
| Akap11 | 0.039374 | 0.63937 | 1 |
| Pigk | 0.039373 | 0.447802 | 1 |
| Lactb | 0.039344 | 0.274258 | 1 |
| Ciao3 | 0.039312 | 0.435403 | 1 |
| Impa1 | 0.039311 | 0.451873 | 1 |
| Serinc1 | 0.039306 | 0.863478 | 1 |
| Eci1 | 0.039304 | 0.037973 | 1 |
| Kctd13 | 0.039268 | 0.386167 | 1 |
| Ccdc97 | 0.039107 | 0.574489 | 1 |
| Med24 | 0.039076 | 0.760406 | 1 |
| Bdp1 | 0.039023 | 0.7963 | 1 |
| Anks1 | 0.038987 | 0.891811 | 1 |
| Med11 | 0.038939 | 0.749473 | 1 |
| Parg | 0.038889 | 0.896142 | 1 |
| Marf1 | 0.038874 | 0.965188 | 1 |
| Pigm | 0.038869 | 0.665326 | 1 |
| Trmt10c | 0.038752 | 0.678667 | 1 |
| Sesn3 | 0.038745 | 0.159417 | 1 |
| Suds3 | 0.03872 | 0.49646 | 1 |
| Aimp1 | 0.038717 | 0.89953 | 1 |
| Ten1 | 0.038677 | 0.606714 | 1 |
| Agpat2 | 0.038675 | 0.258006 | 1 |
| Asl | 0.038661 | 0.222342 | 1 |
| Gosr2 | 0.038642 | 0.749078 | 1 |
| Gm34794 | 0.038635 | 0.618325 | 1 |
| Slc25a25 | 0.038566 | 0.761016 | 1 |
| Pno1 | 0.038558 | 0.370301 | 1 |
| H13 | 0.038549 | 0.409928 | 1 |
| Mtor | 0.038526 | 0.708682 | 1 |

| | | | |
|---|---|---|---|
| Terf2 | 0.038514 | 0.827365 | 1 |
| Nedd8 | 0.038438 | 0.319423 | 1 |
| 4932438A13 | 0.038402 | 0.841787 | 1 |
| Cpeb4 | 0.038398 | 0.710274 | 1 |
| Camk2d | 0.038369 | 0.939999 | 1 |
| Sri | 0.038299 | 0.863568 | 1 |
| Fam210a | 0.038194 | 0.645297 | 1 |
| Coq9 | 0.038163 | 0.42143 | 1 |
| Pced1b | 0.038159 | 0.787211 | 1 |
| Maco1 | 0.03811 | 0.329487 | 1 |
| Gpn3 | 0.037968 | 0.734097 | 1 |
| Rrp7a | 0.037911 | 0.606853 | 1 |
| Rnf13 | 0.037892 | 0.57534 | 1 |
| Cpsf1 | 0.037788 | 0.807064 | 1 |
| Snw1 | 0.037691 | 0.371556 | 1 |
| Ndufs1 | 0.03766 | 0.6871 | 1 |
| Dcakd | 0.037575 | 0.52838 | 1 |
| Chuk | 0.037531 | 0.988096 | 1 |
| Tma16 | 0.037492 | 0.691124 | 1 |
| Zpr1 | 0.037446 | 0.397784 | 1 |
| Mtfr1 | 0.037377 | 0.506047 | 1 |
| Exosc5 | 0.03736 | 0.619467 | 1 |
| Papolg | 0.037328 | 0.842249 | 1 |
| Timm10b | 0.037289 | 0.334662 | 1 |
| Slc25a19 | 0.037264 | 0.793195 | 1 |
| Cox18 | 0.037166 | 0.592078 | 1 |
| Eif3c | 0.037157 | 0.830821 | 1 |
| Scaf11 | 0.037136 | 0.466115 | 1 |
| Sp3 | 0.037098 | 0.461363 | 1 |
| Trp53i13 | 0.037095 | 0.786984 | 1 |
| Mosmo | 0.037094 | 0.538847 | 1 |
| Zfp706 | 0.037079 | 0.448466 | 1 |
| Zfp260 | 0.036996 | 0.804279 | 1 |
| Mrpl32 | 0.036942 | 0.520325 | 1 |
| Gm35037 | 0.03693 | 0.448577 | 1 |
| Cbx6 | 0.036871 | 0.339134 | 1 |
| Zfyve26 | 0.0368 | 0.522398 | 1 |
| Tmcc1 | 0.036768 | 0.391963 | 1 |
| Vwa8 | 0.03676 | 0.418079 | 1 |
| Pradc1 | 0.036679 | 0.363012 | 1 |
| Zmat2 | 0.036596 | 0.424084 | 1 |
| Icosl | 0.036521 | 0.317009 | 1 |
| Cbx4 | 0.036495 | 0.978612 | 1 |
| Rft1 | 0.036452 | 0.411292 | 1 |
| Dnlz | 0.036409 | 0.951504 | 1 |
| Exoc1 | 0.036403 | 0.751727 | 1 |
| Stard4 | 0.036399 | 0.415091 | 1 |
| Hccs | 0.036336 | 0.451291 | 1 |

| | | | |
|---|---|---|---|
| Dimt1 | 0.036331 | 0.929516 | 1 |
| Apex2 | 0.03628 | 0.852739 | 1 |
| Mxd1 | 0.036276 | 0.233136 | 1 |
| Ostc | 0.03626 | 0.666104 | 1 |
| Fam173a | 0.036227 | 0.559182 | 1 |
| Vps26a | 0.036189 | 0.450012 | 1 |
| Tbp | 0.036167 | 0.436768 | 1 |
| Pstpip1 | 0.036151 | 0.643497 | 1 |
| Snap23 | 0.036119 | 0.921436 | 1 |
| Acadsb | 0.036117 | 0.559879 | 1 |
| Fasn | 0.036094 | 0.576941 | 1 |
| A930015D03 | 0.036073 | 0.466245 | 1 |
| Hivep1 | 0.035944 | 0.885812 | 1 |
| Smpdl3a | 0.035722 | 0.583035 | 1 |
| Pea15a | 0.035715 | 0.820124 | 1 |
| Sipa1l3 | 0.035714 | 0.800826 | 1 |
| Kmt2a | 0.035706 | 0.343166 | 1 |
| Rbm18 | 0.035691 | 0.176816 | 1 |
| Vps52 | 0.03566 | 0.973599 | 1 |
| Larp1b | 0.035598 | 0.33806 | 1 |
| Nbn | 0.035596 | 0.356073 | 1 |
| Zfand5 | 0.035591 | 0.629529 | 1 |
| Rcor1 | 0.035554 | 0.264339 | 1 |
| Styx | 0.03553 | 0.401694 | 1 |
| Otud6b | 0.035501 | 0.258663 | 1 |
| Tmem161a | 0.035494 | 0.755436 | 1 |
| Herpud2 | 0.035458 | 0.990427 | 1 |
| Tsc1 | 0.035427 | 0.858245 | 1 |
| Slc35a3 | 0.035367 | 0.887833 | 1 |
| Traf3 | 0.035311 | 0.425911 | 1 |
| Mllt6 | 0.035141 | 0.836002 | 1 |
| Mcph1 | 0.035127 | 0.581336 | 1 |
| Hsph1 | 0.035107 | 0.494312 | 1 |
| Zfp282 | 0.035106 | 0.83761 | 1 |
| Hif1an | 0.035082 | 0.367158 | 1 |
| Dexi | 0.035077 | 0.812263 | 1 |
| Erp44 | 0.035059 | 0.50537 | 1 |
| Mast3 | 0.03501 | 0.763592 | 1 |
| Dennd6a | 0.03494 | 0.374496 | 1 |
| Timm21 | 0.034892 | 0.715749 | 1 |
| Paip2 | 0.034889 | 0.419473 | 1 |
| Slc35d1 | 0.034883 | 0.920594 | 1 |
| Zmynd11 | 0.03482 | 0.694682 | 1 |
| Rab5a | 0.034796 | 0.842309 | 1 |
| Zcchc9 | 0.034775 | 0.02607 | 1 |
| mt-Nd4l | 0.034774 | 0.477501 | 1 |
| Rnf10 | 0.034743 | 0.585511 | 1 |
| Ccdc84 | 0.034691 | 0.393374 | 1 |

| | | | |
|---|---|---|---|
| Miip | 0.034582 | 0.722356 | 1 |
| Ep400 | 0.034571 | 0.747364 | 1 |
| Mrps27 | 0.034554 | 0.706324 | 1 |
| Snrnp25 | 0.034542 | 0.356938 | 1 |
| Bloc1s6 | 0.034515 | 0.727832 | 1 |
| Fem1b | 0.034493 | 0.760074 | 1 |
| C030006K11 | 0.034477 | 0.966522 | 1 |
| Smim26 | 0.034476 | 0.899059 | 1 |
| Casp8 | 0.034454 | 0.646192 | 1 |
| Edem3 | 0.034423 | 0.708102 | 1 |
| Pdpk1 | 0.034352 | 0.948711 | 1 |
| Skap2 | 0.03431 | 0.439242 | 1 |
| Sh3kbp1 | 0.034268 | 0.845817 | 1 |
| Gtf3c5 | 0.03425 | 0.492064 | 1 |
| Ppp1r14b | 0.034245 | 0.609827 | 1 |
| Cds2 | 0.034237 | 0.670017 | 1 |
| Zfp668 | 0.034212 | 0.40837 | 1 |
| Xpo7 | 0.034207 | 0.983529 | 1 |
| Hnrnpa1 | 0.034122 | 0.676738 | 1 |
| Nol6 | 0.034071 | 0.692461 | 1 |
| Zfp866 | 0.034064 | 0.735155 | 1 |
| Etf1 | 0.034061 | 0.390975 | 1 |
| Mrpl34 | 0.034059 | 0.493808 | 1 |
| Immt | 0.03404 | 0.554671 | 1 |
| Zdhhc5 | 0.033994 | 0.664232 | 1 |
| Retreg2 | 0.033983 | 0.463546 | 1 |
| Abce1 | 0.033949 | 0.506068 | 1 |
| Ercc4 | 0.033866 | 0.962755 | 1 |
| Map3k7 | 0.033812 | 0.770428 | 1 |
| Coa6 | 0.033786 | 0.949882 | 1 |
| Rad54l2 | 0.033751 | 0.71379 | 1 |
| Mrps24 | 0.033643 | 0.972481 | 1 |
| Jade2 | 0.0336 | 0.488525 | 1 |
| Ntpcr | 0.033467 | 0.412229 | 1 |
| Klf6 | 0.033458 | 0.991125 | 1 |
| Polr3f | 0.033421 | 0.837251 | 1 |
| Ddx58 | 0.033371 | 0.987582 | 1 |
| Ubfd1 | 0.033363 | 0.632835 | 1 |
| Tmem183a | 0.033363 | 0.736998 | 1 |
| Mak16 | 0.033323 | 0.887412 | 1 |
| Dnajc2 | 0.033272 | 0.966087 | 1 |
| Gdap2 | 0.033252 | 0.559612 | 1 |
| Gm5914 | 0.033207 | 0.556426 | 1 |
| Gabarapl1 | 0.033178 | 0.861976 | 1 |
| Coq3 | 0.033164 | 0.623017 | 1 |
| Ubr3 | 0.033137 | 0.424782 | 1 |
| Gpsm3 | 0.033133 | 0.745029 | 1 |
| Tmem50b | 0.033055 | 0.43767 | 1 |

| | | | |
|---|---|---|---|
| Mrps9 | 0.033026 | 0.566123 | 1 |
| Brf2 | 0.03301 | 0.511527 | 1 |
| Mrm1 | 0.033 | 0.867965 | 1 |
| Zc3h10 | 0.032995 | 0.314127 | 1 |
| Dclre1c | 0.032892 | 0.848578 | 1 |
| Ddx20 | 0.032879 | 0.762975 | 1 |
| Ibtk | 0.032841 | 0.888617 | 1 |
| Kcna3 | 0.032829 | 0.606767 | 1 |
| Pnkp | 0.03277 | 0.964957 | 1 |
| Tomm7 | 0.032705 | 0.75692 | 1 |
| Clpp | 0.032688 | 0.809936 | 1 |
| Mtg1 | 0.032661 | 0.727753 | 1 |
| Flcn | 0.032634 | 0.521158 | 1 |
| Mri1 | 0.032599 | 0.586716 | 1 |
| Cmc1 | 0.032529 | 0.809081 | 1 |
| Cep162 | 0.032505 | 0.92537 | 1 |
| Uqcc1 | 0.032482 | 0.855724 | 1 |
| Gpatch2 | 0.032449 | 0.835769 | 1 |
| Sys1 | 0.032415 | 0.221505 | 1 |
| Rpap3 | 0.032332 | 0.961435 | 1 |
| Il21r | 0.03221 | 0.660131 | 1 |
| Exo5 | 0.03217 | 0.477249 | 1 |
| Taf1a | 0.032138 | 0.426964 | 1 |
| Qdpr | 0.032126 | 0.963983 | 1 |
| Cul5 | 0.032121 | 0.529373 | 1 |
| 2700097O09 | 0.032015 | 0.809516 | 1 |
| Higd2a | 0.031878 | 0.72578 | 1 |
| Alkbh1 | 0.031844 | 0.831207 | 1 |
| Brd9 | 0.031799 | 0.614735 | 1 |
| Suclg2 | 0.031766 | 0.526834 | 1 |
| Arhgap39 | 0.031762 | 0.214758 | 1 |
| Prr7 | 0.031752 | 0.590697 | 1 |
| Rnf138 | 0.031719 | 0.861409 | 1 |
| Spty2d1 | 0.031704 | 0.36299 | 1 |
| Lrch1 | 0.031677 | 0.867776 | 1 |
| Mrpl48 | 0.031671 | 0.77803 | 1 |
| Slc8b1 | 0.031666 | 0.866657 | 1 |
| Bpgm | 0.031611 | 0.644083 | 1 |
| Gpr180 | 0.03159 | 0.7858 | 1 |
| Prep | 0.031578 | 0.950704 | 1 |
| Yeats2 | 0.031572 | 0.884531 | 1 |
| Itfg2 | 0.031554 | 0.763142 | 1 |
| Zfp263 | 0.031506 | 0.694186 | 1 |
| Vps72 | 0.0315 | 0.934121 | 1 |
| Ppie | 0.031486 | 0.46903 | 1 |
| Paxbp1 | 0.031412 | 0.905897 | 1 |
| Fam160b2 | 0.031376 | 0.476604 | 1 |
| Micall1 | 0.031327 | 0.391983 | 1 |

| | | | |
|---|---|---|---|
| Iqce | 0.031313 | 0.85282 | 1 |
| Sppl2b | 0.031313 | 0.808979 | 1 |
| Glo1 | 0.031291 | 0.932122 | 1 |
| Tmem260 | 0.031273 | 0.624069 | 1 |
| Mettl4 | 0.031273 | 0.359756 | 1 |
| Nle1 | 0.031235 | 0.677964 | 1 |
| Polr3a | 0.031229 | 0.757099 | 1 |
| Sympk | 0.031227 | 0.993068 | 1 |
| Upf3a | 0.031186 | 0.546459 | 1 |
| Dpf2 | 0.03117 | 0.81131 | 1 |
| Cdc25a | 0.031146 | 0.666186 | 1 |
| Shmt2 | 0.031083 | 0.597892 | 1 |
| Atp2a2 | 0.031075 | 0.433527 | 1 |
| Dhx35 | 0.031072 | 0.859759 | 1 |
| Pus3 | 0.031067 | 0.620215 | 1 |
| Cox6c | 0.031032 | 0.795505 | 1 |
| Oard1 | 0.031029 | 0.97742 | 1 |
| Dhdds | 0.031001 | 0.947497 | 1 |
| Alkbh7 | 0.030905 | 0.580047 | 1 |
| Agps | 0.030826 | 0.947555 | 1 |
| Gclm | 0.030725 | 0.45883 | 1 |
| Nudt16 | 0.030724 | 0.35893 | 1 |
| Nol9 | 0.030685 | 0.375283 | 1 |
| Dhx29 | 0.030683 | 0.559937 | 1 |
| Pot1a | 0.030589 | 0.703342 | 1 |
| Mga | 0.030587 | 0.776876 | 1 |
| Timm8a1 | 0.030569 | 0.888137 | 1 |
| Thap4 | 0.030488 | 0.488142 | 1 |
| Gsr | 0.030485 | 0.254276 | 1 |
| Twf1 | 0.030474 | 0.567535 | 1 |
| Ppp2cb | 0.030468 | 0.785091 | 1 |
| Rlf | 0.030441 | 0.281433 | 1 |
| Asna1 | 0.030397 | 0.885719 | 1 |
| Tmem179b | 0.030295 | 0.45918 | 1 |
| Whamm | 0.030236 | 0.944216 | 1 |
| Dnmt3a | 0.030205 | 0.776403 | 1 |
| Slc17a9 | 0.030202 | 0.688496 | 1 |
| Tsfm | 0.030197 | 0.998198 | 1 |
| Kpna1 | 0.030075 | 0.666563 | 1 |
| Prickle3 | 0.030052 | 0.319957 | 1 |
| Arhgap12 | 0.030031 | 0.446557 | 1 |
| Ccdc171 | 0.02983 | 0.753823 | 1 |
| Uhmk1 | 0.029821 | 0.940959 | 1 |
| Cyth2 | 0.029734 | 0.912572 | 1 |
| Arpc1a | 0.029718 | 0.209544 | 1 |
| Rpl39 | 0.029706 | 0.521858 | 1 |
| Txndc11 | 0.029681 | 0.858022 | 1 |
| Retreg3 | 0.029657 | 0.738782 | 1 |

| | | | |
|---|---|---|---|
| Tacc2 | 0.029583 | 0.372522 | 1 |
| Atg4b | 0.029546 | 0.905784 | 1 |
| Arfgef2 | 0.029544 | 0.658267 | 1 |
| Pik3r4 | 0.029417 | 0.933308 | 1 |
| Xcl1 | 0.029384 | 0.211936 | 1 |
| Naa60 | 0.029186 | 0.973717 | 1 |
| Nploc4 | 0.029162 | 0.59131 | 1 |
| Sap30bp | 0.029159 | 0.572716 | 1 |
| Snapc1 | 0.029126 | 0.803718 | 1 |
| Cnot4 | 0.029125 | 0.748096 | 1 |
| Zfp983 | 0.029092 | 0.742023 | 1 |
| Pigf | 0.029049 | 0.936205 | 1 |
| Krit1 | 0.029014 | 0.42807 | 1 |
| Phf7 | 0.028988 | 0.847421 | 1 |
| Acadvl | 0.028963 | 0.986162 | 1 |
| Gm4876 | 0.028875 | 0.818144 | 1 |
| Kmt2d | 0.028862 | 0.512405 | 1 |
| Uap1 | 0.028833 | 0.331782 | 1 |
| Pi4ka | 0.02874 | 0.79812 | 1 |
| Sap130 | 0.028722 | 0.564657 | 1 |
| Mpst | 0.028638 | 0.873602 | 1 |
| Zdhhc13 | 0.028614 | 0.810236 | 1 |
| Gfi1 | 0.028584 | 0.99388 | 1 |
| Ahsa2 | 0.028575 | 0.887787 | 1 |
| Hnrnph2 | 0.028509 | 0.52017 | 1 |
| Gt(ROSA)26Sc | 0.028471 | 0.460378 | 1 |
| Ogg1 | 0.028421 | 0.428003 | 1 |
| Ube2q2 | 0.028402 | 0.40572 | 1 |
| Pcgf6 | 0.028386 | 0.58367 | 1 |
| Rpap1 | 0.028358 | 0.637239 | 1 |
| Sqle | 0.028333 | 0.425265 | 1 |
| Stx17 | 0.028318 | 0.810499 | 1 |
| Ankrd54 | 0.02828 | 0.345951 | 1 |
| Nat2 | 0.028238 | 0.620753 | 1 |
| Ppcdc | 0.028174 | 0.979976 | 1 |
| Snx27 | 0.028173 | 0.640992 | 1 |
| 0610030E20I | 0.028172 | 0.440719 | 1 |
| Ilrun | 0.028135 | 0.392133 | 1 |
| Wdr61 | 0.028127 | 0.813618 | 1 |
| Zfp410 | 0.028127 | 0.913714 | 1 |
| Nkapd1 | 0.028087 | 0.928622 | 1 |
| Nceh1 | 0.028075 | 0.647878 | 1 |
| Hint3 | 0.028068 | 0.815412 | 1 |
| Alg3 | 0.028009 | 0.723163 | 1 |
| 5830428M24 | 0.028009 | 0.887365 | 1 |
| Topors | 0.027966 | 0.851346 | 1 |
| Eif1ad | 0.027897 | 0.835678 | 1 |
| Sem1 | 0.027875 | 0.916426 | 1 |

| | | | |
|---|---|---|---|
| Cmas | 0.027825 | 0.65162 | 1 |
| Tmem223 | 0.027785 | 0.63521 | 1 |
| Polr1e | 0.027752 | 0.890519 | 1 |
| Dera | 0.02774 | 0.591405 | 1 |
| Mipol1 | 0.027679 | 0.50563 | 1 |
| 0610009B22 | 0.027571 | 0.653685 | 1 |
| Glrx2 | 0.027569 | 0.566646 | 1 |
| Mphosph6 | 0.027558 | 0.747878 | 1 |
| Nacc1 | 0.027502 | 0.941367 | 1 |
| Rnasek | 0.027378 | 0.205846 | 1 |
| Taco1 | 0.02731 | 0.921853 | 1 |
| Rab10 | 0.027304 | 0.448986 | 1 |
| Msra | 0.027274 | 0.794669 | 1 |
| Sumo3 | 0.027273 | 0.143637 | 1 |
| Zfp84 | 0.027191 | 0.802764 | 1 |
| Sergef | 0.027177 | 0.12512 | 1 |
| Rnf25 | 0.027173 | 0.758557 | 1 |
| Heatr5a | 0.02711 | 0.245948 | 1 |
| Btg3 | 0.027089 | 0.533688 | 1 |
| Echs1 | 0.027079 | 0.632444 | 1 |
| M6pr | 0.027045 | 0.531618 | 1 |
| Dyrk2 | 0.027003 | 0.866824 | 1 |
| Slc35a2 | 0.026992 | 0.979019 | 1 |
| Jade1 | 0.026859 | 0.367814 | 1 |
| Gm16033 | 0.026841 | 0.868412 | 1 |
| Ythdf3 | 0.026818 | 0.412473 | 1 |
| Srrm1 | 0.026785 | 0.953328 | 1 |
| Tbc1d23 | 0.026783 | 0.60015 | 1 |
| Coq7 | 0.026707 | 0.363193 | 1 |
| Gm15821 | 0.026687 | 0.953165 | 1 |
| Lzic | 0.026652 | 0.629705 | 1 |
| Atr | 0.026637 | 0.431701 | 1 |
| Emc6 | 0.026597 | 0.961779 | 1 |
| Cars2 | 0.026517 | 0.693652 | 1 |
| Ranbp9 | 0.026515 | 0.956035 | 1 |
| Fam53c | 0.026503 | 0.840364 | 1 |
| Entr1 | 0.026419 | 0.28699 | 1 |
| Smad2 | 0.026401 | 0.793405 | 1 |
| Gosr1 | 0.026377 | 0.917463 | 1 |
| Vamp4 | 0.026308 | 0.691481 | 1 |
| Hmgcl | 0.026293 | 0.951379 | 1 |
| Rab8a | 0.026266 | 0.903428 | 1 |
| Synrg | 0.026248 | 0.847082 | 1 |
| Sel1l | 0.026244 | 0.538414 | 1 |
| Narf | 0.026186 | 0.563859 | 1 |
| Cnih1 | 0.026123 | 0.086631 | 1 |
| Rbm47 | 0.026113 | 0.735663 | 1 |
| Med27 | 0.026107 | 0.934726 | 1 |

| | | | |
|---|---|---|---|
| Mrpl36 | 0.026082 | 0.818922 | 1 |
| Isy1 | 0.02598 | 0.907494 | 1 |
| Nop9 | 0.02593 | 0.293449 | 1 |
| Nrbf2 | 0.025908 | 0.800614 | 1 |
| Zdhhc7 | 0.025799 | 0.653115 | 1 |
| Ncoa7 | 0.025709 | 0.793566 | 1 |
| Tfb2m | 0.025689 | 0.618021 | 1 |
| Sbno2 | 0.025653 | 0.854877 | 1 |
| Ltn1 | 0.02563 | 0.802832 | 1 |
| Pus1 | 0.025597 | 0.931655 | 1 |
| Jakmip1 | 0.025596 | 0.896794 | 1 |
| Spopl | 0.025575 | 0.553482 | 1 |
| Ing5 | 0.025564 | 0.393694 | 1 |
| Trappc13 | 0.02552 | 0.888111 | 1 |
| Etnk1 | 0.02552 | 0.766968 | 1 |
| Aifm1 | 0.025479 | 0.611572 | 1 |
| Ccdc134 | 0.025473 | 0.791351 | 1 |
| Ctdsp2 | 0.025469 | 0.430399 | 1 |
| Aopep | 0.025446 | 0.3668 | 1 |
| Atad3a | 0.025357 | 0.309399 | 1 |
| Fbxo32 | 0.025328 | 0.672318 | 1 |
| Rexo2 | 0.025296 | 0.467952 | 1 |
| Cradd | 0.025241 | 0.609967 | 1 |
| 2810004N23 | 0.025234 | 0.78856 | 1 |
| Mettl21a | 0.025205 | 0.422943 | 1 |
| Mtpap | 0.025145 | 0.601356 | 1 |
| Sacs | 0.025103 | 0.600273 | 1 |
| Coq6 | 0.02501 | 0.787425 | 1 |
| Ighmbp2 | 0.024996 | 0.618577 | 1 |
| Tgfbr2 | 0.024974 | 0.917743 | 1 |
| Nsd3 | 0.024956 | 0.576626 | 1 |
| Rnf38 | 0.024824 | 0.985603 | 1 |
| Fbrs | 0.024821 | 0.279357 | 1 |
| Zfp740 | 0.024782 | 0.830187 | 1 |
| Tspo | 0.024654 | 0.945157 | 1 |
| Fam214a | 0.024613 | 0.575832 | 1 |
| Unkl | 0.024595 | 0.670912 | 1 |
| Tut1 | 0.024534 | 0.377987 | 1 |
| Pold3 | 0.024504 | 0.8195 | 1 |
| Kars | 0.024503 | 0.705059 | 1 |
| Psmd2 | 0.024486 | 0.494229 | 1 |
| Nrd1 | 0.024449 | 0.930171 | 1 |
| Sacm1l | 0.024444 | 0.966582 | 1 |
| Hcfc2 | 0.02443 | 0.909227 | 1 |
| Zfand1 | 0.024411 | 0.777939 | 1 |
| Ndufb5 | 0.02439 | 0.954803 | 1 |
| Ncoa1 | 0.024347 | 0.697211 | 1 |
| Pisd | 0.024157 | 0.950333 | 1 |

| | | | |
|---|---|---|---|
| Nr2c2ap | 0.024155 | 0.572213 | 1 |
| Hmbs | 0.024117 | 0.917695 | 1 |
| Vmn2r96 | 0.024104 | 0.652253 | 1 |
| Hdhd5 | 0.024084 | 0.757895 | 1 |
| Pan2 | 0.02405 | 0.599353 | 1 |
| Lcmt1 | 0.024049 | 0.563412 | 1 |
| Rundc1 | 0.02399 | 0.868714 | 1 |
| Trp53bp1 | 0.023939 | 0.744035 | 1 |
| Mrps25 | 0.023922 | 0.873282 | 1 |
| Zbtb44 | 0.02389 | 0.643484 | 1 |
| Scnm1 | 0.023848 | 0.447776 | 1 |
| Zfp639 | 0.023837 | 0.617868 | 1 |
| Rnf40 | 0.023801 | 0.961993 | 1 |
| Pptc7 | 0.023777 | 0.948459 | 1 |
| Mms19 | 0.023614 | 0.83804 | 1 |
| Nlrc3 | 0.023569 | 0.901959 | 1 |
| Eloc | 0.023536 | 0.959376 | 1 |
| Pdcd5 | 0.023527 | 0.600587 | 1 |
| 4930469K13 | 0.023522 | 0.680788 | 1 |
| 5330438D12 | 0.023498 | 0.650284 | 1 |
| Gtf2h1 | 0.023448 | 0.870498 | 1 |
| Foxred1 | 0.023346 | 0.614271 | 1 |
| Gvin1 | 0.023314 | 0.852599 | 1 |
| Qars | 0.023247 | 0.936309 | 1 |
| 1110059G10 | 0.023166 | 0.977605 | 1 |
| Yipf6 | 0.023124 | 0.800779 | 1 |
| Klhdc3 | 0.023036 | 0.920763 | 1 |
| Acsl5 | 0.023013 | 0.690583 | 1 |
| Elp3 | 0.023005 | 0.838545 | 1 |
| Dcp1a | 0.022993 | 0.690903 | 1 |
| Actr5 | 0.022935 | 0.874739 | 1 |
| Zmym4 | 0.02287 | 0.658587 | 1 |
| Mad2l2 | 0.022865 | 0.755156 | 1 |
| Impa2 | 0.022838 | 0.425252 | 1 |
| Pfdn4 | 0.022805 | 0.87656 | 1 |
| Ngdn | 0.022712 | 0.901701 | 1 |
| Psmd1 | 0.022706 | 0.841749 | 1 |
| Hnrnph3 | 0.022687 | 0.688833 | 1 |
| Tcf20 | 0.022606 | 0.801467 | 1 |
| Elk3 | 0.022564 | 0.937438 | 1 |
| Rftn1 | 0.022563 | 0.271493 | 1 |
| Mrpl27 | 0.022514 | 0.790603 | 1 |
| Ssh1 | 0.0225 | 0.960987 | 1 |
| Gtf2f1 | 0.022449 | 0.761075 | 1 |
| Pms1 | 0.022407 | 0.823871 | 1 |
| Sdr39u1 | 0.022399 | 0.792392 | 1 |
| Slc39a3 | 0.022366 | 0.777302 | 1 |
| D3Ertd751e | 0.022359 | 0.866065 | 1 |

| | | | |
|---|---|---|---|
| Exoc5 | 0.022275 | 0.42068 | 1 |
| Ankrd13a | 0.022203 | 0.730209 | 1 |
| Zfp687 | 0.022074 | 0.791429 | 1 |
| Pom121 | 0.02205 | 0.357539 | 1 |
| Rrm2b | 0.022038 | 0.797344 | 1 |
| Polr2b | 0.022038 | 0.817103 | 1 |
| Mynn | 0.022019 | 0.584292 | 1 |
| Ybx1 | 0.022001 | 0.713257 | 1 |
| Ccdc51 | 0.021917 | 0.541004 | 1 |
| Cbx8 | 0.021861 | 0.691921 | 1 |
| Zfp142 | 0.021852 | 0.833226 | 1 |
| Tmed5 | 0.021835 | 0.985785 | 1 |
| Rbm8a | 0.021821 | 0.820735 | 1 |
| Mfap1a | 0.021814 | 0.837775 | 1 |
| Prmt9 | 0.021806 | 0.744939 | 1 |
| Rrnad1 | 0.021769 | 0.82742 | 1 |
| Nf2 | 0.021726 | 0.715803 | 1 |
| Acbd4 | 0.021713 | 0.637094 | 1 |
| Sirt3 | 0.021693 | 0.278926 | 1 |
| Rnaseh2a | 0.0216 | 0.233183 | 1 |
| Brms1l | 0.021595 | 0.750462 | 1 |
| Mgat4a | 0.021571 | 0.851054 | 1 |
| Cdk19 | 0.02154 | 0.992747 | 1 |
| Emsy | 0.021534 | 0.68764 | 1 |
| Csgalnact2 | 0.021497 | 0.503111 | 1 |
| Ddhd2 | 0.02142 | 0.718925 | 1 |
| Rbmxl1 | 0.0214 | 0.393695 | 1 |
| Eef1e1 | 0.021395 | 0.914415 | 1 |
| Ppid | 0.021379 | 0.985646 | 1 |
| Mfhas1 | 0.021296 | 0.587811 | 1 |
| Ift20 | 0.021281 | 0.631142 | 1 |
| Pafah1b2 | 0.021273 | 0.356285 | 1 |
| Polr3gl | 0.021257 | 0.902906 | 1 |
| Rnf19b | 0.021212 | 0.496964 | 1 |
| Strada | 0.02105 | 0.98937 | 1 |
| Mon1b | 0.021041 | 0.735712 | 1 |
| Gm28935 | 0.021013 | 0.942734 | 1 |
| Epn1 | 0.02101 | 0.232368 | 1 |
| Vkorc1 | 0.020933 | 0.092602 | 1 |
| Rrp36 | 0.02092 | 0.926412 | 1 |
| Eif1ax | 0.020831 | 0.738519 | 1 |
| Fam160b1 | 0.020809 | 0.929455 | 1 |
| Pcnx3 | 0.020787 | 0.685904 | 1 |
| Nfs1 | 0.020776 | 0.963549 | 1 |
| Brd7 | 0.020763 | 0.986386 | 1 |
| Tmtc4 | 0.020729 | 0.775811 | 1 |
| Paqr7 | 0.020724 | 0.457489 | 1 |
| 2510002D24 | 0.0207 | 0.517654 | 1 |

| | | | |
|---|---|---|---|
| Tmem208 | 0.020687 | 0.760074 | 1 |
| Fgd3 | 0.020685 | 0.988838 | 1 |
| Map4k4 | 0.020676 | 0.28519 | 1 |
| Tnfrsf1b | 0.020673 | 0.963944 | 1 |
| Zfp444 | 0.02064 | 0.960757 | 1 |
| Dram2 | 0.020626 | 0.672221 | 1 |
| Syap1 | 0.020617 | 0.646581 | 1 |
| Dolpp1 | 0.020594 | 0.940911 | 1 |
| Ankrd52 | 0.020572 | 0.769347 | 1 |
| Pabpc4 | 0.020542 | 0.778865 | 1 |
| Xrcc5 | 0.020481 | 0.778402 | 1 |
| Cops6 | 0.020459 | 0.624292 | 1 |
| Dctn4 | 0.020452 | 0.569622 | 1 |
| Golph3l | 0.020404 | 0.939374 | 1 |
| Ttc7b | 0.020355 | 0.830497 | 1 |
| Mrtfb | 0.02032 | 0.712706 | 1 |
| Tmem219 | 0.020296 | 0.777584 | 1 |
| Slc25a13 | 0.020177 | 0.97907 | 1 |
| Zfp524 | 0.020142 | 0.856595 | 1 |
| Mien1 | 0.02013 | 0.829541 | 1 |
| Plekhm1 | 0.020088 | 0.320175 | 1 |
| Tti2 | 0.02005 | 0.637605 | 1 |
| Tab1 | 0.02005 | 0.777237 | 1 |
| Tmx4 | 0.020021 | 0.572843 | 1 |
| Hip1r | 0.019972 | 0.54881 | 1 |
| Enox2 | 0.019897 | 0.931461 | 1 |
| Rad51d | 0.019757 | 0.847355 | 1 |
| Ywhag | 0.019728 | 0.880314 | 1 |
| Tpgs1 | 0.019587 | 0.476321 | 1 |
| Mospd2 | 0.019569 | 0.723131 | 1 |
| Rpp21 | 0.019557 | 0.534435 | 1 |
| Qrsl1 | 0.019484 | 0.378067 | 1 |
| Gtf2ird2 | 0.019474 | 0.854916 | 1 |
| Stk24 | 0.019393 | 0.988965 | 1 |
| Zcchc8 | 0.019368 | 0.984039 | 1 |
| Farsb | 0.019365 | 0.724552 | 1 |
| Dffb | 0.019345 | 0.995915 | 1 |
| Fam98a | 0.019311 | 0.840026 | 1 |
| Fam114a2 | 0.019276 | 0.958415 | 1 |
| Ift80 | 0.019256 | 0.93635 | 1 |
| Fbxo46 | 0.019233 | 0.858587 | 1 |
| Gm3448 | 0.019209 | 0.493543 | 1 |
| Usp7 | 0.019068 | 0.969105 | 1 |
| Dcaf11 | 0.019008 | 0.795559 | 1 |
| Mrpl57 | 0.018953 | 0.930255 | 1 |
| Gpr174 | 0.018911 | 0.920358 | 1 |
| Ptar1 | 0.018864 | 0.466718 | 1 |
| Polr2j | 0.018815 | 0.788823 | 1 |

| | | | |
|---|---|---|---|
| Igf2r | 0.018715 | 0.743989 | 1 |
| Pik3ap1 | 0.018701 | 0.950775 | 1 |
| Siae | 0.018693 | 0.815153 | 1 |
| Arl6ip4 | 0.018661 | 0.207313 | 1 |
| Mul1 | 0.018602 | 0.875238 | 1 |
| Immp1l | 0.018597 | 0.989678 | 1 |
| Cactin | 0.018582 | 0.621559 | 1 |
| Ttc9c | 0.018565 | 0.681921 | 1 |
| Zfp346 | 0.018434 | 0.516438 | 1 |
| Mphosph8 | 0.018428 | 0.560236 | 1 |
| Nfkbie | 0.018389 | 0.265601 | 1 |
| Eif2ak4 | 0.018383 | 0.440898 | 1 |
| Slc39a9 | 0.018287 | 0.765391 | 1 |
| 2310033P09I | 0.018273 | 0.63244 | 1 |
| Clk1 | 0.018255 | 0.725656 | 1 |
| Sh3bp2 | 0.018244 | 0.844863 | 1 |
| Prmt1 | 0.018242 | 0.391148 | 1 |
| Ankib1 | 0.018202 | 0.736064 | 1 |
| Gtf2e1 | 0.018202 | 0.697233 | 1 |
| Zfp322a | 0.018189 | 0.876777 | 1 |
| Pogz | 0.018178 | 0.53391 | 1 |
| Zcchc4 | 0.018172 | 0.898549 | 1 |
| Czib | 0.018172 | 0.920268 | 1 |
| Man2b2 | 0.018153 | 0.917306 | 1 |
| Mphosph10 | 0.018129 | 0.850525 | 1 |
| Gemin2 | 0.018096 | 0.389873 | 1 |
| Trbc1 | 0.017963 | 0.275472 | 1 |
| Nus1 | 0.017944 | 0.658792 | 1 |
| Ctdp1 | 0.017939 | 0.951616 | 1 |
| Irf3 | 0.017934 | 0.790462 | 1 |
| Platr25 | 0.01787 | 0.47019 | 1 |
| Ubp1 | 0.017848 | 0.838461 | 1 |
| Atxn7l1 | 0.017835 | 0.495554 | 1 |
| Clcn7 | 0.0178 | 0.445224 | 1 |
| Gm2245 | 0.017754 | 0.770814 | 1 |
| Sec22b | 0.017711 | 0.852618 | 1 |
| Nipa2 | 0.017634 | 0.654796 | 1 |
| Smim14 | 0.017573 | 0.631417 | 1 |
| Txndc17 | 0.017563 | 0.368939 | 1 |
| Btbd8 | 0.017553 | 0.631894 | 1 |
| Pum3 | 0.017526 | 0.843844 | 1 |
| Stx8 | 0.017524 | 0.990531 | 1 |
| Ccdc174 | 0.01748 | 0.880656 | 1 |
| Prmt2 | 0.017479 | 0.394526 | 1 |
| Ankrd11 | 0.017438 | 0.704514 | 1 |
| Med29 | 0.017412 | 0.799305 | 1 |
| Neil1 | 0.017372 | 0.550312 | 1 |
| Cdk12 | 0.017325 | 0.69042 | 1 |

| | | | |
|---|---|---|---|
| Lonp1 | 0.017318 | 0.490442 | 1 |
| Mybbp1a | 0.017259 | 0.336057 | 1 |
| Bcl2l1 | 0.017236 | 0.678877 | 1 |
| Ppa2 | 0.017136 | 0.652029 | 1 |
| Usf3 | 0.017135 | 0.889514 | 1 |
| Rpe | 0.017113 | 0.890135 | 1 |
| Ip6k2 | 0.017109 | 0.605428 | 1 |
| Rprd1b | 0.017088 | 0.681745 | 1 |
| Ftx | 0.017087 | 0.832397 | 1 |
| Tbccd1 | 0.01707 | 0.693979 | 1 |
| Tom1 | 0.017065 | 0.731742 | 1 |
| Calhm2 | 0.017017 | 0.829024 | 1 |
| Tsc22d3 | 0.016823 | 0.219073 | 1 |
| Cstb | 0.016784 | 0.683297 | 1 |
| Pcgf1 | 0.016757 | 0.699773 | 1 |
| Mvb12b | 0.016755 | 0.23916 | 1 |
| Mrpl16 | 0.016733 | 0.88314 | 1 |
| Fbxo21 | 0.01669 | 0.916837 | 1 |
| Psmd13 | 0.016592 | 0.874971 | 1 |
| Dcxr | 0.01657 | 0.994564 | 1 |
| Borcs6 | 0.016456 | 0.63522 | 1 |
| Lrig2 | 0.016452 | 0.494005 | 1 |
| Wdr41 | 0.016434 | 0.039628 | 1 |
| Prelid2 | 0.01643 | 0.819829 | 1 |
| Pdcl3 | 0.016427 | 0.975361 | 1 |
| Otud7b | 0.016381 | 0.714613 | 1 |
| Atp5md | 0.016359 | 0.998324 | 1 |
| Tada2a | 0.016353 | 0.385343 | 1 |
| Atg2b | 0.016339 | 0.984957 | 1 |
| Pacs1 | 0.016324 | 0.918706 | 1 |
| Etfdh | 0.016311 | 0.780416 | 1 |
| Eif3g | 0.016229 | 0.844363 | 1 |
| Tmem70 | 0.016195 | 0.511443 | 1 |
| Tnip1 | 0.016176 | 0.935247 | 1 |
| Rps6ka4 | 0.01617 | 0.685081 | 1 |
| Patz1 | 0.01612 | 0.851687 | 1 |
| Rnf11 | 0.01611 | 0.773876 | 1 |
| Bud23 | 0.016053 | 0.678794 | 1 |
| Srsf4 | 0.016044 | 0.812975 | 1 |
| Dvl2 | 0.01598 | 0.748287 | 1 |
| Ubl5 | 0.015972 | 0.901382 | 1 |
| Ahdc1 | 0.015881 | 0.935356 | 1 |
| Dnajb1 | 0.015852 | 0.798063 | 1 |
| Ripk3 | 0.015766 | 0.707653 | 1 |
| Map2k7 | 0.015707 | 0.93275 | 1 |
| 5430427O19 | 0.015671 | 0.151453 | 1 |
| Top1 | 0.015649 | 0.964085 | 1 |
| 2510039O18 | 0.01556 | 0.429382 | 1 |

| | | | |
|---|---|---|---|
| Klhl25 | 0.015539 | 0.896633 | 1 |
| Trmt2a | 0.01545 | 0.887735 | 1 |
| Vps37c | 0.015435 | 0.651797 | 1 |
| Tubgcp6 | 0.015343 | 0.711027 | 1 |
| Upf3b | 0.015339 | 0.870856 | 1 |
| Bcat2 | 0.015337 | 0.727605 | 1 |
| Llgl1 | 0.015336 | 0.697744 | 1 |
| Gnpnat1 | 0.015316 | 0.592908 | 1 |
| Atl3 | 0.015315 | 0.349922 | 1 |
| Herc2 | 0.015243 | 0.947643 | 1 |
| Fgfr1op | 0.015222 | 0.870937 | 1 |
| Xdh | 0.01514 | 0.683972 | 1 |
| Ankle2 | 0.01507 | 0.923943 | 1 |
| Eif2a | 0.01506 | 0.999107 | 1 |
| Slirp | 0.014958 | 0.841884 | 1 |
| Sin3b | 0.014908 | 0.597031 | 1 |
| Trim26 | 0.014893 | 0.796988 | 1 |
| Pym1 | 0.014885 | 0.602664 | 1 |
| Ehmt2 | 0.014842 | 0.759348 | 1 |
| Cep63 | 0.014811 | 0.732106 | 1 |
| Commd9 | 0.014808 | 0.775439 | 1 |
| Tmem245 | 0.014804 | 0.307785 | 1 |
| Serpine2 | 0.014739 | 0.077892 | 1 |
| Brwd1 | 0.014728 | 0.732354 | 1 |
| Ikbke | 0.014608 | 0.938568 | 1 |
| Vps45 | 0.014595 | 0.86601 | 1 |
| Mrpl1 | 0.014375 | 0.964903 | 1 |
| Ube2j1 | 0.014362 | 0.903879 | 1 |
| Atp6v0c | 0.014358 | 0.910147 | 1 |
| Prr14 | 0.014356 | 0.789738 | 1 |
| Tmx3 | 0.014354 | 0.71947 | 1 |
| Ccdc127 | 0.014352 | 0.775179 | 1 |
| Pias2 | 0.014264 | 0.87298 | 1 |
| Ppif | 0.014253 | 0.973763 | 1 |
| Atraid | 0.01418 | 0.644483 | 1 |
| Spout1 | 0.014113 | 0.980693 | 1 |
| Spast | 0.014112 | 0.83257 | 1 |
| Oxct1 | 0.014079 | 0.999823 | 1 |
| Mysm1 | 0.014075 | 0.972465 | 1 |
| Zfp146 | 0.014031 | 0.396804 | 1 |
| Atg16l1 | 0.014023 | 0.821449 | 1 |
| Cnst | 0.014004 | 0.193972 | 1 |
| Chm | 0.013967 | 0.854204 | 1 |
| Paxx | 0.013953 | 0.954094 | 1 |
| Eef2kmt | 0.013952 | 0.589533 | 1 |
| Pkn1 | 0.013915 | 0.235932 | 1 |
| Emc1 | 0.013843 | 0.913999 | 1 |
| Rbm7 | 0.013841 | 0.756059 | 1 |

| | | | |
|---|---|---|---|
| Dkc1 | 0.013802 | 0.962068 | 1 |
| Neu1 | 0.013794 | 0.243312 | 1 |
| Zbtb9 | 0.013779 | 0.687827 | 1 |
| Letmd1 | 0.013751 | 0.957829 | 1 |
| Prpsap2 | 0.013729 | 0.88067 | 1 |
| Dynlt1c | 0.013714 | 0.948395 | 1 |
| Cib1 | 0.013699 | 0.910383 | 1 |
| Rbfa | 0.013696 | 0.339064 | 1 |
| Vav3 | 0.01361 | 0.768536 | 1 |
| Pvt1 | 0.013604 | 0.781045 | 1 |
| Cox16 | 0.01352 | 0.569055 | 1 |
| Fadd | 0.013505 | 0.963007 | 1 |
| Tmem185b | 0.013445 | 0.77605 | 1 |
| Mterf1a | 0.013432 | 0.400271 | 1 |
| Dnase2a | 0.013397 | 0.492355 | 1 |
| Zfp335 | 0.013395 | 0.473833 | 1 |
| Trp53i11 | 0.013345 | 0.60591 | 1 |
| Nt5c2 | 0.013345 | 0.913826 | 1 |
| Nt5c3 | 0.01333 | 0.333707 | 1 |
| Kcnn4 | 0.013286 | 0.488494 | 1 |
| Prdx2 | 0.013268 | 0.450525 | 1 |
| C1galt1c1 | 0.013232 | 0.994203 | 1 |
| Mrto4 | 0.013212 | 0.998543 | 1 |
| Smad4 | 0.013175 | 0.430303 | 1 |
| Rab3ip | 0.013151 | 0.982766 | 1 |
| Crybg3 | 0.013136 | 0.383941 | 1 |
| Dld | 0.013135 | 0.816793 | 1 |
| Sik1 | 0.01306 | 0.189645 | 1 |
| Golga2 | 0.013043 | 0.828803 | 1 |
| Pik3c3 | 0.013026 | 0.706331 | 1 |
| Ada | 0.013021 | 0.507007 | 1 |
| Otulin | 0.012982 | 0.806833 | 1 |
| Aga | 0.012873 | 0.974027 | 1 |
| Zswim1 | 0.01282 | 0.947789 | 1 |
| Tspan14 | 0.012794 | 0.391652 | 1 |
| Gpank1 | 0.01278 | 0.731483 | 1 |
| Rmnd1 | 0.012774 | 0.784966 | 1 |
| Ppp1r8 | 0.012762 | 0.884752 | 1 |
| Tnpo3 | 0.012748 | 0.912654 | 1 |
| Mettl14 | 0.012695 | 0.897517 | 1 |
| Thap1 | 0.01269 | 0.701634 | 1 |
| Ube2w | 0.012659 | 0.721811 | 1 |
| AI413582 | 0.01262 | 0.637983 | 1 |
| Ddx28 | 0.012574 | 0.952061 | 1 |
| Kat8 | 0.012523 | 0.176135 | 1 |
| Armc8 | 0.012509 | 0.956961 | 1 |
| Acap2 | 0.012476 | 0.945945 | 1 |
| Nudt8 | 0.012419 | 0.577557 | 1 |

| | | | |
|---|---|---|---|
| Tjap1 | 0.012401 | 0.811446 | 1 |
| Slc25a10 | 0.012315 | 0.718447 | 1 |
| Cep41 | 0.012286 | 0.464184 | 1 |
| Ppp1r2 | 0.012264 | 0.812573 | 1 |
| Ripk1 | 0.012245 | 0.501905 | 1 |
| Mgat2 | 0.012242 | 0.693144 | 1 |
| Dnajc10 | 0.012142 | 0.795164 | 1 |
| Wdr83 | 0.012045 | 0.906045 | 1 |
| Gm16286 | 0.01204 | 0.731096 | 1 |
| Fdft1 | 0.012022 | 0.676862 | 1 |
| Ddx31 | 0.012005 | 0.806055 | 1 |
| Gm29243 | 0.011996 | 0.405836 | 1 |
| Gm9856 | 0.011993 | 0.526096 | 1 |
| Cog8 | 0.011989 | 0.659798 | 1 |
| Letm1 | 0.011977 | 0.885175 | 1 |
| Cep95 | 0.011963 | 0.673488 | 1 |
| Cdc14a | 0.011962 | 0.607739 | 1 |
| Tbc1d2b | 0.011948 | 0.899148 | 1 |
| Tpra1 | 0.011857 | 0.933491 | 1 |
| Prmt5 | 0.011836 | 0.840635 | 1 |
| Usp32 | 0.011734 | 0.692203 | 1 |
| Tbcd | 0.011727 | 0.991836 | 1 |
| Zfp871 | 0.011662 | 0.866903 | 1 |
| Clasp1 | 0.011648 | 0.9555 | 1 |
| Tssc4 | 0.011587 | 0.917117 | 1 |
| 1600010M07 | 0.011499 | 0.90392 | 1 |
| Rasa1 | 0.01149 | 0.483259 | 1 |
| Il4ra | 0.011486 | 0.809322 | 1 |
| Utp6 | 0.011365 | 0.725573 | 1 |
| Kansl3 | 0.011342 | 0.513832 | 1 |
| Wdr46 | 0.011286 | 0.828872 | 1 |
| Nars2 | 0.011274 | 0.841856 | 1 |
| Cela1 | 0.011264 | 0.880137 | 1 |
| Ubap2 | 0.011235 | 0.962233 | 1 |
| Pygo2 | 0.011189 | 0.98902 | 1 |
| Ciao2b | 0.01115 | 0.910496 | 1 |
| Tomm70a | 0.011136 | 0.621182 | 1 |
| Wdr37 | 0.011045 | 0.914633 | 1 |
| Fxr1 | 0.011023 | 0.651637 | 1 |
| Dapp1 | 0.011011 | 0.840024 | 1 |
| Cebpzos | 0.01098 | 0.18223 | 1 |
| Ss18l2 | 0.010933 | 0.877545 | 1 |
| Fnip1 | 0.010917 | 0.652414 | 1 |
| Adprhl2 | 0.010904 | 0.784926 | 1 |
| Smim7 | 0.010802 | 0.953632 | 1 |
| Tent5a | 0.010781 | 0.502216 | 1 |
| Myg1 | 0.010733 | 0.9177 | 1 |
| Ddx49 | 0.010731 | 0.910334 | 1 |

| | | | |
|---|---|---|---|
| Sft2d2 | 0.01062 | 0.836884 | 1 |
| Tfb1m | 0.010583 | 0.690822 | 1 |
| Appl1 | 0.010555 | 0.382622 | 1 |
| Rpl38 | 0.010536 | 0.825236 | 1 |
| Fam117b | 0.010503 | 0.701554 | 1 |
| Urm1 | 0.010499 | 0.270266 | 1 |
| Pja1 | 0.010387 | 0.831617 | 1 |
| Snap29 | 0.010335 | 0.494038 | 1 |
| Asb1 | 0.010286 | 0.667021 | 1 |
| Drg2 | 0.010237 | 0.665159 | 1 |
| Fam222b | 0.010212 | 0.655971 | 1 |
| Noc2l | 0.0102 | 0.589161 | 1 |
| Zswim8 | 0.0102 | 0.297366 | 1 |
| Otud4 | 0.010176 | 0.986537 | 1 |
| Cdc40 | 0.010136 | 0.140071 | 1 |
| Gemin8 | 0.010109 | 0.563424 | 1 |
| Apc | 0.010025 | 0.573717 | 1 |
| Slain2 | 0.010014 | 0.543193 | 1 |
| Zfp808 | -0.01 | 0.332212 | 1 |
| Osbp | -0.01004 | 0.653745 | 1 |
| Itgal | -0.01012 | 0.772146 | 1 |
| Prpf19 | -0.01015 | 0.538447 | 1 |
| Brap | -0.01015 | 0.616785 | 1 |
| Ddx19a | -0.01026 | 0.807524 | 1 |
| Gphn | -0.01028 | 0.377175 | 1 |
| Dicer1 | -0.01029 | 0.569382 | 1 |
| Fem1c | -0.01034 | 0.712198 | 1 |
| Map2k5 | -0.01034 | 0.537259 | 1 |
| Naga | -0.01035 | 0.460046 | 1 |
| Ap1ar | -0.01035 | 0.079419 | 1 |
| Hspa4 | -0.01041 | 0.971789 | 1 |
| Wrap73 | -0.01041 | 0.495174 | 1 |
| Ablim1 | -0.01042 | 0.956612 | 1 |
| Bms1 | -0.01042 | 0.814538 | 1 |
| Ube3b | -0.01044 | 0.583138 | 1 |
| Clk3 | -0.01044 | 0.694719 | 1 |
| Cox11 | -0.01045 | 0.520503 | 1 |
| Borcs8 | -0.01057 | 0.985993 | 1 |
| Cfap20 | -0.01064 | 0.986256 | 1 |
| Scaper | -0.01066 | 0.750069 | 1 |
| Kdm1b | -0.0107 | 0.445833 | 1 |
| Phf5a | -0.0107 | 0.831737 | 1 |
| Cdc23 | -0.01075 | 0.447675 | 1 |
| Dnajc21 | -0.01077 | 0.641607 | 1 |
| Abhd14a | -0.01086 | 0.336397 | 1 |
| Smchd1 | -0.01087 | 0.69863 | 1 |
| Dglucy | -0.01089 | 0.46477 | 1 |
| G3bp2 | -0.0109 | 0.570111 | 1 |

| | | | |
|---|---|---|---|
| Ube2n | -0.01091 | 0.828066 | 1 |
| Rps27rt | -0.01092 | 0.956743 | 1 |
| Gm9844 | -0.01097 | 0.666755 | 1 |
| Ide | -0.01099 | 0.927065 | 1 |
| Polr2h | -0.011 | 0.584569 | 1 |
| Gtf2a1 | -0.01103 | 0.686585 | 1 |
| Lrrc41 | -0.01103 | 0.886825 | 1 |
| Naxe | -0.01114 | 0.635614 | 1 |
| Grn | -0.01116 | 0.0583 | 1 |
| Exd2 | -0.0112 | 0.576797 | 1 |
| Rapgef2 | -0.01126 | 0.85461 | 1 |
| Shfl | -0.01126 | 0.779728 | 1 |
| Gm15472 | -0.01129 | 0.531034 | 1 |
| Mob3a | -0.01133 | 0.905397 | 1 |
| Gspt1 | -0.01135 | 0.961163 | 1 |
| Atxn10 | -0.01136 | 0.634485 | 1 |
| Nup214 | -0.01138 | 0.90457 | 1 |
| Bop1 | -0.01143 | 0.654793 | 1 |
| Tnfrsf13b | -0.01145 | 0.058131 | 1 |
| Ankhd1 | -0.01146 | 0.609323 | 1 |
| Fam193a | -0.01146 | 0.812375 | 1 |
| Lcp1 | -0.01155 | 0.828736 | 1 |
| Esco1 | -0.01159 | 0.639314 | 1 |
| Usp50 | -0.01162 | 0.391719 | 1 |
| Xrcc1 | -0.01166 | 0.517528 | 1 |
| Dnajc11 | -0.01171 | 0.553458 | 1 |
| Gm15489 | -0.0118 | 0.455937 | 1 |
| Slc2a1 | -0.01182 | 0.993145 | 1 |
| Rab11fip4 | -0.01187 | 0.654632 | 1 |
| Wdr44 | -0.01194 | 0.908528 | 1 |
| Fem1a | -0.01196 | 0.786129 | 1 |
| Dele1 | -0.01199 | 0.877032 | 1 |
| Timm44 | -0.01199 | 0.625476 | 1 |
| Bak1 | -0.012 | 0.924079 | 1 |
| Prrc1 | -0.01203 | 0.442743 | 1 |
| Rfesd | -0.01207 | 0.974198 | 1 |
| Polr1c | -0.01207 | 0.985142 | 1 |
| Zfp934 | -0.01219 | 0.810535 | 1 |
| Lmf1 | -0.01221 | 0.84299 | 1 |
| Sap30 | -0.01221 | 0.953605 | 1 |
| Slc28a2 | -0.01224 | 0.560933 | 1 |
| Tmem80 | -0.01224 | 0.441419 | 1 |
| Man1a2 | -0.01229 | 0.850173 | 1 |
| Rab1a | -0.01229 | 0.553109 | 1 |
| Supv3l1 | -0.01231 | 0.617912 | 1 |
| Mtrf1l | -0.01238 | 0.565266 | 1 |
| Itgb1bp1 | -0.01242 | 0.876926 | 1 |
| Usp36 | -0.01243 | 0.666559 | 1 |

| | | | |
|---|---|---|---|
| Fyttd1 | -0.01248 | 0.784253 | 1 |
| H2-T3 | -0.0125 | 0.565775 | 1 |
| Clptm1 | -0.01252 | 0.836245 | 1 |
| Xpnpep1 | -0.01256 | 0.62762 | 1 |
| Hk2 | -0.01257 | 0.309242 | 1 |
| Gpat4 | -0.01259 | 0.997181 | 1 |
| Phkb | -0.01268 | 0.975743 | 1 |
| Nomo1 | -0.0127 | 0.593011 | 1 |
| Gatad2a | -0.01271 | 0.340642 | 1 |
| Preb | -0.01276 | 0.957474 | 1 |
| Zfp287 | -0.01279 | 0.991532 | 1 |
| Fdx1 | -0.01293 | 0.649975 | 1 |
| Klc2 | -0.01294 | 0.920185 | 1 |
| Aco2 | -0.01295 | 0.847464 | 1 |
| Osgep | -0.01296 | 0.467946 | 1 |
| Ptp4a3 | -0.01303 | 0.764326 | 1 |
| Zfp326 | -0.01303 | 0.728804 | 1 |
| Anxa7 | -0.01306 | 0.739796 | 1 |
| Sdad1 | -0.01311 | 0.937174 | 1 |
| 6430590A07 | -0.01324 | 0.99506 | 1 |
| Tmem181a | -0.0133 | 0.562233 | 1 |
| Lin37 | -0.01331 | 0.906265 | 1 |
| Txlng | -0.01335 | 0.588784 | 1 |
| Gpn2 | -0.01336 | 0.659263 | 1 |
| Alkbh5 | -0.0134 | 0.569756 | 1 |
| Parp3 | -0.01343 | 0.877861 | 1 |
| Rpl35 | -0.01344 | 0.98423 | 1 |
| Med17 | -0.01345 | 0.464393 | 1 |
| Creb3 | -0.01347 | 0.892577 | 1 |
| Usp47 | -0.01347 | 0.939769 | 1 |
| Psmc1 | -0.01352 | 0.983983 | 1 |
| Cox19 | -0.01352 | 0.675738 | 1 |
| Mydgf | -0.01356 | 0.011413 | 1 |
| Tbc1d8b | -0.01357 | 0.470903 | 1 |
| H2afj | -0.01359 | 0.873576 | 1 |
| Rgl2 | -0.01363 | 0.547156 | 1 |
| Tex261 | -0.01363 | 0.629104 | 1 |
| Acin1 | -0.01373 | 0.877336 | 1 |
| Zkscan6 | -0.01376 | 0.578534 | 1 |
| Anapc10 | -0.01379 | 0.797019 | 1 |
| Nsrp1 | -0.01382 | 0.614223 | 1 |
| Adora2a | -0.01383 | 0.85092 | 1 |
| Ptpn2 | -0.01384 | 0.774388 | 1 |
| Ncl | -0.01384 | 0.516882 | 1 |
| Cep57l1 | -0.01386 | 0.287391 | 1 |
| Dynlrb1 | -0.01401 | 0.733383 | 1 |
| Mettl5 | -0.01402 | 0.633676 | 1 |
| C1galt1 | -0.01403 | 0.588274 | 1 |

| | | | |
|---|---|---|---|
| 2310057M21 | -0.01407 | 0.981608 | 1 |
| Cyren | -0.01408 | 0.432518 | 1 |
| Vrk1 | -0.01413 | 0.609652 | 1 |
| Clcf1 | -0.01421 | 0.551156 | 1 |
| Cd200r1 | -0.01424 | 0.626599 | 1 |
| Zfp511 | -0.01428 | 0.829119 | 1 |
| Rhbdf2 | -0.01432 | 0.858483 | 1 |
| Ddt | -0.01435 | 0.34744 | 1 |
| Eif2d | -0.01436 | 0.297569 | 1 |
| Fars2 | -0.01441 | 0.682594 | 1 |
| Abcc4 | -0.01444 | 0.642487 | 1 |
| Ube2i | -0.01447 | 0.554657 | 1 |
| Nr1h2 | -0.01451 | 0.371348 | 1 |
| Zbtb41 | -0.01458 | 0.325981 | 1 |
| Limk1 | -0.01458 | 0.507032 | 1 |
| Fbxo33 | -0.01463 | 0.991552 | 1 |
| Dcaf13 | -0.01472 | 0.868915 | 1 |
| Vma21 | -0.01481 | 0.475059 | 1 |
| Lamtor5 | -0.01489 | 0.755559 | 1 |
| Gng5 | -0.01495 | 0.83121 | 1 |
| Kat5 | -0.01496 | 0.411949 | 1 |
| H3f3b | -0.01497 | 0.506599 | 1 |
| Sidt2 | -0.015 | 0.53273 | 1 |
| Rpf1 | -0.015 | 0.578951 | 1 |
| Golt1b | -0.015 | 0.12788 | 1 |
| D11Wsu47e | -0.01502 | 0.333137 | 1 |
| Nop16 | -0.01505 | 0.996496 | 1 |
| Alg2 | -0.01505 | 0.767862 | 1 |
| Mocs2 | -0.01506 | 0.878607 | 1 |
| Mlf2 | -0.01506 | 0.761686 | 1 |
| Atp6v0d1 | -0.01507 | 0.974092 | 1 |
| Uxs1 | -0.0151 | 0.761318 | 1 |
| Slc9a1 | -0.01512 | 0.247644 | 1 |
| Aasdhppt | -0.01516 | 0.592455 | 1 |
| Grpel1 | -0.01523 | 0.695701 | 1 |
| Fkbp1a | -0.01523 | 0.716848 | 1 |
| Twsg1 | -0.01532 | 0.895324 | 1 |
| Rars2 | -0.01533 | 0.923757 | 1 |
| Supt7l | -0.01534 | 0.45661 | 1 |
| Naa10 | -0.01537 | 0.695746 | 1 |
| Amfr | -0.01548 | 0.633816 | 1 |
| Bmp2k | -0.01553 | 0.953931 | 1 |
| Nsmce4a | -0.01557 | 0.279154 | 1 |
| Mrpl3 | -0.01558 | 0.402777 | 1 |
| Gm48719 | -0.01558 | 0.719981 | 1 |
| Kat7 | -0.01559 | 0.858174 | 1 |
| Cwf19l1 | -0.01562 | 0.52699 | 1 |
| Prkcz | -0.01568 | 0.121003 | 1 |

| | | | |
|---|---|---|---|
| Fbxl6 | -0.01576 | 0.235964 | 1 |
| Timmdc1 | -0.01581 | 0.685563 | 1 |
| Dcun1d2 | -0.01582 | 0.871408 | 1 |
| mt-Nd6 | -0.01584 | 0.561618 | 1 |
| Emc4 | -0.01588 | 0.683326 | 1 |
| Fam118b | -0.01607 | 0.584253 | 1 |
| D230025D16 | -0.01607 | 0.341649 | 1 |
| Usp24 | -0.0161 | 0.322779 | 1 |
| Phf23 | -0.01611 | 0.258509 | 1 |
| Sec13 | -0.01618 | 0.821359 | 1 |
| R3hdm4 | -0.01619 | 0.816768 | 1 |
| Gm27008 | -0.01625 | 0.711478 | 1 |
| Sirt1 | -0.01627 | 0.356517 | 1 |
| Tspyl1 | -0.0163 | 0.679035 | 1 |
| Washc3 | -0.01639 | 0.308614 | 1 |
| Gtf2f2 | -0.0164 | 0.978773 | 1 |
| Trrap | -0.01642 | 0.952882 | 1 |
| Srcap | -0.0165 | 0.55106 | 1 |
| Rac2 | -0.01651 | 0.402626 | 1 |
| Jazf1 | -0.01653 | 0.528352 | 1 |
| Ccny | -0.01653 | 0.904199 | 1 |
| Cfdp1 | -0.01659 | 0.703181 | 1 |
| Mief1 | -0.01667 | 0.469625 | 1 |
| Nelfcd | -0.01673 | 0.586807 | 1 |
| Ehd3 | -0.01675 | 0.854837 | 1 |
| Fibp | -0.01675 | 0.60697 | 1 |
| Tufm | -0.01684 | 0.828957 | 1 |
| Cdk8 | -0.01689 | 0.003589 | 1 |
| Cfap298 | -0.01692 | 0.889613 | 1 |
| Snx19 | -0.01695 | 0.267304 | 1 |
| Arf6 | -0.01696 | 0.807167 | 1 |
| Acss2 | -0.01703 | 0.170541 | 1 |
| Trpm7 | -0.01705 | 0.846134 | 1 |
| Gtf2h5 | -0.01709 | 0.767258 | 1 |
| Atg2a | -0.01712 | 0.64945 | 1 |
| Atg13 | -0.01713 | 0.972047 | 1 |
| Siah2 | -0.01721 | 0.85195 | 1 |
| Pcgf2 | -0.01722 | 0.170736 | 1 |
| Pdxdc1 | -0.01728 | 0.994816 | 1 |
| Tor1b | -0.01735 | 0.435978 | 1 |
| Pcnt | -0.01736 | 0.770346 | 1 |
| Wac | -0.01742 | 0.447457 | 1 |
| Ss18 | -0.01757 | 0.306068 | 1 |
| Atp5o | -0.01762 | 0.918013 | 1 |
| Irf4 | -0.01765 | 0.362108 | 1 |
| Ppp2r3d | -0.0177 | 0.776662 | 1 |
| Bsdc1 | -0.01772 | 0.286826 | 1 |
| Utp18 | -0.01784 | 0.660923 | 1 |

| | | | |
|---|---|---|---|
| Chn2 | -0.01786 | 0.631874 | 1 |
| Keap1 | -0.01789 | 0.720721 | 1 |
| Zfyve19 | -0.01789 | 0.40544 | 1 |
| Trap1 | -0.01793 | 0.828042 | 1 |
| Stx12 | -0.01796 | 0.500484 | 1 |
| Arid3b | -0.01801 | 0.783601 | 1 |
| Stt3b | -0.01803 | 0.531126 | 1 |
| Prnp | -0.01807 | 0.296977 | 1 |
| Zbtb11os1 | -0.01811 | 0.774141 | 1 |
| Fancl | -0.01815 | 0.571376 | 1 |
| Taf5 | -0.0182 | 0.372439 | 1 |
| Ostm1 | -0.01823 | 0.778744 | 1 |
| Zfp120 | -0.01823 | 0.7515 | 1 |
| Smurf2 | -0.01824 | 0.754508 | 1 |
| Utrn | -0.01824 | 0.926104 | 1 |
| Tamm41 | -0.01826 | 0.848184 | 1 |
| Vps11 | -0.0183 | 0.964029 | 1 |
| Smg6 | -0.0183 | 0.993389 | 1 |
| Abl2 | -0.01831 | 0.496997 | 1 |
| Nupl2 | -0.01836 | 0.68337 | 1 |
| Tia1 | -0.01837 | 0.229979 | 1 |
| Tmem214 | -0.01839 | 0.587582 | 1 |
| Manf | -0.01842 | 0.880355 | 1 |
| Nudc | -0.01865 | 0.641978 | 1 |
| Vps16 | -0.01873 | 0.209187 | 1 |
| Zfp646 | -0.01874 | 0.738569 | 1 |
| Ncbp3 | -0.01877 | 0.395742 | 1 |
| Ndufb8 | -0.01884 | 0.612316 | 1 |
| Dus2 | -0.01887 | 0.742713 | 1 |
| Ddx10 | -0.01887 | 0.497047 | 1 |
| Ankfy1 | -0.01888 | 0.40024 | 1 |
| Clk4 | -0.01895 | 0.465735 | 1 |
| Cisd3 | -0.01903 | 0.957239 | 1 |
| Ate1 | -0.01903 | 0.209465 | 1 |
| Dnajb11 | -0.01905 | 0.665711 | 1 |
| Adck2 | -0.01934 | 0.903301 | 1 |
| Zfp715 | -0.01935 | 0.82245 | 1 |
| Pkn2 | -0.01937 | 0.837551 | 1 |
| Tm9sf1 | -0.01937 | 0.890555 | 1 |
| Rab29 | -0.0194 | 0.383029 | 1 |
| Prpsap1 | -0.0194 | 0.610521 | 1 |
| Fam207a | -0.01942 | 0.50038 | 1 |
| Polr3d | -0.01944 | 0.535026 | 1 |
| Naa20 | -0.01955 | 0.889133 | 1 |
| Man2a1 | -0.01963 | 0.739661 | 1 |
| Mrps2 | -0.01965 | 0.655964 | 1 |
| Xpo6 | -0.01967 | 0.99741 | 1 |
| Dhx30 | -0.01967 | 0.586627 | 1 |

| | | | |
|---|---|---|---|
| Tlnrd1 | -0.01968 | 0.542302 | 1 |
| Faap24 | -0.01972 | 0.665194 | 1 |
| Mroh1 | -0.01973 | 0.706843 | 1 |
| Slc35b4 | -0.01978 | 0.935688 | 1 |
| Abitram | -0.01979 | 0.442506 | 1 |
| Stoml1 | -0.0198 | 0.723781 | 1 |
| Socs2 | -0.0198 | 0.0553 | 1 |
| Slc9a8 | -0.01982 | 0.873544 | 1 |
| Mrps23 | -0.01984 | 0.775849 | 1 |
| AI504432 | -0.0199 | 0.572749 | 1 |
| Ubqln4 | -0.01998 | 0.853914 | 1 |
| Ebna1bp2 | -0.02 | 0.963531 | 1 |
| Rnf111 | -0.02003 | 0.574354 | 1 |
| Cc2d1b | -0.02007 | 0.996852 | 1 |
| 2610001J05I | -0.0201 | 0.816687 | 1 |
| Tmem242 | -0.02014 | 0.640648 | 1 |
| Abraxas1 | -0.02014 | 0.640285 | 1 |
| Zadh2 | -0.02014 | 0.500379 | 1 |
| G6pdx | -0.02019 | 0.61873 | 1 |
| Ube2l3 | -0.02023 | 0.983727 | 1 |
| Frg1 | -0.02024 | 0.951385 | 1 |
| Cep97 | -0.0203 | 0.432584 | 1 |
| Zfp780b | -0.02032 | 0.412794 | 1 |
| Thap11 | -0.02034 | 0.992295 | 1 |
| Prkx | -0.02037 | 0.452977 | 1 |
| Ccdc43 | -0.02038 | 0.469284 | 1 |
| Tmem19 | -0.0204 | 0.989504 | 1 |
| Dennd1a | -0.02042 | 0.608652 | 1 |
| Trim41 | -0.02047 | 0.422072 | 1 |
| Cep76 | -0.02049 | 0.276202 | 1 |
| Amz2 | -0.02054 | 0.789963 | 1 |
| Smim8 | -0.02057 | 0.357622 | 1 |
| Idh2 | -0.02061 | 0.72727 | 1 |
| Itgb1 | -0.02064 | 0.750343 | 1 |
| Gcc2 | -0.02065 | 0.684357 | 1 |
| Hgs | -0.02066 | 0.503131 | 1 |
| Fig4 | -0.02069 | 0.570329 | 1 |
| Zfp160 | -0.02071 | 0.267548 | 1 |
| Sgpp1 | -0.02071 | 0.476846 | 1 |
| Slc12a9 | -0.02074 | 0.865863 | 1 |
| Apaf1 | -0.02075 | 0.491862 | 1 |
| Mettl3 | -0.02078 | 0.580553 | 1 |
| Lsm7 | -0.02079 | 0.850832 | 1 |
| Gabpb1 | -0.02079 | 0.436155 | 1 |
| Kmt2c | -0.0208 | 0.83183 | 1 |
| Nhlrc3 | -0.02082 | 0.590685 | 1 |
| Pigq | -0.02082 | 0.425702 | 1 |
| Fiz1 | -0.02083 | 0.348766 | 1 |

| | | | |
|---|---|---|---|
| Taf1d | -0.02083 | 0.899485 | 1 |
| Slc17a5 | -0.02095 | 0.418083 | 1 |
| Akirin1 | -0.02098 | 0.763101 | 1 |
| Tpk1 | -0.02101 | 0.80134 | 1 |
| Cdk2ap2 | -0.02101 | 0.983497 | 1 |
| Dph5 | -0.02107 | 0.732182 | 1 |
| Bcl3 | -0.02109 | 0.579059 | 1 |
| Ptger4 | -0.02111 | 0.240439 | 1 |
| Zfp869 | -0.02118 | 0.918842 | 1 |
| Ciz1 | -0.02127 | 0.725031 | 1 |
| Brcc3 | -0.02127 | 0.768087 | 1 |
| Trim59 | -0.02127 | 0.688557 | 1 |
| Pelp1 | -0.02128 | 0.369022 | 1 |
| BC023719 | -0.02142 | 0.908196 | 1 |
| Rnf20 | -0.02147 | 0.427036 | 1 |
| Selenoi | -0.0215 | 0.320721 | 1 |
| Ppp6r1 | -0.02156 | 0.481724 | 1 |
| Inpp1 | -0.02165 | 0.857164 | 1 |
| Clasrp | -0.02171 | 0.88998 | 1 |
| Ric8b | -0.02172 | 0.472198 | 1 |
| Spcs1 | -0.02172 | 0.675528 | 1 |
| N6amt1 | -0.02173 | 0.421261 | 1 |
| Ctsb | -0.02173 | 0.56502 | 1 |
| Hspa4l | -0.02178 | 0.981408 | 1 |
| Pitpnb | -0.02178 | 0.837526 | 1 |
| 1110012L19I | -0.0218 | 0.833098 | 1 |
| Wdr33 | -0.02181 | 0.748879 | 1 |
| Axin1 | -0.02197 | 0.21574 | 1 |
| Lbh | -0.02202 | 0.400157 | 1 |
| Trappc5 | -0.02204 | 0.576904 | 1 |
| Acox1 | -0.02209 | 0.598784 | 1 |
| Daglb | -0.02212 | 0.281957 | 1 |
| Eif5a | -0.02215 | 0.989405 | 1 |
| Mis12 | -0.02222 | 0.858905 | 1 |
| Snhg4 | -0.02223 | 0.326849 | 1 |
| Cep152 | -0.02224 | 0.722615 | 1 |
| Vti1b | -0.02225 | 0.553876 | 1 |
| Fbrsl1 | -0.0223 | 0.928768 | 1 |
| Tspan3 | -0.02231 | 0.973316 | 1 |
| Ubr2 | -0.02233 | 0.643645 | 1 |
| Slc6a6 | -0.02234 | 0.305616 | 1 |
| Pola2 | -0.02236 | 0.664919 | 1 |
| Ulk3 | -0.02244 | 0.975709 | 1 |
| Ctr9 | -0.0225 | 0.803436 | 1 |
| Med6 | -0.02254 | 0.504129 | 1 |
| Twistnb | -0.02264 | 0.445052 | 1 |
| Tmem184c | -0.02265 | 0.417938 | 1 |
| Timm22 | -0.02265 | 0.291665 | 1 |

| | | | |
|---|---|---|---|
| Chd8 | -0.02266 | 0.510876 | 1 |
| 2410004B18I | -0.02268 | 0.81804 | 1 |
| Tcea1 | -0.02268 | 0.673855 | 1 |
| Arih1 | -0.0227 | 0.951893 | 1 |
| Usp38 | -0.02273 | 0.729613 | 1 |
| Mecp2 | -0.02282 | 0.833741 | 1 |
| Klf7 | -0.02287 | 0.628243 | 1 |
| Med20 | -0.02291 | 0.247704 | 1 |
| Washc5 | -0.02292 | 0.505247 | 1 |
| Zfand4 | -0.02293 | 0.449027 | 1 |
| Oip5os1 | -0.023 | 0.767965 | 1 |
| Mtrr | -0.023 | 0.892062 | 1 |
| Rdh14 | -0.02301 | 0.810512 | 1 |
| Vsir | -0.02302 | 0.408134 | 1 |
| Hdac4 | -0.02303 | 0.890613 | 1 |
| Ptk2b | -0.02304 | 0.955137 | 1 |
| Matr3 | -0.02308 | 0.913146 | 1 |
| Nemf | -0.02315 | 0.590382 | 1 |
| Gm17018 | -0.02327 | 0.646867 | 1 |
| Vps41 | -0.0233 | 0.707204 | 1 |
| Hdgfl2 | -0.02334 | 0.703253 | 1 |
| Cep83os | -0.02334 | 0.848792 | 1 |
| Tmem127 | -0.02338 | 0.624529 | 1 |
| S1pr2 | -0.02339 | 0.387873 | 1 |
| Pdss1 | -0.02339 | 0.271846 | 1 |
| Phf2 | -0.02345 | 0.769964 | 1 |
| 9430038I01F | -0.02345 | 0.141524 | 1 |
| Fbh1 | -0.02349 | 0.7058 | 1 |
| Nudt19 | -0.02361 | 0.772944 | 1 |
| Zfp729b | -0.02363 | 0.351747 | 1 |
| Mrps30 | -0.02366 | 0.924618 | 1 |
| Cep68 | -0.02368 | 0.529104 | 1 |
| Cad | -0.02371 | 0.640267 | 1 |
| Lamtor1 | -0.02377 | 0.74726 | 1 |
| Arl13b | -0.02379 | 0.517333 | 1 |
| Trpc4ap | -0.02379 | 0.550535 | 1 |
| Mtch2 | -0.02381 | 0.607196 | 1 |
| Rpf2 | -0.02381 | 0.793052 | 1 |
| Nudt16l1 | -0.02382 | 0.901007 | 1 |
| Imp3 | -0.02383 | 0.671954 | 1 |
| Spred1 | -0.02395 | 0.274414 | 1 |
| Spata2 | -0.02398 | 0.581416 | 1 |
| Pigb | -0.024 | 0.765755 | 1 |
| Mrpl20 | -0.024 | 0.985673 | 1 |
| Cndp2 | -0.024 | 0.698157 | 1 |
| 2900076A07I | -0.02401 | 0.522885 | 1 |
| Heca | -0.02403 | 0.804711 | 1 |
| Vps4b | -0.02403 | 0.558744 | 1 |

| | | | |
|---|---|---|---|
| Zfp362 | -0.02407 | 0.804549 | 1 |
| Gabbr1 | -0.02407 | 0.940444 | 1 |
| Nup54 | -0.02412 | 0.727441 | 1 |
| Mpp1 | -0.02421 | 0.435608 | 1 |
| Faap100 | -0.02422 | 0.225801 | 1 |
| Cetn3 | -0.02424 | 0.328602 | 1 |
| Cryl1 | -0.02424 | 0.254956 | 1 |
| Pim2 | -0.02426 | 0.957506 | 1 |
| Tprkb | -0.02429 | 0.318764 | 1 |
| Setdb2 | -0.02433 | 0.147174 | 1 |
| Skp1a | -0.02435 | 0.974964 | 1 |
| Elof1 | -0.0244 | 0.515826 | 1 |
| Specc1 | -0.02445 | 0.212851 | 1 |
| Gars | -0.0245 | 0.537589 | 1 |
| Tbc1d20 | -0.0245 | 0.643501 | 1 |
| Jak2 | -0.02451 | 0.905519 | 1 |
| Rce1 | -0.02456 | 0.740322 | 1 |
| Emc3 | -0.02457 | 0.87747 | 1 |
| Syf2 | -0.02461 | 0.92141 | 1 |
| Kdm6b | -0.02463 | 0.458052 | 1 |
| Mtmr2 | -0.02464 | 0.467839 | 1 |
| Tdrd3 | -0.02468 | 0.264651 | 1 |
| Churc1 | -0.0247 | 0.419617 | 1 |
| Pitpnc1 | -0.02471 | 0.449358 | 1 |
| Prdx1 | -0.02478 | 0.61559 | 1 |
| N4bp3 | -0.02481 | 0.240578 | 1 |
| Vps18 | -0.02483 | 0.656237 | 1 |
| Eif5 | -0.02489 | 0.838923 | 1 |
| Slc25a46 | -0.02491 | 0.574171 | 1 |
| Tspyl4 | -0.02492 | 0.6181 | 1 |
| Zfp52 | -0.02506 | 0.667803 | 1 |
| Cyp20a1 | -0.02511 | 0.389255 | 1 |
| Nom1 | -0.02513 | 0.798208 | 1 |
| Heatr3 | -0.02513 | 0.655524 | 1 |
| Prmt3 | -0.02514 | 0.973053 | 1 |
| Alg14 | -0.02515 | 0.191798 | 1 |
| Psmc2 | -0.02519 | 0.460226 | 1 |
| Ubr7 | -0.0252 | 0.96911 | 1 |
| Cyhr1 | -0.02521 | 0.431542 | 1 |
| Lgals8 | -0.02522 | 0.977996 | 1 |
| Nsf | -0.02523 | 0.296126 | 1 |
| Csde1 | -0.02536 | 0.872698 | 1 |
| Cyp4v3 | -0.02537 | 0.233512 | 1 |
| Nagpa | -0.02538 | 0.750838 | 1 |
| Erap1 | -0.0254 | 0.944283 | 1 |
| Lmbrd1 | -0.02541 | 0.404972 | 1 |
| Synj2bp | -0.02544 | 0.567017 | 1 |
| Stx5a | -0.02546 | 0.787369 | 1 |

| | | | |
|---|---|---|---|
| R3hcc1l | -0.02553 | 0.682027 | 1 |
| Ubxn2b | -0.02555 | 0.339823 | 1 |
| Zfp770 | -0.02558 | 0.217251 | 1 |
| C2cd2l | -0.02561 | 0.665 | 1 |
| Pdcd7 | -0.02569 | 0.34441 | 1 |
| Samd8 | -0.02569 | 0.521513 | 1 |
| Arhgef7 | -0.02575 | 0.675221 | 1 |
| Gpr107 | -0.02578 | 0.962249 | 1 |
| Ccdc6 | -0.02581 | 0.747159 | 1 |
| Cdan1 | -0.02585 | 0.08859 | 1 |
| Fhod1 | -0.02592 | 0.732389 | 1 |
| Tcrg-C2 | -0.02592 | 0.984497 | 1 |
| Cope | -0.02595 | 0.855794 | 1 |
| Ppp4r2 | -0.02597 | 0.559737 | 1 |
| Tbc1d13 | -0.02602 | 0.781933 | 1 |
| Fancf | -0.0261 | 0.452201 | 1 |
| Pdcd10 | -0.02612 | 0.684593 | 1 |
| Rpa3 | -0.02612 | 0.717887 | 1 |
| Rad50 | -0.02614 | 0.931839 | 1 |
| Ncoa5 | -0.02615 | 0.760777 | 1 |
| Acox3 | -0.02616 | 0.470619 | 1 |
| Pi4kb | -0.02617 | 0.829958 | 1 |
| Glmp | -0.02618 | 0.774126 | 1 |
| Dhrs7b | -0.0262 | 0.733811 | 1 |
| Lactb2 | -0.0262 | 0.230471 | 1 |
| Psma1 | -0.02623 | 0.951646 | 1 |
| Limd1 | -0.02627 | 0.374705 | 1 |
| Cisd1 | -0.02629 | 0.702252 | 1 |
| Hivep3 | -0.02632 | 0.404688 | 1 |
| Sars2 | -0.02634 | 0.886557 | 1 |
| Trim28 | -0.0264 | 0.726869 | 1 |
| Cpsf6 | -0.0264 | 0.530368 | 1 |
| Prkag2 | -0.02647 | 0.380264 | 1 |
| Ubqln2 | -0.0265 | 0.887444 | 1 |
| Zfand2b | -0.02654 | 0.333102 | 1 |
| Mtr | -0.02654 | 0.608996 | 1 |
| Prpf8 | -0.02656 | 0.836074 | 1 |
| Dhrs11 | -0.02663 | 0.462686 | 1 |
| Cog5 | -0.02666 | 0.33639 | 1 |
| Wdr43 | -0.02671 | 0.995783 | 1 |
| Ube4b | -0.02681 | 0.615505 | 1 |
| Gm7072 | -0.02683 | 0.9206 | 1 |
| Rpusd4 | -0.0269 | 0.447166 | 1 |
| Ppp1r12b | -0.02691 | 0.928047 | 1 |
| Snapin | -0.02694 | 0.850911 | 1 |
| Triobp | -0.02698 | 0.74332 | 1 |
| Cetn2 | -0.02708 | 0.489306 | 1 |
| D630008O14 | -0.0271 | 0.441082 | 1 |

| | | | |
|---|---|---|---|
| Cipc | -0.02716 | 0.516002 | 1 |
| Abhd10 | -0.02718 | 0.589084 | 1 |
| Tmem11 | -0.0272 | 0.654191 | 1 |
| Zfp865 | -0.02722 | 0.933358 | 1 |
| Asrgl1 | -0.02725 | 0.329592 | 1 |
| Reps1 | -0.02727 | 0.646857 | 1 |
| Ube2b | -0.0273 | 0.762923 | 1 |
| Mre11a | -0.02734 | 0.513617 | 1 |
| Hipk2 | -0.02738 | 0.730734 | 1 |
| AU040320 | -0.02743 | 0.773943 | 1 |
| Ctc1 | -0.02743 | 0.571235 | 1 |
| Ugp2 | -0.02747 | 0.233575 | 1 |
| Mettl7a1 | -0.0275 | 0.623104 | 1 |
| Ccdc66 | -0.02751 | 0.651311 | 1 |
| Abtb1 | -0.0276 | 0.907363 | 1 |
| Ess2 | -0.02762 | 0.908114 | 1 |
| Seh1l | -0.02763 | 0.577646 | 1 |
| Naa25 | -0.02765 | 0.35818 | 1 |
| Inpp5e | -0.02768 | 0.377536 | 1 |
| Rbmx2 | -0.0277 | 0.326479 | 1 |
| Creld2 | -0.02777 | 0.785362 | 1 |
| Ndufb4 | -0.02786 | 0.789733 | 1 |
| Ppp2r5d | -0.02787 | 0.295934 | 1 |
| Atxn2l | -0.02788 | 0.941856 | 1 |
| Cdkl3 | -0.02789 | 0.616174 | 1 |
| Btla | -0.02792 | 0.111336 | 1 |
| Fosl2 | -0.02799 | 0.601904 | 1 |
| Rfxank | -0.02802 | 0.645635 | 1 |
| Traf3ip3 | -0.02806 | 0.635559 | 1 |
| Armh3 | -0.02818 | 0.364475 | 1 |
| Lsm12 | -0.02822 | 0.827712 | 1 |
| Trib1 | -0.02831 | 0.720036 | 1 |
| Dis3l | -0.02833 | 0.560317 | 1 |
| Enpp1 | -0.02835 | 0.412415 | 1 |
| Tmed10 | -0.02838 | 0.0664 | 1 |
| Glt8d1 | -0.02843 | 0.240531 | 1 |
| Actr1b | -0.02844 | 0.814353 | 1 |
| Cebpg | -0.02847 | 0.474711 | 1 |
| Gm14305 | -0.02849 | 0.264341 | 1 |
| Znrd1 | -0.02852 | 0.647831 | 1 |
| Hacd2 | -0.02854 | 0.708652 | 1 |
| Aldh9a1 | -0.02854 | 0.88147 | 1 |
| Sugp2 | -0.02855 | 0.935228 | 1 |
| Sec61b | -0.02856 | 0.449654 | 1 |
| Lrrc4 | -0.02858 | 0.08659 | 1 |
| Thoc5 | -0.02862 | 0.474334 | 1 |
| Epc1 | -0.02865 | 0.996954 | 1 |
| 0610010K14 | -0.0287 | 0.943593 | 1 |

| | | | |
|---|---|---|---|
| Zmym3 | -0.02873 | 0.522565 | 1 |
| Atox1 | -0.02882 | 0.740354 | 1 |
| Klhdc10 | -0.02885 | 0.256322 | 1 |
| Syce2 | -0.02887 | 0.269831 | 1 |
| Chmp7 | -0.02889 | 0.181083 | 1 |
| Tapt1 | -0.02891 | 0.397898 | 1 |
| Myl12b | -0.02894 | 0.776253 | 1 |
| Cryzl1 | -0.02895 | 0.860761 | 1 |
| Ubxn11 | -0.02902 | 0.995796 | 1 |
| Wdr18 | -0.02903 | 0.57739 | 1 |
| Tmem129 | -0.02905 | 0.457455 | 1 |
| Lpp | -0.02914 | 0.709613 | 1 |
| Chfr | -0.02917 | 0.629817 | 1 |
| Atpaf2 | -0.02921 | 0.201771 | 1 |
| Fbxo3 | -0.02924 | 0.283014 | 1 |
| Mapk1ip1l | -0.02925 | 0.911484 | 1 |
| Helb | -0.02927 | 0.39252 | 1 |
| Ppp2r5a | -0.02931 | 0.779101 | 1 |
| Flvcr1 | -0.02936 | 0.211555 | 1 |
| Zfp131 | -0.02936 | 0.756742 | 1 |
| Insig2 | -0.02937 | 0.641117 | 1 |
| Luc7l | -0.02941 | 0.594716 | 1 |
| Scyl2 | -0.02941 | 0.438778 | 1 |
| Ttf1 | -0.02942 | 0.255421 | 1 |
| Crkl | -0.02943 | 0.989274 | 1 |
| Tomm34 | -0.02948 | 0.28754 | 1 |
| Pip5k1a | -0.02953 | 0.732175 | 1 |
| H2-K1 | -0.02969 | 0.788213 | 1 |
| Higd1a | -0.02971 | 0.751666 | 1 |
| Banp | -0.02981 | 0.913404 | 1 |
| Pop5 | -0.0299 | 0.952901 | 1 |
| Mcmbp | -0.02995 | 0.501216 | 1 |
| Map4k2 | -0.03 | 0.954414 | 1 |
| Mapk6 | -0.03002 | 0.183866 | 1 |
| Pitpnm2 | -0.03003 | 0.745967 | 1 |
| Pced1a | -0.03022 | 0.173518 | 1 |
| Tm2d3 | -0.03023 | 0.342157 | 1 |
| Gprasp1 | -0.03024 | 0.464101 | 1 |
| Cmtm3 | -0.03026 | 0.315296 | 1 |
| Trim5 | -0.03027 | 0.094012 | 1 |
| Ints9 | -0.03029 | 0.706268 | 1 |
| Eral1 | -0.0303 | 0.66647 | 1 |
| Chtop | -0.03037 | 0.302912 | 1 |
| Sprtn | -0.03038 | 0.328999 | 1 |
| Ppp6r2 | -0.0304 | 0.180389 | 1 |
| Nprl2 | -0.03041 | 0.282507 | 1 |
| Ampd2 | -0.03043 | 0.596098 | 1 |
| Mrps36 | -0.03052 | 0.417676 | 1 |

| | | | |
|---|---|---|---|
| Dhps | -0.03062 | 0.524146 | 1 |
| Sdhaf2 | -0.03075 | 0.531339 | 1 |
| Ufsp2 | -0.03076 | 0.923206 | 1 |
| Gpr108 | -0.03081 | 0.617668 | 1 |
| Mcm10 | -0.03092 | 0.033073 | 1 |
| Trabd | -0.03093 | 0.596012 | 1 |
| Zfp398 | -0.03104 | 0.326592 | 1 |
| Ccdc117 | -0.03108 | 0.545966 | 1 |
| Hs2st1 | -0.03111 | 0.90392 | 1 |
| Psmd10 | -0.0312 | 0.566511 | 1 |
| Gle1 | -0.03126 | 0.606149 | 1 |
| Ppp1r11 | -0.03126 | 0.821629 | 1 |
| Spryd3 | -0.03127 | 0.916948 | 1 |
| Usp5 | -0.03127 | 0.622566 | 1 |
| Hus1 | -0.03137 | 0.474251 | 1 |
| Alg1 | -0.03141 | 0.968652 | 1 |
| Zbtb25 | -0.03145 | 0.746262 | 1 |
| Rufy1 | -0.03147 | 0.921178 | 1 |
| Entpd6 | -0.0315 | 0.976961 | 1 |
| Slc25a40 | -0.03155 | 0.835487 | 1 |
| Zscan29 | -0.03178 | 0.577127 | 1 |
| Snrpb | -0.03178 | 0.591917 | 1 |
| Derl1 | -0.03179 | 0.842581 | 1 |
| Gm20324 | -0.03182 | 0.400928 | 1 |
| Abhd16a | -0.03185 | 0.93465 | 1 |
| Glb1 | -0.03187 | 0.461384 | 1 |
| Smarcad1 | -0.03198 | 0.362357 | 1 |
| Snrpd2 | -0.03206 | 0.779264 | 1 |
| Coro1b | -0.03213 | 0.692626 | 1 |
| Prr12 | -0.03216 | 0.895092 | 1 |
| Abhd17b | -0.03221 | 0.273353 | 1 |
| Mtdh | -0.03222 | 0.974545 | 1 |
| Tmem156 | -0.03224 | 0.267302 | 1 |
| Tbl1x | -0.03225 | 0.287899 | 1 |
| Taok2 | -0.03231 | 0.312164 | 1 |
| Trappc12 | -0.03231 | 0.575487 | 1 |
| R3hdm2 | -0.03237 | 0.486408 | 1 |
| Usp16 | -0.03239 | 0.523352 | 1 |
| Fut7 | -0.03248 | 0.282541 | 1 |
| Fez2 | -0.03259 | 0.375914 | 1 |
| Impad1 | -0.03268 | 0.278041 | 1 |
| Zfc3h1 | -0.03269 | 0.541977 | 1 |
| Slc35a5 | -0.03271 | 0.228265 | 1 |
| Sass6 | -0.03275 | 0.363031 | 1 |
| Dnmt3b | -0.0328 | 0.243978 | 1 |
| Mark3 | -0.03282 | 0.465172 | 1 |
| Foxk2 | -0.03283 | 0.352473 | 1 |
| D030056L22I | -0.03285 | 0.950232 | 1 |

| | | | |
|---|---|---|---|
| Gatad1 | -0.03287 | 0.303726 | 1 |
| Heatr5b | -0.03292 | 0.292569 | 1 |
| Atp6ap1 | -0.03296 | 0.30862 | 1 |
| Cacul1 | -0.03297 | 0.821646 | 1 |
| Lysmd3 | -0.03298 | 0.144986 | 1 |
| Rsl24d1 | -0.03314 | 0.296143 | 1 |
| Fhl2 | -0.03315 | 0.262037 | 1 |
| Adap1 | -0.03315 | 0.646838 | 1 |
| Lin7c | -0.03319 | 0.753149 | 1 |
| Uck1 | -0.03331 | 0.641127 | 1 |
| Ptgr2 | -0.03332 | 0.568342 | 1 |
| Abca2 | -0.03333 | 0.469508 | 1 |
| Aste1 | -0.03333 | 0.208903 | 1 |
| Ccdc93 | -0.03335 | 0.569321 | 1 |
| 4930522L14I | -0.03336 | 0.293824 | 1 |
| Slc26a2 | -0.03339 | 0.908227 | 1 |
| Smap2 | -0.03341 | 0.554107 | 1 |
| Atp23 | -0.03346 | 0.445618 | 1 |
| Cops4 | -0.03352 | 0.619844 | 1 |
| Crcp | -0.03352 | 0.410655 | 1 |
| Nsmf | -0.03352 | 0.30187 | 1 |
| Ank | -0.03355 | 0.392854 | 1 |
| Wbp1l | -0.03357 | 0.785288 | 1 |
| Shoc2 | -0.03359 | 0.055526 | 1 |
| Abcf1 | -0.03361 | 0.395843 | 1 |
| Lats2 | -0.03361 | 0.041926 | 1 |
| Cop1 | -0.03361 | 0.35083 | 1 |
| St7 | -0.03366 | 0.768287 | 1 |
| Rab8b | -0.03379 | 0.934581 | 1 |
| Fos | -0.03385 | 0.566537 | 1 |
| 5031425E22I | -0.03388 | 0.501914 | 1 |
| Slc4a1ap | -0.0339 | 0.250138 | 1 |
| Rnf170 | -0.03393 | 0.078971 | 1 |
| Fam126b | -0.03398 | 0.53183 | 1 |
| Tra2a | -0.034 | 0.488201 | 1 |
| Mrpl41 | -0.03403 | 0.495057 | 1 |
| Zfp330 | -0.03408 | 0.981109 | 1 |
| Mthfd1 | -0.03409 | 0.352881 | 1 |
| Rabep2 | -0.03416 | 0.894745 | 1 |
| Dffa | -0.03418 | 0.059557 | 1 |
| Mybl2 | -0.03419 | 0.265767 | 1 |
| Pcsk7 | -0.03433 | 0.44268 | 1 |
| B4galt1 | -0.03436 | 0.186167 | 1 |
| Prr3 | -0.03436 | 0.946726 | 1 |
| Cnot6l | -0.03444 | 0.569597 | 1 |
| Dnajc13 | -0.03455 | 0.511424 | 1 |
| Sp3os | -0.03459 | 0.449189 | 1 |
| Zfp930 | -0.03462 | 0.207188 | 1 |

| | | | |
|---|---|---|---|
| Gm10076 | -0.03462 | 0.49284 | 1 |
| Tmsb15b1 | -0.03463 | 0.594875 | 1 |
| Scd2 | -0.03465 | 0.34551 | 1 |
| Ass1 | -0.03466 | 0.772993 | 1 |
| Rprd1a | -0.0347 | 0.867781 | 1 |
| Sirt2 | -0.03477 | 0.999506 | 1 |
| Nipal3 | -0.03484 | 0.425132 | 1 |
| Tkt | -0.0349 | 0.519134 | 1 |
| Paf1 | -0.0349 | 0.309992 | 1 |
| Csnk1e | -0.03496 | 0.325848 | 1 |
| Blzf1 | -0.03509 | 0.882445 | 1 |
| Slc25a1 | -0.03509 | 0.245989 | 1 |
| Sos1 | -0.03512 | 0.527518 | 1 |
| Swi5 | -0.03513 | 0.587575 | 1 |
| Ppip5k2 | -0.03515 | 0.982984 | 1 |
| Wdtc1 | -0.03518 | 0.882973 | 1 |
| Mob4 | -0.03524 | 0.28995 | 1 |
| Gsap | -0.03525 | 0.766508 | 1 |
| Ppp1r16a | -0.0353 | 0.708752 | 1 |
| Zfp148 | -0.03532 | 0.444168 | 1 |
| Ube2d1 | -0.03537 | 0.44769 | 1 |
| Atp6v1e1 | -0.03539 | 0.661501 | 1 |
| 4930581F22I | -0.03539 | 0.450949 | 1 |
| Map1lc3b | -0.03539 | 0.692538 | 1 |
| Zfp26 | -0.03541 | 0.219733 | 1 |
| Rsrp1 | -0.03558 | 0.692616 | 1 |
| Mgat1 | -0.03559 | 0.551636 | 1 |
| Gm10282 | -0.03562 | 0.207275 | 1 |
| Msl1 | -0.03565 | 0.498928 | 1 |
| Rab1b | -0.03567 | 0.426352 | 1 |
| Hsf1 | -0.03571 | 0.880534 | 1 |
| Dnpep | -0.03573 | 0.692437 | 1 |
| Tmem168 | -0.03574 | 0.044979 | 1 |
| Rps6ka3 | -0.03591 | 0.989079 | 1 |
| Nek9 | -0.03597 | 0.162697 | 1 |
| Srebf1 | -0.03598 | 0.993752 | 1 |
| Ppp4r3a | -0.03602 | 0.530088 | 1 |
| Uxt | -0.03603 | 0.332592 | 1 |
| Abl1 | -0.03603 | 0.674502 | 1 |
| Stx4a | -0.03605 | 0.168013 | 1 |
| Sephs1 | -0.03606 | 0.272723 | 1 |
| Desi1 | -0.03607 | 0.959426 | 1 |
| Zfp319 | -0.03608 | 0.616637 | 1 |
| Dnaja3 | -0.03609 | 0.40663 | 1 |
| Tubg1 | -0.03612 | 0.407346 | 1 |
| Nemp2 | -0.03614 | 0.283244 | 1 |
| 5033430I15F | -0.03615 | 0.974256 | 1 |
| Dcaf15 | -0.03617 | 0.166325 | 1 |

| | | | |
|---|---|---|---|
| Acbd5 | -0.03626 | 0.39972 | 1 |
| Rnps1 | -0.03627 | 0.660325 | 1 |
| Tbce | -0.03627 | 0.426463 | 1 |
| Cog3 | -0.0363 | 0.548595 | 1 |
| B3glct | -0.03641 | 0.46142 | 1 |
| Cyfip1 | -0.03642 | 0.887741 | 1 |
| Cdc5l | -0.03652 | 0.332974 | 1 |
| Abt1 | -0.03653 | 0.628842 | 1 |
| Prkra | -0.03655 | 0.472668 | 1 |
| Dtnbp1 | -0.03657 | 0.990098 | 1 |
| 1700097N02 | -0.03665 | 0.17964 | 1 |
| Fbxw5 | -0.0367 | 0.986044 | 1 |
| Pdcd4 | -0.0367 | 0.265597 | 1 |
| Pcmt1 | -0.03673 | 0.328054 | 1 |
| Cacybp | -0.03676 | 0.706207 | 1 |
| Qsox1 | -0.0369 | 0.509863 | 1 |
| Cct4 | -0.03696 | 0.619826 | 1 |
| Ciao2a | -0.03698 | 0.868744 | 1 |
| Zfp280d | -0.03699 | 0.792428 | 1 |
| Ufl1 | -0.037 | 0.746364 | 1 |
| Zfp395 | -0.03701 | 0.150542 | 1 |
| Trp53rka | -0.03701 | 0.182348 | 1 |
| Slmap | -0.03703 | 0.623887 | 1 |
| Rmi1 | -0.03706 | 0.323821 | 1 |
| Pdpr | -0.03708 | 0.917891 | 1 |
| Lage3 | -0.03715 | 0.16644 | 1 |
| Ilf2 | -0.0372 | 0.952485 | 1 |
| Thap12 | -0.03726 | 0.410606 | 1 |
| Cdyl | -0.03727 | 0.375694 | 1 |
| Polk | -0.03732 | 0.495888 | 1 |
| Cwc27 | -0.03739 | 0.46111 | 1 |
| Tsen15 | -0.03747 | 0.335054 | 1 |
| Nup188 | -0.0375 | 0.150908 | 1 |
| Cenps | -0.03751 | 0.20597 | 1 |
| Ppp1r7 | -0.03755 | 0.324667 | 1 |
| Srp19 | -0.03766 | 0.684908 | 1 |
| Acadm | -0.03773 | 0.788012 | 1 |
| Tgs1 | -0.0378 | 0.342338 | 1 |
| Ivns1abp | -0.0378 | 0.836247 | 1 |
| Atpsckmt | -0.03785 | 0.649851 | 1 |
| Peak1 | -0.03798 | 0.445912 | 1 |
| Snx1 | -0.03799 | 0.795752 | 1 |
| Cep19 | -0.03803 | 0.043194 | 1 |
| Sf3a1 | -0.03806 | 0.60352 | 1 |
| Diaph1 | -0.03806 | 0.693409 | 1 |
| Vrk2 | -0.0381 | 0.858055 | 1 |
| Nck2 | -0.03811 | 0.860087 | 1 |
| Slc31a2 | -0.03819 | 0.12408 | 1 |

| | | | |
|---|---|---|---|
| Dpy19l1 | -0.0382 | 0.81801 | 1 |
| Nat10 | -0.03825 | 0.852502 | 1 |
| Tnk2 | -0.03827 | 0.458582 | 1 |
| Trp53bp2 | -0.03834 | 0.387417 | 1 |
| Psmg3 | -0.03836 | 0.86048 | 1 |
| Arid4a | -0.03843 | 0.377747 | 1 |
| Dhx34 | -0.03844 | 0.282449 | 1 |
| Tbc1d10b | -0.03844 | 0.362687 | 1 |
| Tusc3 | -0.03846 | 0.234184 | 1 |
| Htatsf1 | -0.03856 | 0.958557 | 1 |
| Carmil2 | -0.03861 | 0.90213 | 1 |
| Ing4 | -0.03867 | 0.653724 | 1 |
| Dis3 | -0.03872 | 0.509143 | 1 |
| Sirt7 | -0.03888 | 0.474505 | 1 |
| Dcps | -0.03888 | 0.445694 | 1 |
| Cdc34 | -0.039 | 0.641013 | 1 |
| Polr3g | -0.03904 | 0.331526 | 1 |
| Ccdc12 | -0.03904 | 0.449322 | 1 |
| Dync1li2 | -0.03905 | 0.561789 | 1 |
| Kif1b | -0.03905 | 0.143139 | 1 |
| Fundc2 | -0.0391 | 0.721984 | 1 |
| Med8 | -0.03915 | 0.452722 | 1 |
| Gm2000 | -0.03916 | 0.827596 | 1 |
| Cenpj | -0.03917 | 0.130343 | 1 |
| Jagn1 | -0.0392 | 0.265215 | 1 |
| Scp2 | -0.0392 | 0.788905 | 1 |
| Klhl5 | -0.03922 | 0.123111 | 1 |
| Mtch1 | -0.03923 | 0.792176 | 1 |
| Dr1 | -0.03924 | 0.668659 | 1 |
| Lair1 | -0.03925 | 0.123379 | 1 |
| Tbrg4 | -0.03926 | 0.804029 | 1 |
| Iars2 | -0.03927 | 0.466686 | 1 |
| Tmem167 | -0.03942 | 0.724022 | 1 |
| Mllt10 | -0.03947 | 0.943834 | 1 |
| Larp4 | -0.03952 | 0.912782 | 1 |
| Aamdc | -0.03956 | 0.223673 | 1 |
| Cab39 | -0.03956 | 0.549914 | 1 |
| Traf7 | -0.03961 | 0.379554 | 1 |
| Nbeal2 | -0.03965 | 0.215616 | 1 |
| Trim25 | -0.03966 | 0.798911 | 1 |
| Actl6a | -0.0398 | 0.426139 | 1 |
| Pef1 | -0.03981 | 0.117681 | 1 |
| Thyn1 | -0.03984 | 0.370408 | 1 |
| Paics | -0.03992 | 0.868192 | 1 |
| Msrb2 | -0.04001 | 0.335618 | 1 |
| Fam168b | -0.04003 | 0.431937 | 1 |
| Ift140 | -0.04009 | 0.599572 | 1 |
| Nudt4 | -0.04009 | 0.838418 | 1 |

| | | | |
|---|---|---|---|
| Gtf2h3 | -0.04022 | 0.12632 | 1 |
| B9d2 | -0.04026 | 0.702633 | 1 |
| Gimap1os | -0.04033 | 0.596682 | 1 |
| Ccdc166 | -0.04035 | 0.530941 | 1 |
| Elp1 | -0.04036 | 0.915641 | 1 |
| Agtpbp1 | -0.04038 | 0.46971 | 1 |
| Pvrig | -0.04044 | 0.417209 | 1 |
| Prkar2a | -0.04048 | 0.624275 | 1 |
| Mus81 | -0.0405 | 0.33069 | 1 |
| Gmpr2 | -0.04053 | 0.691126 | 1 |
| Glod4 | -0.04054 | 0.762803 | 1 |
| Tmco4 | -0.04057 | 0.264319 | 1 |
| Smn1 | -0.04059 | 0.904783 | 1 |
| Phka2 | -0.04061 | 0.889978 | 1 |
| Zkscan3 | -0.04073 | 0.882197 | 1 |
| Ints8 | -0.04074 | 0.240432 | 1 |
| Haus2 | -0.04081 | 0.209962 | 1 |
| Tmem60 | -0.04083 | 0.435716 | 1 |
| Bcl9l | -0.04088 | 0.374483 | 1 |
| Ptcd1 | -0.04089 | 0.190136 | 1 |
| Slc36a3os | -0.04097 | 0.074682 | 1 |
| Tbc1d5 | -0.04097 | 0.971222 | 1 |
| Snrnp27 | -0.04104 | 0.766506 | 1 |
| Smim20 | -0.04107 | 0.5693 | 1 |
| Fam3c | -0.0411 | 0.560094 | 1 |
| Slc1a5 | -0.04111 | 0.713932 | 1 |
| Fus | -0.0412 | 0.766724 | 1 |
| Cnpy2 | -0.0412 | 0.571418 | 1 |
| Ttll5 | -0.04134 | 0.341969 | 1 |
| Gmppb | -0.04139 | 0.272071 | 1 |
| Zfp422 | -0.04142 | 0.206188 | 1 |
| Slc37a1 | -0.04158 | 0.311404 | 1 |
| Tars2 | -0.04161 | 0.588252 | 1 |
| Agtrap | -0.04165 | 0.869174 | 1 |
| Ercc3 | -0.04169 | 0.160108 | 1 |
| Tom1l2 | -0.0417 | 0.632862 | 1 |
| Scmh1 | -0.04182 | 0.394426 | 1 |
| Dlg2 | -0.04186 | 0.533902 | 1 |
| Abcg1 | -0.04187 | 0.837424 | 1 |
| Pgd | -0.04191 | 0.831695 | 1 |
| Ube2e3 | -0.04194 | 0.151822 | 1 |
| Smpd1 | -0.04197 | 0.977071 | 1 |
| Snrpf | -0.042 | 0.61415 | 1 |
| Ppp4r3b | -0.042 | 0.288645 | 1 |
| Bcl2 | -0.04202 | 0.851312 | 1 |
| Dbt | -0.04207 | 0.915839 | 1 |
| Rfx5 | -0.04207 | 0.30151 | 1 |
| Galnt11 | -0.04208 | 0.882066 | 1 |

| | | | |
|---|---|---|---|
| Ankrd17 | -0.04212 | 0.38037 | 1 |
| Adprh | -0.04225 | 0.743106 | 1 |
| Mrpl50 | -0.04226 | 0.99556 | 1 |
| Qtrt1 | -0.0423 | 0.484948 | 1 |
| Coq10a | -0.04232 | 0.025557 | 1 |
| Ecsit | -0.04234 | 0.611779 | 1 |
| Sec24d | -0.04234 | 0.922438 | 1 |
| Sgta | -0.04248 | 0.52772 | 1 |
| Ndufs8 | -0.04251 | 0.632398 | 1 |
| Primpol | -0.04254 | 0.260393 | 1 |
| Agpat4 | -0.04256 | 0.533177 | 1 |
| Dnm1l | -0.04279 | 0.824077 | 1 |
| Gm48678 | -0.04293 | 0.064518 | 1 |
| Gmcl1 | -0.04294 | 0.432796 | 1 |
| Piezo1 | -0.04298 | 0.638853 | 1 |
| Fuz | -0.043 | 0.10111 | 1 |
| 2610020C07I | -0.04305 | 0.094687 | 1 |
| Ssr3 | -0.04306 | 0.506417 | 1 |
| Ap4b1 | -0.0431 | 0.440651 | 1 |
| Ncoa6 | -0.0431 | 0.321845 | 1 |
| Yipf3 | -0.04312 | 0.934331 | 1 |
| Ciao1 | -0.04314 | 0.44917 | 1 |
| Appl2 | -0.04319 | 0.352127 | 1 |
| Nxf1 | -0.04321 | 0.386736 | 1 |
| 4921524J17I | -0.04321 | 0.9281 | 1 |
| Mvd | -0.04327 | 0.131306 | 1 |
| Snx6 | -0.04333 | 0.914056 | 1 |
| Tsen54 | -0.0434 | 0.049912 | 1 |
| Sf3b5 | -0.04354 | 0.333012 | 1 |
| Ube2v2 | -0.04358 | 0.928153 | 1 |
| Nadk | -0.04363 | 0.802966 | 1 |
| Eif1 | -0.04363 | 0.521339 | 1 |
| Ap5m1 | -0.04364 | 0.666796 | 1 |
| Uqcrb | -0.04365 | 0.702152 | 1 |
| Srsf9 | -0.04366 | 0.032207 | 1 |
| Prps1l3 | -0.04371 | 0.259933 | 1 |
| Lsm8 | -0.04377 | 0.49717 | 1 |
| Wrap53 | -0.04387 | 0.242902 | 1 |
| Ptger2 | -0.04389 | 0.650616 | 1 |
| Pik3cd | -0.04392 | 0.829074 | 1 |
| Slc25a53 | -0.04393 | 0.273649 | 1 |
| Hps6 | -0.04396 | 0.103627 | 1 |
| Zmat3 | -0.04399 | 0.709416 | 1 |
| Szt2 | -0.04413 | 0.075084 | 1 |
| Vcp | -0.04414 | 0.673462 | 1 |
| Sec61a1 | -0.04425 | 0.618767 | 1 |
| Zfp106 | -0.0444 | 0.848619 | 1 |
| Dhx37 | -0.04446 | 0.281342 | 1 |

| | | | |
|---|---|---|---|
| Psmd3 | -0.04447 | 0.280524 | 1 |
| Mdc1 | -0.04449 | 0.660037 | 1 |
| Sfr1 | -0.0445 | 0.478824 | 1 |
| Zmat5 | -0.04451 | 0.559556 | 1 |
| Parp2 | -0.04453 | 0.60291 | 1 |
| Gm11802 | -0.04455 | 0.010613 | 1 |
| Abhd8 | -0.04458 | 0.925544 | 1 |
| Acaa1a | -0.04462 | 0.347198 | 1 |
| Nampt | -0.04471 | 0.634448 | 1 |
| Natd1 | -0.04473 | 0.174771 | 1 |
| Cab39l | -0.04476 | 0.221771 | 1 |
| Nup160 | -0.04481 | 0.607668 | 1 |
| Lmtk2 | -0.04485 | 0.804067 | 1 |
| Ephx1 | -0.04486 | 0.798603 | 1 |
| Faah | -0.04488 | 0.507003 | 1 |
| Poc1b | -0.04501 | 0.210258 | 1 |
| Vps26c | -0.04503 | 0.688534 | 1 |
| Mrpl33 | -0.04509 | 0.482471 | 1 |
| Asb6 | -0.04513 | 0.23637 | 1 |
| Pear1 | -0.04522 | 0.50253 | 1 |
| Zfp386 | -0.04524 | 0.806407 | 1 |
| E330020D12I | -0.04524 | 0.59957 | 1 |
| Fbxl12 | -0.04526 | 0.641068 | 1 |
| Ammecr1l | -0.04527 | 0.140657 | 1 |
| Zbtb7a | -0.04528 | 0.350591 | 1 |
| Chpf2 | -0.04529 | 0.166799 | 1 |
| Cln5 | -0.04531 | 0.933606 | 1 |
| Gtf2h4 | -0.04533 | 0.129058 | 1 |
| Crtap | -0.04541 | 0.339767 | 1 |
| Sac3d1 | -0.04543 | 0.653574 | 1 |
| Gm17173 | -0.04547 | 0.768792 | 1 |
| Sh3glb2 | -0.04549 | 0.301062 | 1 |
| G3bp1 | -0.04549 | 0.654951 | 1 |
| Galnt1 | -0.04551 | 0.383702 | 1 |
| Ppib | -0.04552 | 0.841046 | 1 |
| Exoc8 | -0.04554 | 0.270561 | 1 |
| Tmem18 | -0.04556 | 0.045637 | 1 |
| Ube2f | -0.04562 | 0.143883 | 1 |
| Pcyt1a | -0.04562 | 0.236996 | 1 |
| Dlgap4 | -0.04567 | 0.717303 | 1 |
| Nbeal1 | -0.04574 | 0.684959 | 1 |
| Kdm5c | -0.04576 | 0.930026 | 1 |
| Znrf3 | -0.0458 | 0.778737 | 1 |
| Aida | -0.04581 | 0.355173 | 1 |
| Taf6l | -0.04586 | 0.180019 | 1 |
| Casp1 | -0.04591 | 0.486609 | 1 |
| Fastk | -0.04592 | 0.683852 | 1 |
| Smad5 | -0.04594 | 0.07567 | 1 |

| | | | |
|---|---|---|---|
| Pum2 | -0.04599 | 0.149337 | 1 |
| Dcaf1 | -0.04604 | 0.314 | 1 |
| Tpcn2 | -0.04605 | 0.144159 | 1 |
| Arl16 | -0.0462 | 0.072275 | 1 |
| Glcci1 | -0.04621 | 0.388203 | 1 |
| Hscb | -0.04628 | 0.408479 | 1 |
| Slc9a6 | -0.04631 | 0.589773 | 1 |
| Nmd3 | -0.04643 | 0.681366 | 1 |
| Med23 | -0.04645 | 0.318438 | 1 |
| Senp1 | -0.04646 | 0.356767 | 1 |
| Ttc4 | -0.04648 | 0.402168 | 1 |
| Maea | -0.04652 | 0.753976 | 1 |
| Wdr81 | -0.04655 | 0.554812 | 1 |
| Coro1c | -0.04668 | 0.793966 | 1 |
| Paip2b | -0.04669 | 0.867878 | 1 |
| Cutc | -0.0467 | 0.746341 | 1 |
| Galns | -0.04672 | 0.698475 | 1 |
| Taf4 | -0.04674 | 0.314577 | 1 |
| Tmem243 | -0.04676 | 0.732509 | 1 |
| Fbxl5 | -0.04677 | 0.220808 | 1 |
| Aftph | -0.04677 | 0.256476 | 1 |
| Snx9 | -0.04682 | 0.65542 | 1 |
| Nup98 | -0.04683 | 0.917371 | 1 |
| Rab5b | -0.04688 | 0.445513 | 1 |
| Comtd1 | -0.04692 | 0.527626 | 1 |
| Kat6a | -0.04698 | 0.526825 | 1 |
| 4833420G17 | -0.047 | 0.921535 | 1 |
| Eif2s3y | -0.04705 | 0.098685 | 1 |
| Cog2 | -0.04706 | 0.525958 | 1 |
| Fastkd1 | -0.04706 | 0.156852 | 1 |
| Itpa | -0.04708 | 0.080512 | 1 |
| Rsl1d1 | -0.04714 | 0.737623 | 1 |
| Ankrd12 | -0.04715 | 0.721564 | 1 |
| Aldh16a1 | -0.04718 | 0.590379 | 1 |
| Tmem62 | -0.04731 | 0.800926 | 1 |
| Nelfa | -0.04741 | 0.641949 | 1 |
| Nipsnap3b | -0.04742 | 0.569248 | 1 |
| Snrpa | -0.04744 | 0.996655 | 1 |
| Ranbp3 | -0.04745 | 0.18339 | 1 |
| Pole3 | -0.04745 | 0.63543 | 1 |
| Extl3 | -0.04749 | 0.56222 | 1 |
| Eif4a1 | -0.04755 | 0.588165 | 1 |
| Anapc16 | -0.04756 | 0.302142 | 1 |
| Ptrh2 | -0.04762 | 0.439273 | 1 |
| Utp20 | -0.04764 | 0.473141 | 1 |
| Rell1 | -0.04773 | 0.176844 | 1 |
| Tsr3 | -0.0478 | 0.404197 | 1 |
| Zgpat | -0.04801 | 0.58988 | 1 |

| | | | |
|---|---|---|---|
| Scaf8 | -0.04804 | 0.571043 | 1 |
| Kcnq1ot1 | -0.04807 | 0.610196 | 1 |
| Atp5e | -0.04811 | 0.445242 | 1 |
| Zfp688 | -0.04831 | 0.396369 | 1 |
| Zdhhc2 | -0.04835 | 0.15483 | 1 |
| Ppp2r5e | -0.04837 | 0.503994 | 1 |
| Tex30 | -0.04839 | 0.32921 | 1 |
| 2210016F16I | -0.04844 | 0.182796 | 1 |
| Dbr1 | -0.04853 | 0.427705 | 1 |
| Pxylp1 | -0.04866 | 0.964882 | 1 |
| Rad52 | -0.04873 | 0.059907 | 1 |
| Akap9 | -0.04878 | 0.524924 | 1 |
| Zfp91 | -0.04881 | 0.436331 | 1 |
| Wdr24 | -0.04895 | 0.156431 | 1 |
| Itpripl1 | -0.04901 | 0.40386 | 1 |
| Tmsb10 | -0.04903 | 0.656591 | 1 |
| Pex7 | -0.04903 | 0.506541 | 1 |
| Vhl | -0.04904 | 0.234696 | 1 |
| Gpatch8 | -0.04908 | 0.634557 | 1 |
| Gcnt1 | -0.04915 | 0.749963 | 1 |
| Gnptab | -0.04916 | 0.409589 | 1 |
| Ankrd16 | -0.04918 | 0.250247 | 1 |
| Dtx2 | -0.0492 | 0.352533 | 1 |
| Atrx | -0.04923 | 0.467655 | 1 |
| Cdk5 | -0.04928 | 0.34176 | 1 |
| Maml1 | -0.04933 | 0.146042 | 1 |
| Cep85l | -0.04933 | 0.004104 | 1 |
| Mettl9 | -0.04937 | 0.924823 | 1 |
| Trpv2 | -0.04938 | 0.277029 | 1 |
| Copz1 | -0.04939 | 0.338497 | 1 |
| Tars | -0.04942 | 0.481079 | 1 |
| Ankrd39 | -0.04945 | 0.192817 | 1 |
| Lypla1 | -0.04949 | 0.339061 | 1 |
| Asah1 | -0.04949 | 0.960697 | 1 |
| Sfmbt1 | -0.04951 | 0.284765 | 1 |
| Champ1 | -0.04953 | 0.566122 | 1 |
| Gipc1 | -0.04955 | 0.92304 | 1 |
| Plekhf2 | -0.04967 | 0.841311 | 1 |
| Phykpl | -0.0497 | 0.240916 | 1 |
| Zhx1 | -0.04972 | 0.266791 | 1 |
| Fam20b | -0.04975 | 0.124726 | 1 |
| Pepd | -0.04975 | 0.290474 | 1 |
| Usp33 | -0.04977 | 0.113131 | 1 |
| Faim | -0.04979 | 0.591176 | 1 |
| Rab40c | -0.04979 | 0.254961 | 1 |
| Zfp771 | -0.0498 | 0.790673 | 1 |
| Tmco6 | -0.04981 | 0.453603 | 1 |
| Thoc7 | -0.04983 | 0.362409 | 1 |

| | | | |
|---|---|---|---|
| Sphk2 | -0.04985 | 0.680524 | 1 |
| Nsmce2 | -0.04989 | 0.151585 | 1 |
| Wdr26 | -0.04991 | 0.587922 | 1 |
| Erf | -0.04992 | 0.145974 | 1 |
| Ppp2ca | -0.05002 | 0.231988 | 1 |
| Rsbn1l | -0.05003 | 0.376843 | 1 |
| Spin1 | -0.05007 | 0.397036 | 1 |
| Elk4 | -0.05007 | 0.942534 | 1 |
| Csrnp1 | -0.05019 | 0.79689 | 1 |
| Fam185a | -0.05027 | 0.926528 | 1 |
| Ndufa2 | -0.05031 | 0.560672 | 1 |
| March5 | -0.05032 | 0.997311 | 1 |
| Ehmt1 | -0.05033 | 0.906738 | 1 |
| Tnpo2 | -0.05039 | 0.114891 | 1 |
| Ipo13 | -0.05046 | 0.237971 | 1 |
| 1110008P14 | -0.05056 | 0.626259 | 1 |
| Camkmt | -0.05057 | 0.663273 | 1 |
| Bloc1s3 | -0.05068 | 0.783228 | 1 |
| Ahctf1 | -0.05084 | 0.74207 | 1 |
| Stxbp3 | -0.05091 | 0.210405 | 1 |
| Pin1 | -0.05096 | 0.228675 | 1 |
| Immp2l | -0.05097 | 0.541317 | 1 |
| Fam8a1 | -0.05112 | 0.099267 | 1 |
| Gtdc1 | -0.05115 | 0.862378 | 1 |
| Bid | -0.0512 | 0.258052 | 1 |
| Dohh | -0.05121 | 0.378558 | 1 |
| Lnpep | -0.05121 | 0.529329 | 1 |
| Arpc1b | -0.05121 | 0.293548 | 1 |
| Trim36 | -0.05123 | 0.253937 | 1 |
| Zbtb38 | -0.05127 | 0.553369 | 1 |
| Btrc | -0.05141 | 0.533408 | 1 |
| Pet100 | -0.05152 | 0.357129 | 1 |
| Bcl2l12 | -0.05154 | 0.740977 | 1 |
| Agbl5 | -0.05163 | 0.006095 | 1 |
| Emc9 | -0.05166 | 0.401435 | 1 |
| Vps33a | -0.05168 | 0.22406 | 1 |
| Tnfaip8 | -0.05174 | 0.686904 | 1 |
| Arfgap2 | -0.05184 | 0.504457 | 1 |
| H2-T23 | -0.05186 | 0.44538 | 1 |
| Mnt | -0.05187 | 0.438708 | 1 |
| Ddx1 | -0.0519 | 0.623672 | 1 |
| Scamp2 | -0.05194 | 0.516821 | 1 |
| Cept1 | -0.05198 | 0.19046 | 1 |
| Traf2 | -0.05198 | 0.839179 | 1 |
| Depdc5 | -0.052 | 0.334432 | 1 |
| Mrpl54 | -0.05207 | 0.511235 | 1 |
| Mocs1 | -0.05215 | 0.849658 | 1 |
| Zfp574 | -0.05221 | 0.070799 | 1 |

| | | | |
|---|---|---|---|
| Clpx | -0.05221 | 0.411711 | 1 |
| Cnot6 | -0.05225 | 0.728748 | 1 |
| Rxrb | -0.05226 | 0.557157 | 1 |
| Cul1 | -0.0523 | 0.943277 | 1 |
| Wscd2 | -0.0523 | 0.279115 | 1 |
| Isca1 | -0.05236 | 0.408674 | 1 |
| Elovl1 | -0.05241 | 0.37777 | 1 |
| Pign | -0.05245 | 0.14353 | 1 |
| Mepce | -0.05246 | 0.286335 | 1 |
| Slc20a1 | -0.0525 | 0.459334 | 1 |
| Traf5 | -0.05254 | 0.832822 | 1 |
| Tmco3 | -0.05272 | 0.028959 | 1 |
| Timm8b | -0.05283 | 0.404682 | 1 |
| Zfp800 | -0.05284 | 0.632188 | 1 |
| Myef2 | -0.05293 | 0.392834 | 1 |
| Tiam1 | -0.05296 | 0.415685 | 1 |
| Mrpl35 | -0.05301 | 0.981811 | 1 |
| Mmut | -0.05305 | 0.178393 | 1 |
| Gm45716 | -0.0531 | 0.522553 | 1 |
| Sgcb | -0.05312 | 0.303016 | 1 |
| 8030462N17 | -0.05314 | 0.298433 | 1 |
| Acot2 | -0.05316 | 0.34128 | 1 |
| Nudt14 | -0.05325 | 0.248088 | 1 |
| P4hb | -0.05327 | 0.855988 | 1 |
| Abhd6 | -0.05328 | 0.738803 | 1 |
| Tbcc | -0.05331 | 0.079489 | 1 |
| Slc38a10 | -0.05335 | 0.44021 | 1 |
| Lpar6 | -0.05343 | 0.879503 | 1 |
| Uba3 | -0.05351 | 0.287776 | 1 |
| Nipsnap2 | -0.05352 | 0.362769 | 1 |
| Fbxo6 | -0.05352 | 0.042413 | 1 |
| Sppl3 | -0.05357 | 0.78227 | 1 |
| Paip1 | -0.05358 | 0.487371 | 1 |
| Actr2 | -0.05363 | 0.118853 | 1 |
| Kdm3b | -0.05372 | 0.741728 | 1 |
| Dym | -0.05374 | 0.332126 | 1 |
| Vamp5 | -0.05375 | 0.763056 | 1 |
| Baz2b | -0.05384 | 0.159312 | 1 |
| Lsm4 | -0.05388 | 0.608157 | 1 |
| Gtpbp6 | -0.05389 | 0.675795 | 1 |
| Ddb2 | -0.05389 | 0.389651 | 1 |
| Tspan2 | -0.05391 | 0.054453 | 1 |
| Gm33104 | -0.05393 | 0.88467 | 1 |
| Magoh | -0.05397 | 0.506746 | 1 |
| BC002059 | -0.05397 | 0.805078 | 1 |
| Gigyf2 | -0.05399 | 0.452638 | 1 |
| Mier2 | -0.05402 | 0.052188 | 1 |
| Rcc2 | -0.05403 | 0.801815 | 1 |

| | | | |
|---|---|---|---|
| Prdm10 | -0.05407 | 0.065212 | 1 |
| Chst11 | -0.0541 | 0.928527 | 1 |
| Rint1 | -0.05411 | 0.138178 | 1 |
| Parn | -0.05412 | 0.267407 | 1 |
| Upf1 | -0.05415 | 0.196693 | 1 |
| Ash1l | -0.05427 | 0.427258 | 1 |
| Mrps35 | -0.05428 | 0.212281 | 1 |
| Fh1 | -0.05438 | 0.463746 | 1 |
| Tcf25 | -0.05442 | 0.830056 | 1 |
| Aspscr1 | -0.05443 | 0.728324 | 1 |
| Enpp4 | -0.05443 | 0.347422 | 1 |
| Pcnp | -0.05448 | 0.614878 | 1 |
| Scml4 | -0.05449 | 0.267316 | 1 |
| Ncdn | -0.05455 | 0.039068 | 1 |
| Snrpd3 | -0.05455 | 0.817944 | 1 |
| Ube2q1 | -0.05457 | 0.360583 | 1 |
| Rad54b | -0.05461 | 0.008882 | 1 |
| Set | -0.05465 | 0.534674 | 1 |
| Trnt1 | -0.05467 | 0.692329 | 1 |
| Ube3c | -0.05472 | 0.426916 | 1 |
| Efcab14 | -0.05479 | 0.110795 | 1 |
| Prdm2 | -0.05479 | 0.815801 | 1 |
| Trip11 | -0.05485 | 0.465089 | 1 |
| Saal1 | -0.05497 | 0.566524 | 1 |
| Tmem263 | -0.05499 | 0.097911 | 1 |
| Med31 | -0.05502 | 0.344284 | 1 |
| Cog1 | -0.05507 | 0.862359 | 1 |
| Oaz1 | -0.0551 | 0.27836 | 1 |
| Aamp | -0.05512 | 0.384645 | 1 |
| Stard3 | -0.05513 | 0.656082 | 1 |
| Kdm7a | -0.05515 | 0.509076 | 1 |
| Srfbp1 | -0.05516 | 0.373 | 1 |
| Dcaf10 | -0.0553 | 0.547408 | 1 |
| Exosc8 | -0.05532 | 0.51569 | 1 |
| Ppm1m | -0.05536 | 0.660971 | 1 |
| Zbtb22 | -0.05538 | 0.007804 | 1 |
| Ssr1 | -0.05539 | 0.372057 | 1 |
| Btbd9 | -0.05541 | 0.116411 | 1 |
| Sh2d3c | -0.05543 | 0.129084 | 1 |
| Pex11b | -0.05544 | 0.807006 | 1 |
| Ccdc71l | -0.05549 | 0.061387 | 1 |
| Tubgcp3 | -0.05551 | 0.37384 | 1 |
| Casp8ap2 | -0.05562 | 0.405005 | 1 |
| Ggta1 | -0.05562 | 0.069119 | 1 |
| Ftsj1 | -0.05567 | 0.335117 | 1 |
| Ptcd2 | -0.05572 | 0.897243 | 1 |
| Rbm4 | -0.05576 | 0.182726 | 1 |
| Polg | -0.05582 | 0.039668 | 1 |

| | | | |
|---|---|---|---|
| Rbm28 | -0.05589 | 0.793352 | 1 |
| Dstn | -0.05592 | 0.06947 | 1 |
| Cklf | -0.05592 | 0.435184 | 1 |
| Pggt1b | -0.05593 | 0.344076 | 1 |
| Brox | -0.05593 | 0.483557 | 1 |
| Gm16845 | -0.05594 | 0.092174 | 1 |
| Hist3h2ba | -0.05599 | 0.244403 | 1 |
| Slc36a1 | -0.05601 | 0.053712 | 1 |
| Xrcc6 | -0.05603 | 0.340771 | 1 |
| Cdpf1 | -0.05606 | 0.843787 | 1 |
| Spata13 | -0.05606 | 0.140499 | 1 |
| Hbs1l | -0.05609 | 0.448163 | 1 |
| Naa35 | -0.05611 | 0.321799 | 1 |
| Topbp1 | -0.05627 | 0.068534 | 1 |
| Azi2 | -0.05628 | 0.794462 | 1 |
| Tmem234 | -0.05629 | 0.923241 | 1 |
| Abhd13 | -0.05633 | 0.055901 | 1 |
| Zfp719 | -0.05635 | 0.048088 | 1 |
| Vkorc1l1 | -0.05641 | 0.066115 | 1 |
| Limk2 | -0.05656 | 0.361563 | 1 |
| Nrarp | -0.05657 | 0.010041 | 1 |
| 1700021F05I | -0.0566 | 0.253071 | 1 |
| Rragc | -0.05664 | 0.28501 | 1 |
| Mtx1 | -0.0567 | 0.216942 | 1 |
| Plscr3 | -0.05672 | 0.224647 | 1 |
| Bcl2l11 | -0.05673 | 0.678413 | 1 |
| Brf1 | -0.05673 | 0.229676 | 1 |
| Ctnnbl1 | -0.05679 | 0.351843 | 1 |
| Haus5 | -0.05679 | 0.644535 | 1 |
| Fbxo7 | -0.05684 | 0.277931 | 1 |
| Rrp15 | -0.05684 | 0.412431 | 1 |
| Rbm15b | -0.05687 | 0.131024 | 1 |
| Ticam1 | -0.05689 | 0.646882 | 1 |
| Nmb | -0.0569 | 0.389915 | 1 |
| Ripor1 | -0.05692 | 0.271207 | 1 |
| Myo9a | -0.05692 | 0.871276 | 1 |
| Sdccag8 | -0.05697 | 0.683949 | 1 |
| Snf8 | -0.057 | 0.321388 | 1 |
| Mad2l1bp | -0.05702 | 0.266843 | 1 |
| Zscan25 | -0.05703 | 0.2922 | 1 |
| Rassf7 | -0.05704 | 0.118612 | 1 |
| Atp5g3 | -0.05706 | 0.707911 | 1 |
| Katna1 | -0.05706 | 0.462616 | 1 |
| Parvg | -0.05706 | 0.774223 | 1 |
| Tfe3 | -0.05706 | 0.640223 | 1 |
| Abraxas2 | -0.05709 | 0.545861 | 1 |
| Ccs | -0.05714 | 0.840356 | 1 |
| Efl1 | -0.05716 | 0.455764 | 1 |

| | | | |
|---|---|---|---|
| Dyrk1a | -0.05717 | 0.59251 | 1 |
| Dot1l | -0.05717 | 0.991665 | 1 |
| Wwp2 | -0.05718 | 0.851577 | 1 |
| P2ry14 | -0.0572 | 0.690362 | 1 |
| Polr1a | -0.05723 | 0.766871 | 1 |
| Golga3 | -0.05724 | 0.164122 | 1 |
| Mrpl47 | -0.05731 | 0.23027 | 1 |
| Trim23 | -0.05743 | 0.036374 | 1 |
| Tsga10 | -0.05745 | 0.018119 | 1 |
| Mycbp | -0.05746 | 0.014968 | 1 |
| Ube2g1 | -0.0575 | 0.738271 | 1 |
| Ttf2 | -0.05751 | 0.688163 | 1 |
| Cenpo | -0.05755 | 0.310416 | 1 |
| Scpep1 | -0.05755 | 0.956995 | 1 |
| Sytl3 | -0.0576 | 0.498102 | 1 |
| Ciapin1 | -0.05761 | 0.397867 | 1 |
| Togaram1 | -0.05762 | 0.404557 | 1 |
| Plin2 | -0.05762 | 0.472845 | 1 |
| Cmpk1 | -0.05764 | 0.576122 | 1 |
| Nup155 | -0.05766 | 0.528328 | 1 |
| Fancm | -0.05771 | 0.058054 | 1 |
| Rhot2 | -0.05775 | 0.346654 | 1 |
| Mtbp | -0.05776 | 0.369988 | 1 |
| Rrp8 | -0.05778 | 0.522514 | 1 |
| Acap1 | -0.05787 | 0.852424 | 1 |
| Rmdn3 | -0.05791 | 0.08914 | 1 |
| Syngr2 | -0.05793 | 0.716974 | 1 |
| Ddx50 | -0.05793 | 0.175905 | 1 |
| Tubgcp4 | -0.05797 | 0.612462 | 1 |
| 9930111J21F | -0.05797 | 0.652679 | 1 |
| Xpa | -0.05805 | 0.816987 | 1 |
| Mrpl49 | -0.0581 | 0.254123 | 1 |
| Cdipt | -0.0581 | 0.418767 | 1 |
| E130307A14I | -0.05812 | 0.181888 | 1 |
| Galnt3 | -0.05813 | 0.350636 | 1 |
| Ddx54 | -0.05816 | 0.729836 | 1 |
| Rab14 | -0.05817 | 0.21373 | 1 |
| Vgll4 | -0.05817 | 0.427954 | 1 |
| Stk11ip | -0.05818 | 0.620981 | 1 |
| Zfp827 | -0.05819 | 0.213694 | 1 |
| Cemip2 | -0.05819 | 0.555796 | 1 |
| Rttn | -0.05823 | 0.046839 | 1 |
| Gale | -0.05825 | 0.007938 | 1 |
| Elk1 | -0.05825 | 0.150529 | 1 |
| Cbll1 | -0.05826 | 0.318965 | 1 |
| Nop56 | -0.0583 | 0.299661 | 1 |
| Prdx3 | -0.05832 | 0.691017 | 1 |
| BC052040 | -0.05834 | 0.283614 | 1 |

| | | | |
|---|---|---|---|
| Endov | -0.05836 | 0.336915 | 1 |
| Copb1 | -0.05839 | 0.264364 | 1 |
| Washc2 | -0.05848 | 0.315535 | 1 |
| Ldb1 | -0.05849 | 0.268176 | 1 |
| Pot1b | -0.0586 | 0.175161 | 1 |
| Med1 | -0.05861 | 0.46018 | 1 |
| 6720427I07F | -0.05862 | 0.254546 | 1 |
| Pygb | -0.05863 | 0.446402 | 1 |
| Ino80d | -0.05868 | 0.977356 | 1 |
| Skp2 | -0.05874 | 0.009485 | 1 |
| Zscan21 | -0.05878 | 0.581761 | 1 |
| Urgcp | -0.0588 | 0.198098 | 1 |
| Hyou1 | -0.05887 | 0.237846 | 1 |
| Arnt | -0.05887 | 0.121613 | 1 |
| Nsfl1c | -0.05888 | 0.715912 | 1 |
| Clec2i | -0.05903 | 0.502855 | 1 |
| Gins1 | -0.05916 | 0.016008 | 1 |
| Ppp2r3c | -0.0593 | 0.402369 | 1 |
| Gng10 | -0.05934 | 0.906144 | 1 |
| Vps4a | -0.05935 | 0.34539 | 1 |
| Etaa1 | -0.05938 | 0.221268 | 1 |
| Slc4a8 | -0.05941 | 0.020541 | 1 |
| Hmg20a | -0.05947 | 0.583933 | 1 |
| Anp32b | -0.05949 | 0.908889 | 1 |
| Cep78 | -0.05955 | 0.063081 | 1 |
| Rragb | -0.05956 | 0.748396 | 1 |
| Sharpin | -0.05967 | 0.838155 | 1 |
| Pglyrp2 | -0.05973 | 0.388346 | 1 |
| Setd5 | -0.05975 | 0.654143 | 1 |
| G2e3 | -0.05977 | 0.012985 | 1 |
| Ints5 | -0.05978 | 0.359223 | 1 |
| Cybc1 | -0.05982 | 0.827095 | 1 |
| Rbm26 | -0.05983 | 0.361212 | 1 |
| Ormdl1 | -0.05987 | 0.19872 | 1 |
| Rab3d | -0.05992 | 0.815152 | 1 |
| Pibf1 | -0.05994 | 0.502109 | 1 |
| D130040H23 | -0.05994 | 0.183709 | 1 |
| Med7 | -0.05997 | 0.107163 | 1 |
| Hnrnph1 | -0.06001 | 0.851328 | 1 |
| Arhgap15 | -0.06002 | 0.418948 | 1 |
| Hmbox1 | -0.06005 | 0.121172 | 1 |
| 1700037H04 | -0.0601 | 0.122788 | 1 |
| Rfc3 | -0.06013 | 0.238181 | 1 |
| Rbm48 | -0.06016 | 0.225125 | 1 |
| Cd80 | -0.0602 | 0.69733 | 1 |
| Ruvbl1 | -0.06025 | 0.224989 | 1 |
| Pnpla2 | -0.06031 | 0.194033 | 1 |
| Rap1gds1 | -0.06037 | 0.73759 | 1 |

| | | | |
|---|---|---|---|
| Ube2k | -0.06041 | 0.49541 | 1 |
| Sh3bgrl | -0.06043 | 0.46472 | 1 |
| Zfp24 | -0.06046 | 0.146756 | 1 |
| Mknk1 | -0.06047 | 0.049006 | 1 |
| Wdr48 | -0.06051 | 0.286203 | 1 |
| Mfap3 | -0.06052 | 0.090178 | 1 |
| Oga | -0.06057 | 0.302968 | 1 |
| Pbx4 | -0.0606 | 0.105226 | 1 |
| Dnaja2 | -0.06064 | 0.477198 | 1 |
| Hist1h4c | -0.0607 | 0.133458 | 1 |
| B4galt3 | -0.06071 | 0.066061 | 1 |
| Sde2 | -0.06078 | 0.224857 | 1 |
| Psmb2 | -0.06085 | 0.810032 | 1 |
| Tmem199 | -0.0609 | 0.114816 | 1 |
| Arl3 | -0.06096 | 0.266192 | 1 |
| Uqcrc1 | -0.06097 | 0.41945 | 1 |
| Kin | -0.06102 | 0.31225 | 1 |
| Pgm2l1 | -0.06103 | 0.040434 | 1 |
| Pold2 | -0.06108 | 0.305226 | 1 |
| Orc2 | -0.06108 | 0.563156 | 1 |
| Gm37494 | -0.06111 | 0.457324 | 1 |
| Mlx | -0.06113 | 0.130324 | 1 |
| Ints13 | -0.06116 | 0.386234 | 1 |
| Ranbp2 | -0.06119 | 0.764179 | 1 |
| Hps4 | -0.06122 | 0.344977 | 1 |
| Ccnd3 | -0.06124 | 0.508566 | 1 |
| Cdkn1a | -0.06125 | 0.44279 | 1 |
| Uty | -0.0614 | 0.542664 | 1 |
| Cherp | -0.06143 | 0.279841 | 1 |
| Gm44175 | -0.06153 | 0.370408 | 1 |
| Ipo5 | -0.06153 | 0.663291 | 1 |
| Wtap | -0.06154 | 0.759356 | 1 |
| Wee1 | -0.06157 | 0.334451 | 1 |
| Cd2bp2 | -0.0616 | 0.367871 | 1 |
| Disc1 | -0.06161 | 0.00338 | 1 |
| Nelfb | -0.06172 | 0.571656 | 1 |
| Cct5 | -0.06181 | 0.344353 | 1 |
| Cdc26 | -0.06183 | 0.345366 | 1 |
| Eml4 | -0.06185 | 0.917886 | 1 |
| Kdm4a | -0.06188 | 0.437277 | 1 |
| Bap1 | -0.06191 | 0.293803 | 1 |
| Ccdc50 | -0.06191 | 0.187082 | 1 |
| Adsl | -0.06196 | 0.926624 | 1 |
| Psmb1 | -0.06201 | 0.479493 | 1 |
| Tango2 | -0.06223 | 0.098172 | 1 |
| Stard3nl | -0.06224 | 0.527834 | 1 |
| Cd2ap | -0.0623 | 0.591799 | 1 |
| Wls | -0.06242 | 0.228901 | 1 |

| | | | |
|---|---|---|---|
| Ptms | -0.06244 | 0.583834 | 1 |
| Brat1 | -0.06253 | 0.985805 | 1 |
| Gemin5 | -0.06257 | 0.47801 | 1 |
| Gtf3c4 | -0.06259 | 0.003768 | 1 |
| Ubxn6 | -0.06273 | 0.904321 | 1 |
| Cd151 | -0.06273 | 0.308775 | 1 |
| Rock1 | -0.06276 | 0.268424 | 1 |
| Ppp3r1 | -0.0628 | 0.726197 | 1 |
| Zfp597 | -0.06284 | 0.414963 | 1 |
| Unc45a | -0.06309 | 0.129653 | 1 |
| Als2cl | -0.06311 | 0.010776 | 1 |
| Rad9a | -0.06319 | 0.210636 | 1 |
| Glipr1 | -0.06321 | 0.309474 | 1 |
| Fam133b | -0.06321 | 0.422733 | 1 |
| Tial1 | -0.06325 | 0.985775 | 1 |
| Nsun2 | -0.06326 | 0.549876 | 1 |
| Leo1 | -0.06327 | 0.243651 | 1 |
| Trappc2 | -0.06343 | 0.439747 | 1 |
| Zfp62 | -0.06345 | 0.202189 | 1 |
| Naip2 | -0.06357 | 0.098284 | 1 |
| Orai3 | -0.06358 | 0.723121 | 1 |
| Zwint | -0.06359 | 0.864044 | 1 |
| Ctps | -0.06361 | 0.723337 | 1 |
| Fancg | -0.06369 | 0.029056 | 1 |
| Hars | -0.06374 | 0.271768 | 1 |
| Btbd6 | -0.0638 | 0.161348 | 1 |
| Ccdc32 | -0.06398 | 0.250445 | 1 |
| Zbtb11 | -0.06399 | 0.774107 | 1 |
| Slc38a9 | -0.06401 | 0.914898 | 1 |
| Fanca | -0.06404 | 0.014435 | 1 |
| Tbk1 | -0.06405 | 0.445582 | 1 |
| Pwwp3a | -0.06411 | 0.08293 | 1 |
| Fam120a | -0.06413 | 0.421947 | 1 |
| Enkd1 | -0.0643 | 0.003494 | 1 |
| Tdg | -0.06432 | 0.229477 | 1 |
| Mrpl53 | -0.06434 | 0.087037 | 1 |
| Numb | -0.06438 | 0.92073 | 1 |
| Zfp868 | -0.06444 | 0.587437 | 1 |
| Txnl4a | -0.06458 | 0.190765 | 1 |
| Zfyve27 | -0.06459 | 0.491074 | 1 |
| Plekha5 | -0.06467 | 0.143713 | 1 |
| Cspp1 | -0.06472 | 0.466219 | 1 |
| Plaa | -0.06476 | 0.323166 | 1 |
| Tmem29 | -0.06481 | 0.680712 | 1 |
| Supt6 | -0.06481 | 0.845731 | 1 |
| Asxl2 | -0.06485 | 0.352181 | 1 |
| Slc12a7 | -0.06493 | 0.109682 | 1 |
| Ago3 | -0.06506 | 0.723069 | 1 |

| | | | |
|---|---|---|---|
| Cd68 | -0.0651 | 0.112234 | 1 |
| A930005H10 | -0.06528 | 0.082428 | 1 |
| Gnpat | -0.06529 | 0.486903 | 1 |
| Sec11c | -0.06534 | 0.233744 | 1 |
| Tut7 | -0.06535 | 0.559918 | 1 |
| Gm15232 | -0.06537 | 0.13082 | 1 |
| Rassf2 | -0.06539 | 0.384892 | 1 |
| Papola | -0.06539 | 0.547313 | 1 |
| Slc30a7 | -0.0654 | 0.559266 | 1 |
| Atg101 | -0.06549 | 0.229064 | 1 |
| Arid2 | -0.06551 | 0.094335 | 1 |
| Myo9b | -0.06558 | 0.270168 | 1 |
| Lin54 | -0.06559 | 0.466588 | 1 |
| Bloc1s1 | -0.0656 | 0.737823 | 1 |
| Psen1 | -0.06563 | 0.21521 | 1 |
| Zfp207 | -0.06563 | 0.341425 | 1 |
| Bloc1s4 | -0.06564 | 0.43109 | 1 |
| Eif4h | -0.06567 | 0.208192 | 1 |
| Zdhhc4 | -0.06576 | 0.886266 | 1 |
| Klhl2 | -0.06581 | 0.023553 | 1 |
| Supt5 | -0.06592 | 0.176237 | 1 |
| Rfc5 | -0.06595 | 0.077615 | 1 |
| Iffo1 | -0.066 | 0.661572 | 1 |
| Cep170 | -0.06601 | 0.179034 | 1 |
| Cct6a | -0.06607 | 0.410021 | 1 |
| Zcchc10 | -0.06607 | 0.440622 | 1 |
| BE692007 | -0.06617 | 0.945811 | 1 |
| Pgm3 | -0.06624 | 0.421448 | 1 |
| Rabep1 | -0.06634 | 0.352333 | 1 |
| Gtpbp10 | -0.0664 | 0.074735 | 1 |
| Cox6a1 | -0.06644 | 0.44225 | 1 |
| Atp5a1 | -0.06647 | 0.186717 | 1 |
| Mcoln2 | -0.06654 | 0.322021 | 1 |
| Ccnd2 | -0.06659 | 0.489944 | 1 |
| Chd9 | -0.06659 | 0.296198 | 1 |
| Ptp4a2 | -0.06665 | 0.441482 | 1 |
| Ormdl3 | -0.06666 | 0.430391 | 1 |
| Polr2i | -0.06666 | 0.553213 | 1 |
| Atp8b2 | -0.06672 | 0.066855 | 1 |
| Cenpq | -0.06676 | 0.167895 | 1 |
| Apeh | -0.06677 | 0.24969 | 1 |
| Mat2a | -0.06683 | 0.317692 | 1 |
| Stau1 | -0.06697 | 0.58928 | 1 |
| Smim24 | -0.06701 | 0.307523 | 1 |
| Sbno1 | -0.06701 | 0.478837 | 1 |
| A330023F24I | -0.06707 | 0.049692 | 1 |
| Trappc10 | -0.06707 | 0.123447 | 1 |
| Ppp6c | -0.06718 | 0.254245 | 1 |

| | | | |
|---|---|---|---|
| Med26 | -0.06726 | 0.079673 | 1 |
| Ulk2 | -0.06728 | 0.189645 | 1 |
| Zfp592 | -0.0673 | 0.808042 | 1 |
| Ric8a | -0.0673 | 0.411266 | 1 |
| U2af2 | -0.06735 | 0.405523 | 1 |
| Alkbh8 | -0.06736 | 0.333225 | 1 |
| Nsun6 | -0.06739 | 0.922663 | 1 |
| Agk | -0.06739 | 0.110377 | 1 |
| Smyd4 | -0.06745 | 0.168384 | 1 |
| Pitrm1 | -0.06748 | 0.695814 | 1 |
| B3galnt2 | -0.0675 | 0.414563 | 1 |
| Smc5 | -0.06752 | 0.210828 | 1 |
| Kansl2 | -0.06757 | 0.919382 | 1 |
| Nudt22 | -0.06761 | 0.738764 | 1 |
| Mical1 | -0.06764 | 0.23439 | 1 |
| Bcl10 | -0.06767 | 0.349512 | 1 |
| Gm11110 | -0.0677 | 0.470627 | 1 |
| Slc41a1 | -0.06771 | 0.129378 | 1 |
| Asxl1 | -0.06776 | 0.128199 | 1 |
| C1d | -0.06785 | 0.223546 | 1 |
| Ctbp1 | -0.06789 | 0.319439 | 1 |
| Nubp1 | -0.0679 | 0.267042 | 1 |
| Zfp51 | -0.06794 | 0.659507 | 1 |
| Tepsin | -0.06797 | 0.079478 | 1 |
| Calcoco1 | -0.06798 | 0.444552 | 1 |
| Pwwp2a | -0.06801 | 0.159895 | 1 |
| Mad1l1 | -0.06808 | 0.608865 | 1 |
| Rnf214 | -0.0681 | 0.118981 | 1 |
| Setd1b | -0.06816 | 0.090724 | 1 |
| Tm2d2 | -0.06823 | 0.49859 | 1 |
| Zc3h14 | -0.06824 | 0.02854 | 1 |
| Fbxw4 | -0.06828 | 0.864132 | 1 |
| Tsg101 | -0.06832 | 0.163138 | 1 |
| Tcf3 | -0.06841 | 0.573601 | 1 |
| Rmc1 | -0.06848 | 0.821585 | 1 |
| Rbm5 | -0.06853 | 0.299191 | 1 |
| Tmem126a | -0.06854 | 0.389825 | 1 |
| Uqcr11 | -0.06858 | 0.730949 | 1 |
| Rnf169 | -0.06861 | 0.332307 | 1 |
| Rps17 | -0.06864 | 0.647181 | 1 |
| Naaa | -0.06874 | 0.021425 | 1 |
| Gaa | -0.06875 | 0.645608 | 1 |
| Ccdc186 | -0.06887 | 0.955934 | 1 |
| Snx25 | -0.06891 | 0.328014 | 1 |
| Cbwd1 | -0.06899 | 0.466266 | 1 |
| Ccdc138 | -0.06901 | 0.041631 | 1 |
| Rcn1 | -0.06902 | 0.691895 | 1 |
| Xpc | -0.06902 | 0.163025 | 1 |

| | | | |
|---|---|---|---|
| Sec61a2 | -0.06905 | 0.074284 | 1 |
| Mxd4 | -0.06907 | 0.324263 | 1 |
| Popdc2 | -0.06909 | 0.055075 | 1 |
| Kdm8 | -0.06915 | 0.113622 | 1 |
| Elmo1 | -0.06917 | 0.382114 | 1 |
| Dazap2 | -0.0692 | 0.12758 | 1 |
| Nxn | -0.06922 | 0.288648 | 1 |
| Anapc1 | -0.06922 | 0.761335 | 1 |
| Bloc1s2 | -0.06924 | 0.43404 | 1 |
| Mef2d | -0.06928 | 0.17331 | 1 |
| Naa50 | -0.06938 | 0.620759 | 1 |
| Rabif | -0.06944 | 0.124885 | 1 |
| Dxo | -0.06946 | 0.197424 | 1 |
| Dctn1 | -0.06947 | 0.340373 | 1 |
| Arpp19 | -0.06947 | 0.233276 | 1 |
| Gm5141 | -0.06953 | 0.221491 | 1 |
| Rsph3a | -0.06961 | 0.529564 | 1 |
| Smpd2 | -0.06967 | 0.040585 | 1 |
| Abcb7 | -0.0697 | 0.623576 | 1 |
| Oxr1 | -0.06982 | 0.768426 | 1 |
| Bcas2 | -0.06984 | 0.132675 | 1 |
| Zfp7 | -0.06986 | 0.009554 | 1 |
| Wdr4 | -0.06992 | 0.40998 | 1 |
| C330007P06I | -0.07003 | 0.470948 | 1 |
| Usp10 | -0.07004 | 0.381135 | 1 |
| Timm17a | -0.07004 | 0.527772 | 1 |
| Tmem9b | -0.07015 | 0.727571 | 1 |
| Ddx3y | -0.07016 | 0.155856 | 1 |
| Ddx59 | -0.07018 | 0.185243 | 1 |
| Tnfrsf1a | -0.07023 | 0.142977 | 1 |
| Ext2 | -0.07028 | 0.580158 | 1 |
| Gigyf1 | -0.07033 | 0.091433 | 1 |
| Tm9sf2 | -0.07033 | 0.377857 | 1 |
| Usp4 | -0.07041 | 0.293249 | 1 |
| Hadh | -0.07042 | 0.825003 | 1 |
| Ranbp1 | -0.07044 | 0.62146 | 1 |
| Hdac3 | -0.07058 | 0.707758 | 1 |
| Scoc | -0.07062 | 0.736816 | 1 |
| Cers6 | -0.07062 | 0.371289 | 1 |
| Borcs5 | -0.07062 | 0.337541 | 1 |
| Zbtb5 | -0.07065 | 0.038764 | 1 |
| Pxmp2 | -0.07075 | 0.03598 | 1 |
| Vdac3 | -0.07082 | 0.3376 | 1 |
| Eef1akmt2 | -0.07086 | 0.252567 | 1 |
| Taok1 | -0.0709 | 0.162654 | 1 |
| Lime1 | -0.07092 | 0.134415 | 1 |
| Mbnl3 | -0.07102 | 0.368702 | 1 |
| Sart3 | -0.07105 | 0.898598 | 1 |

| | | | |
|---|---|---|---|
| Cwf19l2 | -0.07106 | 0.365058 | 1 |
| Tmem115 | -0.0711 | 0.723839 | 1 |
| Akt3 | -0.07113 | 0.349821 | 1 |
| Fam57a | -0.07114 | 0.778256 | 1 |
| Gadd45g | -0.07129 | 0.88119 | 1 |
| Emc8 | -0.07139 | 0.477892 | 1 |
| Dusp5 | -0.07141 | 0.23435 | 1 |
| Auh | -0.07142 | 0.180506 | 1 |
| Sec24a | -0.07146 | 0.677098 | 1 |
| Pigv | -0.07148 | 0.12675 | 1 |
| Fbxw7 | -0.07155 | 0.297417 | 1 |
| Msl2 | -0.07157 | 0.743535 | 1 |
| St13 | -0.0717 | 0.788205 | 1 |
| Phc2 | -0.0718 | 0.98293 | 1 |
| Ccdc71 | -0.07181 | 0.175883 | 1 |
| Uspl1 | -0.07184 | 0.399218 | 1 |
| Nudt5 | -0.07194 | 0.423939 | 1 |
| Bcr | -0.07195 | 0.225801 | 1 |
| Mpc1 | -0.07195 | 0.779156 | 1 |
| Mis18a | -0.07199 | 0.093664 | 1 |
| Rap1b | -0.07206 | 0.42861 | 1 |
| Adk | -0.07216 | 0.651484 | 1 |
| Gfm2 | -0.07218 | 0.785512 | 1 |
| Pelo | -0.07223 | 0.86411 | 1 |
| Plpbp | -0.0723 | 0.108391 | 1 |
| Pgrmc1 | -0.0724 | 0.521685 | 1 |
| Zfand3 | -0.07243 | 0.912109 | 1 |
| Morf4l1 | -0.07249 | 0.719283 | 1 |
| Washc1 | -0.0726 | 0.601493 | 1 |
| Polr2l | -0.07274 | 0.240156 | 1 |
| Sec23ip | -0.0729 | 0.36488 | 1 |
| Tspyl3 | -0.0729 | 0.064359 | 1 |
| Pigyl | -0.07304 | 0.289239 | 1 |
| Scai | -0.07307 | 0.215964 | 1 |
| Ing1 | -0.07307 | 0.063197 | 1 |
| Rbm42 | -0.0731 | 0.462071 | 1 |
| Pnrc2 | -0.07312 | 0.103238 | 1 |
| 2810403D21 | -0.07316 | 0.574136 | 1 |
| Vta1 | -0.07317 | 0.522726 | 1 |
| Celf1 | -0.07322 | 0.164251 | 1 |
| Thrap3 | -0.07327 | 0.740266 | 1 |
| 4833439L19I | -0.07329 | 0.279411 | 1 |
| Cdk2ap1 | -0.07339 | 0.045157 | 1 |
| Mllt3 | -0.07342 | 0.123616 | 1 |
| Zbtb45 | -0.07343 | 0.127807 | 1 |
| Bicral | -0.07357 | 0.154977 | 1 |
| Alyref2 | -0.07361 | 0.72355 | 1 |
| Fopnl | -0.07376 | 0.184867 | 1 |

| | | | |
|---|---|---|---|
| Trip12 | -0.07384 | 0.096214 | 1 |
| Ldlrad4 | -0.07386 | 0.97771 | 1 |
| Zbed4 | -0.07387 | 0.068728 | 1 |
| Lrch4 | -0.07388 | 0.191686 | 1 |
| Zzef1 | -0.07389 | 0.615716 | 1 |
| Gns | -0.07395 | 0.068577 | 1 |
| Gins3 | -0.074 | 0.138492 | 1 |
| Mrgbp | -0.07404 | 0.506865 | 1 |
| Apobec1 | -0.07406 | 0.527285 | 1 |
| Rab7 | -0.07408 | 0.235626 | 1 |
| Tm2d1 | -0.07409 | 0.322688 | 1 |
| Chmp5 | -0.0741 | 0.646393 | 1 |
| Ltv1 | -0.0741 | 0.743594 | 1 |
| A930006K02I | -0.07412 | 0.180905 | 1 |
| Coasy | -0.07419 | 0.395648 | 1 |
| Ppm1d | -0.0742 | 0.358992 | 1 |
| 4930453N24 | -0.07429 | 0.12699 | 1 |
| Zfp958 | -0.07432 | 0.551681 | 1 |
| Cox7c | -0.07432 | 0.444662 | 1 |
| Dvl3 | -0.07437 | 0.027209 | 1 |
| E130308A19I | -0.07441 | 0.122379 | 1 |
| Fkbp8 | -0.07449 | 0.272755 | 1 |
| Cyp51 | -0.07452 | 0.177441 | 1 |
| Dclk2 | -0.07469 | 0.054211 | 1 |
| Bckdha | -0.07469 | 0.378688 | 1 |
| Itprip | -0.07472 | 0.170188 | 1 |
| Pms2 | -0.07479 | 0.365901 | 1 |
| Exo1 | -0.07485 | 0.01563 | 1 |
| Erp29 | -0.07485 | 0.972332 | 1 |
| Slc10a3 | -0.07487 | 0.228434 | 1 |
| Mbnl2 | -0.0749 | 0.627411 | 1 |
| Zc3h7b | -0.07499 | 0.187612 | 1 |
| Btd | -0.07506 | 0.039925 | 1 |
| Ost4 | -0.07512 | 0.142721 | 1 |
| Pten | -0.07518 | 0.261687 | 1 |
| Lsm5 | -0.07522 | 0.799226 | 1 |
| Npat | -0.07525 | 0.099792 | 1 |
| Rnf4 | -0.07528 | 0.5488 | 1 |
| Ntmt1 | -0.07534 | 0.792225 | 1 |
| mt-Nd3 | -0.07538 | 0.307144 | 1 |
| Ergic3 | -0.0754 | 0.601344 | 1 |
| Mpdu1 | -0.07543 | 0.467822 | 1 |
| Tasor | -0.07547 | 0.364612 | 1 |
| Antxr2 | -0.07556 | 0.039676 | 1 |
| Dhx16 | -0.07558 | 0.139028 | 1 |
| Bicra | -0.07562 | 0.373601 | 1 |
| Cep295 | -0.07563 | 0.137294 | 1 |
| Akr1b3 | -0.07566 | 0.613621 | 1 |

| | | | |
|---|---|---|---|
| Riok1 | -0.0757 | 0.118458 | 1 |
| Fut11 | -0.0757 | 0.080161 | 1 |
| Btaf1 | -0.07572 | 0.596661 | 1 |
| Zfp553 | -0.07576 | 0.723718 | 1 |
| Vav1 | -0.07585 | 0.224828 | 1 |
| Brd1 | -0.07586 | 0.245346 | 1 |
| Ap3m1 | -0.0759 | 0.125618 | 1 |
| Lemd3 | -0.07595 | 0.375334 | 1 |
| Btbd7 | -0.07602 | 0.284656 | 1 |
| Gtpbp3 | -0.07611 | 0.1706 | 1 |
| Nbr1 | -0.07619 | 0.755862 | 1 |
| Casp9 | -0.07623 | 0.432944 | 1 |
| Gm13212 | -0.07623 | 0.17354 | 1 |
| Atp6v1f | -0.07624 | 0.107793 | 1 |
| Eif6 | -0.07627 | 0.230065 | 1 |
| Ctsz | -0.07632 | 0.249593 | 1 |
| Akap10 | -0.07637 | 0.901369 | 1 |
| Mrps17 | -0.07643 | 0.939436 | 1 |
| Naxd | -0.0765 | 0.186195 | 1 |
| Arhgap27 | -0.07653 | 0.454457 | 1 |
| Wbp2 | -0.07656 | 0.240369 | 1 |
| Lrrfip2 | -0.0766 | 0.133692 | 1 |
| Arl5b | -0.07662 | 0.265781 | 1 |
| Snhg3 | -0.07664 | 0.437198 | 1 |
| Wrnip1 | -0.07664 | 0.11902 | 1 |
| Bri3bp | -0.07681 | 0.8682 | 1 |
| Ubc | -0.07683 | 0.549824 | 1 |
| Gen1 | -0.07685 | 0.002304 | 1 |
| Fance | -0.07686 | 0.051289 | 1 |
| Svbp | -0.07689 | 0.929879 | 1 |
| Senp3 | -0.0769 | 0.076992 | 1 |
| Ykt6 | -0.0769 | 0.189916 | 1 |
| Frrs1 | -0.07697 | 0.086053 | 1 |
| Med25 | -0.07703 | 0.177754 | 1 |
| Coil | -0.07708 | 0.270605 | 1 |
| Sdhaf4 | -0.07711 | 0.606762 | 1 |
| 5830408C22I | -0.07713 | 0.211837 | 1 |
| Vps53 | -0.07715 | 0.096172 | 1 |
| Kif19a | -0.07721 | 0.002345 | 1 |
| Ppox | -0.0773 | 0.036301 | 1 |
| Spdl1 | -0.07742 | 0.053532 | 1 |
| Rab12 | -0.07745 | 0.101547 | 1 |
| Tpm1 | -0.07746 | 0.272564 | 1 |
| Iws1 | -0.07746 | 0.256199 | 1 |
| Uchl3 | -0.07748 | 0.492412 | 1 |
| Micu1 | -0.07752 | 0.513584 | 1 |
| Mxi1 | -0.07752 | 0.103194 | 1 |
| Wdr82 | -0.07754 | 0.831335 | 1 |

| | | | |
|---|---|---|---|
| Jdp2 | -0.07754 | 0.308721 | 1 |
| Gata3 | -0.07755 | 0.789661 | 1 |
| Rab6a | -0.07757 | 0.679592 | 1 |
| Tdp1 | -0.07765 | 0.010661 | 1 |
| Ash2l | -0.07767 | 0.216809 | 1 |
| Htatip2 | -0.07774 | 0.319255 | 1 |
| BC005624 | -0.07779 | 0.978574 | 1 |
| 1700096K18I | -0.07783 | 0.45996 | 1 |
| Adh5 | -0.07784 | 0.331049 | 1 |
| Mbd1 | -0.07785 | 0.879724 | 1 |
| Frmd4a | -0.07791 | 0.812305 | 1 |
| Nudcd3 | -0.07791 | 0.688147 | 1 |
| Pcm1 | -0.07796 | 0.209359 | 1 |
| B4galt7 | -0.07799 | 0.881513 | 1 |
| Ptpn13 | -0.07799 | 0.432738 | 1 |
| Sms | -0.07804 | 0.384034 | 1 |
| Donson | -0.07804 | 0.095585 | 1 |
| Ipp | -0.07805 | 0.072847 | 1 |
| Srsf10 | -0.07807 | 0.158219 | 1 |
| Epg5 | -0.07807 | 0.816773 | 1 |
| Rcl1 | -0.07809 | 0.604132 | 1 |
| Rnf219 | -0.07817 | 0.685165 | 1 |
| Sec61g | -0.07828 | 0.11745 | 1 |
| Cuedc1 | -0.07829 | 0.071516 | 1 |
| Lrch3 | -0.0783 | 0.591058 | 1 |
| Hsf4 | -0.07835 | 0.07651 | 1 |
| Gnptg | -0.07836 | 0.128916 | 1 |
| Col4a3bp | -0.07844 | 0.241647 | 1 |
| March7 | -0.07847 | 0.976312 | 1 |
| Ptrhd1 | -0.07849 | 0.522665 | 1 |
| Lasp1 | -0.07873 | 0.493724 | 1 |
| Rpain | -0.0788 | 0.584585 | 1 |
| Nsd1 | -0.07888 | 0.313928 | 1 |
| Klf16 | -0.0789 | 0.269438 | 1 |
| Arhgap35 | -0.07891 | 0.124893 | 1 |
| Ndufaf7 | -0.07892 | 0.580933 | 1 |
| Frmd4b | -0.07901 | 0.949798 | 1 |
| Fam49b | -0.07904 | 0.512258 | 1 |
| Ttc37 | -0.07911 | 0.011902 | 1 |
| Psmd5 | -0.07912 | 0.934317 | 1 |
| Them6 | -0.0792 | 0.257999 | 1 |
| Sertad2 | -0.07929 | 0.252676 | 1 |
| Nfatc2ip | -0.07933 | 0.432301 | 1 |
| Smarcd1 | -0.07942 | 0.143823 | 1 |
| Zfp329 | -0.07945 | 0.173258 | 1 |
| Ilvbl | -0.07948 | 0.105222 | 1 |
| Slc12a4 | -0.0795 | 0.040932 | 1 |
| Bptf | -0.07951 | 0.404984 | 1 |

| | | | |
|---|---|---|---|
| Ier2 | -0.07967 | 0.659886 | 1 |
| Rab35 | -0.0797 | 0.025312 | 1 |
| Rad17 | -0.07971 | 0.328092 | 1 |
| Sumf2 | -0.07977 | 0.073313 | 1 |
| Arpc3 | -0.07978 | 0.099318 | 1 |
| Ccdc47 | -0.07985 | 0.433399 | 1 |
| Clcn3 | -0.07988 | 0.2622 | 1 |
| Tbpl1 | -0.07988 | 0.041718 | 1 |
| Mbtps2 | -0.07997 | 0.06517 | 1 |
| Ankle1 | -0.08002 | 0.000348 | 1 |
| Tradd | -0.08002 | 0.205788 | 1 |
| Zhx3 | -0.08023 | 0.014563 | 1 |
| Bin3 | -0.08024 | 0.765573 | 1 |
| Cobll1 | -0.08035 | 0.24824 | 1 |
| Stk4 | -0.08039 | 0.267552 | 1 |
| Capn7 | -0.0804 | 0.294311 | 1 |
| Morc3 | -0.08044 | 0.882286 | 1 |
| Sema4b | -0.08054 | 0.264687 | 1 |
| Gm5165 | -0.08056 | 0.535836 | 1 |
| Iqsec1 | -0.08057 | 0.278535 | 1 |
| Gemin6 | -0.08059 | 0.403052 | 1 |
| Prcp | -0.08064 | 0.763805 | 1 |
| Dars2 | -0.08071 | 0.016358 | 1 |
| Psmb4 | -0.08071 | 0.824727 | 1 |
| Cyb561d2 | -0.08085 | 0.060995 | 1 |
| Cfl2 | -0.0809 | 0.794815 | 1 |
| Ahcyl2 | -0.08099 | 0.524015 | 1 |
| Rtca | -0.081 | 0.423654 | 1 |
| Zfp809 | -0.08101 | 0.088148 | 1 |
| Rbm15 | -0.08111 | 0.172361 | 1 |
| Tmem63a | -0.08113 | 0.845268 | 1 |
| Nhp2 | -0.08122 | 0.404173 | 1 |
| Akip1 | -0.08124 | 0.129559 | 1 |
| Gss | -0.0813 | 0.160334 | 1 |
| Ypel2 | -0.08146 | 0.076059 | 1 |
| Srgap3 | -0.08146 | 0.444231 | 1 |
| Drg1 | -0.08151 | 0.470877 | 1 |
| Ppp4r1 | -0.08158 | 0.024219 | 1 |
| Rbm14 | -0.08175 | 0.080425 | 1 |
| Lnx2 | -0.08176 | 0.346012 | 1 |
| Rab6b | -0.08176 | 0.005207 | 1 |
| Pde12 | -0.08188 | 0.426774 | 1 |
| Tram1 | -0.08188 | 0.528095 | 1 |
| Rpgrip1 | -0.08193 | 0.047334 | 1 |
| Sf3a2 | -0.08193 | 0.072732 | 1 |
| Eef1akmt1 | -0.08197 | 0.420921 | 1 |
| Gdi2 | -0.082 | 0.430805 | 1 |
| Ssbp1 | -0.08211 | 0.92667 | 1 |

| | | | |
|---|---|---|---|
| Chst2 | -0.08219 | 0.00654 | 1 |
| Cul4b | -0.08235 | 0.08457 | 1 |
| Bcdin3d | -0.08253 | 0.242923 | 1 |
| Mtif3 | -0.08258 | 0.594043 | 1 |
| Ccar2 | -0.08259 | 0.071385 | 1 |
| Telo2 | -0.08263 | 0.40887 | 1 |
| Surf4 | -0.08266 | 0.440329 | 1 |
| Mppe1 | -0.08271 | 0.187422 | 1 |
| Zfp69 | -0.08273 | 0.012818 | 1 |
| Mbnl1 | -0.0828 | 0.205762 | 1 |
| Rcn2 | -0.08291 | 0.832181 | 1 |
| Tmc8 | -0.08294 | 0.601611 | 1 |
| Cdkn2aip | -0.08294 | 0.223778 | 1 |
| Agpat1 | -0.08296 | 0.159303 | 1 |
| Spns1 | -0.08296 | 0.07229 | 1 |
| Gga1 | -0.08299 | 0.106991 | 1 |
| Men1 | -0.08306 | 0.123137 | 1 |
| Hdac5 | -0.08321 | 0.328179 | 1 |
| Phlpp2 | -0.08322 | 0.038595 | 1 |
| Tfdp1 | -0.08329 | 0.161284 | 1 |
| Msh5 | -0.08341 | 6.3E-05 | 1 |
| Blmh | -0.08351 | 0.873324 | 1 |
| Csnk1g2 | -0.08354 | 0.528709 | 1 |
| 4930503L19I | -0.08362 | 0.095986 | 1 |
| Hmces | -0.08364 | 0.836501 | 1 |
| Btbd10 | -0.08368 | 0.352073 | 1 |
| Taf9b | -0.08372 | 0.078101 | 1 |
| Tnfaip8l2 | -0.08381 | 0.974789 | 1 |
| Nit1 | -0.08384 | 0.83517 | 1 |
| Ccdc59 | -0.08385 | 0.299954 | 1 |
| Znrd2 | -0.0839 | 0.276081 | 1 |
| Rpia | -0.08393 | 0.713106 | 1 |
| Fbxl14 | -0.08401 | 0.155483 | 1 |
| Mfsd10 | -0.08409 | 0.41934 | 1 |
| Rab21 | -0.08412 | 0.689429 | 1 |
| Pip4k2c | -0.08416 | 0.259857 | 1 |
| Cd81 | -0.08424 | 0.625254 | 1 |
| Dynll2 | -0.08424 | 0.191743 | 1 |
| Cpsf2 | -0.08425 | 0.173904 | 1 |
| Vps35 | -0.08425 | 0.273541 | 1 |
| Vbp1 | -0.08426 | 0.045842 | 1 |
| Capn15 | -0.08433 | 0.072218 | 1 |
| Pmvk | -0.08438 | 0.087149 | 1 |
| Gm43813 | -0.08445 | 0.022447 | 1 |
| Ppp1r21 | -0.08448 | 0.101517 | 1 |
| Rabgef1 | -0.08452 | 0.015007 | 1 |
| Zbtb8os | -0.08456 | 0.233204 | 1 |
| Ikbkg | -0.08458 | 0.318311 | 1 |

| | | | |
|---|---|---|---|
| 2700049A03I | -0.08463 | 0.141028 | 1 |
| Lmf2 | -0.08464 | 0.328817 | 1 |
| Tsc22d4 | -0.08466 | 0.14135 | 1 |
| Utp3 | -0.08475 | 0.11545 | 1 |
| Dnajc18 | -0.08484 | 0.491738 | 1 |
| St6gal1 | -0.08488 | 0.90718 | 1 |
| Lztr1 | -0.08489 | 0.716774 | 1 |
| Spg11 | -0.08491 | 0.263374 | 1 |
| Dync1h1 | -0.08496 | 0.304998 | 1 |
| Hprt | -0.08507 | 0.454593 | 1 |
| Cbfb | -0.08507 | 0.27847 | 1 |
| Lsm3 | -0.08508 | 0.213924 | 1 |
| Zswim6 | -0.08511 | 0.731003 | 1 |
| Lysmd4 | -0.08515 | 0.207519 | 1 |
| Mrpl40 | -0.08516 | 0.198469 | 1 |
| Ino80c | -0.08527 | 0.972627 | 1 |
| Zfp317 | -0.08529 | 0.029011 | 1 |
| Slc52a3 | -0.08529 | 0.624186 | 1 |
| Sapcd2 | -0.08533 | 0.075776 | 1 |
| Ttc13 | -0.0854 | 0.127243 | 1 |
| Zwilch | -0.08548 | 0.03867 | 1 |
| Ndufv2 | -0.08551 | 0.529992 | 1 |
| Cant1 | -0.08553 | 0.692498 | 1 |
| Nfx1 | -0.08553 | 0.251222 | 1 |
| Smg7 | -0.08554 | 0.60973 | 1 |
| Lpar5 | -0.08557 | 0.220935 | 1 |
| Acsl4 | -0.08562 | 0.147526 | 1 |
| Ddost | -0.08569 | 0.205672 | 1 |
| Dusp10 | -0.08573 | 0.499775 | 1 |
| Acoxl | -0.08582 | 0.194248 | 1 |
| Nup37 | -0.08586 | 0.105926 | 1 |
| Nup205 | -0.08587 | 0.223166 | 1 |
| Vdac1 | -0.08607 | 0.382663 | 1 |
| Prkrip1 | -0.08612 | 0.041368 | 1 |
| Thoc6 | -0.08613 | 0.198158 | 1 |
| Msto1 | -0.08636 | 0.449597 | 1 |
| Pcbp4 | -0.08638 | 0.001856 | 1 |
| Snrnp40 | -0.08647 | 0.075766 | 1 |
| Far1 | -0.08648 | 0.207477 | 1 |
| Nfkb1 | -0.08655 | 0.755512 | 1 |
| Phf8 | -0.08658 | 0.054823 | 1 |
| Ppig | -0.08658 | 0.93508 | 1 |
| Cxxc1 | -0.08661 | 0.391473 | 1 |
| Ppat | -0.08663 | 0.204193 | 1 |
| Ghitm | -0.08671 | 0.607289 | 1 |
| Isoc2a | -0.08672 | 0.108236 | 1 |
| Pip5k1c | -0.08676 | 0.066301 | 1 |
| BC005561 | -0.0868 | 0.078446 | 1 |

| | | | |
|---|---|---|---|
| Dhrs1 | -0.08692 | 0.212173 | 1 |
| Exosc10 | -0.08692 | 0.149452 | 1 |
| Tcrg-C4 | -0.08696 | 0.000189 | 1 |
| Ndufa1 | -0.08703 | 0.545725 | 1 |
| Naa40 | -0.08716 | 0.285888 | 1 |
| Arl5a | -0.08728 | 0.02273 | 1 |
| Panx1 | -0.08741 | 0.060489 | 1 |
| Psmd7 | -0.08743 | 0.888227 | 1 |
| Dnajc8 | -0.08746 | 0.237789 | 1 |
| Mnat1 | -0.08752 | 0.285816 | 1 |
| Vdac2 | -0.08757 | 0.921252 | 1 |
| Luc7l2 | -0.08759 | 0.182971 | 1 |
| Ralgps1 | -0.08762 | 0.049356 | 1 |
| Slx4 | -0.08763 | 0.004421 | 1 |
| Bspry | -0.08772 | 0.077575 | 1 |
| Atl2 | -0.08777 | 0.166641 | 1 |
| Psat1 | -0.08797 | 0.065506 | 1 |
| Kif21b | -0.088 | 0.227101 | 1 |
| Fam92a | -0.08802 | 0.661667 | 1 |
| Snrpe | -0.08807 | 0.232122 | 1 |
| Slc22a15 | -0.08809 | 0.096275 | 1 |
| Rab33b | -0.08817 | 0.057577 | 1 |
| Bysl | -0.08821 | 0.05834 | 1 |
| Yipf5 | -0.08835 | 0.57252 | 1 |
| Zfp518a | -0.08841 | 0.27031 | 1 |
| Tpd52 | -0.08849 | 0.024597 | 1 |
| Rad54l | -0.08852 | 0.154118 | 1 |
| Mllt11 | -0.08856 | 0.047637 | 1 |
| Ints6l | -0.08858 | 0.16639 | 1 |
| Dmxl1 | -0.08861 | 0.708561 | 1 |
| Rtel1 | -0.08864 | 0.324865 | 1 |
| Inpp5d | -0.08865 | 0.289481 | 1 |
| Mapk1 | -0.08865 | 0.244575 | 1 |
| Ppp6r3 | -0.08892 | 0.611628 | 1 |
| Rhno1 | -0.08898 | 0.17628 | 1 |
| Mrps22 | -0.08902 | 0.034541 | 1 |
| Hvcn1 | -0.08903 | 0.472826 | 1 |
| Tmem68 | -0.08903 | 0.036361 | 1 |
| Isyna1 | -0.08921 | 0.956226 | 1 |
| Scyl1 | -0.08942 | 0.569384 | 1 |
| Pitpnm1 | -0.08946 | 0.127912 | 1 |
| Cops5 | -0.0895 | 0.792201 | 1 |
| Fam45a | -0.08962 | 0.450133 | 1 |
| Nol10 | -0.08969 | 0.18519 | 1 |
| Gfpt1 | -0.08971 | 0.477967 | 1 |
| Setx | -0.08971 | 0.186295 | 1 |
| Wipf1 | -0.08972 | 0.430494 | 1 |
| Cul3 | -0.08978 | 0.356308 | 1 |

| | | | |
|---|---|---|---|
| Fancd2 | -0.0898 | 0.042764 | 1 |
| Thg1l | -0.08989 | 0.336103 | 1 |
| Anxa4 | -0.08991 | 0.714283 | 1 |
| Coq8b | -0.08995 | 0.227408 | 1 |
| Dnaaf2 | -0.08995 | 0.088199 | 1 |
| Mthfd1l | -0.09 | 0.794389 | 1 |
| Pphln1 | -0.09003 | 0.050674 | 1 |
| Dscc1 | -0.09007 | 0.113314 | 1 |
| Mrpl39 | -0.0902 | 0.222298 | 1 |
| Rpp30 | -0.0902 | 0.119793 | 1 |
| Cdca7 | -0.09021 | 0.045174 | 1 |
| Fbxo22 | -0.09022 | 0.072277 | 1 |
| Phlda3 | -0.09026 | 7.31E-05 | 1 |
| Srp68 | -0.09026 | 0.751733 | 1 |
| Elp4 | -0.09037 | 0.557623 | 1 |
| Nkap | -0.09042 | 0.375691 | 1 |
| Kif5b | -0.09043 | 0.210402 | 1 |
| Fmr1 | -0.09051 | 0.036834 | 1 |
| Ptpre | -0.09059 | 0.275413 | 1 |
| 9830144P21I | -0.09067 | 0.124575 | 1 |
| Gm26917 | -0.09076 | 0.262862 | 1 |
| Tmem229b | -0.09079 | 0.468095 | 1 |
| 1700056E22I | -0.09087 | 0.054379 | 1 |
| Irf5 | -0.09091 | 0.393094 | 1 |
| Pias4 | -0.09104 | 0.308304 | 1 |
| Psmb5 | -0.09109 | 0.512547 | 1 |
| Cd84 | -0.09127 | 0.191434 | 1 |
| Nckap1l | -0.09131 | 0.283711 | 1 |
| Foxn2 | -0.09142 | 0.163178 | 1 |
| Tmbim6 | -0.09143 | 0.114382 | 1 |
| Lipa | -0.09157 | 0.065134 | 1 |
| Zfp729a | -0.09159 | 0.005842 | 1 |
| L1cam | -0.09159 | 0.257084 | 1 |
| Tcp1 | -0.09159 | 0.341405 | 1 |
| Ganc | -0.0916 | 0.927512 | 1 |
| Senp5 | -0.09173 | 0.158052 | 1 |
| Odf2l | -0.09176 | 0.21727 | 1 |
| Med13 | -0.0918 | 0.571433 | 1 |
| Ugdh | -0.0918 | 0.473921 | 1 |
| Ankrd49 | -0.09186 | 0.108857 | 1 |
| Tcn2 | -0.09192 | 0.300194 | 1 |
| Exoc6 | -0.09193 | 0.13611 | 1 |
| Rad1 | -0.09202 | 0.373931 | 1 |
| Map11 | -0.0922 | 0.119335 | 1 |
| Exosc3 | -0.0922 | 0.390178 | 1 |
| Map1s | -0.09221 | 0.349354 | 1 |
| March2 | -0.09223 | 0.574209 | 1 |
| Cacfd1 | -0.09225 | 0.705979 | 1 |

| | | | |
|---|---|---|---|
| Clcn4 | -0.09225 | 0.214379 | 1 |
| Txlna | -0.09226 | 0.37285 | 1 |
| Chid1 | -0.09226 | 0.052722 | 1 |
| Strn3 | -0.09229 | 0.124794 | 1 |
| Fermt3 | -0.0924 | 0.555165 | 1 |
| Dgcr8 | -0.09244 | 0.055003 | 1 |
| Phpt1 | -0.09246 | 0.134528 | 1 |
| Ubxn7 | -0.0925 | 0.485245 | 1 |
| Smu1 | -0.09261 | 0.176788 | 1 |
| Pidd1 | -0.09269 | 0.042237 | 1 |
| 2310022A10I | -0.09271 | 0.15423 | 1 |
| Dlg1 | -0.09283 | 0.138394 | 1 |
| Slc35f2 | -0.09293 | 0.167569 | 1 |
| Rad51b | -0.09297 | 0.07773 | 1 |
| Ankrd26 | -0.09299 | 0.331127 | 1 |
| Haus6 | -0.09308 | 0.048191 | 1 |
| Anxa11 | -0.09309 | 0.701722 | 1 |
| Foxd2os | -0.09317 | 0.010927 | 1 |
| Slc30a6 | -0.09324 | 0.081559 | 1 |
| Gins2 | -0.09326 | 0.079353 | 1 |
| Mrps14 | -0.09328 | 0.292539 | 1 |
| Usp37 | -0.09342 | 0.311893 | 1 |
| Commd2 | -0.09346 | 0.117751 | 1 |
| Ttc28 | -0.09347 | 0.044001 | 1 |
| Trip4 | -0.09353 | 0.174873 | 1 |
| Acyp1 | -0.09363 | 0.009599 | 1 |
| Ncbp2 | -0.09364 | 0.039168 | 1 |
| Cep104 | -0.09364 | 0.679772 | 1 |
| Fbxo25 | -0.09369 | 0.584709 | 1 |
| Tfip11 | -0.09376 | 0.0174 | 1 |
| Sf3b4 | -0.09393 | 0.548086 | 1 |
| Camkk2 | -0.09395 | 0.087826 | 1 |
| Ythdc2 | -0.09396 | 0.129841 | 1 |
| Cep250 | -0.09399 | 0.266483 | 1 |
| Glrx3 | -0.094 | 0.765659 | 1 |
| Kifc5b | -0.09402 | 0.141935 | 1 |
| Rpl7l1 | -0.09421 | 0.437676 | 1 |
| Pde4a | -0.09422 | 0.136496 | 1 |
| Mrfap1 | -0.09424 | 0.356167 | 1 |
| Mettl17 | -0.09432 | 0.341694 | 1 |
| Pex1 | -0.09451 | 0.031052 | 1 |
| Shld1 | -0.09455 | 0.057023 | 1 |
| Rfk | -0.09458 | 0.883512 | 1 |
| Atp13a1 | -0.09464 | 0.100114 | 1 |
| Nup50 | -0.09467 | 0.247944 | 1 |
| Plrg1 | -0.09482 | 0.038849 | 1 |
| BC030867 | -0.09491 | 0.000374 | 1 |
| Cpox | -0.09492 | 0.408791 | 1 |

| | | | |
|---|---|---|---|
| Arid1b | -0.09495 | 0.309524 | 1 |
| Cyb5d2 | -0.09501 | 0.000922 | 1 |
| Tmem222 | -0.09502 | 0.301149 | 1 |
| Chek1 | -0.09515 | 0.164555 | 1 |
| Cenpv | -0.0952 | 0.00066 | 1 |
| Dtnb | -0.09521 | 0.099994 | 1 |
| Gnai3 | -0.09529 | 0.533025 | 1 |
| Kif2a | -0.09531 | 0.876172 | 1 |
| Gm14029 | -0.09539 | 0.222588 | 1 |
| Prpf40a | -0.09544 | 0.196563 | 1 |
| Wnk1 | -0.09545 | 0.183224 | 1 |
| Umad1 | -0.09558 | 0.678671 | 1 |
| Sf3b3 | -0.09567 | 0.749821 | 1 |
| Cox6b1 | -0.09568 | 0.171719 | 1 |
| Ptch1 | -0.09587 | 0.346551 | 1 |
| Tm9sf4 | -0.0959 | 0.226399 | 1 |
| Gm33782 | -0.09599 | 0.058482 | 1 |
| Xylt1 | -0.09604 | 0.165173 | 1 |
| Cdc42ep3 | -0.09605 | 0.517346 | 1 |
| Creg1 | -0.09605 | 0.982526 | 1 |
| Slf2 | -0.09612 | 0.14655 | 1 |
| Psmc6 | -0.09615 | 0.325497 | 1 |
| Etv3 | -0.09619 | 0.311848 | 1 |
| Mlec | -0.09619 | 0.413416 | 1 |
| Selenot | -0.09621 | 0.505684 | 1 |
| Tmem171 | -0.09633 | 0.097596 | 1 |
| Bud13 | -0.09634 | 0.172446 | 1 |
| Gatad2b | -0.09638 | 0.107248 | 1 |
| Zfp407 | -0.09645 | 0.090611 | 1 |
| Gbf1 | -0.09645 | 0.507988 | 1 |
| Mphosph9 | -0.09651 | 0.138147 | 1 |
| Lmnb2 | -0.09655 | 0.011742 | 1 |
| Pstk | -0.09656 | 0.209535 | 1 |
| Ints10 | -0.09657 | 0.264705 | 1 |
| Idi1 | -0.09666 | 0.524557 | 1 |
| Pik3c2a | -0.09666 | 0.135215 | 1 |
| N4bp2 | -0.09672 | 0.044411 | 1 |
| Ercc6l | -0.09672 | 4.65E-07 | 0.015019 |
| Otub1 | -0.09673 | 0.467178 | 1 |
| Smim11 | -0.09675 | 0.280901 | 1 |
| Gm20069 | -0.09676 | 0.045653 | 1 |
| Cd164 | -0.0968 | 0.542467 | 1 |
| Abhd5 | -0.09682 | 0.01149 | 1 |
| Tstd2 | -0.09692 | 0.078744 | 1 |
| Slc12a6 | -0.09694 | 0.244224 | 1 |
| Sf3a3 | -0.09697 | 0.0964 | 1 |
| Syne2 | -0.09711 | 0.044195 | 1 |
| Puf60 | -0.09719 | 0.186366 | 1 |

| | | | |
|---|---|---|---|
| Asb7 | -0.09719 | 0.01642 | 1 |
| Ralbp1 | -0.0972 | 0.585473 | 1 |
| Grb2 | -0.09726 | 0.254192 | 1 |
| Ythdf2 | -0.09727 | 0.563287 | 1 |
| Moap1 | -0.09733 | 0.008219 | 1 |
| Xndc1 | -0.09745 | 0.028652 | 1 |
| Taf1 | -0.09746 | 0.315024 | 1 |
| Papss1 | -0.09761 | 0.121779 | 1 |
| Med21 | -0.09767 | 0.023868 | 1 |
| Copg2 | -0.09769 | 0.191022 | 1 |
| Heatr9 | -0.09771 | 0.000832 | 1 |
| Fdxr | -0.09772 | 0.017654 | 1 |
| Haus1 | -0.09774 | 0.064194 | 1 |
| Raph1 | -0.09795 | 0.262901 | 1 |
| Pigt | -0.09796 | 0.190401 | 1 |
| Bclaf3 | -0.09799 | 0.008369 | 1 |
| Ddx23 | -0.09802 | 0.283882 | 1 |
| Cxcr4 | -0.09815 | 0.053177 | 1 |
| Smndc1 | -0.09822 | 0.055701 | 1 |
| Usp14 | -0.0983 | 0.046474 | 1 |
| Mtmr6 | -0.09842 | 0.250337 | 1 |
| Yeats4 | -0.09846 | 0.229545 | 1 |
| Inpp5f | -0.09846 | 0.106292 | 1 |
| Gtf2h2 | -0.09851 | 0.03698 | 1 |
| Itih5 | -0.09853 | 0.265685 | 1 |
| Trmt13 | -0.09854 | 0.684432 | 1 |
| 9530068E07I | -0.0988 | 0.071076 | 1 |
| Mrpl44 | -0.09881 | 0.275923 | 1 |
| Setd2 | -0.09906 | 0.274702 | 1 |
| Sept6 | -0.09907 | 0.097269 | 1 |
| Cyb5r3 | -0.09908 | 0.19267 | 1 |
| Elf2 | -0.0991 | 0.108913 | 1 |
| Acer3 | -0.09912 | 0.163954 | 1 |
| Armc1 | -0.09918 | 0.244887 | 1 |
| Gtse1 | -0.09919 | 0.108016 | 1 |
| Nip7 | -0.09921 | 0.671856 | 1 |
| Mfsd14a | -0.09933 | 0.175243 | 1 |
| Pdk3 | -0.09934 | 0.112971 | 1 |
| Cnot7 | -0.09938 | 0.067951 | 1 |
| Acd | -0.09939 | 0.077451 | 1 |
| Slx4ip | -0.09943 | 0.87794 | 1 |
| 2610318N02 | -0.09945 | 0.029908 | 1 |
| Pck2 | -0.09955 | 0.108134 | 1 |
| Zmym5 | -0.09955 | 0.055427 | 1 |
| Gm30054 | -0.09955 | 0.342987 | 1 |
| Dync2h1 | -0.09962 | 0.056049 | 1 |
| Tgoln1 | -0.09964 | 0.751321 | 1 |
| Dgkz | -0.09968 | 0.927189 | 1 |

| | | | |
|---|---|---|---|
| Bard1 | -0.09969 | 0.01176 | 1 |
| Anp32a | -0.09971 | 0.109938 | 1 |
| U2surp | -0.09973 | 0.396989 | 1 |
| Neat1 | -0.09976 | 0.181746 | 1 |
| Ndufa13 | -0.09982 | 0.049993 | 1 |
| Naa16 | -0.09988 | 0.249625 | 1 |
| Cdc27 | -0.09988 | 0.352706 | 1 |
| Ppt1 | -0.09988 | 0.079642 | 1 |
| Rnf31 | -0.09991 | 0.003639 | 1 |
| Yif1a | -0.09993 | 0.030586 | 1 |
| Lcor | -0.09996 | 0.441569 | 1 |
| Pml | -0.10002 | 0.129226 | 1 |
| Gins4 | -0.10007 | 0.222148 | 1 |
| Acsbg1 | -0.10022 | 0.06597 | 1 |
| Adck5 | -0.1003 | 0.351144 | 1 |
| 3110009E18I | -0.10031 | 0.061363 | 1 |
| Lcmt2 | -0.10033 | 0.130821 | 1 |
| Lmbr1l | -0.10033 | 0.201205 | 1 |
| Ttc33 | -0.10036 | 0.10592 | 1 |
| Mindy1 | -0.1004 | 0.202064 | 1 |
| Trp53inp1 | -0.10044 | 0.085403 | 1 |
| Pcbp1 | -0.10046 | 0.094966 | 1 |
| Dennd1c | -0.1005 | 0.019311 | 1 |
| Pif1 | -0.1006 | 0.010307 | 1 |
| Rbm6 | -0.1006 | 0.094529 | 1 |
| Srsf6 | -0.10068 | 0.594012 | 1 |
| Jpt2 | -0.10082 | 0.021698 | 1 |
| Zkscan17 | -0.10087 | 0.119122 | 1 |
| Syne3 | -0.10096 | 0.240567 | 1 |
| Ywhae | -0.10097 | 0.221082 | 1 |
| Rb1 | -0.10124 | 0.202984 | 1 |
| Ccdc22 | -0.10128 | 0.010439 | 1 |
| Pik3cg | -0.10132 | 0.74926 | 1 |
| Zfp710 | -0.10134 | 0.033907 | 1 |
| Umps | -0.10136 | 0.118783 | 1 |
| Gcn1 | -0.10136 | 0.293591 | 1 |
| Wdr91 | -0.1014 | 0.21735 | 1 |
| Zc3h11a | -0.10146 | 0.030606 | 1 |
| Cd1d1 | -0.10151 | 0.017998 | 1 |
| Lrrc42 | -0.10155 | 0.363355 | 1 |
| Dcaf17 | -0.10156 | 0.02556 | 1 |
| Rhot1 | -0.10158 | 0.280366 | 1 |
| Btbd1 | -0.10159 | 0.107168 | 1 |
| Rev3l | -0.10161 | 0.115046 | 1 |
| Polh | -0.10161 | 0.177112 | 1 |
| Mgat5 | -0.10168 | 0.743936 | 1 |
| Hdgfl3 | -0.10177 | 0.402192 | 1 |
| Spidr | -0.10198 | 0.006791 | 1 |

| | | | |
|---|---|---|---|
| Ccdc88b | -0.10202 | 0.19207 | 1 |
| Mrpl43 | -0.10212 | 0.184454 | 1 |
| Mrps15 | -0.10217 | 0.649291 | 1 |
| 1300002E11I | -0.10221 | 0.193244 | 1 |
| Micos10 | -0.10229 | 0.244243 | 1 |
| Recql5 | -0.10233 | 0.143382 | 1 |
| Grpel2 | -0.10236 | 0.012358 | 1 |
| Ensa | -0.10238 | 0.062174 | 1 |
| Ddx47 | -0.10245 | 0.399687 | 1 |
| Cggbp1 | -0.10247 | 0.126412 | 1 |
| Tonsl | -0.10248 | 0.038449 | 1 |
| Ints1 | -0.10252 | 0.4038 | 1 |
| Zfp3 | -0.1026 | 0.004387 | 1 |
| Thap3 | -0.1027 | 0.25309 | 1 |
| S100a1 | -0.10289 | 0.038691 | 1 |
| Pxk | -0.10301 | 0.321341 | 1 |
| Zfp707 | -0.10304 | 0.006569 | 1 |
| Ebp | -0.10306 | 0.043493 | 1 |
| Mapk7 | -0.10308 | 0.129477 | 1 |
| Ddhd1 | -0.10309 | 0.508259 | 1 |
| Tmed9 | -0.1031 | 0.273758 | 1 |
| Mkrn2 | -0.10316 | 0.253572 | 1 |
| Rbms2 | -0.10318 | 0.011341 | 1 |
| Gabpb2 | -0.1032 | 0.287899 | 1 |
| Msrb1 | -0.10324 | 0.311281 | 1 |
| Uchl5 | -0.10329 | 0.504001 | 1 |
| Smad3 | -0.10346 | 0.026984 | 1 |
| Rgs2 | -0.10357 | 0.563447 | 1 |
| Vrk3 | -0.10363 | 0.267424 | 1 |
| Dnttip2 | -0.10364 | 0.464998 | 1 |
| Triap1 | -0.10364 | 0.444092 | 1 |
| Smap1 | -0.10365 | 0.211355 | 1 |
| Eif4a3 | -0.10365 | 0.461764 | 1 |
| Rps6ka5 | -0.10372 | 0.023565 | 1 |
| Zfp367 | -0.10379 | 0.164935 | 1 |
| Rtf1 | -0.10394 | 0.100107 | 1 |
| Prpf38b | -0.10402 | 0.416265 | 1 |
| Asnsd1 | -0.10402 | 0.073602 | 1 |
| Ell | -0.10411 | 0.107978 | 1 |
| Tmc6 | -0.10419 | 0.182344 | 1 |
| 2310009A05I | -0.10421 | 0.211559 | 1 |
| St3gal3 | -0.10422 | 0.18897 | 1 |
| Cog6 | -0.10434 | 0.026868 | 1 |
| Naglu | -0.10434 | 0.005873 | 1 |
| Erlin2 | -0.10454 | 0.004878 | 1 |
| Mrps18b | -0.10479 | 0.839777 | 1 |
| Fuca2 | -0.10482 | 0.266979 | 1 |
| Slc9a7 | -0.10495 | 0.023295 | 1 |

| | | | |
|---|---|---|---|
| Phf13 | -0.10498 | 0.000103 | 1 |
| Ddx52 | -0.10509 | 0.170332 | 1 |
| Cops3 | -0.10511 | 0.100983 | 1 |
| Sp1 | -0.10513 | 0.577573 | 1 |
| Rars | -0.10515 | 0.089156 | 1 |
| Tspan4 | -0.10517 | 0.338451 | 1 |
| Zfp58 | -0.10517 | 0.005271 | 1 |
| Cdc73 | -0.10521 | 0.146344 | 1 |
| Trmt10a | -0.10523 | 0.128488 | 1 |
| Gm26782 | -0.10523 | 0.415034 | 1 |
| Snapc5 | -0.10533 | 0.087491 | 1 |
| Ptdss1 | -0.10552 | 0.052098 | 1 |
| Psmc3 | -0.10557 | 0.276261 | 1 |
| Saraf | -0.10563 | 0.401119 | 1 |
| Casp7 | -0.10566 | 0.213177 | 1 |
| Arl8b | -0.10571 | 0.485321 | 1 |
| Tent2 | -0.10585 | 0.178881 | 1 |
| Traip | -0.10598 | 0.013468 | 1 |
| Zbtb24 | -0.10604 | 0.035954 | 1 |
| Comt | -0.10613 | 0.545218 | 1 |
| Rsrc2 | -0.10613 | 0.253433 | 1 |
| Nutf2 | -0.10615 | 0.035481 | 1 |
| Stip1 | -0.10616 | 0.163036 | 1 |
| Helq | -0.10617 | 0.001055 | 1 |
| Rida | -0.10622 | 0.160371 | 1 |
| Sh2d2a | -0.10623 | 0.693395 | 1 |
| Nqo2 | -0.10635 | 0.025055 | 1 |
| Ublcp1 | -0.10637 | 0.360613 | 1 |
| Slc25a36 | -0.10641 | 0.350868 | 1 |
| Msh3 | -0.10645 | 0.213303 | 1 |
| Zfp266 | -0.1065 | 0.825911 | 1 |
| Hexim1 | -0.10653 | 0.638111 | 1 |
| Ranbp10 | -0.10655 | 0.092842 | 1 |
| Haus8 | -0.10656 | 0.115962 | 1 |
| Gm28901 | -0.1067 | 0.077942 | 1 |
| Tyw1 | -0.10676 | 0.007518 | 1 |
| Hirip3 | -0.10684 | 0.087679 | 1 |
| Snx17 | -0.10689 | 0.195401 | 1 |
| Dctpp1 | -0.10689 | 0.52992 | 1 |
| Zc3h15 | -0.10693 | 0.235795 | 1 |
| Psmd8 | -0.10694 | 0.237912 | 1 |
| Kif1c | -0.107 | 0.014086 | 1 |
| Atp8a1 | -0.10702 | 0.203833 | 1 |
| Tpd52l2 | -0.10702 | 0.869074 | 1 |
| Rnasel | -0.10716 | 0.492094 | 1 |
| Pcyt2 | -0.10716 | 0.075799 | 1 |
| Slc25a38 | -0.10727 | 0.508805 | 1 |
| Ccm2 | -0.10743 | 0.052248 | 1 |

| | | | |
|---|---|---|---|
| Ppdpf | -0.10745 | 0.579195 | 1 |
| Ttc21b | -0.1075 | 0.02836 | 1 |
| Ier3ip1 | -0.10754 | 0.226023 | 1 |
| Smim15 | -0.1076 | 0.07916 | 1 |
| Hexa | -0.10764 | 0.183524 | 1 |
| Wdr20 | -0.1077 | 0.095748 | 1 |
| Chtf18 | -0.10783 | 0.001892 | 1 |
| Ubtf | -0.10791 | 0.737869 | 1 |
| Dazap1 | -0.10797 | 0.509338 | 1 |
| Lyrm7 | -0.10806 | 0.24882 | 1 |
| Wdr95 | -0.10808 | 0.00745 | 1 |
| Eloa | -0.10809 | 0.170044 | 1 |
| Zfp382 | -0.10812 | 0.035562 | 1 |
| Arfip1 | -0.10813 | 0.071391 | 1 |
| Nup35 | -0.10815 | 0.262199 | 1 |
| H2-Q7 | -0.1082 | 0.082628 | 1 |
| Cyb5b | -0.10828 | 0.109043 | 1 |
| Cd2 | -0.10841 | 0.36548 | 1 |
| Serpinb9 | -0.10841 | 0.521226 | 1 |
| Hadha | -0.10845 | 0.211239 | 1 |
| Sec23b | -0.10846 | 0.122 | 1 |
| Oaz2 | -0.1086 | 0.091488 | 1 |
| Trim33 | -0.10877 | 0.379183 | 1 |
| Foxo1 | -0.10878 | 0.816902 | 1 |
| Mcl1 | -0.1089 | 0.417233 | 1 |
| Zfp513 | -0.10906 | 0.002314 | 1 |
| Hist1h3a | -0.10913 | 0.001473 | 1 |
| Hist4h4 | -0.10924 | 0.01626 | 1 |
| Ccdc167 | -0.10928 | 0.117426 | 1 |
| Ccnt2 | -0.1094 | 0.023172 | 1 |
| Snrpa1 | -0.1094 | 0.402024 | 1 |
| Gm33460 | -0.10948 | 2.85E-09 | 9.19E-05 |
| Tmprss13 | -0.10955 | 1.68E-11 | 5.42E-07 |
| Ipo8 | -0.10973 | 0.042062 | 1 |
| Cgas | -0.10975 | 0.002749 | 1 |
| Usp40 | -0.1098 | 0.060571 | 1 |
| Herpud1 | -0.10982 | 0.747871 | 1 |
| Vac14 | -0.10983 | 0.394963 | 1 |
| Tefm | -0.10995 | 0.339991 | 1 |
| Gabpa | -0.10997 | 0.005503 | 1 |
| Tardbp | -0.1101 | 0.287404 | 1 |
| Ddx46 | -0.11011 | 0.255565 | 1 |
| Csnk1g3 | -0.11016 | 0.076714 | 1 |
| Gpn1 | -0.11016 | 0.149359 | 1 |
| Slc25a11 | -0.11018 | 0.124876 | 1 |
| Timeless | -0.11019 | 0.061886 | 1 |
| Usp1 | -0.11027 | 0.06435 | 1 |
| Ap1s1 | -0.11036 | 0.146103 | 1 |

| | | | |
|---|---|---|---|
| Ndufb7 | -0.11038 | 0.289474 | 1 |
| Rtn4 | -0.11064 | 0.532631 | 1 |
| Sp2 | -0.11066 | 0.067273 | 1 |
| Pqbp1 | -0.11072 | 0.21403 | 1 |
| Uqcrc2 | -0.1108 | 0.127477 | 1 |
| Wipf2 | -0.11083 | 0.274341 | 1 |
| Pbdc1 | -0.11087 | 0.087492 | 1 |
| Usp22 | -0.11103 | 0.377944 | 1 |
| Entpd7 | -0.11113 | 0.005062 | 1 |
| Lamp2 | -0.11114 | 0.851096 | 1 |
| Cand1 | -0.11124 | 0.199246 | 1 |
| Stk39 | -0.11136 | 0.38071 | 1 |
| Ulk4 | -0.11137 | 0.004027 | 1 |
| Ddx11 | -0.11143 | 0.008815 | 1 |
| Mtfr2 | -0.11146 | 0.044497 | 1 |
| D6Wsu163e | -0.11148 | 0.009706 | 1 |
| Bcap31 | -0.11162 | 0.118681 | 1 |
| Snx11 | -0.11169 | 0.468684 | 1 |
| Snx12 | -0.11171 | 0.397931 | 1 |
| Ing3 | -0.11175 | 0.732791 | 1 |
| Zdhhc20 | -0.1118 | 0.472433 | 1 |
| Taf6 | -0.11186 | 0.10566 | 1 |
| Ppp3ca | -0.11191 | 0.067978 | 1 |
| 2900026A02 | -0.11192 | 0.430109 | 1 |
| Mdm1 | -0.11192 | 0.028375 | 1 |
| Manea | -0.11195 | 0.073787 | 1 |
| Lsm1 | -0.11195 | 0.111007 | 1 |
| Sucla2 | -0.112 | 0.150654 | 1 |
| Rap2b | -0.11206 | 0.05672 | 1 |
| Itgax | -0.11207 | 0.676264 | 1 |
| Akna | -0.11208 | 0.352603 | 1 |
| Snx4 | -0.11208 | 0.327097 | 1 |
| Anxa5 | -0.11208 | 0.246133 | 1 |
| Usp42 | -0.11214 | 0.150472 | 1 |
| Dhx15 | -0.11215 | 0.313906 | 1 |
| Mfsd14b | -0.1122 | 0.035671 | 1 |
| Lars2 | -0.11223 | 0.000812 | 1 |
| Rin3 | -0.11227 | 0.2465 | 1 |
| Ccnb1ip1 | -0.11227 | 0.000788 | 1 |
| Fam76a | -0.11229 | 0.452903 | 1 |
| B230307C23 | -0.11229 | 0.091925 | 1 |
| Eri2 | -0.11234 | 0.002671 | 1 |
| Rpp25l | -0.11238 | 0.557895 | 1 |
| Nae1 | -0.1124 | 0.036532 | 1 |
| Ggt1 | -0.1125 | 6.9E-07 | 0.022266 |
| Tyk2 | -0.11259 | 0.029693 | 1 |
| Chst10 | -0.11268 | 0.031444 | 1 |
| Ptpa | -0.11269 | 0.443233 | 1 |

| | | | |
|---|---|---|---|
| Txn1 | -0.11285 | 0.716453 | 1 |
| Amd1 | -0.11291 | 0.039464 | 1 |
| Lin9 | -0.11291 | 0.004419 | 1 |
| S100pbp | -0.11294 | 0.303246 | 1 |
| Armc3 | -0.11294 | 0.431832 | 1 |
| Fasl | -0.11296 | 0.360505 | 1 |
| Klhl18 | -0.11297 | 0.069135 | 1 |
| Msh6 | -0.11316 | 0.012305 | 1 |
| Igsf8 | -0.11333 | 0.909626 | 1 |
| Gpcpd1 | -0.11335 | 0.228245 | 1 |
| Egln2 | -0.11343 | 0.079485 | 1 |
| Melk | -0.11347 | 0.011843 | 1 |
| Atp5j2 | -0.11367 | 0.036849 | 1 |
| Ylpm1 | -0.11373 | 0.221989 | 1 |
| Anks1b | -0.11391 | 0.020182 | 1 |
| Ppfia1 | -0.11391 | 0.129535 | 1 |
| Inf2 | -0.11398 | 0.259837 | 1 |
| Tnfaip8l1 | -0.11399 | 0.216201 | 1 |
| Tcp11l2 | -0.11401 | 0.89277 | 1 |
| Zfyve16 | -0.11402 | 0.282074 | 1 |
| Vamp1 | -0.11404 | 0.033985 | 1 |
| Nop58 | -0.11407 | 0.424416 | 1 |
| Eapp | -0.11412 | 0.309567 | 1 |
| Praf2 | -0.11415 | 0.100421 | 1 |
| Gm15564 | -0.11425 | 0.240346 | 1 |
| Gm20663 | -0.11427 | 0.121712 | 1 |
| Tmem135 | -0.11429 | 0.119676 | 1 |
| Zfp384 | -0.11436 | 0.044552 | 1 |
| Arrdc1 | -0.11459 | 0.174734 | 1 |
| Commd4 | -0.11487 | 0.44322 | 1 |
| Szrd1 | -0.11506 | 0.237702 | 1 |
| Ankmy2 | -0.11508 | 0.303596 | 1 |
| Ier5 | -0.11517 | 0.390831 | 1 |
| Ythdf1 | -0.11518 | 0.316044 | 1 |
| Arpc2 | -0.11525 | 0.066165 | 1 |
| Kbtbd4 | -0.11534 | 0.475215 | 1 |
| Scrib | -0.11543 | 0.000744 | 1 |
| Stard5 | -0.11545 | 0.020258 | 1 |
| Entpd1 | -0.11547 | 0.056442 | 1 |
| Gm50287 | -0.11549 | 0.023244 | 1 |
| Cstf2 | -0.1155 | 0.028134 | 1 |
| Ftl1 | -0.1155 | 0.860691 | 1 |
| Snx30 | -0.1156 | 0.009119 | 1 |
| Atf7 | -0.11562 | 0.233697 | 1 |
| Ap1g1 | -0.11565 | 0.116765 | 1 |
| Spag7 | -0.1157 | 0.048705 | 1 |
| Lemd2 | -0.11571 | 0.041264 | 1 |
| Lrrc8a | -0.11573 | 0.065264 | 1 |

| | | | |
|---|---|---|---|
| Nxt1 | -0.11576 | 0.220882 | 1 |
| Slc30a5 | -0.11577 | 0.477952 | 1 |
| Fbxo28 | -0.11579 | 0.163736 | 1 |
| Zfp68 | -0.11584 | 0.050822 | 1 |
| Poc1a | -0.11589 | 0.075223 | 1 |
| Smarca2 | -0.1159 | 0.133399 | 1 |
| Dnajc14 | -0.1159 | 0.104461 | 1 |
| Mtmr14 | -0.11591 | 0.185834 | 1 |
| Hibch | -0.11597 | 0.088156 | 1 |
| Ccdc82 | -0.11602 | 0.320628 | 1 |
| Rabl3 | -0.1161 | 0.110822 | 1 |
| Mrpl17 | -0.11617 | 0.64714 | 1 |
| Inpp5b | -0.11621 | 0.137194 | 1 |
| Mpv17 | -0.11625 | 0.048927 | 1 |
| Eif4g3 | -0.11629 | 0.055664 | 1 |
| Ccdc61 | -0.1163 | 0.025575 | 1 |
| Tfg | -0.1163 | 0.31191 | 1 |
| Ccdc124 | -0.11635 | 0.473333 | 1 |
| Hnrnpdl | -0.1164 | 0.789271 | 1 |
| Mios | -0.11644 | 0.327064 | 1 |
| Fam3a | -0.11646 | 0.012169 | 1 |
| Zranb1 | -0.11657 | 0.077441 | 1 |
| Tox2 | -0.11663 | 0.003742 | 1 |
| Clic4 | -0.11666 | 0.54727 | 1 |
| Cdc14b | -0.11666 | 0.003105 | 1 |
| Armc5 | -0.11668 | 0.027992 | 1 |
| Wbp4 | -0.11669 | 0.262518 | 1 |
| Gusb | -0.11671 | 0.139079 | 1 |
| Pnp | -0.11679 | 0.080202 | 1 |
| Hikeshi | -0.11682 | 0.880718 | 1 |
| Bcor | -0.11698 | 0.267554 | 1 |
| Gpkow | -0.11714 | 0.197359 | 1 |
| Pex5 | -0.1172 | 0.077013 | 1 |
| Prag1 | -0.1172 | 0.017073 | 1 |
| Arhgap25 | -0.11736 | 0.625263 | 1 |
| Gimap7 | -0.11745 | 0.201789 | 1 |
| Zdhhc21 | -0.11751 | 0.435873 | 1 |
| Dctn6 | -0.11752 | 0.26962 | 1 |
| Osbpl5 | -0.11754 | 0.154993 | 1 |
| Bbs9 | -0.11755 | 0.322338 | 1 |
| Gpr137b | -0.11757 | 9.62E-06 | 0.310529 |
| Mbd4 | -0.11766 | 0.003347 | 1 |
| Vamp3 | -0.11785 | 0.079533 | 1 |
| Ctdspl2 | -0.11786 | 0.421929 | 1 |
| Ly9 | -0.11788 | 0.117899 | 1 |
| Gtf3a | -0.11792 | 0.073912 | 1 |
| Sc5d | -0.11796 | 0.06358 | 1 |
| Atp6v0a2 | -0.118 | 0.025672 | 1 |

| | | | |
|---|---|---|---|
| Midn | -0.1181 | 0.078211 | 1 |
| Safb2 | -0.11811 | 0.488659 | 1 |
| Egln1 | -0.11828 | 0.290504 | 1 |
| Eif4b | -0.11829 | 0.15213 | 1 |
| Brd3 | -0.11838 | 0.134667 | 1 |
| Spock2 | -0.1184 | 0.002303 | 1 |
| Alas1 | -0.11843 | 0.092221 | 1 |
| Rassf1 | -0.1185 | 0.704824 | 1 |
| Ctps2 | -0.11852 | 0.12357 | 1 |
| Tmem175 | -0.11853 | 0.079244 | 1 |
| Pqlc3 | -0.11854 | 0.036405 | 1 |
| Cnot3 | -0.11855 | 0.004094 | 1 |
| Zfp113 | -0.11857 | 0.019285 | 1 |
| Kmt5c | -0.11863 | 0.089681 | 1 |
| Il16 | -0.1188 | 0.453405 | 1 |
| Gak | -0.11882 | 0.019078 | 1 |
| Rbm27 | -0.11888 | 0.425227 | 1 |
| Pbrm1 | -0.11905 | 0.640778 | 1 |
| Arhgap18 | -0.11911 | 0.737987 | 1 |
| Foxm1 | -0.11911 | 0.04908 | 1 |
| Brd4 | -0.11915 | 0.141171 | 1 |
| Ikzf1 | -0.11918 | 0.256627 | 1 |
| Ttk | -0.11923 | 0.002103 | 1 |
| Flot1 | -0.11928 | 0.33726 | 1 |
| Rbbp5 | -0.11936 | 0.193034 | 1 |
| Atad2b | -0.11941 | 0.464897 | 1 |
| Fdps | -0.11946 | 0.693894 | 1 |
| Hcfc1 | -0.1195 | 0.069816 | 1 |
| Fubp3 | -0.1195 | 0.004158 | 1 |
| Gps2 | -0.11952 | 0.091559 | 1 |
| Mms22l | -0.11953 | 0.093975 | 1 |
| Chaf1a | -0.11962 | 0.115805 | 1 |
| Cstf3 | -0.11982 | 0.094348 | 1 |
| Uso1 | -0.11988 | 0.046819 | 1 |
| Ireb2 | -0.11991 | 0.216804 | 1 |
| Orai2 | -0.12007 | 0.121333 | 1 |
| Klhl11 | -0.12026 | 0.002908 | 1 |
| Serpinb9b | -0.1203 | 0.747185 | 1 |
| Fam111a | -0.12035 | 0.098321 | 1 |
| Rprd2 | -0.12039 | 0.29265 | 1 |
| Ppm1b | -0.12044 | 0.02733 | 1 |
| Osm | -0.12048 | 0.374475 | 1 |
| Ruvbl2 | -0.12069 | 0.043745 | 1 |
| Dpp8 | -0.12073 | 0.326297 | 1 |
| Arcn1 | -0.12096 | 0.621736 | 1 |
| Morc2a | -0.12097 | 0.540349 | 1 |
| Abhd11 | -0.12101 | 0.156764 | 1 |
| Dcp2 | -0.12105 | 0.297485 | 1 |

| | | | |
|---|---|---|---|
| Taf12 | -0.12107 | 0.011912 | 1 |
| Arrb2 | -0.12127 | 0.651224 | 1 |
| Zfp943 | -0.12131 | 0.131859 | 1 |
| Kdm2a | -0.12136 | 0.152259 | 1 |
| Parp1 | -0.1215 | 0.207494 | 1 |
| Zbtb1 | -0.1215 | 0.041157 | 1 |
| Trappc4 | -0.12162 | 0.101072 | 1 |
| Scly | -0.12165 | 0.063115 | 1 |
| E430024P14I | -0.12174 | 0.109668 | 1 |
| Ppp1r10 | -0.12178 | 0.183544 | 1 |
| Cdc7 | -0.12179 | 0.003039 | 1 |
| Zw10 | -0.12179 | 0.065833 | 1 |
| Scaf1 | -0.12189 | 0.036237 | 1 |
| Shkbp1 | -0.12193 | 0.069828 | 1 |
| Dtx3 | -0.12193 | 0.338243 | 1 |
| Rhof | -0.12193 | 0.229396 | 1 |
| Gm37240 | -0.12206 | 0.029203 | 1 |
| Rfx1 | -0.12209 | 0.000942 | 1 |
| 4930579G24 | -0.12211 | 0.007323 | 1 |
| Bnip2 | -0.12215 | 0.092425 | 1 |
| Smim12 | -0.12229 | 0.8097 | 1 |
| Smarca5 | -0.1223 | 0.088561 | 1 |
| Pspc1 | -0.1224 | 0.264273 | 1 |
| Il17ra | -0.12245 | 0.304567 | 1 |
| U2af1 | -0.12246 | 0.258306 | 1 |
| Kntc1 | -0.12248 | 0.063282 | 1 |
| Dhx8 | -0.12267 | 0.032098 | 1 |
| Ier3 | -0.12271 | 0.009328 | 1 |
| Rnf215 | -0.1228 | 0.260052 | 1 |
| Purb | -0.12283 | 0.067617 | 1 |
| Srp54a | -0.12335 | 0.055626 | 1 |
| Qrich1 | -0.12335 | 0.176996 | 1 |
| 1110004F10I | -0.12344 | 0.5107 | 1 |
| Cacna1d | -0.1235 | 0.009196 | 1 |
| Emd | -0.12356 | 0.490728 | 1 |
| Cltc | -0.12358 | 0.095624 | 1 |
| Sfswap | -0.12366 | 0.100169 | 1 |
| Nfatc3 | -0.12369 | 0.191246 | 1 |
| Pak2 | -0.1237 | 0.233477 | 1 |
| Smg9 | -0.12375 | 0.052958 | 1 |
| Csnk1d | -0.12384 | 0.655924 | 1 |
| Csnk2a1 | -0.12385 | 0.164751 | 1 |
| Ubn1 | -0.12387 | 0.172391 | 1 |
| Xpo1 | -0.12396 | 0.416712 | 1 |
| Ppard | -0.12401 | 0.029508 | 1 |
| Klhdc4 | -0.12404 | 0.252794 | 1 |
| Sh3gl1 | -0.12414 | 0.280916 | 1 |
| Fbxw8 | -0.12429 | 0.01501 | 1 |

| | | | |
|---|---|---|---|
| Prdx4 | -0.12431 | 0.412411 | 1 |
| Rnf6 | -0.1244 | 0.027725 | 1 |
| Abcd1 | -0.1244 | 0.086044 | 1 |
| Syt11 | -0.12447 | 0.02336 | 1 |
| Tcf12 | -0.12452 | 0.268661 | 1 |
| Hnrnpa0 | -0.12471 | 0.009807 | 1 |
| Chd3 | -0.12471 | 0.13214 | 1 |
| Heatr1 | -0.12488 | 0.241726 | 1 |
| Eif3j1 | -0.12488 | 0.024829 | 1 |
| Gmeb1 | -0.12505 | 0.004387 | 1 |
| Stt3a | -0.12519 | 0.11285 | 1 |
| Atp11c | -0.12545 | 0.097218 | 1 |
| Map3k1 | -0.12549 | 0.04853 | 1 |
| Sept4 | -0.12552 | 2.31E-05 | 0.747163 |
| Mir99ahg | -0.12556 | 2.11E-07 | 0.006806 |
| Pdia3 | -0.1257 | 0.303106 | 1 |
| Ncald | -0.1257 | 0.106582 | 1 |
| Vps36 | -0.1258 | 0.34224 | 1 |
| Fkbp4 | -0.1258 | 0.329108 | 1 |
| Mob1a | -0.12603 | 0.159094 | 1 |
| Dapk3 | -0.12613 | 0.193876 | 1 |
| Osbpl2 | -0.12613 | 0.124893 | 1 |
| Plekha2 | -0.12615 | 0.516621 | 1 |
| Tnik | -0.12622 | 0.4675 | 1 |
| Fam107b | -0.12632 | 0.206835 | 1 |
| Msantd2 | -0.12636 | 0.487475 | 1 |
| Psmd11 | -0.12646 | 0.077074 | 1 |
| Dcaf7 | -0.12657 | 0.056204 | 1 |
| Pdzd11 | -0.12659 | 0.691648 | 1 |
| Scaf4 | -0.12668 | 0.049042 | 1 |
| Gpbp1 | -0.1268 | 0.492904 | 1 |
| Zfp931 | -0.12695 | 0.032731 | 1 |
| Brpf1 | -0.12711 | 0.033041 | 1 |
| Ndufa8 | -0.12735 | 0.209457 | 1 |
| Dennd4b | -0.12735 | 0.156868 | 1 |
| Rrp9 | -0.12747 | 0.260484 | 1 |
| Relt | -0.12748 | 0.171628 | 1 |
| Brwd3 | -0.12754 | 0.036431 | 1 |
| Uimc1 | -0.12764 | 0.129807 | 1 |
| Tmem256 | -0.12768 | 0.470445 | 1 |
| Drap1 | -0.1277 | 0.008512 | 1 |
| Mns1 | -0.12774 | 0.012259 | 1 |
| Alg13 | -0.12777 | 0.402798 | 1 |
| Arl2 | -0.12794 | 0.924866 | 1 |
| Nptn | -0.12804 | 0.183863 | 1 |
| Polr2g | -0.12807 | 0.067619 | 1 |
| Taok3 | -0.12819 | 0.32444 | 1 |
| Itsn2 | -0.12819 | 0.122738 | 1 |

| | | | |
|---|---|---|---|
| Gnpda1 | -0.12826 | 0.056312 | 1 |
| Copg1 | -0.12827 | 0.648497 | 1 |
| Snx13 | -0.12853 | 0.123158 | 1 |
| Rbx1 | -0.1287 | 0.051297 | 1 |
| Ints6 | -0.12881 | 0.118488 | 1 |
| Arap1 | -0.12884 | 0.010596 | 1 |
| Mta2 | -0.12895 | 0.435502 | 1 |
| Orc6 | -0.12896 | 0.056737 | 1 |
| Diaph2 | -0.12902 | 0.744222 | 1 |
| Accs | -0.12932 | 0.003708 | 1 |
| Supt3 | -0.12932 | 0.434759 | 1 |
| Bbip1 | -0.1294 | 0.019893 | 1 |
| Pnn | -0.12965 | 0.112164 | 1 |
| Rreb1 | -0.12967 | 0.323421 | 1 |
| Armc4 | -0.12968 | 2.7E-21 | 8.72E-17 |
| Slc43a3 | -0.12989 | 0.019737 | 1 |
| Pak1ip1 | -0.12993 | 0.303686 | 1 |
| Ap1s3 | -0.12994 | 0.036364 | 1 |
| Hectd3 | -0.13002 | 0.009008 | 1 |
| Pdcd2l | -0.13005 | 0.237209 | 1 |
| Sgk1 | -0.13008 | 0.090412 | 1 |
| Mrpl10 | -0.13011 | 0.076586 | 1 |
| Ndufa11 | -0.13023 | 0.210229 | 1 |
| Adam10 | -0.13029 | 0.254225 | 1 |
| Rnf141 | -0.13037 | 0.001534 | 1 |
| Clasp2 | -0.13038 | 0.006978 | 1 |
| Cpne1 | -0.13045 | 0.105874 | 1 |
| Dhrs7 | -0.13047 | 0.09582 | 1 |
| Mospd3 | -0.1305 | 0.028129 | 1 |
| Sae1 | -0.13052 | 0.137519 | 1 |
| Eif4e2 | -0.13055 | 0.63114 | 1 |
| Lmbrd2 | -0.13061 | 0.02419 | 1 |
| Rab3gap2 | -0.13066 | 0.357663 | 1 |
| Ska3 | -0.1307 | 0.003734 | 1 |
| Dap3 | -0.13083 | 0.070755 | 1 |
| Fam98b | -0.13087 | 0.060178 | 1 |
| Zgrf1 | -0.13091 | 0.011713 | 1 |
| Taf1b | -0.13096 | 0.036118 | 1 |
| Itga2 | -0.13104 | 0.002315 | 1 |
| Akt1 | -0.13111 | 0.061097 | 1 |
| Cep70 | -0.13124 | 0.023513 | 1 |
| Nf1 | -0.13133 | 0.185494 | 1 |
| Dpp3 | -0.1314 | 0.020247 | 1 |
| Degs1 | -0.13142 | 0.098151 | 1 |
| Rere | -0.1315 | 0.294803 | 1 |
| Blm | -0.13157 | 0.116733 | 1 |
| Prkaa1 | -0.13169 | 0.009742 | 1 |
| Bscl2 | -0.13178 | 0.011365 | 1 |

| | | | |
|---|---|---|---|
| Polr2a | -0.13188 | 0.545346 | 1 |
| Cap1 | -0.13206 | 0.018519 | 1 |
| Stxbp2 | -0.13223 | 0.407988 | 1 |
| Lrp6 | -0.13239 | 0.569408 | 1 |
| Cenpl | -0.1324 | 0.002693 | 1 |
| Plekhg3 | -0.13241 | 0.664521 | 1 |
| Zfp212 | -0.13241 | 0.106529 | 1 |
| Samsn1 | -0.1326 | 0.200062 | 1 |
| Cdk14 | -0.1329 | 1.36E-06 | 0.043883 |
| Steap3 | -0.13295 | 9.24E-16 | 2.98E-11 |
| Nrros | -0.13295 | 0.099376 | 1 |
| Chpt1 | -0.13301 | 0.041156 | 1 |
| Fbxl2 | -0.13302 | 0.000107 | 1 |
| Vamp7 | -0.13313 | 0.162814 | 1 |
| Rbm43 | -0.13325 | 0.070054 | 1 |
| Tmco1 | -0.1334 | 0.092681 | 1 |
| Cmtm6 | -0.13342 | 0.123651 | 1 |
| Dpm3 | -0.13346 | 0.290637 | 1 |
| Dynlt1a | -0.13346 | 0.008715 | 1 |
| Arl4c | -0.13354 | 0.624785 | 1 |
| Mpnd | -0.13358 | 0.069159 | 1 |
| Ccne1 | -0.13366 | 0.036767 | 1 |
| Wiz | -0.13369 | 0.043539 | 1 |
| Crmp1 | -0.13376 | 0.030899 | 1 |
| Plekhg2 | -0.13416 | 0.012263 | 1 |
| Scfd1 | -0.13424 | 0.42483 | 1 |
| Nelfe | -0.13441 | 0.2947 | 1 |
| Cep55 | -0.13443 | 0.033908 | 1 |
| Cdk11b | -0.13451 | 0.146878 | 1 |
| Chp1 | -0.13455 | 0.049447 | 1 |
| Ccng1 | -0.13476 | 0.183866 | 1 |
| Klhl28 | -0.13483 | 0.074794 | 1 |
| Recql | -0.1349 | 0.174975 | 1 |
| Wdyhv1 | -0.13492 | 0.044477 | 1 |
| Ddx55 | -0.13519 | 0.014889 | 1 |
| Errfi1 | -0.1352 | 0.209572 | 1 |
| Becn1 | -0.13524 | 0.078305 | 1 |
| Zfp180 | -0.1353 | 0.028735 | 1 |
| Dipk1a | -0.13555 | 0.010313 | 1 |
| Gemin7 | -0.13572 | 0.040062 | 1 |
| Mapre1 | -0.13573 | 0.2664 | 1 |
| Stk11 | -0.13578 | 0.072121 | 1 |
| P2ry10 | -0.13585 | 0.0629 | 1 |
| Ptbp2 | -0.13586 | 0.808273 | 1 |
| Cpt1a | -0.13587 | 0.00428 | 1 |
| Tdp2 | -0.13601 | 0.005365 | 1 |
| Rad23a | -0.13613 | 0.125087 | 1 |
| Lsm10 | -0.13626 | 0.811051 | 1 |

| | | | |
|---|---|---|---|
| Dse | -0.13637 | 0.253471 | 1 |
| Nudt1 | -0.13655 | 0.017417 | 1 |
| D1Ertd622e | -0.13657 | 0.007194 | 1 |
| Rbl1 | -0.13671 | 0.03937 | 1 |
| Mapk3 | -0.13677 | 0.008966 | 1 |
| Mtmr12 | -0.13678 | 0.008071 | 1 |
| Cops9 | -0.13704 | 0.072515 | 1 |
| Sgpl1 | -0.13707 | 0.058182 | 1 |
| A630001G21 | -0.13709 | 0.277913 | 1 |
| Hcls1 | -0.13716 | 0.355311 | 1 |
| Nans | -0.13717 | 0.140006 | 1 |
| Mylip | -0.13726 | 0.57904 | 1 |
| Upf2 | -0.13734 | 0.048853 | 1 |
| Strbp | -0.13738 | 0.070858 | 1 |
| Taf2 | -0.13739 | 0.014725 | 1 |
| Chkb | -0.13743 | 0.326294 | 1 |
| Ptbp1 | -0.13751 | 0.129808 | 1 |
| Tsta3 | -0.13758 | 0.181501 | 1 |
| Pdzd8 | -0.13762 | 0.027967 | 1 |
| Hmox2 | -0.13768 | 0.320485 | 1 |
| E2f7 | -0.13768 | 0.031856 | 1 |
| Atm | -0.1377 | 0.047976 | 1 |
| Espl1 | -0.13778 | 0.000376 | 1 |
| Cmc2 | -0.13779 | 0.160279 | 1 |
| Erg28 | -0.1378 | 0.038968 | 1 |
| Tnpo1 | -0.13803 | 0.836829 | 1 |
| Ap5b1 | -0.13813 | 0.092859 | 1 |
| 3110056K07I | -0.13815 | 0.60849 | 1 |
| Smarcd2 | -0.1382 | 0.074912 | 1 |
| Pnrc1 | -0.13822 | 0.563456 | 1 |
| Trmt1l | -0.13823 | 0.212135 | 1 |
| Rdx | -0.1383 | 0.024746 | 1 |
| Gripap1 | -0.13836 | 0.22393 | 1 |
| Camta1 | -0.1384 | 0.015232 | 1 |
| Slc35a4 | -0.13858 | 0.01336 | 1 |
| Ptprc | -0.13864 | 0.01387 | 1 |
| Tapbpl | -0.13873 | 0.100179 | 1 |
| Wdr36 | -0.13875 | 0.052216 | 1 |
| Cdc45 | -0.13881 | 0.000443 | 1 |
| Anapc4 | -0.13884 | 0.019785 | 1 |
| Ermp1 | -0.13887 | 0.187015 | 1 |
| Naa15 | -0.13888 | 0.066244 | 1 |
| Fam98c | -0.1389 | 0.004192 | 1 |
| Cops7a | -0.13892 | 0.15336 | 1 |
| Lnpk | -0.13901 | 0.028103 | 1 |
| Hnrnpc | -0.13905 | 0.117802 | 1 |
| Cdca4 | -0.13919 | 0.406161 | 1 |
| Kif18a | -0.13924 | 0.042107 | 1 |

| | | | |
|---|---|---|---|
| Lman1 | -0.13943 | 0.495113 | 1 |
| Mr1 | -0.13944 | 0.115652 | 1 |
| Ndrg3 | -0.13948 | 0.207527 | 1 |
| Lsm6 | -0.13956 | 0.132403 | 1 |
| Klhl22 | -0.13956 | 5.24E-06 | 0.169093 |
| Pimreg | -0.13958 | 0.076635 | 1 |
| Ociad1 | -0.13962 | 0.067875 | 1 |
| Katnb1 | -0.13966 | 0.024719 | 1 |
| Gm20721 | -0.13967 | 0.096979 | 1 |
| Hist1h2bg | -0.13974 | 0.004079 | 1 |
| Brix1 | -0.13982 | 0.038817 | 1 |
| Srrt | -0.13993 | 0.031453 | 1 |
| Polr2c | -0.13994 | 0.128923 | 1 |
| Ganab | -0.13997 | 0.131807 | 1 |
| Srsf1 | -0.13999 | 0.102046 | 1 |
| Pdia4 | -0.14026 | 0.362765 | 1 |
| Chaf1b | -0.14028 | 0.224399 | 1 |
| Kbtbd2 | -0.14032 | 0.069536 | 1 |
| Tbx21 | -0.14045 | 0.889815 | 1 |
| Tmf1 | -0.14048 | 0.044643 | 1 |
| Commd10 | -0.14053 | 0.02043 | 1 |
| Cerkl | -0.14053 | 0.009235 | 1 |
| Rnf8 | -0.14059 | 0.201907 | 1 |
| Chst12 | -0.14068 | 0.091223 | 1 |
| Derl2 | -0.14077 | 0.061224 | 1 |
| Kpnb1 | -0.14083 | 0.172844 | 1 |
| Pura | -0.14086 | 0.093654 | 1 |
| Pcif1 | -0.14087 | 0.036203 | 1 |
| Ptprcap | -0.14097 | 0.013781 | 1 |
| 1810026B05I | -0.14098 | 0.247648 | 1 |
| Slf1 | -0.14101 | 0.121414 | 1 |
| Atp7a | -0.14105 | 0.041649 | 1 |
| Cep57 | -0.14105 | 0.057738 | 1 |
| Pts | -0.14107 | 0.632528 | 1 |
| Gm48099 | -0.14122 | 0.102708 | 1 |
| Sept1 | -0.14123 | 0.022316 | 1 |
| Gdi1 | -0.14126 | 0.065655 | 1 |
| Fryl | -0.14133 | 0.162335 | 1 |
| Prkcsh | -0.14137 | 0.182934 | 1 |
| Pop4 | -0.14137 | 0.078012 | 1 |
| Wdr6 | -0.14166 | 0.020251 | 1 |
| Ptpra | -0.14183 | 0.047051 | 1 |
| Hsdl2 | -0.1419 | 0.000582 | 1 |
| Gm47882 | -0.14216 | 0.168586 | 1 |
| Trip13 | -0.14233 | 0.002223 | 1 |
| Nrip1 | -0.14234 | 0.093605 | 1 |
| Cenpn | -0.1426 | 1.59E-05 | 0.512938 |
| Cers5 | -0.14265 | 0.489395 | 1 |

| | | | |
|---|---|---|---|
| Tedc1 | -0.14296 | 0.000811 | 1 |
| Vamp8 | -0.14301 | 0.611518 | 1 |
| Ino80e | -0.14327 | 0.125388 | 1 |
| Naa38 | -0.14334 | 0.279003 | 1 |
| Cdc6 | -0.14342 | 1.05E-06 | 0.033793 |
| Usp34 | -0.14343 | 0.035425 | 1 |
| Cenpc1 | -0.14353 | 0.026781 | 1 |
| Mto1 | -0.14359 | 0.075345 | 1 |
| Cramp1l | -0.14361 | 0.001081 | 1 |
| D10Wsu102e | -0.14369 | 0.568893 | 1 |
| Sarnp | -0.14383 | 0.091487 | 1 |
| AI506816 | -0.14386 | 0.082752 | 1 |
| Gxylt1 | -0.14406 | 0.016863 | 1 |
| Aff1 | -0.1441 | 0.965096 | 1 |
| Nfe2l2 | -0.14433 | 0.026833 | 1 |
| Knop1 | -0.14438 | 0.185427 | 1 |
| Tmem37 | -0.14443 | 0.005454 | 1 |
| Kpna3 | -0.14443 | 0.057388 | 1 |
| Cog4 | -0.14446 | 0.4582 | 1 |
| Tagap | -0.14459 | 0.008394 | 1 |
| Kif3b | -0.14463 | 0.017123 | 1 |
| Htra2 | -0.14471 | 0.481408 | 1 |
| Ero1lb | -0.14471 | 0.916049 | 1 |
| Hmgxb4 | -0.14474 | 0.030762 | 1 |
| Dbnl | -0.14481 | 0.154281 | 1 |
| Atp13a3 | -0.14487 | 0.015249 | 1 |
| Ccnb1 | -0.14489 | 0.518474 | 1 |
| Pdia6 | -0.1449 | 0.567162 | 1 |
| Tln1 | -0.1451 | 0.081807 | 1 |
| Lamtor2 | -0.14512 | 0.094683 | 1 |
| Zfp956 | -0.14513 | 0.022538 | 1 |
| Arl6ip5 | -0.14536 | 0.189082 | 1 |
| Tsc22d2 | -0.14538 | 0.677486 | 1 |
| Ska2 | -0.14547 | 0.019386 | 1 |
| Ndufb2 | -0.14557 | 0.26442 | 1 |
| Zfp36 | -0.14567 | 0.022723 | 1 |
| Pan3 | -0.14571 | 0.07133 | 1 |
| Prps2 | -0.14575 | 0.081094 | 1 |
| Dedd | -0.14593 | 0.3088 | 1 |
| Ap4s1 | -0.14608 | 0.020886 | 1 |
| Ncoa4 | -0.14618 | 0.666057 | 1 |
| Mon2 | -0.14621 | 0.060305 | 1 |
| Ing2 | -0.14639 | 0.003266 | 1 |
| Tmem161b | -0.14641 | 0.05028 | 1 |
| Stoml2 | -0.14643 | 0.031607 | 1 |
| Son | -0.14655 | 0.011339 | 1 |
| Phf20 | -0.14656 | 0.043299 | 1 |
| Abca7 | -0.14669 | 0.030765 | 1 |

| | | | |
|---|---|---|---|
| Nipbl | -0.14674 | 0.056582 | 1 |
| Mcub | -0.14685 | 4.86E-06 | 0.15702 |
| Dut | -0.14693 | 0.273956 | 1 |
| Uba52 | -0.14695 | 0.033165 | 1 |
| Pou2f2 | -0.14696 | 0.67013 | 1 |
| Tmem71 | -0.147 | 0.77366 | 1 |
| Mdm2 | -0.147 | 0.00636 | 1 |
| Smpdl3b | -0.14704 | 0.035901 | 1 |
| Jkamp | -0.14725 | 0.008517 | 1 |
| Gm10130 | -0.14727 | 0.01688 | 1 |
| Mvb12a | -0.14728 | 0.125069 | 1 |
| Gtf2e2 | -0.14729 | 0.118428 | 1 |
| Larp7 | -0.14745 | 0.359149 | 1 |
| Smarcb1 | -0.14746 | 0.052456 | 1 |
| Smim4 | -0.14749 | 0.123497 | 1 |
| Fam83d | -0.1477 | 0.003116 | 1 |
| Cd82 | -0.1479 | 0.142443 | 1 |
| 1810009A15l | -0.14796 | 0.081045 | 1 |
| Kctd20 | -0.14799 | 0.014268 | 1 |
| Ly86 | -0.14804 | 0.139417 | 1 |
| Hsd11b1 | -0.14813 | 0.313872 | 1 |
| Gsn | -0.14815 | 0.596411 | 1 |
| Fnta | -0.14815 | 0.289165 | 1 |
| Ankra2 | -0.14818 | 0.171592 | 1 |
| Cdc37 | -0.14819 | 0.005087 | 1 |
| Rdm1 | -0.14822 | 0.011732 | 1 |
| Flii | -0.14822 | 0.033304 | 1 |
| Adcy3 | -0.14827 | 0.000865 | 1 |
| Pdlim5 | -0.14835 | 0.175951 | 1 |
| H2-DMa | -0.14843 | 0.31706 | 1 |
| Anapc15 | -0.14844 | 0.249056 | 1 |
| Vopp1 | -0.14855 | 0.2935 | 1 |
| Fip1l1 | -0.14866 | 0.055621 | 1 |
| Raf1 | -0.14867 | 0.094603 | 1 |
| Ica1 | -0.14868 | 0.026095 | 1 |
| Aaas | -0.14874 | 0.046753 | 1 |
| Erlec1 | -0.14874 | 0.755399 | 1 |
| Tbc1d10c | -0.1488 | 0.049758 | 1 |
| Nod1 | -0.14901 | 0.008382 | 1 |
| Toe1 | -0.14904 | 0.212944 | 1 |
| Urod | -0.14917 | 0.073638 | 1 |
| Rab11a | -0.1493 | 0.060734 | 1 |
| Cdca5 | -0.14959 | 0.007299 | 1 |
| 4930402H24 | -0.1496 | 0.145836 | 1 |
| Was | -0.14973 | 0.028451 | 1 |
| Csrp1 | -0.14978 | 0.040861 | 1 |
| Rfx3 | -0.14979 | 0.006736 | 1 |
| Gga3 | -0.14989 | 0.010187 | 1 |

| | | | |
|---|---|---|---|
| Tspan32 | -0.1499 | 0.142906 | 1 |
| Snrk | -0.15001 | 0.022355 | 1 |
| Psmd14 | -0.15017 | 0.150627 | 1 |
| Sept7 | -0.15026 | 0.261287 | 1 |
| Usp21 | -0.1503 | 0.013669 | 1 |
| Tle4 | -0.15031 | 0.106575 | 1 |
| D130009I18F | -0.15045 | 0.000601 | 1 |
| Tbck | -0.15058 | 0.016813 | 1 |
| Tatdn1 | -0.15062 | 0.312461 | 1 |
| Tomm6 | -0.15066 | 0.044511 | 1 |
| Timp2 | -0.15076 | 0.998266 | 1 |
| Calr | -0.15081 | 0.300042 | 1 |
| Prpf38a | -0.15087 | 0.019288 | 1 |
| Itga6 | -0.15117 | 0.004405 | 1 |
| Hmgb3 | -0.15117 | 0.015542 | 1 |
| Sorbs1 | -0.15124 | 0.006889 | 1 |
| Opa1 | -0.15132 | 0.006694 | 1 |
| Ipo9 | -0.15139 | 0.037241 | 1 |
| Cdk5rap2 | -0.15165 | 0.208001 | 1 |
| Brca1 | -0.15166 | 0.038335 | 1 |
| Mypopos | -0.15171 | 2.15E-05 | 0.692696 |
| Haus4 | -0.15172 | 0.082268 | 1 |
| Zranb2 | -0.15188 | 0.248332 | 1 |
| Fanci | -0.15195 | 0.001403 | 1 |
| Kdelr2 | -0.15197 | 0.429997 | 1 |
| Atp5mpl | -0.152 | 0.383416 | 1 |
| Dpp9 | -0.15201 | 0.031876 | 1 |
| Prkd2 | -0.15209 | 0.22928 | 1 |
| Stub1 | -0.15213 | 0.137348 | 1 |
| Txndc16 | -0.15217 | 0.00672 | 1 |
| Cblb | -0.15217 | 0.045471 | 1 |
| Lrmda | -0.15257 | 0.005175 | 1 |
| Pxmp4 | -0.15257 | 0.31996 | 1 |
| Atad1 | -0.15269 | 0.113039 | 1 |
| Nup85 | -0.15269 | 0.095154 | 1 |
| Nedd1 | -0.15274 | 0.033069 | 1 |
| Slu7 | -0.15292 | 0.000938 | 1 |
| Rnf168 | -0.15303 | 0.023268 | 1 |
| Cep83 | -0.15324 | 0.020125 | 1 |
| Mmgt2 | -0.15329 | 0.164487 | 1 |
| Oat | -0.15331 | 0.949561 | 1 |
| Selenof | -0.15331 | 0.079466 | 1 |
| Cercam | -0.15342 | 0.000673 | 1 |
| Arhgap19 | -0.15357 | 0.000201 | 1 |
| Necap2 | -0.15358 | 0.137051 | 1 |
| Wdr5 | -0.15374 | 0.199159 | 1 |
| Rapgef1 | -0.15377 | 0.139927 | 1 |
| Ramac | -0.15378 | 0.02468 | 1 |

| | | | |
|---|---|---|---|
| Khsrp | -0.15385 | 0.006293 | 1 |
| Golim4 | -0.15397 | 0.512428 | 1 |
| mt-Cytb | -0.15402 | 0.117003 | 1 |
| Impdh1 | -0.15416 | 0.072441 | 1 |
| Prr14l | -0.15417 | 0.133099 | 1 |
| Hmgcr | -0.15425 | 0.059771 | 1 |
| Tshz1 | -0.15432 | 0.019985 | 1 |
| Tlk1 | -0.15443 | 0.019062 | 1 |
| Atp6v0b | -0.15451 | 0.399862 | 1 |
| Eif2b4 | -0.15482 | 0.012561 | 1 |
| Wdr62 | -0.15486 | 0.000133 | 1 |
| Pds5b | -0.15496 | 0.041072 | 1 |
| Syne1 | -0.15496 | 0.307282 | 1 |
| Zfp652 | -0.15516 | 0.166506 | 1 |
| Tbc1d1 | -0.15543 | 0.001477 | 1 |
| Zfp961 | -0.1555 | 6.21E-05 | 1 |
| Brca2 | -0.15556 | 0.023752 | 1 |
| Arhgap45 | -0.15591 | 0.097864 | 1 |
| Pigp | -0.15596 | 0.052211 | 1 |
| Zc3h18 | -0.15598 | 0.199411 | 1 |
| Capns1 | -0.15608 | 0.029876 | 1 |
| Cnot10 | -0.1561 | 0.107386 | 1 |
| Gpr34 | -0.15628 | 0.163274 | 1 |
| Srprb | -0.1563 | 0.291535 | 1 |
| Lrpap1 | -0.15633 | 0.426281 | 1 |
| Lrrc57 | -0.15635 | 0.076446 | 1 |
| Lsm2 | -0.15655 | 0.24162 | 1 |
| Tbc1d10a | -0.15662 | 0.013087 | 1 |
| Pdik1l | -0.15668 | 0.01682 | 1 |
| Akap8l | -0.15679 | 0.795156 | 1 |
| Stat6 | -0.1568 | 0.47905 | 1 |
| Rp2 | -0.15697 | 0.00185 | 1 |
| Eif4e | -0.15712 | 0.020044 | 1 |
| Rnaset2a | -0.15734 | 0.040718 | 1 |
| Tra2b | -0.15748 | 0.144114 | 1 |
| Spg20 | -0.15754 | 0.773191 | 1 |
| Gm31597 | -0.15759 | 0.003473 | 1 |
| Cuedc2 | -0.15777 | 0.069789 | 1 |
| Sp100 | -0.15787 | 0.068813 | 1 |
| Mkln1 | -0.15793 | 0.055208 | 1 |
| Nup133 | -0.15809 | 0.001842 | 1 |
| Il31ra | -0.15817 | 0.009997 | 1 |
| Elob | -0.15839 | 0.065789 | 1 |
| AI467606 | -0.15851 | 0.067614 | 1 |
| Bag6 | -0.15863 | 0.023793 | 1 |
| Nrbp1 | -0.1588 | 0.185477 | 1 |
| Mindy2 | -0.15911 | 0.34999 | 1 |
| Ap2s1 | -0.15918 | 0.147164 | 1 |

| | | | |
|---|---|---|---|
| Nup88 | -0.15934 | 0.018762 | 1 |
| Ccar1 | -0.15947 | 0.169764 | 1 |
| Ccser2 | -0.15947 | 0.03093 | 1 |
| Dock2 | -0.15947 | 0.04034 | 1 |
| Acad9 | -0.15951 | 0.008235 | 1 |
| Polr2k | -0.15953 | 0.023439 | 1 |
| Ncoa3 | -0.15957 | 0.074225 | 1 |
| Ppm1g | -0.1597 | 0.143715 | 1 |
| Rptor | -0.15987 | 0.033048 | 1 |
| Haus3 | -0.15993 | 0.022466 | 1 |
| Ttc17 | -0.16014 | 0.035327 | 1 |
| Nhlrc2 | -0.16017 | 0.11259 | 1 |
| Snrpc | -0.16025 | 0.027167 | 1 |
| Scfd2 | -0.16032 | 0.166846 | 1 |
| Sfxn1 | -0.16038 | 0.116229 | 1 |
| Slc25a12 | -0.16041 | 0.023443 | 1 |
| Gm26827 | -0.16054 | 0.005439 | 1 |
| Sun1 | -0.16063 | 0.002249 | 1 |
| Bckdk | -0.16081 | 0.345133 | 1 |
| Ncaph2 | -0.16113 | 0.239931 | 1 |
| Hnrnpul1 | -0.16126 | 0.095874 | 1 |
| Ggnbp2 | -0.16133 | 0.046307 | 1 |
| Fgfr1op2 | -0.16148 | 0.011867 | 1 |
| Gtf3c6 | -0.1615 | 0.413883 | 1 |
| Irf2bpl | -0.16164 | 0.374548 | 1 |
| Kmt5a | -0.16165 | 0.075223 | 1 |
| Sult2b1 | -0.16185 | 0.00881 | 1 |
| Pias3 | -0.16186 | 0.157167 | 1 |
| 1500009L16I | -0.16206 | 0.00596 | 1 |
| Mtrex | -0.16216 | 0.016868 | 1 |
| Poldip3 | -0.16221 | 0.088293 | 1 |
| Rhoq | -0.16232 | 0.002703 | 1 |
| Dpysl2 | -0.16245 | 0.021824 | 1 |
| Stim2 | -0.16281 | 0.042106 | 1 |
| Fam149b | -0.16301 | 0.239954 | 1 |
| Trappc1 | -0.16307 | 0.03402 | 1 |
| Nup93 | -0.16322 | 0.013806 | 1 |
| Pde4dip | -0.16351 | 0.080792 | 1 |
| Atf1 | -0.16359 | 0.083901 | 1 |
| Brip1 | -0.16362 | 0.017525 | 1 |
| Ppp2r1a | -0.16362 | 0.016539 | 1 |
| Srbd1 | -0.16364 | 0.012992 | 1 |
| 4930444A19I | -0.16376 | 0.015825 | 1 |
| Kcnk5 | -0.16411 | 0.004397 | 1 |
| Prim2 | -0.1643 | 0.017882 | 1 |
| Uba6 | -0.16443 | 0.018151 | 1 |
| Pgap1 | -0.1645 | 0.273927 | 1 |
| Rgs14 | -0.1645 | 0.08968 | 1 |

| | | | |
|---|---|---|---|
| Lax1 | -0.16454 | 0.2098 | 1 |
| Ndufc1 | -0.1647 | 0.036047 | 1 |
| Psph | -0.16471 | 0.066153 | 1 |
| Senp7 | -0.16476 | 0.068406 | 1 |
| Pdlim7 | -0.16478 | 0.121536 | 1 |
| Pum1 | -0.16485 | 0.011264 | 1 |
| Pa2g4 | -0.16492 | 0.07707 | 1 |
| Gm10131 | -0.16574 | 0.324464 | 1 |
| Agpat5 | -0.16583 | 0.006798 | 1 |
| Actr1a | -0.16597 | 0.073685 | 1 |
| Tmem163 | -0.16602 | 0.210652 | 1 |
| Kif14 | -0.16607 | 0.000449 | 1 |
| Atp11b | -0.16617 | 0.306691 | 1 |
| Nek2 | -0.16629 | 0.002317 | 1 |
| Baz1a | -0.16633 | 0.106713 | 1 |
| Cenpm | -0.16639 | 0.000122 | 1 |
| Pds5a | -0.1665 | 0.015377 | 1 |
| Klc1 | -0.16651 | 0.018614 | 1 |
| Ermard | -0.16666 | 0.012712 | 1 |
| Commd3 | -0.1668 | 0.064354 | 1 |
| Rbm10 | -0.16703 | 0.003413 | 1 |
| Dsn1 | -0.16703 | 1.95E-05 | 0.62922 |
| Atf7ip | -0.16727 | 0.130674 | 1 |
| Ist1 | -0.1673 | 0.013985 | 1 |
| Mrpl4 | -0.1673 | 0.230706 | 1 |
| Fam76b | -0.16749 | 0.01891 | 1 |
| Herc1 | -0.1676 | 0.031979 | 1 |
| Chchd1 | -0.16767 | 0.00619 | 1 |
| Efcab11 | -0.16771 | 0.003016 | 1 |
| Dda1 | -0.16792 | 0.016252 | 1 |
| Lypla2 | -0.16833 | 0.018797 | 1 |
| Edf1 | -0.16835 | 0.029531 | 1 |
| Zpbp | -0.16849 | 0.019912 | 1 |
| Fcho2 | -0.16876 | 0.042917 | 1 |
| Stn1 | -0.16881 | 0.005203 | 1 |
| Cyfip2 | -0.16885 | 0.204491 | 1 |
| Kdelr1 | -0.16886 | 0.027991 | 1 |
| Myo18a | -0.16892 | 0.00712 | 1 |
| Ckap2 | -0.16911 | 0.026447 | 1 |
| Arl6ip6 | -0.16919 | 0.010462 | 1 |
| Spcs3 | -0.16922 | 0.036694 | 1 |
| Lman2 | -0.16927 | 0.258952 | 1 |
| Psenen | -0.16948 | 0.029703 | 1 |
| Qk | -0.16957 | 0.157397 | 1 |
| Plekhj1 | -0.1697 | 0.008722 | 1 |
| Spag5 | -0.16997 | 0.016908 | 1 |
| Elf1 | -0.17015 | 0.02214 | 1 |
| Npc1 | -0.17016 | 0.141957 | 1 |

| | | | |
|---|---|---|---|
| Lrwd1 | -0.17019 | 0.168277 | 1 |
| Ifnar2 | -0.17046 | 0.086768 | 1 |
| Rrbp1 | -0.17052 | 0.399771 | 1 |
| Net1 | -0.17067 | 0.001952 | 1 |
| Abcg3 | -0.17071 | 0.014975 | 1 |
| Klrd1 | -0.17094 | 0.83378 | 1 |
| Vps13c | -0.17095 | 0.021641 | 1 |
| Prkacb | -0.17115 | 0.253225 | 1 |
| Cnp | -0.17119 | 0.715421 | 1 |
| Tcerg1 | -0.17125 | 0.084626 | 1 |
| Sumo2 | -0.17174 | 0.111994 | 1 |
| BC035044 | -0.17177 | 0.000674 | 1 |
| Hpf1 | -0.17183 | 0.276597 | 1 |
| Hes6 | -0.17183 | 0.007433 | 1 |
| Bad | -0.17199 | 0.040798 | 1 |
| Pogk | -0.17208 | 0.00029 | 1 |
| Cdc16 | -0.17236 | 0.010418 | 1 |
| Brip1os | -0.17238 | 0.073886 | 1 |
| Itm2a | -0.17239 | 0.433016 | 1 |
| Cuta | -0.17257 | 0.024365 | 1 |
| Med12 | -0.17263 | 0.103055 | 1 |
| Btf3l4 | -0.17266 | 3.81E-05 | 1 |
| Eif2s1 | -0.1727 | 0.069555 | 1 |
| Hat1 | -0.17282 | 0.141086 | 1 |
| Def6 | -0.17295 | 0.120087 | 1 |
| Supt16 | -0.17307 | 0.15014 | 1 |
| Ncf1 | -0.17314 | 0.655515 | 1 |
| Ik | -0.17322 | 0.126734 | 1 |
| Pabpn1 | -0.17326 | 0.10249 | 1 |
| Hspd1 | -0.17334 | 0.350074 | 1 |
| 1700001O22 | -0.17351 | 1.11E-13 | 3.57E-09 |
| Mast2 | -0.17352 | 0.039363 | 1 |
| Hmgcs1 | -0.17352 | 0.103866 | 1 |
| Rfc4 | -0.17362 | 0.052073 | 1 |
| Ewsr1 | -0.17369 | 0.015315 | 1 |
| Knstrn | -0.17377 | 0.006063 | 1 |
| Phactr4 | -0.1738 | 0.307713 | 1 |
| Arhgap30 | -0.17386 | 0.052011 | 1 |
| Gnai2 | -0.17422 | 0.002371 | 1 |
| Lonp2 | -0.17447 | 0.077734 | 1 |
| Hist1h2be | -0.1746 | 0.063013 | 1 |
| Trim8 | -0.17475 | 0.242291 | 1 |
| Prpf3 | -0.17518 | 0.006349 | 1 |
| Gtf2i | -0.17519 | 0.033397 | 1 |
| Snx3 | -0.1755 | 0.036977 | 1 |
| Psme3 | -0.17564 | 0.024798 | 1 |
| Tle3 | -0.17566 | 0.00884 | 1 |
| Slc29a3 | -0.17603 | 0.601969 | 1 |

| | | | |
|---|---|---|---|
| Aqr | -0.17629 | 0.029548 | 1 |
| Syncrip | -0.17632 | 0.126919 | 1 |
| Stk38 | -0.17647 | 0.024312 | 1 |
| Dpy30 | -0.17662 | 0.206312 | 1 |
| Carhsp1 | -0.1769 | 0.02969 | 1 |
| Abracl | -0.177 | 0.021788 | 1 |
| Cnep1r1 | -0.1771 | 0.027039 | 1 |
| Serp1 | -0.17717 | 0.776485 | 1 |
| Sdhb | -0.1775 | 0.409549 | 1 |
| Top3b | -0.17761 | 0.0129 | 1 |
| Ppp1cb | -0.17766 | 0.008587 | 1 |
| Sf1 | -0.17782 | 0.048937 | 1 |
| Strn | -0.17802 | 0.001547 | 1 |
| Camta2 | -0.17821 | 0.070673 | 1 |
| Cdk2 | -0.17843 | 0.000916 | 1 |
| Sema4a | -0.17847 | 0.056924 | 1 |
| Pon2 | -0.1785 | 0.307643 | 1 |
| Slamf6 | -0.17857 | 0.000218 | 1 |
| Ctdsp1 | -0.17857 | 0.000287 | 1 |
| Phf14 | -0.17884 | 0.055239 | 1 |
| Cox20 | -0.17895 | 0.064994 | 1 |
| Phrf1 | -0.1793 | 0.02745 | 1 |
| Lamtor3 | -0.17948 | 0.008502 | 1 |
| Msn | -0.17975 | 0.002127 | 1 |
| Rnf216 | -0.17991 | 0.001769 | 1 |
| Svil | -0.18 | 0.068908 | 1 |
| Ppp1r15b | -0.18015 | 0.026967 | 1 |
| Pkmyt1 | -0.18023 | 0.00232 | 1 |
| Copb2 | -0.18051 | 0.065202 | 1 |
| Plcl2 | -0.18061 | 0.230928 | 1 |
| Lpcat3 | -0.18062 | 0.221417 | 1 |
| Commd1 | -0.18067 | 0.135989 | 1 |
| Wdhd1 | -0.18084 | 0.012386 | 1 |
| Dync1i2 | -0.18087 | 0.012039 | 1 |
| Prrc2a | -0.18091 | 0.174699 | 1 |
| Kctd10 | -0.18101 | 0.083315 | 1 |
| Cpsf7 | -0.18101 | 0.045276 | 1 |
| Sap18 | -0.1811 | 0.01146 | 1 |
| Ubl3 | -0.1811 | 0.122785 | 1 |
| Ldlrap1 | -0.18113 | 2.57E-06 | 0.082935 |
| Spen | -0.18113 | 0.127632 | 1 |
| Smarcc2 | -0.18118 | 0.164458 | 1 |
| Ap3b1 | -0.18134 | 0.079273 | 1 |
| Eny2 | -0.18153 | 0.743636 | 1 |
| Psmc4 | -0.18194 | 0.018556 | 1 |
| Hjurp | -0.18198 | 0.03992 | 1 |
| Ticrr | -0.18259 | 0.135325 | 1 |
| Mrpl18 | -0.18268 | 0.111519 | 1 |

| | | | |
|---|---|---|---|
| Abcb9 | -0.18269 | 0.022953 | 1 |
| Lfng | -0.1828 | 0.22048 | 1 |
| Eif3l | -0.18292 | 0.048003 | 1 |
| Tsc2 | -0.18313 | 0.000139 | 1 |
| Dido1 | -0.18313 | 0.009446 | 1 |
| Hdhd2 | -0.18338 | 0.037463 | 1 |
| Klrc2 | -0.18355 | 0.2162 | 1 |
| Gm38394 | -0.18378 | 0.003106 | 1 |
| Xrcc4 | -0.18415 | 0.336227 | 1 |
| Adrb1 | -0.18416 | 1.69E-05 | 0.545678 |
| Zfhx3 | -0.18442 | 9.4E-06 | 0.303411 |
| Cyb561a3 | -0.18465 | 0.032208 | 1 |
| Rnpep | -0.18485 | 0.076327 | 1 |
| Uckl1 | -0.18511 | 0.02235 | 1 |
| Ctse | -0.18517 | 0.306787 | 1 |
| Arpc5l | -0.18534 | 0.310353 | 1 |
| Neurl3 | -0.18538 | 0.135376 | 1 |
| Nrf1 | -0.18553 | 0.002947 | 1 |
| Ninj1 | -0.18558 | 0.457502 | 1 |
| Sf3b1 | -0.18561 | 0.001837 | 1 |
| Tbxa2r | -0.18561 | 0.077241 | 1 |
| Nt5c | -0.18582 | 0.199062 | 1 |
| Mea1 | -0.18582 | 0.027122 | 1 |
| Tgfb1 | -0.18628 | 0.01638 | 1 |
| Sat1 | -0.18638 | 0.522571 | 1 |
| Pacsin2 | -0.18665 | 0.078754 | 1 |
| Igfbp7 | -0.18671 | 7.68E-05 | 1 |
| Pde1c | -0.18679 | 1.07E-10 | 3.46E-06 |
| Tbl2 | -0.18679 | 0.001027 | 1 |
| I730030J21R | -0.18697 | 0.026364 | 1 |
| Dynlt1f | -0.18709 | 0.022023 | 1 |
| Ndufb1-ps | -0.18723 | 0.012465 | 1 |
| Prkcd | -0.18725 | 0.177745 | 1 |
| Ska1 | -0.18725 | 0.003043 | 1 |
| Adrm1 | -0.18747 | 0.014011 | 1 |
| Rnaseh2c | -0.18751 | 0.109852 | 1 |
| Zfx | -0.18751 | 0.006436 | 1 |
| Brk1 | -0.18761 | 0.012227 | 1 |
| Rpn2 | -0.18765 | 0.039001 | 1 |
| Ttc3 | -0.18775 | 0.059931 | 1 |
| Safb | -0.18789 | 0.003572 | 1 |
| Cbl | -0.18801 | 0.016757 | 1 |
| Spn | -0.18874 | 0.056583 | 1 |
| Myo1e | -0.18889 | 0.145952 | 1 |
| Hist1h2bh | -0.18908 | 0.000405 | 1 |
| Osbpl8 | -0.1891 | 0.008716 | 1 |
| C2cd5 | -0.18915 | 0.005409 | 1 |
| Phtf2 | -0.18947 | 0.000593 | 1 |

| | | | |
|---|---|---|---|
| Eps15 | -0.18958 | 0.028672 | 1 |
| Filip1l | -0.18962 | 0.238891 | 1 |
| Faf1 | -0.18963 | 0.000751 | 1 |
| Cep350 | -0.19011 | 0.083663 | 1 |
| Tma7 | -0.19018 | 0.022034 | 1 |
| Map4k1 | -0.19045 | 0.027258 | 1 |
| Ppp1ca | -0.19049 | 0.000956 | 1 |
| Ube2g2 | -0.19052 | 0.196115 | 1 |
| Msh2 | -0.19056 | 0.07391 | 1 |
| Etfb | -0.19073 | 0.024185 | 1 |
| Calcrl | -0.19076 | 0.27724 | 1 |
| Hdlbp | -0.19086 | 0.130413 | 1 |
| Sin3a | -0.19111 | 0.023844 | 1 |
| Bub1 | -0.19112 | 0.000243 | 1 |
| Prr5 | -0.19144 | 0.553406 | 1 |
| Osbpl9 | -0.19144 | 0.012151 | 1 |
| Fam89b | -0.19152 | 0.091926 | 1 |
| Dgka | -0.19153 | 0.057249 | 1 |
| Snx2 | -0.19157 | 0.916447 | 1 |
| Nde1 | -0.19166 | 0.002295 | 1 |
| Adam17 | -0.19183 | 0.010555 | 1 |
| Ifnar1 | -0.19211 | 0.026893 | 1 |
| Dnaaf5 | -0.19248 | 0.000222 | 1 |
| Eif3a | -0.19249 | 0.053072 | 1 |
| Timm17b | -0.19252 | 0.082194 | 1 |
| Man1c1 | -0.19258 | 1.13E-10 | 3.65E-06 |
| Acot8 | -0.1926 | 0.001501 | 1 |
| Ano6 | -0.19268 | 0.02246 | 1 |
| Hnrnpul2 | -0.19285 | 0.016589 | 1 |
| Frs2 | -0.19286 | 0.054759 | 1 |
| Ppp1r18 | -0.19299 | 0.024687 | 1 |
| Ddx17 | -0.19302 | 0.028421 | 1 |
| Bsg | -0.1931 | 0.902873 | 1 |
| Spred2 | -0.19328 | 0.071432 | 1 |
| Srek1 | -0.1933 | 0.017387 | 1 |
| Fcho1 | -0.19352 | 0.019792 | 1 |
| Ddx43 | -0.19355 | 2.74E-13 | 8.84E-09 |
| Rhog | -0.1937 | 0.05405 | 1 |
| Hook3 | -0.19373 | 0.023529 | 1 |
| Srgap2 | -0.19414 | 0.362162 | 1 |
| Atp5b | -0.19451 | 0.005714 | 1 |
| Exoc2 | -0.19456 | 0.023244 | 1 |
| Themis2 | -0.19465 | 0.006102 | 1 |
| Plekha1 | -0.1948 | 0.000165 | 1 |
| Eif4g1 | -0.19499 | 0.030108 | 1 |
| Prr11 | -0.19499 | 0.003133 | 1 |
| Laptm4a | -0.19535 | 0.451104 | 1 |
| Ssr2 | -0.19535 | 0.035523 | 1 |

| | | | |
|---|---|---|---|
| Eri1 | -0.19541 | 0.013913 | 1 |
| Atp6v1c1 | -0.19564 | 0.012037 | 1 |
| Sft2d1 | -0.19592 | 0.016003 | 1 |
| Rexo1 | -0.19608 | 0.010293 | 1 |
| Pmm1 | -0.19643 | 0.07156 | 1 |
| Prelid3b | -0.1967 | 0.08211 | 1 |
| Dapk2 | -0.19676 | 0.010555 | 1 |
| Fam13b | -0.1971 | 0.004556 | 1 |
| Rab3a | -0.19718 | 0.004066 | 1 |
| Ap2b1 | -0.19771 | 0.033826 | 1 |
| Pdlim1 | -0.19803 | 0.011781 | 1 |
| Ap2m1 | -0.19803 | 0.075408 | 1 |
| Cc2d1a | -0.19822 | 0.003585 | 1 |
| Heatr6 | -0.1985 | 0.077619 | 1 |
| Dnajc7 | -0.1985 | 0.8015 | 1 |
| Meaf6 | -0.19865 | 0.002685 | 1 |
| Tmtc2 | -0.19867 | 0.001275 | 1 |
| Klf12 | -0.19872 | 1.55E-05 | 0.49974 |
| Vps29 | -0.19875 | 0.040376 | 1 |
| Babam1 | -0.19878 | 0.009735 | 1 |
| Rbl2 | -0.19879 | 0.025684 | 1 |
| Cs | -0.19884 | 0.132889 | 1 |
| Selplg | -0.19888 | 0.042982 | 1 |
| Bzw1 | -0.19888 | 0.004774 | 1 |
| Ndc80 | -0.19947 | 0.032204 | 1 |
| Ncapg2 | -0.19979 | 0.008814 | 1 |
| Gucd1 | -0.1999 | 0.000354 | 1 |
| Kmt2e | -0.20014 | 0.020615 | 1 |
| Eps15l1 | -0.20066 | 0.157216 | 1 |
| Ddb1 | -0.20074 | 0.019798 | 1 |
| Dock5 | -0.20092 | 0.010698 | 1 |
| Zmiz2 | -0.20095 | 0.018767 | 1 |
| Pias1 | -0.20098 | 0.129636 | 1 |
| Trub2 | -0.20141 | 0.001254 | 1 |
| Pigc | -0.2015 | 0.021892 | 1 |
| Actn1 | -0.20162 | 0.000806 | 1 |
| B3gnt5 | -0.20184 | 0.012974 | 1 |
| Ppm1j | -0.20213 | 0.014549 | 1 |
| Dok2 | -0.20263 | 0.206254 | 1 |
| Tnks2 | -0.20269 | 0.039815 | 1 |
| Smarcc1 | -0.20277 | 0.026788 | 1 |
| Gm36738 | -0.20376 | 0.148872 | 1 |
| Gnb1 | -0.20386 | 0.141459 | 1 |
| Golga4 | -0.20398 | 0.003064 | 1 |
| Park7 | -0.20431 | 0.015697 | 1 |
| Cct8 | -0.20433 | 0.076853 | 1 |
| Prkab1 | -0.20442 | 0.039158 | 1 |
| Ugcg | -0.20457 | 0.005039 | 1 |

| | | | |
|---|---|---|---|
| Ssb | -0.2047 | 0.013286 | 1 |
| Dbf4 | -0.20475 | 0.137961 | 1 |
| Cdt1 | -0.20485 | 0.0021 | 1 |
| Ahcy | -0.2049 | 0.222159 | 1 |
| AB124611 | -0.20526 | 0.006756 | 1 |
| Sgo1 | -0.20539 | 0.000367 | 1 |
| Rcc1 | -0.20564 | 0.011646 | 1 |
| Mrps21 | -0.2061 | 0.014748 | 1 |
| Numa1 | -0.20641 | 0.030495 | 1 |
| Upk1a | -0.20648 | 1.66E-05 | 0.53462 |
| Mid1ip1 | -0.20655 | 0.125817 | 1 |
| Parpbp | -0.2068 | 0.105482 | 1 |
| Cdkn2aipnl | -0.20691 | 0.005368 | 1 |
| Zcrb1 | -0.20699 | 0.453061 | 1 |
| Hnrnpl | -0.20714 | 0.008225 | 1 |
| Ncbp1 | -0.20717 | 0.063027 | 1 |
| Rock2 | -0.20728 | 0.004127 | 1 |
| Gm37401 | -0.20739 | 0.000381 | 1 |
| Slc44a2 | -0.20742 | 0.010547 | 1 |
| Napsa | -0.2078 | 0.697295 | 1 |
| Ncstn | -0.20792 | 0.115881 | 1 |
| Hist1h2ap | -0.20812 | 0.000272 | 1 |
| Dock11 | -0.20819 | 0.014634 | 1 |
| Srsf7 | -0.20824 | 0.068522 | 1 |
| Tbca | -0.20852 | 0.009428 | 1 |
| Kctd18 | -0.20857 | 0.024829 | 1 |
| Nck1 | -0.20868 | 0.016298 | 1 |
| Psmb6 | -0.20868 | 0.008235 | 1 |
| Canx | -0.20905 | 0.230625 | 1 |
| Sf3b2 | -0.20927 | 0.010039 | 1 |
| Map3k3 | -0.20954 | 0.130827 | 1 |
| Capza1 | -0.20973 | 0.034287 | 1 |
| Shc1 | -0.2098 | 0.061558 | 1 |
| Gapvd1 | -0.21017 | 0.046245 | 1 |
| Cd38 | -0.21019 | 0.00055 | 1 |
| Rps27l | -0.21026 | 0.154467 | 1 |
| Casp3 | -0.21055 | 0.034546 | 1 |
| Pfdn6 | -0.21071 | 0.02257 | 1 |
| Sppl2a | -0.21144 | 0.039492 | 1 |
| Arpc4 | -0.2116 | 0.000805 | 1 |
| Psen2 | -0.21198 | 0.071428 | 1 |
| Nup62 | -0.212 | 0.008501 | 1 |
| Ppp2r1b | -0.21219 | 0.007851 | 1 |
| Dtl | -0.21222 | 0.007371 | 1 |
| Ndufa3 | -0.21273 | 0.022657 | 1 |
| Anapc11 | -0.21286 | 0.006886 | 1 |
| Polr3c | -0.21287 | 0.550373 | 1 |
| Kif20a | -0.21299 | 0.002256 | 1 |

| | | | |
|---|---|---|---|
| Tmpo | -0.21315 | 0.237426 | 1 |
| Dusp22 | -0.21317 | 0.312472 | 1 |
| Rexo5 | -0.21359 | 4.71E-05 | 1 |
| Stag2 | -0.2136 | 0.020721 | 1 |
| Tinf2 | -0.21363 | 0.000373 | 1 |
| Pgm2 | -0.21368 | 0.018744 | 1 |
| 1700037C18I | -0.21407 | 0.103672 | 1 |
| Angel2 | -0.21409 | 0.155621 | 1 |
| Ppil1 | -0.2141 | 0.064589 | 1 |
| Klf10 | -0.21425 | 0.000748 | 1 |
| Ak3 | -0.21434 | 0.001229 | 1 |
| Myadm | -0.21454 | 7.78E-09 | 0.000251 |
| Trim37 | -0.2147 | 0.010325 | 1 |
| Plk4 | -0.21493 | 0.006027 | 1 |
| Kifc1 | -0.21518 | 0.107412 | 1 |
| Arhgap4 | -0.21529 | 0.070525 | 1 |
| Atad5 | -0.2155 | 0.0027 | 1 |
| Nsd2 | -0.21553 | 0.014337 | 1 |
| Fzr1 | -0.216 | 0.176775 | 1 |
| Stx7 | -0.21602 | 0.007665 | 1 |
| Il10rb | -0.21616 | 0.00297 | 1 |
| Stil | -0.21647 | 1.84E-06 | 0.059302 |
| Otulinl | -0.2166 | 0.005711 | 1 |
| Xrn1 | -0.21662 | 0.023966 | 1 |
| Hsd17b4 | -0.21688 | 0.124264 | 1 |
| Dlgap5 | -0.21694 | 0.006389 | 1 |
| Wdr76 | -0.21698 | 0.00021 | 1 |
| Dtymk | -0.21719 | 0.015175 | 1 |
| Map3k11 | -0.21737 | 0.000393 | 1 |
| Myb | -0.2179 | 0.004231 | 1 |
| Fxn | -0.2179 | 0.008015 | 1 |
| Nln | -0.21803 | 0.229358 | 1 |
| Cbx5 | -0.21833 | 0.028677 | 1 |
| Ppia | -0.21846 | 0.00347 | 1 |
| Myl12a | -0.21872 | 0.003137 | 1 |
| Fuca1 | -0.21947 | 0.137108 | 1 |
| Pfdn1 | -0.2196 | 0.020292 | 1 |
| Dynll1 | -0.21964 | 0.146493 | 1 |
| Map4 | -0.21969 | 0.005376 | 1 |
| Zbtb20 | -0.21977 | 0.720709 | 1 |
| Pdap1 | -0.22 | 0.058239 | 1 |
| Cdk4 | -0.22002 | 0.083479 | 1 |
| Psap | -0.2202 | 0.005429 | 1 |
| Xiap | -0.22042 | 0.002839 | 1 |
| Cwc15 | -0.22061 | 0.07599 | 1 |
| Atg4d | -0.22081 | 0.029264 | 1 |
| 1500004A13I | -0.22091 | 0.004386 | 1 |
| Calm3 | -0.22103 | 0.020183 | 1 |

| | | | |
|---|---|---|---|
| Tfeb | -0.22124 | 0.227124 | 1 |
| Rpa2 | -0.22151 | 0.100794 | 1 |
| Dhfr | -0.22178 | 0.002375 | 1 |
| Gramd4 | -0.22196 | 4.72E-05 | 1 |
| Esd | -0.22199 | 0.00993 | 1 |
| Ascc2 | -0.22211 | 0.006495 | 1 |
| Myo5a | -0.22215 | 0.043615 | 1 |
| Fchsd2 | -0.22225 | 0.580755 | 1 |
| Epb41l2 | -0.22235 | 0.181378 | 1 |
| Slamf7 | -0.22238 | 0.042412 | 1 |
| Rab11b | -0.22247 | 0.053515 | 1 |
| Cse1l | -0.22261 | 0.240716 | 1 |
| Psmb7 | -0.22278 | 0.223398 | 1 |
| Cenph | -0.22288 | 9.79E-06 | 0.316188 |
| Cpm | -0.22299 | 0.000563 | 1 |
| Ssna1 | -0.22306 | 0.027844 | 1 |
| Fads1 | -0.22357 | 3.19E-06 | 0.102835 |
| Serinc3 | -0.22399 | 0.381135 | 1 |
| Fen1 | -0.22402 | 0.062136 | 1 |
| Tmcc3 | -0.2241 | 0.001786 | 1 |
| E2f8 | -0.22424 | 0.006024 | 1 |
| Por | -0.22428 | 0.732876 | 1 |
| Alms1 | -0.2243 | 1.72E-05 | 0.554358 |
| Rbmx | -0.22495 | 0.092556 | 1 |
| Tgfbr1 | -0.22511 | 0.043803 | 1 |
| Rad51ap1 | -0.22588 | 7.46E-06 | 0.240936 |
| Kif2c | -0.22604 | 2.17E-05 | 0.70135 |
| Rtn4rl1 | -0.22631 | 0.000627 | 1 |
| Pfdn2 | -0.22637 | 0.007104 | 1 |
| Slc35b1 | -0.22649 | 0.102945 | 1 |
| Borcs7 | -0.2267 | 0.000684 | 1 |
| Virma | -0.22685 | 0.001232 | 1 |
| Cox5b | -0.22686 | 0.026009 | 1 |
| Ets1 | -0.22736 | 0.004183 | 1 |
| Irf2 | -0.22738 | 0.032394 | 1 |
| Ttc7 | -0.2277 | 0.000796 | 1 |
| B020010K11l | -0.22792 | 0.000315 | 1 |
| Rap1a | -0.22796 | 0.00677 | 1 |
| Atp5k | -0.22809 | 0.004341 | 1 |
| Rbm22 | -0.22823 | 0.039857 | 1 |
| Zcchc18 | -0.22827 | 3.82E-09 | 0.000123 |
| Glud1 | -0.2283 | 0.004961 | 1 |
| Arhgap1 | -0.22844 | 9.5E-05 | 1 |
| Cers2 | -0.22851 | 0.016308 | 1 |
| Dbi | -0.22853 | 0.003323 | 1 |
| Reep4 | -0.22859 | 0.001353 | 1 |
| Bub1b | -0.22872 | 0.003175 | 1 |
| Dhx32 | -0.22927 | 0.000577 | 1 |

| | | | |
|---|---|---|---|
| Nr4a2 | -0.2294 | 0.302245 | 1 |
| Ccndbp1 | -0.22953 | 0.00948 | 1 |
| Hnrnpd | -0.22955 | 0.004371 | 1 |
| Lta4h | -0.2299 | 0.07783 | 1 |
| Rif1 | -0.23012 | 0.097213 | 1 |
| Fgr | -0.23052 | 8.1E-08 | 0.002615 |
| Ube2d2a | -0.23071 | 0.000616 | 1 |
| Nudt21 | -0.23094 | 0.005422 | 1 |
| Dync1li1 | -0.23105 | 0.001035 | 1 |
| Dnm2 | -0.23111 | 0.006812 | 1 |
| Selenoh | -0.23129 | 0.086981 | 1 |
| Slc9a3r1 | -0.23141 | 0.005933 | 1 |
| Nab2 | -0.23172 | 0.066714 | 1 |
| Slc39a10 | -0.23218 | 0.000223 | 1 |
| Cnot8 | -0.23233 | 0.052737 | 1 |
| Vps35l | -0.23236 | 0.038483 | 1 |
| Ap1s2 | -0.23248 | 0.000928 | 1 |
| Tspan13 | -0.23308 | 0.391561 | 1 |
| Hdgf | -0.23346 | 0.158689 | 1 |
| Hist1h2bn | -0.23348 | 0.000779 | 1 |
| Arid1a | -0.23351 | 0.004803 | 1 |
| Nmral1 | -0.23369 | 2.95E-05 | 0.950856 |
| Ube2t | -0.23388 | 0.000778 | 1 |
| Glyr1 | -0.23392 | 0.025589 | 1 |
| Ywhaz | -0.23463 | 0.000368 | 1 |
| Ube2a | -0.23478 | 0.003816 | 1 |
| Nono | -0.23489 | 0.007095 | 1 |
| Hdac7 | -0.23503 | 0.000538 | 1 |
| Sigmar1 | -0.23531 | 0.020548 | 1 |
| Lpcat4 | -0.23536 | 0.000749 | 1 |
| Bub3 | -0.23579 | 0.469927 | 1 |
| Actn4 | -0.23638 | 0.023157 | 1 |
| C1qtnf6 | -0.23645 | 0.001787 | 1 |
| Coro1a | -0.23663 | 3.71E-05 | 1 |
| Haao | -0.23679 | 0.001418 | 1 |
| Usp19 | -0.23709 | 0.022178 | 1 |
| Haspin | -0.23734 | 0.000346 | 1 |
| Laptm5 | -0.23735 | 0.000486 | 1 |
| Ccdc69 | -0.2375 | 0.001649 | 1 |
| Runx3 | -0.23761 | 0.034583 | 1 |
| Gse1 | -0.23769 | 0.003601 | 1 |
| Snx14 | -0.23776 | 0.001716 | 1 |
| Dlst | -0.23791 | 0.009781 | 1 |
| Msl3 | -0.23803 | 0.019322 | 1 |
| Tax1bp3 | -0.23822 | 0.00179 | 1 |
| Srsf3 | -0.23844 | 0.068628 | 1 |
| Mbd6 | -0.23876 | 0.393456 | 1 |
| Prpf4b | -0.23877 | 0.0057 | 1 |

| | | | |
|---|---|---|---|
| Eno3 | -0.23889 | 0.000577 | 1 |
| Suv39h1 | -0.23932 | 0.000982 | 1 |
| Afmid | -0.23949 | 0.003375 | 1 |
| Sdf2l1 | -0.23968 | 0.107795 | 1 |
| 5830432E09I | -0.23969 | 1.2E-06 | 0.038898 |
| Nap1l1 | -0.23978 | 0.01968 | 1 |
| Cdkn2a | -0.23978 | 3.1E-05 | 1 |
| Hist1h3g | -0.23991 | 2.1E-05 | 0.679424 |
| Agfg2 | -0.24017 | 0.000335 | 1 |
| Tmem131l | -0.24056 | 0.008284 | 1 |
| Hist1h3f | -0.24122 | 0.000951 | 1 |
| Pnkd | -0.24127 | 0.002509 | 1 |
| Med10 | -0.24152 | 0.077682 | 1 |
| Pbk | -0.24173 | 0.0001 | 1 |
| Magt1 | -0.24174 | 0.002096 | 1 |
| Hist1h2bj | -0.24175 | 1.04E-06 | 0.033731 |
| Pja2 | -0.2418 | 0.002498 | 1 |
| Mastl | -0.24208 | 0.000171 | 1 |
| Pttg1 | -0.24227 | 0.012986 | 1 |
| Elf4 | -0.24259 | 0.003465 | 1 |
| Dap | -0.24262 | 0.020117 | 1 |
| Gapdh | -0.24276 | 0.002046 | 1 |
| Pole2 | -0.24292 | 1.36E-07 | 0.00439 |
| Myh9 | -0.24316 | 0.003538 | 1 |
| Stk32c | -0.24328 | 0.000179 | 1 |
| Akt2 | -0.24333 | 0.025059 | 1 |
| Itpr3 | -0.24359 | 0.005947 | 1 |
| Hist1h4h | -0.24372 | 0.000725 | 1 |
| Emc2 | -0.2439 | 0.009889 | 1 |
| Rnf130 | -0.24409 | 5.03E-08 | 0.001625 |
| Rcbtb2 | -0.24445 | 0.002932 | 1 |
| Sh3bp1 | -0.24504 | 0.019954 | 1 |
| Ogdh | -0.24517 | 0.000573 | 1 |
| Odf2 | -0.24612 | 0.000668 | 1 |
| Eif4g2 | -0.24664 | 0.000193 | 1 |
| Ncf2 | -0.24664 | 9.02E-05 | 1 |
| Plxdc1 | -0.24701 | 5.63E-11 | 1.82E-06 |
| Hmgn5 | -0.24719 | 0.000605 | 1 |
| Wdr1 | -0.24743 | 0.000403 | 1 |
| Capn1 | -0.24748 | 0.010021 | 1 |
| Mpc2 | -0.24779 | 0.4567 | 1 |
| Cyba | -0.24804 | 1.44E-05 | 0.463839 |
| Hist1h2an | -0.24805 | 8E-06 | 0.258132 |
| Elmo2 | -0.24817 | 0.000462 | 1 |
| Brd8 | -0.24828 | 0.032829 | 1 |
| Spc25 | -0.24874 | 1.13E-05 | 0.363396 |
| Atp1b3 | -0.24906 | 0.000208 | 1 |
| Dleu2 | -0.24914 | 0.003047 | 1 |

| | | | |
|---|---|---|---|
| Kremen2 | -0.24949 | 0.005262 | 1 |
| Arid3a | -0.24963 | 0.236817 | 1 |
| Banf1 | -0.24996 | 0.119754 | 1 |
| Fignl1 | -0.25039 | 5.89E-06 | 0.190102 |
| Kif20b | -0.25086 | 0.104254 | 1 |
| Unc13d | -0.25122 | 0.008452 | 1 |
| Icam2 | -0.25136 | 5.77E-06 | 0.186374 |
| Cep128 | -0.25147 | 0.003909 | 1 |
| Pdgfb | -0.25205 | 2.69E-09 | 8.69E-05 |
| Slc39a6 | -0.25248 | 0.003512 | 1 |
| Rhoa | -0.25273 | 3.48E-05 | 1 |
| Sipa1 | -0.25355 | 0.005294 | 1 |
| Ubl4a | -0.25362 | 0.017072 | 1 |
| mt-Nd4 | -0.25373 | 0.012022 | 1 |
| Ssrp1 | -0.25411 | 0.005223 | 1 |
| Snrpg | -0.25468 | 0.000532 | 1 |
| Selenop | -0.25491 | 0.809854 | 1 |
| Mxd3 | -0.25512 | 1.95E-06 | 0.062947 |
| Copa | -0.25527 | 0.001268 | 1 |
| Wasf2 | -0.25537 | 0.000613 | 1 |
| Taf15 | -0.25619 | 0.002083 | 1 |
| 1700025G04 | -0.25656 | 4.9E-07 | 0.015826 |
| Tcf4 | -0.25712 | 0.72586 | 1 |
| Flnb | -0.2575 | 2.17E-05 | 0.699692 |
| Cenpp | -0.25761 | 0.019295 | 1 |
| Gnb4 | -0.25767 | 1.03E-08 | 0.000334 |
| Atp5h | -0.2581 | 0.000913 | 1 |
| Slc37a2 | -0.25854 | 0.001411 | 1 |
| Hint1 | -0.25888 | 0.002469 | 1 |
| Cldnd1 | -0.25895 | 0.001813 | 1 |
| Atp2c1 | -0.25916 | 0.000245 | 1 |
| Prkag1 | -0.2592 | 0.002401 | 1 |
| Ube2m | -0.25967 | 0.002424 | 1 |
| Plk1 | -0.25976 | 0.024577 | 1 |
| Gsk3b | -0.26101 | 0.001892 | 1 |
| Zdhhc14 | -0.26122 | 2.03E-08 | 0.000655 |
| Hspa14 | -0.26186 | 8.43E-05 | 1 |
| Tecr | -0.26189 | 0.002715 | 1 |
| Ppp2r5c | -0.26193 | 0.008123 | 1 |
| R3hdm1 | -0.26211 | 0.027972 | 1 |
| Esco2 | -0.26253 | 0.000777 | 1 |
| Ifitm10 | -0.26258 | 0.03285 | 1 |
| Slc35c1 | -0.26264 | 0.000109 | 1 |
| Snrpd1 | -0.26271 | 0.007949 | 1 |
| Ncor1 | -0.26278 | 0.000446 | 1 |
| Memo1 | -0.26344 | 0.003369 | 1 |
| Twf2 | -0.2635 | 0.058837 | 1 |
| Tec | -0.26364 | 0.00116 | 1 |

| | | | |
|---|---|---|---|
| Baz2a | -0.26369 | 0.010397 | 1 |
| Vcl | -0.26376 | 0.000201 | 1 |
| Chit1 | -0.26387 | 4.92E-12 | 1.59E-07 |
| Prim1 | -0.26406 | 0.000325 | 1 |
| Cbx3 | -0.26426 | 0.033723 | 1 |
| Plcb2 | -0.26477 | 0.001749 | 1 |
| Cenpk | -0.26488 | 1.38E-05 | 0.446533 |
| Rbbp8 | -0.26505 | 0.000107 | 1 |
| Serpini1 | -0.26556 | 2.96E-18 | 9.56E-14 |
| Acss1 | -0.26563 | 0.028292 | 1 |
| Slc14a1 | -0.26584 | 0.001561 | 1 |
| Prf1 | -0.26602 | 0.000494 | 1 |
| Mcm7 | -0.26627 | 0.001774 | 1 |
| Atp6v1g1 | -0.26644 | 0.027215 | 1 |
| Pafah1b1 | -0.26644 | 0.001838 | 1 |
| Cdca7l | -0.26688 | 0.014542 | 1 |
| Gpd1l | -0.26691 | 9.2E-05 | 1 |
| Sfpq | -0.26696 | 0.068952 | 1 |
| Vezf1 | -0.26698 | 0.001729 | 1 |
| Ctsc | -0.26815 | 0.000516 | 1 |
| Mcee | -0.26874 | 0.042521 | 1 |
| Slc4a2 | -0.26899 | 0.091497 | 1 |
| Nasp | -0.26963 | 0.005037 | 1 |
| Sbf1 | -0.27006 | 0.001169 | 1 |
| Gfod1 | -0.27008 | 0.000216 | 1 |
| Ywhab | -0.2701 | 0.000498 | 1 |
| Cenpx | -0.27129 | 0.045527 | 1 |
| Tmem258 | -0.2717 | 0.000485 | 1 |
| H2-Oa | -0.27202 | 0.043013 | 1 |
| Bcl2l13 | -0.27208 | 3.22E-05 | 1 |
| Ccng2 | -0.27225 | 0.010554 | 1 |
| Nfu1 | -0.27234 | 0.069677 | 1 |
| St3gal4 | -0.27257 | 0.003869 | 1 |
| 1810037I17F | -0.27271 | 0.009711 | 1 |
| Pole | -0.27335 | 0.000165 | 1 |
| Rad51 | -0.27389 | 0.000452 | 1 |
| Tjp2 | -0.27399 | 2.73E-05 | 0.879949 |
| Rras2 | -0.27412 | 1.98E-06 | 0.063986 |
| Ubash3b | -0.27413 | 0.025105 | 1 |
| Luc7l3 | -0.2757 | 0.000447 | 1 |
| Ccr5 | -0.2761 | 0.321462 | 1 |
| Ctnnb1 | -0.27641 | 0.000214 | 1 |
| Cdkn2c | -0.2766 | 8.77E-05 | 1 |
| Scarb2 | -0.27683 | 0.019766 | 1 |
| Cast | -0.27684 | 0.001448 | 1 |
| Tmsb4x | -0.2769 | 9.83E-08 | 0.003172 |
| Arhgap17 | -0.27727 | 0.004045 | 1 |
| Sgo2a | -0.27733 | 0.000286 | 1 |

| | | | |
|---|---|---|---|
| Hist1h2bc | -0.27795 | 0.088428 | 1 |
| Aip | -0.27796 | 0.012834 | 1 |
| Xrn2 | -0.27824 | 0.000606 | 1 |
| Ap3s1 | -0.27826 | 0.004848 | 1 |
| Camk2n1 | -0.27845 | 0.001479 | 1 |
| Tmod3 | -0.27847 | 0.001566 | 1 |
| Depdc1a | -0.27913 | 0.001119 | 1 |
| Cenpw | -0.27963 | 0.000675 | 1 |
| Ppp1r35 | -0.27971 | 0.000752 | 1 |
| Macf1 | -0.27986 | 0.014255 | 1 |
| Rps6ka1 | -0.27993 | 0.003024 | 1 |
| Alyref | -0.28043 | 0.005326 | 1 |
| Golm1 | -0.28074 | 0.006628 | 1 |
| Cat | -0.28116 | 0.01826 | 1 |
| Rbm34 | -0.28134 | 0.720701 | 1 |
| Dhx9 | -0.28149 | 0.010734 | 1 |
| Ndufs2 | -0.28159 | 8.95E-05 | 1 |
| Pkp4 | -0.28192 | 2.13E-05 | 0.687065 |
| Snx18 | -0.28209 | 3.39E-05 | 1 |
| Suox | -0.28298 | 0.000162 | 1 |
| Hdac1 | -0.28368 | 0.000823 | 1 |
| Ngly1 | -0.28384 | 0.00128 | 1 |
| Sfxn3 | -0.28409 | 8.6E-06 | 0.277656 |
| Top2b | -0.28438 | 0.00061 | 1 |
| Aurka | -0.28579 | 0.000176 | 1 |
| Atp6v1b2 | -0.28584 | 0.000444 | 1 |
| Sike1 | -0.28656 | 0.000945 | 1 |
| Adgrg1 | -0.28666 | 0.002796 | 1 |
| Mmd | -0.28672 | 0.001061 | 1 |
| Ddx39b | -0.28728 | 0.001306 | 1 |
| Cd200r4 | -0.28827 | 0.006502 | 1 |
| Cnn2 | -0.28858 | 0.000485 | 1 |
| Ccdc88c | -0.28979 | 0.002158 | 1 |
| Rangap1 | -0.28992 | 0.001257 | 1 |
| Ankrd10 | -0.29 | 0.022711 | 1 |
| Smarca4 | -0.2901 | 0.121183 | 1 |
| Cdkn2d | -0.29022 | 0.012259 | 1 |
| Slc41a3 | -0.29058 | 0.016334 | 1 |
| Txnl1 | -0.29117 | 0.000915 | 1 |
| Skiv2l | -0.2913 | 8.89E-05 | 1 |
| Smc1a | -0.29169 | 0.010633 | 1 |
| Dgkd | -0.29176 | 0.00047 | 1 |
| Apobec2 | -0.29178 | 5.51E-18 | 1.78E-13 |
| Card11 | -0.29235 | 0.001347 | 1 |
| Lrmp | -0.2925 | 0.000699 | 1 |
| Anp32e | -0.29289 | 0.275657 | 1 |
| Slc23a2 | -0.293 | 0.003877 | 1 |
| Glipr2 | -0.29305 | 0.00108 | 1 |

| | | | |
|---|---|---|---|
| Coro7 | -0.29344 | 4.93E-05 | 1 |
| Sytl2 | -0.29366 | 0.067775 | 1 |
| Tpp1 | -0.29374 | 0.004927 | 1 |
| Cks2 | -0.29381 | 0.094947 | 1 |
| Taldo1 | -0.2939 | 0.003628 | 1 |
| Hnrnpr | -0.29397 | 2.81E-05 | 0.906549 |
| Ep300 | -0.29402 | 0.000208 | 1 |
| Acly | -0.29421 | 0.000155 | 1 |
| Lsp1 | -0.29585 | 0.000192 | 1 |
| Shcbp1 | -0.29675 | 0.000414 | 1 |
| Tacc3 | -0.29697 | 0.002448 | 1 |
| Ehbp1l1 | -0.29746 | 0.001557 | 1 |
| S100a11 | -0.29925 | 0.000145 | 1 |
| B3gat3 | -0.29946 | 3.32E-06 | 0.107072 |
| Galnt7 | -0.29965 | 0.000293 | 1 |
| Arhgdib | -0.29979 | 1.25E-06 | 0.040374 |
| Ankrd44 | -0.29989 | 6.54E-05 | 1 |
| Anln | -0.30063 | 0.000142 | 1 |
| S1pr4 | -0.30165 | 0.000602 | 1 |
| Rtn3 | -0.30184 | 0.000367 | 1 |
| Pole4 | -0.30203 | 0.002567 | 1 |
| Nup107 | -0.30235 | 0.001028 | 1 |
| Fubp1 | -0.30268 | 0.001604 | 1 |
| Add1 | -0.30281 | 0.004698 | 1 |
| Gltp | -0.30326 | 0.000496 | 1 |
| Kif15 | -0.30332 | 0.324372 | 1 |
| Atpif1 | -0.30342 | 0.007549 | 1 |
| Nfia | -0.30351 | 0.020372 | 1 |
| Mcu | -0.30365 | 4.89E-05 | 1 |
| Capn2 | -0.30382 | 0.002597 | 1 |
| Adamts10 | -0.30426 | 0.000197 | 1 |
| Mrtfa | -0.3046 | 0.000866 | 1 |
| Tipin | -0.30513 | 0.001147 | 1 |
| Cd27 | -0.30518 | 0.004692 | 1 |
| Ppp4c | -0.30571 | 1.37E-05 | 0.440838 |
| Aplp2 | -0.30571 | 0.000233 | 1 |
| Tnfrsf9 | -0.30601 | 0.000487 | 1 |
| Chchd10 | -0.3064 | 0.229532 | 1 |
| Paqr4 | -0.30671 | 8.69E-06 | 0.280633 |
| Atad2 | -0.30699 | 0.023248 | 1 |
| Sdcbp | -0.30728 | 0.00881 | 1 |
| Gmnn | -0.3083 | 0.009851 | 1 |
| Mad2l1 | -0.30886 | 0.140787 | 1 |
| Hnrnpk | -0.30956 | 9.37E-06 | 0.302421 |
| Man1a | -0.31041 | 0.003213 | 1 |
| Plp2 | -0.31164 | 0.02793 | 1 |
| Hnrnpa3 | -0.31177 | 6.32E-05 | 1 |
| Cfl1 | -0.31177 | 1.45E-07 | 0.004675 |

| | | | |
|---|---|---|---|
| Grk2 | -0.31182 | 0.000625 | 1 |
| Ect2 | -0.31209 | 0.000907 | 1 |
| Ubap2l | -0.31213 | 0.001681 | 1 |
| Arhgef2 | -0.31252 | 0.000203 | 1 |
| Prkcb | -0.31378 | 0.001994 | 1 |
| Tanc2 | -0.3142 | 4.99E-08 | 0.001612 |
| Trf | -0.31481 | 0.00039 | 1 |
| Mdh1 | -0.31531 | 0.000599 | 1 |
| Arhgdia | -0.31565 | 4.17E-06 | 0.134573 |
| Tsen34 | -0.31601 | 0.02303 | 1 |
| Cux1 | -0.31631 | 0.013749 | 1 |
| Pkig | -0.3174 | 0.278429 | 1 |
| Smc6 | -0.31816 | 0.0119 | 1 |
| Plcg2 | -0.31844 | 0.002925 | 1 |
| Fkbp2 | -0.31877 | 0.009024 | 1 |
| Eed | -0.31892 | 0.005329 | 1 |
| Nuf2 | -0.31902 | 0.000699 | 1 |
| Pdcd6ip | -0.31929 | 6.33E-05 | 1 |
| Tpm3 | -0.31953 | 7.91E-06 | 0.255324 |
| Cdca2 | -0.31965 | 0.001767 | 1 |
| Cdc20 | -0.31972 | 0.006976 | 1 |
| Samd3 | -0.31988 | 2.11E-08 | 0.000681 |
| Baz1b | -0.32103 | 0.002789 | 1 |
| Acot7 | -0.32128 | 0.000746 | 1 |
| Ccdc34 | -0.322 | 0.019182 | 1 |
| Sypl | -0.32213 | 6.63E-07 | 0.021413 |
| Cd72 | -0.32245 | 1.34E-06 | 0.043136 |
| H2afv | -0.32251 | 0.15156 | 1 |
| Pycrl | -0.3237 | 0.007456 | 1 |
| Dtx1 | -0.32486 | 1.16E-05 | 0.373561 |
| Arpc5 | -0.32505 | 6.94E-05 | 1 |
| Myo1c | -0.32527 | 0.278154 | 1 |
| Depdc1b | -0.32546 | 4.4E-06 | 0.142034 |
| Kif18b | -0.32599 | 5.44E-05 | 1 |
| Idh3a | -0.3264 | 5.05E-05 | 1 |
| Smc3 | -0.32643 | 0.000122 | 1 |
| Unc93b1 | -0.32663 | 0.103331 | 1 |
| Mirt1 | -0.32687 | 5.54E-07 | 0.01787 |
| Gpx1 | -0.32734 | 0.300566 | 1 |
| Crip2 | -0.32763 | 2.27E-14 | 7.34E-10 |
| Mcm2 | -0.32874 | 0.004039 | 1 |
| Pold1 | -0.32894 | 7.33E-05 | 1 |
| Hnrnpm | -0.32895 | 0.007482 | 1 |
| Slc25a4 | -0.3293 | 0.000181 | 1 |
| Ergic2 | -0.33062 | 0.011273 | 1 |
| Vasp | -0.33166 | 0.00011 | 1 |
| C1qb | -0.33222 | 0.718043 | 1 |
| Galnt2 | -0.33226 | 1.72E-06 | 0.055686 |

| | | | |
|---|---|---|---|
| Chchd2 | -0.33226 | 0.000517 | 1 |
| Atp2b1 | -0.33341 | 0.001179 | 1 |
| Hnrnpu | -0.33379 | 6.6E-05 | 1 |
| Icam1 | -0.33379 | 0.075126 | 1 |
| Rfc1 | -0.33406 | 3.19E-05 | 1 |
| Clspn | -0.33511 | 0.000104 | 1 |
| Hist1h2bm | -0.33525 | 5.76E-05 | 1 |
| Rnaseh2b | -0.33563 | 0.02613 | 1 |
| Rpa1 | -0.33586 | 0.000127 | 1 |
| Rps18 | -0.33599 | 0.131055 | 1 |
| Bax | -0.33687 | 0.002992 | 1 |
| Rpn1 | -0.33767 | 0.005787 | 1 |
| Spp1 | -0.33839 | 0.006486 | 1 |
| Rfc2 | -0.34147 | 0.028317 | 1 |
| Ap1g2 | -0.34383 | 1.02E-06 | 0.032844 |
| Tank | -0.34391 | 0.002189 | 1 |
| Rbm38 | -0.34519 | 0.000124 | 1 |
| Rfwd3 | -0.34562 | 0.000717 | 1 |
| Cers4 | -0.34574 | 0.000113 | 1 |
| Arhgef1 | -0.34635 | 1.38E-06 | 0.044458 |
| Rbbp4 | -0.34658 | 0.000593 | 1 |
| Adamts14 | -0.34697 | 5.32E-07 | 0.017178 |
| Rbbp7 | -0.34837 | 0.005019 | 1 |
| Susd1 | -0.34845 | 3.65E-06 | 0.117976 |
| Ilk | -0.34848 | 2.06E-05 | 0.663762 |
| Myl6 | -0.34861 | 1.23E-08 | 0.000396 |
| Slc25a24 | -0.34894 | 7.15E-07 | 0.023096 |
| Abi1 | -0.34927 | 2.22E-05 | 0.715869 |
| Bex3 | -0.34937 | 0.000637 | 1 |
| Cdc42se1 | -0.34967 | 0.000664 | 1 |
| Slc38a2 | -0.34968 | 0.073571 | 1 |
| Tsc22d1 | -0.35009 | 1.23E-06 | 0.03963 |
| Ralgps2 | -0.35015 | 0.743744 | 1 |
| Sell | -0.3502 | 1.06E-10 | 3.41E-06 |
| Cdk6 | -0.35049 | 0.000703 | 1 |
| Anapc5 | -0.35053 | 0.004751 | 1 |
| Uhrf1 | -0.35066 | 0.000766 | 1 |
| Vps37b | -0.35091 | 6.38E-05 | 1 |
| Hells | -0.35092 | 0.003487 | 1 |
| Stim1 | -0.35103 | 6.12E-05 | 1 |
| Lrrk1 | -0.3517 | 2.87E-05 | 0.926826 |
| Nup210 | -0.35234 | 6.29E-05 | 1 |
| Cenpa | -0.35255 | 0.01231 | 1 |
| Cdkn3 | -0.35344 | 0.003029 | 1 |
| Itgb2 | -0.35441 | 2.83E-06 | 0.091421 |
| Ulbp1 | -0.35521 | 0.001529 | 1 |
| Ikzf3 | -0.35593 | 0.002212 | 1 |
| Mcm4 | -0.35625 | 0.128466 | 1 |

| | | | |
|---|---|---|---|
| Ncapg | -0.35627 | 4.12E-05 | 1 |
| Man2b1 | -0.35685 | 0.00143 | 1 |
| Nr3c1 | -0.3575 | 0.000354 | 1 |
| Chsy1 | -0.35862 | 2.3E-06 | 0.074097 |
| Hnrnpab | -0.35871 | 0.006598 | 1 |
| Setbp1 | -0.36006 | 5.36E-05 | 1 |
| Hnrnpf | -0.36021 | 9.11E-08 | 0.002942 |
| Rnf2 | -0.36068 | 1.61E-06 | 0.051962 |
| Cep85 | -0.36184 | 6.29E-06 | 0.20294 |
| Spc24 | -0.36235 | 0.001583 | 1 |
| Pfn1 | -0.36257 | 9.17E-11 | 2.96E-06 |
| Exoc4 | -0.36289 | 0.005879 | 1 |
| Jak1 | -0.36293 | 0.000134 | 1 |
| Rnaset2b | -0.363 | 0.000111 | 1 |
| Pde6d | -0.36328 | 8.99E-09 | 0.00029 |
| Kif13b | -0.36424 | 0.001113 | 1 |
| Pabpc1 | -0.36428 | 3.2E-08 | 0.001032 |
| Kpna2 | -0.365 | 0.008656 | 1 |
| Kif4 | -0.36543 | 0.001248 | 1 |
| Dcp1b | -0.36562 | 7.66E-05 | 1 |
| Swap70 | -0.36567 | 0.001051 | 1 |
| Gm42418 | -0.36574 | 0.164627 | 1 |
| Aspm | -0.36582 | 6.65E-06 | 0.21467 |
| Adgre5 | -0.36603 | 0.000811 | 1 |
| Ptbp3 | -0.3664 | 3.84E-07 | 0.012407 |
| Map7d1 | -0.36667 | 2.88E-06 | 0.093003 |
| Ncaph | -0.36775 | 3.04E-05 | 0.981988 |
| Mcm6 | -0.36878 | 0.0117 | 1 |
| Abtb2 | -0.36887 | 0.006976 | 1 |
| Ctsl | -0.36901 | 0.368355 | 1 |
| Dmac1 | -0.36957 | 7.59E-05 | 1 |
| Cdca8 | -0.36965 | 0.00069 | 1 |
| Ncapd3 | -0.36993 | 0.008824 | 1 |
| Lclat1 | -0.37139 | 8.8E-05 | 1 |
| Tyms | -0.3722 | 0.00016 | 1 |
| Rhoc | -0.37231 | 8.15E-05 | 1 |
| Hnrnpa2b1 | -0.37248 | 2.24E-07 | 0.007244 |
| Rnf43 | -0.37472 | 4.81E-13 | 1.55E-08 |
| Mtpn | -0.37477 | 2.83E-05 | 0.912862 |
| Hist2h4 | -0.37506 | 0.247018 | 1 |
| Ap1m1 | -0.37519 | 3.12E-06 | 0.100828 |
| Irf8 | -0.37583 | 0.001995 | 1 |
| Reep5 | -0.37699 | 1.12E-07 | 0.003609 |
| Cbx1 | -0.37857 | 0.029052 | 1 |
| Nfic | -0.38028 | 5.39E-06 | 0.174152 |
| Arhgap11a | -0.38073 | 0.003861 | 1 |
| Stag1 | -0.38079 | 5.24E-05 | 1 |
| Ywhah | -0.3818 | 0.000225 | 1 |

| | | | |
|---|---|---|---|
| Tcf19 | -0.38414 | 0.001829 | 1 |
| Tpm4 | -0.38429 | 0.000273 | 1 |
| Iqgap1 | -0.38449 | 2.6E-07 | 0.00838 |
| Hist1h2bb | -0.38472 | 7.16E-06 | 0.231092 |
| Hmmr | -0.3855 | 0.000305 | 1 |
| Mis18bp1 | -0.38555 | 6.7E-05 | 1 |
| Flot2 | -0.38603 | 0.001265 | 1 |
| Rad21 | -0.38615 | 0.000579 | 1 |
| Ccnf | -0.38733 | 2.75E-05 | 0.887783 |
| St8sia4 | -0.38744 | 2.12E-07 | 0.006846 |
| S100a13 | -0.38899 | 2.55E-06 | 0.082198 |
| Lrrfip1 | -0.38918 | 3.11E-06 | 0.100469 |
| Hist1h2ac | -0.38935 | 2.37E-05 | 0.764884 |
| Cd9 | -0.39025 | 0.388703 | 1 |
| Ndfip2 | -0.39204 | 0.000371 | 1 |
| Dnajc9 | -0.39355 | 0.001785 | 1 |
| Ran | -0.39387 | 2.73E-05 | 0.880908 |
| Mplkip | -0.39518 | 0.001083 | 1 |
| Eml3 | -0.39534 | 1.89E-09 | 6.09E-05 |
| Gm11808 | -0.39545 | 2.25E-07 | 0.007266 |
| Hist1h3c | -0.39721 | 0.000124 | 1 |
| Foxp1 | -0.3978 | 0.411851 | 1 |
| Jarid2 | -0.39801 | 1.61E-05 | 0.519859 |
| Cenpe | -0.39948 | 0.018826 | 1 |
| Cmklr1 | -0.39975 | 1.06E-21 | 3.42E-17 |
| Prex1 | -0.40008 | 0.013076 | 1 |
| Azin1 | -0.40155 | 0.00373 | 1 |
| Sp4 | -0.40244 | 7.92E-06 | 0.255548 |
| Suz12 | -0.40331 | 0.000141 | 1 |
| Tk1 | -0.40345 | 0.003791 | 1 |
| Pcna | -0.404 | 0.000879 | 1 |
| Hist2h2ac | -0.40492 | 0.019805 | 1 |
| Uba2 | -0.40497 | 0.000109 | 1 |
| Stk10 | -0.40818 | 3.21E-07 | 0.010373 |
| Nucks1 | -0.41067 | 0.012054 | 1 |
| Morrbid | -0.41225 | 4.99E-07 | 0.016099 |
| Ckap2l | -0.41249 | 0.00064 | 1 |
| Gm19951 | -0.41275 | 1.57E-05 | 0.506601 |
| Kif23 | -0.4148 | 7.33E-05 | 1 |
| Neil3 | -0.4157 | 1.75E-05 | 0.563813 |
| Calm2 | -0.41709 | 0.000114 | 1 |
| Pik3r5 | -0.41715 | 1.15E-09 | 3.7E-05 |
| Perp | -0.41737 | 7.03E-10 | 2.27E-05 |
| Aurkb | -0.41886 | 6.02E-05 | 1 |
| Zbtb32 | -0.41941 | 1.51E-05 | 0.487259 |
| Slc27a4 | -0.4198 | 2.37E-19 | 7.67E-15 |
| Hypk | -0.42065 | 2.27E-05 | 0.733377 |
| Serp2 | -0.42151 | 2.64E-12 | 8.51E-08 |

| | | | |
|---|---|---|---|
| Sub1 | -0.42166 | 2.56E-09 | 8.27E-05 |
| Lrrc8c | -0.42167 | 2.46E-06 | 0.079442 |
| Nt5c3b | -0.4218 | 5.06E-06 | 0.163465 |
| Ezr | -0.42188 | 5.44E-07 | 0.017548 |
| Baiap3 | -0.42319 | 2.39E-09 | 7.72E-05 |
| Hmgb1 | -0.42355 | 4.15E-06 | 0.133966 |
| Gnb2 | -0.42467 | 3.38E-08 | 0.001091 |
| Cd244a | -0.4249 | 1.54E-09 | 4.99E-05 |
| Gab2 | -0.42685 | 0.00048 | 1 |
| Dck | -0.42843 | 1.7E-05 | 0.549024 |
| Cks1b | -0.42857 | 0.003515 | 1 |
| Lgals3 | -0.42871 | 0.003876 | 1 |
| C1qa | -0.42902 | 0.61152 | 1 |
| Jpt1 | -0.43085 | 4.16E-06 | 0.134237 |
| Plec | -0.43091 | 0.001225 | 1 |
| Bin1 | -0.43173 | 2.19E-07 | 0.007064 |
| Hist2h2bb | -0.43203 | 0.016259 | 1 |
| Cdca3 | -0.4334 | 0.000356 | 1 |
| Aff3 | -0.43506 | 0.446372 | 1 |
| Capzb | -0.43583 | 1.32E-13 | 4.26E-09 |
| S100a4 | -0.43602 | 0.000577 | 1 |
| Actg1 | -0.43665 | 2.72E-09 | 8.79E-05 |
| Csk | -0.43717 | 2.73E-08 | 0.000881 |
| Srsf2 | -0.43726 | 0.000584 | 1 |
| St3gal6 | -0.43748 | 1.8E-05 | 0.580487 |
| Hmga1 | -0.43977 | 0.000132 | 1 |
| Gnas | -0.44181 | 0.011164 | 1 |
| Ptma | -0.44215 | 0.082586 | 1 |
| Actr3 | -0.44251 | 2.76E-10 | 8.92E-06 |
| Ccnb2 | -0.44252 | 0.000251 | 1 |
| Ckap5 | -0.44601 | 0.064646 | 1 |
| Esyt1 | -0.4463 | 1.23E-08 | 0.000396 |
| Pola1 | -0.44701 | 0.000533 | 1 |
| Cit | -0.44748 | 1.2E-05 | 0.387413 |
| Apbb1ip | -0.44751 | 1.12E-06 | 0.036151 |
| Mef2a | -0.44808 | 0.000248 | 1 |
| Sptan1 | -0.44839 | 0.000232 | 1 |
| Nkg7 | -0.45129 | 3.37E-09 | 0.000109 |
| S1pr1 | -0.45559 | 4.69E-10 | 1.51E-05 |
| Ddx39 | -0.4563 | 5.02E-05 | 1 |
| Gm45552 | -0.4569 | 2.89E-12 | 9.32E-08 |
| Snx10 | -0.45798 | 2.27E-05 | 0.733561 |
| Gm42047 | -0.45829 | 1.39E-06 | 0.044728 |
| Marcks | -0.45858 | 0.000978 | 1 |
| Klf3 | -0.45906 | 3.41E-13 | 1.1E-08 |
| Hsp90b1 | -0.46173 | 9.28E-05 | 1 |
| Slbp | -0.46275 | 1.87E-06 | 0.060486 |
| Sptbn1 | -0.46367 | 0.000575 | 1 |

| | | | |
|---|---|---|---|
| Erh | -0.46383 | 5.06E-07 | 0.016321 |
| Hmgn1 | -0.46437 | 0.000246 | 1 |
| Glul | -0.46453 | 0.005821 | 1 |
| Pde2a | -0.46654 | 7.5E-18 | 2.42E-13 |
| Ehd4 | -0.46675 | 1.37E-06 | 0.044256 |
| Tagln2 | -0.46953 | 0.001128 | 1 |
| Ikzf2 | -0.47069 | 1.73E-08 | 0.000558 |
| Tceal9 | -0.47156 | 5.11E-07 | 0.016482 |
| Actb | -0.47233 | 2.26E-15 | 7.3E-11 |
| Klra3 | -0.47491 | 0.006759 | 1 |
| Me2 | -0.47585 | 4.32E-10 | 1.39E-05 |
| Il10ra | -0.47809 | 7.69E-09 | 0.000248 |
| Lat2 | -0.47948 | 2.81E-06 | 0.09058 |
| H3f3a | -0.48139 | 0.000183 | 1 |
| Itgam | -0.48163 | 9.92E-09 | 0.00032 |
| Fmnl1 | -0.48171 | 2.31E-08 | 0.000746 |
| Hexb | -0.4821 | 0.000223 | 1 |
| Iqgap2 | -0.48386 | 7.09E-05 | 1 |
| Serpinb6b | -0.48508 | 1.92E-05 | 0.621296 |
| Fam129a | -0.48557 | 0.000135 | 1 |
| Syk | -0.48665 | 1.34E-07 | 0.004311 |
| Lbr | -0.48703 | 5.44E-08 | 0.001757 |
| Arsb | -0.48977 | 8.66E-06 | 0.279588 |
| Dek | -0.49126 | 2.34E-05 | 0.75648 |
| Ezh2 | -0.49301 | 0.002172 | 1 |
| Ctsd | -0.4935 | 5.01E-07 | 0.016182 |
| Arhgef18 | -0.49381 | 4.01E-06 | 0.129495 |
| Rcsd1 | -0.49394 | 1.09E-07 | 0.003532 |
| Calm1 | -0.49517 | 3.33E-12 | 1.07E-07 |
| Runx1 | -0.49575 | 1.28E-05 | 0.413581 |
| Gm43305 | -0.49632 | 8.38E-09 | 0.00027 |
| Incenp | -0.49681 | 0.155433 | 1 |
| E2f2 | -0.49697 | 3.66E-10 | 1.18E-05 |
| Vim | -0.49781 | 1.42E-05 | 0.459691 |
| Mef2c | -0.49908 | 2.18E-07 | 0.007025 |
| Ncapd2 | -0.50161 | 0.01424 | 1 |
| Gpd2 | -0.50236 | 3.01E-08 | 0.000971 |
| Pmf1 | -0.50238 | 6.89E-06 | 0.22235 |
| Car2 | -0.50376 | 3.34E-24 | 1.08E-19 |
| Cdc42 | -0.5039 | 5.92E-16 | 1.91E-11 |
| Lig1 | -0.50559 | 0.00015 | 1 |
| Ywhaq | -0.50833 | 1.57E-10 | 5.06E-06 |
| Hist1h3i | -0.50941 | 1.73E-05 | 0.558585 |
| Asf1b | -0.51652 | 0.001472 | 1 |
| Cdk1 | -0.51799 | 1.01E-05 | 0.325968 |
| Nfyb | -0.52056 | 2.78E-11 | 8.99E-07 |
| Pafah1b3 | -0.52119 | 0.00756 | 1 |
| Zeb2 | -0.52424 | 0.011274 | 1 |

| | | | |
|---|---|---|---|
| Add3 | -0.52542 | 2.09E-08 | 0.000675 |
| Kif22 | -0.5281 | 2.1E-05 | 0.676572 |
| Il2rb | -0.52862 | 1.88E-06 | 0.060756 |
| Klrk1 | -0.5323 | 0.002631 | 1 |
| Dnah8 | -0.53234 | 2.08E-10 | 6.71E-06 |
| Kcnab2 | -0.53269 | 7.87E-08 | 0.002539 |
| Sidt1 | -0.53333 | 9.48E-10 | 3.06E-05 |
| Rom1 | -0.53439 | 1.73E-27 | 5.58E-23 |
| Nkain2 | -0.5349 | 1.76E-31 | 5.67E-27 |
| Nsmaf | -0.53515 | 9.36E-11 | 3.02E-06 |
| Lockd | -0.53638 | 2.4E-07 | 0.007762 |
| Arl5c | -0.53707 | 2.25E-16 | 7.25E-12 |
| Fyn | -0.53998 | 2.16E-08 | 0.000696 |
| Tpx2 | -0.54377 | 0.004943 | 1 |
| Mcm5 | -0.54496 | 1.01E-06 | 0.032489 |
| Osbpl3 | -0.54908 | 1.45E-08 | 0.000467 |
| Serf2 | -0.55261 | 2.17E-15 | 6.99E-11 |
| Pik3r1 | -0.55386 | 1.48E-07 | 0.004764 |
| Nusap1 | -0.55724 | 0.001228 | 1 |
| Pglyrp1 | -0.55984 | 1.02E-05 | 0.330526 |
| Gas7 | -0.56022 | 1.29E-25 | 4.15E-21 |
| Kcnq5 | -0.56188 | 2.34E-05 | 0.756602 |
| Mcm3 | -0.56245 | 0.000126 | 1 |
| Tuba1a | -0.56448 | 0.000128 | 1 |
| Lyn | -0.56758 | 0.001144 | 1 |
| Smc4 | -0.5681 | 0.000366 | 1 |
| Rrm1 | -0.57612 | 4.12E-06 | 0.132984 |
| Tubb4b | -0.57718 | 0.201645 | 1 |
| Ube2s | -0.57996 | 0.072221 | 1 |
| Ifi30 | -0.58259 | 0.009566 | 1 |
| Fbxo5 | -0.58377 | 1.31E-05 | 0.423258 |
| Cmah | -0.58508 | 3.1E-10 | 9.99E-06 |
| Racgap1 | -0.59033 | 9.63E-07 | 0.031079 |
| Txk | -0.59532 | 4.21E-10 | 1.36E-05 |
| Bin2 | -0.59832 | 1.87E-10 | 6.04E-06 |
| Fgfr2 | -0.59912 | 0.000632 | 1 |
| Smc2 | -0.60052 | 0.007349 | 1 |
| Ccna2 | -0.60076 | 1.63E-05 | 0.527277 |
| Birc5 | -0.6074 | 6.55E-05 | 1 |
| Lmo4 | -0.61369 | 0.012002 | 1 |
| P2ry12 | -0.61785 | 0.508589 | 1 |
| Pycard | -0.62147 | 5.69E-10 | 1.84E-05 |
| Hmgn2 | -0.62429 | 0.013636 | 1 |
| Sh2d1a | -0.62812 | 1.39E-09 | 4.49E-05 |
| Nrm | -0.63021 | 2.27E-09 | 7.33E-05 |
| Foxn3 | -0.6307 | 5.71E-08 | 0.001844 |
| Kif11 | -0.63087 | 0.001184 | 1 |
| Prc1 | -0.63208 | 1.79E-05 | 0.577466 |

| | | | |
|---|---|---|---|
| Ptpn4 | -0.63249 | 5.11E-11 | 1.65E-06 |
| Cenpf | -0.63314 | 0.000816 | 1 |
| Cd48 | -0.64495 | 9.52E-11 | 3.07E-06 |
| H2-Q6 | -0.64598 | 5.15E-05 | 1 |
| C1qc | -0.64766 | 0.299412 | 1 |
| Hist1h1a | -0.65064 | 0.000886 | 1 |
| Diaph3 | -0.65415 | 0.000533 | 1 |
| Agpat3 | -0.65468 | 3.46E-11 | 1.12E-06 |
| Sept11 | -0.65616 | 6.97E-11 | 2.25E-06 |
| S100a10 | -0.65712 | 8.8E-15 | 2.84E-10 |
| Nedd4 | -0.66451 | 4.47E-08 | 0.001442 |
| Cd55 | -0.6657 | 3.03E-32 | 9.79E-28 |
| H2-D1 | -0.6695 | 0.512226 | 1 |
| Dnmt1 | -0.67911 | 5.7E-07 | 0.018407 |
| Knl1 | -0.68343 | 8.71E-06 | 0.281321 |
| Tubb5 | -0.68615 | 0.46288 | 1 |
| Tm6sf1 | -0.69089 | 8.94E-15 | 2.89E-10 |
| Fcgr3 | -0.69178 | 0.000385 | 1 |
| Anxa6 | -0.69271 | 1.35E-19 | 4.37E-15 |
| Hist1h3e | -0.70203 | 0.002692 | 1 |
| Fry | -0.70371 | 2.61E-24 | 8.41E-20 |
| Txndc5 | -0.71406 | 7.4E-15 | 2.39E-10 |
| Ighm | -0.71824 | 0.000643 | 1 |
| Gm2682 | -0.71887 | 1.1E-05 | 0.35527 |
| Atp2a3 | -0.71899 | 3.9E-15 | 1.26E-10 |
| Cdc25b | -0.72744 | 9.68E-14 | 3.13E-09 |
| Rap1gap2 | -0.73473 | 3.26E-23 | 1.05E-18 |
| Myo1f | -0.73506 | 8.9E-12 | 2.87E-07 |
| Zyx | -0.73792 | 5.58E-15 | 1.8E-10 |
| Cd7 | -0.74393 | 2.22E-05 | 0.71692 |
| Ube2c | -0.75997 | 6E-05 | 1 |
| Tox | -0.76555 | 8.89E-12 | 2.87E-07 |
| Klre1 | -0.77218 | 6.19E-05 | 1 |
| Fcer1g | -0.78066 | 0.565798 | 1 |
| Ndufa4 | -0.78588 | 9.22E-19 | 2.98E-14 |
| Rrm2 | -0.79151 | 1.13E-06 | 0.03634 |
| Myo1g | -0.79238 | 3.28E-19 | 1.06E-14 |
| Klrg1 | -0.8014 | 4.03E-14 | 1.3E-09 |
| Plek | -0.82138 | 7.18E-09 | 0.000232 |
| Pclaf | -0.82429 | 8.67E-05 | 1 |
| Lmnb1 | -0.82732 | 2.64E-05 | 0.852398 |
| Bach2 | -0.82798 | 1.52E-09 | 4.91E-05 |
| Rasa3 | -0.84341 | 6.7E-13 | 2.16E-08 |
| Tuba1c | -0.84625 | 6.47E-06 | 0.208847 |
| H2afz | -0.8596 | 2.63E-08 | 0.000848 |
| Hmgb2 | -0.87301 | 3.88E-06 | 0.125229 |
| Anxa2 | -0.87508 | 5.26E-19 | 1.7E-14 |
| Ahnak | -0.88884 | 4.21E-09 | 0.000136 |

| | | | |
|---|---|---|---|
| Prr5l | -0.91709 | 1.67E-55 | 5.38E-51 |
| Hist1h4d | -0.93112 | 0.00177 | 1 |
| H2afx | -0.93364 | 0.020795 | 1 |
| Gzmk | -0.94066 | 1.73E-13 | 5.58E-09 |
| Igkc | -0.95343 | 2E-28 | 6.47E-24 |
| Top2a | -0.96565 | 0.000936 | 1 |
| Crip1 | -0.98697 | 7.06E-23 | 2.28E-18 |
| Eomes | -0.99099 | 7.99E-41 | 2.58E-36 |
| Stmn1 | -1.00239 | 4.21E-08 | 0.00136 |
| Emp3 | -1.02766 | 3.55E-20 | 1.14E-15 |
| Trbv3 | -1.04778 | 2.52E-22 | 8.14E-18 |
| Lpin1 | -1.05453 | 2.79E-33 | 9E-29 |
| Flna | -1.05758 | 2.29E-20 | 7.41E-16 |
| Hist1h2af | -1.0682 | 1.92E-05 | 0.61956 |
| Trav13-1 | -1.08257 | 1.07E-10 | 3.44E-06 |
| Trbv13-1 | -1.08539 | 0.007027 | 1 |
| Tuba1b | -1.08709 | 0.002087 | 1 |
| Lgals1 | -1.09578 | 1.52E-21 | 4.92E-17 |
| Cx3cr1 | -1.13437 | 2.15E-18 | 6.93E-14 |
| Tyrobp | -1.16074 | 0.000138 | 1 |
| Mki67 | -1.16791 | 8.79E-05 | 1 |
| Gm30211 | -1.18466 | 6.66E-18 | 2.15E-13 |
| Ccl5 | -1.20784 | 5.24E-15 | 1.69E-10 |
| Itga4 | -1.21579 | 8.33E-22 | 2.69E-17 |
| Cst3 | -1.23164 | 2.02E-05 | 0.65309 |
| Ripor2 | -1.28846 | 6.81E-21 | 2.2E-16 |
| Hist1h1c | -1.31309 | 1.84E-05 | 0.594979 |
| Gm20628 | -1.31324 | 0.000102 | 1 |
| Hist1h2ae | -1.31444 | 0.000133 | 1 |
| Hist1h2ab | -1.34462 | 1.4E-05 | 0.452098 |
| Hist1h1e | -1.39219 | 2.49E-05 | 0.805139 |
| Rasgrp2 | -1.48674 | 3.14E-32 | 1.01E-27 |
| Hist1h2ak | -1.50048 | 5.21E-05 | 1 |
| Fah | -1.55221 | 4.34E-56 | 1.4E-51 |
| Hist1h1b | -1.55922 | 4.54E-08 | 0.001467 |
| Hist1h1d | -1.58273 | 6.24E-07 | 0.02015 |
| Trbv1 | -1.69083 | 8.5E-12 | 2.75E-07 |
| Gzma | -1.96533 | 1.62E-17 | 5.24E-13 |
| Klf2 | -1.96821 | 7.37E-36 | 2.38E-31 |
| Cd74 | -2.20923 | 0.00124 | 1 |

### SUPPLEMENTAL TABLE 2. Differentially Regulated Genes in GVHD versus BM Macrophages

| Gene | avg_log2FC | p_val | p_val_adj |
|---|---|---|---|
| Chil3 | 2.903806 | 4.51E-27 | 1.46E-22 |
| Cxcl9 | 2.442837 | 3.37E-31 | 1.09E-26 |
| Ccl5 | 2.033277 | 2.12E-14 | 6.83E-10 |
| Cfb | 1.957911 | 1.67E-37 | 5.38E-33 |
| C1qb | 1.913075 | 2.22E-32 | 7.15E-28 |
| Mif | 1.845336 | 3.04E-46 | 9.8E-42 |
| S100a8 | 1.843425 | 2.51E-20 | 8.12E-16 |
| Arg1 | 1.756947 | 7.01E-15 | 2.26E-10 |
| C1qc | 1.704767 | 1.5E-29 | 4.84E-25 |
| AA467197 | 1.670829 | 2.79E-27 | 9.01E-23 |
| C1qa | 1.625199 | 1.38E-32 | 4.47E-28 |
| Atp1b3 | 1.597196 | 2.9E-23 | 9.35E-19 |
| Cstb | 1.569929 | 5.44E-46 | 1.76E-41 |
| Ly6a | 1.514247 | 4.82E-33 | 1.56E-28 |
| Mt1 | 1.479602 | 5.39E-40 | 1.74E-35 |
| AW11201C | 1.455218 | 5.57E-30 | 1.8E-25 |
| Gbp2b | 1.426488 | 5.26E-23 | 1.7E-18 |
| Naaa | 1.412619 | 1.01E-43 | 3.26E-39 |
| Saa3 | 1.282853 | 6.29E-13 | 2.03E-08 |
| Capg | 1.255354 | 1.75E-38 | 5.65E-34 |
| Upp1 | 1.194638 | 2.57E-28 | 8.29E-24 |
| Cxcl10 | 1.185069 | 1.46E-16 | 4.71E-12 |
| Atp6v0c | 1.16339 | 3.1E-40 | 1E-35 |
| Gbp2 | 1.162065 | 3.46E-22 | 1.12E-17 |
| Fth1 | 1.154309 | 1.24E-26 | 4E-22 |
| Tpi1 | 1.150908 | 5.68E-33 | 1.83E-28 |
| Ctsd | 1.149506 | 3.08E-35 | 9.95E-31 |
| Tbxas1 | 1.144055 | 1.83E-42 | 5.91E-38 |
| Pgam1 | 1.139071 | 2.75E-30 | 8.87E-26 |
| Fcgr2b | 1.113581 | 5.5E-30 | 1.77E-25 |
| Ctsc | 1.108892 | 1.06E-31 | 3.41E-27 |
| S100a11 | 1.095159 | 9.98E-28 | 3.22E-23 |
| Aldoa | 1.087262 | 4.34E-36 | 1.4E-31 |
| Gbp4 | 1.070632 | 2.17E-33 | 7E-29 |
| Cd1d1 | 1.067626 | 1.56E-34 | 5.05E-30 |
| Ifi30 | 1.064698 | 2.34E-34 | 7.57E-30 |
| Socs1 | 1.053212 | 2.67E-31 | 8.63E-27 |
| Gbp7 | 1.04851 | 1.87E-32 | 6.05E-28 |
| Txn1 | 1.047075 | 3.65E-35 | 1.18E-30 |
| Ier3 | 1.024432 | 1.09E-22 | 3.51E-18 |
| Dok2 | 1.024311 | 7.54E-33 | 2.43E-28 |
| Fkbp5 | 1.019976 | 1.44E-27 | 4.66E-23 |
| Sdc4 | 1.006327 | 1.16E-22 | 3.75E-18 |
| H2-Eb1 | 1.000127 | 7.83E-18 | 2.53E-13 |

| | | | |
|---|---|---|---|
| Lcn2 | 0.994687 | 8.96E-10 | 2.89E-05 |
| S100a10 | 0.993781 | 3.18E-26 | 1.03E-21 |
| H2-Ab1 | 0.981721 | 2.67E-23 | 8.63E-19 |
| Shtn1 | 0.972734 | 3.17E-38 | 1.02E-33 |
| Creb5 | 0.971936 | 5.82E-28 | 1.88E-23 |
| Cd38 | 0.963919 | 1.07E-37 | 3.45E-33 |
| Hif1a | 0.960531 | 3.64E-18 | 1.17E-13 |
| Glrx | 0.949105 | 9.01E-26 | 2.91E-21 |
| St7 | 0.944361 | 5.85E-31 | 1.89E-26 |
| Ccr1 | 0.943944 | 1.05E-34 | 3.39E-30 |
| Fcgr3 | 0.94241 | 1.48E-28 | 4.79E-24 |
| Serpina3g | 0.93499 | 1.25E-17 | 4.03E-13 |
| Ninj1 | 0.933248 | 2.01E-22 | 6.49E-18 |
| Ccl6 | 0.92701 | 6.41E-15 | 2.07E-10 |
| C4b | 0.92554 | 6.26E-17 | 2.02E-12 |
| C3 | 0.924908 | 5.18E-14 | 1.67E-09 |
| Sod2 | 0.924902 | 9.94E-15 | 3.21E-10 |
| Gbp8 | 0.924511 | 6.2E-26 | 2E-21 |
| Cox8a | 0.92304 | 9.2E-44 | 2.97E-39 |
| Tspo | 0.89432 | 2.77E-28 | 8.94E-24 |
| Lpl | 0.887176 | 3.61E-17 | 1.16E-12 |
| Pfkp | 0.8857 | 1.8E-24 | 5.81E-20 |
| Agtrap | 0.884959 | 2.33E-24 | 7.53E-20 |
| Fmnl2 | 0.879808 | 2.7E-20 | 8.71E-16 |
| Ass1 | 0.877781 | 7.26E-14 | 2.34E-09 |
| Iigp1 | 0.8684 | 9.04E-21 | 2.92E-16 |
| Gbp5 | 0.867145 | 9.66E-18 | 3.12E-13 |
| Ldha | 0.861597 | 3.46E-23 | 1.12E-18 |
| Bst1 | 0.856423 | 2.31E-30 | 7.45E-26 |
| Clec4n | 0.855922 | 4.8E-14 | 1.55E-09 |
| Atp5g3 | 0.852845 | 2.09E-34 | 6.76E-30 |
| Atp5g1 | 0.8494 | 1.63E-26 | 5.28E-22 |
| Cfp | 0.847871 | 1.25E-27 | 4.03E-23 |
| Ccr5 | 0.841761 | 6.38E-24 | 2.06E-19 |
| F10 | 0.841671 | 2.03E-17 | 6.56E-13 |
| mt-Atp8 | 0.837301 | 1.64E-39 | 5.3E-35 |
| Mertk | 0.834458 | 2.01E-23 | 6.49E-19 |
| Zc3h7a | 0.829482 | 0.079924 | 1 |
| Fbxl5 | 0.824349 | 3.36E-17 | 1.09E-12 |
| Il18bp | 0.816485 | 5E-25 | 1.61E-20 |
| Acod1 | 0.815922 | 2.52E-07 | 0.008144 |
| Dip2c | 0.801431 | 4.45E-27 | 1.44E-22 |
| Gatm | 0.798568 | 1.02E-27 | 3.3E-23 |
| H2-Aa | 0.797788 | 5.24E-19 | 1.69E-14 |
| Ms4a8a | 0.79773 | 1.57E-24 | 5.07E-20 |
| Tagln2 | 0.796502 | 8.54E-24 | 2.76E-19 |
| App | 0.793895 | 1.48E-25 | 4.79E-21 |
| Ifi205 | 0.792014 | 7.31E-20 | 2.36E-15 |

| | | | |
|---|---|---|---|
| Aprt | 0.789039 | 1.13E-25 | 3.64E-21 |
| Gapdh | 0.787903 | 1.89E-29 | 6.1E-25 |
| Frmd4b | 0.786873 | 3.48E-19 | 1.12E-14 |
| Cd164 | 0.782566 | 4.85E-29 | 1.57E-24 |
| Spint1 | 0.781028 | 2.93E-26 | 9.46E-22 |
| Snx5 | 0.77921 | 6.7E-25 | 2.16E-20 |
| Wfdc21 | 0.778131 | 1.69E-11 | 5.45E-07 |
| Ctsl | 0.776484 | 4.3E-21 | 1.39E-16 |
| Tfec | 0.77453 | 1.52E-31 | 4.92E-27 |
| Klhl6 | 0.772336 | 5.71E-19 | 1.85E-14 |
| Gbp6 | 0.765402 | 2.97E-17 | 9.59E-13 |
| Anxa4 | 0.765115 | 4.43E-24 | 1.43E-19 |
| Thbs1 | 0.763252 | 7.76E-07 | 0.025042 |
| Grn | 0.762587 | 6.69E-21 | 2.16E-16 |
| Lilr4b | 0.762263 | 5.98E-19 | 1.93E-14 |
| Pycard | 0.756628 | 4.8E-19 | 1.55E-14 |
| Prdx5 | 0.754031 | 1.65E-22 | 5.34E-18 |
| Prelid1 | 0.752677 | 9.96E-27 | 3.21E-22 |
| Cxcl16 | 0.749569 | 3.61E-16 | 1.17E-11 |
| Jaml | 0.749544 | 3.88E-20 | 1.25E-15 |
| Chchd10 | 0.739599 | 1.08E-33 | 3.5E-29 |
| Rps2 | 0.737055 | 5.79E-30 | 1.87E-25 |
| Rpl12 | 0.736385 | 3.56E-22 | 1.15E-17 |
| Slc31a2 | 0.730658 | 2.41E-19 | 7.79E-15 |
| Cox7b | 0.725499 | 1.65E-20 | 5.34E-16 |
| Lap3 | 0.725312 | 3.98E-22 | 1.28E-17 |
| Anxa3 | 0.723557 | 2.37E-24 | 7.65E-20 |
| Mitf | 0.723515 | 9.12E-19 | 2.95E-14 |
| Sell | 0.72168 | 6.37E-18 | 2.06E-13 |
| Ctsz | 0.721674 | 2.12E-21 | 6.86E-17 |
| Kynu | 0.72151 | 3.64E-19 | 1.18E-14 |
| Gpx4 | 0.721163 | 5.21E-15 | 1.68E-10 |
| Axl | 0.717442 | 4.69E-23 | 1.52E-18 |
| Cd74 | 0.713094 | 3.23E-16 | 1.04E-11 |
| Nme2 | 0.711868 | 6.23E-24 | 2.01E-19 |
| Myof | 0.702245 | 3.91E-19 | 1.26E-14 |
| Rrbp1 | 0.701046 | 6.34E-20 | 2.05E-15 |
| Cxcl2 | 0.695396 | 1.33E-08 | 0.000429 |
| Bcl2a1b | 0.694163 | 6.02E-12 | 1.94E-07 |
| Ecm1 | 0.69223 | 7.2E-21 | 2.33E-16 |
| Wfdc17 | 0.691003 | 4.89E-13 | 1.58E-08 |
| Clec12a | 0.689439 | 2.37E-17 | 7.65E-13 |
| mt-Co2 | 0.689195 | 1.65E-26 | 5.33E-22 |
| mt-Co3 | 0.689164 | 3.3E-21 | 1.07E-16 |
| Tuba1b | 0.689115 | 5.62E-13 | 1.81E-08 |
| Scimp | 0.688973 | 1.59E-24 | 5.12E-20 |
| Ly6i | 0.682683 | 0.090609 | 1 |
| C3ar1 | 0.678725 | 2.73E-23 | 8.82E-19 |

| | | | |
|---|---|---|---|
| Gbp3 | 0.677923 | 6E-16 | 1.94E-11 |
| C1qbp | 0.675363 | 6.06E-19 | 1.96E-14 |
| Plxdc2 | 0.672168 | 5.89E-09 | 0.00019 |
| Pim1 | 0.667078 | 6.45E-16 | 2.08E-11 |
| Rpl14 | 0.666962 | 7.04E-27 | 2.27E-22 |
| Serpina3f | 0.661871 | 9.06E-25 | 2.92E-20 |
| Cd84 | 0.660691 | 4.69E-19 | 1.51E-14 |
| Mpc1 | 0.659 | 1.23E-17 | 3.96E-13 |
| Ran | 0.656861 | 6.8E-16 | 2.2E-11 |
| Vim | 0.656262 | 5.05E-16 | 1.63E-11 |
| Tubb5 | 0.655214 | 1.09E-09 | 3.51E-05 |
| Tarm1 | 0.654578 | 2.6E-24 | 8.4E-20 |
| Il1r2 | 0.652791 | 1.87E-15 | 6.04E-11 |
| Slc7a2 | 0.646684 | 6.07E-14 | 1.96E-09 |
| Rplp1 | 0.645917 | 1.59E-32 | 5.12E-28 |
| Clec5a | 0.645798 | 2.53E-14 | 8.16E-10 |
| Slamf8 | 0.643568 | 9.7E-19 | 3.13E-14 |
| Cox5b | 0.642791 | 2.56E-23 | 8.27E-19 |
| Pkm | 0.642344 | 2.07E-17 | 6.69E-13 |
| Mmp14 | 0.634331 | 1.61E-08 | 0.000519 |
| Vdac2 | 0.629021 | 2.35E-19 | 7.59E-15 |
| Nampt | 0.628468 | 2.14E-16 | 6.9E-12 |
| Dab2 | 0.627984 | 4.06E-32 | 1.31E-27 |
| Hk3 | 0.625818 | 7.92E-12 | 2.56E-07 |
| Lgals3bp | 0.624823 | 2.35E-15 | 7.59E-11 |
| Myo1e | 0.623945 | 5.29E-25 | 1.71E-20 |
| Atp5b | 0.622097 | 1.77E-17 | 5.71E-13 |
| Msr1 | 0.620088 | 2.1E-14 | 6.77E-10 |
| Gm4951 | 0.612996 | 3.14E-18 | 1.02E-13 |
| Lilrb4a | 0.611757 | 2.6E-13 | 8.41E-09 |
| Sh3pxd2b | 0.611354 | 7.71E-23 | 2.49E-18 |
| Akr1a1 | 0.60841 | 1.12E-18 | 3.6E-14 |
| F7 | 0.607049 | 2.04E-20 | 6.58E-16 |
| Eps8 | 0.605841 | 3.7E-15 | 1.19E-10 |
| Npc2 | 0.605629 | 1.19E-18 | 3.83E-14 |
| Atp5o | 0.60409 | 7.97E-18 | 2.57E-13 |
| H2-DMb2 | 0.601906 | 2.56E-16 | 8.27E-12 |
| mt-Co1 | 0.601543 | 2.06E-22 | 6.65E-18 |
| mt-Cytb | 0.600054 | 1.47E-15 | 4.74E-11 |
| Picalm | 0.598295 | 9.14E-11 | 2.95E-06 |
| Cdkn1a | 0.596712 | 2.98E-13 | 9.61E-09 |
| AY036118 | 0.596181 | 0.025351 | 1 |
| Tubb4b | 0.595577 | 3.89E-11 | 1.26E-06 |
| Prdx6 | 0.587734 | 1.02E-14 | 3.3E-10 |
| Rbm3 | 0.585703 | 7.62E-19 | 2.46E-14 |
| H2-DMa | 0.582849 | 1.31E-14 | 4.22E-10 |
| Arhgap24 | 0.581887 | 6.11E-14 | 1.97E-09 |
| mt-Nd1 | 0.580309 | 2.14E-11 | 6.92E-07 |

| | | | |
|---|---|---|---|
| Nos2 | 0.580294 | 1.51E-07 | 0.004864 |
| Pgk1 | 0.578259 | 4.84E-12 | 1.56E-07 |
| Bst2 | 0.577393 | 1.38E-14 | 4.45E-10 |
| Psma5 | 0.577168 | 3.43E-18 | 1.11E-13 |
| Procr | 0.576351 | 1.44E-14 | 4.65E-10 |
| Atp5g2 | 0.57296 | 4.32E-18 | 1.4E-13 |
| S100a1 | 0.571994 | 1.74E-25 | 5.62E-21 |
| Eif5a | 0.571303 | 6.67E-12 | 2.15E-07 |
| mt-Atp6 | 0.568511 | 9.01E-16 | 2.91E-11 |
| Ndufb8 | 0.568142 | 2.26E-16 | 7.29E-12 |
| Aoah | 0.568013 | 1.43E-15 | 4.61E-11 |
| Hopx | 0.565022 | 1.76E-12 | 5.68E-08 |
| Cd300lf | 0.56397 | 7.23E-16 | 2.34E-11 |
| Csf2rb | 0.563155 | 3.89E-12 | 1.26E-07 |
| Fgl2 | 0.561953 | 5.28E-13 | 1.7E-08 |
| Sdcbp | 0.561425 | 1.7E-15 | 5.5E-11 |
| AU020206 | 0.559445 | 1.79E-13 | 5.78E-09 |
| Clcn7 | 0.556461 | 7.6E-18 | 2.45E-13 |
| Lmna | 0.554468 | 1.51E-17 | 4.87E-13 |
| Calr | 0.55421 | 5.11E-11 | 1.65E-06 |
| Htr7 | 0.551978 | 9.82E-21 | 3.17E-16 |
| Ndufa12 | 0.550757 | 8.76E-17 | 2.83E-12 |
| Gns | 0.546499 | 1.75E-12 | 5.66E-08 |
| Igtp | 0.544926 | 1.63E-15 | 5.26E-11 |
| Atpif1 | 0.544554 | 4.67E-16 | 1.51E-11 |
| Cndp2 | 0.544538 | 3.69E-14 | 1.19E-09 |
| Trf | 0.543939 | 0.000121 | 1 |
| Fabp5 | 0.543867 | 0.066047 | 1 |
| Lgals1 | 0.540306 | 2.21E-10 | 7.14E-06 |
| Sdhd | 0.539016 | 2.02E-14 | 6.51E-10 |
| Cox6a1 | 0.537384 | 4.52E-15 | 1.46E-10 |
| Efhd2 | 0.537166 | 5.14E-14 | 1.66E-09 |
| Atp5j | 0.533148 | 1.19E-15 | 3.84E-11 |
| Slc3a2 | 0.532367 | 3.41E-13 | 1.1E-08 |
| Socs3 | 0.529401 | 1.01E-09 | 3.25E-05 |
| Slamf7 | 0.528996 | 4.88E-13 | 1.57E-08 |
| Kcnn4 | 0.528397 | 8.65E-21 | 2.79E-16 |
| Zfp703 | 0.528278 | 8.58E-23 | 2.77E-18 |
| Slco3a1 | 0.527266 | 2.83E-09 | 9.14E-05 |
| Cd24a | 0.524121 | 4.65E-17 | 1.5E-12 |
| Aif1 | 0.522834 | 1.29E-12 | 4.15E-08 |
| Slc49a4 | 0.522296 | 7.85E-16 | 2.53E-11 |
| Cd274 | 0.521216 | 1.14E-08 | 0.000367 |
| Lgals9 | 0.52058 | 1.72E-11 | 5.56E-07 |
| Anpep | 0.519095 | 6.78E-16 | 2.19E-11 |
| mt-Nd4l | 0.518161 | 2.13E-11 | 6.88E-07 |
| Vdac1 | 0.514616 | 1.79E-16 | 5.79E-12 |
| Gda | 0.512384 | 4.6E-12 | 1.49E-07 |

| | | | |
|---|---|---|---|
| P2ry14 | 0.512271 | 1.87E-20 | 6.03E-16 |
| Gpnmb | 0.511499 | 1.64E-12 | 5.31E-08 |
| Clic4 | 0.510517 | 5.12E-08 | 0.001653 |
| Emb | 0.509546 | 7.41E-12 | 2.39E-07 |
| Acly | 0.509095 | 2.43E-09 | 7.84E-05 |
| Arid5b | 0.507417 | 7.91E-09 | 0.000255 |
| Tgfbi | 0.507032 | 5.74E-10 | 1.85E-05 |
| Dram1 | 0.504133 | 4.53E-09 | 0.000146 |
| Crip1 | 0.503286 | 1.31E-08 | 0.000421 |
| 1600014C: | 0.502613 | 8.07E-13 | 2.61E-08 |
| Palld | 0.498993 | 1.83E-23 | 5.9E-19 |
| Ndufb6 | 0.498849 | 7.02E-10 | 2.26E-05 |
| Eepd1 | 0.495928 | 1.24E-18 | 4.02E-14 |
| Uck2 | 0.495739 | 2.52E-23 | 8.15E-19 |
| Ctss | 0.494115 | 1.12E-10 | 3.62E-06 |
| Acsl1 | 0.491726 | 0.000176 | 1 |
| Rps20 | 0.490741 | 8.18E-20 | 2.64E-15 |
| Csf2rb2 | 0.489585 | 5.92E-16 | 1.91E-11 |
| mt-Nd5 | 0.489312 | 1.38E-09 | 4.45E-05 |
| Ppia | 0.489185 | 2.13E-19 | 6.88E-15 |
| Manf | 0.488366 | 8.23E-11 | 2.66E-06 |
| Isg15 | 0.485706 | 9.23E-09 | 0.000298 |
| Psma3 | 0.484158 | 1.11E-12 | 3.58E-08 |
| Nme1 | 0.48361 | 1.73E-12 | 5.57E-08 |
| Oasl1 | 0.480925 | 3.77E-09 | 0.000122 |
| Slfn1 | 0.47929 | 4.18E-11 | 1.35E-06 |
| Htra3 | 0.477639 | 3.07E-11 | 9.91E-07 |
| Pdia3 | 0.477164 | 1.5E-09 | 4.84E-05 |
| Nhp2 | 0.476102 | 3.57E-14 | 1.15E-09 |
| Rnf7 | 0.475616 | 4.82E-14 | 1.56E-09 |
| Wars | 0.47559 | 8.28E-13 | 2.67E-08 |
| Hmgn2 | 0.475076 | 2E-10 | 6.45E-06 |
| Atox1 | 0.473872 | 2.02E-12 | 6.52E-08 |
| Fkbp2 | 0.472653 | 1.3E-12 | 4.21E-08 |
| Rpl10a | 0.472594 | 3.2E-15 | 1.03E-10 |
| Klf9 | 0.47003 | 9.69E-17 | 3.13E-12 |
| Cd14 | 0.469213 | 5.16E-12 | 1.67E-07 |
| Aopep | 0.46843 | 2.43E-16 | 7.85E-12 |
| Hmga1 | 0.467083 | 2.98E-11 | 9.63E-07 |
| Nek6 | 0.466428 | 1.44E-17 | 4.64E-13 |
| Mrpl23 | 0.465838 | 9.63E-14 | 3.11E-09 |
| Coa5 | 0.464495 | 2.85E-14 | 9.22E-10 |
| Ddhd1 | 0.461705 | 7.87E-10 | 2.54E-05 |
| Amdhd2 | 0.461228 | 6.1E-10 | 1.97E-05 |
| Uqcrb | 0.459076 | 1.24E-13 | 4.01E-09 |
| Pde7b | 0.458621 | 3.02E-06 | 0.097539 |
| Ppib | 0.45852 | 9.27E-11 | 2.99E-06 |
| Il1rn | 0.457997 | 4.99E-09 | 0.000161 |

| | | | |
|---|---|---|---|
| Atrnl1 | 0.45655 | 9.2E-12 | 2.97E-07 |
| Sema4a | 0.45613 | 7.9E-09 | 0.000255 |
| Ppargc1b | 0.456102 | 9.27E-19 | 2.99E-14 |
| Mki67 | 0.455267 | 6.73E-09 | 0.000217 |
| Snrpf | 0.45302 | 1.17E-09 | 3.78E-05 |
| Igsf8 | 0.452783 | 9.89E-15 | 3.19E-10 |
| Slc25a3 | 0.450962 | 1.21E-12 | 3.9E-08 |
| Glipr2 | 0.449991 | 7.82E-14 | 2.52E-09 |
| Got1 | 0.448803 | 8.7E-20 | 2.81E-15 |
| Ccdc86 | 0.44871 | 2.33E-17 | 7.53E-13 |
| Sdhb | 0.444462 | 1.04E-12 | 3.35E-08 |
| Acadl | 0.443558 | 2.86E-14 | 9.24E-10 |
| Resf1 | 0.442177 | 4.61E-06 | 0.148965 |
| Ndufs3 | 0.441622 | 4.14E-12 | 1.34E-07 |
| Psen2 | 0.441544 | 2.96E-12 | 9.56E-08 |
| Chn2 | 0.440635 | 4.05E-06 | 0.130794 |
| Micos10 | 0.440582 | 2.39E-12 | 7.7E-08 |
| Ak2 | 0.439955 | 5.08E-10 | 1.64E-05 |
| Slc25a5 | 0.439216 | 4.29E-14 | 1.38E-09 |
| H2-DMb1 | 0.438957 | 9.5E-14 | 3.07E-09 |
| Ndufc1 | 0.438319 | 8.25E-10 | 2.66E-05 |
| Trerf1 | 0.438026 | 4.09E-10 | 1.32E-05 |
| Map2k2 | 0.43669 | 7.77E-13 | 2.51E-08 |
| Ctsh | 0.434559 | 2.44E-10 | 7.86E-06 |
| C1ra | 0.434403 | 2.48E-18 | 8.02E-14 |
| CAAA0114[illegible] | 0.432677 | 3.29E-08 | 0.001063 |
| Atp5d | 0.429769 | 1.85E-11 | 5.97E-07 |
| Gas7 | 0.429542 | 5.35E-15 | 1.73E-10 |
| Card19 | 0.42843 | 3.24E-12 | 1.05E-07 |
| Ndufv3 | 0.426788 | 9.53E-13 | 3.08E-08 |
| Ube2c | 0.426243 | 8.28E-05 | 1 |
| Spon1 | 0.424089 | 5.42E-12 | 1.75E-07 |
| Irf2bp2 | 0.423181 | 4.29E-12 | 1.39E-07 |
| Arrdc4 | 0.421453 | 1.74E-14 | 5.61E-10 |
| Dusp1 | 0.420028 | 2.21E-06 | 0.071503 |
| Pld3 | 0.419684 | 1.01E-12 | 3.25E-08 |
| Ccl2 | 0.419092 | 0.003971 | 1 |
| Ccl4 | 0.418573 | 3.2E-05 | 1 |
| Ttc39b | 0.41844 | 5.04E-13 | 1.63E-08 |
| Tlr4 | 0.417727 | 1.31E-11 | 4.24E-07 |
| Pebp1 | 0.417008 | 4.64E-14 | 1.5E-09 |
| Top2a | 0.41696 | 0.041596 | 1 |
| Stmn1 | 0.416834 | 5.66E-06 | 0.182738 |
| Large1 | 0.416181 | 0.000536 | 1 |
| Slc2a1 | 0.414209 | 2.37E-07 | 0.00766 |
| Scarf1 | 0.414066 | 3.91E-14 | 1.26E-09 |
| Psmb10 | 0.413989 | 1.01E-13 | 3.27E-09 |
| Acp5 | 0.413857 | 1.04E-05 | 0.336813 |

| | | | |
|---|---|---|---|
| Cox6b1 | 0.41324 | 2.03E-13 | 6.56E-09 |
| Alas1 | 0.41255 | 6.51E-06 | 0.210293 |
| Rpl32 | 0.412449 | 7.47E-14 | 2.41E-09 |
| Apoe | 0.412281 | 0.002644 | 1 |
| Mgst1 | 0.412208 | 4.03E-07 | 0.012996 |
| Pacsin2 | 0.411179 | 2.41E-10 | 7.77E-06 |
| Pnpla2 | 0.411161 | 2.02E-10 | 6.51E-06 |
| Qpct | 0.411068 | 2.01E-08 | 0.00065 |
| Nrp2 | 0.411006 | 1.23E-15 | 3.98E-11 |
| Rasgef1b | 0.410204 | 5.61E-10 | 1.81E-05 |
| Ptpmt1 | 0.410169 | 1.25E-13 | 4.03E-09 |
| Eno1 | 0.409666 | 3.72E-10 | 1.2E-05 |
| Rps17 | 0.409539 | 1.53E-11 | 4.93E-07 |
| Psma6 | 0.409538 | 2.11E-10 | 6.82E-06 |
| Gng2 | 0.409327 | 2.71E-10 | 8.74E-06 |
| Ly6c1 | 0.408845 | 1.68E-06 | 0.054194 |
| Ndufa1 | 0.407172 | 1.34E-08 | 0.000433 |
| Cnn3 | 0.407102 | 2.82E-10 | 9.09E-06 |
| Bnip3 | 0.40709 | 1.25E-12 | 4.02E-08 |
| Csrp1 | 0.406414 | 8.91E-12 | 2.88E-07 |
| Ifi47 | 0.406308 | 5.65E-08 | 0.001825 |
| Idh3b | 0.404463 | 1.09E-09 | 3.52E-05 |
| Snx2 | 0.404166 | 2.78E-08 | 0.000898 |
| Tmem154 | 0.403543 | 2.78E-13 | 8.97E-09 |
| Psme2 | 0.403378 | 3.26E-09 | 0.000105 |
| Rps27l | 0.40091 | 1.31E-08 | 0.000422 |
| 0610012G( | 0.400341 | 3.74E-14 | 1.21E-09 |
| F830016B( | 0.398744 | 1.77E-15 | 5.71E-11 |
| Uqcrq | 0.398729 | 1.3E-10 | 4.21E-06 |
| Gng12 | 0.397732 | 9.37E-11 | 3.02E-06 |
| Ppa1 | 0.397021 | 6.63E-17 | 2.14E-12 |
| Ndufa13 | 0.397003 | 4.05E-08 | 0.001307 |
| Rab7b | 0.395599 | 4.71E-11 | 1.52E-06 |
| Anp32b | 0.395442 | 4.99E-11 | 1.61E-06 |
| Atp5a1 | 0.394656 | 1.53E-09 | 4.94E-05 |
| Tcirg1 | 0.393587 | 1.15E-11 | 3.73E-07 |
| Sla | 0.393291 | 2.05E-06 | 0.066282 |
| Atp6ap2 | 0.393045 | 5.65E-10 | 1.83E-05 |
| Litaf | 0.392755 | 3.39E-09 | 0.000109 |
| Sdf2l1 | 0.392091 | 3.28E-08 | 0.00106 |
| P4hb | 0.391655 | 1.87E-09 | 6.03E-05 |
| Hsd3b7 | 0.391371 | 3.31E-10 | 1.07E-05 |
| Cox6c | 0.391369 | 3.12E-11 | 1.01E-06 |
| Rpl15 | 0.3911 | 4.24E-11 | 1.37E-06 |
| Cox4i1 | 0.389911 | 2.5E-14 | 8.09E-10 |
| Cox5a | 0.38915 | 7.66E-11 | 2.47E-06 |
| Lamp1 | 0.389029 | 6.39E-11 | 2.06E-06 |
| Cycs | 0.388945 | 4.12E-09 | 0.000133 |

| | | | |
|---|---|---|---|
| Cd86 | 0.388157 | 9.32E-07 | 0.030097 |
| Ndufb5 | 0.388021 | 6.27E-09 | 0.000202 |
| Chmp4b | 0.386225 | 2.55E-10 | 8.25E-06 |
| Gpi1 | 0.385082 | 8.7E-09 | 0.000281 |
| Ppt1 | 0.38422 | 1.09E-06 | 0.035031 |
| Cd200r1 | 0.383146 | 6.18E-16 | 1.99E-11 |
| Pmf1 | 0.382105 | 1.49E-15 | 4.82E-11 |
| Plaat3 | 0.38194 | 3.64E-10 | 1.17E-05 |
| Eci2 | 0.38158 | 5.3E-08 | 0.001711 |
| Pnp | 0.381328 | 1.4E-08 | 0.000453 |
| Lrrc59 | 0.380953 | 9.45E-17 | 3.05E-12 |
| Rps6 | 0.380901 | 2.91E-11 | 9.39E-07 |
| Elob | 0.379229 | 6.52E-12 | 2.1E-07 |
| Comtd1 | 0.378563 | 7.46E-15 | 2.41E-10 |
| Pfkl | 0.378133 | 2.87E-11 | 9.25E-07 |
| Srsf3 | 0.377963 | 3.63E-07 | 0.011712 |
| Per1 | 0.377475 | 8.1E-11 | 2.61E-06 |
| Ndufs5 | 0.377411 | 2.01E-09 | 6.5E-05 |
| Cnih4 | 0.377255 | 1.05E-08 | 0.000339 |
| Ccrl2 | 0.376943 | 2.62E-05 | 0.845735 |
| Timm23 | 0.376768 | 2.26E-13 | 7.31E-09 |
| Vdr | 0.376315 | 5.22E-18 | 1.68E-13 |
| C5ar1 | 0.375183 | 1.17E-13 | 3.77E-09 |
| Cep83 | 0.375069 | 7.55E-12 | 2.44E-07 |
| Mrps36 | 0.374627 | 1.05E-12 | 3.4E-08 |
| Eif3i | 0.374261 | 2.08E-09 | 6.72E-05 |
| Hsp90b1 | 0.373952 | 2.76E-06 | 0.088947 |
| Rexo2 | 0.373117 | 9.21E-10 | 2.97E-05 |
| Timd4 | 0.37297 | 2.34E-05 | 0.756099 |
| Cox7a2 | 0.372489 | 8.58E-12 | 2.77E-07 |
| Pdia6 | 0.372023 | 8.2E-07 | 0.026462 |
| Hspa9 | 0.371947 | 2.31E-11 | 7.44E-07 |
| H2afz | 0.371671 | 1.36E-08 | 0.000439 |
| Cyp4v3 | 0.371634 | 4E-10 | 1.29E-05 |
| H2-Q2 | 0.371594 | 2.16E-06 | 0.069887 |
| Ifitm1 | 0.370186 | 0.008746 | 1 |
| Cped1 | 0.369529 | 1.09E-15 | 3.51E-11 |
| Txndc17 | 0.369338 | 2.43E-10 | 7.83E-06 |
| Tmem173 | 0.368539 | 4.65E-11 | 1.5E-06 |
| Ppp1r14b | 0.368327 | 1.85E-17 | 5.97E-13 |
| Ndufa5 | 0.367659 | 8.43E-10 | 2.72E-05 |
| Socs2 | 0.367607 | 1.6E-18 | 5.17E-14 |
| Nucb1 | 0.365227 | 3.23E-09 | 0.000104 |
| Tma16 | 0.364698 | 4.12E-17 | 1.33E-12 |
| Cyc1 | 0.364329 | 1.32E-07 | 0.004265 |
| Gm4258 | 0.364265 | 8.19E-13 | 2.64E-08 |
| Magohb | 0.363486 | 7.57E-08 | 0.002445 |
| Ndufb9 | 0.363087 | 7.48E-09 | 0.000241 |

| | | | |
|---|---|---|---|
| Ndufab1 | 0.362556 | 2.66E-10 | 8.57E-06 |
| Swi5 | 0.358316 | 8.9E-06 | 0.287215 |
| Grina | 0.357951 | 6.61E-05 | 1 |
| Ptms | 0.357418 | 4.58E-10 | 1.48E-05 |
| Cish | 0.357102 | 2.5E-17 | 8.06E-13 |
| Ptgs1 | 0.356898 | 1.39E-13 | 4.49E-09 |
| Idh3a | 0.356447 | 8.66E-15 | 2.8E-10 |
| Pa2g4 | 0.355868 | 1.24E-10 | 4.02E-06 |
| Saraf | 0.355657 | 6.4E-07 | 0.020663 |
| Il1a | 0.355345 | 3.76E-09 | 0.000121 |
| Ciita | 0.355297 | 1.01E-08 | 0.000325 |
| Cyba | 0.35445 | 6.19E-11 | 2E-06 |
| Clic5 | 0.354343 | 4.82E-11 | 1.56E-06 |
| Uqcr11 | 0.3541 | 5.2E-06 | 0.167878 |
| Mdh2 | 0.354018 | 8.51E-09 | 0.000275 |
| 1110008P1 | 0.353903 | 5.42E-07 | 0.01749 |
| Hmgb1 | 0.353581 | 3.66E-06 | 0.118103 |
| Ccl9 | 0.352616 | 6.39E-05 | 1 |
| Ssr4 | 0.352275 | 3.39E-10 | 1.1E-05 |
| Colgalt1 | 0.352078 | 6.08E-11 | 1.96E-06 |
| Cd40 | 0.351929 | 1.86E-10 | 6.01E-06 |
| Pbx1 | 0.351691 | 3.11E-05 | 1 |
| Ftl1 | 0.351501 | 1.03E-14 | 3.33E-10 |
| Ptgs2 | 0.351499 | 2.49E-09 | 8.04E-05 |
| Gna13 | 0.351399 | 9.94E-10 | 3.21E-05 |
| Furin | 0.351305 | 3.67E-09 | 0.000119 |
| Vps29 | 0.350817 | 1.95E-09 | 6.31E-05 |
| Higd1a | 0.350588 | 2.94E-10 | 9.5E-06 |
| Krtcap2 | 0.350484 | 7.63E-08 | 0.002464 |
| Ltb4r1 | 0.349932 | 9.24E-07 | 0.029843 |
| Snrpd2 | 0.349453 | 3.08E-10 | 9.95E-06 |
| Hvcn1 | 0.349362 | 2.2E-13 | 7.1E-09 |
| Nrg2 | 0.348996 | 8.48E-14 | 2.74E-09 |
| Flt1 | 0.348099 | 6.23E-07 | 0.02011 |
| Mrpl13 | 0.347341 | 1.34E-13 | 4.34E-09 |
| Mrps28 | 0.34716 | 4.05E-13 | 1.31E-08 |
| Tbc1d2 | 0.346627 | 1.14E-13 | 3.68E-09 |
| Trem2 | 0.346148 | 4.33E-10 | 1.4E-05 |
| Pdcd6 | 0.345778 | 2.89E-10 | 9.34E-06 |
| Dnajc15 | 0.345106 | 3E-08 | 0.000968 |
| Sectm1a | 0.344769 | 1.43E-17 | 4.61E-13 |
| Rab20 | 0.343807 | 9.46E-11 | 3.05E-06 |
| Got2 | 0.343807 | 8.72E-07 | 0.028151 |
| Stx3 | 0.34359 | 1.72E-15 | 5.56E-11 |
| Gm10260 | 0.343359 | 3.27E-08 | 0.001057 |
| Tomm20 | 0.343035 | 6.98E-08 | 0.002254 |
| Fbxo6 | 0.342751 | 1.48E-11 | 4.76E-07 |
| Nfia | 0.342506 | 2.15E-10 | 6.94E-06 |

| | | | |
|---|---|---|---|
| C1rl | 0.342252 | 1E-13 | 3.23E-09 |
| Lrp12 | 0.342216 | 1.31E-15 | 4.23E-11 |
| Rbpj | 0.341154 | 3.51E-08 | 0.001132 |
| Uap1l1 | 0.341103 | 8.47E-15 | 2.73E-10 |
| Psma4 | 0.34108 | 2.96E-08 | 0.000956 |
| AI506816 | 0.340456 | 9.11E-11 | 2.94E-06 |
| Npm1 | 0.340287 | 1.38E-05 | 0.446962 |
| Mpv17l2 | 0.340235 | 3.95E-12 | 1.27E-07 |
| Mrpl12 | 0.338632 | 3.35E-10 | 1.08E-05 |
| Rnh1 | 0.337095 | 1.23E-07 | 0.003956 |
| Ski | 0.337068 | 6.17E-12 | 1.99E-07 |
| Hcar2 | 0.33683 | 2.73E-11 | 8.83E-07 |
| Tnfaip3 | 0.336319 | 3.95E-07 | 0.012749 |
| Ranbp1 | 0.33591 | 1.7E-07 | 0.005494 |
| Hspe1 | 0.335848 | 1.58E-06 | 0.05099 |
| Ms4a6d | 0.335712 | 1.17E-07 | 0.003774 |
| Psmb2 | 0.335576 | 1.79E-08 | 0.000577 |
| Plgrkt | 0.334249 | 5E-06 | 0.161542 |
| Psmb6 | 0.332601 | 1.63E-08 | 0.000528 |
| Cope | 0.332067 | 8.17E-07 | 0.026373 |
| Eno1b | 0.331909 | 3.47E-09 | 0.000112 |
| Ly6c2 | 0.331318 | 1.03E-06 | 0.033143 |
| Mrpl42 | 0.331049 | 1.14E-09 | 3.68E-05 |
| Tmed2 | 0.329849 | 8.07E-08 | 0.002606 |
| Olfm1 | 0.329314 | 3.67E-12 | 1.18E-07 |
| Cpq | 0.329171 | 3.15E-09 | 0.000102 |
| Rpl23a | 0.32884 | 2.9E-07 | 0.009357 |
| Il1b | 0.328159 | 0.270517 | 1 |
| Nudt19 | 0.327899 | 8.82E-10 | 2.85E-05 |
| Fuca1 | 0.327274 | 2.03E-06 | 0.065646 |
| Rpl29 | 0.325392 | 5.74E-10 | 1.85E-05 |
| Tmem14c | 0.325388 | 4.1E-08 | 0.001323 |
| Cct6a | 0.324904 | 8.22E-07 | 0.026532 |
| Gnaq | 0.324333 | 2.23E-07 | 0.007199 |
| Psmb4 | 0.324131 | 6.02E-09 | 0.000194 |
| Cenpb | 0.322968 | 4.43E-12 | 1.43E-07 |
| Selenom | 0.322821 | 1E-07 | 0.00324 |
| Ddt | 0.321516 | 1.25E-10 | 4.05E-06 |
| Pla2g7 | 0.319638 | 0.00038 | 1 |
| Rpl13 | 0.319373 | 1.73E-10 | 5.58E-06 |
| Cd200r4 | 0.319085 | 6.67E-13 | 2.15E-08 |
| Ndufa8 | 0.317572 | 3.09E-07 | 0.009964 |
| Eva1b | 0.317539 | 2.31E-14 | 7.46E-10 |
| Atp5f1 | 0.316593 | 5.29E-07 | 0.017069 |
| Gde1 | 0.316259 | 1.99E-09 | 6.42E-05 |
| Rpl22l1 | 0.315446 | 4.88E-05 | 1 |
| Hk1 | 0.314408 | 1.98E-06 | 0.063783 |
| Tgm2 | 0.313266 | 0.024184 | 1 |

| | | | |
|---|---|---|---|
| Cd33 | 0.31296 | 6.31E-17 | 2.04E-12 |
| Def8 | 0.312218 | 2.77E-09 | 8.95E-05 |
| Fam162a | 0.311802 | 1.12E-09 | 3.62E-05 |
| Fcgr1 | 0.311207 | 1.05E-08 | 0.000339 |
| Adam8 | 0.310694 | 1.06E-06 | 0.034136 |
| Cebpa | 0.310667 | 8.43E-08 | 0.002722 |
| Clec4a2 | 0.30915 | 4.02E-05 | 1 |
| Calhm6 | 0.307838 | 7.23E-06 | 0.23357 |
| Hscb | 0.307482 | 3.21E-07 | 0.010352 |
| Prdx4 | 0.307404 | 1.64E-10 | 5.29E-06 |
| Rpl31 | 0.306751 | 4.9E-07 | 0.015811 |
| Phf11b | 0.306356 | 1.26E-05 | 0.40673 |
| Ly86 | 0.306296 | 0.000122 | 1 |
| Pitpna | 0.306002 | 3.56E-07 | 0.011487 |
| Zfp804a | 0.305983 | 1.66E-06 | 0.053567 |
| Slc29a3 | 0.305792 | 2.47E-10 | 7.97E-06 |
| Sash1 | 0.303695 | 1.59E-08 | 0.000513 |
| Lgals3 | 0.302511 | 7.16E-06 | 0.231233 |
| Hipk2 | 0.302269 | 8.74E-07 | 0.028215 |
| Nenf | 0.301804 | 6.13E-14 | 1.98E-09 |
| Snx10 | 0.301656 | 1.79E-06 | 0.05793 |
| Adap2 | 0.301524 | 3.47E-12 | 1.12E-07 |
| Cited2 | 0.301329 | 3.32E-06 | 0.107102 |
| Atp6v1g1 | 0.300841 | 5.08E-07 | 0.016392 |
| Vcp | 0.30075 | 3.72E-05 | 1 |
| Tapbp | 0.299708 | 2.63E-05 | 0.850149 |
| Pgd | 0.298826 | 4.3E-07 | 0.013896 |
| Rplp2 | 0.298139 | 3.66E-10 | 1.18E-05 |
| Dnase2a | 0.297865 | 3.29E-07 | 0.01062 |
| Lrpap1 | 0.297564 | 6.75E-07 | 0.02178 |
| Irgm1 | 0.29697 | 1.13E-05 | 0.363824 |
| Cox7a2l | 0.29693 | 0.000466 | 1 |
| Snrpc | 0.296792 | 1.13E-06 | 0.036501 |
| Gsn | 0.296748 | 4.02E-05 | 1 |
| Cmklr1 | 0.296691 | 2.85E-10 | 9.2E-06 |
| Lamtor1 | 0.295373 | 0.000156 | 1 |
| Ndst1 | 0.295069 | 4.83E-11 | 1.56E-06 |
| Tmem256 | 0.294446 | 1.27E-06 | 0.041032 |
| Ptafr | 0.294361 | 1.92E-09 | 6.18E-05 |
| Fam20c | 0.294181 | 8.28E-11 | 2.67E-06 |
| Hspa5 | 0.293924 | 0.000941 | 1 |
| Pvt1 | 0.293018 | 4.69E-10 | 1.51E-05 |
| Nrip1 | 0.292875 | 1.1E-06 | 0.035612 |
| Arf3 | 0.292818 | 1.26E-07 | 0.004082 |
| Aco2 | 0.292233 | 6.33E-09 | 0.000204 |
| Rps26 | 0.291787 | 1.95E-07 | 0.006303 |
| S100a6 | 0.291483 | 0.000119 | 1 |
| Junos | 0.291417 | 3.61E-08 | 0.001167 |

| | | | |
|---|---|---|---|
| Basp1 | 0.291206 | 4.49E-05 | 1 |
| Stx7 | 0.291072 | 1.37E-05 | 0.442586 |
| Rpn2 | 0.290699 | 1.1E-06 | 0.035355 |
| Pgls | 0.290454 | 6.42E-05 | 1 |
| Mtln | 0.288536 | 1.07E-15 | 3.46E-11 |
| Cacybp | 0.288522 | 1.64E-07 | 0.005293 |
| Slc25a25 | 0.287553 | 2.21E-09 | 7.14E-05 |
| 1110038F1 | 0.28731 | 2.41E-09 | 7.79E-05 |
| Ubald1 | 0.286808 | 0.000384 | 1 |
| Hint1 | 0.28674 | 1.04E-05 | 0.335264 |
| Eef1g | 0.286507 | 5.82E-06 | 0.188004 |
| Txn2 | 0.286436 | 1.88E-07 | 0.00608 |
| Rps12 | 0.286241 | 7.31E-07 | 0.023585 |
| Psmd7 | 0.28542 | 1.56E-09 | 5.03E-05 |
| Sdhc | 0.283901 | 8.72E-12 | 2.82E-07 |
| Il12rb1 | 0.283641 | 2.8E-17 | 9.03E-13 |
| Sdad1 | 0.282985 | 6E-07 | 0.019384 |
| Timm13 | 0.282814 | 4.32E-07 | 0.013945 |
| Ncl | 0.282728 | 4.31E-05 | 1 |
| Ndufv2 | 0.282523 | 4.96E-05 | 1 |
| Rpl3 | 0.281949 | 3.16E-07 | 0.010208 |
| Ifit2 | 0.281561 | 0.030914 | 1 |
| Fbl | 0.281516 | 1.11E-06 | 0.035818 |
| Rnf217 | 0.2806 | 9.45E-13 | 3.05E-08 |
| Ntan1 | 0.280552 | 2.85E-07 | 0.00921 |
| Tomm40 | 0.28035 | 8.89E-10 | 2.87E-05 |
| Vps54 | 0.28009 | 1.32E-07 | 0.004263 |
| Parvg | 0.279385 | 3.45E-08 | 0.001115 |
| Itga5 | 0.279132 | 2.76E-07 | 0.008895 |
| Eif4a1 | 0.278944 | 0.000212 | 1 |
| Btf3 | 0.278917 | 1.17E-06 | 0.037844 |
| Nus1 | 0.278688 | 1.05E-08 | 0.000338 |
| Antxr2 | 0.278425 | 3.7E-06 | 0.119383 |
| Dnase1l3 | 0.278231 | 1.99E-06 | 0.064162 |
| Rwdd1 | 0.277952 | 2.61E-06 | 0.084249 |
| Il2rg | 0.277573 | 0.001203 | 1 |
| Tbcb | 0.277436 | 5.33E-05 | 1 |
| H2-T22 | 0.276161 | 0.000568 | 1 |
| Slc4a8 | 0.276159 | 3.74E-15 | 1.21E-10 |
| Slfn4 | 0.275126 | 7.51E-05 | 1 |
| Vcan | 0.27474 | 0.000404 | 1 |
| Reep3 | 0.274347 | 2.48E-06 | 0.080179 |
| Bcl2l11 | 0.274267 | 3E-09 | 9.7E-05 |
| Rtraf | 0.273121 | 5.22E-06 | 0.168526 |
| Odc1 | 0.273019 | 2.79E-13 | 9.01E-09 |
| Sidt2 | 0.272658 | 3.97E-11 | 1.28E-06 |
| Eif1a | 0.272377 | 3.97E-07 | 0.012825 |
| Chchd1 | 0.272193 | 8.04E-05 | 1 |

| | | | |
|---|---|---|---|
| Pclaf | 0.272116 | 5.74E-05 | 1 |
| Serp1 | 0.272063 | 0.000704 | 1 |
| Ptma | 0.271933 | 0.00015 | 1 |
| Rps25 | 0.271853 | 4.32E-07 | 0.013934 |
| Dst | 0.271701 | 1.11E-05 | 0.358197 |
| Abhd16a | 0.271355 | 1.84E-08 | 0.000594 |
| Cemip2 | 0.271352 | 9.44E-10 | 3.05E-05 |
| Mkrn1 | 0.270217 | 5.27E-05 | 1 |
| Mxd1 | 0.269748 | 2.26E-06 | 0.073028 |
| Mtx1 | 0.269727 | 3.06E-07 | 0.009893 |
| Ndufv1 | 0.269122 | 3.91E-08 | 0.001262 |
| Cotl1 | 0.269109 | 8.66E-05 | 1 |
| Tmem86a | 0.268847 | 1.13E-07 | 0.003642 |
| Tspan3 | 0.268736 | 7.05E-11 | 2.28E-06 |
| Hdlbp | 0.26864 | 3.48E-05 | 1 |
| Snx24 | 0.268495 | 5.25E-15 | 1.7E-10 |
| Aldoc | 0.268179 | 2.82E-11 | 9.12E-07 |
| mt-Nd2 | 0.268152 | 2.59E-05 | 0.837504 |
| Mfsd1 | 0.268125 | 6.4E-06 | 0.206516 |
| Ddost | 0.267922 | 5.45E-08 | 0.001759 |
| Fundc2 | 0.267858 | 1.38E-10 | 4.46E-06 |
| Tmem123 | 0.267731 | 3.71E-05 | 1 |
| Chd7 | 0.267642 | 5.54E-05 | 1 |
| Tap2 | 0.267454 | 0.000179 | 1 |
| Rai14 | 0.267362 | 1.59E-10 | 5.13E-06 |
| Eif6 | 0.267213 | 1.05E-06 | 0.033993 |
| Cerk | 0.267007 | 1.72E-06 | 0.055549 |
| Mrap | 0.266282 | 6.04E-08 | 0.001951 |
| D5Ertd579 | 0.26618 | 1.92E-06 | 0.062101 |
| AI413582 | 0.266176 | 0.00034 | 1 |
| Psme1 | 0.266108 | 2.04E-06 | 0.065705 |
| Junb | 0.265603 | 0.004478 | 1 |
| Net1 | 0.26545 | 1.27E-07 | 0.004099 |
| Slc52a3 | 0.265238 | 9.44E-12 | 3.05E-07 |
| Ap1s1 | 0.265225 | 2.99E-07 | 0.009653 |
| Larp1 | 0.26439 | 2.35E-07 | 0.007595 |
| Tex264 | 0.264094 | 4.01E-11 | 1.29E-06 |
| Aff1 | 0.263565 | 5.61E-08 | 0.001812 |
| Cd80 | 0.263527 | 2.82E-09 | 9.1E-05 |
| Tubb6 | 0.263207 | 4.89E-11 | 1.58E-06 |
| Klrk1 | 0.263146 | 6.51E-06 | 0.210192 |
| Slc31a1 | 0.26291 | 2.24E-07 | 0.007246 |
| Atp5j2 | 0.262712 | 6.47E-05 | 1 |
| Rps15 | 0.262655 | 8.46E-06 | 0.273094 |
| Ube2j2 | 0.262429 | 7.38E-06 | 0.238184 |
| Batf2 | 0.261999 | 2.94E-10 | 9.5E-06 |
| Psma2 | 0.261934 | 4.69E-05 | 1 |
| Rab5if | 0.261901 | 8.2E-08 | 0.002647 |

| | | | |
|---|---|---|---|
| Akap7 | 0.26089 | 6.94E-07 | 0.02242 |
| Etfb | 0.26045 | 7.41E-05 | 1 |
| Hk2 | 0.260406 | 1.56E-06 | 0.050468 |
| Ripk3 | 0.260166 | 8.63E-11 | 2.79E-06 |
| Mafb | 0.260041 | 0.000868 | 1 |
| Uqcc2 | 0.26004 | 2.51E-06 | 0.081094 |
| Txndc5 | 0.259622 | 1.46E-09 | 4.7E-05 |
| Xbp1 | 0.259612 | 0.000114 | 1 |
| Mrpl2 | 0.259011 | 6.05E-05 | 1 |
| mt-Nd4 | 0.259001 | 0.000177 | 1 |
| Nutf2 | 0.258424 | 4.37E-09 | 0.000141 |
| Tspan4 | 0.258378 | 9.77E-13 | 3.16E-08 |
| Pim3 | 0.258043 | 1.98E-09 | 6.39E-05 |
| Echs1 | 0.256359 | 3E-07 | 0.009686 |
| Tmem147 | 0.255994 | 2.9E-06 | 0.093782 |
| Tap1 | 0.25581 | 0.000196 | 1 |
| Eif3c | 0.255598 | 0.000569 | 1 |
| Gadd45g | 0.255512 | 1.43E-07 | 0.004619 |
| Csnk2b | 0.255384 | 0.000105 | 1 |
| Rpl23 | 0.255191 | 1.21E-06 | 0.038987 |
| Rpn1 | 0.254919 | 5.29E-06 | 0.170898 |
| Timm10 | 0.254832 | 9.7E-09 | 0.000313 |
| Itgam | 0.254192 | 0.005232 | 1 |
| Gm15283 | 0.253928 | 8.09E-08 | 0.002611 |
| Rpl7a | 0.253589 | 3.67E-07 | 0.011846 |
| Mtch2 | 0.253019 | 1.01E-07 | 0.003258 |
| Timm8b | 0.252495 | 0.000287 | 1 |
| Slc20a1 | 0.251829 | 3.71E-08 | 0.001198 |
| Nceh1 | 0.251628 | 4.66E-09 | 0.000151 |
| Ralgds | 0.251392 | 3.18E-11 | 1.03E-06 |
| Slirp | 0.251062 | 3.86E-06 | 0.124693 |
| Atp5l | 0.250481 | 1.42E-05 | 0.459326 |
| Tle5 | 0.250475 | 0.0001 | 1 |
| Micos13 | 0.250304 | 9.47E-07 | 0.030572 |
| Nolc1 | 0.250292 | 4.25E-05 | 1 |
| Ftl1-ps1 | 0.249988 | 6.04E-08 | 0.00195 |
| Polr2i | 0.249812 | 5.88E-07 | 0.018982 |
| Hnrnpab | 0.249438 | 0.000468 | 1 |
| Gbp9 | 0.24879 | 4.8E-05 | 1 |
| 1700017B( | 0.248286 | 1.1E-10 | 3.56E-06 |
| Rtp4 | 0.247928 | 2.29E-06 | 0.073914 |
| Syngr2 | 0.247765 | 0.003961 | 1 |
| Clta | 0.247705 | 7.06E-06 | 0.227876 |
| Ybx1 | 0.247514 | 7.75E-05 | 1 |
| Drap1 | 0.247474 | 2.02E-06 | 0.065181 |
| Rgcc | 0.247317 | 7.45E-08 | 0.002405 |
| Nudt21 | 0.247209 | 5.44E-06 | 0.175514 |
| Tnfaip2 | 0.247058 | 0.000182 | 1 |

| | | | |
|---|---|---|---|
| Prtn3 | 0.246871 | 3.64E-08 | 0.001175 |
| Ksr1 | 0.246714 | 6.61E-09 | 0.000213 |
| Myc | 0.246087 | 7.61E-06 | 0.245763 |
| Fkbp1a | 0.245528 | 4.21E-05 | 1 |
| Cass4 | 0.245467 | 0.000304 | 1 |
| Klrb1b | 0.24546 | 2.1E-06 | 0.067774 |
| Pdia4 | 0.24545 | 2.53E-10 | 8.15E-06 |
| Htt | 0.245133 | 1.63E-07 | 0.005253 |
| Comt | 0.245015 | 5.33E-07 | 0.01722 |
| Ncf1 | 0.244648 | 3.48E-05 | 1 |
| Slc25a39 | 0.244584 | 5.29E-07 | 0.017068 |
| Mrc1 | 0.244209 | 2.58E-11 | 8.34E-07 |
| Prkca | 0.243959 | 3.52E-07 | 0.01138 |
| Thyn1 | 0.243788 | 1.14E-12 | 3.69E-08 |
| Aifm2 | 0.243295 | 1.01E-10 | 3.26E-06 |
| Abhd17a | 0.243256 | 2.05E-05 | 0.663121 |
| Smarcb1 | 0.243047 | 5.55E-07 | 0.017925 |
| Cfl1 | 0.242919 | 1.3E-06 | 0.041855 |
| Ncln | 0.242366 | 8.67E-07 | 0.027993 |
| Gabarap | 0.241994 | 2.16E-07 | 0.006978 |
| Gusb | 0.241939 | 0.000139 | 1 |
| Banf1 | 0.241364 | 2.71E-05 | 0.874577 |
| Phb | 0.241338 | 4.23E-07 | 0.013641 |
| Svbp | 0.241245 | 1.18E-07 | 0.0038 |
| Arf5 | 0.240734 | 4.3E-06 | 0.138724 |
| Tbca | 0.240679 | 5.21E-05 | 1 |
| Ndufa11 | 0.240409 | 1.13E-05 | 0.365805 |
| Hspd1 | 0.240015 | 3.24E-05 | 1 |
| Dapk1 | 0.23991 | 6.35E-05 | 1 |
| Snrpd3 | 0.239606 | 1.31E-05 | 0.422388 |
| Atp5h | 0.239529 | 8.47E-06 | 0.273557 |
| 4930430E1 | 0.239385 | 2.99E-13 | 9.65E-09 |
| Tmem189 | 0.239384 | 1.01E-08 | 0.000327 |
| Gm50237 | 0.239296 | 2.62E-13 | 8.47E-09 |
| Tor3a | 0.239023 | 1.57E-05 | 0.506139 |
| Hnrnpa1 | 0.238969 | 0.000209 | 1 |
| Eif4ebp1 | 0.23895 | 0.005636 | 1 |
| Rgl1 | 0.238825 | 0.000104 | 1 |
| Ndufb10 | 0.238704 | 3.48E-05 | 1 |
| Mrpl20 | 0.238517 | 0.000186 | 1 |
| Rap2c | 0.238431 | 7.72E-05 | 1 |
| Pgm1 | 0.238339 | 8.71E-10 | 2.81E-05 |
| Stx4a | 0.238014 | 0.000119 | 1 |
| Ppil1 | 0.237579 | 0.000859 | 1 |
| Rgs1 | 0.237188 | 2.43E-06 | 0.078372 |
| Ric1 | 0.237028 | 1.49E-08 | 0.000483 |
| H2afy | 0.23682 | 0.000179 | 1 |
| Mrps33 | 0.236756 | 1.12E-05 | 0.361435 |

| | | | |
|---|---|---|---|
| Cfap410 | 0.236737 | 2.87E-07 | 0.009268 |
| Eef1d | 0.236458 | 0.000107 | 1 |
| Tmem192 | 0.236437 | 1.48E-06 | 0.047711 |
| Spcs1 | 0.235854 | 3.36E-05 | 1 |
| Mrpl11 | 0.235537 | 1.71E-07 | 0.005525 |
| Snap29 | 0.235391 | 1.76E-05 | 0.567447 |
| Rtcb | 0.235366 | 0.000159 | 1 |
| Pnrc1 | 0.235251 | 0.000436 | 1 |
| Nubp1 | 0.235159 | 1.23E-05 | 0.395772 |
| Sdc3 | 0.234885 | 1.42E-05 | 0.457522 |
| Mrps16 | 0.234826 | 1.15E-06 | 0.036989 |
| Il15ra | 0.234729 | 5.87E-08 | 0.001894 |
| Arhgap31 | 0.234555 | 3.88E-05 | 1 |
| Plod1 | 0.234111 | 9.84E-09 | 0.000318 |
| Atg5 | 0.234084 | 1.6E-06 | 0.051714 |
| Ranbp2 | 0.233696 | 5.49E-05 | 1 |
| Renbp | 0.233694 | 2.93E-10 | 9.46E-06 |
| Tlr12 | 0.233013 | 4E-12 | 1.29E-07 |
| Gnl3 | 0.232785 | 1.3E-09 | 4.2E-05 |
| Cacna1d | 0.23277 | 0.000131 | 1 |
| Creld2 | 0.231819 | 1.7E-09 | 5.5E-05 |
| Vcam1 | 0.231665 | 4.01E-06 | 0.129388 |
| Slc16a6 | 0.231547 | 2.53E-10 | 8.18E-06 |
| Yif1b | 0.231246 | 5.81E-09 | 0.000188 |
| Pik3r6 | 0.230867 | 4.55E-06 | 0.14681 |
| Cct8 | 0.230636 | 0.0008 | 1 |
| Sulf2 | 0.23045 | 1.4E-05 | 0.451088 |
| Apex1 | 0.230246 | 4.95E-09 | 0.00016 |
| Fbxw17 | 0.229941 | 1.55E-09 | 5E-05 |
| Psmd8 | 0.229917 | 0.00139 | 1 |
| Ndufc2 | 0.229854 | 5.35E-07 | 0.017262 |
| Edf1 | 0.229794 | 0.000169 | 1 |
| Impa2 | 0.229783 | 1.99E-08 | 0.000641 |
| Mrpl54 | 0.229677 | 4.08E-07 | 0.013172 |
| Coro2a | 0.229607 | 0.000396 | 1 |
| Trim25 | 0.229575 | 0.002706 | 1 |
| mt-Nd3 | 0.229387 | 7.42E-05 | 1 |
| Sys1 | 0.229382 | 0.000114 | 1 |
| Bri3 | 0.22936 | 1.51E-07 | 0.004889 |
| Bcat2 | 0.228614 | 4.03E-08 | 0.0013 |
| Peak1 | 0.228588 | 2E-05 | 0.645478 |
| Pip5k1c | 0.228481 | 2.58E-06 | 0.083263 |
| Siglec1 | 0.22826 | 3.24E-10 | 1.05E-05 |
| Cdca8 | 0.228184 | 1.87E-07 | 0.006043 |
| Adgb | 0.228132 | 8.29E-07 | 0.026762 |
| Ndufb2 | 0.228125 | 2.51E-05 | 0.809075 |
| Ergic1 | 0.228073 | 7.29E-09 | 0.000235 |
| Dennd1b | 0.228042 | 0.000418 | 1 |

| | | | |
|---|---|---|---|
| Ybx3 | 0.227788 | 1.12E-07 | 0.003605 |
| Cmc1 | 0.227761 | 0.000295 | 1 |
| Plxnd1 | 0.227657 | 1.54E-10 | 4.98E-06 |
| Slc25a4 | 0.227539 | 1.05E-07 | 0.003374 |
| Fcgrt | 0.227523 | 1.55E-06 | 0.049971 |
| Gpx3 | 0.227237 | 4.36E-08 | 0.001408 |
| Rpl4 | 0.227223 | 1.24E-05 | 0.399754 |
| Icam1 | 0.227104 | 0.005487 | 1 |
| Galnt7 | 0.227083 | 2.02E-09 | 6.51E-05 |
| Rab7 | 0.227041 | 0.000145 | 1 |
| Plekho1 | 0.22608 | 1.7E-08 | 0.000549 |
| Iscu | 0.226024 | 0.015944 | 1 |
| Lrrc58 | 0.225749 | 6.11E-06 | 0.19717 |
| Psmb3 | 0.225648 | 8.3E-05 | 1 |
| Anapc11 | 0.225599 | 0.000322 | 1 |
| Actg1 | 0.225095 | 4.71E-07 | 0.015192 |
| Mapk6 | 0.225074 | 7.29E-07 | 0.023521 |
| Itgb1bp1 | 0.225025 | 1.34E-07 | 0.004322 |
| Cenpa | 0.224957 | 0.006964 | 1 |
| Rps8 | 0.224935 | 6.68E-05 | 1 |
| Eef1b2 | 0.224505 | 0.000164 | 1 |
| 9130401M | 0.224469 | 2.89E-05 | 0.93364 |
| Ubb | 0.224403 | 1.37E-06 | 0.044324 |
| Herc6 | 0.224387 | 0.000352 | 1 |
| Rmdn3 | 0.224191 | 1.02E-09 | 3.29E-05 |
| Surf4 | 0.224028 | 1.02E-05 | 0.330135 |
| Emg1 | 0.223999 | 1.34E-06 | 0.043225 |
| Hivep3 | 0.22398 | 4.83E-06 | 0.155824 |
| A930007I1 | 0.223977 | 8.53E-09 | 0.000275 |
| Mlkl | 0.223281 | 1.39E-08 | 0.000449 |
| Rpl27 | 0.22323 | 3E-06 | 0.096702 |
| Slc16a3 | 0.22271 | 0.002674 | 1 |
| Spcs2 | 0.222703 | 0.000443 | 1 |
| Mrpl35 | 0.222478 | 2.18E-09 | 7.05E-05 |
| Hist1h1e | 0.222152 | 0.065817 | 1 |
| Ndufa2 | 0.222093 | 0.000319 | 1 |
| Kif1b | 0.221901 | 4.63E-06 | 0.149501 |
| Fdx2 | 0.221824 | 1.23E-07 | 0.003968 |
| 1700037H | 0.221364 | 2.85E-05 | 0.920396 |
| Abhd11 | 0.221221 | 1.1E-07 | 0.003564 |
| Suclg1 | 0.221169 | 1.86E-06 | 0.059921 |
| Rpl17 | 0.221034 | 0.0004 | 1 |
| Scpep1 | 0.221028 | 3.94E-05 | 1 |
| Snu13 | 0.220895 | 8.09E-05 | 1 |
| Dut | 0.220887 | 8.05E-06 | 0.259975 |
| Nudcd2 | 0.220878 | 1.45E-06 | 0.046737 |
| Pfdn1 | 0.220813 | 1.45E-05 | 0.466829 |
| Nagk | 0.220484 | 5.02E-07 | 0.016202 |

| | | | |
|---|---|---|---|
| Ttc39c | 0.220445 | 1.04E-06 | 0.033483 |
| Batf3 | 0.220407 | 2.32E-05 | 0.750374 |
| Mrps23 | 0.220167 | 1.45E-07 | 0.004684 |
| Rabggtb | 0.219694 | 5.86E-05 | 1 |
| Birc5 | 0.219625 | 3.71E-05 | 1 |
| Zdhhc4 | 0.219507 | 0.000293 | 1 |
| Hsp90ab1 | 0.219483 | 2.4E-05 | 0.774964 |
| Cd302 | 0.218934 | 0.000115 | 1 |
| Rpl24 | 0.218629 | 0.000154 | 1 |
| Fkbp1b | 0.218537 | 0.001936 | 1 |
| Heatr1 | 0.218012 | 6.49E-09 | 0.000209 |
| Bsg | 0.217307 | 1.41E-05 | 0.45439 |
| Ifi203 | 0.217266 | 4.53E-06 | 0.146202 |
| Tmed10 | 0.217145 | 2.5E-05 | 0.807863 |
| Anapc15 | 0.216585 | 2.4E-08 | 0.000774 |
| Nudt3 | 0.216169 | 1.15E-08 | 0.000371 |
| Cpne2 | 0.216 | 1.31E-12 | 4.22E-08 |
| Tmem140 | 0.215796 | 0.000306 | 1 |
| Desi2 | 0.215686 | 2.02E-07 | 0.006534 |
| Oxct1 | 0.215632 | 6.14E-06 | 0.198173 |
| Il12rb2 | 0.215474 | 1.49E-06 | 0.048194 |
| Il1rl2 | 0.215298 | 1.52E-11 | 4.9E-07 |
| Mdfic | 0.215132 | 3.51E-07 | 0.011316 |
| Cd151 | 0.215092 | 8.35E-07 | 0.02695 |
| Mdm2 | 0.214952 | 0.001521 | 1 |
| Prdx1 | 0.214828 | 1.21E-05 | 0.390619 |
| Smc2 | 0.214688 | 2.28E-05 | 0.7372 |
| Fmn1 | 0.214265 | 7.82E-07 | 0.025242 |
| Dmxl2 | 0.214157 | 2.11E-08 | 0.000682 |
| Cnih1 | 0.214008 | 2.68E-06 | 0.086408 |
| Esyt2 | 0.213665 | 4.6E-06 | 0.148527 |
| Rpl28 | 0.213271 | 6.56E-05 | 1 |
| Vat1 | 0.213218 | 5.42E-07 | 0.017502 |
| Gm12865 | 0.213216 | 6.23E-10 | 2.01E-05 |
| Pmvk | 0.212944 | 5.83E-08 | 0.001881 |
| Esd | 0.212664 | 1.46E-05 | 0.470954 |
| Usp14 | 0.212651 | 4.91E-07 | 0.015865 |
| Mroh1 | 0.212238 | 1.73E-08 | 0.000558 |
| Cd93 | 0.212235 | 0.000434 | 1 |
| Hadh | 0.212219 | 4.88E-07 | 0.015743 |
| Kars | 0.212209 | 1.21E-07 | 0.003894 |
| Spg21 | 0.211706 | 2.75E-05 | 0.887935 |
| Rps5 | 0.211589 | 0.000164 | 1 |
| Aph1a | 0.211389 | 0.000174 | 1 |
| Stard3nl | 0.210884 | 1.25E-07 | 0.004024 |
| Cct3 | 0.210754 | 1.5E-05 | 0.485607 |
| Adam9 | 0.210684 | 3.57E-06 | 0.115144 |
| Rab5c | 0.210391 | 8.31E-05 | 1 |

| | | | |
|---|---|---|---|
| Sec13 | 0.210306 | 0.000495 | 1 |
| Prpf31 | 0.210282 | 6.83E-08 | 0.002206 |
| Wwox | 0.210091 | 2.91E-05 | 0.938418 |
| Aig1 | 0.210081 | 4.02E-05 | 1 |
| Rad23a | 0.209873 | 1.1E-05 | 0.356499 |
| Osm | 0.209315 | 7.15E-08 | 0.00231 |
| Mid1 | 0.209236 | 2.87E-08 | 0.000926 |
| Psmd2 | 0.209211 | 0.001492 | 1 |
| 1110038B1 | 0.208935 | 2.18E-06 | 0.070383 |
| Abracl | 0.208869 | 0.001421 | 1 |
| Lsm7 | 0.208743 | 5.91E-05 | 1 |
| Plxna1 | 0.208412 | 1.93E-10 | 6.23E-06 |
| Pla2g4a | 0.20834 | 0.002646 | 1 |
| Myo7a | 0.20822 | 4.83E-07 | 0.015581 |
| Gfer | 0.207781 | 2.45E-07 | 0.007924 |
| Mrps18a | 0.207434 | 0.000124 | 1 |
| Nup93 | 0.207379 | 8.44E-07 | 0.027262 |
| Usp18 | 0.207204 | 2.61E-05 | 0.843723 |
| Ptpn7 | 0.207029 | 5.82E-07 | 0.018788 |
| Eif2b2 | 0.206917 | 0.000129 | 1 |
| Tcp11l1 | 0.206739 | 3.19E-07 | 0.010288 |
| Paics | 0.206668 | 7.41E-06 | 0.239102 |
| Gas2l1 | 0.206165 | 8.95E-10 | 2.89E-05 |
| Tusc3 | 0.205725 | 1.66E-07 | 0.00536 |
| Mvb12b | 0.205566 | 3.7E-08 | 0.001195 |
| Cdk4 | 0.205207 | 1.4E-05 | 0.453571 |
| Egr2 | 0.205122 | 2.56E-07 | 0.008255 |
| Fuom | 0.204935 | 9.18E-10 | 2.96E-05 |
| Khk | 0.204618 | 8.57E-06 | 0.276545 |
| Wdr83os | 0.204307 | 9.64E-05 | 1 |
| Cks1b | 0.203928 | 4.64E-05 | 1 |
| Gars | 0.203486 | 8.42E-05 | 1 |
| Cenpw | 0.203397 | 6.33E-08 | 0.002044 |
| Tes | 0.20324 | 2.91E-05 | 0.939403 |
| Gpr171 | 0.203161 | 2.18E-09 | 7.04E-05 |
| Lamtor2 | 0.203127 | 0.002825 | 1 |
| Ndufs7 | 0.202888 | 1.81E-05 | 0.585709 |
| Uqcrfs1 | 0.202846 | 1.67E-05 | 0.538668 |
| Rnf141 | 0.202809 | 6.52E-08 | 0.002106 |
| Uqcrc2 | 0.202676 | 0.012958 | 1 |
| Ehd1 | 0.202553 | 0.003286 | 1 |
| Taf10 | 0.20235 | 8.31E-06 | 0.268378 |
| Rplp0 | 0.202171 | 8.61E-05 | 1 |
| Lrpprc | 0.201897 | 3.61E-06 | 0.116611 |
| Pkig | 0.201859 | 6.31E-07 | 0.020365 |
| Nrg1 | 0.201701 | 1.55E-07 | 0.004989 |
| Mycbp2 | 0.201562 | 0.002509 | 1 |
| Abcg1 | 0.201009 | 3.13E-05 | 1 |

| | | | |
|---|---|---|---|
| Rps7 | 0.200986 | 0.000313 | 1 |
| Dnajb11 | 0.200984 | 1.44E-05 | 0.466459 |
| Impdh1 | 0.200901 | 3.35E-08 | 0.001081 |
| Jdp2 | 0.200891 | 1.89E-08 | 0.00061 |
| Lmnb1 | 0.20087 | 7.99E-05 | 1 |
| Nans | 0.200774 | 7.27E-06 | 0.234685 |
| Rer1 | 0.200605 | 5.51E-05 | 1 |
| Srp9 | 0.200573 | 0.000354 | 1 |
| B3gnt2 | 0.200498 | 4.65E-09 | 0.00015 |
| Spink2 | 0.20034 | 3.27E-07 | 0.010562 |
| Rgmb | 0.200338 | 2.22E-06 | 0.07164 |
| Naxe | 0.200267 | 0.000221 | 1 |
| Acat1 | 0.200207 | 2.09E-06 | 0.067623 |
| Nsd2 | 0.200156 | 1.5E-10 | 4.85E-06 |
| Dnpep | 0.200143 | 5.01E-07 | 0.01618 |
| Tnfsf12 | 0.200011 | 3.99E-05 | 1 |
| Znrd2 | 0.199368 | 5.98E-09 | 0.000193 |
| Cyb5r3 | 0.199141 | 3.44E-06 | 0.111112 |
| Tmod3 | 0.198291 | 0.000284 | 1 |
| Plau | 0.197988 | 5.27E-09 | 0.00017 |
| Psmg4 | 0.197762 | 1.07E-05 | 0.346235 |
| Ndufs6 | 0.197753 | 8.84E-05 | 1 |
| Metap2 | 0.197501 | 9.13E-07 | 0.029482 |
| Inpp5j | 0.197034 | 1.79E-08 | 0.000579 |
| Naa25 | 0.196964 | 2.11E-06 | 0.068239 |
| Plekha1 | 0.196927 | 2.29E-07 | 0.007392 |
| Pgap2 | 0.196327 | 0.004757 | 1 |
| Timp2 | 0.195925 | 6.08E-07 | 0.01962 |
| Sestd1 | 0.195883 | 3.18E-08 | 0.001026 |
| Atg3 | 0.195869 | 6.49E-06 | 0.209414 |
| Sra1 | 0.195844 | 0.000928 | 1 |
| 9430038I0 | 0.195773 | 9.68E-08 | 0.003125 |
| Mrpl38 | 0.19521 | 5.63E-07 | 0.018172 |
| Hmbs | 0.195098 | 4.24E-05 | 1 |
| Gmnn | 0.194852 | 1.93E-06 | 0.062417 |
| Atp5md | 0.194731 | 0.0002 | 1 |
| AI662270 | 0.194712 | 0.000181 | 1 |
| Prm1 | 0.194679 | 2.77E-13 | 8.93E-09 |
| Ube2s | 0.194147 | 0.001052 | 1 |
| Tns1 | 0.194083 | 4.45E-07 | 0.014353 |
| Eif3m | 0.19403 | 0.00061 | 1 |
| Set | 0.193993 | 0.000457 | 1 |
| Lman2 | 0.193885 | 0.000938 | 1 |
| Atp13a2 | 0.1938 | 2.52E-06 | 0.081504 |
| Gtf2h5 | 0.193641 | 0.000135 | 1 |
| Rnps1 | 0.193274 | 0.001284 | 1 |
| Kdm3a | 0.193081 | 5.61E-08 | 0.001811 |
| Rnasel | 0.193026 | 2.6E-06 | 0.083877 |

| | | | |
|---|---|---|---|
| Mrps14 | 0.192983 | 0.001325 | 1 |
| Acp1 | 0.192959 | 4.12E-05 | 1 |
| Rps10 | 0.192596 | 0.0001 | 1 |
| Impdh2 | 0.192563 | 7.2E-07 | 0.023232 |
| Rasa4 | 0.19247 | 8.98E-05 | 1 |
| Naa20 | 0.192325 | 4.07E-08 | 0.001315 |
| Eng | 0.192319 | 3.91E-06 | 0.126221 |
| Cnpy2 | 0.192222 | 0.001017 | 1 |
| Pygl | 0.191899 | 0.000458 | 1 |
| Tmbim1 | 0.191558 | 1.78E-05 | 0.573153 |
| Snhg6 | 0.191555 | 2.44E-07 | 0.007865 |
| Slc36a1 | 0.191457 | 0.000111 | 1 |
| Sult1a1 | 0.191243 | 2.9E-05 | 0.937116 |
| 2900097C1 | 0.19124 | 1.76E-06 | 0.056982 |
| Insl6 | 0.191167 | 5.78E-10 | 1.87E-05 |
| Coq7 | 0.190939 | 8.47E-10 | 2.73E-05 |
| Uhrf1bp1 | 0.19093 | 4.27E-09 | 0.000138 |
| Gyg | 0.190879 | 0.003523 | 1 |
| Srpr | 0.190738 | 0.000616 | 1 |
| Ccnb2 | 0.190705 | 2.92E-05 | 0.943933 |
| Smn1 | 0.190408 | 9.95E-07 | 0.032123 |
| Psmb8 | 0.190403 | 2.69E-05 | 0.86995 |
| Lactb | 0.190079 | 7.19E-07 | 0.023221 |
| Psmb1 | 0.190026 | 0.000456 | 1 |
| Exoc3l4 | 0.189787 | 4.87E-12 | 1.57E-07 |
| Canx | 0.189517 | 0.001138 | 1 |
| Bola3 | 0.189427 | 7.93E-08 | 0.002561 |
| Mdn1 | 0.189302 | 1.17E-07 | 0.003783 |
| H2-Q1 | 0.189057 | 9.33E-05 | 1 |
| Gm42962 | 0.188999 | 7.98E-09 | 0.000258 |
| Reep5 | 0.188891 | 0.00446 | 1 |
| Prelid3b | 0.188607 | 3.12E-05 | 1 |
| Atp5e | 0.188508 | 0.000219 | 1 |
| Gm46563 | 0.188458 | 1.38E-08 | 0.000447 |
| Sbf2 | 0.188445 | 3.89E-08 | 0.001255 |
| Gar1 | 0.188331 | 1.04E-06 | 0.033534 |
| Eid1 | 0.188287 | 1.57E-06 | 0.050669 |
| Cpped1 | 0.188139 | 1.03E-07 | 0.003328 |
| Ostc | 0.188032 | 0.001706 | 1 |
| Fgd6 | 0.187792 | 9.24E-14 | 2.98E-09 |
| Cox7c | 0.187778 | 0.001112 | 1 |
| Idh3g | 0.187495 | 6.94E-05 | 1 |
| Noc2l | 0.187447 | 3.61E-05 | 1 |
| Ifi44 | 0.18739 | 1.7E-06 | 0.05479 |
| Galk1 | 0.187247 | 4.59E-09 | 0.000148 |
| Cyth1 | 0.18718 | 0.001463 | 1 |
| Dnajc5 | 0.18714 | 0.000305 | 1 |
| Aarsd1 | 0.187086 | 1.04E-05 | 0.335534 |

| | | | |
|---|---|---|---|
| Slc23a2 | 0.186852 | 6.98E-06 | 0.225212 |
| Rps18 | 0.186613 | 0.00171 | 1 |
| Rhoc | 0.18642 | 1.7E-06 | 0.054937 |
| Marcksl1 | 0.186398 | 0.00026 | 1 |
| Dock1 | 0.186308 | 5.15E-05 | 1 |
| Zfas1 | 0.186078 | 0.000801 | 1 |
| Dohh | 0.186055 | 5.41E-09 | 0.000175 |
| Psmb7 | 0.185986 | 0.000162 | 1 |
| Sec11a | 0.185893 | 2.12E-07 | 0.006847 |
| Baiap2 | 0.185791 | 7.91E-08 | 0.002553 |
| Carhsp1 | 0.185658 | 2.79E-08 | 0.0009 |
| Wdfy1 | 0.185356 | 0.000261 | 1 |
| Parvb | 0.185295 | 6.77E-07 | 0.021868 |
| Eprs | 0.185081 | 3.32E-06 | 0.107204 |
| Sco2 | 0.185058 | 1.58E-12 | 5.09E-08 |
| Commd1 | 0.185002 | 0.000553 | 1 |
| Eif3g | 0.184936 | 0.000698 | 1 |
| Ssna1 | 0.184791 | 0.000396 | 1 |
| Kcnq1ot1 | 0.184722 | 2.88E-06 | 0.092912 |
| Slamf6 | 0.184475 | 1.01E-11 | 3.25E-07 |
| Bmp2k | 0.184469 | 6.08E-08 | 0.001962 |
| Inf2 | 0.184308 | 0.000195 | 1 |
| Tmem268 | 0.184258 | 2E-07 | 0.006459 |
| Rsl1d1 | 0.184178 | 2.21E-06 | 0.071211 |
| Trnau1ap | 0.18415 | 2.98E-06 | 0.09631 |
| Il18rap | 0.184081 | 3.64E-08 | 0.001174 |
| Egr1 | 0.18402 | 0.000146 | 1 |
| Phf11d | 0.183949 | 2.15E-07 | 0.006941 |
| Eif3k | 0.18337 | 0.003918 | 1 |
| Atp5mpl | 0.183295 | 0.000297 | 1 |
| Prkcsh | 0.183204 | 3.67E-05 | 1 |
| Acbd6 | 0.183028 | 1.37E-05 | 0.440712 |
| Tnfrsf14 | 0.182568 | 1.23E-09 | 3.98E-05 |
| Ewsr1 | 0.182445 | 0.000683 | 1 |
| Pira2 | 0.182199 | 4.24E-05 | 1 |
| Tax1bp1 | 0.182132 | 0.006119 | 1 |
| Vkorc1 | 0.182118 | 2.96E-06 | 0.095502 |
| Dhrs3 | 0.182019 | 6.29E-09 | 0.000203 |
| Kctd17 | 0.181275 | 3.89E-09 | 0.000126 |
| Casp9 | 0.181187 | 0.000273 | 1 |
| Ubr4 | 0.180728 | 0.000986 | 1 |
| Eef1a1 | 0.180685 | 8.38E-05 | 1 |
| Rps3 | 0.180503 | 0.000285 | 1 |
| P2rx4 | 0.180275 | 1.9E-06 | 0.0615 |
| Tgif1 | 0.180031 | 1.71E-06 | 0.055166 |
| Lage3 | 0.179902 | 1.08E-06 | 0.034807 |
| Eri3 | 0.179897 | 3.77E-06 | 0.121617 |
| Uqcrc1 | 0.179863 | 0.000923 | 1 |

| | | | |
|---|---|---|---|
| Plscr1 | 0.179388 | 4.32E-07 | 0.013955 |
| Rpl8 | 0.179388 | 0.00087 | 1 |
| Selenow | 0.179241 | 0.01417 | 1 |
| Glrx3 | 0.179219 | 0.001681 | 1 |
| Atp11a | 0.179064 | 3.07E-07 | 0.009916 |
| Npm3 | 0.179064 | 0.000829 | 1 |
| Rpl10 | 0.178939 | 0.000408 | 1 |
| Rpl19 | 0.178863 | 5.29E-06 | 0.170776 |
| Clybl | 0.178818 | 1.69E-08 | 0.000546 |
| Prxl2b | 0.178789 | 8.13E-06 | 0.262387 |
| Smim20 | 0.17877 | 0.000416 | 1 |
| Fndc3b | 0.178738 | 4.39E-05 | 1 |
| Psmb9 | 0.178689 | 0.000774 | 1 |
| Rsph3a | 0.178667 | 6.93E-05 | 1 |
| Eif4e2 | 0.178573 | 0.000171 | 1 |
| Nab2 | 0.178407 | 3.01E-07 | 0.009707 |
| Tfg | 0.178359 | 6.78E-05 | 1 |
| Ndufs8 | 0.178358 | 0.000138 | 1 |
| Tpx2 | 0.178275 | 0.000744 | 1 |
| Mrpl15 | 0.178226 | 0.000112 | 1 |
| Asf1b | 0.178138 | 0.000135 | 1 |
| Hs6st1 | 0.17812 | 1.51E-05 | 0.487256 |
| Aimp2 | 0.177954 | 1.95E-06 | 0.063057 |
| Notch1 | 0.177439 | 1.04E-06 | 0.033503 |
| Camk1 | 0.177302 | 0.000389 | 1 |
| Tnfsf13 | 0.17714 | 0.000762 | 1 |
| Mcfd2 | 0.177091 | 9.2E-06 | 0.296898 |
| Tomm7 | 0.177077 | 0.001434 | 1 |
| Camk2g | 0.176977 | 4.41E-05 | 1 |
| Hilpda | 0.176877 | 4.89E-05 | 1 |
| Atp2a2 | 0.176746 | 2.61E-05 | 0.843646 |
| Rpl36a | 0.176652 | 0.001377 | 1 |
| Crlf2 | 0.176563 | 0.078821 | 1 |
| Eif3b | 0.176546 | 2.01E-05 | 0.649538 |
| Stx2 | 0.176525 | 1.72E-09 | 5.55E-05 |
| Ubl4a | 0.176469 | 6.12E-07 | 0.019772 |
| H2-Q10 | 0.176165 | 2.76E-05 | 0.889621 |
| Sec61b | 0.176067 | 0.028981 | 1 |
| Hist1h2ae | 0.175763 | 0.480869 | 1 |
| Slc12a7 | 0.175486 | 1.6E-07 | 0.005167 |
| Atp5c1 | 0.175378 | 0.001314 | 1 |
| Ywhah | 0.17528 | 0.001638 | 1 |
| Crtc3 | 0.17508 | 4.98E-07 | 0.016091 |
| Gas2l3 | 0.174804 | 3.07E-11 | 9.9E-07 |
| Zcchc24 | 0.174614 | 4.03E-05 | 1 |
| Kdelr2 | 0.174612 | 0.000694 | 1 |
| Pno1 | 0.174562 | 3.76E-08 | 0.001215 |
| Dpy19l1 | 0.174455 | 0.000801 | 1 |

| | | | |
|---|---|---|---|
| Ece1 | 0.174376 | 5.79E-05 | 1 |
| Mybbp1a | 0.174116 | 3.14E-06 | 0.101338 |
| Immt | 0.173801 | 4.75E-05 | 1 |
| Eea1 | 0.173594 | 4.4E-05 | 1 |
| Tubb2a | 0.17348 | 0.000164 | 1 |
| Wdr62 | 0.173252 | 1.44E-11 | 4.64E-07 |
| Cst7 | 0.173178 | 9.23E-05 | 1 |
| Gltp | 0.173067 | 0.001411 | 1 |
| Actn1 | 0.172935 | 2.6E-08 | 0.00084 |
| Morrbid | 0.172651 | 0.052795 | 1 |
| Naca | 0.172576 | 0.001255 | 1 |
| Kifc3 | 0.172317 | 0.003549 | 1 |
| Cdt1 | 0.172028 | 4.66E-05 | 1 |
| Treml2 | 0.172005 | 1.95E-06 | 0.062861 |
| Mrps15 | 0.171787 | 0.029258 | 1 |
| Gm4841 | 0.17175 | 1.39E-08 | 0.000449 |
| Plpp2 | 0.17172 | 6.55E-08 | 0.002116 |
| Psap | 0.171549 | 0.000109 | 1 |
| Hs3st3b1 | 0.171536 | 9.45E-10 | 3.05E-05 |
| Mvd | 0.171344 | 1.52E-06 | 0.049152 |
| Luzp1 | 0.171143 | 9.05E-07 | 0.029231 |
| Edem1 | 0.170931 | 7.16E-06 | 0.23131 |
| Gm7030 | 0.170834 | 0.000815 | 1 |
| Rbis | 0.17081 | 0.000265 | 1 |
| Gipc1 | 0.170683 | 3.16E-06 | 0.102128 |
| Snx6 | 0.170501 | 5.86E-05 | 1 |
| Farsa | 0.170244 | 2.15E-06 | 0.069349 |
| Dhx29 | 0.170222 | 4.87E-05 | 1 |
| Rpl26 | 0.170041 | 6.04E-05 | 1 |
| Ets2 | 0.169493 | 2.18E-06 | 0.070321 |
| Csrnp1 | 0.169486 | 1.99E-05 | 0.642625 |
| Cetn3 | 0.169437 | 7.99E-05 | 1 |
| Dynll1 | 0.169397 | 0.001544 | 1 |
| Hcfc1r1 | 0.169346 | 0.000104 | 1 |
| Bub1b | 0.169228 | 4.58E-07 | 0.014782 |
| Asah1 | 0.169121 | 0.000269 | 1 |
| Qdpr | 0.169059 | 1.57E-07 | 0.005065 |
| Slc7a8 | 0.169038 | 6.39E-05 | 1 |
| Nip7 | 0.168861 | 0.000181 | 1 |
| Tpt1 | 0.168848 | 5.13E-06 | 0.165498 |
| Phb2 | 0.168788 | 0.001282 | 1 |
| Hint2 | 0.16871 | 0.000489 | 1 |
| Fam3c | 0.168599 | 5.15E-07 | 0.016626 |
| Sfxn1 | 0.168569 | 3.53E-05 | 1 |
| Ap2s1 | 0.168563 | 0.000109 | 1 |
| Psmc4 | 0.168319 | 0.000624 | 1 |
| Tcp1 | 0.168297 | 0.000287 | 1 |
| Lpxn | 0.16825 | 8.34E-05 | 1 |

| | | | |
|---|---|---|---|
| Mfsd12 | 0.168007 | 2.07E-06 | 0.06696 |
| Mib1 | 0.167769 | 5.99E-06 | 0.193494 |
| Epb41l1 | 0.167413 | 1.37E-07 | 0.004437 |
| Slfn8 | 0.167315 | 0.016986 | 1 |
| Ffar2 | 0.167287 | 3.27E-08 | 0.001055 |
| Nucks1 | 0.167019 | 0.002203 | 1 |
| Pgam5 | 0.16683 | 6.37E-07 | 0.02057 |
| Urm1 | 0.166557 | 0.000113 | 1 |
| Cnbp | 0.166549 | 0.000687 | 1 |
| Dynll2 | 0.166494 | 3.57E-08 | 0.001153 |
| Anapc5 | 0.166429 | 0.002026 | 1 |
| Wdr75 | 0.166415 | 1.46E-05 | 0.471574 |
| Anln | 0.166254 | 6.74E-06 | 0.217736 |
| Dstn | 0.166051 | 0.00198 | 1 |
| Aars | 0.166026 | 4.59E-07 | 0.014814 |
| Endog | 0.165934 | 1.8E-08 | 0.00058 |
| Lxn | 0.165766 | 3.76E-07 | 0.012149 |
| Tmem106a | 0.165728 | 0.001748 | 1 |
| Slc27a1 | 0.165642 | 6.47E-09 | 0.000209 |
| Arl1 | 0.165529 | 0.003446 | 1 |
| Chchd6 | 0.16549 | 1.78E-06 | 0.057317 |
| Rab3d | 0.165399 | 5.11E-06 | 0.165116 |
| Ndufa10 | 0.165202 | 7.55E-06 | 0.243726 |
| Glmp | 0.165041 | 0.003186 | 1 |
| Pomp | 0.164985 | 0.014261 | 1 |
| Crat | 0.164984 | 3.6E-07 | 0.011637 |
| Fh1 | 0.164875 | 2.04E-05 | 0.658584 |
| Wdr41 | 0.164741 | 2.29E-05 | 0.738491 |
| Atp5k | 0.164641 | 0.003386 | 1 |
| Capn5 | 0.164429 | 2.3E-06 | 0.074098 |
| Mmp19 | 0.164379 | 2.06E-07 | 0.006658 |
| M6pr | 0.164147 | 0.003432 | 1 |
| Mrps24 | 0.164051 | 3.79E-06 | 0.122431 |
| Mapk7 | 0.164014 | 6.13E-06 | 0.197775 |
| Psmd3 | 0.16398 | 7.21E-06 | 0.232708 |
| Rrm1 | 0.163907 | 0.002275 | 1 |
| Mgst3 | 0.163845 | 5.63E-07 | 0.018177 |
| Bcl2l1 | 0.163713 | 0.000197 | 1 |
| Snhg1 | 0.163501 | 3.86E-07 | 0.012462 |
| Srsf6 | 0.16346 | 0.017237 | 1 |
| Lyz2 | 0.163363 | 0.00314 | 1 |
| Rpl5 | 0.162856 | 0.001526 | 1 |
| Nsun2 | 0.162717 | 5.66E-05 | 1 |
| Ndufa9 | 0.162358 | 6.19E-05 | 1 |
| Ccni | 0.162332 | 0.001607 | 1 |
| Clpp | 0.162305 | 2.27E-07 | 0.007344 |
| Chchd3 | 0.162278 | 5.17E-05 | 1 |
| BC004004 | 0.162151 | 0.001474 | 1 |

| | | | |
|---|---|---|---|
| Timm50 | 0.162007 | 3.57E-05 | 1 |
| Arl6ip4 | 0.161757 | 0.000188 | 1 |
| B2m | 0.161652 | 0.003393 | 1 |
| 2610001J0 | 0.16114 | 3.25E-06 | 0.105002 |
| Skp1a | 0.160848 | 0.004984 | 1 |
| Hp1bp3 | 0.160835 | 0.001697 | 1 |
| Uqcr10 | 0.160682 | 9.92E-05 | 1 |
| Mcm5 | 0.160552 | 0.01501 | 1 |
| Id2 | 0.160374 | 0.05344 | 1 |
| Pwp1 | 0.160315 | 6.4E-06 | 0.206525 |
| Isyna1 | 0.160296 | 0.000261 | 1 |
| Mrps12 | 0.160164 | 0.00118 | 1 |
| Slc43a2 | 0.160047 | 0.001069 | 1 |
| Myo10 | 0.160035 | 4.36E-06 | 0.140797 |
| Polr2h | 0.159977 | 9.17E-05 | 1 |
| Dnajc19 | 0.159898 | 0.002045 | 1 |
| Plin3 | 0.159656 | 2.85E-05 | 0.919659 |
| Mis18a | 0.159482 | 0.000771 | 1 |
| Rnf103 | 0.159314 | 0.002001 | 1 |
| F11r | 0.159313 | 9.64E-05 | 1 |
| Zfp560 | 0.158826 | 5.44E-06 | 0.175469 |
| Chchd4 | 0.158797 | 3.94E-08 | 0.001272 |
| Dctpp1 | 0.158779 | 2.11E-06 | 0.068124 |
| Casp4 | 0.158728 | 0.001213 | 1 |
| Pdlim7 | 0.158459 | 0.002936 | 1 |
| Ndufb7 | 0.158458 | 0.004554 | 1 |
| Psmd1 | 0.158339 | 0.000771 | 1 |
| Hsbp1 | 0.158102 | 0.001305 | 1 |
| Foxp4 | 0.158069 | 5.03E-08 | 0.001624 |
| Hspa8 | 0.157979 | 0.006473 | 1 |
| Rab2a | 0.157967 | 0.030278 | 1 |
| Sh3tc1 | 0.157923 | 5.88E-07 | 0.018985 |
| Rtca | 0.15792 | 3.68E-05 | 1 |
| Entpd1 | 0.157736 | 0.012122 | 1 |
| Ctdnep1 | 0.15773 | 1.6E-05 | 0.516669 |
| Arap2 | 0.157439 | 0.000141 | 1 |
| Ccl3 | 0.157205 | 0.114481 | 1 |
| Zfp593 | 0.157164 | 2.87E-06 | 0.092638 |
| Smdt1 | 0.157119 | 0.012947 | 1 |
| Plbd2 | 0.156776 | 0.000681 | 1 |
| Tmem160 | 0.156775 | 5.06E-07 | 0.016333 |
| Cmtm7 | 0.156762 | 0.085436 | 1 |
| Rbm7 | 0.156648 | 0.003309 | 1 |
| Mrps17 | 0.156642 | 5.39E-06 | 0.17397 |
| Mcm3 | 0.156502 | 0.001201 | 1 |
| Tmem170b | 0.156275 | 1.54E-06 | 0.049653 |
| Ormdl2 | 0.15624 | 0.000889 | 1 |
| Ube2l3 | 0.156067 | 0.002114 | 1 |

| | | | |
|---|---|---|---|
| Smg9 | 0.155962 | 2.16E-06 | 0.069819 |
| Acvr2a | 0.155855 | 3.86E-05 | 1 |
| Vapa | 0.155632 | 9.37E-05 | 1 |
| Vamp8 | 0.155544 | 0.00023 | 1 |
| Utp20 | 0.155309 | 1.46E-05 | 0.470763 |
| Snx4 | 0.154813 | 0.000297 | 1 |
| Cisd1 | 0.154637 | 0.000122 | 1 |
| Snx3 | 0.154631 | 0.001128 | 1 |
| H3f3b | 0.154513 | 0.01258 | 1 |
| Tmem205 | 0.154458 | 7.74E-05 | 1 |
| Bcap29 | 0.154374 | 1.56E-05 | 0.505221 |
| Esyt1 | 0.154357 | 0.001665 | 1 |
| Ero1l | 0.154243 | 1.86E-05 | 0.601575 |
| Pmpcb | 0.154221 | 6.56E-06 | 0.211921 |
| Cysltr1 | 0.154146 | 2.41E-09 | 7.77E-05 |
| Lpar1 | 0.154057 | 1.01E-05 | 0.325312 |
| Cct2 | 0.153992 | 0.015354 | 1 |
| Homer1 | 0.153786 | 1.13E-05 | 0.36411 |
| Alkbh7 | 0.153204 | 1.98E-07 | 0.006391 |
| Tmem258 | 0.15319 | 0.075426 | 1 |
| Hnrnpll | 0.153167 | 0.000157 | 1 |
| Psmc6 | 0.153025 | 0.001043 | 1 |
| Lamp2 | 0.152962 | 0.012633 | 1 |
| Mllt6 | 0.152938 | 2.99E-06 | 0.096452 |
| Ddhd2 | 0.152837 | 2.02E-07 | 0.006518 |
| Txnrd1 | 0.152533 | 7.67E-07 | 0.024749 |
| Rpl36al | 0.152342 | 0.00375 | 1 |
| Rilpl2 | 0.152195 | 0.000268 | 1 |
| Ksr2 | 0.152106 | 4.02E-07 | 0.012981 |
| Mrps34 | 0.152079 | 5.34E-05 | 1 |
| Pik3ip1 | 0.152013 | 4.08E-05 | 1 |
| Abce1 | 0.151904 | 3.54E-06 | 0.11435 |
| Snrpe | 0.151791 | 0.092047 | 1 |
| Ccdc34 | 0.151411 | 3.01E-07 | 0.009721 |
| Pdxk | 0.151325 | 2.91E-05 | 0.939874 |
| Ssr2 | 0.151133 | 0.000219 | 1 |
| Dennd4c | 0.150778 | 0.002532 | 1 |
| Galnt6 | 0.150736 | 8.24E-08 | 0.00266 |
| Ebp | 0.150698 | 0.000103 | 1 |
| Phgdh | 0.150295 | 5.3E-05 | 1 |
| Pes1 | 0.150282 | 1.4E-05 | 0.450729 |
| Nhlrc3 | 0.150175 | 0.000136 | 1 |
| Dbi | 0.149918 | 0.025472 | 1 |
| Hacd2 | 0.149918 | 3.78E-05 | 1 |
| Mydgf | 0.149892 | 0.000429 | 1 |
| Rasgrp1 | 0.149776 | 9.1E-06 | 0.293787 |
| Rnaseh2a | 0.149574 | 6.33E-07 | 0.02043 |
| Gadd45b | 0.149475 | 0.018397 | 1 |

| | | | |
|---|---|---|---|
| Adh5 | 0.149364 | 4.78E-05 | 1 |
| Tuba1a | 0.149254 | 0.002101 | 1 |
| Guk1 | 0.149062 | 5.09E-09 | 0.000164 |
| Npepl1 | 0.149 | 3.29E-06 | 0.106159 |
| Yipf1 | 0.148901 | 0.000336 | 1 |
| Fuca2 | 0.148836 | 1.29E-05 | 0.417498 |
| Zbtb8os | 0.148811 | 0.000426 | 1 |
| Ift43 | 0.148361 | 1.41E-08 | 0.000456 |
| H2-Q7 | 0.148301 | 0.007092 | 1 |
| Adprh | 0.148283 | 0.000989 | 1 |
| Dip2b | 0.148154 | 7.16E-05 | 1 |
| Gba | 0.148005 | 5.28E-05 | 1 |
| Mpp7 | 0.147775 | 5.6E-05 | 1 |
| Ndfip1 | 0.147749 | 0.018231 | 1 |
| Acot13 | 0.147709 | 4.06E-05 | 1 |
| Rpf2 | 0.147546 | 4.83E-05 | 1 |
| Sec24a | 0.147343 | 2.73E-05 | 0.882331 |
| Siva1 | 0.147326 | 0.000652 | 1 |
| Insig1 | 0.147054 | 0.000153 | 1 |
| Rbx1 | 0.147016 | 0.002486 | 1 |
| Trappc6a | 0.146938 | 1.58E-05 | 0.508719 |
| Erp29 | 0.146922 | 0.014119 | 1 |
| Lgals8 | 0.146831 | 6.51E-05 | 1 |
| Kif13b | 0.146624 | 0.000106 | 1 |
| A530040E1 | 0.146521 | 1.64E-06 | 0.053081 |
| Enah | 0.146396 | 2.61E-07 | 0.008415 |
| Ufm1 | 0.146384 | 0.00051 | 1 |
| Stoml2 | 0.14633 | 5.1E-08 | 0.001645 |
| Ppp1cc | 0.146198 | 6.88E-06 | 0.222012 |
| Card11 | 0.146148 | 0.000644 | 1 |
| Gm10134 | 0.146104 | 5.12E-10 | 1.65E-05 |
| Dpp9 | 0.146088 | 4.79E-06 | 0.154747 |
| Apobec3 | 0.145855 | 0.003841 | 1 |
| Rtf2 | 0.145832 | 0.011267 | 1 |
| Clec9a | 0.145755 | 4.17E-08 | 0.001346 |
| Rpl27a | 0.14548 | 0.012762 | 1 |
| Klf6 | 0.144674 | 0.013915 | 1 |
| Rab22a | 0.144437 | 0.000338 | 1 |
| Ergic3 | 0.144395 | 6.67E-05 | 1 |
| Gpatch4 | 0.144286 | 4.16E-07 | 0.013443 |
| Cdca3 | 0.144146 | 0.000241 | 1 |
| Gnl1 | 0.144012 | 3.01E-06 | 0.09725 |
| Snrpd1 | 0.143996 | 0.001937 | 1 |
| Slc15a3 | 0.143959 | 0.00092 | 1 |
| Nudt14 | 0.143551 | 1.43E-05 | 0.460313 |
| Mpst | 0.1435 | 4.11E-05 | 1 |
| Anxa5 | 0.143427 | 0.009713 | 1 |
| Gm36043 | 0.143186 | 6.05E-05 | 1 |

| | | | |
|---|---|---|---|
| Nubp2 | 0.142379 | 7.65E-05 | 1 |
| Atp6v1d | 0.142171 | 0.000361 | 1 |
| Gm32089 | 0.142128 | 1.82E-09 | 5.86E-05 |
| 1500011B( | 0.142105 | 1.02E-05 | 0.329258 |
| Edem2 | 0.142018 | 7.95E-05 | 1 |
| Ddx39b | 0.14154 | 0.012223 | 1 |
| Hnrnpa0 | 0.141428 | 5.82E-05 | 1 |
| Mtif2 | 0.141221 | 2.84E-06 | 0.091597 |
| Igf2r | 0.141213 | 3.1E-08 | 0.001001 |
| Acaa1a | 0.141211 | 0.002009 | 1 |
| Coro1c | 0.141177 | 0.003406 | 1 |
| Plk2 | 0.141104 | 2.32E-05 | 0.749523 |
| Nudt9 | 0.141029 | 1.12E-05 | 0.361648 |
| Nt5m | 0.140955 | 3.56E-05 | 1 |
| Mapk13 | 0.140942 | 1.31E-07 | 0.004235 |
| Tmbim4 | 0.140539 | 0.106825 | 1 |
| Ifi211 | 0.140397 | 0.000456 | 1 |
| Plekhf2 | 0.140346 | 0.000179 | 1 |
| Ccdc115 | 0.140344 | 0.000332 | 1 |
| Abcd3 | 0.140313 | 9.33E-06 | 0.301276 |
| Wdr36 | 0.140311 | 3.63E-05 | 1 |
| Atf1 | 0.140271 | 0.000106 | 1 |
| Mbd2 | 0.140119 | 2.43E-05 | 0.783586 |
| Arl11 | 0.140045 | 3.25E-05 | 1 |
| Pop7 | 0.139997 | 0.007008 | 1 |
| C1d | 0.139953 | 0.000249 | 1 |
| Ap2m1 | 0.139889 | 0.007464 | 1 |
| Rpl41 | 0.139882 | 0.008772 | 1 |
| Itm2c | 0.139731 | 0.000858 | 1 |
| Nudc | 0.139591 | 0.004688 | 1 |
| Nelfe | 0.139496 | 0.000862 | 1 |
| Lamtor3 | 0.13943 | 0.00161 | 1 |
| Srm | 0.139373 | 2.52E-09 | 8.14E-05 |
| Wsb2 | 0.139214 | 1.94E-07 | 0.006266 |
| Tdg | 0.13869 | 7.81E-06 | 0.252213 |
| Clic1 | 0.138603 | 0.013545 | 1 |
| Ggt5 | 0.138413 | 3.62E-05 | 1 |
| Dusp23 | 0.138316 | 3.05E-09 | 9.84E-05 |
| Ppp5c | 0.137959 | 0.000714 | 1 |
| Hibadh | 0.137928 | 3.7E-05 | 1 |
| Dtymk | 0.137774 | 0.000127 | 1 |
| Apba3 | 0.137688 | 1.06E-05 | 0.343747 |
| Golt1b | 0.13744 | 3.33E-06 | 0.107433 |
| Ppp2r1a | 0.137422 | 0.001951 | 1 |
| Cyb561d2 | 0.13742 | 4.53E-06 | 0.14619 |
| Gab1 | 0.137317 | 3.01E-07 | 0.009717 |
| Rex1bd | 0.137189 | 5.01E-05 | 1 |
| Ilf3 | 0.137133 | 1.5E-06 | 0.048569 |

| | | | |
|---|---|---|---|
| Dhx9 | 0.137092 | 0.002628 | 1 |
| Naglu | 0.137026 | 6.85E-08 | 0.002212 |
| Mtfr1 | 0.13699 | 0.141105 | 1 |
| Mrpl17 | 0.136792 | 1.79E-05 | 0.578244 |
| Pcmtd1 | 0.136767 | 2.57E-07 | 0.00829 |
| Nfkbia | 0.136408 | 0.152635 | 1 |
| Ssbp4 | 0.136116 | 0.000305 | 1 |
| Slc28a2 | 0.136002 | 0.026592 | 1 |
| Fkbp8 | 0.135968 | 0.005977 | 1 |
| Lsm2 | 0.135835 | 4.03E-05 | 1 |
| Trmt6 | 0.135712 | 0.000152 | 1 |
| Ndufs4 | 0.135464 | 0.000268 | 1 |
| March5 | 0.135394 | 0.000264 | 1 |
| Tsn | 0.135142 | 0.055425 | 1 |
| Ptpn9 | 0.135124 | 0.000464 | 1 |
| Aldh1l1 | 0.135008 | 0.001384 | 1 |
| Arid5a | 0.134947 | 2.58E-08 | 0.000832 |
| mt-Nd6 | 0.134898 | 0.000669 | 1 |
| Tmem104 | 0.134818 | 0.000219 | 1 |
| Rhoh | 0.134743 | 0.000161 | 1 |
| Psmc1 | 0.134679 | 0.000265 | 1 |
| Bola1 | 0.134565 | 0.005093 | 1 |
| Fbxl2 | 0.134559 | 2.21E-05 | 0.713305 |
| Tbrg1 | 0.134378 | 2.32E-07 | 0.007492 |
| Aen | 0.13435 | 0.000181 | 1 |
| Calm3 | 0.134165 | 0.008318 | 1 |
| Zcchc17 | 0.134079 | 0.000176 | 1 |
| Mrpl30 | 0.134076 | 0.001909 | 1 |
| Malsu1 | 0.133898 | 2.22E-05 | 0.715352 |
| Gna15 | 0.133872 | 2.8E-06 | 0.090355 |
| Htatsf1 | 0.133869 | 0.002393 | 1 |
| Ndufb11 | 0.133814 | 0.001995 | 1 |
| Med25 | 0.133766 | 0.004052 | 1 |
| Ndufa4 | 0.133747 | 0.247269 | 1 |
| Hebp1 | 0.133743 | 0.000123 | 1 |
| Taf1d | 0.133696 | 0.000188 | 1 |
| Pirb | 0.133692 | 0.253813 | 1 |
| Mgat4a | 0.133604 | 2.62E-06 | 0.084634 |
| Hnrnpa3 | 0.133564 | 0.018499 | 1 |
| Mrpl55 | 0.133455 | 4.18E-06 | 0.134817 |
| Fam78b | 0.133391 | 0.00012 | 1 |
| Wrap73 | 0.13328 | 0.000497 | 1 |
| Ptgs2os2 | 0.133207 | 1.44E-07 | 0.004663 |
| Nt5c | 0.133135 | 0.003985 | 1 |
| Vopp1 | 0.132794 | 1.96E-05 | 0.631496 |
| Selenos | 0.132766 | 0.00026 | 1 |
| Il11ra1 | 0.132693 | 1.17E-06 | 0.037815 |
| Arf1 | 0.132651 | 0.018058 | 1 |

| | | | |
|---|---|---|---|
| Copz2 | 0.13194 | 2.65E-08 | 0.000856 |
| Gpn1 | 0.131899 | 3.73E-05 | 1 |
| AU022252 | 0.131583 | 1.11E-06 | 0.035994 |
| Pcbp1 | 0.131564 | 0.000435 | 1 |
| Tmem33 | 0.131544 | 0.000425 | 1 |
| Pusl1 | 0.131297 | 1.03E-06 | 0.033231 |
| Lifr | 0.130926 | 0.0308 | 1 |
| Casp3 | 0.130856 | 0.004997 | 1 |
| Map1lc3b | 0.130777 | 0.007326 | 1 |
| Tcf12 | 0.130621 | 0.002243 | 1 |
| Kif13a | 0.130435 | 5.74E-06 | 0.185338 |
| Ech1 | 0.13041 | 0.000224 | 1 |
| Atic | 0.130393 | 0.000113 | 1 |
| Atpaf2 | 0.130309 | 0.000306 | 1 |
| Cnot10 | 0.13019 | 0.004313 | 1 |
| Stk11 | 0.130155 | 0.001162 | 1 |
| Alms1 | 0.129984 | 0.000577 | 1 |
| Ube2l6 | 0.129927 | 0.042218 | 1 |
| Plekhn1 | 0.1299 | 1.99E-08 | 0.000643 |
| Fxyd5 | 0.12964 | 0.038213 | 1 |
| Fam174a | 0.129444 | 0.006076 | 1 |
| Fcor | 0.12944 | 8.97E-07 | 0.028967 |
| Gm42047 | 0.129427 | 0.027853 | 1 |
| Lamtor5 | 0.129363 | 0.013509 | 1 |
| Lsr | 0.129256 | 1.05E-05 | 0.338829 |
| Eif1ax | 0.129126 | 4.86E-06 | 0.156955 |
| Dctn3 | 0.129113 | 0.002545 | 1 |
| Iars | 0.128871 | 6.46E-06 | 0.208436 |
| St3gal3 | 0.128742 | 0.000278 | 1 |
| Pdcd5 | 0.128622 | 0.000892 | 1 |
| Mettl16 | 0.12837 | 0.002199 | 1 |
| Snrnp25 | 0.128225 | 2.94E-05 | 0.949268 |
| Ift27 | 0.128172 | 8.19E-06 | 0.264502 |
| Arhgdia | 0.128157 | 0.024045 | 1 |
| Rars | 0.128111 | 3.98E-05 | 1 |
| Acadm | 0.128046 | 0.000126 | 1 |
| 0610030E2 | 0.128043 | 0.001938 | 1 |
| Sh3bgrl3 | 0.127984 | 0.001109 | 1 |
| Gpr108 | 0.127954 | 0.000901 | 1 |
| Fnbp1 | 0.127908 | 0.055513 | 1 |
| Dtd1 | 0.127776 | 1.37E-05 | 0.443008 |
| Tnp2 | 0.127611 | 3.7E-08 | 0.001195 |
| Anapc1 | 0.127574 | 0.021075 | 1 |
| Slc45a4 | 0.12744 | 3.68E-06 | 0.118877 |
| Slc39a1 | 0.127327 | 1.85E-06 | 0.05981 |
| Sri | 0.127285 | 0.068792 | 1 |
| Tomm40l | 0.127205 | 0.001543 | 1 |
| Pald1 | 0.127191 | 1.14E-06 | 0.036799 |

| | | | |
|---|---|---|---|
| Sin3b | 0.127123 | 0.003844 | 1 |
| Nxn | 0.127113 | 0.000506 | 1 |
| Sdha | 0.127097 | 0.008072 | 1 |
| Magoh | 0.126568 | 3.33E-05 | 1 |
| Gm40645 | 0.126541 | 9.42E-05 | 1 |
| Fcho1 | 0.126527 | 0.000346 | 1 |
| Imp4 | 0.126415 | 0.001668 | 1 |
| Cinp | 0.12632 | 0.000174 | 1 |
| Med9 | 0.126252 | 9.53E-07 | 0.030761 |
| Ncapd2 | 0.126208 | 0.000807 | 1 |
| Tbl1x | 0.126195 | 0.002016 | 1 |
| Orai1 | 0.126195 | 4.1E-05 | 1 |
| Ntmt1 | 0.126168 | 0.000424 | 1 |
| Atf4 | 0.126081 | 0.000171 | 1 |
| Mrpl51 | 0.125945 | 0.000246 | 1 |
| Aldh1b1 | 0.125931 | 3.3E-05 | 1 |
| Mrps7 | 0.12593 | 3.41E-05 | 1 |
| Ptges3 | 0.125898 | 0.002795 | 1 |
| Nop58 | 0.125527 | 0.000493 | 1 |
| Pi4ka | 0.125473 | 0.000932 | 1 |
| Bop1 | 0.125419 | 0.004456 | 1 |
| Lyar | 0.125284 | 0.000167 | 1 |
| Slc41a2 | 0.12517 | 6.12E-07 | 0.019751 |
| Smc4 | 0.12511 | 0.030206 | 1 |
| Ube2n | 0.125105 | 0.003641 | 1 |
| Tuba4a | 0.125063 | 0.012328 | 1 |
| Fam136a | 0.12505 | 4.9E-05 | 1 |
| Ddb1 | 0.125046 | 0.006958 | 1 |
| Nub1 | 0.124821 | 0.023224 | 1 |
| Ssu72 | 0.124786 | 0.03863 | 1 |
| Tmem273 | 0.12442 | 8.33E-08 | 0.00269 |
| Bag1 | 0.124409 | 4.5E-05 | 1 |
| Glul | 0.124326 | 0.010759 | 1 |
| Mrpl43 | 0.124297 | 0.02421 | 1 |
| Dnajc11 | 0.124294 | 3.22E-05 | 1 |
| Galk2 | 0.124149 | 0.030709 | 1 |
| Gpr146 | 0.124004 | 2.21E-05 | 0.714234 |
| Tigd2 | 0.123898 | 0.014164 | 1 |
| Ghitm | 0.123873 | 0.00622 | 1 |
| Rnpepl1 | 0.123873 | 4.08E-06 | 0.131648 |
| Caprin1 | 0.123831 | 0.019672 | 1 |
| Mtch1 | 0.12381 | 0.000116 | 1 |
| Slc35b1 | 0.123723 | 0.000426 | 1 |
| Atp8a1 | 0.123277 | 0.040456 | 1 |
| Rpl13a | 0.123248 | 0.004844 | 1 |
| Gtpbp8 | 0.123204 | 3.61E-06 | 0.116443 |
| Mrpl22 | 0.123192 | 8.14E-06 | 0.262686 |
| Zfp995 | 0.123174 | 0.002333 | 1 |

| | | | |
|---|---|---|---|
| Sumo3 | 0.123045 | 1.8E-05 | 0.582019 |
| Klhdc4 | 0.123035 | 4.24E-05 | 1 |
| Gtpbp4 | 0.122984 | 1.35E-05 | 0.436482 |
| Gbgt1 | 0.122574 | 6.53E-08 | 0.002107 |
| Vma21 | 0.122539 | 0.000288 | 1 |
| Ethe1 | 0.122486 | 0.000119 | 1 |
| Alg9 | 0.122444 | 0.001405 | 1 |
| Tcerg1 | 0.122435 | 0.014201 | 1 |
| Cct5 | 0.122383 | 0.024911 | 1 |
| Jpt2 | 0.122214 | 1.24E-05 | 0.400799 |
| Metap1d | 0.121982 | 0.078821 | 1 |
| Ube2d3 | 0.121865 | 0.21706 | 1 |
| Tsen15 | 0.121716 | 9.98E-06 | 0.322267 |
| Psmd13 | 0.121687 | 0.004764 | 1 |
| Rit1 | 0.12164 | 6.75E-05 | 1 |
| Atad3a | 0.121568 | 1.42E-07 | 0.004597 |
| Rbms2 | 0.121528 | 0.002411 | 1 |
| Tfdp1 | 0.121249 | 2.93E-05 | 0.94654 |
| Wdr5 | 0.121186 | 2.22E-05 | 0.717632 |
| Eef1akmt4 | 0.121185 | 1.32E-08 | 0.000427 |
| Fos | 0.121136 | 0.01007 | 1 |
| Ppme1 | 0.121083 | 0.000138 | 1 |
| Synj2 | 0.121075 | 8.01E-06 | 0.258493 |
| Grhpr | 0.121034 | 7.21E-07 | 0.023276 |
| Tnfaip8l3 | 0.120934 | 1.62E-06 | 0.052399 |
| Ap5s1 | 0.120889 | 2.14E-07 | 0.006905 |
| Vps28 | 0.120757 | 0.036592 | 1 |
| Srprb | 0.120754 | 0.001593 | 1 |
| Flywch1 | 0.120707 | 0.001237 | 1 |
| Atp13a3 | 0.120473 | 0.000202 | 1 |
| Selenoh | 0.120386 | 0.000474 | 1 |
| Nlrp3 | 0.120372 | 0.078245 | 1 |
| Rabgef1 | 0.120315 | 7.76E-06 | 0.25069 |
| Ktn1 | 0.120121 | 1.97E-05 | 0.635109 |
| Mgst2 | 0.120095 | 0.003352 | 1 |
| Txndc11 | 0.119913 | 3.91E-05 | 1 |
| Mettl1 | 0.119558 | 1.89E-06 | 0.061166 |
| Gm31718 | 0.119557 | 3.71E-06 | 0.119857 |
| Ndufaf4 | 0.119025 | 9.08E-07 | 0.029321 |
| Adam17 | 0.118972 | 0.005812 | 1 |
| Dhx58 | 0.118816 | 0.000164 | 1 |
| Nfil3 | 0.118658 | 4.1E-05 | 1 |
| Snapc1 | 0.118641 | 0.000158 | 1 |
| Fundc1 | 0.118595 | 0.003687 | 1 |
| Azi2 | 0.118549 | 0.008655 | 1 |
| Tmed9 | 0.118459 | 0.00135 | 1 |
| Ddx31 | 0.118386 | 4.96E-06 | 0.160157 |
| Rrp1 | 0.118356 | 0.006877 | 1 |

| | | | |
|---|---|---|---|
| Shmt2 | 0.118327 | 5.28E-07 | 0.017054 |
| Slpi | 0.118305 | 0.004129 | 1 |
| Atp23 | 0.118173 | 0.000124 | 1 |
| Sar1a | 0.118069 | 0.00034 | 1 |
| Dnajc3 | 0.118057 | 0.0004 | 1 |
| Tmem208 | 0.117859 | 0.010017 | 1 |
| Ctu2 | 0.11782 | 6.03E-05 | 1 |
| Abcf2 | 0.117803 | 0.000262 | 1 |
| Nr4a2 | 0.117765 | 0.005809 | 1 |
| Usp10 | 0.117684 | 0.000489 | 1 |
| Chchd5 | 0.11768 | 6.83E-05 | 1 |
| Pgs1 | 0.11767 | 0.000108 | 1 |
| AB124611 | 0.117447 | 0.02756 | 1 |
| Secisbp2l | 0.11739 | 0.000298 | 1 |
| Oat | 0.1173 | 0.000475 | 1 |
| Nectin2 | 0.117259 | 6.7E-08 | 0.002164 |
| Scamp3 | 0.117211 | 0.000526 | 1 |
| Atp6v1h | 0.117211 | 0.002859 | 1 |
| Pold2 | 0.117173 | 0.000462 | 1 |
| Mpeg1 | 0.117013 | 0.065026 | 1 |
| Ctnnb1 | 0.116847 | 0.060876 | 1 |
| Hspbp1 | 0.116831 | 6.74E-05 | 1 |
| Hmgn1 | 0.116747 | 0.000317 | 1 |
| Rbm17 | 0.116743 | 1.94E-06 | 0.062722 |
| Mcrip2 | 0.116592 | 3.09E-06 | 0.09979 |
| Eif2s1 | 0.116551 | 0.000233 | 1 |
| Cryl1 | 0.116504 | 0.006308 | 1 |
| Dglucy | 0.116425 | 0.001071 | 1 |
| Tmem241 | 0.11642 | 0.000408 | 1 |
| Fpr2 | 0.116329 | 4.27E-06 | 0.137881 |
| Acox1 | 0.116144 | 0.009802 | 1 |
| Plin2 | 0.116052 | 0.080855 | 1 |
| Smim13 | 0.115992 | 3.26E-05 | 1 |
| Rnf180 | 0.115864 | 2.25E-05 | 0.724819 |
| Sdhaf4 | 0.115657 | 7.6E-06 | 0.24522 |
| Gabarapl1 | 0.115551 | 5.28E-06 | 0.170608 |
| H2afv | 0.115363 | 0.073339 | 1 |
| Nisch | 0.115345 | 0.017816 | 1 |
| Tpst1 | 0.115182 | 0.000346 | 1 |
| Hdac2 | 0.115134 | 0.000128 | 1 |
| C1qtnf6 | 0.115089 | 3.1E-08 | 0.001002 |
| Med23 | 0.115032 | 2.08E-06 | 0.067017 |
| Ephx1 | 0.114799 | 0.00059 | 1 |
| Churc1 | 0.114674 | 0.007208 | 1 |
| Tmem87b | 0.114559 | 0.000348 | 1 |
| Pafah1b3 | 0.114551 | 8.48E-05 | 1 |
| Dym | 0.114422 | 0.006005 | 1 |
| Tipin | 0.114335 | 0.016285 | 1 |

| | | | |
|---|---|---|---|
| Iqgap2 | 0.114183 | 0.000113 | 1 |
| Fosl2 | 0.114141 | 0.000276 | 1 |
| Pole4 | 0.114121 | 5.64E-05 | 1 |
| Rab9 | 0.114092 | 0.000388 | 1 |
| Synpo | 0.114029 | 4.59E-07 | 0.014826 |
| Actr8 | 0.113816 | 0.025828 | 1 |
| Atp6v0d1 | 0.113765 | 0.01493 | 1 |
| Chmp1a | 0.1137 | 0.000667 | 1 |
| Snhg12 | 0.113642 | 8.84E-05 | 1 |
| Smim3 | 0.113509 | 0.011173 | 1 |
| Spr | 0.113432 | 0.000124 | 1 |
| Tmem245 | 0.113304 | 0.047139 | 1 |
| Hnrnpdl | 0.1133 | 0.178428 | 1 |
| Kpna1 | 0.113281 | 0.06989 | 1 |
| Enpp4 | 0.113201 | 8.19E-06 | 0.264437 |
| Ms4a4b | 0.11314 | 0.598785 | 1 |
| Ddrgk1 | 0.113086 | 8.56E-05 | 1 |
| Gcsh | 0.112811 | 6.06E-05 | 1 |
| Tmem63a | 0.112756 | 2.29E-07 | 0.007377 |
| Malt1 | 0.112514 | 0.929241 | 1 |
| Prkar1a | 0.112196 | 0.146538 | 1 |
| Tomm22 | 0.112119 | 0.00436 | 1 |
| Utp15 | 0.112085 | 6.35E-06 | 0.205162 |
| Park7 | 0.11201 | 0.006829 | 1 |
| Rabl6 | 0.111957 | 0.000181 | 1 |
| Nup85 | 0.111775 | 7.94E-05 | 1 |
| Tmem106b | 0.111757 | 7.29E-06 | 0.23538 |
| Slc37a2 | 0.111755 | 1.22E-06 | 0.039316 |
| Ctla2b | 0.111732 | 0.000222 | 1 |
| Anapc4 | 0.11173 | 4.53E-05 | 1 |
| Akr1b10 | 0.111647 | 0.000131 | 1 |
| Nol12 | 0.111641 | 1.81E-06 | 0.058279 |
| Sept8 | 0.111584 | 5.87E-05 | 1 |
| Zswim7 | 0.111405 | 3.51E-05 | 1 |
| Arfgap3 | 0.111307 | 1.86E-06 | 0.060207 |
| Slc25a51 | 0.111287 | 0.023259 | 1 |
| Eif3d | 0.111273 | 0.009297 | 1 |
| Hbp1 | 0.111252 | 0.004367 | 1 |
| Heatr6 | 0.111227 | 5.09E-05 | 1 |
| Idnk | 0.111033 | 0.037763 | 1 |
| Tmem9b | 0.110981 | 0.021102 | 1 |
| Knstrn | 0.110723 | 0.003605 | 1 |
| Blvrb | 0.110616 | 0.024997 | 1 |
| Psmd11 | 0.110509 | 0.061562 | 1 |
| Pdcd2 | 0.110387 | 0.001197 | 1 |
| Srp19 | 0.110347 | 0.00312 | 1 |
| Gaa | 0.110342 | 4.61E-05 | 1 |
| Mknk2 | 0.110336 | 0.033464 | 1 |

| | | | |
|---|---|---|---|
| Arrdc3 | 0.110329 | 0.00153 | 1 |
| Mob2 | 0.110168 | 0.000704 | 1 |
| Syvn1 | 0.110114 | 0.001008 | 1 |
| Glod4 | 0.110108 | 0.039564 | 1 |
| Hdgf | 0.110056 | 0.005819 | 1 |
| 2610044O | 0.109566 | 0.012738 | 1 |
| Rpsa | 0.109529 | 0.058615 | 1 |
| Gramd3 | 0.10948 | 0.022191 | 1 |
| Acadsb | 0.109444 | 2.96E-05 | 0.955359 |
| Rrp7a | 0.109399 | 0.010053 | 1 |
| Peli2 | 0.109195 | 2.35E-06 | 0.075986 |
| Rpl11 | 0.109011 | 0.019592 | 1 |
| Gclm | 0.1089 | 3.75E-08 | 0.001211 |
| Fads1 | 0.108788 | 4E-05 | 1 |
| Ptbp1 | 0.10876 | 0.021426 | 1 |
| Slc48a1 | 0.108697 | 0.001589 | 1 |
| Exosc8 | 0.108631 | 0.000183 | 1 |
| Elp1 | 0.108544 | 3.28E-06 | 0.106048 |
| Psmd9 | 0.108419 | 4.43E-05 | 1 |
| Ciapin1 | 0.108415 | 0.005672 | 1 |
| Trim30c | 0.108385 | 0.000458 | 1 |
| Dusp10 | 0.108101 | 3.85E-05 | 1 |
| Mvk | 0.108079 | 0.003262 | 1 |
| Eif2b1 | 0.108006 | 0.000891 | 1 |
| Eef1e1 | 0.107802 | 0.00019 | 1 |
| Rps6kc1 | 0.107791 | 4.11E-05 | 1 |
| Vill | 0.107765 | 0.131033 | 1 |
| Rack1 | 0.107723 | 0.036008 | 1 |
| Tor2a | 0.107715 | 5.7E-05 | 1 |
| Lztfl1 | 0.107523 | 0.00285 | 1 |
| Coro7 | 0.10729 | 5.23E-05 | 1 |
| Ubac1 | 0.107237 | 1.09E-05 | 0.352283 |
| Ndufb4 | 0.107207 | 0.012044 | 1 |
| Asna1 | 0.107195 | 1.83E-05 | 0.590132 |
| Lamtor4 | 0.107174 | 0.112093 | 1 |
| Tuba1c | 0.107119 | 0.026991 | 1 |
| Vps35 | 0.107077 | 0.041752 | 1 |
| Eif3a | 0.107068 | 0.020974 | 1 |
| Ints2 | 0.10681 | 7.67E-06 | 0.247651 |
| Sdhaf1 | 0.106779 | 0.000174 | 1 |
| Eif2s2 | 0.106751 | 0.021114 | 1 |
| Mtm1 | 0.106742 | 0.001336 | 1 |
| Gm47283 | 0.106718 | 0.014879 | 1 |
| Atm | 0.106706 | 0.000223 | 1 |
| Slc7a11 | 0.106704 | 2.63E-05 | 0.848521 |
| Scamp1 | 0.106626 | 3.71E-06 | 0.119916 |
| Pin1 | 0.106624 | 1.9E-05 | 0.612813 |
| Ube2v1 | 0.106577 | 0.026173 | 1 |

| | | | |
|---|---|---|---|
| Mrpl48 | 0.106527 | 0.004663 | 1 |
| Tchh | 0.106476 | 1.64E-07 | 0.005279 |
| B230219D: | 0.106455 | 0.001127 | 1 |
| Cd69 | 0.106316 | 0.000553 | 1 |
| Polr1d | 0.106254 | 0.01308 | 1 |
| Gm37240 | 0.106206 | 0.000241 | 1 |
| Tgtp1 | 0.106155 | 6.95E-05 | 1 |
| Cdkn3 | 0.106148 | 0.000307 | 1 |
| Eif3l | 0.106059 | 0.000503 | 1 |
| Aldh9a1 | 0.10604 | 0.008174 | 1 |
| Flot2 | 0.105894 | 0.000971 | 1 |
| Vti1b | 0.105669 | 0.003468 | 1 |
| 2310022A: | 0.105457 | 0.130413 | 1 |
| Ccng2 | 0.105308 | 0.004358 | 1 |
| Ctnnd1 | 0.10527 | 0.015943 | 1 |
| Kif23 | 0.104937 | 0.147498 | 1 |
| Dcaf1 | 0.104937 | 1.61E-06 | 0.051839 |
| Ostm1 | 0.104878 | 1.45E-05 | 0.469108 |
| Dctn2 | 0.10486 | 0.067072 | 1 |
| Leprotl1 | 0.104725 | 0.000367 | 1 |
| G3bp1 | 0.104719 | 0.031235 | 1 |
| Ezh2 | 0.104683 | 0.020709 | 1 |
| Dad1 | 0.104562 | 0.035139 | 1 |
| Bin1 | 0.104508 | 0.002161 | 1 |
| Adora3 | 0.104501 | 5.75E-05 | 1 |
| Pik3r1 | 0.104481 | 0.014878 | 1 |
| Casp12 | 0.104362 | 2.5E-06 | 0.080606 |
| Wdr48 | 0.104352 | 0.000513 | 1 |
| Golgb1 | 0.104322 | 0.001005 | 1 |
| Esr1 | 0.104307 | 0.003607 | 1 |
| Zfp1 | 0.104293 | 0.000752 | 1 |
| Abl2 | 0.104209 | 2.37E-06 | 0.076392 |
| Icosl | 0.10415 | 0.000108 | 1 |
| Rps15a | 0.104019 | 0.106088 | 1 |
| 3830406C: | 0.103978 | 0.005281 | 1 |
| Src | 0.103977 | 1.5E-05 | 0.485629 |
| Rnpep | 0.103954 | 0.404501 | 1 |
| Poglut1 | 0.103917 | 0.011556 | 1 |
| Acd | 0.103886 | 0.000598 | 1 |
| Tor1b | 0.103613 | 0.002229 | 1 |
| Vezt | 0.10358 | 0.000373 | 1 |
| Hspa4l | 0.10353 | 0.000126 | 1 |
| Atrn | 0.103451 | 0.019265 | 1 |
| E330020D: | 0.103424 | 4.97E-05 | 1 |
| Mgrn1 | 0.103399 | 0.006444 | 1 |
| Tnfsf13b | 0.103338 | 0.000886 | 1 |
| Sec24d | 0.103226 | 1.42E-05 | 0.459264 |
| Acpp | 0.103179 | 1E-05 | 0.323229 |

| | | | |
|---|---|---|---|
| Ssbp1 | 0.103118 | 0.001587 | 1 |
| Usp5 | 0.102963 | 0.000215 | 1 |
| Bscl2 | 0.102877 | 0.015415 | 1 |
| Rxra | 0.102868 | 0.000993 | 1 |
| Rcan3 | 0.102822 | 0.000327 | 1 |
| Ccdc122 | 0.10279 | 2.06E-05 | 0.66485 |
| Chst14 | 0.102714 | 3.04E-06 | 0.09807 |
| Lonp1 | 0.102617 | 0.001741 | 1 |
| Mphosph6 | 0.102567 | 3.52E-05 | 1 |
| Cyth2 | 0.10242 | 0.000235 | 1 |
| Galt | 0.10218 | 0.025053 | 1 |
| Pon3 | 0.102143 | 1.5E-05 | 0.484915 |
| Alkbh2 | 0.10201 | 0.000133 | 1 |
| Rmi2 | 0.101933 | 2.14E-08 | 0.000691 |
| Tmem199 | 0.101667 | 0.000651 | 1 |
| Fez2 | 0.101596 | 8.1E-05 | 1 |
| Wdr13 | 0.101576 | 0.000837 | 1 |
| Osbpl3 | 0.101449 | 0.011213 | 1 |
| Ubxn1 | 0.101446 | 0.029117 | 1 |
| Pacsin1 | 0.101371 | 3.68E-06 | 0.118869 |
| Rps4x | 0.10133 | 0.083042 | 1 |
| Imp3 | 0.101312 | 0.003845 | 1 |
| Gnb4 | 0.101266 | 0.000747 | 1 |
| Zpr1 | 0.101178 | 0.000437 | 1 |
| Pi4k2a | 0.101169 | 0.000375 | 1 |
| Fkbp4 | 0.101167 | 0.009346 | 1 |
| Rufy2 | 0.101114 | 3.31E-05 | 1 |
| Chmp6 | 0.101081 | 0.001331 | 1 |
| Hdhd2 | 0.100985 | 1.55E-05 | 0.500623 |
| Mlf2 | 0.100727 | 0.030703 | 1 |
| Otub1 | 0.100672 | 0.012132 | 1 |
| Fam129b | 0.10057 | 0.000196 | 1 |
| Gpat3 | 0.100255 | 4.13E-06 | 0.133454 |
| Mrps9 | 0.10016 | 0.000602 | 1 |
| Cuta | 0.100147 | 0.015102 | 1 |
| Sclt1 | 0.100026 | 0.038896 | 1 |
| Mrpl3 | 0.100022 | 0.003577 | 1 |
| Dnmt1 | 0.099933 | 0.00449 | 1 |
| Maff | 0.099925 | 0.000117 | 1 |
| Vapb | 0.099775 | 0.000141 | 1 |
| Slc10a3 | 0.099717 | 0.000121 | 1 |
| Glrx2 | 0.099672 | 0.00646 | 1 |
| Brip1 | 0.099653 | 0.000136 | 1 |
| Smarcc1 | 0.099583 | 2.68E-05 | 0.865461 |
| Prdx3 | 0.099567 | 0.000166 | 1 |
| Mcph1 | 0.099401 | 0.000753 | 1 |
| Gpkow | 0.09931 | 0.021829 | 1 |
| Ubald2 | 0.09926 | 0.049136 | 1 |

| | | | |
|---|---|---|---|
| Ptges | 0.099156 | 6.87E-06 | 0.22183 |
| Cdk16 | 0.098958 | 2.4E-05 | 0.775223 |
| Gm21860 | 0.09889 | 2.61E-05 | 0.842105 |
| Tep1 | 0.098877 | 0.021715 | 1 |
| Mecr | 0.098798 | 0.011235 | 1 |
| Pcyox1 | 0.098713 | 0.002649 | 1 |
| Akt1 | 0.0985 | 0.0001 | 1 |
| Ruvbl1 | 0.098392 | 2.14E-05 | 0.689693 |
| Lcmt1 | 0.098288 | 0.00373 | 1 |
| Slc38a6 | 0.098256 | 0.000163 | 1 |
| Polr2m | 0.098189 | 0.000792 | 1 |
| Ifit1 | 0.098142 | 5.07E-05 | 1 |
| Arhgap10 | 0.098118 | 1.4E-06 | 0.045311 |
| Ccdc85b | 0.098094 | 4.37E-05 | 1 |
| Srp72 | 0.09799 | 0.000197 | 1 |
| Timm10b | 0.097917 | 0.027793 | 1 |
| Ttc27 | 0.097864 | 0.002241 | 1 |
| Rpia | 0.097725 | 0.000105 | 1 |
| Pdss1 | 0.097659 | 1.18E-05 | 0.380007 |
| Ccdc32 | 0.097653 | 0.007186 | 1 |
| Xpa | 0.097652 | 7.19E-05 | 1 |
| Uba2 | 0.097473 | 0.003474 | 1 |
| Dap3 | 0.097377 | 8.77E-05 | 1 |
| Cops9 | 0.097351 | 0.022655 | 1 |
| Parp12 | 0.097332 | 0.003224 | 1 |
| Dync1h1 | 0.097327 | 0.06966 | 1 |
| Carmil1 | 0.097238 | 1.44E-05 | 0.465828 |
| Tmem97 | 0.09719 | 0.002225 | 1 |
| B4galt1 | 0.097135 | 0.020674 | 1 |
| Gosr2 | 0.097095 | 0.045952 | 1 |
| Ints11 | 0.097027 | 4.15E-05 | 1 |
| Zbtb42 | 0.096999 | 0.009993 | 1 |
| Cfap20 | 0.09696 | 0.000108 | 1 |
| Tk1 | 0.096941 | 0.000574 | 1 |
| March3 | 0.096786 | 0.000726 | 1 |
| Usf2 | 0.096718 | 0.000735 | 1 |
| Pip4p2 | 0.09661 | 0.00017 | 1 |
| Tnfsf10 | 0.096592 | 8.85E-05 | 1 |
| Rfc1 | 0.096571 | 0.240966 | 1 |
| Il4ra | 0.096469 | 0.005911 | 1 |
| Apol9b | 0.09644 | 7.17E-06 | 0.231527 |
| Srr | 0.096425 | 0.002059 | 1 |
| Chrac1 | 0.096413 | 0.009917 | 1 |
| Mapk9 | 0.096376 | 0.003168 | 1 |
| Cox10 | 0.096326 | 0.03696 | 1 |
| Enpp5 | 0.096284 | 0.000407 | 1 |
| Zmat2 | 0.096261 | 0.001256 | 1 |
| Tmem106c | 0.096204 | 2.21E-05 | 0.712935 |

| | | | |
|---|---|---|---|
| Anp32e | 0.096184 | 0.025271 | 1 |
| Micall2 | 0.095931 | 3.81E-05 | 1 |
| Tiparp | 0.095862 | 6.99E-05 | 1 |
| Trmt11 | 0.095846 | 0.060474 | 1 |
| Nup88 | 0.095779 | 0.019236 | 1 |
| Iars2 | 0.095716 | 0.021068 | 1 |
| Gtf2h4 | 0.095485 | 0.015394 | 1 |
| Hras | 0.095393 | 0.011302 | 1 |
| Fnip1 | 0.095382 | 0.000587 | 1 |
| Per2 | 0.095378 | 3.19E-06 | 0.102856 |
| Morf4l2 | 0.095358 | 0.008969 | 1 |
| Smyd2 | 0.095347 | 9.68E-06 | 0.312453 |
| Minpp1 | 0.09533 | 0.002586 | 1 |
| Nfkbie | 0.095313 | 0.210749 | 1 |
| Bhlhe40 | 0.095175 | 0.002604 | 1 |
| Cops6 | 0.095069 | 0.004748 | 1 |
| Mrps18b | 0.095062 | 0.001968 | 1 |
| Trmt2a | 0.094944 | 0.000179 | 1 |
| Adgrb1 | 0.094935 | 4.23E-07 | 0.013659 |
| 1110012L1 | 0.094846 | 0.043305 | 1 |
| St6galnac4 | 0.094821 | 0.000955 | 1 |
| Tyms | 0.094717 | 2.27E-05 | 0.734483 |
| Tsta3 | 0.094627 | 2.16E-06 | 0.069693 |
| A930037H( | 0.09461 | 0.023441 | 1 |
| Wdr43 | 0.094606 | 0.000614 | 1 |
| Optn | 0.09457 | 1.41E-06 | 0.04555 |
| Bad | 0.094479 | 2.32E-05 | 0.748872 |
| Txndc15 | 0.094347 | 0.015214 | 1 |
| Nln | 0.094273 | 0.003009 | 1 |
| Msl3 | 0.094161 | 0.000971 | 1 |
| Hyou1 | 0.094148 | 0.014799 | 1 |
| Tbrg4 | 0.094119 | 0.006476 | 1 |
| Nufip1 | 0.094024 | 3.98E-05 | 1 |
| Mxd4 | 0.094016 | 0.000751 | 1 |
| Tradd | 0.093732 | 0.009386 | 1 |
| Pinx1 | 0.093656 | 0.000246 | 1 |
| Ruvbl2 | 0.093605 | 3.81E-05 | 1 |
| Prkaa1 | 0.093503 | 0.022839 | 1 |
| Rmc1 | 0.093483 | 0.002472 | 1 |
| Mavs | 0.093452 | 0.000601 | 1 |
| Neu1 | 0.093416 | 0.001375 | 1 |
| Dmxl1 | 0.093398 | 0.000127 | 1 |
| Mrto4 | 0.09327 | 0.002026 | 1 |
| Prkar2a | 0.093216 | 7.74E-05 | 1 |
| 2310039H( | 0.09321 | 0.000301 | 1 |
| Tomm5 | 0.093035 | 0.01275 | 1 |
| Klhl2 | 0.092892 | 0.129135 | 1 |
| Lsm4 | 0.092854 | 0.056168 | 1 |

| | | | |
|---|---|---|---|
| Stip1 | 0.092852 | 0.005633 | 1 |
| Ankrd40 | 0.092852 | 0.000242 | 1 |
| Psat1 | 0.092786 | 0.000221 | 1 |
| Mepce | 0.092717 | 0.042738 | 1 |
| Kpna3 | 0.092704 | 0.000117 | 1 |
| Rps23 | 0.092694 | 0.139873 | 1 |
| Dlat | 0.092647 | 0.000111 | 1 |
| Ppp1r12c | 0.092523 | 0.018136 | 1 |
| Rps19bp1 | 0.092457 | 0.000288 | 1 |
| Als2cl | 0.092435 | 2.4E-06 | 0.077471 |
| Cggbp1 | 0.092393 | 0.060231 | 1 |
| Ckap4 | 0.092232 | 0.012833 | 1 |
| Ddx54 | 0.092223 | 0.006783 | 1 |
| Ei24 | 0.092159 | 0.000289 | 1 |
| Slc35f6 | 0.092098 | 0.000255 | 1 |
| Trmt61a | 0.092054 | 0.000227 | 1 |
| Ell2 | 0.092032 | 0.005928 | 1 |
| Uba5 | 0.091979 | 0.000191 | 1 |
| Tpcn2 | 0.091968 | 4E-06 | 0.129255 |
| Dld | 0.091858 | 0.006352 | 1 |
| Zfp64 | 0.091854 | 1.31E-05 | 0.423766 |
| Clip2 | 0.09176 | 1.64E-05 | 0.528644 |
| Ado | 0.091731 | 0.000502 | 1 |
| Fam214a | 0.091709 | 0.163408 | 1 |
| Cops5 | 0.091638 | 7.95E-05 | 1 |
| Dnlz | 0.091566 | 0.022009 | 1 |
| Mat2b | 0.091539 | 0.009515 | 1 |
| Cd52 | 0.091504 | 0.227807 | 1 |
| Trappc3 | 0.091458 | 0.001216 | 1 |
| Pym1 | 0.091431 | 0.000302 | 1 |
| Mta1 | 0.091406 | 0.000241 | 1 |
| Psmc3 | 0.091363 | 0.061775 | 1 |
| Raly | 0.091177 | 0.005681 | 1 |
| Cox14 | 0.090975 | 0.03965 | 1 |
| Adamtsl4 | 0.090909 | 2.45E-06 | 0.079208 |
| Prps1 | 0.090895 | 7.67E-05 | 1 |
| Maz | 0.090698 | 0.003212 | 1 |
| Rbck1 | 0.090679 | 0.00277 | 1 |
| Afg1l | 0.090632 | 0.000105 | 1 |
| Zfp281 | 0.090498 | 0.000359 | 1 |
| Stard9 | 0.090426 | 0.001582 | 1 |
| Cdc16 | 0.090278 | 0.000111 | 1 |
| Gsto1 | 0.090233 | 0.005878 | 1 |
| Rest | 0.090227 | 0.000386 | 1 |
| Polr2f | 0.090125 | 0.0003 | 1 |
| Rhou | 0.089867 | 7.02E-05 | 1 |
| Mdh1 | 0.089855 | 0.21577 | 1 |
| Sik3 | 0.089855 | 0.003772 | 1 |

| | | | |
|---|---|---|---|
| Atf2 | 0.089837 | 0.001483 | 1 |
| Cpsf1 | 0.089837 | 5.05E-05 | 1 |
| Trap1 | 0.089807 | 8.44E-06 | 0.272632 |
| Gpr180 | 0.089766 | 0.000313 | 1 |
| Pld2 | 0.089687 | 6.08E-05 | 1 |
| Rpain | 0.089684 | 0.000351 | 1 |
| Pkib | 0.089593 | 1.01E-06 | 0.032661 |
| Ncapg2 | 0.089567 | 0.003969 | 1 |
| Tmem183a | 0.089437 | 0.000694 | 1 |
| Fam120a | 0.08943 | 0.047692 | 1 |
| Smurf1 | 0.089375 | 1.42E-05 | 0.459481 |
| Hells | 0.089332 | 0.00211 | 1 |
| Iah1 | 0.089312 | 0.000625 | 1 |
| Itm2b | 0.089246 | 0.034165 | 1 |
| Pigk | 0.089169 | 1.03E-05 | 0.332288 |
| Hspb11 | 0.089096 | 9.98E-05 | 1 |
| Ncbp2 | 0.089046 | 0.001529 | 1 |
| Gtf2a2 | 0.089044 | 0.039189 | 1 |
| Uqcc3 | 0.08904 | 0.002273 | 1 |
| Nmi | 0.088869 | 0.008258 | 1 |
| Ftsj1 | 0.088861 | 9.04E-05 | 1 |
| Pola1 | 0.088843 | 0.000297 | 1 |
| Fggy | 0.088813 | 1.97E-06 | 0.063647 |
| Diaph3 | 0.088744 | 0.027244 | 1 |
| Ints1 | 0.088634 | 0.001622 | 1 |
| Rpl18a | 0.088626 | 0.031021 | 1 |
| E130309D( | 0.088572 | 3.36E-05 | 1 |
| E2f4 | 0.088467 | 0.002527 | 1 |
| Mpg | 0.088391 | 0.014487 | 1 |
| Slx4ip | 0.088302 | 0.043406 | 1 |
| Aurkaip1 | 0.088284 | 0.119949 | 1 |
| Gm37233 | 0.08827 | 2.33E-06 | 0.07521 |
| Trnt1 | 0.088142 | 0.034492 | 1 |
| Asl | 0.088103 | 0.000217 | 1 |
| Hmgb2 | 0.088068 | 0.649722 | 1 |
| Crbn | 0.088065 | 0.005309 | 1 |
| Hikeshi | 0.088023 | 0.004774 | 1 |
| Manbal | 0.088014 | 6.39E-06 | 0.20614 |
| Eif4g1 | 0.087923 | 0.013536 | 1 |
| Slc25a46 | 0.087827 | 3.79E-05 | 1 |
| Fbxo34 | 0.087739 | 0.012974 | 1 |
| Ppt2 | 0.087642 | 0.015418 | 1 |
| Gm4117 | 0.087591 | 0.04551 | 1 |
| Rab3ip | 0.087529 | 1.64E-05 | 0.528607 |
| Pttg1ip | 0.087373 | 0.000269 | 1 |
| Ptges2 | 0.087343 | 0.000433 | 1 |
| Arf4 | 0.087092 | 0.028059 | 1 |
| Znhit2 | 0.087024 | 9.13E-05 | 1 |

| | | | |
|---|---|---|---|
| Rpa2 | 0.086958 | 0.14361 | 1 |
| Strap | 0.086771 | 0.030736 | 1 |
| Zfand2a | 0.086675 | 0.001339 | 1 |
| Mea1 | 0.086458 | 0.040654 | 1 |
| Rbm28 | 0.086273 | 0.013728 | 1 |
| Rusc1 | 0.08623 | 0.403541 | 1 |
| Agap3 | 0.086209 | 5.57E-05 | 1 |
| Itpk1 | 0.086181 | 0.002071 | 1 |
| Ifi35 | 0.086172 | 0.01068 | 1 |
| Spata5 | 0.086046 | 0.053688 | 1 |
| Ubl7 | 0.085988 | 0.008189 | 1 |
| Mrps5 | 0.085977 | 1.37E-05 | 0.441232 |
| Nfu1 | 0.085873 | 0.000333 | 1 |
| Tra2a | 0.085869 | 0.031052 | 1 |
| Osbpl1a | 0.085852 | 3.78E-06 | 0.122092 |
| 1110051M | 0.085806 | 5.15E-05 | 1 |
| Polr3d | 0.085735 | 0.000314 | 1 |
| Mag | 0.085707 | 3.98E-06 | 0.128467 |
| Rfc5 | 0.085598 | 3.64E-05 | 1 |
| Polr2e | 0.085479 | 0.001017 | 1 |
| Snx8 | 0.085424 | 2.45E-06 | 0.079224 |
| Rrs1 | 0.085324 | 1.5E-05 | 0.485167 |
| Gla | 0.085194 | 0.000166 | 1 |
| Tmem101 | 0.085134 | 0.009997 | 1 |
| Lamc1 | 0.085052 | 0.00026 | 1 |
| Hdgfl2 | 0.084999 | 0.000251 | 1 |
| Acvrl1 | 0.084984 | 0.002002 | 1 |
| Ccdc71l | 0.084837 | 0.013854 | 1 |
| Tagap | 0.084741 | 0.030615 | 1 |
| Arsg | 0.084684 | 2.26E-05 | 0.729837 |
| Rab11fip5 | 0.084654 | 0.000205 | 1 |
| Slc35a3 | 0.084646 | 0.003264 | 1 |
| Sgta | 0.084609 | 0.000248 | 1 |
| Cmtm6 | 0.084607 | 0.025704 | 1 |
| Rcc1 | 0.084512 | 0.002788 | 1 |
| Rcc1l | 0.084506 | 2.41E-05 | 0.778319 |
| Prps1l3 | 0.084426 | 1.21E-05 | 0.391398 |
| Thap7 | 0.08439 | 0.02531 | 1 |
| Hpf1 | 0.084352 | 0.002199 | 1 |
| Zfp622 | 0.084337 | 0.010187 | 1 |
| Gys1 | 0.084278 | 0.00037 | 1 |
| Sh3glb2 | 0.084057 | 0.000393 | 1 |
| Kat14 | 0.084023 | 0.000509 | 1 |
| Pofut2 | 0.083962 | 0.006472 | 1 |
| H2-M3 | 0.083942 | 0.160273 | 1 |
| 1110032A( | 0.083937 | 0.015255 | 1 |
| Abhd4 | 0.083926 | 0.004241 | 1 |
| Kif1c | 0.083885 | 5.21E-06 | 0.168259 |

| | | | |
|---|---|---|---|
| Maf | 0.083879 | 8.39E-06 | 0.270814 |
| Mrpl28 | 0.08387 | 0.000123 | 1 |
| Ltbr | 0.083834 | 0.007403 | 1 |
| Clec4d | 0.083805 | 8.44E-05 | 1 |
| Chd4 | 0.083767 | 0.355206 | 1 |
| Cox17 | 0.083723 | 0.067531 | 1 |
| Gm49359 | 0.083705 | 0.001446 | 1 |
| Tmem126a | 0.083604 | 0.0006 | 1 |
| Fxn | 0.083506 | 0.000955 | 1 |
| Carm1 | 0.083458 | 0.010275 | 1 |
| Hint3 | 0.083411 | 6.17E-05 | 1 |
| Slc7a5 | 0.083323 | 5.5E-06 | 0.177611 |
| Ciao2a | 0.083321 | 0.093986 | 1 |
| Meaf6 | 0.083201 | 0.0005 | 1 |
| Psma7 | 0.083194 | 0.116533 | 1 |
| Mrrf | 0.083156 | 0.000165 | 1 |
| Vps18 | 0.083141 | 8.71E-05 | 1 |
| Mbd3 | 0.08306 | 0.000237 | 1 |
| Atp6v1b2 | 0.082977 | 0.064374 | 1 |
| Egln2 | 0.082878 | 0.00019 | 1 |
| Txnl1 | 0.082858 | 0.006255 | 1 |
| Eif1ad | 0.08278 | 0.005683 | 1 |
| Mfn2 | 0.082774 | 0.000277 | 1 |
| Mcm7 | 0.082731 | 0.00468 | 1 |
| Cep164 | 0.082685 | 0.00326 | 1 |
| Ptrh2 | 0.082664 | 0.000569 | 1 |
| Cluh | 0.082662 | 0.009169 | 1 |
| Rab6a | 0.082657 | 0.028874 | 1 |
| Gorasp2 | 0.082644 | 0.020017 | 1 |
| Polr2d | 0.082637 | 0.000444 | 1 |
| Htatip2 | 0.08258 | 0.001003 | 1 |
| Elof1 | 0.082565 | 7.67E-05 | 1 |
| Cdc34 | 0.082549 | 0.000101 | 1 |
| Rab28 | 0.08249 | 4.98E-05 | 1 |
| Pcm1 | 0.082461 | 0.00193 | 1 |
| Bckdhb | 0.082412 | 0.001736 | 1 |
| Kdsr | 0.082407 | 0.003559 | 1 |
| Slc39a3 | 0.082319 | 5.65E-05 | 1 |
| Alg5 | 0.08229 | 0.00017 | 1 |
| Ddx21 | 0.082232 | 0.03804 | 1 |
| Cenpm | 0.082227 | 1.03E-05 | 0.331641 |
| Eif4e3 | 0.082199 | 0.018143 | 1 |
| Tex14 | 0.082141 | 0.000346 | 1 |
| Mfhas1 | 0.082084 | 0.000659 | 1 |
| Bcl2a1d | 0.082083 | 0.028584 | 1 |
| Gtf2f1 | 0.082037 | 0.002248 | 1 |
| Npc1 | 0.081953 | 0.001809 | 1 |
| Serbp1 | 0.08192 | 0.088288 | 1 |

| | | | |
|---|---|---|---|
| Bcl2a1a | 0.081881 | 0.000126 | 1 |
| Elp3 | 0.081818 | 2.06E-05 | 0.665684 |
| Mcm4 | 0.08177 | 0.077972 | 1 |
| Rragc | 0.081763 | 0.002914 | 1 |
| Urb1 | 0.081719 | 0.000105 | 1 |
| Zdhhc1 | 0.081567 | 0.003157 | 1 |
| Sqle | 0.081454 | 0.052074 | 1 |
| Actr1b | 0.081437 | 0.000396 | 1 |
| Aco1 | 0.081417 | 0.001515 | 1 |
| Ppm1m | 0.081363 | 0.00295 | 1 |
| Ubxn6 | 0.08131 | 0.000129 | 1 |
| Pdcd1lg2 | 0.081281 | 0.00198 | 1 |
| Gnai3 | 0.081268 | 0.072729 | 1 |
| Trappc5 | 0.081184 | 0.001487 | 1 |
| Wdr12 | 0.081074 | 0.000501 | 1 |
| Snhg4 | 0.081044 | 0.000489 | 1 |
| Trir | 0.081012 | 0.114147 | 1 |
| Dnajc25 | 0.080984 | 7.23E-05 | 1 |
| Sav1 | 0.080945 | 6.03E-05 | 1 |
| Exosc5 | 0.080899 | 1.87E-05 | 0.604709 |
| Tma7 | 0.080732 | 0.45422 | 1 |
| Ints4 | 0.08065 | 0.037284 | 1 |
| Ocstamp | 0.080501 | 0.046481 | 1 |
| Arl8b | 0.080452 | 0.004236 | 1 |
| Plk3 | 0.080449 | 0.001225 | 1 |
| Ctns | 0.080325 | 0.000268 | 1 |
| Psme4 | 0.080321 | 0.026768 | 1 |
| Slfn2 | 0.080098 | 0.232206 | 1 |
| Mrpl37 | 0.080042 | 0.000418 | 1 |
| Blvra | 0.080006 | 0.009927 | 1 |
| Lancl1 | 0.079959 | 0.063986 | 1 |
| Thap4 | 0.079933 | 1.47E-05 | 0.475206 |
| Romo1 | 0.079923 | 0.008738 | 1 |
| Ptpn18 | 0.079756 | 0.127455 | 1 |
| Rab24 | 0.079756 | 0.000438 | 1 |
| Slc9a3r1 | 0.079678 | 0.010219 | 1 |
| B4galt6 | 0.079609 | 0.000505 | 1 |
| Pink1 | 0.079402 | 0.016035 | 1 |
| Dtx3 | 0.079307 | 2.45E-06 | 0.079051 |
| Nedd8 | 0.079291 | 0.21107 | 1 |
| Zcchc2 | 0.079238 | 0.038978 | 1 |
| Wipi1 | 0.079125 | 0.000125 | 1 |
| Tubgcp3 | 0.079124 | 0.056241 | 1 |
| Dag1 | 0.07904 | 7.47E-05 | 1 |
| Uggt1 | 0.079039 | 0.018781 | 1 |
| Gatd3a | 0.078989 | 0.000725 | 1 |
| Focad | 0.078959 | 0.001274 | 1 |
| St14 | 0.07885 | 5.75E-05 | 1 |

| | | | |
|---|---|---|---|
| Mvp | 0.078769 | 0.016289 | 1 |
| Mocs2 | 0.078737 | 5.68E-05 | 1 |
| Dis3l2 | 0.078713 | 0.000529 | 1 |
| Acyp2 | 0.078702 | 0.001286 | 1 |
| Wdr4 | 0.078695 | 0.000352 | 1 |
| Mcub | 0.078636 | 0.013475 | 1 |
| Atxn10 | 0.078615 | 0.019878 | 1 |
| Gmds | 0.078538 | 0.001175 | 1 |
| Tmem65 | 0.078414 | 9E-05 | 1 |
| Clec4e | 0.078388 | 0.655322 | 1 |
| Gm13203 | 0.078356 | 6.7E-06 | 0.21625 |
| Hsdl2 | 0.07818 | 0.005139 | 1 |
| Bcs1l | 0.078159 | 0.005768 | 1 |
| Cltb | 0.078089 | 4.55E-05 | 1 |
| Gnptab | 0.078077 | 0.008327 | 1 |
| Ehmt2 | 0.077978 | 0.008903 | 1 |
| Noc4l | 0.077954 | 0.00067 | 1 |
| Timm17a | 0.07788 | 0.003622 | 1 |
| Tmx1 | 0.077871 | 0.059214 | 1 |
| Dexi | 0.077833 | 0.000385 | 1 |
| Ptpn23 | 0.077822 | 2.97E-05 | 0.960224 |
| Ecsit | 0.077817 | 0.001714 | 1 |
| Dpy30 | 0.077745 | 0.006221 | 1 |
| Ccz1 | 0.077652 | 0.020908 | 1 |
| Eif4h | 0.077646 | 0.045473 | 1 |
| Hspbap1 | 0.07761 | 0.001514 | 1 |
| Cldnd1 | 0.077569 | 5.07E-05 | 1 |
| Fgfr1op | 0.077556 | 0.015147 | 1 |
| Ssrp1 | 0.077499 | 0.010897 | 1 |
| Smarca4 | 0.077488 | 0.1423 | 1 |
| Dera | 0.077437 | 0.01297 | 1 |
| Aacs | 0.077434 | 0.000275 | 1 |
| Uxt | 0.077409 | 0.001004 | 1 |
| Ipo5 | 0.077306 | 3.5E-05 | 1 |
| Msantd2 | 0.077178 | 0.004119 | 1 |
| Trp53inp2 | 0.077165 | 0.00028 | 1 |
| BC147527 | 0.077147 | 0.397873 | 1 |
| Slc25a13 | 0.077122 | 7.99E-06 | 0.257975 |
| Rnf213 | 0.077095 | 0.274707 | 1 |
| 2410022M | 0.077028 | 0.004954 | 1 |
| Osbpl7 | 0.077006 | 0.007035 | 1 |
| Psma1 | 0.077005 | 0.019305 | 1 |
| Rps11 | 0.076962 | 0.100689 | 1 |
| Fbh1 | 0.076949 | 0.001669 | 1 |
| Parn | 0.076929 | 0.001151 | 1 |
| Dcun1d5 | 0.076859 | 0.007011 | 1 |
| Ltv1 | 0.076799 | 0.000431 | 1 |
| Rad17 | 0.076794 | 0.05418 | 1 |

| | | | |
|---|---|---|---|
| Commd9 | 0.076791 | 0.000248 | 1 |
| Gm33370 | 0.076637 | 0.000843 | 1 |
| Chpt1 | 0.076562 | 0.000637 | 1 |
| Srgn | 0.076433 | 0.78997 | 1 |
| Yeats4 | 0.076315 | 0.001407 | 1 |
| Fn1 | 0.076261 | 2.41E-05 | 0.776494 |
| Slc25a12 | 0.07626 | 0.002487 | 1 |
| Glt8d1 | 0.076242 | 0.006755 | 1 |
| Lrrc42 | 0.076231 | 0.001756 | 1 |
| Phf5a | 0.076198 | 0.004107 | 1 |
| Calm2 | 0.075704 | 0.201605 | 1 |
| Kcnk13 | 0.075628 | 0.000846 | 1 |
| Gspt1 | 0.075625 | 0.006797 | 1 |
| Wdr44 | 0.075619 | 0.034197 | 1 |
| Pfdn4 | 0.075613 | 0.012781 | 1 |
| Tbl3 | 0.075603 | 0.006763 | 1 |
| Tent5c | 0.075573 | 3.87E-05 | 1 |
| Trim3 | 0.075512 | 2.83E-05 | 0.91319 |
| Sap18 | 0.075475 | 0.213628 | 1 |
| Rnf187 | 0.075466 | 0.002056 | 1 |
| Nudcd1 | 0.075322 | 4.56E-06 | 0.147349 |
| 6330562C2 | 0.075301 | 6.85E-06 | 0.221228 |
| Ifi208 | 0.075289 | 0.000131 | 1 |
| Pip4p1 | 0.075286 | 0.00252 | 1 |
| Ten1 | 0.075268 | 0.022625 | 1 |
| Rab12 | 0.075252 | 3.59E-05 | 1 |
| Mrps18c | 0.07521 | 0.010881 | 1 |
| Zc3h10 | 0.075197 | 0.033737 | 1 |
| Eef1aknmt | 0.075162 | 0.002797 | 1 |
| Spryd3 | 0.075131 | 5.13E-05 | 1 |
| Zfp787 | 0.075117 | 0.001441 | 1 |
| Asb11 | 0.075117 | 1.24E-06 | 0.040175 |
| Shq1 | 0.075117 | 0.001029 | 1 |
| Setd1a | 0.075114 | 0.020643 | 1 |
| Faf2 | 0.075027 | 0.001627 | 1 |
| Eny2 | 0.075003 | 0.001245 | 1 |
| Aamdc | 0.074961 | 0.001057 | 1 |
| Mrpl27 | 0.074904 | 2.13E-05 | 0.686311 |
| Dnajc9 | 0.074901 | 0.077247 | 1 |
| Lman1 | 0.074851 | 7.5E-05 | 1 |
| Nos1ap | 0.074788 | 2.42E-05 | 0.781332 |
| Dcaf4 | 0.074777 | 7.44E-05 | 1 |
| Ptpn2 | 0.074749 | 0.000609 | 1 |
| Lrrc4 | 0.074734 | 2.81E-05 | 0.907166 |
| Agpat2 | 0.074651 | 0.000462 | 1 |
| Ctdspl | 0.074647 | 0.000621 | 1 |
| Brap | 0.074636 | 0.032581 | 1 |
| Ilf2 | 0.074629 | 0.011705 | 1 |

| | | | |
|---|---|---|---|
| Ap4b1 | 0.074623 | 0.013938 | 1 |
| Drg1 | 0.074605 | 0.033916 | 1 |
| Gadd45gip | 0.074584 | 9.43E-05 | 1 |
| Timm8a1 | 0.074571 | 9.93E-05 | 1 |
| Rnf145 | 0.074481 | 0.004921 | 1 |
| Gca | 0.074391 | 4.57E-05 | 1 |
| Rhbdd2 | 0.07439 | 0.005472 | 1 |
| Psmb5 | 0.074323 | 0.072134 | 1 |
| Ccdc58 | 0.074268 | 0.001117 | 1 |
| Washc1 | 0.074224 | 0.003724 | 1 |
| 5430416N( | 0.07415 | 0.002106 | 1 |
| Ppp1r7 | 0.074113 | 0.055763 | 1 |
| Blcap | 0.074098 | 0.002843 | 1 |
| Cars | 0.074072 | 0.000557 | 1 |
| Mrpl50 | 0.074005 | 0.000112 | 1 |
| Rfxank | 0.073869 | 0.004731 | 1 |
| Galnt11 | 0.073817 | 0.004323 | 1 |
| Fdxr | 0.073804 | 0.025478 | 1 |
| Twsg1 | 0.073722 | 3.41E-05 | 1 |
| Nipsnap2 | 0.073707 | 0.002215 | 1 |
| Lars | 0.073684 | 0.000654 | 1 |
| Cep120 | 0.073637 | 0.000601 | 1 |
| Alg8 | 0.073446 | 0.007916 | 1 |
| 4930405A1 | 0.073369 | 0.004946 | 1 |
| Tg | 0.07335 | 0.003959 | 1 |
| Emc6 | 0.073335 | 0.015218 | 1 |
| Ufc1 | 0.073212 | 0.023625 | 1 |
| Mmp8 | 0.073147 | 0.093328 | 1 |
| Tacc3 | 0.073142 | 0.036678 | 1 |
| Arl4a | 0.073121 | 0.000113 | 1 |
| Mterf4 | 0.073116 | 0.000504 | 1 |
| Kifap3 | 0.073097 | 3.59E-05 | 1 |
| Tmem67 | 0.073018 | 0.000281 | 1 |
| Rnf169 | 0.072987 | 0.101931 | 1 |
| Gm17494 | 0.072936 | 0.00623 | 1 |
| Pld1 | 0.072853 | 0.063711 | 1 |
| Sf3b4 | 0.072814 | 0.078989 | 1 |
| Mrpl47 | 0.072768 | 0.000495 | 1 |
| Fam104a | 0.072738 | 0.009541 | 1 |
| Foxk2 | 0.072587 | 0.004552 | 1 |
| Zfp691 | 0.07245 | 0.01664 | 1 |
| Pard6a | 0.072395 | 0.000527 | 1 |
| Adsl | 0.072393 | 0.000122 | 1 |
| Psmd6 | 0.072345 | 0.003987 | 1 |
| Slc30a7 | 0.072343 | 0.000453 | 1 |
| Poldip2 | 0.07229 | 3.81E-05 | 1 |
| Mov10 | 0.072202 | 0.000368 | 1 |
| Aga | 0.071971 | 0.016425 | 1 |

| | | | |
|---|---|---|---|
| Agfg2 | 0.071971 | 0.000286 | 1 |
| Sdf4 | 0.071926 | 0.178172 | 1 |
| Thoc3 | 0.071915 | 0.020962 | 1 |
| Rcor1 | 0.071874 | 0.002482 | 1 |
| Get4 | 0.071868 | 3.02E-05 | 0.974054 |
| Pik3r2 | 0.071831 | 0.002145 | 1 |
| Zmym6 | 0.07171 | 0.022015 | 1 |
| Zfp810 | 0.07165 | 0.000768 | 1 |
| Gmps | 0.071601 | 0.014171 | 1 |
| Ankrd37 | 0.071543 | 0.002524 | 1 |
| Clcn5 | 0.071539 | 0.00418 | 1 |
| Tmem134 | 0.071527 | 0.139538 | 1 |
| Dpysl2 | 0.071478 | 0.002457 | 1 |
| Asah2 | 0.071379 | 0.001244 | 1 |
| Psmc5 | 0.071267 | 0.035795 | 1 |
| Irs2 | 0.071261 | 0.116271 | 1 |
| Pmm1 | 0.071235 | 0.000146 | 1 |
| Nup205 | 0.071098 | 0.001572 | 1 |
| Txndc9 | 0.071045 | 0.002655 | 1 |
| Riok3 | 0.071003 | 0.095467 | 1 |
| Plekhm2 | 0.070974 | 0.000234 | 1 |
| Slc52a2 | 0.070968 | 0.000122 | 1 |
| Rdm1 | 0.07088 | 0.221267 | 1 |
| Mul1 | 0.070869 | 0.001087 | 1 |
| Tbc1d14 | 0.07085 | 0.011701 | 1 |
| Zdhhc21 | 0.070804 | 4.5E-05 | 1 |
| Tnfaip8l2 | 0.070718 | 0.125543 | 1 |
| Arl2 | 0.070685 | 0.000992 | 1 |
| Riok1 | 0.070584 | 0.081249 | 1 |
| Rnaseh2b | 0.070535 | 0.000418 | 1 |
| Nubpl | 0.070525 | 0.00041 | 1 |
| Nol7 | 0.070469 | 0.000536 | 1 |
| Ext2 | 0.070369 | 0.010781 | 1 |
| Atad5 | 0.070357 | 0.008472 | 1 |
| Ccnd1 | 0.070303 | 5.1E-06 | 0.164519 |
| Ndufs2 | 0.070291 | 0.04083 | 1 |
| Stk19 | 0.070285 | 0.001882 | 1 |
| Stx12 | 0.070234 | 0.000564 | 1 |
| Epg5 | 0.070219 | 0.002218 | 1 |
| Cp | 0.070214 | 0.011245 | 1 |
| Gmppb | 0.070178 | 0.042448 | 1 |
| Poc1a | 0.07017 | 0.028306 | 1 |
| Chka | 0.070124 | 5.39E-05 | 1 |
| Exosc2 | 0.07009 | 0.000638 | 1 |
| Arl10 | 0.070054 | 0.281914 | 1 |
| Rbbp8 | 0.069954 | 0.137851 | 1 |
| Grwd1 | 0.069948 | 0.00012 | 1 |
| Clmp | 0.069914 | 0.000289 | 1 |

| | | | |
|---|---|---|---|
| Alkbh1 | 0.069902 | 0.037661 | 1 |
| Cebpe | 0.069899 | 0.008918 | 1 |
| Hsp90aa1 | 0.069896 | 0.084608 | 1 |
| Relt | 0.069866 | 0.116638 | 1 |
| Fam3a | 0.069866 | 0.008057 | 1 |
| Washc5 | 0.069829 | 0.002331 | 1 |
| Lsm12 | 0.069743 | 0.012468 | 1 |
| Ndst2 | 0.069726 | 6.92E-05 | 1 |
| Nfkbil1 | 0.06964 | 0.025927 | 1 |
| Nomo1 | 0.069566 | 0.000712 | 1 |
| Rnf220 | 0.069562 | 0.014591 | 1 |
| Pold4 | 0.069556 | 0.016944 | 1 |
| Pias4 | 0.069513 | 0.002435 | 1 |
| Tmed1 | 0.069414 | 0.00142 | 1 |
| Slc6a6 | 0.069401 | 0.003905 | 1 |
| Nmt2 | 0.06931 | 0.000226 | 1 |
| Mrm1 | 0.069181 | 0.002514 | 1 |
| Stard7 | 0.069118 | 0.006208 | 1 |
| Sf3a2 | 0.069042 | 0.031078 | 1 |
| Pml | 0.068965 | 0.00698 | 1 |
| Znfx1 | 0.068916 | 0.000965 | 1 |
| Slc30a9 | 0.068736 | 6.78E-06 | 0.218903 |
| Tecpr2 | 0.068661 | 0.002198 | 1 |
| Trp53i13 | 0.068659 | 0.012053 | 1 |
| Pop4 | 0.068585 | 6.51E-05 | 1 |
| Bbs9 | 0.068528 | 0.090382 | 1 |
| Hs2st1 | 0.068527 | 0.005756 | 1 |
| Ino80e | 0.06851 | 0.006097 | 1 |
| Cttnbp2nl | 0.068447 | 3.77E-06 | 0.12159 |
| Hyal2 | 0.068436 | 0.000662 | 1 |
| Mmadhc | 0.068429 | 0.000485 | 1 |
| Armc10 | 0.068425 | 0.044237 | 1 |
| Smc5 | 0.068414 | 0.007659 | 1 |
| Rab18 | 0.068356 | 0.002299 | 1 |
| Them6 | 0.06816 | 0.010683 | 1 |
| Tspan32 | 0.068138 | 0.00152 | 1 |
| Cad | 0.068103 | 0.0002 | 1 |
| Bcar3 | 0.068058 | 0.000132 | 1 |
| Ch25h | 0.068019 | 2.13E-05 | 0.686504 |
| Psme3 | 0.067924 | 0.001275 | 1 |
| Plppr3 | 0.06791 | 0.000144 | 1 |
| Gm5577 | 0.067867 | 0.001103 | 1 |
| Abtb2 | 0.067823 | 0.003888 | 1 |
| Gfm1 | 0.067782 | 5.04E-06 | 0.162727 |
| Ogfod3 | 0.067763 | 5.55E-05 | 1 |
| Srrd | 0.067762 | 0.006624 | 1 |
| Mrps11 | 0.06776 | 0.011716 | 1 |
| Pdk3 | 0.067749 | 0.002743 | 1 |

| | | | |
|---|---|---|---|
| Tpd52l2 | 0.067717 | 0.206004 | 1 |
| Unc45a | 0.067683 | 0.001429 | 1 |
| Dip2a | 0.067677 | 0.009408 | 1 |
| Uba1 | 0.067622 | 0.035696 | 1 |
| Cs | 0.06757 | 0.176642 | 1 |
| Hnrnpc | 0.067485 | 0.168322 | 1 |
| Eif3e | 0.067347 | 0.002527 | 1 |
| Ctbp1 | 0.067288 | 0.000706 | 1 |
| Pbdc1 | 0.067204 | 0.01355 | 1 |
| Mvb12a | 0.067163 | 0.001295 | 1 |
| Nudt6 | 0.067098 | 0.001803 | 1 |
| Nudt8 | 0.067065 | 0.000909 | 1 |
| Dph1 | 0.066948 | 0.032373 | 1 |
| Zc3h12c | 0.066918 | 0.301146 | 1 |
| Znhit6 | 0.066864 | 6.46E-05 | 1 |
| Batf | 0.06676 | 0.134948 | 1 |
| Fam117a | 0.066739 | 0.01134 | 1 |
| Cd81 | 0.066684 | 0.001691 | 1 |
| Bcl2l2 | 0.06661 | 0.000287 | 1 |
| Ola1 | 0.066583 | 0.006205 | 1 |
| Txnl4a | 0.066579 | 0.008411 | 1 |
| Ebna1bp2 | 0.066559 | 0.065928 | 1 |
| Fdx1 | 0.066551 | 0.001764 | 1 |
| Apool | 0.066523 | 6.28E-06 | 0.202821 |
| Castor2 | 0.066466 | 2.27E-05 | 0.732762 |
| Prpf38a | 0.066444 | 0.00287 | 1 |
| Rala | 0.066414 | 0.003334 | 1 |
| Ubxn4 | 0.066408 | 0.049562 | 1 |
| Wdr73 | 0.066363 | 0.0014 | 1 |
| Trappc2l | 0.066282 | 0.008951 | 1 |
| Racgap1 | 0.066226 | 0.309931 | 1 |
| Emc4 | 0.066191 | 0.007572 | 1 |
| Chmp7 | 0.066104 | 0.005386 | 1 |
| 2310040G: | 0.066076 | 0.001077 | 1 |
| Dis3 | 0.066009 | 9.66E-06 | 0.31172 |
| Rp9 | 0.065985 | 0.083615 | 1 |
| Rac1 | 0.065791 | 0.289108 | 1 |
| Keap1 | 0.065767 | 0.052422 | 1 |
| Stk39 | 0.065748 | 0.000288 | 1 |
| Rsl24d1 | 0.065663 | 0.009727 | 1 |
| Rps14 | 0.065611 | 0.194149 | 1 |
| Wdr1 | 0.065592 | 0.242374 | 1 |
| Brox | 0.065576 | 0.047503 | 1 |
| Fbxo33 | 0.065567 | 0.000566 | 1 |
| Iqsec1 | 0.065546 | 1.04E-05 | 0.336681 |
| Dpm3 | 0.065491 | 0.109417 | 1 |
| Sae1 | 0.065445 | 0.00948 | 1 |
| Palm | 0.065424 | 0.003009 | 1 |

| | | | |
|---|---|---|---|
| Tulp4 | 0.065229 | 0.001126 | 1 |
| Plpp5 | 0.065143 | 0.000235 | 1 |
| Mad2l1 | 0.065137 | 0.000533 | 1 |
| Mapkbp1 | 0.065086 | 2.71E-05 | 0.87573 |
| Cby1 | 0.064939 | 0.001007 | 1 |
| Ptger2 | 0.064907 | 3.17E-06 | 0.102409 |
| Fen1 | 0.064851 | 0.01202 | 1 |
| Zswim8 | 0.064821 | 0.003636 | 1 |
| Rpl6 | 0.064819 | 0.072401 | 1 |
| Ik | 0.06477 | 0.043502 | 1 |
| Kdm4b | 0.064694 | 0.000269 | 1 |
| Dicer1 | 0.064615 | 0.002383 | 1 |
| Mob3c | 0.0646 | 0.00031 | 1 |
| Gart | 0.064591 | 3.53E-05 | 1 |
| Ipo9 | 0.064583 | 0.000276 | 1 |
| Yars | 0.064542 | 0.061682 | 1 |
| Mcemp1 | 0.064472 | 0.006038 | 1 |
| Dnajc4 | 0.064352 | 0.001567 | 1 |
| Wdr77 | 0.064309 | 0.003542 | 1 |
| Rbm45 | 0.064264 | 0.199239 | 1 |
| Jazf1 | 0.064232 | 0.003144 | 1 |
| AI504432 | 0.064143 | 0.006724 | 1 |
| Ctps | 0.064019 | 8.34E-05 | 1 |
| Tmco3 | 0.064018 | 0.001338 | 1 |
| Wipi2 | 0.063949 | 0.003476 | 1 |
| Zfp771 | 0.06393 | 0.004714 | 1 |
| Idh1 | 0.063928 | 0.035052 | 1 |
| Tmem248 | 0.063857 | 0.003769 | 1 |
| Klhdc3 | 0.063849 | 0.009363 | 1 |
| Cherp | 0.063769 | 0.016869 | 1 |
| Ubfd1 | 0.063673 | 0.001075 | 1 |
| Paqr7 | 0.063661 | 0.001289 | 1 |
| Ttpal | 0.063632 | 0.00425 | 1 |
| Ergic2 | 0.063617 | 0.002645 | 1 |
| Sepsecs | 0.063545 | 5.71E-05 | 1 |
| Rpl22 | 0.063539 | 0.035479 | 1 |
| E230016K2 | 0.063531 | 0.014369 | 1 |
| Ppih | 0.063512 | 0.001195 | 1 |
| Fech | 0.063507 | 0.002234 | 1 |
| Cfap36 | 0.063504 | 9.07E-05 | 1 |
| Kdm2b | 0.063399 | 0.040506 | 1 |
| Abca1 | 0.063303 | 0.001367 | 1 |
| Bax | 0.063245 | 0.014254 | 1 |
| Trit1 | 0.063203 | 0.001233 | 1 |
| Gpr160 | 0.063187 | 0.013181 | 1 |
| Dcp2 | 0.063179 | 0.003777 | 1 |
| Mcoln1 | 0.063113 | 0.010393 | 1 |
| Galnt2 | 0.063088 | 0.001568 | 1 |

| | | | |
|---|---|---|---|
| Sppl2b | 0.06306 | 0.000374 | 1 |
| Ttll5 | 0.062999 | 0.004688 | 1 |
| Aptx | 0.062903 | 0.024855 | 1 |
| Hdac1 | 0.062872 | 0.027022 | 1 |
| Tmem37 | 0.06287 | 0.014195 | 1 |
| Erg28 | 0.062826 | 0.006352 | 1 |
| Man2a2 | 0.062823 | 0.010839 | 1 |
| Mapre2 | 0.062813 | 0.173486 | 1 |
| Nudt2 | 0.062791 | 0.000113 | 1 |
| Ddx18 | 0.062785 | 0.00072 | 1 |
| Kat2a | 0.062763 | 0.003737 | 1 |
| Dusp19 | 0.062714 | 5.48E-05 | 1 |
| Msh3 | 0.062624 | 0.001117 | 1 |
| Wdr11 | 0.062519 | 0.002127 | 1 |
| Tmem167 | 0.062511 | 0.022511 | 1 |
| Odf2 | 0.06241 | 0.000252 | 1 |
| Ift57 | 0.062403 | 0.00538 | 1 |
| Galm | 0.062367 | 0.000393 | 1 |
| Tmem260 | 0.062281 | 0.000152 | 1 |
| Gtpbp2 | 0.06227 | 0.007031 | 1 |
| Lemd2 | 0.06223 | 0.001181 | 1 |
| Atpaf1 | 0.062221 | 0.04025 | 1 |
| Tgs1 | 0.062217 | 0.01067 | 1 |
| Ints12 | 0.062176 | 0.010927 | 1 |
| Tnfsf9 | 0.062063 | 0.00018 | 1 |
| Myh10 | 0.062018 | 0.000131 | 1 |
| Mapk1ip1l | 0.061921 | 0.037597 | 1 |
| Bzw2 | 0.061919 | 0.004578 | 1 |
| Srp68 | 0.061848 | 0.001844 | 1 |
| Nit1 | 0.061759 | 0.022276 | 1 |
| Rab35 | 0.061708 | 0.040648 | 1 |
| Tmtc4 | 0.061638 | 5.58E-05 | 1 |
| Kdelr1 | 0.061635 | 0.018022 | 1 |
| Zfyve19 | 0.061624 | 0.001339 | 1 |
| Nmnat3 | 0.061593 | 0.005721 | 1 |
| Unc93b1 | 0.061571 | 0.118827 | 1 |
| Incenp | 0.061434 | 0.011632 | 1 |
| Rpp21 | 0.061411 | 0.111431 | 1 |
| Sugt1 | 0.061392 | 0.024148 | 1 |
| Rnf121 | 0.061347 | 0.008689 | 1 |
| Tubg1 | 0.061346 | 0.002416 | 1 |
| Ide | 0.061329 | 0.002398 | 1 |
| Trpm4 | 0.061326 | 0.003022 | 1 |
| Spryd7 | 0.061311 | 0.001035 | 1 |
| Gas8 | 0.061295 | 0.000403 | 1 |
| Coa6 | 0.061258 | 0.020159 | 1 |
| Fnip2 | 0.061251 | 2.75E-05 | 0.888583 |
| Arfgap1 | 0.061212 | 0.019948 | 1 |

| | | | |
|---|---|---|---|
| Dcxr | 0.061144 | 0.045571 | 1 |
| Agpat5 | 0.061112 | 3.63E-05 | 1 |
| Mfsd7a | 0.061096 | 0.013788 | 1 |
| Polr1a | 0.061047 | 4.63E-05 | 1 |
| Natd1 | 0.061028 | 0.000677 | 1 |
| Cd3eap | 0.060947 | 0.003295 | 1 |
| Nom1 | 0.060936 | 0.000561 | 1 |
| Mcrs1 | 0.060869 | 0.020773 | 1 |
| Dph6 | 0.060838 | 0.003542 | 1 |
| Zcchc9 | 0.06079 | 0.002148 | 1 |
| Uri1 | 0.06077 | 0.000171 | 1 |
| Fam102b | 0.060762 | 0.004031 | 1 |
| Suv39h1 | 0.060713 | 0.000771 | 1 |
| Oas1g | 0.060637 | 0.019437 | 1 |
| Zbtb5 | 0.060609 | 0.000291 | 1 |
| Atp6v1c1 | 0.060552 | 0.000158 | 1 |
| As3mt | 0.060531 | 0.002067 | 1 |
| Megf8 | 0.060391 | 0.001397 | 1 |
| Arpc1a | 0.06039 | 0.050627 | 1 |
| Ctdp1 | 0.060284 | 0.000264 | 1 |
| Zfp951 | 0.060215 | 0.023339 | 1 |
| Mzt1 | 0.0602 | 0.002896 | 1 |
| Jagn1 | 0.060177 | 0.008959 | 1 |
| Seh1l | 0.060132 | 0.000132 | 1 |
| Nsmce1 | 0.060094 | 0.007248 | 1 |
| Gtf2i | 0.06 | 0.031508 | 1 |
| Rce1 | 0.059977 | 2.23E-05 | 0.718943 |
| Emp1 | 0.059959 | 0.006078 | 1 |
| Smim14 | 0.059922 | 0.048764 | 1 |
| Dusp3 | 0.0599 | 0.002931 | 1 |
| Fer | 0.05989 | 0.000239 | 1 |
| 5730480H( | 0.059885 | 0.001383 | 1 |
| Triobp | 0.05982 | 0.005413 | 1 |
| Leo1 | 0.059799 | 0.027184 | 1 |
| Adi1 | 0.059715 | 2.63E-05 | 0.849483 |
| Gm14636 | 0.059696 | 0.000101 | 1 |
| Ccl25 | 0.059673 | 0.057109 | 1 |
| Epn1 | 0.059593 | 0.116055 | 1 |
| Sdr39u1 | 0.059461 | 0.001894 | 1 |
| Dnajc2 | 0.059456 | 0.002813 | 1 |
| Pex11a | 0.059434 | 0.000304 | 1 |
| Rps19 | 0.059433 | 0.097374 | 1 |
| Qrsl1 | 0.059311 | 0.048346 | 1 |
| Entr1 | 0.059306 | 0.244066 | 1 |
| Chid1 | 0.059187 | 1.76E-05 | 0.566885 |
| Etf1 | 0.059184 | 0.350421 | 1 |
| Calu | 0.05917 | 0.00142 | 1 |
| Elovl1 | 0.059041 | 0.048379 | 1 |

| | | | |
|---|---|---|---|
| Fam122a | 0.059007 | 0.00097 | 1 |
| Thop1 | 0.059007 | 0.009342 | 1 |
| Rrp9 | 0.058924 | 0.000175 | 1 |
| Tmem129 | 0.058874 | 0.000155 | 1 |
| Smim26 | 0.058818 | 0.000323 | 1 |
| Hectd3 | 0.058761 | 0.031456 | 1 |
| Pex16 | 0.058713 | 0.001845 | 1 |
| Ada | 0.058702 | 0.00205 | 1 |
| Mast2 | 0.058647 | 0.001498 | 1 |
| Rabac1 | 0.058618 | 0.060556 | 1 |
| R3hcc1 | 0.058553 | 0.000564 | 1 |
| Orai3 | 0.058529 | 0.019659 | 1 |
| Kcnk6 | 0.058511 | 0.00652 | 1 |
| Tmem126b | 0.058435 | 0.003216 | 1 |
| Ccnc | 0.058324 | 0.088418 | 1 |
| Mipep | 0.058298 | 0.009791 | 1 |
| Akip1 | 0.058219 | 0.003652 | 1 |
| Farp2 | 0.058182 | 0.000175 | 1 |
| Cdk10 | 0.058178 | 0.000844 | 1 |
| Smap1 | 0.058166 | 0.031988 | 1 |
| Slc4a1ap | 0.058056 | 0.015835 | 1 |
| Znhit1 | 0.057971 | 0.040391 | 1 |
| Fam110a | 0.057944 | 3.45E-06 | 0.111391 |
| Cpt2 | 0.057922 | 0.070246 | 1 |
| Mllt3 | 0.057911 | 0.015156 | 1 |
| St6galnac6 | 0.05791 | 0.001597 | 1 |
| Acot7 | 0.057829 | 0.000475 | 1 |
| Dazap1 | 0.057819 | 0.01651 | 1 |
| Lrrc14 | 0.057784 | 9.24E-05 | 1 |
| Parp6 | 0.057759 | 7.76E-05 | 1 |
| Syap1 | 0.057703 | 0.230447 | 1 |
| Rpl7l1 | 0.057602 | 0.079261 | 1 |
| Twf1 | 0.057595 | 0.002693 | 1 |
| Acad8 | 0.057581 | 0.045825 | 1 |
| Trp53rka | 0.057553 | 0.020763 | 1 |
| Dkc1 | 0.057486 | 0.000596 | 1 |
| Clptm1 | 0.057467 | 0.026477 | 1 |
| Blzf1 | 0.057337 | 0.002747 | 1 |
| Mrpl9 | 0.05731 | 0.009921 | 1 |
| Selenbp1 | 0.057298 | 0.023857 | 1 |
| Pole3 | 0.057282 | 0.002287 | 1 |
| Alg13 | 0.05725 | 0.00767 | 1 |
| Mrpl45 | 0.057194 | 0.000665 | 1 |
| Eef2 | 0.057191 | 0.057739 | 1 |
| Dgcr8 | 0.057135 | 0.000678 | 1 |
| Tlr1 | 0.056968 | 0.005509 | 1 |
| Pafah1b2 | 0.056957 | 0.002955 | 1 |
| Selenon | 0.056913 | 0.017237 | 1 |

| | | | |
|---|---|---|---|
| Snapc2 | 0.056888 | 0.001499 | 1 |
| Taf4b | 0.056879 | 0.000431 | 1 |
| Rab43 | 0.056824 | 0.029322 | 1 |
| 1110059E2 | 0.056754 | 0.001 | 1 |
| Xpo1 | 0.056621 | 0.044447 | 1 |
| Zfp664 | 0.0566 | 0.089514 | 1 |
| Phospho2 | 0.056595 | 0.000158 | 1 |
| Nol6 | 0.056586 | 0.00175 | 1 |
| Slc35e4 | 0.056585 | 0.023828 | 1 |
| Pkd2 | 0.056537 | 0.000249 | 1 |
| Tardbp | 0.056445 | 0.011256 | 1 |
| Fam78a | 0.056408 | 0.005806 | 1 |
| Eif4e | 0.056343 | 0.11012 | 1 |
| Cnot9 | 0.056343 | 0.000693 | 1 |
| Sigmar1 | 0.056311 | 0.001664 | 1 |
| Cebpzos | 0.05625 | 0.000932 | 1 |
| Dimt1 | 0.056243 | 0.004101 | 1 |
| Ift22 | 0.056214 | 0.002025 | 1 |
| Rusc2 | 0.056172 | 0.001979 | 1 |
| Acy1 | 0.056165 | 5.63E-05 | 1 |
| Pfkfb2 | 0.056062 | 0.002289 | 1 |
| Psmd10 | 0.056022 | 2.56E-05 | 0.826006 |
| Atraid | 0.056008 | 0.013917 | 1 |
| Dennd1a | 0.055984 | 0.320898 | 1 |
| Dmap1 | 0.055957 | 0.001119 | 1 |
| H2-Oa | 0.055937 | 0.000518 | 1 |
| Gas5 | 0.055922 | 0.858359 | 1 |
| Ubl5 | 0.055915 | 0.358134 | 1 |
| H2afx | 0.055809 | 0.322635 | 1 |
| Apoc2 | 0.05569 | 0.568736 | 1 |
| Tecpr1 | 0.055685 | 0.002997 | 1 |
| Klc4 | 0.05562 | 1.32E-06 | 0.042516 |
| Nvl | 0.055604 | 0.002022 | 1 |
| Zwint | 0.055539 | 0.00052 | 1 |
| Polr1b | 0.055483 | 0.005186 | 1 |
| Clcn6 | 0.055468 | 0.010325 | 1 |
| Kdm1a | 0.055414 | 0.036933 | 1 |
| Eif2b5 | 0.055398 | 0.00126 | 1 |
| Gmcl1 | 0.055343 | 0.006458 | 1 |
| Zfp202 | 0.055334 | 0.000104 | 1 |
| Dbt | 0.0553 | 0.01573 | 1 |
| Slc35e1 | 0.055268 | 0.005963 | 1 |
| Med12l | 0.055218 | 0.001013 | 1 |
| Tm2d2 | 0.055198 | 0.008769 | 1 |
| Glb1 | 0.055192 | 0.000676 | 1 |
| Ptov1 | 0.05519 | 0.190184 | 1 |
| Abcf1 | 0.055103 | 0.092574 | 1 |
| D430042O | 0.05501 | 0.0023 | 1 |

| | | | |
|---|---|---|---|
| Tlr3 | 0.05494 | 0.000683 | 1 |
| Tarbp2 | 0.054891 | 0.000128 | 1 |
| Mpv17 | 0.054875 | 0.077384 | 1 |
| Pgrmc2 | 0.054838 | 0.000255 | 1 |
| Mzt2 | 0.054798 | 0.014663 | 1 |
| Mrps2 | 0.054792 | 0.002597 | 1 |
| Ift88 | 0.054766 | 3.52E-05 | 1 |
| Farsb | 0.05473 | 0.005163 | 1 |
| Ttc13 | 0.054675 | 0.001507 | 1 |
| Shb | 0.054666 | 0.010105 | 1 |
| Ftsj3 | 0.054647 | 0.000165 | 1 |
| Rcbtb2 | 0.054601 | 0.071329 | 1 |
| Bud23 | 0.054591 | 0.006325 | 1 |
| Pigq | 0.054579 | 0.005276 | 1 |
| Grpel1 | 0.05455 | 0.007641 | 1 |
| Napa | 0.0545 | 0.011577 | 1 |
| Slc22a18 | 0.054486 | 0.000154 | 1 |
| Ccdc90b | 0.054477 | 0.001273 | 1 |
| Coa7 | 0.054474 | 2.75E-05 | 0.88795 |
| Tjp2 | 0.054429 | 0.009838 | 1 |
| Cables2 | 0.054429 | 0.072081 | 1 |
| Birc2 | 0.05442 | 0.002561 | 1 |
| Mrpl21 | 0.054383 | 0.006735 | 1 |
| Rabepk | 0.054341 | 0.00085 | 1 |
| Gins2 | 0.054266 | 0.001288 | 1 |
| Tmem63b | 0.054265 | 0.001043 | 1 |
| Pprc1 | 0.054092 | 0.000772 | 1 |
| Coro1b | 0.054053 | 0.282282 | 1 |
| Eif2b4 | 0.054019 | 0.008202 | 1 |
| Hmg20b | 0.053983 | 0.062055 | 1 |
| Dusp22 | 0.05398 | 0.032638 | 1 |
| Stoml1 | 0.053972 | 0.007385 | 1 |
| Wdr91 | 0.053892 | 0.00028 | 1 |
| Atp6v0a2 | 0.053874 | 0.02199 | 1 |
| Coq6 | 0.053839 | 0.18525 | 1 |
| Cmtm3 | 0.053807 | 0.007274 | 1 |
| Gm11361 | 0.053746 | 0.000892 | 1 |
| Tmco4 | 0.053665 | 0.00071 | 1 |
| Mfsd8 | 0.053632 | 0.110142 | 1 |
| Gga2 | 0.053525 | 0.002053 | 1 |
| Cnep1r1 | 0.053492 | 0.006257 | 1 |
| Rxrb | 0.053491 | 0.001539 | 1 |
| Nudt5 | 0.053471 | 6.03E-05 | 1 |
| 5031425F1 | 0.053469 | 0.005911 | 1 |
| Sgsm3 | 0.053424 | 0.000384 | 1 |
| Tasp1 | 0.053362 | 0.000246 | 1 |
| Guf1 | 0.053291 | 4.19E-05 | 1 |
| Prmt7 | 0.053174 | 0.00032 | 1 |

| | | | |
|---|---|---|---|
| Ndrg1 | 0.053173 | 0.062417 | 1 |
| BC029722 | 0.053146 | 0.013841 | 1 |
| Emc7 | 0.05314 | 0.038692 | 1 |
| Pdap1 | 0.053122 | 0.033756 | 1 |
| Ppcs | 0.053096 | 0.005225 | 1 |
| F5 | 0.053068 | 0.113697 | 1 |
| Elp5 | 0.05305 | 0.021796 | 1 |
| Exoc5 | 0.052925 | 0.001023 | 1 |
| Mir22hg | 0.052793 | 0.01978 | 1 |
| Katnb1 | 0.052759 | 7.08E-06 | 0.228738 |
| Nup62 | 0.052692 | 0.00287 | 1 |
| Gm17018 | 0.052687 | 0.001501 | 1 |
| Ahctf1 | 0.05264 | 0.000962 | 1 |
| Ctdsp2 | 0.052628 | 0.046187 | 1 |
| Cdc37 | 0.052626 | 0.183011 | 1 |
| Bloc1s6 | 0.052578 | 0.001858 | 1 |
| G6pc3 | 0.052552 | 0.010245 | 1 |
| Adat2 | 0.052511 | 0.009219 | 1 |
| Sf3a3 | 0.05232 | 0.010324 | 1 |
| Mbnl2 | 0.052192 | 0.377724 | 1 |
| Samd1 | 0.052171 | 0.034503 | 1 |
| Cds1 | 0.052058 | 0.000725 | 1 |
| Oasl2 | 0.051969 | 0.095406 | 1 |
| Lyrm9 | 0.051956 | 0.009378 | 1 |
| Slc12a4 | 0.051907 | 0.000257 | 1 |
| Ndufa7 | 0.051889 | 0.092151 | 1 |
| Tbc1d15 | 0.051888 | 0.015736 | 1 |
| March8 | 0.051843 | 0.000819 | 1 |
| Dnajc12 | 0.051827 | 0.002097 | 1 |
| Josd2 | 0.051765 | 0.077539 | 1 |
| Mtrf1l | 0.051713 | 0.033028 | 1 |
| Hagh | 0.051697 | 0.002478 | 1 |
| Pced1b | 0.051647 | 0.000374 | 1 |
| Ciao1 | 0.051566 | 0.014621 | 1 |
| Gm46224 | 0.051552 | 0.054357 | 1 |
| Prdx2 | 0.051492 | 0.30939 | 1 |
| B020010K1 | 0.05146 | 0.002011 | 1 |
| Dnaja1 | 0.05136 | 0.029652 | 1 |
| Golph3 | 0.051306 | 0.056325 | 1 |
| Rab44 | 0.051145 | 0.005771 | 1 |
| Zzz3 | 0.051106 | 0.010714 | 1 |
| Mier2 | 0.051074 | 0.386392 | 1 |
| Rps9 | 0.051074 | 0.553004 | 1 |
| Pfdn2 | 0.051071 | 0.008684 | 1 |
| Gpatch1 | 0.050997 | 0.003733 | 1 |
| Xrcc6 | 0.050975 | 0.010339 | 1 |
| Taf12 | 0.050894 | 0.009388 | 1 |
| Rps16 | 0.050834 | 0.30359 | 1 |

| | | | |
|---|---|---|---|
| Tsnax | 0.050783 | 0.006459 | 1 |
| Zfp384 | 0.05069 | 0.000112 | 1 |
| Oxa1l | 0.050676 | 0.008893 | 1 |
| Taf11 | 0.050631 | 0.005785 | 1 |
| Ercc6l2 | 0.05056 | 0.002172 | 1 |
| Snx25 | 0.050544 | 0.000731 | 1 |
| Polr1e | 0.050495 | 0.001327 | 1 |
| Rtl5 | 0.050422 | 0.004374 | 1 |
| Fnbp1l | 0.050402 | 0.002052 | 1 |
| Usp31 | 0.050377 | 0.004286 | 1 |
| P2rx7 | 0.050346 | 0.133835 | 1 |
| 2210016L2 | 0.050314 | 0.005793 | 1 |
| Psmg2 | 0.050301 | 8.17E-05 | 1 |
| Cep162 | 0.050294 | 0.01343 | 1 |
| Mrpl18 | 0.050286 | 0.014014 | 1 |
| Mob1b | 0.050219 | 0.020452 | 1 |
| Crtap | 0.050188 | 0.012512 | 1 |
| Nmd3 | 0.050086 | 0.025185 | 1 |
| Tsg101 | 0.050082 | 0.031576 | 1 |
| Ptdss2 | 0.050059 | 0.044076 | 1 |
| Dnajc16 | 0.050054 | 0.000333 | 1 |
| Cul9 | 0.050043 | 0.002899 | 1 |
| Znhit3 | 0.050003 | 0.000178 | 1 |
| Trim47 | 0.05 | 0.004803 | 1 |
| Slc44a1 | 0.049996 | 0.067516 | 1 |
| Tns3 | 0.049993 | 0.169104 | 1 |
| Tsfm | 0.049991 | 0.002342 | 1 |
| Tnfsf13os | 0.049982 | 0.166033 | 1 |
| Rnmt | 0.04993 | 0.005642 | 1 |
| Fmr1 | 0.049863 | 0.002565 | 1 |
| Pcbp2 | 0.04986 | 0.28687 | 1 |
| Chsy1 | 0.04984 | 0.003712 | 1 |
| Rsph3b | 0.049807 | 0.031263 | 1 |
| Tbc1d9 | 0.049771 | 0.069691 | 1 |
| H13 | 0.049768 | 0.542162 | 1 |
| Taf1 | 0.049654 | 0.032484 | 1 |
| Mthfr | 0.049631 | 0.0344 | 1 |
| Tmem165 | 0.049613 | 0.013927 | 1 |
| Cables1 | 0.049585 | 0.000909 | 1 |
| Rnd1 | 0.049524 | 0.000906 | 1 |
| Rfx5 | 0.049513 | 0.000604 | 1 |
| Tmem250-| | 0.049434 | 0.0292 | 1 |
| Llph | 0.049397 | 0.063945 | 1 |
| Bcl2a1c | 0.049373 | 0.001162 | 1 |
| Usp28 | 0.049324 | 0.000173 | 1 |
| Gcdh | 0.049295 | 0.001264 | 1 |
| Pi4k2b | 0.049257 | 0.004454 | 1 |
| Mapk1ip1 | 0.049205 | 0.01474 | 1 |

| | | | |
|---|---|---|---|
| 2300009A( | 0.049155 | 0.023034 | 1 |
| Capn10 | 0.049136 | 0.003768 | 1 |
| Mtfmt | 0.049108 | 5.56E-07 | 0.017953 |
| Anapc13 | 0.049081 | 0.177046 | 1 |
| Fahd2a | 0.049048 | 0.027724 | 1 |
| Ganab | 0.04901 | 0.025956 | 1 |
| Pstpip2 | 0.048923 | 0.090527 | 1 |
| Adora2b | 0.048916 | 0.201249 | 1 |
| Mprip | 0.048906 | 0.06893 | 1 |
| Gpr137 | 0.048838 | 0.005175 | 1 |
| Abcc3 | 0.048821 | 0.002676 | 1 |
| Arrb2 | 0.048771 | 0.10289 | 1 |
| Zfp358 | 0.048706 | 0.003603 | 1 |
| Adk | 0.048491 | 0.01471 | 1 |
| Fam173a | 0.048443 | 0.012343 | 1 |
| Phc2 | 0.048438 | 0.000225 | 1 |
| Ddx56 | 0.048425 | 0.00055 | 1 |
| Btg1 | 0.048411 | 0.589025 | 1 |
| P3h1 | 0.048408 | 0.004028 | 1 |
| Prkrip1 | 0.048394 | 0.019806 | 1 |
| Glo1 | 0.048377 | 0.00083 | 1 |
| Hspa14 | 0.04834 | 0.027896 | 1 |
| Zbtb38 | 0.048316 | 0.003105 | 1 |
| Atp6v1f | 0.048302 | 0.375545 | 1 |
| Tars2 | 0.048227 | 0.000222 | 1 |
| Zfp319 | 0.048221 | 0.000805 | 1 |
| Med4 | 0.048216 | 0.011268 | 1 |
| Ankzf1 | 0.048213 | 0.284734 | 1 |
| Kti12 | 0.048142 | 0.042615 | 1 |
| Cenpx | 0.048134 | 0.097494 | 1 |
| Nit2 | 0.048109 | 0.003625 | 1 |
| Shld2 | 0.048098 | 0.040058 | 1 |
| Tmem42 | 0.048076 | 0.005482 | 1 |
| Snrnp27 | 0.047978 | 0.032406 | 1 |
| H6pd | 0.047917 | 0.008439 | 1 |
| Sap18b | 0.047878 | 0.01674 | 1 |
| Gadd45a | 0.04781 | 0.008546 | 1 |
| Pdcl3 | 0.047744 | 0.013413 | 1 |
| Ipo13 | 0.047735 | 0.031099 | 1 |
| Rpap3 | 0.047682 | 0.001086 | 1 |
| Dhrs7 | 0.047664 | 0.592143 | 1 |
| Mcee | 0.047657 | 0.004759 | 1 |
| Pcna | 0.047609 | 0.205676 | 1 |
| Kank3 | 0.0476 | 0.001324 | 1 |
| Ube2a | 0.047576 | 0.32428 | 1 |
| Slc11a2 | 0.047531 | 0.004476 | 1 |
| Vps33a | 0.047486 | 0.010547 | 1 |
| Dtnbp1 | 0.047453 | 0.010748 | 1 |

| | | | |
|---|---|---|---|
| Osgin1 | 0.047447 | 0.006021 | 1 |
| Ptpn4 | 0.047435 | 0.031799 | 1 |
| Mycbp | 0.047428 | 0.019568 | 1 |
| Ilvbl | 0.047416 | 0.000804 | 1 |
| Mrpl41 | 0.047399 | 0.001909 | 1 |
| Decr2 | 0.047286 | 0.012857 | 1 |
| Mrpl36 | 0.047253 | 0.039133 | 1 |
| Tsc1 | 0.047078 | 0.021044 | 1 |
| Fntb | 0.046943 | 0.004449 | 1 |
| Zfp472 | 0.046866 | 0.091288 | 1 |
| Sco1 | 0.046836 | 0.004826 | 1 |
| Adck2 | 0.046723 | 0.196825 | 1 |
| Atp8b2 | 0.046715 | 0.152625 | 1 |
| Tnfsf14 | 0.04662 | 0.003369 | 1 |
| Sac3d1 | 0.046507 | 0.020594 | 1 |
| Ube2i | 0.046485 | 0.316591 | 1 |
| Gid4 | 0.04645 | 0.002868 | 1 |
| Gtf2f2 | 0.046444 | 0.008 | 1 |
| Ppfia1 | 0.046402 | 0.012084 | 1 |
| Pex19 | 0.046347 | 0.077773 | 1 |
| Heatr5a | 0.046311 | 0.08835 | 1 |
| Cct7 | 0.046311 | 0.208099 | 1 |
| Pla2g15 | 0.046268 | 0.001625 | 1 |
| Ndc1 | 0.046227 | 0.006607 | 1 |
| Commd3 | 0.046215 | 0.052956 | 1 |
| Champ1 | 0.046144 | 0.031835 | 1 |
| Tmem9 | 0.046132 | 0.00752 | 1 |
| Chtop | 0.046114 | 0.001881 | 1 |
| Fam76a | 0.046106 | 2.6E-05 | 0.838891 |
| Trim35 | 0.046043 | 0.005422 | 1 |
| Wiz | 0.046028 | 0.000285 | 1 |
| Fastk | 0.046025 | 0.000353 | 1 |
| Sypl | 0.046016 | 0.267421 | 1 |
| Nasp | 0.045992 | 0.050679 | 1 |
| Khdrbs1 | 0.045981 | 0.54887 | 1 |
| Mtg1 | 0.045966 | 0.030905 | 1 |
| Trio | 0.045918 | 0.043472 | 1 |
| Max | 0.045888 | 0.021628 | 1 |
| Lpgat1 | 0.045835 | 0.000402 | 1 |
| Rpl18 | 0.045757 | 0.207232 | 1 |
| Ube4b | 0.045745 | 0.031611 | 1 |
| Map3k4 | 0.045732 | 0.001522 | 1 |
| Wrnip1 | 0.045697 | 0.000657 | 1 |
| Cdk6 | 0.045653 | 0.074668 | 1 |
| Rpa3 | 0.045618 | 0.007575 | 1 |
| Proser1 | 0.045612 | 0.002854 | 1 |
| Gtf3c5 | 0.045575 | 0.006241 | 1 |
| Wdr92 | 0.045571 | 0.033788 | 1 |

| | | | |
|---|---|---|---|
| Irf7 | 0.045513 | 0.291459 | 1 |
| Gm47882 | 0.045498 | 0.007868 | 1 |
| Tceanc2 | 0.045481 | 0.01341 | 1 |
| Rps3a1 | 0.045452 | 0.469966 | 1 |
| Hps6 | 0.045388 | 0.001731 | 1 |
| Tent2 | 0.045368 | 0.386474 | 1 |
| Ccdc127 | 0.045238 | 0.026928 | 1 |
| Scap | 0.045163 | 0.007692 | 1 |
| Zfyve21 | 0.045127 | 0.000956 | 1 |
| Rbm43 | 0.045097 | 0.007624 | 1 |
| Vmp1 | 0.045088 | 0.301141 | 1 |
| Cd99l2 | 0.044941 | 0.158044 | 1 |
| Fbxl14 | 0.04493 | 0.054503 | 1 |
| Xpot | 0.044918 | 0.003207 | 1 |
| Zdhhc5 | 0.044868 | 0.014045 | 1 |
| Khsrp | 0.044866 | 0.331448 | 1 |
| Rad54l2 | 0.044793 | 0.007006 | 1 |
| Galc | 0.044771 | 0.006821 | 1 |
| Tnfrsf12a | 0.044689 | 0.000741 | 1 |
| 0610010FC | 0.044656 | 0.002113 | 1 |
| Nup160 | 0.044633 | 0.082532 | 1 |
| Nup37 | 0.044535 | 0.001087 | 1 |
| Nat10 | 0.044502 | 0.001856 | 1 |
| Tvp23b | 0.044444 | 0.052939 | 1 |
| Nqo2 | 0.044346 | 0.001401 | 1 |
| Uqcc1 | 0.044251 | 0.000854 | 1 |
| Sptssa | 0.04417 | 0.013943 | 1 |
| Rexo4 | 0.044098 | 0.04672 | 1 |
| Rnf19b | 0.044091 | 0.031982 | 1 |
| Gypc | 0.044018 | 0.011463 | 1 |
| Plp2 | 0.043984 | 0.010366 | 1 |
| Arhgap5 | 0.043951 | 0.06104 | 1 |
| Ccs | 0.043934 | 0.001554 | 1 |
| Tmem138 | 0.043893 | 0.003819 | 1 |
| Tpp1 | 0.043886 | 0.001368 | 1 |
| Cox19 | 0.043872 | 0.004054 | 1 |
| Uros | 0.043754 | 0.017371 | 1 |
| Vps72 | 0.043629 | 0.183798 | 1 |
| Eif4ebp3 | 0.043609 | 0.10525 | 1 |
| B3gntl1 | 0.043575 | 0.005122 | 1 |
| Rpp14 | 0.043574 | 0.009872 | 1 |
| Mesd | 0.043482 | 0.00175 | 1 |
| Ahsa1 | 0.043404 | 0.025527 | 1 |
| Cabin1 | 0.043376 | 0.005699 | 1 |
| Runx1 | 0.04336 | 0.291082 | 1 |
| Rfxap | 0.043298 | 0.002337 | 1 |
| Abhd5 | 0.043258 | 0.002285 | 1 |
| Itpripl1 | 0.043227 | 0.002827 | 1 |

| | | | |
|---|---|---|---|
| Prmt1 | 0.043224 | 0.022102 | 1 |
| Mrps10 | 0.043204 | 0.047486 | 1 |
| Cav2 | 0.04319 | 0.007771 | 1 |
| Qsox1 | 0.043148 | 0.041705 | 1 |
| Agk | 0.043132 | 0.000226 | 1 |
| Acaca | 0.043109 | 0.04183 | 1 |
| Cuedc2 | 0.042966 | 0.06971 | 1 |
| Prr14l | 0.04296 | 0.091145 | 1 |
| Mrps26 | 0.042959 | 0.027693 | 1 |
| Ifrd1 | 0.04294 | 0.001137 | 1 |
| Chchd7 | 0.042915 | 0.009102 | 1 |
| Spout1 | 0.042871 | 0.001064 | 1 |
| Cdpf1 | 0.042864 | 0.000238 | 1 |
| Snhg17 | 0.042811 | 0.001423 | 1 |
| Atp10a | 0.042805 | 0.063416 | 1 |
| Rbbp4 | 0.042675 | 0.044814 | 1 |
| Zfp768 | 0.042607 | 0.012654 | 1 |
| Fam207a | 0.042597 | 0.00224 | 1 |
| Ccdc107 | 0.042557 | 0.003128 | 1 |
| Ppp2ca | 0.042529 | 0.029097 | 1 |
| 2510009E( | 0.04237 | 0.012149 | 1 |
| Tmem161b | 0.042232 | 0.326908 | 1 |
| Nfix | 0.042214 | 0.029957 | 1 |
| Alg11 | 0.042093 | 0.001253 | 1 |
| Pde4d | 0.042069 | 0.016034 | 1 |
| Ost4 | 0.041993 | 0.384622 | 1 |
| Paox | 0.041989 | 0.000888 | 1 |
| Habp4 | 0.041952 | 0.034259 | 1 |
| Traf6 | 0.041913 | 0.023809 | 1 |
| Sc5d | 0.041864 | 0.048126 | 1 |
| Mrps6 | 0.041847 | 0.00236 | 1 |
| Hrh2 | 0.041839 | 0.001332 | 1 |
| Tram2 | 0.041796 | 0.01652 | 1 |
| Sfmbt1 | 0.041744 | 0.00095 | 1 |
| Bckdk | 0.0417 | 0.003779 | 1 |
| Stxbp1 | 0.041616 | 0.000304 | 1 |
| Stx17 | 0.041596 | 0.178788 | 1 |
| Chaf1a | 0.041325 | 0.004607 | 1 |
| Rps13 | 0.041267 | 0.406759 | 1 |
| Foxred1 | 0.041264 | 0.184366 | 1 |
| Ankmy2 | 0.041232 | 0.163102 | 1 |
| Errfi1 | 0.041224 | 0.008932 | 1 |
| Trmt61b | 0.041197 | 0.028188 | 1 |
| Agap1 | 0.041156 | 0.040764 | 1 |
| Nup214 | 0.041121 | 0.038373 | 1 |
| Pycr2 | 0.0411 | 0.026146 | 1 |
| Alkbh6 | 0.041015 | 0.01189 | 1 |
| Pin4 | 0.041014 | 0.030202 | 1 |

| | | | |
|---|---|---|---|
| Tmem161a | 0.040915 | 0.024353 | 1 |
| Akr7a5 | 0.040908 | 0.217282 | 1 |
| Kif3b | 0.040877 | 0.001886 | 1 |
| Cops7a | 0.04082 | 0.016153 | 1 |
| Isoc1 | 0.040818 | 0.011277 | 1 |
| Thra | 0.040748 | 0.01625 | 1 |
| Snrpa1 | 0.040731 | 0.014168 | 1 |
| Tsr1 | 0.040644 | 0.01137 | 1 |
| Ptbp2 | 0.040592 | 0.009654 | 1 |
| Flad1 | 0.040472 | 0.000417 | 1 |
| Slc39a7 | 0.040451 | 0.023085 | 1 |
| Tpmt | 0.04043 | 0.044125 | 1 |
| Tlr11 | 0.040374 | 0.008792 | 1 |
| Letmd1 | 0.040373 | 0.046877 | 1 |
| Mboat1 | 0.040367 | 0.015103 | 1 |
| Ccdc22 | 0.040252 | 0.061505 | 1 |
| Mogs | 0.040222 | 0.043884 | 1 |
| Ppp1r21 | 0.040132 | 0.248217 | 1 |
| Clec10a | 0.040044 | 0.000123 | 1 |
| Eml4 | 0.039996 | 0.282711 | 1 |
| Ccnk | 0.039994 | 0.004999 | 1 |
| Ldb1 | 0.039983 | 0.009298 | 1 |
| Bak1 | 0.039967 | 0.328578 | 1 |
| Nudt16l1 | 0.039936 | 0.00051 | 1 |
| Pdhx | 0.039874 | 0.000953 | 1 |
| Zc3h6 | 0.039746 | 0.075816 | 1 |
| AI837181 | 0.039694 | 0.008963 | 1 |
| Bub3 | 0.039674 | 0.034609 | 1 |
| Srek1 | 0.039672 | 0.003915 | 1 |
| Pskh1 | 0.039631 | 0.0442 | 1 |
| Nol8 | 0.039546 | 0.013304 | 1 |
| Utp18 | 0.039494 | 0.012953 | 1 |
| 3110040N | 0.039478 | 0.005941 | 1 |
| Zfp36l1 | 0.039383 | 0.228048 | 1 |
| Prr12 | 0.03926 | 0.013698 | 1 |
| Dhx32 | 0.039128 | 0.012997 | 1 |
| Faap100 | 0.039094 | 0.021421 | 1 |
| Samm50 | 0.039047 | 0.270271 | 1 |
| Cluap1 | 0.038863 | 0.002764 | 1 |
| Tmem237 | 0.038826 | 9.58E-05 | 1 |
| Mettl6 | 0.038825 | 0.000802 | 1 |
| Babam1 | 0.038801 | 0.002905 | 1 |
| Tent5a | 0.038756 | 0.031806 | 1 |
| Ofd1 | 0.038665 | 0.000993 | 1 |
| Sqstm1 | 0.038637 | 0.051032 | 1 |
| Sfxn3 | 0.038635 | 0.117953 | 1 |
| Ube2q2 | 0.038621 | 0.044044 | 1 |
| 2310057M | 0.038562 | 0.029589 | 1 |

| | | | |
|---|---|---|---|
| Snf8 | 0.03852 | 0.123401 | 1 |
| Gabpa | 0.038487 | 0.024005 | 1 |
| Prpf19 | 0.038485 | 0.078065 | 1 |
| Prmt5 | 0.038428 | 0.005616 | 1 |
| Cbr1 | 0.038424 | 0.049066 | 1 |
| Bpnt1 | 0.038396 | 0.011895 | 1 |
| Hook2 | 0.038388 | 0.035401 | 1 |
| Unc119b | 0.038354 | 0.003565 | 1 |
| Ercc3 | 0.038348 | 0.021713 | 1 |
| Wdr74 | 0.038319 | 0.003939 | 1 |
| Tmem80 | 0.038275 | 0.007324 | 1 |
| Zfp800 | 0.03803 | 0.330566 | 1 |
| Acp6 | 0.037904 | 0.150513 | 1 |
| Tpm3 | 0.037833 | 0.176042 | 1 |
| Sertad1 | 0.037785 | 0.003623 | 1 |
| Gstk1 | 0.037743 | 0.00114 | 1 |
| D530033B | 0.037742 | 0.027702 | 1 |
| Dennd3 | 0.037733 | 0.003198 | 1 |
| Tmco6 | 0.03769 | 0.012879 | 1 |
| Tut4 | 0.037676 | 0.557991 | 1 |
| Haus3 | 0.037632 | 0.029212 | 1 |
| Scarb2 | 0.037606 | 0.048016 | 1 |
| Sdhaf2 | 0.037602 | 0.016047 | 1 |
| Dnaja2 | 0.037573 | 0.062633 | 1 |
| Fbxw9 | 0.03752 | 0.021192 | 1 |
| Gm10131 | 0.037495 | 0.011144 | 1 |
| Uap1 | 0.037491 | 0.00406 | 1 |
| Il3ra | 0.037404 | 0.032849 | 1 |
| Mfsd11 | 0.0374 | 0.019684 | 1 |
| 2010320M | 0.037306 | 0.000407 | 1 |
| Acadvl | 0.037229 | 0.000808 | 1 |
| Nek1 | 0.037195 | 0.014525 | 1 |
| Hsd17b4 | 0.037171 | 0.630196 | 1 |
| Map1lc3a | 0.037149 | 0.017197 | 1 |
| Ehbp1 | 0.037143 | 0.010099 | 1 |
| Ift172 | 0.037139 | 0.00121 | 1 |
| Snap47 | 0.037127 | 0.015236 | 1 |
| Slc30a1 | 0.037117 | 0.013893 | 1 |
| Actr1a | 0.037072 | 0.228946 | 1 |
| BC002059 | 0.037063 | 0.011735 | 1 |
| Brf1 | 0.03702 | 0.046258 | 1 |
| Abitram | 0.037016 | 0.007029 | 1 |
| Tada2a | 0.036945 | 0.095206 | 1 |
| Spns1 | 0.036898 | 0.009125 | 1 |
| Cbx1 | 0.036896 | 0.005063 | 1 |
| Polr3e | 0.036841 | 0.005031 | 1 |
| Marc2 | 0.036838 | 0.031113 | 1 |
| Fasn | 0.036738 | 0.023206 | 1 |

| | | | |
|---|---|---|---|
| Tpra1 | 0.036549 | 0.01502 | 1 |
| Lipo3 | 0.036478 | 0.003904 | 1 |
| Smg8 | 0.036469 | 0.02591 | 1 |
| Eefsec | 0.036392 | 0.17132 | 1 |
| Nupl2 | 0.03637 | 0.102526 | 1 |
| Cib2 | 0.036357 | 0.21135 | 1 |
| Tlr2 | 0.036344 | 0.006042 | 1 |
| Dmac1 | 0.036303 | 0.026415 | 1 |
| Zfp456 | 0.036233 | 0.001625 | 1 |
| Dbr1 | 0.036227 | 0.003469 | 1 |
| Rft1 | 0.036214 | 0.00799 | 1 |
| Dpp7 | 0.0362 | 0.000453 | 1 |
| Zc3h7b | 0.036165 | 0.039482 | 1 |
| Tgm1 | 0.03616 | 0.00021 | 1 |
| Txnip | 0.036125 | 0.673102 | 1 |
| Usb1 | 0.036046 | 0.002942 | 1 |
| Ppp1r11 | 0.035992 | 0.134026 | 1 |
| Pnpt1 | 0.035987 | 0.063991 | 1 |
| Nudt18 | 0.035982 | 0.005335 | 1 |
| Tubgcp5 | 0.035972 | 0.005857 | 1 |
| Bccip | 0.035954 | 0.004964 | 1 |
| Tmem39a | 0.035904 | 0.016637 | 1 |
| Thoc6 | 0.035901 | 0.007462 | 1 |
| Necap2 | 0.035816 | 0.226073 | 1 |
| Zfp444 | 0.035807 | 0.023838 | 1 |
| Ascc2 | 0.035799 | 0.022727 | 1 |
| Yif1a | 0.035736 | 4.33E-06 | 0.13995 |
| Apoo | 0.035726 | 0.04688 | 1 |
| Gatd1 | 0.035725 | 0.004427 | 1 |
| Tom1l2 | 0.035677 | 0.017143 | 1 |
| Top3b | 0.035653 | 0.183613 | 1 |
| Praf2 | 0.035649 | 3.35E-05 | 1 |
| Rbm15b | 0.03563 | 0.008815 | 1 |
| Atp6ap1 | 0.035623 | 0.4563 | 1 |
| Dbf4 | 0.035529 | 0.004108 | 1 |
| Yipf4 | 0.0355 | 0.189153 | 1 |
| Rabif | 0.035488 | 0.680769 | 1 |
| Zxdc | 0.035451 | 0.011474 | 1 |
| Alg14 | 0.035435 | 0.001615 | 1 |
| Pef1 | 0.035433 | 0.002064 | 1 |
| Pcgf5 | 0.035401 | 0.017382 | 1 |
| Asf1a | 0.035368 | 0.060572 | 1 |
| Pdcl | 0.035366 | 0.57133 | 1 |
| Mmp25 | 0.035336 | 0.000639 | 1 |
| Sub1 | 0.035295 | 0.148563 | 1 |
| Slfn3 | 0.03528 | 0.01976 | 1 |
| Mrpl32 | 0.035275 | 0.01919 | 1 |
| Glrx5 | 0.035246 | 0.066872 | 1 |

| | | | |
|---|---|---|---|
| Rpl30 | 0.035243 | 0.482217 | 1 |
| Clcc1 | 0.035239 | 0.044297 | 1 |
| Taf4 | 0.035172 | 0.069605 | 1 |
| Maip1 | 0.035166 | 0.010534 | 1 |
| Fgd2 | 0.03509 | 0.400688 | 1 |
| Dnajb4 | 0.035086 | 0.056986 | 1 |
| Stt3a | 0.035083 | 0.021379 | 1 |
| Mcm6 | 0.035066 | 0.013623 | 1 |
| Psme2b | 0.03506 | 0.015134 | 1 |
| Tyk2 | 0.035027 | 0.306882 | 1 |
| Pdhb | 0.035012 | 0.000439 | 1 |
| Cstf1 | 0.034881 | 0.0013 | 1 |
| Uckl1 | 0.03486 | 0.004781 | 1 |
| Eftud2 | 0.03484 | 0.115229 | 1 |
| Nars2 | 0.034811 | 0.030091 | 1 |
| Adam15 | 0.034676 | 0.052413 | 1 |
| Pwwp2b | 0.034613 | 0.002515 | 1 |
| Ddx46 | 0.034608 | 0.0431 | 1 |
| Usp46 | 0.03451 | 0.00414 | 1 |
| Inpp5b | 0.034505 | 0.001413 | 1 |
| Pkp4 | 0.034336 | 0.003653 | 1 |
| Caml | 0.034323 | 0.000146 | 1 |
| Arsk | 0.034322 | 0.004708 | 1 |
| Cmc2 | 0.034314 | 0.003639 | 1 |
| Ogfr | 0.034309 | 0.262902 | 1 |
| Ypel5 | 0.0343 | 0.026751 | 1 |
| Vps26a | 0.034279 | 0.082101 | 1 |
| Zbtb25 | 0.034221 | 0.021246 | 1 |
| Zfp318 | 0.034195 | 0.006916 | 1 |
| Rgs10 | 0.034163 | 0.008857 | 1 |
| Rnf157 | 0.034154 | 0.003502 | 1 |
| Dus1l | 0.034141 | 0.004728 | 1 |
| Hnrnpa2b1 | 0.034127 | 0.368186 | 1 |
| Pde2a | 0.03412 | 0.153803 | 1 |
| Arl2bp | 0.034078 | 0.020636 | 1 |
| Gm14325 | 0.034021 | 0.074572 | 1 |
| Ndufa3 | 0.033997 | 0.16662 | 1 |
| Tmem185b | 0.033985 | 0.004469 | 1 |
| Atp6v0b | 0.033934 | 0.202415 | 1 |
| Ttyh3 | 0.033874 | 0.075495 | 1 |
| Csf1 | 0.033873 | 0.021339 | 1 |
| Slc25a19 | 0.033842 | 0.028096 | 1 |
| Telo2 | 0.033807 | 0.003305 | 1 |
| Ap2a1 | 0.033806 | 0.012768 | 1 |
| Cdk5rap3 | 0.033776 | 0.0006 | 1 |
| Hmox2 | 0.033733 | 0.137462 | 1 |
| Czib | 0.033715 | 0.001883 | 1 |
| Abt1 | 0.033656 | 0.006397 | 1 |

| | | | |
|---|---|---|---|
| Srxn1 | 0.033643 | 0.005363 | 1 |
| Ucp2 | 0.033639 | 0.734661 | 1 |
| Galnt3 | 0.03361 | 0.060949 | 1 |
| Upf1 | 0.033564 | 0.000636 | 1 |
| Ginm1 | 0.033455 | 0.005038 | 1 |
| Uck1 | 0.033384 | 0.007459 | 1 |
| Rwdd4a | 0.033374 | 0.003649 | 1 |
| Scrn3 | 0.033342 | 0.476356 | 1 |
| Gtf3c3 | 0.033322 | 0.03456 | 1 |
| Brix1 | 0.033316 | 0.005922 | 1 |
| Srebf1 | 0.033285 | 0.022226 | 1 |
| Ciz1 | 0.033241 | 0.056671 | 1 |
| Ndufaf2 | 0.033227 | 0.004622 | 1 |
| Abcd4 | 0.033208 | 0.045761 | 1 |
| Gm11131 | 0.033112 | 0.053939 | 1 |
| Csnk2a2 | 0.033107 | 0.002281 | 1 |
| Lpp | 0.033104 | 0.201419 | 1 |
| Srbd1 | 0.033099 | 0.156291 | 1 |
| Rnf10 | 0.032968 | 0.046804 | 1 |
| Taf1c | 0.032918 | 0.010361 | 1 |
| Atp11b | 0.032838 | 0.032495 | 1 |
| Styx | 0.032817 | 0.025714 | 1 |
| Cfl2 | 0.032802 | 0.034238 | 1 |
| Use1 | 0.032775 | 0.011597 | 1 |
| Eif4b | 0.032752 | 0.064641 | 1 |
| Cln6 | 0.032722 | 0.004763 | 1 |
| Etfrf1 | 0.032707 | 0.015687 | 1 |
| Bnip2 | 0.032687 | 0.197387 | 1 |
| Nsfl1c | 0.032601 | 0.003971 | 1 |
| Fermt3 | 0.032598 | 0.301705 | 1 |
| Nup155 | 0.032582 | 0.021285 | 1 |
| Tasor2 | 0.03257 | 0.133996 | 1 |
| 6330418K( | 0.03255 | 0.027903 | 1 |
| 6530409C: | 0.032511 | 0.019136 | 1 |
| Kif19a | 0.032481 | 0.000911 | 1 |
| Wnk1 | 0.032425 | 0.606432 | 1 |
| Atg4b | 0.032393 | 0.051661 | 1 |
| Fars2 | 0.032393 | 0.005086 | 1 |
| Fam53a | 0.032378 | 0.028506 | 1 |
| Cep78 | 0.032376 | 0.100855 | 1 |
| Rnf185 | 0.032368 | 0.004175 | 1 |
| Ccdc25 | 0.032316 | 0.006931 | 1 |
| Efcab14 | 0.032311 | 0.015793 | 1 |
| Golim4 | 0.032263 | 0.016211 | 1 |
| Dffb | 0.032258 | 0.105778 | 1 |
| Rps6ka4 | 0.032182 | 0.000717 | 1 |
| Sirt6 | 0.032122 | 0.049657 | 1 |
| Supv3l1 | 0.032059 | 0.023439 | 1 |

| | | | |
|---|---|---|---|
| Abcb8 | 0.032009 | 0.0022 | 1 |
| Aasdhppt | 0.031978 | 0.021551 | 1 |
| Sf3b6 | 0.031869 | 0.205481 | 1 |
| Mpi | 0.031818 | 0.015417 | 1 |
| Ascc1 | 0.031808 | 0.872755 | 1 |
| Ckap5 | 0.031805 | 0.005577 | 1 |
| Ciao3 | 0.031775 | 0.000217 | 1 |
| Elp4 | 0.031757 | 0.001755 | 1 |
| Med21 | 0.031751 | 0.023226 | 1 |
| Oprm1 | 0.031749 | 0.016976 | 1 |
| Zc3hc1 | 0.031745 | 0.096538 | 1 |
| Arhgef7 | 0.031711 | 0.002922 | 1 |
| Lin54 | 0.031664 | 0.003355 | 1 |
| Zkscan6 | 0.031629 | 0.010989 | 1 |
| Fcf1 | 0.031614 | 0.069116 | 1 |
| Ncoa6 | 0.031597 | 0.002383 | 1 |
| Pacs2 | 0.031593 | 0.000737 | 1 |
| Rffl | 0.031584 | 0.011434 | 1 |
| Slc36a4 | 0.03156 | 0.197944 | 1 |
| Ddah1 | 0.031523 | 0.004964 | 1 |
| Pgm2 | 0.03145 | 0.006223 | 1 |
| Ash2l | 0.031254 | 0.01702 | 1 |
| Ddx10 | 0.031214 | 0.009697 | 1 |
| Rbm8a | 0.031193 | 0.308279 | 1 |
| Nprl3 | 0.031161 | 0.040345 | 1 |
| Adprhl2 | 0.031155 | 0.075611 | 1 |
| Ciao2b | 0.031008 | 0.039766 | 1 |
| 2610020C( | 0.030996 | 0.030332 | 1 |
| Zfand1 | 0.030976 | 0.042151 | 1 |
| Spata5l1 | 0.030933 | 0.039353 | 1 |
| Vps9d1 | 0.030845 | 0.098296 | 1 |
| Tmem141 | 0.030822 | 0.012198 | 1 |
| Saal1 | 0.030811 | 0.041015 | 1 |
| Rdh14 | 0.030775 | 0.000648 | 1 |
| Rbfa | 0.030773 | 0.015374 | 1 |
| Dhdh | 0.030772 | 0.009809 | 1 |
| Mgat4b | 0.030761 | 0.00044 | 1 |
| Egln1 | 0.030729 | 0.009313 | 1 |
| Ivd | 0.030669 | 0.001395 | 1 |
| Mtmr9 | 0.030572 | 0.016438 | 1 |
| Setd3 | 0.030559 | 0.020584 | 1 |
| Srd5a3 | 0.030509 | 0.011466 | 1 |
| Ift46 | 0.030489 | 0.033528 | 1 |
| Slc25a16 | 0.030462 | 0.001223 | 1 |
| Mms22l | 0.030461 | 0.040961 | 1 |
| Psmd14 | 0.030379 | 0.006237 | 1 |
| Cisd3 | 0.030363 | 0.00153 | 1 |
| Tceal9 | 0.030355 | 0.104409 | 1 |

| | | | |
|---|---|---|---|
| Fbxl15 | 0.030293 | 0.005533 | 1 |
| Traf2 | 0.030234 | 0.014431 | 1 |
| Lrch1 | 0.030191 | 0.091509 | 1 |
| Hpcal1 | 0.03016 | 0.246881 | 1 |
| Dnajc21 | 0.030052 | 0.006466 | 1 |
| Tent4a | 0.030013 | 0.015151 | 1 |
| Dusp28 | 0.030006 | 0.008755 | 1 |
| Myl12a | 0.030005 | 0.125597 | 1 |
| Flot1 | 0.029903 | 0.093033 | 1 |
| Fastkd1 | 0.029897 | 0.040406 | 1 |
| Snapc4 | 0.029829 | 0.046696 | 1 |
| E430024P1 | 0.029727 | 0.0715 | 1 |
| Fbxw7 | 0.029727 | 0.001715 | 1 |
| Nsmf | 0.029721 | 0.033028 | 1 |
| Map2k3 | 0.029696 | 0.037604 | 1 |
| Tmem185a | 0.029562 | 0.021104 | 1 |
| Hspa4 | 0.029512 | 0.18141 | 1 |
| Mks1 | 0.02951 | 0.017919 | 1 |
| Aamp | 0.029371 | 0.003743 | 1 |
| Prorp | 0.029357 | 0.013071 | 1 |
| Eif2ak1 | 0.029252 | 0.445742 | 1 |
| Cep152 | 0.029224 | 0.487239 | 1 |
| Polr3a | 0.029174 | 0.088209 | 1 |
| Zswim4 | 0.029131 | 0.046833 | 1 |
| Gm16229 | 0.029121 | 0.012832 | 1 |
| Yju2 | 0.029104 | 0.011055 | 1 |
| Dock7 | 0.029053 | 0.002248 | 1 |
| Arhgef11 | 0.028999 | 0.008123 | 1 |
| Rnf5 | 0.028997 | 0.016332 | 1 |
| Exo5 | 0.028968 | 0.003027 | 1 |
| Cep70 | 0.028962 | 0.10057 | 1 |
| Rab11fip1 | 0.028961 | 0.036867 | 1 |
| Pcid2 | 0.028943 | 0.122654 | 1 |
| Tysnd1 | 0.028917 | 0.000323 | 1 |
| Narf | 0.02889 | 0.038898 | 1 |
| Sirt2 | 0.028874 | 0.004127 | 1 |
| Abhd17c | 0.02883 | 0.018864 | 1 |
| Tmem184c | 0.028771 | 0.293849 | 1 |
| Hadha | 0.028747 | 0.120864 | 1 |
| Smim24 | 0.028666 | 0.015499 | 1 |
| Dgcr6 | 0.02862 | 0.03215 | 1 |
| 2310061I0 | 0.028583 | 0.029199 | 1 |
| Jmjd1c | 0.028582 | 0.107856 | 1 |
| Nap1l4 | 0.028527 | 0.187702 | 1 |
| Dars | 0.028518 | 0.170914 | 1 |
| D10Wsu10 | 0.028517 | 0.111815 | 1 |
| Ocrl | 0.028499 | 0.126469 | 1 |
| Cstf2 | 0.028496 | 0.044245 | 1 |

| | | | |
|---|---|---|---|
| Etfbkmt | 0.028444 | 0.130317 | 1 |
| Tgds | 0.028414 | 0.004822 | 1 |
| Thap12 | 0.028402 | 0.007077 | 1 |
| Gm9725 | 0.028383 | 0.055295 | 1 |
| 9530082P2 | 0.028279 | 0.000393 | 1 |
| Gna12 | 0.028264 | 0.005566 | 1 |
| Gm15821 | 0.028253 | 0.046733 | 1 |
| Szt2 | 0.02824 | 0.018062 | 1 |
| Tmbim6 | 0.028226 | 0.171675 | 1 |
| Amz2 | 0.028184 | 0.016529 | 1 |
| Zbtb11os1 | 0.028114 | 0.021758 | 1 |
| Mfge8 | 0.028035 | 0.051432 | 1 |
| Tmed8 | 0.028034 | 0.015369 | 1 |
| Fam149b | 0.028029 | 0.003355 | 1 |
| Gmppa | 0.028023 | 0.093977 | 1 |
| Pnpla6 | 0.028003 | 0.057317 | 1 |
| 2700062C( | 0.027983 | 0.029937 | 1 |
| Nsdhl | 0.027958 | 0.001393 | 1 |
| Dynlt3 | 0.027952 | 0.0432 | 1 |
| Rcbtb1 | 0.027921 | 0.004203 | 1 |
| Pex3 | 0.027872 | 0.472539 | 1 |
| Ykt6 | 0.027854 | 0.040302 | 1 |
| 1600002K( | 0.027827 | 0.001521 | 1 |
| Mpdu1 | 0.027724 | 0.040297 | 1 |
| Rab1b | 0.027722 | 0.162407 | 1 |
| Ap2a2 | 0.027709 | 0.109413 | 1 |
| Atrip | 0.027637 | 0.02043 | 1 |
| Dtx3l | 0.027535 | 0.029955 | 1 |
| Gbe1 | 0.027432 | 0.013597 | 1 |
| Tmem219 | 0.027428 | 0.03967 | 1 |
| Cog7 | 0.027398 | 0.357457 | 1 |
| Ube2e1 | 0.027394 | 0.021004 | 1 |
| Zfp157 | 0.02738 | 0.129905 | 1 |
| Itpa | 0.027376 | 0.060712 | 1 |
| Zmpste24 | 0.027375 | 0.029638 | 1 |
| Rps6ka5 | 0.027357 | 0.012249 | 1 |
| Nepro | 0.027356 | 0.026277 | 1 |
| Tlr6 | 0.027306 | 0.016902 | 1 |
| Gm4316 | 0.027285 | 0.069436 | 1 |
| Osbpl11 | 0.027265 | 0.072134 | 1 |
| Reep4 | 0.027221 | 0.003457 | 1 |
| Ccdc167 | 0.027214 | 0.024027 | 1 |
| Zscan21 | 0.027142 | 0.035916 | 1 |
| Rab5a | 0.02711 | 0.034713 | 1 |
| Nacc2 | 0.027084 | 0.018026 | 1 |
| Abcb10 | 0.027078 | 0.028729 | 1 |
| Mief1 | 0.026915 | 0.174505 | 1 |
| Casz1 | 0.026904 | 0.055927 | 1 |

| | | | |
|---|---|---|---|
| Atr | 0.026864 | 0.00388 | 1 |
| Csnk1g3 | 0.026863 | 0.047968 | 1 |
| Vamp7 | 0.026813 | 0.001223 | 1 |
| Slc22a4 | 0.026809 | 0.01722 | 1 |
| Zfp654 | 0.026702 | 0.042414 | 1 |
| Eogt | 0.026667 | 0.155853 | 1 |
| Gstz1 | 0.026662 | 0.001293 | 1 |
| Gnptg | 0.026588 | 0.04453 | 1 |
| Gnl3l | 0.026565 | 0.051052 | 1 |
| Prdm1 | 0.026558 | 0.030457 | 1 |
| Med20 | 0.02653 | 0.148258 | 1 |
| Zbtb17 | 0.026514 | 0.023953 | 1 |
| Kyat3 | 0.026508 | 0.073541 | 1 |
| Coq10a | 0.026383 | 0.020484 | 1 |
| Katna1 | 0.026335 | 0.211444 | 1 |
| Sf3b5 | 0.026259 | 0.110324 | 1 |
| 2610002M | 0.026215 | 0.000374 | 1 |
| Dnaja3 | 0.02621 | 0.002048 | 1 |
| Scfd1 | 0.026199 | 0.064207 | 1 |
| Rhbdd3 | 0.026153 | 0.054249 | 1 |
| Nt5dc3 | 0.026149 | 0.015659 | 1 |
| Dhx34 | 0.026108 | 0.113095 | 1 |
| Tmx2 | 0.026095 | 0.004271 | 1 |
| Trub2 | 0.026048 | 0.050702 | 1 |
| Lrrc47 | 0.026023 | 0.004052 | 1 |
| 1110065P2 | 0.025938 | 0.006818 | 1 |
| Gemin7 | 0.025861 | 0.128208 | 1 |
| Pank4 | 0.025837 | 0.015015 | 1 |
| Ndufb3 | 0.025707 | 0.173799 | 1 |
| Ulbp1 | 0.025694 | 0.00305 | 1 |
| Btrc | 0.025647 | 0.153243 | 1 |
| Ubqln1 | 0.025617 | 0.02541 | 1 |
| Vac14 | 0.025613 | 0.011458 | 1 |
| Sh2b1 | 0.02558 | 0.069305 | 1 |
| Polr1c | 0.025561 | 0.001135 | 1 |
| Xpnpep1 | 0.025531 | 0.068693 | 1 |
| Ikbke | 0.025405 | 3.18E-05 | 1 |
| Dpagt1 | 0.025384 | 0.001973 | 1 |
| Pign | 0.025369 | 0.052281 | 1 |
| Gdap2 | 0.025341 | 0.343346 | 1 |
| Serhl | 0.025221 | 0.029951 | 1 |
| Usp24 | 0.025182 | 0.030227 | 1 |
| Zfp993 | 0.025169 | 0.009373 | 1 |
| Cpne8 | 0.025088 | 0.008677 | 1 |
| Nemp1 | 0.025074 | 0.153986 | 1 |
| Bcl2l12 | 0.025045 | 0.339391 | 1 |
| Neil1 | 0.025041 | 0.021139 | 1 |
| Ccdc6 | 0.02504 | 0.004945 | 1 |

| | | | |
|---|---|---|---|
| Tsr3 | 0.024974 | 0.004614 | 1 |
| Bcl9l | 0.02496 | 0.10805 | 1 |
| Tpgs1 | 0.024948 | 0.105258 | 1 |
| Zfp934 | 0.024907 | 0.102017 | 1 |
| Stk35 | 0.024862 | 0.837285 | 1 |
| Creld1 | 0.024856 | 0.042492 | 1 |
| Git1 | 0.024811 | 0.019116 | 1 |
| Ss18l2 | 0.024811 | 0.003811 | 1 |
| Trmt10a | 0.024796 | 0.082352 | 1 |
| Gm8739 | 0.024753 | 0.030691 | 1 |
| P4ha1 | 0.024728 | 0.246772 | 1 |
| Fbxl8 | 0.024689 | 0.377312 | 1 |
| Exoc8 | 0.024681 | 0.014076 | 1 |
| BC005561 | 0.024625 | 0.033126 | 1 |
| Panx1 | 0.024595 | 0.027198 | 1 |
| Stom | 0.024541 | 0.088495 | 1 |
| Snd1 | 0.024528 | 0.181488 | 1 |
| Naa40 | 0.024516 | 0.001981 | 1 |
| Rmnd5a | 0.024494 | 0.046095 | 1 |
| Lnpk | 0.024447 | 0.098684 | 1 |
| Ubxn2b | 0.024355 | 0.087185 | 1 |
| Thumpd2 | 0.024344 | 0.018003 | 1 |
| Epm2aip1 | 0.024212 | 0.03267 | 1 |
| Manba | 0.024112 | 0.048499 | 1 |
| Tirap | 0.024111 | 0.101311 | 1 |
| Samd8 | 0.024102 | 0.229555 | 1 |
| Fance | 0.024095 | 0.005163 | 1 |
| Desi1 | 0.024091 | 0.272717 | 1 |
| Poc5 | 0.024031 | 0.116012 | 1 |
| Tpgs2 | 0.024021 | 0.013394 | 1 |
| Xpo4 | 0.024007 | 0.010316 | 1 |
| Atn1 | 0.023953 | 0.033179 | 1 |
| Lpcat1 | 0.023953 | 0.005534 | 1 |
| Ssh3 | 0.02384 | 0.013302 | 1 |
| Cflar | 0.023837 | 0.042014 | 1 |
| Exd2 | 0.023746 | 0.018696 | 1 |
| Slc37a4 | 0.023727 | 0.152387 | 1 |
| Rprd1b | 0.023611 | 0.021021 | 1 |
| Nle1 | 0.02356 | 0.021332 | 1 |
| Maea | 0.023435 | 0.101185 | 1 |
| Ahcy | 0.023402 | 0.029244 | 1 |
| Zfp24 | 0.02339 | 0.043498 | 1 |
| Nup50 | 0.023338 | 0.005586 | 1 |
| Copz1 | 0.023336 | 0.422222 | 1 |
| Trip10 | 0.023311 | 0.023848 | 1 |
| Rpl21 | 0.023305 | 0.707055 | 1 |
| Platr25 | 0.023218 | 0.205988 | 1 |
| A330023F2 | 0.023199 | 0.009129 | 1 |

| | | | |
|---|---|---|---|
| Spaca9 | 0.023191 | 0.001403 | 1 |
| Tiprl | 0.0231 | 0.046243 | 1 |
| Slc2a6 | 0.023083 | 0.128422 | 1 |
| Nras | 0.023063 | 0.263414 | 1 |
| Ubap2l | 0.022987 | 0.291203 | 1 |
| Cul4a | 0.022919 | 0.005066 | 1 |
| 1700021F( | 0.022897 | 0.025401 | 1 |
| Utp25 | 0.022896 | 0.002714 | 1 |
| Fem1b | 0.022896 | 0.038266 | 1 |
| Fbxo42 | 0.022823 | 0.276644 | 1 |
| Gpat4 | 0.022755 | 0.059656 | 1 |
| Fam114a1 | 0.022718 | 0.019492 | 1 |
| Tmem214 | 0.022654 | 0.008949 | 1 |
| Adipor2 | 0.022592 | 0.537839 | 1 |
| Ecd | 0.022583 | 0.011529 | 1 |
| Mphosph9 | 0.022576 | 0.028147 | 1 |
| Kansl2 | 0.022525 | 0.041229 | 1 |
| Eif5b | 0.022481 | 0.091955 | 1 |
| Gstt2 | 0.022474 | 0.190539 | 1 |
| Cops2 | 0.022428 | 0.111057 | 1 |
| Armh3 | 0.022409 | 0.027938 | 1 |
| Tbk1 | 0.022322 | 0.006684 | 1 |
| Golga5 | 0.022243 | 0.002151 | 1 |
| Mmut | 0.022228 | 0.135723 | 1 |
| 1190007I0 | 0.022213 | 0.001564 | 1 |
| Aven | 0.022159 | 0.049689 | 1 |
| St13 | 0.022137 | 0.015713 | 1 |
| Tnip3 | 0.022135 | 0.229504 | 1 |
| Dcaf15 | 0.022131 | 0.011195 | 1 |
| Clock | 0.022091 | 0.033767 | 1 |
| Myef2 | 0.022071 | 0.005301 | 1 |
| Gcc1 | 0.022051 | 0.122167 | 1 |
| Ap3s1 | 0.02204 | 0.060298 | 1 |
| Prpf40a | 0.022039 | 0.10752 | 1 |
| Vps35l | 0.022036 | 0.097239 | 1 |
| Pip5k1a | 0.022007 | 0.052434 | 1 |
| Ppip5k1 | 0.021973 | 0.015051 | 1 |
| Lig1 | 0.021968 | 0.094175 | 1 |
| 0610010K1 | 0.021872 | 0.00292 | 1 |
| Hist1h1c | 0.021868 | 0.483643 | 1 |
| G3bp2 | 0.021841 | 0.363364 | 1 |
| Erf | 0.021841 | 0.008292 | 1 |
| Spg20 | 0.021817 | 0.035392 | 1 |
| Snrnp35 | 0.021783 | 0.009023 | 1 |
| Prmt6 | 0.021776 | 0.074412 | 1 |
| Tbcd | 0.021722 | 0.00677 | 1 |
| Acp2 | 0.021659 | 0.01941 | 1 |
| Arl5b | 0.021641 | 0.031622 | 1 |

| | | | |
|---|---|---|---|
| Tomm70a | 0.021609 | 0.001676 | 1 |
| 2310022B( | 0.021581 | 0.03911 | 1 |
| Mtmr14 | 0.021566 | 0.177997 | 1 |
| Sh3bp5l | 0.021502 | 0.027087 | 1 |
| Tusc2 | 0.021464 | 0.007016 | 1 |
| Nacc1 | 0.021348 | 0.231934 | 1 |
| Nudt1 | 0.021268 | 0.092234 | 1 |
| Abhd14b | 0.021259 | 0.096918 | 1 |
| Psmc2 | 0.021254 | 0.003628 | 1 |
| Timm29 | 0.021252 | 0.239967 | 1 |
| Ccnl1 | 0.021235 | 0.127955 | 1 |
| Eaf1 | 0.021231 | 0.012073 | 1 |
| Smim8 | 0.021222 | 0.00107 | 1 |
| Sec22a | 0.021221 | 0.021295 | 1 |
| 4930402H: | 0.021212 | 0.042952 | 1 |
| Pmpca | 0.021164 | 0.025906 | 1 |
| Arhgap22 | 0.021099 | 0.604873 | 1 |
| Taf1a | 0.021077 | 0.003427 | 1 |
| Vdac3 | 0.020909 | 0.02865 | 1 |
| Rapgef2 | 0.020899 | 0.082842 | 1 |
| Fam219a | 0.020892 | 0.005112 | 1 |
| Eif4a3 | 0.020803 | 0.012018 | 1 |
| Trmt2b | 0.020745 | 0.000567 | 1 |
| Lmf2 | 0.020736 | 0.010162 | 1 |
| Dnttip2 | 0.020731 | 0.054629 | 1 |
| Med10 | 0.020688 | 0.004289 | 1 |
| Gm16286 | 0.020636 | 0.178602 | 1 |
| Mnat1 | 0.020597 | 0.069791 | 1 |
| Commd2 | 0.02057 | 0.159096 | 1 |
| Ercc8 | 0.020488 | 0.001048 | 1 |
| Smim4 | 0.020471 | 0.076914 | 1 |
| Sars2 | 0.020456 | 0.001046 | 1 |
| Sft2d3 | 0.020453 | 0.093702 | 1 |
| Ttf2 | 0.020402 | 0.004072 | 1 |
| Kdm6a | 0.020383 | 0.010711 | 1 |
| Prkra | 0.020377 | 0.571096 | 1 |
| Fam217b | 0.02029 | 0.059908 | 1 |
| Eif2s3x | 0.020281 | 0.001766 | 1 |
| Nipa2 | 0.020263 | 0.258425 | 1 |
| Usp19 | 0.020259 | 0.104128 | 1 |
| Mnt | 0.020195 | 0.061296 | 1 |
| Borcs7 | 0.020194 | 0.061383 | 1 |
| Twf2 | 0.02019 | 0.057643 | 1 |
| Tm9sf1 | 0.020185 | 0.003972 | 1 |
| Scyl1 | 0.020105 | 0.028461 | 1 |
| Lsm3 | 0.020082 | 0.025727 | 1 |
| Gm12216 | 0.019897 | 0.009149 | 1 |
| Cbwd1 | 0.019863 | 0.167047 | 1 |

| | | | |
|---|---|---|---|
| Cpsf2 | 0.019842 | 0.108634 | 1 |
| Gatad1 | 0.019814 | 0.03665 | 1 |
| Mrpl49 | 0.019785 | 0.01446 | 1 |
| Msto1 | 0.019652 | 0.054184 | 1 |
| Pfkfb3 | 0.019632 | 0.071036 | 1 |
| Dhcr24 | 0.019621 | 0.697295 | 1 |
| Nfs1 | 0.019549 | 0.014431 | 1 |
| Ogg1 | 0.019441 | 0.016537 | 1 |
| Stx6 | 0.01938 | 0.313998 | 1 |
| Zfat | 0.019362 | 0.000681 | 1 |
| Acsf2 | 0.019354 | 0.033213 | 1 |
| Nucb2 | 0.019321 | 0.009983 | 1 |
| Crtc2 | 0.019299 | 0.032478 | 1 |
| Ilkap | 0.019294 | 0.042967 | 1 |
| Aida | 0.019268 | 0.02411 | 1 |
| Mccc1 | 0.019247 | 0.217626 | 1 |
| Kdm4a | 0.019243 | 0.019753 | 1 |
| Trp53inp1 | 0.019102 | 0.136136 | 1 |
| Flnb | 0.019074 | 0.003601 | 1 |
| Rbmx | 0.018991 | 0.050616 | 1 |
| 2410002F2 | 0.018926 | 0.182631 | 1 |
| Hirip3 | 0.018923 | 0.002873 | 1 |
| Kctd5 | 0.01884 | 0.051934 | 1 |
| Rpl36 | 0.018837 | 0.896432 | 1 |
| Paxip1 | 0.018821 | 0.011053 | 1 |
| Fam114a2 | 0.018794 | 0.015501 | 1 |
| Dnal4 | 0.018744 | 0.003788 | 1 |
| Wdr61 | 0.018672 | 0.034258 | 1 |
| Nabp2 | 0.018669 | 0.404484 | 1 |
| Galns | 0.018668 | 0.094266 | 1 |
| Rheb | 0.018657 | 0.068781 | 1 |
| Nosip | 0.018652 | 0.011945 | 1 |
| Ube2k | 0.018585 | 0.090887 | 1 |
| Ing2 | 0.018467 | 0.954231 | 1 |
| Top3a | 0.018459 | 0.020678 | 1 |
| Washc2 | 0.01844 | 0.055179 | 1 |
| Afg3l2 | 0.018431 | 0.007868 | 1 |
| Il1rap | 0.018377 | 0.054307 | 1 |
| Zdhhc13 | 0.01832 | 0.017131 | 1 |
| Tbcel | 0.018195 | 0.004121 | 1 |
| Lacc1 | 0.018127 | 0.028598 | 1 |
| Nr2c2 | 0.01812 | 0.045007 | 1 |
| Vps13d | 0.018034 | 0.090239 | 1 |
| Plaa | 0.017921 | 0.041532 | 1 |
| Xab2 | 0.017881 | 0.744691 | 1 |
| Camsap1 | 0.017831 | 0.070779 | 1 |
| Noa1 | 0.017823 | 0.126314 | 1 |
| Msh6 | 0.017809 | 0.128308 | 1 |

| | | | |
|---|---|---|---|
| Syncrip | 0.017713 | 0.125217 | 1 |
| Pramef8 | 0.017697 | 0.059011 | 1 |
| Isg20l2 | 0.017687 | 0.012018 | 1 |
| Pola2 | 0.017654 | 0.019282 | 1 |
| Itprid2 | 0.017625 | 0.095102 | 1 |
| Tmem29 | 0.017624 | 0.028843 | 1 |
| Ago3 | 0.017578 | 0.265121 | 1 |
| Hddc2 | 0.017552 | 0.000968 | 1 |
| Cenpt | 0.017544 | 0.009252 | 1 |
| Mrps25 | 0.017537 | 0.001407 | 1 |
| Gm2629 | 0.017523 | 0.003026 | 1 |
| 4932438A1 | 0.01752 | 0.123931 | 1 |
| Sesn2 | 0.017516 | 0.008821 | 1 |
| Decr1 | 0.017435 | 0.071349 | 1 |
| Ppp2cb | 0.017406 | 0.072959 | 1 |
| Nop9 | 0.017372 | 0.011573 | 1 |
| Tcf3 | 0.017344 | 0.078771 | 1 |
| Tmf1 | 0.017329 | 0.240931 | 1 |
| Pabpc4 | 0.017318 | 0.028511 | 1 |
| Tnpo3 | 0.017311 | 0.089989 | 1 |
| Maco1 | 0.017301 | 0.157397 | 1 |
| Csnk2a1 | 0.017287 | 0.38644 | 1 |
| Cul4b | 0.017266 | 0.020185 | 1 |
| Llgl1 | 0.017262 | 0.006857 | 1 |
| Bud13 | 0.01723 | 0.021577 | 1 |
| Gpr84 | 0.017117 | 0.007531 | 1 |
| Nop16 | 0.017116 | 0.01118 | 1 |
| Pisd | 0.017104 | 0.188772 | 1 |
| Srsf2 | 0.017067 | 0.582789 | 1 |
| Klhdc10 | 0.01706 | 0.252045 | 1 |
| Fzd7 | 0.017042 | 0.184668 | 1 |
| Mrps31 | 0.01703 | 0.005367 | 1 |
| Syce2 | 0.017018 | 0.003977 | 1 |
| Nkapd1 | 0.016995 | 0.050752 | 1 |
| Cyp20a1 | 0.016985 | 0.01598 | 1 |
| Bloc1s5 | 0.016928 | 0.008755 | 1 |
| E2f1 | 0.01692 | 0.063153 | 1 |
| Mcm3ap | 0.016891 | 0.100786 | 1 |
| G2e3 | 0.016878 | 0.027637 | 1 |
| Mtfr1l | 0.01681 | 0.012629 | 1 |
| Gm9844 | 0.016786 | 0.132366 | 1 |
| Ccdc47 | 0.016736 | 0.32974 | 1 |
| Hsd17b7 | 0.016734 | 0.134745 | 1 |
| Nanp | 0.016649 | 0.005921 | 1 |
| Zc3h14 | 0.016576 | 0.012973 | 1 |
| Slc25a10 | 0.016541 | 0.040582 | 1 |
| Asnsd1 | 0.016356 | 0.021738 | 1 |
| Osbp | 0.016344 | 0.024336 | 1 |

| | | | |
|---|---|---|---|
| Phf10 | 0.016303 | 0.002119 | 1 |
| Rad18 | 0.016233 | 0.004341 | 1 |
| Denr | 0.016212 | 0.058987 | 1 |
| 4931406C( | 0.016179 | 0.014696 | 1 |
| Pip4k2c | 0.016114 | 0.114702 | 1 |
| Fkrp | 0.016079 | 0.029505 | 1 |
| Rpusd1 | 0.016072 | 0.007259 | 1 |
| Mrpl10 | 0.016069 | 0.160304 | 1 |
| Chtf8 | 0.016056 | 0.445521 | 1 |
| Cast | 0.016053 | 0.174055 | 1 |
| Cask | 0.016042 | 0.384057 | 1 |
| Plcb1 | 0.016027 | 0.187228 | 1 |
| Ap3s2 | 0.015976 | 0.036655 | 1 |
| Fmc1 | 0.015918 | 0.007512 | 1 |
| Kctd3 | 0.015893 | 0.01634 | 1 |
| Nup35 | 0.015871 | 0.003369 | 1 |
| Bahd1 | 0.015806 | 0.119486 | 1 |
| Zfp598 | 0.015794 | 0.004758 | 1 |
| Sertad3 | 0.015789 | 0.010313 | 1 |
| Ampd2 | 0.015739 | 0.040447 | 1 |
| Chd2 | 0.01572 | 0.240072 | 1 |
| Sertad2 | 0.015674 | 0.000772 | 1 |
| Arhgef6 | 0.015653 | 0.042867 | 1 |
| Tatdn2 | 0.015535 | 0.002423 | 1 |
| Jmjd7 | 0.015509 | 0.008919 | 1 |
| Med15 | 0.015503 | 0.063992 | 1 |
| Ywhae | 0.015451 | 0.255936 | 1 |
| Mcm2 | 0.015442 | 0.052714 | 1 |
| Atp6v1e1 | 0.015435 | 0.246512 | 1 |
| Surf6 | 0.015433 | 0.282166 | 1 |
| Tbc1d20 | 0.015335 | 0.029862 | 1 |
| Sec23b | 0.015281 | 0.009286 | 1 |
| Mthfd1 | 0.015261 | 0.011918 | 1 |
| Slc25a26 | 0.015219 | 0.007567 | 1 |
| Fbxo25 | 0.01519 | 0.032248 | 1 |
| Akap11 | 0.015136 | 0.032088 | 1 |
| Atg14 | 0.015116 | 0.438368 | 1 |
| Cplane1 | 0.015097 | 0.059206 | 1 |
| Tyw5 | 0.015062 | 0.018977 | 1 |
| Ndufaf6 | 0.01504 | 0.071464 | 1 |
| Coasy | 0.015034 | 0.042131 | 1 |
| Nme7 | 0.015001 | 0.04489 | 1 |
| Crem | 0.014975 | 0.090049 | 1 |
| Zdhhc16 | 0.014971 | 0.008746 | 1 |
| Ube2v2 | 0.014956 | 0.217628 | 1 |
| Asb8 | 0.01495 | 0.008184 | 1 |
| Deaf1 | 0.014942 | 0.005459 | 1 |
| Rars2 | 0.014916 | 0.123658 | 1 |

| | | | |
|---|---|---|---|
| Ints7 | 0.014827 | 0.106149 | 1 |
| Inpp5a | 0.01481 | 0.061622 | 1 |
| Mtf1 | 0.014724 | 0.082772 | 1 |
| Cbll1 | 0.014702 | 0.038502 | 1 |
| Car9 | 0.014681 | 0.11889 | 1 |
| Phactr4 | 0.014677 | 0.000842 | 1 |
| Prcc | 0.01466 | 0.075121 | 1 |
| Pdpr | 0.014644 | 0.005167 | 1 |
| Mrpl58 | 0.014643 | 0.016529 | 1 |
| Lancl2 | 0.014637 | 0.068832 | 1 |
| Creb3 | 0.014574 | 0.200889 | 1 |
| Kat5 | 0.014564 | 0.931028 | 1 |
| 1810030O( | 0.014542 | 0.330239 | 1 |
| Smyd5 | 0.014504 | 0.000241 | 1 |
| Ell | 0.01448 | 0.022984 | 1 |
| Hax1 | 0.014435 | 0.06695 | 1 |
| Mtus1 | 0.014388 | 0.045843 | 1 |
| Ddx20 | 0.01437 | 0.010905 | 1 |
| Tax1bp3 | 0.014334 | 0.006315 | 1 |
| Gale | 0.014252 | 0.025248 | 1 |
| Pdcd6ip | 0.014247 | 0.371166 | 1 |
| Bmi1 | 0.014227 | 0.008918 | 1 |
| Tex2 | 0.014213 | 0.031668 | 1 |
| Mars | 0.014128 | 0.089552 | 1 |
| Mif4gd | 0.014117 | 0.051278 | 1 |
| Mtor | 0.014089 | 0.015092 | 1 |
| Tti2 | 0.014081 | 0.229867 | 1 |
| Oxld1 | 0.014078 | 0.060703 | 1 |
| Rae1 | 0.014029 | 0.276512 | 1 |
| Uchl5 | 0.013973 | 0.010089 | 1 |
| Atad2 | 0.013971 | 0.031414 | 1 |
| Pigf | 0.013915 | 0.023398 | 1 |
| Tmem109 | 0.013879 | 0.352827 | 1 |
| Hotairm1 | 0.013876 | 0.00547 | 1 |
| Dapk3 | 0.013872 | 0.093943 | 1 |
| Otud6b | 0.013849 | 0.003194 | 1 |
| Ttc19 | 0.013838 | 0.030987 | 1 |
| Gm8797 | 0.013791 | 0.029731 | 1 |
| Kxd1 | 0.013778 | 0.120181 | 1 |
| Eef1akmt1 | 0.013696 | 0.04448 | 1 |
| Uqcrh | 0.01369 | 0.990188 | 1 |
| Elp2 | 0.013674 | 0.025647 | 1 |
| Tex30 | 0.013624 | 0.029539 | 1 |
| Vkorc1l1 | 0.01357 | 0.105899 | 1 |
| Inppl1 | 0.013533 | 0.000184 | 1 |
| Mtx2 | 0.013403 | 0.001635 | 1 |
| Slc1a5 | 0.013401 | 0.003651 | 1 |
| Ccdc43 | 0.013299 | 0.266156 | 1 |

| | | | |
|---|---|---|---|
| Pms2 | 0.013268 | 0.020042 | 1 |
| Sec63 | 0.013265 | 0.004687 | 1 |
| Virma | 0.013258 | 0.015701 | 1 |
| Rnf38 | 0.013244 | 0.291323 | 1 |
| Wdr19 | 0.01321 | 0.022488 | 1 |
| Bnip1 | 0.013199 | 0.003609 | 1 |
| Sfxn5 | 0.013162 | 0.010618 | 1 |
| Ap1ar | 0.013141 | 0.02477 | 1 |
| Pdk1 | 0.013138 | 0.029118 | 1 |
| Tcn2 | 0.013103 | 0.019013 | 1 |
| Aph1c | 0.013102 | 0.197325 | 1 |
| Heatr3 | 0.013077 | 0.036521 | 1 |
| Zfpl1 | 0.013039 | 0.00408 | 1 |
| Cxxc1 | 0.013034 | 0.277242 | 1 |
| Map2k4 | 0.012994 | 0.074846 | 1 |
| Cops8 | 0.012963 | 0.032284 | 1 |
| Ppie | 0.012933 | 0.108496 | 1 |
| Stk25 | 0.012901 | 0.038811 | 1 |
| Tars | 0.012853 | 0.080613 | 1 |
| Slc25a43 | 0.012803 | 0.036099 | 1 |
| Ctu1 | 0.012727 | 0.062985 | 1 |
| Tmem251 | 0.012719 | 0.006425 | 1 |
| Med16 | 0.012712 | 0.07493 | 1 |
| Iffo2 | 0.012703 | 0.165369 | 1 |
| Psmg3 | 0.012697 | 0.123616 | 1 |
| Hs1bp3 | 0.012656 | 0.001951 | 1 |
| Sptlc1 | 0.012567 | 0.001377 | 1 |
| Pnp2 | 0.012501 | 0.007032 | 1 |
| Ubap2 | 0.012404 | 0.147957 | 1 |
| 1110004F1 | 0.012402 | 0.449227 | 1 |
| Rbsn | 0.012364 | 0.033998 | 1 |
| Mafk | 0.012335 | 0.131965 | 1 |
| Necap1 | 0.012259 | 0.002225 | 1 |
| Rnf8 | 0.012236 | 0.009183 | 1 |
| Zfp266 | 0.012191 | 0.730797 | 1 |
| Hacd4 | 0.012015 | 0.149759 | 1 |
| Sdhaf3 | 0.01198 | 0.00872 | 1 |
| Alyref2 | 0.011946 | 0.045958 | 1 |
| Dnaaf3 | 0.011907 | 0.012793 | 1 |
| Eral1 | 0.011879 | 0.244335 | 1 |
| Eif2b3 | 0.011857 | 0.005046 | 1 |
| Ccar1 | 0.01185 | 0.044132 | 1 |
| Rfc4 | 0.011846 | 0.073589 | 1 |
| Myl6 | 0.01171 | 0.599453 | 1 |
| Alg3 | 0.011675 | 0.050003 | 1 |
| Srpk2 | 0.011611 | 0.070747 | 1 |
| Ghdc | 0.011589 | 0.028878 | 1 |
| Med6 | 0.011546 | 0.014846 | 1 |

| | | | |
|---|---|---|---|
| Dennd2d | 0.011501 | 0.011677 | 1 |
| Irf2 | 0.011478 | 0.667223 | 1 |
| Alad | 0.01142 | 0.002431 | 1 |
| Dync2h1 | 0.011377 | 0.053798 | 1 |
| Armc8 | 0.011308 | 0.01566 | 1 |
| Sephs1 | 0.011302 | 0.072284 | 1 |
| Pus10 | 0.011281 | 0.137869 | 1 |
| Gnpda2 | 0.011255 | 0.018511 | 1 |
| Adrm1 | 0.011247 | 0.027624 | 1 |
| Cand1 | 0.0112 | 0.029255 | 1 |
| Zfp236 | 0.011198 | 0.140314 | 1 |
| Klf7 | 0.011193 | 9.66E-05 | 1 |
| Polrmt | 0.01119 | 0.010968 | 1 |
| Usp36 | 0.011156 | 0.00121 | 1 |
| 2700049A( | 0.011149 | 0.069436 | 1 |
| Dcps | 0.011116 | 0.094322 | 1 |
| Eef2kmt | 0.011078 | 0.014747 | 1 |
| Dhdds | 0.011025 | 0.039835 | 1 |
| Ppif | 0.011023 | 0.25527 | 1 |
| Tmx4 | 0.010903 | 0.089489 | 1 |
| Rpusd4 | 0.010893 | 0.014151 | 1 |
| Pcca | 0.010841 | 0.025894 | 1 |
| Ddx19a | 0.010782 | 0.005496 | 1 |
| Cdc37l1 | 0.010776 | 0.004823 | 1 |
| Haus7 | 0.010737 | 0.020069 | 1 |
| Sec23ip | 0.010718 | 0.107478 | 1 |
| Fbxo9 | 0.01068 | 0.039033 | 1 |
| Mrtfb | 0.010646 | 0.497446 | 1 |
| Coa4 | 0.010608 | 0.003087 | 1 |
| Ccdc71 | 0.010602 | 0.071586 | 1 |
| D16Ertd47 | 0.010581 | 0.019313 | 1 |
| Srpk1 | 0.010561 | 0.004961 | 1 |
| Msh2 | 0.01055 | 0.051476 | 1 |
| Zcchc10 | 0.010505 | 0.117127 | 1 |
| Tor1aip2 | 0.010493 | 0.102398 | 1 |
| 2310015A: | 0.010475 | 0.164768 | 1 |
| Naf1 | 0.010433 | 0.031019 | 1 |
| Zfp451 | 0.010429 | 0.150252 | 1 |
| Mast3 | 0.010428 | 0.044846 | 1 |
| Sfswap | 0.010372 | 0.161309 | 1 |
| Siah2 | 0.010359 | 0.020721 | 1 |
| Taf6l | 0.010356 | 0.085626 | 1 |
| Rasal3 | 0.010349 | 0.030113 | 1 |
| Ly9 | 0.010337 | 0.062267 | 1 |
| Hnrnpm | 0.01032 | 0.126966 | 1 |
| Snx11 | 0.010239 | 0.050606 | 1 |
| Bag4 | 0.010213 | 0.136901 | 1 |
| Gnpat | 0.010096 | 0.328723 | 1 |

| | | | |
|---|---|---|---|
| Mctp2 | 0.010073 | 0.031012 | 1 |
| Ankrd52 | 0.010039 | 0.003359 | 1 |
| Aph1b | 0.010011 | 0.194709 | 1 |
| Strn4 | -0.01006 | 0.001055 | 1 |
| Pde8a | -0.01007 | 0.050173 | 1 |
| Jpt1 | -0.01011 | 0.915481 | 1 |
| Zfp932 | -0.01013 | 0.010176 | 1 |
| Cpd | -0.01017 | 0.062478 | 1 |
| Polr2k | -0.01019 | 0.014656 | 1 |
| Tbkbp1 | -0.01019 | 1.61E-05 | 0.520158 |
| Cdk2 | -0.01025 | 0.595488 | 1 |
| AC154200. | -0.01026 | 0.333719 | 1 |
| Exosc7 | -0.0103 | 0.230315 | 1 |
| Ahsa2 | -0.0103 | 0.500683 | 1 |
| Tbc1d10b | -0.01034 | 0.343634 | 1 |
| Coq9 | -0.01036 | 0.187996 | 1 |
| Cog2 | -0.01039 | 0.308028 | 1 |
| Mmgt2 | -0.01043 | 0.454059 | 1 |
| Thap11 | -0.01046 | 0.106784 | 1 |
| Cul2 | -0.01048 | 0.045412 | 1 |
| Ndufb1-ps | -0.01048 | 0.94187 | 1 |
| Dop1b | -0.01049 | 0.024241 | 1 |
| Cep19 | -0.0105 | 0.599431 | 1 |
| Sdf2 | -0.01062 | 0.108152 | 1 |
| Zkscan8 | -0.01066 | 0.09306 | 1 |
| Tspyl2 | -0.01067 | 0.039969 | 1 |
| Pstpip1 | -0.01079 | 0.57646 | 1 |
| Tmem94 | -0.01085 | 0.101498 | 1 |
| Tk2 | -0.01088 | 0.181692 | 1 |
| Aldh4a1 | -0.01091 | 0.919499 | 1 |
| Cyb5a | -0.01098 | 0.004455 | 1 |
| Slc25a44 | -0.01101 | 0.102916 | 1 |
| C9orf72 | -0.01101 | 0.08749 | 1 |
| Sept10 | -0.01103 | 0.1154 | 1 |
| Recql | -0.01105 | 0.00777 | 1 |
| Cog6 | -0.01106 | 0.440766 | 1 |
| Actb | -0.01108 | 0.645018 | 1 |
| Mrps30 | -0.01109 | 0.1004 | 1 |
| Glmn | -0.01111 | 0.003903 | 1 |
| Zfp280d | -0.01116 | 0.244525 | 1 |
| Cpsf4 | -0.01117 | 0.024928 | 1 |
| Ube2q1 | -0.01118 | 0.452673 | 1 |
| Ap1s3 | -0.01119 | 0.467093 | 1 |
| Tbc1d22a | -0.01119 | 0.036831 | 1 |
| Polk | -0.01121 | 0.277075 | 1 |
| Arl8a | -0.01125 | 0.051542 | 1 |
| Yrdc | -0.0113 | 0.041045 | 1 |
| Cyren | -0.01135 | 0.755546 | 1 |

| | | | |
|---|---|---|---|
| F730043M | -0.01139 | 0.131494 | 1 |
| Dis3l | -0.0114 | 0.397529 | 1 |
| Bcr | -0.01141 | 0.958582 | 1 |
| Mbd1 | -0.01141 | 0.107186 | 1 |
| B9d2 | -0.01144 | 0.477547 | 1 |
| Mbip | -0.01153 | 0.05283 | 1 |
| Zfp90 | -0.01154 | 0.152306 | 1 |
| Blmh | -0.01157 | 0.057669 | 1 |
| Rpl34 | -0.01157 | 0.662546 | 1 |
| Tmem11 | -0.01158 | 0.00172 | 1 |
| Arl3 | -0.01164 | 0.583906 | 1 |
| Nsmce3 | -0.01164 | 0.09106 | 1 |
| Pam | -0.01165 | 0.237835 | 1 |
| Abhd10 | -0.01169 | 0.195693 | 1 |
| Chp1 | -0.01169 | 0.079902 | 1 |
| U2af2 | -0.01178 | 0.019924 | 1 |
| Cdc25a | -0.01179 | 0.021612 | 1 |
| Akap9 | -0.0118 | 0.100026 | 1 |
| Met | -0.01188 | 0.027871 | 1 |
| Depdc5 | -0.01188 | 0.170697 | 1 |
| BC003965 | -0.01198 | 0.192115 | 1 |
| Apex2 | -0.01198 | 0.478139 | 1 |
| Erap1 | -0.01198 | 0.06949 | 1 |
| Hipk3 | -0.01201 | 0.192527 | 1 |
| Tmem43 | -0.01202 | 0.331936 | 1 |
| Nup54 | -0.01202 | 0.10669 | 1 |
| Lcmt2 | -0.01208 | 0.231319 | 1 |
| Zyg11b | -0.01209 | 0.948418 | 1 |
| Cnot8 | -0.0121 | 0.541462 | 1 |
| Rps24 | -0.01213 | 0.428816 | 1 |
| Prim2 | -0.01217 | 0.178604 | 1 |
| Stard3 | -0.01218 | 0.163469 | 1 |
| Gtf3a | -0.0122 | 0.150799 | 1 |
| Ddit3 | -0.01223 | 0.020842 | 1 |
| Arhgef10 | -0.01227 | 0.555592 | 1 |
| Smarcd1 | -0.01231 | 0.009212 | 1 |
| Zbtb33 | -0.01238 | 0.48769 | 1 |
| Cwf19l1 | -0.01244 | 0.282166 | 1 |
| Chmp2a | -0.01245 | 0.939293 | 1 |
| Gatb | -0.01247 | 0.019933 | 1 |
| Pus7l | -0.01254 | 0.134421 | 1 |
| Uevld | -0.01257 | 0.308882 | 1 |
| Setd7 | -0.01258 | 0.258492 | 1 |
| Aaas | -0.01263 | 0.149182 | 1 |
| Ppat | -0.01265 | 0.084857 | 1 |
| Ankrd49 | -0.01272 | 0.199454 | 1 |
| Capn7 | -0.01273 | 0.058321 | 1 |
| Esf1 | -0.01277 | 0.018307 | 1 |

| | | | |
|---|---|---|---|
| Lsm8 | -0.01283 | 0.006064 | 1 |
| 1110059G1 | -0.01283 | 0.284037 | 1 |
| Slc37a3 | -0.01287 | 0.121926 | 1 |
| Dusp7 | -0.01289 | 0.033602 | 1 |
| Aggf1 | -0.01291 | 0.066102 | 1 |
| Pogz | -0.01291 | 0.282637 | 1 |
| C87436 | -0.01297 | 0.141003 | 1 |
| Upf3a | -0.01299 | 0.067334 | 1 |
| Tjap1 | -0.01311 | 0.438084 | 1 |
| Deptor | -0.01313 | 0.142876 | 1 |
| Atxn7l3 | -0.01315 | 0.055756 | 1 |
| Rap2b | -0.01318 | 0.173594 | 1 |
| Qser1 | -0.01319 | 0.320449 | 1 |
| Trim65 | -0.01324 | 0.007733 | 1 |
| Pdcd10 | -0.0133 | 0.632195 | 1 |
| Atxn3 | -0.01331 | 0.203823 | 1 |
| Prpf18 | -0.01343 | 0.32619 | 1 |
| Det1 | -0.01344 | 0.453254 | 1 |
| Pilra | -0.01349 | 0.368042 | 1 |
| Sumf1 | -0.0135 | 0.094157 | 1 |
| Slfn9 | -0.01354 | 0.07326 | 1 |
| Zfp53 | -0.01354 | 0.042117 | 1 |
| Kcmf1 | -0.01355 | 0.054859 | 1 |
| Srrt | -0.01366 | 0.15213 | 1 |
| Tfpt | -0.01366 | 0.011603 | 1 |
| Ddx41 | -0.01366 | 0.096062 | 1 |
| Yars2 | -0.01368 | 0.059969 | 1 |
| Ube2z | -0.01375 | 0.053495 | 1 |
| Tnfaip1 | -0.01376 | 0.113112 | 1 |
| Zfp386 | -0.0138 | 0.187515 | 1 |
| Il15 | -0.01381 | 0.045122 | 1 |
| Prrc2b | -0.01384 | 0.850628 | 1 |
| Ddx52 | -0.01386 | 0.086008 | 1 |
| Map2k6 | -0.01388 | 0.081239 | 1 |
| Emc8 | -0.01392 | 0.09392 | 1 |
| Pigp | -0.01392 | 0.198167 | 1 |
| Mib2 | -0.01392 | 0.346606 | 1 |
| Wdsub1 | -0.01393 | 0.182508 | 1 |
| Alox5ap | -0.01397 | 0.91568 | 1 |
| Gtf2h2 | -0.01399 | 0.127013 | 1 |
| Alkbh8 | -0.01403 | 0.17551 | 1 |
| Taf8 | -0.01405 | 0.083042 | 1 |
| Pacc1 | -0.01408 | 0.47296 | 1 |
| Bclaf1 | -0.01412 | 0.146881 | 1 |
| B4gat1 | -0.01413 | 0.415011 | 1 |
| Mad2l1bp | -0.01413 | 0.092796 | 1 |
| Actl6a | -0.01413 | 0.056316 | 1 |
| Isca2 | -0.01414 | 0.052601 | 1 |

| | | | |
|---|---|---|---|
| Dnajc10 | -0.01416 | 0.03365 | 1 |
| Mrpl46 | -0.01419 | 0.092267 | 1 |
| Casp7 | -0.01421 | 0.030351 | 1 |
| Parp10 | -0.01423 | 0.072993 | 1 |
| Golph3l | -0.01426 | 0.186878 | 1 |
| Pex11b | -0.0143 | 0.060147 | 1 |
| Rnf135 | -0.01431 | 0.072391 | 1 |
| Mfsd5 | -0.01431 | 0.285039 | 1 |
| Plekha2 | -0.01434 | 0.138808 | 1 |
| Spata2 | -0.01437 | 0.231765 | 1 |
| Skiv2l | -0.01437 | 0.275169 | 1 |
| Foxj2 | -0.01437 | 0.031164 | 1 |
| Mrps35 | -0.01437 | 0.119744 | 1 |
| Mrpl34 | -0.01444 | 0.091195 | 1 |
| Acsf3 | -0.01452 | 0.553304 | 1 |
| Cdca4 | -0.01467 | 0.050076 | 1 |
| Chmp5 | -0.01478 | 0.33462 | 1 |
| Med31 | -0.01479 | 0.566505 | 1 |
| Klhl24 | -0.0148 | 0.31648 | 1 |
| Eif2d | -0.01485 | 0.055464 | 1 |
| Thada | -0.01486 | 0.023355 | 1 |
| Ppm1g | -0.01489 | 0.159888 | 1 |
| Fbxw5 | -0.01497 | 0.135977 | 1 |
| Klhl7 | -0.01503 | 0.549696 | 1 |
| Tmem41b | -0.01506 | 0.010854 | 1 |
| Zmat3 | -0.01507 | 0.032445 | 1 |
| Zfp830 | -0.01508 | 0.254459 | 1 |
| Runx3 | -0.0151 | 0.16308 | 1 |
| Ahcyl1 | -0.01512 | 0.000621 | 1 |
| Pex14 | -0.01514 | 0.160805 | 1 |
| Dcun1d3 | -0.01517 | 0.085038 | 1 |
| Aar2 | -0.01519 | 0.041675 | 1 |
| Rrp1b | -0.01525 | 0.072617 | 1 |
| Rnf214 | -0.01527 | 0.152222 | 1 |
| Zfp692 | -0.01531 | 0.630458 | 1 |
| Pik3ca | -0.01538 | 0.070367 | 1 |
| Dop1a | -0.0154 | 0.027074 | 1 |
| Drosha | -0.01542 | 0.38702 | 1 |
| Erc1 | -0.01546 | 0.098975 | 1 |
| Ctnnbip1 | -0.01552 | 0.148595 | 1 |
| Copb2 | -0.01554 | 0.068148 | 1 |
| Yeats2 | -0.01555 | 0.034261 | 1 |
| Ttc4 | -0.01559 | 0.071868 | 1 |
| 1810037I1 | -0.01561 | 0.07083 | 1 |
| Brk1 | -0.01564 | 0.386856 | 1 |
| Spag7 | -0.01568 | 0.32422 | 1 |
| Rnf13 | -0.01572 | 0.784803 | 1 |
| Gdi2 | -0.01575 | 0.695589 | 1 |

| | | | |
|---|---|---|---|
| Hsh2d | -0.01576 | 0.815963 | 1 |
| Gpn3 | -0.01584 | 0.24766 | 1 |
| Las1l | -0.01592 | 0.311733 | 1 |
| Rhot1 | -0.01598 | 0.204023 | 1 |
| Lrrfip2 | -0.01601 | 0.506332 | 1 |
| Casd1 | -0.01606 | 0.25537 | 1 |
| Alg12 | -0.01609 | 0.387801 | 1 |
| Jmjd6 | -0.01611 | 0.196385 | 1 |
| Was | -0.01613 | 0.806143 | 1 |
| Hjurp | -0.01615 | 0.285647 | 1 |
| Idua | -0.01617 | 0.017058 | 1 |
| Slc7a6os | -0.01618 | 0.354192 | 1 |
| Lrp6 | -0.01624 | 0.174179 | 1 |
| Coq8b | -0.01626 | 0.291351 | 1 |
| Oas1c | -0.01635 | 0.17378 | 1 |
| Pde6d | -0.01636 | 0.524844 | 1 |
| Tbccd1 | -0.01637 | 0.056795 | 1 |
| Polr2g | -0.01638 | 0.379655 | 1 |
| Zfp639 | -0.01645 | 0.011697 | 1 |
| Sp140 | -0.01648 | 0.429989 | 1 |
| Nxt1 | -0.01652 | 0.902372 | 1 |
| Ing5 | -0.01652 | 0.048573 | 1 |
| Casp8 | -0.01658 | 0.290336 | 1 |
| Commd8 | -0.01662 | 0.109847 | 1 |
| Cmas | -0.01667 | 0.177934 | 1 |
| Yipf6 | -0.01668 | 0.027063 | 1 |
| Hp | -0.01669 | 0.220066 | 1 |
| Tet3 | -0.01672 | 0.119665 | 1 |
| St3gal5 | -0.01675 | 0.135926 | 1 |
| Exoc7 | -0.01684 | 0.008368 | 1 |
| Acot2 | -0.01684 | 0.375494 | 1 |
| Coq10b | -0.01685 | 0.092987 | 1 |
| Clp1 | -0.01686 | 0.496929 | 1 |
| Ddx27 | -0.01693 | 0.033356 | 1 |
| Glb1l | -0.01694 | 0.786015 | 1 |
| Afmid | -0.01697 | 0.077322 | 1 |
| Dmac2 | -0.01703 | 0.544477 | 1 |
| Mcur1 | -0.01709 | 0.123078 | 1 |
| Hdhd5 | -0.0171 | 0.579466 | 1 |
| Ddx55 | -0.01713 | 0.182021 | 1 |
| Trpv2 | -0.01726 | 0.901539 | 1 |
| Adar | -0.01737 | 0.247207 | 1 |
| D2hgdh | -0.01742 | 0.626169 | 1 |
| Rufy3 | -0.01742 | 0.022424 | 1 |
| Sars | -0.01744 | 0.253252 | 1 |
| Serpinb9 | -0.01746 | 0.073416 | 1 |
| Ddx24 | -0.01746 | 0.743218 | 1 |
| Mboat7 | -0.01751 | 0.132846 | 1 |

| | | | |
|---|---|---|---|
| Zfp516 | -0.01757 | 0.101916 | 1 |
| Stk11ip | -0.01759 | 0.523816 | 1 |
| Ankrd16 | -0.0176 | 0.638941 | 1 |
| Epb41l4aos | -0.01767 | 0.057919 | 1 |
| Mapre1 | -0.01767 | 0.070343 | 1 |
| Lrrc41 | -0.01773 | 0.015893 | 1 |
| Nedd1 | -0.01778 | 0.022514 | 1 |
| Sec22b | -0.01786 | 0.076497 | 1 |
| Rbmx2 | -0.01788 | 0.664836 | 1 |
| Flii | -0.01791 | 0.226144 | 1 |
| Cnpy4 | -0.01792 | 0.050151 | 1 |
| Gpd2 | -0.01793 | 0.10202 | 1 |
| 2410004B1 | -0.018 | 0.061419 | 1 |
| Gtf3c6 | -0.01801 | 0.238624 | 1 |
| Prkacb | -0.01804 | 0.471006 | 1 |
| Myg1 | -0.0181 | 0.019793 | 1 |
| Ccdc117 | -0.0181 | 0.4798 | 1 |
| Jund | -0.01818 | 0.990417 | 1 |
| Tdp2 | -0.01819 | 0.27385 | 1 |
| Phf23 | -0.01821 | 0.094555 | 1 |
| Fkbp3 | -0.01825 | 0.142304 | 1 |
| Pias2 | -0.0184 | 0.167037 | 1 |
| Fkbpl | -0.01842 | 0.798567 | 1 |
| Dnase1l1 | -0.01844 | 0.247973 | 1 |
| Oxsm | -0.01845 | 0.100838 | 1 |
| Endod1 | -0.01845 | 0.541772 | 1 |
| Ropn1l | -0.01847 | 0.104346 | 1 |
| Fnbp4 | -0.0185 | 0.813495 | 1 |
| Shisa5 | -0.01852 | 0.719082 | 1 |
| Emc10 | -0.01856 | 0.051759 | 1 |
| Mterf3 | -0.01862 | 0.221233 | 1 |
| Crkl | -0.01872 | 0.135298 | 1 |
| Pdik1l | -0.01876 | 0.310457 | 1 |
| Slc22a5 | -0.01877 | 0.291928 | 1 |
| Ice1 | -0.01879 | 0.276696 | 1 |
| Fas | -0.01884 | 0.03542 | 1 |
| Dhcr7 | -0.01885 | 0.583162 | 1 |
| Brcc3 | -0.01886 | 0.022046 | 1 |
| Rbm34 | -0.0189 | 0.059902 | 1 |
| Pfn1 | -0.01894 | 0.855772 | 1 |
| Ankrd39 | -0.01902 | 0.084237 | 1 |
| Nckipsd | -0.01907 | 0.220264 | 1 |
| Ptcd1 | -0.01909 | 0.492366 | 1 |
| Gm8909 | -0.01912 | 0.190006 | 1 |
| Zfp617 | -0.01913 | 0.457749 | 1 |
| Smarcal1 | -0.01936 | 0.135292 | 1 |
| Cars2 | -0.01938 | 0.001309 | 1 |
| Zfp729b | -0.0195 | 0.098748 | 1 |

| | | | |
|---|---|---|---|
| Tnip2 | -0.01955 | 0.012975 | 1 |
| Atf6b | -0.01957 | 0.350802 | 1 |
| Ppp2r3c | -0.01958 | 0.301051 | 1 |
| S100a13 | -0.01962 | 0.717978 | 1 |
| Tmem115 | -0.01963 | 0.416366 | 1 |
| Smpd1 | -0.01976 | 0.176638 | 1 |
| 1700025G( | -0.01983 | 0.322187 | 1 |
| Gmfg | -0.01995 | 0.864209 | 1 |
| Srsf7 | -0.02005 | 0.357241 | 1 |
| Ufd1 | -0.0201 | 0.015328 | 1 |
| Cwc15 | -0.02027 | 0.347045 | 1 |
| Txnrd2 | -0.02027 | 0.043366 | 1 |
| Ano10 | -0.02027 | 0.507185 | 1 |
| Pja1 | -0.02034 | 0.691915 | 1 |
| Fdxacb1 | -0.02047 | 0.074547 | 1 |
| Rbm19 | -0.02057 | 0.115188 | 1 |
| Carnmt1 | -0.02057 | 0.562964 | 1 |
| Gm26917 | -0.0206 | 0.530292 | 1 |
| Kctd12 | -0.02064 | 0.246261 | 1 |
| Ice2 | -0.02068 | 0.841716 | 1 |
| Supt7l | -0.0207 | 0.084584 | 1 |
| Brd3 | -0.02071 | 0.08328 | 1 |
| Gtf2h1 | -0.02073 | 0.070488 | 1 |
| Ulk1 | -0.02075 | 0.766074 | 1 |
| Tns4 | -0.02076 | 0.383354 | 1 |
| Cdk7 | -0.02082 | 0.054697 | 1 |
| Myo9b | -0.02094 | 0.273551 | 1 |
| Arl15 | -0.02099 | 0.073915 | 1 |
| Tamm41 | -0.02103 | 0.215224 | 1 |
| Ptrhd1 | -0.02103 | 0.253424 | 1 |
| Gosr1 | -0.02103 | 0.124281 | 1 |
| Acap3 | -0.02103 | 0.322747 | 1 |
| Sfr1 | -0.02106 | 0.664072 | 1 |
| Slc25a37 | -0.02107 | 0.005654 | 1 |
| Zfp668 | -0.0211 | 0.07669 | 1 |
| Plrg1 | -0.02112 | 0.13575 | 1 |
| Idh2 | -0.02115 | 0.000955 | 1 |
| Sbf1 | -0.02135 | 0.188672 | 1 |
| Zfp868 | -0.02135 | 0.091124 | 1 |
| 2810402E2 | -0.02136 | 0.439585 | 1 |
| Babam2 | -0.02142 | 0.570131 | 1 |
| Trmt5 | -0.02143 | 0.060252 | 1 |
| Tcta | -0.02147 | 0.260625 | 1 |
| Slf1 | -0.02147 | 0.011307 | 1 |
| Fam193a | -0.0215 | 0.097691 | 1 |
| Mettl5 | -0.02153 | 0.239914 | 1 |
| Kbtbd3 | -0.02155 | 0.106239 | 1 |
| Mmgt1 | -0.02164 | 0.29502 | 1 |

| | | | |
|---|---|---|---|
| Oser1 | -0.02171 | 0.180849 | 1 |
| Klc1 | -0.02174 | 0.194636 | 1 |
| Dffa | -0.02175 | 0.382908 | 1 |
| Ormdl3 | -0.02175 | 0.738295 | 1 |
| Mthfsd | -0.02175 | 0.127842 | 1 |
| Wdr24 | -0.0218 | 0.217198 | 1 |
| Ppp4r1 | -0.02181 | 0.059803 | 1 |
| Hist2h2ac | -0.02187 | 0.425678 | 1 |
| Unk | -0.02197 | 0.253911 | 1 |
| Neurl3 | -0.02206 | 0.383893 | 1 |
| Pias3 | -0.02206 | 0.204615 | 1 |
| Isoc2b | -0.02208 | 0.612221 | 1 |
| Senp5 | -0.02214 | 0.047104 | 1 |
| Orc5 | -0.0222 | 0.106925 | 1 |
| H2-Q4 | -0.02226 | 0.860714 | 1 |
| Il21r | -0.02226 | 0.045554 | 1 |
| Togaram1 | -0.0223 | 0.005781 | 1 |
| Topors | -0.02235 | 0.161971 | 1 |
| Cep250 | -0.02239 | 0.246786 | 1 |
| Zfp518a | -0.02241 | 0.025035 | 1 |
| Arl13b | -0.02245 | 0.454159 | 1 |
| Mospd3 | -0.02245 | 0.288698 | 1 |
| Enoph1 | -0.02255 | 0.155375 | 1 |
| Slc11a1 | -0.02257 | 0.244513 | 1 |
| Zfp330 | -0.0226 | 0.020513 | 1 |
| Mocos | -0.02266 | 0.371667 | 1 |
| Uty | -0.02267 | 0.295049 | 1 |
| Cutc | -0.02272 | 0.679491 | 1 |
| Rnaset2b | -0.02272 | 0.602632 | 1 |
| Mrpl24 | -0.02276 | 0.257071 | 1 |
| Polr3k | -0.02278 | 0.085515 | 1 |
| Scly | -0.02281 | 0.000219 | 1 |
| Memo1 | -0.02287 | 0.002505 | 1 |
| Cipc | -0.02293 | 0.439969 | 1 |
| Agap2 | -0.023 | 0.510562 | 1 |
| Gm14548 | -0.02306 | 0.048472 | 1 |
| Mettl25 | -0.02312 | 0.272952 | 1 |
| Pxmp4 | -0.02313 | 0.233856 | 1 |
| Zfp160 | -0.02316 | 0.419351 | 1 |
| Smim15 | -0.02319 | 0.204094 | 1 |
| Pick1 | -0.0232 | 0.386034 | 1 |
| Zfp943 | -0.0232 | 0.161923 | 1 |
| Atg2a | -0.0232 | 0.44586 | 1 |
| Rab37 | -0.02327 | 0.191696 | 1 |
| Nadk2 | -0.02329 | 0.867926 | 1 |
| Zfp653 | -0.02342 | 0.23388 | 1 |
| Ppp6r2 | -0.02342 | 0.82726 | 1 |
| Fto | -0.02343 | 0.023665 | 1 |

| | | | |
|---|---|---|---|
| Mettl17 | -0.02348 | 0.254847 | 1 |
| Pdzd11 | -0.02351 | 0.108193 | 1 |
| Itpripl2 | -0.02352 | 0.016191 | 1 |
| Matk | -0.02357 | 0.283173 | 1 |
| Ppa2 | -0.0236 | 0.228685 | 1 |
| Axin1 | -0.02363 | 0.456561 | 1 |
| BC023719 | -0.02364 | 0.091908 | 1 |
| Ncapd3 | -0.02364 | 0.012048 | 1 |
| Ube2f | -0.02366 | 0.904697 | 1 |
| Cep57 | -0.02377 | 0.094126 | 1 |
| Cep104 | -0.02378 | 0.005988 | 1 |
| Bicd2 | -0.02378 | 0.126155 | 1 |
| Pja2 | -0.0238 | 0.679957 | 1 |
| Stxbp2 | -0.02382 | 0.37086 | 1 |
| Nr1h3 | -0.02388 | 0.744118 | 1 |
| Kctd9 | -0.0239 | 0.388975 | 1 |
| Gm39469 | -0.02392 | 0.498122 | 1 |
| Ccdc88b | -0.02393 | 0.780668 | 1 |
| Acbd3 | -0.02394 | 0.231239 | 1 |
| Ercc4 | -0.02398 | 0.22189 | 1 |
| Zfp512 | -0.02402 | 0.046882 | 1 |
| Padi4 | -0.02403 | 0.743366 | 1 |
| Tnpo2 | -0.02405 | 0.400284 | 1 |
| Smg5 | -0.02412 | 0.124588 | 1 |
| 2810403D2 | -0.02412 | 0.944754 | 1 |
| Crls1 | -0.02423 | 0.052522 | 1 |
| Dolk | -0.02425 | 0.502706 | 1 |
| Hmox1 | -0.02429 | 0.001304 | 1 |
| Slc35a1 | -0.02433 | 0.364785 | 1 |
| Surf1 | -0.0244 | 0.059825 | 1 |
| Zfp429 | -0.02449 | 0.279609 | 1 |
| Fig4 | -0.02468 | 0.09835 | 1 |
| Hmgxb4 | -0.02469 | 0.341064 | 1 |
| Arel1 | -0.02469 | 0.063029 | 1 |
| Agtpbp1 | -0.02477 | 0.003345 | 1 |
| Znrf1 | -0.02481 | 0.096013 | 1 |
| Fut11 | -0.02484 | 0.027741 | 1 |
| Ly6e | -0.02491 | 0.25116 | 1 |
| Rnf113a2 | -0.02494 | 0.753636 | 1 |
| Trim37 | -0.02498 | 0.902982 | 1 |
| Arhgap23 | -0.02503 | 0.441899 | 1 |
| Larp4 | -0.02508 | 0.211822 | 1 |
| Tor4a | -0.02508 | 0.976193 | 1 |
| Brms1l | -0.02517 | 0.969999 | 1 |
| Aldh18a1 | -0.02521 | 0.395419 | 1 |
| Tmem259 | -0.02523 | 0.227592 | 1 |
| Gapt | -0.02544 | 0.545045 | 1 |
| Cpox | -0.02544 | 0.085711 | 1 |

| | | | |
|---|---|---|---|
| Ctps2 | -0.02544 | 0.194988 | 1 |
| Usp16 | -0.02546 | 0.004921 | 1 |
| Selenoo | -0.02549 | 0.358071 | 1 |
| Ints10 | -0.0255 | 0.127532 | 1 |
| Dus3l | -0.02553 | 0.471359 | 1 |
| Ikzf5 | -0.02556 | 0.166012 | 1 |
| Strip1 | -0.02557 | 0.11162 | 1 |
| Zscan26 | -0.02564 | 0.333517 | 1 |
| Erlin2 | -0.02567 | 0.131264 | 1 |
| Pgpep1 | -0.02568 | 0.06753 | 1 |
| Patl1 | -0.02573 | 0.193279 | 1 |
| Ccdc77 | -0.02576 | 0.100464 | 1 |
| 2610507I0 | -0.02576 | 0.997677 | 1 |
| Elk3 | -0.02584 | 0.063334 | 1 |
| Clpb | -0.02585 | 0.274941 | 1 |
| Scd2 | -0.02587 | 0.15987 | 1 |
| Gm49336 | -0.0259 | 0.120646 | 1 |
| Polr3f | -0.02593 | 0.458206 | 1 |
| Vamp4 | -0.02593 | 0.058856 | 1 |
| Cox18 | -0.02604 | 0.556306 | 1 |
| Rev3l | -0.02608 | 0.148778 | 1 |
| Polr2j | -0.02608 | 0.225756 | 1 |
| Ddi2 | -0.02608 | 0.21332 | 1 |
| Trmt44 | -0.02613 | 0.216386 | 1 |
| Fbxo8 | -0.02617 | 0.092863 | 1 |
| Slc25a24 | -0.02618 | 0.285205 | 1 |
| Abhd18 | -0.02626 | 0.436342 | 1 |
| Dock6 | -0.02629 | 0.066178 | 1 |
| Rhot2 | -0.02631 | 0.634082 | 1 |
| Cops3 | -0.02639 | 0.272701 | 1 |
| Dhodh | -0.02639 | 0.139308 | 1 |
| Crnkl1 | -0.0264 | 0.425822 | 1 |
| 5830432E( | -0.02641 | 0.851957 | 1 |
| Phpt1 | -0.02647 | 0.753572 | 1 |
| Ankrd54 | -0.02648 | 0.501775 | 1 |
| Fra10ac1 | -0.02652 | 0.038128 | 1 |
| Rnf146 | -0.02652 | 0.089521 | 1 |
| Gan | -0.02654 | 0.382537 | 1 |
| Sec31a | -0.02654 | 0.212613 | 1 |
| Ppp2r1b | -0.02655 | 0.222766 | 1 |
| Cenpc1 | -0.02659 | 0.135828 | 1 |
| Rab11a | -0.02661 | 0.327483 | 1 |
| Rpap1 | -0.02663 | 0.236932 | 1 |
| Zfp91 | -0.02665 | 0.053611 | 1 |
| Ggh | -0.02666 | 0.669459 | 1 |
| Zbtb45 | -0.02671 | 0.368892 | 1 |
| Orc2 | -0.02675 | 0.222142 | 1 |
| Trrap | -0.02675 | 0.075887 | 1 |

| | | | |
|---|---|---|---|
| Ubxn8 | -0.02676 | 0.131552 | 1 |
| Prkaca | -0.02682 | 0.167023 | 1 |
| 2210016F1 | -0.02691 | 0.076055 | 1 |
| Mtmr3 | -0.02692 | 0.245112 | 1 |
| Vps51 | -0.02694 | 0.045077 | 1 |
| Ttll4 | -0.02696 | 0.374535 | 1 |
| Igbp1 | -0.02703 | 0.334195 | 1 |
| 5730455P1 | -0.0271 | 0.570517 | 1 |
| Ppan | -0.02711 | 0.49711 | 1 |
| Dcun1d2 | -0.02711 | 0.101708 | 1 |
| Akt1s1 | -0.02711 | 0.068872 | 1 |
| Pcmtd2 | -0.02722 | 0.547952 | 1 |
| 1700084C0 | -0.02726 | 0.416587 | 1 |
| Zfand4 | -0.0273 | 0.185882 | 1 |
| Sec61g | -0.02734 | 0.464558 | 1 |
| Brf2 | -0.02738 | 0.009304 | 1 |
| Clpx | -0.02743 | 0.014537 | 1 |
| Rttn | -0.02745 | 0.662723 | 1 |
| Phrf1 | -0.02748 | 0.030223 | 1 |
| Psph | -0.02751 | 0.501215 | 1 |
| Noc3l | -0.02752 | 0.00719 | 1 |
| Tm2d1 | -0.02752 | 0.539368 | 1 |
| Mak16 | -0.02755 | 0.090403 | 1 |
| Gm31597 | -0.02756 | 0.124702 | 1 |
| Ppp2r2d | -0.02756 | 0.027227 | 1 |
| Phtf1 | -0.02758 | 0.120431 | 1 |
| Dcp1b | -0.02759 | 0.223712 | 1 |
| Mgat1 | -0.02766 | 0.159071 | 1 |
| Gcn1 | -0.02767 | 0.211683 | 1 |
| Dnajc30 | -0.02769 | 0.024635 | 1 |
| Nars | -0.0277 | 0.221058 | 1 |
| Spcs3 | -0.02775 | 0.49722 | 1 |
| Ccdc130 | -0.02776 | 0.24103 | 1 |
| Scyl2 | -0.02779 | 0.016127 | 1 |
| Ndrg3 | -0.02779 | 0.248359 | 1 |
| Zdhhc9 | -0.02779 | 0.211638 | 1 |
| Lyrm2 | -0.02781 | 0.405225 | 1 |
| Ptar1 | -0.02782 | 0.475936 | 1 |
| Stub1 | -0.02783 | 0.301402 | 1 |
| Actr5 | -0.02785 | 0.371162 | 1 |
| Ttc38 | -0.02787 | 0.370336 | 1 |
| Gmfb | -0.02788 | 0.222821 | 1 |
| Ptdss1 | -0.02801 | 0.18402 | 1 |
| Wdr46 | -0.02803 | 0.017011 | 1 |
| Usp21 | -0.02805 | 0.23305 | 1 |
| Gm36738 | -0.02805 | 0.358741 | 1 |
| Med24 | -0.02808 | 0.159713 | 1 |
| Itpr3 | -0.02822 | 0.10249 | 1 |

| | | | |
|---|---|---|---|
| Nudt22 | -0.02822 | 0.554256 | 1 |
| Trem1 | -0.02823 | 0.382192 | 1 |
| Topbp1 | -0.02823 | 0.200411 | 1 |
| Trappc11 | -0.02829 | 0.089955 | 1 |
| Pank3 | -0.02829 | 0.076434 | 1 |
| Apip | -0.0283 | 0.232189 | 1 |
| Scmh1 | -0.02835 | 0.38412 | 1 |
| Nmral1 | -0.02836 | 0.451036 | 1 |
| Edrf1 | -0.02843 | 0.886892 | 1 |
| Ndufaf1 | -0.02844 | 0.817573 | 1 |
| Usp39 | -0.02851 | 0.116237 | 1 |
| Odr4 | -0.02853 | 0.194098 | 1 |
| Surf2 | -0.02856 | 0.09982 | 1 |
| Cops7b | -0.02856 | 0.185564 | 1 |
| Prpsap1 | -0.02858 | 0.174067 | 1 |
| Parp3 | -0.02863 | 0.121649 | 1 |
| Higd2a | -0.02863 | 0.588865 | 1 |
| Bag5 | -0.02865 | 0.518773 | 1 |
| Mga | -0.02865 | 0.025073 | 1 |
| Mex3c | -0.02866 | 0.185467 | 1 |
| AU040320 | -0.02868 | 0.181628 | 1 |
| Fem1a | -0.02875 | 0.750872 | 1 |
| Gclc | -0.02887 | 0.041351 | 1 |
| Daxx | -0.02891 | 0.232554 | 1 |
| Med8 | -0.02893 | 0.051678 | 1 |
| Rinl | -0.02897 | 0.978643 | 1 |
| Sap30bp | -0.02899 | 0.086511 | 1 |
| Dnm2 | -0.02904 | 0.684016 | 1 |
| Crebzf | -0.02907 | 0.249213 | 1 |
| Pde12 | -0.0291 | 0.478649 | 1 |
| Smad5 | -0.0292 | 0.862142 | 1 |
| Zfp655 | -0.02921 | 0.067953 | 1 |
| Wdyhv1 | -0.02927 | 0.166815 | 1 |
| Gne | -0.02927 | 0.912984 | 1 |
| Mrfap1 | -0.02936 | 0.806434 | 1 |
| Stard4 | -0.0294 | 0.275583 | 1 |
| Phf6 | -0.02948 | 0.689636 | 1 |
| Tcp11l2 | -0.02951 | 0.894242 | 1 |
| Slamf9 | -0.02962 | 0.036688 | 1 |
| Mettl26 | -0.02962 | 0.847564 | 1 |
| Atg16l1 | -0.02974 | 0.027799 | 1 |
| Micu3 | -0.02974 | 0.449149 | 1 |
| Zdhhc18 | -0.02982 | 0.005578 | 1 |
| Emc1 | -0.02987 | 0.121198 | 1 |
| Psmf1 | -0.02988 | 0.323431 | 1 |
| Rnf114 | -0.02992 | 0.904021 | 1 |
| Mtmr10 | -0.02997 | 0.841885 | 1 |
| Fbxo46 | -0.03007 | 0.813834 | 1 |

| | | | |
|---|---|---|---|
| Lsm14b | -0.0302 | 0.564582 | 1 |
| Snx12 | -0.0303 | 0.478205 | 1 |
| Rnf4 | -0.03031 | 0.585544 | 1 |
| Taf1b | -0.03039 | 0.779861 | 1 |
| Hist1h2bc | -0.03062 | 0.051948 | 1 |
| Ogdh | -0.03068 | 0.049633 | 1 |
| Gnpda1 | -0.03069 | 0.033895 | 1 |
| Mitd1 | -0.03071 | 0.092555 | 1 |
| Pcmt1 | -0.03071 | 0.373467 | 1 |
| Nfyb | -0.03074 | 0.344514 | 1 |
| Prorsd1 | -0.03078 | 0.123963 | 1 |
| Mrpl19 | -0.03087 | 0.026679 | 1 |
| Adnp2 | -0.0309 | 0.082566 | 1 |
| Dnajc24 | -0.03091 | 0.379015 | 1 |
| Wipf1 | -0.03091 | 0.752894 | 1 |
| Pds5a | -0.03092 | 0.036943 | 1 |
| Arfip1 | -0.031 | 0.032314 | 1 |
| Avpi1 | -0.031 | 0.101185 | 1 |
| Dctn6 | -0.03102 | 0.529791 | 1 |
| Rab3il1 | -0.03114 | 0.062753 | 1 |
| Cetn2 | -0.03116 | 0.050419 | 1 |
| Gemin5 | -0.03124 | 0.309854 | 1 |
| Dus2 | -0.03135 | 0.193685 | 1 |
| Acsl4 | -0.03136 | 0.119976 | 1 |
| Zfp738 | -0.03139 | 0.108411 | 1 |
| Acox3 | -0.0314 | 0.659317 | 1 |
| 1600020E( | -0.03141 | 0.065376 | 1 |
| Pstk | -0.03147 | 0.817894 | 1 |
| Wrap53 | -0.03148 | 0.791307 | 1 |
| Qtrt1 | -0.03148 | 0.590022 | 1 |
| Bap1 | -0.03148 | 0.560247 | 1 |
| Rtel1 | -0.03156 | 0.307405 | 1 |
| Arpp19 | -0.03156 | 0.512116 | 1 |
| Mon1b | -0.03157 | 0.370187 | 1 |
| Ddx60 | -0.03167 | 0.066283 | 1 |
| Asb13 | -0.03174 | 0.738589 | 1 |
| Lsm1 | -0.03176 | 0.020903 | 1 |
| Zfp687 | -0.03178 | 0.892043 | 1 |
| Ap4e1 | -0.03178 | 0.076329 | 1 |
| Wdr7 | -0.0318 | 0.14099 | 1 |
| Paf1 | -0.0319 | 0.263199 | 1 |
| Cds2 | -0.03215 | 0.33266 | 1 |
| Borcs5 | -0.03218 | 0.09585 | 1 |
| Lbr | -0.03221 | 0.567045 | 1 |
| Ocel1 | -0.03223 | 0.05151 | 1 |
| Taf6 | -0.03241 | 0.105636 | 1 |
| Zbtb9 | -0.03242 | 0.155007 | 1 |
| Aplp2 | -0.03244 | 0.970967 | 1 |

| | | | |
|---|---|---|---|
| Anks1 | -0.03245 | 0.040845 | 1 |
| Pou2f1 | -0.03248 | 0.201767 | 1 |
| Fzr1 | -0.03249 | 0.139876 | 1 |
| Slc35a2 | -0.03255 | 0.594337 | 1 |
| Osgep | -0.03255 | 0.316608 | 1 |
| Etv3 | -0.03257 | 0.198789 | 1 |
| Mrpl57 | -0.03263 | 0.095781 | 1 |
| Rangap1 | -0.03265 | 0.043183 | 1 |
| Anapc7 | -0.03275 | 0.033473 | 1 |
| Rpp30 | -0.03278 | 0.301529 | 1 |
| Zswim6 | -0.03289 | 0.08788 | 1 |
| Zfp688 | -0.03289 | 0.179379 | 1 |
| Cln5 | -0.03309 | 0.048855 | 1 |
| Daam1 | -0.0331 | 0.412924 | 1 |
| Accs | -0.03313 | 0.457203 | 1 |
| Manea | -0.03319 | 0.8345 | 1 |
| Psmg1 | -0.03319 | 0.152044 | 1 |
| Rab1a | -0.03329 | 0.282863 | 1 |
| Mettl2 | -0.03335 | 0.833361 | 1 |
| Sgpl1 | -0.0334 | 0.002214 | 1 |
| Mpzl3 | -0.03342 | 0.575152 | 1 |
| Nt5c3 | -0.03344 | 0.231394 | 1 |
| Csrp2 | -0.03348 | 0.317576 | 1 |
| Ercc6 | -0.03354 | 0.493041 | 1 |
| Snrpb2 | -0.03355 | 0.009672 | 1 |
| Cnp | -0.03355 | 0.043876 | 1 |
| Helz | -0.0336 | 0.036387 | 1 |
| Dgcr2 | -0.03362 | 0.184798 | 1 |
| Fbxw8 | -0.03364 | 0.019881 | 1 |
| 3110056K( | -0.03367 | 0.60094 | 1 |
| Sucla2 | -0.03375 | 0.239412 | 1 |
| Elk4 | -0.03376 | 0.319141 | 1 |
| Gt(ROSA)2( | -0.03376 | 0.771333 | 1 |
| Zfp709 | -0.03377 | 0.254002 | 1 |
| Rpl7 | -0.03377 | 0.389673 | 1 |
| Sergef | -0.03378 | 0.770586 | 1 |
| Stk38l | -0.03382 | 0.864797 | 1 |
| Cul5 | -0.03388 | 0.630791 | 1 |
| Hinfp | -0.03391 | 0.567562 | 1 |
| Polr2c | -0.03394 | 0.584345 | 1 |
| Fam222b | -0.03397 | 0.186784 | 1 |
| Lactb2 | -0.034 | 0.269048 | 1 |
| Mcrip1 | -0.03402 | 0.578962 | 1 |
| Pdrg1 | -0.03406 | 0.6971 | 1 |
| Ppig | -0.03411 | 0.047399 | 1 |
| Fam118b | -0.03419 | 0.441889 | 1 |
| Pspc1 | -0.03419 | 0.171344 | 1 |
| Mbd6 | -0.03423 | 0.107058 | 1 |

| | | | |
|---|---|---|---|
| Gm36486 | -0.03426 | 0.195006 | 1 |
| Fibp | -0.03428 | 0.231053 | 1 |
| Ctbp2 | -0.03428 | 0.064836 | 1 |
| Npat | -0.0343 | 0.988567 | 1 |
| Sirt3 | -0.03437 | 0.338003 | 1 |
| Klhl22 | -0.03438 | 0.018408 | 1 |
| Rdh5 | -0.03443 | 0.341655 | 1 |
| Rai1 | -0.03448 | 0.795562 | 1 |
| Borcs8 | -0.03452 | 0.134091 | 1 |
| 2010315B( | -0.03454 | 0.351105 | 1 |
| Rbm22 | -0.03457 | 0.468288 | 1 |
| Klhl26 | -0.03457 | 0.321048 | 1 |
| Zkscan1 | -0.03457 | 0.300178 | 1 |
| Rfk | -0.03458 | 0.541203 | 1 |
| Uvssa | -0.03464 | 0.099168 | 1 |
| Hbs1l | -0.03466 | 0.441111 | 1 |
| Rhno1 | -0.03467 | 0.799422 | 1 |
| Ppp2r5d | -0.03478 | 0.299169 | 1 |
| Gm10076 | -0.0348 | 0.05646 | 1 |
| Thoc7 | -0.03489 | 0.020873 | 1 |
| Aldh3b1 | -0.03492 | 0.422586 | 1 |
| Fbxw4 | -0.03493 | 0.756108 | 1 |
| Ipo11 | -0.03494 | 0.723669 | 1 |
| Tbp | -0.03499 | 0.255067 | 1 |
| Cnot11 | -0.03505 | 0.043192 | 1 |
| Trim41 | -0.0351 | 0.18078 | 1 |
| Zfp260 | -0.03512 | 0.645233 | 1 |
| Zfp35 | -0.03517 | 0.850288 | 1 |
| Zfp865 | -0.03522 | 0.16339 | 1 |
| Ranbp9 | -0.03523 | 0.306111 | 1 |
| Slc2a9 | -0.03524 | 0.088161 | 1 |
| Bclaf3 | -0.03534 | 0.037856 | 1 |
| Gak | -0.03536 | 0.1104 | 1 |
| Pon2 | -0.03537 | 0.727005 | 1 |
| Fhod1 | -0.03544 | 0.57138 | 1 |
| Cep290 | -0.03549 | 0.130412 | 1 |
| Golga3 | -0.0355 | 0.382663 | 1 |
| Dubr | -0.03555 | 0.022813 | 1 |
| Cmtr1 | -0.03556 | 0.104487 | 1 |
| Sptlc2 | -0.03557 | 0.537819 | 1 |
| Zfp574 | -0.03559 | 0.442048 | 1 |
| Mlxip | -0.03559 | 0.173188 | 1 |
| Abl1 | -0.0356 | 0.932515 | 1 |
| Pigb | -0.03562 | 0.23287 | 1 |
| Rnf41 | -0.03563 | 0.251497 | 1 |
| Tent4b | -0.03581 | 0.026601 | 1 |
| Ap1b1 | -0.03591 | 0.169923 | 1 |
| Braf | -0.03593 | 0.033884 | 1 |

| | | | |
|---|---|---|---|
| Uhmk1 | -0.03593 | 0.054242 | 1 |
| G430095P1 | -0.03601 | 0.1676 | 1 |
| Etfa | -0.03605 | 0.112482 | 1 |
| Cdk2ap1 | -0.03607 | 0.395158 | 1 |
| Faap24 | -0.03613 | 0.139588 | 1 |
| Frmd4a | -0.03615 | 0.6078 | 1 |
| Map7d1 | -0.03617 | 0.220939 | 1 |
| Xaf1 | -0.03617 | 0.40462 | 1 |
| Grsf1 | -0.0362 | 0.060185 | 1 |
| Galnt9 | -0.03625 | 0.592625 | 1 |
| Cks2 | -0.03632 | 0.368593 | 1 |
| Srp14 | -0.03635 | 0.514277 | 1 |
| Taf15 | -0.03636 | 0.222213 | 1 |
| Cant1 | -0.03642 | 0.02102 | 1 |
| Pip4k2b | -0.03646 | 0.042861 | 1 |
| Ncdn | -0.03651 | 0.282898 | 1 |
| Dpf2 | -0.03651 | 0.854002 | 1 |
| Map11 | -0.03652 | 0.019428 | 1 |
| Tbc1d22b | -0.03654 | 0.204137 | 1 |
| Ufl1 | -0.03655 | 0.114547 | 1 |
| Kansl1l | -0.03658 | 0.120855 | 1 |
| Arhgap1 | -0.03661 | 0.401552 | 1 |
| Rpap2 | -0.03662 | 0.128805 | 1 |
| Arl6ip1 | -0.03664 | 0.607649 | 1 |
| Larp4b | -0.03664 | 0.307219 | 1 |
| Tufm | -0.03667 | 0.052689 | 1 |
| Cyhr1 | -0.03668 | 0.196788 | 1 |
| Trappc12 | -0.03669 | 0.302702 | 1 |
| Plscr3 | -0.03672 | 0.318585 | 1 |
| Arfip2 | -0.03673 | 0.037799 | 1 |
| Eif5 | -0.03678 | 0.360721 | 1 |
| Ppp1r13b | -0.03679 | 0.744833 | 1 |
| Mindy3 | -0.03686 | 0.072861 | 1 |
| Yipf5 | -0.03695 | 0.035981 | 1 |
| Tbc1d17 | -0.03696 | 0.263245 | 1 |
| Orc3 | -0.03696 | 0.647587 | 1 |
| Trmt12 | -0.037 | 0.622085 | 1 |
| Ero1lb | -0.03709 | 0.177025 | 1 |
| Ttc32 | -0.0371 | 0.259368 | 1 |
| Ints13 | -0.03713 | 0.34959 | 1 |
| Usp2 | -0.03721 | 0.60915 | 1 |
| Phkg2 | -0.03721 | 0.566079 | 1 |
| Setd5 | -0.03724 | 0.737316 | 1 |
| Pggt1b | -0.03725 | 0.760723 | 1 |
| Gle1 | -0.03732 | 0.618159 | 1 |
| Mettl23 | -0.03735 | 0.38498 | 1 |
| Vps4a | -0.03737 | 0.508675 | 1 |
| Camsap2 | -0.03742 | 0.023338 | 1 |

| | | | |
|---|---|---|---|
| Tm7sf3 | -0.03745 | 0.158092 | 1 |
| Dync1li2 | -0.03745 | 0.287527 | 1 |
| Pex1 | -0.03746 | 0.10313 | 1 |
| Jak2 | -0.0375 | 0.347294 | 1 |
| Rpp25l | -0.03751 | 0.096792 | 1 |
| Ramac | -0.03753 | 0.327067 | 1 |
| Dnttip1 | -0.03755 | 0.013346 | 1 |
| Synrg | -0.03759 | 0.589546 | 1 |
| Trpm7 | -0.03761 | 0.102923 | 1 |
| Hprt | -0.03771 | 0.228668 | 1 |
| Klhl36 | -0.03785 | 0.255609 | 1 |
| Pik3c2a | -0.03793 | 0.100106 | 1 |
| Prrg2 | -0.03794 | 0.102926 | 1 |
| Zfp146 | -0.03799 | 0.955583 | 1 |
| Irf8 | -0.03801 | 0.298342 | 1 |
| Synj2bp | -0.03807 | 0.063561 | 1 |
| Msantd3 | -0.03813 | 0.221458 | 1 |
| Rbl2 | -0.03814 | 0.910868 | 1 |
| Immp1l | -0.03816 | 0.10702 | 1 |
| Apmap | -0.03817 | 0.017688 | 1 |
| Arpc3 | -0.03818 | 0.98947 | 1 |
| Elovl6 | -0.03821 | 0.721326 | 1 |
| Haus5 | -0.03823 | 0.151954 | 1 |
| Sil1 | -0.03828 | 0.025852 | 1 |
| Zbtb34 | -0.03831 | 0.625308 | 1 |
| Prdm4 | -0.03831 | 0.4127 | 1 |
| Ftx | -0.03833 | 0.015329 | 1 |
| Ube2g2 | -0.03839 | 0.377313 | 1 |
| Dpm1 | -0.03845 | 0.660129 | 1 |
| Ptpa | -0.03862 | 0.463569 | 1 |
| Ctse | -0.03863 | 0.700583 | 1 |
| Wdr45b | -0.03868 | 0.525555 | 1 |
| Atxn1l | -0.03869 | 0.647631 | 1 |
| Ric8a | -0.03873 | 0.376201 | 1 |
| Arl5a | -0.03873 | 0.05047 | 1 |
| Dtd2 | -0.03874 | 0.097814 | 1 |
| Qrich1 | -0.03881 | 0.070482 | 1 |
| Srf | -0.03881 | 0.452882 | 1 |
| Trim33 | -0.03882 | 0.118736 | 1 |
| Bcas3 | -0.03883 | 0.128948 | 1 |
| Bcap31 | -0.03884 | 0.707269 | 1 |
| Trmt10c | -0.039 | 0.326769 | 1 |
| Dnajc17 | -0.03902 | 0.31586 | 1 |
| Rnaseh2c | -0.03912 | 0.446007 | 1 |
| Zfp282 | -0.03915 | 0.040131 | 1 |
| Clk2 | -0.03933 | 0.283408 | 1 |
| Hexa | -0.03941 | 0.309259 | 1 |
| Serinc1 | -0.03941 | 0.02619 | 1 |

| | | | |
|---|---|---|---|
| Rbm14 | -0.03945 | 0.270304 | 1 |
| Lats1 | -0.03948 | 0.434552 | 1 |
| Sgcb | -0.03953 | 0.286434 | 1 |
| Prr3 | -0.03958 | 0.063822 | 1 |
| Oma1 | -0.03959 | 0.498363 | 1 |
| Tmub2 | -0.0396 | 0.710004 | 1 |
| Rtn4ip1 | -0.03961 | 0.42193 | 1 |
| Slc38a10 | -0.03962 | 0.164957 | 1 |
| Ighmbp2 | -0.03964 | 0.71174 | 1 |
| Vipas39 | -0.03966 | 0.015099 | 1 |
| Pigyl | -0.03968 | 0.375002 | 1 |
| Tmem209 | -0.03973 | 0.063405 | 1 |
| Eif2s3y | -0.03973 | 0.063921 | 1 |
| Cdkn2aipnl | -0.03976 | 0.456548 | 1 |
| Tmed7 | -0.03978 | 0.53619 | 1 |
| Hadhb | -0.03982 | 0.081019 | 1 |
| Fdps | -0.03984 | 0.10623 | 1 |
| Vps37c | -0.03987 | 0.573626 | 1 |
| Nkap | -0.03987 | 0.496487 | 1 |
| Rif1 | -0.03992 | 0.07233 | 1 |
| Fam20b | -0.03998 | 0.639224 | 1 |
| Mkrn2 | -0.04004 | 0.673203 | 1 |
| Med12 | -0.04005 | 0.033379 | 1 |
| Slc30a5 | -0.04013 | 0.039717 | 1 |
| Zcchc4 | -0.04018 | 0.302888 | 1 |
| Sumo2 | -0.0402 | 0.968611 | 1 |
| Cep85 | -0.04027 | 0.207658 | 1 |
| 2510002D | -0.04028 | 0.746995 | 1 |
| Zfp326 | -0.04029 | 0.157468 | 1 |
| Man2c1 | -0.04029 | 0.347671 | 1 |
| Wdr81 | -0.04032 | 0.283541 | 1 |
| Edc4 | -0.04038 | 0.228014 | 1 |
| Wbp1 | -0.04038 | 0.098554 | 1 |
| Snapin | -0.04038 | 0.030201 | 1 |
| Pbrm1 | -0.04041 | 0.748481 | 1 |
| Cog1 | -0.04055 | 0.089399 | 1 |
| Rtn3 | -0.04057 | 0.211577 | 1 |
| Slc25a1 | -0.04058 | 0.079821 | 1 |
| Dctn4 | -0.04058 | 0.124691 | 1 |
| Stambp | -0.04059 | 0.131233 | 1 |
| Tbcc | -0.04063 | 0.746717 | 1 |
| Kbtbd11 | -0.04072 | 0.929755 | 1 |
| Krr1 | -0.04082 | 0.555185 | 1 |
| Taf2 | -0.04084 | 0.168159 | 1 |
| Zfp385a | -0.04085 | 0.087425 | 1 |
| Irak2 | -0.0409 | 0.158204 | 1 |
| Thoc1 | -0.04093 | 0.054062 | 1 |
| Hcfc1 | -0.04098 | 0.235124 | 1 |

| | | | |
|---|---|---|---|
| Dyrk2 | -0.04102 | 0.18869 | 1 |
| Asb7 | -0.04109 | 0.346356 | 1 |
| Nsun4 | -0.04111 | 0.813292 | 1 |
| Stimate | -0.04111 | 0.311334 | 1 |
| Xpo5 | -0.04116 | 0.173947 | 1 |
| Ank | -0.04116 | 0.94235 | 1 |
| Zbtb6 | -0.04119 | 0.574841 | 1 |
| Rap2a | -0.0412 | 0.204829 | 1 |
| Dnajb1 | -0.04122 | 0.287831 | 1 |
| Slc33a1 | -0.04126 | 0.321118 | 1 |
| Man2a1 | -0.04132 | 0.054568 | 1 |
| Fam241a | -0.04151 | 0.136671 | 1 |
| Slc35b4 | -0.04152 | 0.209848 | 1 |
| Ndfip2 | -0.04152 | 0.263005 | 1 |
| Uvrag | -0.04155 | 0.534246 | 1 |
| S1pr2 | -0.04158 | 0.050374 | 1 |
| Ing3 | -0.04167 | 0.095269 | 1 |
| Gm11944 | -0.04173 | 0.900525 | 1 |
| Zfp422 | -0.04189 | 0.960962 | 1 |
| Snhg20 | -0.04193 | 0.406006 | 1 |
| Cfdp1 | -0.04195 | 0.070599 | 1 |
| Thrap3 | -0.04196 | 0.836182 | 1 |
| Pdzd8 | -0.04198 | 0.180047 | 1 |
| Fpr1 | -0.04202 | 0.009959 | 1 |
| Ints9 | -0.04203 | 0.74526 | 1 |
| Stk16 | -0.04219 | 0.043026 | 1 |
| Ndufs1 | -0.04222 | 0.11129 | 1 |
| Rnf138 | -0.04223 | 0.114238 | 1 |
| Fam210a | -0.04224 | 0.37844 | 1 |
| Wdfy2 | -0.04225 | 0.438497 | 1 |
| Zbtb43 | -0.04239 | 0.134571 | 1 |
| Zfp983 | -0.04239 | 0.783661 | 1 |
| Nagpa | -0.0424 | 0.173025 | 1 |
| Gpn2 | -0.04241 | 0.045796 | 1 |
| Ap1g2 | -0.04244 | 0.279832 | 1 |
| Miip | -0.04246 | 0.362442 | 1 |
| Gpr89 | -0.04252 | 0.392252 | 1 |
| Prxl2c | -0.04254 | 0.053733 | 1 |
| Smu1 | -0.04258 | 0.221802 | 1 |
| Usp4 | -0.0426 | 0.60873 | 1 |
| Dr1 | -0.0426 | 0.224065 | 1 |
| Rrn3 | -0.04262 | 0.956921 | 1 |
| Zfp212 | -0.04263 | 0.156484 | 1 |
| Trappc13 | -0.04265 | 0.471734 | 1 |
| Rab4b | -0.04267 | 0.209231 | 1 |
| Sun2 | -0.04268 | 0.005232 | 1 |
| Zfp346 | -0.04269 | 0.292149 | 1 |
| Rad9a | -0.04269 | 0.638253 | 1 |

| | | | |
|---|---|---|---|
| Brd9 | -0.0427 | 0.284426 | 1 |
| Tmem128 | -0.04271 | 0.271804 | 1 |
| Ppil4 | -0.04275 | 0.320929 | 1 |
| Mtrex | -0.04279 | 0.138546 | 1 |
| Gm13012 | -0.0428 | 0.505736 | 1 |
| Emsy | -0.04285 | 0.525592 | 1 |
| Pcyox1l | -0.04287 | 0.002567 | 1 |
| Pde4a | -0.04287 | 0.464596 | 1 |
| Atp9b | -0.04289 | 0.332014 | 1 |
| CAAA0111{ | -0.0429 | 0.246501 | 1 |
| Faap20 | -0.04296 | 0.395841 | 1 |
| Wtap | -0.04303 | 0.182338 | 1 |
| Armcx3 | -0.04304 | 0.861579 | 1 |
| Srek1ip1 | -0.04314 | 0.028658 | 1 |
| Suclg2 | -0.04315 | 0.157027 | 1 |
| Zfp148 | -0.04315 | 0.92682 | 1 |
| Ap3m1 | -0.04316 | 0.241929 | 1 |
| Tle3 | -0.04317 | 0.050545 | 1 |
| Tbl1xr1 | -0.0432 | 0.208356 | 1 |
| Gtpbp1 | -0.0432 | 0.724729 | 1 |
| Eif4enif1 | -0.04322 | 0.17235 | 1 |
| Srfbp1 | -0.04325 | 0.037314 | 1 |
| Tnks | -0.04327 | 0.039814 | 1 |
| Tctex1d2 | -0.04332 | 0.490569 | 1 |
| Eif2ak4 | -0.04334 | 0.000139 | 1 |
| Stat2 | -0.0434 | 0.169611 | 1 |
| Sacm1l | -0.04341 | 0.588897 | 1 |
| Trappc6b | -0.04348 | 0.109412 | 1 |
| Pfdn6 | -0.04354 | 0.088285 | 1 |
| 4931406P1 | -0.04355 | 0.320488 | 1 |
| Rlf | -0.04376 | 0.130931 | 1 |
| Pctp | -0.04377 | 0.954017 | 1 |
| Tefm | -0.0438 | 0.98101 | 1 |
| Rab21 | -0.04381 | 0.412106 | 1 |
| Usp37 | -0.04383 | 0.15874 | 1 |
| Ngrn | -0.04385 | 0.362326 | 1 |
| Pdcd2l | -0.04389 | 0.141485 | 1 |
| Sass6 | -0.04398 | 0.097402 | 1 |
| Tfcp2 | -0.04405 | 0.404482 | 1 |
| Fam160b2 | -0.04406 | 0.230856 | 1 |
| Slc25a17 | -0.0441 | 0.081793 | 1 |
| Ss18 | -0.04413 | 0.132721 | 1 |
| Rnasek | -0.04413 | 0.431857 | 1 |
| Dbndd2 | -0.04413 | 0.099411 | 1 |
| Fdft1 | -0.04421 | 0.692675 | 1 |
| Phf20 | -0.04426 | 0.066406 | 1 |
| Top1 | -0.04435 | 0.65126 | 1 |
| Cspp1 | -0.04443 | 0.297203 | 1 |

| | | | |
|---|---|---|---|
| Selenoi | -0.04448 | 0.642948 | 1 |
| Trp53bp2 | -0.0445 | 0.307776 | 1 |
| Cenpq | -0.04452 | 0.482678 | 1 |
| Hectd4 | -0.04456 | 0.006517 | 1 |
| Csnk1a1 | -0.04463 | 0.958733 | 1 |
| Mon1a | -0.04464 | 0.452123 | 1 |
| Hdac9 | -0.04467 | 0.914424 | 1 |
| Toe1 | -0.04469 | 0.125937 | 1 |
| L3mbtl2 | -0.04474 | 0.540138 | 1 |
| Cir1 | -0.04475 | 0.21572 | 1 |
| Pex7 | -0.04485 | 0.247554 | 1 |
| Ivns1abp | -0.04496 | 0.193422 | 1 |
| Arid3b | -0.04498 | 0.975298 | 1 |
| Dnajb9 | -0.04499 | 0.697415 | 1 |
| Hif1an | -0.0451 | 0.455966 | 1 |
| Gm16599 | -0.04511 | 0.653408 | 1 |
| Wdr25 | -0.04511 | 0.330391 | 1 |
| Vrk2 | -0.04512 | 0.357467 | 1 |
| Fam160b1 | -0.04516 | 0.402788 | 1 |
| Nprl2 | -0.0452 | 0.027081 | 1 |
| Rgp1 | -0.04521 | 0.182427 | 1 |
| Dhx57 | -0.04526 | 0.679395 | 1 |
| Usp22 | -0.04531 | 0.398925 | 1 |
| 5031439G( | -0.04533 | 0.652078 | 1 |
| Abca2 | -0.04534 | 0.175933 | 1 |
| Prrc1 | -0.04536 | 0.165834 | 1 |
| Arglu1 | -0.04536 | 0.19788 | 1 |
| Fcho2 | -0.04539 | 0.820059 | 1 |
| Zfp780b | -0.04545 | 0.972009 | 1 |
| Anapc2 | -0.04554 | 0.284694 | 1 |
| Kiz | -0.04554 | 0.114741 | 1 |
| Ripk1 | -0.04561 | 0.089954 | 1 |
| Plekhg1 | -0.04563 | 0.323747 | 1 |
| Dgat1 | -0.04576 | 7.4E-05 | 1 |
| Thumpd3 | -0.04584 | 0.061393 | 1 |
| Arf2 | -0.04585 | 0.332885 | 1 |
| Btf3l4 | -0.04587 | 0.455166 | 1 |
| Cactin | -0.04593 | 0.041795 | 1 |
| Cnppd1 | -0.04594 | 0.492372 | 1 |
| Rnf139 | -0.0461 | 0.970962 | 1 |
| Ccdc134 | -0.0461 | 0.15251 | 1 |
| Trdmt1 | -0.04613 | 0.126021 | 1 |
| Appl2 | -0.04613 | 0.009306 | 1 |
| Dele1 | -0.04619 | 0.283043 | 1 |
| Kmt5c | -0.04626 | 0.370105 | 1 |
| Slc50a1 | -0.04626 | 0.70987 | 1 |
| Parp1 | -0.04627 | 0.41635 | 1 |
| Tspan17 | -0.04629 | 0.011018 | 1 |

| | | | |
|---|---|---|---|
| Tada1 | -0.0463 | 0.323913 | 1 |
| Scoc | -0.04631 | 0.755305 | 1 |
| Atg12 | -0.04632 | 0.391395 | 1 |
| Golga7 | -0.04633 | 0.152442 | 1 |
| Ufsp2 | -0.04635 | 0.39263 | 1 |
| Nup188 | -0.04636 | 0.730095 | 1 |
| Ncbp3 | -0.04639 | 0.117897 | 1 |
| Gskip | -0.04644 | 0.461818 | 1 |
| Tcea1 | -0.04654 | 0.489979 | 1 |
| Acads | -0.04655 | 0.379806 | 1 |
| Fadd | -0.04669 | 0.505101 | 1 |
| Ubr2 | -0.0467 | 0.146059 | 1 |
| Snx13 | -0.04674 | 0.481607 | 1 |
| Pcgf1 | -0.04694 | 0.312433 | 1 |
| Trappc10 | -0.04695 | 0.15578 | 1 |
| Dcaf13 | -0.04697 | 0.182172 | 1 |
| Ubqln4 | -0.04707 | 0.858089 | 1 |
| Sec23a | -0.04726 | 0.795261 | 1 |
| Dcaf8 | -0.04726 | 0.51531 | 1 |
| Zfp994 | -0.04727 | 0.778342 | 1 |
| Ccdc174 | -0.04744 | 0.721906 | 1 |
| Lpcat3 | -0.04748 | 0.203595 | 1 |
| Cd2ap | -0.04748 | 0.810815 | 1 |
| Ddit4 | -0.04758 | 0.422824 | 1 |
| Plcb2 | -0.04762 | 0.707996 | 1 |
| Rnf126 | -0.04765 | 0.3938 | 1 |
| Tmem19 | -0.04771 | 0.290696 | 1 |
| Clptm1l | -0.04778 | 0.059661 | 1 |
| Kdm1b | -0.04779 | 0.979284 | 1 |
| Xylt2 | -0.04782 | 0.479192 | 1 |
| Rassf2 | -0.04785 | 0.560529 | 1 |
| Ddx19b | -0.04793 | 0.469714 | 1 |
| Me2 | -0.04797 | 0.901094 | 1 |
| Selenof | -0.048 | 0.507057 | 1 |
| Alg10b | -0.04803 | 0.228532 | 1 |
| Gpatch3 | -0.04804 | 0.293853 | 1 |
| Ier3ip1 | -0.0481 | 0.334614 | 1 |
| Tmem222 | -0.04821 | 0.116479 | 1 |
| Sar1b | -0.04823 | 0.0599 | 1 |
| Mdc1 | -0.04831 | 0.746483 | 1 |
| Polr2l | -0.04846 | 0.340076 | 1 |
| Siglece | -0.04855 | 0.150486 | 1 |
| Xpnpep3 | -0.04857 | 0.289084 | 1 |
| Dcaf11 | -0.04858 | 0.263534 | 1 |
| Mettl14 | -0.04859 | 0.346542 | 1 |
| Eif2ak2 | -0.04866 | 0.324537 | 1 |
| Elac2 | -0.04869 | 0.001332 | 1 |
| Nkiras1 | -0.04875 | 0.386221 | 1 |

| | | | |
|---|---|---|---|
| Slc35a5 | -0.04876 | 0.100134 | 1 |
| Ptcd2 | -0.04877 | 0.345868 | 1 |
| Lmo2 | -0.04881 | 0.612064 | 1 |
| Vps45 | -0.049 | 0.224546 | 1 |
| Ythdf2 | -0.04904 | 0.311738 | 1 |
| March7 | -0.04907 | 0.136576 | 1 |
| Dhrs4 | -0.04927 | 0.629545 | 1 |
| Erp44 | -0.04935 | 0.353021 | 1 |
| Clint1 | -0.04936 | 0.865669 | 1 |
| Epb41 | -0.04939 | 0.151643 | 1 |
| Rfc2 | -0.0494 | 0.259218 | 1 |
| Snrnp200 | -0.04941 | 0.32947 | 1 |
| Tmed3 | -0.04941 | 0.686539 | 1 |
| Rnf34 | -0.04951 | 0.210806 | 1 |
| Strbp | -0.04956 | 0.496168 | 1 |
| Ccnt2 | -0.04965 | 0.277506 | 1 |
| Hcst | -0.04967 | 0.582771 | 1 |
| 1700037C1 | -0.04971 | 0.509184 | 1 |
| Zbed4 | -0.04971 | 0.802119 | 1 |
| Itpkb | -0.04976 | 0.76368 | 1 |
| Kctd2 | -0.04978 | 0.60659 | 1 |
| Sirpb1c | -0.04979 | 0.884504 | 1 |
| Arfgap2 | -0.04982 | 0.221671 | 1 |
| Prpf4 | -0.04988 | 0.269168 | 1 |
| Gm48678 | -0.04989 | 0.860632 | 1 |
| Tcf25 | -0.04993 | 0.900346 | 1 |
| Rab2b | -0.04996 | 0.832699 | 1 |
| Ddx42 | -0.04996 | 0.693917 | 1 |
| Calcoco1 | -0.05003 | 0.703179 | 1 |
| Ankrd10 | -0.05005 | 0.738917 | 1 |
| Ipmk | -0.05006 | 0.361684 | 1 |
| Trim14 | -0.05008 | 0.240742 | 1 |
| Nudt16 | -0.05008 | 0.0886 | 1 |
| Elmod2 | -0.05012 | 0.585941 | 1 |
| Atg2b | -0.05014 | 0.107841 | 1 |
| Med28 | -0.05015 | 0.367985 | 1 |
| Lat2 | -0.05023 | 0.373932 | 1 |
| Hsdl1 | -0.05024 | 0.067897 | 1 |
| C2cd3 | -0.05038 | 0.854683 | 1 |
| Ppard | -0.0504 | 0.011495 | 1 |
| Supt4a | -0.05042 | 0.924585 | 1 |
| Wbp11 | -0.05046 | 0.406159 | 1 |
| Invs | -0.05047 | 0.329344 | 1 |
| A430005L1 | -0.05048 | 0.337475 | 1 |
| Slx1b | -0.05053 | 0.127969 | 1 |
| Utp6 | -0.05061 | 0.217152 | 1 |
| Tmem175 | -0.05076 | 0.568231 | 1 |
| Rab3gap2 | -0.05076 | 0.295132 | 1 |

| | | | |
|---|---|---|---|
| Bet1l | -0.05078 | 0.752838 | 1 |
| Aste1 | -0.05079 | 0.287775 | 1 |
| Ssh1 | -0.0508 | 0.109995 | 1 |
| Aak1 | -0.05083 | 0.06979 | 1 |
| Lrrc8b | -0.05086 | 0.701155 | 1 |
| Lysmd3 | -0.05095 | 0.650692 | 1 |
| Tec | -0.051 | 0.311695 | 1 |
| Cdc73 | -0.05103 | 0.318639 | 1 |
| Usp6nl | -0.05106 | 0.190883 | 1 |
| Gm20513 | -0.05106 | 0.17482 | 1 |
| Polr3c | -0.05106 | 0.564837 | 1 |
| Gm16845 | -0.05114 | 0.831636 | 1 |
| Stt3b | -0.05117 | 0.499522 | 1 |
| Dnajb14 | -0.05118 | 0.065071 | 1 |
| Ap4m1 | -0.05118 | 0.087222 | 1 |
| Exoc1 | -0.05118 | 0.021644 | 1 |
| Pdcd11 | -0.05122 | 0.256721 | 1 |
| Dvl3 | -0.05122 | 0.363039 | 1 |
| Lrrc57 | -0.05122 | 0.238306 | 1 |
| Usp38 | -0.05123 | 0.542062 | 1 |
| Sft2d2 | -0.05134 | 0.050496 | 1 |
| Nde1 | -0.05136 | 0.0385 | 1 |
| Kctd13 | -0.05138 | 0.228239 | 1 |
| Atf7 | -0.05141 | 0.869999 | 1 |
| Pak1 | -0.05141 | 0.106085 | 1 |
| Ireb2 | -0.05154 | 0.485342 | 1 |
| Ttf1 | -0.05163 | 0.036148 | 1 |
| Zfp672 | -0.0517 | 0.445011 | 1 |
| Mark3 | -0.05175 | 0.293077 | 1 |
| Ap5b1 | -0.05184 | 0.571466 | 1 |
| Morf4l1 | -0.05192 | 0.967919 | 1 |
| Mtr | -0.05192 | 0.245929 | 1 |
| Pdss2 | -0.05195 | 0.190275 | 1 |
| Tpst2 | -0.05195 | 0.906395 | 1 |
| Mtf2 | -0.05196 | 0.145267 | 1 |
| Traf7 | -0.05204 | 0.9319 | 1 |
| Recql5 | -0.05205 | 0.840799 | 1 |
| Otud4 | -0.05207 | 0.050765 | 1 |
| Cd2bp2 | -0.05208 | 0.300649 | 1 |
| Hsph1 | -0.05213 | 0.404389 | 1 |
| Cry1 | -0.05218 | 0.277706 | 1 |
| Idi1 | -0.05222 | 0.282892 | 1 |
| Cfap97 | -0.05226 | 0.291228 | 1 |
| Ehmt1 | -0.05228 | 0.497313 | 1 |
| Spty2d1 | -0.05231 | 0.368055 | 1 |
| Xrcc4 | -0.05233 | 0.430132 | 1 |
| Gopc | -0.05233 | 0.453377 | 1 |
| Dedd | -0.05234 | 0.729289 | 1 |

| | | | |
|---|---|---|---|
| Ring1 | -0.05241 | 0.320771 | 1 |
| Arih2 | -0.05245 | 0.986275 | 1 |
| Hmg20a | -0.05249 | 0.233955 | 1 |
| LTO1 | -0.0525 | 0.536253 | 1 |
| 4930523C( | -0.05255 | 0.903636 | 1 |
| Entpd6 | -0.05259 | 0.570412 | 1 |
| Senp3 | -0.05259 | 0.308447 | 1 |
| Sumf2 | -0.05261 | 0.144403 | 1 |
| Sh3bp4 | -0.05263 | 0.391503 | 1 |
| Golga4 | -0.05267 | 0.019087 | 1 |
| Gigyf1 | -0.05268 | 0.270558 | 1 |
| Rab14 | -0.05271 | 0.760056 | 1 |
| Cog4 | -0.05279 | 0.332146 | 1 |
| Txndc16 | -0.05282 | 0.454538 | 1 |
| Rps28 | -0.0529 | 0.217345 | 1 |
| Gm20732 | -0.05295 | 0.635168 | 1 |
| Rnf40 | -0.05296 | 0.207544 | 1 |
| Pnkd | -0.05308 | 0.191128 | 1 |
| Rassf1 | -0.05312 | 0.293 | 1 |
| Ube3b | -0.05316 | 0.128511 | 1 |
| Lrrc61 | -0.05317 | 0.955361 | 1 |
| Nr1d2 | -0.05324 | 0.737536 | 1 |
| Mlh3 | -0.05335 | 0.713397 | 1 |
| Lrch4 | -0.05336 | 0.542759 | 1 |
| Prpsap2 | -0.05339 | 0.414141 | 1 |
| Iqce | -0.05345 | 0.122211 | 1 |
| Fuz | -0.05354 | 0.884131 | 1 |
| Disc1 | -0.0536 | 0.395241 | 1 |
| Lrrc40 | -0.05361 | 0.394994 | 1 |
| Rab10os | -0.05372 | 0.338629 | 1 |
| Rpl39 | -0.05386 | 0.169912 | 1 |
| Xpo7 | -0.05388 | 0.687318 | 1 |
| Trappc4 | -0.05394 | 0.357627 | 1 |
| Tssc4 | -0.05398 | 0.159758 | 1 |
| Mosmo | -0.05409 | 0.262502 | 1 |
| Vhl | -0.05411 | 0.049777 | 1 |
| Tmed4 | -0.05421 | 0.90084 | 1 |
| Cyb5b | -0.05422 | 0.187471 | 1 |
| Dctn1 | -0.05423 | 0.341748 | 1 |
| Yy1 | -0.05425 | 0.060634 | 1 |
| Vav2 | -0.05434 | 0.411775 | 1 |
| Btbd10 | -0.05435 | 0.636778 | 1 |
| Ywhab | -0.0544 | 0.813977 | 1 |
| Ubr5 | -0.05445 | 0.07189 | 1 |
| Lrp10 | -0.05451 | 0.48325 | 1 |
| Cdk9 | -0.05456 | 0.102438 | 1 |
| Ston2 | -0.05457 | 0.233077 | 1 |
| Bphl | -0.05468 | 0.981597 | 1 |

| | | | |
|---|---|---|---|
| Orai2 | -0.05472 | 0.371469 | 1 |
| Ppil2 | -0.05475 | 0.535621 | 1 |
| Nectin1 | -0.05479 | 0.705665 | 1 |
| Pwwp2a | -0.05485 | 0.354686 | 1 |
| Scnm1 | -0.0549 | 0.389214 | 1 |
| Mtmr4 | -0.05494 | 0.808561 | 1 |
| Ubr3 | -0.05499 | 0.255582 | 1 |
| Nktr | -0.055 | 0.10812 | 1 |
| Ifit1bl1 | -0.05504 | 0.529945 | 1 |
| Snip1 | -0.05506 | 0.301948 | 1 |
| Aup1 | -0.05509 | 0.940043 | 1 |
| Gsk3a | -0.05509 | 0.066246 | 1 |
| Cnot4 | -0.0551 | 0.895858 | 1 |
| Reps1 | -0.05518 | 0.269562 | 1 |
| Rabgap1 | -0.05518 | 0.535008 | 1 |
| Dhx35 | -0.05521 | 0.152821 | 1 |
| Pigc | -0.05527 | 0.09921 | 1 |
| 4732471J0 | -0.0553 | 0.451617 | 1 |
| Lyrm1 | -0.05532 | 0.528332 | 1 |
| Zbtb1 | -0.05533 | 0.212533 | 1 |
| Rnf123 | -0.05546 | 0.886746 | 1 |
| Dhx30 | -0.05557 | 0.052955 | 1 |
| Cnnm4 | -0.05557 | 0.727229 | 1 |
| Exoc6 | -0.05559 | 0.775803 | 1 |
| Caap1 | -0.05562 | 0.406046 | 1 |
| Ankrd27 | -0.05568 | 0.305501 | 1 |
| Spsb2 | -0.05582 | 0.867356 | 1 |
| Sprtn | -0.05583 | 0.389325 | 1 |
| U2af1l4 | -0.05583 | 0.863915 | 1 |
| Man1c1 | -0.05584 | 0.285886 | 1 |
| Irgc1 | -0.05588 | 0.528915 | 1 |
| Cstf3 | -0.05588 | 0.175126 | 1 |
| Rxylt1 | -0.05599 | 0.057532 | 1 |
| Uba3 | -0.05601 | 0.295144 | 1 |
| Col4a3bp | -0.05601 | 0.735484 | 1 |
| Slc4a2 | -0.05605 | 0.116338 | 1 |
| Peli1 | -0.05611 | 0.696033 | 1 |
| Map2k7 | -0.05614 | 0.06886 | 1 |
| Slc39a11 | -0.05621 | 0.327517 | 1 |
| Cd300lg | -0.0564 | 0.765019 | 1 |
| Nrde2 | -0.05646 | 0.694823 | 1 |
| Hdac4 | -0.05649 | 0.230301 | 1 |
| Eci1 | -0.05649 | 0.106608 | 1 |
| Lipa | -0.05652 | 0.195794 | 1 |
| Wasl | -0.05655 | 0.879296 | 1 |
| Smox | -0.05661 | 0.422898 | 1 |
| Adck1 | -0.05663 | 0.620483 | 1 |
| Tmem120a | -0.05666 | 0.112458 | 1 |

| | | | |
|---|---|---|---|
| Impa1 | -0.05667 | 0.72241 | 1 |
| Pde3b | -0.05672 | 0.056584 | 1 |
| Gmeb1 | -0.05674 | 0.516103 | 1 |
| Sike1 | -0.05675 | 0.913653 | 1 |
| Hdac3 | -0.05681 | 0.860186 | 1 |
| Uba7 | -0.05685 | 0.51027 | 1 |
| Fam120b | -0.05698 | 0.460922 | 1 |
| Slc41a3 | -0.05699 | 0.298519 | 1 |
| Ccdc102a | -0.05704 | 0.805753 | 1 |
| Mgme1 | -0.05707 | 0.141592 | 1 |
| Nxt2 | -0.05709 | 0.845301 | 1 |
| Setdb1 | -0.05709 | 0.068466 | 1 |
| Ang | -0.05718 | 0.202657 | 1 |
| Ino80b | -0.05718 | 0.068414 | 1 |
| Otulin | -0.05723 | 0.723261 | 1 |
| Rps6kb1 | -0.05726 | 0.204444 | 1 |
| Csnk1d | -0.05737 | 0.637083 | 1 |
| Nlrc5 | -0.05738 | 0.276434 | 1 |
| Srsf1 | -0.05743 | 0.028187 | 1 |
| Gm14326 | -0.05745 | 0.500439 | 1 |
| Lonp2 | -0.05748 | 0.75339 | 1 |
| Exoc2 | -0.05753 | 0.770281 | 1 |
| Morc2a | -0.05755 | 0.229671 | 1 |
| Slc39a6 | -0.05755 | 0.282273 | 1 |
| Bcas2 | -0.05764 | 0.257275 | 1 |
| Kptn | -0.0578 | 0.828493 | 1 |
| Capza2 | -0.05784 | 0.436495 | 1 |
| Ssbp3 | -0.05788 | 0.253913 | 1 |
| Rab29 | -0.0579 | 0.576495 | 1 |
| Ddx1 | -0.05791 | 0.136557 | 1 |
| Pmm2 | -0.05806 | 0.977568 | 1 |
| Sfpq | -0.05807 | 0.644129 | 1 |
| Ppp2r3d | -0.05812 | 0.463457 | 1 |
| Nbr1 | -0.05812 | 0.071462 | 1 |
| Atp6v1a | -0.05814 | 0.77775 | 1 |
| Ppp1r15b | -0.05827 | 0.400413 | 1 |
| Trim56 | -0.05827 | 0.288597 | 1 |
| Spg7 | -0.05833 | 0.345782 | 1 |
| Ap5m1 | -0.05834 | 0.861267 | 1 |
| Mrpl39 | -0.05837 | 0.53904 | 1 |
| Dtx2 | -0.05842 | 0.490495 | 1 |
| Bcl9 | -0.05849 | 0.209446 | 1 |
| Rbm47 | -0.05852 | 0.466729 | 1 |
| Ccdc84 | -0.05854 | 0.870493 | 1 |
| Nrp1 | -0.05856 | 0.751416 | 1 |
| Pak4 | -0.05857 | 0.527984 | 1 |
| Ern1 | -0.05857 | 0.174755 | 1 |
| Lpcat2 | -0.05864 | 0.79959 | 1 |

| | | | |
|---|---|---|---|
| Tmem159 | -0.05865 | 0.556428 | 1 |
| Igf1r | -0.05866 | 0.753938 | 1 |
| Mbtps2 | -0.05869 | 0.392223 | 1 |
| Izumo4 | -0.0587 | 0.393961 | 1 |
| Derl1 | -0.05887 | 0.910216 | 1 |
| Prpf3 | -0.05888 | 0.088429 | 1 |
| Tmem223 | -0.05888 | 0.079095 | 1 |
| Fam98b | -0.0589 | 0.790253 | 1 |
| Upf3b | -0.05892 | 0.114938 | 1 |
| Cbfa2t2 | -0.05894 | 0.48146 | 1 |
| Nsmaf | -0.05894 | 0.491368 | 1 |
| Rreb1 | -0.05903 | 0.871822 | 1 |
| Gtpbp3 | -0.05904 | 0.230853 | 1 |
| Raet1e | -0.05912 | 0.965346 | 1 |
| Nob1 | -0.05914 | 0.374352 | 1 |
| Saysd1 | -0.05922 | 0.137472 | 1 |
| Msl1 | -0.05928 | 0.261773 | 1 |
| Zer1 | -0.05934 | 0.67398 | 1 |
| Eml2 | -0.05936 | 0.400599 | 1 |
| Zkscan14 | -0.05943 | 0.719474 | 1 |
| Atl2 | -0.05948 | 0.747641 | 1 |
| Commd6 | -0.05948 | 0.160595 | 1 |
| Klhl11 | -0.05956 | 0.324105 | 1 |
| Khdc4 | -0.05957 | 0.523897 | 1 |
| Aimp1 | -0.0596 | 0.769412 | 1 |
| D11Wsu47 | -0.05971 | 0.22237 | 1 |
| Herc3 | -0.05973 | 0.598261 | 1 |
| Dda1 | -0.05982 | 0.008482 | 1 |
| Rfwd3 | -0.05987 | 0.942851 | 1 |
| Asb6 | -0.05999 | 0.769104 | 1 |
| Wbp1l | -0.06004 | 0.829246 | 1 |
| Zfp36 | -0.06007 | 0.452443 | 1 |
| Metap1 | -0.06007 | 0.275984 | 1 |
| Pop5 | -0.06012 | 0.634583 | 1 |
| Tmtc3 | -0.06035 | 0.819156 | 1 |
| Pigv | -0.06047 | 0.678592 | 1 |
| Pnpla7 | -0.0605 | 0.974308 | 1 |
| Zfp280c | -0.06056 | 0.924039 | 1 |
| Lsg1 | -0.06066 | 0.06949 | 1 |
| Ercc5 | -0.0607 | 0.712748 | 1 |
| Zc3h18 | -0.0607 | 0.408657 | 1 |
| Efl1 | -0.06074 | 0.932935 | 1 |
| Traf3ip2 | -0.06081 | 0.295097 | 1 |
| Nat2 | -0.06085 | 0.72189 | 1 |
| Ctcf | -0.06088 | 0.074843 | 1 |
| Nono | -0.06088 | 0.599669 | 1 |
| Fam76b | -0.06089 | 0.273103 | 1 |
| Ctsb | -0.06094 | 0.051154 | 1 |

| | | | |
|---|---|---|---|
| Srsf4 | -0.06094 | 0.212684 | 1 |
| Rsad2 | -0.06102 | 0.616601 | 1 |
| Vgll4 | -0.06114 | 0.020777 | 1 |
| Pam16 | -0.06119 | 0.060845 | 1 |
| Tbce | -0.06123 | 0.722414 | 1 |
| Paip2 | -0.06125 | 0.79294 | 1 |
| Hipk1 | -0.06127 | 0.124551 | 1 |
| Avl9 | -0.06128 | 0.480588 | 1 |
| Haghl | -0.06131 | 0.419654 | 1 |
| Snx19 | -0.06137 | 0.652249 | 1 |
| Vwa5a | -0.06142 | 0.461835 | 1 |
| Pithd1 | -0.06146 | 0.397923 | 1 |
| Fxr1 | -0.06157 | 0.277154 | 1 |
| Ddx49 | -0.06157 | 0.069661 | 1 |
| Ikbip | -0.06158 | 0.5631 | 1 |
| Mtap | -0.06177 | 0.676714 | 1 |
| Vps13a | -0.06179 | 0.766125 | 1 |
| Ndufa6 | -0.06179 | 0.359159 | 1 |
| Slc9a1 | -0.06183 | 0.116717 | 1 |
| Cwc27 | -0.06194 | 0.031325 | 1 |
| Sf3b2 | -0.06196 | 0.739267 | 1 |
| St7l | -0.06205 | 0.485502 | 1 |
| Rtn4 | -0.06208 | 0.487716 | 1 |
| Vps11 | -0.06208 | 0.006869 | 1 |
| Shprh | -0.06217 | 0.352885 | 1 |
| Cerkl | -0.0622 | 0.706411 | 1 |
| Ogfod2 | -0.06224 | 0.946968 | 1 |
| B3galt4 | -0.06226 | 0.953903 | 1 |
| Smad1 | -0.06227 | 0.114337 | 1 |
| Rrnad1 | -0.06228 | 0.728767 | 1 |
| Mrs2 | -0.06233 | 0.280725 | 1 |
| Pold3 | -0.06239 | 0.126786 | 1 |
| Plekha3 | -0.06253 | 0.60295 | 1 |
| Fopnl | -0.06272 | 0.902549 | 1 |
| Ppp1r37 | -0.06282 | 0.619204 | 1 |
| Crot | -0.06288 | 0.448668 | 1 |
| Fam50a | -0.06309 | 0.806485 | 1 |
| Lpin1 | -0.06311 | 0.266239 | 1 |
| Appbp2 | -0.06316 | 0.334236 | 1 |
| Xpr1 | -0.06322 | 0.367976 | 1 |
| Pigm | -0.06328 | 0.846839 | 1 |
| Cdk5 | -0.0633 | 0.259623 | 1 |
| Snx27 | -0.0633 | 0.178219 | 1 |
| Bag6 | -0.06351 | 0.049143 | 1 |
| Commd10 | -0.06355 | 0.087982 | 1 |
| Cisd2 | -0.06358 | 0.206571 | 1 |
| Exosc10 | -0.0637 | 0.70043 | 1 |
| Fxr2 | -0.06373 | 0.219013 | 1 |

| | | | |
|---|---|---|---|
| Arap3 | -0.06378 | 0.03884 | 1 |
| Snap23 | -0.06384 | 0.782834 | 1 |
| Diaph1 | -0.06385 | 0.544283 | 1 |
| Pptc7 | -0.06396 | 0.178162 | 1 |
| Coq2 | -0.06396 | 0.839286 | 1 |
| Arpc1b | -0.06406 | 0.334605 | 1 |
| Cyp4f17 | -0.06406 | 0.940306 | 1 |
| Npepps | -0.06408 | 0.14197 | 1 |
| Relb | -0.06411 | 0.418409 | 1 |
| Ldlrad4 | -0.06417 | 0.070689 | 1 |
| Suds3 | -0.06427 | 0.117059 | 1 |
| R3hcc1l | -0.06437 | 0.319647 | 1 |
| Gm45894 | -0.06448 | 0.764478 | 1 |
| Alg1 | -0.06454 | 0.280397 | 1 |
| Ap3m2 | -0.06454 | 0.609148 | 1 |
| Mcu | -0.06456 | 0.989129 | 1 |
| Acad9 | -0.06467 | 0.814611 | 1 |
| Emc3 | -0.06468 | 0.157289 | 1 |
| Mrpl53 | -0.06468 | 0.044514 | 1 |
| Slc25a36 | -0.06478 | 0.683906 | 1 |
| Mtmr2 | -0.06481 | 0.10496 | 1 |
| Prr14 | -0.06486 | 0.133025 | 1 |
| Gm2000 | -0.0649 | 0.816967 | 1 |
| Fam204a | -0.06498 | 0.349732 | 1 |
| Ncoa5 | -0.06502 | 0.253851 | 1 |
| Pts | -0.06503 | 0.891743 | 1 |
| Pum1 | -0.06508 | 0.42787 | 1 |
| C2cd2 | -0.06508 | 0.623285 | 1 |
| Gga1 | -0.06512 | 0.131828 | 1 |
| Bms1 | -0.06514 | 0.143136 | 1 |
| Agpat1 | -0.06521 | 0.422436 | 1 |
| Fancl | -0.06531 | 0.713755 | 1 |
| Mtmr12 | -0.06531 | 0.252592 | 1 |
| Lrrc45 | -0.06537 | 0.805215 | 1 |
| Haus6 | -0.06544 | 0.153092 | 1 |
| Dnajb12 | -0.0655 | 0.474906 | 1 |
| Nf1 | -0.06552 | 0.228409 | 1 |
| Ptpn11 | -0.0656 | 0.51396 | 1 |
| Ccdc186 | -0.06562 | 0.392412 | 1 |
| Spg11 | -0.0657 | 0.478627 | 1 |
| Tapbpl | -0.06575 | 0.234207 | 1 |
| Med14 | -0.06584 | 0.154645 | 1 |
| Prps2 | -0.06584 | 0.475188 | 1 |
| Lin7c | -0.06591 | 0.180879 | 1 |
| Eif3h | -0.06596 | 0.725983 | 1 |
| Utp3 | -0.06596 | 0.943975 | 1 |
| Gm43813 | -0.06597 | 0.620293 | 1 |
| Vps39 | -0.066 | 0.24486 | 1 |

| | | | |
|---|---|---|---|
| Selenok | -0.06601 | 0.900305 | 1 |
| Dcaf17 | -0.06614 | 0.392325 | 1 |
| Acvr1 | -0.06615 | 0.719388 | 1 |
| Ifi206 | -0.06615 | 0.044221 | 1 |
| Zfp839 | -0.0662 | 0.119197 | 1 |
| Usp1 | -0.06621 | 0.468722 | 1 |
| Cacul1 | -0.06624 | 0.479468 | 1 |
| Rev1 | -0.0663 | 0.719473 | 1 |
| Csnk1g1 | -0.06636 | 0.487285 | 1 |
| Ago1 | -0.06646 | 0.285184 | 1 |
| Cstf2t | -0.06647 | 0.596988 | 1 |
| Hps1 | -0.06649 | 0.489834 | 1 |
| B230307C2 | -0.06652 | 0.06069 | 1 |
| Whamm | -0.06655 | 0.180129 | 1 |
| Mfn1 | -0.06655 | 0.316275 | 1 |
| Sel1l | -0.06661 | 0.304419 | 1 |
| Gm38394 | -0.06662 | 0.563398 | 1 |
| Plcl1 | -0.06686 | 0.511746 | 1 |
| Sphk2 | -0.06688 | 0.640716 | 1 |
| Stam | -0.06706 | 0.601642 | 1 |
| Hsd11b1 | -0.06711 | 0.383894 | 1 |
| Ulk2 | -0.06713 | 0.072053 | 1 |
| Gsdme | -0.06724 | 0.665112 | 1 |
| Mfsd10 | -0.06728 | 0.363432 | 1 |
| Plekhg2 | -0.0673 | 0.885271 | 1 |
| Gse1 | -0.06731 | 0.702944 | 1 |
| Ip6k2 | -0.06732 | 0.150817 | 1 |
| Blm | -0.06733 | 0.701194 | 1 |
| Map4 | -0.06735 | 0.829959 | 1 |
| Tab3 | -0.06736 | 0.551575 | 1 |
| 4833438C( | -0.06749 | 0.223438 | 1 |
| Prepl | -0.06751 | 0.600158 | 1 |
| Ppp1r8 | -0.06753 | 0.684168 | 1 |
| Sbno1 | -0.06754 | 0.571366 | 1 |
| Gm15448 | -0.06761 | 0.335213 | 1 |
| Sde2 | -0.06772 | 0.919239 | 1 |
| Rbm41 | -0.06772 | 0.763473 | 1 |
| Map4k1 | -0.06774 | 0.692329 | 1 |
| Ssb | -0.06774 | 0.86631 | 1 |
| Slc35c1 | -0.06779 | 0.467924 | 1 |
| Man2b1 | -0.06783 | 0.216152 | 1 |
| Scyl3 | -0.06784 | 0.259934 | 1 |
| Zscan29 | -0.06789 | 0.205904 | 1 |
| Zfp131 | -0.06797 | 0.673429 | 1 |
| Tmem150b | -0.06801 | 0.80367 | 1 |
| Ptcd3 | -0.06802 | 0.440172 | 1 |
| Ppp1r12b | -0.06828 | 0.005593 | 1 |
| Mre11a | -0.06835 | 0.231058 | 1 |

| | | | |
|---|---|---|---|
| Nfatc2ip | -0.06837 | 0.875715 | 1 |
| Gatc | -0.06839 | 0.600158 | 1 |
| Riox2 | -0.06841 | 0.623564 | 1 |
| Ranbp3 | -0.06841 | 0.264911 | 1 |
| Hexim1 | -0.06841 | 0.661253 | 1 |
| Pcyt2 | -0.06847 | 0.620384 | 1 |
| Diablo | -0.06847 | 0.979843 | 1 |
| Cklf | -0.06851 | 0.869707 | 1 |
| Cog3 | -0.06859 | 0.240015 | 1 |
| Lnpep | -0.06862 | 0.577839 | 1 |
| Smad7 | -0.06873 | 0.793167 | 1 |
| Cers6 | -0.0688 | 0.555071 | 1 |
| Ulk3 | -0.06882 | 0.243046 | 1 |
| Timm17b | -0.06886 | 0.077248 | 1 |
| Wdr53 | -0.069 | 0.781843 | 1 |
| Mrpl4 | -0.06904 | 0.961835 | 1 |
| Fbxo7 | -0.06905 | 0.218072 | 1 |
| Cryzl1 | -0.06909 | 0.341223 | 1 |
| Cyb561d1 | -0.06926 | 0.377828 | 1 |
| Uimc1 | -0.06932 | 0.410465 | 1 |
| Gm44686 | -0.06932 | 0.648628 | 1 |
| Trak1 | -0.06942 | 0.13133 | 1 |
| Rbm48 | -0.06954 | 0.781912 | 1 |
| Cep128 | -0.06958 | 0.952759 | 1 |
| Lias | -0.06971 | 0.363849 | 1 |
| Esrra | -0.06976 | 0.052862 | 1 |
| Ercc1 | -0.06977 | 0.475478 | 1 |
| Ugcg | -0.0698 | 0.498037 | 1 |
| Naa60 | -0.06984 | 0.706421 | 1 |
| Ggps1 | -0.06986 | 0.193472 | 1 |
| Kidins220 | -0.06994 | 0.179636 | 1 |
| 9030025P2 | -0.07004 | 0.898495 | 1 |
| Rabggta | -0.07004 | 0.938645 | 1 |
| Pcnt | -0.0702 | 0.375951 | 1 |
| Rnf144b | -0.07026 | 0.012887 | 1 |
| Zbtb44 | -0.07034 | 0.834498 | 1 |
| Lclat1 | -0.07037 | 0.916795 | 1 |
| Zfp715 | -0.07043 | 0.456686 | 1 |
| Gm16675 | -0.07044 | 0.529808 | 1 |
| Clec16a | -0.07048 | 0.906474 | 1 |
| Pepd | -0.07056 | 0.727236 | 1 |
| Srrm1 | -0.07057 | 0.30343 | 1 |
| Hectd1 | -0.07058 | 0.903346 | 1 |
| Capn15 | -0.07058 | 0.708072 | 1 |
| Slc35f5 | -0.07066 | 0.009008 | 1 |
| P2ry10b | -0.07067 | 0.962076 | 1 |
| Rogdi | -0.07068 | 0.420205 | 1 |
| Xpc | -0.07072 | 0.426149 | 1 |

| | | | |
|---|---|---|---|
| Zfp729a | -0.07075 | 0.881284 | 1 |
| Ncoa1 | -0.07081 | 0.501561 | 1 |
| Flvcr1 | -0.07082 | 0.242263 | 1 |
| Rab32 | -0.07086 | 0.729319 | 1 |
| Mios | -0.0709 | 0.234685 | 1 |
| Milr1 | -0.07096 | 0.820254 | 1 |
| Zfp87 | -0.071 | 0.59794 | 1 |
| Zdhhc7 | -0.07105 | 0.077255 | 1 |
| Mrps27 | -0.0711 | 0.50774 | 1 |
| Plpbp | -0.0711 | 0.505381 | 1 |
| Plec | -0.07118 | 0.940768 | 1 |
| Psmd5 | -0.07125 | 0.419554 | 1 |
| Jkamp | -0.07129 | 0.093566 | 1 |
| Limk1 | -0.07147 | 0.927013 | 1 |
| Lemd3 | -0.07156 | 0.187863 | 1 |
| Eri1 | -0.07156 | 0.443266 | 1 |
| Nol10 | -0.0716 | 0.588364 | 1 |
| Rmnd1 | -0.07161 | 0.190103 | 1 |
| Snx21 | -0.07161 | 0.590009 | 1 |
| Aebp2 | -0.07178 | 0.146633 | 1 |
| Gpalpp1 | -0.07179 | 0.373724 | 1 |
| Emp3 | -0.07183 | 0.621594 | 1 |
| Gm6034 | -0.07188 | 0.874861 | 1 |
| Ebag9 | -0.07193 | 0.498213 | 1 |
| Snhg3 | -0.07197 | 0.069426 | 1 |
| Cep44 | -0.07197 | 0.612561 | 1 |
| Rock1 | -0.07198 | 0.579661 | 1 |
| Nifk | -0.07199 | 0.056755 | 1 |
| Cbx5 | -0.07199 | 0.235173 | 1 |
| March6 | -0.07204 | 0.17608 | 1 |
| Fubp1 | -0.07205 | 0.755393 | 1 |
| Cdk5rap1 | -0.07209 | 0.853896 | 1 |
| Mbtps1 | -0.07209 | 0.135078 | 1 |
| Mto1 | -0.07225 | 0.798694 | 1 |
| Mfap1a | -0.07226 | 0.325602 | 1 |
| Sumo1 | -0.07227 | 0.359881 | 1 |
| Ankrd17 | -0.07236 | 0.727754 | 1 |
| Csnk1g2 | -0.07241 | 0.375607 | 1 |
| Ankib1 | -0.0725 | 0.180687 | 1 |
| Mapk8ip3 | -0.07256 | 0.405197 | 1 |
| Gamt | -0.07257 | 0.457897 | 1 |
| Pum3 | -0.07259 | 0.324712 | 1 |
| Lrif1 | -0.0726 | 0.888756 | 1 |
| Scrib | -0.07268 | 0.912339 | 1 |
| Uba6 | -0.07271 | 0.566755 | 1 |
| Nup133 | -0.07277 | 0.178748 | 1 |
| Numb | -0.0728 | 0.13054 | 1 |
| AI987944 | -0.07282 | 0.624144 | 1 |

| | | | |
|---|---|---|---|
| Coq5 | -0.07311 | 0.032104 | 1 |
| Pkd1 | -0.07316 | 0.707637 | 1 |
| Ifi209 | -0.0732 | 0.314939 | 1 |
| Ak3 | -0.07323 | 0.981248 | 1 |
| Cyfip1 | -0.07326 | 0.873545 | 1 |
| Tsc2 | -0.07326 | 0.737629 | 1 |
| AW549877 | -0.07326 | 0.136504 | 1 |
| Ids | -0.07347 | 0.57319 | 1 |
| Pcyt1a | -0.07355 | 0.555472 | 1 |
| Vrk3 | -0.07356 | 0.466399 | 1 |
| Coq3 | -0.07357 | 0.980273 | 1 |
| Leprot | -0.07364 | 0.234666 | 1 |
| Prkab1 | -0.07367 | 0.40611 | 1 |
| Rundc1 | -0.0737 | 0.13333 | 1 |
| Tnpo1 | -0.07371 | 0.875789 | 1 |
| Btbd1 | -0.07377 | 0.637953 | 1 |
| Wwc2 | -0.07401 | 0.477911 | 1 |
| Lcorl | -0.07402 | 0.281173 | 1 |
| Angel2 | -0.07432 | 0.901055 | 1 |
| Cnbd2 | -0.07446 | 0.390359 | 1 |
| Acsl5 | -0.07448 | 0.931423 | 1 |
| BC031181 | -0.07452 | 0.443107 | 1 |
| Flt3l | -0.07452 | 0.931161 | 1 |
| Ppfibp1 | -0.07454 | 0.191471 | 1 |
| Ints5 | -0.07455 | 0.344904 | 1 |
| Rlim | -0.0747 | 0.888588 | 1 |
| Casp6 | -0.07474 | 0.031833 | 1 |
| Dot1l | -0.07476 | 0.185216 | 1 |
| Cd53 | -0.07477 | 0.521238 | 1 |
| Ranbp10 | -0.07477 | 0.678828 | 1 |
| Tbck | -0.07478 | 0.314886 | 1 |
| Nup98 | -0.07482 | 0.532229 | 1 |
| Pibf1 | -0.0749 | 0.197355 | 1 |
| Hibch | -0.0749 | 0.956849 | 1 |
| Slc35b2 | -0.075 | 0.552788 | 1 |
| Hes6 | -0.07503 | 0.203776 | 1 |
| Rfesd | -0.07505 | 0.990205 | 1 |
| Retreg1 | -0.07508 | 0.133395 | 1 |
| Gpatch8 | -0.07509 | 0.283565 | 1 |
| Cnot6 | -0.07512 | 0.184694 | 1 |
| Kbtbd2 | -0.07515 | 0.716471 | 1 |
| Lypla1 | -0.07523 | 0.731211 | 1 |
| Dck | -0.07529 | 0.561728 | 1 |
| Dpm2 | -0.07535 | 0.317872 | 1 |
| Ino80c | -0.0754 | 0.391209 | 1 |
| Rpl9 | -0.0754 | 0.160951 | 1 |
| Oxsr1 | -0.0754 | 0.793659 | 1 |
| Mtmr1 | -0.07544 | 0.808667 | 1 |

| | | | |
|---|---|---|---|
| Lzic | -0.07545 | 0.223808 | 1 |
| Zfyve1 | -0.07549 | 0.023642 | 1 |
| Tusc1 | -0.07561 | 0.970132 | 1 |
| Gm32036 | -0.07566 | 0.447814 | 1 |
| Ankrd13c | -0.07577 | 0.153153 | 1 |
| Fiz1 | -0.07577 | 0.787066 | 1 |
| Atg101 | -0.07577 | 0.438351 | 1 |
| Dhps | -0.07585 | 0.256468 | 1 |
| Zfp637 | -0.0759 | 0.449906 | 1 |
| Timmdc1 | -0.0759 | 0.660342 | 1 |
| Papolg | -0.07592 | 0.691215 | 1 |
| Pak1ip1 | -0.07603 | 0.392865 | 1 |
| Cltc | -0.07604 | 0.639993 | 1 |
| Cdkal1 | -0.07605 | 0.276422 | 1 |
| Gtf2b | -0.07605 | 0.929092 | 1 |
| Bptf | -0.07606 | 0.445085 | 1 |
| Atxn7l3b | -0.07613 | 0.910179 | 1 |
| Mpp6 | -0.07614 | 0.034267 | 1 |
| Nelfcd | -0.07615 | 0.358637 | 1 |
| Tdrd3 | -0.07619 | 0.711746 | 1 |
| Tmem184b | -0.0762 | 0.190526 | 1 |
| Cdadc1 | -0.07628 | 0.700318 | 1 |
| Abcf3 | -0.07636 | 0.91711 | 1 |
| Mthfs | -0.07637 | 0.27715 | 1 |
| Ccdc93 | -0.07647 | 0.565713 | 1 |
| Ammecr1l | -0.07651 | 0.088855 | 1 |
| Rabep2 | -0.07657 | 0.842598 | 1 |
| Primpol | -0.07668 | 0.905633 | 1 |
| 9930111J2 | -0.07669 | 0.328148 | 1 |
| Aff4 | -0.07675 | 0.399236 | 1 |
| Rgs19 | -0.07685 | 0.417548 | 1 |
| Spata6 | -0.0769 | 0.608152 | 1 |
| Tatdn1 | -0.07691 | 0.539832 | 1 |
| Crebrf | -0.07719 | 0.561123 | 1 |
| Hmgn5 | -0.0772 | 0.897946 | 1 |
| Hhex | -0.07721 | 0.433487 | 1 |
| Bnip3l | -0.07721 | 0.777483 | 1 |
| Mllt10 | -0.07723 | 0.975659 | 1 |
| Trim5 | -0.07726 | 0.753437 | 1 |
| Psmd12 | -0.07733 | 0.872722 | 1 |
| Ccdc9 | -0.07735 | 0.484831 | 1 |
| Rcl1 | -0.07746 | 0.877676 | 1 |
| B4galt4 | -0.0775 | 0.156376 | 1 |
| Cd72 | -0.07753 | 0.011745 | 1 |
| Zfp322a | -0.07753 | 0.574188 | 1 |
| Med17 | -0.0776 | 0.543014 | 1 |
| Brd8 | -0.0776 | 0.281193 | 1 |
| Sgf29 | -0.07763 | 0.505472 | 1 |

| | | | |
|---|---|---|---|
| Flcn | -0.07764 | 0.580425 | 1 |
| Vasp | -0.07768 | 0.939919 | 1 |
| Wdfy3 | -0.07769 | 0.350813 | 1 |
| Sap130 | -0.07769 | 0.226515 | 1 |
| Vars | -0.07771 | 0.852318 | 1 |
| 2310011J0 | -0.07771 | 0.923969 | 1 |
| Stk26 | -0.07772 | 0.093727 | 1 |
| Krcc1 | -0.07776 | 0.471012 | 1 |
| Akirin1 | -0.07778 | 0.707774 | 1 |
| Eya3 | -0.07781 | 0.008831 | 1 |
| Dnaaf2 | -0.07788 | 0.356246 | 1 |
| Zbtb11 | -0.07789 | 0.311239 | 1 |
| Pan2 | -0.07797 | 0.964961 | 1 |
| Fbxo4 | -0.07799 | 0.922562 | 1 |
| Rnf115 | -0.07802 | 0.590195 | 1 |
| Fbxl20 | -0.07813 | 0.315607 | 1 |
| Naa30 | -0.07843 | 0.332015 | 1 |
| Rmi1 | -0.07843 | 0.667183 | 1 |
| Szrd1 | -0.07871 | 0.500244 | 1 |
| Edc3 | -0.07874 | 0.863728 | 1 |
| Ccdc124 | -0.07874 | 0.386896 | 1 |
| Parp2 | -0.07874 | 0.300669 | 1 |
| Phf2 | -0.07882 | 0.752829 | 1 |
| Map4k5 | -0.0789 | 0.619564 | 1 |
| Sord | -0.07898 | 0.439115 | 1 |
| Rbm15 | -0.07901 | 0.695072 | 1 |
| Zfp646 | -0.07903 | 0.258775 | 1 |
| Snx14 | -0.07906 | 0.281697 | 1 |
| Trappc9 | -0.07915 | 0.120202 | 1 |
| Smyd3 | -0.07935 | 0.104498 | 1 |
| Scamp2 | -0.07935 | 0.24757 | 1 |
| Smc3 | -0.07944 | 0.708081 | 1 |
| Foxo1 | -0.07946 | 0.577352 | 1 |
| Cxcr4 | -0.07954 | 0.584526 | 1 |
| Arhgap12 | -0.07974 | 0.032852 | 1 |
| Usp12 | -0.07978 | 0.697078 | 1 |
| Ppwd1 | -0.07979 | 0.251697 | 1 |
| Ralb | -0.07995 | 0.259496 | 1 |
| Map3k20 | -0.08001 | 0.512018 | 1 |
| Zfp397 | -0.08005 | 0.735007 | 1 |
| Ywhaq | -0.08014 | 0.74502 | 1 |
| Snx18 | -0.08014 | 0.709615 | 1 |
| Zfp597 | -0.08017 | 0.956849 | 1 |
| Zranb2 | -0.08017 | 0.219783 | 1 |
| Iws1 | -0.08035 | 0.418118 | 1 |
| Rbmxl1 | -0.08039 | 0.393347 | 1 |
| Polr3gl | -0.08039 | 0.74832 | 1 |
| Trim23 | -0.0804 | 0.327078 | 1 |

| | | | |
|---|---|---|---|
| Smarcad1 | -0.08043 | 0.734508 | 1 |
| Tecr | -0.08044 | 0.870187 | 1 |
| Zfp106 | -0.08049 | 0.696652 | 1 |
| Mlx | -0.0805 | 0.388102 | 1 |
| Larp7 | -0.0805 | 0.228014 | 1 |
| Senp6 | -0.08051 | 0.639819 | 1 |
| Lonrf1 | -0.08052 | 0.880421 | 1 |
| Fam45a | -0.08069 | 0.461603 | 1 |
| Vps26c | -0.08075 | 0.22264 | 1 |
| Gm15417 | -0.08077 | 0.735643 | 1 |
| Pdgfb | -0.08083 | 0.230073 | 1 |
| Gpatch11 | -0.08091 | 0.602717 | 1 |
| Fam219b | -0.08092 | 0.767753 | 1 |
| Mynn | -0.08101 | 0.06457 | 1 |
| Snapc3 | -0.08105 | 0.599812 | 1 |
| Ecpas | -0.08114 | 0.317203 | 1 |
| Fam193b | -0.08116 | 0.897628 | 1 |
| Irak4 | -0.08116 | 0.747337 | 1 |
| Becn1 | -0.08118 | 0.94502 | 1 |
| Hsf2 | -0.08119 | 0.70207 | 1 |
| Wwp1 | -0.08132 | 0.631455 | 1 |
| N6amt1 | -0.08134 | 0.565101 | 1 |
| 6530402F1 | -0.08135 | 0.234484 | 1 |
| Banp | -0.08139 | 0.923308 | 1 |
| 4930453N1 | -0.0814 | 0.333748 | 1 |
| Mbtd1 | -0.08142 | 0.206373 | 1 |
| Sgsh | -0.08165 | 0.343789 | 1 |
| Snrnp40 | -0.08166 | 0.684095 | 1 |
| Vps52 | -0.08194 | 0.957788 | 1 |
| Zbtb18 | -0.08199 | 0.591787 | 1 |
| Dnajc13 | -0.08199 | 0.101103 | 1 |
| Terf2ip | -0.08207 | 0.918891 | 1 |
| Gtf2e2 | -0.08214 | 0.44232 | 1 |
| Erlec1 | -0.08215 | 0.22206 | 1 |
| Smarcc2 | -0.0823 | 0.818297 | 1 |
| Vps37a | -0.08232 | 0.402544 | 1 |
| Timm44 | -0.08233 | 0.847447 | 1 |
| Exosc3 | -0.08239 | 0.826975 | 1 |
| Inpp4a | -0.08248 | 0.818687 | 1 |
| Sirt7 | -0.08252 | 0.545779 | 1 |
| Apopt1 | -0.08257 | 0.748045 | 1 |
| Rhbdd1 | -0.08266 | 0.774445 | 1 |
| Fam160a2 | -0.08271 | 0.626272 | 1 |
| Smad4 | -0.08272 | 0.801806 | 1 |
| Mien1 | -0.08275 | 0.870384 | 1 |
| Rad50 | -0.08281 | 0.204999 | 1 |
| Slc25a32 | -0.08282 | 0.875993 | 1 |
| Rb1cc1 | -0.08296 | 0.933174 | 1 |

| | | | |
|---|---|---|---|
| Zfp329 | -0.08306 | 0.967476 | 1 |
| Dynlt1c | -0.08308 | 0.834915 | 1 |
| Cd68 | -0.08311 | 0.571428 | 1 |
| Hspa1a | -0.08316 | 0.154523 | 1 |
| Pex2 | -0.08325 | 0.018959 | 1 |
| Cnot7 | -0.08336 | 0.622811 | 1 |
| Tut1 | -0.08342 | 0.988051 | 1 |
| Polm | -0.08342 | 0.587356 | 1 |
| Mindy1 | -0.08342 | 0.27288 | 1 |
| Terf2 | -0.08344 | 0.556461 | 1 |
| Nup153 | -0.08373 | 0.333482 | 1 |
| Gpbp1l1 | -0.08375 | 0.537488 | 1 |
| Chuk | -0.08375 | 0.96695 | 1 |
| Nol11 | -0.08384 | 0.549465 | 1 |
| Gm5165 | -0.08386 | 0.660088 | 1 |
| Dedd2 | -0.08399 | 0.112476 | 1 |
| Wbp4 | -0.08402 | 0.462321 | 1 |
| Trim34a | -0.08405 | 0.628822 | 1 |
| Trafd1 | -0.08408 | 0.851482 | 1 |
| Plod3 | -0.08417 | 0.717623 | 1 |
| Snx17 | -0.08423 | 0.574008 | 1 |
| Yipf2 | -0.08425 | 0.178674 | 1 |
| Skap2 | -0.08427 | 0.128913 | 1 |
| Pot1a | -0.0843 | 0.492149 | 1 |
| Cpne3 | -0.08434 | 0.670044 | 1 |
| Clk3 | -0.08438 | 0.795208 | 1 |
| Retreg3 | -0.0844 | 0.814559 | 1 |
| Bloc1s3 | -0.08443 | 0.2624 | 1 |
| Nsun3 | -0.08451 | 0.559775 | 1 |
| Chic2 | -0.08463 | 0.748912 | 1 |
| Naa80 | -0.08465 | 0.561757 | 1 |
| Ctdspl2 | -0.08466 | 0.405057 | 1 |
| 1810013L2 | -0.08473 | 0.582977 | 1 |
| Kif2a | -0.08495 | 0.527197 | 1 |
| Ankfy1 | -0.08496 | 0.673653 | 1 |
| Rptor | -0.08496 | 0.711216 | 1 |
| Erh | -0.08499 | 0.941651 | 1 |
| Poc1b | -0.08513 | 0.44635 | 1 |
| Cpsf6 | -0.08518 | 0.070195 | 1 |
| Fosb | -0.0852 | 0.54087 | 1 |
| Akirin2 | -0.08526 | 0.612289 | 1 |
| Trip12 | -0.08527 | 0.783255 | 1 |
| Xrn1 | -0.08527 | 0.597571 | 1 |
| Bcl2l13 | -0.08537 | 0.938363 | 1 |
| Zfp961 | -0.08545 | 0.617969 | 1 |
| Wdr82 | -0.08551 | 0.835056 | 1 |
| Ap4s1 | -0.08552 | 0.942603 | 1 |
| Tgfbrap1 | -0.08554 | 0.959051 | 1 |

| | | | |
|---|---|---|---|
| Prkag1 | -0.08554 | 0.311495 | 1 |
| 5530601H( | -0.08555 | 0.521658 | 1 |
| Dgat2 | -0.08555 | 0.161717 | 1 |
| Dhx37 | -0.08556 | 0.754301 | 1 |
| Cers2 | -0.08577 | 0.94953 | 1 |
| Pphln1 | -0.0858 | 0.56008 | 1 |
| Trim26 | -0.08582 | 0.02129 | 1 |
| Ubn2 | -0.08583 | 0.213014 | 1 |
| Hnrnpk | -0.08585 | 0.343652 | 1 |
| Bag3 | -0.08585 | 0.292186 | 1 |
| Grb2 | -0.08589 | 0.813254 | 1 |
| Ube2d2a | -0.08598 | 0.372663 | 1 |
| Ociad1 | -0.0862 | 0.69528 | 1 |
| Rspo1 | -0.08622 | 0.381646 | 1 |
| Stambpl1 | -0.08622 | 0.067032 | 1 |
| Ccdc82 | -0.08623 | 0.709447 | 1 |
| Snrpa | -0.08631 | 0.516781 | 1 |
| Gpaa1 | -0.08631 | 0.872391 | 1 |
| Pitrm1 | -0.08636 | 0.939781 | 1 |
| Srsf9 | -0.08638 | 0.422708 | 1 |
| Zdhhc14 | -0.08644 | 0.578113 | 1 |
| Yme1l1 | -0.08648 | 0.521554 | 1 |
| Raph1 | -0.08651 | 0.267567 | 1 |
| Ppp1r9b | -0.08652 | 0.723699 | 1 |
| Zc3h11a | -0.08654 | 0.121212 | 1 |
| Txlng | -0.08657 | 0.825022 | 1 |
| Adnp | -0.08658 | 0.175605 | 1 |
| Klhl18 | -0.08661 | 0.558993 | 1 |
| Nae1 | -0.08669 | 0.25838 | 1 |
| Mtmr6 | -0.08681 | 0.529203 | 1 |
| Rdx | -0.0871 | 0.492958 | 1 |
| Rbm25 | -0.08712 | 0.33934 | 1 |
| E2f3 | -0.08716 | 0.502292 | 1 |
| Anks1b | -0.08717 | 0.979479 | 1 |
| Myo5a | -0.08721 | 0.37787 | 1 |
| Trak2 | -0.08722 | 0.587526 | 1 |
| Tpm1 | -0.08722 | 0.983225 | 1 |
| Kat6b | -0.08723 | 0.16145 | 1 |
| Cdc27 | -0.08723 | 0.047957 | 1 |
| Gzf1 | -0.08734 | 0.896172 | 1 |
| Tsga10 | -0.08741 | 0.831366 | 1 |
| Irgm2 | -0.08749 | 0.286216 | 1 |
| Cic | -0.08756 | 0.870105 | 1 |
| Faim | -0.0876 | 0.207141 | 1 |
| Gm20528 | -0.08762 | 0.760825 | 1 |
| Nudt4 | -0.08763 | 0.481288 | 1 |
| Dcaf12 | -0.08766 | 0.921236 | 1 |
| Rad51d | -0.08777 | 0.632983 | 1 |

| | | | |
|---|---|---|---|
| Elavl1 | -0.08782 | 0.456042 | 1 |
| Parp14 | -0.08788 | 0.826421 | 1 |
| Trmt1l | -0.0879 | 0.993438 | 1 |
| Cdk11b | -0.08796 | 0.660282 | 1 |
| Aspscr1 | -0.08796 | 0.497356 | 1 |
| Ppp6c | -0.08803 | 0.970123 | 1 |
| Rnf11 | -0.08804 | 0.35883 | 1 |
| Vamp2 | -0.08806 | 0.13508 | 1 |
| Nfxl1 | -0.08808 | 0.697219 | 1 |
| Atp2b4 | -0.0882 | 0.058398 | 1 |
| Cnot1 | -0.08825 | 0.766671 | 1 |
| Ube2g1 | -0.08826 | 0.585405 | 1 |
| Zfhx3 | -0.08827 | 0.043635 | 1 |
| Hsf1 | -0.08827 | 0.541564 | 1 |
| Abhd13 | -0.08835 | 0.476204 | 1 |
| Cdc23 | -0.08849 | 0.797526 | 1 |
| Cradd | -0.08851 | 0.263901 | 1 |
| Rraga | -0.08857 | 0.343097 | 1 |
| Abhd17b | -0.08862 | 0.473569 | 1 |
| Epc2 | -0.08866 | 0.328937 | 1 |
| Mau2 | -0.08869 | 0.776219 | 1 |
| Kdm2a | -0.08871 | 0.634864 | 1 |
| Ppp3r1 | -0.08874 | 0.722878 | 1 |
| Nipsnap3b | -0.08877 | 0.080918 | 1 |
| Vps53 | -0.08878 | 0.73376 | 1 |
| Pnpla8 | -0.08878 | 0.73144 | 1 |
| Nfrkb | -0.08881 | 0.715304 | 1 |
| Impad1 | -0.08883 | 0.762182 | 1 |
| Sf3b3 | -0.08883 | 0.642039 | 1 |
| Rnf219 | -0.089 | 0.542801 | 1 |
| Mthfsl | -0.08901 | 0.697246 | 1 |
| 2010013B2 | -0.08902 | 0.022621 | 1 |
| Nploc4 | -0.08909 | 0.049973 | 1 |
| Tango2 | -0.08915 | 0.106417 | 1 |
| Icmt | -0.08916 | 0.078942 | 1 |
| U2surp | -0.0892 | 0.718778 | 1 |
| Kif16b | -0.08923 | 0.572082 | 1 |
| Ppil3 | -0.08923 | 0.121469 | 1 |
| Naa15 | -0.08928 | 0.624547 | 1 |
| Smim27 | -0.0893 | 0.769637 | 1 |
| Prep | -0.08939 | 0.541877 | 1 |
| Golga2 | -0.08957 | 0.370562 | 1 |
| Stat5a | -0.08963 | 0.736823 | 1 |
| Usp7 | -0.08969 | 0.370115 | 1 |
| Pde7a | -0.08973 | 0.536871 | 1 |
| Sesn1 | -0.0898 | 0.606519 | 1 |
| Polr3b | -0.08986 | 0.897008 | 1 |
| Mfng | -0.08986 | 0.347259 | 1 |

| | | | |
|---|---|---|---|
| Pcgf3 | -0.08988 | 0.297688 | 1 |
| Cdkl4 | -0.08989 | 0.542727 | 1 |
| Ccng1 | -0.08999 | 0.462667 | 1 |
| Prkd2 | -0.09005 | 0.254882 | 1 |
| Rfx7 | -0.09008 | 0.77514 | 1 |
| AI467606 | -0.09011 | 0.18517 | 1 |
| Gm6377 | -0.09012 | 0.771284 | 1 |
| Copa | -0.09033 | 0.943755 | 1 |
| Sh2b2 | -0.09035 | 0.24134 | 1 |
| Mrps21 | -0.09046 | 0.640253 | 1 |
| 4921524J1 | -0.09046 | 0.513955 | 1 |
| Sos1 | -0.09048 | 0.513888 | 1 |
| Nkiras2 | -0.09056 | 0.104163 | 1 |
| Ly96 | -0.09059 | 0.649445 | 1 |
| Tspan31 | -0.0906 | 0.221409 | 1 |
| Ccdc28b | -0.09061 | 0.398734 | 1 |
| Dync1li1 | -0.09065 | 0.409013 | 1 |
| Terf1 | -0.09078 | 0.45936 | 1 |
| Stard5 | -0.09082 | 0.460346 | 1 |
| Ptpra | -0.09082 | 0.328744 | 1 |
| Sh3gl1 | -0.09098 | 0.446031 | 1 |
| Acot8 | -0.09114 | 0.675532 | 1 |
| Utp11 | -0.09118 | 0.109734 | 1 |
| Ilk | -0.09118 | 0.662621 | 1 |
| Rnft1 | -0.09121 | 0.724047 | 1 |
| Eloa | -0.09123 | 0.991598 | 1 |
| Sipa1l2 | -0.09125 | 0.878374 | 1 |
| Tubgcp4 | -0.09145 | 0.907057 | 1 |
| Srcap | -0.09148 | 0.514597 | 1 |
| Xxylt1 | -0.09149 | 0.674361 | 1 |
| Cers5 | -0.09149 | 0.647874 | 1 |
| Clns1a | -0.0915 | 0.239596 | 1 |
| Arsa | -0.09158 | 0.437449 | 1 |
| Hlx | -0.09163 | 0.917108 | 1 |
| Mcts1 | -0.09165 | 0.652183 | 1 |
| Rin3 | -0.09187 | 0.57053 | 1 |
| Soat1 | -0.09194 | 0.513267 | 1 |
| Gm26510 | -0.09196 | 0.123944 | 1 |
| Prpf38b | -0.09206 | 0.686032 | 1 |
| Zmym5 | -0.09206 | 0.315137 | 1 |
| Dhx38 | -0.09211 | 0.446147 | 1 |
| Ccndbp1 | -0.09212 | 0.447658 | 1 |
| Letm1 | -0.09216 | 0.950918 | 1 |
| Zdhhc20 | -0.09219 | 0.889417 | 1 |
| 1700123O: | -0.09221 | 0.149889 | 1 |
| Mfap1b | -0.0923 | 0.345763 | 1 |
| Dclre1c | -0.09233 | 0.449906 | 1 |
| Ssbp2 | -0.09233 | 0.23175 | 1 |

| | | | |
|---|---|---|---|
| Usp42 | -0.09234 | 0.697726 | 1 |
| Tifa | -0.09237 | 0.978304 | 1 |
| Tprn | -0.09237 | 0.385015 | 1 |
| Slc26a2 | -0.09239 | 0.737702 | 1 |
| Ttc1 | -0.09241 | 0.608481 | 1 |
| Tex261 | -0.09246 | 0.530077 | 1 |
| Chst15 | -0.09247 | 0.121422 | 1 |
| Tesk2 | -0.0925 | 0.318642 | 1 |
| Samd4b | -0.09254 | 0.227392 | 1 |
| Ncbp1 | -0.09277 | 0.266481 | 1 |
| A930015D( | -0.09289 | 0.977698 | 1 |
| Zfand2b | -0.09291 | 0.427027 | 1 |
| Zfp592 | -0.09292 | 0.976763 | 1 |
| Dusp12 | -0.09296 | 0.722347 | 1 |
| Exosc1 | -0.09303 | 0.532936 | 1 |
| Frs2 | -0.09305 | 0.58321 | 1 |
| Commd7 | -0.09308 | 0.773825 | 1 |
| Thoc5 | -0.09314 | 0.890486 | 1 |
| N4bp2 | -0.09315 | 0.899942 | 1 |
| Exosc4 | -0.09318 | 0.06214 | 1 |
| Ctso | -0.09318 | 0.362048 | 1 |
| Rubcn | -0.0932 | 0.411237 | 1 |
| Srp54a | -0.09328 | 0.47741 | 1 |
| Actr6 | -0.09332 | 0.82609 | 1 |
| Gabpb1 | -0.09336 | 0.772356 | 1 |
| Ube2d1 | -0.09338 | 0.693046 | 1 |
| Rad21 | -0.09349 | 0.858583 | 1 |
| Ifi27 | -0.09349 | 0.008008 | 1 |
| Bcl7b | -0.0935 | 0.523124 | 1 |
| Ywhag | -0.09351 | 0.76299 | 1 |
| Brat1 | -0.09354 | 0.895627 | 1 |
| 5330438D1 | -0.09357 | 0.489897 | 1 |
| Zfp207 | -0.09357 | 0.944792 | 1 |
| D130009I1 | -0.09365 | 0.531651 | 1 |
| Bcdin3d | -0.09374 | 0.273554 | 1 |
| 5430405H( | -0.0938 | 0.698433 | 1 |
| Polg | -0.09391 | 0.690955 | 1 |
| Trip4 | -0.09393 | 0.343499 | 1 |
| Ddx39 | -0.09397 | 0.994283 | 1 |
| Strn | -0.09398 | 0.72877 | 1 |
| Dcun1d1 | -0.09402 | 0.659272 | 1 |
| Ift20 | -0.09403 | 0.617671 | 1 |
| Dcaf10 | -0.0941 | 0.656319 | 1 |
| Edem3 | -0.09421 | 0.231824 | 1 |
| Ythdf1 | -0.09433 | 0.956047 | 1 |
| Mgat5 | -0.09438 | 0.94649 | 1 |
| Agfg1 | -0.0944 | 0.939031 | 1 |
| Snrnp70 | -0.09441 | 0.602835 | 1 |

| | | | |
|---|---|---|---|
| H1f0 | -0.09443 | 0.381359 | 1 |
| Impact | -0.09443 | 0.641739 | 1 |
| Lasp1 | -0.09457 | 0.702501 | 1 |
| Clasrp | -0.09467 | 0.584873 | 1 |
| Zfp410 | -0.09469 | 0.795814 | 1 |
| Rbm18 | -0.09475 | 0.895616 | 1 |
| Cog8 | -0.09483 | 0.496101 | 1 |
| Rngtt | -0.09484 | 0.450564 | 1 |
| Cdc42bpb | -0.09485 | 0.576464 | 1 |
| Nr2c1 | -0.09497 | 0.897628 | 1 |
| Zfp277 | -0.0953 | 0.480461 | 1 |
| Rnaseh1 | -0.09531 | 0.286554 | 1 |
| Hypk | -0.09537 | 0.784551 | 1 |
| Dnajc7 | -0.09542 | 0.727517 | 1 |
| Dynlt1a | -0.09549 | 0.712685 | 1 |
| Fam210b | -0.0955 | 0.133299 | 1 |
| Hdac8 | -0.09552 | 0.649064 | 1 |
| Atp11c | -0.09566 | 0.508842 | 1 |
| Bsdc1 | -0.09567 | 0.595675 | 1 |
| Chd8 | -0.09567 | 0.536712 | 1 |
| Nbeal1 | -0.09583 | 0.285277 | 1 |
| Sirt1 | -0.09584 | 0.593645 | 1 |
| Dock4 | -0.09588 | 0.516777 | 1 |
| Kdm5b | -0.09589 | 0.076379 | 1 |
| Pde1b | -0.09591 | 0.157241 | 1 |
| Lmf1 | -0.09597 | 0.563144 | 1 |
| Csnk1e | -0.09607 | 0.444895 | 1 |
| Irf1 | -0.09609 | 0.919447 | 1 |
| Rgl2 | -0.09615 | 0.753124 | 1 |
| Fam234a | -0.09616 | 0.2256 | 1 |
| Isy1 | -0.09619 | 0.121602 | 1 |
| Naxd | -0.0962 | 0.911638 | 1 |
| Supt16 | -0.09621 | 0.926156 | 1 |
| Abraxas2 | -0.09631 | 0.890561 | 1 |
| BC005624 | -0.09634 | 0.557027 | 1 |
| Zbtb24 | -0.09636 | 0.950642 | 1 |
| Tm6sf1 | -0.09645 | 0.296979 | 1 |
| Il6st | -0.09649 | 0.303197 | 1 |
| Khnyn | -0.09649 | 0.716587 | 1 |
| Ip6k1 | -0.09658 | 0.171399 | 1 |
| St3gal2 | -0.09659 | 0.045177 | 1 |
| Pck2 | -0.09661 | 0.989584 | 1 |
| Dnajc8 | -0.09672 | 0.787803 | 1 |
| Otud5 | -0.09673 | 0.70928 | 1 |
| Parp9 | -0.09678 | 0.652729 | 1 |
| Rpe | -0.09682 | 0.342009 | 1 |
| Kctd20 | -0.09692 | 0.921718 | 1 |
| Acin1 | -0.09706 | 0.765661 | 1 |

| | | | |
|---|---|---|---|
| Mospd2 | -0.0971 | 0.723496 | 1 |
| Cntln | -0.09712 | 0.81296 | 1 |
| Sept11 | -0.09717 | 0.875911 | 1 |
| Ralgps1 | -0.09724 | 0.981969 | 1 |
| Chst12 | -0.09725 | 0.625976 | 1 |
| Tmem60 | -0.09729 | 0.00433 | 1 |
| Tdrd7 | -0.0974 | 0.323768 | 1 |
| Gba2 | -0.09742 | 0.501096 | 1 |
| Tmem234 | -0.09743 | 0.31546 | 1 |
| Snx30 | -0.09751 | 0.331924 | 1 |
| Gcc2 | -0.09764 | 0.480207 | 1 |
| Ssr1 | -0.09772 | 0.760889 | 1 |
| Scai | -0.09773 | 0.042654 | 1 |
| Naa16 | -0.09777 | 0.152505 | 1 |
| Lypla2 | -0.09792 | 0.674658 | 1 |
| Rnd3 | -0.098 | 0.83618 | 1 |
| Taf7 | -0.09807 | 0.879812 | 1 |
| Slc25a11 | -0.09808 | 0.695464 | 1 |
| Numa1 | -0.09812 | 0.874398 | 1 |
| Fis1 | -0.09821 | 0.453135 | 1 |
| Hmgcr | -0.09826 | 0.474557 | 1 |
| Trim8 | -0.09838 | 0.311528 | 1 |
| Parl | -0.09841 | 0.837655 | 1 |
| Itprip | -0.09847 | 0.48386 | 1 |
| Asxl1 | -0.09849 | 0.56444 | 1 |
| Ubn1 | -0.09849 | 0.728806 | 1 |
| Wars2 | -0.09862 | 0.780021 | 1 |
| Fbxo11 | -0.09867 | 0.831371 | 1 |
| Hnrnpu | -0.09871 | 0.061953 | 1 |
| Pcnx3 | -0.09879 | 0.981615 | 1 |
| Ctc1 | -0.09889 | 0.340545 | 1 |
| Casp1 | -0.09894 | 0.985503 | 1 |
| Vps8 | -0.09895 | 0.671795 | 1 |
| Cgas | -0.09898 | 0.720082 | 1 |
| Itsn1 | -0.099 | 0.339341 | 1 |
| Lrig2 | -0.099 | 0.829783 | 1 |
| Arhgef18 | -0.09901 | 0.583603 | 1 |
| Mark4 | -0.09901 | 0.861177 | 1 |
| Slf2 | -0.09914 | 0.624339 | 1 |
| Sntb2 | -0.09927 | 0.420234 | 1 |
| Actn4 | -0.09929 | 0.592423 | 1 |
| Smim12 | -0.0993 | 0.280344 | 1 |
| Cwc25 | -0.09941 | 0.416325 | 1 |
| Men1 | -0.09948 | 0.419075 | 1 |
| Bbip1 | -0.09953 | 0.404282 | 1 |
| Vps4b | -0.09964 | 0.779926 | 1 |
| Evi2 | -0.09977 | 0.009186 | 1 |
| Zfp984 | -0.09992 | 0.687736 | 1 |

| | | | |
|---|---|---|---|
| Casc3 | -0.09994 | 0.793405 | 1 |
| Dnajc14 | -0.09996 | 0.08065 | 1 |
| Tor1a | -0.1 | 0.562839 | 1 |
| B4galt3 | -0.10004 | 0.682634 | 1 |
| Gapvd1 | -0.10027 | 0.323491 | 1 |
| Tmem181a | -0.10037 | 0.279843 | 1 |
| Calcrl | -0.10038 | 0.673522 | 1 |
| Slc17a5 | -0.10038 | 0.74684 | 1 |
| C330007P( | -0.10039 | 0.918155 | 1 |
| Ap1g1 | -0.1006 | 0.51584 | 1 |
| Arnt | -0.1006 | 0.779046 | 1 |
| Fkbp15 | -0.10065 | 0.621351 | 1 |
| Cyth4 | -0.1007 | 0.572669 | 1 |
| Brd7 | -0.10073 | 0.905517 | 1 |
| Atg9a | -0.10082 | 0.244789 | 1 |
| Armc1 | -0.1009 | 0.050507 | 1 |
| Tnf | -0.10091 | 0.918937 | 1 |
| Ube3c | -0.10142 | 0.474283 | 1 |
| Pex6 | -0.10143 | 0.294821 | 1 |
| Med13l | -0.10153 | 0.79435 | 1 |
| Rybp | -0.10153 | 0.074614 | 1 |
| Zfp809 | -0.1017 | 0.511517 | 1 |
| Tpcn1 | -0.1017 | 0.647397 | 1 |
| Auh | -0.10204 | 0.175522 | 1 |
| Bach2 | -0.10205 | 0.166065 | 1 |
| Bmf | -0.10217 | 0.956779 | 1 |
| Ttc9c | -0.10223 | 0.849121 | 1 |
| Rps6kb2 | -0.10226 | 0.717337 | 1 |
| Btaf1 | -0.10227 | 0.179823 | 1 |
| Tgtp2 | -0.10249 | 0.52911 | 1 |
| Sarnp | -0.10257 | 0.892493 | 1 |
| Ugt1a7c | -0.10258 | 0.624351 | 1 |
| Pygo2 | -0.10265 | 0.263393 | 1 |
| Casp8ap2 | -0.10278 | 0.899431 | 1 |
| Slc35b3 | -0.10281 | 0.618354 | 1 |
| Tbc1d13 | -0.10287 | 0.394534 | 1 |
| Med1 | -0.10293 | 0.166875 | 1 |
| Smim11 | -0.10297 | 0.964133 | 1 |
| Efr3a | -0.10298 | 0.144518 | 1 |
| Gm13212 | -0.10319 | 0.942275 | 1 |
| Acvr1b | -0.10327 | 0.884879 | 1 |
| Sntb1 | -0.10328 | 0.013021 | 1 |
| Gm47162 | -0.10331 | 0.01125 | 1 |
| Nf2 | -0.10339 | 0.578255 | 1 |
| Gm27017 | -0.10345 | 0.542265 | 1 |
| Hcfc2 | -0.10349 | 0.937112 | 1 |
| Tmem229b | -0.10353 | 0.892159 | 1 |
| Acot11 | -0.10357 | 0.131989 | 1 |

| | | | |
|---|---|---|---|
| Rasa1 | -0.10362 | 0.27436 | 1 |
| Supt3 | -0.10364 | 0.69035 | 1 |
| Marf1 | -0.10364 | 0.979028 | 1 |
| Snrnp48 | -0.10368 | 0.581174 | 1 |
| Wdr3 | -0.10373 | 0.660789 | 1 |
| Zfp991 | -0.10376 | 0.132076 | 1 |
| Syk | -0.10384 | 0.37182 | 1 |
| Zfp110 | -0.104 | 0.512034 | 1 |
| Safb | -0.1041 | 0.928017 | 1 |
| Exoc6b | -0.10412 | 0.498205 | 1 |
| Spin1 | -0.10412 | 0.522493 | 1 |
| Oas2 | -0.10416 | 0.917337 | 1 |
| Psip1 | -0.10416 | 0.484008 | 1 |
| Cep63 | -0.10421 | 0.630204 | 1 |
| Brpf3 | -0.10433 | 0.201831 | 1 |
| Utp23 | -0.10448 | 0.450897 | 1 |
| Sowahc | -0.10448 | 0.831334 | 1 |
| Nelfb | -0.1045 | 0.813477 | 1 |
| Frg1 | -0.10461 | 0.432876 | 1 |
| Pik3r4 | -0.10473 | 0.915806 | 1 |
| Zfx | -0.10476 | 0.607306 | 1 |
| Zfp644 | -0.10488 | 0.806824 | 1 |
| Zfp945 | -0.10494 | 0.717597 | 1 |
| Hpse | -0.1051 | 0.868449 | 1 |
| Btk | -0.10514 | 0.887224 | 1 |
| B430306N( | -0.10528 | 0.309574 | 1 |
| Ncstn | -0.10541 | 0.828568 | 1 |
| Opa3 | -0.1055 | 0.463813 | 1 |
| Stam2 | -0.1055 | 0.491867 | 1 |
| Znrf2 | -0.10558 | 0.783963 | 1 |
| Isg20 | -0.10573 | 0.325575 | 1 |
| Acaa2 | -0.10574 | 0.303939 | 1 |
| Bckdha | -0.10585 | 0.243641 | 1 |
| Ticam1 | -0.10591 | 0.520687 | 1 |
| Nfyc | -0.10593 | 0.204381 | 1 |
| Man1b1 | -0.10593 | 0.206104 | 1 |
| Cep350 | -0.10595 | 0.485373 | 1 |
| Zfp866 | -0.10598 | 0.955663 | 1 |
| Rpa1 | -0.10599 | 0.583386 | 1 |
| 4833439L1 | -0.10612 | 0.6793 | 1 |
| Fam98a | -0.10613 | 0.832409 | 1 |
| Gm5617 | -0.10617 | 0.456551 | 1 |
| Lilra6 | -0.10619 | 0.435064 | 1 |
| Disp1 | -0.10623 | 0.481294 | 1 |
| Hoxb4 | -0.10627 | 0.116849 | 1 |
| Arih1 | -0.10646 | 0.757586 | 1 |
| Trpc4ap | -0.10649 | 0.587124 | 1 |
| Rnf25 | -0.10656 | 0.986697 | 1 |

| | | | |
|---|---|---|---|
| Vps36 | -0.10661 | 0.68411 | 1 |
| Sp4 | -0.1067 | 0.399296 | 1 |
| Itpr2 | -0.10674 | 0.983795 | 1 |
| Prrc2c | -0.10676 | 0.583402 | 1 |
| Ezh1 | -0.1068 | 0.74936 | 1 |
| Wbp2 | -0.1068 | 0.997232 | 1 |
| Rbm27 | -0.1069 | 0.759593 | 1 |
| Leng9 | -0.10697 | 0.716167 | 1 |
| Chmp1b | -0.10698 | 0.983705 | 1 |
| Hnrnpf | -0.10698 | 0.079682 | 1 |
| Ndufaf3 | -0.10699 | 0.409284 | 1 |
| Camta2 | -0.10707 | 0.909883 | 1 |
| Hars2 | -0.10711 | 0.581506 | 1 |
| Pdcd4 | -0.10724 | 0.22184 | 1 |
| Chordc1 | -0.10734 | 0.346678 | 1 |
| Rpl37 | -0.10741 | 0.020587 | 1 |
| Prcp | -0.10766 | 0.303873 | 1 |
| Lin37 | -0.10776 | 0.702615 | 1 |
| Pqlc3 | -0.1078 | 0.469326 | 1 |
| Dclre1a | -0.1079 | 0.654134 | 1 |
| Mob4 | -0.1079 | 0.958871 | 1 |
| Tra2b | -0.10798 | 0.39909 | 1 |
| Uba52 | -0.10813 | 0.347325 | 1 |
| Pxk | -0.10818 | 0.988734 | 1 |
| Hmbox1 | -0.10818 | 0.656726 | 1 |
| Ano6 | -0.1082 | 0.468969 | 1 |
| Skil | -0.1083 | 0.259261 | 1 |
| Pih1d1 | -0.10835 | 0.78118 | 1 |
| Abtb1 | -0.10836 | 0.999105 | 1 |
| Gpatch2 | -0.10841 | 0.830064 | 1 |
| Ugdh | -0.10843 | 0.565788 | 1 |
| Bcl10 | -0.10843 | 0.116027 | 1 |
| Map3k7 | -0.10846 | 0.626041 | 1 |
| Rnf44 | -0.10865 | 0.887937 | 1 |
| Aldh16a1 | -0.10874 | 0.524971 | 1 |
| Fgd3 | -0.10874 | 0.953469 | 1 |
| Mylip | -0.1088 | 0.717509 | 1 |
| Akr1b3 | -0.10882 | 0.619224 | 1 |
| Rbbp7 | -0.10892 | 0.600448 | 1 |
| Lmbr1l | -0.10895 | 0.365094 | 1 |
| Zkscan3 | -0.10896 | 0.155163 | 1 |
| Ifi213 | -0.10904 | 0.027146 | 1 |
| Mff | -0.10928 | 0.938981 | 1 |
| Tmem167b | -0.10941 | 0.976903 | 1 |
| Adss | -0.10946 | 0.658437 | 1 |
| Naa10 | -0.10948 | 0.491961 | 1 |
| Ppp1r35 | -0.10955 | 0.735531 | 1 |
| Kpna6 | -0.10958 | 0.833404 | 1 |

| | | | |
|---|---|---|---|
| Abr | -0.10959 | 0.698229 | 1 |
| Amfr | -0.10961 | 0.103767 | 1 |
| Utp14a | -0.10968 | 0.76264 | 1 |
| Ubap1 | -0.10969 | 0.332265 | 1 |
| Twistnb | -0.10976 | 0.673284 | 1 |
| Frrs1 | -0.10987 | 0.794876 | 1 |
| Uxs1 | -0.10997 | 0.991654 | 1 |
| Hps4 | -0.11003 | 0.715418 | 1 |
| AW011738 | -0.11018 | 0.032056 | 1 |
| 4930581F2 | -0.11019 | 0.053186 | 1 |
| Gmpr2 | -0.11024 | 0.452622 | 1 |
| Gm15964 | -0.11025 | 0.131923 | 1 |
| Dync1i2 | -0.11028 | 0.339606 | 1 |
| Tmem59 | -0.11033 | 0.246735 | 1 |
| Snhg15 | -0.11042 | 0.984773 | 1 |
| Emilin2 | -0.11045 | 0.375709 | 1 |
| Rab5b | -0.11048 | 0.548272 | 1 |
| Elf2 | -0.11051 | 0.325915 | 1 |
| Zmym4 | -0.11053 | 0.683103 | 1 |
| Usp45 | -0.11065 | 0.473421 | 1 |
| Eif4ebp2 | -0.11066 | 0.411562 | 1 |
| A730081D( | -0.11073 | 0.268123 | 1 |
| Eef1akmt2 | -0.11078 | 0.224428 | 1 |
| Tgoln1 | -0.11084 | 0.78545 | 1 |
| Mafg | -0.11109 | 0.709467 | 1 |
| Irf2bpl | -0.11113 | 0.925894 | 1 |
| Plekhm1 | -0.1113 | 0.840028 | 1 |
| Stim1 | -0.1113 | 0.388925 | 1 |
| Faf1 | -0.11142 | 0.127044 | 1 |
| Tnks2 | -0.11142 | 0.853851 | 1 |
| Lsm10 | -0.11145 | 0.683201 | 1 |
| Cyb561a3 | -0.11147 | 0.573769 | 1 |
| Grcc10 | -0.11152 | 0.259372 | 1 |
| Rab8b | -0.11159 | 0.817682 | 1 |
| Bicra | -0.11162 | 0.589235 | 1 |
| Gm15832 | -0.11162 | 0.081166 | 1 |
| Atp1a1 | -0.11188 | 0.626283 | 1 |
| D6Wsu163 | -0.11194 | 0.70391 | 1 |
| Plcl2 | -0.11195 | 0.821257 | 1 |
| Trim24 | -0.11202 | 0.12267 | 1 |
| Nelfa | -0.11205 | 0.606249 | 1 |
| Entpd7 | -0.1122 | 0.699125 | 1 |
| Gm3448 | -0.11224 | 0.255426 | 1 |
| Herpud2 | -0.11248 | 0.45849 | 1 |
| Dhx40 | -0.11272 | 0.907928 | 1 |
| Gm16556 | -0.11282 | 0.166484 | 1 |
| Ttc14 | -0.11293 | 0.575701 | 1 |
| Peg13 | -0.11293 | 0.006626 | 1 |

| | | | |
|---|---|---|---|
| Zgpat | -0.11297 | 0.922726 | 1 |
| Hgs | -0.11298 | 0.423559 | 1 |
| Cbx3 | -0.113 | 0.004772 | 1 |
| Ifi207 | -0.11306 | 0.664153 | 1 |
| Slc7a1 | -0.11319 | 0.700544 | 1 |
| Dhx16 | -0.1132 | 0.690915 | 1 |
| Capns1 | -0.11339 | 0.90522 | 1 |
| Smndc1 | -0.11368 | 0.385132 | 1 |
| Mms19 | -0.11373 | 0.820881 | 1 |
| Rnf20 | -0.11386 | 0.702343 | 1 |
| Kmt5a | -0.11388 | 0.330603 | 1 |
| Gsdmd | -0.11393 | 0.029004 | 1 |
| Lsm6 | -0.11399 | 0.728713 | 1 |
| Zfp607a | -0.11417 | 0.074401 | 1 |
| Pitpnc1 | -0.11423 | 0.965646 | 1 |
| Phkb | -0.11424 | 0.556916 | 1 |
| Ikbkg | -0.11429 | 0.475821 | 1 |
| Rmnd5b | -0.1143 | 0.92123 | 1 |
| Nek7 | -0.11435 | 0.454741 | 1 |
| Ogfod1 | -0.11435 | 0.108023 | 1 |
| Brms1 | -0.11456 | 0.865127 | 1 |
| 2610507B1 | -0.11463 | 0.781462 | 1 |
| Zc3h3 | -0.11464 | 0.618205 | 1 |
| Fbxl6 | -0.11473 | 0.51313 | 1 |
| Sf3a1 | -0.11474 | 0.56921 | 1 |
| Tm9sf2 | -0.11481 | 0.825026 | 1 |
| Zbtb22 | -0.11492 | 0.395711 | 1 |
| C1galt1c1 | -0.11492 | 0.905324 | 1 |
| Dcaf6 | -0.11512 | 0.63892 | 1 |
| Socs5 | -0.1152 | 0.087043 | 1 |
| Nbeal2 | -0.11521 | 0.579637 | 1 |
| BC005537 | -0.11524 | 0.322894 | 1 |
| Zfp26 | -0.11527 | 0.583971 | 1 |
| Zfp652 | -0.11531 | 0.24983 | 1 |
| Zfp524 | -0.11534 | 0.472869 | 1 |
| Ppp4r2 | -0.11548 | 0.868815 | 1 |
| Matr3 | -0.11549 | 0.64635 | 1 |
| Borcs6 | -0.1156 | 0.683691 | 1 |
| Nfya | -0.11563 | 0.80504 | 1 |
| Lrwd1 | -0.11587 | 0.604894 | 1 |
| Zc3h15 | -0.11588 | 0.65028 | 1 |
| Oaz2 | -0.11599 | 0.256077 | 1 |
| Crybg3 | -0.11616 | 0.077954 | 1 |
| Ppp6r3 | -0.11625 | 0.840732 | 1 |
| Mapkapk3 | -0.11627 | 0.985028 | 1 |
| Rrp36 | -0.11644 | 0.484022 | 1 |
| Zfp335 | -0.11649 | 0.825339 | 1 |
| Ythdc2 | -0.11658 | 0.767727 | 1 |

| | | | |
|---|---|---|---|
| Lrrc20 | -0.1168 | 0.119413 | 1 |
| Chkb | -0.11688 | 0.731371 | 1 |
| Appl1 | -0.11692 | 0.698785 | 1 |
| Itgb2 | -0.11729 | 0.037758 | 1 |
| Ndor1 | -0.11737 | 0.721828 | 1 |
| Wdr26 | -0.11743 | 0.556547 | 1 |
| Ralgapb | -0.11744 | 0.117935 | 1 |
| Ttc37 | -0.1175 | 0.996556 | 1 |
| Slc9a8 | -0.11759 | 0.271431 | 1 |
| Specc1l | -0.1176 | 0.733056 | 1 |
| Serf2 | -0.11764 | 0.005786 | 1 |
| Hira | -0.11777 | 0.147609 | 1 |
| Ilrun | -0.11781 | 0.720097 | 1 |
| Ints3 | -0.11797 | 0.93979 | 1 |
| Scaf1 | -0.11802 | 0.51552 | 1 |
| Adora2a | -0.11808 | 0.030184 | 1 |
| 1600012H( | -0.11821 | 0.772744 | 1 |
| Morc3 | -0.11826 | 0.761887 | 1 |
| Sap25 | -0.11842 | 0.590965 | 1 |
| Ltn1 | -0.11849 | 0.301954 | 1 |
| Rc3h2 | -0.11851 | 0.415845 | 1 |
| Ralbp1 | -0.11853 | 0.535379 | 1 |
| Ubqln2 | -0.11854 | 0.574869 | 1 |
| Prr5l | -0.11883 | 0.741455 | 1 |
| Creg1 | -0.11891 | 0.770663 | 1 |
| Taf5 | -0.11892 | 0.053051 | 1 |
| Prpf6 | -0.11899 | 0.756094 | 1 |
| Cwf19l2 | -0.11911 | 0.707688 | 1 |
| Pigs | -0.11938 | 0.320136 | 1 |
| Ppp1r10 | -0.1194 | 0.494476 | 1 |
| Supt20 | -0.11949 | 0.961872 | 1 |
| Clk4 | -0.11949 | 0.948496 | 1 |
| Agl | -0.11951 | 0.767195 | 1 |
| Yipf3 | -0.1196 | 0.861629 | 1 |
| Arfrp1 | -0.11962 | 0.350572 | 1 |
| Elovl5 | -0.11966 | 0.458064 | 1 |
| Gga3 | -0.11976 | 0.505818 | 1 |
| Os9 | -0.11987 | 0.711799 | 1 |
| Pitpnm1 | -0.1199 | 0.987485 | 1 |
| Fbxl4 | -0.11992 | 0.214894 | 1 |
| Fam168b | -0.11996 | 0.813564 | 1 |
| Bid | -0.12 | 0.467294 | 1 |
| Rela | -0.12003 | 0.175917 | 1 |
| Haus8 | -0.12034 | 0.941322 | 1 |
| Nlrp1b | -0.12035 | 0.993584 | 1 |
| Tanc2 | -0.12035 | 0.620251 | 1 |
| Jak1 | -0.12042 | 0.674981 | 1 |
| Ankrd11 | -0.12053 | 0.324852 | 1 |

| | | | |
|---|---|---|---|
| Ctr9 | -0.12055 | 0.344192 | 1 |
| Azin1 | -0.12059 | 0.500684 | 1 |
| Ctnna1 | -0.12063 | 0.303477 | 1 |
| Tmem168 | -0.12066 | 0.12209 | 1 |
| Emc2 | -0.12074 | 0.956624 | 1 |
| Tmtc2 | -0.12079 | 0.037347 | 1 |
| Insr | -0.12086 | 0.482952 | 1 |
| Pds5b | -0.12089 | 0.594231 | 1 |
| Dclre1b | -0.12089 | 0.146845 | 1 |
| Tial1 | -0.1209 | 0.865472 | 1 |
| Zfp398 | -0.12102 | 0.537301 | 1 |
| Gucd1 | -0.12105 | 0.742755 | 1 |
| Tti1 | -0.12105 | 0.921101 | 1 |
| Trappc1 | -0.12105 | 0.825696 | 1 |
| Asxl2 | -0.12107 | 0.838939 | 1 |
| Commd4 | -0.12116 | 0.705443 | 1 |
| Iqsec2 | -0.12122 | 0.106765 | 1 |
| Atmin | -0.12125 | 0.795281 | 1 |
| Cirbp | -0.12126 | 0.714029 | 1 |
| Ppp1r18 | -0.1213 | 0.099606 | 1 |
| Mccc2 | -0.1214 | 0.122096 | 1 |
| Dcp1a | -0.12147 | 0.694674 | 1 |
| Vrk1 | -0.12148 | 0.306569 | 1 |
| Ddx58 | -0.12149 | 0.658902 | 1 |
| Luc7l3 | -0.1216 | 0.292653 | 1 |
| Gxylt1 | -0.12161 | 0.227408 | 1 |
| Zmiz2 | -0.12178 | 0.80226 | 1 |
| Dcaf7 | -0.12184 | 0.587452 | 1 |
| Prickle3 | -0.12191 | 0.184935 | 1 |
| Nbas | -0.12191 | 0.146843 | 1 |
| Tia1 | -0.12196 | 0.318893 | 1 |
| Cd177 | -0.12201 | 0.21166 | 1 |
| Smarca5 | -0.12208 | 0.982391 | 1 |
| Rbm42 | -0.12211 | 0.3897 | 1 |
| Umad1 | -0.12216 | 0.970622 | 1 |
| Rgs14 | -0.12219 | 0.79619 | 1 |
| Dhx15 | -0.12227 | 0.841864 | 1 |
| 4933412E1 | -0.12229 | 0.3656 | 1 |
| Gpr65 | -0.1224 | 0.558077 | 1 |
| Map3k15 | -0.12253 | 0.995959 | 1 |
| Sppl3 | -0.12255 | 0.880183 | 1 |
| Smad2 | -0.12273 | 0.617019 | 1 |
| Rab3gap1 | -0.12288 | 0.579529 | 1 |
| Hdac5 | -0.12288 | 0.181879 | 1 |
| Mid1ip1 | -0.12305 | 0.966728 | 1 |
| Gm49797 | -0.12312 | 0.583702 | 1 |
| Ormdl1 | -0.12315 | 0.406612 | 1 |
| Psd4 | -0.12318 | 0.821631 | 1 |

| | | | |
|---|---|---|---|
| Chd6 | -0.12324 | 0.995455 | 1 |
| Gmeb2 | -0.12325 | 0.644593 | 1 |
| Slu7 | -0.12326 | 0.635074 | 1 |
| Pdlim2 | -0.12342 | 0.759609 | 1 |
| Eif3f | -0.12345 | 0.135758 | 1 |
| Kat7 | -0.12352 | 0.516919 | 1 |
| Oas1a | -0.12354 | 0.641934 | 1 |
| Cab39 | -0.12362 | 0.444963 | 1 |
| Zfp871 | -0.12362 | 0.939293 | 1 |
| Ddx3y | -0.12394 | 0.751011 | 1 |
| Wapl | -0.12407 | 0.9311 | 1 |
| Ltc4s | -0.12408 | 0.087915 | 1 |
| Zfp942 | -0.1241 | 0.996782 | 1 |
| Btbd7 | -0.1241 | 0.539707 | 1 |
| Zc3h4 | -0.12426 | 0.845291 | 1 |
| Susd6 | -0.12434 | 0.382317 | 1 |
| Tyw1 | -0.12435 | 0.503199 | 1 |
| Rgs18 | -0.12436 | 0.002335 | 1 |
| Mrpl1 | -0.12442 | 0.520907 | 1 |
| Ric8b | -0.12443 | 0.851154 | 1 |
| Cnr2 | -0.12448 | 0.803289 | 1 |
| Uhrf2 | -0.1245 | 0.77274 | 1 |
| Mtdh | -0.12457 | 0.10213 | 1 |
| Rprd2 | -0.1246 | 0.991432 | 1 |
| Zfr | -0.12462 | 0.551498 | 1 |
| Pkn2 | -0.12465 | 0.583443 | 1 |
| Nsmce4a | -0.12466 | 0.901333 | 1 |
| Dusp11 | -0.12467 | 0.937296 | 1 |
| Agpat3 | -0.1247 | 0.473156 | 1 |
| Lrp1 | -0.12471 | 0.353872 | 1 |
| Vcpip1 | -0.12481 | 0.155566 | 1 |
| Shld1 | -0.12488 | 0.233776 | 1 |
| Tcof1 | -0.12489 | 0.425783 | 1 |
| Zfand3 | -0.12501 | 0.737937 | 1 |
| Cpsf3 | -0.12517 | 0.424882 | 1 |
| Top2b | -0.12517 | 0.265551 | 1 |
| Tfam | -0.12525 | 0.939138 | 1 |
| Ppm1d | -0.12539 | 0.259043 | 1 |
| Nt5c2 | -0.12539 | 0.663473 | 1 |
| Arl6ip6 | -0.12564 | 0.404472 | 1 |
| Arid2 | -0.12565 | 0.184336 | 1 |
| Setdb2 | -0.12577 | 0.648121 | 1 |
| Ppp4r3a | -0.1259 | 0.723622 | 1 |
| Ramp1 | -0.12606 | 0.587799 | 1 |
| Cramp1l | -0.12608 | 0.985528 | 1 |
| Zfp511 | -0.12613 | 0.216808 | 1 |
| Swt1 | -0.12614 | 0.204956 | 1 |
| Mettl3 | -0.12617 | 0.74408 | 1 |

| | | | |
|---|---|---|---|
| Dnajc1 | -0.12632 | 0.611364 | 1 |
| Fanca | -0.12635 | 0.920008 | 1 |
| Rpl35a | -0.12644 | 0.034089 | 1 |
| Copb1 | -0.12646 | 0.51501 | 1 |
| Inip | -0.1265 | 0.27804 | 1 |
| Tmem179b | -0.12693 | 0.316504 | 1 |
| Slbp | -0.12703 | 0.949841 | 1 |
| Tmem243 | -0.12703 | 0.357626 | 1 |
| Usp48 | -0.1271 | 0.330135 | 1 |
| Kansl3 | -0.12721 | 0.483531 | 1 |
| Nap1l1 | -0.12744 | 0.116087 | 1 |
| Ddx28 | -0.12747 | 0.624484 | 1 |
| Zfyve26 | -0.1275 | 0.93903 | 1 |
| Slc38a1 | -0.12756 | 0.792517 | 1 |
| Srrm2 | -0.12759 | 0.283701 | 1 |
| Sharpin | -0.12781 | 0.743353 | 1 |
| Slc22a15 | -0.12791 | 0.286298 | 1 |
| Ticam2 | -0.12791 | 0.821806 | 1 |
| Elac1 | -0.12793 | 0.142419 | 1 |
| 1810026B( | -0.12804 | 0.665024 | 1 |
| Uhrf1bp1l | -0.12808 | 0.804955 | 1 |
| Myd88 | -0.1281 | 0.50573 | 1 |
| Nsrp1 | -0.12811 | 0.891393 | 1 |
| Pan3 | -0.12814 | 0.179365 | 1 |
| Mocs1 | -0.12818 | 0.248498 | 1 |
| Kpnb1 | -0.12819 | 0.375758 | 1 |
| Ube2e3 | -0.12835 | 0.91294 | 1 |
| Tprgl | -0.12842 | 0.675298 | 1 |
| Gtf3c1 | -0.12843 | 0.218348 | 1 |
| Gbf1 | -0.12847 | 0.890599 | 1 |
| Brpf1 | -0.12851 | 0.847691 | 1 |
| 2310033P( | -0.12857 | 0.803376 | 1 |
| Nlrx1 | -0.1287 | 0.129929 | 1 |
| Prdm10 | -0.12893 | 0.152995 | 1 |
| Nfic | -0.12897 | 0.449344 | 1 |
| Smg6 | -0.12898 | 0.713438 | 1 |
| Smc1a | -0.12902 | 0.44233 | 1 |
| Mdp1 | -0.12926 | 0.265401 | 1 |
| Dhx8 | -0.1294 | 0.983486 | 1 |
| Golm1 | -0.12943 | 0.259215 | 1 |
| Thoc2 | -0.12947 | 0.586084 | 1 |
| Fbxw2 | -0.12954 | 0.680997 | 1 |
| Spast | -0.12957 | 0.421672 | 1 |
| Micu2 | -0.12962 | 0.585306 | 1 |
| Mri1 | -0.12965 | 0.847938 | 1 |
| Plbd1 | -0.12968 | 0.510048 | 1 |
| Oip5os1 | -0.12974 | 0.358854 | 1 |
| Parg | -0.12989 | 0.614021 | 1 |

| | | | |
|---|---|---|---|
| Sympk | -0.13007 | 0.262851 | 1 |
| Bud31 | -0.13009 | 0.831681 | 1 |
| Araf | -0.13012 | 0.919325 | 1 |
| Midn | -0.13028 | 0.455971 | 1 |
| Mon2 | -0.1305 | 0.624092 | 1 |
| Hmgcl | -0.13063 | 0.289404 | 1 |
| Pcf11 | -0.13066 | 0.623451 | 1 |
| Rnf215 | -0.13074 | 0.116849 | 1 |
| Snhg8 | -0.13082 | 0.510729 | 1 |
| Tlr8 | -0.13084 | 0.767876 | 1 |
| Serpinb8 | -0.13097 | 0.124253 | 1 |
| Nsd1 | -0.13112 | 0.312127 | 1 |
| Kmt2b | -0.13115 | 0.472627 | 1 |
| Evi5 | -0.13116 | 0.951325 | 1 |
| Afdn | -0.13122 | 0.860329 | 1 |
| Dph5 | -0.13125 | 0.081421 | 1 |
| Dgkz | -0.13131 | 0.562517 | 1 |
| Emd | -0.1314 | 0.527053 | 1 |
| Plxdc1 | -0.13158 | 0.024733 | 1 |
| Suz12 | -0.1316 | 0.47668 | 1 |
| Tpk1 | -0.1317 | 0.139379 | 1 |
| Slc25a28 | -0.13174 | 0.995728 | 1 |
| Cpt1a | -0.13187 | 0.892293 | 1 |
| Rab33b | -0.1319 | 0.189412 | 1 |
| Cab39l | -0.13206 | 0.82257 | 1 |
| Nedd4l | -0.13213 | 0.112799 | 1 |
| Shoc2 | -0.13223 | 0.932453 | 1 |
| Ube4a | -0.13226 | 0.483537 | 1 |
| Chfr | -0.13233 | 0.149251 | 1 |
| Prkag2 | -0.13235 | 0.156034 | 1 |
| Arfgef1 | -0.13238 | 0.585391 | 1 |
| Spag9 | -0.13242 | 0.713213 | 1 |
| Wdr20 | -0.13249 | 0.363828 | 1 |
| Klf10 | -0.13273 | 0.403386 | 1 |
| Rufy1 | -0.13275 | 0.991651 | 1 |
| Ppp6r1 | -0.13289 | 0.82321 | 1 |
| Prpf8 | -0.13324 | 0.412776 | 1 |
| Paxbp1 | -0.13325 | 0.941324 | 1 |
| Rsu1 | -0.13327 | 0.418185 | 1 |
| Zmym2 | -0.13343 | 0.457401 | 1 |
| Mbd5 | -0.13345 | 0.963663 | 1 |
| Cfh | -0.13347 | 0.121401 | 1 |
| 2610035D1 | -0.13349 | 0.00917 | 1 |
| Dynlt1b | -0.13355 | 0.097288 | 1 |
| Pltp | -0.13359 | 0.010503 | 1 |
| Hnrnpr | -0.13373 | 0.706815 | 1 |
| Setx | -0.13373 | 0.94868 | 1 |
| Gps1 | -0.1341 | 0.802808 | 1 |

| | | | |
|---|---|---|---|
| Ppm1a | -0.13415 | 0.398359 | 1 |
| Fyttd1 | -0.13433 | 0.342375 | 1 |
| Lmbrd1 | -0.13434 | 0.919759 | 1 |
| Mcm9 | -0.13441 | 0.487327 | 1 |
| Arntl | -0.13442 | 0.126673 | 1 |
| Apc | -0.13453 | 0.797955 | 1 |
| Atg4a | -0.13454 | 0.50646 | 1 |
| Glyr1 | -0.13457 | 0.42953 | 1 |
| Spop | -0.13479 | 0.22207 | 1 |
| Exoc3 | -0.13483 | 0.98273 | 1 |
| Ehd4 | -0.13512 | 0.399946 | 1 |
| Phf1 | -0.13514 | 0.206115 | 1 |
| Fam98c | -0.13534 | 0.841798 | 1 |
| Supt5 | -0.13537 | 0.847056 | 1 |
| Pqlc2 | -0.1354 | 0.58898 | 1 |
| Gls | -0.13542 | 0.755836 | 1 |
| Med7 | -0.13554 | 0.53171 | 1 |
| Filip1l | -0.13555 | 4.69E-07 | 0.015152 |
| Rps27a | -0.13573 | 0.003023 | 1 |
| Ppp4c | -0.13577 | 0.093647 | 1 |
| Klhl28 | -0.13585 | 0.394033 | 1 |
| Rab27a | -0.13628 | 0.339757 | 1 |
| Taf13 | -0.1363 | 0.923262 | 1 |
| Hgf | -0.13632 | 0.897355 | 1 |
| Vav1 | -0.13637 | 0.523897 | 1 |
| Fam117b | -0.13655 | 0.212048 | 1 |
| Nemp2 | -0.13659 | 0.004998 | 1 |
| Vps41 | -0.13663 | 0.068227 | 1 |
| Tmem50a | -0.13667 | 0.022282 | 1 |
| Acyp1 | -0.13697 | 0.102766 | 1 |
| Immp2l | -0.137 | 0.64676 | 1 |
| Ifnar1 | -0.1371 | 0.694759 | 1 |
| Cnpy3 | -0.13714 | 0.443604 | 1 |
| Rnpc3 | -0.13726 | 0.891012 | 1 |
| Lgmn | -0.1374 | 0.165685 | 1 |
| Jtb | -0.13743 | 0.528152 | 1 |
| Arpc5l | -0.13747 | 0.422863 | 1 |
| Ubr1 | -0.1376 | 0.740135 | 1 |
| Osgin2 | -0.13771 | 0.038478 | 1 |
| Nek9 | -0.13793 | 0.682409 | 1 |
| Tomm34 | -0.138 | 0.473958 | 1 |
| Casp2 | -0.13809 | 0.299493 | 1 |
| Aagab | -0.13816 | 0.599401 | 1 |
| Zfyve27 | -0.13833 | 0.55319 | 1 |
| Gng5 | -0.13847 | 0.031082 | 1 |
| Akap8l | -0.13856 | 0.732054 | 1 |
| Med30 | -0.13859 | 0.792739 | 1 |
| Map3k2 | -0.13863 | 0.328124 | 1 |

| | | | |
|---|---|---|---|
| Nsun6 | -0.13867 | 0.838018 | 1 |
| AI839979 | -0.13889 | 0.003606 | 1 |
| Gm35154 | -0.13901 | 0.120336 | 1 |
| Nrbf2 | -0.13913 | 0.912228 | 1 |
| Eps15l1 | -0.13915 | 0.663047 | 1 |
| Atrx | -0.13916 | 0.161494 | 1 |
| Prpf39 | -0.13917 | 0.978885 | 1 |
| Cep85l | -0.13924 | 0.191859 | 1 |
| 3300002I0 | -0.13925 | 0.084436 | 1 |
| March1 | -0.13928 | 0.076985 | 1 |
| Hsd17b10 | -0.13934 | 0.271047 | 1 |
| Chm | -0.13958 | 0.842068 | 1 |
| Osbpl2 | -0.13966 | 0.866232 | 1 |
| Dapp1 | -0.13977 | 0.135715 | 1 |
| Naa35 | -0.13978 | 0.739214 | 1 |
| Dyrk1a | -0.13979 | 0.314473 | 1 |
| Stn1 | -0.13991 | 0.272 | 1 |
| Sik1 | -0.13992 | 0.900801 | 1 |
| Nhlrc2 | -0.13995 | 0.739902 | 1 |
| Paip1 | -0.14018 | 0.77632 | 1 |
| Thap3 | -0.14022 | 0.767646 | 1 |
| Dennd1c | -0.14023 | 0.941056 | 1 |
| Mtss1 | -0.14024 | 0.610375 | 1 |
| Ubl3 | -0.14025 | 0.350448 | 1 |
| Wdr37 | -0.14027 | 0.238645 | 1 |
| Gfm2 | -0.14036 | 0.152099 | 1 |
| Zfp84 | -0.14045 | 0.115943 | 1 |
| Zfp120 | -0.14046 | 0.274738 | 1 |
| Mphosph8 | -0.14066 | 0.69915 | 1 |
| Tmem127 | -0.14076 | 0.493606 | 1 |
| Rbbp5 | -0.14098 | 0.220699 | 1 |
| Bfar | -0.14134 | 0.464588 | 1 |
| Ubxn2a | -0.14139 | 0.337893 | 1 |
| Plekhb2 | -0.14149 | 0.759446 | 1 |
| Anxa1 | -0.14166 | 0.870984 | 1 |
| Rbks | -0.1417 | 0.606763 | 1 |
| Amz1 | -0.14174 | 0.036301 | 1 |
| Ccnt1 | -0.1418 | 0.48096 | 1 |
| Pigx | -0.14187 | 0.555944 | 1 |
| Cops4 | -0.14214 | 0.801857 | 1 |
| Ppp2r5e | -0.14225 | 0.515785 | 1 |
| Gripap1 | -0.14229 | 0.050792 | 1 |
| Ube2b | -0.1423 | 0.400184 | 1 |
| Ccdc61 | -0.14231 | 0.037945 | 1 |
| Oga | -0.14234 | 0.766565 | 1 |
| Clasp1 | -0.14244 | 0.63687 | 1 |
| Nfkbib | -0.14247 | 0.455645 | 1 |
| Tmco1 | -0.14256 | 0.602939 | 1 |

| | | | |
|---|---|---|---|
| Ankrd12 | -0.14265 | 0.589959 | 1 |
| Mpc2 | -0.14271 | 0.366969 | 1 |
| Gid8 | -0.14284 | 0.988609 | 1 |
| Bcor | -0.14296 | 0.857583 | 1 |
| Stau1 | -0.14296 | 0.928215 | 1 |
| Hnrnpl | -0.14298 | 0.333511 | 1 |
| Snrk | -0.14302 | 0.354652 | 1 |
| Rbm10 | -0.14332 | 0.220599 | 1 |
| Sema4b | -0.14333 | 0.336996 | 1 |
| Trappc8 | -0.14346 | 0.786993 | 1 |
| Upf2 | -0.14356 | 0.305036 | 1 |
| Zbtb16 | -0.14359 | 0.008585 | 1 |
| Grap | -0.14371 | 0.168192 | 1 |
| Parp4 | -0.14371 | 0.426906 | 1 |
| Tbc1d8b | -0.14392 | 0.038493 | 1 |
| 2610037D( | -0.14408 | 0.381569 | 1 |
| H2-T23 | -0.14417 | 0.463282 | 1 |
| Safb2 | -0.14419 | 0.78758 | 1 |
| 6030458C1 | -0.1442 | 0.483678 | 1 |
| Fchsd2 | -0.14429 | 0.970312 | 1 |
| Nsmce2 | -0.14435 | 0.895731 | 1 |
| Mthfd2 | -0.1445 | 0.503701 | 1 |
| Vamp1 | -0.1445 | 0.109262 | 1 |
| Gpr35 | -0.14461 | 0.083775 | 1 |
| Lipe | -0.14463 | 0.127961 | 1 |
| Zmat5 | -0.14466 | 0.508399 | 1 |
| Jup | -0.14467 | 0.00386 | 1 |
| Ubtf | -0.14474 | 0.128524 | 1 |
| Txlna | -0.14482 | 0.728458 | 1 |
| Ccpg1 | -0.14483 | 0.218486 | 1 |
| Sec24c | -0.14484 | 0.984107 | 1 |
| Epb41l2 | -0.14492 | 0.517967 | 1 |
| Usp33 | -0.145 | 0.662518 | 1 |
| Sbds | -0.14503 | 0.246509 | 1 |
| E130307A1 | -0.14504 | 0.307946 | 1 |
| Arpc2 | -0.14516 | 0.011418 | 1 |
| Cyp51 | -0.14535 | 0.301886 | 1 |
| Lta4h | -0.14548 | 0.489162 | 1 |
| Sh3glb1 | -0.14561 | 0.209721 | 1 |
| Fbrsl1 | -0.14586 | 0.585217 | 1 |
| Zfp513 | -0.14592 | 0.340346 | 1 |
| Patz1 | -0.14601 | 0.783387 | 1 |
| Amd1 | -0.14612 | 0.719247 | 1 |
| Eed | -0.14652 | 0.621617 | 1 |
| Camkmt | -0.14654 | 0.33706 | 1 |
| Zfp971 | -0.14654 | 0.872487 | 1 |
| Rarg | -0.14664 | 0.027755 | 1 |
| Cnst | -0.1467 | 0.059609 | 1 |

| | | | |
|---|---|---|---|
| Zdhhc17 | -0.14679 | 0.663396 | 1 |
| Rnf111 | -0.14683 | 0.331096 | 1 |
| Pdpk1 | -0.14688 | 0.803154 | 1 |
| Slain2 | -0.147 | 0.4669 | 1 |
| Rpl10-ps3 | -0.147 | 0.734833 | 1 |
| Kdm4c | -0.14716 | 0.25485 | 1 |
| Bicral | -0.14726 | 0.31099 | 1 |
| Brip1os | -0.14728 | 0.645848 | 1 |
| Vta1 | -0.14735 | 0.346644 | 1 |
| Oas3 | -0.14745 | 0.896788 | 1 |
| Cse1l | -0.14747 | 0.68171 | 1 |
| Atp2b1 | -0.14751 | 0.207764 | 1 |
| Tmcc3 | -0.14765 | 0.003294 | 1 |
| Hip1r | -0.14778 | 0.108013 | 1 |
| Gng10 | -0.14779 | 0.332974 | 1 |
| Slc7a7 | -0.14785 | 0.644613 | 1 |
| Brwd3 | -0.14811 | 0.506378 | 1 |
| Tkt | -0.14814 | 0.060563 | 1 |
| D230025D | -0.14819 | 0.513248 | 1 |
| Lfng | -0.14828 | 0.580364 | 1 |
| Zc3h13 | -0.14836 | 0.703025 | 1 |
| Cep135 | -0.1486 | 0.760781 | 1 |
| Rbbp6 | -0.14863 | 0.859184 | 1 |
| Fbxo3 | -0.14868 | 0.481391 | 1 |
| Tnfrsf1a | -0.14879 | 0.374176 | 1 |
| Tbc1d1 | -0.14906 | 0.362736 | 1 |
| P2ry6 | -0.1491 | 0.529071 | 1 |
| Ube2h | -0.14919 | 0.881627 | 1 |
| St6galnac3 | -0.14925 | 0.001917 | 1 |
| Pex13 | -0.14931 | 0.296908 | 1 |
| Rab31 | -0.14932 | 0.018996 | 1 |
| Bmyc | -0.14935 | 0.053721 | 1 |
| Mplkip | -0.14948 | 0.231816 | 1 |
| Pnrc2 | -0.14971 | 0.047225 | 1 |
| Eif3j1 | -0.14975 | 0.859068 | 1 |
| Xrn2 | -0.14983 | 0.121309 | 1 |
| Rtf1 | -0.14986 | 0.866355 | 1 |
| Vamp3 | -0.15001 | 0.787173 | 1 |
| Zfp790 | -0.15008 | 0.01689 | 1 |
| Sec62 | -0.15021 | 0.082571 | 1 |
| Rhbdf2 | -0.15022 | 0.706299 | 1 |
| Mppe1 | -0.15033 | 0.51674 | 1 |
| Ensa | -0.1504 | 0.553009 | 1 |
| Znrd1 | -0.15043 | 0.191717 | 1 |
| Ammecr1 | -0.15068 | 0.193918 | 1 |
| Tsen34 | -0.15076 | 0.991704 | 1 |
| Nfx1 | -0.15088 | 0.593123 | 1 |
| Kctd10 | -0.15102 | 0.247489 | 1 |

| | | | |
|---|---|---|---|
| Atxn2l | -0.15102 | 0.211026 | 1 |
| Samhd1 | -0.1511 | 0.055983 | 1 |
| Dram2 | -0.15129 | 0.63293 | 1 |
| Zmynd11 | -0.1513 | 0.811225 | 1 |
| Tbc1d9b | -0.15134 | 0.1442 | 1 |
| Adgrg5 | -0.1514 | 0.367556 | 1 |
| Ppp1r2 | -0.15159 | 0.668905 | 1 |
| Fgfr1op2 | -0.1516 | 0.189553 | 1 |
| Slc36a3os | -0.15171 | 0.11938 | 1 |
| Arid4b | -0.15199 | 0.570193 | 1 |
| Ppp1r15a | -0.15201 | 0.838422 | 1 |
| Brd1 | -0.15226 | 0.914725 | 1 |
| Chmp2b | -0.15246 | 0.334351 | 1 |
| Gm20658 | -0.15246 | 0.853155 | 1 |
| Eipr1 | -0.15254 | 0.368221 | 1 |
| Cnot3 | -0.15263 | 0.478817 | 1 |
| 2810013P( | -0.15265 | 0.643811 | 1 |
| Arhgap25 | -0.15268 | 0.845904 | 1 |
| Nsf | -0.15273 | 0.356806 | 1 |
| Wac | -0.15284 | 0.313299 | 1 |
| Gm39556 | -0.15316 | 0.207016 | 1 |
| Glg1 | -0.15333 | 0.792665 | 1 |
| Golga1 | -0.15341 | 0.199479 | 1 |
| Adap1 | -0.15344 | 0.570134 | 1 |
| Kcnj2 | -0.15385 | 1.43E-06 | 0.046011 |
| Inpp5f | -0.15404 | 0.844133 | 1 |
| Nck2 | -0.15413 | 0.022278 | 1 |
| Ppip5k2 | -0.15431 | 0.602624 | 1 |
| Rnf2 | -0.15434 | 0.040106 | 1 |
| Rnf166 | -0.1544 | 0.955923 | 1 |
| Sgpp1 | -0.15443 | 0.286775 | 1 |
| Msmo1 | -0.1545 | 0.752004 | 1 |
| Socs4 | -0.15489 | 0.605454 | 1 |
| Zfp770 | -0.15495 | 0.001766 | 1 |
| Socs6 | -0.15503 | 0.815599 | 1 |
| Asb3 | -0.15508 | 0.485051 | 1 |
| Pbx3 | -0.15516 | 0.789516 | 1 |
| Cul3 | -0.15524 | 0.786796 | 1 |
| Shkbp1 | -0.15538 | 0.216295 | 1 |
| Bloc1s4 | -0.15546 | 0.9047 | 1 |
| Cdkn2d | -0.1555 | 0.269232 | 1 |
| Gm5431 | -0.15563 | 0.690758 | 1 |
| Nrbp1 | -0.15571 | 0.234364 | 1 |
| Il10rb | -0.15577 | 0.775127 | 1 |
| Cdc42se2 | -0.15578 | 0.184896 | 1 |
| Nlrp1a | -0.15598 | 0.738209 | 1 |
| Suco | -0.15608 | 0.739186 | 1 |
| Dnmt3a | -0.1561 | 0.979968 | 1 |

| | | | |
|---|---|---|---|
| Cd79b | -0.15622 | 0.000727 | 1 |
| Dcaf5 | -0.15625 | 0.550527 | 1 |
| Chd9 | -0.15638 | 0.173838 | 1 |
| Pdlim1 | -0.15646 | 0.051314 | 1 |
| Rcsd1 | -0.15653 | 0.739215 | 1 |
| Vbp1 | -0.15668 | 0.992209 | 1 |
| Med27 | -0.15671 | 0.396324 | 1 |
| Slc16a7 | -0.15671 | 0.069328 | 1 |
| Kmt2d | -0.15683 | 0.55034 | 1 |
| Cpeb4 | -0.15686 | 0.888424 | 1 |
| N4bp2l2 | -0.15688 | 0.284101 | 1 |
| Mfsd14a | -0.157 | 0.66932 | 1 |
| Tob2 | -0.1574 | 0.768688 | 1 |
| Gnpnat1 | -0.15764 | 0.067747 | 1 |
| Cr1l | -0.15802 | 0.488792 | 1 |
| R3hdm4 | -0.15817 | 0.116114 | 1 |
| Rprd1a | -0.15828 | 0.12102 | 1 |
| Cog5 | -0.15839 | 0.641669 | 1 |
| Sik2 | -0.15855 | 0.392045 | 1 |
| Zmynd8 | -0.15893 | 0.46899 | 1 |
| Vps16 | -0.15906 | 0.166182 | 1 |
| Fus | -0.15907 | 0.177135 | 1 |
| Gm47754 | -0.15932 | 0.111484 | 1 |
| Fbxo38 | -0.15934 | 0.47045 | 1 |
| Syne3 | -0.15939 | 0.017047 | 1 |
| Pea15a | -0.15947 | 0.118576 | 1 |
| Fbxl12 | -0.15956 | 0.857521 | 1 |
| Brwd1 | -0.15989 | 0.32404 | 1 |
| Zhx1 | -0.15989 | 0.089395 | 1 |
| Osbpl9 | -0.15993 | 0.317026 | 1 |
| Phka2 | -0.16 | 0.721696 | 1 |
| Spopl | -0.16013 | 0.943582 | 1 |
| Oard1 | -0.1602 | 0.814876 | 1 |
| Hgsnat | -0.16025 | 0.285419 | 1 |
| E230029C( | -0.16028 | 0.997133 | 1 |
| Pwwp3a | -0.16046 | 0.009391 | 1 |
| Nufip2 | -0.16053 | 0.313454 | 1 |
| Cox20 | -0.16071 | 0.363815 | 1 |
| Stat1 | -0.16122 | 0.04384 | 1 |
| Dna2 | -0.16143 | 0.563754 | 1 |
| Dynlt1f | -0.16172 | 0.187657 | 1 |
| Tmem30a | -0.16184 | 0.563955 | 1 |
| Dennd4b | -0.16197 | 0.334303 | 1 |
| Cep192 | -0.16206 | 0.591094 | 1 |
| Smcr8 | -0.16209 | 0.071849 | 1 |
| C1galt1 | -0.16219 | 0.817541 | 1 |
| Phlpp1 | -0.16225 | 0.171967 | 1 |
| Tmem230 | -0.16236 | 0.6198 | 1 |

| | | | |
|---|---|---|---|
| Anapc16 | -0.1624 | 0.539579 | 1 |
| Eif1b | -0.16242 | 0.199077 | 1 |
| Zrsr2 | -0.16253 | 0.4214 | 1 |
| Rapgef6 | -0.16258 | 0.474575 | 1 |
| Ahcyl2 | -0.16278 | 0.278999 | 1 |
| Ate1 | -0.16286 | 0.916439 | 1 |
| Gtf3c2 | -0.16298 | 0.491275 | 1 |
| Erich1 | -0.16307 | 0.096498 | 1 |
| Cyp4f13 | -0.1632 | 0.573173 | 1 |
| Nr1h2 | -0.16332 | 0.653937 | 1 |
| Gpank1 | -0.16336 | 0.281751 | 1 |
| Fip1l1 | -0.16339 | 0.289361 | 1 |
| Gatad2b | -0.1634 | 0.89761 | 1 |
| Kin | -0.16341 | 0.478804 | 1 |
| Pcsk7 | -0.16346 | 0.811637 | 1 |
| Ascc3 | -0.16352 | 0.430184 | 1 |
| Ngdn | -0.16359 | 0.532753 | 1 |
| Man1a2 | -0.1636 | 0.881642 | 1 |
| Pycrl | -0.16362 | 0.470586 | 1 |
| Dbnl | -0.16373 | 0.041216 | 1 |
| Pgap1 | -0.16395 | 0.64812 | 1 |
| Nemf | -0.16409 | 0.463642 | 1 |
| Eml3 | -0.16416 | 0.101645 | 1 |
| Cntrl | -0.16435 | 0.575574 | 1 |
| Gdpd3 | -0.16441 | 0.216914 | 1 |
| Nlk | -0.16445 | 0.88127 | 1 |
| Poldip3 | -0.1645 | 0.26547 | 1 |
| Wipf2 | -0.16454 | 0.529057 | 1 |
| Herc2 | -0.16506 | 0.440049 | 1 |
| Hnrnpul2 | -0.16507 | 0.087939 | 1 |
| Pqlc1 | -0.16519 | 0.979505 | 1 |
| Crybg1 | -0.16521 | 0.858345 | 1 |
| Ints6l | -0.16522 | 0.925254 | 1 |
| Lrp5 | -0.16525 | 0.388373 | 1 |
| Rhog | -0.16529 | 0.008047 | 1 |
| Ptp4a3 | -0.16537 | 0.6058 | 1 |
| Brd4 | -0.16537 | 0.360405 | 1 |
| Arpc4 | -0.16542 | 0.007592 | 1 |
| Mettl9 | -0.16544 | 0.29008 | 1 |
| Acss2 | -0.16575 | 0.136524 | 1 |
| Cyb5r4 | -0.16579 | 0.280148 | 1 |
| Ssr3 | -0.16585 | 0.184527 | 1 |
| H2-Ke6 | -0.16612 | 0.309583 | 1 |
| Pknox1 | -0.1662 | 0.088491 | 1 |
| Ddx23 | -0.16621 | 0.288384 | 1 |
| Prkn | -0.16621 | 0.152971 | 1 |
| Vsir | -0.16627 | 0.264213 | 1 |
| Chd3 | -0.16666 | 0.660884 | 1 |

| | | | |
|---|---|---|---|
| Arrdc2 | -0.16673 | 0.017537 | 1 |
| Ryr1 | -0.16703 | 0.302966 | 1 |
| Fbxo28 | -0.16706 | 0.305827 | 1 |
| Gm15564 | -0.16707 | 0.010761 | 1 |
| Rsrc2 | -0.16744 | 0.247545 | 1 |
| Gm7072 | -0.16747 | 0.213987 | 1 |
| Fmnl3 | -0.16749 | 0.481543 | 1 |
| Senp2 | -0.16754 | 0.408686 | 1 |
| Traf5 | -0.16759 | 0.495527 | 1 |
| Tspyl1 | -0.16759 | 0.352343 | 1 |
| Zfp182 | -0.16759 | 0.085279 | 1 |
| Tbc1d23 | -0.16778 | 0.310448 | 1 |
| Atp6v0e | -0.16789 | 0.017731 | 1 |
| Zfp467 | -0.16792 | 0.18052 | 1 |
| Pura | -0.16793 | 0.380913 | 1 |
| Myo9a | -0.16809 | 0.919368 | 1 |
| Washc4 | -0.16812 | 0.635741 | 1 |
| Srsf10 | -0.16814 | 0.502736 | 1 |
| Gm35769 | -0.16817 | 0.451706 | 1 |
| Chchd2 | -0.1682 | 0.011322 | 1 |
| Kcnip1 | -0.16859 | 0.033538 | 1 |
| Arhgap11a | -0.1686 | 0.347919 | 1 |
| Prkdc | -0.16866 | 0.728402 | 1 |
| Hnrnph1 | -0.16868 | 0.284403 | 1 |
| Cpsf7 | -0.16869 | 0.295818 | 1 |
| Parp11 | -0.1687 | 0.406153 | 1 |
| Ints6 | -0.16871 | 0.489912 | 1 |
| Plekhg3 | -0.16912 | 0.04566 | 1 |
| Stx18 | -0.16933 | 0.579333 | 1 |
| Icam2 | -0.16939 | 0.210913 | 1 |
| Phip | -0.16953 | 0.444448 | 1 |
| Pak2 | -0.16961 | 0.040562 | 1 |
| Lcor | -0.16966 | 0.856232 | 1 |
| Fam43a | -0.16973 | 0.00817 | 1 |
| Itgb5 | -0.16976 | 0.02505 | 1 |
| Fam53c | -0.16991 | 0.172633 | 1 |
| Pnn | -0.16994 | 0.189481 | 1 |
| Atg10 | -0.17003 | 0.324817 | 1 |
| Pcif1 | -0.17018 | 0.9714 | 1 |
| Lrch3 | -0.17019 | 0.71536 | 1 |
| Mtpap | -0.1702 | 0.777849 | 1 |
| Prdm2 | -0.17025 | 0.393076 | 1 |
| Ifih1 | -0.17038 | 0.631534 | 1 |
| Sec11c | -0.17041 | 0.182401 | 1 |
| Ganc | -0.17049 | 0.104533 | 1 |
| Bmpr2 | -0.17057 | 0.511853 | 1 |
| Arhgef2 | -0.17075 | 0.587767 | 1 |
| Rnf31 | -0.17077 | 0.879013 | 1 |

| | | | |
|---|---|---|---|
| Eif2a | -0.17104 | 0.300207 | 1 |
| Uspl1 | -0.17115 | 0.835246 | 1 |
| Tlr9 | -0.17134 | 0.1437 | 1 |
| Sf1 | -0.17143 | 0.190135 | 1 |
| Kcnab2 | -0.17144 | 0.911977 | 1 |
| Ing4 | -0.17167 | 0.58254 | 1 |
| Ankle2 | -0.1717 | 0.6603 | 1 |
| Cln8 | -0.17184 | 0.235191 | 1 |
| Abca3 | -0.17191 | 0.161156 | 1 |
| Gtf2a1 | -0.17212 | 0.529546 | 1 |
| Atp1a3 | -0.17214 | 0.210947 | 1 |
| N4bp1 | -0.17229 | 0.999515 | 1 |
| Sephs2 | -0.1724 | 0.228009 | 1 |
| Ythdc1 | -0.17242 | 0.973132 | 1 |
| Dlg1 | -0.1725 | 0.484066 | 1 |
| Nck1 | -0.17265 | 0.908535 | 1 |
| Gps2 | -0.17266 | 0.464743 | 1 |
| Tmem87a | -0.1729 | 0.84753 | 1 |
| Rasa2 | -0.17308 | 0.549 | 1 |
| Anxa11 | -0.1731 | 0.513393 | 1 |
| Spred2 | -0.17312 | 0.647509 | 1 |
| Ophn1 | -0.17321 | 0.243983 | 1 |
| Mob3a | -0.17325 | 0.453656 | 1 |
| Ermard | -0.17332 | 0.191961 | 1 |
| Ankrd13a | -0.17338 | 0.272381 | 1 |
| Fbxw11 | -0.17352 | 0.516154 | 1 |
| Zfp143 | -0.1736 | 0.263296 | 1 |
| Pfdn5 | -0.1736 | 0.004709 | 1 |
| Sin3a | -0.17403 | 0.86677 | 1 |
| Ube3a | -0.17417 | 0.503301 | 1 |
| Dguok | -0.17421 | 0.339588 | 1 |
| Ublcp1 | -0.17422 | 0.867332 | 1 |
| Abcb7 | -0.17427 | 0.004463 | 1 |
| Pyroxd1 | -0.17451 | 0.016054 | 1 |
| Kmt2a | -0.17453 | 0.077974 | 1 |
| Cox16 | -0.17467 | 0.712473 | 1 |
| Slfn5 | -0.1747 | 0.065917 | 1 |
| Dhrs7b | -0.17475 | 0.955405 | 1 |
| Pnkp | -0.17481 | 0.159797 | 1 |
| Sh3pxd2a | -0.17483 | 0.02471 | 1 |
| Kri1 | -0.17483 | 0.444449 | 1 |
| Rictor | -0.17486 | 0.300289 | 1 |
| Setd1b | -0.17495 | 0.445696 | 1 |
| Tram1 | -0.17504 | 0.132947 | 1 |
| 9930021J0 | -0.1751 | 0.139773 | 1 |
| Tinf2 | -0.17513 | 0.318628 | 1 |
| Gm9949 | -0.17533 | 0.000634 | 1 |
| Nxf1 | -0.17586 | 0.766411 | 1 |

| | | | |
|---|---|---|---|
| Hltf | -0.1759 | 0.263065 | 1 |
| Sqor | -0.17598 | 0.581884 | 1 |
| Svip | -0.17601 | 0.000723 | 1 |
| Pdxdc1 | -0.17603 | 0.820953 | 1 |
| Pde4dip | -0.17625 | 0.307863 | 1 |
| Pde4b | -0.17629 | 0.371397 | 1 |
| Strada | -0.17648 | 0.285601 | 1 |
| Polr2b | -0.17689 | 0.108354 | 1 |
| Sptan1 | -0.17696 | 0.227619 | 1 |
| Gpr155 | -0.17717 | 0.00405 | 1 |
| Psen1 | -0.17721 | 0.941481 | 1 |
| Ankra2 | -0.17722 | 0.849615 | 1 |
| Cdk12 | -0.17739 | 0.639239 | 1 |
| St6gal1 | -0.1777 | 0.073722 | 1 |
| P2ry13 | -0.17795 | 0.153438 | 1 |
| Hivep2 | -0.17805 | 0.160069 | 1 |
| Gigyf2 | -0.17812 | 0.309651 | 1 |
| Stab1 | -0.17836 | 3.23E-05 | 1 |
| Pgghg | -0.17843 | 0.133751 | 1 |
| Sp2 | -0.17847 | 0.233051 | 1 |
| Tspyl3 | -0.17896 | 0.003737 | 1 |
| Ipcef1 | -0.17923 | 0.506961 | 1 |
| Rftn1 | -0.17928 | 0.158034 | 1 |
| Usp34 | -0.17932 | 0.366147 | 1 |
| Clec2i | -0.17936 | 0.243954 | 1 |
| Tsc22d2 | -0.17942 | 0.262989 | 1 |
| Hmgxb3 | -0.17946 | 0.94215 | 1 |
| Nfkbid | -0.17947 | 0.760435 | 1 |
| Usf3 | -0.17955 | 0.05705 | 1 |
| R3hdm2 | -0.17955 | 0.689767 | 1 |
| Slc25a20 | -0.17985 | 0.564334 | 1 |
| Ubac2 | -0.17987 | 0.340245 | 1 |
| Nod2 | -0.18009 | 0.854803 | 1 |
| Gphn | -0.18031 | 0.002993 | 1 |
| Micu1 | -0.1804 | 0.548688 | 1 |
| Maf1 | -0.18069 | 0.276722 | 1 |
| Armc3 | -0.18078 | 0.000754 | 1 |
| Zfp950 | -0.18108 | 0.107761 | 1 |
| Sp3 | -0.18113 | 0.703251 | 1 |
| 2810004N[illegible] | -0.18117 | 0.9045 | 1 |
| Trim21 | -0.18145 | 0.621738 | 1 |
| Pik3cb | -0.18154 | 0.721563 | 1 |
| Smarcd2 | -0.18161 | 0.029645 | 1 |
| Eif4g2 | -0.18178 | 0.100433 | 1 |
| Rnf6 | -0.18203 | 0.49139 | 1 |
| Lsm14a | -0.1821 | 0.993047 | 1 |
| Zcrb1 | -0.18216 | 0.271341 | 1 |
| Med13 | -0.18219 | 0.183355 | 1 |

| | | | |
|---|---|---|---|
| Spic | -0.18233 | 0.031198 | 1 |
| Pi4kb | -0.18262 | 0.408741 | 1 |
| Stard8 | -0.18282 | 0.96482 | 1 |
| Cybc1 | -0.18295 | 0.095725 | 1 |
| Tmem71 | -0.18318 | 0.120439 | 1 |
| Slc8b1 | -0.18397 | 0.3656 | 1 |
| Sod1 | -0.18403 | 0.814153 | 1 |
| Pdha1 | -0.18405 | 0.529603 | 1 |
| Snx29 | -0.1844 | 0.824776 | 1 |
| Mpp1 | -0.18441 | 0.862176 | 1 |
| Ppfibp2 | -0.18455 | 0.000222 | 1 |
| Heca | -0.18465 | 0.766843 | 1 |
| Gm20559 | -0.18466 | 0.044366 | 1 |
| Mta2 | -0.1852 | 0.073536 | 1 |
| Lrba | -0.18554 | 0.059175 | 1 |
| Ndufaf7 | -0.1857 | 0.042737 | 1 |
| Nrf1 | -0.1858 | 0.497246 | 1 |
| Ncoa3 | -0.18585 | 0.033844 | 1 |
| Fbrs | -0.18587 | 0.294419 | 1 |
| Ldlrap1 | -0.18609 | 0.052875 | 1 |
| Mia3 | -0.18646 | 0.159931 | 1 |
| Fnta | -0.18697 | 0.500437 | 1 |
| Copg1 | -0.18719 | 0.577552 | 1 |
| Fam133b | -0.18733 | 0.091366 | 1 |
| Gon4l | -0.18748 | 0.433543 | 1 |
| Creb1 | -0.18771 | 0.740172 | 1 |
| Smg1 | -0.18776 | 0.1505 | 1 |
| Rcan1 | -0.18778 | 0.332234 | 1 |
| Pabpc1 | -0.18779 | 0.024401 | 1 |
| A530064D( | -0.18781 | 0.001378 | 1 |
| Ggct | -0.18783 | 0.428054 | 1 |
| Zfp263 | -0.18795 | 0.189612 | 1 |
| Ifi27l2a | -0.18809 | 0.267213 | 1 |
| Scaf4 | -0.1881 | 0.316019 | 1 |
| Fli1 | -0.18837 | 0.086136 | 1 |
| Senp1 | -0.18864 | 0.357893 | 1 |
| Tmpo | -0.18886 | 0.066925 | 1 |
| Aqr | -0.18888 | 0.670688 | 1 |
| Wls | -0.18911 | 0.299431 | 1 |
| Bod1l | -0.18918 | 0.178737 | 1 |
| Nup107 | -0.18949 | 0.040901 | 1 |
| Plpp1 | -0.18952 | 0.090488 | 1 |
| Arhgap27o | -0.18965 | 0.002784 | 1 |
| Paxx | -0.18973 | 0.038929 | 1 |
| Galnt10 | -0.18978 | 0.803988 | 1 |
| Mindy2 | -0.18992 | 0.67161 | 1 |
| Ralgps2 | -0.19004 | 0.194046 | 1 |
| Nrm | -0.19019 | 0.08704 | 1 |

| | | | |
|---|---|---|---|
| Baz2a | -0.19028 | 0.329683 | 1 |
| Tmsb10 | -0.19031 | 0.231647 | 1 |
| Phf12 | -0.19038 | 0.269571 | 1 |
| Adipor1 | -0.19084 | 0.10336 | 1 |
| S100a4 | -0.19096 | 0.030862 | 1 |
| Rock2 | -0.19097 | 0.1666 | 1 |
| Tmc8 | -0.19109 | 0.002849 | 1 |
| Gss | -0.19115 | 0.717232 | 1 |
| L3mbtl3 | -0.1912 | 0.61443 | 1 |
| Rab8a | -0.19133 | 0.07536 | 1 |
| Maml2 | -0.19141 | 0.618257 | 1 |
| 4833420G1 | -0.19144 | 0.620482 | 1 |
| Mefv | -0.19157 | 0.202611 | 1 |
| Tug1 | -0.19158 | 0.142415 | 1 |
| Lmbrd2 | -0.19158 | 0.957066 | 1 |
| Agps | -0.19169 | 0.887734 | 1 |
| Ubash3b | -0.19218 | 0.1432 | 1 |
| Shc1 | -0.19238 | 0.654589 | 1 |
| Cdk19 | -0.19238 | 0.891661 | 1 |
| Plxnc1 | -0.19248 | 0.051405 | 1 |
| Ahr | -0.1927 | 0.37029 | 1 |
| Phc3 | -0.19339 | 0.10159 | 1 |
| Cdc26 | -0.19366 | 0.344935 | 1 |
| Gfpt1 | -0.19391 | 0.377681 | 1 |
| Rassf3 | -0.19431 | 0.888994 | 1 |
| Herpud1 | -0.19439 | 0.518046 | 1 |
| Zfp62 | -0.19461 | 0.051899 | 1 |
| Rbm26 | -0.19472 | 0.115108 | 1 |
| Triap1 | -0.19507 | 0.144741 | 1 |
| AW554918 | -0.19512 | 0.588668 | 1 |
| Irf3 | -0.19514 | 0.437204 | 1 |
| Jrkl | -0.19548 | 0.048705 | 1 |
| Cdipt | -0.19551 | 0.935246 | 1 |
| Slc38a2 | -0.19553 | 0.400299 | 1 |
| Huwe1 | -0.19562 | 0.910801 | 1 |
| Bbc3 | -0.19572 | 0.054751 | 1 |
| Fcgr4 | -0.19584 | 0.298308 | 1 |
| Pdlim5 | -0.19589 | 0.114581 | 1 |
| Gm17767 | -0.19597 | 0.000588 | 1 |
| Lman2l | -0.19606 | 0.779699 | 1 |
| Serpinb6a | -0.19657 | 0.566135 | 1 |
| Scaf8 | -0.19657 | 0.270659 | 1 |
| Il27 | -0.19722 | 0.935334 | 1 |
| Serinc3 | -0.19724 | 0.127096 | 1 |
| Il18 | -0.19739 | 0.195384 | 1 |
| Dpp8 | -0.19739 | 0.205926 | 1 |
| Prrc2a | -0.19757 | 0.408772 | 1 |
| Osbpl8 | -0.19773 | 0.511688 | 1 |

| | | | |
|---|---|---|---|
| Pygb | -0.19816 | 0.13786 | 1 |
| Kdm5c | -0.19852 | 0.067272 | 1 |
| Sft2d1 | -0.19869 | 0.154044 | 1 |
| Usp8 | -0.19869 | 0.328166 | 1 |
| Arf6 | -0.19871 | 0.242897 | 1 |
| Anxa7 | -0.19902 | 0.885677 | 1 |
| Ogfrl1 | -0.19938 | 0.947813 | 1 |
| Selenot | -0.19948 | 0.050759 | 1 |
| Def6 | -0.19958 | 0.209798 | 1 |
| Purb | -0.19998 | 0.127505 | 1 |
| Slc29a1 | -0.20008 | 0.441867 | 1 |
| Cln3 | -0.201 | 0.242152 | 1 |
| 4930444A1 | -0.20107 | 0.001275 | 1 |
| Dock10 | -0.20112 | 0.288968 | 1 |
| Abcd2 | -0.20137 | 0.230341 | 1 |
| Mark2 | -0.20138 | 0.06006 | 1 |
| Gpr18 | -0.20151 | 0.501426 | 1 |
| Hnrnpd | -0.20156 | 0.017103 | 1 |
| Zup1 | -0.20163 | 0.13138 | 1 |
| Ube2r2 | -0.20168 | 0.518915 | 1 |
| Tle4 | -0.20183 | 0.528234 | 1 |
| Tspan5 | -0.20192 | 0.373452 | 1 |
| Kdm7a | -0.20216 | 0.051324 | 1 |
| Senp7 | -0.20227 | 0.326212 | 1 |
| BE692007 | -0.2023 | 0.32614 | 1 |
| Laptm4a | -0.20241 | 0.04839 | 1 |
| Ago2 | -0.20243 | 0.307457 | 1 |
| Irf9 | -0.20243 | 0.01532 | 1 |
| Amn1 | -0.20255 | 0.846808 | 1 |
| Crebl2 | -0.20259 | 4.8E-05 | 1 |
| Hps5 | -0.2027 | 0.644044 | 1 |
| Tut7 | -0.2028 | 0.028827 | 1 |
| Abi1 | -0.20282 | 0.002839 | 1 |
| Slmap | -0.20284 | 0.523972 | 1 |
| Trp53 | -0.20292 | 0.275213 | 1 |
| Gnb1 | -0.203 | 0.231757 | 1 |
| Tor1aip1 | -0.203 | 0.011778 | 1 |
| Sppl2a | -0.20311 | 0.096908 | 1 |
| Wdr33 | -0.20324 | 0.068367 | 1 |
| N4bp2l1 | -0.20337 | 0.210997 | 1 |
| Nudcd3 | -0.20347 | 0.789642 | 1 |
| Baz1b | -0.20356 | 0.160209 | 1 |
| Ube2w | -0.20363 | 0.155577 | 1 |
| Spen | -0.20364 | 0.357487 | 1 |
| Phf8 | -0.20373 | 0.083924 | 1 |
| Ncoa7 | -0.20381 | 0.061331 | 1 |
| Nlrc4 | -0.2039 | 0.008841 | 1 |
| Rbm6 | -0.20396 | 0.219986 | 1 |

| | | | |
|---|---|---|---|
| Apaf1 | -0.20397 | 0.52258 | 1 |
| Prkx | -0.20407 | 0.086881 | 1 |
| Mrpl14 | -0.20413 | 0.459626 | 1 |
| Hpgds | -0.2043 | 0.114463 | 1 |
| 0610040J0 | -0.20438 | 0.082512 | 1 |
| Rfx3 | -0.20467 | 0.024077 | 1 |
| Hivep1 | -0.20473 | 0.726977 | 1 |
| Trim30b | -0.20477 | 0.323536 | 1 |
| Map2k5 | -0.20488 | 0.915176 | 1 |
| Ccdc69 | -0.20502 | 0.010055 | 1 |
| Relch | -0.20505 | 0.173312 | 1 |
| Atxn7l1 | -0.20505 | 0.364542 | 1 |
| Prkd3 | -0.20516 | 0.19888 | 1 |
| Bloc1s2 | -0.20542 | 0.151358 | 1 |
| Mical1 | -0.2057 | 0.004554 | 1 |
| Pikfyve | -0.2058 | 0.151154 | 1 |
| Pxn | -0.20602 | 0.008469 | 1 |
| Map4k3 | -0.20637 | 0.196846 | 1 |
| Atad2b | -0.20645 | 0.02471 | 1 |
| Gpbp1 | -0.20655 | 0.291803 | 1 |
| Atp6v0a1 | -0.2067 | 0.750115 | 1 |
| Nsa2 | -0.20706 | 0.018756 | 1 |
| Abcc4 | -0.20714 | 0.309371 | 1 |
| Fam13b | -0.20734 | 0.539095 | 1 |
| Stim2 | -0.20745 | 0.008428 | 1 |
| Ccdc171 | -0.20765 | 0.047841 | 1 |
| Ralgapa1 | -0.20794 | 0.707497 | 1 |
| Maml1 | -0.20795 | 0.017266 | 1 |
| Exoc4 | -0.20821 | 0.989238 | 1 |
| Gm20663 | -0.20831 | 0.016355 | 1 |
| Capzb | -0.20847 | 1.62E-05 | 0.521867 |
| 3110082I1 | -0.20859 | 0.200925 | 1 |
| Gpr137b | -0.20867 | 0.259827 | 1 |
| Bdp1 | -0.2096 | 0.811654 | 1 |
| Nostrin | -0.20961 | 0.111033 | 1 |
| Tpp2 | -0.2097 | 0.083556 | 1 |
| Cdyl | -0.20991 | 0.354169 | 1 |
| 4632427E1 | -0.20993 | 0.056608 | 1 |
| Tbc1d5 | -0.21053 | 0.126457 | 1 |
| Cyb5r1 | -0.21057 | 0.597288 | 1 |
| Gm32401 | -0.2106 | 0.000124 | 1 |
| Rtn1 | -0.21121 | 1.62E-06 | 0.052353 |
| Sptbn1 | -0.21126 | 0.598396 | 1 |
| Cbfb | -0.21128 | 0.560407 | 1 |
| Map4k2 | -0.21133 | 0.341144 | 1 |
| Cux1 | -0.21157 | 0.170516 | 1 |
| Aip | -0.2117 | 0.188045 | 1 |
| Wwp2 | -0.2119 | 0.090948 | 1 |

| | | | |
|---|---|---|---|
| Unc13d | -0.21193 | 0.222679 | 1 |
| Rbm33 | -0.21194 | 0.116101 | 1 |
| Arid1b | -0.21201 | 0.104876 | 1 |
| Ubxn7 | -0.21203 | 0.037853 | 1 |
| Akt3 | -0.2121 | 0.306875 | 1 |
| Rpl38 | -0.21215 | 5.41E-05 | 1 |
| Prkab2 | -0.21219 | 0.005371 | 1 |
| Cop1 | -0.21224 | 0.147056 | 1 |
| Pnisr | -0.21244 | 0.304584 | 1 |
| Tmx3 | -0.21253 | 0.145182 | 1 |
| Atg7 | -0.21263 | 0.14399 | 1 |
| 5430427O: | -0.21272 | 0.004977 | 1 |
| Leng8 | -0.21279 | 0.054559 | 1 |
| Pofut1 | -0.21298 | 0.665705 | 1 |
| Rnf168 | -0.21303 | 0.223757 | 1 |
| Slc12a9 | -0.21314 | 0.42024 | 1 |
| Sept7 | -0.21317 | 0.063588 | 1 |
| Zcchc8 | -0.21353 | 0.026149 | 1 |
| Sp110 | -0.21374 | 0.018326 | 1 |
| Sos2 | -0.21377 | 0.102336 | 1 |
| Tmem50b | -0.21382 | 0.082448 | 1 |
| 4921511I1 | -0.21433 | 0.000376 | 1 |
| Eapp | -0.21435 | 0.86171 | 1 |
| Fyco1 | -0.21439 | 0.470714 | 1 |
| Cpne1 | -0.21445 | 0.105031 | 1 |
| Glud1 | -0.21467 | 0.000662 | 1 |
| H2-D1 | -0.2147 | 0.014689 | 1 |
| Tnrc6c | -0.21506 | 0.52342 | 1 |
| Cbx4 | -0.21512 | 0.025466 | 1 |
| Nmt1 | -0.21514 | 0.750334 | 1 |
| Slc4a7 | -0.21521 | 0.780378 | 1 |
| Taf3 | -0.21527 | 0.080416 | 1 |
| Puf60 | -0.21561 | 0.341369 | 1 |
| Kat2b | -0.21587 | 0.398867 | 1 |
| Zhx2 | -0.21592 | 6.36E-05 | 1 |
| Gm37494 | -0.21603 | 0.04693 | 1 |
| Gabarapl2 | -0.21617 | 0.353644 | 1 |
| Kmt5b | -0.21619 | 0.193615 | 1 |
| Tbc1d32 | -0.21622 | 0.443237 | 1 |
| Adgre1 | -0.21631 | 0.912721 | 1 |
| Rsbn1l | -0.21631 | 0.460766 | 1 |
| Gm13562 | -0.21653 | 0.072049 | 1 |
| Trmt112 | -0.21677 | 0.058809 | 1 |
| Lrrk1 | -0.21692 | 0.12719 | 1 |
| Scaper | -0.21732 | 0.967809 | 1 |
| Rp2 | -0.21747 | 0.045032 | 1 |
| Mfap3 | -0.21773 | 0.115296 | 1 |
| Itpr1 | -0.21829 | 0.20489 | 1 |

| | | | |
|---|---|---|---|
| Ap2b1 | -0.21846 | 0.326256 | 1 |
| Preb | -0.2185 | 0.012324 | 1 |
| Adamdec1 | -0.21852 | 0.123395 | 1 |
| Atp2c1 | -0.21878 | 0.222686 | 1 |
| Mta3 | -0.21894 | 0.38175 | 1 |
| Chd1 | -0.21904 | 0.557027 | 1 |
| Ugp2 | -0.21905 | 0.246411 | 1 |
| Cdk2ap2 | -0.2194 | 0.010845 | 1 |
| Hook3 | -0.21961 | 0.174796 | 1 |
| 4930469K1 | -0.21962 | 0.359614 | 1 |
| Iffo1 | -0.21963 | 0.190593 | 1 |
| Rmdn1 | -0.21973 | 0.080446 | 1 |
| Cebpd | -0.21994 | 0.248376 | 1 |
| Pld4 | -0.22058 | 0.006685 | 1 |
| Zfp944 | -0.22151 | 0.040733 | 1 |
| Akap8 | -0.22154 | 0.462339 | 1 |
| Cyld | -0.22155 | 0.035548 | 1 |
| Zeb2 | -0.22156 | 0.009291 | 1 |
| Mkln1 | -0.22195 | 0.104654 | 1 |
| Mapkapk2 | -0.22215 | 0.003475 | 1 |
| Tnrc6b | -0.22269 | 0.028806 | 1 |
| Arhgdib | -0.22315 | 0.000427 | 1 |
| Tfe3 | -0.22319 | 0.104347 | 1 |
| Elmsan1 | -0.22327 | 0.604259 | 1 |
| Tstd3 | -0.22347 | 0.059009 | 1 |
| Armc7 | -0.22366 | 0.024599 | 1 |
| Hdac7 | -0.22368 | 0.107109 | 1 |
| F630028O1 | -0.22377 | 0.003537 | 1 |
| Rpl3l | -0.22388 | 0.055614 | 1 |
| Arpc5 | -0.22408 | 0.00081 | 1 |
| Mrpl52 | -0.22409 | 0.007555 | 1 |
| Snx1 | -0.22426 | 0.035107 | 1 |
| Tpr | -0.22452 | 0.013266 | 1 |
| Dgka | -0.22474 | 0.001569 | 1 |
| Rhob | -0.22482 | 0.853153 | 1 |
| Ist1 | -0.22486 | 0.121568 | 1 |
| Ldah | -0.22508 | 0.020626 | 1 |
| Traf3ip3 | -0.22521 | 0.046331 | 1 |
| Rspry1 | -0.22533 | 0.531095 | 1 |
| Gm10130 | -0.22593 | 0.032869 | 1 |
| Piezo1 | -0.22597 | 0.254472 | 1 |
| Gm16120 | -0.22607 | 1.89E-06 | 0.061095 |
| Rrm2b | -0.22608 | 0.113705 | 1 |
| Zranb1 | -0.2262 | 0.158098 | 1 |
| Gab3 | -0.22646 | 0.022371 | 1 |
| Tmsb4x | -0.22685 | 2.5E-08 | 0.000807 |
| Arhgap9 | -0.22733 | 0.009237 | 1 |
| Gch1 | -0.22743 | 0.090863 | 1 |

| | | | |
|---|---|---|---|
| Prex1 | -0.22748 | 0.041339 | 1 |
| Baz1a | -0.22778 | 0.034318 | 1 |
| Lrmp | -0.22788 | 0.618347 | 1 |
| Kif21b | -0.22791 | 0.145474 | 1 |
| Herc1 | -0.22808 | 0.154215 | 1 |
| Sh3bp2 | -0.22812 | 0.037247 | 1 |
| Abhd2 | -0.22849 | 0.321119 | 1 |
| Rsrc1 | -0.22906 | 0.481912 | 1 |
| 9030404E1 | -0.22921 | 7.6E-07 | 0.024528 |
| Gdi1 | -0.22934 | 0.139844 | 1 |
| Ntng2 | -0.22943 | 0.268723 | 1 |
| Trip11 | -0.22958 | 0.13845 | 1 |
| Etnk1 | -0.22976 | 0.451595 | 1 |
| Gm4070 | -0.22977 | 0.211115 | 1 |
| Vegfc | -0.23148 | 1.29E-08 | 0.000417 |
| Mettl7a1 | -0.23164 | 0.000341 | 1 |
| Bola2 | -0.23172 | 0.13798 | 1 |
| Gtdc1 | -0.23221 | 0.013081 | 1 |
| Ppp3ca | -0.23229 | 0.045021 | 1 |
| Rac2 | -0.23291 | 0.004299 | 1 |
| Card9 | -0.23303 | 0.081036 | 1 |
| Stk40 | -0.23306 | 0.017969 | 1 |
| Tsc22d3 | -0.2334 | 0.048029 | 1 |
| Smagp | -0.2335 | 0.001298 | 1 |
| Ptpn1 | -0.23361 | 0.000512 | 1 |
| Ccdc180 | -0.23362 | 0.036896 | 1 |
| Gm11651 | -0.23381 | 1.37E-08 | 0.000443 |
| Ifit3b | -0.23383 | 7.36E-05 | 1 |
| Gm42418 | -0.23448 | 0.037331 | 1 |
| Far1 | -0.23452 | 0.002256 | 1 |
| Cdc42ep2 | -0.23456 | 0.430475 | 1 |
| Bin3 | -0.23463 | 0.11586 | 1 |
| Mink1 | -0.23485 | 0.568706 | 1 |
| Capza1 | -0.23487 | 0.002281 | 1 |
| Stat6 | -0.2351 | 0.167531 | 1 |
| Ccdc97 | -0.23525 | 0.106946 | 1 |
| Ppfia4 | -0.23537 | 0.026488 | 1 |
| Mad1l1 | -0.23547 | 0.299226 | 1 |
| Zfp52 | -0.23548 | 0.002893 | 1 |
| Dazap2 | -0.23575 | 7.52E-05 | 1 |
| Ambra1 | -0.23592 | 0.042054 | 1 |
| Rab11b | -0.23643 | 0.14883 | 1 |
| Phf3 | -0.23655 | 0.935767 | 1 |
| 2310009A( | -0.2369 | 0.12966 | 1 |
| Rnf167 | -0.23721 | 0.082502 | 1 |
| Dap | -0.23769 | 0.114667 | 1 |
| Nudt13 | -0.23794 | 0.028919 | 1 |
| Mbnl1 | -0.23797 | 0.001101 | 1 |

| | | | |
|---|---|---|---|
| Myo1c | -0.23805 | 0.163648 | 1 |
| Csf2ra | -0.23815 | 0.000274 | 1 |
| Lats2 | -0.2382 | 0.235694 | 1 |
| Arrb1 | -0.23844 | 0.338662 | 1 |
| Bloc1s1 | -0.2385 | 0.054582 | 1 |
| Slc25a38 | -0.23891 | 0.087038 | 1 |
| Rere | -0.23923 | 0.21055 | 1 |
| Pik3cg | -0.23927 | 0.027397 | 1 |
| Slc38a9 | -0.2394 | 0.112467 | 1 |
| Srsf5 | -0.23958 | 0.014685 | 1 |
| Phyh | -0.23961 | 0.484088 | 1 |
| Gabpb2 | -0.23963 | 0.030096 | 1 |
| Fryl | -0.2398 | 0.033576 | 1 |
| Nrd1 | -0.23984 | 0.031947 | 1 |
| Fndc3a | -0.24032 | 0.03111 | 1 |
| Ampd3 | -0.24054 | 0.321615 | 1 |
| Fut8 | -0.24056 | 0.559189 | 1 |
| Slc39a13 | -0.24059 | 0.018277 | 1 |
| Parp8 | -0.24066 | 0.022839 | 1 |
| Gm37401 | -0.24082 | 0.005498 | 1 |
| Gm7644 | -0.24111 | 0.000464 | 1 |
| Tcf4 | -0.24303 | 0.34159 | 1 |
| Zfand6 | -0.24307 | 0.026732 | 1 |
| Slc35c2 | -0.24314 | 0.005718 | 1 |
| Nsd3 | -0.24328 | 0.00237 | 1 |
| Adamts10 | -0.24329 | 0.144535 | 1 |
| Tmem156 | -0.24334 | 0.00163 | 1 |
| Rbm38 | -0.24345 | 0.011456 | 1 |
| Cd82 | -0.24349 | 0.164352 | 1 |
| Gab2 | -0.2435 | 0.19848 | 1 |
| Dmtf1 | -0.24353 | 0.048312 | 1 |
| Tppp3 | -0.24365 | 0.084216 | 1 |
| Hnrnph3 | -0.24369 | 0.284064 | 1 |
| Hmgcs1 | -0.2437 | 0.424455 | 1 |
| Zfp407 | -0.2437 | 0.549487 | 1 |
| Ip6k3 | -0.24398 | 1.47E-06 | 0.04751 |
| Ptger4 | -0.24419 | 0.109069 | 1 |
| Setd2 | -0.24475 | 0.069518 | 1 |
| Zbtb7b | -0.24503 | 0.001874 | 1 |
| Helz2 | -0.24506 | 0.041216 | 1 |
| Rchy1 | -0.24514 | 0.720443 | 1 |
| Lima1 | -0.24515 | 0.047859 | 1 |
| Sipa1 | -0.24518 | 0.051805 | 1 |
| Gpsm3 | -0.24518 | 0.000747 | 1 |
| Vps13c | -0.24531 | 0.077955 | 1 |
| Gpr183 | -0.24553 | 0.008473 | 1 |
| Supt6 | -0.24554 | 0.269508 | 1 |
| Mapk1 | -0.24602 | 0.110186 | 1 |

| | | | |
|---|---|---|---|
| Cdc40 | -0.24618 | 0.078236 | 1 |
| Pten | -0.24672 | 0.077172 | 1 |
| Mcmbp | -0.2474 | 0.04455 | 1 |
| Rbms1 | -0.24745 | 0.015301 | 1 |
| Itch | -0.24811 | 0.003734 | 1 |
| Cdc5l | -0.24839 | 0.046239 | 1 |
| Srgap2 | -0.2484 | 0.293994 | 1 |
| Atxn7 | -0.24855 | 0.060854 | 1 |
| 5031425E2 | -0.24856 | 0.012106 | 1 |
| Usf1 | -0.24857 | 0.067083 | 1 |
| Zfp608 | -0.24878 | 0.004248 | 1 |
| Enox2 | -0.24882 | 0.028618 | 1 |
| Tmed5 | -0.24882 | 0.023928 | 1 |
| AC149090. | -0.24897 | 0.118386 | 1 |
| Erlin1 | -0.24926 | 0.225655 | 1 |
| Setbp1 | -0.24986 | 0.000383 | 1 |
| Macf1 | -0.25032 | 0.00108 | 1 |
| Erbin | -0.25057 | 0.027361 | 1 |
| Arhgap15 | -0.25072 | 0.002742 | 1 |
| A130014A( | -0.25074 | 0.000115 | 1 |
| Clcn3 | -0.25088 | 0.404395 | 1 |
| H2-T24 | -0.2509 | 0.006071 | 1 |
| Mrtfa | -0.2509 | 0.017132 | 1 |
| Pias1 | -0.251 | 0.018088 | 1 |
| Phf20l1 | -0.2511 | 0.401934 | 1 |
| Wrn | -0.25229 | 0.352943 | 1 |
| Plekhj1 | -0.25273 | 0.070581 | 1 |
| Plek | -0.25281 | 0.027992 | 1 |
| Rc3h1 | -0.2529 | 0.274284 | 1 |
| Nckap1l | -0.25331 | 0.069843 | 1 |
| Hsd17b11 | -0.25343 | 0.057294 | 1 |
| Acbd5 | -0.2535 | 0.102839 | 1 |
| Hnrnpul1 | -0.25355 | 0.018135 | 1 |
| Zfp869 | -0.25356 | 0.017419 | 1 |
| Lysmd4 | -0.25401 | 0.080799 | 1 |
| Msra | -0.25406 | 0.089686 | 1 |
| Sgk1 | -0.25422 | 0.012402 | 1 |
| Rsbn1 | -0.25479 | 0.044149 | 1 |
| Map3k3 | -0.25489 | 0.071039 | 1 |
| Mdm4 | -0.25489 | 0.107012 | 1 |
| Ppm1b | -0.25497 | 0.153473 | 1 |
| Camk1d | -0.25527 | 0.002313 | 1 |
| Clcn4 | -0.25608 | 0.427525 | 1 |
| Kdm5a | -0.25609 | 0.541488 | 1 |
| Luc7l2 | -0.25623 | 0.005873 | 1 |
| Map3k11 | -0.25633 | 0.00949 | 1 |
| BC028528 | -0.25653 | 0.046092 | 1 |
| Pik3r5 | -0.25655 | 0.0478 | 1 |

| | | | |
|---|---|---|---|
| Zfp445 | -0.25659 | 0.154316 | 1 |
| Syf2 | -0.25667 | 0.019857 | 1 |
| Wasf2 | -0.2567 | 0.000981 | 1 |
| Tmc6 | -0.25673 | 0.000228 | 1 |
| G6pdx | -0.25674 | 0.01816 | 1 |
| Vezf1 | -0.25675 | 0.005441 | 1 |
| Extl3 | -0.25678 | 0.148047 | 1 |
| Msl2 | -0.25683 | 0.010147 | 1 |
| Alpk1 | -0.25704 | 0.169611 | 1 |
| Atp2a3 | -0.25748 | 0.002992 | 1 |
| Nav2 | -0.25766 | 0.013495 | 1 |
| Arl6ip5 | -0.2579 | 0.001419 | 1 |
| Zfyve16 | -0.25851 | 0.216335 | 1 |
| Nfkb2 | -0.25855 | 0.213058 | 1 |
| Capn2 | -0.25895 | 0.011744 | 1 |
| Cept1 | -0.25926 | 0.194161 | 1 |
| Ms4a7 | -0.25947 | 0.153132 | 1 |
| R3hdm1 | -0.26074 | 0.050228 | 1 |
| Ctdsp1 | -0.26084 | 0.815724 | 1 |
| Msrb1 | -0.26157 | 0.000546 | 1 |
| Csk | -0.26159 | 0.003848 | 1 |
| Actr10 | -0.26179 | 0.121122 | 1 |
| Asph | -0.26196 | 0.001801 | 1 |
| Cat | -0.26209 | 0.425439 | 1 |
| Secisbp2 | -0.26243 | 0.061595 | 1 |
| Ezr | -0.26252 | 0.104989 | 1 |
| Ifngr2 | -0.26276 | 0.00084 | 1 |
| Gmip | -0.26301 | 0.000908 | 1 |
| Phtf2 | -0.26317 | 0.049983 | 1 |
| Mknk1 | -0.26376 | 0.030692 | 1 |
| Vps37b | -0.26417 | 0.027568 | 1 |
| Klf13 | -0.26417 | 0.001207 | 1 |
| Rps21 | -0.26437 | 1.26E-06 | 0.040608 |
| Api5 | -0.26451 | 0.030126 | 1 |
| Cdk13 | -0.26457 | 0.433054 | 1 |
| Arrdc1 | -0.26473 | 0.145198 | 1 |
| Tsc22d4 | -0.26507 | 0.000999 | 1 |
| Usp15 | -0.26553 | 0.078637 | 1 |
| Calm1 | -0.26569 | 0.000107 | 1 |
| Sms | -0.26596 | 0.087522 | 1 |
| H2-K1 | -0.26606 | 2.55E-06 | 0.082313 |
| Dok1 | -0.26606 | 0.134192 | 1 |
| Gm26877 | -0.26646 | 5.62E-06 | 0.181503 |
| Dido1 | -0.26662 | 0.013224 | 1 |
| Dhx36 | -0.26665 | 0.068242 | 1 |
| A330040F1 | -0.26688 | 0.098853 | 1 |
| Git2 | -0.26711 | 0.002628 | 1 |
| Cd83 | -0.26726 | 0.988335 | 1 |

| | | | |
|---|---|---|---|
| Phactr2 | -0.26734 | 0.405602 | 1 |
| Ccl12 | -0.2675 | 0.049263 | 1 |
| Taz | -0.26754 | 0.105717 | 1 |
| Pum2 | -0.26772 | 0.080636 | 1 |
| Scaf11 | -0.26832 | 0.687798 | 1 |
| Papola | -0.26843 | 0.018406 | 1 |
| Zeb2os | -0.26848 | 0.024057 | 1 |
| Kansl1 | -0.26854 | 0.008963 | 1 |
| Haao | -0.26926 | 0.001128 | 1 |
| Syne1 | -0.26937 | 0.122465 | 1 |
| Sh3bgrl | -0.26985 | 0.003321 | 1 |
| B4galnt1 | -0.27096 | 0.02658 | 1 |
| Nbn | -0.27107 | 0.02463 | 1 |
| Dcakd | -0.27183 | 0.035492 | 1 |
| Ostf1 | -0.27197 | 8.97E-05 | 1 |
| Sec14l1 | -0.27206 | 0.185193 | 1 |
| Tmem135 | -0.2723 | 0.056168 | 1 |
| Dusp2 | -0.27237 | 0.10608 | 1 |
| Cdc42 | -0.27238 | 2.67E-07 | 0.008615 |
| Stat3 | -0.27242 | 0.003359 | 1 |
| Ino80d | -0.27244 | 0.00678 | 1 |
| Fam214b | -0.27269 | 0.011105 | 1 |
| Fam172a | -0.2735 | 0.084344 | 1 |
| Smchd1 | -0.27419 | 0.004696 | 1 |
| Bank1 | -0.27459 | 0.014617 | 1 |
| Usp47 | -0.27469 | 0.120343 | 1 |
| Ylpm1 | -0.27484 | 0.056258 | 1 |
| Phf14 | -0.27523 | 0.016818 | 1 |
| Evi2a | -0.27558 | 0.083557 | 1 |
| Fam126a | -0.27574 | 0.040813 | 1 |
| Nop10 | -0.27685 | 0.030604 | 1 |
| B3galnt2 | -0.27706 | 0.000734 | 1 |
| Sh3bp1 | -0.27746 | 0.016638 | 1 |
| Prpf4b | -0.27784 | 0.070598 | 1 |
| Kat6a | -0.27827 | 0.00343 | 1 |
| Cmpk2 | -0.27882 | 0.01682 | 1 |
| Pet100 | -0.27884 | 0.029146 | 1 |
| Msi2 | -0.27886 | 0.273754 | 1 |
| Sp1 | -0.27917 | 0.177293 | 1 |
| Btbd9 | -0.27934 | 0.062141 | 1 |
| E2f2 | -0.27966 | 0.050109 | 1 |
| Ldlr | -0.27969 | 0.019497 | 1 |
| Gvin1 | -0.28001 | 0.031505 | 1 |
| Frmd8 | -0.28028 | 0.008564 | 1 |
| Cd44 | -0.28068 | 6.43E-05 | 1 |
| Dennd4a | -0.2807 | 0.266193 | 1 |
| Gm17749 | -0.28092 | 9.6E-08 | 0.0031 |
| Copg2 | -0.2826 | 0.097644 | 1 |

| | | | |
|---|---|---|---|
| Mier1 | -0.28286 | 0.004128 | 1 |
| Srsf11 | -0.28328 | 0.026717 | 1 |
| Anxa2 | -0.2835 | 0.00052 | 1 |
| Cd47 | -0.28381 | 0.000126 | 1 |
| Cnot2 | -0.28396 | 0.198716 | 1 |
| 1600010M | -0.28396 | 0.00281 | 1 |
| Rnf216 | -0.28419 | 0.073875 | 1 |
| Zfand5 | -0.28488 | 0.011755 | 1 |
| C2cd5 | -0.28492 | 0.206959 | 1 |
| Nrarp | -0.28515 | 8.18E-07 | 0.026414 |
| Pros1 | -0.2856 | 0.003813 | 1 |
| Mia2 | -0.28562 | 0.034222 | 1 |
| Arid4a | -0.28611 | 0.000214 | 1 |
| Rab10 | -0.28632 | 0.002082 | 1 |
| Trim30d | -0.28637 | 0.188537 | 1 |
| Apobr | -0.28701 | 0.081264 | 1 |
| Crk | -0.28726 | 0.101805 | 1 |
| Srebf2 | -0.28745 | 0.046101 | 1 |
| Bcl11a | -0.28835 | 5.02E-06 | 0.162056 |
| Snx20 | -0.28836 | 0.045256 | 1 |
| Swap70 | -0.28873 | 0.000376 | 1 |
| Efcab11 | -0.28873 | 0.027134 | 1 |
| Gm13919 | -0.28876 | 1.24E-07 | 0.003996 |
| St3gal6 | -0.28877 | 1.83E-05 | 0.592337 |
| Zdhhc6 | -0.28899 | 0.005261 | 1 |
| Klf3 | -0.28917 | 0.019059 | 1 |
| Lpin2 | -0.28919 | 0.027474 | 1 |
| Fcer1g | -0.28969 | 4.32E-05 | 1 |
| Pik3cd | -0.29014 | 0.000462 | 1 |
| Rps6ka1 | -0.29017 | 0.000362 | 1 |
| Tnfrsf21 | -0.29036 | 0.022226 | 1 |
| Zfc3h1 | -0.29044 | 0.000852 | 1 |
| Nt5dc1 | -0.29142 | 0.101771 | 1 |
| Thbd | -0.29153 | 1.07E-05 | 0.344166 |
| 8030462N | -0.29184 | 0.000303 | 1 |
| Cbfa2t3 | -0.29313 | 0.001847 | 1 |
| Magt1 | -0.29364 | 0.002767 | 1 |
| 2210408F | -0.29366 | 0.048226 | 1 |
| Zc3h12d | -0.29386 | 1.14E-05 | 0.36934 |
| Tom1 | -0.29448 | 0.015825 | 1 |
| Tnfrsf13b | -0.2945 | 0.057258 | 1 |
| Lilra5 | -0.29465 | 0.042996 | 1 |
| Tmem38b | -0.29471 | 0.016397 | 1 |
| Tnfaip8 | -0.29539 | 0.001197 | 1 |
| Jun | -0.29568 | 0.164686 | 1 |
| Gm2245 | -0.29581 | 0.00771 | 1 |
| Celf1 | -0.29587 | 0.231711 | 1 |
| Zfp217 | -0.29621 | 0.009382 | 1 |

| | | | |
|---|---|---|---|
| Degs1 | -0.29635 | 0.000639 | 1 |
| Trpm2 | -0.29649 | 0.618104 | 1 |
| Tnrc18 | -0.29651 | 0.00728 | 1 |
| Itgb7 | -0.29664 | 0.001864 | 1 |
| Foxk1 | -0.29668 | 0.003128 | 1 |
| Bmt2 | -0.29668 | 0.024539 | 1 |
| Vamp5 | -0.29728 | 0.034095 | 1 |
| Rexo1 | -0.29729 | 0.007595 | 1 |
| Abca9 | -0.29734 | 0.00561 | 1 |
| Mdm1 | -0.29819 | 9.1E-07 | 0.029375 |
| Usp9x | -0.2982 | 0.013636 | 1 |
| Sfi1 | -0.29827 | 0.083465 | 1 |
| Spidr | -0.29898 | 0.058657 | 1 |
| Trim12c | -0.29906 | 0.016231 | 1 |
| Slc16a10 | -0.29922 | 0.008691 | 1 |
| Cib1 | -0.29934 | 0.021869 | 1 |
| Ptgir | -0.2998 | 0.000161 | 1 |
| Madd | -0.30002 | 0.001904 | 1 |
| Ndel1 | -0.30005 | 0.068275 | 1 |
| H2afj | -0.30072 | 0.004674 | 1 |
| St3gal1 | -0.30074 | 0.000285 | 1 |
| Lpcat4 | -0.30078 | 0.000933 | 1 |
| Actr2 | -0.30119 | 1.11E-05 | 0.358301 |
| Mxi1 | -0.30122 | 0.130207 | 1 |
| Rnf149 | -0.30129 | 0.004137 | 1 |
| B4galt5 | -0.30164 | 0.042539 | 1 |
| Fam168a | -0.3022 | 0.135226 | 1 |
| Tasor | -0.30267 | 0.036422 | 1 |
| Zeb1 | -0.3037 | 0.000452 | 1 |
| Samd9l | -0.30372 | 0.016509 | 1 |
| Phf21a | -0.30391 | 0.004737 | 1 |
| Ccnl2 | -0.30408 | 0.012453 | 1 |
| Hcls1 | -0.30435 | 0.000249 | 1 |
| Mcl1 | -0.30436 | 4.83E-05 | 1 |
| Acot9 | -0.30603 | 0.005809 | 1 |
| Stxbp6 | -0.30666 | 2.62E-05 | 0.845464 |
| Cmpk1 | -0.30696 | 0.000903 | 1 |
| Cyp27a1 | -0.30734 | 0.002609 | 1 |
| Dennd5a | -0.30748 | 0.000314 | 1 |
| Ifnar2 | -0.30749 | 0.012118 | 1 |
| Clk1 | -0.30768 | 0.001807 | 1 |
| Ralgapa2 | -0.30777 | 0.061722 | 1 |
| Sept9 | -0.30785 | 0.001161 | 1 |
| Kif5b | -0.3079 | 0.000578 | 1 |
| Apobec1 | -0.30813 | 0.002902 | 1 |
| Rabgap1l | -0.30851 | 0.084629 | 1 |
| Apba1 | -0.30853 | 0.00333 | 1 |
| Myl12b | -0.3092 | 1.4E-07 | 0.004515 |

| | | | |
|---|---|---|---|
| Oxr1 | -0.3094 | 0.004143 | 1 |
| Mapk14 | -0.3097 | 0.002566 | 1 |
| Arid1a | -0.30986 | 0.005053 | 1 |
| Rps6ka2 | -0.31006 | 1.76E-06 | 0.056725 |
| Gm32633 | -0.3103 | 6.43E-06 | 0.207623 |
| Ep300 | -0.31056 | 0.016754 | 1 |
| Naip2 | -0.31062 | 0.004802 | 1 |
| Galnt1 | -0.3108 | 0.098095 | 1 |
| Ythdf3 | -0.31096 | 0.011363 | 1 |
| Irf5 | -0.31159 | 1.96E-05 | 0.633403 |
| Dgkd | -0.31182 | 0.000979 | 1 |
| Synj1 | -0.31231 | 0.000525 | 1 |
| Ddx6 | -0.31233 | 0.001547 | 1 |
| Zcchc7 | -0.31253 | 0.052465 | 1 |
| Mapkap1 | -0.31281 | 0.020877 | 1 |
| Zfp638 | -0.31304 | 0.006834 | 1 |
| Tmem131l | -0.31336 | 0.00302 | 1 |
| Ccr2 | -0.31338 | 0.392748 | 1 |
| Tpm4 | -0.31348 | 0.001334 | 1 |
| Ngly1 | -0.31359 | 0.107789 | 1 |
| Tmem51 | -0.31456 | 0.000649 | 1 |
| Anxa6 | -0.31472 | 0.013914 | 1 |
| Rnase4 | -0.31546 | 0.968243 | 1 |
| Itsn2 | -0.31567 | 0.046016 | 1 |
| Nop53 | -0.31597 | 0.075826 | 1 |
| Spred1 | -0.316 | 0.278647 | 1 |
| Prkcd | -0.31601 | 2.37E-06 | 0.076438 |
| Polr2a | -0.31629 | 0.000478 | 1 |
| Sltm | -0.31655 | 0.064651 | 1 |
| Inpp4b | -0.31664 | 6.14E-05 | 1 |
| Ddx5 | -0.3177 | 1.95E-06 | 0.06289 |
| Limd1 | -0.31771 | 0.00153 | 1 |
| Stx16 | -0.31807 | 0.001129 | 1 |
| Hes1 | -0.31809 | 4.94E-05 | 1 |
| Vcl | -0.31947 | 0.000347 | 1 |
| Foxo3 | -0.31982 | 0.005378 | 1 |
| Sdccag8 | -0.31993 | 0.029061 | 1 |
| Smpdl3a | -0.32 | 0.000229 | 1 |
| Tfeb | -0.32054 | 0.006833 | 1 |
| Eif4a2 | -0.32108 | 0.01522 | 1 |
| Sipa1l3 | -0.32134 | 0.008604 | 1 |
| Lims1 | -0.32306 | 0.000189 | 1 |
| Pkn1 | -0.32484 | 0.002281 | 1 |
| Ash1l | -0.32487 | 0.00192 | 1 |
| Fam32a | -0.32543 | 0.000703 | 1 |
| Zzef1 | -0.32655 | 0.056282 | 1 |
| Tank | -0.32661 | 0.025956 | 1 |
| Cnot6l | -0.32666 | 0.003044 | 1 |

| | | | |
|---|---|---|---|
| Ckb | -0.32675 | 0.015612 | 1 |
| Trim12a | -0.32696 | 0.042434 | 1 |
| Sh2b3 | -0.32724 | 0.004184 | 1 |
| Tiam1 | -0.32881 | 0.190825 | 1 |
| Ncor1 | -0.32887 | 0.003172 | 1 |
| Cebpb | -0.3293 | 0.002069 | 1 |
| Pid1 | -0.3294 | 0.000103 | 1 |
| Mef2d | -0.32969 | 0.01772 | 1 |
| Rbm39 | -0.33006 | 2.65E-06 | 0.085455 |
| Ext1 | -0.33238 | 0.107743 | 1 |
| Ttc17 | -0.33264 | 0.014028 | 1 |
| Rnf130 | -0.33322 | 0.005348 | 1 |
| Arsb | -0.3336 | 0.00164 | 1 |
| Naga | -0.33362 | 0.000115 | 1 |
| Tbc1d2b | -0.33397 | 0.134733 | 1 |
| Abcd1 | -0.33412 | 0.034584 | 1 |
| Stag1 | -0.33474 | 0.000629 | 1 |
| Ddx3x | -0.33477 | 0.000446 | 1 |
| Evl | -0.33479 | 0.000779 | 1 |
| Specc1 | -0.33495 | 0.001008 | 1 |
| Gstm1 | -0.33525 | 0.108602 | 1 |
| Gsk3b | -0.33564 | 0.002301 | 1 |
| Myh9 | -0.33637 | 1.55E-05 | 0.501375 |
| A630001G: | -0.33655 | 0.025766 | 1 |
| Crlf3 | -0.33684 | 0.018567 | 1 |
| Ggta1 | -0.33694 | 0.00617 | 1 |
| Fau | -0.33727 | 4.69E-13 | 1.51E-08 |
| Birc6 | -0.33746 | 0.003892 | 1 |
| Cep170 | -0.3375 | 0.000423 | 1 |
| Epc1 | -0.33753 | 0.008349 | 1 |
| Krit1 | -0.33811 | 0.012357 | 1 |
| Mfsd6 | -0.33833 | 4.68E-08 | 0.001512 |
| Ncoa2 | -0.33935 | 0.096612 | 1 |
| 4833407H: | -0.34004 | 2E-08 | 0.000647 |
| Tlr13 | -0.34082 | 0.003002 | 1 |
| Fes | -0.34162 | 0.001212 | 1 |
| Cd300ld3 | -0.34163 | 5.63E-11 | 1.82E-06 |
| Tspan14 | -0.34243 | 0.000156 | 1 |
| Pilrb1 | -0.34265 | 0.002098 | 1 |
| Ebi3 | -0.34295 | 0.006147 | 1 |
| Rab19 | -0.34345 | 0.000374 | 1 |
| Cbl | -0.3435 | 0.001057 | 1 |
| Tgm3 | -0.34468 | 1.42E-08 | 0.000458 |
| Kdm3b | -0.34508 | 0.004385 | 1 |
| Elf1 | -0.34532 | 0.000203 | 1 |
| Tmem119 | -0.3455 | 0.089042 | 1 |
| Gpr34 | -0.34578 | 0.636778 | 1 |
| Stx11 | -0.34654 | 0.074242 | 1 |

| | | | |
|---|---|---|---|
| Atp8b4 | -0.34708 | 0.002561 | 1 |
| Sash3 | -0.34739 | 2.58E-05 | 0.833174 |
| Fcrl1 | -0.34786 | 3.45E-08 | 0.001115 |
| Nhsl2 | -0.34834 | 0.005358 | 1 |
| Ppdpf | -0.34849 | 9.94E-05 | 1 |
| Ppp4r3b | -0.34888 | 0.003338 | 1 |
| Eif1 | -0.34941 | 4.85E-09 | 0.000156 |
| Itgb1 | -0.35053 | 0.001704 | 1 |
| Snrpb | -0.35086 | 4.06E-06 | 0.130945 |
| Ifit3 | -0.35125 | 0.077896 | 1 |
| Dpp4 | -0.35136 | 6.14E-07 | 0.019814 |
| Mecp2 | -0.35154 | 0.002347 | 1 |
| Trib1 | -0.35154 | 0.08754 | 1 |
| Gstp1 | -0.35217 | 0.000136 | 1 |
| Ogt | -0.35252 | 0.001689 | 1 |
| Dstyk | -0.35382 | 0.000224 | 1 |
| Lbh | -0.35397 | 0.002527 | 1 |
| Sgms1 | -0.3546 | 0.010484 | 1 |
| Nptn | -0.3547 | 1.85E-05 | 0.598187 |
| Hspa1b | -0.35478 | 0.026073 | 1 |
| Unc119 | -0.35478 | 0.105625 | 1 |
| Megf9 | -0.35498 | 5.72E-05 | 1 |
| Zc3hav1 | -0.35512 | 0.000301 | 1 |
| Glipr1 | -0.35515 | 0.010882 | 1 |
| Kdm6b | -0.35521 | 0.008921 | 1 |
| Strn3 | -0.35566 | 0.205187 | 1 |
| Cyp4f18 | -0.35754 | 0.000312 | 1 |
| Gcnt1 | -0.35844 | 0.074913 | 1 |
| Daglb | -0.3595 | 0.001177 | 1 |
| Gramd4 | -0.35964 | 6.3E-05 | 1 |
| Taldo1 | -0.3607 | 2.58E-05 | 0.832366 |
| Atf7ip | -0.36087 | 0.012641 | 1 |
| Tnrc6a | -0.36115 | 0.088433 | 1 |
| Lyst | -0.36158 | 0.034014 | 1 |
| Polb | -0.36174 | 0.000572 | 1 |
| Dock11 | -0.36175 | 0.003028 | 1 |
| Usp25 | -0.36179 | 0.050847 | 1 |
| Inpp1 | -0.36296 | 8.58E-05 | 1 |
| Padi2 | -0.36518 | 7.66E-08 | 0.002474 |
| Slc15a4 | -0.36539 | 0.000597 | 1 |
| Rras | -0.36548 | 0.001651 | 1 |
| Rpl37a | -0.36601 | 5.69E-12 | 1.84E-07 |
| Lncpint | -0.36659 | 0.017158 | 1 |
| Gm5547 | -0.367 | 3.76E-09 | 0.000121 |
| Cdc14a | -0.36803 | 0.000408 | 1 |
| Nipbl | -0.36867 | 0.000548 | 1 |
| Nin | -0.36924 | 0.003573 | 1 |
| Rassf4 | -0.36972 | 1.13E-05 | 0.364205 |

| | | | |
|---|---|---|---|
| Adssl1 | -0.37053 | 0.000402 | 1 |
| Crebbp | -0.37077 | 0.001574 | 1 |
| Lrrc25 | -0.37112 | 3.23E-07 | 0.010413 |
| Rgs3 | -0.37127 | 8.78E-05 | 1 |
| Taok3 | -0.37202 | 5.95E-06 | 0.191991 |
| Sh3bp5 | -0.37204 | 0.000274 | 1 |
| Cd37 | -0.37241 | 0.015502 | 1 |
| Lars2 | -0.37267 | 1.58E-08 | 0.00051 |
| Ccdc125 | -0.3729 | 0.006235 | 1 |
| Smad3 | -0.37307 | 0.008314 | 1 |
| Cap1 | -0.3737 | 3.84E-10 | 1.24E-05 |
| Rhoq | -0.37377 | 0.431048 | 1 |
| Rps29 | -0.37379 | 1.23E-12 | 3.97E-08 |
| Grk5 | -0.37495 | 0.023132 | 1 |
| Sept6 | -0.37658 | 3E-05 | 0.968123 |
| 3110021N1 | -0.37835 | 1.31E-07 | 0.004225 |
| Rhoa | -0.37837 | 1.77E-10 | 5.71E-06 |
| Rbpms | -0.37853 | 0.001629 | 1 |
| Sort1 | -0.37878 | 0.002841 | 1 |
| Acap2 | -0.37912 | 4.06E-06 | 0.131051 |
| Arl4c | -0.38004 | 0.00636 | 1 |
| Cmss1 | -0.38163 | 2.83E-10 | 9.15E-06 |
| Trps1 | -0.38193 | 0.001759 | 1 |
| Atg4d | -0.38198 | 4.61E-05 | 1 |
| Il10ra | -0.38349 | 1.85E-06 | 0.059664 |
| Ccnd2 | -0.38364 | 0.671513 | 1 |
| Stk3 | -0.38378 | 0.007725 | 1 |
| Wdfy4 | -0.38384 | 0.037932 | 1 |
| Ptbp3 | -0.38401 | 1.66E-08 | 0.000534 |
| Tgfbr2 | -0.38403 | 0.00276 | 1 |
| Nfat5 | -0.38438 | 0.072773 | 1 |
| Myo18a | -0.38465 | 0.000165 | 1 |
| Iqgap1 | -0.38477 | 5.36E-08 | 0.00173 |
| Birc3 | -0.3849 | 8.67E-05 | 1 |
| Stag2 | -0.38529 | 0.000398 | 1 |
| Arhgap17 | -0.38585 | 1.25E-05 | 0.403493 |
| Actr3 | -0.38598 | 7.47E-11 | 2.41E-06 |
| Ankhd1 | -0.38605 | 0.000192 | 1 |
| Rin2 | -0.38644 | 0.000332 | 1 |
| Acer3 | -0.38652 | 0.000211 | 1 |
| Map2k1 | -0.38735 | 1.12E-06 | 0.0361 |
| Rsf1 | -0.38792 | 0.003796 | 1 |
| Igsf6 | -0.38806 | 3.86E-06 | 0.124585 |
| Abcc5 | -0.38826 | 0.085705 | 1 |
| Inpp5d | -0.38838 | 0.00012 | 1 |
| Abca7 | -0.38871 | 0.000404 | 1 |
| St8sia4 | -0.38889 | 0.001205 | 1 |
| Tgfb1 | -0.38895 | 1.41E-06 | 0.045509 |

| | | | |
|---|---|---|---|
| Bri3bp | -0.38911 | 8.22E-06 | 0.265343 |
| Psenen | -0.39061 | 4.86E-06 | 0.157051 |
| Baz2b | -0.39105 | 3.11E-06 | 0.100251 |
| Havcr2 | -0.39113 | 0.000887 | 1 |
| Nfatc1 | -0.39115 | 7.74E-05 | 1 |
| Map3k1 | -0.39171 | 0.000355 | 1 |
| D1Ertd622 | -0.39195 | 0.000362 | 1 |
| Tifab | -0.39207 | 1.86E-08 | 0.000599 |
| Plac8 | -0.39218 | 1.21E-06 | 0.039154 |
| 9930111J2 | -0.39245 | 5.09E-06 | 0.16424 |
| Ncor2 | -0.39293 | 0.001051 | 1 |
| Akap13 | -0.39345 | 7.18E-06 | 0.231827 |
| Fam49b | -0.39409 | 1.44E-06 | 0.046435 |
| Gnb2 | -0.39441 | 1.75E-07 | 0.005653 |
| Kras | -0.39453 | 0.000933 | 1 |
| Pfkfb4 | -0.39597 | 5.06E-05 | 1 |
| Rnf144a | -0.39874 | 2.07E-12 | 6.67E-08 |
| Cmip | -0.40044 | 0.000414 | 1 |
| Bach1 | -0.40067 | 4.83E-07 | 0.01558 |
| Rps6ka3 | -0.40155 | 0.000169 | 1 |
| Tacc1 | -0.40161 | 9.73E-05 | 1 |
| Selenop | -0.40177 | 0.017728 | 1 |
| Kmt2c | -0.40258 | 3.22E-05 | 1 |
| Scarb1 | -0.40276 | 1.29E-06 | 0.041496 |
| Myadm | -0.40348 | 0.000288 | 1 |
| Rps27rt | -0.40379 | 4.02E-06 | 0.129782 |
| 6430548M | -0.40383 | 0.000189 | 1 |
| Lmo4 | -0.40431 | 0.022835 | 1 |
| Pcnx | -0.40548 | 2.8E-06 | 0.090312 |
| Nup210 | -0.40554 | 2.23E-05 | 0.720204 |
| Ggnbp2 | -0.40563 | 3.21E-06 | 0.103549 |
| Qk | -0.4063 | 0.000746 | 1 |
| Zmiz1 | -0.40672 | 0.001365 | 1 |
| Ccnd3 | -0.40725 | 4.57E-07 | 0.014749 |
| Rel | -0.40813 | 0.000212 | 1 |
| Gpr132 | -0.40986 | 9.74E-05 | 1 |
| Fbxl17 | -0.41065 | 0.000581 | 1 |
| Lrrc8c | -0.4113 | 0.002493 | 1 |
| Mrpl33 | -0.41301 | 1.96E-06 | 0.063281 |
| Mob1a | -0.41327 | 3.49E-07 | 0.011273 |
| Alcam | -0.41371 | 0.001661 | 1 |
| Irak3 | -0.41468 | 0.008231 | 1 |
| Nuak2 | -0.41479 | 0.000668 | 1 |
| Ptp4a2 | -0.41528 | 2.8E-06 | 0.090383 |
| Ncf2 | -0.41548 | 1.88E-07 | 0.006073 |
| Eps15 | -0.41701 | 0.000647 | 1 |
| Cybb | -0.42022 | 2.62E-11 | 8.47E-07 |
| Dse | -0.42046 | 0.171014 | 1 |

| | | | |
|---|---|---|---|
| Tm9sf4 | -0.4205 | 1.89E-05 | 0.611431 |
| Pafah1b1 | -0.42055 | 1.04E-05 | 0.336379 |
| Dusp5 | -0.42075 | 2.55E-07 | 0.008236 |
| I830077J0: | -0.42101 | 0.001777 | 1 |
| Ifitm3 | -0.42165 | 2.22E-11 | 7.16E-07 |
| Cdc42se1 | -0.42179 | 1.98E-06 | 0.063861 |
| Insyn2b | -0.42235 | 4.04E-05 | 1 |
| Ypel3 | -0.42325 | 9.54E-07 | 0.030802 |
| Usp32 | -0.42437 | 4.46E-05 | 1 |
| Gpatch2l | -0.42503 | 0.000233 | 1 |
| Ahnak | -0.42631 | 0.251593 | 1 |
| Ptpn6 | -0.42777 | 1.31E-09 | 4.24E-05 |
| Ncf4 | -0.42816 | 6.35E-05 | 1 |
| Rsrp1 | -0.42849 | 1.24E-07 | 0.004007 |
| Htra2 | -0.4288 | 0.001939 | 1 |
| Tnip1 | -0.42884 | 0.008722 | 1 |
| Traf3 | -0.42895 | 0.001934 | 1 |
| Rara | -0.429 | 2.64E-06 | 0.0852 |
| Ptpn22 | -0.43006 | 0.000713 | 1 |
| Map3k8 | -0.43007 | 0.00066 | 1 |
| Ddx17 | -0.43041 | 2.93E-05 | 0.94648 |
| Ier2 | -0.43063 | 1.21E-06 | 0.03892 |
| Vps13b | -0.43166 | 8.87E-05 | 1 |
| Fam111a | -0.43188 | 1.96E-05 | 0.633814 |
| Clip1 | -0.43194 | 2.62E-05 | 0.845817 |
| Nfe2 | -0.43345 | 2.76E-07 | 0.008906 |
| Gpx1 | -0.4336 | 5.88E-12 | 1.9E-07 |
| Tmem176a | -0.43378 | 0.038691 | 1 |
| Rbm5 | -0.43402 | 1.11E-06 | 0.035947 |
| Stk4 | -0.43515 | 2.58E-06 | 0.083424 |
| Scp2 | -0.43578 | 2.27E-08 | 0.000732 |
| Kpna4 | -0.43626 | 3.8E-05 | 1 |
| Abcg3 | -0.43693 | 0.043661 | 1 |
| Tyrobp | -0.43719 | 2.59E-19 | 8.35E-15 |
| Taok1 | -0.4375 | 0.000327 | 1 |
| Slk | -0.438 | 4.72E-06 | 0.152299 |
| Plxnb2 | -0.43804 | 1.05E-08 | 0.000338 |
| Mfsd14b | -0.43848 | 0.00118 | 1 |
| Elf4 | -0.43901 | 9.4E-07 | 0.030336 |
| Nfatc3 | -0.43966 | 0.000483 | 1 |
| Ms4a4c | -0.44107 | 9.86E-05 | 1 |
| Gm48099 | -0.44126 | 0.000509 | 1 |
| Tcf7l2 | -0.44182 | 0.039165 | 1 |
| Zfp36l2 | -0.44391 | 5.68E-06 | 0.183335 |
| Ikbkb | -0.44423 | 6.68E-09 | 0.000216 |
| Diaph2 | -0.44515 | 9.5E-05 | 1 |
| Btg2 | -0.44517 | 0.000319 | 1 |
| Dgkh | -0.44654 | 1.05E-05 | 0.338342 |

| | | | |
|---|---|---|---|
| Arhgap39 | -0.44692 | 0.000434 | 1 |
| Pmaip1 | -0.44829 | 1.08E-05 | 0.350115 |
| Zfp706 | -0.44843 | 1.86E-06 | 0.06007 |
| Atf6 | -0.45198 | 8.69E-06 | 0.280434 |
| Napg | -0.45233 | 9.5E-06 | 0.306749 |
| Csgalnact2 | -0.45254 | 5.12E-08 | 0.001654 |
| Mapk3 | -0.45264 | 3.2E-05 | 1 |
| 2310001H1 | -0.45285 | 2.01E-06 | 0.064971 |
| Aftph | -0.453 | 1.48E-05 | 0.477035 |
| Sh2d1b1 | -0.45337 | 0.00407 | 1 |
| Cdkn1b | -0.45441 | 3.64E-05 | 1 |
| Xylt1 | -0.45445 | 8.5E-08 | 0.002744 |
| Gm2a | -0.45457 | 6E-08 | 0.001936 |
| Stk17b | -0.45568 | 5.88E-05 | 1 |
| Kmt2e | -0.45659 | 9.55E-09 | 0.000308 |
| Gm12185 | -0.45746 | 6.92E-07 | 0.02234 |
| Ccdc192 | -0.45825 | 2.79E-09 | 9.02E-05 |
| Gm34084 | -0.45949 | 0.004611 | 1 |
| Tmem176b | -0.4596 | 0.475555 | 1 |
| Nab1 | -0.45982 | 0.000196 | 1 |
| Cd9 | -0.46005 | 0.005274 | 1 |
| Rapgef1 | -0.46049 | 2.07E-06 | 0.066762 |
| Il16 | -0.46258 | 0.000138 | 1 |
| Lrrk2 | -0.46266 | 0.000134 | 1 |
| Ripk2 | -0.46283 | 0.000704 | 1 |
| Smarca2 | -0.46481 | 0.00017 | 1 |
| Xiap | -0.46511 | 6.03E-06 | 0.194602 |
| Eml6 | -0.46553 | 1.83E-06 | 0.059204 |
| Akap10 | -0.46643 | 2.52E-05 | 0.813624 |
| Arhgap27 | -0.46921 | 5.63E-05 | 1 |
| Il6ra | -0.46956 | 2.23E-09 | 7.2E-05 |
| Gm11808 | -0.47046 | 1.48E-08 | 0.000478 |
| Arhgap4 | -0.4709 | 2.27E-07 | 0.007321 |
| Son | -0.47126 | 3.74E-09 | 0.000121 |
| Nr3c1 | -0.47163 | 1.07E-05 | 0.34703 |
| Ppp1r12a | -0.47199 | 8.83E-10 | 2.85E-05 |
| Prkce | -0.47269 | 1.35E-06 | 0.043683 |
| Sf3b1 | -0.47279 | 2.41E-11 | 7.78E-07 |
| Arhgap45 | -0.47309 | 6.93E-08 | 0.002238 |
| Gk | -0.47392 | 0.026247 | 1 |
| Naip5 | -0.47408 | 2.2E-06 | 0.071105 |
| Cd300lb | -0.47439 | 8.41E-08 | 0.002716 |
| Epsti1 | -0.47664 | 3.56E-10 | 1.15E-05 |
| Sp100 | -0.47703 | 2.74E-09 | 8.86E-05 |
| BC035044 | -0.47722 | 5.15E-11 | 1.66E-06 |
| Ttc7b | -0.47759 | 2.52E-05 | 0.815102 |
| Adrb2 | -0.4781 | 0.000246 | 1 |
| Fry | -0.47955 | 2.68E-05 | 0.865888 |

| | | | |
|---|---|---|---|
| Susd3 | -0.48289 | 5.3E-08 | 0.001711 |
| Gnas | -0.48418 | 6.18E-09 | 0.0002 |
| Ptpro | -0.48444 | 3.16E-07 | 0.010211 |
| Wsb1 | -0.48458 | 2.14E-07 | 0.006896 |
| Plekhm3 | -0.48918 | 1.4E-06 | 0.045319 |
| Klhl5 | -0.48931 | 2.98E-08 | 0.000962 |
| Cdc42ep3 | -0.4907 | 2.61E-09 | 8.43E-05 |
| Arap1 | -0.49252 | 1.86E-07 | 0.00602 |
| Tgfbr3 | -0.49315 | 1.82E-12 | 5.87E-08 |
| Neat1 | -0.4934 | 1.73E-05 | 0.559677 |
| Ehbp1l1 | -0.4938 | 1.49E-07 | 0.004817 |
| Sh2d3c | -0.49381 | 1.7E-07 | 0.005491 |
| Scnn1a | -0.49597 | 7.39E-09 | 0.000238 |
| Cst3 | -0.49697 | 3.65E-16 | 1.18E-11 |
| Tab2 | -0.4972 | 4.16E-09 | 0.000134 |
| Foxn2 | -0.4983 | 1.3E-05 | 0.418864 |
| Pacs1 | -0.49859 | 5.14E-06 | 0.165794 |
| Lyl1 | -0.4988 | 4.69E-08 | 0.001514 |
| Zbtb7a | -0.49893 | 1.33E-07 | 0.0043 |
| C130050O1 | -0.50133 | 5.08E-06 | 0.163998 |
| Xdh | -0.50267 | 6.85E-10 | 2.21E-05 |
| Tln1 | -0.50275 | 2.74E-13 | 8.83E-09 |
| Zbtb20 | -0.50346 | 3.09E-05 | 0.998837 |
| Csf3r | -0.50415 | 1.04E-05 | 0.334588 |
| Sorl1 | -0.50641 | 3.79E-06 | 0.122319 |
| Atg4c | -0.50826 | 1.22E-07 | 0.00393 |
| Cd180 | -0.50925 | 2.88E-06 | 0.09294 |
| Cd48 | -0.50975 | 2.28E-13 | 7.36E-09 |
| Laptm5 | -0.51005 | 3.03E-22 | 9.78E-18 |
| Man1a | -0.51045 | 1.11E-05 | 0.359023 |
| Mob3b | -0.51127 | 1.32E-05 | 0.425513 |
| Pbxip1 | -0.51128 | 5.69E-07 | 0.018378 |
| Ccdc12 | -0.51253 | 1.29E-09 | 4.17E-05 |
| Adgre5 | -0.51306 | 7.44E-10 | 2.4E-05 |
| Aim2 | -0.51555 | 4.23E-06 | 0.136551 |
| Utrn | -0.5158 | 6.05E-08 | 0.001952 |
| Hexb | -0.51582 | 0.213836 | 1 |
| Arid3a | -0.51608 | 2.77E-05 | 0.894167 |
| Clec2g | -0.51879 | 2.41E-12 | 7.8E-08 |
| Tbc1d8 | -0.52038 | 0.000386 | 1 |
| Cdk17 | -0.52153 | 2.88E-09 | 9.31E-05 |
| Sipa1l1 | -0.52228 | 4.2E-05 | 1 |
| Calhm2 | -0.5223 | 4.99E-08 | 0.001612 |
| Arhgap30 | -0.52286 | 2.47E-11 | 7.97E-07 |
| 2410006H1 | -0.5251 | 1.42E-07 | 0.004597 |
| Rubcnl | -0.52651 | 1.45E-07 | 0.004665 |
| Sirpa | -0.52828 | 2.22E-12 | 7.18E-08 |
| Itgax | -0.52971 | 3E-05 | 0.968813 |

| | | | |
|---|---|---|---|
| Gm35853 | -0.53067 | 2.41E-13 | 7.79E-09 |
| Camkk2 | -0.53255 | 2.94E-10 | 9.49E-06 |
| Mtpn | -0.53283 | 6.52E-12 | 2.11E-07 |
| Cebpz | -0.53491 | 0.001098 | 1 |
| Fgfr2 | -0.5359 | 3.3E-09 | 0.000106 |
| Traf1 | -0.53604 | 0.002724 | 1 |
| Gsr | -0.53626 | 1.85E-09 | 5.98E-05 |
| Cyp2ab1 | -0.53839 | 2.91E-14 | 9.41E-10 |
| Stk24 | -0.53947 | 2.15E-07 | 0.006945 |
| Spi1 | -0.54072 | 2.6E-17 | 8.38E-13 |
| Lyz1 | -0.54092 | 0.120595 | 1 |
| Rap1gds1 | -0.54248 | 8.78E-06 | 0.283609 |
| Limd2 | -0.54372 | 5.98E-08 | 0.00193 |
| Selplg | -0.54609 | 1.03E-08 | 0.000334 |
| Etv6 | -0.54629 | 7.28E-10 | 2.35E-05 |
| Atp7a | -0.54635 | 6.79E-05 | 1 |
| Notch2 | -0.54915 | 1.57E-07 | 0.00506 |
| Nfkb1 | -0.54972 | 7.62E-09 | 0.000246 |
| Elmo2 | -0.54973 | 9.19E-07 | 0.029679 |
| Slc8a1 | -0.5526 | 1.6E-06 | 0.051698 |
| Arhgap26 | -0.55302 | 3.36E-06 | 0.10833 |
| Ccdc50 | -0.55453 | 3.71E-07 | 0.011976 |
| Grk2 | -0.55625 | 6.33E-07 | 0.02043 |
| Aldh2 | -0.55649 | 1.03E-07 | 0.003315 |
| Lrrc8d | -0.55838 | 3.38E-08 | 0.00109 |
| Pilrb2 | -0.55895 | 1.38E-05 | 0.444717 |
| Map4k4 | -0.5598 | 3.23E-06 | 0.104312 |
| Tbpl1 | -0.56027 | 0.000155 | 1 |
| Lair1 | -0.56209 | 9.12E-06 | 0.294427 |
| Nrros | -0.56287 | 1.71E-12 | 5.53E-08 |
| Slc9a9 | -0.56739 | 1.63E-05 | 0.526257 |
| Lcp2 | -0.56957 | 4.4E-10 | 1.42E-05 |
| Ier5 | -0.57136 | 3.88E-08 | 0.001252 |
| H3f3a | -0.5716 | 7E-22 | 2.26E-17 |
| Gpr141 | -0.57311 | 2.64E-09 | 8.51E-05 |
| Arhgef3 | -0.57479 | 2.34E-06 | 0.075518 |
| Hps3 | -0.57882 | 9.22E-08 | 0.002978 |
| Hlcs | -0.58325 | 4.37E-06 | 0.141061 |
| Cnn2 | -0.58406 | 1.52E-09 | 4.91E-05 |
| Rnase6 | -0.58443 | 4.71E-09 | 0.000152 |
| Adcy7 | -0.58739 | 1.03E-08 | 0.000332 |
| Asap1 | -0.58801 | 1.98E-09 | 6.41E-05 |
| Lpar6 | -0.58898 | 1.07E-08 | 0.000347 |
| Ms4a6b | -0.58978 | 1.15E-10 | 3.72E-06 |
| Jarid2 | -0.59177 | 1.17E-13 | 3.79E-09 |
| Celf2 | -0.59272 | 1.94E-11 | 6.26E-07 |
| Heg1 | -0.59478 | 1.34E-08 | 0.000434 |
| Lrrfip1 | -0.5953 | 1.39E-08 | 0.000449 |

| | | | |
|---|---|---|---|
| Sparc | -0.59635 | 0.206848 | 1 |
| Marcks | -0.59753 | 3.74E-09 | 0.000121 |
| Vav3 | -0.59879 | 2.63E-07 | 0.008492 |
| Tspan13 | -0.59891 | 3.31E-08 | 0.001068 |
| Slc12a6 | -0.6005 | 3.58E-07 | 0.011572 |
| Ptpn12 | -0.60139 | 1.4E-05 | 0.452569 |
| Plekho2 | -0.60398 | 1.75E-10 | 5.65E-06 |
| L1cam | -0.60409 | 3.73E-14 | 1.2E-09 |
| Ppm1h | -0.60514 | 1.34E-06 | 0.043399 |
| Themis2 | -0.6057 | 6.06E-09 | 0.000196 |
| Eif4g3 | -0.60608 | 1.76E-11 | 5.69E-07 |
| Lcp1 | -0.60657 | 7.29E-21 | 2.35E-16 |
| Fabp4 | -0.60935 | 7.09E-12 | 2.29E-07 |
| Ap1s2 | -0.60956 | 1.91E-11 | 6.16E-07 |
| Cdk8 | -0.61158 | 5.75E-10 | 1.86E-05 |
| Ppp2r5c | -0.61169 | 4.48E-08 | 0.001445 |
| Herc4 | -0.61279 | 1.3E-09 | 4.2E-05 |
| Mctp1 | -0.61944 | 7.1E-10 | 2.29E-05 |
| Gm31814 | -0.62056 | 1E-09 | 3.23E-05 |
| Klf4 | -0.62158 | 5.57E-08 | 0.001797 |
| Ifngr1 | -0.62184 | 3.77E-15 | 1.22E-10 |
| Grap2 | -0.62244 | 4.4E-08 | 0.00142 |
| Gpcpd1 | -0.62283 | 1.98E-07 | 0.006402 |
| Smurf2 | -0.6256 | 0.001222 | 1 |
| Sh3kbp1 | -0.62582 | 5.73E-11 | 1.85E-06 |
| Dipk1a | -0.62808 | 8.92E-09 | 0.000288 |
| Sgk3 | -0.62827 | 1.89E-14 | 6.12E-10 |
| Ms4a6c | -0.63096 | 4.04E-19 | 1.31E-14 |
| Cyp4f16 | -0.63109 | 2.19E-09 | 7.06E-05 |
| Plcg2 | -0.63193 | 4.01E-15 | 1.29E-10 |
| P2ry12 | -0.63402 | 0.574125 | 1 |
| Akna | -0.63566 | 1.05E-12 | 3.38E-08 |
| Abhd12 | -0.63736 | 2.2E-11 | 7.1E-07 |
| Svil | -0.6418 | 5.73E-06 | 0.18497 |
| Rasgrp2 | -0.64565 | 1.28E-11 | 4.13E-07 |
| Ttr | -0.64611 | 4.31E-09 | 0.000139 |
| Tpd52 | -0.64776 | 1.39E-11 | 4.49E-07 |
| Agpat4 | -0.65131 | 2.15E-06 | 0.069469 |
| St3gal4 | -0.6527 | 2.92E-07 | 0.009419 |
| Plagl2 | -0.65422 | 1.07E-11 | 3.45E-07 |
| Cblb | -0.65451 | 2.89E-05 | 0.932659 |
| Ccdc88a | -0.65549 | 4.04E-11 | 1.3E-06 |
| Il31ra | -0.65867 | 1.5E-06 | 0.048441 |
| Rb1 | -0.65967 | 1.27E-06 | 0.040884 |
| Arhgef37 | -0.66384 | 1.11E-08 | 0.000359 |
| Klra2 | -0.66705 | 2.4E-15 | 7.75E-11 |
| Nfe2l2 | -0.66886 | 4.82E-16 | 1.56E-11 |
| Lyn | -0.67068 | 1.49E-22 | 4.8E-18 |

| | | | |
|---|---|---|---|
| Tiam2 | -0.6709 | 3.3E-10 | 1.07E-05 |
| Rasgrp4 | -0.67167 | 8.91E-15 | 2.88E-10 |
| Nod1 | -0.67201 | 1.11E-07 | 0.003581 |
| Anp32a | -0.67201 | 7.14E-15 | 2.31E-10 |
| Tet2 | -0.67299 | 4.38E-08 | 0.001413 |
| Ttc7 | -0.68216 | 5.63E-14 | 1.82E-09 |
| Rap1b | -0.68255 | 1.7E-12 | 5.47E-08 |
| Acss1 | -0.68541 | 9.34E-17 | 3.02E-12 |
| Foxp1 | -0.6877 | 6.59E-13 | 2.13E-08 |
| Nav1 | -0.69273 | 6.55E-14 | 2.11E-09 |
| Ccm2 | -0.69667 | 1.64E-07 | 0.005295 |
| Rps27 | -0.69912 | 3.28E-28 | 1.06E-23 |
| Fgd4 | -0.70104 | 2.71E-09 | 8.75E-05 |
| Tgfbr1 | -0.70172 | 6.8E-08 | 0.002194 |
| Raf1 | -0.7034 | 2.27E-11 | 7.32E-07 |
| Smc6 | -0.70532 | 6.12E-08 | 0.001976 |
| Mirt1 | -0.70781 | 9.57E-16 | 3.09E-11 |
| Tmem131 | -0.71052 | 1.28E-11 | 4.13E-07 |
| Gcnt2 | -0.71225 | 9.42E-14 | 3.04E-09 |
| Smpdl3b | -0.71419 | 9E-06 | 0.290656 |
| Metrnl | -0.71734 | 6.18E-15 | 1.99E-10 |
| Ppp1cb | -0.71744 | 1.52E-11 | 4.9E-07 |
| Rasa3 | -0.71851 | 6.72E-15 | 2.17E-10 |
| Tmem164 | -0.71975 | 1.48E-12 | 4.77E-08 |
| Slc46a3 | -0.72205 | 2.52E-19 | 8.12E-15 |
| Gm21188 | -0.72289 | 5.53E-17 | 1.79E-12 |
| Vti1a | -0.72632 | 1.61E-12 | 5.21E-08 |
| Ifitm2 | -0.72709 | 2.28E-13 | 7.36E-09 |
| Dusp6 | -0.7318 | 8.43E-12 | 2.72E-07 |
| Itgav | -0.73231 | 3.41E-10 | 1.1E-05 |
| Arl5c | -0.73411 | 1.31E-11 | 4.22E-07 |
| Naip6 | -0.73434 | 5.68E-10 | 1.84E-05 |
| Ppp2r5a | -0.73475 | 1.21E-16 | 3.92E-12 |
| Gsap | -0.73665 | 7.08E-10 | 2.29E-05 |
| Tlr7 | -0.73795 | 1.08E-12 | 3.49E-08 |
| Stat5b | -0.74347 | 5.16E-11 | 1.67E-06 |
| Sirpb1b | -0.7445 | 2.08E-14 | 6.73E-10 |
| Ywhaz | -0.74687 | 5.86E-26 | 1.89E-21 |
| Otulinl | -0.74875 | 4.99E-19 | 1.61E-14 |
| Pecam1 | -0.74984 | 3.17E-10 | 1.02E-05 |
| Flna | -0.75063 | 1.37E-20 | 4.43E-16 |
| Nfam1 | -0.75145 | 1.7E-22 | 5.48E-18 |
| Msn | -0.75227 | 7.73E-25 | 2.5E-20 |
| Gm9733 | -0.75381 | 1.42E-05 | 0.458003 |
| Cdk14 | -0.75515 | 1.56E-12 | 5.05E-08 |
| F13a1 | -0.75876 | 0.595464 | 1 |
| Fyn | -0.76013 | 1.78E-09 | 5.75E-05 |
| Add3 | -0.76088 | 8.16E-14 | 2.63E-09 |

| | | | |
|---|---|---|---|
| Trem3 | -0.76143 | 2.45E-11 | 7.9E-07 |
| Rassf5 | -0.76337 | 4.91E-14 | 1.59E-09 |
| Mef2a | -0.76514 | 8.24E-17 | 2.66E-12 |
| Csf1r | -0.76975 | 3.17E-20 | 1.02E-15 |
| Nupr1 | -0.77176 | 1.69E-12 | 5.45E-08 |
| Dok3 | -0.77188 | 3.21E-16 | 1.04E-11 |
| Dock8 | -0.77798 | 2.69E-21 | 8.69E-17 |
| Elmo1 | -0.77899 | 2.6E-16 | 8.4E-12 |
| Fmnl1 | -0.78105 | 1.38E-23 | 4.44E-19 |
| Dgkg | -0.78393 | 9.65E-18 | 3.11E-13 |
| Lsp1 | -0.7874 | 6.84E-27 | 2.21E-22 |
| Dnah12 | -0.78871 | 3.2E-15 | 1.03E-10 |
| Sirpb1a | -0.7892 | 1.96E-10 | 6.32E-06 |
| Runx2 | -0.79115 | 1.12E-15 | 3.62E-11 |
| Map3k14 | -0.79159 | 6.29E-15 | 2.03E-10 |
| Clec4a1 | -0.79468 | 1.55E-18 | 5.02E-14 |
| Gm5150 | -0.79504 | 1.45E-11 | 4.69E-07 |
| Hip1 | -0.80381 | 1.99E-10 | 6.41E-06 |
| Napsa | -0.80959 | 2.18E-24 | 7.05E-20 |
| Gm36161 | -0.81009 | 2.82E-17 | 9.12E-13 |
| Ltb | -0.82933 | 2.43E-11 | 7.85E-07 |
| Ighm | -0.83057 | 5.51E-17 | 1.78E-12 |
| Arhgef1 | -0.83492 | 7.33E-21 | 2.37E-16 |
| Nabp1 | -0.83569 | 1.96E-14 | 6.34E-10 |
| Tmcc1 | -0.83584 | 4.22E-13 | 1.36E-08 |
| Cd300c2 | -0.83585 | 3.74E-15 | 1.21E-10 |
| Clec4a3 | -0.83711 | 8.93E-17 | 2.88E-12 |
| Dleu2 | -0.8458 | 5.18E-20 | 1.67E-15 |
| Prr13 | -0.85378 | 3.68E-19 | 1.19E-14 |
| Rgs2 | -0.85468 | 6.1E-16 | 1.97E-11 |
| Id3 | -0.8553 | 1.35E-17 | 4.36E-13 |
| Clec2d | -0.86192 | 2.8E-17 | 9.03E-13 |
| Fam107b | -0.86279 | 5.72E-24 | 1.85E-19 |
| Gm26740 | -0.87096 | 7.34E-20 | 2.37E-15 |
| Malat1 | -0.8759 | 2.79E-23 | 9E-19 |
| Map3k5 | -0.87862 | 8.71E-16 | 2.81E-11 |
| Nkain2 | -0.88321 | 4.37E-29 | 1.41E-24 |
| Mef2c | -0.88449 | 2.94E-15 | 9.5E-11 |
| Itga4 | -0.88458 | 1.83E-17 | 5.92E-13 |
| Mir142hg | -0.88612 | 7.32E-24 | 2.36E-19 |
| Coro1a | -0.88703 | 2.62E-38 | 8.46E-34 |
| Zfp710 | -0.88752 | 8.29E-20 | 2.68E-15 |
| Gm19951 | -0.89014 | 2.43E-16 | 7.84E-12 |
| Ccdc88c | -0.8903 | 2.49E-18 | 8.05E-14 |
| Ikzf1 | -0.89322 | 7.92E-17 | 2.56E-12 |
| Bin2 | -0.90201 | 1.41E-23 | 4.55E-19 |
| Hfe | -0.90202 | 5.92E-16 | 1.91E-11 |
| Fam129a | -0.90858 | 2.14E-19 | 6.91E-15 |

| | | | |
|---|---|---|---|
| Stk38 | -0.91672 | 1.99E-25 | 6.43E-21 |
| Hck | -0.92346 | 8.77E-25 | 2.83E-20 |
| Nxpe4 | -0.93278 | 4.41E-24 | 1.42E-19 |
| Skint3 | -0.93482 | 1.19E-15 | 3.84E-11 |
| Camk2d | -0.93907 | 1.78E-18 | 5.75E-14 |
| Ptprc | -0.94995 | 4.74E-33 | 1.53E-28 |
| Nfkbiz | -0.95325 | 1.8E-12 | 5.81E-08 |
| Sema4d | -0.96043 | 7.18E-23 | 2.32E-18 |
| Serpinb2 | -0.96339 | 7.18E-23 | 2.32E-18 |
| Arhgef10l | -0.9642 | 1.25E-21 | 4.04E-17 |
| Prkcb | -0.96538 | 1.95E-18 | 6.29E-14 |
| Cyth3 | -0.96893 | 3.09E-17 | 9.96E-13 |
| Krt80 | -0.97187 | 9.04E-17 | 2.92E-12 |
| Rap1a | -0.97846 | 1.78E-30 | 5.73E-26 |
| Pip4k2a | -0.98232 | 1.99E-22 | 6.41E-18 |
| Myo1f | -0.99971 | 1.97E-33 | 6.38E-29 |
| Ptk2b | -1.00172 | 1.08E-20 | 3.48E-16 |
| Spata13 | -1.00891 | 4.6E-16 | 1.48E-11 |
| Dock2 | -1.01103 | 1.77E-27 | 5.72E-23 |
| Foxn3 | -1.01208 | 1.48E-18 | 4.79E-14 |
| Ms4a4a | -1.01578 | 2.18E-16 | 7.05E-12 |
| Slc44a2 | -1.01626 | 2.54E-29 | 8.19E-25 |
| Pik3ap1 | -1.01729 | 3.34E-20 | 1.08E-15 |
| Maml3 | -1.02775 | 2.74E-17 | 8.85E-13 |
| Tnfrsf1b | -1.03809 | 7.31E-26 | 2.36E-21 |
| Ear2 | -1.04483 | 5.21E-11 | 1.68E-06 |
| Dock5 | -1.04836 | 5.83E-20 | 1.88E-15 |
| Ripor2 | -1.0496 | 8.96E-12 | 2.89E-07 |
| Samsn1 | -1.05546 | 9.16E-23 | 2.96E-18 |
| Ceacam1 | -1.05701 | 1.23E-22 | 3.96E-18 |
| Zfyve9 | -1.05902 | 1.62E-23 | 5.23E-19 |
| Lst1 | -1.0627 | 1.69E-35 | 5.46E-31 |
| S1pr5 | -1.06466 | 5.88E-24 | 1.9E-19 |
| Ankrd44 | -1.0666 | 2.11E-23 | 6.81E-19 |
| Fgr | -1.07056 | 1.92E-27 | 6.2E-23 |
| Bcl6 | -1.07857 | 1E-21 | 3.23E-17 |
| Mbp | -1.07982 | 5.02E-22 | 1.62E-17 |
| Klf2 | -1.08464 | 9.21E-24 | 2.97E-19 |
| Pag1 | -1.10772 | 3.18E-12 | 1.03E-07 |
| Cd244a | -1.1143 | 3.73E-28 | 1.2E-23 |
| Spn | -1.12777 | 7.98E-19 | 2.58E-14 |
| Prkch | -1.13812 | 1.02E-24 | 3.31E-20 |
| Sat1 | -1.1441 | 5.36E-23 | 1.73E-18 |
| Ifitm6 | -1.1468 | 1.71E-26 | 5.53E-22 |
| Nadk | -1.1473 | 3.68E-34 | 1.19E-29 |
| Ptprj | -1.15706 | 2.86E-23 | 9.23E-19 |
| Ldlrad3 | -1.1606 | 4.62E-27 | 1.49E-22 |
| Pparg | -1.16734 | 1.02E-23 | 3.28E-19 |

| | | | |
|---|---|---|---|
| Cd300a | -1.17538 | 2.19E-34 | 7.07E-30 |
| Myo1g | -1.20861 | 6.11E-27 | 1.97E-22 |
| Fyb | -1.21373 | 1.88E-36 | 6.06E-32 |
| Ssh2 | -1.22099 | 3.05E-28 | 9.84E-24 |
| Abi3 | -1.22647 | 8.43E-31 | 2.72E-26 |
| Apbb1ip | -1.23368 | 9.49E-35 | 3.06E-30 |
| Fam49a | -1.24109 | 3.3E-27 | 1.07E-22 |
| Il17ra | -1.24203 | 8.46E-37 | 2.73E-32 |
| Ptpre | -1.24842 | 9.36E-26 | 3.02E-21 |
| Gm15987 | -1.26678 | 1.57E-30 | 5.07E-26 |
| Cytip | -1.29882 | 2.05E-33 | 6.61E-29 |
| Pot1b | -1.30744 | 5.85E-24 | 1.89E-19 |
| Cyfip2 | -1.34281 | 2.25E-18 | 7.27E-14 |
| Itgal | -1.34611 | 1.88E-30 | 6.08E-26 |
| Treml4 | -1.34634 | 1.04E-23 | 3.36E-19 |
| Slc12a2 | -1.37068 | 9.81E-27 | 3.17E-22 |
| Stap1 | -1.40906 | 2.44E-38 | 7.87E-34 |
| Cd36 | -1.43029 | 6.15E-17 | 1.99E-12 |
| Rap1gap2 | -1.4423 | 2.61E-19 | 8.41E-15 |
| Cd300e | -1.45568 | 2.21E-26 | 7.12E-22 |
| Grk3 | -1.48962 | 1.04E-28 | 3.35E-24 |
| Dusp16 | -1.49659 | 1.52E-17 | 4.91E-13 |
| Nedd9 | -1.51741 | 8.78E-27 | 2.84E-22 |
| Stk10 | -1.5274 | 2.7E-37 | 8.73E-33 |
| Cd300ld | -1.54779 | 3.82E-23 | 1.23E-18 |
| Gngt2 | -1.6334 | 2.11E-31 | 6.8E-27 |
| Pglyrp1 | -1.65274 | 1.8E-27 | 5.82E-23 |
| Eno3 | -1.68341 | 9.97E-27 | 3.22E-22 |
| Bcl2 | -1.69121 | 1.53E-20 | 4.93E-16 |
| Cx3cr1 | -1.72454 | 4.64E-42 | 1.5E-37 |
| Adgre4 | -1.79619 | 5.91E-30 | 1.91E-25 |
| Nr4a1 | -1.8488 | 9.41E-39 | 3.04E-34 |
| Ace | -1.89424 | 1.08E-33 | 3.49E-29 |
| Fah | -2.07008 | 6.64E-65 | 2.14E-60 |
| Pou2f2 | -2.20573 | 1.89E-50 | 6.11E-46 |

### SUPPLEMENTAL TABLE 3. Differentially Regulated Genes in GVHD versus BM Microglia

| Gene | avg_log2FC | p_val | p_val_adj |
|---|---|---|---|
| Cd74 | 4.631676 | 0 | 0 |
| H2-Ab1 | 3.315752 | 0 | 0 |
| H2-Aa | 3.308687 | 0 | 0 |
| H2-Eb1 | 3.032956 | 0 | 0 |
| Apoe | 2.954148 | 2.2E-259 | 7E-255 |
| Ccl2 | 2.36626 | 4.1E-221 | 1.3E-216 |
| Cxcl10 | 2.245663 | 7.3E-80 | 2.36E-75 |
| Cxcl9 | 2.115001 | 5.41E-76 | 1.75E-71 |
| Ifitm3 | 2.013149 | 7E-299 | 2.3E-294 |
| Ccl5 | 1.985545 | 1.64E-77 | 5.31E-73 |
| Gbp2 | 1.962679 | 5.9E-297 | 1.9E-292 |
| Ccl7 | 1.883627 | 9.5E-147 | 3.1E-142 |
| Ccl12 | 1.875255 | 4.9E-144 | 1.6E-139 |
| Ly6a | 1.699295 | 2.9E-169 | 9.4E-165 |
| H2-Q7 | 1.680514 | 2.9E-280 | 9.5E-276 |
| Cd300lf | 1.672374 | 2.9E-289 | 9.4E-285 |
| H2-K1 | 1.669221 | 0 | 0 |
| Iigp1 | 1.627557 | 5.8E-148 | 1.9E-143 |
| H2-T23 | 1.500082 | 6.3E-291 | 2E-286 |
| Fcgr4 | 1.4871 | 3.9E-248 | 1.2E-243 |
| H2-Q4 | 1.469189 | 1.2E-247 | 3.8E-243 |
| Stat1 | 1.460111 | 2.6E-224 | 8.5E-220 |
| Gbp7 | 1.450896 | 3.6E-246 | 1.2E-241 |
| Ccl4 | 1.446982 | 5.46E-64 | 1.76E-59 |
| H2-D1 | 1.436948 | 0 | 0 |
| Fgl2 | 1.412627 | 2.8E-259 | 9.1E-255 |
| Gbp5 | 1.411745 | 1.5E-175 | 5E-171 |
| Ifi27l2a | 1.400547 | 5.3E-161 | 1.7E-156 |
| Ddit4 | 1.391013 | 2.42E-46 | 7.82E-42 |
| Hcar2 | 1.379885 | 1.8E-211 | 5.9E-207 |
| Cd274 | 1.371381 | 1.4E-164 | 4.5E-160 |
| Lgals3bp | 1.368695 | 8.6E-251 | 2.8E-246 |
| H2-Q6 | 1.367432 | 1.7E-218 | 5.3E-214 |
| Cd52 | 1.301768 | 6.2E-199 | 2E-194 |
| Il2rg | 1.29776 | 2.8E-134 | 8.9E-130 |
| Ifi207 | 1.294544 | 9.1E-210 | 2.9E-205 |
| Bst2 | 1.289813 | 3.8E-197 | 1.2E-192 |
| Oasl2 | 1.279411 | 2E-194 | 6.6E-190 |
| Cst7 | 1.271549 | 6.3E-114 | 2E-109 |
| Zbp1 | 1.263794 | 5.7E-235 | 1.8E-230 |
| Ifi30 | 1.254503 | 1.2E-202 | 3.8E-198 |
| Isg15 | 1.245352 | 7.1E-149 | 2.3E-144 |
| Ifi204 | 1.235391 | 3.1E-217 | 1E-212 |
| Irf1 | 1.229177 | 2E-125 | 6.4E-121 |

| | | | |
|---|---|---|---|
| Ctsc | 1.196438 | 5.8E-199 | 1.9E-194 |
| Nlrc5 | 1.183851 | 1.3E-195 | 4.2E-191 |
| Lilrb4a | 1.183265 | 6.4E-111 | 2.1E-106 |
| Cybb | 1.170243 | 2E-140 | 6.6E-136 |
| C4b | 1.168057 | 1.1E-188 | 3.6E-184 |
| Gm4951 | 1.152232 | 2.7E-161 | 8.6E-157 |
| Srgn | 1.141687 | 1.1E-145 | 3.6E-141 |
| Rpl32 | 1.137283 | 4E-248 | 1.3E-243 |
| Rps20 | 1.122064 | 1.3E-257 | 4.1E-253 |
| B2m | 1.119388 | 3.1E-297 | 1E-292 |
| Tap1 | 1.117568 | 1.1E-183 | 3.5E-179 |
| Tspo | 1.1104 | 1.1E-139 | 3.7E-135 |
| Rpsa | 1.095168 | 1.7E-219 | 5.4E-215 |
| Ccl3 | 1.064149 | 2.51E-30 | 8.11E-26 |
| Rnf213 | 1.055791 | 8.5E-123 | 2.7E-118 |
| Irf7 | 1.048852 | 1.9E-151 | 6.3E-147 |
| AW11201C | 1.048045 | 3.5E-120 | 1.1E-115 |
| Fkbp5 | 1.040891 | 4.9E-100 | 1.6E-95 |
| Samhd1 | 1.040362 | 2.6E-119 | 8.3E-115 |
| Clec2d | 1.026459 | 4.9E-143 | 1.6E-138 |
| Rps2 | 1.026277 | 4.8E-209 | 1.5E-204 |
| Cd36 | 1.010312 | 6.2E-151 | 2E-146 |
| Gbp4 | 1.009687 | 9.1E-170 | 2.9E-165 |
| Stat2 | 0.990243 | 3.4E-148 | 1.1E-143 |
| Spint1 | 0.980676 | 2.4E-152 | 7.7E-148 |
| Rps18 | 0.974699 | 9.5E-183 | 3.1E-178 |
| Ifit2 | 0.973332 | 7.68E-97 | 2.48E-92 |
| Slfn2 | 0.97086 | 1.3E-138 | 4.2E-134 |
| Fth1 | 0.955782 | 1.4E-185 | 4.4E-181 |
| Cebpb | 0.953734 | 1.7E-125 | 5.6E-121 |
| Plaur | 0.953622 | 2.72E-84 | 8.77E-80 |
| Psme1 | 0.952682 | 1.8E-154 | 5.8E-150 |
| Axl | 0.952592 | 5.5E-140 | 1.8E-135 |
| Irgm1 | 0.944915 | 6.6E-127 | 2.1E-122 |
| AU020206 | 0.939862 | 6.1E-128 | 2E-123 |
| Nfkbia | 0.933597 | 4.19E-75 | 1.35E-70 |
| Rps12 | 0.931308 | 8.2E-192 | 2.6E-187 |
| Arhgap15 | 0.927296 | 1.83E-32 | 5.92E-28 |
| Lyz2 | 0.919916 | 2.5E-126 | 8.2E-122 |
| Rps24 | 0.919428 | 9.3E-211 | 3E-206 |
| Rplp0 | 0.917267 | 1.7E-184 | 5.5E-180 |
| Il18bp | 0.912049 | 1.2E-168 | 4E-164 |
| Gbp8 | 0.900554 | 7.9E-170 | 2.6E-165 |
| Rps5 | 0.895888 | 4.7E-184 | 1.5E-179 |
| Tnf | 0.89455 | 1.54E-43 | 4.97E-39 |
| Gbp6 | 0.886408 | 7.7E-138 | 2.5E-133 |
| Sp100 | 0.876176 | 4.8E-128 | 1.5E-123 |
| Rps7 | 0.875812 | 3.4E-179 | 1.1E-174 |

| | | | |
|---|---|---|---|
| mt-Atp8 | 0.872948 | 1.9E-147 | 6E-143 |
| Ms4a6c | 0.86972 | 1.4E-76 | 4.52E-72 |
| Rpl19 | 0.863279 | 2.3E-185 | 7.3E-181 |
| Cxcl16 | 0.857896 | 1.02E-69 | 3.29E-65 |
| Rpl10a | 0.854533 | 6.5E-166 | 2.1E-161 |
| Fxyd5 | 0.847605 | 3.5E-104 | 1.1E-99 |
| Il15ra | 0.845558 | 9.3E-130 | 3E-125 |
| Psap | 0.834524 | 1.8E-228 | 5.9E-224 |
| Rps11 | 0.833966 | 6.9E-177 | 2.2E-172 |
| H2-DMa | 0.833954 | 2.5E-116 | 8.2E-112 |
| Pim1 | 0.822244 | 1.5E-123 | 4.9E-119 |
| Csf1 | 0.813464 | 6.65E-83 | 2.15E-78 |
| Gbp9 | 0.810874 | 4E-114 | 1.3E-109 |
| Cd83 | 0.807978 | 3.76E-48 | 1.21E-43 |
| H2-T22 | 0.804287 | 1.4E-107 | 4.7E-103 |
| Rpl13 | 0.800608 | 4.3E-184 | 1.4E-179 |
| Rps26 | 0.797089 | 3.7E-132 | 1.2E-127 |
| Rpl27a | 0.796682 | 4.8E-162 | 1.6E-157 |
| Socs1 | 0.795728 | 2.61E-86 | 8.43E-82 |
| Psmb8 | 0.7929 | 1.9E-108 | 6.2E-104 |
| Prdx5 | 0.787641 | 1.3E-88 | 4.21E-84 |
| Rps8 | 0.786649 | 2E-160 | 6.6E-156 |
| Rpl12 | 0.783156 | 8.7E-126 | 2.8E-121 |
| Rps15a | 0.781702 | 8.2E-155 | 2.6E-150 |
| Rpl23 | 0.778918 | 2.5E-152 | 7.9E-148 |
| Rpl17 | 0.777323 | 6.1E-142 | 2E-137 |
| Rpl14 | 0.768931 | 4.6E-113 | 1.5E-108 |
| Ifit3 | 0.764504 | 1.94E-90 | 6.25E-86 |
| mt-Co1 | 0.761493 | 2.3E-136 | 7.5E-132 |
| Rps27a | 0.759381 | 8E-164 | 2.6E-159 |
| Rpl24 | 0.758691 | 1.4E-124 | 4.6E-120 |
| Rps13 | 0.755327 | 4.2E-140 | 1.4E-135 |
| Arid5b | 0.740637 | 3.54E-44 | 1.14E-39 |
| Ifi211 | 0.740093 | 1.5E-125 | 4.7E-121 |
| Psme2 | 0.738871 | 5.45E-87 | 1.76E-82 |
| Ucp2 | 0.737073 | 4.14E-68 | 1.34E-63 |
| Rps16 | 0.735663 | 4.7E-137 | 1.5E-132 |
| Rtp4 | 0.735518 | 5.54E-90 | 1.79E-85 |
| Ccdc86 | 0.734403 | 7.7E-100 | 2.47E-95 |
| Atp8a1 | 0.730706 | 1.41E-86 | 4.57E-82 |
| Ctss | 0.726126 | 6E-221 | 1.9E-216 |
| Rpl35 | 0.725271 | 4E-118 | 1.3E-113 |
| Rack1 | 0.723405 | 7.5E-107 | 2.4E-102 |
| Rps6 | 0.721232 | 1.25E-95 | 4.05E-91 |
| Rps19 | 0.719262 | 2.7E-129 | 8.8E-125 |
| Gas5 | 0.71803 | 1.52E-87 | 4.9E-83 |
| Rps10 | 0.716517 | 2.5E-138 | 8.2E-134 |
| Tor3a | 0.713377 | 7.53E-83 | 2.43E-78 |

| | | | |
|---|---|---|---|
| Vcam1 | 0.710246 | 8.44E-82 | 2.72E-77 |
| Rbm3 | 0.710211 | 4.37E-93 | 1.41E-88 |
| Bcl2a1a | 0.707149 | 6.17E-90 | 1.99E-85 |
| Rpl26 | 0.705508 | 3.1E-121 | 1E-116 |
| Parp14 | 0.704187 | 6.98E-70 | 2.25E-65 |
| Tpt1 | 0.701956 | 2.4E-135 | 7.9E-131 |
| Igtp | 0.701926 | 7.82E-64 | 2.53E-59 |
| Rpl28 | 0.701777 | 6.5E-118 | 2.1E-113 |
| Nme2 | 0.7013 | 1.43E-84 | 4.62E-80 |
| Bcl2a1b | 0.698919 | 2.17E-76 | 7E-72 |
| Slfn5 | 0.695787 | 5.03E-85 | 1.62E-80 |
| Rpl18a | 0.695488 | 1.1E-137 | 3.5E-133 |
| Siglecf | 0.695396 | 3.73E-67 | 1.2E-62 |
| Ly86 | 0.693107 | 6.3E-121 | 2E-116 |
| mt-Co2 | 0.6931 | 3.2E-112 | 1E-107 |
| Bcl2a1d | 0.689157 | 6.3E-103 | 2.02E-98 |
| Rpl22l1 | 0.688197 | 1.76E-77 | 5.67E-73 |
| Gbp3 | 0.684802 | 7.7E-114 | 2.5E-109 |
| Id2 | 0.684458 | 2.1E-57 | 6.78E-53 |
| Ccnd3 | 0.683708 | 1.13E-32 | 3.64E-28 |
| Lap3 | 0.683495 | 1.51E-72 | 4.87E-68 |
| Pkm | 0.678331 | 3.89E-50 | 1.26E-45 |
| Rpl27 | 0.677376 | 2.1E-100 | 6.63E-96 |
| Rpl29 | 0.676322 | 1.3E-104 | 4.3E-100 |
| Eif2ak2 | 0.673275 | 2.27E-74 | 7.34E-70 |
| Herc6 | 0.672027 | 3E-53 | 9.68E-49 |
| Mif | 0.672007 | 2.75E-43 | 8.87E-39 |
| Slamf8 | 0.667789 | 1.7E-84 | 5.48E-80 |
| Rpl11 | 0.666305 | 1.7E-121 | 5.6E-117 |
| Rpl15 | 0.665657 | 6.8E-102 | 2.2E-97 |
| Sgip1 | 0.664556 | 8.02E-77 | 2.59E-72 |
| Socs2 | 0.662734 | 1.1E-116 | 3.6E-112 |
| Rpl13a | 0.659035 | 2E-100 | 6.44E-96 |
| Ier3 | 0.658685 | 2.97E-29 | 9.59E-25 |
| Ifi213 | 0.655639 | 1.35E-86 | 4.36E-82 |
| Rps23 | 0.652719 | 1.1E-113 | 3.7E-109 |
| Xaf1 | 0.651004 | 1.11E-88 | 3.57E-84 |
| Il1b | 0.648757 | 1.16E-32 | 3.76E-28 |
| Rplp2 | 0.6466 | 3.4E-103 | 1.08E-98 |
| Ly6e | 0.645646 | 8.5E-125 | 2.7E-120 |
| Rpl37 | 0.645536 | 3E-112 | 9.5E-108 |
| Il1a | 0.641159 | 2.67E-31 | 8.61E-27 |
| Phf11b | 0.63923 | 2.01E-72 | 6.49E-68 |
| Dna2 | 0.638955 | 2.95E-95 | 9.53E-91 |
| Csf2rb | 0.635706 | 3.51E-60 | 1.13E-55 |
| Slc11a1 | 0.6337 | 4.96E-71 | 1.6E-66 |
| Rpl9 | 0.629875 | 5.4E-103 | 1.75E-98 |
| Npc2 | 0.629181 | 7.2E-109 | 2.3E-104 |

| | | | |
|---|---|---|---|
| Oas1a | 0.625228 | 5.9E-74 | 1.91E-69 |
| Rpl39 | 0.624107 | 2.2E-105 | 7.2E-101 |
| Selenow | 0.624022 | 3.54E-72 | 1.14E-67 |
| Rpl36 | 0.623959 | 4.04E-92 | 1.3E-87 |
| Naaa | 0.623671 | 2.15E-62 | 6.94E-58 |
| Oasl1 | 0.622786 | 3.43E-79 | 1.11E-74 |
| Rps4x | 0.622494 | 3.8E-107 | 1.2E-102 |
| Rpl3 | 0.620644 | 2.2E-106 | 7.2E-102 |
| Tgtp2 | 0.615951 | 8.4E-100 | 2.7E-95 |
| Trim30a | 0.612562 | 4.23E-66 | 1.37E-61 |
| Rpl41 | 0.611855 | 2.4E-100 | 7.73E-96 |
| Rpl6 | 0.611618 | 1.7E-104 | 5.5E-100 |
| H2-DMb1 | 0.610001 | 9.09E-35 | 2.93E-30 |
| Ifi206 | 0.608203 | 4.56E-93 | 1.47E-88 |
| mt-Co3 | 0.604885 | 1.49E-93 | 4.83E-89 |
| Ifi47 | 0.602366 | 9.56E-78 | 3.09E-73 |
| Psmb9 | 0.595356 | 1.53E-69 | 4.93E-65 |
| Rpl35a | 0.595152 | 9.9E-104 | 3.2E-99 |
| Rtcb | 0.594767 | 9.17E-58 | 2.96E-53 |
| Rpl34 | 0.594724 | 1.36E-96 | 4.39E-92 |
| Dusp1 | 0.594444 | 1.58E-18 | 5.09E-14 |
| Rpl36a | 0.594228 | 3.92E-70 | 1.26E-65 |
| Cd72 | 0.59264 | 1.3E-43 | 4.21E-39 |
| Rpl30 | 0.591658 | 1.5E-102 | 4.75E-98 |
| Rpl18 | 0.590741 | 1.18E-95 | 3.82E-91 |
| Eef1b2 | 0.587563 | 2.57E-72 | 8.29E-68 |
| Coro2a | 0.585648 | 1.22E-54 | 3.94E-50 |
| Rps9 | 0.583611 | 1.2E-102 | 3.74E-98 |
| Csf2ra | 0.583051 | 1.34E-58 | 4.32E-54 |
| Tsc22d3 | 0.577174 | 1.62E-24 | 5.23E-20 |
| Efhd2 | 0.576475 | 1.87E-66 | 6.05E-62 |
| Gm26917 | 0.573047 | 1.58E-61 | 5.12E-57 |
| Eef1a1 | 0.570923 | 3.8E-127 | 1.2E-122 |
| Gpr65 | 0.570493 | 1.09E-67 | 3.51E-63 |
| Rps14 | 0.564747 | 1.4E-84 | 4.51E-80 |
| Rgs1 | 0.563393 | 2.66E-16 | 8.6E-12 |
| Nampt | 0.563249 | 6.86E-62 | 2.21E-57 |
| Fau | 0.561115 | 1.2E-111 | 3.9E-107 |
| Socs3 | 0.560591 | 2.73E-29 | 8.83E-25 |
| 2410006H | 0.553591 | 6E-47 | 1.94E-42 |
| Rpl8 | 0.551779 | 1.08E-74 | 3.49E-70 |
| Ms4a4c | 0.54919 | 5.77E-86 | 1.86E-81 |
| Hacd2 | 0.548755 | 3.59E-47 | 1.16E-42 |
| Gm43305 | 0.545413 | 1.77E-11 | 5.72E-07 |
| Rps3 | 0.543702 | 2.22E-83 | 7.18E-79 |
| Epsti1 | 0.542933 | 1.49E-43 | 4.81E-39 |
| Itgax | 0.542787 | 7.77E-72 | 2.51E-67 |
| Rpl7a | 0.542633 | 1.42E-82 | 4.57E-78 |

| | | | |
|---|---|---|---|
| Ldha | 0.54197 | 2.33E-39 | 7.52E-35 |
| Cd69 | 0.541957 | 3.81E-40 | 1.23E-35 |
| Psmb10 | 0.540451 | 1.18E-51 | 3.8E-47 |
| C3ar1 | 0.539311 | 3.61E-50 | 1.16E-45 |
| Tlr2 | 0.537563 | 1.22E-22 | 3.95E-18 |
| Gm42601 | 0.537036 | 1.31E-51 | 4.22E-47 |
| Etv6 | 0.535861 | 8.64E-51 | 2.79E-46 |
| Ifih1 | 0.533715 | 4.87E-38 | 1.57E-33 |
| Rps28 | 0.531313 | 2.73E-72 | 8.82E-68 |
| Rad51b | 0.530893 | 9.69E-39 | 3.13E-34 |
| Zbtb16 | 0.528596 | 2.27E-47 | 7.32E-43 |
| Apobec3 | 0.527082 | 1.18E-55 | 3.8E-51 |
| Rps25 | 0.526598 | 7.03E-57 | 2.27E-52 |
| Parp9 | 0.526545 | 6.83E-60 | 2.2E-55 |
| Rplp1 | 0.525387 | 1.51E-95 | 4.86E-91 |
| Gatm | 0.52226 | 5.07E-33 | 1.64E-28 |
| Rpl31 | 0.518145 | 3.38E-53 | 1.09E-48 |
| Ier2 | 0.517808 | 5.77E-30 | 1.86E-25 |
| Rab43 | 0.514514 | 5.32E-53 | 1.72E-48 |
| Sp110 | 0.514218 | 8.59E-51 | 2.77E-46 |
| Rpl21 | 0.513965 | 2.81E-88 | 9.07E-84 |
| Usp18 | 0.513041 | 3.79E-48 | 1.22E-43 |
| Egr1 | 0.510878 | 1.68E-24 | 5.43E-20 |
| Rps3a1 | 0.510734 | 9.29E-82 | 3E-77 |
| Gm46224 | 0.509773 | 4.22E-43 | 1.36E-38 |
| Cyp4f18 | 0.508032 | 2.13E-88 | 6.89E-84 |
| Myc | 0.506748 | 1.03E-31 | 3.32E-27 |
| Zfas1 | 0.502367 | 1.37E-69 | 4.44E-65 |
| Chka | 0.502202 | 1.03E-42 | 3.32E-38 |
| Rpl23a | 0.501613 | 5.48E-51 | 1.77E-46 |
| Grn | 0.500038 | 4.47E-95 | 1.44E-90 |
| Eif3f | 0.499968 | 1.15E-51 | 3.71E-47 |
| Per1 | 0.496775 | 1.45E-42 | 4.68E-38 |
| Cfb | 0.495975 | 1.18E-68 | 3.81E-64 |
| Apobec1 | 0.49489 | 2.01E-37 | 6.48E-33 |
| Tnfsf13b | 0.493836 | 1.99E-45 | 6.41E-41 |
| Cstb | 0.489776 | 3.2E-37 | 1.03E-32 |
| Fmnl2 | 0.489549 | 5.99E-43 | 1.94E-38 |
| Rpl37a | 0.488962 | 1.47E-69 | 4.74E-65 |
| Ifi209 | 0.488665 | 2.32E-66 | 7.51E-62 |
| Capg | 0.488381 | 1.44E-45 | 4.64E-41 |
| Rnf169 | 0.486985 | 5.9E-23 | 1.9E-18 |
| Pitpna | 0.484431 | 8.05E-45 | 2.6E-40 |
| Slc2a6 | 0.482406 | 8.95E-76 | 2.89E-71 |
| mt-Atp6 | 0.481501 | 5.68E-53 | 1.84E-48 |
| Ranbp2 | 0.477771 | 1.03E-38 | 3.34E-34 |
| Fcgr2b | 0.475718 | 2.94E-39 | 9.49E-35 |
| Rpl10 | 0.475526 | 1.44E-73 | 4.65E-69 |

| | | | |
|---|---|---|---|
| Znfx1 | 0.474485 | 1.3E-44 | 4.19E-40 |
| Treml2 | 0.472048 | 1.58E-77 | 5.1E-73 |
| Rpl5 | 0.469765 | 3.36E-46 | 1.09E-41 |
| Ccnl1 | 0.469136 | 4.71E-43 | 1.52E-38 |
| Tagln2 | 0.461906 | 6.57E-36 | 2.12E-31 |
| Gm2a | 0.460983 | 2.01E-43 | 6.5E-39 |
| Cpd | 0.459876 | 8.88E-31 | 2.87E-26 |
| Clic4 | 0.45981 | 1.86E-44 | 6E-40 |
| Icam1 | 0.459378 | 4.81E-19 | 1.55E-14 |
| Gpi1 | 0.458718 | 1.21E-36 | 3.91E-32 |
| Lat2 | 0.458001 | 1.12E-44 | 3.6E-40 |
| Ccrl2 | 0.456222 | 7.48E-34 | 2.42E-29 |
| Slc31a2 | 0.456196 | 8.27E-39 | 2.67E-34 |
| Npm1 | 0.454669 | 1.12E-41 | 3.62E-37 |
| Lpp | 0.451481 | 1E-34 | 3.24E-30 |
| Fcgr1 | 0.450788 | 2.4E-48 | 7.75E-44 |
| Gm33370 | 0.449228 | 8.53E-53 | 2.75E-48 |
| Rps15 | 0.448111 | 3.77E-45 | 1.22E-40 |
| Atp6v0c | 0.447225 | 1.23E-41 | 3.96E-37 |
| Cox8a | 0.446042 | 3.58E-44 | 1.16E-39 |
| Ctsh | 0.445537 | 1.73E-55 | 5.58E-51 |
| Rpl7 | 0.445177 | 1.01E-53 | 3.25E-49 |
| Scimp | 0.44364 | 4.91E-61 | 1.59E-56 |
| Pik3ip1 | 0.439971 | 2.01E-33 | 6.5E-29 |
| Rpl22 | 0.439343 | 3.02E-50 | 9.75E-46 |
| Ifi35 | 0.435763 | 6.02E-38 | 1.94E-33 |
| Pml | 0.434485 | 5.2E-38 | 1.68E-33 |
| Hsp90ab1 | 0.431976 | 2.02E-34 | 6.51E-30 |
| Trpm2 | 0.431429 | 2.55E-39 | 8.24E-35 |
| Tap2 | 0.428301 | 1.03E-35 | 3.33E-31 |
| Ripor2 | 0.428198 | 4.61E-39 | 1.49E-34 |
| Plaat3 | 0.427339 | 1.59E-54 | 5.13E-50 |
| 2210408F2 | 0.426988 | 8.71E-42 | 2.81E-37 |
| Birc3 | 0.42629 | 2.94E-34 | 9.48E-30 |
| Cyba | 0.426 | 4.82E-66 | 1.56E-61 |
| Zfp36 | 0.424861 | 3.54E-10 | 1.14E-05 |
| Ctsb | 0.423436 | 1.27E-53 | 4.11E-49 |
| Cep85 | 0.423357 | 1.26E-11 | 4.05E-07 |
| Ssr4 | 0.420408 | 6.64E-40 | 2.14E-35 |
| Vps54 | 0.41567 | 1.72E-30 | 5.56E-26 |
| Rpl36al | 0.415559 | 3.38E-38 | 1.09E-33 |
| Rpl38 | 0.415154 | 1.98E-43 | 6.38E-39 |
| B4galt1 | 0.41413 | 8.32E-20 | 2.69E-15 |
| Tpst1 | 0.41181 | 3.21E-60 | 1.04E-55 |
| Serp1 | 0.411707 | 3.42E-24 | 1.1E-19 |
| Gcnt2 | 0.411271 | 3.75E-29 | 1.21E-24 |
| Phf11d | 0.410474 | 3.47E-41 | 1.12E-36 |
| H2-M3 | 0.407967 | 1.66E-40 | 5.37E-36 |

| | | | |
|---|---|---|---|
| Cited2 | 0.405673 | 4.2E-20 | 1.36E-15 |
| Sdc3 | 0.405385 | 5.96E-47 | 1.92E-42 |
| Creb5 | 0.399635 | 4.6E-26 | 1.49E-21 |
| 1110038B: | 0.398445 | 5.38E-41 | 1.74E-36 |
| Gpr84 | 0.397897 | 3.38E-06 | 0.109133 |
| Smim3 | 0.395422 | 1.83E-43 | 5.91E-39 |
| Eif5a | 0.393564 | 1.56E-28 | 5.03E-24 |
| Eef1g | 0.390073 | 2.07E-30 | 6.68E-26 |
| Zup1 | 0.389259 | 4.69E-27 | 1.51E-22 |
| Ifit1 | 0.388288 | 2.1E-39 | 6.79E-35 |
| Hspe1 | 0.388027 | 1.48E-32 | 4.77E-28 |
| Gnl3 | 0.387654 | 9.16E-34 | 2.96E-29 |
| Vim | 0.387108 | 2.84E-23 | 9.18E-19 |
| Gm12216 | 0.386652 | 3.74E-51 | 1.21E-46 |
| Tpi1 | 0.386471 | 5.81E-18 | 1.88E-13 |
| Pvt1 | 0.386179 | 2.81E-33 | 9.06E-29 |
| Itgal | 0.384791 | 4.96E-58 | 1.6E-53 |
| Ctsz | 0.383125 | 6.24E-62 | 2.01E-57 |
| Tapbpl | 0.382223 | 2.38E-43 | 7.69E-39 |
| Nuak2 | 0.382105 | 2.59E-41 | 8.35E-37 |
| Rsad2 | 0.380068 | 3.41E-39 | 1.1E-34 |
| Rasgef1b | 0.379057 | 1.5E-19 | 4.85E-15 |
| Man2a1 | 0.378906 | 3.47E-34 | 1.12E-29 |
| Rps17 | 0.374468 | 8.97E-31 | 2.9E-26 |
| Akr1a1 | 0.374182 | 6.73E-28 | 2.17E-23 |
| Eif4a1 | 0.37409 | 4.08E-32 | 1.32E-27 |
| Zc3hav1 | 0.373315 | 6.04E-27 | 1.95E-22 |
| Cd244a | 0.373156 | 6.55E-60 | 2.12E-55 |
| Morrbid | 0.372832 | 1.49E-34 | 4.81E-30 |
| Iqgap1 | 0.372434 | 5.56E-18 | 1.8E-13 |
| Chd7 | 0.370817 | 6.22E-28 | 2.01E-23 |
| Lgals3 | 0.370574 | 1.75E-29 | 5.64E-25 |
| Arhgap24 | 0.368598 | 6.78E-16 | 2.19E-11 |
| Rpl4 | 0.368358 | 1.4E-27 | 4.52E-23 |
| Ybx1 | 0.367938 | 3.44E-31 | 1.11E-26 |
| Slc31a1 | 0.366908 | 1.38E-32 | 4.46E-28 |
| Trafd1 | 0.365877 | 4.38E-24 | 1.41E-19 |
| Ube2l6 | 0.365365 | 3.25E-46 | 1.05E-41 |
| Gnptab | 0.365292 | 3.22E-37 | 1.04E-32 |
| Ccr1 | 0.365148 | 5.49E-19 | 1.77E-14 |
| Klf6 | 0.364729 | 4.38E-15 | 1.42E-10 |
| Coro7 | 0.364691 | 3.37E-40 | 1.09E-35 |
| Atp5g2 | 0.363726 | 2.11E-32 | 6.82E-28 |
| Rabgap1l | 0.363487 | 0.25711 | 1 |
| Fbxw17 | 0.363124 | 6.46E-48 | 2.09E-43 |
| Tapbp | 0.363087 | 1.12E-26 | 3.62E-22 |
| Sorl1 | 0.362343 | 5.64E-25 | 1.82E-20 |
| Cept1 | 0.362167 | 2.51E-27 | 8.11E-23 |

| | | | |
|---|---|---|---|
| Slc25a3 | 0.361867 | 1.53E-26 | 4.95E-22 |
| Mmp14 | 0.360308 | 3.2E-32 | 1.03E-27 |
| Klf9 | 0.360109 | 3E-40 | 9.67E-36 |
| Map3k8 | 0.35957 | 2.55E-36 | 8.23E-32 |
| Bcl2 | 0.359426 | 0.001874 | 1 |
| Sema4a | 0.359263 | 4.61E-50 | 1.49E-45 |
| Plac8 | 0.357987 | 1.56E-40 | 5.03E-36 |
| mt-Nd1 | 0.357634 | 9.82E-24 | 3.17E-19 |
| Lpl | 0.357473 | 3.33E-19 | 1.07E-14 |
| Camk1d | 0.35738 | 2.09E-28 | 6.75E-24 |
| Msr1 | 0.357072 | 1.01E-57 | 3.27E-53 |
| Cd63 | 0.355991 | 2.48E-19 | 8.01E-15 |
| Eif3h | 0.355182 | 2E-22 | 6.45E-18 |
| Ehd1 | 0.35489 | 5.45E-48 | 1.76E-43 |
| Siglec1 | 0.35202 | 4.46E-53 | 1.44E-48 |
| Klf13 | 0.351948 | 5.76E-21 | 1.86E-16 |
| Rbm47 | 0.35127 | 7.01E-14 | 2.26E-09 |
| Psma5 | 0.351201 | 1.62E-23 | 5.23E-19 |
| Acsl4 | 0.350132 | 9.11E-28 | 2.94E-23 |
| Ly9 | 0.349293 | 1.77E-37 | 5.73E-33 |
| Lacc1 | 0.349214 | 8.11E-26 | 2.62E-21 |
| Rab5if | 0.348381 | 2.56E-36 | 8.26E-32 |
| Rps21 | 0.347884 | 1.64E-40 | 5.28E-36 |
| Glrx | 0.347676 | 8.79E-35 | 2.84E-30 |
| AB124611 | 0.345486 | 1.41E-50 | 4.55E-46 |
| Gm15964 | 0.344374 | 9.1E-48 | 2.94E-43 |
| Prr5l | 0.344356 | 1.13E-37 | 3.65E-33 |
| Dhx58 | 0.343818 | 3.32E-38 | 1.07E-33 |
| Ccnd2 | 0.342518 | 2.16E-62 | 6.98E-58 |
| Prdx1 | 0.342226 | 1.73E-24 | 5.57E-20 |
| Cd40 | 0.342135 | 1.71E-37 | 5.52E-33 |
| Sh3bgrl3 | 0.341876 | 6.61E-30 | 2.13E-25 |
| Uqcrh | 0.340557 | 2.47E-24 | 7.97E-20 |
| Trim30d | 0.340551 | 3.05E-28 | 9.84E-24 |
| Parp12 | 0.338097 | 1.02E-32 | 3.28E-28 |
| Mllt3 | 0.337379 | 3.4E-19 | 1.1E-14 |
| Zc3h7a | 0.336937 | 6.17E-15 | 1.99E-10 |
| C1qa | 0.335173 | 4.72E-71 | 1.52E-66 |
| Lrrfip1 | 0.335014 | 4.42E-21 | 1.43E-16 |
| 1600014C: | 0.334348 | 2.41E-26 | 7.78E-22 |
| Cblb | 0.334297 | 0.000146 | 1 |
| Txn1 | 0.333494 | 8.08E-26 | 2.61E-21 |
| Abr | 0.33277 | 6.13E-21 | 1.98E-16 |
| Ppp1r15a | 0.332723 | 7.32E-22 | 2.36E-17 |
| Ppp1r21 | 0.332705 | 6.21E-17 | 2.01E-12 |
| Hsp90aa1 | 0.331434 | 6.87E-12 | 2.22E-07 |
| Rassf3 | 0.331057 | 7.91E-39 | 2.55E-34 |
| Wwox | 0.32986 | 7.02E-16 | 2.27E-11 |

| | | | |
|---|---|---|---|
| Npnt | 0.329601 | 2.03E-30 | 6.54E-26 |
| Dgkz | 0.32957 | 1.72E-22 | 5.56E-18 |
| B3gnt2 | 0.329422 | 1.02E-24 | 3.28E-20 |
| Xdh | 0.328736 | 7.33E-47 | 2.37E-42 |
| Hnrnpa1 | 0.328446 | 1.84E-26 | 5.95E-22 |
| Sat1 | 0.328198 | 1.1E-15 | 3.55E-11 |
| Csf2rb2 | 0.327223 | 2.87E-24 | 9.26E-20 |
| Chmp4b | 0.32713 | 1.69E-31 | 5.46E-27 |
| Snhg12 | 0.324271 | 1.63E-27 | 5.27E-23 |
| Oas2 | 0.323573 | 1.19E-50 | 3.85E-46 |
| Arid5a | 0.323036 | 7.24E-32 | 2.34E-27 |
| Cd48 | 0.322795 | 9.9E-25 | 3.2E-20 |
| Aldoa | 0.322286 | 5.79E-14 | 1.87E-09 |
| Slfn8 | 0.320265 | 1.05E-21 | 3.37E-17 |
| C1qb | 0.320169 | 2.24E-68 | 7.24E-64 |
| March5 | 0.31995 | 1.09E-33 | 3.51E-29 |
| Itpr1 | 0.319352 | 2.34E-20 | 7.55E-16 |
| Casp4 | 0.319301 | 3.28E-27 | 1.06E-22 |
| Pilra | 0.318337 | 5.15E-23 | 1.66E-18 |
| Phf11a | 0.318277 | 5.73E-43 | 1.85E-38 |
| Naca | 0.318001 | 9.76E-29 | 3.15E-24 |
| Snhg1 | 0.31701 | 1.33E-31 | 4.28E-27 |
| Uqcrq | 0.316558 | 1.22E-22 | 3.95E-18 |
| Rps29 | 0.316128 | 9.49E-37 | 3.06E-32 |
| Kcnk6 | 0.31541 | 3.51E-23 | 1.13E-18 |
| Tmem106a | 0.314781 | 3.41E-29 | 1.1E-24 |
| Trim14 | 0.313399 | 4.79E-22 | 1.55E-17 |
| Scpep1 | 0.313252 | 5.71E-23 | 1.84E-18 |
| Echs1 | 0.313061 | 1.51E-22 | 4.86E-18 |
| Ddx60 | 0.312372 | 6.07E-20 | 1.96E-15 |
| Tmem104 | 0.310864 | 1.58E-17 | 5.09E-13 |
| Siglecg | 0.310046 | 1.44E-35 | 4.65E-31 |
| F830016B( | 0.308998 | 4.36E-34 | 1.41E-29 |
| Rnase6 | 0.308528 | 6.03E-32 | 1.95E-27 |
| Tns1 | 0.30703 | 1.27E-25 | 4.1E-21 |
| Ston1 | 0.30702 | 1.97E-40 | 6.37E-36 |
| Tagap | 0.306632 | 1.93E-12 | 6.24E-08 |
| Fcgr3 | 0.306058 | 5.23E-21 | 1.69E-16 |
| Dhx9 | 0.305282 | 1.33E-18 | 4.3E-14 |
| Tkt | 0.304726 | 2.07E-21 | 6.67E-17 |
| P2rx4 | 0.304022 | 1.21E-21 | 3.9E-17 |
| Ifit3b | 0.30378 | 2.35E-32 | 7.59E-28 |
| Lrmda | 0.303551 | 4.47E-31 | 1.44E-26 |
| Coro1c | 0.303079 | 1.11E-17 | 3.6E-13 |
| Btf3 | 0.302279 | 3.57E-23 | 1.15E-18 |
| Slc2a1 | 0.302223 | 4.09E-23 | 1.32E-18 |
| Hck | 0.300141 | 3.15E-21 | 1.02E-16 |
| Zfand5 | 0.299582 | 5.59E-12 | 1.81E-07 |

| | | | |
|---|---|---|---|
| Snx2 | 0.299353 | 6.51E-19 | 2.1E-14 |
| Taf1d | 0.298828 | 8.63E-22 | 2.79E-17 |
| C1qbp | 0.298303 | 9.48E-23 | 3.06E-18 |
| Myo5a | 0.298105 | 1.17E-24 | 3.79E-20 |
| Trim25 | 0.298015 | 2.05E-23 | 6.62E-19 |
| Cox5b | 0.297471 | 5.27E-22 | 1.7E-17 |
| Pip4p1 | 0.297323 | 1.79E-23 | 5.78E-19 |
| Slc1a3 | 0.296496 | 4.96E-16 | 1.6E-11 |
| Gsap | 0.296127 | 1.97E-17 | 6.36E-13 |
| Cox7b | 0.295477 | 1.95E-25 | 6.29E-21 |
| Calm1 | 0.294974 | 3.06E-15 | 9.87E-11 |
| Minpp1 | 0.293602 | 6.54E-30 | 2.11E-25 |
| Atp5e | 0.292448 | 4.2E-22 | 1.36E-17 |
| C3 | 0.292061 | 1.85E-32 | 5.97E-28 |
| Dtx3l | 0.291337 | 5.87E-28 | 1.9E-23 |
| Milr1 | 0.291197 | 7.09E-32 | 2.29E-27 |
| Lncpint | 0.290575 | 1.17E-16 | 3.79E-12 |
| Neat1 | 0.290388 | 2.25E-19 | 7.26E-15 |
| Naa25 | 0.289832 | 2.67E-26 | 8.61E-22 |
| Utrn | 0.288572 | 4.47E-05 | 1 |
| Aprt | 0.28743 | 8.53E-27 | 2.75E-22 |
| CAAA0114 | 0.286888 | 2.95E-18 | 9.53E-14 |
| Kpna4 | 0.286305 | 6.89E-15 | 2.22E-10 |
| Sp140 | 0.285167 | 2.77E-25 | 8.95E-21 |
| mt-Nd4l | 0.284356 | 2.09E-21 | 6.74E-17 |
| Fnbp4 | 0.284161 | 4.16E-18 | 1.34E-13 |
| Pgs1 | 0.283865 | 1.48E-27 | 4.78E-23 |
| Gns | 0.283258 | 5.05E-18 | 1.63E-13 |
| Eif3i | 0.283062 | 1.55E-17 | 5E-13 |
| Xrn2 | 0.281953 | 7.21E-24 | 2.33E-19 |
| Pomp | 0.281719 | 9.57E-17 | 3.09E-12 |
| Mapkapk2 | 0.281685 | 1.8E-17 | 5.8E-13 |
| Ripk2 | 0.280237 | 5.56E-17 | 1.8E-12 |
| Prpf38b | 0.279727 | 1.2E-20 | 3.88E-16 |
| Cd86 | 0.279242 | 2.13E-13 | 6.89E-09 |
| Ncl | 0.278622 | 3.85E-14 | 1.24E-09 |
| Ftl1 | 0.277964 | 9.39E-16 | 3.03E-11 |
| Gm47283 | 0.277822 | 2.12E-12 | 6.85E-08 |
| Atp5mpl | 0.277609 | 2.27E-18 | 7.33E-14 |
| Hspa8 | 0.276592 | 5.06E-22 | 1.63E-17 |
| Btaf1 | 0.275739 | 7.72E-18 | 2.49E-13 |
| Uba7 | 0.274475 | 2.37E-18 | 7.65E-14 |
| Calhm6 | 0.274336 | 9.26E-23 | 2.99E-18 |
| Ttc39b | 0.27408 | 5.34E-15 | 1.72E-10 |
| Tnfaip3 | 0.273991 | 3.59E-09 | 0.000116 |
| Ifrd1 | 0.272954 | 1.91E-18 | 6.18E-14 |
| B4galnt1 | 0.271763 | 4.69E-20 | 1.51E-15 |
| Rhbdf2 | 0.271073 | 2.77E-25 | 8.94E-21 |

| | | | |
|---|---|---|---|
| Hint1 | 0.270491 | 1.24E-18 | 4E-14 |
| AI506816 | 0.26966 | 7.87E-20 | 2.54E-15 |
| Pgam1 | 0.2688 | 3.85E-10 | 1.24E-05 |
| Gls | 0.268729 | 1.35E-21 | 4.35E-17 |
| Daxx | 0.268605 | 4.85E-21 | 1.56E-16 |
| Tbc1d1 | 0.268006 | 7.36E-23 | 2.38E-18 |
| Sh3bp2 | 0.267938 | 5.24E-20 | 1.69E-15 |
| Nod1 | 0.267879 | 4.79E-23 | 1.55E-18 |
| Abcg3 | 0.26777 | 9.7E-22 | 3.13E-17 |
| Cox4i1 | 0.267299 | 3.91E-21 | 1.26E-16 |
| Ppargc1b | 0.26713 | 7.36E-34 | 2.38E-29 |
| Slc25a22 | 0.266731 | 6.75E-30 | 2.18E-25 |
| Ppan | 0.266157 | 5.2E-23 | 1.68E-18 |
| Nsd2 | 0.266045 | 6.17E-16 | 1.99E-11 |
| Slc2a3 | 0.266001 | 6.47E-18 | 2.09E-13 |
| Igsf6 | 0.26551 | 1.46E-18 | 4.7E-14 |
| Osbpl9 | 0.265489 | 2.83E-14 | 9.15E-10 |
| Cox5a | 0.265397 | 6.61E-20 | 2.13E-15 |
| Arhgef3 | 0.265265 | 7E-12 | 2.26E-07 |
| Adrb1 | 0.264446 | 3.76E-16 | 1.21E-11 |
| Tmod3 | 0.263605 | 2.46E-15 | 7.93E-11 |
| Lgals9 | 0.263516 | 3.18E-22 | 1.03E-17 |
| Plin2 | 0.263448 | 1.63E-20 | 5.25E-16 |
| Fyn | 0.263016 | 0.543318 | 1 |
| Diaph1 | 0.262387 | 8.67E-27 | 2.8E-22 |
| Atp1a3 | 0.262262 | 1.81E-29 | 5.85E-25 |
| Rtraf | 0.262058 | 2.75E-13 | 8.88E-09 |
| Eepd1 | 0.261867 | 8.93E-16 | 2.88E-11 |
| Myo1e | 0.261814 | 3.91E-12 | 1.26E-07 |
| Arpc1b | 0.260173 | 6.22E-22 | 2.01E-17 |
| Pla2g7 | 0.259887 | 1.62E-34 | 5.24E-30 |
| Cmtr1 | 0.259808 | 1.16E-15 | 3.74E-11 |
| Gnl2 | 0.259611 | 2.01E-17 | 6.49E-13 |
| Coq10b | 0.259154 | 7.03E-18 | 2.27E-13 |
| Pnpla2 | 0.258293 | 1.5E-13 | 4.83E-09 |
| Ppm1h | 0.258204 | 2.31E-12 | 7.46E-08 |
| Rmdn3 | 0.257806 | 8.95E-23 | 2.89E-18 |
| Foxn2 | 0.257805 | 1.33E-18 | 4.28E-14 |
| Atp1a1 | 0.257316 | 2.69E-15 | 8.68E-11 |
| AY036118 | 0.257191 | 0.02155 | 1 |
| Ncoa1 | 0.257165 | 8.07E-15 | 2.61E-10 |
| Adar | 0.256628 | 5.22E-23 | 1.69E-18 |
| H2-Oa | 0.256335 | 1.67E-10 | 5.39E-06 |
| Eif3e | 0.256171 | 2.53E-17 | 8.16E-13 |
| Pstpip1 | 0.2558 | 3.53E-25 | 1.14E-20 |
| Anxa5 | 0.255348 | 1.08E-17 | 3.47E-13 |
| Pgk1 | 0.255254 | 7E-18 | 2.26E-13 |
| Hivep2 | 0.255228 | 3.17E-11 | 1.02E-06 |

| | | | |
|---|---|---|---|
| Tank | 0.255108 | 9.17E-17 | 2.96E-12 |
| Il27 | 0.254847 | 7.1E-29 | 2.29E-24 |
| Colgalt1 | 0.253937 | 1.77E-14 | 5.72E-10 |
| Eif3k | 0.253859 | 8.86E-19 | 2.86E-14 |
| Ptma | 0.25245 | 4.35E-13 | 1.4E-08 |
| Dst | 0.252412 | 1.38E-21 | 4.46E-17 |
| Dnajc13 | 0.252376 | 4.6E-13 | 1.49E-08 |
| Rbpj | 0.252374 | 2.39E-18 | 7.7E-14 |
| Rftn1 | 0.252078 | 6.37E-20 | 2.06E-15 |
| Atp5l | 0.25203 | 1.69E-16 | 5.46E-12 |
| Tomm20 | 0.251324 | 3.15E-16 | 1.02E-11 |
| Cd47 | 0.249563 | 1.18E-11 | 3.81E-07 |
| Plekho2 | 0.249436 | 1.49E-22 | 4.8E-18 |
| Helz2 | 0.248246 | 8.93E-33 | 2.88E-28 |
| Ldlr | 0.247047 | 9.98E-26 | 3.22E-21 |
| Relb | 0.246988 | 4.29E-18 | 1.38E-13 |
| Ifitm2 | 0.246081 | 5.98E-38 | 1.93E-33 |
| Eif4ebp1 | 0.245837 | 1.13E-21 | 3.65E-17 |
| Mlxip | 0.245133 | 4.74E-15 | 1.53E-10 |
| Sharpin | 0.244492 | 5.21E-19 | 1.68E-14 |
| Hs3st3b1 | 0.244207 | 2.46E-29 | 7.95E-25 |
| Nenf | 0.243995 | 6.84E-22 | 2.21E-17 |
| Hspa1a | 0.243621 | 9.46E-18 | 3.05E-13 |
| Cep152 | 0.243521 | 2.66E-10 | 8.6E-06 |
| Taldo1 | 0.242477 | 2.81E-12 | 9.08E-08 |
| Pnpt1 | 0.242337 | 2E-22 | 6.46E-18 |
| Ly6i | 0.241929 | 2.8E-38 | 9.05E-34 |
| Got1 | 0.241928 | 1.58E-18 | 5.11E-14 |
| Gpx1 | 0.241883 | 1.23E-18 | 3.98E-14 |
| Gramd1b | 0.241699 | 0.000126 | 1 |
| Max | 0.241195 | 1.82E-18 | 5.88E-14 |
| Pik3ap1 | 0.241026 | 1.74E-14 | 5.62E-10 |
| E230001N( | 0.241016 | 2.75E-33 | 8.88E-29 |
| Ms4a6b | 0.240205 | 3.26E-10 | 1.05E-05 |
| Ddx39 | 0.240031 | 1.06E-14 | 3.42E-10 |
| Oas1g | 0.239975 | 3.4E-39 | 1.1E-34 |
| Tmed2 | 0.239733 | 1.12E-15 | 3.6E-11 |
| Atp5g3 | 0.23914 | 3.15E-12 | 1.02E-07 |
| C030034L1 | 0.238826 | 1.88E-26 | 6.08E-22 |
| A330040F1 | 0.238586 | 1.68E-16 | 5.42E-12 |
| Junb | 0.238205 | 3.07E-10 | 9.91E-06 |
| Cpq | 0.237808 | 2.2E-19 | 7.11E-15 |
| Cd44 | 0.237639 | 6.85E-15 | 2.21E-10 |
| C1galt1c1 | 0.237559 | 3.16E-14 | 1.02E-09 |
| Tmsb10 | 0.236345 | 6.18E-07 | 0.019939 |
| Anxa3 | 0.236307 | 3.23E-15 | 1.04E-10 |
| Nmi | 0.235219 | 2.5E-15 | 8.07E-11 |
| Rab37 | 0.235107 | 2.23E-39 | 7.2E-35 |

| | | | |
|---|---|---|---|
| Slc25a25 | 0.235085 | 2.85E-26 | 9.19E-22 |
| H2-T24 | 0.234815 | 6.57E-17 | 2.12E-12 |
| Rab20 | 0.234803 | 2.08E-11 | 6.73E-07 |
| Tmem140 | 0.234738 | 1.54E-13 | 4.98E-09 |
| Clcn7 | 0.234438 | 5.82E-15 | 1.88E-10 |
| Slfn9 | 0.234247 | 2.71E-24 | 8.75E-20 |
| Hspd1 | 0.233959 | 4.2E-12 | 1.35E-07 |
| Sdf4 | 0.233922 | 2.65E-18 | 8.55E-14 |
| Mbd2 | 0.233707 | 6.18E-15 | 2E-10 |
| Parp10 | 0.233523 | 2.21E-21 | 7.13E-17 |
| Icosl | 0.233195 | 2.55E-16 | 8.24E-12 |
| Tgif1 | 0.232562 | 1.81E-17 | 5.84E-13 |
| Ccdc102a | 0.232251 | 1.24E-32 | 3.99E-28 |
| Eif3a | 0.231997 | 1.18E-12 | 3.82E-08 |
| mt-Cytb | 0.231492 | 2.07E-21 | 6.7E-17 |
| St14 | 0.231422 | 1.11E-23 | 3.57E-19 |
| Nceh1 | 0.2303 | 7.14E-27 | 2.31E-22 |
| Acot9 | 0.229894 | 2.15E-23 | 6.96E-19 |
| Zc3h12a | 0.229843 | 5.46E-13 | 1.76E-08 |
| B430306N( | 0.229727 | 7.36E-28 | 2.38E-23 |
| Eif1a | 0.228796 | 5.78E-19 | 1.87E-14 |
| Rcan1 | 0.228395 | 0.031109 | 1 |
| Flna | 0.228323 | 2.17E-09 | 7.01E-05 |
| Lyst | 0.227817 | 1.7E-07 | 0.005486 |
| Mt1 | 0.227237 | 3.27E-08 | 0.001057 |
| Gars | 0.227072 | 6.61E-14 | 2.13E-09 |
| Mdfic | 0.227004 | 1.07E-11 | 3.46E-07 |
| Etv3 | 0.22654 | 6.16E-14 | 1.99E-09 |
| Alkbh1 | 0.226516 | 2.14E-20 | 6.9E-16 |
| Tent5a | 0.226506 | 1.25E-13 | 4.04E-09 |
| Lrp10 | 0.226461 | 2.2E-17 | 7.11E-13 |
| Mat2a | 0.22635 | 1.04E-09 | 3.36E-05 |
| Herc4 | 0.2255 | 1.3E-12 | 4.19E-08 |
| Uvrag | 0.225264 | 2.8E-14 | 9.05E-10 |
| Fcgrt | 0.224993 | 2.51E-14 | 8.11E-10 |
| Zdhhc21 | 0.224959 | 7.92E-21 | 2.56E-16 |
| Atp6v0e | 0.22472 | 1.9E-13 | 6.15E-09 |
| Il10ra | 0.224662 | 2.6E-10 | 8.39E-06 |
| Rnf114 | 0.224481 | 8.37E-15 | 2.7E-10 |
| Srm | 0.224423 | 1.07E-27 | 3.46E-23 |
| Tor1aip1 | 0.223688 | 5.86E-13 | 1.89E-08 |
| Ddhd1 | 0.223555 | 1.54E-21 | 4.96E-17 |
| Rasa4 | 0.222828 | 8.46E-12 | 2.73E-07 |
| Cfl1 | 0.222371 | 1.01E-13 | 3.28E-09 |
| Morc3 | 0.222168 | 4.8E-13 | 1.55E-08 |
| Dram2 | 0.221921 | 1.87E-08 | 0.000603 |
| Gm50237 | 0.221855 | 2.36E-34 | 7.63E-30 |
| Vmp1 | 0.221531 | 1.73E-11 | 5.58E-07 |

| | | | |
|---|---|---|---|
| H2afy | 0.221386 | 1.13E-20 | 3.64E-16 |
| Rbms1 | 0.220867 | 2.61E-14 | 8.44E-10 |
| Cox6c | 0.220646 | 3.12E-14 | 1.01E-09 |
| Slc9a3r1 | 0.220528 | 3.51E-14 | 1.13E-09 |
| Dek | 0.220365 | 2.87E-14 | 9.27E-10 |
| Anxa6 | 0.219615 | 3.69E-15 | 1.19E-10 |
| Slc1a5 | 0.219605 | 2.86E-21 | 9.24E-17 |
| Fam174a | 0.219416 | 3.33E-17 | 1.07E-12 |
| Mrpl30 | 0.218988 | 5.78E-15 | 1.87E-10 |
| Gapdh | 0.218789 | 1.44E-10 | 4.66E-06 |
| Ulbp1 | 0.218648 | 1.56E-15 | 5.02E-11 |
| Tor1aip2 | 0.218264 | 5.69E-12 | 1.84E-07 |
| Crybg1 | 0.218124 | 1.6E-23 | 5.16E-19 |
| Bbx | 0.217788 | 6.72E-11 | 2.17E-06 |
| Dbnl | 0.217777 | 1.2E-10 | 3.88E-06 |
| Wdr43 | 0.217459 | 6E-17 | 1.94E-12 |
| Gpr18 | 0.21674 | 3.08E-28 | 9.94E-24 |
| Mapre2 | 0.216524 | 1.06E-09 | 3.41E-05 |
| Coa5 | 0.215601 | 2.25E-19 | 7.26E-15 |
| Tmem154 | 0.215338 | 3.44E-25 | 1.11E-20 |
| Atf3 | 0.214777 | 1.69E-07 | 0.005471 |
| Gas7 | 0.213299 | 2.59E-31 | 8.35E-27 |
| Tent2 | 0.212779 | 2.34E-09 | 7.56E-05 |
| Wars | 0.212489 | 2.98E-14 | 9.63E-10 |
| Asah1 | 0.212363 | 6.44E-17 | 2.08E-12 |
| Lmln | 0.211779 | 5.93E-18 | 1.91E-13 |
| Abhd11 | 0.211566 | 9.56E-19 | 3.09E-14 |
| Actr3 | 0.211469 | 8.63E-12 | 2.78E-07 |
| Nfkbiz | 0.211438 | 3.76E-10 | 1.21E-05 |
| Ogfr | 0.210944 | 8.17E-15 | 2.64E-10 |
| Cox6a1 | 0.210663 | 1.53E-12 | 4.93E-08 |
| Acer3 | 0.210238 | 6.12E-13 | 1.98E-08 |
| Cd180 | 0.209869 | 4.42E-06 | 0.142767 |
| Galc | 0.209568 | 1E-12 | 3.24E-08 |
| Cct2 | 0.209336 | 1.29E-11 | 4.15E-07 |
| Akr1b3 | 0.208773 | 1.42E-11 | 4.58E-07 |
| Fcer1g | 0.208472 | 1.24E-18 | 3.99E-14 |
| Sla | 0.207905 | 7.9E-06 | 0.255137 |
| Atp5c1 | 0.207545 | 3.79E-13 | 1.22E-08 |
| Tlr12 | 0.207372 | 2.33E-13 | 7.52E-09 |
| Shisa5 | 0.206751 | 4.66E-15 | 1.51E-10 |
| Gsto1 | 0.206415 | 1.83E-23 | 5.91E-19 |
| Prkch | 0.206163 | 7.07E-07 | 0.022816 |
| Samd9l | 0.205485 | 2.22E-22 | 7.17E-18 |
| Tlr1 | 0.20548 | 5.38E-16 | 1.74E-11 |
| Map2k2 | 0.205273 | 2.76E-13 | 8.89E-09 |
| Nsa2 | 0.205198 | 1.1E-10 | 3.55E-06 |
| Atp5b | 0.204143 | 1.22E-09 | 3.94E-05 |

| | | | |
|---|---|---|---|
| Psma4 | 0.203964 | 1.77E-11 | 5.72E-07 |
| Gvin1 | 0.203413 | 1.36E-13 | 4.39E-09 |
| Trim12a | 0.203268 | 7.57E-12 | 2.44E-07 |
| Nsmce2 | 0.202933 | 1.53E-09 | 4.95E-05 |
| A930007I1 | 0.202767 | 8.41E-18 | 2.71E-13 |
| Hpcal1 | 0.202452 | 3.52E-18 | 1.14E-13 |
| Uba52 | 0.202191 | 1.25E-17 | 4.05E-13 |
| Timm23 | 0.201975 | 1.65E-13 | 5.33E-09 |
| Rac2 | 0.201945 | 2.01E-09 | 6.5E-05 |
| Gsdmd | 0.2019 | 3.02E-14 | 9.74E-10 |
| Isg20 | 0.20169 | 9.2E-29 | 2.97E-24 |
| Tnfsf12 | 0.201658 | 2.73E-15 | 8.82E-11 |
| Lrrk1 | 0.201609 | 9.66E-11 | 3.12E-06 |
| Eef1d | 0.20151 | 1.67E-13 | 5.4E-09 |
| H2-DMb2 | 0.2012 | 2.13E-21 | 6.88E-17 |
| Ms4a6d | 0.201179 | 2.23E-09 | 7.19E-05 |
| Tmem251 | 0.200994 | 2.22E-17 | 7.17E-13 |
| Art3 | 0.200693 | 4.66E-20 | 1.5E-15 |
| Pfdn5 | 0.200517 | 9.8E-14 | 3.16E-09 |
| Itgb2 | 0.200517 | 4.42E-11 | 1.43E-06 |
| Prelid1 | 0.200494 | 2.08E-14 | 6.7E-10 |
| Myo9b | 0.200473 | 6.15E-13 | 1.99E-08 |
| Hnrnpdl | 0.200403 | 1.6E-08 | 0.000516 |
| Dclre1c | 0.200087 | 1.66E-10 | 5.37E-06 |
| Rab2a | 0.199555 | 1.98E-13 | 6.39E-09 |
| Slc7a1 | 0.199437 | 3.33E-18 | 1.08E-13 |
| Atxn1 | 0.198986 | 0.004559 | 1 |
| Mapkapk3 | 0.198823 | 3.41E-11 | 1.1E-06 |
| Ndst2 | 0.198493 | 3.02E-15 | 9.74E-11 |
| Psma7 | 0.198255 | 1.52E-13 | 4.91E-09 |
| Hsd17b12 | 0.197978 | 3.42E-16 | 1.11E-11 |
| Rab27a | 0.197807 | 5.27E-14 | 1.7E-09 |
| Znrf3 | 0.197603 | 2.81E-16 | 9.07E-12 |
| Nfe2l2 | 0.197551 | 4.16E-11 | 1.34E-06 |
| Cers6 | 0.197354 | 5.81E-07 | 0.01877 |
| Ndufa13 | 0.196679 | 3.49E-10 | 1.13E-05 |
| Snrpg | 0.196485 | 5.09E-10 | 1.64E-05 |
| Otud4 | 0.196017 | 1.32E-11 | 4.25E-07 |
| Tbc1d8 | 0.195481 | 1.03E-16 | 3.34E-12 |
| Rspo1 | 0.195374 | 8.57E-41 | 2.77E-36 |
| Plek | 0.19479 | 9.09E-06 | 0.2936 |
| Gm4070 | 0.194661 | 4.06E-13 | 1.31E-08 |
| Arl5c | 0.194092 | 1.95E-15 | 6.3E-11 |
| Nhp2 | 0.19408 | 1.21E-15 | 3.91E-11 |
| Slc25a5 | 0.193552 | 1.4E-10 | 4.51E-06 |
| Srsf2 | 0.193345 | 5.48E-10 | 1.77E-05 |
| Cycs | 0.19328 | 4.88E-09 | 0.000158 |
| Usp25 | 0.19297 | 4E-11 | 1.29E-06 |

| | | | |
|---|---|---|---|
| Luzp1 | 0.192356 | 4.64E-16 | 1.5E-11 |
| Wdfy1 | 0.192354 | 5.74E-11 | 1.85E-06 |
| Ap3b1 | 0.192272 | 1.01E-10 | 3.26E-06 |
| Slc30a7 | 0.192053 | 1.23E-10 | 3.98E-06 |
| Tnfsf8 | 0.19197 | 4.45E-16 | 1.44E-11 |
| Rnf19b | 0.191671 | 2.49E-16 | 8.04E-12 |
| Nudt9 | 0.191283 | 7.17E-15 | 2.32E-10 |
| Kif3b | 0.191123 | 9.02E-13 | 2.91E-08 |
| Scfd2 | 0.191065 | 3.59E-09 | 0.000116 |
| Ran | 0.190563 | 9.19E-06 | 0.296573 |
| Snx10 | 0.190179 | 4.51E-09 | 0.000146 |
| Idnk | 0.190085 | 4.65E-14 | 1.5E-09 |
| Tpr | 0.189542 | 6.53E-14 | 2.11E-09 |
| Irgm2 | 0.189443 | 7.96E-08 | 0.002571 |
| Rilpl2 | 0.189382 | 1.34E-16 | 4.31E-12 |
| Mfsd12 | 0.189304 | 3.67E-21 | 1.18E-16 |
| Golgb1 | 0.189113 | 4.84E-13 | 1.56E-08 |
| Etf1 | 0.188649 | 1.92E-11 | 6.19E-07 |
| Prkca | 0.188597 | 7.18E-05 | 1 |
| Uri1 | 0.188119 | 1.05E-13 | 3.37E-09 |
| C1qc | 0.187954 | 1.95E-28 | 6.3E-24 |
| Xpo6 | 0.187743 | 1.92E-12 | 6.19E-08 |
| Pgd | 0.18745 | 1.31E-12 | 4.23E-08 |
| Hk3 | 0.187248 | 1.53E-07 | 0.004949 |
| Ndufa1 | 0.187181 | 4.71E-14 | 1.52E-09 |
| Vps13c | 0.186861 | 1.18E-13 | 3.82E-09 |
| Nt5c | 0.186775 | 4.37E-17 | 1.41E-12 |
| Sdad1 | 0.18668 | 6.03E-13 | 1.95E-08 |
| Alkbh2 | 0.186054 | 1.4E-26 | 4.51E-22 |
| Epb41l4os | 0.185941 | 1.94E-15 | 6.26E-11 |
| Setdb2 | 0.185743 | 1.28E-14 | 4.15E-10 |
| Mfsd1 | 0.184774 | 5.45E-14 | 1.76E-09 |
| Gpr160 | 0.184179 | 3.34E-09 | 0.000108 |
| Ndufb1-ps | 0.183989 | 7.16E-12 | 2.31E-07 |
| Ncf1 | 0.18391 | 1.97E-09 | 6.34E-05 |
| A930015D( | 0.183836 | 2.65E-13 | 8.57E-09 |
| Pabpc1 | 0.183798 | 6.47E-09 | 0.000209 |
| Sap30 | 0.183785 | 1.44E-17 | 4.64E-13 |
| Spop | 0.183759 | 8.61E-12 | 2.78E-07 |
| Slc49a4 | 0.183219 | 6.14E-09 | 0.000198 |
| Msrb1 | 0.182862 | 4.59E-15 | 1.48E-10 |
| 2810013P( | 0.182781 | 7.52E-09 | 0.000243 |
| Hebp1 | 0.182525 | 3.24E-14 | 1.04E-09 |
| Ppa1 | 0.18241 | 5.43E-21 | 1.75E-16 |
| Ak2 | 0.182324 | 3E-09 | 9.68E-05 |
| Il18 | 0.181986 | 3.85E-10 | 1.24E-05 |
| Lfng | 0.181463 | 1.39E-11 | 4.48E-07 |
| Pde7a | 0.181231 | 2.33E-05 | 0.75097 |

| | | | |
|---|---|---|---|
| Slc43a2 | 0.181194 | 0.000204 | 1 |
| D16Ertd47 | 0.180945 | 2.29E-15 | 7.41E-11 |
| Gm7030 | 0.18012 | 6.42E-25 | 2.07E-20 |
| Bola2 | 0.18001 | 4.66E-14 | 1.51E-09 |
| Hivep1 | 0.17996 | 1.96E-13 | 6.32E-09 |
| Bri3bp | 0.179769 | 1.22E-18 | 3.94E-14 |
| Pirb | 0.17938 | 2.13E-13 | 6.88E-09 |
| Mphosph1( | 0.178513 | 2.72E-13 | 8.79E-09 |
| A930037H( | 0.178312 | 2.35E-22 | 7.58E-18 |
| Nop56 | 0.178215 | 5.34E-09 | 0.000173 |
| Sgcb | 0.177894 | 1.28E-18 | 4.13E-14 |
| Atp5h | 0.177625 | 3.38E-11 | 1.09E-06 |
| Tubb5 | 0.177153 | 1.79E-07 | 0.005793 |
| Sub1 | 0.177006 | 1.31E-07 | 0.004233 |
| Pi4ka | 0.176995 | 6.26E-13 | 2.02E-08 |
| Fam241a | 0.176885 | 1.57E-29 | 5.07E-25 |
| Cox6b1 | 0.176871 | 5.08E-11 | 1.64E-06 |
| Mrpl54 | 0.176768 | 2.06E-11 | 6.65E-07 |
| Dennd1b | 0.176767 | 2.68E-08 | 0.000865 |
| Slc38a6 | 0.17628 | 5.61E-10 | 1.81E-05 |
| Phf20l1 | 0.176105 | 3.65E-10 | 1.18E-05 |
| Raly | 0.175718 | 2.77E-14 | 8.94E-10 |
| Hspa5 | 0.175706 | 2.33E-08 | 0.000751 |
| Pced1b | 0.175423 | 1.54E-13 | 4.96E-09 |
| Dop1b | 0.175362 | 7.28E-13 | 2.35E-08 |
| Trpv4 | 0.175222 | 1.87E-23 | 6.05E-19 |
| Parp11 | 0.175081 | 3.71E-14 | 1.2E-09 |
| Crybb1 | 0.174982 | 0.001431 | 1 |
| Grina | 0.174205 | 6.18E-12 | 2E-07 |
| Psme4 | 0.174182 | 4.39E-11 | 1.42E-06 |
| Anp32b | 0.173773 | 1.38E-14 | 4.45E-10 |
| Atf6 | 0.173127 | 2.06E-10 | 6.66E-06 |
| Pdk3 | 0.173026 | 5.55E-12 | 1.79E-07 |
| Sfxn1 | 0.172362 | 5.37E-14 | 1.73E-09 |
| Atg3 | 0.172198 | 5.07E-13 | 1.64E-08 |
| Tyk2 | 0.171969 | 3.43E-11 | 1.11E-06 |
| Psmb2 | 0.171651 | 6.48E-08 | 0.002093 |
| Nfkbib | 0.17119 | 1.33E-10 | 4.28E-06 |
| Casp1 | 0.170947 | 4.19E-12 | 1.35E-07 |
| Plbd2 | 0.170278 | 2.25E-11 | 7.26E-07 |
| Rps27 | 0.170266 | 6.27E-11 | 2.02E-06 |
| Rbm17 | 0.169636 | 1.03E-18 | 3.32E-14 |
| Pfkfb3 | 0.169385 | 1.85E-08 | 0.000598 |
| Slc15a4 | 0.169301 | 1.69E-15 | 5.45E-11 |
| Relt | 0.16915 | 1.94E-18 | 6.25E-14 |
| Eef2 | 0.168918 | 4.86E-10 | 1.57E-05 |
| Cox7a2 | 0.168547 | 3.69E-10 | 1.19E-05 |
| Trim34a | 0.168479 | 4.09E-13 | 1.32E-08 |

| | | | |
|---|---|---|---|
| Arf4 | 0.168452 | 6.16E-07 | 0.019893 |
| Nktr | 0.167778 | 9.03E-11 | 2.92E-06 |
| Mvp | 0.167463 | 8.2E-12 | 2.65E-07 |
| Snhg6 | 0.167393 | 7.78E-21 | 2.51E-16 |
| Pld4 | 0.167007 | 6.94E-11 | 2.24E-06 |
| Gm17268 | 0.166881 | 3.94E-15 | 1.27E-10 |
| Lcp2 | 0.1668 | 5.92E-06 | 0.190992 |
| M6pr | 0.166509 | 1.1E-08 | 0.000355 |
| Pfkl | 0.166327 | 2.52E-11 | 8.13E-07 |
| Ctsl | 0.166201 | 6.37E-09 | 0.000206 |
| Coq7 | 0.166091 | 5.92E-18 | 1.91E-13 |
| Taok3 | 0.165687 | 1.78E-10 | 5.73E-06 |
| Hmgb2 | 0.16557 | 0.000384 | 1 |
| Dnm1l | 0.165288 | 1.37E-12 | 4.42E-08 |
| Hnrnpll | 0.164926 | 3.49E-08 | 0.001127 |
| Higd1a | 0.164873 | 9.67E-10 | 3.12E-05 |
| Svbp | 0.164283 | 2.18E-09 | 7.03E-05 |
| Mcl1 | 0.164024 | 4.51E-07 | 0.014553 |
| Atp5o | 0.163406 | 7.6E-09 | 0.000246 |
| Cep128 | 0.163141 | 3.69E-11 | 1.19E-06 |
| Haao | 0.162053 | 4.11E-19 | 1.33E-14 |
| Vars | 0.162013 | 3.61E-08 | 0.001167 |
| Adipor2 | 0.162004 | 2.29E-13 | 7.38E-09 |
| Lrpap1 | 0.161916 | 2.18E-08 | 0.000704 |
| Ppfia4 | 0.161568 | 9.36E-07 | 0.030222 |
| Gm37240 | 0.161357 | 2.59E-07 | 0.008361 |
| Slc12a9 | 0.161345 | 2.41E-09 | 7.78E-05 |
| Nmd3 | 0.161038 | 1.67E-09 | 5.38E-05 |
| Cct5 | 0.160899 | 1.81E-09 | 5.85E-05 |
| Uqcc2 | 0.160592 | 2.55E-08 | 0.000824 |
| Bak1 | 0.16059 | 3.05E-12 | 9.84E-08 |
| Ptrhd1 | 0.160582 | 1.44E-11 | 4.66E-07 |
| Eps8 | 0.160394 | 1.06E-10 | 3.41E-06 |
| Atrnl1 | 0.160285 | 3.33E-18 | 1.08E-13 |
| Vps13a | 0.160083 | 1.69E-10 | 5.47E-06 |
| Dynll1 | 0.160025 | 1.64E-06 | 0.052949 |
| Dnajb14 | 0.159912 | 4.03E-09 | 0.00013 |
| 0610012G( | 0.159658 | 1.37E-10 | 4.42E-06 |
| Tpd52 | 0.15934 | 1.29E-08 | 0.000417 |
| Osm | 0.158773 | 1.12E-08 | 0.000363 |
| Hcst | 0.158112 | 5.51E-12 | 1.78E-07 |
| Mthfr | 0.157767 | 1.29E-11 | 4.17E-07 |
| Tex2 | 0.157227 | 3.05E-08 | 0.000984 |
| Plekhf2 | 0.157046 | 2.48E-15 | 8.01E-11 |
| Ppib | 0.156924 | 2.27E-06 | 0.073242 |
| Rnh1 | 0.156898 | 9.58E-08 | 0.003092 |
| Fbl | 0.156755 | 2.62E-07 | 0.008462 |
| Gusb | 0.156738 | 1.17E-08 | 0.000378 |

| | | | |
|---|---|---|---|
| Abl2 | 0.156594 | 8.56E-09 | 0.000276 |
| Atp5j2 | 0.156487 | 3.21E-06 | 0.103608 |
| Man2b1 | 0.155841 | 8.17E-12 | 2.64E-07 |
| Ola1 | 0.155603 | 1.3E-10 | 4.2E-06 |
| Capn1 | 0.155157 | 1.31E-14 | 4.24E-10 |
| Psma2 | 0.155155 | 9.93E-08 | 0.003205 |
| Rras2 | 0.154956 | 1.18E-13 | 3.8E-09 |
| 2410002F2 | 0.154841 | 1.89E-05 | 0.611695 |
| Aatf | 0.154798 | 5.42E-12 | 1.75E-07 |
| Ndufv3 | 0.154753 | 2.08E-08 | 0.000673 |
| Plekha1 | 0.154714 | 5.14E-11 | 1.66E-06 |
| Pde4b | 0.154452 | 0.266336 | 1 |
| Kxd1 | 0.154442 | 3.59E-08 | 0.00116 |
| C5ar1 | 0.15437 | 5.26E-06 | 0.169758 |
| Lrp12 | 0.154313 | 1.23E-08 | 0.000397 |
| Spata5 | 0.154037 | 1.14E-09 | 3.68E-05 |
| Ftl1-ps1 | 0.153922 | 2.64E-17 | 8.52E-13 |
| Tmem184b | 0.153816 | 9.66E-12 | 3.12E-07 |
| Vapa | 0.153809 | 1.58E-05 | 0.51024 |
| Ankle2 | 0.153791 | 9.25E-09 | 0.000299 |
| Snd1 | 0.153733 | 1.81E-09 | 5.84E-05 |
| Gyg | 0.152838 | 1.13E-07 | 0.003656 |
| Cp | 0.152642 | 7.89E-11 | 2.55E-06 |
| Gm14023 | 0.152539 | 1.22E-10 | 3.94E-06 |
| Synj1 | 0.152532 | 3.77E-08 | 0.001216 |
| Nop14 | 0.152243 | 5.3E-08 | 0.001711 |
| Htt | 0.152219 | 5.99E-10 | 1.93E-05 |
| Exoc6 | 0.152011 | 4.99E-10 | 1.61E-05 |
| Anapc5 | 0.151817 | 4.49E-08 | 0.00145 |
| Tubb6 | 0.151791 | 1.83E-11 | 5.89E-07 |
| Limd2 | 0.151617 | 4.91E-06 | 0.158566 |
| Prmt3 | 0.151549 | 4.66E-16 | 1.5E-11 |
| Larp1 | 0.151361 | 2.09E-11 | 6.75E-07 |
| Neu1 | 0.151297 | 1.05E-08 | 0.000339 |
| Pik3r1 | 0.150924 | 0.000713 | 1 |
| Aoah | 0.15087 | 3.69E-20 | 1.19E-15 |
| Ecpas | 0.150318 | 2.95E-09 | 9.52E-05 |
| Hat1 | 0.150177 | 3.48E-09 | 0.000112 |
| Sys1 | 0.15011 | 4.54E-07 | 0.014653 |
| Tpm4 | 0.1501 | 8.17E-09 | 0.000264 |
| Cyb5r3 | 0.149626 | 4.04E-09 | 0.00013 |
| Tlr7 | 0.149444 | 0.004613 | 1 |
| Psmg4 | 0.149178 | 2.76E-11 | 8.9E-07 |
| Dnttip1 | 0.149024 | 1.27E-16 | 4.09E-12 |
| Cntrl | 0.148642 | 7.09E-08 | 0.00229 |
| Txndc17 | 0.148417 | 4.65E-08 | 0.001503 |
| Sri | 0.148403 | 7.53E-11 | 2.43E-06 |
| Elovl1 | 0.148107 | 1.44E-06 | 0.046625 |

| | | | |
|---|---|---|---|
| Nvl | 0.147426 | 1.69E-11 | 5.46E-07 |
| Ndufb8 | 0.147341 | 3.61E-09 | 0.000117 |
| mt-Nd2 | 0.146885 | 5.4E-07 | 0.017436 |
| Eea1 | 0.14687 | 1.35E-09 | 4.37E-05 |
| Ndufb9 | 0.146846 | 5.82E-07 | 0.018781 |
| Ccdc134 | 0.14684 | 2.26E-12 | 7.29E-08 |
| Slc3a2 | 0.146686 | 8.53E-08 | 0.002755 |
| Hmox1 | 0.146671 | 8.12E-05 | 1 |
| Gm40645 | 0.146638 | 1.51E-13 | 4.87E-09 |
| Gm31718 | 0.146178 | 1.15E-15 | 3.71E-11 |
| Anapc16 | 0.145651 | 3.36E-10 | 1.09E-05 |
| Ptk2b | 0.145485 | 1.9E-05 | 0.614944 |
| Zfp106 | 0.145381 | 2.01E-11 | 6.5E-07 |
| Acadl | 0.144812 | 5.58E-06 | 0.180284 |
| 4930469K1 | 0.144671 | 1.83E-07 | 0.005907 |
| Hps5 | 0.144623 | 1.22E-10 | 3.94E-06 |
| Cct8 | 0.144415 | 2.41E-07 | 0.007784 |
| Micu2 | 0.144415 | 1.46E-09 | 4.73E-05 |
| Atp5g1 | 0.144413 | 7.04E-08 | 0.002272 |
| Creld2 | 0.144393 | 6.05E-07 | 0.01952 |
| Sdhb | 0.144302 | 7.5E-07 | 0.02423 |
| Snrpe | 0.144272 | 3.59E-06 | 0.115837 |
| Srsf7 | 0.144193 | 0.000286 | 1 |
| Rbm26 | 0.144072 | 4.11E-07 | 0.013283 |
| Lrch3 | 0.143868 | 1.46E-06 | 0.047135 |
| Psmb4 | 0.14361 | 7.4E-08 | 0.002388 |
| Gpr35 | 0.143571 | 2.78E-20 | 8.98E-16 |
| Stx16 | 0.143433 | 1.48E-07 | 0.004779 |
| Dennd4a | 0.143166 | 0.347554 | 1 |
| Frrs1 | 0.143084 | 3.64E-05 | 1 |
| Jdp2 | 0.142981 | 6.36E-17 | 2.05E-12 |
| Irak2 | 0.14279 | 0.005285 | 1 |
| Fryl | 0.142743 | 0.000111 | 1 |
| Gng5 | 0.142615 | 5.36E-07 | 0.017308 |
| 2310001H1 | 0.142583 | 2.56E-16 | 8.25E-12 |
| Ipcef1 | 0.14251 | 8.61E-10 | 2.78E-05 |
| Hspa4 | 0.142435 | 1.37E-09 | 4.42E-05 |
| Rbm25 | 0.142426 | 1.5E-10 | 4.86E-06 |
| Lrrc59 | 0.142281 | 3.96E-12 | 1.28E-07 |
| Tnfsf10 | 0.142258 | 7.05E-14 | 2.28E-09 |
| Coro1a | 0.142214 | 4.29E-06 | 0.138342 |
| Il6 | 0.141883 | 1.46E-10 | 4.71E-06 |
| Naip2 | 0.141709 | 2E-08 | 0.000646 |
| Pcna | 0.141629 | 2.58E-05 | 0.833 |
| Ppia | 0.141569 | 3.32E-05 | 1 |
| Cfap410 | 0.141337 | 4.04E-11 | 1.31E-06 |
| Nob1 | 0.140895 | 4.32E-08 | 0.001395 |
| Sod2 | 0.140802 | 2.31E-10 | 7.47E-06 |

| | | | |
|---|---|---|---|
| Cd81 | 0.140743 | 2.22E-08 | 0.000716 |
| Psmd8 | 0.14073 | 4.91E-06 | 0.158548 |
| Tmbim1 | 0.140607 | 4.89E-12 | 1.58E-07 |
| Eif2s1 | 0.140232 | 2.16E-08 | 0.000697 |
| Nedd4l | 0.140219 | 3.58E-11 | 1.16E-06 |
| Mrps24 | 0.139942 | 1.46E-09 | 4.7E-05 |
| H2afj | 0.139645 | 6.49E-06 | 0.209588 |
| Gtf2f1 | 0.139336 | 2.77E-07 | 0.008932 |
| Eif4e3 | 0.139266 | 5.47E-10 | 1.77E-05 |
| Rufy3 | 0.139166 | 5.46E-13 | 1.76E-08 |
| Slamf6 | 0.138931 | 3.55E-10 | 1.15E-05 |
| Nucb2 | 0.138766 | 3.65E-13 | 1.18E-08 |
| Eva1b | 0.138678 | 1.32E-12 | 4.26E-08 |
| Alpk1 | 0.13859 | 1.68E-11 | 5.42E-07 |
| Inpp1 | 0.138465 | 8.53E-15 | 2.75E-10 |
| Trim21 | 0.138353 | 3.8E-12 | 1.23E-07 |
| Litaf | 0.138275 | 8.36E-07 | 0.027004 |
| Nab1 | 0.138225 | 1.8E-09 | 5.82E-05 |
| C9orf72 | 0.138152 | 5.95E-09 | 0.000192 |
| Csnk1g1 | 0.137745 | 2.9E-06 | 0.093655 |
| Snrpd2 | 0.137504 | 8.46E-10 | 2.73E-05 |
| Nudt19 | 0.137291 | 7.1E-20 | 2.29E-15 |
| Lair1 | 0.13716 | 4.84E-09 | 0.000156 |
| Arhgef7 | 0.136901 | 2.83E-07 | 0.009147 |
| Kntc1 | 0.136861 | 1.7E-09 | 5.5E-05 |
| Sptlc2 | 0.136807 | 6.1E-06 | 0.196958 |
| Farsa | 0.136592 | 4.14E-10 | 1.34E-05 |
| Gde1 | 0.136507 | 2.09E-10 | 6.76E-06 |
| Snhg16 | 0.136116 | 6.67E-12 | 2.15E-07 |
| Rassf4 | 0.135918 | 1.74E-07 | 0.005614 |
| Sass6 | 0.135813 | 3.49E-11 | 1.13E-06 |
| Aim2 | 0.135582 | 8.14E-06 | 0.26274 |
| B4galt5 | 0.135354 | 3.62E-06 | 0.116976 |
| Dapk1 | 0.135337 | 1.87E-07 | 0.006029 |
| Rapgef2 | 0.135123 | 3.35E-08 | 0.00108 |
| Rnf149 | 0.134943 | 5.65E-08 | 0.001823 |
| Slirp | 0.134817 | 7.93E-11 | 2.56E-06 |
| Psma3 | 0.134742 | 5.45E-07 | 0.017608 |
| Pmm2 | 0.134546 | 8.77E-09 | 0.000283 |
| Apmap | 0.134417 | 4.61E-10 | 1.49E-05 |
| Bclaf1 | 0.134078 | 1.41E-06 | 0.045594 |
| Lifr | 0.133888 | 0.005844 | 1 |
| Aopep | 0.133821 | 0.000352 | 1 |
| Cox7a2l | 0.13374 | 2.48E-08 | 0.000799 |
| Uqcr11 | 0.133707 | 4.15E-10 | 1.34E-05 |
| Ilrun | 0.13353 | 3.21E-11 | 1.04E-06 |
| Atp13a1 | 0.13335 | 3.92E-12 | 1.27E-07 |
| Tomm40 | 0.133025 | 4.08E-07 | 0.013171 |

| | | | |
|---|---|---|---|
| Peli1 | 0.132999 | 0.000283 | 1 |
| Bnip2 | 0.132642 | 4.53E-06 | 0.14614 |
| Nsd3 | 0.132603 | 0.00041 | 1 |
| Slc6a6 | 0.132598 | 1.1E-08 | 0.000355 |
| Synj2 | 0.132197 | 1.99E-11 | 6.42E-07 |
| Alg5 | 0.132123 | 2.55E-05 | 0.821869 |
| Prpf40a | 0.13174 | 1.28E-07 | 0.004126 |
| Heatr5a | 0.131525 | 4.86E-06 | 0.157035 |
| Eed | 0.131474 | 2.71E-06 | 0.087446 |
| Chd1 | 0.131351 | 6.04E-08 | 0.001951 |
| Hsph1 | 0.131225 | 1.59E-08 | 0.000512 |
| Psen2 | 0.131224 | 9.23E-08 | 0.002981 |
| Slc16a3 | 0.131008 | 0.000136 | 1 |
| Mrto4 | 0.130653 | 1.56E-08 | 0.000503 |
| Ripk1 | 0.130651 | 3.01E-10 | 9.71E-06 |
| Srsf6 | 0.130636 | 3.48E-07 | 0.011229 |
| Eif3m | 0.130612 | 0.000175 | 1 |
| Zdhhc3 | 0.130569 | 1.33E-07 | 0.004278 |
| Ranbp1 | 0.130553 | 0.000107 | 1 |
| Acvrl1 | 0.130486 | 2.83E-07 | 0.009145 |
| Txn2 | 0.130479 | 1.02E-06 | 0.032793 |
| Asb13 | 0.130285 | 4.19E-09 | 0.000135 |
| Nap1l1 | 0.130198 | 3.98E-05 | 1 |
| Ddx58 | 0.1301 | 3.19E-07 | 0.010299 |
| Mrps18b | 0.12991 | 4.99E-09 | 0.000161 |
| Susd6 | 0.129749 | 9.46E-05 | 1 |
| mt-Nd5 | 0.129669 | 3.97E-07 | 0.012824 |
| Psmd12 | 0.129197 | 1.38E-05 | 0.445528 |
| Sept9 | 0.129108 | 0.000289 | 1 |
| Gm2000 | 0.128958 | 6.33E-14 | 2.04E-09 |
| Cryba4 | 0.128901 | 1.67E-07 | 0.00539 |
| Il4ra | 0.128896 | 3.85E-07 | 0.012425 |
| Ctla2b | 0.128891 | 3.26E-11 | 1.05E-06 |
| Chuk | 0.128753 | 5.62E-08 | 0.001815 |
| Emc4 | 0.12874 | 4.42E-06 | 0.142722 |
| Ndufs3 | 0.128287 | 3.36E-08 | 0.001084 |
| Pola1 | 0.128176 | 0.006905 | 1 |
| Cbwd1 | 0.127946 | 3.03E-09 | 9.8E-05 |
| Gla | 0.127579 | 2.15E-17 | 6.95E-13 |
| Pfn1 | 0.127495 | 0.000673 | 1 |
| Usp30 | 0.127384 | 1.5E-17 | 4.84E-13 |
| Pign | 0.127178 | 2.64E-09 | 8.54E-05 |
| Mcm6 | 0.126744 | 0.000964 | 1 |
| Slc7a6 | 0.126712 | 1.16E-10 | 3.74E-06 |
| Atp11b | 0.126682 | 2.1E-07 | 0.006778 |
| mt-Nd4 | 0.126425 | 2.52E-07 | 0.008146 |
| Tbc1d10a | 0.126405 | 1.13E-06 | 0.036608 |
| Rpia | 0.126403 | 4.4E-08 | 0.00142 |

| | | | |
|---|---|---|---|
| Ube2d3 | 0.126353 | 2.62E-05 | 0.844889 |
| Top1 | 0.126196 | 0.000133 | 1 |
| Ubr4 | 0.126083 | 5.74E-07 | 0.018534 |
| Zfp810 | 0.126061 | 6.81E-06 | 0.219879 |
| Zfp593 | 0.125918 | 3.15E-10 | 1.02E-05 |
| Ets2 | 0.125835 | 2.57E-05 | 0.829106 |
| Sqstm1 | 0.125563 | 0.000915 | 1 |
| Btg2 | 0.125392 | 0.054724 | 1 |
| Rabggtb | 0.125141 | 8.31E-09 | 0.000268 |
| Aig1 | 0.124897 | 9.06E-09 | 0.000292 |
| Gstcd | 0.124711 | 9.05E-10 | 2.92E-05 |
| Myh10 | 0.124686 | 8.88E-15 | 2.87E-10 |
| Pi4k2a | 0.124551 | 3.85E-08 | 0.001244 |
| Cmpk2 | 0.124383 | 2.63E-22 | 8.5E-18 |
| Stim2 | 0.124332 | 2.95E-10 | 9.54E-06 |
| Calr | 0.1241 | 0.000661 | 1 |
| Tmem87b | 0.124084 | 2.32E-05 | 0.749602 |
| Pdia5 | 0.124073 | 3.58E-08 | 0.001155 |
| Ndufa6 | 0.124067 | 2.43E-06 | 0.078591 |
| Notch4 | 0.124021 | 7.12E-10 | 2.3E-05 |
| Tmem67 | 0.124007 | 6.06E-12 | 1.96E-07 |
| Mark2 | 0.123985 | 2.52E-05 | 0.813081 |
| Zfp330 | 0.123796 | 1.43E-10 | 4.62E-06 |
| Irs2 | 0.123598 | 1.61E-08 | 0.000521 |
| Zc3h12c | 0.123493 | 5.88E-09 | 0.00019 |
| Zeb2 | 0.123355 | 3.11E-05 | 1 |
| Rassf1 | 0.122898 | 2.42E-06 | 0.078225 |
| Kctd5 | 0.122857 | 1.98E-09 | 6.4E-05 |
| Snhg15 | 0.122821 | 1.08E-08 | 0.000348 |
| Wdr12 | 0.122477 | 6.2E-07 | 0.02001 |
| Mov10 | 0.122427 | 1.77E-07 | 0.005701 |
| Tpcn2 | 0.122247 | 8.64E-13 | 2.79E-08 |
| Impdh2 | 0.122148 | 5.48E-08 | 0.001768 |
| Zfp622 | 0.122101 | 2.13E-05 | 0.68712 |
| Dnajc15 | 0.121761 | 5.14E-06 | 0.165805 |
| Rnasel | 0.121365 | 5.56E-08 | 0.001794 |
| Kdm7a | 0.121286 | 5.25E-07 | 0.016965 |
| Gtf2f2 | 0.121277 | 2.1E-07 | 0.006782 |
| Orai1 | 0.121219 | 4.84E-06 | 0.156289 |
| Safb2 | 0.121176 | 1.48E-06 | 0.047628 |
| Snu13 | 0.121136 | 1.83E-05 | 0.590303 |
| Erap1 | 0.121051 | 2.42E-06 | 0.078221 |
| Reep4 | 0.120983 | 2.04E-05 | 0.658367 |
| Atp6v1a | 0.120828 | 0.000136 | 1 |
| Utp14a | 0.120745 | 4.46E-06 | 0.144131 |
| Slc22a17 | 0.120567 | 7.42E-07 | 0.023961 |
| Fgfr1op | 0.120521 | 1.93E-09 | 6.24E-05 |
| Cmc1 | 0.120503 | 5.55E-09 | 0.000179 |

| | | | |
|---|---|---|---|
| Wdr55 | 0.120466 | 1.14E-07 | 0.003679 |
| Dnajc3 | 0.120382 | 1.6E-07 | 0.005152 |
| Jak3 | 0.120222 | 9.65E-10 | 3.11E-05 |
| Keap1 | 0.119958 | 1.75E-05 | 0.565505 |
| Abhd15 | 0.119882 | 0.261343 | 1 |
| Elob | 0.119842 | 0.000116 | 1 |
| Sdf2l1 | 0.119841 | 5.39E-07 | 0.017397 |
| Psmd14 | 0.119836 | 4.31E-06 | 0.139133 |
| Taf10 | 0.119649 | 1.97E-07 | 0.006345 |
| Pacsin2 | 0.119616 | 1.94E-07 | 0.006255 |
| Serf2 | 0.119112 | 2.08E-06 | 0.06723 |
| Daam1 | 0.1191 | 1.2E-12 | 3.88E-08 |
| Nop16 | 0.119061 | 6.78E-10 | 2.19E-05 |
| Abhd16a | 0.119006 | 0.000177 | 1 |
| Trim56 | 0.118951 | 3.07E-06 | 0.099131 |
| BC147527 | 0.118804 | 6.02E-16 | 1.94E-11 |
| St7 | 0.118777 | 3.97E-05 | 1 |
| Tnfrsf14 | 0.118488 | 1.66E-09 | 5.35E-05 |
| Cisd2 | 0.118228 | 5.67E-08 | 0.001831 |
| Sec61b | 0.118073 | 7.56E-08 | 0.002441 |
| Abracl | 0.117988 | 1.31E-05 | 0.421741 |
| Resf1 | 0.117826 | 1.15E-06 | 0.037058 |
| Batf3 | 0.117816 | 3.37E-07 | 0.010891 |
| Zfyve26 | 0.11772 | 2.91E-07 | 0.009389 |
| Plin3 | 0.117578 | 1.38E-12 | 4.44E-08 |
| Tra2b | 0.11757 | 1.29E-06 | 0.041649 |
| Bag3 | 0.1175 | 3.51E-06 | 0.113405 |
| Tomm70a | 0.117499 | 3.51E-08 | 0.001132 |
| Csnk2b | 0.117377 | 0.00061 | 1 |
| Masp1 | 0.117263 | 3.59E-15 | 1.16E-10 |
| Ppp1r14b | 0.117215 | 6.89E-09 | 0.000223 |
| Gch1 | 0.117179 | 4.42E-12 | 1.43E-07 |
| Abhd17a | 0.117172 | 1.49E-07 | 0.004808 |
| Sec61g | 0.117093 | 8.42E-06 | 0.271857 |
| Rrs1 | 0.117053 | 9.73E-07 | 0.031401 |
| Nubp1 | 0.116947 | 3.52E-07 | 0.011357 |
| Sidt2 | 0.116914 | 9.72E-08 | 0.003139 |
| Tspan32 | 0.116729 | 1.53E-10 | 4.95E-06 |
| Polr1d | 0.116724 | 9.57E-07 | 0.030886 |
| Rcn1 | 0.116686 | 1.75E-08 | 0.000565 |
| Gmppb | 0.116546 | 6.29E-09 | 0.000203 |
| Nfkbid | 0.116474 | 2.97E-05 | 0.958291 |
| Rbbp8 | 0.116244 | 1.22E-06 | 0.039287 |
| Atad2 | 0.116074 | 1.19E-05 | 0.385013 |
| Noc4l | 0.116028 | 7.35E-13 | 2.37E-08 |
| Scarf2 | 0.115963 | 3.75E-10 | 1.21E-05 |
| Ccdc25 | 0.115681 | 1.08E-08 | 0.000348 |
| Bzw2 | 0.11535 | 8.31E-10 | 2.68E-05 |

| | | | |
|---|---|---|---|
| Slc35b1 | 0.115272 | 5.48E-05 | 1 |
| Eftud2 | 0.115163 | 4.68E-07 | 0.015117 |
| Mrpl32 | 0.115139 | 7.85E-08 | 0.002534 |
| Elf4 | 0.115134 | 0.000566 | 1 |
| Bach2os | 0.115109 | 5.25E-12 | 1.7E-07 |
| AC113595. | 0.115064 | 1.76E-11 | 5.68E-07 |
| Cacybp | 0.114979 | 1.07E-07 | 0.003453 |
| Ist1 | 0.114839 | 6.7E-07 | 0.021639 |
| Rap2b | 0.114723 | 3.2E-10 | 1.03E-05 |
| Gbgt1 | 0.114658 | 5.36E-14 | 1.73E-09 |
| Pdss1 | 0.11459 | 1.09E-10 | 3.53E-06 |
| Rab24 | 0.114589 | 7.39E-09 | 0.000239 |
| Cyp51 | 0.114583 | 2.18E-07 | 0.007028 |
| Tpp2 | 0.114173 | 4.01E-07 | 0.012949 |
| Akr1b10 | 0.113985 | 0.000207 | 1 |
| Snhg4 | 0.113839 | 1.1E-13 | 3.54E-09 |
| Ccnl2 | 0.113598 | 8.3E-06 | 0.268091 |
| Qars | 0.113417 | 2.77E-07 | 0.008944 |
| Incenp | 0.113246 | 2.08E-05 | 0.671738 |
| Adam9 | 0.113093 | 1.9E-07 | 0.006144 |
| Tmco4 | 0.113077 | 1.27E-06 | 0.040946 |
| Pcgf5 | 0.113011 | 2.74E-09 | 8.84E-05 |
| Sf3b1 | 0.112951 | 7.66E-07 | 0.02474 |
| Nfxl1 | 0.112706 | 1.27E-07 | 0.004115 |
| Mafk | 0.11262 | 2.81E-09 | 9.08E-05 |
| Pdcd2 | 0.112556 | 1.07E-09 | 3.46E-05 |
| Cnr2 | 0.112511 | 1.81E-09 | 5.84E-05 |
| Dcp2 | 0.112448 | 3.99E-11 | 1.29E-06 |
| Eno1b | 0.112431 | 1.32E-08 | 0.000425 |
| Ciao2a | 0.112263 | 2.01E-05 | 0.648861 |
| Elac2 | 0.112172 | 2.57E-07 | 0.008297 |
| Mif4gd | 0.111985 | 9.15E-06 | 0.295303 |
| Pgap2 | 0.111979 | 1E-10 | 3.24E-06 |
| Ndufb5 | 0.111951 | 7.19E-07 | 0.023203 |
| Psmb1 | 0.111827 | 1.39E-06 | 0.044868 |
| Arf6 | 0.111525 | 8.48E-06 | 0.273632 |
| Txnrd1 | 0.111417 | 0.000406 | 1 |
| Gtpbp4 | 0.111182 | 3.99E-06 | 0.128839 |
| Anxa4 | 0.111138 | 7.44E-08 | 0.002402 |
| Arih1 | 0.111121 | 1.56E-05 | 0.503955 |
| Gm11808 | 0.111115 | 3.16E-06 | 0.101884 |
| Tuba1c | 0.111001 | 1.11E-05 | 0.357284 |
| Azi2 | 0.11097 | 6.65E-07 | 0.02148 |
| Tes | 0.110898 | 1.4E-06 | 0.04519 |
| Timm10b | 0.110716 | 1.54E-07 | 0.00496 |
| Arpc3 | 0.11043 | 0.000194 | 1 |
| Rfc2 | 0.110411 | 1.88E-05 | 0.60539 |
| Atox1 | 0.110355 | 1.3E-06 | 0.041816 |

| | | | |
|---|---|---|---|
| Tcp1 | 0.110313 | 2.13E-05 | 0.686885 |
| Vamp8 | 0.110286 | 0.000152 | 1 |
| Jpt1 | 0.110199 | 2.93E-06 | 0.094512 |
| B4galt7 | 0.110177 | 1.86E-07 | 0.005997 |
| Dph5 | 0.109887 | 1.77E-07 | 0.005706 |
| Gm11084 | 0.109886 | 5.91E-09 | 0.000191 |
| Snrnp48 | 0.109856 | 0.000487 | 1 |
| Get4 | 0.109833 | 2.29E-08 | 0.00074 |
| Atp2b1 | 0.109715 | 0.000126 | 1 |
| Xrn1 | 0.109539 | 1.95E-06 | 0.063088 |
| Myo1c | 0.109346 | 5.01E-06 | 0.161793 |
| Mpi | 0.109184 | 1.42E-05 | 0.458351 |
| Atp5d | 0.108989 | 6.26E-06 | 0.202263 |
| Mospd2 | 0.108551 | 5.32E-08 | 0.001719 |
| Irf2bp2 | 0.108512 | 0.000512 | 1 |
| Afg3l2 | 0.108361 | 9.71E-06 | 0.313578 |
| Ube2j2 | 0.108296 | 1.64E-07 | 0.005292 |
| Irf8 | 0.108138 | 0.002038 | 1 |
| Nipa2 | 0.108061 | 0.000101 | 1 |
| Atp7a | 0.107751 | 6.04E-06 | 0.194852 |
| Gk | 0.10765 | 7.14E-11 | 2.3E-06 |
| Rabepk | 0.107619 | 1.01E-10 | 3.27E-06 |
| Pgap1 | 0.107618 | 2.79E-07 | 0.008995 |
| Tbc1d14 | 0.107546 | 3.5E-06 | 0.112891 |
| Cdk14 | 0.107532 | 2.06E-05 | 0.665788 |
| Naa50 | 0.107441 | 4.57E-07 | 0.014767 |
| Ddx50 | 0.107417 | 9.18E-05 | 1 |
| Mthfd1l | 0.107405 | 9.3E-06 | 0.300244 |
| Themis2 | 0.107284 | 0.000645 | 1 |
| Csrnp1 | 0.10728 | 0.000239 | 1 |
| Sin3b | 0.107235 | 0.000346 | 1 |
| Mfsd10 | 0.10666 | 4.41E-07 | 0.014222 |
| Ndufa12 | 0.106603 | 5.4E-09 | 0.000174 |
| Atm | 0.10654 | 1.07E-07 | 0.003456 |
| Ncoa7 | 0.106226 | 1.88E-05 | 0.607113 |
| Ddi2 | 0.106146 | 6.34E-07 | 0.020483 |
| Ptpn1 | 0.106028 | 0.000465 | 1 |
| Hcfc1r1 | 0.105984 | 0.000794 | 1 |
| Sf3b3 | 0.105931 | 2.63E-05 | 0.849549 |
| Tasor2 | 0.105771 | 9.56E-06 | 0.308676 |
| Hspa9 | 0.105618 | 1.85E-05 | 0.598577 |
| Pa2g4 | 0.10559 | 5.49E-05 | 1 |
| Esyt2 | 0.105541 | 0.011207 | 1 |
| Mia2 | 0.105523 | 1.92E-05 | 0.621326 |
| Igbp1 | 0.105387 | 9.56E-10 | 3.08E-05 |
| Stxbp2 | 0.105279 | 1.53E-09 | 4.92E-05 |
| A330023F2 | 0.104999 | 3.28E-08 | 0.00106 |
| Tmpo | 0.104714 | 1.23E-06 | 0.039786 |

| | | | |
|---|---|---|---|
| Elk4 | 0.104603 | 1.37E-05 | 0.441219 |
| Fam49a | 0.104508 | 5.74E-05 | 1 |
| Psmc4 | 0.104372 | 1.4E-05 | 0.451504 |
| 1810013L2 | 0.104284 | 4.69E-05 | 1 |
| Tra2a | 0.104081 | 0.000283 | 1 |
| Parl | 0.10385 | 4.54E-06 | 0.146693 |
| Srpk2 | 0.103834 | 3.14E-05 | 1 |
| Sra1 | 0.103699 | 0.001334 | 1 |
| Mre11a | 0.103528 | 1E-06 | 0.032279 |
| Tmbim6 | 0.103527 | 3.92E-06 | 0.126426 |
| Dck | 0.103435 | 3.18E-10 | 1.03E-05 |
| Ppig | 0.103434 | 7.73E-08 | 0.002497 |
| Lclat1 | 0.103416 | 0.000259 | 1 |
| Pld3 | 0.103227 | 0.002136 | 1 |
| Ankrd16 | 0.103192 | 3.4E-08 | 0.001097 |
| Mettl17 | 0.103093 | 1.92E-05 | 0.618517 |
| Aimp1 | 0.102791 | 9.49E-09 | 0.000306 |
| Rars | 0.102734 | 5.22E-09 | 0.000168 |
| Chd4 | 0.102664 | 0.000198 | 1 |
| Mdn1 | 0.10254 | 4.62E-07 | 0.0149 |
| Ndufa8 | 0.102433 | 3.19E-06 | 0.103136 |
| Hnrnpab | 0.102391 | 0.000986 | 1 |
| Mocs1 | 0.102236 | 3.25E-07 | 0.010486 |
| Ggnbp2 | 0.102037 | 2.34E-07 | 0.007539 |
| Sdhd | 0.101782 | 3.51E-07 | 0.011326 |
| Uqcr10 | 0.101714 | 5.9E-06 | 0.190361 |
| Cgas | 0.101563 | 6.58E-05 | 1 |
| Eif5b | 0.101461 | 1.86E-08 | 0.000599 |
| Tmbim4 | 0.101412 | 0.00044 | 1 |
| Nufip2 | 0.101239 | 3.32E-05 | 1 |
| Fam219a | 0.101222 | 1.66E-05 | 0.535361 |
| Ndufc1 | 0.101072 | 1.66E-07 | 0.005371 |
| Runx1 | 0.100845 | 0.000156 | 1 |
| Mrpl12 | 0.100761 | 3.26E-07 | 0.010528 |
| Nfs1 | 0.100757 | 1.24E-05 | 0.399352 |
| Mdh2 | 0.100751 | 2.56E-05 | 0.82766 |
| Nemf | 0.100584 | 7.93E-07 | 0.025595 |
| Gtf2h1 | 0.100558 | 2.9E-08 | 0.000936 |
| Sergef | 0.100445 | 2.41E-06 | 0.077693 |
| Mybbp1a | 0.100328 | 0.000135 | 1 |
| St5 | 0.100226 | 2.03E-06 | 0.065524 |
| Birc2 | 0.100188 | 2.15E-06 | 0.069572 |
| Supt16 | 0.100105 | 7.5E-05 | 1 |
| Tubb4b | 0.100036 | 9.25E-06 | 0.298606 |
| Serbp1 | 0.099987 | 1.51E-05 | 0.485973 |
| Rab22a | 0.099736 | 2.7E-05 | 0.872881 |
| Hmga1 | 0.099618 | 1.7E-10 | 5.48E-06 |
| Myd88 | 0.099466 | 0.000124 | 1 |

| | | | |
|---|---|---|---|
| Phb2 | 0.099446 | 5.88E-06 | 0.189866 |
| Lypla2 | 0.099351 | 3.11E-05 | 1 |
| Suco | 0.099314 | 0.000421 | 1 |
| Clint1 | 0.099286 | 8.17E-05 | 1 |
| Dcun1d5 | 0.099229 | 2.06E-06 | 0.066653 |
| Arhgef18 | 0.099195 | 0.01508 | 1 |
| Eva1a | 0.099147 | 0.000163 | 1 |
| Smchd1 | 0.099076 | 0.00228 | 1 |
| Farsb | 0.098924 | 1.5E-07 | 0.004838 |
| Fen1 | 0.098777 | 4.68E-05 | 1 |
| Srek1 | 0.098678 | 0.001437 | 1 |
| Cyb5b | 0.098661 | 6.76E-08 | 0.002183 |
| Cpeb2 | 0.09848 | 4.88E-05 | 1 |
| Sppl2a | 0.098446 | 0.000182 | 1 |
| Gng10 | 0.098421 | 0.002212 | 1 |
| Washc2 | 0.098377 | 4.55E-06 | 0.146745 |
| Snhg17 | 0.098081 | 1.55E-09 | 5.01E-05 |
| Calm3 | 0.098041 | 1.27E-05 | 0.410441 |
| 4933421O | 0.098018 | 8.81E-10 | 2.84E-05 |
| Fundc2 | 0.097874 | 2.1E-08 | 0.000676 |
| Atp6v1b2 | 0.097845 | 3.33E-05 | 1 |
| Adcy7 | 0.097796 | 1.25E-05 | 0.402898 |
| Mecr | 0.097693 | 6.57E-08 | 0.002122 |
| Tomm7 | 0.097659 | 8.45E-06 | 0.272902 |
| Stk24 | 0.097605 | 0.021622 | 1 |
| Atg16l1 | 0.09736 | 0.000426 | 1 |
| Tcerg1 | 0.097351 | 4.24E-05 | 1 |
| Usp16 | 0.097246 | 6.99E-06 | 0.225784 |
| Ston2 | 0.097135 | 8.23E-07 | 0.026582 |
| Nlrp1b | 0.096909 | 7.39E-08 | 0.002386 |
| Gm12185 | 0.096706 | 1.01E-05 | 0.327645 |
| Ddx21 | 0.096615 | 2.18E-05 | 0.703793 |
| Banf1 | 0.096614 | 1.38E-06 | 0.044588 |
| Slc30a6 | 0.096602 | 3.79E-05 | 1 |
| Rnps1 | 0.096582 | 1.74E-05 | 0.563091 |
| Rnf34 | 0.096031 | 5.25E-06 | 0.16962 |
| Rlim | 0.096007 | 0.000152 | 1 |
| Llph | 0.095887 | 4.62E-06 | 0.149277 |
| Wtap | 0.095877 | 2.91E-06 | 0.094084 |
| Dnaja1 | 0.095476 | 1.76E-06 | 0.056839 |
| Hdac2 | 0.095397 | 1.89E-08 | 0.000612 |
| Hipk1 | 0.09537 | 7.51E-05 | 1 |
| Slc11a2 | 0.0953 | 4.87E-06 | 0.157207 |
| Nfrkb | 0.095112 | 1.08E-05 | 0.347284 |
| Gatad2a | 0.094935 | 5.21E-06 | 0.168228 |
| Apaf1 | 0.094852 | 4.66E-05 | 1 |
| Ptpn2 | 0.094636 | 1.14E-05 | 0.367568 |
| Rrp7a | 0.094607 | 1.09E-06 | 0.035341 |

| | | | |
|---|---|---|---|
| Ube2l3 | 0.094582 | 0.01725 | 1 |
| Arf3 | 0.094557 | 8.21E-06 | 0.265188 |
| Trim27 | 0.094365 | 9.38E-06 | 0.302859 |
| Nedd8 | 0.094352 | 3.59E-05 | 1 |
| Elovl5 | 0.094274 | 5.67E-05 | 1 |
| Ndel1 | 0.094254 | 5.42E-05 | 1 |
| Stx7 | 0.094254 | 9.04E-05 | 1 |
| Hscb | 0.094223 | 3.4E-08 | 0.001099 |
| Lrrfip2 | 0.094183 | 5.66E-06 | 0.182794 |
| Nme1 | 0.094092 | 5.72E-06 | 0.184778 |
| Atad2b | 0.094073 | 1.01E-06 | 0.032575 |
| Srrd | 0.09364 | 1.49E-08 | 0.000482 |
| Ndufb7 | 0.093168 | 0.002058 | 1 |
| Pus10 | 0.093099 | 6.63E-05 | 1 |
| Cmklr1 | 0.093085 | 1.32E-05 | 0.426655 |
| Psma1 | 0.09306 | 0.000307 | 1 |
| Hps3 | 0.093041 | 0.000574 | 1 |
| Ptpn11 | 0.092792 | 9.67E-07 | 0.031234 |
| 1500011B( | 0.092738 | 1.12E-07 | 0.003624 |
| Tecpr1 | 0.092692 | 0.029639 | 1 |
| Pts | 0.092576 | 0.000465 | 1 |
| Pcsk7 | 0.092507 | 2.72E-07 | 0.008794 |
| Shmt2 | 0.092404 | 1.71E-07 | 0.005523 |
| Aif1 | 0.092251 | 0.00423 | 1 |
| Insig1 | 0.092237 | 4.6E-09 | 0.000149 |
| Cdk9 | 0.092198 | 1.28E-06 | 0.041321 |
| Plk3 | 0.092096 | 4.12E-07 | 0.013299 |
| Crybg3 | 0.092082 | 9.51E-06 | 0.306985 |
| Phf5a | 0.091862 | 0.004143 | 1 |
| Snrpd1 | 0.091832 | 3.87E-07 | 0.012502 |
| Rhoh | 0.091569 | 0.009278 | 1 |
| Parp3 | 0.091519 | 1.01E-08 | 0.000327 |
| Psmb6 | 0.091391 | 0.014148 | 1 |
| Mrpl57 | 0.091233 | 0.00086 | 1 |
| Esd | 0.091076 | 1.87E-05 | 0.602822 |
| Bcl2l1 | 0.091044 | 0.000415 | 1 |
| Maco1 | 0.090923 | 4.57E-06 | 0.147593 |
| Zbtb11 | 0.09089 | 1.75E-06 | 0.056617 |
| Tsta3 | 0.090801 | 2.58E-06 | 0.083236 |
| Ndufs6 | 0.090748 | 6.13E-06 | 0.197848 |
| Insyn2b | 0.090566 | 0.000133 | 1 |
| Slc7a5 | 0.09033 | 3.7E-09 | 0.00012 |
| Sbno2 | 0.090315 | 3.68E-05 | 1 |
| Prdx6 | 0.090265 | 2.97E-07 | 0.009593 |
| Cotl1 | 0.090186 | 0.001493 | 1 |
| Ormdl2 | 0.090106 | 1.1E-06 | 0.035387 |
| Ccdc9 | 0.090086 | 1.83E-06 | 0.059067 |
| Olfr111 | 0.090053 | 0.004501 | 1 |

| | | | |
|---|---|---|---|
| Limd1 | 0.089946 | 1.84E-07 | 0.005955 |
| Setd7 | 0.08993 | 2.4E-08 | 0.000776 |
| Ddx27 | 0.08982 | 1.6E-05 | 0.517086 |
| Ubfd1 | 0.089812 | 1.96E-08 | 0.000634 |
| Galns | 0.089734 | 5.58E-10 | 1.8E-05 |
| Trnau1ap | 0.089633 | 1.21E-07 | 0.003908 |
| Glcci1 | 0.089616 | 0.000328 | 1 |
| Nars2 | 0.089495 | 3.87E-06 | 0.12504 |
| Timm13 | 0.089279 | 7.23E-05 | 1 |
| Nol10 | 0.08908 | 4.92E-05 | 1 |
| Ifngr2 | 0.089064 | 0.002485 | 1 |
| Fars2 | 0.088602 | 3.3E-06 | 0.106584 |
| Ppt2 | 0.08849 | 0.000115 | 1 |
| Ndufaf6 | 0.088437 | 1.34E-08 | 0.000432 |
| Fndc3a | 0.088378 | 0.006213 | 1 |
| Mapk1ip1l | 0.088369 | 2.45E-05 | 0.790047 |
| Mrps9 | 0.088151 | 1.46E-05 | 0.470639 |
| Mvb12a | 0.088098 | 2.98E-07 | 0.00961 |
| Cdk6 | 0.088053 | 0.01018 | 1 |
| Wdr41 | 0.087999 | 1.91E-07 | 0.00617 |
| Riok3 | 0.08784 | 2.91E-05 | 0.938131 |
| Wdr74 | 0.087619 | 5.11E-06 | 0.164923 |
| Nt5m | 0.087595 | 1.59E-05 | 0.513859 |
| Ino80 | 0.087475 | 1.29E-09 | 4.16E-05 |
| Pdhb | 0.087439 | 0.000196 | 1 |
| Smg5 | 0.087361 | 4.78E-10 | 1.54E-05 |
| Nr2c2ap | 0.08727 | 1.27E-07 | 0.004101 |
| Sept6 | 0.087265 | 0.000449 | 1 |
| Limk1 | 0.087107 | 0.000189 | 1 |
| Heatr1 | 0.08699 | 1.07E-05 | 0.345777 |
| Sltm | 0.086957 | 4.21E-05 | 1 |
| S100a1 | 0.086956 | 1.15E-08 | 0.000371 |
| Thoc1 | 0.086869 | 0.00041 | 1 |
| Hhat | 0.086805 | 3.51E-06 | 0.113216 |
| Pdia3 | 0.086753 | 0.000183 | 1 |
| Magohb | 0.086727 | 2.14E-09 | 6.9E-05 |
| Lsm4 | 0.086633 | 2.52E-05 | 0.814015 |
| Scand1 | 0.086631 | 3.54E-05 | 1 |
| 4932438A1 | 0.086622 | 4.51E-06 | 0.145574 |
| Slc15a3 | 0.08662 | 6.46E-06 | 0.208481 |
| Rp9 | 0.086506 | 0.000516 | 1 |
| Anxa11 | 0.086499 | 0.004867 | 1 |
| Selenoh | 0.086443 | 3.04E-06 | 0.098089 |
| Tle5 | 0.086391 | 2.06E-05 | 0.666593 |
| Eef1akmt4 | 0.086316 | 3.15E-11 | 1.02E-06 |
| Mpeg1 | 0.086284 | 0.000154 | 1 |
| Dnaaf3 | 0.086259 | 2.92E-11 | 9.44E-07 |
| Pex16 | 0.086139 | 2.74E-07 | 0.008856 |

| | | | |
|---|---|---|---|
| Nin | 0.086066 | 5.73E-06 | 0.184974 |
| Cluh | 0.085917 | 2.66E-11 | 8.59E-07 |
| Pgp | 0.085733 | 1.19E-05 | 0.383497 |
| Kif5b | 0.085722 | 0.000651 | 1 |
| Dlst | 0.085651 | 6.23E-05 | 1 |
| Zfp367 | 0.085641 | 7.52E-07 | 0.024275 |
| Ftsj3 | 0.085519 | 1.45E-07 | 0.004677 |
| Ddost | 0.085489 | 0.001489 | 1 |
| Thoc6 | 0.085417 | 5.04E-10 | 1.63E-05 |
| Trit1 | 0.085401 | 2.35E-05 | 0.759996 |
| Rabl6 | 0.085353 | 0.000348 | 1 |
| Coq2 | 0.085323 | 1.72E-07 | 0.005567 |
| Ndufa2 | 0.085312 | 9.34E-05 | 1 |
| Gm17106 | 0.085306 | 3.47E-06 | 0.112105 |
| Rnf115 | 0.085271 | 0.000633 | 1 |
| Riox1 | 0.085102 | 4.04E-06 | 0.130328 |
| Rad50 | 0.08488 | 1.35E-05 | 0.435114 |
| Spata6 | 0.084826 | 0.000407 | 1 |
| Srsf10 | 0.084809 | 0.000143 | 1 |
| Relch | 0.084765 | 4.33E-05 | 1 |
| Sertad1 | 0.084762 | 8.05E-06 | 0.259889 |
| Cdk11b | 0.084716 | 1.45E-06 | 0.046962 |
| Psat1 | 0.084568 | 1.36E-05 | 0.440656 |
| Setd3 | 0.084409 | 0.000125 | 1 |
| Txndc5 | 0.084336 | 1.91E-07 | 0.006167 |
| Naa20 | 0.084292 | 5.01E-05 | 1 |
| Upf3b | 0.084287 | 0.000421 | 1 |
| Mrpl52 | 0.084178 | 0.000278 | 1 |
| Bmyc | 0.08409 | 0.013777 | 1 |
| Peli2 | 0.083944 | 0.000538 | 1 |
| Cdv3 | 0.083914 | 0.003492 | 1 |
| Scly | 0.083858 | 1.4E-05 | 0.453154 |
| Creb3 | 0.083795 | 0.001365 | 1 |
| Tlk1 | 0.08362 | 8.52E-06 | 0.275227 |
| Pes1 | 0.083557 | 3.79E-07 | 0.012241 |
| Ap2s1 | 0.083388 | 5.78E-06 | 0.18651 |
| Tmem128 | 0.083364 | 8.49E-05 | 1 |
| Sem1 | 0.08317 | 0.000178 | 1 |
| Ube2w | 0.083158 | 0.000136 | 1 |
| Mrpl33 | 0.083157 | 7.85E-05 | 1 |
| Eno1 | 0.083018 | 0.001024 | 1 |
| Psmb7 | 0.082937 | 0.000636 | 1 |
| Zfp945 | 0.08293 | 2.03E-07 | 0.006544 |
| Tomm6 | 0.082743 | 0.000296 | 1 |
| 4933412E1 | 0.082652 | 4.15E-09 | 0.000134 |
| Wdr36 | 0.082647 | 6.86E-05 | 1 |
| Huwe1 | 0.082555 | 0.000627 | 1 |
| Atad3a | 0.082549 | 6.21E-07 | 0.020037 |

| | | | |
|---|---|---|---|
| Med1 | 0.08246 | 0.000117 | 1 |
| Tiparp | 0.082254 | 7.77E-06 | 0.250758 |
| Lamtor5 | 0.082242 | 9.78E-06 | 0.315859 |
| Mak16 | 0.082238 | 0.000127 | 1 |
| Rev3l | 0.081974 | 9.72E-06 | 0.313732 |
| Uimc1 | 0.081946 | 0.003028 | 1 |
| Ndufb6 | 0.081936 | 0.000563 | 1 |
| Ddx10 | 0.081842 | 5.78E-06 | 0.186632 |
| Tmed1 | 0.081133 | 1.66E-08 | 0.000537 |
| Fkbp1a | 0.080932 | 3.01E-05 | 0.973387 |
| Birc6 | 0.080915 | 0.000806 | 1 |
| Ndufs5 | 0.080877 | 0.000718 | 1 |
| Mrps16 | 0.080869 | 9.99E-05 | 1 |
| Tiam1 | 0.080852 | 0.000478 | 1 |
| Prrc2c | 0.080732 | 0.002657 | 1 |
| Grwd1 | 0.080713 | 1.65E-06 | 0.053374 |
| Tut7 | 0.080706 | 0.00012 | 1 |
| Atg2a | 0.080679 | 0.003045 | 1 |
| Tec | 0.080421 | 0.000677 | 1 |
| Szrd1 | 0.080344 | 2.99E-05 | 0.964195 |
| Batf | 0.080264 | 1.9E-10 | 6.14E-06 |
| Igf2bp3 | 0.080252 | 1.99E-11 | 6.43E-07 |
| Kri1 | 0.080242 | 3.87E-07 | 0.012492 |
| Mydgf | 0.080121 | 0.000428 | 1 |
| Ubp1 | 0.08007 | 4.62E-08 | 0.001492 |
| Trib1 | 0.08007 | 0.000521 | 1 |
| Cast | 0.080034 | 0.001142 | 1 |
| Arf5 | 0.079981 | 2.27E-05 | 0.73239 |
| Pum3 | 0.079875 | 1.08E-06 | 0.034842 |
| Psma6 | 0.079871 | 4.11E-05 | 1 |
| Nrd1 | 0.079776 | 2.55E-06 | 0.082372 |
| Gm43813 | 0.079768 | 8.41E-08 | 0.002716 |
| Sertad2 | 0.079642 | 1.9E-06 | 0.06142 |
| Tfpi | 0.079625 | 0.003383 | 1 |
| Swt1 | 0.07961 | 0.000308 | 1 |
| Stx8 | 0.07948 | 0.000414 | 1 |
| Ttll5 | 0.079476 | 2.54E-06 | 0.082131 |
| Cops9 | 0.079021 | 1.03E-06 | 0.033206 |
| Tmem97 | 0.079001 | 1.43E-06 | 0.046318 |
| Srsf11 | 0.078992 | 0.000325 | 1 |
| Med21 | 0.078964 | 6.82E-05 | 1 |
| Fdps | 0.078863 | 0.000236 | 1 |
| Abca7 | 0.078837 | 0.000226 | 1 |
| Gar1 | 0.078792 | 3.35E-06 | 0.108062 |
| Rabgef1 | 0.078753 | 0.000309 | 1 |
| Parp4 | 0.078629 | 0.000346 | 1 |
| Rps27l | 0.07858 | 0.000198 | 1 |
| Slc20a1 | 0.078555 | 0.000491 | 1 |

| | | | |
|---|---|---|---|
| Myl6 | 0.078411 | 0.000177 | 1 |
| Tent5c | 0.078355 | 0.045067 | 1 |
| Mob3c | 0.078327 | 0.000146 | 1 |
| Snrpd3 | 0.078223 | 0.000181 | 1 |
| Fos | 0.078133 | 1.74E-06 | 0.056131 |
| Tyms | 0.07809 | 0.028414 | 1 |
| Lsm14a | 0.07807 | 0.000298 | 1 |
| Gm15283 | 0.07803 | 0.001498 | 1 |
| Slc25a38 | 0.077999 | 7.31E-07 | 0.023613 |
| Mta3 | 0.077884 | 0.002341 | 1 |
| Chd8 | 0.077827 | 0.015921 | 1 |
| Cops7a | 0.077795 | 0.000132 | 1 |
| Cog4 | 0.077641 | 0.000148 | 1 |
| Ipo7 | 0.077639 | 0.008315 | 1 |
| Cox17 | 0.077256 | 0.001586 | 1 |
| Arel1 | 0.07718 | 2.92E-06 | 0.094171 |
| Psmc6 | 0.077105 | 0.00229 | 1 |
| Tbcd | 0.077027 | 0.00013 | 1 |
| Wrn | 0.077019 | 7.79E-05 | 1 |
| Rnf168 | 0.076971 | 0.000423 | 1 |
| Mycbp2 | 0.076958 | 0.002633 | 1 |
| Acy1 | 0.076926 | 1.65E-06 | 0.053252 |
| Vdac2 | 0.076898 | 0.004128 | 1 |
| Usp21 | 0.076826 | 0.000311 | 1 |
| Eif2ak4 | 0.076821 | 1.4E-05 | 0.451387 |
| Klhl2 | 0.07682 | 4.85E-09 | 0.000157 |
| Myo1g | 0.076786 | 7.44E-09 | 0.00024 |
| Ten1 | 0.076712 | 0.00013 | 1 |
| Thrap3 | 0.076616 | 0.001026 | 1 |
| Lztfl1 | 0.076532 | 0.001804 | 1 |
| Supt20 | 0.076354 | 0.002334 | 1 |
| Tmem181a | 0.076353 | 0.002368 | 1 |
| Htatip2 | 0.076172 | 1.06E-10 | 3.42E-06 |
| Ccar1 | 0.076042 | 9.69E-05 | 1 |
| Sept11 | 0.076036 | 0.121353 | 1 |
| Cd34 | 0.075983 | 2.5E-06 | 0.080866 |
| Cdt1 | 0.075918 | 0.001665 | 1 |
| Ywhag | 0.075676 | 0.012214 | 1 |
| Mapk6 | 0.075662 | 0.000182 | 1 |
| Fhad1 | 0.075632 | 7.46E-06 | 0.240964 |
| Atrx | 0.07547 | 9.65E-06 | 0.311419 |
| 0610030E2 | 0.075424 | 0.0005 | 1 |
| Timm17a | 0.075393 | 0.0037 | 1 |
| Tmem242 | 0.075367 | 2.92E-07 | 0.009421 |
| Nop58 | 0.075164 | 2.06E-05 | 0.663547 |
| Myo9a | 0.075136 | 0.000118 | 1 |
| Cyth1 | 0.075088 | 4.23E-05 | 1 |
| Ltv1 | 0.074986 | 8.39E-07 | 0.027099 |

| | | | |
|---|---|---|---|
| Slc38a1 | 0.07497 | 0.001265 | 1 |
| Prune2 | 0.074872 | 3.1E-06 | 0.099997 |
| Lmbr1l | 0.074866 | 0.000105 | 1 |
| Phb | 0.074833 | 0.000166 | 1 |
| Arl6ip4 | 0.074817 | 0.000277 | 1 |
| Kdm4b | 0.074807 | 1.17E-05 | 0.378684 |
| Atg7 | 0.074701 | 1.68E-06 | 0.054127 |
| Epb41 | 0.07467 | 0.003593 | 1 |
| Wdr75 | 0.074632 | 0.000446 | 1 |
| Dera | 0.074406 | 0.000224 | 1 |
| Zkscan6 | 0.074305 | 1.83E-05 | 0.59147 |
| Fdx2 | 0.074277 | 3.56E-07 | 0.011485 |
| Cyp2r1 | 0.074205 | 9.74E-07 | 0.031435 |
| Ccdc58 | 0.074093 | 0.002876 | 1 |
| Myl12a | 0.074017 | 0.000809 | 1 |
| Vti1a | 0.073973 | 0.000497 | 1 |
| Pbdc1 | 0.073871 | 6.91E-07 | 0.022322 |
| B3gntl1 | 0.073746 | 0.00097 | 1 |
| Inpp5b | 0.073686 | 0.000473 | 1 |
| Mad2l2 | 0.073654 | 5.19E-05 | 1 |
| Fbrs | 0.073597 | 0.000134 | 1 |
| Meis3 | 0.073484 | 0.000467 | 1 |
| Prpf38a | 0.073396 | 5.34E-07 | 0.017226 |
| Urm1 | 0.073394 | 4.05E-06 | 0.130877 |
| Dctpp1 | 0.073387 | 9.98E-05 | 1 |
| Nup153 | 0.073328 | 3.07E-06 | 0.099238 |
| Kmt2a | 0.073285 | 0.000846 | 1 |
| Jak2 | 0.073211 | 5.12E-06 | 0.165245 |
| Cracr2a | 0.073156 | 0.004334 | 1 |
| Chchd1 | 0.072851 | 5.7E-06 | 0.183869 |
| Itpk1 | 0.072766 | 0.000331 | 1 |
| Stat3 | 0.072675 | 0.002833 | 1 |
| Arrdc4 | 0.072618 | 7.24E-07 | 0.023386 |
| Snrpa1 | 0.072596 | 2.79E-06 | 0.089918 |
| Srp9 | 0.072451 | 0.025417 | 1 |
| Ugcg | 0.072372 | 5.36E-05 | 1 |
| Gltp | 0.072297 | 0.003356 | 1 |
| Nle1 | 0.07223 | 1.52E-05 | 0.489728 |
| Becn1 | 0.072223 | 0.000215 | 1 |
| 5430427O | 0.072179 | 1.29E-05 | 0.415564 |
| Ikzf2 | 0.072123 | 5.16E-05 | 1 |
| Gadd45gip | 0.072011 | 0.00011 | 1 |
| Fam89a | 0.071957 | 3.65E-08 | 0.00118 |
| Rab8a | 0.07189 | 0.01175 | 1 |
| Stx12 | 0.071786 | 2.69E-05 | 0.868103 |
| Gm42031 | 0.071578 | 0.000171 | 1 |
| Mettl6 | 0.071425 | 9.61E-06 | 0.310358 |
| Nasp | 0.071415 | 0.002947 | 1 |

| | | | |
|---|---|---|---|
| Tmem39a | 0.071348 | 1.85E-05 | 0.598154 |
| H2afz | 0.071277 | 0.028301 | 1 |
| Uqcrb | 0.071277 | 0.000167 | 1 |
| Ssr3 | 0.071252 | 0.006064 | 1 |
| Mcph1 | 0.071185 | 0.002704 | 1 |
| Notch1 | 0.071125 | 0.001023 | 1 |
| Rpf2 | 0.071049 | 1.5E-06 | 0.048322 |
| Ell2 | 0.071011 | 3.22E-05 | 1 |
| Lsm12 | 0.070919 | 0.002744 | 1 |
| Ctdnep1 | 0.070649 | 0.000306 | 1 |
| B230219D2 | 0.070603 | 2.8E-05 | 0.903113 |
| Srsf3 | 0.070576 | 0.064053 | 1 |
| Rnf4 | 0.070573 | 0.014397 | 1 |
| Bcat2 | 0.070534 | 7.9E-07 | 0.025516 |
| Srpk1 | 0.07032 | 0.008412 | 1 |
| Pik3r6 | 0.070242 | 0.000175 | 1 |
| Ccdc84 | 0.070209 | 6.01E-07 | 0.019417 |
| Rsl24d1 | 0.070188 | 3.4E-05 | 1 |
| Ei24 | 0.070173 | 2.15E-05 | 0.694419 |
| Manba | 0.070166 | 0.000788 | 1 |
| Agpat5 | 0.070066 | 4.7E-06 | 0.151804 |
| Bola3 | 0.069971 | 5.28E-11 | 1.71E-06 |
| Mrpl20 | 0.069783 | 1.31E-05 | 0.422401 |
| Fam222b | 0.069693 | 4.34E-05 | 1 |
| Phc3 | 0.069642 | 0.062515 | 1 |
| Ndor1 | 0.069617 | 0.00012 | 1 |
| Ywhab | 0.069512 | 0.014835 | 1 |
| Plaa | 0.069465 | 1.25E-06 | 0.040269 |
| Aurkaip1 | 0.069333 | 1.1E-05 | 0.355838 |
| Cacnb2 | 0.069323 | 0.000332 | 1 |
| Eps15 | 0.069276 | 0.001117 | 1 |
| Exosc1 | 0.069251 | 7.73E-07 | 0.024945 |
| Lsg1 | 0.069192 | 0.000158 | 1 |
| Igf1r | 0.06915 | 0.030218 | 1 |
| Hipk2 | 0.069138 | 5.99E-06 | 0.193268 |
| Tmem123 | 0.069053 | 2.78E-08 | 0.000899 |
| Tma7 | 0.06894 | 0.003988 | 1 |
| Tecpr2 | 0.068885 | 5.2E-10 | 1.68E-05 |
| Atic | 0.068848 | 0.009816 | 1 |
| Nol12 | 0.068637 | 1.95E-06 | 0.062893 |
| Ube2s | 0.068424 | 0.000247 | 1 |
| Aco2 | 0.068398 | 4.14E-05 | 1 |
| Msh3 | 0.068393 | 9.23E-06 | 0.298087 |
| Aldh18a1 | 0.068226 | 1.48E-05 | 0.477415 |
| Dhps | 0.068099 | 9.73E-08 | 0.003142 |
| Il13ra1 | 0.068014 | 0.001119 | 1 |
| Utp3 | 0.067873 | 0.000293 | 1 |
| Strn4 | 0.06784 | 8.04E-05 | 1 |

| | | | |
|---|---|---|---|
| Osbpl8 | 0.06783 | 0.007786 | 1 |
| Vipas39 | 0.067817 | 0.000255 | 1 |
| Agfg2 | 0.067549 | 4.95E-05 | 1 |
| Utp4 | 0.06753 | 8.1E-09 | 0.000261 |
| Slc25a39 | 0.067503 | 0.001046 | 1 |
| Ctu2 | 0.067416 | 4.59E-05 | 1 |
| Nolc1 | 0.067321 | 0.000245 | 1 |
| Tcirg1 | 0.067189 | 0.000658 | 1 |
| Arap2 | 0.067148 | 0.001109 | 1 |
| Med15 | 0.067121 | 1.11E-05 | 0.359932 |
| Aak1 | 0.067075 | 0.555672 | 1 |
| Cmas | 0.066959 | 0.000106 | 1 |
| Morf4l2 | 0.066908 | 0.035502 | 1 |
| Nip7 | 0.066838 | 0.000114 | 1 |
| Rpe | 0.066838 | 2.06E-05 | 0.663472 |
| Cyb5r4 | 0.06676 | 0.001208 | 1 |
| Eif1ad | 0.066733 | 2.04E-05 | 0.659964 |
| Tnip1 | 0.066726 | 0.000114 | 1 |
| Ddx46 | 0.066645 | 0.000622 | 1 |
| Rlf | 0.066558 | 0.019241 | 1 |
| Kdm6b | 0.066479 | 0.101761 | 1 |
| Trap1 | 0.066417 | 0.00522 | 1 |
| Eif2b5 | 0.066332 | 5.42E-05 | 1 |
| Mical1 | 0.06631 | 0.005295 | 1 |
| Foxo1 | 0.066251 | 0.392785 | 1 |
| Cndp2 | 0.066169 | 0.000647 | 1 |
| Ints9 | 0.066087 | 8.35E-06 | 0.269683 |
| Serpinb9 | 0.065928 | 0.000104 | 1 |
| Dnajc11 | 0.065806 | 5.59E-05 | 1 |
| Cdc37 | 0.065793 | 0.004246 | 1 |
| Rngtt | 0.065792 | 0.007838 | 1 |
| Baiap2 | 0.065712 | 0.000945 | 1 |
| Paox | 0.065708 | 8.1E-06 | 0.261537 |
| Nubp2 | 0.065706 | 1.03E-06 | 0.033225 |
| Dusp11 | 0.065628 | 0.000289 | 1 |
| Nup98 | 0.065616 | 0.000237 | 1 |
| Psmb3 | 0.065607 | 0.000901 | 1 |
| Rab29 | 0.065596 | 1.35E-05 | 0.436895 |
| Sult1a1 | 0.065474 | 0.609741 | 1 |
| Nop9 | 0.065445 | 0.000313 | 1 |
| Ap1g2 | 0.065202 | 0.000162 | 1 |
| Gfer | 0.065188 | 0.002063 | 1 |
| Prdx2 | 0.065174 | 0.00011 | 1 |
| Tgs1 | 0.065087 | 3.69E-05 | 1 |
| Dhodh | 0.065081 | 4.34E-07 | 0.014004 |
| Ap1s1 | 0.064904 | 0.004877 | 1 |
| Osgin2 | 0.064868 | 2.66E-05 | 0.858292 |
| Tg | 0.064835 | 0.015173 | 1 |

| | | | |
|---|---|---|---|
| Ankrd17 | 0.064753 | 0.00313 | 1 |
| Aff1 | 0.064711 | 0.002995 | 1 |
| Bri3 | 0.064644 | 4.77E-08 | 0.00154 |
| Ndufa7 | 0.064616 | 0.000784 | 1 |
| Slc9a8 | 0.06455 | 6.01E-05 | 1 |
| Rbx1 | 0.064527 | 0.012899 | 1 |
| Nupl2 | 0.064522 | 2.43E-06 | 0.078402 |
| Ddx24 | 0.064456 | 9.55E-05 | 1 |
| Atg4a | 0.064421 | 0.000214 | 1 |
| Msto1 | 0.064397 | 0.000196 | 1 |
| Snhg8 | 0.064191 | 3.91E-05 | 1 |
| Tbk1 | 0.064186 | 0.000981 | 1 |
| Tsr3 | 0.064141 | 0.002939 | 1 |
| Ethe1 | 0.064037 | 5.32E-06 | 0.171808 |
| Abcf1 | 0.063993 | 0.000456 | 1 |
| D530033B1 | 0.063992 | 3.04E-06 | 0.098296 |
| Jade2 | 0.063902 | 0.017427 | 1 |
| Ahctf1 | 0.063874 | 0.000733 | 1 |
| Dhdh | 0.063661 | 1.23E-05 | 0.395738 |
| Lsm5 | 0.063646 | 0.013436 | 1 |
| Gmeb2 | 0.063532 | 0.006014 | 1 |
| Mrpl21 | 0.063462 | 2.51E-05 | 0.811508 |
| Zfp655 | 0.063409 | 0.002577 | 1 |
| Map2k7 | 0.063382 | 1.09E-05 | 0.350441 |
| Kat2b | 0.063244 | 0.000156 | 1 |
| Wdr18 | 0.063185 | 3.63E-05 | 1 |
| Aph1a | 0.063087 | 0.00032 | 1 |
| Ndufa11 | 0.063073 | 0.001085 | 1 |
| Foxj3 | 0.06306 | 0.000112 | 1 |
| CAAA0111 | 0.063038 | 2.58E-05 | 0.832074 |
| Ogfrl1 | 0.063027 | 0.004316 | 1 |
| Ears2 | 0.062931 | 0.017947 | 1 |
| Nat10 | 0.062921 | 1.12E-05 | 0.361867 |
| Srp14 | 0.062839 | 0.002758 | 1 |
| Tmem268 | 0.062705 | 0.00072 | 1 |
| Slc10a7 | 0.062655 | 0.030846 | 1 |
| Ankrd12 | 0.062629 | 0.001545 | 1 |
| Timm9 | 0.062432 | 0.00579 | 1 |
| Rrp15 | 0.062424 | 5.56E-06 | 0.179519 |
| Ppm1g | 0.062235 | 0.00939 | 1 |
| Mrpl18 | 0.062196 | 3.21E-06 | 0.103547 |
| Ptges2 | 0.062194 | 7.96E-05 | 1 |
| N4bp1 | 0.062108 | 0.001704 | 1 |
| Hnrnpa3 | 0.062087 | 0.000515 | 1 |
| Arfgef1 | 0.062085 | 0.001585 | 1 |
| Tmem189 | 0.06206 | 0.004717 | 1 |
| Ascc3 | 0.06191 | 0.003762 | 1 |
| Gm20275 | 0.061695 | 6.93E-05 | 1 |

| | | | |
|---|---|---|---|
| Acd | 0.061682 | 4.83E-05 | 1 |
| Dnaja2 | 0.061643 | 0.001783 | 1 |
| Med14 | 0.061582 | 0.002976 | 1 |
| Tmem164 | 0.061484 | 0.000436 | 1 |
| Nufip1 | 0.061478 | 0.000239 | 1 |
| Zfp36l2 | 0.061376 | 0.743397 | 1 |
| Vwa8 | 0.06135 | 2.22E-05 | 0.718317 |
| Ptprj | 0.061311 | 0.002481 | 1 |
| Ivd | 0.061083 | 0.000974 | 1 |
| Mthfd2 | 0.061069 | 8.82E-06 | 0.284614 |
| Klf2 | 0.060896 | 7.11E-07 | 0.02297 |
| Ero1l | 0.060896 | 4.88E-06 | 0.157392 |
| Nsun6 | 0.060884 | 0.074854 | 1 |
| Pms2 | 0.060878 | 0.009094 | 1 |
| Tsr1 | 0.060788 | 4.11E-05 | 1 |
| Bop1 | 0.060698 | 1.99E-05 | 0.643017 |
| Mafb | 0.060548 | 0.269344 | 1 |
| Pold4 | 0.06051 | 0.000713 | 1 |
| Ipo4 | 0.060454 | 0.004925 | 1 |
| Lsp1 | 0.060211 | 0.141901 | 1 |
| Manf | 0.060069 | 0.004041 | 1 |
| Mppe1 | 0.059885 | 0.000797 | 1 |
| Slc39a14 | 0.05973 | 1.32E-05 | 0.426517 |
| Etnk1 | 0.059687 | 0.005301 | 1 |
| Ppp1r12c | 0.059674 | 7.05E-05 | 1 |
| Inpp4a | 0.059641 | 8.17E-06 | 0.263849 |
| Zc3h13 | 0.059639 | 6.54E-07 | 0.021109 |
| Mpp6 | 0.059465 | 0.000135 | 1 |
| Cep78 | 0.059336 | 0.000669 | 1 |
| Snrnp35 | 0.059326 | 0.000104 | 1 |
| Pno1 | 0.059299 | 2.39E-05 | 0.772989 |
| Fam3c | 0.059294 | 0.001174 | 1 |
| Mdm2 | 0.059281 | 0.060758 | 1 |
| Mfsd5 | 0.059153 | 0.002206 | 1 |
| Slc35b3 | 0.059139 | 0.000173 | 1 |
| Nckipsd | 0.058788 | 0.002399 | 1 |
| Src | 0.058656 | 2.26E-06 | 0.073017 |
| Ascc2 | 0.058655 | 4.14E-06 | 0.133737 |
| Mrpl2 | 0.058622 | 0.000268 | 1 |
| Stx18 | 0.058614 | 0.000152 | 1 |
| Fyb | 0.05857 | 0.019247 | 1 |
| Lrrc4 | 0.05855 | 0.000149 | 1 |
| Tmed8 | 0.058546 | 0.00016 | 1 |
| Prkx | 0.058453 | 0.011631 | 1 |
| Rer1 | 0.058336 | 0.000183 | 1 |
| Mrpl23 | 0.058258 | 0.000378 | 1 |
| Gm16286 | 0.058195 | 0.004699 | 1 |
| Fam162a | 0.058193 | 7.41E-07 | 0.023922 |

| | | | |
|---|---|---|---|
| Tet2 | 0.058157 | 0.004037 | 1 |
| Ppif | 0.058154 | 3.51E-05 | 1 |
| Itpr3 | 0.058093 | 8.8E-05 | 1 |
| Rnft1 | 0.057982 | 0.001302 | 1 |
| Dazap1 | 0.057868 | 0.003373 | 1 |
| Erp44 | 0.057867 | 0.000514 | 1 |
| Slc35d2 | 0.057819 | 8.73E-05 | 1 |
| Ints6 | 0.057792 | 5.12E-05 | 1 |
| Slc12a7 | 0.057754 | 6.26E-05 | 1 |
| Wapl | 0.057724 | 0.002296 | 1 |
| 2610507B1 | 0.057666 | 0.00018 | 1 |
| Rnaseh2b | 0.05765 | 9.87E-06 | 0.318751 |
| Snap23 | 0.057636 | 0.001058 | 1 |
| Zmiz2 | 0.057611 | 0.00133 | 1 |
| Tspan13 | 0.057582 | 0.374455 | 1 |
| March2 | 0.057546 | 2.95E-05 | 0.953624 |
| Rab8b | 0.057507 | 0.051483 | 1 |
| Rbck1 | 0.057344 | 0.003138 | 1 |
| Nipsnap3b | 0.057326 | 0.0002 | 1 |
| Pphln1 | 0.057325 | 0.000216 | 1 |
| Fbxw2 | 0.057233 | 0.004733 | 1 |
| Ncf4 | 0.057215 | 0.019867 | 1 |
| Snx1 | 0.057194 | 0.000679 | 1 |
| Uhrf1bp1 | 0.057154 | 1.31E-09 | 4.24E-05 |
| Golga4 | 0.057133 | 0.000123 | 1 |
| Pgls | 0.057075 | 0.001514 | 1 |
| Commd4 | 0.057038 | 0.002124 | 1 |
| Trim12c | 0.057009 | 0.000389 | 1 |
| Dhcr7 | 0.056948 | 3.41E-06 | 0.110004 |
| Pycard | 0.05688 | 0.056705 | 1 |
| Mink1 | 0.056768 | 0.013618 | 1 |
| Stk19 | 0.056729 | 0.000277 | 1 |
| Akt3 | 0.056681 | 2.6E-05 | 0.839509 |
| Hmbs | 0.056492 | 0.001266 | 1 |
| Trmt61a | 0.056465 | 0.000433 | 1 |
| Tmsb4x | 0.056432 | 0.41423 | 1 |
| Trappc6a | 0.056397 | 0.000308 | 1 |
| Mob1b | 0.05639 | 0.000779 | 1 |
| Dnajb1 | 0.05637 | 0.004448 | 1 |
| Dkc1 | 0.056348 | 8.3E-05 | 1 |
| Tmem167 | 0.056323 | 0.010761 | 1 |
| Ambra1 | 0.056246 | 0.00509 | 1 |
| Pde4a | 0.056159 | 7.19E-05 | 1 |
| Nus1 | 0.05614 | 4.57E-05 | 1 |
| Mtmr14 | 0.05612 | 0.00038 | 1 |
| Rab32 | 0.056083 | 0.010123 | 1 |
| Ttc27 | 0.05603 | 0.000506 | 1 |
| Mrpl45 | 0.055741 | 0.008591 | 1 |

| | | | |
|---|---|---|---|
| Hgsnat | 0.055716 | 0.000285 | 1 |
| Cyp4v3 | 0.055631 | 0.019302 | 1 |
| Tox4 | 0.055622 | 0.000138 | 1 |
| Eif3j1 | 0.055603 | 0.001008 | 1 |
| Ppm1b | 0.055573 | 0.355813 | 1 |
| Gipc1 | 0.055559 | 0.000428 | 1 |
| Arv1 | 0.055515 | 3.88E-07 | 0.012543 |
| Plekha5 | 0.055477 | 0.000535 | 1 |
| Cct6a | 0.055335 | 0.086295 | 1 |
| Trub2 | 0.05521 | 0.000814 | 1 |
| Tor1b | 0.055059 | 0.011674 | 1 |
| Rrp9 | 0.05493 | 1.57E-05 | 0.505286 |
| Lmnb1 | 0.054875 | 0.000118 | 1 |
| Odf2 | 0.054781 | 0.006758 | 1 |
| Nol9 | 0.054737 | 0.000952 | 1 |
| Churc1 | 0.054713 | 0.000417 | 1 |
| Trmt10c | 0.054652 | 0.003414 | 1 |
| Sik3 | 0.054635 | 7.63E-05 | 1 |
| Snx6 | 0.054632 | 0.010369 | 1 |
| Bcl6 | 0.054615 | 0.031822 | 1 |
| Spout1 | 0.054551 | 8.71E-05 | 1 |
| Mri1 | 0.054512 | 5.73E-06 | 0.185013 |
| St8sia1 | 0.054411 | 1.59E-06 | 0.051249 |
| Micos10 | 0.054315 | 0.005342 | 1 |
| Utp18 | 0.054255 | 2.37E-06 | 0.076519 |
| Elof1 | 0.05422 | 0.000119 | 1 |
| Gdap2 | 0.05413 | 0.000275 | 1 |
| Rnf180 | 0.054034 | 0.002051 | 1 |
| Serhl | 0.053959 | 0.000538 | 1 |
| Triobp | 0.053909 | 0.001075 | 1 |
| Smdt1 | 0.053849 | 0.000199 | 1 |
| Tent4b | 0.053842 | 0.000485 | 1 |
| Cox14 | 0.053801 | 0.000995 | 1 |
| Ddx39b | 0.053542 | 0.001731 | 1 |
| Guk1 | 0.05349 | 0.000139 | 1 |
| Mbp | 0.053406 | 0.000411 | 1 |
| Rras | 0.053358 | 3.96E-07 | 0.012773 |
| Polr3c | 0.053346 | 0.005439 | 1 |
| Acaca | 0.053078 | 1.96E-05 | 0.631857 |
| Lrrc8d | 0.053035 | 4.89E-05 | 1 |
| Nt5c3 | 0.052991 | 0.000607 | 1 |
| Sart3 | 0.052833 | 0.001649 | 1 |
| Ints12 | 0.052799 | 0.001646 | 1 |
| Iars2 | 0.052736 | 0.002046 | 1 |
| Zc3h15 | 0.052728 | 0.034512 | 1 |
| Jtb | 0.052659 | 0.017005 | 1 |
| Cope | 0.052626 | 0.292504 | 1 |
| Mrps10 | 0.052615 | 0.001124 | 1 |

| | | | |
|---|---|---|---|
| Ccz1 | 0.052566 | 0.003286 | 1 |
| Baz1a | 0.052507 | 0.000343 | 1 |
| Pdcd5 | 0.052452 | 0.012167 | 1 |
| Pitpnm1 | 0.052343 | 0.000789 | 1 |
| Heatr6 | 0.052318 | 0.001579 | 1 |
| 3110082I1 | 0.052241 | 0.007235 | 1 |
| Tfdp1 | 0.052241 | 0.004388 | 1 |
| Selenoo | 0.052112 | 0.001526 | 1 |
| Dnajc17 | 0.052016 | 0.00076 | 1 |
| Alg3 | 0.051945 | 0.000153 | 1 |
| Agpat2 | 0.05188 | 2.05E-06 | 0.066262 |
| Raet1e | 0.051865 | 0.00021 | 1 |
| Ece1 | 0.051863 | 0.00112 | 1 |
| Mrpl35 | 0.051737 | 0.000228 | 1 |
| Lnpep | 0.051644 | 0.020021 | 1 |
| Arhgdib | 0.051637 | 0.058643 | 1 |
| Fam111a | 0.051605 | 0.000168 | 1 |
| Cyld | 0.051499 | 0.001909 | 1 |
| Nln | 0.051448 | 5.49E-05 | 1 |
| Ewsr1 | 0.051438 | 0.002544 | 1 |
| Ppm1m | 0.05134 | 0.000435 | 1 |
| Brms1 | 0.051336 | 0.000124 | 1 |
| Nsun2 | 0.05116 | 0.003632 | 1 |
| Ube2v1 | 0.051084 | 0.015345 | 1 |
| Esyt1 | 0.051053 | 0.029408 | 1 |
| Adpgk | 0.050916 | 2.44E-05 | 0.788346 |
| Cbx5 | 0.050884 | 0.000637 | 1 |
| Cdyl2 | 0.050783 | 0.000521 | 1 |
| C1rl | 0.050724 | 6.04E-05 | 1 |
| Ube2f | 0.050537 | 0.003903 | 1 |
| Rnpep | 0.050487 | 0.012965 | 1 |
| Hlx | 0.05048 | 2.2E-05 | 0.711406 |
| Elf2 | 0.050439 | 0.006104 | 1 |
| Herpud1 | 0.050333 | 0.202961 | 1 |
| Yars2 | 0.050104 | 0.000862 | 1 |
| Nploc4 | 0.050027 | 6.3E-05 | 1 |
| Taf13 | 0.049885 | 4.01E-06 | 0.129362 |
| Plagl2 | 0.049826 | 0.00101 | 1 |
| Bloc1s2 | 0.04982 | 0.008454 | 1 |
| Cdk12 | 0.049646 | 0.021624 | 1 |
| Bin3 | 0.049634 | 0.000106 | 1 |
| Tars2 | 0.049476 | 0.000521 | 1 |
| Cd2bp2 | 0.049459 | 0.000972 | 1 |
| Prpf39 | 0.049438 | 0.000994 | 1 |
| Tmem126a | 0.049337 | 0.019907 | 1 |
| Pcyt2 | 0.049322 | 0.012313 | 1 |
| Fam104a | 0.049302 | 0.002694 | 1 |
| Mxd1 | 0.049295 | 0.005227 | 1 |

| | | | |
|---|---|---|---|
| Lrpprc | 0.04918 | 0.018566 | 1 |
| Pgm1 | 0.049118 | 0.000604 | 1 |
| Smc2 | 0.049108 | 2.31E-05 | 0.747323 |
| Ube2d1 | 0.049091 | 0.000571 | 1 |
| Cwc27 | 0.049066 | 2.23E-05 | 0.720518 |
| Ptprc | 0.049012 | 0.381522 | 1 |
| Mms19 | 0.048892 | 0.001545 | 1 |
| Sprydd7 | 0.048841 | 0.034775 | 1 |
| Prpf4 | 0.048729 | 3.93E-05 | 1 |
| Os9 | 0.048631 | 0.006568 | 1 |
| Usp3 | 0.048608 | 0.061485 | 1 |
| Nudt21 | 0.048551 | 0.002502 | 1 |
| Ube2g2 | 0.048524 | 0.000891 | 1 |
| Soat1 | 0.048516 | 0.016799 | 1 |
| Pola2 | 0.048505 | 0.00144 | 1 |
| Mier1 | 0.048428 | 0.00124 | 1 |
| Arid4b | 0.048349 | 0.019726 | 1 |
| Ern1 | 0.048269 | 0.002912 | 1 |
| Kars | 0.04822 | 0.000202 | 1 |
| Mcoln1 | 0.047915 | 6.25E-05 | 1 |
| Ttc7b | 0.047884 | 0.095799 | 1 |
| Prelid3b | 0.04788 | 0.013112 | 1 |
| Sp4 | 0.047839 | 0.019237 | 1 |
| Pgam5 | 0.047825 | 0.002523 | 1 |
| Mapk9 | 0.047811 | 5.41E-07 | 0.017456 |
| Rbm28 | 0.047789 | 0.008606 | 1 |
| Rap1gap2 | 0.047693 | 0.070225 | 1 |
| Mrpl15 | 0.047636 | 0.001158 | 1 |
| Mitf | 0.047587 | 0.057552 | 1 |
| Eef1e1 | 0.047567 | 0.00659 | 1 |
| Dap3 | 0.047561 | 0.000637 | 1 |
| Jmjd6 | 0.047502 | 0.002774 | 1 |
| Ncor1 | 0.047472 | 0.055344 | 1 |
| Taf3 | 0.047414 | 0.014824 | 1 |
| Chchd4 | 0.047365 | 0.000643 | 1 |
| Ildr1 | 0.047235 | 0.006788 | 1 |
| Tmem87a | 0.047213 | 0.000223 | 1 |
| Vapb | 0.04721 | 0.001203 | 1 |
| Psmd1 | 0.047186 | 0.000258 | 1 |
| Eif1ax | 0.047147 | 0.000673 | 1 |
| Def6 | 0.047139 | 0.038964 | 1 |
| Htatsf1 | 0.047101 | 0.000527 | 1 |
| Trim30c | 0.046947 | 0.014514 | 1 |
| Vps50 | 0.046894 | 0.007377 | 1 |
| Hook2 | 0.046827 | 0.00013 | 1 |
| Slamf9 | 0.046725 | 0.000585 | 1 |
| Acsl5 | 0.046714 | 0.009263 | 1 |
| Pkig | 0.046575 | 0.011445 | 1 |

| | | | |
|---|---|---|---|
| Cenpx | 0.046479 | 0.004743 | 1 |
| Brix1 | 0.046434 | 0.000408 | 1 |
| Glb1 | 0.046393 | 0.006002 | 1 |
| Znhit6 | 0.046361 | 0.005032 | 1 |
| Mrpl11 | 0.046204 | 0.016089 | 1 |
| Acox3 | 0.046202 | 0.200774 | 1 |
| Zdhhc12 | 0.046191 | 0.000228 | 1 |
| Cmip | 0.046113 | 2.8E-05 | 0.905447 |
| Afg1l | 0.045927 | 0.00798 | 1 |
| Pkn2 | 0.045926 | 0.0015 | 1 |
| Glyr1 | 0.045909 | 0.137056 | 1 |
| Acin1 | 0.045894 | 0.014237 | 1 |
| Ndufaf4 | 0.045792 | 0.016351 | 1 |
| Traf3 | 0.045726 | 0.000484 | 1 |
| Irak1 | 0.045558 | 0.000331 | 1 |
| Limk2 | 0.045545 | 0.04766 | 1 |
| Acbd6 | 0.045542 | 0.002978 | 1 |
| Akap9 | 0.045452 | 0.005359 | 1 |
| Gm17018 | 0.045434 | 0.000214 | 1 |
| Nadk | 0.045405 | 0.032619 | 1 |
| Shfl | 0.045363 | 0.021087 | 1 |
| Hpn | 0.045339 | 6.23E-06 | 0.201097 |
| Psmd9 | 0.045283 | 0.000389 | 1 |
| Scaf4 | 0.045201 | 0.000463 | 1 |
| Gins1 | 0.045188 | 8.82E-05 | 1 |
| Eif4e2 | 0.04518 | 0.000394 | 1 |
| Heatr3 | 0.045132 | 0.000879 | 1 |
| Mtif2 | 0.045058 | 0.026139 | 1 |
| Dtymk | 0.045032 | 0.019642 | 1 |
| Grsf1 | 0.044738 | 3.54E-06 | 0.114187 |
| Sdhaf2 | 0.044644 | 0.000705 | 1 |
| Cct7 | 0.044639 | 0.03922 | 1 |
| Sh3bgrl | 0.044584 | 0.00104 | 1 |
| Pitrm1 | 0.044565 | 0.004517 | 1 |
| Zcchc7 | 0.044527 | 0.019438 | 1 |
| Snrpf | 0.044458 | 0.001687 | 1 |
| Naa38 | 0.044438 | 0.00058 | 1 |
| Gab3 | 0.044438 | 0.27786 | 1 |
| Wdr3 | 0.044212 | 0.003526 | 1 |
| Polr2f | 0.044168 | 0.001225 | 1 |
| Secisbp2 | 0.044137 | 0.055323 | 1 |
| Fam217b | 0.044032 | 0.002286 | 1 |
| Rnf121 | 0.04401 | 0.001391 | 1 |
| C2cd5 | 0.044001 | 0.102674 | 1 |
| Fam13b | 0.043989 | 0.002771 | 1 |
| D130040H | 0.043874 | 0.00033 | 1 |
| Gabpb2 | 0.043861 | 0.200355 | 1 |
| Pnpla7 | 0.043821 | 0.691004 | 1 |

| | | | |
|---|---|---|---|
| Riox2 | 0.043758 | 0.000433 | 1 |
| Usp7 | 0.043727 | 0.000496 | 1 |
| Nif3l1 | 0.043636 | 0.000905 | 1 |
| Chfr | 0.043562 | 0.006632 | 1 |
| Zfand2a | 0.043542 | 0.00056 | 1 |
| Inafm1 | 0.043501 | 0.266603 | 1 |
| Pdia4 | 0.043465 | 0.03604 | 1 |
| Nfkb2 | 0.043331 | 0.000176 | 1 |
| Gtf2h5 | 0.043307 | 0.000563 | 1 |
| Cep164 | 0.043298 | 0.00074 | 1 |
| Ep400 | 0.043288 | 0.000463 | 1 |
| Prmt1 | 0.043238 | 0.01357 | 1 |
| Fermt3 | 0.043212 | 0.093431 | 1 |
| Hltf | 0.043209 | 0.004717 | 1 |
| Zfp64 | 0.043182 | 0.051321 | 1 |
| Mtch2 | 0.043153 | 0.005851 | 1 |
| Trak2 | 0.04305 | 0.000902 | 1 |
| Twf1 | 0.043029 | 0.007517 | 1 |
| Fbxo6 | 0.043008 | 0.011939 | 1 |
| Ppid | 0.042888 | 0.004818 | 1 |
| Tcf20 | 0.042868 | 0.05343 | 1 |
| Nifk | 0.042761 | 0.003664 | 1 |
| Uchl5 | 0.042716 | 0.00731 | 1 |
| Rnf19a | 0.042689 | 0.388682 | 1 |
| Mea1 | 0.042672 | 0.001087 | 1 |
| Tex10 | 0.042491 | 0.000325 | 1 |
| Fbxo34 | 0.042485 | 0.000307 | 1 |
| Speg | 0.042476 | 0.00016 | 1 |
| Slc36a4 | 0.042475 | 0.009963 | 1 |
| Tbl3 | 0.04242 | 0.006147 | 1 |
| Enpp4 | 0.042275 | 0.000276 | 1 |
| Zdhhc13 | 0.042267 | 0.011433 | 1 |
| Pon2 | 0.042189 | 0.215669 | 1 |
| Dynll2 | 0.042167 | 0.031447 | 1 |
| Mtfmt | 0.042121 | 0.000528 | 1 |
| Msra | 0.041993 | 0.000366 | 1 |
| Trmt6 | 0.041943 | 0.057034 | 1 |
| Eif2b3 | 0.041801 | 0.004385 | 1 |
| Poli | 0.041785 | 7.6E-05 | 1 |
| Cep162 | 0.041732 | 0.00812 | 1 |
| Stoml1 | 0.041727 | 0.000599 | 1 |
| Tomm5 | 0.041707 | 0.000185 | 1 |
| Slc25a19 | 0.041677 | 0.00038 | 1 |
| Pkib | 0.041671 | 0.001149 | 1 |
| Adora3 | 0.041669 | 0.012382 | 1 |
| Rae1 | 0.041659 | 0.019893 | 1 |
| Dtnbp1 | 0.041589 | 0.000195 | 1 |
| Prpf31 | 0.041579 | 0.000966 | 1 |

| | | | |
|---|---|---|---|
| Zpr1 | 0.041336 | 0.002136 | 1 |
| Zfp992 | 0.041277 | 2.63E-05 | 0.849385 |
| Btbd19 | 0.041261 | 0.021967 | 1 |
| Pinx1 | 0.041257 | 2.86E-05 | 0.921776 |
| Trmo | 0.041242 | 0.001615 | 1 |
| Sco2 | 0.041162 | 0.010533 | 1 |
| Clec5a | 0.041137 | 0.062432 | 1 |
| Dus2 | 0.041029 | 0.241813 | 1 |
| Paf1 | 0.040968 | 0.004672 | 1 |
| Dmxl2 | 0.040878 | 0.000267 | 1 |
| Sae1 | 0.040833 | 0.142243 | 1 |
| Tmub1 | 0.040829 | 0.00019 | 1 |
| Elp1 | 0.040809 | 0.006846 | 1 |
| Trip12 | 0.040715 | 0.011593 | 1 |
| Ccdc82 | 0.040623 | 0.049523 | 1 |
| Adprm | 0.040617 | 0.004841 | 1 |
| Fam120a | 0.040565 | 0.005358 | 1 |
| Katnbl1 | 0.040516 | 0.001504 | 1 |
| Slk | 0.040461 | 0.000103 | 1 |
| Ints2 | 0.040442 | 0.012861 | 1 |
| Ptcd1 | 0.040409 | 0.019083 | 1 |
| Srfbp1 | 0.040252 | 0.006116 | 1 |
| Rexo4 | 0.040232 | 0.001781 | 1 |
| Pwp1 | 0.040122 | 0.003014 | 1 |
| Wdr83os | 0.040117 | 0.004539 | 1 |
| Plekhn1 | 0.040093 | 0.001453 | 1 |
| Taf15 | 0.040001 | 0.015662 | 1 |
| Ufm1 | 0.039976 | 0.000229 | 1 |
| 9430038I0 | 0.039952 | 0.000187 | 1 |
| Sertad3 | 0.039881 | 0.000337 | 1 |
| Gpr89 | 0.039839 | 0.000918 | 1 |
| Dtd1 | 0.039795 | 0.035959 | 1 |
| Isy1 | 0.039794 | 0.012631 | 1 |
| Ago2 | 0.039784 | 0.004552 | 1 |
| Phf6 | 0.03971 | 0.004266 | 1 |
| Acot8 | 0.039692 | 0.005759 | 1 |
| Cyc1 | 0.039677 | 0.062509 | 1 |
| Uqcrfs1 | 0.039567 | 0.000883 | 1 |
| Galk1 | 0.039494 | 0.058745 | 1 |
| Mrps28 | 0.03947 | 0.000249 | 1 |
| Yeats2 | 0.039463 | 0.000135 | 1 |
| Hmgcl | 0.039373 | 0.028126 | 1 |
| Ppfia1 | 0.039291 | 0.250789 | 1 |
| Nudt3 | 0.039256 | 2.05E-05 | 0.661848 |
| Cisd1 | 0.039254 | 0.045546 | 1 |
| Spata5l1 | 0.039225 | 0.000792 | 1 |
| Fnbp1 | 0.039102 | 0.035206 | 1 |
| Arpc5 | 0.0391 | 0.13718 | 1 |

| | | | |
|---|---|---|---|
| Asxl2 | 0.039052 | 0.479285 | 1 |
| Dctn3 | 0.03898 | 0.015439 | 1 |
| Pank2 | 0.038968 | 0.001287 | 1 |
| Haus8 | 0.038955 | 0.000131 | 1 |
| Gart | 0.038924 | 0.002743 | 1 |
| Cyb561d2 | 0.038917 | 0.00011 | 1 |
| Alkbh6 | 0.038857 | 0.043701 | 1 |
| Fblim1 | 0.038856 | 5.12E-06 | 0.16522 |
| Zmat3 | 0.038772 | 0.017891 | 1 |
| Dcaf13 | 0.038716 | 0.033612 | 1 |
| Itgb1bp1 | 0.038693 | 0.240533 | 1 |
| Arfip2 | 0.03861 | 0.006477 | 1 |
| Rint1 | 0.038578 | 0.019938 | 1 |
| Tmem219 | 0.038518 | 0.023449 | 1 |
| Taf11 | 0.038491 | 0.057483 | 1 |
| Qrsl1 | 0.038484 | 0.000995 | 1 |
| Atp5md | 0.038484 | 0.006112 | 1 |
| Bag1 | 0.038413 | 3.39E-06 | 0.109412 |
| Rabep1 | 0.038406 | 0.001626 | 1 |
| Gab2 | 0.038379 | 0.006768 | 1 |
| Ppme1 | 0.03832 | 0.208001 | 1 |
| Eci2 | 0.038315 | 0.065162 | 1 |
| Mrps35 | 0.038269 | 6.51E-05 | 1 |
| Aimp2 | 0.038158 | 0.000125 | 1 |
| Cs | 0.038157 | 0.009672 | 1 |
| Ppp1r13b | 0.038145 | 0.019055 | 1 |
| Zrsr2 | 0.03813 | 0.000236 | 1 |
| Chd3 | 0.038073 | 0.004939 | 1 |
| Atl3 | 0.038026 | 0.007795 | 1 |
| Sugp1 | 0.038012 | 0.000729 | 1 |
| Dnajc2 | 0.038006 | 2.68E-05 | 0.865062 |
| Ikbkb | 0.038002 | 0.009088 | 1 |
| Gm16118 | 0.037977 | 0.000339 | 1 |
| Micos13 | 0.037926 | 0.001931 | 1 |
| Rbpms | 0.037853 | 0.000379 | 1 |
| Snap29 | 0.037769 | 0.0012 | 1 |
| 2310061I0 | 0.037765 | 0.079436 | 1 |
| Fam214b | 0.037688 | 0.000358 | 1 |
| Klhdc4 | 0.037621 | 0.046323 | 1 |
| Exosc8 | 0.037486 | 0.032283 | 1 |
| Fnip1 | 0.03741 | 0.059165 | 1 |
| Ncapd3 | 0.037388 | 0.024118 | 1 |
| Smarca4 | 0.037347 | 0.001304 | 1 |
| Pprc1 | 0.037311 | 0.007245 | 1 |
| Tmem161b | 0.037223 | 0.027401 | 1 |
| Scfd1 | 0.037182 | 0.015015 | 1 |
| Eif3l | 0.037164 | 0.072188 | 1 |
| Ankib1 | 0.037135 | 0.077253 | 1 |

| | | | |
|---|---|---|---|
| Fcf1 | 0.036986 | 0.027169 | 1 |
| Hinfp | 0.036858 | 0.000136 | 1 |
| Ube2g1 | 0.03684 | 0.331553 | 1 |
| Gmds | 0.036827 | 0.000955 | 1 |
| Nsmaf | 0.036805 | 0.003194 | 1 |
| Trip11 | 0.036775 | 0.001153 | 1 |
| Eipr1 | 0.036762 | 0.000131 | 1 |
| Atp6v1d | 0.036623 | 0.012067 | 1 |
| Pik3cb | 0.036542 | 0.00385 | 1 |
| Marc2 | 0.036505 | 0.054275 | 1 |
| Golga5 | 0.036477 | 0.000818 | 1 |
| Eri2 | 0.036468 | 0.00231 | 1 |
| Caprin1 | 0.036457 | 0.007103 | 1 |
| mt-Nd3 | 0.036401 | 0.001774 | 1 |
| Mier2 | 0.036338 | 0.001521 | 1 |
| Ankrd11 | 0.036296 | 0.001094 | 1 |
| Nup205 | 0.036263 | 0.000318 | 1 |
| Pttg1 | 0.036233 | 0.003477 | 1 |
| Mdm4 | 0.036225 | 0.003005 | 1 |
| Prkar2a | 0.036204 | 0.132102 | 1 |
| Tdrd7 | 0.036151 | 2.43E-05 | 0.78438 |
| Tango2 | 0.035994 | 0.002582 | 1 |
| Esf1 | 0.035968 | 0.000591 | 1 |
| Osbpl2 | 0.035733 | 0.123649 | 1 |
| Eri1 | 0.035571 | 0.001788 | 1 |
| Ints7 | 0.035505 | 0.024211 | 1 |
| Arl5a | 0.035482 | 0.002826 | 1 |
| G3bp1 | 0.035437 | 0.003505 | 1 |
| Gm9993 | 0.035366 | 0.006656 | 1 |
| A630089N | 0.03535 | 0.004691 | 1 |
| Med8 | 0.035334 | 0.032966 | 1 |
| Stimate | 0.035141 | 0.001025 | 1 |
| Arfgap1 | 0.035117 | 0.017226 | 1 |
| March8 | 0.035084 | 1.27E-05 | 0.409258 |
| Pold3 | 0.035039 | 0.001839 | 1 |
| Cbfb | 0.034965 | 0.000983 | 1 |
| Rwdd4a | 0.034954 | 0.380758 | 1 |
| Atad5 | 0.034884 | 0.000447 | 1 |
| Trim30b | 0.034816 | 0.001003 | 1 |
| Ecsit | 0.034782 | 0.014459 | 1 |
| Smurf2 | 0.034748 | 0.048219 | 1 |
| Ssr2 | 0.034717 | 0.031388 | 1 |
| Rnf123 | 0.034556 | 0.099499 | 1 |
| Phlpp1 | 0.034518 | 0.39833 | 1 |
| Dnmt1 | 0.034459 | 0.885098 | 1 |
| Rnf214 | 0.034406 | 0.004658 | 1 |
| Ndufb2 | 0.034335 | 0.017598 | 1 |
| Slc13a3 | 0.034185 | 0.014903 | 1 |

| | | | |
|---|---|---|---|
| Gin1 | 0.034129 | 0.114885 | 1 |
| Tigd2 | 0.033966 | 0.001749 | 1 |
| Ifnar1 | 0.033926 | 0.023206 | 1 |
| Lin54 | 0.033924 | 0.021968 | 1 |
| Golt1b | 0.03379 | 0.000482 | 1 |
| Gnaq | 0.033776 | 1.6E-05 | 0.517907 |
| Sec24b | 0.03373 | 0.001359 | 1 |
| Coq5 | 0.033689 | 0.004854 | 1 |
| Tnip2 | 0.033649 | 0.002754 | 1 |
| Pop7 | 0.033527 | 0.000244 | 1 |
| Rab5c | 0.033455 | 0.109222 | 1 |
| U2af1 | 0.033453 | 0.017308 | 1 |
| Mrps17 | 0.033363 | 0.044469 | 1 |
| Noc3l | 0.033352 | 0.029253 | 1 |
| Uqcc1 | 0.03323 | 0.003623 | 1 |
| Arid4a | 0.033221 | 0.129072 | 1 |
| Guca1a | 0.033207 | 0.003794 | 1 |
| Ndufb10 | 0.033201 | 0.000839 | 1 |
| Eif4b | 0.033153 | 0.004505 | 1 |
| 6720427I0 | 0.032994 | 0.008324 | 1 |
| Mrps31 | 0.032851 | 0.020102 | 1 |
| Ampd2 | 0.032831 | 0.032402 | 1 |
| Osbpl11 | 0.032821 | 0.147643 | 1 |
| Inpp5f | 0.03281 | 0.000493 | 1 |
| Erg28 | 0.03278 | 0.000923 | 1 |
| Xbp1 | 0.032769 | 0.065116 | 1 |
| Mrps15 | 0.032752 | 0.003532 | 1 |
| Metap1d | 0.032666 | 0.01492 | 1 |
| Msn | 0.032624 | 0.211352 | 1 |
| Timm10 | 0.032598 | 0.001123 | 1 |
| Rcbtb1 | 0.032432 | 0.012428 | 1 |
| Traf2 | 0.032427 | 0.035291 | 1 |
| Heca | 0.032339 | 0.066862 | 1 |
| Tm9sf1 | 0.032333 | 0.007275 | 1 |
| Ptrh2 | 0.032299 | 0.022106 | 1 |
| Ppdpf | 0.032298 | 0.104076 | 1 |
| Mfge8 | 0.032283 | 0.00073 | 1 |
| Uchl3 | 0.032271 | 0.10028 | 1 |
| Mroh1 | 0.032255 | 0.010202 | 1 |
| Siva1 | 0.032203 | 0.007282 | 1 |
| Rala | 0.032166 | 0.000118 | 1 |
| Exosc4 | 0.032086 | 0.011684 | 1 |
| Phip | 0.032057 | 0.077334 | 1 |
| Lamp1 | 0.031996 | 0.14652 | 1 |
| Prmt7 | 0.031944 | 0.041288 | 1 |
| Ercc8 | 0.03191 | 0.003068 | 1 |
| Gnai2 | 0.031867 | 0.316571 | 1 |
| Fbxw11 | 0.031748 | 0.000498 | 1 |

| | | | |
|---|---|---|---|
| Abce1 | 0.031716 | 0.002041 | 1 |
| Cyp20a1 | 0.03168 | 0.012997 | 1 |
| Cebpg | 0.03168 | 0.004778 | 1 |
| Lars | 0.031572 | 0.007841 | 1 |
| Nup93 | 0.031548 | 0.015193 | 1 |
| Adss | 0.031516 | 0.354987 | 1 |
| Senp2 | 0.03149 | 0.000494 | 1 |
| Seh1l | 0.031454 | 0.000332 | 1 |
| Ndufs4 | 0.031314 | 0.012946 | 1 |
| Mettl1 | 0.031271 | 0.004646 | 1 |
| Slc35e2 | 0.03112 | 0.114663 | 1 |
| Slc30a5 | 0.031082 | 0.015655 | 1 |
| Mrpl41 | 0.031056 | 0.000144 | 1 |
| Gm16599 | 0.030836 | 0.00172 | 1 |
| 4930402H | 0.030815 | 0.198591 | 1 |
| Ythdc1 | 0.030693 | 0.010463 | 1 |
| Umps | 0.030676 | 0.028383 | 1 |
| Recql5 | 0.03064 | 0.011271 | 1 |
| Mrpl27 | 0.0306 | 0.007178 | 1 |
| Me2 | 0.030547 | 0.102236 | 1 |
| Surf4 | 0.030545 | 0.034547 | 1 |
| Mien1 | 0.030503 | 0.005208 | 1 |
| Psmf1 | 0.030438 | 0.006829 | 1 |
| Etfb | 0.030339 | 0.005484 | 1 |
| B3galt6 | 0.030329 | 0.008051 | 1 |
| Psmb5 | 0.030303 | 0.154642 | 1 |
| Dlg1 | 0.030274 | 0.003039 | 1 |
| Lman2 | 0.030222 | 0.009602 | 1 |
| 1110032A | 0.030221 | 0.021299 | 1 |
| Arfgap3 | 0.030209 | 0.246202 | 1 |
| Ubr5 | 0.030044 | 0.05404 | 1 |
| Cdk7 | 0.03003 | 0.011497 | 1 |
| Pofut2 | 0.029991 | 0.031433 | 1 |
| Abhd18 | 0.029967 | 0.048569 | 1 |
| Cog5 | 0.029958 | 0.021689 | 1 |
| Pym1 | 0.029892 | 0.016439 | 1 |
| Bnip1 | 0.029871 | 0.000122 | 1 |
| Eps15l1 | 0.029772 | 0.050867 | 1 |
| Rab4b | 0.029711 | 0.019282 | 1 |
| Golph3 | 0.029681 | 0.028144 | 1 |
| Itga5 | 0.029567 | 0.00629 | 1 |
| Dtd2 | 0.029524 | 0.200742 | 1 |
| Dis3l2 | 0.029483 | 0.008462 | 1 |
| Rpp21 | 0.029433 | 0.035414 | 1 |
| Ubl4a | 0.029316 | 0.170642 | 1 |
| Ythdc2 | 0.029307 | 0.050612 | 1 |
| Rgs14 | 0.029307 | 0.076909 | 1 |
| Ppp2r2a | 0.029305 | 0.051793 | 1 |

| | | | |
|---|---|---|---|
| Lyar | 0.029301 | 0.010512 | 1 |
| Tfam | 0.029285 | 0.569087 | 1 |
| H13 | 0.029203 | 0.024282 | 1 |
| Dpp3 | 0.029054 | 0.005415 | 1 |
| Inip | 0.029043 | 0.000975 | 1 |
| Opa1 | 0.028957 | 0.010602 | 1 |
| Tbca | 0.028944 | 8.59E-05 | 1 |
| Ddx41 | 0.028932 | 0.015809 | 1 |
| Slf1 | 0.0289 | 0.000813 | 1 |
| Naa15 | 0.028894 | 0.003651 | 1 |
| Atp6v1f | 0.028866 | 0.019426 | 1 |
| Etfa | 0.028822 | 0.023276 | 1 |
| Necap2 | 0.028757 | 0.009672 | 1 |
| Rnpc3 | 0.028751 | 0.211142 | 1 |
| Cpsf4 | 0.028712 | 0.030194 | 1 |
| Gmps | 0.02871 | 0.029801 | 1 |
| Use1 | 0.028602 | 0.010059 | 1 |
| Usp14 | 0.028561 | 0.008114 | 1 |
| Yif1b | 0.028557 | 0.031082 | 1 |
| U2af2 | 0.02853 | 0.127026 | 1 |
| Nrbf2 | 0.028523 | 0.002143 | 1 |
| Park7 | 0.028506 | 0.07076 | 1 |
| Lig3 | 0.028473 | 0.003773 | 1 |
| Lonp1 | 0.028428 | 7.69E-05 | 1 |
| Pigs | 0.028417 | 0.005713 | 1 |
| Trnt1 | 0.028407 | 0.08828 | 1 |
| Cul4b | 0.028283 | 0.02288 | 1 |
| Dyrk4 | 0.028166 | 0.170363 | 1 |
| B230307C2 | 0.028161 | 0.004882 | 1 |
| Ddt | 0.028085 | 0.001414 | 1 |
| Rce1 | 0.028074 | 0.005708 | 1 |
| Fis1 | 0.027956 | 0.039566 | 1 |
| Vps29 | 0.02788 | 0.008498 | 1 |
| Nub1 | 0.027666 | 0.029082 | 1 |
| Tmem11 | 0.027625 | 0.050019 | 1 |
| Uap1 | 0.027624 | 0.010429 | 1 |
| Ssbp4 | 0.027435 | 0.064561 | 1 |
| Thyn1 | 0.027427 | 0.006306 | 1 |
| Knop1 | 0.02724 | 0.008735 | 1 |
| Havcr2 | 0.027239 | 0.683486 | 1 |
| Srsf4 | 0.027225 | 0.011631 | 1 |
| Ndufa10 | 0.027157 | 0.018992 | 1 |
| Ap3m2 | 0.02713 | 0.090072 | 1 |
| Sirt3 | 0.027107 | 0.000889 | 1 |
| Zfp131 | 0.027058 | 0.004566 | 1 |
| Hectd3 | 0.02703 | 0.011745 | 1 |
| Clk2 | 0.027028 | 0.015253 | 1 |
| Gba | 0.026979 | 0.004411 | 1 |

| | | | |
|---|---|---|---|
| Rsrc1 | 0.026969 | 0.05009 | 1 |
| Cnot10 | 0.026863 | 0.012013 | 1 |
| Wnk1 | 0.026824 | 0.014757 | 1 |
| Ints13 | 0.026777 | 0.017053 | 1 |
| Tatdn1 | 0.026763 | 0.005481 | 1 |
| Ero1lb | 0.02667 | 0.003089 | 1 |
| Mgst3 | 0.026574 | 0.001453 | 1 |
| Sms | 0.026553 | 0.041988 | 1 |
| Gm26510 | 0.026491 | 0.008489 | 1 |
| Ppp1cc | 0.026449 | 0.030226 | 1 |
| Utp6 | 0.026158 | 0.017033 | 1 |
| Rfxap | 0.026071 | 0.040327 | 1 |
| Diaph2 | 0.026034 | 0.014318 | 1 |
| Rsph3a | 0.026006 | 0.018442 | 1 |
| Sugt1 | 0.025952 | 0.070985 | 1 |
| Stk38 | 0.025938 | 0.004432 | 1 |
| Baz2a | 0.025927 | 0.006012 | 1 |
| 2210016L2 | 0.025886 | 0.000841 | 1 |
| Tubgcp4 | 0.025841 | 0.05452 | 1 |
| D230025D | 0.025827 | 0.214003 | 1 |
| Mbd3 | 0.025809 | 0.000648 | 1 |
| Ppp1r3d | 0.025807 | 0.000915 | 1 |
| Ing5 | 0.025789 | 0.009132 | 1 |
| Nras | 0.025703 | 0.03005 | 1 |
| Bms1 | 0.025699 | 0.037764 | 1 |
| Rnf166 | 0.025696 | 0.002857 | 1 |
| Fubp1 | 0.025591 | 0.036821 | 1 |
| Sgf29 | 0.025558 | 0.023683 | 1 |
| Ndufv2 | 0.025476 | 0.016393 | 1 |
| Gm16337 | 0.025428 | 0.162336 | 1 |
| Acaa1a | 0.025376 | 0.112556 | 1 |
| Mrpl19 | 0.025363 | 0.034925 | 1 |
| Tep1 | 0.025348 | 0.001908 | 1 |
| Plekhm2 | 0.025339 | 8.73E-05 | 1 |
| Pctp | 0.025333 | 0.004814 | 1 |
| Tceanc2 | 0.025269 | 0.013731 | 1 |
| Smn1 | 0.025236 | 6.05E-06 | 0.195352 |
| Slc25a12 | 0.025181 | 0.019025 | 1 |
| Nkapd1 | 0.025149 | 0.000949 | 1 |
| Xrcc6 | 0.025106 | 0.253248 | 1 |
| Psd4 | 0.02507 | 0.10593 | 1 |
| Crlf2 | 0.025065 | 0.011411 | 1 |
| Zfp346 | 0.024992 | 0.021231 | 1 |
| Crk | 0.024943 | 0.00449 | 1 |
| Nhlrc2 | 0.024864 | 0.031689 | 1 |
| Slc39a6 | 0.024839 | 0.003827 | 1 |
| Mpp1 | 0.024828 | 0.110282 | 1 |
| Ndufab1 | 0.024811 | 0.035234 | 1 |

| | | | |
|---|---|---|---|
| Dnajc5 | 0.024798 | 0.010022 | 1 |
| Tsen2 | 0.024753 | 0.080671 | 1 |
| Pdcd4 | 0.024733 | 0.835457 | 1 |
| Stmn1 | 0.024686 | 0.005704 | 1 |
| Rbl1 | 0.02468 | 0.070045 | 1 |
| Zdhhc4 | 0.024551 | 0.00444 | 1 |
| Qtrt1 | 0.024445 | 0.022988 | 1 |
| Tsc1 | 0.024414 | 0.138175 | 1 |
| 3830406C1 | 0.024318 | 0.014803 | 1 |
| Gmppa | 0.024228 | 0.079297 | 1 |
| Smg7 | 0.024165 | 0.0211 | 1 |
| Eef2kmt | 0.024051 | 0.00071 | 1 |
| 9130401M | 0.024018 | 0.011248 | 1 |
| Mrpl46 | 0.02399 | 0.005241 | 1 |
| Fiz1 | 0.023984 | 0.070026 | 1 |
| Meaf6 | 0.023966 | 0.019581 | 1 |
| Ube2k | 0.023897 | 0.00145 | 1 |
| Ric1 | 0.023886 | 0.000863 | 1 |
| Ilkap | 0.023875 | 0.01841 | 1 |
| Toe1 | 0.023866 | 0.244694 | 1 |
| Polr3h | 0.023739 | 0.053539 | 1 |
| Denr | 0.023637 | 0.000397 | 1 |
| Dph6 | 0.023635 | 0.0186 | 1 |
| Plekhb2 | 0.023552 | 0.002452 | 1 |
| Sars2 | 0.023516 | 0.002599 | 1 |
| Eif3g | 0.023499 | 0.000564 | 1 |
| 4921524J1 | 0.023434 | 0.010603 | 1 |
| Hnrnpm | 0.023422 | 0.069816 | 1 |
| Ccnh | 0.023419 | 0.047688 | 1 |
| Safb | 0.023382 | 0.012067 | 1 |
| Ip6k1 | 0.023351 | 0.000843 | 1 |
| Camk2d | 0.023312 | 0.065657 | 1 |
| Snupn | 0.023281 | 0.002371 | 1 |
| AI413582 | 0.023209 | 0.043349 | 1 |
| Mzt2 | 0.02317 | 0.04804 | 1 |
| Hdac4 | 0.023117 | 0.045984 | 1 |
| Adipor1 | 0.022952 | 0.585147 | 1 |
| Miga1 | 0.022847 | 0.05387 | 1 |
| AU022793 | 0.022805 | 0.029333 | 1 |
| Simc1 | 0.022792 | 0.038169 | 1 |
| Set | 0.022617 | 0.000573 | 1 |
| Flad1 | 0.022611 | 0.011977 | 1 |
| Fam98c | 0.022602 | 0.055066 | 1 |
| Trpm7 | 0.022579 | 0.025647 | 1 |
| Kitl | 0.022423 | 0.011407 | 1 |
| Mrpl17 | 0.022402 | 0.024949 | 1 |
| Prkrip1 | 0.022381 | 0.010501 | 1 |
| Fam114a1 | 0.022272 | 0.000952 | 1 |

| | | | |
|---|---|---|---|
| Cars2 | 0.022239 | 0.001093 | 1 |
| Nup54 | 0.022125 | 0.033126 | 1 |
| N4bp2l2 | 0.022088 | 0.184107 | 1 |
| Rtca | 0.022057 | 0.01612 | 1 |
| Sumf1 | 0.022057 | 0.00485 | 1 |
| Tmem41b | 0.021996 | 0.1157 | 1 |
| Cdc40 | 0.021981 | 0.118487 | 1 |
| Tbce | 0.021954 | 0.042471 | 1 |
| Tmem161a | 0.021946 | 0.023899 | 1 |
| Cd2ap | 0.021823 | 0.037413 | 1 |
| Cdk17 | 0.021711 | 0.251226 | 1 |
| Rap1b | 0.021569 | 0.011816 | 1 |
| Hmgcr | 0.021478 | 0.003159 | 1 |
| Polr3d | 0.021379 | 0.000637 | 1 |
| Gpatch3 | 0.021271 | 0.018663 | 1 |
| Ralgapa2 | 0.021198 | 0.42819 | 1 |
| Gemin7 | 0.021145 | 0.046008 | 1 |
| Ddx23 | 0.02114 | 0.060068 | 1 |
| Ak6 | 0.021132 | 0.05271 | 1 |
| Cog3 | 0.021041 | 0.076269 | 1 |
| Ccm2 | 0.020998 | 0.040212 | 1 |
| Atp5a1 | 0.02099 | 0.340931 | 1 |
| Pex13 | 0.020978 | 0.285558 | 1 |
| Bdp1 | 0.020945 | 0.000605 | 1 |
| Pin4 | 0.020941 | 0.04784 | 1 |
| Sart1 | 0.020936 | 0.038091 | 1 |
| Nectin2 | 0.020822 | 0.000332 | 1 |
| Mbtd1 | 0.020819 | 0.016353 | 1 |
| Mrps26 | 0.020812 | 0.042107 | 1 |
| Rrbp1 | 0.020811 | 0.022277 | 1 |
| Ppm1k | 0.020776 | 0.137078 | 1 |
| Ubxn4 | 0.020776 | 0.027292 | 1 |
| Mrtfb | 0.020732 | 0.082108 | 1 |
| Tmf1 | 0.020708 | 0.235757 | 1 |
| Ccdc186 | 0.0207 | 0.013963 | 1 |
| Pds5b | 0.020627 | 0.689942 | 1 |
| Gm17066 | 0.020607 | 0.008177 | 1 |
| Kif13b | 0.020599 | 0.031094 | 1 |
| Prep | 0.020567 | 0.087158 | 1 |
| Pcyox1 | 0.020554 | 0.030806 | 1 |
| Nom1 | 0.020538 | 0.008293 | 1 |
| Jund | 0.020517 | 0.356759 | 1 |
| Fbxw7 | 0.02051 | 0.002217 | 1 |
| Sun1 | 0.02051 | 0.100039 | 1 |
| Tctex1d2 | 0.020482 | 0.019985 | 1 |
| Rpp30 | 0.020411 | 0.13951 | 1 |
| Zfp654 | 0.020371 | 0.073608 | 1 |
| H2afx | 0.020363 | 0.000806 | 1 |

| | | | |
|---|---|---|---|
| Atf4 | 0.020353 | 0.012622 | 1 |
| Ciapin1 | 0.020344 | 0.047033 | 1 |
| Scyl3 | 0.020296 | 0.032453 | 1 |
| Prim2 | 0.020284 | 0.87842 | 1 |
| Csnk1g3 | 0.020225 | 0.040093 | 1 |
| Hspbap1 | 0.020149 | 0.016804 | 1 |
| Nsun4 | 0.02009 | 0.003887 | 1 |
| Tars | 0.019932 | 0.022015 | 1 |
| Ttll4 | 0.019913 | 0.000822 | 1 |
| Selenof | 0.01985 | 0.307657 | 1 |
| Brd2 | 0.019787 | 0.401472 | 1 |
| March1 | 0.019757 | 0.433764 | 1 |
| Notch2 | 0.019752 | 0.079385 | 1 |
| Naip6 | 0.019746 | 0.001113 | 1 |
| Ddx56 | 0.01973 | 0.031646 | 1 |
| Mtap | 0.019696 | 0.042662 | 1 |
| Eif4enif1 | 0.019678 | 0.00399 | 1 |
| Ndufa4 | 0.019652 | 0.058918 | 1 |
| Fam160a2 | 0.019639 | 0.486392 | 1 |
| Coq9 | 0.019496 | 0.017502 | 1 |
| Supv3l1 | 0.019378 | 0.406302 | 1 |
| Gcsh | 0.019158 | 0.018723 | 1 |
| Gm27017 | 0.019144 | 0.024921 | 1 |
| Rpa1 | 0.019143 | 0.1251 | 1 |
| Fastkd2 | 0.019075 | 0.001966 | 1 |
| Dbi | 0.019056 | 0.00889 | 1 |
| Cox10 | 0.018925 | 0.019748 | 1 |
| Cox18 | 0.018903 | 0.002819 | 1 |
| Znhit1 | 0.018834 | 0.004524 | 1 |
| Tipin | 0.01881 | 0.073372 | 1 |
| Eif2b2 | 0.018776 | 0.144964 | 1 |
| Btbd7 | 0.018774 | 0.01138 | 1 |
| Wdr70 | 0.018754 | 0.003423 | 1 |
| Csnk2a2 | 0.018669 | 0.011142 | 1 |
| Sumo3 | 0.018655 | 0.117905 | 1 |
| Epn1 | 0.01857 | 0.004036 | 1 |
| Chst14 | 0.018521 | 0.261594 | 1 |
| Dars | 0.018484 | 0.170168 | 1 |
| Telo2 | 0.018476 | 0.000109 | 1 |
| Ubb | 0.018456 | 0.945108 | 1 |
| Zfp53 | 0.018451 | 0.086577 | 1 |
| Dpy19l4 | 0.01843 | 0.051289 | 1 |
| Sec61a2 | 0.018391 | 0.023384 | 1 |
| Wdr83 | 0.018386 | 0.001309 | 1 |
| Ftx | 0.018295 | 0.281391 | 1 |
| Psmd13 | 0.018202 | 0.119959 | 1 |
| Rbm39 | 0.018156 | 0.836377 | 1 |
| Polg | 0.018107 | 0.055998 | 1 |

| | | | |
|---|---|---|---|
| Tex14 | 0.018044 | 0.228082 | 1 |
| Tysnd1 | 0.018022 | 0.25748 | 1 |
| Xpa | 0.018002 | 0.112268 | 1 |
| Cnot4 | 0.017991 | 0.015071 | 1 |
| Wdr73 | 0.017954 | 0.023141 | 1 |
| Eprs | 0.017866 | 0.039995 | 1 |
| Tradd | 0.017837 | 0.059254 | 1 |
| Ints11 | 0.01782 | 0.25218 | 1 |
| Tsfm | 0.017682 | 0.001325 | 1 |
| Rusc1 | 0.017628 | 0.001247 | 1 |
| Pip5k1c | 0.017627 | 0.04338 | 1 |
| Psme3 | 0.017623 | 0.019315 | 1 |
| Lsm1 | 0.017606 | 0.006986 | 1 |
| Gm48678 | 0.017508 | 0.146949 | 1 |
| Atxn7 | 0.017485 | 0.195348 | 1 |
| Stk40 | 0.017445 | 0.043751 | 1 |
| Spryd3 | 0.017381 | 0.003414 | 1 |
| Ergic2 | 0.017306 | 0.035641 | 1 |
| Nudt7 | 0.017263 | 0.035767 | 1 |
| Msmo1 | 0.017247 | 0.28081 | 1 |
| Cul2 | 0.01724 | 0.173732 | 1 |
| Trir | 0.017195 | 0.159308 | 1 |
| Dgcr8 | 0.017144 | 0.098411 | 1 |
| Prr3 | 0.017059 | 0.000282 | 1 |
| Atp5j | 0.017046 | 0.009915 | 1 |
| Qpctl | 0.016911 | 0.022068 | 1 |
| Mrpl10 | 0.016889 | 0.173557 | 1 |
| Hmgn2 | 0.016761 | 0.192845 | 1 |
| Cpeb3 | 0.016744 | 0.541764 | 1 |
| Emc8 | 0.016683 | 0.011887 | 1 |
| Ranbp9 | 0.016655 | 0.180148 | 1 |
| Gna13 | 0.016596 | 0.088991 | 1 |
| Exosc9 | 0.016585 | 0.012112 | 1 |
| Slbp | 0.016582 | 0.276495 | 1 |
| Ust | 0.016309 | 0.993677 | 1 |
| Dedd | 0.016222 | 0.163284 | 1 |
| Nudcd2 | 0.016204 | 0.004923 | 1 |
| Htra3 | 0.016152 | 0.001728 | 1 |
| Med26 | 0.016107 | 0.017705 | 1 |
| Hmg20b | 0.016078 | 0.071829 | 1 |
| Katna1 | 0.016072 | 0.002761 | 1 |
| Npm3 | 0.016072 | 0.009082 | 1 |
| Epb41l3 | 0.016067 | 0.029856 | 1 |
| Napg | 0.016024 | 0.032074 | 1 |
| Rragc | 0.016019 | 0.425308 | 1 |
| Tbcel | 0.015953 | 0.308805 | 1 |
| Rab12 | 0.015924 | 0.011451 | 1 |
| Tnfaip8 | 0.015904 | 0.009393 | 1 |

| | | | |
|---|---|---|---|
| Utp15 | 0.015899 | 0.04705 | 1 |
| Zfp958 | 0.015855 | 0.023253 | 1 |
| Mrps34 | 0.015845 | 0.331486 | 1 |
| Vac14 | 0.015798 | 0.000677 | 1 |
| Asf1a | 0.015704 | 0.073624 | 1 |
| Glrx3 | 0.015631 | 0.064004 | 1 |
| Agtrap | 0.015627 | 0.004601 | 1 |
| Prxl2c | 0.015543 | 0.03527 | 1 |
| Atp6v1h | 0.015516 | 0.342466 | 1 |
| Tomm22 | 0.015496 | 0.075605 | 1 |
| Smarce1 | 0.015459 | 0.041607 | 1 |
| Borcs5 | 0.01536 | 0.105207 | 1 |
| Azin1 | 0.015314 | 0.005211 | 1 |
| Sugp2 | 0.015291 | 0.022923 | 1 |
| Slc27a1 | 0.015264 | 0.000538 | 1 |
| Pnn | 0.015243 | 0.048293 | 1 |
| Bad | 0.015166 | 0.019441 | 1 |
| Fem1b | 0.015059 | 0.21499 | 1 |
| Mon1a | 0.015005 | 0.149938 | 1 |
| Mad1l1 | 0.014922 | 0.208188 | 1 |
| Mlh3 | 0.014863 | 0.031336 | 1 |
| Isca2 | 0.014845 | 0.039466 | 1 |
| Rps27rt | 0.014757 | 0.007547 | 1 |
| Cryzl2 | 0.014743 | 0.010579 | 1 |
| Rtf1 | 0.014662 | 0.021815 | 1 |
| Acly | 0.014625 | 0.163342 | 1 |
| Zfp516 | 0.014618 | 7.16E-06 | 0.231287 |
| Polr3k | 0.014601 | 0.163009 | 1 |
| Map3k7 | 0.014587 | 0.002181 | 1 |
| Copg2 | 0.014551 | 0.153176 | 1 |
| Skiv2l | 0.014503 | 0.052368 | 1 |
| Polr1c | 0.014485 | 0.039064 | 1 |
| Fastkd1 | 0.01443 | 0.00696 | 1 |
| Nkiras2 | 0.014421 | 0.002718 | 1 |
| Timmdc1 | 0.01435 | 0.016889 | 1 |
| Zdhhc16 | 0.014277 | 0.073961 | 1 |
| Sfxn2 | 0.014222 | 0.054445 | 1 |
| Mpst | 0.014211 | 0.007045 | 1 |
| Prkag2 | 0.014189 | 0.070194 | 1 |
| Ndufa3 | 0.014183 | 0.000458 | 1 |
| Ggh | 0.014172 | 0.096341 | 1 |
| Usf3 | 0.014067 | 0.013008 | 1 |
| Ddx54 | 0.014064 | 0.008014 | 1 |
| Hddc2 | 0.013984 | 0.038828 | 1 |
| Snhg3 | 0.01392 | 0.002442 | 1 |
| Ppp1r12a | 0.013814 | 0.888964 | 1 |
| Tmem192 | 0.013809 | 0.010853 | 1 |
| Trappc2l | 0.013753 | 0.026068 | 1 |

| | | | |
|---|---|---|---|
| Tmem131 | 0.013652 | 0.139312 | 1 |
| Fxr1 | 0.013611 | 0.014603 | 1 |
| Rhbdd1 | 0.013604 | 0.296417 | 1 |
| Fbxo7 | 0.01352 | 0.00802 | 1 |
| Sos1 | 0.013497 | 0.079797 | 1 |
| Glud1 | 0.013454 | 0.077277 | 1 |
| Zfp609 | 0.013407 | 0.018154 | 1 |
| Hrh2 | 0.01339 | 0.001005 | 1 |
| Pdrg1 | 0.01338 | 0.024671 | 1 |
| Plgrkt | 0.013361 | 0.457207 | 1 |
| Rnf24 | 0.013259 | 0.046218 | 1 |
| Rad9a | 0.013218 | 0.102748 | 1 |
| Il1rl2 | 0.013159 | 0.005645 | 1 |
| Acp1 | 0.013148 | 0.004462 | 1 |
| Tsen15 | 0.013123 | 0.055337 | 1 |
| 1810026B( | 0.013062 | 0.052221 | 1 |
| Mrpl28 | 0.013054 | 0.008668 | 1 |
| Pold2 | 0.012987 | 0.005381 | 1 |
| Syncrip | 0.012945 | 0.137822 | 1 |
| Slc39a11 | 0.012929 | 0.019987 | 1 |
| Mrpl51 | 0.012865 | 0.009374 | 1 |
| Spred2 | 0.012807 | 0.145153 | 1 |
| Taf7 | 0.012759 | 0.610831 | 1 |
| Ufc1 | 0.012729 | 0.008606 | 1 |
| 1700021F( | 0.012652 | 0.015964 | 1 |
| Smc5 | 0.012601 | 0.184225 | 1 |
| Mfsd8 | 0.012591 | 0.036538 | 1 |
| Fbxo33 | 0.012545 | 0.111299 | 1 |
| Kif1b | 0.012499 | 0.043904 | 1 |
| Klhdc10 | 0.012459 | 0.070067 | 1 |
| Cd101 | 0.012456 | 0.189828 | 1 |
| Dimt1 | 0.012425 | 0.002488 | 1 |
| Ddx49 | 0.012414 | 0.001895 | 1 |
| Ergic1 | 0.012384 | 0.026048 | 1 |
| Ric3 | 0.012317 | 0.089631 | 1 |
| Med11 | 0.012216 | 0.000439 | 1 |
| Mir142hg | 0.0122 | 0.278862 | 1 |
| Nbdy | 0.01213 | 0.09994 | 1 |
| Atpaf2 | 0.012129 | 0.10313 | 1 |
| Nup155 | 0.012098 | 0.002835 | 1 |
| Traf5 | 0.012095 | 0.143363 | 1 |
| Zfp40 | 0.011923 | 0.302257 | 1 |
| Mob2 | 0.011917 | 0.152212 | 1 |
| Mief1 | 0.011909 | 0.158496 | 1 |
| Eif2a | 0.011778 | 0.004443 | 1 |
| Dnajc4 | 0.011776 | 0.691638 | 1 |
| Parvb | 0.011634 | 0.005953 | 1 |
| Rpusd4 | 0.011588 | 0.006939 | 1 |

| | | | |
|---|---|---|---|
| Flnb | 0.011522 | 0.403083 | 1 |
| BC004004 | 0.011516 | 0.182435 | 1 |
| Phtf1 | 0.011449 | 0.000326 | 1 |
| Metap1 | 0.011408 | 0.012093 | 1 |
| Rrm1 | 0.01139 | 0.309066 | 1 |
| Rev1 | 0.01138 | 0.12381 | 1 |
| Cuedc2 | 0.011349 | 0.067344 | 1 |
| Flvcr1 | 0.011325 | 0.014238 | 1 |
| Crlf3 | 0.01131 | 0.367245 | 1 |
| Wdr91 | 0.011243 | 0.001732 | 1 |
| Rnaseh1 | 0.011237 | 0.01752 | 1 |
| Nono | 0.011228 | 0.350158 | 1 |
| Tusc2 | 0.011185 | 0.053294 | 1 |
| Wdr5 | 0.011152 | 0.166876 | 1 |
| L3mbtl3 | 0.01115 | 0.018498 | 1 |
| Atr | 0.011146 | 0.000735 | 1 |
| Gng12 | 0.01114 | 0.022353 | 1 |
| Mrps27 | 0.011082 | 0.018389 | 1 |
| Tma16 | 0.011028 | 0.001687 | 1 |
| Ubl5 | 0.011009 | 0.008537 | 1 |
| Synrg | 0.011004 | 0.023449 | 1 |
| Ankrd39 | 0.010982 | 0.011254 | 1 |
| Zfp281 | 0.01094 | 0.032692 | 1 |
| Zswim8 | 0.010816 | 0.300602 | 1 |
| Hs2st1 | 0.010791 | 0.012082 | 1 |
| Cabin1 | 0.010782 | 0.018609 | 1 |
| Ilf2 | 0.010733 | 0.038478 | 1 |
| Eif4e | 0.010731 | 0.041422 | 1 |
| Ppp2r5e | 0.010706 | 0.01583 | 1 |
| Arl6ip6 | 0.01056 | 0.219732 | 1 |
| Washc1 | 0.010457 | 0.111031 | 1 |
| Ubac2 | 0.010456 | 0.980841 | 1 |
| Sdc4 | 0.010428 | 0.016958 | 1 |
| Rpain | 0.010384 | 0.015317 | 1 |
| Tut1 | 0.010383 | 0.075226 | 1 |
| Zfp472 | 0.010382 | 0.097375 | 1 |
| Ehbp1l1 | 0.010325 | 0.000307 | 1 |
| Txndc16 | 0.010263 | 0.000269 | 1 |
| Akap8 | 0.010202 | 0.091153 | 1 |
| Traf6 | 0.010171 | 0.002032 | 1 |
| Sec24a | 0.010147 | 0.039848 | 1 |
| Hsd3b7 | 0.010129 | 0.25942 | 1 |
| Setd5 | 0.01012 | 0.30045 | 1 |
| Fdx1 | 0.010099 | 0.002806 | 1 |
| Kat2a | 0.010058 | 0.030785 | 1 |
| Ptar1 | 0.010041 | 0.006203 | 1 |
| Chd2 | 0.010029 | 0.092083 | 1 |
| Rab7 | -0.01003 | 0.127083 | 1 |

| | | | |
|---|---|---|---|
| Rnf8 | -0.01005 | 0.057545 | 1 |
| Nprl3 | -0.01006 | 0.098269 | 1 |
| Nab2 | -0.01017 | 0.038815 | 1 |
| Mag | -0.01022 | 0.064738 | 1 |
| Tefm | -0.01025 | 0.039575 | 1 |
| Gpkow | -0.01029 | 0.185185 | 1 |
| Actl6a | -0.01031 | 0.191202 | 1 |
| Hif1an | -0.01041 | 0.852002 | 1 |
| Rnf31 | -0.01043 | 0.005781 | 1 |
| Gnptg | -0.01043 | 0.163103 | 1 |
| Ufd1 | -0.01044 | 0.185259 | 1 |
| Ing3 | -0.01044 | 0.972826 | 1 |
| Mospd3 | -0.01046 | 0.140457 | 1 |
| Zfp429 | -0.01047 | 0.099748 | 1 |
| Zfp91 | -0.01053 | 0.006925 | 1 |
| Mrpl37 | -0.01055 | 0.119139 | 1 |
| Rufy1 | -0.01067 | 0.262268 | 1 |
| Zfp326 | -0.0107 | 0.339841 | 1 |
| Rnf215 | -0.01073 | 0.14739 | 1 |
| Socs4 | -0.01074 | 0.037607 | 1 |
| Usp47 | -0.01075 | 0.331748 | 1 |
| Rars2 | -0.01077 | 0.660113 | 1 |
| Nelfe | -0.0108 | 0.515136 | 1 |
| Pgrmc2 | -0.01088 | 0.826686 | 1 |
| Tarbp2 | -0.01089 | 0.754355 | 1 |
| Vrk2 | -0.0109 | 0.290366 | 1 |
| Fam207a | -0.01094 | 0.036631 | 1 |
| Ubr7 | -0.01096 | 0.471425 | 1 |
| Stk11ip | -0.01098 | 0.305917 | 1 |
| Uxs1 | -0.01107 | 0.043423 | 1 |
| Supt6 | -0.01114 | 0.311012 | 1 |
| Ankrd52 | -0.01115 | 0.544138 | 1 |
| Mrpl3 | -0.01117 | 0.066259 | 1 |
| Zcchc9 | -0.0112 | 0.065749 | 1 |
| Las1l | -0.01131 | 0.574518 | 1 |
| Ppp2r3c | -0.01134 | 0.022829 | 1 |
| Mysm1 | -0.01137 | 0.983832 | 1 |
| Ndufaf7 | -0.01142 | 0.582193 | 1 |
| Hif1a | -0.01145 | 0.524419 | 1 |
| Zc3h18 | -0.01145 | 0.149573 | 1 |
| Slc12a4 | -0.0115 | 0.014821 | 1 |
| Ccdc34 | -0.0115 | 0.146689 | 1 |
| Tmem209 | -0.01152 | 0.012288 | 1 |
| Rad51d | -0.01152 | 0.280457 | 1 |
| Bccip | -0.01153 | 0.439038 | 1 |
| Mrps18a | -0.01154 | 0.164359 | 1 |
| Tmem68 | -0.01155 | 0.096603 | 1 |
| Nbn | -0.01161 | 0.361604 | 1 |

| | | | |
|---|---|---|---|
| Gphn | -0.01164 | 0.05709 | 1 |
| Oaz1 | -0.01168 | 0.89465 | 1 |
| Armc5 | -0.01168 | 0.341693 | 1 |
| Tmem9b | -0.01168 | 0.333622 | 1 |
| Ntmt1 | -0.01174 | 0.034087 | 1 |
| Abcc4 | -0.01177 | 0.109883 | 1 |
| Faim | -0.01179 | 0.124464 | 1 |
| Ddx42 | -0.01187 | 0.045747 | 1 |
| Galnt7 | -0.01197 | 0.030515 | 1 |
| Exog | -0.01203 | 0.205413 | 1 |
| Ogfod2 | -0.01206 | 0.505561 | 1 |
| Asb6 | -0.0121 | 0.814009 | 1 |
| Zfp410 | -0.01211 | 0.330903 | 1 |
| Zfp141 | -0.0122 | 0.407546 | 1 |
| Spns2 | -0.01221 | 0.001711 | 1 |
| Nomo1 | -0.01222 | 0.352898 | 1 |
| Map3k2 | -0.01225 | 0.994177 | 1 |
| Dda1 | -0.01232 | 0.110257 | 1 |
| Gle1 | -0.01235 | 0.038727 | 1 |
| Washc3 | -0.01237 | 0.124463 | 1 |
| Rpap1 | -0.0124 | 0.550782 | 1 |
| Smap1 | -0.01241 | 0.203678 | 1 |
| Wdyhv1 | -0.01243 | 0.3626 | 1 |
| Epg5 | -0.01244 | 0.069016 | 1 |
| Nup50 | -0.01247 | 0.047342 | 1 |
| Galnt12 | -0.0125 | 0.111608 | 1 |
| Rabggta | -0.01261 | 0.017585 | 1 |
| Ddx55 | -0.01264 | 0.383439 | 1 |
| Galnt11 | -0.01267 | 0.06163 | 1 |
| Trmu | -0.01275 | 0.974772 | 1 |
| Ripor1 | -0.01277 | 0.185424 | 1 |
| Fastk | -0.01278 | 0.641259 | 1 |
| Cbfa2t2 | -0.01284 | 0.156747 | 1 |
| Tpd52l2 | -0.01287 | 0.303997 | 1 |
| Snx15 | -0.01287 | 0.204836 | 1 |
| BC017158 | -0.01292 | 0.41178 | 1 |
| Tada2a | -0.01297 | 0.509531 | 1 |
| Bckdhb | -0.01298 | 0.023299 | 1 |
| Poglut2 | -0.01299 | 0.596915 | 1 |
| Kif2a | -0.013 | 0.417941 | 1 |
| Foxred1 | -0.01301 | 0.197236 | 1 |
| Terf2 | -0.01306 | 0.344346 | 1 |
| Luc7l | -0.01312 | 0.181527 | 1 |
| Rnf220 | -0.01317 | 0.885094 | 1 |
| Sdhaf1 | -0.01317 | 0.00734 | 1 |
| Xpnpep3 | -0.01319 | 0.235518 | 1 |
| Utp20 | -0.01319 | 0.024267 | 1 |
| P4hb | -0.01319 | 0.43053 | 1 |

| | | | |
|---|---|---|---|
| Xpnpep1 | -0.0132 | 0.109891 | 1 |
| Trub1 | -0.01322 | 0.113982 | 1 |
| Sec22a | -0.01325 | 0.451921 | 1 |
| Ccnk | -0.01326 | 0.71732 | 1 |
| Cox7c | -0.01332 | 0.304007 | 1 |
| Laptm4a | -0.01347 | 0.814665 | 1 |
| Gstt2 | -0.0135 | 0.178918 | 1 |
| Pcyt1a | -0.01356 | 0.431045 | 1 |
| Ftsj1 | -0.01361 | 0.661167 | 1 |
| Fnbp1l | -0.01363 | 0.022798 | 1 |
| Parp2 | -0.01371 | 0.141671 | 1 |
| Cdk5rap2 | -0.01385 | 0.417163 | 1 |
| Scaf8 | -0.01385 | 0.249729 | 1 |
| Fbxo38 | -0.01388 | 0.111264 | 1 |
| Dhrs1 | -0.01401 | 0.198836 | 1 |
| 5730480H( | -0.01404 | 0.323846 | 1 |
| Rab10 | -0.01405 | 0.929051 | 1 |
| Flot1 | -0.01405 | 0.402162 | 1 |
| Ldlrap1 | -0.01406 | 0.182214 | 1 |
| Gm2245 | -0.01412 | 0.364085 | 1 |
| Prkd2 | -0.01414 | 0.124393 | 1 |
| Trmt1 | -0.01414 | 0.127187 | 1 |
| Cds1 | -0.01417 | 0.275299 | 1 |
| Ppil2 | -0.01418 | 0.538364 | 1 |
| Rbm7 | -0.01418 | 0.46752 | 1 |
| Ubxn1 | -0.01425 | 0.036313 | 1 |
| Xpr1 | -0.01432 | 0.870625 | 1 |
| Thoc5 | -0.01433 | 0.073389 | 1 |
| Otud6b | -0.01435 | 0.512948 | 1 |
| Fzr1 | -0.01439 | 0.797438 | 1 |
| Fuca2 | -0.01441 | 0.094126 | 1 |
| Naa40 | -0.01445 | 0.143754 | 1 |
| Rpf1 | -0.01446 | 0.151464 | 1 |
| Psmd6 | -0.01449 | 0.184514 | 1 |
| Tvp23b | -0.01455 | 0.588985 | 1 |
| Esr1 | -0.0146 | 0.124544 | 1 |
| Ift88 | -0.0147 | 0.171654 | 1 |
| Polr3a | -0.01477 | 0.754181 | 1 |
| Zfp160 | -0.01478 | 0.382598 | 1 |
| Sephs2 | -0.01481 | 0.648727 | 1 |
| Zgpat | -0.01481 | 0.713771 | 1 |
| Exosc5 | -0.01484 | 0.531023 | 1 |
| Nup160 | -0.01488 | 0.544062 | 1 |
| Fam53b | -0.01489 | 0.162863 | 1 |
| Cluap1 | -0.01491 | 0.826089 | 1 |
| Zkscan14 | -0.01496 | 0.965508 | 1 |
| Pfdn1 | -0.01497 | 0.092349 | 1 |
| Atg4d | -0.01501 | 0.307412 | 1 |

| | | | |
|---|---|---|---|
| Ctif | -0.01501 | 0.511112 | 1 |
| Braf | -0.01501 | 0.101896 | 1 |
| Ankrd27 | -0.01508 | 0.777055 | 1 |
| Tmem41a | -0.01508 | 0.460613 | 1 |
| Sh3bp5 | -0.0151 | 0.674198 | 1 |
| Dop1a | -0.01512 | 0.235964 | 1 |
| Psmd2 | -0.01519 | 0.702263 | 1 |
| Osbp | -0.01521 | 0.293759 | 1 |
| Aggf1 | -0.01527 | 0.292201 | 1 |
| Gdpd3 | -0.01539 | 0.24321 | 1 |
| Hectd1 | -0.01543 | 0.362914 | 1 |
| Trrap | -0.01545 | 0.070406 | 1 |
| Rsph3b | -0.0156 | 0.044353 | 1 |
| Paxbp1 | -0.0156 | 0.039265 | 1 |
| Ndufs8 | -0.01571 | 0.304408 | 1 |
| Tbpl1 | -0.01572 | 0.450917 | 1 |
| Sumo2 | -0.01582 | 0.611181 | 1 |
| Usp12 | -0.01583 | 0.000849 | 1 |
| Mdp1 | -0.01584 | 0.223069 | 1 |
| Dcaf10 | -0.01587 | 0.150605 | 1 |
| Dpy19l1 | -0.01596 | 0.090248 | 1 |
| Ccdc32 | -0.01598 | 0.670996 | 1 |
| Cep350 | -0.01607 | 0.164295 | 1 |
| Wdr77 | -0.01609 | 0.599276 | 1 |
| Zfp36l1 | -0.0161 | 0.011569 | 1 |
| Vkorc1l1 | -0.0161 | 0.068452 | 1 |
| Ldah | -0.01614 | 0.808437 | 1 |
| Dlat | -0.01618 | 0.201 | 1 |
| 6030458C1 | -0.01622 | 0.577748 | 1 |
| Ino80c | -0.01627 | 0.465886 | 1 |
| Zfp292 | -0.01628 | 0.306556 | 1 |
| Gtpbp1 | -0.01636 | 0.163205 | 1 |
| Sod1 | -0.01637 | 0.045472 | 1 |
| Crbn | -0.0164 | 0.75381 | 1 |
| Nagk | -0.01643 | 0.366815 | 1 |
| Cers5 | -0.01649 | 0.416458 | 1 |
| Lmbrd1 | -0.01654 | 0.319917 | 1 |
| Erlin1 | -0.01657 | 0.945885 | 1 |
| Itsn2 | -0.01659 | 0.273618 | 1 |
| Dctn4 | -0.01661 | 0.545596 | 1 |
| Fbxw8 | -0.01664 | 0.313528 | 1 |
| Mpg | -0.0167 | 0.428332 | 1 |
| Gosr1 | -0.01674 | 0.560355 | 1 |
| Gabarap | -0.01684 | 0.43591 | 1 |
| Tubg1 | -0.01687 | 0.757942 | 1 |
| Esco1 | -0.01694 | 0.179952 | 1 |
| Rab6a | -0.01698 | 0.232833 | 1 |
| Ccdc115 | -0.01698 | 0.387442 | 1 |

| | | | |
|---|---|---|---|
| Cct3 | -0.01698 | 0.062852 | 1 |
| Ess2 | -0.01713 | 0.448806 | 1 |
| Rps19bp1 | -0.01714 | 0.620114 | 1 |
| Zfp809 | -0.01714 | 0.08386 | 1 |
| Rnf138 | -0.01719 | 0.846606 | 1 |
| Fh1 | -0.01721 | 0.255593 | 1 |
| mt-Nd6 | -0.01723 | 0.259989 | 1 |
| Glipr1 | -0.01725 | 0.396892 | 1 |
| Ermard | -0.01728 | 0.580987 | 1 |
| Tjap1 | -0.01729 | 0.179 | 1 |
| Casp7 | -0.01731 | 0.009687 | 1 |
| Polr2i | -0.01732 | 0.102984 | 1 |
| Slc23a2 | -0.01734 | 0.305402 | 1 |
| Lsm6 | -0.01741 | 0.075026 | 1 |
| Uspl1 | -0.01746 | 0.758477 | 1 |
| Acbd5 | -0.01748 | 0.121926 | 1 |
| Selenoi | -0.0175 | 0.89141 | 1 |
| Ppm1a | -0.01751 | 0.191775 | 1 |
| Lbh | -0.01754 | 0.59429 | 1 |
| Anxa7 | -0.01755 | 0.22305 | 1 |
| Tspyl2 | -0.01766 | 0.368945 | 1 |
| Kdm2a | -0.01767 | 0.697399 | 1 |
| Btg1 | -0.0177 | 0.701953 | 1 |
| Sf3b5 | -0.0177 | 0.281833 | 1 |
| Thap4 | -0.01782 | 0.292559 | 1 |
| Rbm19 | -0.01782 | 0.204351 | 1 |
| Pfdn4 | -0.01788 | 0.262309 | 1 |
| Iws1 | -0.0179 | 0.121755 | 1 |
| Scamp1 | -0.01791 | 0.397211 | 1 |
| Tmem234 | -0.01791 | 0.155717 | 1 |
| Cds2 | -0.01791 | 0.246677 | 1 |
| Faap20 | -0.01793 | 0.17752 | 1 |
| Cops8 | -0.018 | 0.768007 | 1 |
| Erp29 | -0.01808 | 0.89137 | 1 |
| Siae | -0.01811 | 0.231911 | 1 |
| Clpp | -0.01826 | 0.185509 | 1 |
| Ammecr1l | -0.01829 | 0.143956 | 1 |
| Rtf2 | -0.01831 | 0.846283 | 1 |
| Fam98b | -0.01833 | 0.763044 | 1 |
| Chic2 | -0.01834 | 0.717974 | 1 |
| Isoc2a | -0.01834 | 0.622393 | 1 |
| Hdac8 | -0.01837 | 0.098919 | 1 |
| Ndufb4 | -0.01837 | 0.284704 | 1 |
| Ccdc88a | -0.01839 | 0.00382 | 1 |
| Nmt2 | -0.01844 | 0.082114 | 1 |
| Pdhx | -0.01844 | 0.019692 | 1 |
| A430005L1 | -0.01848 | 0.163463 | 1 |
| Fdft1 | -0.01851 | 0.400246 | 1 |

| | | | |
|---|---|---|---|
| Kpna2 | -0.01858 | 0.789387 | 1 |
| Traf7 | -0.01859 | 0.170243 | 1 |
| Khk | -0.01861 | 0.476068 | 1 |
| Uvssa | -0.01869 | 0.135526 | 1 |
| Gnb1 | -0.01869 | 0.050927 | 1 |
| Tmcc1 | -0.01875 | 0.032216 | 1 |
| Uba3 | -0.01881 | 0.638656 | 1 |
| Zwint | -0.01881 | 0.321231 | 1 |
| Arntl | -0.01883 | 0.633854 | 1 |
| Smarca5 | -0.01887 | 0.009615 | 1 |
| Tmem208 | -0.01888 | 0.371491 | 1 |
| Pcgf3 | -0.01894 | 0.291667 | 1 |
| Eml3 | -0.01896 | 0.975324 | 1 |
| Aspscr1 | -0.01896 | 0.164809 | 1 |
| Mrps30 | -0.01903 | 0.154212 | 1 |
| Pole4 | -0.01907 | 0.177743 | 1 |
| Dnttip2 | -0.0191 | 0.731475 | 1 |
| Txndc11 | -0.01914 | 0.529253 | 1 |
| Timm8b | -0.0192 | 0.266974 | 1 |
| Fam20b | -0.01924 | 0.11475 | 1 |
| Smg1 | -0.0193 | 0.061804 | 1 |
| Dmap1 | -0.01932 | 0.203825 | 1 |
| Pmpca | -0.01933 | 0.701015 | 1 |
| Shc1 | -0.01933 | 0.147761 | 1 |
| C2cd2l | -0.01934 | 0.599056 | 1 |
| Rhbdd3 | -0.01934 | 0.333026 | 1 |
| Sc5d | -0.01942 | 0.280946 | 1 |
| Maip1 | -0.01943 | 0.220191 | 1 |
| Fam49b | -0.01945 | 0.310199 | 1 |
| Nudt4 | -0.01953 | 0.106468 | 1 |
| Asnsd1 | -0.01954 | 0.32224 | 1 |
| Mapk7 | -0.01955 | 0.942765 | 1 |
| Gskip | -0.01956 | 0.931353 | 1 |
| Slc8b1 | -0.01958 | 0.133108 | 1 |
| Ksr1 | -0.01958 | 0.549829 | 1 |
| Fbxo11 | -0.01959 | 0.022898 | 1 |
| Zfp780b | -0.01959 | 0.455342 | 1 |
| Alg11 | -0.01963 | 0.467031 | 1 |
| Rsrc2 | -0.01964 | 0.378393 | 1 |
| Agfg1 | -0.01965 | 0.870938 | 1 |
| Asl | -0.01965 | 0.137822 | 1 |
| Pdcd7 | -0.01967 | 0.403761 | 1 |
| Ccl25 | -0.0197 | 0.087535 | 1 |
| Zbtb8os | -0.01974 | 0.263555 | 1 |
| Eif2s2 | -0.01975 | 0.48389 | 1 |
| Mrps7 | -0.01977 | 0.687317 | 1 |
| Nipsnap2 | -0.0198 | 0.125386 | 1 |
| Rmnd5b | -0.01982 | 0.707076 | 1 |

| | | | |
|---|---|---|---|
| Dxo | -0.01985 | 0.880462 | 1 |
| Ptms | -0.01986 | 0.431747 | 1 |
| Tmem184c | -0.01988 | 0.138403 | 1 |
| Brms1l | -0.01995 | 0.479588 | 1 |
| Atxn7l1 | -0.01997 | 0.586119 | 1 |
| Acsf2 | -0.01999 | 0.324694 | 1 |
| Rnf20 | -0.02001 | 0.569334 | 1 |
| Naa16 | -0.02004 | 0.135364 | 1 |
| Hnrnpul2 | -0.02006 | 0.709154 | 1 |
| Cul4a | -0.02012 | 0.233039 | 1 |
| Opa3 | -0.02012 | 0.591257 | 1 |
| Ppt1 | -0.02013 | 0.497272 | 1 |
| E330020D1 | -0.02017 | 0.24179 | 1 |
| Abhd10 | -0.02018 | 0.569917 | 1 |
| U2surp | -0.02031 | 0.115236 | 1 |
| Cenpo | -0.02034 | 0.131173 | 1 |
| Sp3os | -0.02035 | 0.261814 | 1 |
| Rcor1 | -0.02038 | 0.029545 | 1 |
| Atg2b | -0.02045 | 0.545467 | 1 |
| Mettl2 | -0.0205 | 0.165918 | 1 |
| Hgs | -0.0205 | 0.5038 | 1 |
| Rfx5 | -0.02052 | 0.087955 | 1 |
| Rrad | -0.02058 | 0.783249 | 1 |
| Zfp335 | -0.0207 | 0.608668 | 1 |
| Samd4b | -0.02075 | 0.457595 | 1 |
| Mrpl49 | -0.02075 | 0.981493 | 1 |
| Bnip3 | -0.02075 | 0.826533 | 1 |
| Fam126a | -0.02076 | 0.75662 | 1 |
| Ranbp3 | -0.02077 | 0.605561 | 1 |
| Stim1 | -0.0208 | 0.324538 | 1 |
| Rrp8 | -0.02085 | 0.658197 | 1 |
| Slc35a2 | -0.02094 | 0.232209 | 1 |
| Als2 | -0.02096 | 0.338447 | 1 |
| Fam133b | -0.02099 | 0.118177 | 1 |
| Med29 | -0.02106 | 0.174945 | 1 |
| Usp5 | -0.02108 | 0.294113 | 1 |
| Mrpl24 | -0.02108 | 0.224328 | 1 |
| Cnot1 | -0.02114 | 0.256277 | 1 |
| Trp53inp1 | -0.02114 | 0.391233 | 1 |
| Nans | -0.02117 | 0.123305 | 1 |
| Stk38l | -0.02118 | 0.156403 | 1 |
| Pebp1 | -0.02127 | 0.146888 | 1 |
| Psmg1 | -0.02133 | 0.115451 | 1 |
| Dbt | -0.02137 | 0.736798 | 1 |
| Lsm3 | -0.02138 | 0.338059 | 1 |
| Aida | -0.02138 | 0.202141 | 1 |
| Rab28 | -0.02141 | 0.432983 | 1 |
| Myl12b | -0.02143 | 0.694284 | 1 |

| | | | |
|---|---|---|---|
| Thumpd1 | -0.02144 | 0.779295 | 1 |
| Cab39l | -0.02152 | 0.401022 | 1 |
| Ankrd13a | -0.02154 | 0.199676 | 1 |
| Trim26 | -0.02163 | 0.9601 | 1 |
| Hipk3 | -0.02163 | 0.678889 | 1 |
| Mtx1 | -0.02164 | 0.319529 | 1 |
| Pan2 | -0.02167 | 0.460321 | 1 |
| Drap1 | -0.02168 | 0.631179 | 1 |
| Zscan29 | -0.02174 | 0.493746 | 1 |
| Usp6nl | -0.0218 | 0.350772 | 1 |
| Chpf2 | -0.02181 | 0.89228 | 1 |
| Pnkp | -0.02183 | 0.154806 | 1 |
| Snf8 | -0.02186 | 0.710293 | 1 |
| Lias | -0.02188 | 0.311425 | 1 |
| Nsun3 | -0.02193 | 0.110432 | 1 |
| Usp42 | -0.02208 | 0.448122 | 1 |
| Znhit2 | -0.02209 | 0.510386 | 1 |
| Mcu | -0.0221 | 0.237013 | 1 |
| Iffo2 | -0.02211 | 0.41795 | 1 |
| Tmem179b | -0.02212 | 0.160112 | 1 |
| Stt3b | -0.02215 | 0.141713 | 1 |
| S100a13 | -0.02216 | 0.041118 | 1 |
| Zfp574 | -0.0222 | 0.502261 | 1 |
| Aldh3b1 | -0.02224 | 0.515266 | 1 |
| S100pbp | -0.02226 | 0.998776 | 1 |
| Natd1 | -0.02226 | 0.766119 | 1 |
| Hist1h1c | -0.02228 | 0.404868 | 1 |
| Gtf2ird2 | -0.02228 | 0.815937 | 1 |
| Ttc1 | -0.02232 | 0.457677 | 1 |
| Pik3cd | -0.02234 | 0.264796 | 1 |
| Snrpc | -0.02246 | 0.17799 | 1 |
| Lsm2 | -0.0225 | 0.23506 | 1 |
| Ap5b1 | -0.0225 | 0.533766 | 1 |
| Gm7072 | -0.02252 | 0.859999 | 1 |
| Zfp282 | -0.02253 | 0.927065 | 1 |
| Rrp1 | -0.02266 | 0.126403 | 1 |
| Vps16 | -0.02266 | 0.344873 | 1 |
| Ahsa1 | -0.02272 | 0.402465 | 1 |
| 2700097O0 | -0.02275 | 0.301424 | 1 |
| Mkrn2 | -0.0228 | 0.182713 | 1 |
| Clasrp | -0.02282 | 0.478516 | 1 |
| Edf1 | -0.02285 | 0.10345 | 1 |
| Eef1akmt1 | -0.02287 | 0.514044 | 1 |
| Ctdp1 | -0.02292 | 0.335358 | 1 |
| Rfwd3 | -0.02297 | 0.349379 | 1 |
| Map4 | -0.02299 | 0.240331 | 1 |
| Rnf41 | -0.02305 | 0.222852 | 1 |
| Ttc37 | -0.02313 | 0.027768 | 1 |

| | | | |
|---|---|---|---|
| Fancc | -0.02313 | 0.170872 | 1 |
| Nfyc | -0.02314 | 0.845625 | 1 |
| Fbh1 | -0.02321 | 0.08043 | 1 |
| Ino80b | -0.02325 | 0.648803 | 1 |
| Btbd1 | -0.02348 | 0.224617 | 1 |
| Pkd1 | -0.0235 | 0.696003 | 1 |
| Lrp6 | -0.0235 | 0.876285 | 1 |
| Bclaf3 | -0.02359 | 0.880008 | 1 |
| Cdc14a | -0.02361 | 0.16867 | 1 |
| Mtpap | -0.02363 | 0.417452 | 1 |
| Snrpb2 | -0.02363 | 0.237912 | 1 |
| Mib1 | -0.02364 | 0.210107 | 1 |
| Sv2a | -0.02368 | 0.263972 | 1 |
| Usp15 | -0.02374 | 0.234111 | 1 |
| Auh | -0.02374 | 0.940679 | 1 |
| Setx | -0.02375 | 0.590305 | 1 |
| Pcnt | -0.02377 | 0.726298 | 1 |
| Riok1 | -0.02391 | 0.082419 | 1 |
| Gucd1 | -0.02392 | 0.816531 | 1 |
| Fuca1 | -0.02397 | 0.738195 | 1 |
| Spopl | -0.024 | 0.183548 | 1 |
| Irf3 | -0.02403 | 0.08724 | 1 |
| Med12 | -0.02403 | 0.227853 | 1 |
| Akap11 | -0.02405 | 0.239109 | 1 |
| Mrgbp | -0.02406 | 0.14225 | 1 |
| Rel | -0.02406 | 0.936261 | 1 |
| Slc39a3 | -0.02414 | 0.583532 | 1 |
| Srcap | -0.02417 | 0.924427 | 1 |
| Ints1 | -0.02431 | 0.284022 | 1 |
| 1110038F1 | -0.02435 | 0.378462 | 1 |
| Akap10 | -0.02436 | 0.28958 | 1 |
| Strn3 | -0.02442 | 0.122357 | 1 |
| Chchd5 | -0.02444 | 0.345427 | 1 |
| Nacc1 | -0.02446 | 0.707533 | 1 |
| Camta1 | -0.02447 | 0.042338 | 1 |
| Kpna1 | -0.02449 | 0.927758 | 1 |
| Plcg2 | -0.02453 | 0.618354 | 1 |
| Kin | -0.02455 | 0.401457 | 1 |
| Parg | -0.02457 | 0.366562 | 1 |
| Enc1 | -0.02458 | 0.775626 | 1 |
| Ubxn6 | -0.02461 | 0.766952 | 1 |
| Rffl | -0.02468 | 0.178832 | 1 |
| Ddx47 | -0.02472 | 0.530658 | 1 |
| Rps6kb1 | -0.02478 | 0.124197 | 1 |
| Prkaa1 | -0.0248 | 0.305601 | 1 |
| Hsbp1 | -0.02483 | 0.54065 | 1 |
| Fance | -0.02484 | 0.463962 | 1 |
| Cwc25 | -0.02486 | 0.155899 | 1 |

| | | | |
|---|---|---|---|
| Psmc5 | -0.0249 | 0.616025 | 1 |
| Tlnrd1 | -0.02491 | 0.663126 | 1 |
| Mllt10 | -0.02491 | 0.979087 | 1 |
| Ttc33 | -0.02491 | 0.920001 | 1 |
| Sipa1l3 | -0.02494 | 0.066618 | 1 |
| Snx20 | -0.02495 | 0.23261 | 1 |
| Hint2 | -0.02499 | 0.07711 | 1 |
| Phka2 | -0.02509 | 0.116142 | 1 |
| Taf1 | -0.02511 | 0.252004 | 1 |
| Rttn | -0.02513 | 0.321439 | 1 |
| Ppp5c | -0.02513 | 0.429045 | 1 |
| Snx24 | -0.02514 | 0.528669 | 1 |
| Zfp407 | -0.02519 | 0.035506 | 1 |
| Tmem222 | -0.0252 | 0.41394 | 1 |
| Lsm7 | -0.02526 | 0.055525 | 1 |
| Rex1bd | -0.0253 | 0.125836 | 1 |
| Slc35a1 | -0.02533 | 0.861007 | 1 |
| Dcaf6 | -0.02536 | 0.951241 | 1 |
| Zfp800 | -0.02539 | 0.831342 | 1 |
| Zfp983 | -0.02546 | 0.937644 | 1 |
| Taf2 | -0.0255 | 0.54728 | 1 |
| Araf | -0.02551 | 0.95309 | 1 |
| Sde2 | -0.02551 | 0.476881 | 1 |
| Psmd4 | -0.02553 | 0.197156 | 1 |
| Fam234a | -0.02558 | 0.351069 | 1 |
| Ppp1r37 | -0.02561 | 0.141331 | 1 |
| Ube2n | -0.02566 | 0.201708 | 1 |
| Ncbp2 | -0.02566 | 0.491451 | 1 |
| Rexo2 | -0.0257 | 0.724136 | 1 |
| Anks1 | -0.02572 | 0.076753 | 1 |
| Elp5 | -0.02579 | 0.348857 | 1 |
| Spty2d1 | -0.02583 | 0.941756 | 1 |
| Mettl9 | -0.02588 | 0.265174 | 1 |
| Crtc2 | -0.0259 | 0.910308 | 1 |
| Gbe1 | -0.02591 | 0.792724 | 1 |
| Rdm1 | -0.02593 | 0.257936 | 1 |
| Atp13a2 | -0.02599 | 0.027466 | 1 |
| Taf6 | -0.02599 | 0.622729 | 1 |
| Bag4 | -0.02602 | 0.998995 | 1 |
| Lpgat1 | -0.02608 | 0.326386 | 1 |
| Zfp771 | -0.02608 | 0.249137 | 1 |
| Klc2 | -0.02622 | 0.318603 | 1 |
| Gpbp1l1 | -0.02625 | 0.613619 | 1 |
| Primpol | -0.02629 | 0.144811 | 1 |
| Mtln | -0.02641 | 0.968707 | 1 |
| Uqcrc2 | -0.02647 | 0.933606 | 1 |
| Thumpd3 | -0.02648 | 0.230455 | 1 |
| Kras | -0.02649 | 0.792928 | 1 |

| | | | |
|---|---|---|---|
| Kmt2d | -0.02653 | 0.452855 | 1 |
| Trpv2 | -0.02655 | 0.327276 | 1 |
| Gga3 | -0.02657 | 0.60624 | 1 |
| Saal1 | -0.0266 | 0.784304 | 1 |
| Bcl2l12 | -0.02663 | 0.947734 | 1 |
| Cmpk1 | -0.02665 | 0.342239 | 1 |
| Uty | -0.02665 | 0.251682 | 1 |
| Rapsn | -0.02666 | 0.0445 | 1 |
| Grpel1 | -0.02666 | 0.911807 | 1 |
| Mast4 | -0.02674 | 0.053623 | 1 |
| Eif2s3y | -0.02678 | 0.251207 | 1 |
| Vps33a | -0.02683 | 0.293563 | 1 |
| Sprtn | -0.02689 | 0.781253 | 1 |
| Pdzd11 | -0.0269 | 0.403181 | 1 |
| Hmgxb3 | -0.02694 | 0.168234 | 1 |
| Stk16 | -0.02698 | 0.095486 | 1 |
| Ciao2b | -0.02702 | 0.333804 | 1 |
| Dhx29 | -0.02702 | 0.808038 | 1 |
| Mphosph6 | -0.02702 | 0.079681 | 1 |
| Vti1b | -0.02703 | 0.285917 | 1 |
| Rc3h1 | -0.02705 | 0.221376 | 1 |
| Galnt1 | -0.02707 | 0.289745 | 1 |
| Jmjd8 | -0.02707 | 0.370999 | 1 |
| Ric8a | -0.02713 | 0.355265 | 1 |
| Atp5f1 | -0.02719 | 0.912908 | 1 |
| Zhx1 | -0.02719 | 0.113304 | 1 |
| Tmtc3 | -0.02723 | 0.559044 | 1 |
| Anapc15 | -0.02724 | 0.002264 | 1 |
| Ncoa6 | -0.02738 | 0.867456 | 1 |
| Grpel2 | -0.02743 | 0.971787 | 1 |
| Gigyf2 | -0.02755 | 0.283146 | 1 |
| Mtrex | -0.02759 | 0.924502 | 1 |
| Amz1 | -0.02762 | 0.059164 | 1 |
| Selenot | -0.02763 | 0.395183 | 1 |
| Zfp146 | -0.02764 | 0.569188 | 1 |
| Tmem243 | -0.02765 | 0.455189 | 1 |
| Paxx | -0.02778 | 0.429101 | 1 |
| Akap13 | -0.02779 | 0.263649 | 1 |
| Pitpnc1 | -0.02785 | 0.126373 | 1 |
| Nudt5 | -0.02788 | 0.354459 | 1 |
| Stk17b | -0.02791 | 0.190845 | 1 |
| Setd2 | -0.02792 | 0.69115 | 1 |
| Hspa14 | -0.02794 | 0.364774 | 1 |
| Phyh | -0.02795 | 0.243373 | 1 |
| Aifm1 | -0.02798 | 0.149506 | 1 |
| Nutf2 | -0.028 | 0.459373 | 1 |
| Imp3 | -0.02802 | 0.941194 | 1 |
| Abt1 | -0.02802 | 0.764769 | 1 |

| | | | |
|---|---|---|---|
| Sac3d1 | -0.02812 | 0.49694 | 1 |
| Tcp11l1 | -0.02818 | 0.307622 | 1 |
| Fra10ac1 | -0.02825 | 0.854799 | 1 |
| Pdcd2l | -0.02826 | 0.99319 | 1 |
| Ints5 | -0.02831 | 0.987184 | 1 |
| Them6 | -0.02832 | 0.708956 | 1 |
| Amn1 | -0.02843 | 0.664285 | 1 |
| Dhx33 | -0.02853 | 0.255328 | 1 |
| Ptbp2 | -0.02857 | 0.430562 | 1 |
| Imp4 | -0.0286 | 0.107967 | 1 |
| Wdr82 | -0.02862 | 0.798653 | 1 |
| Klhl26 | -0.02863 | 0.704967 | 1 |
| Hpf1 | -0.02865 | 0.262522 | 1 |
| Hsf1 | -0.02865 | 0.963294 | 1 |
| Chchd7 | -0.02866 | 0.52297 | 1 |
| Usp40 | -0.02866 | 0.441172 | 1 |
| Dctn6 | -0.02866 | 0.64827 | 1 |
| Tasp1 | -0.02867 | 0.110234 | 1 |
| Snx8 | -0.02868 | 0.79979 | 1 |
| Cep120 | -0.02881 | 0.612936 | 1 |
| Pqlc2 | -0.02884 | 0.339488 | 1 |
| Commd1 | -0.02885 | 0.34986 | 1 |
| Golph3l | -0.02894 | 0.594745 | 1 |
| C330018D1 | -0.02896 | 0.93425 | 1 |
| Ptdss1 | -0.02904 | 0.100382 | 1 |
| Rnmt | -0.02905 | 0.199317 | 1 |
| Dph7 | -0.02905 | 0.844474 | 1 |
| Brwd3 | -0.0291 | 0.370823 | 1 |
| Phrf1 | -0.02915 | 0.555908 | 1 |
| Depdc5 | -0.02922 | 0.472149 | 1 |
| Nmnat3 | -0.02923 | 0.391069 | 1 |
| Lrrc57 | -0.02926 | 0.688292 | 1 |
| Pdpk1 | -0.02937 | 0.513077 | 1 |
| Pfdn2 | -0.02942 | 0.52482 | 1 |
| Kansl3 | -0.02944 | 0.008007 | 1 |
| Smarcad1 | -0.0295 | 0.882222 | 1 |
| Nxt1 | -0.02951 | 0.576159 | 1 |
| Tmem18 | -0.02953 | 0.999543 | 1 |
| Nectin4 | -0.02954 | 0.030392 | 1 |
| Rnf40 | -0.0296 | 0.482572 | 1 |
| Tti2 | -0.02961 | 0.940846 | 1 |
| Yif1a | -0.02964 | 0.485935 | 1 |
| Mul1 | -0.02966 | 0.326569 | 1 |
| Ndufs7 | -0.02968 | 0.613375 | 1 |
| Anks3 | -0.02968 | 0.701787 | 1 |
| Slc41a3 | -0.02973 | 0.951413 | 1 |
| Fgfr1op2 | -0.02977 | 0.510536 | 1 |
| Pop5 | -0.02978 | 0.42087 | 1 |

| | | | |
|---|---|---|---|
| Asah2 | -0.02987 | 0.184764 | 1 |
| Fip1l1 | -0.02996 | 0.841503 | 1 |
| Pafah1b2 | -0.02997 | 0.400235 | 1 |
| Smim4 | -0.03003 | 0.257604 | 1 |
| Acadvl | -0.03003 | 0.45452 | 1 |
| Copg1 | -0.0301 | 0.065212 | 1 |
| Tmem185a | -0.03018 | 0.411881 | 1 |
| Ibtk | -0.03019 | 0.206225 | 1 |
| Nlrp1a | -0.03021 | 0.008865 | 1 |
| Cct4 | -0.03022 | 0.881924 | 1 |
| Rnf10 | -0.03023 | 0.144497 | 1 |
| Med4 | -0.03031 | 0.546597 | 1 |
| Ptcd2 | -0.03043 | 0.792512 | 1 |
| Fam53a | -0.03043 | 0.676847 | 1 |
| Pdlim2 | -0.03044 | 0.631068 | 1 |
| Rnf185 | -0.03044 | 0.157785 | 1 |
| Haus3 | -0.03051 | 0.459101 | 1 |
| Paip1 | -0.03053 | 0.508116 | 1 |
| Cep135 | -0.03054 | 0.27789 | 1 |
| E2f4 | -0.03062 | 0.990154 | 1 |
| Rrn3 | -0.03063 | 0.168273 | 1 |
| Cwc22 | -0.03068 | 0.610168 | 1 |
| Prr14l | -0.03069 | 0.653615 | 1 |
| Dhx36 | -0.03071 | 0.362011 | 1 |
| Uggt1 | -0.03071 | 0.709025 | 1 |
| Ecd | -0.03073 | 0.500057 | 1 |
| Osbpl7 | -0.03078 | 0.325998 | 1 |
| Fcho2 | -0.0308 | 0.747549 | 1 |
| Ino80e | -0.03082 | 0.292642 | 1 |
| Gm1976 | -0.03089 | 0.547934 | 1 |
| 1810030O0 | -0.0309 | 0.526959 | 1 |
| Zfp87 | -0.0309 | 0.937166 | 1 |
| Hdac3 | -0.03095 | 0.863552 | 1 |
| Decr1 | -0.03095 | 0.92163 | 1 |
| 9330159M | -0.03101 | 0.106071 | 1 |
| Wdr61 | -0.03105 | 0.101105 | 1 |
| Chst12 | -0.03106 | 0.40006 | 1 |
| Sccpdh | -0.0311 | 0.540043 | 1 |
| Cactin | -0.03117 | 0.594884 | 1 |
| Arfrp1 | -0.03119 | 0.176942 | 1 |
| Ddx3y | -0.03121 | 0.674035 | 1 |
| Nudt13 | -0.03122 | 0.303878 | 1 |
| Nde1 | -0.03122 | 0.322891 | 1 |
| Pias4 | -0.0313 | 0.265522 | 1 |
| Rnasek | -0.03133 | 0.902755 | 1 |
| Ash2l | -0.03136 | 0.962731 | 1 |
| BC005561 | -0.03139 | 0.133804 | 1 |
| Dnajb9 | -0.03142 | 0.548513 | 1 |

| | | | |
|---|---|---|---|
| Npepps | -0.03144 | 0.314038 | 1 |
| Zfp597 | -0.03147 | 0.926436 | 1 |
| Pigf | -0.0315 | 0.074159 | 1 |
| Rgs3 | -0.03153 | 0.004127 | 1 |
| Mettl16 | -0.03157 | 0.537536 | 1 |
| Ap2m1 | -0.03158 | 0.875303 | 1 |
| 2510039O | -0.03162 | 0.711545 | 1 |
| Cox19 | -0.03167 | 0.897827 | 1 |
| Actr5 | -0.03168 | 0.911789 | 1 |
| Cdc16 | -0.0317 | 0.448291 | 1 |
| Eci1 | -0.03171 | 0.730761 | 1 |
| Miip | -0.03174 | 0.28826 | 1 |
| Mrpl53 | -0.03179 | 0.667651 | 1 |
| Cfap36 | -0.03183 | 0.514904 | 1 |
| Taf1b | -0.03185 | 0.329084 | 1 |
| Slc36a1 | -0.03187 | 0.102478 | 1 |
| Pycrl | -0.0319 | 0.174931 | 1 |
| Nfe2l1 | -0.03193 | 0.460931 | 1 |
| Lamtor2 | -0.03196 | 0.176782 | 1 |
| Phykpl | -0.03199 | 0.975018 | 1 |
| Sbf1 | -0.032 | 0.463037 | 1 |
| Fyco1 | -0.03212 | 0.533045 | 1 |
| Cuta | -0.03216 | 0.412044 | 1 |
| Rbm45 | -0.03218 | 0.735753 | 1 |
| Gpr157 | -0.03219 | 0.854055 | 1 |
| Gfpt1 | -0.03221 | 0.146576 | 1 |
| Hspbp1 | -0.03222 | 0.14776 | 1 |
| Slx4ip | -0.03231 | 0.083309 | 1 |
| Mnat1 | -0.03248 | 0.053038 | 1 |
| Cdc37l1 | -0.03248 | 0.716223 | 1 |
| Mrps14 | -0.03249 | 0.448742 | 1 |
| Nup107 | -0.0325 | 0.639029 | 1 |
| Recql | -0.03253 | 0.456607 | 1 |
| Trim33 | -0.03255 | 0.967854 | 1 |
| Gng2 | -0.03257 | 0.920307 | 1 |
| Preb | -0.03279 | 0.434762 | 1 |
| Caap1 | -0.03279 | 0.814512 | 1 |
| Tbc1d32 | -0.03283 | 0.954678 | 1 |
| Brd4 | -0.03285 | 0.283228 | 1 |
| Pptc7 | -0.03292 | 0.206876 | 1 |
| Adprhl2 | -0.03297 | 0.953585 | 1 |
| Dpm1 | -0.03297 | 0.320009 | 1 |
| Nsf | -0.03302 | 0.458833 | 1 |
| Cpt1a | -0.03308 | 0.382884 | 1 |
| Scai | -0.0331 | 0.005894 | 1 |
| Dhx16 | -0.03311 | 0.954131 | 1 |
| Eif2b1 | -0.03319 | 0.791784 | 1 |
| Snrpb | -0.03319 | 0.843717 | 1 |

| | | | |
|---|---|---|---|
| Zfp26 | -0.0332 | 0.256412 | 1 |
| Wdr11 | -0.03322 | 0.802409 | 1 |
| Sec24d | -0.03324 | 0.637783 | 1 |
| Mrps6 | -0.03327 | 0.529703 | 1 |
| Vps53 | -0.03339 | 0.120638 | 1 |
| Dynlt1c | -0.03344 | 0.213741 | 1 |
| 8030462N1 | -0.03344 | 0.985303 | 1 |
| Mtmr6 | -0.03345 | 0.445579 | 1 |
| Rinl | -0.03353 | 0.033606 | 1 |
| Taf8 | -0.03356 | 0.695373 | 1 |
| Yipf6 | -0.03365 | 0.998357 | 1 |
| Mtch1 | -0.03366 | 0.246315 | 1 |
| Zfp709 | -0.03371 | 0.160132 | 1 |
| Fam98a | -0.03374 | 0.93335 | 1 |
| Zfp943 | -0.03378 | 0.703985 | 1 |
| Zfand1 | -0.03381 | 0.912265 | 1 |
| Tomm34 | -0.03383 | 0.425188 | 1 |
| Trappc1 | -0.03383 | 0.330465 | 1 |
| Rufy2 | -0.03384 | 0.945244 | 1 |
| Cnih1 | -0.03384 | 0.196083 | 1 |
| Zdhhc9 | -0.03388 | 0.975584 | 1 |
| Trappc5 | -0.03389 | 0.599325 | 1 |
| Cyth2 | -0.03393 | 0.755832 | 1 |
| Nr1d2 | -0.03395 | 0.107696 | 1 |
| Necap1 | -0.03397 | 0.952363 | 1 |
| Spata1 | -0.03399 | 0.677605 | 1 |
| Rictor | -0.03399 | 0.377919 | 1 |
| Kat6b | -0.03409 | 0.102098 | 1 |
| Cdk4 | -0.0341 | 0.663327 | 1 |
| Polr2g | -0.0341 | 0.364278 | 1 |
| Stk25 | -0.03415 | 0.982567 | 1 |
| Fkrp | -0.03421 | 0.799198 | 1 |
| Ift140 | -0.03427 | 0.806587 | 1 |
| Nol7 | -0.03429 | 0.225088 | 1 |
| Agtpbp1 | -0.0343 | 0.108178 | 1 |
| Rad54l2 | -0.03432 | 0.874756 | 1 |
| Ap5m1 | -0.03433 | 0.74873 | 1 |
| Rfk | -0.03434 | 0.960128 | 1 |
| Zfp991 | -0.03436 | 0.513818 | 1 |
| Trim44 | -0.03442 | 0.222767 | 1 |
| Ghitm | -0.03442 | 0.312764 | 1 |
| Setd1a | -0.03442 | 0.161921 | 1 |
| Tmem19 | -0.03444 | 0.923477 | 1 |
| Chtop | -0.03449 | 0.700656 | 1 |
| Chst1 | -0.03449 | 0.403395 | 1 |
| Arhgap30 | -0.03454 | 0.043607 | 1 |
| Foxj2 | -0.03458 | 0.748982 | 1 |
| Tsga10 | -0.0346 | 0.459442 | 1 |

| | | | |
|---|---|---|---|
| Bfar | -0.0346 | 0.02833 | 1 |
| Efr3a | -0.03461 | 0.853542 | 1 |
| Tcf25 | -0.03463 | 0.760406 | 1 |
| Clk1 | -0.0347 | 0.657802 | 1 |
| Orc2 | -0.03471 | 0.482629 | 1 |
| Furin | -0.03474 | 0.112643 | 1 |
| Cant1 | -0.03488 | 0.685204 | 1 |
| Mfn2 | -0.03491 | 0.586512 | 1 |
| Rasa1 | -0.03505 | 0.362618 | 1 |
| Pex7 | -0.03507 | 0.706345 | 1 |
| Cpt2 | -0.03507 | 0.378506 | 1 |
| Tbc1d13 | -0.03513 | 0.386224 | 1 |
| Mettl25 | -0.03513 | 0.366377 | 1 |
| Chmp2a | -0.03514 | 0.915219 | 1 |
| Foxk1 | -0.03515 | 0.844483 | 1 |
| Mpdu1 | -0.03515 | 0.981422 | 1 |
| Rnf181 | -0.03518 | 0.205781 | 1 |
| Gtpbp8 | -0.03519 | 0.541526 | 1 |
| Hes6 | -0.03521 | 0.862413 | 1 |
| Fkbp3 | -0.03525 | 0.800594 | 1 |
| Cbr4 | -0.03526 | 0.637373 | 1 |
| Zfp207 | -0.03532 | 0.498175 | 1 |
| Zfyve1 | -0.03533 | 0.895444 | 1 |
| Sgpl1 | -0.03535 | 0.27551 | 1 |
| Stxbp1 | -0.03543 | 0.946714 | 1 |
| Khnyn | -0.03548 | 0.6128 | 1 |
| Cdc42bpb | -0.0355 | 0.007914 | 1 |
| Malt1 | -0.03551 | 0.089364 | 1 |
| Terf2ip | -0.03552 | 0.068571 | 1 |
| Sap30bp | -0.03556 | 0.221133 | 1 |
| Anp32e | -0.03556 | 0.832914 | 1 |
| Gpr108 | -0.03556 | 0.138548 | 1 |
| Pigx | -0.03561 | 0.658736 | 1 |
| Zfp688 | -0.03569 | 0.618821 | 1 |
| Zfp182 | -0.03573 | 0.514776 | 1 |
| Krit1 | -0.03578 | 0.135753 | 1 |
| Hdac1 | -0.0358 | 0.502162 | 1 |
| Stat5a | -0.03581 | 0.407148 | 1 |
| Zfp444 | -0.03581 | 0.733384 | 1 |
| Cep192 | -0.03587 | 0.322337 | 1 |
| Mrpl44 | -0.03588 | 0.525088 | 1 |
| Ddx52 | -0.03598 | 0.75497 | 1 |
| Med27 | -0.03601 | 0.284117 | 1 |
| Nacc2 | -0.03607 | 0.515418 | 1 |
| Coa6 | -0.0361 | 0.858156 | 1 |
| Uck1 | -0.03615 | 0.675686 | 1 |
| Dtx2 | -0.03623 | 0.215563 | 1 |
| Mus81 | -0.03632 | 0.142061 | 1 |

| | | | |
|---|---|---|---|
| Sf3a3 | -0.03633 | 0.826991 | 1 |
| Eaf1 | -0.03637 | 0.406607 | 1 |
| Trmt2b | -0.03639 | 0.69075 | 1 |
| Psmg2 | -0.03641 | 0.474452 | 1 |
| Foxk2 | -0.03642 | 0.798191 | 1 |
| Abca3 | -0.03644 | 0.319915 | 1 |
| Vps45 | -0.03647 | 0.323806 | 1 |
| Bet1 | -0.03651 | 0.996914 | 1 |
| Mettl3 | -0.03659 | 0.955878 | 1 |
| 1810037I1 | -0.03661 | 0.94263 | 1 |
| Ttc14 | -0.03668 | 0.638689 | 1 |
| Gmcl1 | -0.03671 | 0.882484 | 1 |
| 4931406P1 | -0.03672 | 0.439276 | 1 |
| Lin7c | -0.03676 | 0.731297 | 1 |
| Dicer1 | -0.03676 | 0.21864 | 1 |
| Zfyve16 | -0.03687 | 0.919141 | 1 |
| Ipo5 | -0.03692 | 0.329341 | 1 |
| Gapvd1 | -0.03694 | 0.83843 | 1 |
| Poglut1 | -0.03698 | 0.518859 | 1 |
| Osbpl1a | -0.037 | 0.788635 | 1 |
| Eya3 | -0.03703 | 0.342089 | 1 |
| Phactr4 | -0.03716 | 0.508632 | 1 |
| Ppp6r3 | -0.03722 | 0.664686 | 1 |
| Usp48 | -0.03727 | 0.274129 | 1 |
| Map2k5 | -0.03728 | 0.503164 | 1 |
| Slc4a1ap | -0.03728 | 0.837728 | 1 |
| Eny2 | -0.03729 | 0.462175 | 1 |
| Mtmr9 | -0.03735 | 0.876313 | 1 |
| Ptp4a2 | -0.03736 | 0.285061 | 1 |
| Brpf1 | -0.03741 | 0.206809 | 1 |
| Klhl18 | -0.03742 | 0.370125 | 1 |
| Malat1 | -0.03743 | 0.00096 | 1 |
| Tmem147 | -0.03746 | 0.221437 | 1 |
| Slc35b4 | -0.03747 | 0.882514 | 1 |
| Dnajc30 | -0.03752 | 0.468307 | 1 |
| 2900097C1 | -0.03754 | 0.322904 | 1 |
| Rcl1 | -0.0376 | 0.2934 | 1 |
| 1700037C1 | -0.03762 | 0.551166 | 1 |
| Mllt11 | -0.03766 | 0.570463 | 1 |
| Tmem127 | -0.03768 | 0.792246 | 1 |
| Slc35a4 | -0.0378 | 0.302852 | 1 |
| Med10 | -0.03789 | 0.481119 | 1 |
| Clns1a | -0.03791 | 0.658103 | 1 |
| Ap4e1 | -0.03793 | 0.682856 | 1 |
| Cops5 | -0.03794 | 0.289257 | 1 |
| Pelp1 | -0.03796 | 0.210479 | 1 |
| Ngdn | -0.03799 | 0.799701 | 1 |
| Lsm8 | -0.038 | 0.827232 | 1 |

| | | | |
|---|---|---|---|
| Rbm8a | -0.03803 | 0.796482 | 1 |
| Mrps5 | -0.03811 | 0.528809 | 1 |
| Man2c1 | -0.03822 | 0.870173 | 1 |
| 4833420G: | -0.03827 | 0.380815 | 1 |
| Atf1 | -0.03828 | 0.168952 | 1 |
| Fbxw9 | -0.03831 | 0.801231 | 1 |
| Pdpr | -0.03833 | 0.588324 | 1 |
| Gtf3c6 | -0.03836 | 0.130368 | 1 |
| Zbtb43 | -0.0385 | 0.587572 | 1 |
| Tmem260 | -0.03854 | 0.034059 | 1 |
| Rdh14 | -0.03863 | 0.288206 | 1 |
| Actr10 | -0.03864 | 0.892259 | 1 |
| Zfp738 | -0.03865 | 0.834395 | 1 |
| Ttc9c | -0.03866 | 0.718417 | 1 |
| Pum1 | -0.03868 | 0.971831 | 1 |
| Zfp592 | -0.03869 | 0.625038 | 1 |
| Srsf1 | -0.03869 | 0.251018 | 1 |
| Lrrc8a | -0.0387 | 0.508591 | 1 |
| Ercc6l2 | -0.03875 | 0.75152 | 1 |
| Cox11 | -0.03882 | 0.393034 | 1 |
| Snhg20 | -0.03885 | 0.141125 | 1 |
| Zfp512 | -0.03885 | 0.728855 | 1 |
| Nlrx1 | -0.03889 | 0.780999 | 1 |
| Nr2c1 | -0.03892 | 0.276584 | 1 |
| Tlr4 | -0.03892 | 0.210167 | 1 |
| Alas1 | -0.03892 | 0.846595 | 1 |
| Zfp692 | -0.03902 | 0.706895 | 1 |
| Lcor | -0.03906 | 0.228654 | 1 |
| Leng1 | -0.03907 | 0.444258 | 1 |
| Exoc2 | -0.03911 | 0.573994 | 1 |
| Smyd5 | -0.03919 | 0.624287 | 1 |
| Gm9725 | -0.03921 | 0.917648 | 1 |
| Abcd4 | -0.03925 | 0.602 | 1 |
| Cspp1 | -0.03927 | 0.073746 | 1 |
| Usp45 | -0.03928 | 0.775898 | 1 |
| Tab1 | -0.03931 | 0.484726 | 1 |
| Sar1b | -0.03932 | 0.88767 | 1 |
| Dcaf1 | -0.03942 | 0.775956 | 1 |
| Pds5a | -0.03946 | 0.490105 | 1 |
| Rnf25 | -0.03946 | 0.345939 | 1 |
| Tmem14c | -0.03948 | 0.729208 | 1 |
| Wipi2 | -0.03953 | 0.733086 | 1 |
| Sptlc1 | -0.03962 | 0.795242 | 1 |
| Nfkbie | -0.03964 | 0.303972 | 1 |
| Tbl1x | -0.03965 | 0.828922 | 1 |
| Mrps18c | -0.03965 | 0.199777 | 1 |
| Zfp839 | -0.03967 | 0.877009 | 1 |
| 3110040N: | -0.03967 | 0.943035 | 1 |

| | | | |
|---|---|---|---|
| Zranb1 | -0.03968 | 0.668382 | 1 |
| Cstf2 | -0.03969 | 0.357702 | 1 |
| Nupr1 | -0.0397 | 0.307487 | 1 |
| Rps6kc1 | -0.03971 | 0.058147 | 1 |
| Tmem131l | -0.03971 | 0.310926 | 1 |
| Terf1 | -0.03975 | 0.481402 | 1 |
| Uhmk1 | -0.03985 | 0.702064 | 1 |
| Eri3 | -0.03991 | 0.072235 | 1 |
| Ostm1 | -0.03997 | 0.31646 | 1 |
| Desi1 | -0.03998 | 0.858798 | 1 |
| Rapgef6 | -0.04002 | 0.255194 | 1 |
| Fbxl6 | -0.04011 | 0.90001 | 1 |
| Mrtfa | -0.04016 | 0.846619 | 1 |
| Chmp7 | -0.04017 | 0.074417 | 1 |
| Nubpl | -0.04017 | 0.842547 | 1 |
| Serpinb6a | -0.04018 | 0.877562 | 1 |
| Cep83os | -0.04025 | 0.943041 | 1 |
| Pfdn6 | -0.04026 | 0.173821 | 1 |
| St3gal3 | -0.04028 | 0.165282 | 1 |
| Eif2d | -0.04029 | 0.889345 | 1 |
| Scnm1 | -0.04032 | 0.502443 | 1 |
| Lcp1 | -0.04043 | 0.092585 | 1 |
| Ap4b1 | -0.04044 | 0.366287 | 1 |
| Cops3 | -0.04045 | 0.873132 | 1 |
| Nup133 | -0.04051 | 0.805137 | 1 |
| Dpp9 | -0.04053 | 0.463564 | 1 |
| Gm34280 | -0.04055 | 0.327842 | 1 |
| Mtx2 | -0.04057 | 0.653679 | 1 |
| Ube2q1 | -0.04063 | 0.555097 | 1 |
| Acaa2 | -0.04064 | 0.785932 | 1 |
| Wdr92 | -0.0407 | 0.290648 | 1 |
| Desi2 | -0.0407 | 0.210799 | 1 |
| Cops6 | -0.04071 | 0.437186 | 1 |
| Gpd1l | -0.04077 | 0.291912 | 1 |
| Mdh1 | -0.04079 | 0.527487 | 1 |
| Lrrc3 | -0.0408 | 0.621493 | 1 |
| Shld1 | -0.04081 | 0.975551 | 1 |
| 1600010M | -0.04082 | 0.658563 | 1 |
| Lcorl | -0.04085 | 0.318014 | 1 |
| Ankrd40 | -0.04088 | 0.890688 | 1 |
| Sh2b1 | -0.04092 | 0.515841 | 1 |
| Zfp944 | -0.04096 | 0.574169 | 1 |
| Zyx | -0.04099 | 0.196868 | 1 |
| Hadhb | -0.04102 | 0.421085 | 1 |
| Tpm1 | -0.04111 | 0.182042 | 1 |
| Timm44 | -0.04125 | 0.121233 | 1 |
| Zzz3 | -0.04125 | 0.831561 | 1 |
| Nt5dc1 | -0.04127 | 0.772111 | 1 |

| | | | |
|---|---|---|---|
| Golim4 | -0.04131 | 0.23306 | 1 |
| Xpo4 | -0.04138 | 0.860698 | 1 |
| Trp53i13 | -0.04149 | 0.406592 | 1 |
| Bmpr1a | -0.0415 | 0.156813 | 1 |
| Smim11 | -0.0415 | 0.719643 | 1 |
| Jagn1 | -0.04151 | 0.300043 | 1 |
| 5730455P1 | -0.04152 | 0.860681 | 1 |
| 5530601H0 | -0.04153 | 0.656537 | 1 |
| Ubald2 | -0.04155 | 0.951047 | 1 |
| Med25 | -0.04155 | 0.369704 | 1 |
| Ppp4r3b | -0.0416 | 0.983813 | 1 |
| Tpgs1 | -0.04169 | 0.709573 | 1 |
| Zfp639 | -0.04169 | 0.963973 | 1 |
| Btbd10 | -0.04172 | 0.728527 | 1 |
| Tifa | -0.04173 | 0.577052 | 1 |
| Trabd | -0.04179 | 0.586776 | 1 |
| Arhgap1 | -0.04181 | 0.843574 | 1 |
| Uqcrc1 | -0.04186 | 0.860675 | 1 |
| Thoc7 | -0.04188 | 0.703354 | 1 |
| Map2k3 | -0.04191 | 0.392252 | 1 |
| Vcpip1 | -0.04195 | 0.848032 | 1 |
| Rmnd1 | -0.04204 | 0.194052 | 1 |
| Ppil4 | -0.04209 | 0.51981 | 1 |
| Ypel5 | -0.04212 | 0.760578 | 1 |
| Apopt1 | -0.04214 | 0.949744 | 1 |
| Plcb2 | -0.0422 | 0.622306 | 1 |
| Pck2 | -0.04222 | 0.493071 | 1 |
| 1600020E0 | -0.04227 | 0.070257 | 1 |
| Pigg | -0.04229 | 0.750066 | 1 |
| Mrpl39 | -0.04238 | 0.409604 | 1 |
| Dld | -0.04241 | 0.735521 | 1 |
| Mbip | -0.04242 | 0.836991 | 1 |
| Nrp2 | -0.04244 | 0.550006 | 1 |
| Marcksl1 | -0.04245 | 0.865128 | 1 |
| Ltn1 | -0.04249 | 0.829119 | 1 |
| 1110059E2 | -0.04263 | 0.410136 | 1 |
| Alox5 | -0.04285 | 0.3893 | 1 |
| Gcc2 | -0.04289 | 0.107174 | 1 |
| Sgpp1 | -0.04294 | 0.497573 | 1 |
| Dynlt3 | -0.04296 | 0.371953 | 1 |
| Atpsckmt | -0.04309 | 0.981641 | 1 |
| Lemd3 | -0.04313 | 0.482098 | 1 |
| Tdrd3 | -0.04314 | 0.729463 | 1 |
| Crtc3 | -0.04315 | 0.086442 | 1 |
| Mrpl22 | -0.04326 | 0.810261 | 1 |
| Malsu1 | -0.04328 | 0.295977 | 1 |
| Herpud2 | -0.04333 | 0.170157 | 1 |
| Wbp1l | -0.04333 | 0.58256 | 1 |

| | | | |
|---|---|---|---|
| St7l | -0.04334 | 0.209216 | 1 |
| Yipf2 | -0.04337 | 0.345304 | 1 |
| Acad8 | -0.04342 | 0.862083 | 1 |
| Larp7 | -0.04348 | 0.175085 | 1 |
| Cacul1 | -0.0435 | 0.74896 | 1 |
| Med30 | -0.04354 | 0.751089 | 1 |
| Vamp7 | -0.04355 | 0.991857 | 1 |
| Lin52 | -0.04356 | 0.895798 | 1 |
| Znrd1 | -0.04357 | 0.948188 | 1 |
| Invs | -0.04361 | 0.870347 | 1 |
| Mindy3 | -0.04363 | 0.678753 | 1 |
| Vps51 | -0.04372 | 0.808364 | 1 |
| Orc4 | -0.04383 | 0.959011 | 1 |
| Mthfd1 | -0.04384 | 0.891891 | 1 |
| Cetn3 | -0.0439 | 0.829536 | 1 |
| Nsmce3 | -0.04394 | 0.559686 | 1 |
| Ccdc28b | -0.04395 | 0.955855 | 1 |
| Trappc10 | -0.04396 | 0.794958 | 1 |
| Lipo3 | -0.04399 | 0.464983 | 1 |
| Rbbp5 | -0.04407 | 0.874534 | 1 |
| BC052040 | -0.04409 | 0.952875 | 1 |
| Ncbp1 | -0.04417 | 0.927232 | 1 |
| Eya4 | -0.04423 | 0.112628 | 1 |
| Erlec1 | -0.04424 | 0.771109 | 1 |
| Hdgfl2 | -0.04425 | 0.746661 | 1 |
| Ehmt1 | -0.0443 | 0.046982 | 1 |
| Vdac1 | -0.04431 | 0.520494 | 1 |
| Firre | -0.04431 | 0.237837 | 1 |
| Dnajc10 | -0.04436 | 0.754169 | 1 |
| Kdm3a | -0.04442 | 0.856982 | 1 |
| Mnt | -0.04443 | 0.439419 | 1 |
| Psmd10 | -0.04451 | 0.967463 | 1 |
| Specc1l | -0.04453 | 0.061116 | 1 |
| Ncstn | -0.04463 | 0.817696 | 1 |
| Hadh | -0.04464 | 0.775834 | 1 |
| Mprip | -0.04471 | 0.347371 | 1 |
| Ostc | -0.04478 | 0.517674 | 1 |
| 2310033P( | -0.04479 | 0.626042 | 1 |
| Zfp74 | -0.04484 | 0.075877 | 1 |
| Nudt6 | -0.04484 | 0.438931 | 1 |
| Brd1 | -0.04488 | 0.947865 | 1 |
| Bud13 | -0.04488 | 0.975462 | 1 |
| Snx25 | -0.0449 | 0.996533 | 1 |
| Actb | -0.0449 | 0.000187 | 1 |
| Ift172 | -0.04491 | 0.359562 | 1 |
| Atxn10 | -0.04494 | 0.866995 | 1 |
| Gm14325 | -0.04497 | 0.994871 | 1 |
| Pias2 | -0.04498 | 0.066556 | 1 |

| | | | |
|---|---|---|---|
| Adck5 | -0.04505 | 0.633254 | 1 |
| Ube2q2 | -0.04506 | 0.265689 | 1 |
| Atf2 | -0.04506 | 0.887271 | 1 |
| Hmces | -0.04507 | 0.666374 | 1 |
| Tmem241 | -0.04512 | 0.828716 | 1 |
| Piezo1 | -0.04512 | 0.958947 | 1 |
| Stam2 | -0.04522 | 0.448347 | 1 |
| Wrap73 | -0.04523 | 0.098004 | 1 |
| Herc1 | -0.04523 | 0.488334 | 1 |
| Neurl1a | -0.04527 | 0.583627 | 1 |
| Cnp | -0.04529 | 0.572831 | 1 |
| Tnpo3 | -0.04531 | 0.308225 | 1 |
| Exosc3 | -0.04531 | 0.739263 | 1 |
| Ctnnbl1 | -0.04534 | 0.332022 | 1 |
| Hccs | -0.04536 | 0.285663 | 1 |
| 5430405H( | -0.04538 | 0.860713 | 1 |
| Ndufa9 | -0.04539 | 0.267992 | 1 |
| Socs6 | -0.0454 | 0.758571 | 1 |
| Psmg3 | -0.04541 | 0.601394 | 1 |
| Med6 | -0.04542 | 0.194885 | 1 |
| Smim26 | -0.04545 | 0.821169 | 1 |
| Kbtbd4 | -0.04545 | 0.106374 | 1 |
| Arhgap9 | -0.04546 | 0.827888 | 1 |
| Uckl1 | -0.04547 | 0.992895 | 1 |
| Mis12 | -0.04549 | 0.239614 | 1 |
| Pspc1 | -0.04555 | 0.435923 | 1 |
| Snx9 | -0.04557 | 0.166197 | 1 |
| Apex2 | -0.04558 | 0.84692 | 1 |
| Pxdc1 | -0.04565 | 0.902885 | 1 |
| Ccdc124 | -0.0457 | 0.386679 | 1 |
| Tom1 | -0.04576 | 0.712606 | 1 |
| Klc1 | -0.04584 | 0.021313 | 1 |
| Ap2a2 | -0.04589 | 0.3194 | 1 |
| Cc2d1a | -0.04591 | 0.425469 | 1 |
| Rmdn1 | -0.04591 | 0.853974 | 1 |
| Ubxn8 | -0.04594 | 0.874349 | 1 |
| Emd | -0.04595 | 0.236548 | 1 |
| Ino80d | -0.04598 | 0.947335 | 1 |
| Pcnx | -0.04599 | 0.962686 | 1 |
| Cse1l | -0.04599 | 0.441017 | 1 |
| 2810004N: | -0.04602 | 0.55238 | 1 |
| Mcm3ap | -0.04608 | 0.723623 | 1 |
| Wdr37 | -0.04612 | 0.638672 | 1 |
| Arl2 | -0.04613 | 0.607569 | 1 |
| App | -0.04618 | 0.763837 | 1 |
| Sbds | -0.04622 | 0.605796 | 1 |
| Edem3 | -0.04625 | 0.506588 | 1 |
| Clcn6 | -0.04627 | 0.519577 | 1 |

| | | | |
|---|---|---|---|
| Klhl28 | -0.04629 | 0.597613 | 1 |
| Xrcc1 | -0.04631 | 0.662049 | 1 |
| Proser1 | -0.04632 | 0.77251 | 1 |
| Ehmt2 | -0.04634 | 0.649185 | 1 |
| Thada | -0.04634 | 0.641073 | 1 |
| Adh5 | -0.04635 | 0.894773 | 1 |
| Prr13 | -0.04637 | 0.849675 | 1 |
| Dock7 | -0.0465 | 0.198758 | 1 |
| Prmt5 | -0.04661 | 0.204293 | 1 |
| Sepsecs | -0.04663 | 0.496393 | 1 |
| Tcf12 | -0.04666 | 0.698599 | 1 |
| Ezh1 | -0.04668 | 0.915019 | 1 |
| Rnf146 | -0.04671 | 0.553004 | 1 |
| Yeats4 | -0.04672 | 0.415586 | 1 |
| Mpv17l2 | -0.04677 | 0.566708 | 1 |
| Dag1 | -0.04682 | 0.540067 | 1 |
| Klhl20 | -0.04685 | 0.421773 | 1 |
| Dnajc1 | -0.04689 | 0.964806 | 1 |
| Cherp | -0.04694 | 0.20284 | 1 |
| Zbtb45 | -0.04694 | 0.549255 | 1 |
| Ccdc91 | -0.04698 | 0.577508 | 1 |
| Hnrnpl | -0.04699 | 0.190339 | 1 |
| Zyg11b | -0.04707 | 0.394945 | 1 |
| Sike1 | -0.04709 | 0.350722 | 1 |
| Ascc1 | -0.0471 | 0.857969 | 1 |
| Ddx19a | -0.04714 | 0.949014 | 1 |
| Zfp276 | -0.0472 | 0.690909 | 1 |
| Ubn2 | -0.04721 | 0.878143 | 1 |
| Snw1 | -0.04728 | 0.493373 | 1 |
| Dusp12 | -0.04733 | 0.839499 | 1 |
| Zfp619 | -0.04736 | 0.276481 | 1 |
| Pank3 | -0.04737 | 0.773573 | 1 |
| Smg8 | -0.04738 | 0.584975 | 1 |
| Dnajc19 | -0.04741 | 0.181854 | 1 |
| Cwf19l2 | -0.04743 | 0.072762 | 1 |
| Rhbdd2 | -0.04744 | 0.281417 | 1 |
| Ogdh | -0.04745 | 0.995625 | 1 |
| Mrpl58 | -0.04746 | 0.192148 | 1 |
| Ubl7 | -0.04754 | 0.711681 | 1 |
| Arrdc1 | -0.0476 | 0.563855 | 1 |
| Dus1l | -0.04768 | 0.592011 | 1 |
| Clpb | -0.0477 | 0.703115 | 1 |
| Fbxl5 | -0.04777 | 0.533197 | 1 |
| Dap | -0.04777 | 0.711397 | 1 |
| Arfgef2 | -0.04782 | 0.196552 | 1 |
| Nmral1 | -0.04783 | 0.904486 | 1 |
| G6pdx | -0.04788 | 0.807375 | 1 |
| Washc5 | -0.04794 | 0.172015 | 1 |

| | | | |
|---|---|---|---|
| Taf1c | -0.04796 | 0.926453 | 1 |
| Ipo9 | -0.04797 | 0.488209 | 1 |
| Pacs2 | -0.048 | 0.093274 | 1 |
| Zfp445 | -0.04801 | 0.890557 | 1 |
| Nudt22 | -0.04806 | 0.974669 | 1 |
| Ppm1d | -0.04807 | 0.547684 | 1 |
| Skap2 | -0.04808 | 0.36724 | 1 |
| Ppp3cb | -0.04818 | 0.742598 | 1 |
| Rps6ka3 | -0.04818 | 0.336858 | 1 |
| Zmat2 | -0.04822 | 0.445947 | 1 |
| Asb7 | -0.04829 | 0.77071 | 1 |
| Cep250 | -0.04832 | 0.840584 | 1 |
| Hdgf | -0.04835 | 0.519981 | 1 |
| A530088E( | -0.04843 | 0.808926 | 1 |
| Naglu | -0.04848 | 0.776543 | 1 |
| Arpc2 | -0.04848 | 0.110813 | 1 |
| Gabpa | -0.0485 | 0.771971 | 1 |
| Akt2 | -0.04852 | 0.772637 | 1 |
| Apc | -0.04856 | 0.528945 | 1 |
| Adk | -0.04857 | 0.588904 | 1 |
| Tmem214 | -0.04866 | 0.533688 | 1 |
| Arhgap21 | -0.04869 | 0.983884 | 1 |
| Bcap31 | -0.04872 | 0.394945 | 1 |
| Smad7 | -0.04874 | 0.293349 | 1 |
| Ralgapa1 | -0.04876 | 0.70295 | 1 |
| Spen | -0.04877 | 0.967711 | 1 |
| Zfp524 | -0.04878 | 0.891982 | 1 |
| Vrk3 | -0.04878 | 0.859409 | 1 |
| Fbxo9 | -0.04879 | 0.642901 | 1 |
| Srsf9 | -0.04888 | 0.721261 | 1 |
| Atp9b | -0.0489 | 0.561014 | 1 |
| Prdm1 | -0.04891 | 0.205787 | 1 |
| Eif4a2 | -0.04897 | 0.762068 | 1 |
| Elavl4 | -0.04897 | 0.551374 | 1 |
| Sarnp | -0.04898 | 0.841116 | 1 |
| Gm42962 | -0.049 | 0.584463 | 1 |
| Afap1l1 | -0.049 | 0.852984 | 1 |
| Ndufs1 | -0.04909 | 0.715817 | 1 |
| Ap1m1 | -0.0491 | 0.503592 | 1 |
| Sec23ip | -0.04911 | 0.932348 | 1 |
| Leo1 | -0.04911 | 0.948695 | 1 |
| Edc3 | -0.04916 | 0.542717 | 1 |
| Synj2bp | -0.04917 | 0.67235 | 1 |
| Stx5a | -0.0492 | 0.710622 | 1 |
| D8Ertd738 | -0.04921 | 0.636795 | 1 |
| Mtmr4 | -0.04928 | 0.888443 | 1 |
| Ccnt1 | -0.04929 | 0.867261 | 1 |
| Ptprs | -0.0493 | 0.868875 | 1 |

| | | | |
|---|---|---|---|
| Elf1 | -0.04932 | 0.231368 | 1 |
| Arhgap18 | -0.04933 | 0.558515 | 1 |
| Nt5c2 | -0.04933 | 0.781972 | 1 |
| Abhd17b | -0.04936 | 0.71093 | 1 |
| Ppp1r15b | -0.04941 | 0.827927 | 1 |
| Armc1 | -0.04954 | 0.757345 | 1 |
| Smim12 | -0.04954 | 0.042939 | 1 |
| Ppp1r12b | -0.04955 | 0.952791 | 1 |
| Zfp942 | -0.04961 | 0.774082 | 1 |
| Cd9 | -0.04962 | 0.158904 | 1 |
| Ddx31 | -0.04963 | 0.054556 | 1 |
| Gm13562 | -0.04966 | 0.866171 | 1 |
| Gsg1 | -0.04971 | 0.657639 | 1 |
| B3gat3 | -0.04975 | 0.814853 | 1 |
| Ints8 | -0.04978 | 0.779665 | 1 |
| Scyl1 | -0.04979 | 0.664911 | 1 |
| Phtf2 | -0.04981 | 0.831589 | 1 |
| Mgat4b | -0.04983 | 0.98424 | 1 |
| Kcnq1ot1 | -0.04995 | 0.036093 | 1 |
| Fam149b | -0.05006 | 0.822393 | 1 |
| Alg6 | -0.05013 | 0.620792 | 1 |
| Hsd17b10 | -0.05013 | 0.820778 | 1 |
| Cetn2 | -0.05015 | 0.904234 | 1 |
| Nup214 | -0.05027 | 0.748676 | 1 |
| Ssrp1 | -0.0503 | 0.629242 | 1 |
| Atg10 | -0.05032 | 0.552235 | 1 |
| Usp8 | -0.05033 | 0.550724 | 1 |
| Inppl1 | -0.05034 | 0.450268 | 1 |
| 2610020C( | -0.05038 | 0.164318 | 1 |
| Zdhhc24 | -0.05038 | 0.71234 | 1 |
| Zc3h14 | -0.05041 | 0.345668 | 1 |
| Fam76a | -0.05043 | 0.583042 | 1 |
| Ube2b | -0.05044 | 0.521357 | 1 |
| Zscan25 | -0.05056 | 0.155655 | 1 |
| Capn7 | -0.05059 | 0.652803 | 1 |
| Irf9 | -0.05062 | 0.98241 | 1 |
| Nipbl | -0.05067 | 0.834461 | 1 |
| Cir1 | -0.05071 | 0.506659 | 1 |
| Klc4 | -0.05071 | 0.794628 | 1 |
| Gm5617 | -0.05074 | 0.362851 | 1 |
| Smc1a | -0.05077 | 0.249822 | 1 |
| Pef1 | -0.05083 | 0.477109 | 1 |
| Mtmr2 | -0.05087 | 0.76065 | 1 |
| Rheb | -0.05102 | 0.482135 | 1 |
| Uevld | -0.05103 | 0.665475 | 1 |
| Atg13 | -0.05119 | 0.409496 | 1 |
| Cdk5 | -0.05126 | 0.697631 | 1 |
| Prkcd | -0.0513 | 0.983128 | 1 |

| | | | |
|---|---|---|---|
| Lnpk | -0.05132 | 0.615981 | 1 |
| 2410022M | -0.05139 | 0.270212 | 1 |
| Ntan1 | -0.0514 | 0.537054 | 1 |
| Lyn | -0.05144 | 0.492377 | 1 |
| Zfp511 | -0.05153 | 0.865272 | 1 |
| Ndufaf3 | -0.05155 | 0.420994 | 1 |
| Impact | -0.0517 | 0.040844 | 1 |
| Cyth3 | -0.05172 | 0.256325 | 1 |
| Nkap | -0.05173 | 0.683843 | 1 |
| Tceal9 | -0.05173 | 0.824894 | 1 |
| Lemd2 | -0.05175 | 0.299687 | 1 |
| Gtf3a | -0.05183 | 0.392497 | 1 |
| Rab11fip5 | -0.05187 | 0.903195 | 1 |
| Ppp1r10 | -0.05189 | 0.851328 | 1 |
| D330023K | -0.05192 | 0.231909 | 1 |
| Dmac2 | -0.05193 | 0.290267 | 1 |
| Cfl2 | -0.05194 | 0.910701 | 1 |
| Alkbh3 | -0.05199 | 0.909713 | 1 |
| Ttpal | -0.05203 | 0.759324 | 1 |
| Nhlrc3 | -0.05205 | 0.873555 | 1 |
| Zzef1 | -0.05207 | 0.623096 | 1 |
| Mcee | -0.05207 | 0.964312 | 1 |
| Pex19 | -0.05208 | 0.783382 | 1 |
| Pcgf1 | -0.05208 | 0.782003 | 1 |
| Rcan3 | -0.0521 | 0.647257 | 1 |
| Rspry1 | -0.05211 | 0.616082 | 1 |
| Slc7a6os | -0.05217 | 0.670197 | 1 |
| Gpbp1 | -0.05219 | 0.712907 | 1 |
| Nek9 | -0.05225 | 0.882568 | 1 |
| Zdhhc6 | -0.05225 | 0.995652 | 1 |
| Cnbp | -0.05225 | 0.177198 | 1 |
| Ccs | -0.05225 | 0.338593 | 1 |
| Dpagt1 | -0.05231 | 0.960234 | 1 |
| Wasl | -0.05233 | 0.718856 | 1 |
| Akr1e1 | -0.05234 | 0.314534 | 1 |
| Irf5 | -0.05238 | 0.199956 | 1 |
| Rbm15 | -0.0524 | 0.620988 | 1 |
| Ice2 | -0.05247 | 0.791258 | 1 |
| Phf8 | -0.05254 | 0.944609 | 1 |
| Slc17a9 | -0.05255 | 0.207369 | 1 |
| Gm49336 | -0.05268 | 0.528815 | 1 |
| Actg1 | -0.05275 | 0.023683 | 1 |
| Myo6 | -0.05286 | 0.682115 | 1 |
| Copb2 | -0.0529 | 0.699695 | 1 |
| Chst11 | -0.05291 | 0.002865 | 1 |
| Sfxn3 | -0.05293 | 0.479561 | 1 |
| Frs2 | -0.05297 | 0.687955 | 1 |
| Lrif1 | -0.05298 | 0.890766 | 1 |

| | | | |
|---|---|---|---|
| Ttc32 | -0.0531 | 0.506968 | 1 |
| Slc30a9 | -0.0531 | 0.757025 | 1 |
| Prpsap1 | -0.05313 | 0.235028 | 1 |
| Mynn | -0.05321 | 0.20459 | 1 |
| Hsdl1 | -0.05324 | 0.302303 | 1 |
| Dnajc8 | -0.05332 | 0.665212 | 1 |
| Orc5 | -0.05333 | 0.550856 | 1 |
| Ormdl3 | -0.05333 | 0.196885 | 1 |
| Crcp | -0.05338 | 0.87369 | 1 |
| Acat1 | -0.05341 | 0.656364 | 1 |
| Shkbp1 | -0.0535 | 0.611329 | 1 |
| Ttc8 | -0.05351 | 0.276727 | 1 |
| Strada | -0.05353 | 0.860771 | 1 |
| Ccar2 | -0.05357 | 0.533187 | 1 |
| Arhgap35 | -0.05365 | 0.863798 | 1 |
| Ttc4 | -0.05366 | 0.684459 | 1 |
| Trim28 | -0.0537 | 0.929188 | 1 |
| Cdc27 | -0.05373 | 0.736822 | 1 |
| Hnrnpa2b1 | -0.05375 | 0.410665 | 1 |
| Rnf2 | -0.05389 | 0.108258 | 1 |
| Zscan21 | -0.0539 | 0.224933 | 1 |
| Mccc2 | -0.05391 | 0.726144 | 1 |
| Mtg2 | -0.05392 | 0.678876 | 1 |
| Aqr | -0.05393 | 0.813602 | 1 |
| Sumf2 | -0.05397 | 0.917599 | 1 |
| Cltb | -0.05398 | 0.289673 | 1 |
| Polr3f | -0.05399 | 0.156041 | 1 |
| Tirap | -0.05405 | 0.167932 | 1 |
| Vdac3 | -0.05406 | 0.778584 | 1 |
| Sec61a1 | -0.05409 | 0.497753 | 1 |
| Galk2 | -0.05416 | 0.83561 | 1 |
| Tmem229b | -0.05417 | 0.658633 | 1 |
| Pitpnb | -0.05419 | 0.845461 | 1 |
| Abhd2 | -0.0542 | 0.393387 | 1 |
| Smad2 | -0.05431 | 0.823006 | 1 |
| Rpp25l | -0.05436 | 0.805605 | 1 |
| Tbrg1 | -0.05437 | 0.194779 | 1 |
| Vps52 | -0.05438 | 0.637195 | 1 |
| Aco1 | -0.05445 | 0.781314 | 1 |
| Atp13a3 | -0.05449 | 0.974433 | 1 |
| Elavl1 | -0.05452 | 0.713382 | 1 |
| Rmi1 | -0.05453 | 0.471575 | 1 |
| Spag9 | -0.05453 | 0.933618 | 1 |
| Pex3 | -0.05455 | 0.96389 | 1 |
| Lrrc41 | -0.05456 | 0.578384 | 1 |
| Uap1l1 | -0.05459 | 0.938421 | 1 |
| Suz12 | -0.05459 | 0.566997 | 1 |
| Gpr146 | -0.0546 | 0.91243 | 1 |

| | | | |
|---|---|---|---|
| 2700062C( | -0.05461 | 0.993696 | 1 |
| Camsap2 | -0.05466 | 0.423623 | 1 |
| Cnih4 | -0.05469 | 0.864202 | 1 |
| Top3a | -0.05473 | 0.180243 | 1 |
| Rogdi | -0.05476 | 0.487011 | 1 |
| Rnf38 | -0.05477 | 0.255129 | 1 |
| Chp1 | -0.05477 | 0.568554 | 1 |
| Gstz1 | -0.0548 | 0.480286 | 1 |
| St3gal4 | -0.05483 | 0.66004 | 1 |
| Lrrc58 | -0.05486 | 0.778767 | 1 |
| Fam192a | -0.05486 | 0.624463 | 1 |
| Ckap5 | -0.05491 | 0.601132 | 1 |
| Txnrd2 | -0.05496 | 0.57783 | 1 |
| Cpsf7 | -0.05498 | 0.452507 | 1 |
| Slc39a9 | -0.05499 | 0.469012 | 1 |
| Higd2a | -0.05503 | 0.653977 | 1 |
| Abcb7 | -0.05505 | 0.939013 | 1 |
| Cdkn1a | -0.05506 | 0.732592 | 1 |
| Fbxw4 | -0.05509 | 0.928597 | 1 |
| Ttc5 | -0.0551 | 0.818763 | 1 |
| Rnf14 | -0.05512 | 0.996804 | 1 |
| Gm20528 | -0.05512 | 0.373622 | 1 |
| Gatd1 | -0.05512 | 0.86517 | 1 |
| Kdm5d | -0.05513 | 0.531718 | 1 |
| Hdac11 | -0.05517 | 0.19588 | 1 |
| Ndufaf8 | -0.05519 | 0.32501 | 1 |
| Calu | -0.05524 | 0.839674 | 1 |
| Pigm | -0.05525 | 0.82121 | 1 |
| Ncaph2 | -0.05527 | 0.919604 | 1 |
| Hcfc1 | -0.05544 | 0.386533 | 1 |
| Usp37 | -0.05548 | 0.670963 | 1 |
| Mmadhc | -0.05555 | 0.830449 | 1 |
| Stk4 | -0.05555 | 0.069835 | 1 |
| Zfp322a | -0.05557 | 0.924208 | 1 |
| Hars2 | -0.05559 | 0.133588 | 1 |
| Fam126b | -0.05566 | 0.318573 | 1 |
| Mfap1a | -0.0557 | 0.128257 | 1 |
| Zfp868 | -0.05572 | 0.853723 | 1 |
| Homer3 | -0.05573 | 0.426421 | 1 |
| Mettl15 | -0.05576 | 0.668195 | 1 |
| Ap4s1 | -0.05577 | 0.558579 | 1 |
| Eif2s3x | -0.05583 | 0.911988 | 1 |
| Wdr13 | -0.05584 | 0.874398 | 1 |
| Trappc6b | -0.05585 | 0.211362 | 1 |
| Fggy | -0.05587 | 0.105176 | 1 |
| Ube3b | -0.05589 | 0.396617 | 1 |
| Chid1 | -0.05601 | 0.793759 | 1 |
| Zfp438 | -0.05604 | 0.701282 | 1 |

| | | | |
|---|---|---|---|
| Urod | -0.05606 | 0.300846 | 1 |
| 9930111J2 | -0.05611 | 0.157063 | 1 |
| Ik | -0.05612 | 0.488004 | 1 |
| Phka1 | -0.05614 | 0.369261 | 1 |
| Itm2b | -0.05615 | 0.000205 | 1 |
| Pde12 | -0.05618 | 0.671966 | 1 |
| Dolpp1 | -0.05622 | 0.434977 | 1 |
| Ugdh | -0.05626 | 0.570337 | 1 |
| Dhx30 | -0.05628 | 0.832017 | 1 |
| Bid | -0.05629 | 0.780357 | 1 |
| Rgmb | -0.05632 | 0.253353 | 1 |
| Klhl5 | -0.05641 | 0.347127 | 1 |
| Plcg1 | -0.05642 | 0.39466 | 1 |
| Rbm14 | -0.05642 | 0.91746 | 1 |
| Fbxl12 | -0.05643 | 0.794131 | 1 |
| Eid1 | -0.05645 | 0.596659 | 1 |
| Gtf3c1 | -0.05655 | 0.837128 | 1 |
| Trp53bp2 | -0.05666 | 0.710632 | 1 |
| Tbc1d20 | -0.05667 | 0.523373 | 1 |
| Rnf103 | -0.05678 | 0.30223 | 1 |
| Ghdc | -0.05679 | 0.260669 | 1 |
| Apex1 | -0.05681 | 0.823438 | 1 |
| Mtmr10 | -0.0569 | 0.836353 | 1 |
| Zfp280d | -0.05691 | 0.430628 | 1 |
| Ninj1 | -0.05691 | 0.511312 | 1 |
| Map2k4 | -0.05694 | 0.609047 | 1 |
| E130307A1 | -0.05694 | 0.72706 | 1 |
| Rpp14 | -0.05695 | 0.622327 | 1 |
| Tmc6 | -0.05697 | 0.721121 | 1 |
| Rexo1 | -0.05698 | 0.554491 | 1 |
| Adck1 | -0.05706 | 0.606468 | 1 |
| Arid2 | -0.05707 | 0.463129 | 1 |
| Pmpcb | -0.05709 | 0.672005 | 1 |
| Emc2 | -0.05712 | 0.231736 | 1 |
| Kif1bp | -0.05713 | 0.262175 | 1 |
| Tmem199 | -0.05722 | 0.749661 | 1 |
| Ttc17 | -0.05725 | 0.619445 | 1 |
| Snrk | -0.05725 | 0.369243 | 1 |
| Golga7 | -0.0573 | 0.700143 | 1 |
| C1d | -0.0573 | 0.659782 | 1 |
| Pde4dip | -0.05732 | 0.177804 | 1 |
| Tnrc6a | -0.05736 | 0.669113 | 1 |
| Tom1l2 | -0.0574 | 0.308125 | 1 |
| Papolg | -0.0575 | 0.442103 | 1 |
| Med16 | -0.05752 | 0.957489 | 1 |
| Rtl8c | -0.05759 | 0.167798 | 1 |
| Dym | -0.05761 | 0.785558 | 1 |
| Strap | -0.05767 | 0.112555 | 1 |

| | | | |
|---|---|---|---|
| Rnf187 | -0.05768 | 0.306791 | 1 |
| Echdc1 | -0.05769 | 0.450806 | 1 |
| Pcm1 | -0.05769 | 0.28485 | 1 |
| Kctd12 | -0.0578 | 0.006931 | 1 |
| Fktn | -0.05781 | 0.832988 | 1 |
| Tnfaip1 | -0.05793 | 0.481359 | 1 |
| Catspere2 | -0.05796 | 0.918633 | 1 |
| Gxylt1 | -0.05796 | 0.803468 | 1 |
| Trappc3 | -0.05796 | 0.931946 | 1 |
| Gm26782 | -0.05797 | 0.642114 | 1 |
| Gtf3c3 | -0.05801 | 0.928069 | 1 |
| Triap1 | -0.05802 | 0.408745 | 1 |
| Vps9d1 | -0.05803 | 0.711268 | 1 |
| Pip5k1a | -0.05811 | 0.952494 | 1 |
| Rrp36 | -0.05814 | 0.541729 | 1 |
| Zfp951 | -0.05814 | 0.991525 | 1 |
| Snrnp40 | -0.05814 | 0.47804 | 1 |
| Sephs1 | -0.05823 | 0.950914 | 1 |
| Psmc3 | -0.05837 | 0.470542 | 1 |
| Ube2h | -0.05847 | 0.371959 | 1 |
| Mark3 | -0.05847 | 0.273994 | 1 |
| Ndufb3 | -0.05847 | 0.918112 | 1 |
| Ube2j1 | -0.05847 | 0.349364 | 1 |
| Cpped1 | -0.05854 | 0.479664 | 1 |
| Maz | -0.05861 | 0.738113 | 1 |
| Map3k4 | -0.05866 | 0.676663 | 1 |
| Clec16a | -0.05868 | 0.066582 | 1 |
| Aen | -0.05874 | 0.721532 | 1 |
| Pet100 | -0.05881 | 0.467428 | 1 |
| Twistnb | -0.05885 | 0.949611 | 1 |
| Zfr | -0.05897 | 0.691401 | 1 |
| Thumpd2 | -0.05898 | 0.663692 | 1 |
| C130050O1 | -0.05899 | 0.353657 | 1 |
| Tef | -0.05904 | 0.860468 | 1 |
| Cep70 | -0.05905 | 0.420324 | 1 |
| Mark4 | -0.05915 | 0.561136 | 1 |
| Hnrnpa0 | -0.05924 | 0.531047 | 1 |
| Prpf3 | -0.05924 | 0.542891 | 1 |
| Myg1 | -0.05925 | 0.708849 | 1 |
| Tfpt | -0.05926 | 0.944224 | 1 |
| Plekho1 | -0.05931 | 0.976227 | 1 |
| Tcaim | -0.05933 | 0.228344 | 1 |
| Acad9 | -0.05933 | 0.295308 | 1 |
| Entpd7 | -0.05937 | 0.788772 | 1 |
| Lamp2 | -0.05939 | 0.241303 | 1 |
| 9530068E0 | -0.0594 | 0.429366 | 1 |
| Drosha | -0.05941 | 0.96423 | 1 |
| Bub3 | -0.05943 | 0.177702 | 1 |

| | | | |
|---|---|---|---|
| Fbxo3 | -0.05943 | 0.066297 | 1 |
| Ppp4c | -0.05945 | 0.935416 | 1 |
| Mettl21a | -0.05949 | 0.495797 | 1 |
| Usp32 | -0.05954 | 0.802223 | 1 |
| Mlf2 | -0.05958 | 0.774902 | 1 |
| Pisd | -0.05961 | 0.575779 | 1 |
| Smarcal1 | -0.05963 | 0.969925 | 1 |
| Spin1 | -0.05968 | 0.407699 | 1 |
| Uso1 | -0.05971 | 0.857848 | 1 |
| Slc17a5 | -0.05974 | 0.737101 | 1 |
| Pex14 | -0.05975 | 0.802435 | 1 |
| Nedd1 | -0.05986 | 0.618223 | 1 |
| Cdc42se1 | -0.05991 | 0.168694 | 1 |
| Lipe | -0.05993 | 0.791145 | 1 |
| Tor2a | -0.05995 | 0.883872 | 1 |
| Atg5 | -0.05996 | 0.593955 | 1 |
| Actr1b | -0.06005 | 0.715548 | 1 |
| Nbas | -0.06008 | 0.209126 | 1 |
| Pigb | -0.06017 | 0.756157 | 1 |
| Nr2c2 | -0.06019 | 0.557071 | 1 |
| Med20 | -0.06021 | 0.476674 | 1 |
| Tyrobp | -0.06022 | 0.033198 | 1 |
| Zfp799 | -0.06023 | 0.250579 | 1 |
| Ube2v2 | -0.06023 | 0.939348 | 1 |
| Psmd5 | -0.06024 | 0.640889 | 1 |
| Stub1 | -0.06024 | 0.515071 | 1 |
| Acbd3 | -0.06034 | 0.546585 | 1 |
| Armt1 | -0.06035 | 0.035218 | 1 |
| Axin1 | -0.06036 | 0.899596 | 1 |
| Dcaf4 | -0.06036 | 0.200614 | 1 |
| Kdm5c | -0.06046 | 0.707398 | 1 |
| Vamp5 | -0.06047 | 0.677693 | 1 |
| Ankmy2 | -0.06052 | 0.191989 | 1 |
| Gkap1 | -0.06053 | 0.854557 | 1 |
| Snapc3 | -0.06054 | 0.819547 | 1 |
| Sgta | -0.06055 | 0.422824 | 1 |
| Magoh | -0.06058 | 0.654653 | 1 |
| Uba1 | -0.06062 | 0.990039 | 1 |
| Cpox | -0.06067 | 0.588626 | 1 |
| Mphosph9 | -0.06069 | 0.112279 | 1 |
| Arnt | -0.06071 | 0.342397 | 1 |
| Zfpl1 | -0.06077 | 0.929119 | 1 |
| Dhx35 | -0.0608 | 0.123766 | 1 |
| Rfc1 | -0.06093 | 0.290236 | 1 |
| Eef1akmt2 | -0.06095 | 0.349044 | 1 |
| Tsg101 | -0.06097 | 0.897886 | 1 |
| Evi2 | -0.06105 | 0.039712 | 1 |
| Aar2 | -0.06111 | 0.163716 | 1 |

| | | | |
|---|---|---|---|
| Thoc3 | -0.06116 | 0.115238 | 1 |
| Arl11 | -0.06121 | 0.977701 | 1 |
| Ubald1 | -0.06121 | 0.172584 | 1 |
| Ell | -0.06126 | 0.349619 | 1 |
| Snrnp27 | -0.06127 | 0.615563 | 1 |
| Ankzf1 | -0.06129 | 0.087679 | 1 |
| Slc38a2 | -0.06131 | 0.2888 | 1 |
| Sh3tc1 | -0.06138 | 0.18728 | 1 |
| Arl14ep | -0.06139 | 0.55709 | 1 |
| Zbtb2 | -0.06139 | 0.262456 | 1 |
| Adap2os | -0.06141 | 0.618419 | 1 |
| Tmem9 | -0.06141 | 0.741725 | 1 |
| Golga1 | -0.06141 | 0.97001 | 1 |
| Tm9sf4 | -0.06153 | 0.623519 | 1 |
| Mks1 | -0.06163 | 0.463202 | 1 |
| Sigmar1 | -0.06177 | 0.137815 | 1 |
| Nudt18 | -0.06184 | 0.075059 | 1 |
| Scarb1 | -0.0619 | 0.347389 | 1 |
| Klhl6 | -0.06191 | 0.823543 | 1 |
| B430010I2 | -0.06195 | 0.53245 | 1 |
| Strbp | -0.06197 | 0.875939 | 1 |
| Ccdc127 | -0.06198 | 0.633799 | 1 |
| Mrps33 | -0.06206 | 0.809568 | 1 |
| Elac1 | -0.06209 | 0.215949 | 1 |
| Ago3 | -0.06212 | 0.392242 | 1 |
| Kctd13 | -0.06214 | 0.475197 | 1 |
| Nfatc3 | -0.06214 | 0.273388 | 1 |
| Dhx15 | -0.06215 | 0.620432 | 1 |
| Tmem65 | -0.06218 | 0.801055 | 1 |
| Yme1l1 | -0.06221 | 0.590067 | 1 |
| Blvra | -0.06224 | 0.253387 | 1 |
| Pibf1 | -0.06225 | 0.627533 | 1 |
| Tcf3 | -0.06225 | 0.813279 | 1 |
| Stambp | -0.06227 | 0.838421 | 1 |
| Rbmxl1 | -0.0623 | 0.524211 | 1 |
| BC031181 | -0.06235 | 0.845496 | 1 |
| Sbno1 | -0.06256 | 0.79919 | 1 |
| Ifnar2 | -0.06263 | 0.417268 | 1 |
| Ccdc62 | -0.06267 | 0.591355 | 1 |
| Klhl22 | -0.06269 | 0.036769 | 1 |
| Pcid2 | -0.0627 | 0.634294 | 1 |
| Cbll1 | -0.0627 | 0.898553 | 1 |
| Ciao1 | -0.06271 | 0.34014 | 1 |
| Fndc3b | -0.06275 | 0.016683 | 1 |
| Hyal2 | -0.06279 | 0.244527 | 1 |
| Hnrnpc | -0.06279 | 0.802992 | 1 |
| Slc4a2 | -0.06279 | 0.592669 | 1 |
| Luc7l3 | -0.06282 | 0.267733 | 1 |

| | | | |
|---|---|---|---|
| Utp11 | -0.06289 | 0.70252 | 1 |
| Polr2e | -0.06291 | 0.846761 | 1 |
| Tnfrsf11a | -0.06293 | 0.223978 | 1 |
| Lancl1 | -0.06294 | 0.981421 | 1 |
| Adssl1 | -0.06296 | 0.687398 | 1 |
| Smcr8 | -0.06304 | 0.475097 | 1 |
| Gga1 | -0.06304 | 0.903074 | 1 |
| Ccdc85b | -0.06305 | 0.306456 | 1 |
| Cdkl3 | -0.06311 | 0.915629 | 1 |
| Dad1 | -0.06313 | 0.618358 | 1 |
| Chmp5 | -0.06317 | 0.484524 | 1 |
| Flii | -0.06331 | 0.424384 | 1 |
| Sf3a2 | -0.06336 | 0.581242 | 1 |
| Pdik1l | -0.06337 | 0.497318 | 1 |
| Gdi2 | -0.06343 | 0.033815 | 1 |
| Cog6 | -0.06348 | 0.022451 | 1 |
| 1110059G | -0.06357 | 0.50765 | 1 |
| Lypla1 | -0.06361 | 0.191965 | 1 |
| Asb8 | -0.06364 | 0.544479 | 1 |
| Cops7b | -0.06366 | 0.045318 | 1 |
| Letm1 | -0.0637 | 0.91906 | 1 |
| Emc1 | -0.0637 | 0.916923 | 1 |
| Nelfa | -0.06373 | 0.88619 | 1 |
| Rap2a | -0.06378 | 0.375909 | 1 |
| Prxl2b | -0.06381 | 0.900916 | 1 |
| Abcb1b | -0.06386 | 0.279919 | 1 |
| Ubqln1 | -0.06391 | 0.460439 | 1 |
| Rab9 | -0.06397 | 0.740737 | 1 |
| Slc16a10 | -0.06408 | 0.693327 | 1 |
| Rnf141 | -0.06409 | 0.922211 | 1 |
| Sord | -0.0641 | 0.286796 | 1 |
| Tbc1d9 | -0.06413 | 0.59329 | 1 |
| Chrac1 | -0.06418 | 0.252125 | 1 |
| Fam193b | -0.06421 | 0.022927 | 1 |
| Rpap2 | -0.06427 | 0.201094 | 1 |
| Ptpn4 | -0.06431 | 0.565073 | 1 |
| Med31 | -0.06435 | 0.079243 | 1 |
| Rab11a | -0.06438 | 0.84686 | 1 |
| Pbx3 | -0.06439 | 0.683573 | 1 |
| Tmem258 | -0.06442 | 0.477759 | 1 |
| Rprd1a | -0.06452 | 0.994601 | 1 |
| Drg1 | -0.06453 | 0.325636 | 1 |
| Ctps2 | -0.06453 | 0.283949 | 1 |
| Mtpn | -0.06455 | 0.439935 | 1 |
| 2610507I0 | -0.06458 | 0.236936 | 1 |
| Mgme1 | -0.06459 | 0.174721 | 1 |
| Fut11 | -0.06459 | 0.728582 | 1 |
| Mau2 | -0.06461 | 0.943288 | 1 |

| | | | |
|---|---|---|---|
| Sacm1l | -0.06464 | 0.983499 | 1 |
| Tecr | -0.06467 | 0.299084 | 1 |
| Ankrd46 | -0.0647 | 0.072616 | 1 |
| Rnf113a2 | -0.06471 | 0.140373 | 1 |
| Dab2 | -0.06471 | 0.647049 | 1 |
| Trappc11 | -0.06472 | 0.819778 | 1 |
| Exosc7 | -0.06483 | 0.859574 | 1 |
| Vps37c | -0.06484 | 0.931456 | 1 |
| Sirt7 | -0.06484 | 0.802497 | 1 |
| Clic1 | -0.0649 | 0.024321 | 1 |
| Trmt11 | -0.065 | 0.948112 | 1 |
| Ostf1 | -0.06506 | 0.394186 | 1 |
| Mapk8 | -0.06509 | 0.970665 | 1 |
| Rbm18 | -0.06512 | 0.80188 | 1 |
| Eloa | -0.06517 | 0.809119 | 1 |
| Atp2a2 | -0.06519 | 0.568394 | 1 |
| Rest | -0.06525 | 0.765305 | 1 |
| Trmt13 | -0.06533 | 0.481404 | 1 |
| Psip1 | -0.06536 | 0.117088 | 1 |
| Gon4l | -0.06539 | 0.937709 | 1 |
| Sdhaf4 | -0.06546 | 0.318172 | 1 |
| Vps13b | -0.06548 | 0.550245 | 1 |
| Dennd4b | -0.06551 | 0.709247 | 1 |
| Rab11fip2 | -0.06552 | 0.676673 | 1 |
| Dnlz | -0.06555 | 0.24152 | 1 |
| Zbtb9 | -0.06556 | 0.381546 | 1 |
| Dynlrb1 | -0.06558 | 0.615274 | 1 |
| Tlr6 | -0.0656 | 0.836817 | 1 |
| Fkbp2 | -0.06566 | 0.875649 | 1 |
| Egln2 | -0.06567 | 0.287469 | 1 |
| Arl15 | -0.06568 | 0.029354 | 1 |
| Lats2 | -0.06572 | 0.174107 | 1 |
| Phf3 | -0.06584 | 0.971489 | 1 |
| Edc4 | -0.06585 | 0.139572 | 1 |
| Mrps11 | -0.06593 | 0.406513 | 1 |
| E4f1 | -0.06598 | 0.993305 | 1 |
| Usp28 | -0.06599 | 0.361676 | 1 |
| Rbbp6 | -0.066 | 0.342436 | 1 |
| Efcab14 | -0.06603 | 0.566485 | 1 |
| Vps8 | -0.06607 | 0.649565 | 1 |
| Slc35c1 | -0.06607 | 0.337859 | 1 |
| Pld2 | -0.06615 | 0.150445 | 1 |
| Dmac1 | -0.06621 | 0.680412 | 1 |
| Dgcr2 | -0.06625 | 0.536544 | 1 |
| Commd10 | -0.06626 | 0.987427 | 1 |
| Zfp994 | -0.0663 | 0.821727 | 1 |
| Xrcc4 | -0.06631 | 0.541829 | 1 |
| Trmt2a | -0.06634 | 0.295883 | 1 |

| | | | |
|---|---|---|---|
| Hagh | -0.06637 | 0.288735 | 1 |
| Afg3l1 | -0.0665 | 0.627661 | 1 |
| Ssna1 | -0.06653 | 0.559832 | 1 |
| AU041133 | -0.0666 | 0.229519 | 1 |
| Aga | -0.06667 | 0.907088 | 1 |
| Gtf2b | -0.0667 | 0.753035 | 1 |
| BC002059 | -0.06671 | 0.096566 | 1 |
| Nup188 | -0.06672 | 0.728596 | 1 |
| Rock1 | -0.06675 | 0.964655 | 1 |
| Vgll4 | -0.06685 | 0.241959 | 1 |
| Derl2 | -0.06692 | 0.704193 | 1 |
| Rbm6 | -0.06692 | 0.955816 | 1 |
| Tada1 | -0.06699 | 0.711869 | 1 |
| Smarcb1 | -0.067 | 0.982957 | 1 |
| Sclt1 | -0.06707 | 0.784157 | 1 |
| Dcaf5 | -0.06714 | 0.628618 | 1 |
| Tm2d1 | -0.06716 | 0.771785 | 1 |
| Slc35f5 | -0.06717 | 0.414603 | 1 |
| Dcp1a | -0.0672 | 0.786171 | 1 |
| Ercc5 | -0.06721 | 0.828861 | 1 |
| Actn4 | -0.06722 | 0.226578 | 1 |
| Ppp2ca | -0.06728 | 0.757789 | 1 |
| Zscan22 | -0.0673 | 0.085631 | 1 |
| Cpeb4 | -0.06741 | 0.565645 | 1 |
| Cramp1l | -0.06742 | 0.950218 | 1 |
| Ppa2 | -0.0675 | 0.304714 | 1 |
| Il21r | -0.06762 | 0.344711 | 1 |
| Gimap1 | -0.06764 | 0.009856 | 1 |
| Ninl | -0.06767 | 0.734362 | 1 |
| Nfkbil1 | -0.06768 | 0.246357 | 1 |
| Pofut1 | -0.06771 | 0.752791 | 1 |
| Srd5a3 | -0.06777 | 0.74216 | 1 |
| Gps1 | -0.06788 | 0.244238 | 1 |
| C330007P( | -0.06794 | 0.447881 | 1 |
| Smad4 | -0.06794 | 0.762093 | 1 |
| Tmem170b | -0.06801 | 0.773353 | 1 |
| Eogt | -0.06807 | 0.020948 | 1 |
| Rab10os | -0.06811 | 0.408043 | 1 |
| Wdr7 | -0.06822 | 0.88364 | 1 |
| Blvrb | -0.06822 | 0.177152 | 1 |
| Actr6 | -0.06832 | 0.097502 | 1 |
| Oser1 | -0.06838 | 0.314982 | 1 |
| Ap3m1 | -0.06848 | 0.988581 | 1 |
| Naa80 | -0.0685 | 0.231778 | 1 |
| Ngly1 | -0.06859 | 0.924476 | 1 |
| Scap | -0.06861 | 0.389369 | 1 |
| Larp4 | -0.06863 | 0.846042 | 1 |
| Fgd4 | -0.06864 | 0.966504 | 1 |

| | | | |
|---|---|---|---|
| Ccdc59 | -0.0687 | 0.736372 | 1 |
| Mesd | -0.0687 | 0.776439 | 1 |
| Edrf1 | -0.06872 | 0.609042 | 1 |
| Ctcf | -0.06879 | 0.858972 | 1 |
| Ccpg1 | -0.06882 | 0.591554 | 1 |
| Capza2 | -0.06885 | 0.052711 | 1 |
| Gpalpp1 | -0.06885 | 0.178897 | 1 |
| Bloc1s1 | -0.06887 | 0.731394 | 1 |
| Usp35 | -0.0689 | 0.07623 | 1 |
| Smarcd2 | -0.06892 | 0.001749 | 1 |
| Snapc5 | -0.06894 | 0.358717 | 1 |
| Rft1 | -0.06895 | 0.924074 | 1 |
| Cdkn2aipnl | -0.06898 | 0.729446 | 1 |
| Hadha | -0.06899 | 0.448511 | 1 |
| Ccndbp1 | -0.06905 | 0.247745 | 1 |
| B3galt4 | -0.06912 | 0.609316 | 1 |
| Slc9a1 | -0.06913 | 0.209803 | 1 |
| Tspan3 | -0.06913 | 0.933221 | 1 |
| Atxn3 | -0.06917 | 0.170691 | 1 |
| Psmd3 | -0.06919 | 0.496748 | 1 |
| Dus3l | -0.06919 | 0.2252 | 1 |
| Pom121 | -0.06923 | 0.269357 | 1 |
| Fam193a | -0.06928 | 0.809106 | 1 |
| Sec16a | -0.06929 | 0.043111 | 1 |
| Msi2 | -0.0694 | 0.591428 | 1 |
| Tm9sf3 | -0.0694 | 0.702146 | 1 |
| Tsnax | -0.06943 | 0.546414 | 1 |
| Prpf18 | -0.06945 | 0.051052 | 1 |
| Mapk1ip1 | -0.06948 | 0.11539 | 1 |
| Med28 | -0.0695 | 0.307839 | 1 |
| Uros | -0.06951 | 0.39177 | 1 |
| Kdm5a | -0.06953 | 0.376619 | 1 |
| Cebpz | -0.06954 | 0.831137 | 1 |
| 2610037D0 | -0.06956 | 0.27372 | 1 |
| Aasdhppt | -0.06961 | 0.875805 | 1 |
| Ptcd3 | -0.06965 | 0.872717 | 1 |
| Sap25 | -0.06967 | 0.727822 | 1 |
| 2210408I2 | -0.06969 | 0.561274 | 1 |
| Rbbp7 | -0.0697 | 0.138744 | 1 |
| Dusp22 | -0.0697 | 0.417241 | 1 |
| H2-Ob | -0.06971 | 0.6012 | 1 |
| Aktip | -0.06974 | 0.124262 | 1 |
| Mfsd14a | -0.06976 | 0.500228 | 1 |
| Rbm41 | -0.06982 | 0.303096 | 1 |
| Coasy | -0.06991 | 0.549988 | 1 |
| Rab35 | -0.06992 | 0.461918 | 1 |
| Cnot7 | -0.07 | 0.616206 | 1 |
| Mbd6 | -0.07005 | 0.021313 | 1 |

| | | | |
|---|---|---|---|
| Ice1 | -0.07008 | 0.819128 | 1 |
| Spast | -0.0701 | 0.652424 | 1 |
| Ccdc191 | -0.0701 | 0.528367 | 1 |
| Tmco3 | -0.07014 | 0.471885 | 1 |
| Tbccd1 | -0.07014 | 0.294361 | 1 |
| Pcbd2 | -0.07015 | 0.170254 | 1 |
| Prmt9 | -0.07017 | 0.154772 | 1 |
| Cfap20 | -0.0702 | 0.02158 | 1 |
| Clcn3 | -0.0702 | 0.231712 | 1 |
| Polr3b | -0.07025 | 0.69279 | 1 |
| Yipf5 | -0.07029 | 0.806984 | 1 |
| AI987944 | -0.0703 | 0.564504 | 1 |
| Dync2h1 | -0.07065 | 0.717839 | 1 |
| Pygo2 | -0.07082 | 0.113634 | 1 |
| Dync1li2 | -0.07087 | 0.06139 | 1 |
| Pih1d1 | -0.0709 | 0.617685 | 1 |
| Mcrip1 | -0.07095 | 0.442674 | 1 |
| Glmp | -0.07096 | 0.521389 | 1 |
| Gimap6 | -0.07097 | 2.18E-05 | 0.704928 |
| Akt1s1 | -0.07098 | 0.252178 | 1 |
| Eif4ebp2 | -0.07103 | 0.167608 | 1 |
| Phf7 | -0.07108 | 0.0395 | 1 |
| Atpaf1 | -0.07121 | 0.909203 | 1 |
| Virma | -0.07124 | 0.687036 | 1 |
| Zfp143 | -0.07128 | 0.821841 | 1 |
| Znrf1 | -0.07132 | 0.124553 | 1 |
| Lgmn | -0.07135 | 5.59E-05 | 1 |
| Fopnl | -0.07139 | 0.563026 | 1 |
| Mttp | -0.07142 | 0.040727 | 1 |
| Pten | -0.07144 | 0.266237 | 1 |
| Map4k5 | -0.07153 | 0.271848 | 1 |
| Sar1a | -0.07153 | 0.117032 | 1 |
| Abcf3 | -0.07154 | 0.35658 | 1 |
| Lyrm2 | -0.07156 | 0.140977 | 1 |
| Fam120c | -0.07156 | 0.33074 | 1 |
| Gtf2a1 | -0.07165 | 0.943337 | 1 |
| Zfp386 | -0.07177 | 0.695322 | 1 |
| Mtmr12 | -0.07177 | 0.059794 | 1 |
| Pip4k2c | -0.07181 | 0.051756 | 1 |
| Dmac2l | -0.07186 | 0.008896 | 1 |
| Oat | -0.07193 | 0.958493 | 1 |
| Gigyf1 | -0.07193 | 0.202664 | 1 |
| Baz2b | -0.07194 | 0.297951 | 1 |
| Arfip1 | -0.072 | 0.092149 | 1 |
| Trappc2 | -0.07202 | 0.022557 | 1 |
| Alg12 | -0.07206 | 0.758134 | 1 |
| Mmut | -0.07208 | 0.962793 | 1 |
| Prdx4 | -0.07208 | 0.83657 | 1 |

| | | | |
|---|---|---|---|
| Ercc1 | -0.07211 | 0.032656 | 1 |
| Rpn2 | -0.07216 | 0.017438 | 1 |
| Fbxo42 | -0.07218 | 0.087723 | 1 |
| Cdyl | -0.07222 | 0.903076 | 1 |
| Prkd3 | -0.07227 | 0.957841 | 1 |
| Pkd2 | -0.0723 | 0.381629 | 1 |
| Fads1 | -0.07233 | 0.288821 | 1 |
| Stip1 | -0.07235 | 0.963493 | 1 |
| BC029722 | -0.0724 | 0.180451 | 1 |
| Sel1l | -0.07241 | 0.329874 | 1 |
| Kdm6a | -0.07247 | 0.605789 | 1 |
| Pdap1 | -0.07247 | 0.357033 | 1 |
| Rnf135 | -0.0725 | 0.805349 | 1 |
| Fmn1 | -0.07251 | 0.410636 | 1 |
| Sec22b | -0.07256 | 0.528413 | 1 |
| Slc10a3 | -0.07262 | 0.222323 | 1 |
| Scaf1 | -0.07275 | 0.343084 | 1 |
| Hibadh | -0.07287 | 0.505465 | 1 |
| Syvn1 | -0.07289 | 0.935034 | 1 |
| Abtb1 | -0.07299 | 0.008634 | 1 |
| Mef2d | -0.07304 | 0.796052 | 1 |
| Krba1 | -0.07308 | 0.040305 | 1 |
| Zfp846 | -0.0731 | 0.102848 | 1 |
| Lmo2 | -0.07318 | 0.834422 | 1 |
| Rnf6 | -0.0732 | 0.848063 | 1 |
| Spsb3 | -0.07327 | 0.693124 | 1 |
| Gm47664 | -0.07328 | 0.145125 | 1 |
| Tcta | -0.07328 | 0.539988 | 1 |
| Mettl23 | -0.07329 | 0.107311 | 1 |
| Mtf2 | -0.07332 | 0.147255 | 1 |
| Grb2 | -0.07336 | 0.018107 | 1 |
| Cog7 | -0.07336 | 0.354769 | 1 |
| Mto1 | -0.07349 | 0.626342 | 1 |
| Ctbs | -0.07363 | 0.663137 | 1 |
| Inpp5k | -0.07364 | 0.643113 | 1 |
| Vamp3 | -0.07367 | 0.363069 | 1 |
| Hbs1l | -0.07368 | 0.477829 | 1 |
| Cers2 | -0.07369 | 0.212851 | 1 |
| Sirt2 | -0.07372 | 0.210122 | 1 |
| Hacd4 | -0.0739 | 0.260022 | 1 |
| Zfp84 | -0.07395 | 0.767621 | 1 |
| Abcc5 | -0.07402 | 0.386289 | 1 |
| Supt4a | -0.07403 | 0.249448 | 1 |
| Stoml2 | -0.07405 | 0.795612 | 1 |
| Zfp24 | -0.07408 | 0.11811 | 1 |
| Fbxl20 | -0.0742 | 0.516442 | 1 |
| Mrps12 | -0.07421 | 0.857373 | 1 |
| Neurl3 | -0.07434 | 0.458081 | 1 |

| | | | |
|---|---|---|---|
| Scaf11 | -0.07435 | 0.153953 | 1 |
| Coq8b | -0.07438 | 0.41395 | 1 |
| Hmbox1 | -0.07451 | 0.651932 | 1 |
| Pcca | -0.07453 | 0.777569 | 1 |
| Wdr33 | -0.07455 | 0.09238 | 1 |
| Crebzf | -0.07457 | 0.191949 | 1 |
| Enoph1 | -0.07459 | 0.176404 | 1 |
| Syt11 | -0.0746 | 0.853988 | 1 |
| Psmc1 | -0.07461 | 0.70807 | 1 |
| Ugp2 | -0.07463 | 0.185947 | 1 |
| Lrrc14 | -0.07466 | 0.285423 | 1 |
| Zbtb24 | -0.07468 | 0.232846 | 1 |
| Trim24 | -0.07468 | 0.429901 | 1 |
| Vamp1 | -0.07476 | 0.083653 | 1 |
| Dexi | -0.07484 | 0.522767 | 1 |
| Iqce | -0.07487 | 0.892794 | 1 |
| Pdgfa | -0.07493 | 0.343345 | 1 |
| Strn | -0.07497 | 0.19656 | 1 |
| Ric8b | -0.07503 | 0.282917 | 1 |
| Ltbr | -0.07507 | 0.156148 | 1 |
| Odr4 | -0.07508 | 0.835914 | 1 |
| Lmo4 | -0.07521 | 0.252013 | 1 |
| Nek1 | -0.07523 | 0.291547 | 1 |
| Ssb | -0.0753 | 0.84784 | 1 |
| 1500004A1 | -0.07534 | 0.712603 | 1 |
| Mterf3 | -0.07537 | 0.105284 | 1 |
| Vash1 | -0.07543 | 0.059372 | 1 |
| Gm5165 | -0.07546 | 0.011384 | 1 |
| BC024978 | -0.07552 | 0.046586 | 1 |
| Ulk3 | -0.07553 | 0.226405 | 1 |
| Cnot11 | -0.07555 | 0.091764 | 1 |
| Zmat5 | -0.07556 | 0.334602 | 1 |
| Pacs1 | -0.07559 | 0.520653 | 1 |
| Acp6 | -0.07561 | 0.104698 | 1 |
| Glt8d1 | -0.07562 | 0.061099 | 1 |
| Rbm22 | -0.07567 | 0.446349 | 1 |
| Gatc | -0.07568 | 0.056081 | 1 |
| Hprt | -0.07579 | 0.344238 | 1 |
| Ciao3 | -0.07584 | 0.658197 | 1 |
| Ccdc61 | -0.07585 | 0.098448 | 1 |
| Hax1 | -0.07586 | 0.723236 | 1 |
| Snrnp70 | -0.07592 | 0.019624 | 1 |
| Bcl10 | -0.07596 | 0.686616 | 1 |
| N4bp2l1 | -0.07596 | 0.314937 | 1 |
| Plcd1 | -0.07603 | 0.456437 | 1 |
| Tnfrsf1a | -0.07607 | 0.163143 | 1 |
| Borcs7 | -0.07608 | 0.686232 | 1 |
| Slc48a1 | -0.07619 | 0.310032 | 1 |

| | | | |
|---|---|---|---|
| Nrf1 | -0.07621 | 0.260501 | 1 |
| Pnrc2 | -0.07624 | 0.151489 | 1 |
| Fam204a | -0.07624 | 0.152569 | 1 |
| Timm50 | -0.07625 | 0.678212 | 1 |
| Zfp719 | -0.0763 | 0.085869 | 1 |
| Rab1b | -0.0763 | 0.278774 | 1 |
| Lats1 | -0.07633 | 0.271709 | 1 |
| Gps2 | -0.07634 | 0.189048 | 1 |
| Rrm2b | -0.07636 | 0.111887 | 1 |
| Pqbp1 | -0.07639 | 0.022948 | 1 |
| Zfp704 | -0.0764 | 0.892242 | 1 |
| Anapc1 | -0.07651 | 0.192676 | 1 |
| Ubr1 | -0.07654 | 0.559271 | 1 |
| Atp6v0a2 | -0.07659 | 0.164424 | 1 |
| Fibp | -0.07665 | 0.953454 | 1 |
| Ccdc130 | -0.07668 | 0.356948 | 1 |
| Tmem50b | -0.07677 | 0.740464 | 1 |
| Iqcb1 | -0.07678 | 0.00133 | 1 |
| Agpat1 | -0.07678 | 0.080366 | 1 |
| Gripap1 | -0.0768 | 0.732541 | 1 |
| Emsy | -0.07687 | 0.326141 | 1 |
| Fhl3 | -0.07688 | 0.002625 | 1 |
| 2610001J0 | -0.07692 | 0.804916 | 1 |
| Cwc15 | -0.07695 | 0.766957 | 1 |
| Creld1 | -0.07702 | 0.06943 | 1 |
| Far1 | -0.07703 | 0.55497 | 1 |
| Zfp317 | -0.07706 | 0.105491 | 1 |
| Etfdh | -0.07709 | 0.161725 | 1 |
| Ube4b | -0.07711 | 0.865813 | 1 |
| Dock2 | -0.07712 | 3.1E-06 | 0.100069 |
| Dnajb12 | -0.07712 | 0.334417 | 1 |
| Rsf1 | -0.07729 | 0.684495 | 1 |
| Kdsr | -0.07729 | 0.193078 | 1 |
| Elmsan1 | -0.0773 | 0.677583 | 1 |
| Khsrp | -0.07733 | 0.287711 | 1 |
| Ppard | -0.07737 | 0.784732 | 1 |
| Erbin | -0.07738 | 0.871289 | 1 |
| Cep95 | -0.07742 | 0.880732 | 1 |
| Msl3 | -0.07744 | 0.094308 | 1 |
| Alg14 | -0.07751 | 0.926323 | 1 |
| Xlr | -0.07753 | 0.5188 | 1 |
| Gm49359 | -0.07756 | 0.183736 | 1 |
| Slc38a7 | -0.07757 | 0.765449 | 1 |
| Armc7 | -0.07762 | 0.401978 | 1 |
| Prpf6 | -0.07762 | 0.127645 | 1 |
| Setd1b | -0.0777 | 0.444431 | 1 |
| Taok2 | -0.07791 | 0.899 | 1 |
| Stx6 | -0.07793 | 0.880785 | 1 |

| | | | |
|---|---|---|---|
| Smg6 | -0.07795 | 0.216017 | 1 |
| Ufsp2 | -0.07802 | 0.698368 | 1 |
| Zscan26 | -0.07804 | 0.973804 | 1 |
| Pbxip1 | -0.07805 | 0.886859 | 1 |
| Gpn1 | -0.07811 | 0.618831 | 1 |
| Ipmk | -0.07811 | 0.971396 | 1 |
| As3mt | -0.07814 | 0.344597 | 1 |
| 5031425E2 | -0.0783 | 0.225416 | 1 |
| Mapk8ip3 | -0.07833 | 0.995706 | 1 |
| Pgm2 | -0.07836 | 0.132727 | 1 |
| Abcd1 | -0.07837 | 0.074041 | 1 |
| Derl1 | -0.07843 | 0.425082 | 1 |
| Vps41 | -0.07844 | 0.831654 | 1 |
| Fam219b | -0.07847 | 0.06582 | 1 |
| Slc33a1 | -0.07847 | 0.265964 | 1 |
| Krtcap2 | -0.07848 | 0.743435 | 1 |
| Nae1 | -0.0785 | 0.556354 | 1 |
| Ruvbl1 | -0.07852 | 0.246521 | 1 |
| Pcnp | -0.07854 | 0.761357 | 1 |
| Trappc12 | -0.0786 | 0.117906 | 1 |
| Slu7 | -0.07864 | 0.947746 | 1 |
| Ncor2 | -0.07867 | 0.378085 | 1 |
| Micu1 | -0.07868 | 0.317077 | 1 |
| Entr1 | -0.07871 | 0.795841 | 1 |
| Supt7l | -0.07873 | 0.090752 | 1 |
| Setdb1 | -0.07877 | 0.501292 | 1 |
| Retreg3 | -0.07882 | 0.126797 | 1 |
| Phf20 | -0.07886 | 0.959666 | 1 |
| Ypel2 | -0.07892 | 0.011279 | 1 |
| Cmtm4 | -0.07893 | 0.57815 | 1 |
| Ash1l | -0.07898 | 0.938557 | 1 |
| Klhdc3 | -0.07901 | 0.283089 | 1 |
| Brd7 | -0.07905 | 0.988686 | 1 |
| Ubap2l | -0.07912 | 0.480411 | 1 |
| E230029C0 | -0.07916 | 0.385032 | 1 |
| Scarb2 | -0.07919 | 0.121786 | 1 |
| Eif2b4 | -0.0792 | 0.417817 | 1 |
| Pstk | -0.07921 | 0.118161 | 1 |
| Prkdc | -0.07924 | 0.962671 | 1 |
| Casp6 | -0.07924 | 0.193324 | 1 |
| Cdca4 | -0.07928 | 0.101927 | 1 |
| Retsat | -0.07936 | 0.233938 | 1 |
| Zfp644 | -0.07939 | 0.807735 | 1 |
| Tardbp | -0.0795 | 0.771339 | 1 |
| Smu1 | -0.07956 | 0.088539 | 1 |
| Cdc73 | -0.07956 | 0.788437 | 1 |
| 3110056K0 | -0.07963 | 0.048721 | 1 |
| Nup88 | -0.07967 | 0.044076 | 1 |

| | | | |
|---|---|---|---|
| Plpp5 | -0.07968 | 0.016687 | 1 |
| Prdx3 | -0.07972 | 0.724452 | 1 |
| Pigt | -0.07973 | 0.684879 | 1 |
| Ap3d1 | -0.07974 | 0.485375 | 1 |
| Npepl1 | -0.07977 | 0.866474 | 1 |
| Ints14 | -0.07984 | 0.149111 | 1 |
| Cux1 | -0.07985 | 0.592675 | 1 |
| Apip | -0.07992 | 0.609207 | 1 |
| Atg4c | -0.07992 | 0.600892 | 1 |
| 4930453N | -0.07994 | 0.132496 | 1 |
| Znrf2 | -0.07995 | 0.803438 | 1 |
| Zc3h3 | -0.07999 | 0.503802 | 1 |
| C2cd3 | -0.08011 | 0.730447 | 1 |
| Gaa | -0.08011 | 0.735194 | 1 |
| Rab18 | -0.08015 | 0.40174 | 1 |
| Nova1 | -0.08018 | 0.132654 | 1 |
| Trim23 | -0.08018 | 0.065236 | 1 |
| Nlrp3 | -0.08018 | 0.491313 | 1 |
| Hps1 | -0.08021 | 0.135173 | 1 |
| Zfp414 | -0.08023 | 0.368082 | 1 |
| Zfp865 | -0.08023 | 0.135168 | 1 |
| Kdelr2 | -0.08026 | 0.426158 | 1 |
| Ublcp1 | -0.08034 | 0.186403 | 1 |
| Bzw1 | -0.08035 | 0.034313 | 1 |
| Ofd1 | -0.08041 | 0.571956 | 1 |
| Pabpn1 | -0.08041 | 0.692833 | 1 |
| Sec23a | -0.08051 | 0.034903 | 1 |
| Acvr2a | -0.08055 | 0.222939 | 1 |
| Ddx20 | -0.08055 | 0.151829 | 1 |
| Pot1a | -0.08058 | 0.131088 | 1 |
| Reep3 | -0.08061 | 0.635795 | 1 |
| Gtpbp2 | -0.08062 | 0.794865 | 1 |
| Spg11 | -0.08063 | 0.952867 | 1 |
| Pam16 | -0.08069 | 0.827236 | 1 |
| Tmem134 | -0.08072 | 0.616535 | 1 |
| Ptdss2 | -0.08076 | 0.202208 | 1 |
| Ppp1r35 | -0.08086 | 0.101161 | 1 |
| Fyttd1 | -0.08088 | 0.212145 | 1 |
| Rbsn | -0.0809 | 0.179749 | 1 |
| Mpnd | -0.08091 | 0.457418 | 1 |
| Smarcc2 | -0.08098 | 0.448055 | 1 |
| Arrdc3 | -0.08098 | 0.131453 | 1 |
| Bmt2 | -0.08103 | 0.892772 | 1 |
| Fam168b | -0.08104 | 0.442668 | 1 |
| Mettl5 | -0.08104 | 0.013713 | 1 |
| Commd2 | -0.08106 | 0.019148 | 1 |
| Lefty1 | -0.08108 | 0.023824 | 1 |
| Dcun1d2 | -0.08109 | 0.13402 | 1 |

| | | | |
|---|---|---|---|
| Fam135a | -0.08112 | 0.134669 | 1 |
| Rnf11 | -0.08119 | 0.681305 | 1 |
| Nup62 | -0.08128 | 0.958251 | 1 |
| Tmem256 | -0.08133 | 0.807296 | 1 |
| Aplp2 | -0.08164 | 0.817784 | 1 |
| Prpf40b | -0.08165 | 0.289007 | 1 |
| Fancl | -0.08167 | 0.17059 | 1 |
| Tmem63b | -0.0817 | 0.24954 | 1 |
| Tm2d3 | -0.08175 | 0.373868 | 1 |
| Gpr107 | -0.08176 | 0.899967 | 1 |
| Zfp35 | -0.08177 | 0.061573 | 1 |
| Ebp | -0.08189 | 0.352303 | 1 |
| Sap130 | -0.08189 | 0.814307 | 1 |
| Kdm4c | -0.0819 | 0.947076 | 1 |
| Trappc13 | -0.0819 | 0.008045 | 1 |
| Scd2 | -0.0819 | 0.013254 | 1 |
| Zfp703 | -0.08194 | 2.51E-05 | 0.809722 |
| Nsrp1 | -0.08196 | 0.565659 | 1 |
| Gpcpd1 | -0.08199 | 0.022239 | 1 |
| Tspan9 | -0.08203 | 0.523845 | 1 |
| Marf1 | -0.08205 | 0.461515 | 1 |
| Dcps | -0.08209 | 0.092 | 1 |
| Wac | -0.08217 | 0.885257 | 1 |
| Prrc2b | -0.08218 | 0.143292 | 1 |
| Dhx34 | -0.08223 | 0.211226 | 1 |
| Cul9 | -0.08238 | 0.260165 | 1 |
| Smarcc1 | -0.08248 | 0.600996 | 1 |
| Tmem203 | -0.08249 | 0.005166 | 1 |
| Rad23b | -0.08257 | 0.628051 | 1 |
| Srp54a | -0.08258 | 0.002921 | 1 |
| Mpp5 | -0.08271 | 0.010289 | 1 |
| Dhrs4 | -0.08276 | 0.43167 | 1 |
| Ap2b1 | -0.0828 | 0.531685 | 1 |
| Gm20703 | -0.08282 | 0.014052 | 1 |
| Smg9 | -0.0829 | 0.390231 | 1 |
| Tnks2 | -0.08293 | 0.866569 | 1 |
| Tjp2 | -0.08303 | 0.176581 | 1 |
| Akap7 | -0.08311 | 0.884604 | 1 |
| Gm49797 | -0.08322 | 0.163872 | 1 |
| Rab40c | -0.08324 | 0.318611 | 1 |
| Zdhhc17 | -0.08331 | 0.287242 | 1 |
| Sesn2 | -0.08343 | 0.160955 | 1 |
| Arhgef11 | -0.08344 | 0.555735 | 1 |
| Ogt | -0.08362 | 0.020866 | 1 |
| Tug1 | -0.08364 | 0.111515 | 1 |
| Pde6d | -0.08376 | 0.103577 | 1 |
| Vps18 | -0.08377 | 0.254974 | 1 |
| Zmpste24 | -0.08379 | 0.999055 | 1 |

| | | | |
|---|---|---|---|
| Anapc4 | -0.0838 | 0.383858 | 1 |
| Ganab | -0.08389 | 0.518991 | 1 |
| Il7r | -0.0839 | 0.050811 | 1 |
| Srp19 | -0.08393 | 0.32573 | 1 |
| Dr1 | -0.08399 | 0.360417 | 1 |
| Brwd1 | -0.08399 | 0.455196 | 1 |
| Cxxc1 | -0.084 | 0.01006 | 1 |
| Ssu72 | -0.08403 | 0.316392 | 1 |
| Rnaseh2c | -0.08406 | 0.146723 | 1 |
| Med23 | -0.08406 | 0.695977 | 1 |
| Cgrrf1 | -0.08414 | 0.306604 | 1 |
| Pogz | -0.08427 | 0.333042 | 1 |
| Ctdsp1 | -0.08429 | 0.017806 | 1 |
| Snrnp200 | -0.08432 | 0.411208 | 1 |
| Srprb | -0.08432 | 0.148267 | 1 |
| Elp3 | -0.08435 | 0.038237 | 1 |
| Gm6712 | -0.08437 | 0.120103 | 1 |
| Mrpl48 | -0.08437 | 0.211919 | 1 |
| Bloc1s6 | -0.0846 | 0.127814 | 1 |
| Elmo2 | -0.08461 | 0.060251 | 1 |
| Xab2 | -0.08462 | 0.417027 | 1 |
| Mcm3 | -0.08466 | 0.452954 | 1 |
| Ankrd10 | -0.08469 | 0.039797 | 1 |
| Aldh6a1 | -0.08473 | 0.032142 | 1 |
| Zfp68 | -0.08483 | 0.037269 | 1 |
| Gm10138 | -0.08489 | 0.082026 | 1 |
| Dcun1d1 | -0.08491 | 0.267036 | 1 |
| Prcc | -0.08493 | 0.167289 | 1 |
| Myo7a | -0.085 | 0.992025 | 1 |
| Disp1 | -0.08506 | 0.012404 | 1 |
| Nabp2 | -0.08511 | 0.23126 | 1 |
| 1190007I0 | -0.08513 | 0.082673 | 1 |
| Msh2 | -0.08515 | 0.371625 | 1 |
| Fam50a | -0.08515 | 0.181911 | 1 |
| Gm39469 | -0.0852 | 0.090891 | 1 |
| Nudcd3 | -0.08524 | 0.427667 | 1 |
| Capn3 | -0.08536 | 0.155937 | 1 |
| Zfp955b | -0.08548 | 0.005452 | 1 |
| Cpsf2 | -0.08549 | 0.377636 | 1 |
| Senp3 | -0.08556 | 0.004259 | 1 |
| Kat7 | -0.08557 | 0.225095 | 1 |
| Jmjd1c | -0.08565 | 0.286786 | 1 |
| Shoc2 | -0.08572 | 0.273023 | 1 |
| Zmym6 | -0.08577 | 0.032317 | 1 |
| D1Ertd622 | -0.08592 | 0.276964 | 1 |
| Sulf2 | -0.08595 | 0.87205 | 1 |
| Ccdc47 | -0.08604 | 0.628085 | 1 |
| Prr12 | -0.08617 | 0.021897 | 1 |

| | | | |
|---|---|---|---|
| Dapp1 | -0.08617 | 0.117032 | 1 |
| Tgfa | -0.0862 | 0.166569 | 1 |
| Nit1 | -0.08623 | 0.091146 | 1 |
| Hectd4 | -0.08628 | 0.813568 | 1 |
| Smndc1 | -0.08629 | 0.427873 | 1 |
| Vps26c | -0.08636 | 0.225869 | 1 |
| Faap100 | -0.08638 | 0.081333 | 1 |
| Ubr3 | -0.08646 | 0.640302 | 1 |
| Tmem106b | -0.08647 | 0.002364 | 1 |
| Ipo13 | -0.08652 | 0.002663 | 1 |
| Mbtps1 | -0.08652 | 0.187417 | 1 |
| Txlna | -0.08661 | 0.909726 | 1 |
| Wipf2 | -0.08666 | 0.177182 | 1 |
| Cdc23 | -0.08671 | 0.484146 | 1 |
| Zfp629 | -0.08672 | 0.002936 | 1 |
| Senp6 | -0.08686 | 0.057292 | 1 |
| Hmgn1 | -0.0869 | 0.233396 | 1 |
| Zfp217 | -0.08694 | 0.652247 | 1 |
| Pex5 | -0.08697 | 0.308981 | 1 |
| Alyref2 | -0.08702 | 0.106334 | 1 |
| Ubtf | -0.08704 | 0.036589 | 1 |
| Fem1a | -0.08705 | 0.009596 | 1 |
| Cdk10 | -0.08707 | 0.299428 | 1 |
| Kbtbd3 | -0.08711 | 0.044139 | 1 |
| Ipo8 | -0.08724 | 0.891373 | 1 |
| Bsg | -0.08724 | 0.027497 | 1 |
| Dbndd2 | -0.0873 | 0.755991 | 1 |
| Renbp | -0.08732 | 0.645505 | 1 |
| Ssbp1 | -0.08737 | 0.259404 | 1 |
| Rb1cc1 | -0.0874 | 0.267884 | 1 |
| Sh3gl1 | -0.08743 | 0.160983 | 1 |
| Atg12 | -0.08747 | 0.257872 | 1 |
| Hnrnpf | -0.08753 | 0.150495 | 1 |
| Lrig2 | -0.08756 | 0.326712 | 1 |
| Klhl25 | -0.08757 | 0.014131 | 1 |
| Itgb3bp | -0.08758 | 0.018278 | 1 |
| Pggt1b | -0.08768 | 0.307145 | 1 |
| Phf23 | -0.0878 | 0.111628 | 1 |
| Stx4a | -0.08782 | 0.601112 | 1 |
| Tbc1d22a | -0.08789 | 0.387182 | 1 |
| Pogk | -0.08792 | 0.058853 | 1 |
| Map11 | -0.08803 | 0.024481 | 1 |
| Slc25a51 | -0.08809 | 0.034125 | 1 |
| Arpc5l | -0.08809 | 0.912702 | 1 |
| Ggps1 | -0.0881 | 0.348557 | 1 |
| Tnpo2 | -0.08814 | 0.288621 | 1 |
| Lpcat2 | -0.08816 | 0.000831 | 1 |
| Ddhd2 | -0.08817 | 0.436018 | 1 |

| | | | |
|---|---|---|---|
| Maea | -0.08819 | 0.537673 | 1 |
| Hdhd2 | -0.0882 | 0.087996 | 1 |
| Dnmbp | -0.08821 | 0.108836 | 1 |
| Eapp | -0.08828 | 0.402289 | 1 |
| Stard5 | -0.08832 | 0.030792 | 1 |
| Taf12 | -0.08837 | 0.396981 | 1 |
| Slc25a17 | -0.08839 | 0.303653 | 1 |
| Gsdme | -0.08842 | 0.020318 | 1 |
| Zfp729b | -0.08848 | 0.975912 | 1 |
| Kansl2 | -0.08852 | 0.094152 | 1 |
| AU040320 | -0.08853 | 0.001815 | 1 |
| Fut8 | -0.0886 | 0.050486 | 1 |
| Npl | -0.08863 | 0.09573 | 1 |
| Zfp384 | -0.0887 | 0.001266 | 1 |
| Trim41 | -0.08877 | 0.60986 | 1 |
| Top3b | -0.08879 | 0.306437 | 1 |
| Pigyl | -0.0888 | 0.074718 | 1 |
| Tex264 | -0.08882 | 0.186135 | 1 |
| Snx19 | -0.08883 | 0.892745 | 1 |
| Slc38a10 | -0.08888 | 0.08073 | 1 |
| Gm37233 | -0.08902 | 0.028279 | 1 |
| Gtf2e2 | -0.08909 | 0.026047 | 1 |
| 1700028E1 | -0.08912 | 0.092167 | 1 |
| Rrnad1 | -0.08919 | 0.762731 | 1 |
| Pygb | -0.08924 | 0.019651 | 1 |
| Chmp6 | -0.08924 | 0.118178 | 1 |
| Ginm1 | -0.08935 | 0.226557 | 1 |
| Rmnd5a | -0.08936 | 0.083848 | 1 |
| Ate1 | -0.08941 | 0.228022 | 1 |
| Clk3 | -0.08958 | 0.606416 | 1 |
| Gtf3c2 | -0.08959 | 0.02514 | 1 |
| Pskh1 | -0.08976 | 0.549128 | 1 |
| Il17ra | -0.08979 | 0.082725 | 1 |
| Alg9 | -0.08983 | 0.307275 | 1 |
| Ap1s2 | -0.08984 | 0.047776 | 1 |
| Napa | -0.0899 | 0.130158 | 1 |
| Fam129a | -0.08996 | 0.136964 | 1 |
| Akirin2 | -0.09001 | 0.054753 | 1 |
| 2210016F1 | -0.09004 | 0.395378 | 1 |
| Ap2a1 | -0.09009 | 0.600231 | 1 |
| Sh2d3c | -0.09013 | 0.016761 | 1 |
| Arcn1 | -0.09022 | 0.217339 | 1 |
| Faf2 | -0.09022 | 0.301407 | 1 |
| Mtif3 | -0.09027 | 0.009613 | 1 |
| Hk2 | -0.09034 | 0.855285 | 1 |
| Ro60 | -0.09039 | 0.023756 | 1 |
| Acox1 | -0.09047 | 0.25241 | 1 |
| Galnt2 | -0.09049 | 0.065402 | 1 |

| | | | |
|---|---|---|---|
| Ylpm1 | -0.09054 | 0.650867 | 1 |
| Mrpl43 | -0.09055 | 0.088713 | 1 |
| Clock | -0.09064 | 0.896359 | 1 |
| Hk1 | -0.09067 | 0.000234 | 1 |
| Angel2 | -0.09077 | 0.881164 | 1 |
| Por | -0.0908 | 0.106435 | 1 |
| Sf3a1 | -0.0908 | 0.558983 | 1 |
| Mmd | -0.09081 | 0.429049 | 1 |
| Kpna6 | -0.09085 | 0.117298 | 1 |
| Rpl7l1 | -0.09091 | 0.339602 | 1 |
| Appl1 | -0.09091 | 0.032621 | 1 |
| Pip4p2 | -0.09096 | 0.114019 | 1 |
| Myh9 | -0.09097 | 0.003208 | 1 |
| Rcn2 | -0.09097 | 0.163345 | 1 |
| Rcc2 | -0.09112 | 0.407184 | 1 |
| Kiz | -0.09113 | 0.092001 | 1 |
| Tnni2 | -0.09122 | 0.002121 | 1 |
| Slc45a4 | -0.09123 | 0.193455 | 1 |
| Fbxo22 | -0.0913 | 0.58932 | 1 |
| Alg2 | -0.09135 | 0.200475 | 1 |
| Ndufs2 | -0.09136 | 0.252612 | 1 |
| Pigu | -0.09138 | 0.456408 | 1 |
| Brox | -0.09145 | 0.144752 | 1 |
| Arpc1a | -0.09149 | 0.335079 | 1 |
| Dennd1a | -0.09155 | 0.809454 | 1 |
| Smim1 | -0.09161 | 0.001284 | 1 |
| Fech | -0.09165 | 0.264917 | 1 |
| Unk | -0.09189 | 0.034063 | 1 |
| Zfyve21 | -0.09193 | 0.016913 | 1 |
| Eif4g3 | -0.09195 | 0.127209 | 1 |
| Rprd1b | -0.09203 | 0.270398 | 1 |
| Cradd | -0.09208 | 0.552726 | 1 |
| Ptpn9 | -0.09217 | 0.469951 | 1 |
| Rbm4 | -0.09223 | 0.128087 | 1 |
| Iars | -0.09225 | 0.871495 | 1 |
| F8a | -0.09226 | 0.002245 | 1 |
| Tyw1 | -0.09228 | 0.591094 | 1 |
| Tmc7 | -0.0923 | 0.002238 | 1 |
| Acsl1 | -0.09231 | 0.453122 | 1 |
| Rbm5 | -0.09234 | 0.228545 | 1 |
| Zhx3 | -0.09234 | 0.047579 | 1 |
| Pdcd6ip | -0.09248 | 0.467036 | 1 |
| Tubgcp5 | -0.09258 | 0.158833 | 1 |
| Hnrnpd | -0.09264 | 0.242058 | 1 |
| Mfap1b | -0.09265 | 0.001056 | 1 |
| Ndfip1 | -0.09273 | 0.022526 | 1 |
| Fto | -0.0928 | 0.995495 | 1 |
| Gm29291 | -0.0928 | 8.83E-05 | 1 |

| | | | |
|---|---|---|---|
| Gm15564 | -0.09285 | 0.141078 | 1 |
| Trmt12 | -0.09286 | 0.032495 | 1 |
| Lrrcc1 | -0.09296 | 0.197798 | 1 |
| Ebpl | -0.09303 | 0.023995 | 1 |
| Itprid2 | -0.09324 | 0.504057 | 1 |
| Gm41764 | -0.09327 | 0.005803 | 1 |
| Ip6k2 | -0.09328 | 0.034077 | 1 |
| Camkk2 | -0.09329 | 0.789206 | 1 |
| Vasp | -0.09333 | 0.437751 | 1 |
| Sympk | -0.09333 | 0.162166 | 1 |
| Orai2 | -0.09339 | 0.549589 | 1 |
| Ank2 | -0.09344 | 0.910878 | 1 |
| Ppp3ca | -0.09351 | 0.117189 | 1 |
| Ankra2 | -0.09354 | 0.043171 | 1 |
| Ctns | -0.09357 | 0.206754 | 1 |
| Msl1 | -0.0936 | 0.432002 | 1 |
| Pdcl3 | -0.09363 | 0.272009 | 1 |
| Josd2 | -0.09368 | 0.69987 | 1 |
| Senp8 | -0.09373 | 0.009498 | 1 |
| Nxf1 | -0.09376 | 0.59381 | 1 |
| Tmem185b | -0.09388 | 0.047642 | 1 |
| Eif2ak1 | -0.09393 | 0.080955 | 1 |
| Sgsm3 | -0.09393 | 0.006637 | 1 |
| Selenos | -0.09401 | 0.080769 | 1 |
| Clcc1 | -0.09402 | 0.182195 | 1 |
| Cdc42se2 | -0.09405 | 0.036021 | 1 |
| Tmem101 | -0.09407 | 0.159289 | 1 |
| Prrg2 | -0.09412 | 0.004962 | 1 |
| Smpd2 | -0.09412 | 0.009385 | 1 |
| Bcl2l11 | -0.09415 | 5.99E-07 | 0.019335 |
| Ppm1f | -0.09426 | 0.423326 | 1 |
| Ganc | -0.0943 | 0.488151 | 1 |
| Smpdl3a | -0.0943 | 0.038757 | 1 |
| Idh3g | -0.09431 | 0.106149 | 1 |
| Tpra1 | -0.09446 | 0.243444 | 1 |
| Ctdspl2 | -0.09459 | 0.036563 | 1 |
| Foxo3 | -0.09461 | 0.021695 | 1 |
| Polr1a | -0.09465 | 0.684021 | 1 |
| Ppp2r5a | -0.09467 | 0.083311 | 1 |
| Fuz | -0.0947 | 0.007552 | 1 |
| Fkbp7 | -0.09478 | 0.015598 | 1 |
| Poldip2 | -0.09483 | 0.746311 | 1 |
| Srr | -0.09488 | 0.093081 | 1 |
| Cdc5l | -0.09488 | 0.083769 | 1 |
| Atp6v1e1 | -0.09493 | 0.172692 | 1 |
| Ddx18 | -0.09499 | 0.341629 | 1 |
| Cul3 | -0.095 | 0.35059 | 1 |
| Hace1 | -0.0951 | 0.039431 | 1 |

| | | | |
|---|---|---|---|
| Crls1 | -0.0952 | 0.00761 | 1 |
| Phax | -0.0953 | 0.602179 | 1 |
| Sap18 | -0.09531 | 0.094006 | 1 |
| Ppp4r1 | -0.09536 | 0.459592 | 1 |
| Elp4 | -0.09541 | 0.464046 | 1 |
| Bud31 | -0.09552 | 0.457193 | 1 |
| Abraxas2 | -0.09561 | 0.134909 | 1 |
| Zfp263 | -0.09574 | 0.250542 | 1 |
| Ndufaf2 | -0.0958 | 0.017624 | 1 |
| Zcchc2 | -0.0958 | 0.018847 | 1 |
| Kif16b | -0.09585 | 0.107343 | 1 |
| Ppp6r2 | -0.09593 | 0.033395 | 1 |
| Shprh | -0.09596 | 0.106091 | 1 |
| Atxn2 | -0.09599 | 0.588249 | 1 |
| Mob3a | -0.096 | 0.25199 | 1 |
| Tnrc6c | -0.09608 | 0.930052 | 1 |
| Sowahc | -0.0961 | 0.032951 | 1 |
| Vps37a | -0.09612 | 0.107682 | 1 |
| Otulin | -0.09617 | 0.739828 | 1 |
| Ilvbl | -0.0962 | 0.700214 | 1 |
| Mpc2 | -0.09624 | 0.339438 | 1 |
| Cdk2ap1 | -0.09631 | 0.002791 | 1 |
| Tlr13 | -0.09632 | 0.136199 | 1 |
| Stam | -0.09634 | 0.097057 | 1 |
| Hint3 | -0.09642 | 0.236802 | 1 |
| Pdss2 | -0.0965 | 0.396409 | 1 |
| Myo1f | -0.09655 | 0.042408 | 1 |
| Map4k3 | -0.09656 | 0.391692 | 1 |
| Nudt16 | -0.09657 | 0.174312 | 1 |
| Dolk | -0.0966 | 0.004457 | 1 |
| Leng8 | -0.09664 | 0.380149 | 1 |
| Psmc2 | -0.09672 | 0.25947 | 1 |
| Zfp672 | -0.09685 | 0.128979 | 1 |
| Zfp933 | -0.09685 | 0.019515 | 1 |
| Zbed3 | -0.09685 | 0.176955 | 1 |
| Rbm27 | -0.09692 | 0.326271 | 1 |
| Pkp4 | -0.09694 | 0.73752 | 1 |
| Suclg1 | -0.09694 | 0.165794 | 1 |
| Tfip11 | -0.09701 | 0.006692 | 1 |
| Epc2 | -0.09709 | 0.105833 | 1 |
| Zfp120 | -0.09711 | 0.186951 | 1 |
| Nrbp1 | -0.09712 | 0.851239 | 1 |
| Il31ra | -0.09718 | 0.000846 | 1 |
| Ticam2 | -0.09721 | 0.010155 | 1 |
| Pigq | -0.09724 | 0.080002 | 1 |
| Mocs2 | -0.09727 | 0.257699 | 1 |
| Clptm1l | -0.0973 | 0.485198 | 1 |
| Gins4 | -0.09739 | 0.001765 | 1 |

| | | | |
|---|---|---|---|
| Vamp4 | -0.09742 | 0.160911 | 1 |
| Tmco6 | -0.09742 | 0.003751 | 1 |
| Lztr1 | -0.09746 | 0.121133 | 1 |
| Ndrg3 | -0.09747 | 0.028064 | 1 |
| Slc37a4 | -0.09756 | 0.149869 | 1 |
| Dync1h1 | -0.0976 | 0.372437 | 1 |
| Stk11 | -0.09762 | 0.395821 | 1 |
| Polr2j | -0.09765 | 0.378063 | 1 |
| Cstf2t | -0.09769 | 2.11E-05 | 0.682096 |
| Smc3 | -0.0977 | 0.081941 | 1 |
| Dgat1 | -0.09775 | 0.019957 | 1 |
| Romo1 | -0.09786 | 0.581167 | 1 |
| Cript | -0.09828 | 0.165689 | 1 |
| Fam214a | -0.09836 | 0.089458 | 1 |
| Kctd21 | -0.09841 | 0.000483 | 1 |
| Pik3r2 | -0.09843 | 0.046679 | 1 |
| Pigh | -0.0985 | 0.002067 | 1 |
| Apeh | -0.09852 | 0.542041 | 1 |
| Zkscan1 | -0.09856 | 0.319308 | 1 |
| Csnk1d | -0.09859 | 0.208344 | 1 |
| Aarsd1 | -0.09867 | 0.071012 | 1 |
| Fubp3 | -0.09878 | 0.119224 | 1 |
| Sdf2 | -0.09881 | 0.250346 | 1 |
| Rab3gap2 | -0.0989 | 0.702905 | 1 |
| Ncoa2 | -0.09892 | 0.077642 | 1 |
| Lysmd3 | -0.09899 | 0.029488 | 1 |
| Ehbp1 | -0.09908 | 0.226967 | 1 |
| Ppwd1 | -0.09914 | 0.265694 | 1 |
| Rbm43 | -0.09916 | 0.005513 | 1 |
| Gopc | -0.09917 | 0.199782 | 1 |
| Gm15726 | -0.0992 | 0.02259 | 1 |
| Ciz1 | -0.0992 | 0.017002 | 1 |
| Nelfb | -0.09924 | 0.016184 | 1 |
| Hvcn1 | -0.09933 | 0.017922 | 1 |
| Tti1 | -0.09941 | 0.482217 | 1 |
| Casp3 | -0.09942 | 0.069729 | 1 |
| Cul5 | -0.09942 | 0.921286 | 1 |
| Hikeshi | -0.09946 | 0.036392 | 1 |
| Stat5b | -0.09953 | 0.011409 | 1 |
| Nf2 | -0.09956 | 0.474114 | 1 |
| Hmgcs1 | -0.09958 | 0.196234 | 1 |
| Cog1 | -0.09959 | 0.107281 | 1 |
| Atg101 | -0.09963 | 0.000823 | 1 |
| Pcmtd1 | -0.09967 | 0.4396 | 1 |
| Eloc | -0.09979 | 0.193992 | 1 |
| Tfg | -0.0998 | 0.868517 | 1 |
| Vkorc1 | -0.09981 | 0.279622 | 1 |
| Nudt16l1 | -0.09982 | 0.001163 | 1 |

| | | | |
|---|---|---|---|
| Cdc42bpa | -0.09985 | 0.598521 | 1 |
| Qdpr | -0.09991 | 0.18435 | 1 |
| Uqcc3 | -0.10011 | 0.039733 | 1 |
| Dpy30 | -0.10015 | 0.411989 | 1 |
| Ccnd1 | -0.10019 | 0.464249 | 1 |
| Zfp740 | -0.10029 | 0.006101 | 1 |
| Crnkl1 | -0.10033 | 0.144753 | 1 |
| Gatb | -0.10039 | 0.663275 | 1 |
| Rap2c | -0.10044 | 0.238226 | 1 |
| Oaf | -0.10054 | 0.549584 | 1 |
| Anapc11 | -0.10057 | 0.102653 | 1 |
| Zfp212 | -0.1006 | 0.009672 | 1 |
| Cdkn2aip | -0.10061 | 0.036751 | 1 |
| Iqsec2 | -0.10069 | 0.682909 | 1 |
| Hnrnpu | -0.1007 | 0.017404 | 1 |
| Orai3 | -0.10075 | 0.036428 | 1 |
| Errfi1 | -0.10101 | 0.006358 | 1 |
| Ing1 | -0.10108 | 0.017851 | 1 |
| Ssr1 | -0.10113 | 0.244775 | 1 |
| Smpd5 | -0.10126 | 0.029839 | 1 |
| Slc39a1 | -0.1013 | 0.328922 | 1 |
| Ppp4r2 | -0.10135 | 0.675445 | 1 |
| Rc3h2 | -0.10138 | 0.464682 | 1 |
| Nsd1 | -0.10141 | 0.101617 | 1 |
| Med19 | -0.10148 | 0.013639 | 1 |
| Gm15537 | -0.10152 | 0.004698 | 1 |
| Polr2b | -0.10168 | 0.160985 | 1 |
| Snx3 | -0.1018 | 0.106007 | 1 |
| Dnajc7 | -0.10192 | 0.373123 | 1 |
| Zmym3 | -0.10194 | 0.007583 | 1 |
| Ptgs1 | -0.10201 | 1.25E-05 | 0.404035 |
| Mrpl36 | -0.10205 | 0.048439 | 1 |
| Agap1 | -0.10221 | 0.17646 | 1 |
| Hspa12a | -0.10225 | 0.112078 | 1 |
| Bola1 | -0.10229 | 0.197415 | 1 |
| Appl2 | -0.10237 | 0.028968 | 1 |
| Ktn1 | -0.10243 | 0.35405 | 1 |
| Ankrd13c | -0.1025 | 0.129247 | 1 |
| Filip1l | -0.10253 | 0.086868 | 1 |
| Ubr2 | -0.10261 | 0.130267 | 1 |
| Rab33b | -0.10262 | 0.001088 | 1 |
| Amz2 | -0.10266 | 0.107777 | 1 |
| Api5 | -0.10267 | 0.053726 | 1 |
| Syap1 | -0.10278 | 0.148664 | 1 |
| Nagpa | -0.1029 | 0.452172 | 1 |
| Ring1 | -0.10297 | 0.0988 | 1 |
| Vezt | -0.10297 | 0.106307 | 1 |
| Prkra | -0.10298 | 0.074166 | 1 |

| | | | |
|---|---|---|---|
| March7 | -0.10302 | 0.022855 | 1 |
| Ube2r2 | -0.10305 | 0.779996 | 1 |
| Med22 | -0.10307 | 0.083325 | 1 |
| Arf2 | -0.10308 | 0.073926 | 1 |
| Cacna1d | -0.1031 | 0.135114 | 1 |
| Cyb5r1 | -0.10311 | 0.319957 | 1 |
| Smc6 | -0.10312 | 0.263262 | 1 |
| Spsb2 | -0.1032 | 0.150853 | 1 |
| Mbd1 | -0.10323 | 0.137733 | 1 |
| Phospho2 | -0.10325 | 0.00198 | 1 |
| E2f3 | -0.10325 | 0.550448 | 1 |
| Fadd | -0.10327 | 0.003883 | 1 |
| Ost4 | -0.10335 | 0.361443 | 1 |
| Otub1 | -0.10336 | 0.687317 | 1 |
| 1600012H( | -0.10342 | 0.328078 | 1 |
| Map1lc3a | -0.10351 | 0.003822 | 1 |
| Rps6ka4 | -0.10352 | 0.160367 | 1 |
| Slx1b | -0.10357 | 0.007229 | 1 |
| Wdr26 | -0.10368 | 0.105382 | 1 |
| Snrpa | -0.10371 | 0.158017 | 1 |
| Nol4l | -0.10372 | 0.711946 | 1 |
| Dapk3 | -0.10376 | 0.144947 | 1 |
| Smarcd1 | -0.10378 | 0.025387 | 1 |
| Psmd11 | -0.10382 | 0.055647 | 1 |
| Xpo5 | -0.10383 | 0.047919 | 1 |
| Sucla2 | -0.10389 | 0.137911 | 1 |
| Nisch | -0.1039 | 0.420278 | 1 |
| Txndc12 | -0.10391 | 0.039724 | 1 |
| Chmp2b | -0.10392 | 0.0044 | 1 |
| Bcas2 | -0.10401 | 0.149737 | 1 |
| Dhrs7b | -0.10411 | 0.680224 | 1 |
| Eefsec | -0.10416 | 0.424939 | 1 |
| Zmym5 | -0.10417 | 0.236435 | 1 |
| Tfeb | -0.10422 | 0.58176 | 1 |
| Dync1li1 | -0.10428 | 0.850168 | 1 |
| Slc2a9 | -0.10431 | 0.737374 | 1 |
| Alkbh5 | -0.10434 | 0.05627 | 1 |
| Yipf4 | -0.10435 | 0.389975 | 1 |
| Vamp2 | -0.10441 | 0.034906 | 1 |
| Smpdl3b | -0.10468 | 0.000282 | 1 |
| Snx4 | -0.10471 | 0.053661 | 1 |
| Tmem159 | -0.10473 | 0.031836 | 1 |
| Polr2k | -0.10474 | 0.289953 | 1 |
| Lpar5 | -0.10483 | 0.013962 | 1 |
| Ints4 | -0.10487 | 0.068637 | 1 |
| Ppp1r9b | -0.1049 | 0.085238 | 1 |
| Map3k14 | -0.10492 | 0.259949 | 1 |
| Ppp6c | -0.10495 | 0.137114 | 1 |

| | | | |
|---|---|---|---|
| Spag7 | -0.10497 | 0.02412 | 1 |
| Platr25 | -0.10508 | 0.053924 | 1 |
| Brip1os | -0.10508 | 0.164214 | 1 |
| Sec23b | -0.10511 | 0.089243 | 1 |
| Vps26a | -0.1052 | 0.126029 | 1 |
| Cstf3 | -0.10524 | 0.273122 | 1 |
| Smim8 | -0.10529 | 0.001485 | 1 |
| 2610528A1 | -0.10532 | 0.18734 | 1 |
| Tcf7l2 | -0.10537 | 0.608695 | 1 |
| Dlgap4 | -0.1054 | 0.859529 | 1 |
| Galnt10 | -0.10541 | 0.870801 | 1 |
| Bap1 | -0.10547 | 0.041685 | 1 |
| Zc3h4 | -0.10548 | 0.056532 | 1 |
| Brd9 | -0.10551 | 0.128851 | 1 |
| Wbp4 | -0.10552 | 0.590634 | 1 |
| Zfp617 | -0.10553 | 0.004649 | 1 |
| Lzic | -0.10553 | 0.008859 | 1 |
| Stard3nl | -0.10561 | 0.099708 | 1 |
| Bcas3 | -0.10562 | 0.568881 | 1 |
| Atn1 | -0.1057 | 0.000745 | 1 |
| Celf2 | -0.10573 | 0.000205 | 1 |
| Kat8 | -0.1058 | 0.012158 | 1 |
| Layn | -0.10584 | 0.008168 | 1 |
| Gak | -0.10585 | 0.7853 | 1 |
| Dusp3 | -0.10589 | 0.043539 | 1 |
| Midn | -0.10611 | 0.100053 | 1 |
| Ccdc22 | -0.10616 | 0.023687 | 1 |
| Pdxdc1 | -0.10618 | 0.704909 | 1 |
| Gatd3a | -0.1062 | 0.039202 | 1 |
| Zc3h6 | -0.10626 | 0.009104 | 1 |
| Slc7a7 | -0.10635 | 0.384322 | 1 |
| Dzip3 | -0.10641 | 0.013225 | 1 |
| Tbc1d12 | -0.10645 | 0.013349 | 1 |
| Map3k5 | -0.10646 | 0.04924 | 1 |
| Yars | -0.10646 | 0.072368 | 1 |
| Dcp1b | -0.10648 | 0.036651 | 1 |
| Btf3l4 | -0.10652 | 0.047979 | 1 |
| Prdm2 | -0.10659 | 0.312959 | 1 |
| Ppp3r1 | -0.10662 | 0.016524 | 1 |
| Kansl1 | -0.10662 | 0.387508 | 1 |
| Spire1 | -0.1067 | 0.564946 | 1 |
| Usf1 | -0.10687 | 0.000849 | 1 |
| 2010013B2 | -0.10691 | 0.002571 | 1 |
| Lman2l | -0.10693 | 0.442446 | 1 |
| Pomt1 | -0.10697 | 0.072853 | 1 |
| Strip1 | -0.10699 | 0.401518 | 1 |
| Nelfcd | -0.10703 | 0.001819 | 1 |
| Pecam1 | -0.10704 | 4.44E-08 | 0.001435 |

| | | | |
|---|---|---|---|
| Usp11 | -0.10709 | 0.00039 | 1 |
| Fbxl14 | -0.1071 | 0.022251 | 1 |
| Wdsub1 | -0.10713 | 0.000406 | 1 |
| Zbtb37 | -0.10715 | 0.014654 | 1 |
| Dnpep | -0.10731 | 0.053228 | 1 |
| Bphl | -0.10741 | 0.000785 | 1 |
| Spns1 | -0.10775 | 0.23053 | 1 |
| Pgpep1 | -0.10786 | 0.00386 | 1 |
| B4galt3 | -0.10792 | 0.021151 | 1 |
| Tollip | -0.10794 | 0.000501 | 1 |
| Pdk1 | -0.10799 | 0.16835 | 1 |
| Ick | -0.10801 | 0.000461 | 1 |
| Bsdc1 | -0.10806 | 0.007598 | 1 |
| A530072M | -0.1081 | 0.169847 | 1 |
| Tmed4 | -0.10812 | 0.052842 | 1 |
| Oga | -0.10813 | 0.124846 | 1 |
| Pigc | -0.10814 | 0.064062 | 1 |
| Idua | -0.10817 | 0.217563 | 1 |
| Nxt2 | -0.10821 | 0.029663 | 1 |
| Mrpl34 | -0.10832 | 0.284392 | 1 |
| Cand1 | -0.10835 | 0.03968 | 1 |
| Cnot6l | -0.10842 | 0.026478 | 1 |
| Srrm1 | -0.10842 | 0.024608 | 1 |
| Trappc4 | -0.10844 | 0.008333 | 1 |
| Ncf2 | -0.10847 | 0.074932 | 1 |
| Aaas | -0.10868 | 0.178559 | 1 |
| Ubtd1 | -0.10869 | 0.40725 | 1 |
| Man2a2 | -0.10873 | 0.01979 | 1 |
| Agpat3 | -0.1089 | 0.221406 | 1 |
| Tmem115 | -0.10898 | 0.033872 | 1 |
| Ppp2r3a | -0.10899 | 0.002481 | 1 |
| Tbck | -0.10903 | 0.612312 | 1 |
| Zbtb18 | -0.10909 | 0.039423 | 1 |
| Zfp277 | -0.1091 | 0.33148 | 1 |
| Zfp422 | -0.10917 | 0.003544 | 1 |
| Rraga | -0.10933 | 0.487153 | 1 |
| Med13 | -0.10946 | 0.308985 | 1 |
| Zfp866 | -0.10951 | 0.002262 | 1 |
| Impa1 | -0.10953 | 0.030027 | 1 |
| Hibch | -0.10961 | 0.004223 | 1 |
| Tm7sf3 | -0.10968 | 0.034925 | 1 |
| Wdfy4 | -0.10973 | 0.420743 | 1 |
| Ikzf1 | -0.10979 | 0.000477 | 1 |
| Gnpda2 | -0.10988 | 0.035598 | 1 |
| Fig4 | -0.10988 | 0.13773 | 1 |
| Maml2 | -0.10991 | 0.445726 | 1 |
| Sec24c | -0.10993 | 0.028485 | 1 |
| Fam118b | -0.11002 | 0.009344 | 1 |

| | | | |
|---|---|---|---|
| Gatad2b | -0.11005 | 0.752825 | 1 |
| Zfp236 | -0.11006 | 0.475314 | 1 |
| Mast2 | -0.11015 | 0.402508 | 1 |
| Nckap5l | -0.11016 | 0.160563 | 1 |
| Ppp1r11 | -0.11019 | 0.352538 | 1 |
| Dstyk | -0.11021 | 0.051076 | 1 |
| Dhx57 | -0.11025 | 0.04784 | 1 |
| Ago1 | -0.11035 | 0.288923 | 1 |
| Kctd20 | -0.1104 | 0.024523 | 1 |
| Dpf2 | -0.1105 | 0.232649 | 1 |
| Dcakd | -0.11056 | 0.561142 | 1 |
| Plxna4os1 | -0.11059 | 0.034156 | 1 |
| Apbb2 | -0.11064 | 0.354326 | 1 |
| Sufu | -0.11069 | 0.275769 | 1 |
| Dalrd3 | -0.11075 | 0.000496 | 1 |
| D030028A( | -0.11081 | 0.144256 | 1 |
| Elp2 | -0.11087 | 0.246337 | 1 |
| Anapc7 | -0.11088 | 0.039324 | 1 |
| Dleu2 | -0.11092 | 0.000402 | 1 |
| Camk2g | -0.11118 | 0.552373 | 1 |
| Qser1 | -0.11127 | 0.158463 | 1 |
| Bhlhe41 | -0.11135 | 0.496919 | 1 |
| Gm15635 | -0.11141 | 0.013421 | 1 |
| Bag5 | -0.11145 | 0.067363 | 1 |
| 1110051M | -0.1115 | 0.6021 | 1 |
| Tmem230 | -0.11152 | 0.018801 | 1 |
| Oxsm | -0.11153 | 0.037182 | 1 |
| Copb1 | -0.11154 | 0.015554 | 1 |
| Fbxl3 | -0.11162 | 0.063675 | 1 |
| Wdr48 | -0.11165 | 0.012457 | 1 |
| Rybp | -0.11169 | 0.001328 | 1 |
| Ldb1 | -0.11169 | 0.04141 | 1 |
| Cbr1 | -0.1117 | 0.055379 | 1 |
| Lactb | -0.11173 | 0.025128 | 1 |
| Lmf1 | -0.11189 | 0.152666 | 1 |
| 0610010K1 | -0.11193 | 0.045718 | 1 |
| Ctnnal1 | -0.11207 | 0.016423 | 1 |
| Trappc9 | -0.11207 | 0.44091 | 1 |
| Cep85l | -0.11208 | 0.271456 | 1 |
| Cyhr1 | -0.11218 | 0.020158 | 1 |
| Prkag1 | -0.1122 | 0.121863 | 1 |
| Poldip3 | -0.11235 | 0.02563 | 1 |
| Caml | -0.11243 | 0.084547 | 1 |
| Ormdl1 | -0.11244 | 0.005028 | 1 |
| Pkn1 | -0.11245 | 0.492056 | 1 |
| Cmtm3 | -0.11246 | 0.345988 | 1 |
| Apobr | -0.11251 | 0.021409 | 1 |
| Dnajc24 | -0.11255 | 0.124301 | 1 |

| | | | |
|---|---|---|---|
| Zfx | -0.11256 | 0.110454 | 1 |
| Spata13 | -0.11259 | 0.115859 | 1 |
| Jkamp | -0.11271 | 0.06702 | 1 |
| Crkl | -0.11274 | 0.095277 | 1 |
| Hypk | -0.11279 | 0.064386 | 1 |
| Cdip1 | -0.11285 | 0.073626 | 1 |
| Acadm | -0.11287 | 0.030019 | 1 |
| Smim7 | -0.11292 | 0.630079 | 1 |
| Vbp1 | -0.11306 | 0.000638 | 1 |
| Man1b1 | -0.11312 | 0.007783 | 1 |
| Narf | -0.11312 | 0.005215 | 1 |
| Nat9 | -0.11318 | 0.078721 | 1 |
| Gnpat | -0.1132 | 0.165882 | 1 |
| Armcx3 | -0.11324 | 0.050724 | 1 |
| Med17 | -0.11327 | 0.225784 | 1 |
| Ppp2r5d | -0.11329 | 0.0005 | 1 |
| Tmed5 | -0.11337 | 0.009999 | 1 |
| Itfg1 | -0.11339 | 0.049456 | 1 |
| Gramd1a | -0.1134 | 0.024201 | 1 |
| Nfkb1 | -0.11345 | 0.032337 | 1 |
| Coro1b | -0.11348 | 0.01498 | 1 |
| Tcp11l2 | -0.11371 | 1.03E-08 | 0.000334 |
| Oxld1 | -0.11376 | 0.004162 | 1 |
| Ift20 | -0.11387 | 0.049223 | 1 |
| Cdipt | -0.11391 | 0.023442 | 1 |
| Srrm2 | -0.11396 | 0.022319 | 1 |
| Rbm10 | -0.114 | 0.000329 | 1 |
| Lonp2 | -0.11401 | 0.043839 | 1 |
| Gclc | -0.11402 | 0.001098 | 1 |
| Cebpzos | -0.11406 | 0.229123 | 1 |
| 2510009E( | -0.11406 | 0.213966 | 1 |
| Ext1 | -0.11407 | 0.627701 | 1 |
| Yaf2 | -0.11409 | 0.324659 | 1 |
| Rprd2 | -0.11412 | 0.031125 | 1 |
| Usp10 | -0.11412 | 0.03962 | 1 |
| Lpin1 | -0.11413 | 0.002015 | 1 |
| Eif4h | -0.11413 | 0.01106 | 1 |
| St6galnac4 | -0.11417 | 0.063556 | 1 |
| Tmed9 | -0.11421 | 0.001209 | 1 |
| Rubcnl | -0.11429 | 0.074211 | 1 |
| Blcap | -0.1144 | 0.097631 | 1 |
| Acot13 | -0.11445 | 0.245301 | 1 |
| Zfp266 | -0.11446 | 0.090586 | 1 |
| Vma21 | -0.11451 | 0.019276 | 1 |
| Supt5 | -0.11455 | 0.085253 | 1 |
| Rhoc | -0.11458 | 0.01214 | 1 |
| Slc39a13 | -0.11458 | 0.061554 | 1 |
| Vps35l | -0.11461 | 0.137523 | 1 |

| | | | |
|---|---|---|---|
| Ppp1r7 | -0.11464 | 0.037909 | 1 |
| Vps39 | -0.11466 | 0.000324 | 1 |
| Ahcyl2 | -0.1147 | 0.116146 | 1 |
| Aip | -0.11471 | 0.343582 | 1 |
| Ccdc71 | -0.11478 | 0.003604 | 1 |
| Nosip | -0.11485 | 0.022047 | 1 |
| Pxmp4 | -0.11494 | 0.003495 | 1 |
| Ube3c | -0.11497 | 0.426788 | 1 |
| 5330438D1 | -0.11501 | 0.009505 | 1 |
| Trmt1l | -0.11503 | 0.005343 | 1 |
| Arid3b | -0.11505 | 0.342172 | 1 |
| P2rx7 | -0.11515 | 0.01504 | 1 |
| Snap47 | -0.11527 | 0.009264 | 1 |
| Rbl2 | -0.11533 | 0.000145 | 1 |
| Dnajb6 | -0.11544 | 0.040149 | 1 |
| Gnpda1 | -0.11563 | 0.059466 | 1 |
| Vps26b | -0.11564 | 0.002644 | 1 |
| Pias3 | -0.11575 | 0.004698 | 1 |
| Rassf2 | -0.11579 | 0.005322 | 1 |
| Ppox | -0.11581 | 0.043006 | 1 |
| Smyd4 | -0.11586 | 0.14547 | 1 |
| Atp6ap1 | -0.11587 | 0.016064 | 1 |
| Dnase2a | -0.11592 | 0.016562 | 1 |
| Cnot3 | -0.11594 | 0.028997 | 1 |
| Atp6v0d1 | -0.11594 | 0.135962 | 1 |
| Isca1 | -0.11599 | 0.050179 | 1 |
| Dis3l | -0.11599 | 0.135286 | 1 |
| Card9 | -0.11611 | 0.010778 | 1 |
| Dock9 | -0.11611 | 0.001672 | 1 |
| Sema4c | -0.11618 | 0.477213 | 1 |
| Armh3 | -0.11624 | 0.1718 | 1 |
| Gpat4 | -0.11627 | 0.086177 | 1 |
| Mcur1 | -0.11637 | 0.675772 | 1 |
| Arhgef12 | -0.11644 | 0.693049 | 1 |
| Pik3ca | -0.1168 | 0.210235 | 1 |
| Fam160b1 | -0.11684 | 0.007771 | 1 |
| Zfp658 | -0.11686 | 0.126193 | 1 |
| Zcchc14 | -0.11686 | 0.001344 | 1 |
| Rrp1b | -0.11687 | 0.129715 | 1 |
| Pyroxd2 | -0.11692 | 0.00015 | 1 |
| Taco1 | -0.11692 | 0.530159 | 1 |
| Cog8 | -0.11705 | 0.002996 | 1 |
| Metap2 | -0.11708 | 0.017451 | 1 |
| Pycr2 | -0.11708 | 0.152239 | 1 |
| Fam107b | -0.11712 | 2.07E-09 | 6.67E-05 |
| Glod4 | -0.1172 | 0.046779 | 1 |
| Camta2 | -0.11722 | 0.43965 | 1 |
| Adrm1 | -0.11723 | 0.225991 | 1 |

| | | | |
|---|---|---|---|
| Hs1bp3 | -0.11733 | 0.061165 | 1 |
| Cops2 | -0.11734 | 0.192837 | 1 |
| Vezf1 | -0.11735 | 0.004284 | 1 |
| Ubap1 | -0.11735 | 0.08186 | 1 |
| G6pc3 | -0.11765 | 0.016671 | 1 |
| Asxl1 | -0.11768 | 0.102665 | 1 |
| Vwa5a | -0.11769 | 0.123511 | 1 |
| Vps11 | -0.11776 | 0.060892 | 1 |
| Lrrc28 | -0.11785 | 0.003922 | 1 |
| Rhog | -0.11787 | 0.031867 | 1 |
| Gpsm1 | -0.11791 | 3.35E-06 | 0.108257 |
| Prr14 | -0.11792 | 0.018657 | 1 |
| Usf2 | -0.11795 | 0.171668 | 1 |
| Tgfbr2 | -0.11801 | 0.008984 | 1 |
| Ube2z | -0.11808 | 0.132751 | 1 |
| Naalad2 | -0.11808 | 0.092889 | 1 |
| Mlph | -0.11811 | 0.000208 | 1 |
| Lrp5 | -0.11812 | 0.29403 | 1 |
| Adck2 | -0.11816 | 0.004551 | 1 |
| Atpif1 | -0.11822 | 0.034076 | 1 |
| Tet3 | -0.11825 | 0.176181 | 1 |
| Fam120b | -0.11829 | 0.017259 | 1 |
| Btbd9 | -0.11837 | 0.460837 | 1 |
| Kmt2e | -0.11838 | 0.067533 | 1 |
| Trp53bp1 | -0.11853 | 0.231369 | 1 |
| Cd82 | -0.11857 | 0.012657 | 1 |
| Tgm2 | -0.1186 | 0.018902 | 1 |
| Ireb2 | -0.11877 | 0.343361 | 1 |
| Commd6 | -0.11879 | 0.31232 | 1 |
| Vta1 | -0.11885 | 0.064457 | 1 |
| Mettl7a1 | -0.1189 | 0.375508 | 1 |
| Fbxw5 | -0.1189 | 0.139862 | 1 |
| Miga2 | -0.11891 | 0.001075 | 1 |
| Sf3b4 | -0.11892 | 0.056183 | 1 |
| B4galt6 | -0.11908 | 0.057596 | 1 |
| Samm50 | -0.11908 | 0.063231 | 1 |
| Tusc3 | -0.11913 | 0.256825 | 1 |
| Eif4a3 | -0.1192 | 0.008236 | 1 |
| R3hdm4 | -0.11924 | 0.112161 | 1 |
| Atg4b | -0.11928 | 0.27252 | 1 |
| Cklf | -0.11931 | 0.040048 | 1 |
| Ccdc107 | -0.11946 | 0.141427 | 1 |
| Tbc1d23 | -0.1195 | 0.503487 | 1 |
| 5033421B( | -0.11954 | 0.020049 | 1 |
| Sun2 | -0.11966 | 0.634692 | 1 |
| Rbfa | -0.11967 | 0.173212 | 1 |
| Oip5os1 | -0.11985 | 0.122812 | 1 |
| Ccdc122 | -0.11986 | 0.241799 | 1 |

| | | | |
|---|---|---|---|
| Fbxo8 | -0.11986 | 0.000269 | 1 |
| Bicra | -0.1199 | 0.384865 | 1 |
| Arid1b | -0.11994 | 0.014599 | 1 |
| Tmed3 | -0.11999 | 0.050186 | 1 |
| Ppp2r3d | -0.12002 | 0.001241 | 1 |
| Xrcc5 | -0.12003 | 9.07E-06 | 0.292755 |
| Babam2 | -0.12004 | 0.132988 | 1 |
| Kmt5c | -0.12011 | 9.05E-05 | 1 |
| Gtdc1 | -0.12032 | 0.492197 | 1 |
| Rhoq | -0.12051 | 0.388564 | 1 |
| Ppp1cb | -0.12054 | 0.015585 | 1 |
| Pik3r4 | -0.12057 | 0.002069 | 1 |
| Memo1 | -0.12065 | 0.517076 | 1 |
| Spcs2 | -0.1207 | 0.003002 | 1 |
| Zfp637 | -0.12071 | 0.001954 | 1 |
| Actr2 | -0.12076 | 0.00355 | 1 |
| Card19 | -0.12076 | 0.019441 | 1 |
| Btd | -0.12082 | 0.000434 | 1 |
| Dock10 | -0.121 | 4.59E-10 | 1.48E-05 |
| Kbtbd2 | -0.12109 | 0.000509 | 1 |
| Smim14 | -0.1211 | 0.056457 | 1 |
| Gm15441 | -0.12113 | 1.02E-05 | 0.329815 |
| Borcs6 | -0.12125 | 0.001371 | 1 |
| Lgals8 | -0.12131 | 0.065184 | 1 |
| Zbtb22 | -0.12134 | 2.17E-06 | 0.070067 |
| Zfp787 | -0.12134 | 0.21305 | 1 |
| Pip4k2b | -0.12135 | 0.265101 | 1 |
| Srbd1 | -0.12139 | 0.075625 | 1 |
| Spred1 | -0.12147 | 0.00107 | 1 |
| Cyp27a1 | -0.12162 | 0.017669 | 1 |
| Rgl3 | -0.12183 | 9.48E-05 | 1 |
| Dennd6a | -0.12184 | 0.028146 | 1 |
| Glo1 | -0.12189 | 0.002039 | 1 |
| Atp11c | -0.12197 | 0.102242 | 1 |
| Pcyox1l | -0.122 | 0.000367 | 1 |
| Prrc1 | -0.122 | 0.085144 | 1 |
| Rnf150 | -0.12208 | 0.069462 | 1 |
| Tsc22d2 | -0.12208 | 0.258828 | 1 |
| Fgf13 | -0.12217 | 0.539261 | 1 |
| Ntpcr | -0.12217 | 0.000387 | 1 |
| Umad1 | -0.1222 | 0.110866 | 1 |
| Usp4 | -0.12223 | 0.230252 | 1 |
| Ppcdc | -0.1223 | 0.009685 | 1 |
| Tppp | -0.12247 | 9.95E-06 | 0.321137 |
| Nfix | -0.12251 | 0.51745 | 1 |
| Lman1 | -0.12254 | 0.012752 | 1 |
| Cdk19 | -0.12255 | 0.177304 | 1 |
| Gpatch2l | -0.1226 | 0.19818 | 1 |

| | | | |
|---|---|---|---|
| Ebi3 | -0.12266 | 0.388155 | 1 |
| Qrich1 | -0.12274 | 0.00855 | 1 |
| Mtr | -0.12283 | 0.007805 | 1 |
| Fkbp4 | -0.12284 | 0.023985 | 1 |
| Cplane1 | -0.1229 | 0.325632 | 1 |
| Pecr | -0.12307 | 0.097561 | 1 |
| Chmp1a | -0.1231 | 0.044261 | 1 |
| Tubgcp2 | -0.12316 | 0.000493 | 1 |
| Cfap97 | -0.12319 | 0.000415 | 1 |
| Nop2 | -0.12322 | 0.103025 | 1 |
| Rbbp9 | -0.12332 | 1.69E-06 | 0.054723 |
| Wbp11 | -0.12352 | 0.024524 | 1 |
| Hyou1 | -0.12354 | 0.056092 | 1 |
| Scamp3 | -0.12354 | 0.011217 | 1 |
| Jun | -0.12357 | 2.67E-05 | 0.863293 |
| Cpne1 | -0.12358 | 0.246071 | 1 |
| Btrc | -0.12361 | 0.045592 | 1 |
| Slc29a3 | -0.12362 | 0.00874 | 1 |
| Insig2 | -0.12366 | 0.011194 | 1 |
| Cldnd1 | -0.12372 | 0.158047 | 1 |
| Uba5 | -0.12394 | 0.021239 | 1 |
| Shld2 | -0.12394 | 0.005563 | 1 |
| Sept10 | -0.12396 | 0.017928 | 1 |
| Cfdp1 | -0.12406 | 0.634107 | 1 |
| Ap1g1 | -0.12411 | 0.205815 | 1 |
| Pik3c3 | -0.12416 | 0.013325 | 1 |
| Ggcx | -0.12416 | 0.000855 | 1 |
| Slco4a1 | -0.12418 | 1.32E-05 | 0.425543 |
| 0610009B2 | -0.1243 | 0.005377 | 1 |
| Sphk2 | -0.12434 | 0.038992 | 1 |
| Mgat4a | -0.12438 | 0.041869 | 1 |
| Tmx2 | -0.1244 | 0.040975 | 1 |
| Impad1 | -0.12449 | 0.113753 | 1 |
| Blmh | -0.1246 | 0.081887 | 1 |
| Bmf | -0.12462 | 0.001578 | 1 |
| Vps4a | -0.12472 | 0.03757 | 1 |
| Cdca7 | -0.12488 | 0.000285 | 1 |
| Cd200r1 | -0.12494 | 0.000752 | 1 |
| Mapre1 | -0.12499 | 0.035904 | 1 |
| Trpm4 | -0.12514 | 0.0453 | 1 |
| Vps72 | -0.12516 | 0.002497 | 1 |
| Tbc1d31 | -0.12522 | 0.051967 | 1 |
| Gspt1 | -0.12532 | 0.013683 | 1 |
| Coa3 | -0.12533 | 0.133208 | 1 |
| Fbxl17 | -0.12536 | 0.23431 | 1 |
| Dhrs7 | -0.12538 | 0.019241 | 1 |
| Usp9x | -0.12544 | 0.136796 | 1 |
| Xpot | -0.12548 | 0.000144 | 1 |

| | | | |
|---|---|---|---|
| Zfp113 | -0.12559 | 0.001514 | 1 |
| D3Ertd751 | -0.12574 | 8.75E-06 | 0.282439 |
| Gnai3 | -0.12577 | 0.018289 | 1 |
| Akr7a5 | -0.1258 | 0.023057 | 1 |
| Mmaa | -0.12586 | 0.000194 | 1 |
| Surf1 | -0.12597 | 0.022355 | 1 |
| Gm33858 | -0.12602 | 0.003366 | 1 |
| Pcf11 | -0.12614 | 0.326181 | 1 |
| Cracr2b | -0.12617 | 0.000363 | 1 |
| Dcaf8 | -0.12618 | 0.012952 | 1 |
| Stau1 | -0.12619 | 0.238945 | 1 |
| Ikbip | -0.12628 | 0.00428 | 1 |
| Wwp1 | -0.12632 | 0.020552 | 1 |
| Naa10 | -0.12632 | 0.021349 | 1 |
| Tfcp2 | -0.12641 | 0.012279 | 1 |
| AW146154 | -0.12654 | 0.000309 | 1 |
| Tssc4 | -0.12658 | 0.0025 | 1 |
| Trim5 | -0.12678 | 0.001275 | 1 |
| Xiap | -0.12681 | 0.004531 | 1 |
| Naxe | -0.12687 | 0.035905 | 1 |
| Gm15523 | -0.12693 | 0.000487 | 1 |
| Pak1 | -0.12705 | 0.225589 | 1 |
| Pdcl | -0.12712 | 0.000141 | 1 |
| March6 | -0.12714 | 0.010171 | 1 |
| Cdk13 | -0.12716 | 0.02667 | 1 |
| Tmx3 | -0.12733 | 0.009727 | 1 |
| Tob2 | -0.12751 | 0.016208 | 1 |
| Wbp1 | -0.12756 | 0.016259 | 1 |
| Tmx4 | -0.12765 | 8.98E-07 | 0.028988 |
| Map4k2 | -0.12772 | 0.001947 | 1 |
| Tmem245 | -0.12778 | 0.022783 | 1 |
| Topors | -0.1279 | 0.0567 | 1 |
| B3galnt1 | -0.12794 | 0.326766 | 1 |
| Tmem198b | -0.12798 | 0.000246 | 1 |
| Banp | -0.12802 | 0.022208 | 1 |
| Gne | -0.12802 | 0.000161 | 1 |
| Ndufc2 | -0.12803 | 0.006689 | 1 |
| Wdpcp | -0.12805 | 0.872901 | 1 |
| Brd8 | -0.12808 | 0.100835 | 1 |
| Phf21a | -0.12816 | 0.045248 | 1 |
| Mars | -0.12819 | 0.031867 | 1 |
| Pex6 | -0.12832 | 5.38E-05 | 1 |
| Gse1 | -0.12838 | 0.186814 | 1 |
| Tdp1 | -0.12838 | 0.068736 | 1 |
| Emc10 | -0.1284 | 0.001692 | 1 |
| Ahdc1 | -0.12845 | 0.014145 | 1 |
| Zcchc8 | -0.12858 | 0.030006 | 1 |
| Rhot1 | -0.1286 | 0.082529 | 1 |

| | | | |
|---|---|---|---|
| Cdk8 | -0.12874 | 1.35E-05 | 0.436492 |
| Tmem120a | -0.12875 | 0.087594 | 1 |
| Arsk | -0.12877 | 0.001798 | 1 |
| Ccnq | -0.12883 | 0.00226 | 1 |
| Pou2f1 | -0.12888 | 0.016617 | 1 |
| Dse | -0.12893 | 0.004089 | 1 |
| Prmt2 | -0.12894 | 0.027849 | 1 |
| Rabgap1 | -0.12901 | 0.024628 | 1 |
| Tsn | -0.12909 | 0.004958 | 1 |
| Yipf3 | -0.12915 | 0.016003 | 1 |
| Copa | -0.12944 | 8.02E-05 | 1 |
| Uba6 | -0.12944 | 0.003619 | 1 |
| Arhgdia | -0.12947 | 0.003919 | 1 |
| Pias1 | -0.12961 | 0.000258 | 1 |
| Smim27 | -0.12965 | 0.000356 | 1 |
| Gm26520 | -0.12969 | 0.019808 | 1 |
| Ergic3 | -0.1297 | 0.013839 | 1 |
| Zfp397 | -0.12978 | 0.022682 | 1 |
| Ap1b1 | -0.12979 | 0.010538 | 1 |
| Nlk | -0.12987 | 0.127357 | 1 |
| Pcbp1 | -0.12998 | 0.044643 | 1 |
| Exosc10 | -0.13 | 0.141439 | 1 |
| Bcdin3d | -0.13019 | 0.003112 | 1 |
| Yae1d1 | -0.13023 | 0.001754 | 1 |
| Wdr20 | -0.13028 | 0.273364 | 1 |
| Ogg1 | -0.13028 | 0.000792 | 1 |
| Gpatch8 | -0.13034 | 0.861454 | 1 |
| Slc39a7 | -0.13034 | 0.01468 | 1 |
| Sgk3 | -0.13044 | 0.151047 | 1 |
| Rgl1 | -0.13046 | 0.013002 | 1 |
| Gm38843 | -0.13054 | 0.012517 | 1 |
| Daglb | -0.13064 | 0.000261 | 1 |
| Epha2 | -0.13069 | 0.011931 | 1 |
| Yes1 | -0.1307 | 0.007937 | 1 |
| Hspa13 | -0.13075 | 0.026348 | 1 |
| Bcl2l13 | -0.131 | 0.000445 | 1 |
| Ctdsp2 | -0.131 | 2.66E-05 | 0.858724 |
| Clcf1 | -0.13102 | 0.000197 | 1 |
| Zmynd11 | -0.13104 | 0.019446 | 1 |
| Zfp157 | -0.13117 | 0.005131 | 1 |
| Hcls1 | -0.13122 | 0.038266 | 1 |
| Tnks | -0.13124 | 0.688229 | 1 |
| Snx30 | -0.13125 | 0.351041 | 1 |
| Pag1 | -0.13126 | 1.85E-05 | 0.597255 |
| Phkb | -0.13135 | 0.104374 | 1 |
| Cpsf1 | -0.13143 | 0.021045 | 1 |
| Brk1 | -0.13146 | 0.0299 | 1 |
| Glrp1 | -0.1315 | 0.004151 | 1 |

| | | | |
|---|---|---|---|
| Gm29488 | -0.13153 | 0.001524 | 1 |
| Gm15503 | -0.13163 | 0.003183 | 1 |
| Gm16023 | -0.13164 | 4.41E-05 | 1 |
| Ralbp1 | -0.13169 | 0.17081 | 1 |
| Phf12 | -0.1317 | 0.252344 | 1 |
| Lrrc39 | -0.1319 | 1.72E-08 | 0.000556 |
| Aldh4a1 | -0.1321 | 0.000129 | 1 |
| Tmed7 | -0.13213 | 0.027822 | 1 |
| Morc2a | -0.13214 | 0.011201 | 1 |
| Spsb1 | -0.13221 | 0.001445 | 1 |
| Rubcn | -0.13222 | 0.014365 | 1 |
| Smurf1 | -0.13222 | 0.247673 | 1 |
| Unc93b1 | -0.13236 | 1.29E-10 | 4.15E-06 |
| Hmgxb4 | -0.13238 | 0.031759 | 1 |
| Tpk1 | -0.13241 | 0.035053 | 1 |
| Ppp4r3a | -0.13242 | 0.395283 | 1 |
| Pigk | -0.13258 | 0.085791 | 1 |
| Pcif1 | -0.13263 | 0.025456 | 1 |
| Zbtb7a | -0.13272 | 0.0131 | 1 |
| Ccdc97 | -0.13272 | 7.85E-05 | 1 |
| Nsmce1 | -0.13279 | 0.016218 | 1 |
| Mplkip | -0.1328 | 0.017169 | 1 |
| H2-Ke6 | -0.1328 | 0.003591 | 1 |
| Brd3 | -0.13282 | 0.000593 | 1 |
| Abi2 | -0.13283 | 0.000414 | 1 |
| Arfgap2 | -0.13286 | 0.014737 | 1 |
| Jpx | -0.13294 | 0.002147 | 1 |
| Ube2m | -0.13294 | 0.026604 | 1 |
| Madd | -0.13302 | 0.003387 | 1 |
| Ddrgk1 | -0.13317 | 0.085049 | 1 |
| Kif13a | -0.13317 | 0.239685 | 1 |
| Scaper | -0.13318 | 0.146381 | 1 |
| Vav1 | -0.13324 | 0.007193 | 1 |
| AI467606 | -0.13329 | 0.01912 | 1 |
| Map3k20 | -0.13333 | 0.004742 | 1 |
| Fes | -0.13338 | 0.049936 | 1 |
| Ift52 | -0.13351 | 0.098647 | 1 |
| Ppp2r5c | -0.13352 | 0.274937 | 1 |
| Haghl | -0.13365 | 0.002705 | 1 |
| Ank | -0.13375 | 0.001665 | 1 |
| Luc7l2 | -0.13375 | 0.002864 | 1 |
| Tada3 | -0.13399 | 0.001517 | 1 |
| Wdr45b | -0.134 | 0.01272 | 1 |
| Fkbp15 | -0.13407 | 0.019483 | 1 |
| Slain2 | -0.13413 | 0.016367 | 1 |
| Mdc1 | -0.13418 | 0.001124 | 1 |
| Epc1 | -0.13437 | 0.000319 | 1 |
| Scrn3 | -0.13438 | 0.002822 | 1 |

| | | | |
|---|---|---|---|
| Oxct1 | -0.13446 | 0.001421 | 1 |
| Sfi1 | -0.13447 | 0.007352 | 1 |
| Dennd1c | -0.1345 | 0.036878 | 1 |
| Cdk2ap2 | -0.13464 | 0.018442 | 1 |
| Foxp1 | -0.13466 | 0.008115 | 1 |
| Txnl1 | -0.13467 | 0.004213 | 1 |
| Zkscan3 | -0.13469 | 0.010707 | 1 |
| Bptf | -0.13471 | 0.076796 | 1 |
| Mafg | -0.13471 | 0.087468 | 1 |
| Dhx32 | -0.13481 | 0.000932 | 1 |
| Fam210a | -0.13481 | 0.003087 | 1 |
| Tbc1d16 | -0.13489 | 0.022603 | 1 |
| Calcoco1 | -0.1349 | 7.48E-05 | 1 |
| Fam168a | -0.13493 | 0.256677 | 1 |
| Tbcc | -0.13497 | 1.86E-05 | 0.601168 |
| Aldh9a1 | -0.13511 | 0.009057 | 1 |
| Timm17b | -0.13512 | 0.0082 | 1 |
| Sdha | -0.13514 | 0.023524 | 1 |
| Stradb | -0.13515 | 3.49E-07 | 0.011283 |
| Tial1 | -0.13532 | 0.072638 | 1 |
| Tle4 | -0.13533 | 0.008999 | 1 |
| Dusp16 | -0.13556 | 0.076827 | 1 |
| Mbtps2 | -0.13572 | 0.098722 | 1 |
| Lrwd1 | -0.13573 | 4.31E-05 | 1 |
| Dhx8 | -0.13577 | 0.127723 | 1 |
| Trim8 | -0.13585 | 0.001871 | 1 |
| Ppp2r1b | -0.13589 | 0.003008 | 1 |
| Cd3eap | -0.13591 | 0.060857 | 1 |
| Pdlim5 | -0.13593 | 0.010581 | 1 |
| Kdm4a | -0.13604 | 0.007224 | 1 |
| Dtx3 | -0.13611 | 0.05321 | 1 |
| Tmub2 | -0.13613 | 0.019562 | 1 |
| Pot1b | -0.13622 | 0.045063 | 1 |
| Arap1 | -0.13626 | 0.110404 | 1 |
| D6Wsu163 | -0.13645 | 0.016117 | 1 |
| Nucks1 | -0.13648 | 0.006315 | 1 |
| Mta2 | -0.13652 | 0.026253 | 1 |
| Gm41071 | -0.13655 | 0.000228 | 1 |
| Gtf3c4 | -0.13659 | 1.19E-05 | 0.384866 |
| Parn | -0.1367 | 0.018831 | 1 |
| Ttyh3 | -0.1367 | 0.002006 | 1 |
| Tnfaip8l2 | -0.13678 | 0.000629 | 1 |
| Tsc2 | -0.1368 | 0.035208 | 1 |
| Gm16124 | -0.13682 | 0.00148 | 1 |
| Ss18 | -0.13685 | 0.000954 | 1 |
| Csnk2a1 | -0.13695 | 0.024009 | 1 |
| Ppp2r1a | -0.13696 | 0.000751 | 1 |
| Prkcb | -0.13697 | 0.700786 | 1 |

| | | | |
|---|---|---|---|
| Zfp646 | -0.13697 | 3.29E-05 | 1 |
| Fgd3 | -0.13703 | 0.243495 | 1 |
| Sipa1l2 | -0.13718 | 0.52231 | 1 |
| Cab39 | -0.13753 | 0.012077 | 1 |
| Pdzd8 | -0.13759 | 0.010932 | 1 |
| Wwc2 | -0.13762 | 0.134888 | 1 |
| Dok1 | -0.13762 | 0.041258 | 1 |
| Ccng1 | -0.13762 | 0.015356 | 1 |
| Nmt1 | -0.13765 | 0.007651 | 1 |
| Vps13d | -0.138 | 0.827197 | 1 |
| D630023F1 | -0.13803 | 9.37E-05 | 1 |
| Cdc123 | -0.13804 | 0.003099 | 1 |
| Ddb1 | -0.1381 | 0.000839 | 1 |
| Msl2 | -0.1381 | 0.009143 | 1 |
| Ywhaq | -0.1382 | 0.002134 | 1 |
| Ier5 | -0.13844 | 8.12E-06 | 0.262161 |
| Appbp2 | -0.13844 | 0.026875 | 1 |
| Akt1 | -0.13873 | 0.006705 | 1 |
| Anapc13 | -0.13875 | 0.09276 | 1 |
| Rab5b | -0.13886 | 0.0026 | 1 |
| Mpv17 | -0.13897 | 0.000497 | 1 |
| Osgep | -0.13898 | 0.017679 | 1 |
| Sema4g | -0.1391 | 6.19E-08 | 0.001998 |
| Lpxn | -0.13912 | 0.000775 | 1 |
| Mcrs1 | -0.1392 | 0.004391 | 1 |
| Cib2 | -0.13921 | 1.72E-06 | 0.055445 |
| Aldh3a2 | -0.13924 | 3.42E-05 | 1 |
| Tmx1 | -0.13935 | 0.000314 | 1 |
| Slc35b2 | -0.13947 | 0.001233 | 1 |
| Insr | -0.13948 | 0.092153 | 1 |
| Gorasp2 | -0.13969 | 0.012754 | 1 |
| Mon2 | -0.13976 | 0.0214 | 1 |
| Slc50a1 | -0.13981 | 0.026366 | 1 |
| Pabpc4 | -0.13985 | 0.00318 | 1 |
| Ilk | -0.14008 | 0.001477 | 1 |
| Gpx4 | -0.14012 | 0.016313 | 1 |
| Zbtb1 | -0.14012 | 0.022522 | 1 |
| Lsm10 | -0.14032 | 0.034107 | 1 |
| Pcnx3 | -0.14044 | 1.2E-05 | 0.387024 |
| St3gal2 | -0.14045 | 0.002711 | 1 |
| Ech1 | -0.14057 | 0.002277 | 1 |
| Champ1 | -0.14057 | 0.024763 | 1 |
| Mpp7 | -0.1407 | 0.003207 | 1 |
| Zfp426 | -0.14071 | 0.018895 | 1 |
| 5031439G0 | -0.14071 | 0.000674 | 1 |
| Mgat1 | -0.14098 | 0.010494 | 1 |
| Smim15 | -0.14114 | 0.000653 | 1 |
| Zfp90 | -0.14119 | 0.003364 | 1 |

| | | | |
|---|---|---|---|
| Dcaf17 | -0.14124 | 0.035312 | 1 |
| Sh3bp1 | -0.14153 | 0.156541 | 1 |
| Rb1 | -0.14157 | 0.408844 | 1 |
| Vps4b | -0.14161 | 0.005574 | 1 |
| Isoc2b | -0.14174 | 4.49E-05 | 1 |
| Sec31a | -0.14179 | 0.005444 | 1 |
| Actr1a | -0.1418 | 0.00312 | 1 |
| Zkscan8 | -0.14182 | 0.002906 | 1 |
| Commd3 | -0.14219 | 0.015607 | 1 |
| Gm16867 | -0.14237 | 5.34E-07 | 0.017241 |
| Plcb3 | -0.14239 | 0.00029 | 1 |
| Erh | -0.14251 | 0.007937 | 1 |
| D130009I1 | -0.14262 | 3.29E-08 | 0.001062 |
| Ier3ip1 | -0.14264 | 0.026415 | 1 |
| Bckdk | -0.14265 | 0.003436 | 1 |
| Ptpa | -0.14265 | 0.013419 | 1 |
| Zkscan17 | -0.14269 | 0.006206 | 1 |
| Srek1ip1 | -0.14272 | 0.128989 | 1 |
| Thap11 | -0.14277 | 9.1E-05 | 1 |
| Ndfip2 | -0.14282 | 0.001492 | 1 |
| Mob4 | -0.14304 | 0.009797 | 1 |
| Actr8 | -0.14356 | 0.000835 | 1 |
| Mfsd11 | -0.14358 | 6.78E-05 | 1 |
| Sec13 | -0.14359 | 0.016696 | 1 |
| Nbeal2 | -0.14376 | 2.42E-09 | 7.8E-05 |
| Atf6b | -0.14376 | 0.032073 | 1 |
| Ngrn | -0.14381 | 1.32E-05 | 0.426572 |
| Uba2 | -0.14399 | 0.008697 | 1 |
| Arhgap32 | -0.14401 | 0.000699 | 1 |
| Zfp110 | -0.14405 | 0.000165 | 1 |
| Parp1 | -0.14413 | 0.007333 | 1 |
| Calcrl | -0.1442 | 9.44E-05 | 1 |
| Puf60 | -0.14432 | 0.001994 | 1 |
| Plau | -0.14434 | 0.000202 | 1 |
| Agbl3 | -0.14437 | 1.37E-07 | 0.004416 |
| Ankrd44 | -0.14442 | 3.78E-10 | 1.22E-05 |
| Rnf44 | -0.14474 | 0.002422 | 1 |
| Csk | -0.14476 | 0.018102 | 1 |
| Stxbp3 | -0.14477 | 0.046142 | 1 |
| Snapin | -0.14488 | 0.003192 | 1 |
| Nucb1 | -0.14489 | 0.033423 | 1 |
| AW549877 | -0.14496 | 0.012501 | 1 |
| Capzb | -0.14502 | 4.34E-05 | 1 |
| Khdrbs1 | -0.14514 | 0.003079 | 1 |
| Tlr9 | -0.14537 | 0.149034 | 1 |
| Ubxn7 | -0.14541 | 0.007908 | 1 |
| Rin3 | -0.14544 | 0.001617 | 1 |
| 1110019D | -0.1456 | 6.47E-05 | 1 |

| | | | |
|---|---|---|---|
| Chmp3 | -0.14575 | 0.007106 | 1 |
| Cyp4f13 | -0.14595 | 0.017828 | 1 |
| St13 | -0.14615 | 0.01862 | 1 |
| Gm15489 | -0.1462 | 3.16E-05 | 1 |
| Hnrnpr | -0.14621 | 0.00279 | 1 |
| Ralgapb | -0.14622 | 0.005667 | 1 |
| Mthfs | -0.14625 | 0.02523 | 1 |
| Chm | -0.14625 | 0.008071 | 1 |
| Zfp518a | -0.14626 | 0.001445 | 1 |
| Plekhm3 | -0.14629 | 0.013125 | 1 |
| Chd6 | -0.1463 | 0.136531 | 1 |
| Golga2 | -0.14631 | 0.003067 | 1 |
| Lima1 | -0.14632 | 0.001685 | 1 |
| Arglu1 | -0.14632 | 0.285057 | 1 |
| Secisbp2l | -0.14645 | 0.021735 | 1 |
| Dstn | -0.14678 | 0.000432 | 1 |
| Kdm5b | -0.14678 | 1.76E-05 | 0.568283 |
| Sfswap | -0.14693 | 0.569884 | 1 |
| Brat1 | -0.14704 | 0.00322 | 1 |
| Tax1bp3 | -0.14724 | 0.001305 | 1 |
| Pnisr | -0.14732 | 0.15268 | 1 |
| Sp2 | -0.14739 | 0.0002 | 1 |
| Tns3 | -0.14758 | 0.054788 | 1 |
| Itgav | -0.14763 | 0.000232 | 1 |
| Lrrc61 | -0.1477 | 2.41E-07 | 0.007787 |
| Rsbn1 | -0.1477 | 0.005978 | 1 |
| Rasal2 | -0.14773 | 0.138779 | 1 |
| Mtfr1l | -0.1479 | 0.001099 | 1 |
| 2310009A( | -0.14792 | 0.003797 | 1 |
| Timm21 | -0.14795 | 1.28E-05 | 0.413129 |
| Oard1 | -0.14803 | 0.000139 | 1 |
| Dipk1a | -0.14804 | 2.83E-07 | 0.009124 |
| 5830432E( | -0.14808 | 3.28E-05 | 1 |
| Nbeal1 | -0.14811 | 0.002281 | 1 |
| Srp68 | -0.14816 | 0.016276 | 1 |
| Pgrmc1 | -0.14817 | 1.25E-05 | 0.40246 |
| Atf7ip | -0.1485 | 0.000949 | 1 |
| Hacd3 | -0.14856 | 0.002571 | 1 |
| Rab3gap1 | -0.14862 | 0.013711 | 1 |
| Thap3 | -0.14862 | 0.001143 | 1 |
| Unc50 | -0.14888 | 0.00094 | 1 |
| Rai1 | -0.14894 | 0.363351 | 1 |
| Ano10 | -0.14896 | 0.154291 | 1 |
| Gbf1 | -0.14897 | 0.155409 | 1 |
| Map2k6 | -0.14898 | 0.000811 | 1 |
| Was | -0.14904 | 0.002521 | 1 |
| Lamtor1 | -0.14914 | 0.000996 | 1 |
| Paqr7 | -0.14935 | 0.023639 | 1 |

| | | | |
|---|---|---|---|
| Naxd | -0.14941 | 0.001818 | 1 |
| 5830444B( | -0.14943 | 2.54E-06 | 0.082075 |
| H3f3b | -0.14945 | 3.3E-07 | 0.010663 |
| Cyb5d2 | -0.1496 | 7.45E-06 | 0.240675 |
| Gmfg | -0.14985 | 0.201336 | 1 |
| Amd1 | -0.15011 | 2.97E-06 | 0.095951 |
| Enox2 | -0.15016 | 0.314254 | 1 |
| Zfp579 | -0.15034 | 4.24E-05 | 1 |
| Pxk | -0.15041 | 0.003692 | 1 |
| Adnp | -0.15048 | 0.223931 | 1 |
| 2610008E1 | -0.15052 | 5.83E-06 | 0.188359 |
| Sptssa | -0.15068 | 0.00097 | 1 |
| Capns1 | -0.15074 | 0.007642 | 1 |
| 4930481A1 | -0.15089 | 2.42E-08 | 0.000781 |
| Polr2a | -0.15095 | 0.001852 | 1 |
| Ube4a | -0.15097 | 0.075252 | 1 |
| Engase | -0.15099 | 4.27E-07 | 0.013777 |
| Usp19 | -0.15102 | 0.005657 | 1 |
| Laptm4b | -0.15121 | 0.010424 | 1 |
| Degs1 | -0.15127 | 0.001652 | 1 |
| Trf | -0.15143 | 2.03E-07 | 0.006555 |
| Stard9 | -0.15155 | 0.024112 | 1 |
| Exoc7 | -0.15162 | 0.012046 | 1 |
| Snx5 | -0.15169 | 2.99E-06 | 0.096669 |
| Hist1h2bc | -0.15193 | 0.005621 | 1 |
| Zmym2 | -0.15199 | 0.026399 | 1 |
| Pja1 | -0.152 | 8.52E-06 | 0.275135 |
| Cd79b | -0.15228 | 0.003769 | 1 |
| Otud7b | -0.15234 | 0.081099 | 1 |
| Immp1l | -0.15242 | 0.091049 | 1 |
| Ccdc106 | -0.15253 | 1.35E-05 | 0.436735 |
| Mmgt2 | -0.15254 | 5.51E-05 | 1 |
| Cybc1 | -0.15266 | 0.001568 | 1 |
| Abi1 | -0.15272 | 0.00439 | 1 |
| Dyrk1a | -0.1528 | 0.067517 | 1 |
| Gpsm3 | -0.15289 | 0.000497 | 1 |
| Abcc1 | -0.15298 | 0.001325 | 1 |
| Zfp606 | -0.15306 | 8.83E-07 | 0.028519 |
| Mpc1 | -0.15317 | 0.000493 | 1 |
| Ncbp3 | -0.15318 | 0.003013 | 1 |
| Gmeb1 | -0.15339 | 0.015405 | 1 |
| Naa60 | -0.15341 | 0.00666 | 1 |
| Ccni | -0.15343 | 0.001856 | 1 |
| Rad23a | -0.15345 | 0.002024 | 1 |
| Dctn1 | -0.15346 | 2.54E-05 | 0.819084 |
| 4930517O1 | -0.15353 | 0.001691 | 1 |
| Nek7 | -0.15353 | 0.008751 | 1 |
| Frg1 | -0.15365 | 0.004845 | 1 |

| | | | |
|---|---|---|---|
| Ssbp2 | -0.15368 | 0.153384 | 1 |
| Cdkn1b | -0.15371 | 0.000176 | 1 |
| A630001G: | -0.15371 | 0.013291 | 1 |
| Padi2 | -0.15377 | 7.02E-05 | 1 |
| Nsfl1c | -0.15378 | 0.033194 | 1 |
| Ampd3 | -0.15402 | 5.07E-06 | 0.163843 |
| Bcl7a | -0.15407 | 1.52E-06 | 0.049203 |
| 2010315B( | -0.15419 | 0.001887 | 1 |
| Cic | -0.15432 | 0.000301 | 1 |
| Tmem35b | -0.15434 | 3.84E-05 | 1 |
| Top2b | -0.15443 | 0.006857 | 1 |
| Cc2d1b | -0.15474 | 0.000189 | 1 |
| Lrmp | -0.15504 | 0.00278 | 1 |
| Hbp1 | -0.1552 | 0.000633 | 1 |
| Rcbtb2 | -0.15521 | 0.000601 | 1 |
| Map3k1 | -0.15525 | 0.000104 | 1 |
| Cnnm2 | -0.15541 | 7.56E-08 | 0.002441 |
| Arsa | -0.15543 | 0.001943 | 1 |
| Hjurp | -0.1556 | 0.002372 | 1 |
| Lamtor4 | -0.15572 | 0.002789 | 1 |
| Pigv | -0.15577 | 2.05E-05 | 0.662899 |
| AW554918 | -0.15607 | 0.013635 | 1 |
| Yy1 | -0.1561 | 0.029947 | 1 |
| Plekha2 | -0.15611 | 0.001617 | 1 |
| Ccdc192 | -0.1562 | 2.85E-05 | 0.921383 |
| Mat2b | -0.15622 | 0.003934 | 1 |
| Cdc26 | -0.15629 | 0.113407 | 1 |
| Trim35 | -0.15661 | 0.000314 | 1 |
| Atg14 | -0.15689 | 1.93E-06 | 0.062167 |
| Hsd17b4 | -0.15696 | 0.001875 | 1 |
| Dcaf7 | -0.15697 | 0.024181 | 1 |
| Cnnm3 | -0.15698 | 2.04E-05 | 0.657519 |
| Kcnk13 | -0.15708 | 0.001795 | 1 |
| Ufl1 | -0.15722 | 0.012598 | 1 |
| Bbip1 | -0.15723 | 0.012128 | 1 |
| Tmlhe | -0.15727 | 0.001462 | 1 |
| Gmpr2 | -0.15727 | 0.020404 | 1 |
| Pak2 | -0.15734 | 3.29E-06 | 0.106153 |
| Rad21 | -0.15736 | 7.08E-05 | 1 |
| Sh3glb1 | -0.15736 | 0.000588 | 1 |
| Maml1 | -0.15749 | 0.000345 | 1 |
| Prex1 | -0.15762 | 0.002399 | 1 |
| Ophn1 | -0.15763 | 0.001328 | 1 |
| Chmp1b | -0.1577 | 9.06E-06 | 0.292477 |
| Edem1 | -0.15773 | 0.002696 | 1 |
| Fkbp9 | -0.15779 | 1.69E-05 | 0.545321 |
| Erlin2 | -0.15803 | 0.013037 | 1 |
| Calhm2 | -0.15806 | 0.001034 | 1 |

| | | | |
|---|---|---|---|
| Bco2 | -0.15808 | 0.002802 | 1 |
| Trmt112 | -0.15829 | 0.016213 | 1 |
| Hnrnph2 | -0.15842 | 0.013755 | 1 |
| Zc3h12b | -0.15862 | 0.000511 | 1 |
| Mapkap1 | -0.15867 | 0.001384 | 1 |
| Nfat5 | -0.15877 | 0.012566 | 1 |
| 2810006K2 | -0.15878 | 0.001159 | 1 |
| Eml4 | -0.15903 | 0.076118 | 1 |
| Letm2 | -0.15931 | 4.42E-06 | 0.142651 |
| Il15 | -0.15973 | 0.000358 | 1 |
| 4931406C( | -0.15986 | 0.002997 | 1 |
| Stk3 | -0.15995 | 0.150065 | 1 |
| Phgdh | -0.16004 | 0.01499 | 1 |
| Hist1h4d | -0.16016 | 9.83E-05 | 1 |
| Ano6 | -0.16019 | 0.000923 | 1 |
| Dcxr | -0.16022 | 0.003549 | 1 |
| Zfp329 | -0.16025 | 0.000112 | 1 |
| Gid8 | -0.16027 | 0.000843 | 1 |
| Xpo7 | -0.1603 | 0.025798 | 1 |
| Ctnna1 | -0.16032 | 0.002053 | 1 |
| Slc26a2 | -0.16039 | 5.49E-05 | 1 |
| Trim65 | -0.16062 | 3.97E-05 | 1 |
| 4732471JC | -0.16071 | 0.005163 | 1 |
| Plxdc1 | -0.16097 | 0.069164 | 1 |
| Lilra5 | -0.1611 | 0.003258 | 1 |
| Rab5a | -0.16113 | 0.13399 | 1 |
| Lhfpl2 | -0.16121 | 0.017659 | 1 |
| Smim10l1 | -0.16127 | 0.0103 | 1 |
| Ptgr2 | -0.16128 | 0.000423 | 1 |
| Mfsd14b | -0.16131 | 0.00035 | 1 |
| Vxn | -0.16154 | 2.72E-08 | 0.00088 |
| Riok2 | -0.16169 | 0.026107 | 1 |
| Fundc1 | -0.16172 | 0.000181 | 1 |
| Smad5 | -0.16174 | 3.04E-06 | 0.098131 |
| Mecp2 | -0.16178 | 0.043483 | 1 |
| Slc38a9 | -0.16183 | 0.001764 | 1 |
| Gm14455 | -0.16184 | 2.09E-11 | 6.74E-07 |
| Ints10 | -0.16203 | 0.001683 | 1 |
| Clpx | -0.16213 | 0.001108 | 1 |
| Polr3gl | -0.16224 | 4.54E-06 | 0.146512 |
| Sort1 | -0.16234 | 0.000346 | 1 |
| Cpsf3 | -0.16247 | 0.008099 | 1 |
| H2afv | -0.16253 | 0.000889 | 1 |
| Atp6v1c1 | -0.16253 | 0.007444 | 1 |
| Dctn2 | -0.16257 | 0.00704 | 1 |
| Bax | -0.16262 | 0.01093 | 1 |
| Smo | -0.16263 | 0.001175 | 1 |
| Fem1c | -0.1629 | 2.65E-05 | 0.854773 |

| | | | |
|---|---|---|---|
| Pnkd | -0.16305 | 0.001006 | 1 |
| Tmem33 | -0.1636 | 0.02365 | 1 |
| Dennd5a | -0.16376 | 0.062639 | 1 |
| Mff | -0.16377 | 0.005623 | 1 |
| Bcl7b | -0.16378 | 0.007409 | 1 |
| Fbxl4 | -0.16389 | 0.003765 | 1 |
| Tln1 | -0.16408 | 0.00818 | 1 |
| Gmip | -0.16413 | 0.000777 | 1 |
| Dguok | -0.16427 | 0.000819 | 1 |
| Gm47754 | -0.1645 | 2.6E-05 | 0.838974 |
| Cebpa | -0.16451 | 0.000543 | 1 |
| Rdx | -0.16452 | 0.002613 | 1 |
| Tia1 | -0.1647 | 0.00104 | 1 |
| Dnajc18 | -0.16476 | 0.00089 | 1 |
| Spcs3 | -0.16483 | 0.000411 | 1 |
| Cebpd | -0.1651 | 2.91E-06 | 0.094014 |
| Agps | -0.16517 | 0.009169 | 1 |
| 1110004F1 | -0.16526 | 0.006534 | 1 |
| Chtf8 | -0.16565 | 5.84E-05 | 1 |
| 6330418K( | -0.16573 | 3.73E-09 | 0.00012 |
| Tcea1 | -0.16584 | 0.002046 | 1 |
| Fnip2 | -0.16593 | 0.069505 | 1 |
| Cox20 | -0.16605 | 0.000313 | 1 |
| Cox16 | -0.16606 | 0.003713 | 1 |
| Camsap1 | -0.16609 | 0.000981 | 1 |
| Ptpn6 | -0.16627 | 7.31E-06 | 0.235856 |
| Bcl9 | -0.16631 | 0.001829 | 1 |
| Tbc1d17 | -0.16645 | 9.65E-06 | 0.311634 |
| Map3k11 | -0.16648 | 0.000229 | 1 |
| Dock11 | -0.16652 | 0.024007 | 1 |
| Ccdc88b | -0.16655 | 0.00029 | 1 |
| Amfr | -0.16665 | 0.000113 | 1 |
| Dazap2 | -0.1667 | 0.001768 | 1 |
| Fam167b | -0.1668 | 0.001275 | 1 |
| Cd300c2 | -0.16686 | 1.77E-06 | 0.057088 |
| Patz1 | -0.16706 | 0.000354 | 1 |
| Cdkal1 | -0.16708 | 0.152709 | 1 |
| Fzd4 | -0.16708 | 2.3E-08 | 0.000744 |
| Gm37401 | -0.16708 | 1.54E-07 | 0.004963 |
| Asb3 | -0.16714 | 0.001555 | 1 |
| Ulk4 | -0.16723 | 0.001119 | 1 |
| Tle3 | -0.16725 | 6.9E-05 | 1 |
| Abhd17c | -0.1673 | 0.000323 | 1 |
| Ankfy1 | -0.16749 | 0.007323 | 1 |
| Czib | -0.16755 | 0.001651 | 1 |
| Spr | -0.16769 | 5.58E-05 | 1 |
| Cnep1r1 | -0.16776 | 8.37E-06 | 0.270359 |
| Ccl9 | -0.16776 | 0.030923 | 1 |

| | | | |
|---|---|---|---|
| Abhd4 | -0.16789 | 0.000105 | 1 |
| Matr3 | -0.16792 | 0.000156 | 1 |
| Tulp4 | -0.16802 | 0.003664 | 1 |
| Stxbp5 | -0.16803 | 0.010212 | 1 |
| Jak1 | -0.16806 | 1.29E-07 | 0.004173 |
| Tgoln1 | -0.16808 | 0.007445 | 1 |
| Skp1a | -0.16808 | 1.98E-05 | 0.640564 |
| Gsk3b | -0.16815 | 1.17E-05 | 0.37902 |
| Gm28417 | -0.16821 | 0.000426 | 1 |
| Letmd1 | -0.16822 | 0.000203 | 1 |
| Mepce | -0.16824 | 2.74E-05 | 0.883814 |
| Cacna1a | -0.16839 | 0.014938 | 1 |
| Slf2 | -0.16841 | 0.001962 | 1 |
| Mtmr3 | -0.1685 | 0.005083 | 1 |
| Rxylt1 | -0.16863 | 0.000252 | 1 |
| Asap3 | -0.16863 | 0.001857 | 1 |
| 1700012D1 | -0.16875 | 7.81E-06 | 0.252004 |
| Cep112 | -0.16878 | 0.083787 | 1 |
| Vcp | -0.16887 | 0.003009 | 1 |
| Zfp148 | -0.16898 | 0.011438 | 1 |
| Rbm33 | -0.16931 | 0.01192 | 1 |
| Kmt2c | -0.16941 | 0.000131 | 1 |
| B4galt4 | -0.1695 | 0.004019 | 1 |
| Sfmbt1 | -0.16956 | 0.056079 | 1 |
| St8sia4 | -0.16964 | 3.33E-10 | 1.07E-05 |
| Camk2n1 | -0.1697 | 0.001284 | 1 |
| Tmem60 | -0.16978 | 0.000506 | 1 |
| Washc4 | -0.16991 | 0.000161 | 1 |
| Taok1 | -0.16993 | 0.001916 | 1 |
| Gm13710 | -0.16994 | 0.001096 | 1 |
| Bnip3l | -0.17024 | 0.000223 | 1 |
| Hs6st1 | -0.17036 | 1.17E-05 | 0.378015 |
| Dido1 | -0.17074 | 0.000245 | 1 |
| Pon3 | -0.17074 | 0.000759 | 1 |
| Lmf2 | -0.17085 | 1.63E-05 | 0.526206 |
| Zfp770 | -0.17105 | 0.006776 | 1 |
| Cdk5rap3 | -0.17109 | 0.000466 | 1 |
| Fam76b | -0.17109 | 0.00424 | 1 |
| Slc2a8 | -0.1712 | 0.000277 | 1 |
| Exoc5 | -0.17134 | 0.002095 | 1 |
| Mrc2 | -0.17138 | 6.76E-05 | 1 |
| Sf1 | -0.17175 | 1.72E-05 | 0.555149 |
| Map7d1 | -0.17183 | 0.000129 | 1 |
| Prorsd1 | -0.17195 | 0.000192 | 1 |
| Ctr9 | -0.17201 | 0.000664 | 1 |
| Scp2 | -0.17202 | 0.000228 | 1 |
| Zbtb38 | -0.17207 | 0.000233 | 1 |
| Snx27 | -0.1722 | 0.002272 | 1 |

| | | | |
|---|---|---|---|
| Gdi1 | -0.1722 | 2.27E-05 | 0.732029 |
| Fam45a | -0.17221 | 8.13E-05 | 1 |
| Reep5 | -0.17228 | 1.01E-05 | 0.324644 |
| Rps6ka5 | -0.17228 | 0.005707 | 1 |
| Espl1 | -0.17229 | 2.38E-05 | 0.769123 |
| Ube3a | -0.1724 | 0.001086 | 1 |
| Maged2 | -0.17271 | 5.08E-08 | 0.001641 |
| 4833439L1 | -0.1728 | 0.002503 | 1 |
| Nr6a1 | -0.17285 | 0.046079 | 1 |
| Gpr137 | -0.17295 | 0.000344 | 1 |
| C87436 | -0.173 | 0.000896 | 1 |
| Bcl9l | -0.17301 | 4.2E-06 | 0.135682 |
| Dyrk2 | -0.17323 | 1.05E-05 | 0.337457 |
| Cflar | -0.17333 | 0.001627 | 1 |
| Ccny | -0.1734 | 0.017314 | 1 |
| Atat1 | -0.1735 | 2.44E-06 | 0.078838 |
| Magi3 | -0.17374 | 8.14E-05 | 1 |
| Ugt1a7c | -0.1738 | 0.000264 | 1 |
| Lbr | -0.17412 | 4.32E-08 | 0.001394 |
| H3f3a | -0.17413 | 3.38E-05 | 1 |
| Ly96 | -0.17421 | 0.001191 | 1 |
| Zfp260 | -0.17439 | 1.7E-05 | 0.549629 |
| 2810403D: | -0.17458 | 0.00127 | 1 |
| Mxi1 | -0.1746 | 0.000731 | 1 |
| Gabpb1 | -0.17499 | 0.000199 | 1 |
| Ptafr | -0.17518 | 2.73E-05 | 0.882637 |
| Pan3 | -0.1752 | 0.056906 | 1 |
| Clip1 | -0.17532 | 0.006357 | 1 |
| Upp2 | -0.17536 | 0.008489 | 1 |
| Pik3c2a | -0.17545 | 0.001285 | 1 |
| Scn1b | -0.17554 | 0.000128 | 1 |
| Ppp1ca | -0.17563 | 0.000126 | 1 |
| Nbr1 | -0.17572 | 2.31E-05 | 0.747371 |
| Tbc1d4 | -0.1761 | 0.000267 | 1 |
| Cd14 | -0.17611 | 1.05E-10 | 3.39E-06 |
| Ubqln2 | -0.17618 | 3.43E-08 | 0.001108 |
| Prcp | -0.17651 | 0.000247 | 1 |
| Armc3 | -0.17653 | 7.65E-11 | 2.47E-06 |
| Mertk | -0.17663 | 2.11E-07 | 0.006805 |
| Tpst2 | -0.17666 | 2.27E-06 | 0.073192 |
| Tmem63a | -0.17671 | 0.005444 | 1 |
| Hdlbp | -0.17684 | 0.006958 | 1 |
| Sfpq | -0.17687 | 0.002755 | 1 |
| Pnpla8 | -0.17721 | 0.000135 | 1 |
| Gm32036 | -0.17741 | 3.23E-06 | 0.104334 |
| Chsy1 | -0.17744 | 1.74E-06 | 0.05606 |
| Kcnj2 | -0.17753 | 5.18E-05 | 1 |
| Zmym4 | -0.17771 | 0.015687 | 1 |

| | | | |
|---|---|---|---|
| Map3k3 | -0.17778 | 0.002289 | 1 |
| Manea | -0.17837 | 0.000544 | 1 |
| Ppm1e | -0.17871 | 8.89E-05 | 1 |
| Mbd5 | -0.17878 | 0.010758 | 1 |
| Zmynd8 | -0.17913 | 0.039348 | 1 |
| Dmtf1 | -0.17934 | 0.002274 | 1 |
| Sppl3 | -0.17945 | 0.051564 | 1 |
| Cd151 | -0.1795 | 0.007056 | 1 |
| Prpf4b | -0.17965 | 0.056066 | 1 |
| Arl3 | -0.17988 | 8.3E-08 | 0.00268 |
| Zmat1 | -0.18013 | 9.3E-08 | 0.003003 |
| Slc25a4 | -0.18038 | 0.000748 | 1 |
| Zfp638 | -0.18039 | 0.002157 | 1 |
| Il11ra1 | -0.18039 | 7.68E-07 | 0.02481 |
| Peg13 | -0.18044 | 7.59E-15 | 2.45E-10 |
| Cfap74 | -0.18059 | 0.04202 | 1 |
| Sf3b2 | -0.18066 | 0.006548 | 1 |
| Ranbp10 | -0.18073 | 0.001406 | 1 |
| Grk3 | -0.18124 | 0.002431 | 1 |
| Tspan14 | -0.18172 | 9.38E-05 | 1 |
| Urgcp | -0.18183 | 0.002974 | 1 |
| Anapc2 | -0.18197 | 0.000206 | 1 |
| Supt3 | -0.18199 | 0.000858 | 1 |
| Bbc3 | -0.18201 | 2.35E-05 | 0.757654 |
| Itpripl1 | -0.18208 | 4.69E-05 | 1 |
| Sdccag8 | -0.18216 | 0.001369 | 1 |
| Pnp | -0.18249 | 6.43E-06 | 0.207683 |
| Phactr2 | -0.1826 | 0.063984 | 1 |
| Fam173a | -0.18278 | 0.000996 | 1 |
| Irf2 | -0.18294 | 3.14E-06 | 0.101275 |
| Tax1bp1 | -0.18307 | 0.00073 | 1 |
| Tubb2a | -0.18329 | 0.000655 | 1 |
| Ceacam1 | -0.18348 | 3.82E-07 | 0.012324 |
| Trim36 | -0.18348 | 6.77E-05 | 1 |
| Magt1 | -0.18349 | 0.008886 | 1 |
| Zfp871 | -0.1835 | 0.00054 | 1 |
| Syngr2 | -0.18372 | 3.05E-07 | 0.009848 |
| Wdfy3 | -0.18396 | 0.001588 | 1 |
| Tmem141 | -0.18405 | 6.8E-05 | 1 |
| Cpsf6 | -0.18412 | 7.84E-05 | 1 |
| Tmtc2 | -0.18431 | 9.33E-11 | 3.01E-06 |
| Spg20 | -0.18436 | 2.34E-06 | 0.075537 |
| Rbm42 | -0.18437 | 0.000229 | 1 |
| Klhl36 | -0.18457 | 5.35E-06 | 0.172673 |
| Pnrc1 | -0.18475 | 6.57E-07 | 0.021209 |
| Celf1 | -0.18476 | 0.003622 | 1 |
| Hmox2 | -0.1848 | 1.23E-05 | 0.395598 |
| Ppfibp2 | -0.18492 | 0.000342 | 1 |

| | | | |
|---|---|---|---|
| Wdr44 | -0.18498 | 0.01728 | 1 |
| Trim2 | -0.18512 | 3.71E-05 | 1 |
| Ramac | -0.18517 | 1.15E-05 | 0.371998 |
| Tnpo1 | -0.18517 | 0.029917 | 1 |
| Nck1 | -0.1858 | 8.88E-05 | 1 |
| Casd1 | -0.18594 | 6.53E-05 | 1 |
| Sema4d | -0.18611 | 8.68E-07 | 0.028039 |
| Crebrf | -0.1862 | 3.96E-05 | 1 |
| B3galt5 | -0.18626 | 0.000427 | 1 |
| Ssh2 | -0.1864 | 5.69E-13 | 1.84E-08 |
| Iscu | -0.18644 | 0.01202 | 1 |
| Slc39a8 | -0.18646 | 9.57E-08 | 0.003088 |
| Rbbp4 | -0.18671 | 1.75E-05 | 0.564754 |
| Clta | -0.18672 | 8.34E-10 | 2.69E-05 |
| Abcg2 | -0.18672 | 0.00518 | 1 |
| Slc35a5 | -0.18712 | 1.81E-05 | 0.584219 |
| Tgfbi | -0.18732 | 0.000159 | 1 |
| Crim1 | -0.18732 | 1.24E-07 | 0.003989 |
| Pura | -0.1874 | 0.006203 | 1 |
| Ing4 | -0.18756 | 5.8E-05 | 1 |
| Zfp467 | -0.18759 | 0.00058 | 1 |
| Zdhhc20 | -0.18784 | 0.000474 | 1 |
| Pcmtd2 | -0.18817 | 9.32E-09 | 0.000301 |
| Dagla | -0.18823 | 0.000346 | 1 |
| Trp53 | -0.18834 | 0.013127 | 1 |
| Vav2 | -0.18835 | 0.010258 | 1 |
| Itm2c | -0.18845 | 1.56E-12 | 5.02E-08 |
| Pepd | -0.18869 | 0.000913 | 1 |
| Syf2 | -0.18875 | 8.04E-06 | 0.259719 |
| Ralgps1 | -0.18892 | 0.019144 | 1 |
| Rnf144b | -0.18956 | 1.39E-06 | 0.044958 |
| Fam89b | -0.18969 | 0.000622 | 1 |
| Rab1a | -0.1899 | 5.35E-05 | 1 |
| Sept7 | -0.19011 | 0.000112 | 1 |
| Clptm1 | -0.19019 | 0.00022 | 1 |
| R3hdm2 | -0.1902 | 0.027145 | 1 |
| Dip2c | -0.1906 | 0.002496 | 1 |
| Ncoa3 | -0.19078 | 0.001356 | 1 |
| Gtf2ird1 | -0.19084 | 2.89E-08 | 0.000934 |
| Hnrnph3 | -0.19087 | 0.000499 | 1 |
| Zfand6 | -0.19089 | 0.000604 | 1 |
| Gngt2 | -0.19111 | 1.77E-05 | 0.570262 |
| Senp7 | -0.19119 | 0.000766 | 1 |
| Tnrc18 | -0.19123 | 0.000836 | 1 |
| Creb1 | -0.1913 | 0.000792 | 1 |
| Arrb2 | -0.19132 | 5.64E-09 | 0.000182 |
| Emc3 | -0.19158 | 2.61E-05 | 0.8427 |
| Ncaph | -0.19165 | 3.7E-10 | 1.19E-05 |

| | | | |
|---|---|---|---|
| Rassf5 | -0.19177 | 0.000175 | 1 |
| Tab2 | -0.1918 | 2.11E-08 | 0.000683 |
| Nrros | -0.19181 | 1.17E-09 | 3.76E-05 |
| Dpm3 | -0.19204 | 2.49E-06 | 0.080392 |
| Herc2 | -0.19212 | 2.51E-05 | 0.8105 |
| Liph | -0.19213 | 0.010438 | 1 |
| Rchy1 | -0.19218 | 2.89E-05 | 0.932359 |
| Usp33 | -0.19238 | 0.000476 | 1 |
| Ralb | -0.19243 | 2.24E-07 | 0.007242 |
| Arhgap17 | -0.19256 | 7.6E-06 | 0.245238 |
| Slc9a7 | -0.19294 | 1.8E-06 | 0.058133 |
| Cd37 | -0.19299 | 1.09E-11 | 3.52E-07 |
| Cd276 | -0.19318 | 3.44E-12 | 1.11E-07 |
| Sp1 | -0.19372 | 1.53E-07 | 0.004929 |
| Bicral | -0.19409 | 0.000247 | 1 |
| Frmd8 | -0.19443 | 0.000268 | 1 |
| Papola | -0.19449 | 1.85E-05 | 0.596688 |
| Stard3 | -0.1947 | 1.73E-05 | 0.559355 |
| Dctn5 | -0.19474 | 9.42E-05 | 1 |
| Acvr1 | -0.19475 | 0.000606 | 1 |
| Aagab | -0.19553 | 6.93E-05 | 1 |
| Parp8 | -0.19554 | 0.022142 | 1 |
| Prps2 | -0.19569 | 6.82E-07 | 0.022024 |
| Oma1 | -0.19583 | 1.74E-05 | 0.560171 |
| Skil | -0.19656 | 0.000354 | 1 |
| Gm3252 | -0.19668 | 4.21E-19 | 1.36E-14 |
| Mfhas1 | -0.19699 | 0.022467 | 1 |
| Bcor | -0.19709 | 8.84E-06 | 0.285451 |
| Bckdha | -0.19739 | 5.04E-05 | 1 |
| Itch | -0.19779 | 0.001121 | 1 |
| Eif1b | -0.19783 | 7.18E-05 | 1 |
| Zfp715 | -0.19797 | 3.37E-06 | 0.108827 |
| Fam91a1 | -0.198 | 8.1E-07 | 0.026149 |
| Sp3 | -0.19825 | 0.000602 | 1 |
| Tcn2 | -0.19836 | 3.34E-08 | 0.001077 |
| Ak1 | -0.19837 | 0.000729 | 1 |
| Ube2d2a | -0.1985 | 5.04E-06 | 0.162689 |
| Maf1 | -0.19862 | 3.2E-07 | 0.010347 |
| Tbxas1 | -0.19917 | 2.24E-06 | 0.072178 |
| Fhod1 | -0.19918 | 2.78E-05 | 0.897 |
| Slc25a45 | -0.19924 | 0.000297 | 1 |
| Mcfd2 | -0.19931 | 5.73E-05 | 1 |
| Commd7 | -0.19942 | 0.0077 | 1 |
| Stt3a | -0.19961 | 1.16E-05 | 0.375602 |
| G3bp2 | -0.19966 | 1.57E-06 | 0.050529 |
| Phlda3 | -0.19984 | 5.84E-08 | 0.001885 |
| Arpc4 | -0.20012 | 9.6E-10 | 3.1E-05 |
| Nf1 | -0.20035 | 0.001819 | 1 |

| | | | |
|---|---|---|---|
| Ctso | -0.20088 | 0.000194 | 1 |
| Lrch1 | -0.20103 | 1.39E-05 | 0.448063 |
| Lpcat1 | -0.20103 | 0.000463 | 1 |
| Trio | -0.20115 | 0.079318 | 1 |
| Zfp950 | -0.20124 | 0.000496 | 1 |
| Idh1 | -0.2014 | 2E-07 | 0.006473 |
| Evi5 | -0.20165 | 0.012841 | 1 |
| Zbtb4 | -0.20166 | 5.38E-06 | 0.173676 |
| Crebbp | -0.20185 | 2.39E-05 | 0.77149 |
| Rgs7bp | -0.20207 | 0.003044 | 1 |
| Ppp1r2 | -0.20245 | 4.28E-07 | 0.013828 |
| Kcnd1 | -0.20249 | 6.2E-06 | 0.200299 |
| Cat | -0.2027 | 7.96E-06 | 0.257006 |
| Tbl1xr1 | -0.203 | 0.000973 | 1 |
| Pbx2 | -0.20307 | 2.23E-08 | 0.000719 |
| Mgmt | -0.20319 | 0.011722 | 1 |
| Fam3a | -0.20355 | 2.34E-07 | 0.007547 |
| Ythdf2 | -0.20393 | 1.53E-05 | 0.492946 |
| Tceal1 | -0.20407 | 6.05E-18 | 1.95E-13 |
| Ube2e3 | -0.20436 | 3.55E-07 | 0.011454 |
| Tasor | -0.20437 | 0.000183 | 1 |
| Tmem223 | -0.20438 | 1.41E-06 | 0.045431 |
| Galnt4 | -0.20445 | 6.55E-09 | 0.000211 |
| Ids | -0.20463 | 1.28E-05 | 0.414661 |
| Flcn | -0.20474 | 0.000172 | 1 |
| Stag2 | -0.20479 | 7.86E-05 | 1 |
| Ppp1r3b | -0.20494 | 7.14E-08 | 0.002305 |
| Tmem106c | -0.20513 | 1.06E-06 | 0.034378 |
| Nek6 | -0.2052 | 0.000384 | 1 |
| Rnf5 | -0.2053 | 1.56E-05 | 0.502566 |
| Pum2 | -0.20538 | 3.05E-05 | 0.98317 |
| Tm2d2 | -0.2058 | 8.42E-06 | 0.271812 |
| Hdac5 | -0.20587 | 1.36E-05 | 0.438565 |
| Tut4 | -0.206 | 5.13E-09 | 0.000166 |
| Ebf3 | -0.20623 | 4.92E-06 | 0.158862 |
| Elmod2 | -0.2064 | 7.1E-07 | 0.022934 |
| Nrm | -0.2065 | 4.51E-09 | 0.000146 |
| Selenok | -0.20693 | 2.63E-08 | 0.000849 |
| Klhl24 | -0.20717 | 3.28E-07 | 0.010576 |
| Rfx3 | -0.20735 | 8.32E-07 | 0.026874 |
| Mkln1 | -0.20738 | 0.001055 | 1 |
| Stag1 | -0.20746 | 0.002116 | 1 |
| Itpr2 | -0.20758 | 5E-05 | 1 |
| Erf | -0.20814 | 6.09E-06 | 0.196711 |
| Trpc4ap | -0.2088 | 2.57E-06 | 0.083016 |
| Kpnb1 | -0.20884 | 1.52E-05 | 0.491187 |
| Arsg | -0.20887 | 0.000443 | 1 |
| Gm3264 | -0.2092 | 1.21E-18 | 3.9E-14 |

| | | | |
|---|---|---|---|
| Aup1 | -0.20921 | 2.22E-08 | 0.000717 |
| Wdr1 | -0.20922 | 1.61E-07 | 0.005206 |
| Rela | -0.20933 | 0.000155 | 1 |
| Kdm3b | -0.20977 | 1.67E-05 | 0.539696 |
| Sesn1 | -0.21013 | 4.96E-08 | 0.001601 |
| Nars | -0.21024 | 4.58E-06 | 0.14772 |
| Mvb12b | -0.21033 | 0.000406 | 1 |
| Kifap3 | -0.21034 | 1.63E-06 | 0.052774 |
| Mbnl2 | -0.21036 | 1.55E-06 | 0.050178 |
| Rac1 | -0.21054 | 1.81E-05 | 0.585472 |
| Canx | -0.21092 | 1.3E-07 | 0.004185 |
| Atxn2l | -0.21122 | 0.000197 | 1 |
| Anp32a | -0.21133 | 9.04E-07 | 0.029179 |
| Rbmx | -0.21136 | 1.14E-05 | 0.367911 |
| Cbx1 | -0.21155 | 9.66E-05 | 1 |
| Spi1 | -0.21163 | 8.8E-12 | 2.84E-07 |
| Ociad1 | -0.21191 | 9.32E-08 | 0.003008 |
| Prkacb | -0.21205 | 1.53E-06 | 0.049359 |
| Pex2 | -0.21207 | 3.41E-06 | 0.110062 |
| Slc39a10 | -0.2123 | 2.75E-07 | 0.008888 |
| Hist1h2be | -0.2128 | 1.15E-09 | 3.7E-05 |
| Creb3l2 | -0.21293 | 1.16E-05 | 0.375806 |
| Tmem43 | -0.21296 | 1.29E-08 | 0.000417 |
| Foxn3 | -0.21302 | 2.67E-11 | 8.62E-07 |
| Zfp62 | -0.21312 | 1.22E-05 | 0.394052 |
| Add1 | -0.21323 | 1.28E-07 | 0.004118 |
| 4933434E2 | -0.21332 | 5.09E-06 | 0.164488 |
| Lpin2 | -0.21406 | 1.79E-08 | 0.000577 |
| Cmss1 | -0.21413 | 9.72E-10 | 3.14E-05 |
| Btk | -0.21425 | 4.07E-06 | 0.131331 |
| Isyna1 | -0.2143 | 5.57E-06 | 0.179682 |
| Igfbp4 | -0.21466 | 1.3E-06 | 0.041973 |
| Arl6ip1 | -0.21466 | 5E-07 | 0.016132 |
| Tubb2b | -0.2147 | 1.18E-06 | 0.038076 |
| Stambpl1 | -0.21488 | 1.69E-06 | 0.054414 |
| Cpne3 | -0.21498 | 0.000177 | 1 |
| Hnrnph1 | -0.2152 | 7.73E-05 | 1 |
| Cops4 | -0.21531 | 1.8E-06 | 0.058048 |
| Cysltr1 | -0.21535 | 3.39E-07 | 0.010961 |
| Srpk3 | -0.21552 | 1.68E-17 | 5.43E-13 |
| Zswim6 | -0.21568 | 0.002446 | 1 |
| Nfatc2 | -0.21591 | 3.52E-05 | 1 |
| Clec4a2 | -0.21621 | 7.49E-10 | 2.42E-05 |
| Fnta | -0.21627 | 9.26E-08 | 0.00299 |
| Scmh1 | -0.21639 | 0.000843 | 1 |
| Grk2 | -0.21651 | 0.005758 | 1 |
| Rpn1 | -0.21655 | 7.62E-08 | 0.002461 |
| Cmya5 | -0.21715 | 1.54E-12 | 4.98E-08 |

| | | | |
|---|---|---|---|
| Adap2 | -0.21719 | 7.21E-08 | 0.002326 |
| Ext2 | -0.21764 | 0.000408 | 1 |
| Ykt6 | -0.21781 | 1.99E-06 | 0.064339 |
| Vps28 | -0.21828 | 0.000742 | 1 |
| Rab11b | -0.21843 | 2.44E-09 | 7.87E-05 |
| Mknk1 | -0.21848 | 7.88E-09 | 0.000254 |
| Zfp691 | -0.21864 | 7.31E-09 | 0.000236 |
| Slc39a12 | -0.21866 | 2.7E-16 | 8.71E-12 |
| Myo18a | -0.21868 | 0.026012 | 1 |
| Tmem86a | -0.21948 | 7.05E-08 | 0.002275 |
| Ipo11 | -0.21962 | 3.02E-07 | 0.009746 |
| Bicd2 | -0.21998 | 4.62E-06 | 0.149011 |
| Apba1 | -0.22031 | 0.000565 | 1 |
| Sash3 | -0.22092 | 4.86E-08 | 0.001569 |
| Helz | -0.22098 | 3.83E-05 | 1 |
| Krcc1 | -0.2213 | 6.37E-06 | 0.205549 |
| Eif1 | -0.22158 | 2.12E-12 | 6.84E-08 |
| Tnrc6b | -0.22196 | 2.09E-09 | 6.75E-05 |
| Nfya | -0.22258 | 0.000336 | 1 |
| Ccdc90b | -0.22278 | 1.09E-08 | 0.000353 |
| Pigp | -0.22279 | 3.07E-06 | 0.099197 |
| Sh2b3 | -0.2235 | 2.09E-06 | 0.067514 |
| Vhl | -0.22363 | 4.22E-05 | 1 |
| Inpp5d | -0.22376 | 2.49E-14 | 8.04E-10 |
| Bmp2k | -0.22404 | 1.78E-07 | 0.00575 |
| Tmed10 | -0.22411 | 7.16E-11 | 2.31E-06 |
| Adam17 | -0.22432 | 1.15E-06 | 0.037012 |
| Csnk1a1 | -0.22448 | 6.54E-08 | 0.002111 |
| Ddx3x | -0.22464 | 1.81E-05 | 0.584143 |
| Cnrip1 | -0.22478 | 9.17E-09 | 0.000296 |
| Tmem52 | -0.2249 | 1.27E-10 | 4.11E-06 |
| Gadd45g | -0.22512 | 2.47E-07 | 0.007983 |
| Ints3 | -0.22521 | 8.17E-08 | 0.002639 |
| Gm35188 | -0.22528 | 8.8E-07 | 0.028406 |
| Poc1b | -0.22531 | 1.78E-06 | 0.057382 |
| Nfic | -0.22533 | 1.28E-06 | 0.04123 |
| Hnrnpk | -0.2254 | 1.84E-11 | 5.94E-07 |
| Tcf4 | -0.22566 | 4.16E-06 | 0.134396 |
| Gm46367 | -0.22568 | 1.1E-08 | 0.000355 |
| Stau2 | -0.22637 | 0.000165 | 1 |
| Dele1 | -0.22638 | 1.36E-06 | 0.043905 |
| Clcn5 | -0.22653 | 2.33E-06 | 0.075317 |
| Trim47 | -0.22679 | 4.57E-05 | 1 |
| Purb | -0.22679 | 1.53E-05 | 0.49342 |
| Slc4a7 | -0.22718 | 0.000186 | 1 |
| Tmem175 | -0.22771 | 3.09E-06 | 0.099911 |
| Slc18b1 | -0.22772 | 3E-13 | 9.69E-09 |
| Ang | -0.22782 | 6.32E-05 | 1 |

| | | | |
|---|---|---|---|
| Syk | -0.22789 | 5.8E-07 | 0.018717 |
| Papss1 | -0.2281 | 7.16E-07 | 0.023105 |
| Adgrl2 | -0.22811 | 4.74E-07 | 0.015292 |
| Arhgef10l | -0.22819 | 3.87E-12 | 1.25E-07 |
| Add3 | -0.22835 | 8.65E-08 | 0.002794 |
| Ptbp1 | -0.2285 | 7.5E-07 | 0.024198 |
| Lasp1 | -0.22864 | 6.38E-10 | 2.06E-05 |
| Tspan4 | -0.22879 | 8.47E-08 | 0.002735 |
| Smad1 | -0.22914 | 2.21E-05 | 0.714245 |
| Tor4a | -0.2296 | 2.87E-06 | 0.092743 |
| Clstn1 | -0.22966 | 4.92E-08 | 0.00159 |
| Rasal3 | -0.22971 | 1.23E-09 | 3.98E-05 |
| Nhsl2 | -0.23001 | 5.93E-18 | 1.92E-13 |
| Dcaf12 | -0.23003 | 3.34E-08 | 0.00108 |
| Kpna3 | -0.23013 | 1.25E-07 | 0.004045 |
| Cyren | -0.23019 | 8.63E-10 | 2.79E-05 |
| Smyd3 | -0.23028 | 0.000356 | 1 |
| Txndc15 | -0.23036 | 3.69E-07 | 0.011922 |
| Gm31243 | -0.23043 | 3.16E-13 | 1.02E-08 |
| Zfp869 | -0.23067 | 2.13E-08 | 0.000689 |
| Ddx17 | -0.23089 | 0.000435 | 1 |
| Fez2 | -0.23121 | 1.79E-06 | 0.057715 |
| Aftph | -0.23126 | 0.000693 | 1 |
| Rtn3 | -0.23136 | 4.18E-11 | 1.35E-06 |
| Man2b2 | -0.23207 | 8.99E-06 | 0.290292 |
| Rsu1 | -0.23225 | 8.78E-07 | 0.028335 |
| Tiprl | -0.2324 | 9.29E-06 | 0.299974 |
| Stard8 | -0.233 | 3.32E-07 | 0.010725 |
| Kansl1l | -0.23365 | 1.68E-05 | 0.541345 |
| Mkrn1 | -0.23375 | 2.09E-06 | 0.067409 |
| 1600022D1 | -0.23423 | 1.05E-15 | 3.37E-11 |
| Gm3164 | -0.23448 | 1.61E-23 | 5.19E-19 |
| Mast3 | -0.23566 | 1.27E-06 | 0.041102 |
| Tspyl1 | -0.23576 | 4.5E-10 | 1.45E-05 |
| Snx14 | -0.23586 | 1.67E-09 | 5.4E-05 |
| Arrb1 | -0.23609 | 5.24E-10 | 1.69E-05 |
| Pafah1b1 | -0.23682 | 5.6E-08 | 0.001807 |
| Prkar1a | -0.23709 | 7.32E-12 | 2.36E-07 |
| Mgrn1 | -0.2371 | 6.2E-08 | 0.002001 |
| Gm9949 | -0.23713 | 5.48E-10 | 1.77E-05 |
| Bend6 | -0.23749 | 6.84E-16 | 2.21E-11 |
| Aars | -0.23759 | 3.62E-08 | 0.001167 |
| Man1a2 | -0.23824 | 1.01E-06 | 0.032513 |
| Il10rb | -0.23845 | 1.65E-09 | 5.32E-05 |
| Cib1 | -0.23888 | 9.15E-09 | 0.000295 |
| Slc12a6 | -0.23896 | 5.75E-05 | 1 |
| Sall3 | -0.23914 | 1.72E-06 | 0.055521 |
| BC035044 | -0.23974 | 2.99E-06 | 0.096527 |

| | | | |
|---|---|---|---|
| Gm3411 | -0.23998 | 3.64E-20 | 1.17E-15 |
| Fam117b | -0.24017 | 0.000278 | 1 |
| Rnf130 | -0.2402 | 8.5E-07 | 0.027429 |
| Vrk1 | -0.24062 | 1.74E-06 | 0.056111 |
| Exoc3 | -0.24074 | 1.18E-05 | 0.382509 |
| Gm43821 | -0.24075 | 5.2E-12 | 1.68E-07 |
| 2900052L1 | -0.24111 | 2.4E-15 | 7.74E-11 |
| Clip2 | -0.24138 | 1.94E-05 | 0.626084 |
| Bag6 | -0.24143 | 7.64E-11 | 2.47E-06 |
| Fam114a2 | -0.24228 | 1.08E-05 | 0.349827 |
| Wdr81 | -0.24232 | 1.6E-10 | 5.16E-06 |
| Cbx3 | -0.24252 | 1.84E-07 | 0.005939 |
| Cerk | -0.24259 | 2.11E-08 | 0.000682 |
| Cul1 | -0.24299 | 1.75E-05 | 0.564769 |
| Mindy2 | -0.24309 | 2.19E-06 | 0.070795 |
| Gm30329 | -0.24338 | 5.2E-20 | 1.68E-15 |
| Gabarapl2 | -0.24358 | 1.11E-05 | 0.357075 |
| Cbfa2t3 | -0.24373 | 4.15E-10 | 1.34E-05 |
| Arhgef6 | -0.24383 | 2.7E-07 | 0.00872 |
| Arhgap25 | -0.24429 | 5.78E-07 | 0.018667 |
| Gm35154 | -0.24446 | 5.34E-06 | 0.172434 |
| Plscr3 | -0.2449 | 8.12E-09 | 0.000262 |
| Phf1 | -0.24493 | 3.66E-13 | 1.18E-08 |
| Itgb3 | -0.24517 | 9.84E-10 | 3.18E-05 |
| Swap70 | -0.24604 | 9.86E-08 | 0.003183 |
| Rp2 | -0.24638 | 5.62E-06 | 0.181306 |
| Hist1h1e | -0.24638 | 1.26E-05 | 0.405696 |
| Cnot2 | -0.24643 | 1.74E-05 | 0.561223 |
| Zfp934 | -0.24653 | 1.09E-09 | 3.51E-05 |
| Gm3173 | -0.2468 | 3.23E-28 | 1.04E-23 |
| Rhoa | -0.24762 | 3.52E-14 | 1.14E-09 |
| Rgs19 | -0.24803 | 1.4E-10 | 4.53E-06 |
| Hmgb1 | -0.24829 | 1.31E-09 | 4.23E-05 |
| Wbp2 | -0.24844 | 1.34E-06 | 0.043315 |
| Htra2 | -0.24861 | 8.53E-08 | 0.002755 |
| 4632427E1 | -0.24865 | 5.93E-07 | 0.01915 |
| Ddx5 | -0.24918 | 3.22E-23 | 1.04E-18 |
| Abcd3 | -0.24928 | 7.1E-10 | 2.29E-05 |
| Glg1 | -0.24936 | 5.67E-07 | 0.018295 |
| Fbrsl1 | -0.24942 | 3.82E-06 | 0.123312 |
| 3222401L1 | -0.24991 | 4.7E-14 | 1.52E-09 |
| Arl4c | -0.24994 | 3.13E-09 | 0.000101 |
| Gstp1 | -0.25025 | 3.17E-08 | 0.001023 |
| Arf1 | -0.25043 | 6.63E-13 | 2.14E-08 |
| Tom1l1 | -0.25146 | 2.23E-14 | 7.19E-10 |
| Gprasp1 | -0.25148 | 2.67E-14 | 8.62E-10 |
| Nrp1 | -0.25153 | 1.67E-06 | 0.053888 |
| Dtx4 | -0.2517 | 3.66E-11 | 1.18E-06 |

| | | | |
|---|---|---|---|
| Rap1a | -0.25192 | 1.86E-08 | 0.0006 |
| Ttc7 | -0.25207 | 9.2E-07 | 0.029701 |
| Ywhah | -0.25327 | 1.96E-10 | 6.31E-06 |
| Crat | -0.25348 | 5.73E-09 | 0.000185 |
| Tifab | -0.25349 | 1.82E-07 | 0.005872 |
| Snx13 | -0.25354 | 4.13E-07 | 0.013323 |
| Ywhaz | -0.2536 | 3.01E-13 | 9.71E-09 |
| Lrch4 | -0.25363 | 1.5E-09 | 4.85E-05 |
| Prpf8 | -0.25373 | 3.32E-08 | 0.001071 |
| Entpd1 | -0.2539 | 2.22E-22 | 7.17E-18 |
| Grcc10 | -0.25403 | 5.53E-08 | 0.001786 |
| Ets1 | -0.25467 | 4.59E-27 | 1.48E-22 |
| Pknox1 | -0.25497 | 8.84E-08 | 0.002854 |
| Inka2 | -0.25511 | 8.98E-08 | 0.002899 |
| Arvcf | -0.25518 | 1.35E-14 | 4.35E-10 |
| Tsen34 | -0.25521 | 3.74E-08 | 0.001208 |
| Arl10 | -0.25539 | 2.47E-11 | 7.97E-07 |
| Rnaset2a | -0.25624 | 1.03E-06 | 0.033256 |
| Rgl2 | -0.25689 | 4.18E-11 | 1.35E-06 |
| Hmg20a | -0.2571 | 7.66E-09 | 0.000247 |
| Dnmt3a | -0.25729 | 0.004584 | 1 |
| Dync1i2 | -0.25757 | 1.5E-08 | 0.000483 |
| Smpd1 | -0.2581 | 8.01E-13 | 2.58E-08 |
| Pik3r5 | -0.25826 | 6.51E-07 | 0.02101 |
| Tpm3 | -0.25843 | 1.57E-14 | 5.08E-10 |
| Slc37a2 | -0.2586 | 4.23E-09 | 0.000137 |
| Immp2l | -0.25877 | 0.004567 | 1 |
| Usp24 | -0.25879 | 1.53E-07 | 0.004929 |
| Klk8 | -0.25888 | 6.4E-09 | 0.000206 |
| Ttc3 | -0.25923 | 1.34E-06 | 0.043178 |
| Accs | -0.25938 | 1.85E-09 | 5.96E-05 |
| Ctnnb1 | -0.25944 | 1.27E-09 | 4.11E-05 |
| Ppp6r1 | -0.25961 | 1.26E-08 | 0.000407 |
| Kmt5b | -0.25973 | 3.16E-06 | 0.101955 |
| Cnpy2 | -0.26009 | 2.24E-07 | 0.007223 |
| I830077J0: | -0.26044 | 1.27E-08 | 0.000411 |
| Cdc42 | -0.26097 | 1.2E-21 | 3.86E-17 |
| Saysd1 | -0.2613 | 3.92E-10 | 1.26E-05 |
| Retreg1 | -0.2616 | 6.07E-07 | 0.01961 |
| Acads | -0.26162 | 3.79E-09 | 0.000122 |
| Zfp608 | -0.26196 | 4.06E-07 | 0.013092 |
| Arid1a | -0.26229 | 5.72E-09 | 0.000185 |
| Adam10 | -0.26234 | 1.52E-09 | 4.9E-05 |
| Arhgap4 | -0.26254 | 1.19E-10 | 3.84E-06 |
| Morf4l1 | -0.26289 | 7.77E-10 | 2.51E-05 |
| Ctbp2 | -0.26349 | 9.52E-11 | 3.08E-06 |
| Zfp385a | -0.2638 | 1.16E-08 | 0.000373 |
| Prag1 | -0.26426 | 2.92E-12 | 9.42E-08 |

| | | | |
|---|---|---|---|
| Arhgef2 | -0.26436 | 6.67E-08 | 0.002154 |
| Slc25a11 | -0.26454 | 6.27E-09 | 0.000202 |
| 9130015G1 | -0.26464 | 5.48E-14 | 1.77E-09 |
| Tpbgl | -0.26494 | 1.05E-06 | 0.033938 |
| Tmem50a | -0.26527 | 8.36E-17 | 2.7E-12 |
| Cap1 | -0.26589 | 1.75E-15 | 5.65E-11 |
| Clcn4 | -0.26597 | 4.99E-07 | 0.016097 |
| Atp6v0a1 | -0.2662 | 1.02E-09 | 3.3E-05 |
| Arl1 | -0.2662 | 8.29E-09 | 0.000268 |
| Efnb1 | -0.26666 | 7.56E-21 | 2.44E-16 |
| Sik2 | -0.26692 | 1.51E-07 | 0.004869 |
| Tk2 | -0.26704 | 1.65E-09 | 5.34E-05 |
| Akna | -0.26721 | 5.96E-10 | 1.93E-05 |
| Arhgap12 | -0.26735 | 3.71E-07 | 0.011973 |
| Gpr137b | -0.26752 | 1.71E-09 | 5.52E-05 |
| Gm37494 | -0.26792 | 2.25E-10 | 7.26E-06 |
| Sec14l1 | -0.2682 | 4.01E-09 | 0.00013 |
| Cln8 | -0.26841 | 1.28E-08 | 0.000413 |
| Kat6a | -0.26857 | 4.32E-07 | 0.013943 |
| Kdelr1 | -0.26927 | 2.92E-09 | 9.44E-05 |
| Ep300 | -0.26944 | 5.97E-10 | 1.93E-05 |
| Sgms1 | -0.26991 | 6.54E-07 | 0.021119 |
| Taz | -0.26992 | 4.92E-11 | 1.59E-06 |
| Rasa3 | -0.26992 | 3.91E-08 | 0.001263 |
| Tmem30a | -0.27026 | 1.12E-10 | 3.6E-06 |
| 2610035D1 | -0.27101 | 3.33E-21 | 1.08E-16 |
| Tgfbrap1 | -0.27118 | 6.48E-11 | 2.09E-06 |
| Mapk14 | -0.27119 | 1.32E-07 | 0.00425 |
| Smox | -0.27131 | 6.4E-11 | 2.07E-06 |
| Vps35 | -0.2716 | 3.8E-08 | 0.001227 |
| Arl8b | -0.27177 | 1.36E-09 | 4.39E-05 |
| Sh3pxd2a | -0.27194 | 0.000836 | 1 |
| Tmem37 | -0.27207 | 2.36E-09 | 7.6E-05 |
| Zbtb20 | -0.27247 | 2.67E-07 | 0.008607 |
| Tmem173 | -0.27254 | 3.7E-14 | 1.19E-09 |
| Hspb3 | -0.27371 | 7.28E-15 | 2.35E-10 |
| Plpp1 | -0.27423 | 4.1E-09 | 0.000132 |
| Chchd2 | -0.27427 | 1.35E-08 | 0.000435 |
| Adam3 | -0.27434 | 5.54E-11 | 1.79E-06 |
| Endod1 | -0.27434 | 3.7E-10 | 1.2E-05 |
| Stat6 | -0.27505 | 1.62E-09 | 5.25E-05 |
| Slc16a7 | -0.27575 | 7.56E-08 | 0.00244 |
| S1pr1 | -0.2763 | 3.53E-24 | 1.14E-19 |
| Aamp | -0.27631 | 2.33E-13 | 7.51E-09 |
| Abcd2 | -0.27688 | 1.88E-08 | 0.000607 |
| Slmap | -0.27747 | 2.48E-08 | 0.000801 |
| Phc2 | -0.27756 | 3.36E-07 | 0.010856 |
| Pttg1ip | -0.27821 | 3.38E-11 | 1.09E-06 |

| | | | |
|---|---|---|---|
| Creg1 | -0.27996 | 2.28E-13 | 7.35E-09 |
| Chst8 | -0.28047 | 7.1E-20 | 2.29E-15 |
| Paip2 | -0.28052 | 3.92E-10 | 1.26E-05 |
| Smad3 | -0.28062 | 2.88E-07 | 0.009311 |
| Rhobtb1 | -0.28079 | 1.37E-11 | 4.44E-07 |
| Dpysl2 | -0.28118 | 4.4E-09 | 0.000142 |
| Casp8 | -0.28127 | 1.81E-08 | 0.000585 |
| Mob1a | -0.28137 | 1.16E-10 | 3.75E-06 |
| Ccdc50 | -0.28162 | 5.04E-08 | 0.001626 |
| Nr6a1os | -0.28168 | 2.16E-09 | 6.97E-05 |
| Pxn | -0.28197 | 1.44E-09 | 4.66E-05 |
| Rab31 | -0.28214 | 1.09E-13 | 3.53E-09 |
| Son | -0.2824 | 5.73E-22 | 1.85E-17 |
| Etv1 | -0.28291 | 2.57E-09 | 8.28E-05 |
| Usp22 | -0.28306 | 1.87E-13 | 6.02E-09 |
| Eml6 | -0.28308 | 1.02E-06 | 0.033084 |
| Dusp7 | -0.28341 | 2.59E-13 | 8.35E-09 |
| Gnas | -0.28367 | 8.75E-10 | 2.82E-05 |
| Cnpy3 | -0.28382 | 6.58E-12 | 2.12E-07 |
| Clasp2 | -0.2839 | 3.46E-13 | 1.12E-08 |
| Hps4 | -0.28401 | 5.06E-11 | 1.63E-06 |
| Ski | -0.28574 | 8.92E-07 | 0.028795 |
| Lactb2 | -0.28594 | 6.16E-10 | 1.99E-05 |
| Stk10 | -0.286 | 1.35E-13 | 4.35E-09 |
| Tbc1d9b | -0.28613 | 1.15E-07 | 0.003714 |
| Rab39 | -0.28614 | 1.69E-11 | 5.45E-07 |
| Tmem109 | -0.28635 | 1.83E-11 | 5.9E-07 |
| Arhgef1 | -0.28745 | 5.24E-15 | 1.69E-10 |
| Gcnt1 | -0.28751 | 7.63E-11 | 2.46E-06 |
| Slc25a37 | -0.28761 | 4.49E-14 | 1.45E-09 |
| Cd302 | -0.28785 | 7.89E-11 | 2.55E-06 |
| B4gat1 | -0.2879 | 1.2E-13 | 3.86E-09 |
| Arid3a | -0.28831 | 1.53E-12 | 4.95E-08 |
| Pja2 | -0.28904 | 5.24E-11 | 1.69E-06 |
| Zranb2 | -0.28906 | 2.97E-06 | 0.095944 |
| Adcy9 | -0.28917 | 8.15E-07 | 0.026317 |
| Inka1 | -0.28968 | 5.37E-09 | 0.000173 |
| Map1s | -0.28991 | 2.29E-10 | 7.4E-06 |
| Pdlim4 | -0.29018 | 6.12E-10 | 1.98E-05 |
| Lims1 | -0.29024 | 1.98E-13 | 6.38E-09 |
| Disc1 | -0.29025 | 1.65E-05 | 0.531107 |
| Slc8a1 | -0.29276 | 4.12E-12 | 1.33E-07 |
| Ctsa | -0.29363 | 2.19E-25 | 7.07E-21 |
| Mboat7 | -0.29391 | 1.06E-10 | 3.42E-06 |
| Nfatc1 | -0.2942 | 6.62E-11 | 2.14E-06 |
| H1f0 | -0.29442 | 3.19E-12 | 1.03E-07 |
| Kdm2b | -0.29447 | 9.1E-13 | 2.94E-08 |
| 6030468B1 | -0.29547 | 5.75E-19 | 1.86E-14 |

| | | | |
|---|---|---|---|
| Ccdc12 | -0.29598 | 3.2E-13 | 1.03E-08 |
| Ccdc162 | -0.29619 | 1.6E-27 | 5.17E-23 |
| Wdfy2 | -0.29666 | 3.34E-08 | 0.001078 |
| Atp6v0b | -0.29667 | 8.55E-25 | 2.76E-20 |
| Maged1 | -0.29676 | 1.35E-22 | 4.37E-18 |
| Scamp2 | -0.29682 | 7.49E-26 | 2.42E-21 |
| Adgre1 | -0.29728 | 1.84E-10 | 5.93E-06 |
| Hivep3 | -0.2973 | 3.34E-06 | 0.107709 |
| Hnrnpul1 | -0.29734 | 2.09E-14 | 6.75E-10 |
| Ramp1 | -0.29811 | 7.97E-09 | 0.000257 |
| Cirbp | -0.29822 | 1.49E-13 | 4.8E-09 |
| Sh2b2 | -0.29849 | 2.19E-11 | 7.08E-07 |
| P4ha1 | -0.2986 | 3.35E-11 | 1.08E-06 |
| Epn2 | -0.29893 | 9.34E-15 | 3.01E-10 |
| Klf7 | -0.29916 | 4.05E-11 | 1.31E-06 |
| Arhgap39 | -0.29929 | 5.49E-06 | 0.177157 |
| Rhob | -0.29951 | 5.79E-33 | 1.87E-28 |
| Tuba1b | -0.29969 | 5.55E-09 | 0.000179 |
| P2ry10b | -0.30038 | 7.27E-20 | 2.35E-15 |
| Cmtm7 | -0.30087 | 1.63E-16 | 5.25E-12 |
| Gna12 | -0.30091 | 8.43E-06 | 0.27216 |
| Lars2 | -0.3014 | 1.73E-13 | 5.58E-09 |
| Tmem59 | -0.30158 | 1.97E-19 | 6.36E-15 |
| Tnfrsf1b | -0.30167 | 1.24E-13 | 4E-09 |
| Macf1 | -0.30172 | 7.78E-13 | 2.51E-08 |
| Samsn1 | -0.30199 | 1.26E-15 | 4.06E-11 |
| Ywhae | -0.30258 | 1.71E-14 | 5.52E-10 |
| Smarca2 | -0.3027 | 6.19E-09 | 0.0002 |
| 1700011L2 | -0.30321 | 4.5E-23 | 1.45E-18 |
| Ppp1r18 | -0.30327 | 1.31E-14 | 4.24E-10 |
| Dock4 | -0.3035 | 1.75E-15 | 5.65E-11 |
| Sema4b | -0.30398 | 1.42E-25 | 4.59E-21 |
| Cables1 | -0.30511 | 4.29E-06 | 0.138517 |
| Itgam | -0.30524 | 4.96E-25 | 1.6E-20 |
| Glis3 | -0.30533 | 4.53E-18 | 1.46E-13 |
| Arhgap22 | -0.30569 | 2.19E-08 | 0.000706 |
| Matk | -0.30569 | 1.4E-21 | 4.53E-17 |
| Laptm5 | -0.30584 | 4.09E-58 | 1.32E-53 |
| Med13l | -0.30596 | 1.53E-09 | 4.95E-05 |
| Ddx6 | -0.30625 | 7.05E-11 | 2.28E-06 |
| Cass4 | -0.30841 | 8.4E-17 | 2.71E-12 |
| Jup | -0.30844 | 1.26E-14 | 4.07E-10 |
| Rere | -0.3087 | 1.46E-11 | 4.73E-07 |
| Ccdc93 | -0.30874 | 1.67E-11 | 5.39E-07 |
| Atraid | -0.30913 | 2.49E-14 | 8.05E-10 |
| Crot | -0.30915 | 1.34E-20 | 4.33E-16 |
| Wsb1 | -0.30938 | 1.71E-13 | 5.51E-09 |
| Ehd4 | -0.30949 | 2.05E-14 | 6.62E-10 |

| | | | |
|---|---|---|---|
| Gm3002 | -0.30974 | 1.99E-34 | 6.41E-30 |
| Mfap3 | -0.31019 | 4.94E-11 | 1.59E-06 |
| Cnot8 | -0.3104 | 3.9E-11 | 1.26E-06 |
| Pid1 | -0.31178 | 1.46E-08 | 0.000473 |
| Clec4a3 | -0.31227 | 5.28E-12 | 1.71E-07 |
| Cd33 | -0.31253 | 6.48E-23 | 2.09E-18 |
| Tram1 | -0.31266 | 4.71E-19 | 1.52E-14 |
| Cyb5a | -0.31292 | 2.97E-13 | 9.58E-09 |
| Prpsap2 | -0.31308 | 1.06E-10 | 3.43E-06 |
| Hhex | -0.31337 | 3.76E-14 | 1.21E-09 |
| Cmtm8 | -0.31369 | 5.83E-16 | 1.88E-11 |
| Mrfap1 | -0.31412 | 1.11E-11 | 3.57E-07 |
| Gmpr | -0.31429 | 2.66E-15 | 8.59E-11 |
| Cbl | -0.31502 | 1.15E-14 | 3.7E-10 |
| Col15a1 | -0.31595 | 7.14E-12 | 2.31E-07 |
| Rab14 | -0.31646 | 5.09E-22 | 1.64E-17 |
| Tmem64 | -0.31653 | 2.87E-15 | 9.26E-11 |
| Tpcn1 | -0.31682 | 7.76E-14 | 2.51E-09 |
| Gm3248 | -0.31706 | 8.63E-28 | 2.79E-23 |
| Zfp532 | -0.31723 | 3.27E-11 | 1.06E-06 |
| Pbx1 | -0.3174 | 1.42E-12 | 4.6E-08 |
| Myo1d | -0.31747 | 1.73E-33 | 5.58E-29 |
| Tspan7 | -0.31848 | 1.2E-13 | 3.87E-09 |
| Mlec | -0.31878 | 5.92E-14 | 1.91E-09 |
| Dock1 | -0.32026 | 1.52E-10 | 4.92E-06 |
| Serinc1 | -0.32068 | 3.92E-16 | 1.27E-11 |
| Rtn4 | -0.32073 | 8.03E-22 | 2.59E-17 |
| Rnf7 | -0.32116 | 3.05E-12 | 9.86E-08 |
| Itgb1 | -0.32133 | 2.43E-17 | 7.85E-13 |
| Ralgps2 | -0.32136 | 8.52E-11 | 2.75E-06 |
| Ubc | -0.3218 | 6.14E-29 | 1.98E-24 |
| Mib2 | -0.32258 | 2.23E-19 | 7.2E-15 |
| Wls | -0.32263 | 8.92E-13 | 2.88E-08 |
| Frmd4b | -0.32334 | 1.49E-19 | 4.81E-15 |
| Fer | -0.32342 | 4.99E-10 | 1.61E-05 |
| Zeb1 | -0.32396 | 2.34E-08 | 0.000755 |
| Tln2 | -0.3241 | 2.42E-08 | 0.000781 |
| Acss1 | -0.32439 | 2.15E-14 | 6.93E-10 |
| Atp2c1 | -0.32464 | 2.12E-13 | 6.83E-09 |
| Zeb2os | -0.32486 | 2.96E-12 | 9.54E-08 |
| Ctnnd2 | -0.32512 | 5.41E-28 | 1.75E-23 |
| Sipa1 | -0.32523 | 1.13E-17 | 3.64E-13 |
| Mgat5 | -0.32537 | 1.05E-08 | 0.000338 |
| Map2k1 | -0.3256 | 7.3E-10 | 2.36E-05 |
| Hsd17b11 | -0.32567 | 1.35E-15 | 4.37E-11 |
| Abcb4 | -0.32574 | 1.29E-23 | 4.17E-19 |
| Abi3 | -0.32687 | 4.1E-26 | 1.32E-21 |
| Sec62 | -0.32719 | 1.27E-11 | 4.11E-07 |

| | | | |
|---|---|---|---|
| Sypl | -0.32804 | 5.79E-15 | 1.87E-10 |
| Rin2 | -0.32865 | 1.56E-20 | 5.05E-16 |
| Mapk3 | -0.32918 | 1.28E-13 | 4.13E-09 |
| Il6ra | -0.32967 | 2.91E-18 | 9.4E-14 |
| Sbf2 | -0.33044 | 1.04E-11 | 3.34E-07 |
| Naa35 | -0.33099 | 8.36E-16 | 2.7E-11 |
| Large1 | -0.33133 | 2.2E-14 | 7.1E-10 |
| Map4k4 | -0.33151 | 2.76E-09 | 8.91E-05 |
| Pik3cg | -0.33182 | 5.08E-16 | 1.64E-11 |
| Hook3 | -0.33234 | 2.61E-09 | 8.43E-05 |
| Acp2 | -0.33237 | 3.8E-14 | 1.23E-09 |
| Rsrp1 | -0.3327 | 1.32E-21 | 4.27E-17 |
| Slc35c2 | -0.33284 | 7.27E-17 | 2.35E-12 |
| Tbc1d5 | -0.3329 | 2.67E-15 | 8.61E-11 |
| Hp1bp3 | -0.33367 | 3.97E-16 | 1.28E-11 |
| Bach2 | -0.33462 | 1.99E-13 | 6.42E-09 |
| Ldhb | -0.33473 | 9.92E-20 | 3.2E-15 |
| Tm9sf2 | -0.33544 | 3.71E-18 | 1.2E-13 |
| Mbnl1 | -0.3356 | 9.44E-36 | 3.05E-31 |
| Slc24a3 | -0.33658 | 3.32E-07 | 0.010725 |
| Atxn7l3b | -0.33684 | 2.14E-11 | 6.91E-07 |
| Lag3 | -0.33805 | 3.21E-13 | 1.04E-08 |
| Psenen | -0.33826 | 2.34E-16 | 7.55E-12 |
| Cdk5r1 | -0.33874 | 1.5E-30 | 4.85E-26 |
| Nr3c2 | -0.33926 | 2.61E-13 | 8.42E-09 |
| Rap1gds1 | -0.33937 | 4.31E-16 | 1.39E-11 |
| Tmcc3 | -0.33971 | 1.02E-13 | 3.31E-09 |
| Fgfr2 | -0.33985 | 5.93E-14 | 1.92E-09 |
| Ubl3 | -0.34041 | 2.34E-18 | 7.57E-14 |
| Pde4d | -0.34046 | 1.79E-13 | 5.78E-09 |
| Lysmd4 | -0.34075 | 7.15E-17 | 2.31E-12 |
| Rilpl1 | -0.34078 | 3.11E-16 | 1E-11 |
| Stab1 | -0.34147 | 6.86E-20 | 2.22E-15 |
| Tacc1 | -0.34166 | 4.86E-17 | 1.57E-12 |
| Gfm2 | -0.34226 | 7.04E-14 | 2.27E-09 |
| Rbfox1 | -0.34337 | 7.05E-15 | 2.28E-10 |
| Dennd2c | -0.34392 | 2.01E-16 | 6.5E-12 |
| Rab6b | -0.34422 | 5.76E-18 | 1.86E-13 |
| Git2 | -0.3443 | 1.74E-22 | 5.63E-18 |
| Tmco1 | -0.34494 | 2.64E-15 | 8.54E-11 |
| Nrip1 | -0.34523 | 1.94E-16 | 6.28E-12 |
| Siglece | -0.34645 | 2.17E-13 | 7E-09 |
| Pla2g4a | -0.3465 | 3.97E-12 | 1.28E-07 |
| Rabac1 | -0.34662 | 1.71E-20 | 5.51E-16 |
| Cadm1 | -0.34665 | 3.23E-08 | 0.001044 |
| Pld1 | -0.3467 | 4.15E-13 | 1.34E-08 |
| Npc1 | -0.34721 | 7.08E-15 | 2.29E-10 |
| Abca1 | -0.34746 | 4.61E-17 | 1.49E-12 |

| | | | |
|---|---|---|---|
| Dglucy | -0.34771 | 7.38E-10 | 2.38E-05 |
| Bmpr2 | -0.34791 | 5.36E-13 | 1.73E-08 |
| Chn2 | -0.34795 | 3.94E-18 | 1.27E-13 |
| Chst7 | -0.34872 | 2.69E-13 | 8.7E-09 |
| Asb2 | -0.34945 | 4.28E-13 | 1.38E-08 |
| Dok3 | -0.35052 | 1.45E-16 | 4.69E-12 |
| Ccl6 | -0.35113 | 7.29E-07 | 0.023539 |
| Comt | -0.35205 | 3.39E-21 | 1.09E-16 |
| Sipa1l1 | -0.35219 | 6.15E-18 | 1.99E-13 |
| Mfng | -0.35229 | 1.03E-18 | 3.31E-14 |
| Ctc1 | -0.3523 | 5.33E-18 | 1.72E-13 |
| Pla2g15 | -0.35234 | 8.75E-21 | 2.83E-16 |
| Csad | -0.35237 | 2.05E-17 | 6.63E-13 |
| Ncoa4 | -0.35242 | 8.49E-17 | 2.74E-12 |
| Hnmt | -0.35341 | 8.75E-27 | 2.83E-22 |
| Gm42418 | -0.35381 | 1.56E-14 | 5.05E-10 |
| Snx17 | -0.35407 | 1.4E-15 | 4.51E-11 |
| Snta1 | -0.35448 | 2.82E-16 | 9.1E-12 |
| Lipa | -0.35733 | 7.7E-17 | 2.49E-12 |
| Csf3r | -0.35917 | 1.13E-16 | 3.64E-12 |
| C5ar2 | -0.3596 | 2.19E-18 | 7.08E-14 |
| Nfam1 | -0.36008 | 3.95E-15 | 1.28E-10 |
| Pald1 | -0.36182 | 1.54E-15 | 4.97E-11 |
| Nav1 | -0.36228 | 1.8E-15 | 5.8E-11 |
| Iffo1 | -0.36271 | 1.53E-15 | 4.95E-11 |
| Twf2 | -0.36299 | 7.12E-19 | 2.3E-14 |
| Man1a | -0.363 | 6.61E-23 | 2.13E-18 |
| Ulk2 | -0.36319 | 1.54E-13 | 4.96E-09 |
| Nr3c1 | -0.36321 | 1.15E-13 | 3.71E-09 |
| Tmem144 | -0.36376 | 4.17E-19 | 1.35E-14 |
| Dusp6 | -0.36398 | 7E-24 | 2.26E-19 |
| Prrc2a | -0.36418 | 9.35E-19 | 3.02E-14 |
| Ptpra | -0.3642 | 1.78E-16 | 5.74E-12 |
| Tal1 | -0.3642 | 3.4E-19 | 1.1E-14 |
| Slc29a1 | -0.36457 | 4.6E-16 | 1.48E-11 |
| Cyb561a3 | -0.36527 | 3.48E-13 | 1.12E-08 |
| Adrb2 | -0.36613 | 1.1E-26 | 3.56E-22 |
| Tmem204 | -0.36676 | 4.24E-36 | 1.37E-31 |
| Sec11c | -0.36795 | 1.8E-21 | 5.81E-17 |
| Nbea | -0.36809 | 7.46E-07 | 0.024086 |
| Mtus1 | -0.36931 | 2.6E-21 | 8.39E-17 |
| Zfp697 | -0.36958 | 1.47E-20 | 4.73E-16 |
| Klhl9 | -0.37308 | 3.96E-21 | 1.28E-16 |
| Dnm2 | -0.37322 | 1.05E-23 | 3.39E-19 |
| Klhl7 | -0.37357 | 8.24E-18 | 2.66E-13 |
| Fcrl1 | -0.37402 | 2.09E-25 | 6.74E-21 |
| Tsc22d4 | -0.37411 | 1.49E-24 | 4.81E-20 |
| Tpp1 | -0.37436 | 2.35E-24 | 7.58E-20 |

| | | | |
|---|---|---|---|
| Rcsd1 | -0.37438 | 3.36E-21 | 1.09E-16 |
| Ccng2 | -0.37467 | 7.39E-20 | 2.39E-15 |
| Tnfrsf17 | -0.37612 | 5.32E-34 | 1.72E-29 |
| Soga1 | -0.37658 | 2.41E-12 | 7.77E-08 |
| Cdh23 | -0.37663 | 9.87E-17 | 3.19E-12 |
| Ggta1 | -0.37729 | 1.01E-18 | 3.24E-14 |
| Adprh | -0.37793 | 5.48E-17 | 1.77E-12 |
| Rps6ka1 | -0.3786 | 5.85E-28 | 1.89E-23 |
| Tm6sf1 | -0.37951 | 2.31E-26 | 7.44E-22 |
| Blnk | -0.38024 | 2.77E-22 | 8.96E-18 |
| Tnfrsf13b | -0.38041 | 8.21E-18 | 2.65E-13 |
| Prkab1 | -0.38079 | 2.4E-18 | 7.76E-14 |
| Nuak1 | -0.3816 | 3.97E-12 | 1.28E-07 |
| Arhgap31 | -0.38222 | 4.56E-10 | 1.47E-05 |
| Slc44a2 | -0.38269 | 1.09E-23 | 3.52E-19 |
| Pou2f2 | -0.38323 | 4.67E-23 | 1.51E-18 |
| Peak1 | -0.38329 | 6.83E-20 | 2.2E-15 |
| Gm2237 | -0.38428 | 1.18E-30 | 3.81E-26 |
| Il16 | -0.38529 | 3.57E-24 | 1.15E-19 |
| Gab1 | -0.38531 | 1.29E-22 | 4.17E-18 |
| Mef2c | -0.38543 | 1.28E-36 | 4.14E-32 |
| Zmiz1 | -0.38688 | 2.86E-20 | 9.24E-16 |
| Acap2 | -0.387 | 1.82E-24 | 5.86E-20 |
| Rnf216 | -0.38729 | 3.81E-21 | 1.23E-16 |
| Gtf2i | -0.38766 | 8.05E-21 | 2.6E-16 |
| Abhd6 | -0.38811 | 2.31E-19 | 7.47E-15 |
| Parvg | -0.38986 | 7.96E-20 | 2.57E-15 |
| Ccr5 | -0.39195 | 2.09E-25 | 6.74E-21 |
| Sgce | -0.39227 | 3.02E-19 | 9.76E-15 |
| Gcn1 | -0.39257 | 9.03E-19 | 2.92E-14 |
| Tmem44 | -0.39347 | 5.28E-35 | 1.71E-30 |
| Bach1 | -0.39402 | 1.4E-21 | 4.51E-17 |
| Tmem273 | -0.39417 | 7.49E-19 | 2.42E-14 |
| Gm4258 | -0.39456 | 4.6E-22 | 1.48E-17 |
| Kcnip3 | -0.39492 | 9.81E-21 | 3.17E-16 |
| Gm3468 | -0.3969 | 1.57E-37 | 5.08E-33 |
| Ptpre | -0.39949 | 1.25E-18 | 4.03E-14 |
| Snx18 | -0.40017 | 1.7E-31 | 5.49E-27 |
| Lrp1 | -0.40018 | 1.12E-27 | 3.62E-23 |
| Med12l | -0.40026 | 3.78E-19 | 1.22E-14 |
| Nkain2 | -0.40057 | 2.65E-42 | 8.55E-38 |
| Klf3 | -0.40082 | 2.32E-20 | 7.5E-16 |
| Timp2 | -0.401 | 2.26E-24 | 7.28E-20 |
| Specc1 | -0.4013 | 2.18E-15 | 7.03E-11 |
| Nid2 | -0.40159 | 3.97E-38 | 1.28E-33 |
| Syngr1 | -0.40165 | 3.3E-29 | 1.06E-24 |
| Saraf | -0.40215 | 3.19E-34 | 1.03E-29 |
| Lpar6 | -0.40254 | 2.31E-17 | 7.46E-13 |

| | | | |
|---|---|---|---|
| Dgkd | -0.40312 | 1.77E-21 | 5.73E-17 |
| Gnb2 | -0.40319 | 7.46E-24 | 2.41E-19 |
| Ptpro | -0.40326 | 6.13E-22 | 1.98E-17 |
| Sft2d2 | -0.40418 | 3.31E-27 | 1.07E-22 |
| Tlr5 | -0.40501 | 1.84E-43 | 5.94E-39 |
| Tmem100 | -0.40528 | 7.02E-23 | 2.26E-18 |
| Pacc1 | -0.4058 | 4.11E-25 | 1.33E-20 |
| Gna15 | -0.40666 | 2.57E-27 | 8.29E-23 |
| Rptor | -0.40708 | 1.06E-19 | 3.41E-15 |
| Olfml2b | -0.40767 | 1.35E-25 | 4.36E-21 |
| Msrb2 | -0.40795 | 2.35E-30 | 7.57E-26 |
| Plod3 | -0.40936 | 1.48E-24 | 4.77E-20 |
| Gabbr1 | -0.40957 | 3.47E-35 | 1.12E-30 |
| Lyzl4 | -0.40981 | 3.37E-25 | 1.09E-20 |
| Zfp69 | -0.41112 | 1.31E-21 | 4.23E-17 |
| Ighm | -0.41295 | 8.88E-22 | 2.87E-17 |
| Edem2 | -0.41311 | 1.38E-25 | 4.46E-21 |
| Mlxipl | -0.41339 | 8.81E-26 | 2.84E-21 |
| Cd53 | -0.41471 | 2.25E-41 | 7.26E-37 |
| Arhgap27 | -0.41521 | 4.83E-24 | 1.56E-19 |
| Lyl1 | -0.41656 | 9.28E-22 | 2.99E-17 |
| Elk3 | -0.41741 | 7.69E-15 | 2.48E-10 |
| Zfp652 | -0.41765 | 6.71E-19 | 2.17E-14 |
| Ephx1 | -0.42034 | 7.05E-24 | 2.28E-19 |
| Ints6l | -0.42036 | 1.09E-19 | 3.53E-15 |
| Gm2629 | -0.4206 | 6.73E-24 | 2.17E-19 |
| Exoc6b | -0.4211 | 1.07E-14 | 3.44E-10 |
| Fmnl1 | -0.42192 | 5.48E-25 | 1.77E-20 |
| St6gal1 | -0.42273 | 2.37E-29 | 7.64E-25 |
| Man1c1 | -0.42331 | 8.6E-21 | 2.77E-16 |
| Rgs2 | -0.42534 | 2.26E-24 | 7.3E-20 |
| Phyhd1 | -0.42567 | 2.5E-26 | 8.08E-22 |
| Gm40841 | -0.42579 | 2.59E-35 | 8.35E-31 |
| Dock8 | -0.4258 | 6.85E-27 | 2.21E-22 |
| Myo18b | -0.42648 | 2.25E-39 | 7.28E-35 |
| Ubash3b | -0.42681 | 6.58E-19 | 2.13E-14 |
| Garnl3 | -0.42709 | 3.18E-17 | 1.03E-12 |
| Rock2 | -0.42819 | 1.05E-20 | 3.4E-16 |
| Asap1 | -0.42845 | 1.13E-18 | 3.66E-14 |
| Lrrc25 | -0.42911 | 1.52E-24 | 4.9E-20 |
| Plod1 | -0.42916 | 2.66E-26 | 8.58E-22 |
| Grap | -0.42978 | 2.98E-24 | 9.62E-20 |
| G530011O | -0.43001 | 2.32E-19 | 7.49E-15 |
| Oxr1 | -0.43015 | 2.96E-15 | 9.57E-11 |
| Nfia | -0.43101 | 1.69E-20 | 5.47E-16 |
| Hfe | -0.43107 | 1.62E-27 | 5.21E-23 |
| Pmepa1 | -0.43136 | 2.43E-18 | 7.84E-14 |
| Nckap1l | -0.43166 | 1.14E-26 | 3.68E-22 |

| | | | |
|---|---|---|---|
| Snx29 | -0.43179 | 2.91E-23 | 9.4E-19 |
| Gm33699 | -0.43306 | 1.02E-28 | 3.29E-24 |
| Ggt5 | -0.43407 | 4.97E-31 | 1.6E-26 |
| Ivns1abp | -0.43511 | 3.13E-35 | 1.01E-30 |
| Csnk1e | -0.43627 | 1.39E-22 | 4.48E-18 |
| Ddah2 | -0.43866 | 1.79E-38 | 5.78E-34 |
| Fli1 | -0.43908 | 1.57E-38 | 5.08E-34 |
| Myadm | -0.43927 | 1.81E-24 | 5.86E-20 |
| Slc46a1 | -0.43942 | 2.63E-20 | 8.49E-16 |
| Serpinf1 | -0.44067 | 4.44E-26 | 1.43E-21 |
| Hexb | -0.44243 | 4.3E-151 | 1.4E-146 |
| Tspan18 | -0.44252 | 2.42E-31 | 7.82E-27 |
| Myo1b | -0.44281 | 5.57E-22 | 1.8E-17 |
| Ptp4a3 | -0.44756 | 4.02E-23 | 1.3E-18 |
| Dip2b | -0.44868 | 9.17E-23 | 2.96E-18 |
| Abhd12 | -0.44884 | 1.65E-60 | 5.33E-56 |
| Comtd1 | -0.4493 | 7.8E-27 | 2.52E-22 |
| Gm34589 | -0.45024 | 6.84E-34 | 2.21E-29 |
| Dlc1 | -0.45103 | 6.13E-23 | 1.98E-18 |
| Fmnl3 | -0.45165 | 4.73E-31 | 1.53E-26 |
| Ppp1r9a | -0.45217 | 4.55E-22 | 1.47E-17 |
| Whrn | -0.45225 | 1.23E-18 | 3.96E-14 |
| Atp8a2 | -0.45293 | 8.51E-22 | 2.75E-17 |
| Nptn | -0.4533 | 1.53E-35 | 4.95E-31 |
| Hdac9 | -0.45344 | 3.2E-14 | 1.03E-09 |
| Itga6 | -0.45419 | 1.49E-28 | 4.82E-24 |
| Numa1 | -0.45484 | 1.07E-24 | 3.44E-20 |
| Tfe3 | -0.4549 | 1.15E-25 | 3.71E-21 |
| Zfp706 | -0.45526 | 1.82E-26 | 5.87E-22 |
| Gas6 | -0.45529 | 6.03E-24 | 1.95E-19 |
| Slc7a8 | -0.45532 | 1.32E-31 | 4.27E-27 |
| Gm42047 | -0.45834 | 0.00049 | 1 |
| Tgfb1 | -0.45974 | 5.23E-37 | 1.69E-32 |
| Ctsf | -0.46144 | 7.4E-34 | 2.39E-29 |
| Thrsp | -0.4617 | 2.12E-44 | 6.85E-40 |
| Pwwp2a | -0.46264 | 1.83E-23 | 5.9E-19 |
| Tnfrsf21 | -0.46309 | 2.97E-35 | 9.59E-31 |
| Tgfbr1 | -0.46485 | 1.42E-49 | 4.59E-45 |
| St3gal5 | -0.46504 | 2E-26 | 6.45E-22 |
| Gm48099 | -0.46605 | 3.86E-18 | 1.25E-13 |
| Itgb5 | -0.46606 | 2.09E-42 | 6.74E-38 |
| Ttc28 | -0.46757 | 3.31E-21 | 1.07E-16 |
| Adam15 | -0.46793 | 1.47E-32 | 4.76E-28 |
| Fat3 | -0.4708 | 5.09E-15 | 1.64E-10 |
| Sqor | -0.47294 | 8.27E-33 | 2.67E-28 |
| Sncaip | -0.47401 | 4.04E-29 | 1.31E-24 |
| Mtdh | -0.47455 | 3.19E-60 | 1.03E-55 |
| Rnf13 | -0.47477 | 2.47E-50 | 7.99E-46 |

| | | | |
|---|---|---|---|
| Prkce | -0.47553 | 4.93E-15 | 1.59E-10 |
| Bin2 | -0.47582 | 8.53E-46 | 2.75E-41 |
| Eif4g2 | -0.47622 | 1.88E-36 | 6.06E-32 |
| Ifngr1 | -0.47727 | 1.2E-65 | 3.87E-61 |
| Tmem135 | -0.47734 | 2.31E-34 | 7.46E-30 |
| Ypel3 | -0.4781 | 9.8E-29 | 3.16E-24 |
| Lpcat3 | -0.48204 | 2.52E-32 | 8.15E-28 |
| A830008E2 | -0.48239 | 4.56E-20 | 1.47E-15 |
| Commd8 | -0.4824 | 1.61E-33 | 5.19E-29 |
| Ppm1l | -0.48394 | 1.55E-26 | 4.99E-22 |
| Gtf2h2 | -0.48406 | 9.62E-39 | 3.11E-34 |
| Irf2bpl | -0.48487 | 3.39E-25 | 1.09E-20 |
| Abl1 | -0.48644 | 2.8E-26 | 9.05E-22 |
| Rreb1 | -0.48783 | 2.96E-27 | 9.56E-23 |
| Frmd4a | -0.48874 | 1.21E-46 | 3.89E-42 |
| Tuba1a | -0.48888 | 1.61E-24 | 5.21E-20 |
| Zdhhc14 | -0.48919 | 8.3E-25 | 2.68E-20 |
| Eef2k | -0.4893 | 8.74E-29 | 2.82E-24 |
| Gm6277 | -0.49259 | 9E-36 | 2.91E-31 |
| Cep170 | -0.49321 | 1.45E-30 | 4.69E-26 |
| Rtn1 | -0.4933 | 1.01E-25 | 3.25E-21 |
| Exoc4 | -0.49352 | 1.59E-24 | 5.13E-20 |
| Glul | -0.49354 | 5.13E-57 | 1.66E-52 |
| Plcl1 | -0.4939 | 3.77E-19 | 1.22E-14 |
| Magi1 | -0.49464 | 7.34E-15 | 2.37E-10 |
| Pard3b | -0.49537 | 1.67E-20 | 5.38E-16 |
| Asph | -0.4967 | 7.57E-36 | 2.44E-31 |
| Khdrbs3 | -0.49742 | 7.75E-44 | 2.5E-39 |
| Wasf2 | -0.49923 | 1.03E-54 | 3.32E-50 |
| Pip4k2a | -0.49953 | 3.75E-35 | 1.21E-30 |
| 1700017B0 | -0.50179 | 8.02E-41 | 2.59E-36 |
| Smap2 | -0.50216 | 4.73E-48 | 1.53E-43 |
| AC149090. | -0.50391 | 2.72E-38 | 8.79E-34 |
| Kcnk12 | -0.50453 | 1.26E-48 | 4.07E-44 |
| Eng | -0.50658 | 9.38E-32 | 3.03E-27 |
| Ctsd | -0.50734 | 7.1E-84 | 2.29E-79 |
| Fry | -0.50934 | 8.27E-32 | 2.67E-27 |
| Col27a1 | -0.51054 | 7.6E-33 | 2.45E-28 |
| Slc12a2 | -0.51106 | 2.06E-28 | 6.65E-24 |
| Pros1 | -0.51161 | 1.39E-46 | 4.5E-42 |
| Slc16a6 | -0.51241 | 7.58E-36 | 2.45E-31 |
| Klhdc8b | -0.51496 | 7.11E-57 | 2.29E-52 |
| Mid1ip1 | -0.51581 | 3.51E-39 | 1.13E-34 |
| Cltc | -0.51604 | 8.59E-49 | 2.77E-44 |
| Dhrs3 | -0.51713 | 9.29E-36 | 3E-31 |
| Upk1b | -0.5179 | 4.61E-16 | 1.49E-11 |
| Otulinl | -0.51856 | 4.63E-60 | 1.5E-55 |
| Trp53cor1 | -0.51921 | 1.57E-30 | 5.06E-26 |

| | | | |
|---|---|---|---|
| Mef2a | -0.52181 | 3.42E-69 | 1.1E-64 |
| Olfml3 | -0.52408 | 9.58E-60 | 3.09E-55 |
| Rab3il1 | -0.52475 | 7.76E-43 | 2.5E-38 |
| Kif21b | -0.52573 | 5.14E-28 | 1.66E-23 |
| Slc46a3 | -0.52684 | 1.91E-32 | 6.18E-28 |
| Cyth4 | -0.52844 | 1.78E-75 | 5.74E-71 |
| Arhgap45 | -0.52942 | 5.53E-52 | 1.78E-47 |
| Etv5 | -0.53021 | 2.71E-37 | 8.74E-33 |
| Mtss1 | -0.53185 | 3.99E-35 | 1.29E-30 |
| Slco2b1 | -0.5333 | 9.52E-63 | 3.07E-58 |
| Phf14 | -0.53334 | 1.54E-31 | 4.96E-27 |
| Kcnma1 | -0.5336 | 1.17E-18 | 3.77E-14 |
| Idh2 | -0.53698 | 6.67E-41 | 2.15E-36 |
| P3h2 | -0.53752 | 2.62E-30 | 8.47E-26 |
| Mgll | -0.53764 | 3.55E-40 | 1.15E-35 |
| Cep68 | -0.53924 | 1.89E-36 | 6.12E-32 |
| Cmtm6 | -0.53958 | 2.15E-51 | 6.94E-47 |
| Cask | -0.54045 | 1.37E-33 | 4.41E-29 |
| Txnip | -0.54164 | 1.34E-49 | 4.32E-45 |
| Gal3st4 | -0.54185 | 1.79E-37 | 5.77E-33 |
| Gm10790 | -0.54311 | 3.84E-43 | 1.24E-38 |
| Evi2a | -0.54329 | 5.25E-43 | 1.69E-38 |
| Ctnnd1 | -0.54457 | 6.03E-36 | 1.95E-31 |
| Pde3b | -0.54498 | 3.92E-46 | 1.27E-41 |
| Abcc3 | -0.54645 | 2.99E-36 | 9.67E-32 |
| Cd300a | -0.54745 | 4.07E-33 | 1.31E-28 |
| Mctp1 | -0.55148 | 2.23E-23 | 7.21E-19 |
| Csf1r | -0.55252 | 1.1E-164 | 3.5E-160 |
| F11r | -0.55252 | 4.91E-72 | 1.59E-67 |
| Serinc3 | -0.55317 | 2.3E-136 | 7.3E-132 |
| Rnf167 | -0.55849 | 1.09E-45 | 3.52E-41 |
| Gpr165 | -0.56352 | 3.58E-63 | 1.16E-58 |
| Inpp4b | -0.56521 | 1.92E-64 | 6.2E-60 |
| Cyfip1 | -0.56577 | 4.12E-85 | 1.33E-80 |
| Bbs9 | -0.56589 | 3.51E-26 | 1.13E-21 |
| Epb41l2 | -0.56741 | 2.17E-75 | 7.01E-71 |
| Gpr155 | -0.57133 | 6.98E-42 | 2.25E-37 |
| Tmem119 | -0.57214 | 1.2E-86 | 3.88E-82 |
| Fam110a | -0.5732 | 3.08E-45 | 9.95E-41 |
| Itga9 | -0.57378 | 7.07E-32 | 2.28E-27 |
| 2610203C2 | -0.57418 | 2.29E-33 | 7.38E-29 |
| Ldlrad4 | -0.57514 | 9.48E-43 | 3.06E-38 |
| Cryl1 | -0.57599 | 6.74E-36 | 2.18E-31 |
| Mid1 | -0.57715 | 1.27E-34 | 4.11E-30 |
| Tjp1 | -0.57807 | 6.86E-41 | 2.21E-36 |
| Jarid2 | -0.57849 | 2.7E-55 | 8.72E-51 |
| Gm26542 | -0.57854 | 8.8E-29 | 2.84E-24 |
| Apbb1ip | -0.57904 | 1.2E-101 | 3.84E-97 |

| | | | |
|---|---|---|---|
| Gpr183 | -0.58109 | 1.15E-52 | 3.72E-48 |
| Usp2 | -0.58352 | 3.53E-47 | 1.14E-42 |
| Scamp5 | -0.58472 | 5.82E-41 | 1.88E-36 |
| Arhgap5 | -0.58484 | 1.37E-64 | 4.42E-60 |
| Cttnbp2nl | -0.59085 | 2.75E-61 | 8.88E-57 |
| Arap3 | -0.59153 | 3.06E-52 | 9.88E-48 |
| Gp9 | -0.59434 | 3.77E-68 | 1.22E-63 |
| Camk1 | -0.5972 | 8.66E-64 | 2.8E-59 |
| Jam2 | -0.60253 | 2.08E-46 | 6.7E-42 |
| Hexa | -0.60539 | 4.81E-74 | 1.55E-69 |
| Pdgfb | -0.60891 | 1.62E-42 | 5.22E-38 |
| 4933406I1 | -0.61493 | 2.31E-42 | 7.47E-38 |
| Sirpa | -0.61506 | 2.3E-102 | 7.37E-98 |
| Plxnb2 | -0.6153 | 3.06E-50 | 9.89E-46 |
| Rgs10 | -0.61676 | 5.2E-89 | 1.68E-84 |
| Slc9a9 | -0.61735 | 1.36E-55 | 4.38E-51 |
| Nos1ap | -0.61764 | 6.12E-32 | 1.97E-27 |
| Chd9 | -0.61944 | 5.07E-85 | 1.64E-80 |
| Mylip | -0.62078 | 3.74E-58 | 1.21E-53 |
| Bin1 | -0.6209 | 2.18E-88 | 7.02E-84 |
| Sgk1 | -0.62103 | 4.2E-61 | 1.36E-56 |
| Rasgrp3 | -0.62108 | 9.05E-54 | 2.92E-49 |
| Gm3739 | -0.62219 | 6.8E-71 | 2.2E-66 |
| Ptprm | -0.63052 | 4.44E-23 | 1.43E-18 |
| Trem2 | -0.63184 | 1.4E-110 | 4.6E-106 |
| Cd164 | -0.64327 | 5.94E-67 | 1.92E-62 |
| Scoc | -0.64595 | 3.05E-54 | 9.86E-50 |
| 0610040J0 | -0.64973 | 4.52E-58 | 1.46E-53 |
| Klf12 | -0.65348 | 4.16E-37 | 1.34E-32 |
| Hpgds | -0.65468 | 1.13E-83 | 3.66E-79 |
| Rapgef5 | -0.65585 | 1.06E-57 | 3.43E-53 |
| Adamts16 | -0.66022 | 2.92E-72 | 9.44E-68 |
| Rnaset2b | -0.66337 | 1.46E-64 | 4.71E-60 |
| Qk | -0.66373 | 2.34E-73 | 7.54E-69 |
| Gm5086 | -0.66738 | 9.58E-48 | 3.09E-43 |
| Fhit | -0.67024 | 6.58E-30 | 2.12E-25 |
| Cd68 | -0.67267 | 1.16E-95 | 3.76E-91 |
| Sft2d1 | -0.67613 | 1E-99 | 3.28E-95 |
| Zfhx3 | -0.67789 | 6.8E-80 | 2.2E-75 |
| Sall1 | -0.68168 | 2.93E-64 | 9.45E-60 |
| Ltc4s | -0.68581 | 1.59E-73 | 5.14E-69 |
| Plcl2 | -0.69481 | 8.92E-51 | 2.88E-46 |
| Siglech | -0.70006 | 1.1E-103 | 3.7E-99 |
| Tanc2 | -0.7043 | 8.74E-87 | 2.82E-82 |
| Snn | -0.70525 | 1.17E-67 | 3.77E-63 |
| Sdk1 | -0.70609 | 4.5E-21 | 1.45E-16 |
| Maml3 | -0.71037 | 3.14E-55 | 1.01E-50 |
| Agmo | -0.71076 | 3.87E-61 | 1.25E-56 |

| | | | |
|---|---|---|---|
| D830030K: | -0.71493 | 4.97E-75 | 1.6E-70 |
| Basp1 | -0.71582 | 4.1E-110 | 1.3E-105 |
| Arhgef40 | -0.71826 | 8.29E-64 | 2.68E-59 |
| Mir99ahg | -0.72716 | 4.03E-67 | 1.3E-62 |
| Fchsd2 | -0.73909 | 2.02E-62 | 6.52E-58 |
| Il6st | -0.74065 | 3.18E-63 | 1.03E-58 |
| Ckb | -0.7438 | 1.4E-102 | 4.58E-98 |
| Plxna4 | -0.74644 | 6.75E-37 | 2.18E-32 |
| Slc40a1 | -0.75423 | 8.64E-72 | 2.79E-67 |
| Serpine2 | -0.75476 | 5.89E-97 | 1.9E-92 |
| St3gal6 | -0.75509 | 6E-99 | 1.92E-94 |
| Srgap2 | -0.75541 | 5.4E-113 | 1.8E-108 |
| Abca9 | -0.75734 | 6.69E-89 | 2.16E-84 |
| Marcks | -0.75991 | 1E-159 | 3.3E-155 |
| Rnase4 | -0.76942 | 5.7E-99 | 1.83E-94 |
| Sparc | -0.77025 | 6.6E-150 | 2.1E-145 |
| Cxxc5 | -0.77109 | 1.22E-79 | 3.93E-75 |
| Fam102b | -0.77149 | 1.54E-68 | 4.96E-64 |
| Zfp710 | -0.77337 | 5.25E-55 | 1.7E-50 |
| Vsir | -0.77507 | 3.1E-152 | 1E-147 |
| Prkn | -0.78715 | 3.6E-51 | 1.16E-46 |
| Ifi27 | -0.80462 | 2.85E-63 | 9.2E-59 |
| Susd3 | -0.80638 | 5.62E-96 | 1.81E-91 |
| Rtn4rl1 | -0.81826 | 8.63E-96 | 2.79E-91 |
| Tmem176a | -0.81865 | 7.12E-65 | 2.3E-60 |
| Calm2 | -0.83628 | 6.6E-106 | 2.1E-101 |
| Nav3 | -0.83824 | 1.66E-75 | 5.34E-71 |
| Plxdc2 | -0.84679 | 2.3E-153 | 7.5E-149 |
| Sox4 | -0.85864 | 2.1E-79 | 6.79E-75 |
| Elmo1 | -0.86367 | 5.9E-139 | 1.9E-134 |
| Adgrg1 | -0.86368 | 2.27E-94 | 7.33E-90 |
| Bank1 | -0.87821 | 1.53E-70 | 4.93E-66 |
| Pmp22 | -0.89087 | 1.1E-108 | 3.6E-104 |
| Slc2a5 | -0.89236 | 6.42E-85 | 2.07E-80 |
| Lrba | -0.89836 | 7.81E-95 | 2.52E-90 |
| Ifitm10 | -0.90419 | 4.69E-95 | 1.51E-90 |
| Selplg | -0.93551 | 5.6E-227 | 1.8E-222 |
| Maf | -0.94589 | 1.3E-122 | 4.1E-118 |
| Numb | -0.96895 | 7E-145 | 2.3E-140 |
| Cfh | -0.97188 | 8.3E-166 | 2.7E-161 |
| Arsb | -0.97925 | 2.7E-112 | 8.6E-108 |
| Selenop | -0.98951 | 2.4E-165 | 7.6E-161 |
| Golm1 | -1.00205 | 1E-166 | 3.4E-162 |
| Cx3cr1 | -1.02735 | 1.5E-249 | 4.8E-245 |
| 8030442B( | -1.03455 | 7.9E-111 | 2.5E-106 |
| P2ry13 | -1.04686 | 7.2E-149 | 2.3E-144 |
| Gm19951 | -1.0733 | 6.07E-52 | 1.96E-47 |
| Cst3 | -1.16119 | 1.7E-300 | 5.6E-296 |

| | | | |
|---|---|---|---|
| Tmem176b | -1.20156 | 7.4E-139 | 2.4E-134 |
| Ecscr | -1.21566 | 1.7E-157 | 5.6E-153 |
| Nav2 | -1.24292 | 2.6E-138 | 8.4E-134 |
| Hpgd | -1.25788 | 1.7E-169 | 5.4E-165 |
| Gpr34 | -1.46005 | 3E-243 | 9.8E-239 |
| Fscn1 | -1.49179 | 1.9E-213 | 6.2E-209 |
| P2ry12 | -1.5932 | 0 | 0 |
| Csmd3 | -1.63133 | 8.3E-158 | 2.7E-153 |
| Fah | -1.64448 | 4.3E-300 | 1.4E-295 |
| Ttr | -1.74041 | 2.44E-32 | 7.88E-28 |
| Fcrls | -2.35806 | 0 | 0 |

#### SUPPLEMENTAL TABLE 4. Differentially Regulated Genes in WT versus CB2R$^{-/-}$ Microglia

| Gene | avg_log2FC | p_val | p_val_adj |
|---|---|---|---|
| Gm47283 | 0.523913 | 7.8E-67 | 2.52E-62 |
| Mir142hg | 0.408831 | 1.58E-54 | 5.1E-50 |
| Zfp36 | 0.406314 | 1.73E-38 | 5.6E-34 |
| G530011O06Rik | 0.40226 | 2.34E-59 | 7.56E-55 |
| H3f3b | 0.400692 | 1.16E-48 | 3.73E-44 |
| Uba52 | 0.368005 | 1.34E-65 | 4.33E-61 |
| Atf3 | 0.364562 | 6.72E-22 | 2.17E-17 |
| Jund | 0.352425 | 4.04E-38 | 1.3E-33 |
| Cst3 | 0.349709 | 1.4E-72 | 4.53E-68 |
| Fos | 0.349141 | 2.25E-16 | 7.26E-12 |
| Hist1h1e | 0.344081 | 3.96E-43 | 1.28E-38 |
| Gm10076 | 0.325133 | 4.46E-48 | 1.44E-43 |
| Zfp36l1 | 0.314942 | 1.66E-16 | 5.37E-12 |
| Gm34455 | 0.308338 | 4.28E-28 | 1.38E-23 |
| Jun | 0.304113 | 3.04E-20 | 9.81E-16 |
| Hist1h2bc | 0.299136 | 4.31E-13 | 1.39E-08 |
| Ccl4 | 0.298441 | 4.81E-08 | 0.001551 |
| Zfand5 | 0.277478 | 2.85E-21 | 9.21E-17 |
| Hist1h2ac | 0.260357 | 3.69E-20 | 1.19E-15 |
| Hist1h1c | 0.259677 | 4.76E-30 | 1.54E-25 |
| Olfml3 | 0.258959 | 2.86E-25 | 9.25E-21 |
| Ltc4s | 0.257464 | 4.36E-18 | 1.41E-13 |
| Timp2 | 0.249441 | 2.45E-26 | 7.9E-22 |
| Btg2 | 0.248186 | 3.9E-10 | 1.26E-05 |
| Tnf | 0.24434 | 5.98E-12 | 1.93E-07 |
| Cd63 | 0.244056 | 2.49E-14 | 8.03E-10 |
| Ccl5 | 0.239295 | 0.900621 | 1 |
| Gpr34 | 0.235808 | 7.14E-20 | 2.3E-15 |
| Tyrobp | 0.229426 | 6.24E-40 | 2.01E-35 |
| mt-Nd4l | 0.227318 | 2.13E-20 | 6.87E-16 |
| Fosb | 0.223403 | 4.91E-23 | 1.58E-18 |
| Trem2 | 0.222643 | 5.86E-33 | 1.89E-28 |
| Mid1 | 0.221497 | 2.95E-26 | 9.52E-22 |
| Nfkbiz | 0.217497 | 2.14E-11 | 6.9E-07 |
| Ctsd | 0.216533 | 1.8E-22 | 5.81E-18 |
| Cd9 | 0.212324 | 7.28E-23 | 2.35E-18 |
| Ccl3 | 0.211851 | 1.21E-09 | 3.92E-05 |
| Fcrls | 0.208982 | 1.29E-17 | 4.18E-13 |
| Siglech | 0.208899 | 7.08E-21 | 2.29E-16 |
| Rgs1 | 0.205949 | 2.26E-07 | 0.007309 |
| Lyz2 | 0.20521 | 4.23E-15 | 1.37E-10 |
| Tmx4 | 0.203751 | 4.99E-10 | 1.61E-05 |
| Ier5 | 0.202569 | 5.29E-18 | 1.71E-13 |

| | | | |
|---|---|---|---|
| Sqstm1 | 0.198842 | 6.39E-15 | 2.06E-10 |
| Selenop | 0.197911 | 7.11E-15 | 2.3E-10 |
| Hexb | 0.197257 | 1.77E-52 | 5.72E-48 |
| mt-Nd5 | 0.196784 | 2.79E-20 | 9.02E-16 |
| 2810013P06Rik | 0.196661 | 7.97E-21 | 2.57E-16 |
| Ddx3x | 0.195149 | 1.41E-09 | 4.55E-05 |
| Fcer1g | 0.19499 | 7.03E-29 | 2.27E-24 |
| Il1a | 0.194291 | 2.95E-10 | 9.52E-06 |
| C1qb | 0.194129 | 5.04E-39 | 1.63E-34 |
| Junb | 0.193173 | 1.04E-09 | 3.36E-05 |
| Btg1 | 0.19146 | 2.25E-14 | 7.26E-10 |
| Pmepa1 | 0.184971 | 1.31E-15 | 4.22E-11 |
| E330020D12Rik | 0.184363 | 1.06E-07 | 0.003412 |
| Rhob | 0.184118 | 2.78E-13 | 8.97E-09 |
| Rpl35 | 0.183194 | 7E-19 | 2.26E-14 |
| Hsp90aa1 | 0.182547 | 1.19E-12 | 3.84E-08 |
| Apoe | 0.182246 | 1.46E-06 | 0.047062 |
| Rpl37a | 0.181229 | 5.31E-25 | 1.72E-20 |
| Prkg1 | 0.181191 | 4.38E-06 | 0.141529 |
| Rgs2 | 0.177521 | 7.57E-10 | 2.44E-05 |
| Itm2b | 0.176651 | 4.14E-32 | 1.34E-27 |
| Mafb | 0.176199 | 3.37E-13 | 1.09E-08 |
| Hist1h2ap | 0.174716 | 2.77E-18 | 8.94E-14 |
| Tpt1 | 0.174694 | 1.19E-31 | 3.83E-27 |
| Egr1 | 0.174648 | 0.025553 | 1 |
| Cd68 | 0.174196 | 2.7E-14 | 8.72E-10 |
| Tent5c | 0.174046 | 3.05E-12 | 9.85E-08 |
| Fam102b | 0.173482 | 1.4E-12 | 4.53E-08 |
| Rps27 | 0.172116 | 9.8E-20 | 3.16E-15 |
| Plek | 0.171905 | 4.83E-13 | 1.56E-08 |
| Kcnma1 | 0.171196 | 4.07E-11 | 1.31E-06 |
| Rpl36a | 0.170757 | 5.28E-18 | 1.7E-13 |
| Ccl12 | 0.169747 | 0.0004 | 1 |
| Prxl2c | 0.169734 | 9.81E-07 | 0.03166 |
| Rps21 | 0.169407 | 3.03E-22 | 9.77E-18 |
| Rps15a | 0.168202 | 3.49E-22 | 1.13E-17 |
| Sgk1 | 0.16377 | 1.13E-05 | 0.365475 |
| Ctsl | 0.163008 | 2.52E-19 | 8.13E-15 |
| Glul | 0.16101 | 0.034026 | 1 |
| Cd83 | 0.160681 | 0.002593 | 1 |
| Creg1 | 0.160357 | 4.97E-16 | 1.6E-11 |
| Actg1 | 0.159525 | 4.5E-13 | 1.45E-08 |
| Ctsz | 0.158701 | 1.55E-21 | 5.02E-17 |
| mt-Nd2 | 0.155615 | 6.03E-14 | 1.95E-09 |
| Gnas | 0.154175 | 2.84E-10 | 9.18E-06 |
| Cx3cr1 | 0.15398 | 2.4E-17 | 7.74E-13 |
| Ctsb | 0.153554 | 9.38E-13 | 3.03E-08 |
| Selenok | 0.15283 | 1.9E-14 | 6.13E-10 |

| | | | |
|---|---|---|---|
| Rpl39 | 0.152724 | 6.95E-20 | 2.24E-15 |
| mt-Cytb | 0.151846 | 6.57E-14 | 2.12E-09 |
| Rasgef1b | 0.151775 | 1.06E-10 | 3.41E-06 |
| Rpl30 | 0.151622 | 6.78E-20 | 2.19E-15 |
| Cd81 | 0.150558 | 2.75E-17 | 8.89E-13 |
| Ubb | 0.150413 | 2.09E-17 | 6.76E-13 |
| Ppib | 0.149814 | 2.34E-12 | 7.57E-08 |
| Lgmn | 0.148436 | 9E-26 | 2.9E-21 |
| Fau | 0.148144 | 8.82E-19 | 2.85E-14 |
| Rps19 | 0.147252 | 3.96E-17 | 1.28E-12 |
| Sparc | 0.147164 | 7.05E-12 | 2.28E-07 |
| Actb | 0.145372 | 2.18E-13 | 7.02E-09 |
| Ssr4 | 0.144988 | 3.91E-13 | 1.26E-08 |
| Ighm | 0.144269 | 1.73E-10 | 5.58E-06 |
| Ckb | 0.144237 | 1.82E-11 | 5.86E-07 |
| Gas6 | 0.143671 | 1.69E-07 | 0.005468 |
| Syngr1 | 0.143598 | 3.78E-09 | 0.000122 |
| mt-Nd4 | 0.143475 | 7.77E-15 | 2.51E-10 |
| Hist1h3d | 0.143372 | 5.24E-17 | 1.69E-12 |
| Gm15726 | 0.142472 | 3.73E-17 | 1.2E-12 |
| Arl6ip1 | 0.141943 | 2.16E-11 | 6.96E-07 |
| Ptma | 0.141514 | 2E-18 | 6.45E-14 |
| Rplp1 | 0.140916 | 1.41E-18 | 4.55E-14 |
| Gpr183 | 0.140771 | 8.88E-06 | 0.286711 |
| Cxcl9 | 0.140579 | 0.924126 | 1 |
| Lamp1 | 0.139146 | 4.71E-14 | 1.52E-09 |
| C1qc | 0.138154 | 6.44E-21 | 2.08E-16 |
| Npc2 | 0.137632 | 7E-15 | 2.26E-10 |
| P2ry12 | 0.137234 | 4.23E-11 | 1.36E-06 |
| Rpl13 | 0.136625 | 8.31E-16 | 2.68E-11 |
| Smad7 | 0.136515 | 2.64E-08 | 0.000854 |
| Atp8a2 | 0.136323 | 8.76E-10 | 2.83E-05 |
| Rps12 | 0.136218 | 1.27E-17 | 4.09E-13 |
| Ubc | 0.13599 | 1.34E-16 | 4.31E-12 |
| Fcgr3 | 0.135367 | 2.03E-09 | 6.57E-05 |
| Abhd12 | 0.134656 | 6.13E-12 | 1.98E-07 |
| Hsp90b1 | 0.134425 | 1.06E-10 | 3.42E-06 |
| Rnase4 | 0.13404 | 1.68E-09 | 5.42E-05 |
| Nav3 | 0.133347 | 3.01E-06 | 0.097334 |
| St3gal6 | 0.133325 | 1.39E-07 | 0.004473 |
| Rpl36 | 0.132634 | 9.59E-13 | 3.1E-08 |
| Cap1 | 0.132049 | 1.3E-13 | 4.18E-09 |
| Rps24 | 0.131813 | 1.97E-16 | 6.36E-12 |
| Rpl11 | 0.131259 | 2.11E-14 | 6.82E-10 |
| Zeb2os | 0.130648 | 1.17E-09 | 3.79E-05 |
| Tmem258 | 0.130315 | 1.28E-09 | 4.14E-05 |
| Hist1h1d | 0.13006 | 4.64E-15 | 1.5E-10 |
| Rps26 | 0.129907 | 2.18E-11 | 7.05E-07 |

| | | | |
|---|---|---|---|
| Kctd12 | 0.129393 | 1.96E-11 | 6.32E-07 |
| Bhlhe41 | 0.128491 | 6.53E-09 | 0.000211 |
| 4933421O10Rik | 0.127689 | 3.7E-12 | 1.2E-07 |
| Hmgb2 | 0.126986 | 2.23E-10 | 7.21E-06 |
| Rnasek | 0.126826 | 2.27E-07 | 0.007344 |
| Rpl34 | 0.126473 | 3.24E-14 | 1.05E-09 |
| Pfdn5 | 0.126125 | 2.85E-11 | 9.21E-07 |
| Ptms | 0.126108 | 0.000398 | 1 |
| Klf4 | 0.12597 | 8.67E-11 | 2.8E-06 |
| Rps7 | 0.12559 | 2.15E-14 | 6.93E-10 |
| Socs3 | 0.125497 | 1.12E-07 | 0.003615 |
| Manf | 0.124632 | 3.2E-08 | 0.001032 |
| Basp1 | 0.124396 | 2.74E-08 | 0.000883 |
| Klf2 | 0.124022 | 1.23E-06 | 0.039859 |
| Fat3 | 0.123855 | 0.010692 | 1 |
| Rpl23a | 0.12382 | 2.29E-11 | 7.4E-07 |
| mt-Nd3 | 0.123463 | 2.68E-10 | 8.67E-06 |
| Bsg | 0.123325 | 1.69E-10 | 5.45E-06 |
| Slc15a3 | 0.123317 | 0.000147 | 1 |
| Arhgap22 | 0.123019 | 1.17E-08 | 0.000379 |
| Efhd2 | 0.122803 | 9.06E-07 | 0.029256 |
| Ifitm10 | 0.122246 | 3.12E-05 | 1 |
| Tmed10 | 0.122052 | 1.29E-09 | 4.18E-05 |
| Rps4x | 0.121723 | 2.95E-12 | 9.52E-08 |
| Hnrnpa3 | 0.121677 | 1.63E-10 | 5.26E-06 |
| Arl4c | 0.121542 | 9E-08 | 0.002905 |
| Cd52 | 0.121495 | 4.94E-07 | 0.015953 |
| Hist1h2ae | 0.121321 | 1.66E-11 | 5.37E-07 |
| Cyba | 0.12117 | 1.35E-10 | 4.37E-06 |
| Pdia3 | 0.120943 | 1.48E-09 | 4.78E-05 |
| Gng5 | 0.120717 | 4.04E-10 | 1.31E-05 |
| Slamf9 | 0.120547 | 1.67E-05 | 0.540209 |
| Pdia6 | 0.120139 | 4.08E-10 | 1.32E-05 |
| Pdgfb | 0.119987 | 1.11E-06 | 0.035679 |
| Rps3 | 0.119738 | 6.17E-13 | 1.99E-08 |
| mt-Atp8 | 0.119336 | 6.01E-08 | 0.001941 |
| Rpl28 | 0.119198 | 2.69E-10 | 8.69E-06 |
| Rpl17 | 0.119181 | 2.79E-12 | 9E-08 |
| Cmtm6 | 0.119015 | 1.11E-08 | 0.000357 |
| Klhl6 | 0.118908 | 1.47E-05 | 0.474291 |
| Dnajb9 | 0.118665 | 7.05E-09 | 0.000228 |
| Tanc2 | 0.118032 | 8.32E-11 | 2.69E-06 |
| Atp6v0c | 0.117983 | 4.81E-07 | 0.015531 |
| Ccr5 | 0.11698 | 0.000852 | 1 |
| Rpl27a | 0.116561 | 1.22E-12 | 3.94E-08 |
| Naca | 0.116377 | 2.11E-09 | 6.81E-05 |
| Ndufa13 | 0.115269 | 2.35E-08 | 0.000758 |
| Hist1h4d | 0.115073 | 2.82E-08 | 0.000911 |

| | | | |
|---|---|---|---|
| Rpl37 | 0.114835 | 6.06E-12 | 1.96E-07 |
| Selenos | 0.114525 | 9.43E-08 | 0.003044 |
| mt-Nd1 | 0.114216 | 1.02E-10 | 3.3E-06 |
| Rps27a | 0.11419 | 6.58E-12 | 2.13E-07 |
| Dad1 | 0.113702 | 5.55E-09 | 0.000179 |
| 4930557K07Rik | 0.11353 | 0.439794 | 1 |
| Calm2 | 0.113306 | 7.52E-09 | 0.000243 |
| Kif21b | 0.11287 | 3.91E-07 | 0.012639 |
| Rpl35a | 0.112262 | 6.25E-12 | 2.02E-07 |
| Wsb1 | 0.112076 | 5.3E-08 | 0.001711 |
| Rps5 | 0.111944 | 1.62E-10 | 5.24E-06 |
| Ppp1r15a | 0.111827 | 3.05E-11 | 9.83E-07 |
| Mrpl52 | 0.111652 | 8.33E-07 | 0.026893 |
| Rpl7 | 0.111565 | 1.5E-11 | 4.85E-07 |
| Uqcrb | 0.111246 | 5.47E-08 | 0.001766 |
| Tmem50a | 0.111185 | 5.14E-08 | 0.001659 |
| Mettl23 | 0.110793 | 2.13E-10 | 6.87E-06 |
| Whrn | 0.110719 | 5.07E-08 | 0.001636 |
| Rpl32 | 0.110507 | 2.09E-12 | 6.75E-08 |
| Rpl26 | 0.110361 | 1.11E-11 | 3.59E-07 |
| Sft2d1 | 0.11019 | 6.85E-09 | 0.000221 |
| Fth1 | 0.110189 | 1.57E-07 | 0.005076 |
| Rpl21 | 0.110003 | 2.14E-13 | 6.9E-09 |
| Rplp2 | 0.10995 | 1.08E-12 | 3.49E-08 |
| Mylip | 0.109517 | 1.15E-06 | 0.037285 |
| Rps16 | 0.108244 | 1.41E-10 | 4.56E-06 |
| Rps20 | 0.10821 | 5.8E-11 | 1.87E-06 |
| Tmed3 | 0.108078 | 2.46E-09 | 7.94E-05 |
| Rpl22 | 0.108001 | 3.92E-08 | 0.001265 |
| Mrc2 | 0.107836 | 9.24E-05 | 1 |
| Cxcl16 | 0.107023 | 0.000862 | 1 |
| Dpm3 | 0.106561 | 3.02E-09 | 9.75E-05 |
| Rps9 | 0.106486 | 1.46E-11 | 4.73E-07 |
| Clta | 0.10635 | 4.07E-10 | 1.31E-05 |
| Golm1 | 0.106287 | 1.77E-07 | 0.005702 |
| A930007I19Rik | 0.106011 | 0.000179 | 1 |
| Cotl1 | 0.105955 | 0.000486 | 1 |
| Ywhaz | 0.105789 | 2.24E-06 | 0.072432 |
| Plxna4 | 0.105247 | 0.000123 | 1 |
| Calr | 0.105194 | 1.31E-07 | 0.004229 |
| Rps3a1 | 0.104946 | 3.55E-10 | 1.15E-05 |
| Sec11c | 0.1048 | 4.31E-07 | 0.013926 |
| Ostc | 0.104649 | 2.37E-07 | 0.007641 |
| Rpl9 | 0.104501 | 2.21E-09 | 7.15E-05 |
| Krtcap2 | 0.104423 | 9.73E-07 | 0.031407 |
| Lpl | 0.104295 | 0.020055 | 1 |
| Pmp22 | 0.104282 | 5.15E-06 | 0.166319 |
| Nr6a1os | 0.104178 | 0.001902 | 1 |

| | | | |
|---|---|---|---|
| Rps15 | 0.103435 | 3.61E-08 | 0.001165 |
| Rpl23 | 0.103422 | 2.03E-12 | 6.54E-08 |
| Nr1d2 | 0.103 | 5.14E-10 | 1.66E-05 |
| Rps6 | 0.102999 | 2.62E-09 | 8.45E-05 |
| Gm11867 | 0.102236 | 1.12E-18 | 3.63E-14 |
| Rps13 | 0.102069 | 1.85E-09 | 5.98E-05 |
| Rpl6 | 0.101866 | 1.1E-07 | 0.003567 |
| Rpl22l1 | 0.10148 | 1.77E-07 | 0.005721 |
| Rps14 | 0.101089 | 4.72E-09 | 0.000152 |
| Ecscr | 0.100892 | 5.95E-08 | 0.001921 |
| Rpl19 | 0.100774 | 4E-11 | 1.29E-06 |
| Rplp0 | 0.100742 | 1.88E-10 | 6.07E-06 |
| Fkbp2 | 0.099716 | 3.13E-06 | 0.1012 |
| mt-Co2 | 0.099689 | 2.46E-09 | 7.95E-05 |
| 2410006H16Rik | 0.099204 | 1.27E-05 | 0.410404 |
| Rpl24 | 0.09916 | 1.55E-09 | 5.02E-05 |
| Cox4i1 | 0.098971 | 9.65E-08 | 0.003115 |
| Ctso | 0.098469 | 1.19E-05 | 0.384564 |
| Tmsb4x | 0.098087 | 4.97E-12 | 1.61E-07 |
| Gm34921 | 0.097858 | 1.1E-11 | 3.56E-07 |
| Rps25 | 0.097836 | 1.51E-09 | 4.87E-05 |
| Slc3a2 | 0.097306 | 7.13E-07 | 0.023031 |
| Ncl | 0.09704 | 1.44E-05 | 0.463775 |
| Ctsf | 0.09686 | 6.66E-07 | 0.021495 |
| Itm2c | 0.096596 | 2.36E-07 | 0.007607 |
| Nptn | 0.096587 | 4.14E-06 | 0.133821 |
| Cd300c2 | 0.096542 | 5.42E-08 | 0.00175 |
| Eif1 | 0.09645 | 3.7E-07 | 0.011938 |
| Sox4 | 0.09636 | 0.000337 | 1 |
| Rps8 | 0.095922 | 6.75E-08 | 0.002181 |
| Ndufc2 | 0.095864 | 1.94E-05 | 0.625987 |
| Eef1a1 | 0.095839 | 4.15E-09 | 0.000134 |
| Spcs2 | 0.095535 | 2.28E-07 | 0.007364 |
| Eif5 | 0.094932 | 2.64E-06 | 0.085237 |
| Kpna4 | 0.094873 | 1.94E-05 | 0.626363 |
| Srgn | 0.094817 | 3.61E-06 | 0.116653 |
| Dnajc3 | 0.094776 | 7.75E-07 | 0.025017 |
| Id2 | 0.094758 | 0.005033 | 1 |
| Rpl38 | 0.094715 | 5.85E-09 | 0.000189 |
| Chsy1 | 0.094479 | 2.63E-05 | 0.849202 |
| Prkn | 0.094456 | 0.009256 | 1 |
| Tpp1 | 0.094379 | 2.13E-07 | 0.006877 |
| Rpl8 | 0.094322 | 3.53E-09 | 0.000114 |
| Qk | 0.094153 | 1.02E-06 | 0.032812 |
| Rpl18 | 0.094091 | 8.64E-08 | 0.002789 |
| Cd164 | 0.09396 | 2.19E-05 | 0.707319 |
| Bst2 | 0.093945 | 5.1E-05 | 1 |
| Rpl5 | 0.093933 | 2.81E-08 | 0.000908 |

| | | | |
|---|---|---|---|
| Ndufa6 | 0.093416 | 6.7E-05 | 1 |
| Atraid | 0.093145 | 7.84E-06 | 0.25303 |
| Ifrd1 | 0.092795 | 5.64E-07 | 0.018218 |
| Rps11 | 0.092538 | 7.86E-10 | 2.54E-05 |
| Zfp706 | 0.092511 | 3.74E-06 | 0.120766 |
| Hexa | 0.091318 | 8.82E-06 | 0.284857 |
| Surf4 | 0.091267 | 9.5E-05 | 1 |
| Ociad1 | 0.090972 | 2.79E-06 | 0.090093 |
| Tgfbr1 | 0.090802 | 4.77E-06 | 0.153904 |
| mt-Atp6 | 0.090718 | 1.14E-08 | 0.000369 |
| Gem | 0.090191 | 2.67E-10 | 8.6E-06 |
| Gm26887 | 0.090166 | 1.09E-18 | 3.53E-14 |
| Rpl12 | 0.08997 | 2.47E-09 | 7.98E-05 |
| Sdf2l1 | 0.089891 | 2.94E-06 | 0.094826 |
| Snrpg | 0.089519 | 2.2E-06 | 0.071033 |
| Klk8 | 0.088941 | 8.96E-07 | 0.028924 |
| Tmem37 | 0.088926 | 3.36E-06 | 0.108488 |
| Tex14 | 0.088888 | 0.000487 | 1 |
| C1qa | 0.088839 | 1.62E-10 | 5.23E-06 |
| Psenen | 0.088797 | 1.29E-07 | 0.004151 |
| Arf4 | 0.088245 | 0.001363 | 1 |
| Plk3 | 0.088048 | 0.000707 | 1 |
| Trmt10a | 0.087861 | 3.05E-07 | 0.00984 |
| Rpl27 | 0.087721 | 1.55E-07 | 0.005019 |
| Dusp1 | 0.086834 | 0.000179 | 1 |
| Rps23 | 0.086766 | 4.47E-10 | 1.44E-05 |
| Ccl9 | 0.086625 | 8.8E-06 | 0.284186 |
| Mef2c | 0.085639 | 3.34E-05 | 1 |
| Oaz1 | 0.085458 | 7.79E-07 | 0.025155 |
| Rpl7a | 0.085442 | 1.31E-05 | 0.422704 |
| Rps29 | 0.085344 | 1.57E-09 | 5.07E-05 |
| Il1b | 0.085162 | 0.052924 | 1 |
| Tmco1 | 0.085035 | 0.00016 | 1 |
| Rpl10 | 0.084944 | 1.59E-07 | 0.005146 |
| Hspa5 | 0.084768 | 4.82E-06 | 0.155486 |
| Rps10 | 0.08435 | 2.05E-08 | 0.000662 |
| Impact | 0.084153 | 0.000131 | 1 |
| Rack1 | 0.083705 | 4.32E-07 | 0.013937 |
| Tma7 | 0.083515 | 0.000201 | 1 |
| Laptm5 | 0.08347 | 1.38E-08 | 0.000446 |
| Rps28 | 0.083425 | 9.35E-08 | 0.003017 |
| Shisa5 | 0.083352 | 0.000692 | 1 |
| App | 0.083288 | 0.000381 | 1 |
| Mtch1 | 0.083006 | 5.95E-05 | 1 |
| 5033421B08Rik | 0.082976 | 0.000286 | 1 |
| Pard3b | 0.082863 | 0.085578 | 1 |
| Gm2629 | 0.08277 | 0.000651 | 1 |
| H1f0 | 0.082672 | 0.002318 | 1 |

| | | | |
|---|---|---|---|
| Hnrnpk | 0.082488 | 1.29E-05 | 0.415836 |
| Nars | 0.081994 | 0.000135 | 1 |
| Ube2j1 | 0.081687 | 2.77E-05 | 0.894899 |
| Calm1 | 0.080933 | 3.91E-05 | 1 |
| Pnrc1 | 0.080891 | 6.1E-06 | 0.197053 |
| Eif3k | 0.080878 | 0.000439 | 1 |
| Cox6b1 | 0.080568 | 3.83E-06 | 0.123718 |
| B2m | 0.080426 | 1.45E-07 | 0.004686 |
| Cd14 | 0.080137 | 0.000415 | 1 |
| Tmed9 | 0.080083 | 2.04E-05 | 0.657258 |
| Mtmr2 | 0.080039 | 0.000509 | 1 |
| Mrfap1 | 0.079789 | 5.46E-05 | 1 |
| Rpl15 | 0.079714 | 2.19E-05 | 0.706802 |
| Cuta | 0.079615 | 7.27E-06 | 0.234616 |
| Snhg8 | 0.079595 | 3.06E-05 | 0.988977 |
| Bcas2 | 0.079582 | 2.05E-05 | 0.660668 |
| Mt1 | 0.079402 | 0.000499 | 1 |
| Gal3st4 | 0.078945 | 0.001364 | 1 |
| Tmem176b | 0.078878 | 0.01657 | 1 |
| Edem1 | 0.078854 | 9.27E-06 | 0.299178 |
| Gabarap | 0.078161 | 6.67E-06 | 0.215237 |
| Rpl29 | 0.07815 | 6.05E-07 | 0.019528 |
| Rpl10a | 0.078108 | 5.91E-06 | 0.190658 |
| Ninj1 | 0.077927 | 2.48E-05 | 0.800379 |
| Selenof | 0.077893 | 1.24E-05 | 0.399504 |
| Alkbh1 | 0.077507 | 0.000322 | 1 |
| Ang | 0.077457 | 0.000475 | 1 |
| Rpl18a | 0.077371 | 1.06E-07 | 0.003425 |
| mt-Co3 | 0.077302 | 5.28E-07 | 0.017047 |
| Uqcrh | 0.077234 | 4.48E-05 | 1 |
| Itgam | 0.077214 | 0.000137 | 1 |
| 1700023H06Rik | 0.077197 | 6.88E-12 | 2.22E-07 |
| Adrb1 | 0.077122 | 0.00303 | 1 |
| Snhg12 | 0.077036 | 0.000265 | 1 |
| Sdf4 | 0.076449 | 1.11E-05 | 0.358661 |
| Saraf | 0.07631 | 4.04E-06 | 0.130468 |
| Mat2a | 0.0762 | 0.007383 | 1 |
| Reep5 | 0.075673 | 1.31E-05 | 0.422373 |
| Dmac2l | 0.075631 | 0.418353 | 1 |
| Rpl41 | 0.075304 | 6.42E-05 | 1 |
| Irf9 | 0.075209 | 0.197684 | 1 |
| Atp5e | 0.075097 | 0.00037 | 1 |
| Syngr2 | 0.074905 | 2.82E-05 | 0.910567 |
| Ptpro | 0.074831 | 0.007219 | 1 |
| Rasgrp3 | 0.074789 | 0.031721 | 1 |
| Tmem119 | 0.074775 | 2.83E-05 | 0.912635 |
| Rabac1 | 0.074396 | 1.43E-05 | 0.460854 |
| Rpl13a | 0.074395 | 3.8E-07 | 0.012264 |

| | | | |
|---|---|---|---|
| Nbea | 0.074329 | 0.002872 | 1 |
| 2010013B24Rik | 0.074027 | 5.03E-07 | 0.01625 |
| Sec61g | 0.073926 | 1.97E-06 | 0.063759 |
| Wdr83os | 0.073917 | 1.94E-05 | 0.626988 |
| Immp2l | 0.073824 | 0.016535 | 1 |
| Ctsa | 0.073789 | 2.84E-05 | 0.915588 |
| Gngt2 | 0.073759 | 1.45E-05 | 0.466864 |
| Dgkd | 0.073604 | 0.00489 | 1 |
| Glmp | 0.073504 | 2.32E-06 | 0.07495 |
| Dusp6 | 0.073478 | 0.004062 | 1 |
| Spint1 | 0.073181 | 0.000616 | 1 |
| Gde1 | 0.072885 | 0.00053 | 1 |
| Tm2d2 | 0.072857 | 0.000174 | 1 |
| Ftl1 | 0.072532 | 0.001612 | 1 |
| Npm1 | 0.07234 | 1.47E-05 | 0.474136 |
| Ccdc134 | 0.072319 | 1.29E-05 | 0.416234 |
| Sall3 | 0.072163 | 0.001168 | 1 |
| Atp5g2 | 0.072023 | 9.3E-05 | 1 |
| Lgals3 | 0.071914 | 0.713169 | 1 |
| Tmem208 | 0.071654 | 0.00015 | 1 |
| Eef1d | 0.071586 | 0.000402 | 1 |
| Srsf2 | 0.071495 | 0.00029 | 1 |
| Nsa2 | 0.070736 | 0.00269 | 1 |
| Atp6v0b | 0.070468 | 0.000158 | 1 |
| Hist1h2be | 0.07009 | 0.000236 | 1 |
| Mctp1 | 0.07005 | 0.021703 | 1 |
| Aif1 | 0.070039 | 5.77E-05 | 1 |
| mt-Co1 | 0.069996 | 2.34E-05 | 0.755484 |
| Adgrg1 | 0.069774 | 0.011028 | 1 |
| Abi3 | 0.069706 | 0.001387 | 1 |
| Khdrbs3 | 0.069646 | 2.36E-06 | 0.076349 |
| Pde3b | 0.069642 | 0.000584 | 1 |
| Plekhf2 | 0.069622 | 0.002456 | 1 |
| Trps1 | 0.069539 | 0.128241 | 1 |
| Gla | 0.069467 | 2.75E-05 | 0.888896 |
| Bcap31 | 0.069376 | 9.27E-05 | 1 |
| Tmed2 | 0.069171 | 5.57E-07 | 0.017997 |
| F11r | 0.069161 | 0.005669 | 1 |
| Scand1 | 0.068882 | 0.000331 | 1 |
| Ctsh | 0.068806 | 1.23E-05 | 0.395577 |
| Hspe1 | 0.068721 | 0.000544 | 1 |
| Rpl14 | 0.068453 | 3.18E-05 | 1 |
| Ifitm3 | 0.068177 | 0.019033 | 1 |
| Pcna | 0.068081 | 8.7E-09 | 0.000281 |
| Son | 0.068022 | 4.06E-06 | 0.131156 |
| Ndufb1-ps | 0.067867 | 0.002139 | 1 |
| Rpn2 | 0.067851 | 0.000106 | 1 |
| Atp5a1 | 0.067813 | 0.000194 | 1 |

| | | | |
|---|---|---|---|
| Arrdc3 | 0.067738 | 5.89E-06 | 0.19003 |
| Vsir | 0.067676 | 0.001734 | 1 |
| Med22 | 0.067613 | 4.27E-06 | 0.137909 |
| Hsp90ab1 | 0.067605 | 1.16E-05 | 0.375964 |
| Plpbp | 0.067479 | 0.000258 | 1 |
| Ssr3 | 0.067446 | 0.001678 | 1 |
| Tcn2 | 0.067398 | 0.000533 | 1 |
| Ndufa4 | 0.067271 | 1.38E-05 | 0.446487 |
| Herpud1 | 0.067268 | 0.001202 | 1 |
| Rer1 | 0.067228 | 0.000488 | 1 |
| Sept7 | 0.067119 | 0.000213 | 1 |
| Canx | 0.067047 | 0.000521 | 1 |
| Gatm | 0.066947 | 0.054951 | 1 |
| H2-Q7 | 0.066845 | 0.00073 | 1 |
| Fhad1 | 0.066722 | 2.78E-05 | 0.897119 |
| Rnf19b | 0.066611 | 0.941157 | 1 |
| Aup1 | 0.066534 | 0.001762 | 1 |
| 1810024B03Rik | 0.066525 | 0.010484 | 1 |
| Dnajb11 | 0.066371 | 0.000263 | 1 |
| Pdhb | 0.066237 | 0.003883 | 1 |
| Fam168b | 0.066181 | 3.25E-05 | 1 |
| Itgb1 | 0.066082 | 0.000261 | 1 |
| Skil | 0.066042 | 0.000115 | 1 |
| Btaf1 | 0.066041 | 0.001694 | 1 |
| Bach2 | 0.066015 | 0.049804 | 1 |
| Tmed7 | 0.065746 | 0.003694 | 1 |
| Pycard | 0.065733 | 0.000435 | 1 |
| Higd2a | 0.065441 | 0.000491 | 1 |
| Sar1b | 0.065425 | 0.001545 | 1 |
| Mydgf | 0.065377 | 0.00052 | 1 |
| Btf3 | 0.065311 | 0.000494 | 1 |
| Samsn1 | 0.065279 | 0.000218 | 1 |
| Jam2 | 0.065211 | 7.91E-05 | 1 |
| Dnase2a | 0.065137 | 0.000103 | 1 |
| Capza2 | 0.06511 | 0.000395 | 1 |
| Zfp202 | 0.064816 | 0.009189 | 1 |
| Lst1 | 0.064619 | 0.000175 | 1 |
| Slc9a9 | 0.064439 | 0.007718 | 1 |
| Chst8 | 0.06419 | 0.001036 | 1 |
| Tex261 | 0.064122 | 0.014495 | 1 |
| Txnl1 | 0.064006 | 0.004396 | 1 |
| AI506816 | 0.063679 | 0.002735 | 1 |
| Ier2 | 0.063578 | 0.190469 | 1 |
| Eef1b2 | 0.063479 | 6.39E-06 | 0.206156 |
| Daglb | 0.063472 | 0.003425 | 1 |
| Fmnl3 | 0.063434 | 0.002057 | 1 |
| Sirpa | 0.063404 | 0.001623 | 1 |
| Slc7a8 | 0.063355 | 0.00212 | 1 |

| | | | |
|---|---|---|---|
| Atp1b3 | 0.063289 | 0.000735 | 1 |
| Cd180 | 0.063201 | 0.102112 | 1 |
| Ldlr | 0.063169 | 0.03592 | 1 |
| Rpl4 | 0.063062 | 0.00019 | 1 |
| Hfe | 0.063054 | 0.000247 | 1 |
| Cbfa2t3 | 0.063033 | 7.42E-05 | 1 |
| Hdac8 | 0.062971 | 0.094565 | 1 |
| Tmem86a | 0.062622 | 0.000616 | 1 |
| Tmem256 | 0.062614 | 0.003332 | 1 |
| Magi1 | 0.062494 | 0.098641 | 1 |
| Taf7 | 0.06226 | 0.000736 | 1 |
| Ifngr2 | 0.062194 | 0.002808 | 1 |
| Thoc7 | 0.062093 | 0.002523 | 1 |
| Lrpap1 | 0.062063 | 0.000822 | 1 |
| Cnpy2 | 0.062055 | 0.000549 | 1 |
| Rbfox1 | 0.061862 | 0.0487 | 1 |
| Atp5d | 0.061827 | 0.000959 | 1 |
| Cables1 | 0.061598 | 0.004428 | 1 |
| Hnrnpdl | 0.061591 | 0.000265 | 1 |
| Rpl3 | 0.061341 | 1.72E-06 | 0.055456 |
| Man1a2 | 0.061339 | 0.000179 | 1 |
| Slc25a5 | 0.061287 | 0.000563 | 1 |
| Ywhab | 0.061065 | 0.007305 | 1 |
| Emc7 | 0.060999 | 0.002509 | 1 |
| Zfp827 | 0.060855 | 2.46E-06 | 0.079404 |
| Clic1 | 0.060815 | 0.000112 | 1 |
| Tspan31 | 0.060786 | 0.003445 | 1 |
| Gna15 | 0.060774 | 0.002268 | 1 |
| Laptm4a | 0.060728 | 0.000155 | 1 |
| Ero1l | 0.060627 | 0.001349 | 1 |
| Evi2a | 0.060387 | 0.007862 | 1 |
| Rps27l | 0.060278 | 0.000214 | 1 |
| Glrp1 | 0.060201 | 0.06279 | 1 |
| Vkorc1 | 0.060167 | 0.001497 | 1 |
| Fcgr2b | 0.06009 | 0.004098 | 1 |
| Tomm7 | 0.060061 | 0.000533 | 1 |
| Mrc1 | 0.059979 | 0.015425 | 1 |
| Tmed5 | 0.059738 | 0.000281 | 1 |
| Bmyc | 0.059458 | 0.017644 | 1 |
| Lyzl4 | 0.059375 | 0.012499 | 1 |
| Ssr2 | 0.059356 | 0.001019 | 1 |
| Alg5 | 0.05933 | 0.001121 | 1 |
| Slc44a1 | 0.059297 | 0.007167 | 1 |
| Ndufb3 | 0.059179 | 0.008745 | 1 |
| Eif2b4 | 0.059169 | 0.038502 | 1 |
| 9530068E07Rik | 0.059133 | 0.000356 | 1 |
| Ggh | 0.058799 | 0.000496 | 1 |
| Gfer | 0.05871 | 0.0034 | 1 |

| | | | |
|---|---|---|---|
| Cnbp | 0.05867 | 0.000464 | 1 |
| Ago1 | 0.058617 | 0.002388 | 1 |
| Cnih4 | 0.058581 | 0.000187 | 1 |
| Rgs10 | 0.058577 | 0.004114 | 1 |
| Rrp1 | 0.058536 | 0.003079 | 1 |
| Nkap | 0.058344 | 0.003255 | 1 |
| Gm26542 | 0.058257 | 0.038883 | 1 |
| Tmbim4 | 0.058246 | 0.00688 | 1 |
| Gnl3 | 0.058174 | 0.000481 | 1 |
| Edem2 | 0.05804 | 0.017463 | 1 |
| Ppp1r10 | 0.058036 | 0.001667 | 1 |
| Saysd1 | 0.057972 | 0.001534 | 1 |
| Cyc1 | 0.057955 | 0.003625 | 1 |
| Igf1r | 0.057873 | 0.002775 | 1 |
| Odc1 | 0.057781 | 7.17E-06 | 0.231451 |
| Sept9 | 0.057736 | 9.85E-06 | 0.318064 |
| Gfm2 | 0.057567 | 0.042564 | 1 |
| Cacnb2 | 0.057531 | 0.200009 | 1 |
| Irf2bpl | 0.057465 | 0.000415 | 1 |
| Gm37168 | 0.057241 | 7.3E-05 | 1 |
| Chd3 | 0.057163 | 0.007652 | 1 |
| Zfp869 | 0.057083 | 0.001391 | 1 |
| Tnrc6c | 0.05693 | 0.008739 | 1 |
| Ell2 | 0.056924 | 0.041212 | 1 |
| Ldhb | 0.056912 | 0.000642 | 1 |
| Emc6 | 0.056906 | 0.001038 | 1 |
| Cst7 | 0.056689 | 0.756047 | 1 |
| Lpxn | 0.056626 | 0.016679 | 1 |
| B3gat3 | 0.05661 | 0.001732 | 1 |
| Snx17 | 0.056533 | 0.006326 | 1 |
| Emc10 | 0.056436 | 0.014565 | 1 |
| Cox6c | 0.056428 | 0.000268 | 1 |
| Naa35 | 0.056368 | 0.015894 | 1 |
| Slc7a5 | 0.056326 | 0.000612 | 1 |
| Pdgfa | 0.056324 | 0.020499 | 1 |
| Ank2 | 0.056168 | 0.008792 | 1 |
| Mcfd2 | 0.056097 | 0.000231 | 1 |
| Serinc1 | 0.055972 | 0.00256 | 1 |
| Klf12 | 0.055746 | 0.065097 | 1 |
| Aurkaip1 | 0.055668 | 0.044722 | 1 |
| Ybx1 | 0.055604 | 0.009186 | 1 |
| Hscb | 0.055601 | 0.000319 | 1 |
| Rel | 0.05553 | 0.20745 | 1 |
| Ppp1r2 | 0.055429 | 0.000979 | 1 |
| Atp5f1 | 0.055324 | 0.003515 | 1 |
| Gcc2 | 0.055261 | 0.011967 | 1 |
| Ehd4 | 0.055208 | 0.0429 | 1 |
| Hcst | 0.055103 | 0.000151 | 1 |

| | | | |
|---|---|---|---|
| Zfp318 | 0.055067 | 0.022125 | 1 |
| Ergic3 | 0.054912 | 0.001265 | 1 |
| Ddx50 | 0.054857 | 0.011799 | 1 |
| Wdr36 | 0.054749 | 0.010632 | 1 |
| Bin2 | 0.054574 | 0.000245 | 1 |
| Cd53 | 0.054501 | 0.000236 | 1 |
| Tef | 0.054439 | 0.016896 | 1 |
| H3f3a | 0.054409 | 4.71E-05 | 1 |
| Rhoa | 0.054353 | 0.000516 | 1 |
| Clptm1l | 0.054343 | 0.000624 | 1 |
| Spty2d1 | 0.054234 | 0.015956 | 1 |
| Gm12703 | 0.054177 | 7.35E-06 | 0.237451 |
| Nrbf2 | 0.054023 | 0.004364 | 1 |
| Agmo | 0.05393 | 0.13562 | 1 |
| Vkorc1l1 | 0.053823 | 0.114548 | 1 |
| Ptpn18 | 0.053673 | 0.003674 | 1 |
| Arhgdia | 0.053604 | 0.007064 | 1 |
| Tmem100 | 0.053548 | 0.000268 | 1 |
| Dazap2 | 0.053482 | 0.022271 | 1 |
| Snrpf | 0.053405 | 0.005615 | 1 |
| Csnk1e | 0.053365 | 0.053262 | 1 |
| Serpine2 | 0.053295 | 0.002977 | 1 |
| Gspt1 | 0.053285 | 0.015675 | 1 |
| Il21r | 0.05321 | 0.057139 | 1 |
| Ndufb11 | 0.053026 | 0.001703 | 1 |
| Sall1 | 0.052946 | 0.013924 | 1 |
| Pttg1ip | 0.05288 | 0.003635 | 1 |
| Maml3 | 0.052843 | 0.009901 | 1 |
| Pet100 | 0.052819 | 0.000677 | 1 |
| Nagpa | 0.052803 | 0.007317 | 1 |
| Pin4 | 0.05278 | 0.031546 | 1 |
| Slc25a4 | 0.052764 | 0.042061 | 1 |
| Fgf13 | 0.052644 | 0.256882 | 1 |
| Tmem44 | 0.052641 | 0.018047 | 1 |
| Atp5g1 | 0.052641 | 0.002846 | 1 |
| Cass4 | 0.052558 | 0.205635 | 1 |
| Spcs1 | 0.052504 | 0.000246 | 1 |
| Cox5b | 0.052503 | 0.000189 | 1 |
| Abca1 | 0.052462 | 0.107153 | 1 |
| Hist1h2bg | 0.052434 | 3.77E-06 | 0.121748 |
| Rpl7l1 | 0.052392 | 0.004335 | 1 |
| Cmtm3 | 0.052243 | 0.013874 | 1 |
| Ly6e | 0.052095 | 0.003466 | 1 |
| H2-D1 | 0.052038 | 0.000282 | 1 |
| Atp5h | 0.052034 | 0.031483 | 1 |
| Neat1 | 0.051952 | 0.024525 | 1 |
| Tecr | 0.051932 | 0.006454 | 1 |
| Sys1 | 0.05193 | 0.000417 | 1 |

| | | | |
|---|---|---|---|
| Sem1 | 0.051866 | 0.001917 | 1 |
| Ptgs1 | 0.051762 | 0.005982 | 1 |
| Mfap3 | 0.051632 | 0.002591 | 1 |
| Ranbp1 | 0.051398 | 0.000396 | 1 |
| Pdia4 | 0.051292 | 0.010733 | 1 |
| Adprh | 0.051061 | 0.010772 | 1 |
| Gm44577 | 0.051013 | 0.000513 | 1 |
| Camk1 | 0.050907 | 0.003316 | 1 |
| Slc37a2 | 0.050851 | 0.102074 | 1 |
| Trf | 0.050811 | 0.000295 | 1 |
| Klf10 | 0.050437 | 0.000337 | 1 |
| Eif3e | 0.050364 | 0.006659 | 1 |
| Tmem14c | 0.050338 | 0.000283 | 1 |
| Arhgap32 | 0.050332 | 0.09243 | 1 |
| Slc39a7 | 0.050277 | 0.003661 | 1 |
| Cd86 | 0.050216 | 0.001405 | 1 |
| Morf4l1 | 0.05006 | 0.006622 | 1 |
| Ccdc107 | 0.050048 | 0.000236 | 1 |
| Tmem59 | 0.050046 | 0.006375 | 1 |
| Acadl | 0.050016 | 0.015178 | 1 |
| Anp32b | 0.050004 | 0.005784 | 1 |
| M6pr | 0.049882 | 0.004789 | 1 |
| Gm2a | 0.049771 | 0.030246 | 1 |
| Ddrgk1 | 0.049754 | 0.006507 | 1 |
| Arpc2 | 0.04974 | 0.116151 | 1 |
| Lcmt1 | 0.049708 | 0.028517 | 1 |
| Sec62 | 0.049675 | 0.001445 | 1 |
| Tnfaip8l2 | 0.049624 | 0.012053 | 1 |
| Selplg | 0.0496 | 0.017768 | 1 |
| Coq10b | 0.049586 | 0.021176 | 1 |
| Bax | 0.049408 | 0.002565 | 1 |
| Ywhah | 0.049278 | 0.000717 | 1 |
| Fads1 | 0.049245 | 0.023151 | 1 |
| Hnrnph2 | 0.049239 | 0.001205 | 1 |
| D630023F18Rik | 0.049202 | 1.93E-05 | 0.621986 |
| Ddost | 0.049199 | 0.000802 | 1 |
| Lmo2 | 0.04918 | 0.042204 | 1 |
| Plau | 0.049119 | 0.005207 | 1 |
| Hnrnpa2b1 | 0.049109 | 0.047814 | 1 |
| Tmem30a | 0.049087 | 0.009323 | 1 |
| Eif4a1 | 0.049045 | 0.002185 | 1 |
| Rpsa | 0.048948 | 0.004162 | 1 |
| Kat7 | 0.048869 | 0.039342 | 1 |
| Hist1h4c | 0.048836 | 1.17E-06 | 0.037897 |
| Arpc4 | 0.048773 | 0.112581 | 1 |
| Etf1 | 0.048766 | 0.017529 | 1 |
| Lsm6 | 0.048711 | 0.081703 | 1 |
| Amfr | 0.048667 | 0.006846 | 1 |

| | | | |
|---|---|---|---|
| Fkbp7 | 0.048636 | 6.98E-05 | 1 |
| Serpinf1 | 0.048585 | 0.025963 | 1 |
| Fam173a | 0.048568 | 0.0026 | 1 |
| Derl1 | 0.048539 | 0.000844 | 1 |
| Slc38a10 | 0.04842 | 0.028106 | 1 |
| Uqcrfs1 | 0.048259 | 0.018863 | 1 |
| Khk | 0.048257 | 0.008905 | 1 |
| Marcks | 0.048257 | 0.004684 | 1 |
| Baiap2 | 0.048226 | 0.049083 | 1 |
| Ndfip2 | 0.048204 | 0.010219 | 1 |
| Otud1 | 0.048051 | 0.017108 | 1 |
| Nenf | 0.048033 | 0.00295 | 1 |
| Ssr1 | 0.047934 | 0.012101 | 1 |
| Cd72 | 0.047815 | 0.869463 | 1 |
| 8030442B05Rik | 0.047797 | 0.019222 | 1 |
| Ncln | 0.047702 | 0.133989 | 1 |
| Pmaip1 | 0.047701 | 0.006125 | 1 |
| Naa20 | 0.047688 | 0.153854 | 1 |
| Pold4 | 0.047607 | 0.008445 | 1 |
| Cebpzos | 0.047515 | 0.003935 | 1 |
| Slc25a25 | 0.047435 | 0.115744 | 1 |
| Slc16a6 | 0.047333 | 0.134097 | 1 |
| Ddx18 | 0.047255 | 0.004551 | 1 |
| Terf1 | 0.047204 | 0.061207 | 1 |
| Cldnd1 | 0.047019 | 0.002708 | 1 |
| Sema4d | 0.04697 | 0.066233 | 1 |
| Larp7 | 0.046965 | 0.002168 | 1 |
| Cask | 0.046961 | 0.053602 | 1 |
| Eef2 | 0.046953 | 0.012961 | 1 |
| Cpd | 0.046915 | 0.185769 | 1 |
| Tgoln1 | 0.046838 | 0.005824 | 1 |
| Slc29a3 | 0.046825 | 0.047825 | 1 |
| Bcl2a1b | 0.046824 | 0.078492 | 1 |
| Arpc3 | 0.046813 | 0.001841 | 1 |
| Gm10138 | 0.046775 | 0.080052 | 1 |
| Mpg | 0.046706 | 0.001434 | 1 |
| Smc5 | 0.046656 | 0.007811 | 1 |
| Brix1 | 0.046635 | 1.55E-05 | 0.500022 |
| Sap18 | 0.046616 | 0.000509 | 1 |
| Uqcrq | 0.046597 | 0.010278 | 1 |
| Tor2a | 0.046445 | 0.00067 | 1 |
| Rpl36al | 0.046379 | 0.000438 | 1 |
| Necap2 | 0.046204 | 0.03652 | 1 |
| Cstb | 0.046137 | 0.001099 | 1 |
| Errfi1 | 0.046091 | 0.011224 | 1 |
| Gimap6 | 0.046065 | 3.49E-05 | 1 |
| Rps18 | 0.046045 | 0.000513 | 1 |
| Chd6 | 0.046041 | 0.197728 | 1 |

| | | | |
|---|---|---|---|
| Hsd17b10 | 0.046032 | 0.009778 | 1 |
| Col27a1 | 0.045983 | 0.089856 | 1 |
| Dapk1 | 0.045932 | 0.044897 | 1 |
| Ppcdc | 0.045926 | 0.13785 | 1 |
| Ptpmt1 | 0.0459 | 0.017393 | 1 |
| Micu3 | 0.045857 | 0.288539 | 1 |
| Iws1 | 0.04582 | 0.030307 | 1 |
| 1700110C19Rik | 0.045799 | 0.010942 | 1 |
| Mfng | 0.045541 | 0.00521 | 1 |
| Camk2n1 | 0.045457 | 0.081223 | 1 |
| Slc39a1 | 0.045447 | 0.017571 | 1 |
| Rfc3 | 0.045418 | 0.000259 | 1 |
| Spast | 0.045396 | 0.174404 | 1 |
| D8Ertd738e | 0.045359 | 0.062587 | 1 |
| Rbm42 | 0.045297 | 0.000485 | 1 |
| Ndufab1 | 0.045253 | 0.036682 | 1 |
| Eif3c | 0.04517 | 0.025629 | 1 |
| Mfsd14a | 0.045036 | 0.007131 | 1 |
| Pde8b | 0.044996 | 0.013752 | 1 |
| Rps17 | 0.044996 | 0.004887 | 1 |
| Hspa4 | 0.04499 | 0.03073 | 1 |
| Tram1 | 0.044892 | 0.036135 | 1 |
| Srp14 | 0.044834 | 0.000663 | 1 |
| Sec61b | 0.044818 | 0.014588 | 1 |
| Cox7a2l | 0.044802 | 0.000595 | 1 |
| Ptges3 | 0.044797 | 0.00376 | 1 |
| Ablim1 | 0.044741 | 0.00866 | 1 |
| Poc1b | 0.044648 | 0.043295 | 1 |
| Commd3 | 0.044551 | 0.001048 | 1 |
| Ifi30 | 0.044512 | 0.003624 | 1 |
| Slamf8 | 0.044455 | 0.033522 | 1 |
| Msmo1 | 0.044378 | 0.004267 | 1 |
| Cops9 | 0.044322 | 0.011226 | 1 |
| Havcr2 | 0.044297 | 0.02218 | 1 |
| Tvp23b | 0.044291 | 0.05728 | 1 |
| Golga5 | 0.044246 | 0.095816 | 1 |
| Cenpb | 0.044165 | 0.000197 | 1 |
| Ebna1bp2 | 0.044123 | 0.013594 | 1 |
| Fdft1 | 0.044073 | 0.002227 | 1 |
| Golt1b | 0.043985 | 0.011516 | 1 |
| Dusp3 | 0.043974 | 0.002743 | 1 |
| Pgk1 | 0.043967 | 0.039185 | 1 |
| Dennd2c | 0.043934 | 0.035512 | 1 |
| Uqcrc2 | 0.043846 | 0.051837 | 1 |
| Micos13 | 0.043825 | 0.02448 | 1 |
| Vmp1 | 0.043807 | 0.008766 | 1 |
| Spg20 | 0.043792 | 0.046443 | 1 |
| Ide | 0.043742 | 0.003027 | 1 |

| | | | |
|---|---|---|---|
| Aga | 0.043736 | 0.006805 | 1 |
| Brip1os | 0.043727 | 0.152494 | 1 |
| Rbm3 | 0.043686 | 0.000287 | 1 |
| Napa | 0.043686 | 0.006016 | 1 |
| Lmnb1 | 0.043686 | 0.004569 | 1 |
| Cox7c | 0.043676 | 0.000196 | 1 |
| Btbd19 | 0.043524 | 0.006684 | 1 |
| Actl6a | 0.043499 | 0.001969 | 1 |
| Timm13 | 0.043422 | 0.015324 | 1 |
| Ndufs8 | 0.043401 | 0.019175 | 1 |
| Zswim8 | 0.043396 | 0.101642 | 1 |
| Dennd4a | 0.043394 | 0.090655 | 1 |
| Sycp2 | 0.043393 | 0.32489 | 1 |
| Insig1 | 0.043335 | 8.96E-05 | 1 |
| Eif4a2 | 0.043299 | 0.015762 | 1 |
| Dek | 0.04314 | 0.006504 | 1 |
| Ndufa7 | 0.043121 | 0.003086 | 1 |
| Hspa9 | 0.043055 | 0.002264 | 1 |
| Adh5 | 0.043008 | 0.005195 | 1 |
| Uqcr11 | 0.042965 | 0.008627 | 1 |
| Eif3h | 0.042947 | 0.023847 | 1 |
| Gm14023 | 0.042906 | 7.36E-05 | 1 |
| Tbl3 | 0.0429 | 0.043143 | 1 |
| Erp29 | 0.042766 | 0.013088 | 1 |
| Csf1 | 0.042694 | 0.474176 | 1 |
| Arhgef40 | 0.042538 | 0.028881 | 1 |
| Golgb1 | 0.042509 | 0.027718 | 1 |
| 1110032A03Rik | 0.042497 | 0.009867 | 1 |
| Nf1 | 0.04248 | 0.023589 | 1 |
| Polr2k | 0.042233 | 0.00134 | 1 |
| Leprotl1 | 0.042153 | 0.003838 | 1 |
| Donson | 0.042122 | 0.000158 | 1 |
| Atg2a | 0.042071 | 0.011222 | 1 |
| Arl8b | 0.042003 | 0.033537 | 1 |
| Hadha | 0.041965 | 0.015333 | 1 |
| C5ar1 | 0.041855 | 0.362957 | 1 |
| Xrcc5 | 0.041795 | 0.661723 | 1 |
| Derl2 | 0.041703 | 0.047506 | 1 |
| Rnf130 | 0.041685 | 0.013304 | 1 |
| Thra | 0.041681 | 0.000271 | 1 |
| Eif2s1 | 0.041637 | 0.152611 | 1 |
| Washc1 | 0.041573 | 0.012148 | 1 |
| Med31 | 0.041539 | 0.002399 | 1 |
| Adss | 0.041509 | 0.00585 | 1 |
| Cox8a | 0.041478 | 0.000656 | 1 |
| Sumo2 | 0.041472 | 0.012145 | 1 |
| Utp18 | 0.041461 | 0.050332 | 1 |
| Med12l | 0.041456 | 0.030712 | 1 |

| | | | |
|---|---|---|---|
| Eif3g | 0.041382 | 0.001412 | 1 |
| Prkar1a | 0.041301 | 0.009782 | 1 |
| Cnih1 | 0.041057 | 0.014637 | 1 |
| Hif1a | 0.040895 | 0.04618 | 1 |
| Rnf157 | 0.040851 | 0.031086 | 1 |
| Hpgd | 0.040843 | 0.323382 | 1 |
| Gatd3a | 0.040826 | 0.000506 | 1 |
| Ets1 | 0.040756 | 0.003089 | 1 |
| Pex1 | 0.040729 | 0.537548 | 1 |
| Ccnd3 | 0.040728 | 0.015624 | 1 |
| Lamtor4 | 0.040637 | 0.000906 | 1 |
| Vps4b | 0.040413 | 0.014081 | 1 |
| Grap | 0.040403 | 0.162928 | 1 |
| Slc20a1 | 0.040327 | 0.01037 | 1 |
| Ildr1 | 0.040294 | 2.18E-05 | 0.705115 |
| Sdcbp | 0.040245 | 0.02356 | 1 |
| Arv1 | 0.040202 | 0.01882 | 1 |
| 1700017B05Rik | 0.040104 | 0.016476 | 1 |
| Acvrl1 | 0.040103 | 0.018056 | 1 |
| Atad5 | 0.039968 | 0.479903 | 1 |
| Rpn1 | 0.039928 | 0.034012 | 1 |
| Ndufs7 | 0.039891 | 0.090999 | 1 |
| Degs1 | 0.039852 | 0.007269 | 1 |
| Tspan9 | 0.03985 | 0.024398 | 1 |
| Cct2 | 0.039827 | 0.082393 | 1 |
| Bin1 | 0.039813 | 0.029024 | 1 |
| Limd2 | 0.039754 | 0.062136 | 1 |
| Fis1 | 0.039717 | 0.003471 | 1 |
| Pld3 | 0.039696 | 0.001331 | 1 |
| Cgrrf1 | 0.039677 | 0.01736 | 1 |
| Edf1 | 0.03967 | 0.019502 | 1 |
| Ilf2 | 0.039657 | 0.402645 | 1 |
| Cyp20a1 | 0.039609 | 0.001752 | 1 |
| Snhg4 | 0.039565 | 0.04809 | 1 |
| Mfge8 | 0.039519 | 0.00421 | 1 |
| Nedd8 | 0.039517 | 0.010235 | 1 |
| Liph | 0.039422 | 0.011003 | 1 |
| Gnai2 | 0.039418 | 0.022628 | 1 |
| Gm16192 | 0.039386 | 0.000726 | 1 |
| Nup205 | 0.03934 | 0.147476 | 1 |
| Ski | 0.039311 | 0.06084 | 1 |
| Sowahc | 0.039242 | 0.234781 | 1 |
| Gm5617 | 0.039228 | 0.054152 | 1 |
| Myl6 | 0.03922 | 0.000927 | 1 |
| Prox2 | 0.039206 | 0.093204 | 1 |
| Rabgap1l | 0.03913 | 0.048825 | 1 |
| Esf1 | 0.039109 | 0.004597 | 1 |
| Bod1l | 0.039107 | 0.013375 | 1 |

| | | | |
|---|---|---|---|
| Nop14 | 0.039105 | 0.054 | 1 |
| Phf5a | 0.039014 | 0.042685 | 1 |
| H2afz | 0.038989 | 8.43E-05 | 1 |
| Scaf11 | 0.038874 | 0.032295 | 1 |
| Gpr84 | 0.038844 | 0.404451 | 1 |
| Naglu | 0.038837 | 0.032334 | 1 |
| Vdac1 | 0.038786 | 0.004722 | 1 |
| Pdcd4 | 0.038783 | 0.064616 | 1 |
| Tmtc3 | 0.038762 | 0.227941 | 1 |
| Chmp5 | 0.038744 | 0.083237 | 1 |
| Map1lc3b | 0.038703 | 0.00132 | 1 |
| Romo1 | 0.038664 | 0.05966 | 1 |
| Fuca1 | 0.038648 | 0.018004 | 1 |
| Cd244a | 0.0386 | 0.092835 | 1 |
| Hcls1 | 0.038598 | 0.0696 | 1 |
| Alg6 | 0.038548 | 0.021256 | 1 |
| Atp5o | 0.038524 | 0.0244 | 1 |
| Usp22 | 0.038518 | 0.002326 | 1 |
| Rin2 | 0.03849 | 0.029742 | 1 |
| Lipa | 0.038459 | 0.127957 | 1 |
| Slc50a1 | 0.038446 | 0.062346 | 1 |
| Stab1 | 0.038437 | 0.254924 | 1 |
| Ldlrad4 | 0.038429 | 0.007114 | 1 |
| Slamf6 | 0.03839 | 0.018055 | 1 |
| Polr2a | 0.038377 | 0.029344 | 1 |
| Mrpl54 | 0.038348 | 0.002887 | 1 |
| Epc1 | 0.03834 | 0.199006 | 1 |
| Zbtb24 | 0.038318 | 0.003161 | 1 |
| Zfp697 | 0.038308 | 0.819235 | 1 |
| mt-Nd6 | 0.038298 | 0.000962 | 1 |
| Srek1 | 0.038068 | 0.028918 | 1 |
| Ndufb2 | 0.038059 | 0.02826 | 1 |
| Pbdc1 | 0.037935 | 0.034811 | 1 |
| Scamp2 | 0.037902 | 0.065478 | 1 |
| Tmem160 | 0.037888 | 0.00323 | 1 |
| Galk1 | 0.037868 | 0.001571 | 1 |
| Sulf2 | 0.037834 | 0.12619 | 1 |
| Nav2 | 0.037832 | 0.087175 | 1 |
| Mnt | 0.037776 | 7.03E-05 | 1 |
| Camk2g | 0.037742 | 0.076822 | 1 |
| Plbd2 | 0.037718 | 0.007338 | 1 |
| Atf4 | 0.037655 | 0.004559 | 1 |
| Epn2 | 0.03764 | 0.708871 | 1 |
| Ccdc115 | 0.03763 | 0.083167 | 1 |
| Acox3 | 0.037573 | 0.075844 | 1 |
| Ifitm2 | 0.037532 | 5.91E-05 | 1 |
| Smpdl3a | 0.037492 | 1.49E-05 | 0.480088 |
| Gnptg | 0.03748 | 0.031162 | 1 |

| | | | |
|---|---|---|---|
| C330007P06Rik | 0.037439 | 0.203693 | 1 |
| Got1 | 0.03741 | 0.005143 | 1 |
| AU022793 | 0.037361 | 0.035732 | 1 |
| Tmem33 | 0.037334 | 0.062496 | 1 |
| Sgta | 0.03732 | 0.009965 | 1 |
| Cks2 | 0.037315 | 0.281358 | 1 |
| Srek1ip1 | 0.03726 | 0.027378 | 1 |
| Jup | 0.037257 | 0.093845 | 1 |
| Nr2c2 | 0.037252 | 0.085894 | 1 |
| Cisd1 | 0.03722 | 0.021648 | 1 |
| Arhgap30 | 0.037149 | 0.120903 | 1 |
| BC005624 | 0.037079 | 0.037535 | 1 |
| Eif3a | 0.037075 | 0.014874 | 1 |
| Rpl31 | 0.037073 | 0.020702 | 1 |
| Gm38843 | 0.037055 | 0.035488 | 1 |
| Exoc5 | 0.037042 | 0.048181 | 1 |
| Ywhaq | 0.037037 | 0.153247 | 1 |
| Slc35e1 | 0.037022 | 0.011879 | 1 |
| Commd10 | 0.03699 | 0.15597 | 1 |
| Vps28 | 0.03696 | 0.040359 | 1 |
| Slc25a3 | 0.036956 | 0.001439 | 1 |
| Dctn2 | 0.036928 | 0.105269 | 1 |
| Arpc5 | 0.036918 | 0.008118 | 1 |
| Eif2ak4 | 0.036885 | 0.196998 | 1 |
| Snw1 | 0.036832 | 0.038875 | 1 |
| Tbc1d15 | 0.036809 | 0.066104 | 1 |
| Erh | 0.036794 | 0.000626 | 1 |
| Gpx4 | 0.03679 | 0.005351 | 1 |
| Rab5a | 0.03673 | 0.01385 | 1 |
| Taz | 0.036729 | 0.01411 | 1 |
| Ywhae | 0.036688 | 0.055945 | 1 |
| Armh3 | 0.036649 | 0.181442 | 1 |
| Sf1 | 0.036635 | 0.06177 | 1 |
| Cops6 | 0.036578 | 0.018758 | 1 |
| Atp6v1a | 0.036559 | 0.280798 | 1 |
| Il6st | 0.036529 | 0.059773 | 1 |
| Grpel1 | 0.036489 | 0.021895 | 1 |
| Slc7a6os | 0.036429 | 0.089723 | 1 |
| D330023K18Rik | 0.036412 | 0.115485 | 1 |
| Bloc1s1 | 0.036399 | 3.18E-06 | 0.10265 |
| Psap | 0.036362 | 0.118119 | 1 |
| Med7 | 0.036328 | 0.018295 | 1 |
| Cacybp | 0.036293 | 0.023672 | 1 |
| Nfkbid | 0.036291 | 0.005248 | 1 |
| Clec4a2 | 0.036275 | 0.038887 | 1 |
| Cox6a1 | 0.036196 | 0.002433 | 1 |
| Pex2 | 0.036144 | 0.031482 | 1 |
| Gtf2h2 | 0.036126 | 0.039769 | 1 |

| | | | |
|---|---|---|---|
| Eif3i | 0.036115 | 0.009115 | 1 |
| Alox5ap | 0.036102 | 0.017315 | 1 |
| Parp1 | 0.036095 | 0.6778 | 1 |
| Ccnq | 0.036081 | 0.002551 | 1 |
| Ppia | 0.036034 | 0.002357 | 1 |
| Emc3 | 0.036026 | 0.005202 | 1 |
| Marc2 | 0.036017 | 0.022986 | 1 |
| Mapre1 | 0.035974 | 0.011997 | 1 |
| Comtd1 | 0.03591 | 0.009195 | 1 |
| Slc25a11 | 0.035881 | 0.01539 | 1 |
| Park7 | 0.035855 | 0.051554 | 1 |
| Med23 | 0.035852 | 0.109786 | 1 |
| Igfbp4 | 0.035734 | 0.0022 | 1 |
| Memo1 | 0.035705 | 0.026078 | 1 |
| Cyfip1 | 0.035657 | 0.117663 | 1 |
| Akr1b3 | 0.035645 | 0.019236 | 1 |
| Fbxl12 | 0.035636 | 0.059148 | 1 |
| Ptp4a3 | 0.035619 | 0.058334 | 1 |
| Elp1 | 0.035606 | 0.076305 | 1 |
| Rtraf | 0.035592 | 0.001166 | 1 |
| Aph1a | 0.035548 | 0.162043 | 1 |
| Mpdu1 | 0.035484 | 0.045591 | 1 |
| Akirin2 | 0.035435 | 0.003419 | 1 |
| Elob | 0.035396 | 0.003449 | 1 |
| Gm9725 | 0.03535 | 0.00177 | 1 |
| Grcc10 | 0.035284 | 0.04349 | 1 |
| Map2k2 | 0.035228 | 0.098254 | 1 |
| Mrps5 | 0.035222 | 0.006167 | 1 |
| Hnrnpu | 0.035215 | 0.047336 | 1 |
| Pigb | 0.035183 | 0.464967 | 1 |
| Snrpa | 0.035127 | 0.024429 | 1 |
| Rrp1b | 0.035106 | 0.171809 | 1 |
| Snrpe | 0.035092 | 0.017715 | 1 |
| Gm47664 | 0.035081 | 0.008377 | 1 |
| Prpf39 | 0.035003 | 0.120642 | 1 |
| Trappc4 | 0.034904 | 0.082245 | 1 |
| Gsg1 | 0.034898 | 0.036359 | 1 |
| Mpc1 | 0.034852 | 0.149356 | 1 |
| Eif2ak3 | 0.034823 | 0.378437 | 1 |
| Ppwd1 | 0.034809 | 0.440171 | 1 |
| Pabpn1 | 0.034729 | 0.05652 | 1 |
| Dlst | 0.034709 | 0.157239 | 1 |
| Dcxr | 0.034694 | 0.096325 | 1 |
| Myl12a | 0.034688 | 0.014885 | 1 |
| Zmat2 | 0.03465 | 0.009261 | 1 |
| Coq2 | 0.0346 | 0.187191 | 1 |
| Atp13a3 | 0.034542 | 0.023434 | 1 |
| Gaa | 0.034527 | 0.010023 | 1 |

| | | | |
|---|---|---|---|
| Lyrm2 | 0.034515 | 0.007929 | 1 |
| Sgce | 0.034498 | 0.49258 | 1 |
| Hmgxb4 | 0.034409 | 0.248166 | 1 |
| Tmem9b | 0.034345 | 0.009227 | 1 |
| Psmc4 | 0.034252 | 0.028457 | 1 |
| Inpp4b | 0.034223 | 0.096768 | 1 |
| Xrn2 | 0.034204 | 0.088269 | 1 |
| BC031181 | 0.03417 | 0.002406 | 1 |
| Ccnl1 | 0.034169 | 0.081193 | 1 |
| Kdelr2 | 0.034145 | 0.019469 | 1 |
| Uevld | 0.034121 | 0.056052 | 1 |
| Zswim4 | 0.03412 | 0.026 | 1 |
| Cdk5r1 | 0.034118 | 0.008887 | 1 |
| Ykt6 | 0.03409 | 0.033827 | 1 |
| Nipsnap3b | 0.034086 | 0.233791 | 1 |
| Gusb | 0.034047 | 0.090824 | 1 |
| Chchd1 | 0.03404 | 0.047361 | 1 |
| Bbip1 | 0.034036 | 0.032675 | 1 |
| Mrpl9 | 0.033982 | 0.548262 | 1 |
| Mrps28 | 0.03388 | 0.041212 | 1 |
| Osm | 0.033879 | 0.087282 | 1 |
| Psmb6 | 0.033871 | 0.170632 | 1 |
| 2610528A11Rik | 0.03386 | 0.000616 | 1 |
| Art3 | 0.033856 | 0.015842 | 1 |
| Esr1 | 0.033844 | 0.883713 | 1 |
| Neurl3 | 0.033777 | 0.162964 | 1 |
| Pgp | 0.03375 | 0.107321 | 1 |
| Gnb1l | 0.033673 | 0.186484 | 1 |
| Asap3 | 0.03362 | 0.031985 | 1 |
| Mrpl41 | 0.033505 | 0.007223 | 1 |
| Spag7 | 0.033472 | 0.019492 | 1 |
| Hnrnpab | 0.033419 | 0.024739 | 1 |
| Clec5a | 0.033406 | 0.09516 | 1 |
| Cirbp | 0.0334 | 0.296979 | 1 |
| Sms | 0.033398 | 0.077553 | 1 |
| Usp2 | 0.03336 | 0.039432 | 1 |
| Elovl1 | 0.033303 | 0.095507 | 1 |
| Dnm1l | 0.033291 | 0.040047 | 1 |
| Nav1 | 0.033269 | 0.203123 | 1 |
| Zfp36l2 | 0.033262 | 0.353769 | 1 |
| Yeats2 | 0.03326 | 0.185907 | 1 |
| Ppig | 0.03326 | 0.106489 | 1 |
| Nop10 | 0.033257 | 0.09076 | 1 |
| Ctnnd1 | 0.033228 | 0.446988 | 1 |
| Fancc | 0.033214 | 0.024177 | 1 |
| Pum3 | 0.033214 | 0.248596 | 1 |
| Lrrc3 | 0.033197 | 0.003517 | 1 |
| Kazn | 0.033188 | 0.008627 | 1 |

| | | | |
|---|---|---|---|
| Fhit | 0.03317 | 0.635386 | 1 |
| Mppe1 | 0.033169 | 0.015864 | 1 |
| Fam217b | 0.03305 | 0.146298 | 1 |
| Alyref2 | 0.033029 | 0.004394 | 1 |
| Chn2 | 0.032998 | 0.037444 | 1 |
| 2410002F23Rik | 0.032926 | 0.005295 | 1 |
| Cndp2 | 0.032914 | 0.042551 | 1 |
| Chka | 0.032904 | 0.035476 | 1 |
| Fbxo8 | 0.032835 | 0.145125 | 1 |
| Pfkfb3 | 0.032806 | 0.636399 | 1 |
| Ggta1 | 0.032769 | 0.075794 | 1 |
| Hist2h2ac | 0.032713 | 0.000245 | 1 |
| Itpripl2 | 0.032689 | 0.04535 | 1 |
| Rps6ka5 | 0.03268 | 0.163006 | 1 |
| Bnip3 | 0.032659 | 0.023114 | 1 |
| Sgpl1 | 0.032647 | 0.067422 | 1 |
| Rell1 | 0.032642 | 0.001967 | 1 |
| Tcea1 | 0.032631 | 0.012276 | 1 |
| Ubl5 | 0.032621 | 0.010605 | 1 |
| Polr2f | 0.032592 | 0.118999 | 1 |
| Usp1 | 0.032591 | 0.58553 | 1 |
| Stau2 | 0.032581 | 0.211876 | 1 |
| Pnkd | 0.032576 | 0.012643 | 1 |
| Top3a | 0.032574 | 0.121901 | 1 |
| Prdx4 | 0.032574 | 0.05403 | 1 |
| Gadd45g | 0.032464 | 0.000751 | 1 |
| Cd37 | 0.032422 | 0.191235 | 1 |
| Fam32a | 0.032422 | 0.12225 | 1 |
| Jmjd1c | 0.032377 | 0.147783 | 1 |
| Cops2 | 0.032362 | 0.300336 | 1 |
| Vps37c | 0.032357 | 0.167464 | 1 |
| Copb1 | 0.03232 | 0.102853 | 1 |
| Cox7b | 0.03232 | 0.011901 | 1 |
| Nop53 | 0.032317 | 0.103462 | 1 |
| Rubcnl | 0.032271 | 0.069805 | 1 |
| Hmgcr | 0.032226 | 0.075892 | 1 |
| Aggf1 | 0.032209 | 0.143065 | 1 |
| Herpud2 | 0.032202 | 0.444088 | 1 |
| Trib1 | 0.03218 | 0.101124 | 1 |
| Phf23 | 0.032115 | 0.416081 | 1 |
| Psma4 | 0.032103 | 0.020486 | 1 |
| Cux1 | 0.03208 | 0.065269 | 1 |
| Mdc1 | 0.032057 | 0.048619 | 1 |
| Gdi2 | 0.032017 | 0.057725 | 1 |
| Mrpl16 | 0.031997 | 0.004638 | 1 |
| Tmem176a | 0.031972 | 0.045876 | 1 |
| Rsl24d1 | 0.031956 | 0.056813 | 1 |
| Eftud2 | 0.031927 | 0.046537 | 1 |

| | | | |
|---|---|---|---|
| Ginm1 | 0.031841 | 0.051181 | 1 |
| Anxa5 | 0.031808 | 0.137316 | 1 |
| Paip1 | 0.031768 | 0.464483 | 1 |
| Ebp | 0.031754 | 0.102712 | 1 |
| Actr3 | 0.031745 | 0.167976 | 1 |
| Dpm1 | 0.031742 | 0.103986 | 1 |
| Mid1ip1 | 0.03167 | 0.028035 | 1 |
| Tmem123 | 0.031663 | 0.10337 | 1 |
| Osbpl3 | 0.031549 | 0.497085 | 1 |
| Slc35b1 | 0.031537 | 0.109866 | 1 |
| Lamp2 | 0.031507 | 0.031928 | 1 |
| Tug1 | 0.031484 | 0.084612 | 1 |
| Plaur | 0.031366 | 0.097041 | 1 |
| Exoc1 | 0.031363 | 0.011591 | 1 |
| Arl2bp | 0.031225 | 0.068766 | 1 |
| Lair1 | 0.031195 | 0.002933 | 1 |
| Tmem50b | 0.031178 | 0.346775 | 1 |
| Tmed4 | 0.031166 | 0.002532 | 1 |
| Klhl7 | 0.031141 | 0.008476 | 1 |
| Lipo3 | 0.031087 | 0.092836 | 1 |
| Nktr | 0.031065 | 0.065719 | 1 |
| Pja2 | 0.031027 | 0.708281 | 1 |
| Txnrd2 | 0.03101 | 0.213911 | 1 |
| Lrch4 | 0.030985 | 0.017587 | 1 |
| Srfbp1 | 0.030979 | 0.180474 | 1 |
| Slc23a2 | 0.030925 | 0.085793 | 1 |
| Midn | 0.030876 | 0.041516 | 1 |
| Spsb1 | 0.030862 | 0.05298 | 1 |
| Gm20186 | 0.0308 | 0.000273 | 1 |
| Cfl1 | 0.030784 | 0.009906 | 1 |
| Fzd4 | 0.030773 | 0.0162 | 1 |
| Med10 | 0.030761 | 0.002338 | 1 |
| Atg12 | 0.030748 | 0.084204 | 1 |
| Mogs | 0.03073 | 0.193192 | 1 |
| Ddx21 | 0.030724 | 0.119099 | 1 |
| 1700028E10Rik | 0.030681 | 0.785574 | 1 |
| Lyl1 | 0.030678 | 0.092518 | 1 |
| Il4ra | 0.030669 | 0.054355 | 1 |
| Rbmxl1 | 0.030622 | 0.208413 | 1 |
| Hnrnpl | 0.030611 | 0.050265 | 1 |
| Atp5k | 0.030581 | 0.03816 | 1 |
| Lman1 | 0.030487 | 0.003186 | 1 |
| Pfdn2 | 0.030475 | 0.109224 | 1 |
| B3gnt2 | 0.030474 | 0.337835 | 1 |
| Ddx54 | 0.030445 | 0.002543 | 1 |
| Kcnip3 | 0.030421 | 0.264926 | 1 |
| Ak1 | 0.030414 | 0.101346 | 1 |
| Spns2 | 0.030396 | 0.117017 | 1 |

| | | | |
|---|---|---|---|
| H2-K1 | 0.030364 | 0.001434 | 1 |
| Pycr2 | 0.030353 | 0.035122 | 1 |
| Atp6v1g1 | 0.030348 | 0.071298 | 1 |
| Gm12905 | 0.030329 | 0.031239 | 1 |
| Atp6ap1 | 0.030279 | 0.024646 | 1 |
| Srr | 0.030268 | 0.464074 | 1 |
| Mpv17l2 | 0.030262 | 0.172959 | 1 |
| P3h2 | 0.030261 | 0.371254 | 1 |
| Pex19 | 0.030255 | 0.102744 | 1 |
| Atp6v0e | 0.030234 | 0.009812 | 1 |
| Bcap29 | 0.030229 | 0.187472 | 1 |
| Tmem109 | 0.0302 | 0.043756 | 1 |
| Nfatc2ip | 0.030181 | 0.06558 | 1 |
| Cep63 | 0.030174 | 0.379921 | 1 |
| Vma21 | 0.030173 | 0.02181 | 1 |
| Mrpl46 | 0.030128 | 0.003307 | 1 |
| Hnrnpm | 0.030075 | 0.172645 | 1 |
| Eif1a | 0.029974 | 0.132245 | 1 |
| Lonrf1 | 0.02993 | 0.003623 | 1 |
| Tnfrsf1a | 0.02992 | 0.027365 | 1 |
| Tpd52 | 0.029874 | 0.014812 | 1 |
| Tmem41b | 0.029851 | 0.024102 | 1 |
| Mgat4a | 0.029845 | 0.1446 | 1 |
| Cdipt | 0.02983 | 0.083269 | 1 |
| Snrnp35 | 0.029815 | 0.02502 | 1 |
| Bag1 | 0.029809 | 0.083555 | 1 |
| Plod3 | 0.029799 | 0.035089 | 1 |
| Ubxn1 | 0.02977 | 0.008566 | 1 |
| Sub1 | 0.029731 | 0.073751 | 1 |
| Snx5 | 0.029718 | 0.109519 | 1 |
| Nme2 | 0.029712 | 0.018648 | 1 |
| Trim7 | 0.029592 | 0.345777 | 1 |
| Smim10l1 | 0.02955 | 0.018204 | 1 |
| Cenpv | 0.029451 | 0.132796 | 1 |
| Hcfc1 | 0.029429 | 0.03532 | 1 |
| Tmem223 | 0.029397 | 0.114107 | 1 |
| Eef1g | 0.029379 | 0.154564 | 1 |
| Glrx3 | 0.02935 | 0.127046 | 1 |
| Tmem141 | 0.029333 | 0.040841 | 1 |
| Hspa14 | 0.029324 | 0.01288 | 1 |
| 4632427E13Rik | 0.029323 | 0.351009 | 1 |
| 2810402E24Rik | 0.029316 | 0.002325 | 1 |
| Dhps | 0.029301 | 0.095024 | 1 |
| Snrpa1 | 0.029288 | 0.039149 | 1 |
| Gm30198 | 0.029279 | 0.063536 | 1 |
| Tomm22 | 0.029242 | 0.106834 | 1 |
| Cnot8 | 0.029238 | 0.547264 | 1 |
| Ipp | 0.029189 | 0.127735 | 1 |

| | | | |
|---|---|---|---|
| Csf1r | 0.029182 | 0.013641 | 1 |
| Bag3 | 0.029173 | 0.056464 | 1 |
| Tbpl1 | 0.029142 | 0.240374 | 1 |
| Fbxo3 | 0.029141 | 0.015059 | 1 |
| Tbc1d9b | 0.029133 | 0.219819 | 1 |
| Mrto4 | 0.029129 | 0.001871 | 1 |
| Arhgap35 | 0.02911 | 0.466243 | 1 |
| Uggt2 | 0.029099 | 0.390374 | 1 |
| Ryk | 0.029077 | 0.400681 | 1 |
| Jtb | 0.029037 | 0.013083 | 1 |
| Taf13 | 0.029028 | 0.065296 | 1 |
| Mfap1a | 0.028965 | 0.073938 | 1 |
| Gm33858 | 0.028941 | 0.032305 | 1 |
| Rwdd1 | 0.028938 | 0.053072 | 1 |
| Tmbim6 | 0.02892 | 0.169858 | 1 |
| Fance | 0.028918 | 0.063999 | 1 |
| Trappc2l | 0.028849 | 0.15115 | 1 |
| Trp53cor1 | 0.028816 | 0.849008 | 1 |
| A630089N07Rik | 0.028743 | 0.651663 | 1 |
| Zmpste24 | 0.028735 | 0.15364 | 1 |
| G3bp1 | 0.028722 | 0.178737 | 1 |
| Clgn | 0.028673 | 0.007455 | 1 |
| Ssna1 | 0.028635 | 0.050324 | 1 |
| Slc8b1 | 0.028624 | 0.041708 | 1 |
| Grina | 0.028618 | 0.009478 | 1 |
| Phgdh | 0.028611 | 0.090559 | 1 |
| Ebi3 | 0.02859 | 0.052352 | 1 |
| Wdpcp | 0.028536 | 0.654067 | 1 |
| Cltb | 0.028473 | 0.104659 | 1 |
| Srrm2 | 0.028467 | 0.007631 | 1 |
| Eif4ebp1 | 0.028457 | 0.005535 | 1 |
| Tm2d3 | 0.028451 | 0.085604 | 1 |
| Qpctl | 0.028433 | 0.573333 | 1 |
| Psat1 | 0.028408 | 0.043221 | 1 |
| Psmc6 | 0.028393 | 0.394087 | 1 |
| Olfr111 | 0.02838 | 0.205762 | 1 |
| Il11ra1 | 0.028355 | 0.140525 | 1 |
| Agpat2 | 0.028331 | 0.026962 | 1 |
| Msrb2 | 0.028315 | 0.136119 | 1 |
| Ndufb5 | 0.028315 | 0.047361 | 1 |
| Rmnd5a | 0.02831 | 0.102498 | 1 |
| Slc17a5 | 0.028282 | 0.408471 | 1 |
| Spi1 | 0.028281 | 0.032738 | 1 |
| H2-M3 | 0.028247 | 0.002966 | 1 |
| Sigmar1 | 0.028206 | 0.011559 | 1 |
| Psmc3 | 0.028186 | 0.060374 | 1 |
| 4933406I18Rik | 0.028167 | 0.4167 | 1 |
| Adi1 | 0.028159 | 0.005515 | 1 |

| | | | |
|---|---|---|---|
| Lefty1 | 0.028053 | 0.000167 | 1 |
| Kpnb1 | 0.028046 | 0.015318 | 1 |
| Crybb1 | 0.028041 | 0.774065 | 1 |
| Leng9 | 0.028035 | 0.043856 | 1 |
| Garnl3 | 0.028005 | 0.342575 | 1 |
| Tmem234 | 0.027985 | 0.14123 | 1 |
| Mrps12 | 0.027983 | 0.011757 | 1 |
| Slmap | 0.027954 | 0.080415 | 1 |
| Fgfr1op2 | 0.027944 | 0.205418 | 1 |
| Supt4a | 0.027905 | 0.060701 | 1 |
| Ndufa1 | 0.027891 | 0.012982 | 1 |
| Sec63 | 0.027827 | 0.628121 | 1 |
| Prkacb | 0.027812 | 0.281409 | 1 |
| Adam3 | 0.027775 | 0.189669 | 1 |
| Polr2j | 0.027766 | 0.175296 | 1 |
| Arl5b | 0.027727 | 0.029427 | 1 |
| Mrs2 | 0.02769 | 0.773356 | 1 |
| Dnajc17 | 0.027639 | 0.041241 | 1 |
| Rnf11 | 0.027588 | 0.066695 | 1 |
| Mrpl12 | 0.027572 | 0.052233 | 1 |
| Ncf4 | 0.027546 | 0.241546 | 1 |
| Fcgr1 | 0.027541 | 0.08595 | 1 |
| AI504432 | 0.027506 | 0.031119 | 1 |
| Dusp16 | 0.027499 | 0.155509 | 1 |
| Hvcn1 | 0.027482 | 0.033083 | 1 |
| Mmd | 0.027465 | 0.215192 | 1 |
| Sf3b5 | 0.027462 | 0.023758 | 1 |
| Lman2 | 0.027453 | 0.018877 | 1 |
| P2ry13 | 0.027398 | 0.090451 | 1 |
| Naxd | 0.027321 | 0.099882 | 1 |
| Fkbp3 | 0.027293 | 0.351399 | 1 |
| Hnrnpa1 | 0.027291 | 0.027272 | 1 |
| Smarcd1 | 0.027226 | 0.091583 | 1 |
| Zfp266 | 0.02721 | 0.339654 | 1 |
| Gm39469 | 0.027207 | 0.641647 | 1 |
| Slc39a12 | 0.027166 | 0.062355 | 1 |
| Slc25a33 | 0.027137 | 0.095751 | 1 |
| Dhrs7 | 0.02712 | 0.005127 | 1 |
| Hp1bp3 | 0.027096 | 0.134516 | 1 |
| Morf4l2 | 0.02709 | 0.049063 | 1 |
| Spcs3 | 0.027058 | 0.050356 | 1 |
| Abcg2 | 0.027049 | 0.178755 | 1 |
| Junos | 0.027036 | 0.076783 | 1 |
| Uba6 | 0.027029 | 0.995596 | 1 |
| Pds5b | 0.027018 | 0.223101 | 1 |
| Svbp | 0.026978 | 0.026925 | 1 |
| Polr2e | 0.026958 | 0.006376 | 1 |
| Arap3 | 0.026901 | 0.311802 | 1 |

| | | | |
|---|---|---|---|
| Ccni | 0.026896 | 0.056239 | 1 |
| Chchd5 | 0.026877 | 0.252169 | 1 |
| Ston2 | 0.026864 | 0.1398 | 1 |
| Tomm20 | 0.026847 | 0.027311 | 1 |
| Ap1m1 | 0.026831 | 0.268887 | 1 |
| Marcksl1 | 0.026825 | 0.058468 | 1 |
| Pdlim4 | 0.026805 | 0.029302 | 1 |
| Sh3bp5 | 0.026795 | 0.145389 | 1 |
| Fcrl1 | 0.026759 | 0.389082 | 1 |
| Slc2a1 | 0.026753 | 0.06851 | 1 |
| Dmac1 | 0.026751 | 0.052656 | 1 |
| Sf3b4 | 0.026739 | 0.042668 | 1 |
| Npc1 | 0.026727 | 0.101952 | 1 |
| Ssu72 | 0.026678 | 0.221842 | 1 |
| Ap2m1 | 0.026645 | 0.292157 | 1 |
| Stub1 | 0.026643 | 0.083192 | 1 |
| Pigm | 0.026609 | 0.5407 | 1 |
| Alyref | 0.026551 | 0.250882 | 1 |
| Clns1a | 0.026526 | 0.272641 | 1 |
| Smc3 | 0.026516 | 0.413753 | 1 |
| Gnpda2 | 0.026507 | 0.051025 | 1 |
| Ecsit | 0.026473 | 0.317639 | 1 |
| Nbas | 0.026462 | 0.158633 | 1 |
| Puf60 | 0.026427 | 0.062624 | 1 |
| Zdhhc3 | 0.026406 | 0.564685 | 1 |
| Bub3 | 0.026398 | 0.198973 | 1 |
| Nudt16l1 | 0.026397 | 0.058077 | 1 |
| Ralgps1 | 0.026387 | 0.416775 | 1 |
| Srsf11 | 0.026387 | 0.011967 | 1 |
| Dpagt1 | 0.026379 | 0.095014 | 1 |
| Fyn | 0.026377 | 0.69839 | 1 |
| Hacd3 | 0.026367 | 0.247801 | 1 |
| D830025C05Rik | 0.026352 | 0.016457 | 1 |
| Rnf13 | 0.026333 | 0.08903 | 1 |
| Rassf1 | 0.026333 | 0.157351 | 1 |
| Naxe | 0.026309 | 0.050615 | 1 |
| Susd3 | 0.026303 | 0.182614 | 1 |
| Cnnm3 | 0.0263 | 0.04067 | 1 |
| Nucb1 | 0.026287 | 0.088379 | 1 |
| Sept10 | 0.026285 | 0.608523 | 1 |
| Ppp1ca | 0.02628 | 0.033856 | 1 |
| Vps52 | 0.026249 | 0.019436 | 1 |
| Tmem11 | 0.026246 | 0.139999 | 1 |
| Hist2h2aa1 | 0.026221 | 0.003349 | 1 |
| Med30 | 0.026178 | 0.020955 | 1 |
| Lamtor3 | 0.026142 | 0.382756 | 1 |
| Slc15a4 | 0.026141 | 0.058698 | 1 |
| Irf2bp2 | 0.026139 | 0.185558 | 1 |

| | | | |
|---|---|---|---|
| Cebpz | 0.026134 | 0.025428 | 1 |
| Snx18 | 0.026127 | 0.082569 | 1 |
| Fuca2 | 0.026107 | 0.165918 | 1 |
| Sec11a | 0.026097 | 0.037516 | 1 |
| Smo | 0.026081 | 0.195969 | 1 |
| Srsf4 | 0.026075 | 0.591612 | 1 |
| Cct4 | 0.026058 | 0.120728 | 1 |
| Rbis | 0.026033 | 0.044322 | 1 |
| Fbxw9 | 0.026005 | 0.653796 | 1 |
| Coa3 | 0.025989 | 0.150728 | 1 |
| Pde4d | 0.025979 | 0.115728 | 1 |
| Blvrb | 0.025965 | 0.257486 | 1 |
| Fam110a | 0.025964 | 0.021714 | 1 |
| Prag1 | 0.025964 | 0.073013 | 1 |
| Smagp | 0.025944 | 0.080984 | 1 |
| Grn | 0.025923 | 0.259761 | 1 |
| Tspan13 | 0.025906 | 0.00935 | 1 |
| Cd302 | 0.025902 | 0.192036 | 1 |
| Pola1 | 0.025901 | 0.560687 | 1 |
| Ckap4 | 0.025893 | 0.028706 | 1 |
| Surf1 | 0.025875 | 0.107663 | 1 |
| Akirin1 | 0.025869 | 0.198686 | 1 |
| Iffo1 | 0.025852 | 0.376342 | 1 |
| Mtfr1l | 0.025852 | 0.635418 | 1 |
| Tmem159 | 0.025834 | 0.438985 | 1 |
| Rab11b | 0.025808 | 0.05944 | 1 |
| Fam167b | 0.025806 | 0.1531 | 1 |
| Pde12 | 0.025772 | 0.140609 | 1 |
| Ankra2 | 0.025769 | 0.019046 | 1 |
| Rcn2 | 0.025758 | 0.012362 | 1 |
| Sf3b1 | 0.02575 | 0.025937 | 1 |
| Gpat4 | 0.025723 | 0.129389 | 1 |
| Ppp2r2d | 0.025653 | 0.289533 | 1 |
| Trappc6a | 0.025628 | 0.05777 | 1 |
| Leng8 | 0.025613 | 0.269291 | 1 |
| Ndufaf2 | 0.025591 | 0.46539 | 1 |
| Tsc1 | 0.025553 | 0.255672 | 1 |
| Abcb4 | 0.025547 | 0.025712 | 1 |
| Nxf1 | 0.025502 | 0.248554 | 1 |
| Zfp653 | 0.025492 | 0.260762 | 1 |
| Mr1 | 0.025483 | 0.08179 | 1 |
| Ctss | 0.025457 | 0.027079 | 1 |
| Decr1 | 0.025443 | 0.254025 | 1 |
| Bag5 | 0.025417 | 0.122547 | 1 |
| Set | 0.025398 | 0.041853 | 1 |
| St6galnac4 | 0.025397 | 0.004658 | 1 |
| Telo2 | 0.025351 | 0.104336 | 1 |
| Tbcb | 0.02533 | 0.477219 | 1 |

| | | | |
|---|---|---|---|
| Sec61a1 | 0.025249 | 0.25519 | 1 |
| Ppm1a | 0.025242 | 0.109259 | 1 |
| Fam114a2 | 0.02522 | 0.127927 | 1 |
| Psmb8 | 0.025183 | 0.127871 | 1 |
| Arhgap31 | 0.025177 | 0.372932 | 1 |
| Disc1 | 0.025172 | 0.270452 | 1 |
| Pex5 | 0.025155 | 0.052811 | 1 |
| Mapk6 | 0.025132 | 0.20349 | 1 |
| Msl3 | 0.025098 | 0.052741 | 1 |
| Cuedc2 | 0.025083 | 0.103895 | 1 |
| Ppp6r1 | 0.025055 | 0.50551 | 1 |
| Isy1 | 0.025046 | 0.133617 | 1 |
| Tm7sf3 | 0.025046 | 0.04055 | 1 |
| Trim65 | 0.025046 | 0.473504 | 1 |
| Zdhhc5 | 0.02502 | 0.151846 | 1 |
| Yif1a | 0.025014 | 0.0057 | 1 |
| Ahsa2 | 0.025006 | 0.071259 | 1 |
| Map2k3 | 0.024983 | 0.116122 | 1 |
| Tpd52l2 | 0.024965 | 0.025535 | 1 |
| Mtch2 | 0.024948 | 0.057577 | 1 |
| Srrm1 | 0.024944 | 0.329009 | 1 |
| Kctd6 | 0.024918 | 0.023481 | 1 |
| Csnk2b | 0.024909 | 0.877618 | 1 |
| Exosc3 | 0.024898 | 0.011539 | 1 |
| Arf2 | 0.024892 | 0.106375 | 1 |
| Selenot | 0.024892 | 0.137013 | 1 |
| Prorsd1 | 0.024853 | 0.092808 | 1 |
| Pcbp1 | 0.02484 | 0.258219 | 1 |
| Erlec1 | 0.024814 | 0.165494 | 1 |
| Nol11 | 0.024802 | 0.512106 | 1 |
| Gba | 0.02475 | 0.425435 | 1 |
| Xylt1 | 0.024749 | 0.022008 | 1 |
| Rnf187 | 0.024726 | 0.327735 | 1 |
| Ly6i | 0.024705 | 0.119538 | 1 |
| Ints11 | 0.024705 | 0.86954 | 1 |
| Mrpl33 | 0.024653 | 0.103599 | 1 |
| Kdsr | 0.024632 | 0.031121 | 1 |
| Cyb5r3 | 0.024577 | 0.226706 | 1 |
| Glipr1 | 0.024561 | 0.162612 | 1 |
| Atp6ap2 | 0.024544 | 0.021199 | 1 |
| Arfip1 | 0.024541 | 0.017982 | 1 |
| Ttc7b | 0.024514 | 0.181738 | 1 |
| Tspyl2 | 0.024503 | 0.00017 | 1 |
| Gm9949 | 0.024484 | 0.024982 | 1 |
| Setd3 | 0.02448 | 0.369207 | 1 |
| Tceal9 | 0.02445 | 0.061767 | 1 |
| Prpf19 | 0.024438 | 0.293555 | 1 |
| Flii | 0.024432 | 0.569488 | 1 |

| | | | |
|---|---|---|---|
| Rnf121 | 0.024382 | 0.7165 | 1 |
| Gm31410 | 0.02438 | 0.898686 | 1 |
| Agk | 0.024338 | 0.079954 | 1 |
| Armcx3 | 0.024337 | 0.070794 | 1 |
| Pdcd5 | 0.024336 | 0.20434 | 1 |
| Ncstn | 0.024236 | 0.032775 | 1 |
| Vwa5a | 0.024185 | 0.269107 | 1 |
| Igsf8 | 0.024138 | 0.309854 | 1 |
| Luc7l2 | 0.024128 | 0.170176 | 1 |
| Rngtt | 0.024128 | 0.165615 | 1 |
| Urb2 | 0.024126 | 0.169837 | 1 |
| Ccl6 | 0.024103 | 0.860165 | 1 |
| Cib1 | 0.024095 | 0.0691 | 1 |
| Ccng2 | 0.024061 | 0.024187 | 1 |
| Cyp51 | 0.024043 | 0.041029 | 1 |
| Cetn2 | 0.024028 | 0.067609 | 1 |
| Slc29a1 | 0.024014 | 0.155146 | 1 |
| Map7 | 0.024012 | 0.03424 | 1 |
| Rcl1 | 0.024005 | 0.048297 | 1 |
| Sdf2 | 0.024003 | 0.362049 | 1 |
| Mad2l2 | 0.023973 | 0.055823 | 1 |
| Uggt1 | 0.023965 | 0.107224 | 1 |
| Mtln | 0.023933 | 0.099715 | 1 |
| Pabpc1 | 0.023927 | 0.226707 | 1 |
| Lag3 | 0.023924 | 0.096078 | 1 |
| Ran | 0.023911 | 0.008967 | 1 |
| Nhlrc3 | 0.023911 | 0.097492 | 1 |
| Vti1b | 0.023909 | 0.069276 | 1 |
| Eif3f | 0.023897 | 0.006634 | 1 |
| Ash2l | 0.023877 | 0.504077 | 1 |
| Eef1e1 | 0.023852 | 0.027369 | 1 |
| Eva1b | 0.023837 | 0.12736 | 1 |
| Gm20721 | 0.023832 | 0.007211 | 1 |
| Timm17b | 0.023811 | 0.001176 | 1 |
| Akap1 | 0.023809 | 0.305298 | 1 |
| Cyth2 | 0.023793 | 0.002115 | 1 |
| Snhg15 | 0.023788 | 0.024521 | 1 |
| Copz1 | 0.023773 | 0.67765 | 1 |
| Atp6v1d | 0.023749 | 0.117802 | 1 |
| Rgs7bp | 0.023711 | 0.105268 | 1 |
| Ntmt1 | 0.02371 | 0.176125 | 1 |
| Zfp846 | 0.02371 | 0.487714 | 1 |
| Fam133b | 0.023709 | 0.100867 | 1 |
| Bclaf3 | 0.023688 | 0.540684 | 1 |
| Arl1 | 0.023656 | 0.008456 | 1 |
| Tnip2 | 0.02364 | 0.149078 | 1 |
| Dhrs3 | 0.023633 | 0.031937 | 1 |
| Ms4a6d | 0.023617 | 0.151777 | 1 |

| | | | |
|---|---|---|---|
| Nudcd2 | 0.023612 | 0.496155 | 1 |
| Syf2 | 0.023602 | 0.1213 | 1 |
| Gm48855 | 0.0236 | 0.851446 | 1 |
| Usp28 | 0.023596 | 0.022418 | 1 |
| Armc10 | 0.023592 | 0.407005 | 1 |
| Atxn7l3b | 0.023584 | 0.476046 | 1 |
| Cdc27 | 0.023564 | 0.717316 | 1 |
| Lgals3bp | 0.023563 | 0.165737 | 1 |
| Dhx29 | 0.023494 | 0.957747 | 1 |
| Rbsn | 0.023482 | 0.084816 | 1 |
| Ndufv3 | 0.02348 | 0.155624 | 1 |
| Per1 | 0.023472 | 0.016212 | 1 |
| Lmbrd2 | 0.023464 | 0.139442 | 1 |
| Zfp335os | 0.023463 | 0.007318 | 1 |
| Gpr155 | 0.02345 | 0.366127 | 1 |
| Vcp | 0.023449 | 0.154237 | 1 |
| Lnpk | 0.02339 | 0.21576 | 1 |
| Lacc1 | 0.023388 | 0.191086 | 1 |
| Tgm2 | 0.023374 | 0.506693 | 1 |
| Snapc3 | 0.02337 | 0.102404 | 1 |
| Xlr | 0.023357 | 0.071389 | 1 |
| Eif4a3 | 0.023356 | 0.082661 | 1 |
| Matr3 | 0.023349 | 0.358048 | 1 |
| Agtrap | 0.023341 | 0.050785 | 1 |
| Tspan18 | 0.02334 | 0.166889 | 1 |
| Mospd2 | 0.023322 | 0.347166 | 1 |
| Nus1 | 0.023296 | 0.501531 | 1 |
| Gm30054 | 0.023274 | 0.016833 | 1 |
| Il6 | 0.023272 | 0.085113 | 1 |
| Magt1 | 0.023259 | 0.055067 | 1 |
| Zfyve21 | 0.02325 | 0.041207 | 1 |
| Atp6v1f | 0.023243 | 0.031602 | 1 |
| Plxnb2 | 0.023232 | 0.077879 | 1 |
| Rabep2 | 0.023187 | 0.065758 | 1 |
| Klhdc4 | 0.023146 | 0.007041 | 1 |
| Myo1b | 0.023145 | 0.414279 | 1 |
| Tmem60 | 0.023143 | 0.054576 | 1 |
| Sap30bp | 0.023115 | 0.276871 | 1 |
| Colgalt1 | 0.023093 | 0.221847 | 1 |
| CAAA01118383.1 | 0.023082 | 0.247961 | 1 |
| Upf3a | 0.023076 | 0.133735 | 1 |
| Rab5if | 0.023059 | 0.106823 | 1 |
| Haghl | 0.02305 | 0.161463 | 1 |
| Ccdc47 | 0.023042 | 0.098522 | 1 |
| 4930581F22Rik | 0.023039 | 0.119951 | 1 |
| Ndufa2 | 0.023034 | 0.049943 | 1 |
| Aprt | 0.022955 | 0.167718 | 1 |
| Khsrp | 0.022954 | 0.913915 | 1 |

| | | | |
|---|---|---|---|
| Fancl | 0.022937 | 0.595171 | 1 |
| Ndufb7 | 0.022925 | 0.060609 | 1 |
| Cnot6 | 0.02292 | 0.116498 | 1 |
| Commd7 | 0.022896 | 0.073375 | 1 |
| Tmem9 | 0.022895 | 0.022103 | 1 |
| Bcl7b | 0.022885 | 0.066091 | 1 |
| Mpc2 | 0.022867 | 0.168613 | 1 |
| Cbx4 | 0.022863 | 0.019677 | 1 |
| Tuba1c | 0.022858 | 0.201124 | 1 |
| Acp1 | 0.022829 | 0.100419 | 1 |
| Top1 | 0.022739 | 0.043852 | 1 |
| Pdcl | 0.02272 | 0.102443 | 1 |
| Zbtb10 | 0.022674 | 0.091778 | 1 |
| Lpcat2 | 0.022641 | 0.041361 | 1 |
| Paqr7 | 0.022609 | 0.263815 | 1 |
| Arhgap19 | 0.022606 | 0.352299 | 1 |
| C1qbp | 0.022589 | 0.210784 | 1 |
| Gsr | 0.022578 | 0.77496 | 1 |
| Tmem251 | 0.022563 | 0.135683 | 1 |
| Tppp | 0.022541 | 0.309057 | 1 |
| H2-Q6 | 0.022535 | 0.150803 | 1 |
| Dtnb | 0.022469 | 0.13313 | 1 |
| Peak1 | 0.022468 | 0.159637 | 1 |
| Ipo5 | 0.022433 | 0.224046 | 1 |
| 5730480H06Rik | 0.022424 | 0.487552 | 1 |
| Sra1 | 0.022409 | 0.021107 | 1 |
| Ndufa11 | 0.022405 | 0.026906 | 1 |
| Trmt112 | 0.022392 | 0.02106 | 1 |
| Exosc10 | 0.022377 | 0.116764 | 1 |
| Csnk1g2 | 0.022349 | 0.2218 | 1 |
| Emc4 | 0.022349 | 0.370776 | 1 |
| Apmap | 0.022344 | 0.122374 | 1 |
| Ppp2r1a | 0.022336 | 0.071475 | 1 |
| Prxl2b | 0.022329 | 0.04345 | 1 |
| Ercc5 | 0.022277 | 0.335182 | 1 |
| Enc1 | 0.022271 | 0.325156 | 1 |
| Hbs1l | 0.022253 | 0.200275 | 1 |
| Glyr1 | 0.022253 | 0.769683 | 1 |
| Fmc1 | 0.022223 | 0.210557 | 1 |
| Wdr76 | 0.022198 | 0.07947 | 1 |
| Epb41l2 | 0.022194 | 0.083462 | 1 |
| Nfe2l1 | 0.022193 | 0.41246 | 1 |
| Actr1b | 0.022137 | 0.515673 | 1 |
| Golga4 | 0.022087 | 0.059947 | 1 |
| Zfp143 | 0.02206 | 0.353185 | 1 |
| Ctbs | 0.022055 | 0.265434 | 1 |
| Sgpp1 | 0.022053 | 0.772817 | 1 |
| Plekhg5 | 0.022052 | 0.076058 | 1 |

| | | | |
|---|---|---|---|
| Naga | 0.022019 | 0.103816 | 1 |
| Klhl9 | 0.022006 | 0.580043 | 1 |
| Spata24 | 0.021999 | 0.028536 | 1 |
| Cybc1 | 0.021992 | 0.097491 | 1 |
| Hmga1 | 0.021972 | 0.021415 | 1 |
| Fopnl | 0.021948 | 0.736609 | 1 |
| Ubald1 | 0.021944 | 0.015631 | 1 |
| Gmpr | 0.021931 | 0.108236 | 1 |
| Tpbgl | 0.021922 | 0.45684 | 1 |
| Srsf6 | 0.02192 | 0.013461 | 1 |
| Aldh9a1 | 0.021889 | 0.273345 | 1 |
| Nlrp3 | 0.021885 | 0.258647 | 1 |
| Rmdn1 | 0.021846 | 0.014305 | 1 |
| Timm50 | 0.021814 | 0.114362 | 1 |
| Eif2s2 | 0.021806 | 0.061482 | 1 |
| Gimap1 | 0.021781 | 0.036316 | 1 |
| Ago4 | 0.021775 | 0.410264 | 1 |
| Jagn1 | 0.021761 | 0.10641 | 1 |
| Ly86 | 0.021725 | 0.606529 | 1 |
| Iah1 | 0.021691 | 0.07132 | 1 |
| Nemp2 | 0.021635 | 0.149386 | 1 |
| 4930522L14Rik | 0.021613 | 0.163469 | 1 |
| Rnf2 | 0.021548 | 0.366857 | 1 |
| Slc39a6 | 0.021543 | 0.184389 | 1 |
| Pou2f2 | 0.02153 | 0.716003 | 1 |
| Arhgap11a | 0.021502 | 0.696805 | 1 |
| Zfp622 | 0.021488 | 0.223552 | 1 |
| Gk | 0.021479 | 0.289176 | 1 |
| Cfap20 | 0.021459 | 0.576588 | 1 |
| Eif4enif1 | 0.021452 | 0.646422 | 1 |
| Idh1 | 0.021445 | 0.050114 | 1 |
| Rhoc | 0.021413 | 0.022392 | 1 |
| Zfp729b | 0.021401 | 0.899625 | 1 |
| Phb2 | 0.02129 | 0.027036 | 1 |
| B3galt6 | 0.021273 | 0.701373 | 1 |
| Dnajc30 | 0.021272 | 0.452398 | 1 |
| Rxylt1 | 0.021265 | 0.007159 | 1 |
| Snu13 | 0.021263 | 0.058132 | 1 |
| Tctex1d2 | 0.021248 | 0.040413 | 1 |
| Serbp1 | 0.021222 | 0.352441 | 1 |
| Manbal | 0.021212 | 0.082354 | 1 |
| Fyttd1 | 0.021183 | 0.276113 | 1 |
| Rmc1 | 0.021182 | 0.132181 | 1 |
| Coro1b | 0.021173 | 0.014559 | 1 |
| Zfand2a | 0.021152 | 0.358186 | 1 |
| Fundc2 | 0.021142 | 0.065951 | 1 |
| Ppp3cb | 0.021113 | 0.177787 | 1 |
| Ddx49 | 0.021103 | 0.605045 | 1 |

| | | | |
|---|---|---|---|
| Ost4 | 0.021087 | 0.124429 | 1 |
| 2610002M06Rik | 0.021077 | 0.112551 | 1 |
| Eif2b1 | 0.021033 | 0.107731 | 1 |
| Slc48a1 | 0.02101 | 0.020698 | 1 |
| Erg28 | 0.021002 | 0.303253 | 1 |
| Hnrnpf | 0.020992 | 0.014082 | 1 |
| Isyna1 | 0.020988 | 0.505268 | 1 |
| Ciao2a | 0.020986 | 0.073894 | 1 |
| Ctdnep1 | 0.020985 | 0.072696 | 1 |
| Adrm1 | 0.020978 | 0.110019 | 1 |
| Prdx6 | 0.020976 | 0.082594 | 1 |
| Lamtor5 | 0.020957 | 0.196655 | 1 |
| Vhl | 0.020951 | 0.783594 | 1 |
| Tle5 | 0.020921 | 0.390049 | 1 |
| Nudt16 | 0.020915 | 0.073407 | 1 |
| Znrd1 | 0.020912 | 0.746308 | 1 |
| A930024E05Rik | 0.020897 | 0.205932 | 1 |
| Pstk | 0.020894 | 0.100693 | 1 |
| Gmfg | 0.020892 | 0.177019 | 1 |
| Rabepk | 0.020884 | 0.034681 | 1 |
| Smpd4 | 0.020884 | 0.039845 | 1 |
| Phlpp1 | 0.020861 | 0.333835 | 1 |
| Oxa1l | 0.020859 | 0.12925 | 1 |
| Yy1 | 0.020854 | 0.300041 | 1 |
| Gps2 | 0.020814 | 0.075903 | 1 |
| Tango6 | 0.020803 | 0.691115 | 1 |
| Lta4h | 0.020802 | 0.719634 | 1 |
| Zfp799 | 0.020781 | 0.139818 | 1 |
| Cmpk1 | 0.020772 | 0.274002 | 1 |
| Fdps | 0.020725 | 0.260122 | 1 |
| Ddx1 | 0.020716 | 0.218208 | 1 |
| Nudt9 | 0.02071 | 0.018859 | 1 |
| Trim24 | 0.020686 | 0.09491 | 1 |
| Klf6 | 0.020662 | 0.620273 | 1 |
| Ostf1 | 0.020648 | 0.069877 | 1 |
| Klhl15 | 0.020629 | 0.374862 | 1 |
| Ndufs5 | 0.020607 | 0.062318 | 1 |
| Noc3l | 0.02058 | 0.662872 | 1 |
| Ctdspl | 0.020567 | 0.202922 | 1 |
| Hspd1 | 0.020556 | 0.067397 | 1 |
| Srpr | 0.020539 | 0.070027 | 1 |
| Ids | 0.020538 | 0.271381 | 1 |
| Fut10 | 0.020502 | 0.447272 | 1 |
| Srsf3 | 0.0205 | 0.112734 | 1 |
| Ndufb10 | 0.020499 | 0.023539 | 1 |
| Mettl14 | 0.020497 | 0.298981 | 1 |
| Srrd | 0.020483 | 0.087268 | 1 |
| Lage3 | 0.020477 | 0.263614 | 1 |

| | | | |
|---|---|---|---|
| Faim | 0.020477 | 0.083197 | 1 |
| Surf2 | 0.020426 | 0.015042 | 1 |
| Dhx36 | 0.020404 | 0.027236 | 1 |
| 2310015A10Rik | 0.020398 | 0.276715 | 1 |
| Trir | 0.020381 | 0.024916 | 1 |
| Zfpl1 | 0.020374 | 0.122543 | 1 |
| Vdac2 | 0.020362 | 0.085495 | 1 |
| Timm22 | 0.020361 | 0.09714 | 1 |
| Zfp704 | 0.020342 | 0.696006 | 1 |
| Psmd4 | 0.020323 | 0.046458 | 1 |
| Bnip1 | 0.020319 | 0.721194 | 1 |
| Ube2s | 0.020299 | 0.032098 | 1 |
| Ebag9 | 0.020289 | 0.478192 | 1 |
| Alkbh2 | 0.020264 | 0.172533 | 1 |
| Tmem173 | 0.020252 | 0.386488 | 1 |
| Cwc15 | 0.020225 | 0.297272 | 1 |
| Senp6 | 0.020172 | 0.189191 | 1 |
| Mea1 | 0.020171 | 0.152374 | 1 |
| Scp2 | 0.020166 | 0.135556 | 1 |
| Sec24c | 0.020161 | 0.019652 | 1 |
| Rpp21 | 0.020149 | 0.089757 | 1 |
| Homer3 | 0.02013 | 0.10973 | 1 |
| Ctbp1 | 0.020129 | 0.435538 | 1 |
| Hmgn2 | 0.020107 | 0.10691 | 1 |
| Nucks1 | 0.020098 | 0.024879 | 1 |
| Nudc | 0.020078 | 0.189564 | 1 |
| Atad3a | 0.020036 | 0.202284 | 1 |
| Naa38 | 0.020019 | 0.099841 | 1 |
| Snapc1 | 0.020013 | 0.63556 | 1 |
| Ufl1 | 0.020006 | 0.76105 | 1 |
| Rapgef5 | 0.019998 | 0.606229 | 1 |
| H2afv | 0.019982 | 0.265545 | 1 |
| P4hb | 0.019979 | 0.143154 | 1 |
| Snrpb2 | 0.019968 | 0.024958 | 1 |
| Ptbp1 | 0.019957 | 0.353481 | 1 |
| Selenoh | 0.019956 | 0.10989 | 1 |
| Aimp2 | 0.019933 | 0.027501 | 1 |
| Lsm1 | 0.019928 | 0.700549 | 1 |
| Nsmce1 | 0.019898 | 0.429024 | 1 |
| Polr2m | 0.019889 | 0.289043 | 1 |
| Siglece | 0.019888 | 0.319209 | 1 |
| Esyt2 | 0.019885 | 0.571534 | 1 |
| Ears2 | 0.019881 | 0.121087 | 1 |
| Thrsp | 0.019872 | 0.275389 | 1 |
| Src | 0.019862 | 0.123076 | 1 |
| Ldb1 | 0.019839 | 0.170623 | 1 |
| Rela | 0.019822 | 0.09735 | 1 |
| Atp5md | 0.019806 | 0.211621 | 1 |

| | | | |
|---|---|---|---|
| Mdm4 | 0.019788 | 0.813606 | 1 |
| Ints12 | 0.019788 | 0.152211 | 1 |
| Socs4 | 0.019753 | 0.122516 | 1 |
| Idua | 0.019725 | 0.119828 | 1 |
| Coa6 | 0.019712 | 0.122838 | 1 |
| Hjurp | 0.01971 | 0.319211 | 1 |
| Ccnd1 | 0.019696 | 0.29745 | 1 |
| Tceal8 | 0.019685 | 0.07445 | 1 |
| Bckdk | 0.019638 | 0.818068 | 1 |
| Slc39a10 | 0.019635 | 0.074622 | 1 |
| Dld | 0.019623 | 0.162272 | 1 |
| Egln2 | 0.019617 | 0.456747 | 1 |
| Zmym3 | 0.019597 | 0.116164 | 1 |
| Btg3 | 0.019557 | 0.019835 | 1 |
| Churc1 | 0.019539 | 0.020653 | 1 |
| Trmt13 | 0.019489 | 0.58528 | 1 |
| Zdhhc20 | 0.019488 | 0.080919 | 1 |
| Gps1 | 0.019466 | 0.239457 | 1 |
| Ramp1 | 0.019444 | 0.783301 | 1 |
| Med24 | 0.019443 | 0.567504 | 1 |
| Gxylt1 | 0.019408 | 0.350245 | 1 |
| Iffo2 | 0.019396 | 0.18838 | 1 |
| Cisd2 | 0.019369 | 0.93386 | 1 |
| Smad4 | 0.019361 | 0.228203 | 1 |
| Nsun6 | 0.019353 | 0.858819 | 1 |
| Pecam1 | 0.019346 | 0.453893 | 1 |
| Cln5 | 0.019344 | 0.171927 | 1 |
| Ipo9 | 0.019341 | 0.042676 | 1 |
| Cops8 | 0.019336 | 0.133507 | 1 |
| Atpif1 | 0.019328 | 0.019369 | 1 |
| Tgfbi | 0.019317 | 0.459445 | 1 |
| Fzr1 | 0.01931 | 0.032697 | 1 |
| Fam78a | 0.019295 | 0.228217 | 1 |
| Iqsec1 | 0.019235 | 0.404388 | 1 |
| Dnajc1 | 0.019224 | 0.362263 | 1 |
| Suds3 | 0.019209 | 0.547287 | 1 |
| Etfdh | 0.019187 | 0.679613 | 1 |
| Srp54a | 0.01915 | 0.276575 | 1 |
| Psmg4 | 0.019123 | 0.087178 | 1 |
| Mcrip1 | 0.019117 | 0.433277 | 1 |
| Prdx3 | 0.019113 | 0.285694 | 1 |
| Slc2a8 | 0.019111 | 0.107908 | 1 |
| Ppp3r1 | 0.019078 | 0.640171 | 1 |
| Haus8 | 0.01906 | 0.016758 | 1 |
| P2ry10b | 0.019053 | 0.190772 | 1 |
| Hsd3b7 | 0.019032 | 0.043157 | 1 |
| Uqcr10 | 0.01902 | 0.441522 | 1 |
| Mars | 0.018987 | 0.194585 | 1 |

| | | | |
|---|---|---|---|
| Gmeb2 | 0.018959 | 0.104563 | 1 |
| Dcun1d4 | 0.018949 | 0.90825 | 1 |
| Fibp | 0.018936 | 0.27234 | 1 |
| Arl10 | 0.018923 | 0.424284 | 1 |
| Slc30a5 | 0.018914 | 0.314527 | 1 |
| Hat1 | 0.018907 | 0.36761 | 1 |
| Trim36 | 0.018906 | 0.32058 | 1 |
| Trp53bp2 | 0.018874 | 0.013918 | 1 |
| Mrpl51 | 0.01886 | 0.347496 | 1 |
| Smim26 | 0.01885 | 0.428995 | 1 |
| Rab3il1 | 0.018847 | 0.128331 | 1 |
| Abhd2 | 0.018819 | 0.274132 | 1 |
| Actr5 | 0.018812 | 0.086813 | 1 |
| Psma3 | 0.018802 | 0.161659 | 1 |
| Adamts10 | 0.018799 | 0.188902 | 1 |
| Wipi1 | 0.018797 | 0.154763 | 1 |
| 2900093K20Rik | 0.018781 | 0.213612 | 1 |
| Ankrd10 | 0.018776 | 0.091876 | 1 |
| Rnf7 | 0.018756 | 0.057995 | 1 |
| Srp9 | 0.018728 | 0.29244 | 1 |
| Sumo3 | 0.018721 | 0.020897 | 1 |
| Tmem43 | 0.018647 | 0.228969 | 1 |
| Plin2 | 0.018637 | 0.520848 | 1 |
| Alg2 | 0.018609 | 0.137771 | 1 |
| Ttc32 | 0.018595 | 0.331654 | 1 |
| Ndfip1 | 0.018582 | 0.239761 | 1 |
| Banf1 | 0.018564 | 0.121717 | 1 |
| Hccs | 0.01855 | 0.423359 | 1 |
| B4gat1 | 0.018549 | 0.06589 | 1 |
| Cnrip1 | 0.01854 | 0.081833 | 1 |
| Chchd2 | 0.018529 | 0.002928 | 1 |
| Cfl2 | 0.018516 | 0.074682 | 1 |
| Tubgcp2 | 0.018505 | 0.269488 | 1 |
| Ccdc82 | 0.018458 | 0.209538 | 1 |
| Sema4g | 0.018457 | 0.190747 | 1 |
| Atp5mpl | 0.018446 | 0.205827 | 1 |
| Ppfibp2 | 0.018439 | 0.121581 | 1 |
| Stt3a | 0.018403 | 0.076069 | 1 |
| Pam16 | 0.018353 | 0.264402 | 1 |
| Ubn1 | 0.018353 | 0.167954 | 1 |
| Lman2l | 0.018334 | 0.413253 | 1 |
| Smn1 | 0.018331 | 0.18931 | 1 |
| Rab35 | 0.018325 | 0.292969 | 1 |
| Dctn3 | 0.01832 | 0.538418 | 1 |
| Ggps1 | 0.018319 | 0.66844 | 1 |
| Tm9sf2 | 0.018314 | 0.180835 | 1 |
| Cwf19l1 | 0.018305 | 0.255793 | 1 |
| Cul2 | 0.018298 | 0.803922 | 1 |

| | | | |
|---|---|---|---|
| Asah1 | 0.018267 | 0.180711 | 1 |
| Ptch1 | 0.018234 | 0.431835 | 1 |
| Lpcat3 | 0.018199 | 0.270873 | 1 |
| Pde8a | 0.018187 | 0.00889 | 1 |
| Htatsf1 | 0.018181 | 0.080322 | 1 |
| Ccdc28b | 0.018169 | 0.504785 | 1 |
| Arsk | 0.018167 | 0.099014 | 1 |
| Cpox | 0.018088 | 0.169069 | 1 |
| Epb41l3 | 0.018087 | 0.325468 | 1 |
| Mpp7 | 0.01808 | 0.317114 | 1 |
| Tcaim | 0.018075 | 0.809414 | 1 |
| Ddt | 0.018066 | 0.04743 | 1 |
| Ighmbp2 | 0.018056 | 0.291574 | 1 |
| Prdx5 | 0.018046 | 0.199424 | 1 |
| Hmgb1 | 0.018043 | 0.046952 | 1 |
| Pou6f1 | 0.018015 | 0.241862 | 1 |
| Aldh6a1 | 0.018014 | 0.044072 | 1 |
| Lrrc59 | 0.018006 | 0.069191 | 1 |
| Acvr1 | 0.017998 | 0.074644 | 1 |
| Noc2l | 0.017988 | 0.112386 | 1 |
| Zswim7 | 0.017978 | 0.012478 | 1 |
| Fnbp1l | 0.017943 | 0.470248 | 1 |
| Eea1 | 0.017932 | 0.168599 | 1 |
| Fam50a | 0.017924 | 0.116176 | 1 |
| Bace1 | 0.01788 | 0.299313 | 1 |
| Eif5a | 0.017872 | 0.02963 | 1 |
| Snip1 | 0.01787 | 0.13167 | 1 |
| Ctnna1 | 0.017841 | 0.943043 | 1 |
| Snrnp27 | 0.017837 | 0.595506 | 1 |
| Tut4 | 0.017829 | 0.403942 | 1 |
| Rlim | 0.017813 | 0.142035 | 1 |
| Mettl5 | 0.017772 | 0.404733 | 1 |
| Fut11 | 0.017745 | 0.193304 | 1 |
| Mrpl30 | 0.017722 | 0.607648 | 1 |
| Tuba1a | 0.01772 | 0.156865 | 1 |
| Flot2 | 0.017697 | 0.852635 | 1 |
| Slc52a2 | 0.01769 | 0.243994 | 1 |
| Lsm7 | 0.017687 | 0.563024 | 1 |
| Lims1 | 0.017679 | 0.407237 | 1 |
| Usp24 | 0.017657 | 0.442276 | 1 |
| Plxna4os1 | 0.01765 | 0.219921 | 1 |
| Clip1 | 0.017628 | 0.345789 | 1 |
| Lamc1 | 0.017606 | 0.230132 | 1 |
| Gsdme | 0.017568 | 0.147343 | 1 |
| Ncapd3 | 0.017558 | 0.589266 | 1 |
| Vps39 | 0.01754 | 0.168863 | 1 |
| Rbx1 | 0.017537 | 0.439572 | 1 |
| Aip | 0.017533 | 0.194927 | 1 |

| | | | |
|---|---|---|---|
| Tsn | 0.017521 | 0.031638 | 1 |
| Ift46 | 0.017513 | 0.197412 | 1 |
| Fus | 0.0175 | 0.29075 | 1 |
| B4galt5 | 0.017443 | 0.244313 | 1 |
| Rab4b | 0.017432 | 0.184561 | 1 |
| Dcakd | 0.017428 | 0.301171 | 1 |
| Was | 0.017411 | 0.211174 | 1 |
| Mrpl18 | 0.017411 | 0.470441 | 1 |
| Brk1 | 0.017403 | 0.013558 | 1 |
| Chmp2a | 0.017401 | 0.377465 | 1 |
| 0610012G03Rik | 0.017359 | 0.110325 | 1 |
| Hexdc | 0.017354 | 0.289697 | 1 |
| Amdhd2 | 0.017338 | 0.043584 | 1 |
| Yif1b | 0.017313 | 0.051062 | 1 |
| Mfsd5 | 0.017311 | 0.012156 | 1 |
| Firre | 0.017302 | 0.337256 | 1 |
| Crem | 0.017291 | 0.166864 | 1 |
| Ncbp2 | 0.01728 | 0.074247 | 1 |
| Prdx2 | 0.017262 | 0.167577 | 1 |
| Atn1 | 0.017255 | 0.802671 | 1 |
| Zfp422 | 0.017247 | 0.095085 | 1 |
| Epm2aip1 | 0.017242 | 0.217172 | 1 |
| Tiprl | 0.017216 | 0.167249 | 1 |
| Thumpd2 | 0.017214 | 0.126536 | 1 |
| Mettl17 | 0.01721 | 0.494489 | 1 |
| Abhd17b | 0.01719 | 0.514246 | 1 |
| Abcd4 | 0.017185 | 0.611058 | 1 |
| Zfp839 | 0.017179 | 0.773026 | 1 |
| Stx7 | 0.017135 | 0.29455 | 1 |
| Slc46a1 | 0.01713 | 0.868737 | 1 |
| Riok1 | 0.017072 | 0.295295 | 1 |
| Golph3 | 0.017058 | 0.259764 | 1 |
| LTO1 | 0.017056 | 0.260912 | 1 |
| St5 | 0.017051 | 0.32865 | 1 |
| Mtss1 | 0.017041 | 0.184616 | 1 |
| Gls | 0.017035 | 0.404819 | 1 |
| Ift22 | 0.017019 | 0.302593 | 1 |
| Plekha4 | 0.017009 | 0.285334 | 1 |
| Dhx16 | 0.016925 | 0.252722 | 1 |
| Gnai3 | 0.016921 | 0.432513 | 1 |
| Lemd2 | 0.016895 | 0.044057 | 1 |
| Ndufa8 | 0.016878 | 0.01574 | 1 |
| Serf2 | 0.016874 | 0.051463 | 1 |
| Pign | 0.016864 | 0.616366 | 1 |
| Sri | 0.016855 | 0.534697 | 1 |
| Gm26759 | 0.016808 | 0.001892 | 1 |
| Elmo1 | 0.016782 | 0.292139 | 1 |
| Mkrn2 | 0.016779 | 0.134632 | 1 |

| | | | |
|---|---|---|---|
| Gpaa1 | 0.016779 | 0.444892 | 1 |
| Imp3 | 0.016773 | 0.117999 | 1 |
| Tmx1 | 0.016767 | 0.219154 | 1 |
| Guca1a | 0.016766 | 0.054316 | 1 |
| Nol7 | 0.016765 | 0.050734 | 1 |
| Ift20 | 0.016765 | 0.204018 | 1 |
| Ap4e1 | 0.016755 | 0.674114 | 1 |
| Gm10790 | 0.016754 | 0.403111 | 1 |
| Phb | 0.01675 | 0.238329 | 1 |
| Capns1 | 0.016744 | 0.454081 | 1 |
| Prpsap1 | 0.016733 | 0.282338 | 1 |
| Tfam | 0.016719 | 0.690569 | 1 |
| Fam160b2 | 0.016718 | 0.419104 | 1 |
| Fbxl5 | 0.016711 | 0.234769 | 1 |
| Mrps22 | 0.016699 | 0.510174 | 1 |
| Rbbp7 | 0.016664 | 0.613052 | 1 |
| Pin1 | 0.016649 | 0.181973 | 1 |
| Psmd8 | 0.016637 | 0.054151 | 1 |
| 9130015G15Rik | 0.016636 | 0.74803 | 1 |
| Cd22 | 0.016633 | 0.389201 | 1 |
| Cycs | 0.016625 | 0.152355 | 1 |
| Ralgps2 | 0.016589 | 0.174232 | 1 |
| Timm10 | 0.016566 | 0.363897 | 1 |
| Las1l | 0.016562 | 0.776326 | 1 |
| Npat | 0.016542 | 0.478948 | 1 |
| 2810405F17Rik | 0.016533 | 0.312858 | 1 |
| Zfp91 | 0.016514 | 0.081112 | 1 |
| Thrap3 | 0.016508 | 0.182135 | 1 |
| Rbm34 | 0.016495 | 0.655337 | 1 |
| Cenpo | 0.016462 | 0.620534 | 1 |
| Zfp24 | 0.016447 | 0.7964 | 1 |
| Tprgl | 0.016435 | 0.621669 | 1 |
| Calm3 | 0.016434 | 0.111645 | 1 |
| Lrp10 | 0.016394 | 0.341325 | 1 |
| Pes1 | 0.016384 | 0.229115 | 1 |
| Pcif1 | 0.016379 | 0.140201 | 1 |
| E430024P14Rik | 0.016354 | 0.633052 | 1 |
| Ero1lb | 0.016353 | 0.175509 | 1 |
| Tax1bp3 | 0.016353 | 0.197149 | 1 |
| Ate1 | 0.016349 | 0.961879 | 1 |
| Hspb3 | 0.016335 | 0.1066 | 1 |
| Cep44 | 0.016304 | 0.055091 | 1 |
| Riox2 | 0.016303 | 0.025775 | 1 |
| Cyb5d2 | 0.016292 | 0.686831 | 1 |
| Tpm1 | 0.016292 | 0.102287 | 1 |
| Pdha1 | 0.016289 | 0.080372 | 1 |
| Arf1 | 0.016286 | 0.030202 | 1 |
| Arid3b | 0.016279 | 0.260183 | 1 |

| | | | |
|---|---|---|---|
| Mfsd1 | 0.016277 | 0.317519 | 1 |
| Ccdc152 | 0.016263 | 0.231768 | 1 |
| Rnase6 | 0.016251 | 0.042623 | 1 |
| Stoml2 | 0.016244 | 0.15719 | 1 |
| Sdc4 | 0.016236 | 0.56043 | 1 |
| Yipf4 | 0.01621 | 0.238631 | 1 |
| Gadd45gip1 | 0.016201 | 0.809391 | 1 |
| Adssl1 | 0.016191 | 0.26193 | 1 |
| Pkp4 | 0.016151 | 0.588117 | 1 |
| Samm50 | 0.01615 | 0.53944 | 1 |
| Gm26782 | 0.016147 | 0.29422 | 1 |
| Mak16 | 0.016131 | 0.218178 | 1 |
| Depdc7 | 0.01612 | 0.137827 | 1 |
| Cct5 | 0.016112 | 0.076643 | 1 |
| Cdc23 | 0.016109 | 0.1984 | 1 |
| Anp32a | 0.016106 | 0.030398 | 1 |
| Nr3c1 | 0.016092 | 0.859565 | 1 |
| Pla2g12a | 0.016092 | 0.940039 | 1 |
| Sar1a | 0.016082 | 0.318272 | 1 |
| Meaf6 | 0.016057 | 0.451101 | 1 |
| Mapk3 | 0.016051 | 0.375351 | 1 |
| Cep57 | 0.016019 | 0.020904 | 1 |
| Frg1 | 0.016017 | 0.125219 | 1 |
| Nol8 | 0.016008 | 0.486946 | 1 |
| Klc4 | 0.016007 | 0.159689 | 1 |
| Klhl12 | 0.016007 | 0.373013 | 1 |
| Naa10 | 0.016001 | 0.151518 | 1 |
| Poglut2 | 0.015992 | 0.267473 | 1 |
| Idh3g | 0.01595 | 0.221516 | 1 |
| Fam98b | 0.01594 | 0.79639 | 1 |
| 2310033P09Rik | 0.015935 | 0.29248 | 1 |
| Eif3j1 | 0.015934 | 0.23103 | 1 |
| Lats2 | 0.015912 | 0.342171 | 1 |
| Aplp2 | 0.015904 | 0.896993 | 1 |
| Rce1 | 0.015901 | 0.110095 | 1 |
| Dynll2 | 0.0159 | 0.23465 | 1 |
| Swi5 | 0.015892 | 0.092157 | 1 |
| Tor1a | 0.015872 | 0.147916 | 1 |
| Tbc1d8b | 0.015857 | 0.243567 | 1 |
| Pdzd11 | 0.015829 | 0.190487 | 1 |
| Tmem97 | 0.015811 | 0.1376 | 1 |
| Prdx1 | 0.0158 | 0.123975 | 1 |
| A430005L14Rik | 0.015742 | 0.122884 | 1 |
| 6030468B19Rik | 0.015699 | 0.218104 | 1 |
| Zfp68 | 0.015698 | 0.430049 | 1 |
| Pomp | 0.015685 | 0.038436 | 1 |
| Cops4 | 0.015669 | 0.897625 | 1 |
| Cenpc1 | 0.015643 | 0.630039 | 1 |

| | | | |
|---|---|---|---|
| Purb | 0.01564 | 0.079491 | 1 |
| Fcgrt | 0.015635 | 0.165956 | 1 |
| Fdxr | 0.015633 | 0.164569 | 1 |
| Tmem268 | 0.015609 | 0.151907 | 1 |
| Denr | 0.015591 | 0.326419 | 1 |
| Ddb1 | 0.015575 | 0.862467 | 1 |
| Snx19 | 0.01555 | 0.878759 | 1 |
| Htatip2 | 0.01555 | 0.190728 | 1 |
| Pxdc1 | 0.01554 | 0.711227 | 1 |
| Timm23 | 0.015539 | 0.478582 | 1 |
| Zcrb1 | 0.015537 | 0.419013 | 1 |
| Tmem185a | 0.015528 | 0.0177 | 1 |
| Zc2hc1a | 0.01552 | 0.731158 | 1 |
| Dnajb4 | 0.015505 | 0.147737 | 1 |
| Dimt1 | 0.015501 | 0.319933 | 1 |
| Cct6a | 0.015497 | 0.257113 | 1 |
| Arhgap21 | 0.015461 | 0.750836 | 1 |
| Atpsckmt | 0.015451 | 0.778809 | 1 |
| Pkib | 0.015445 | 0.146637 | 1 |
| Cfap410 | 0.015443 | 0.151736 | 1 |
| Sept11 | 0.015428 | 0.468578 | 1 |
| Vps36 | 0.015412 | 0.23467 | 1 |
| Coq9 | 0.015393 | 0.965217 | 1 |
| Maf1 | 0.015388 | 0.509021 | 1 |
| Ufc1 | 0.015376 | 0.036622 | 1 |
| Ncbp1 | 0.015375 | 0.318673 | 1 |
| Cyp4v3 | 0.015354 | 0.103406 | 1 |
| Smad5 | 0.015354 | 0.694085 | 1 |
| Ccdc85b | 0.015341 | 0.268716 | 1 |
| Stk35 | 0.01534 | 0.541681 | 1 |
| Yipf3 | 0.015339 | 0.191179 | 1 |
| 1700021F05Rik | 0.015331 | 0.271752 | 1 |
| Zfyve16 | 0.015306 | 0.784273 | 1 |
| Higd1a | 0.015288 | 0.217501 | 1 |
| Caprin1 | 0.015281 | 0.067839 | 1 |
| Fer | 0.015267 | 0.700236 | 1 |
| Tubb2b | 0.015229 | 0.00569 | 1 |
| Scamp5 | 0.015215 | 0.804527 | 1 |
| Gm33699 | 0.015209 | 0.112518 | 1 |
| Papola | 0.015176 | 0.578646 | 1 |
| Npm3 | 0.015168 | 0.078378 | 1 |
| Nr2c2ap | 0.015153 | 0.064787 | 1 |
| Dync1li1 | 0.015151 | 0.727718 | 1 |
| Use1 | 0.01515 | 0.091496 | 1 |
| Fam3c | 0.015135 | 0.152346 | 1 |
| Twistnb | 0.015134 | 0.226893 | 1 |
| Usp49 | 0.01513 | 0.853733 | 1 |
| Itgav | 0.015123 | 0.397142 | 1 |

| | | | |
|---|---|---|---|
| Cdk2ap2 | 0.015078 | 0.03585 | 1 |
| Ly96 | 0.015072 | 0.198479 | 1 |
| Cactin | 0.015072 | 0.115946 | 1 |
| Zfp69 | 0.015041 | 0.216591 | 1 |
| Med12 | 0.015032 | 0.337699 | 1 |
| Man1a | 0.015015 | 0.522163 | 1 |
| Uqcc3 | 0.015012 | 0.214621 | 1 |
| Ralb | 0.014957 | 0.392381 | 1 |
| Edc3 | 0.014951 | 0.674314 | 1 |
| Mif4gd | 0.01495 | 0.27516 | 1 |
| Atg2b | 0.014921 | 0.311732 | 1 |
| Fundc1 | 0.014917 | 0.225139 | 1 |
| Kynu | 0.014915 | 0.522841 | 1 |
| Tmem250-ps | 0.014891 | 0.121234 | 1 |
| Sptssa | 0.014883 | 0.114786 | 1 |
| Ankrd46 | 0.01487 | 0.14287 | 1 |
| Map3k4 | 0.014869 | 0.342208 | 1 |
| Commd4 | 0.014852 | 0.085449 | 1 |
| Fam8a1 | 0.014804 | 0.591804 | 1 |
| Gcnt2 | 0.014802 | 0.150046 | 1 |
| Clcn3 | 0.014794 | 0.163626 | 1 |
| Ctcf | 0.014793 | 0.3353 | 1 |
| Rangap1 | 0.014782 | 0.460557 | 1 |
| Bcor | 0.014774 | 0.522741 | 1 |
| Sin3b | 0.014733 | 0.084669 | 1 |
| Timm10b | 0.014714 | 0.166301 | 1 |
| Eef2kmt | 0.014713 | 0.881477 | 1 |
| Snf8 | 0.014704 | 0.936391 | 1 |
| Mfap1b | 0.014688 | 0.361419 | 1 |
| BC017158 | 0.014671 | 0.381903 | 1 |
| Ubxn2a | 0.014669 | 0.60699 | 1 |
| Selenon | 0.014663 | 0.290326 | 1 |
| Mrps27 | 0.014662 | 0.801817 | 1 |
| Milr1 | 0.014613 | 0.494945 | 1 |
| Ndufs6 | 0.014594 | 0.164933 | 1 |
| Ythdc1 | 0.014574 | 0.887463 | 1 |
| Gltp | 0.014572 | 0.254065 | 1 |
| Capn10 | 0.014571 | 0.268432 | 1 |
| Kcnj2 | 0.014548 | 0.153357 | 1 |
| Smpd5 | 0.014514 | 0.697544 | 1 |
| Washc5 | 0.014496 | 0.061865 | 1 |
| Elp2 | 0.014486 | 0.114486 | 1 |
| B4galt7 | 0.014486 | 0.417021 | 1 |
| Ints13 | 0.014478 | 0.117282 | 1 |
| Apc | 0.014476 | 0.160333 | 1 |
| Pigk | 0.014474 | 0.917148 | 1 |
| 2610035D17Rik | 0.014469 | 0.450393 | 1 |
| Pnkp | 0.014469 | 0.687361 | 1 |

| | | | |
|---|---|---|---|
| Ncoa4 | 0.014468 | 0.817049 | 1 |
| Rtn4 | 0.014455 | 0.098755 | 1 |
| Gm41611 | 0.014442 | 0.116791 | 1 |
| Cd33 | 0.014441 | 0.394152 | 1 |
| Magoh | 0.01443 | 0.631606 | 1 |
| Guk1 | 0.014414 | 0.352218 | 1 |
| Sh3glb2 | 0.014405 | 0.497527 | 1 |
| Slc39a8 | 0.014403 | 0.357593 | 1 |
| Casp1 | 0.014369 | 0.859324 | 1 |
| Gabarapl2 | 0.01436 | 0.158617 | 1 |
| Zdhhc13 | 0.014317 | 0.143308 | 1 |
| Elovl5 | 0.014311 | 0.898592 | 1 |
| Bcl7a | 0.014299 | 0.528417 | 1 |
| Arhgap23 | 0.014297 | 0.348905 | 1 |
| Trmu | 0.014269 | 0.542133 | 1 |
| Rusc2 | 0.014266 | 0.394901 | 1 |
| Oaf | 0.014256 | 0.067771 | 1 |
| Atf1 | 0.014248 | 0.087568 | 1 |
| Hikeshi | 0.014235 | 0.757532 | 1 |
| Osbpl2 | 0.014235 | 0.171775 | 1 |
| Actn4 | 0.014211 | 0.736532 | 1 |
| Pomt1 | 0.014198 | 0.42666 | 1 |
| Zfp275 | 0.014191 | 0.09488 | 1 |
| Gba2 | 0.014175 | 0.862797 | 1 |
| Pomgnt1 | 0.014174 | 0.790688 | 1 |
| Mrpl39 | 0.014171 | 0.17539 | 1 |
| Fam89b | 0.014171 | 0.96879 | 1 |
| Skp1a | 0.014169 | 0.358843 | 1 |
| Vapa | 0.014167 | 0.391139 | 1 |
| Smc1a | 0.014149 | 0.215064 | 1 |
| Ube2d3 | 0.014148 | 0.076356 | 1 |
| Dhx34 | 0.014144 | 0.272961 | 1 |
| Zfp944 | 0.014144 | 0.708111 | 1 |
| Dcun1d2 | 0.01414 | 0.63075 | 1 |
| Birc6 | 0.014128 | 0.644297 | 1 |
| Bmt2 | 0.01411 | 0.89084 | 1 |
| Tsr1 | 0.014103 | 0.981354 | 1 |
| Dstn | 0.0141 | 0.787267 | 1 |
| U2af1 | 0.014079 | 0.48313 | 1 |
| Tagln2 | 0.014078 | 0.038622 | 1 |
| Mthfsd | 0.014064 | 0.305278 | 1 |
| Osgin2 | 0.014062 | 0.045755 | 1 |
| Ssrp1 | 0.014049 | 0.23505 | 1 |
| Cep170 | 0.014026 | 0.247452 | 1 |
| Ndel1 | 0.014015 | 0.916073 | 1 |
| Pcyt2 | 0.014013 | 0.425699 | 1 |
| Tmbim1 | 0.014009 | 0.345556 | 1 |
| Slc35d2 | 0.014003 | 0.18516 | 1 |

| | | | |
|---|---|---|---|
| Fh1 | 0.013987 | 0.143283 | 1 |
| Lbh | 0.013956 | 0.232686 | 1 |
| Fam149b | 0.013925 | 0.79851 | 1 |
| Cct7 | 0.013917 | 0.272764 | 1 |
| Rb1cc1 | 0.013901 | 0.806539 | 1 |
| Adcy3 | 0.013865 | 0.680697 | 1 |
| Lsm3 | 0.013864 | 0.096726 | 1 |
| Srm | 0.013862 | 0.29397 | 1 |
| Polr2d | 0.01386 | 0.098389 | 1 |
| Btf3l4 | 0.013851 | 0.297253 | 1 |
| Ppp2r3d | 0.013851 | 0.875061 | 1 |
| Hnrnpll | 0.013841 | 0.183456 | 1 |
| Slc30a4 | 0.01384 | 0.378209 | 1 |
| Rexo4 | 0.01384 | 0.945403 | 1 |
| Bzw1 | 0.013818 | 0.011483 | 1 |
| 2210016L21Rik | 0.013814 | 0.618645 | 1 |
| Slc2a5 | 0.013771 | 0.323345 | 1 |
| Scai | 0.013762 | 0.450608 | 1 |
| Nfix | 0.013756 | 0.411055 | 1 |
| Laptm4b | 0.013747 | 0.291472 | 1 |
| Pura | 0.013736 | 0.168735 | 1 |
| Psmd7 | 0.01369 | 0.218566 | 1 |
| 1810058I24Rik | 0.013688 | 0.21453 | 1 |
| Inpp1 | 0.013669 | 0.986033 | 1 |
| Bmi1 | 0.013658 | 0.654854 | 1 |
| Dpy19l3 | 0.013646 | 0.402305 | 1 |
| Nek6 | 0.013641 | 0.194361 | 1 |
| Dffa | 0.013626 | 0.12333 | 1 |
| Zfpm1 | 0.013617 | 0.288519 | 1 |
| Ric1 | 0.013617 | 0.985785 | 1 |
| Hyou1 | 0.013608 | 0.667654 | 1 |
| Poldip2 | 0.013593 | 0.785636 | 1 |
| Cnot9 | 0.013582 | 0.401129 | 1 |
| Slx1b | 0.013565 | 0.245278 | 1 |
| Plgrkt | 0.013552 | 0.483467 | 1 |
| Jpt1 | 0.013543 | 0.155446 | 1 |
| St13 | 0.01354 | 0.098463 | 1 |
| Mrpl23 | 0.013517 | 0.242067 | 1 |
| Stk3 | 0.013509 | 0.817668 | 1 |
| Nelfe | 0.013477 | 0.154275 | 1 |
| Ppp1r7 | 0.013444 | 0.339701 | 1 |
| Cdk5rap3 | 0.01344 | 0.15481 | 1 |
| Gm43464 | 0.013433 | 0.309708 | 1 |
| Arhgap27 | 0.013432 | 0.395639 | 1 |
| Farp1 | 0.013418 | 0.742413 | 1 |
| Cmas | 0.013413 | 0.913646 | 1 |
| Adipor1 | 0.013391 | 0.077097 | 1 |
| Ccdc88a | 0.013373 | 0.116612 | 1 |

| | | | |
|---|---|---|---|
| Tmem237 | 0.013371 | 0.222624 | 1 |
| Kif5b | 0.013361 | 0.518509 | 1 |
| Slc16a3 | 0.013351 | 0.08106 | 1 |
| Acsl1 | 0.013347 | 0.491215 | 1 |
| Smim11 | 0.013342 | 0.141073 | 1 |
| Mcts1 | 0.013338 | 0.22003 | 1 |
| Mlxipl | 0.013337 | 0.179798 | 1 |
| Srsf5 | 0.013327 | 0.829394 | 1 |
| Mrps7 | 0.013311 | 0.466338 | 1 |
| Pafah1b3 | 0.013308 | 0.160698 | 1 |
| Pgrmc1 | 0.013308 | 0.197176 | 1 |
| Yipf5 | 0.013306 | 0.238436 | 1 |
| Nphp1 | 0.013289 | 0.30388 | 1 |
| Gm43466 | 0.013265 | 0.796612 | 1 |
| Ndufb4 | 0.013257 | 0.43715 | 1 |
| Kctd18 | 0.01324 | 0.510566 | 1 |
| Etfb | 0.013227 | 0.079592 | 1 |
| Cysltr1 | 0.013213 | 0.633772 | 1 |
| Isoc2a | 0.013212 | 0.569742 | 1 |
| Zfp961 | 0.013194 | 0.62278 | 1 |
| Zhx3 | 0.013186 | 0.958846 | 1 |
| Mef2a | 0.013176 | 0.659625 | 1 |
| Sh3glb1 | 0.013173 | 0.639512 | 1 |
| Arfgap2 | 0.013165 | 0.046777 | 1 |
| Dnajc2 | 0.013163 | 0.168887 | 1 |
| Tyw5 | 0.013142 | 0.126037 | 1 |
| Mrps36 | 0.013131 | 0.223825 | 1 |
| Sclt1 | 0.01313 | 0.413254 | 1 |
| Cpsf7 | 0.01313 | 0.494262 | 1 |
| Rgs19 | 0.013109 | 0.08734 | 1 |
| Stt3b | 0.013096 | 0.278981 | 1 |
| Etaa1 | 0.01309 | 0.919774 | 1 |
| Atxn10 | 0.013061 | 0.20294 | 1 |
| Naaa | 0.013057 | 0.638582 | 1 |
| Scoc | 0.013052 | 0.331908 | 1 |
| Wdr92 | 0.013038 | 0.976069 | 1 |
| Med9 | 0.013038 | 0.709408 | 1 |
| Wrap73 | 0.013038 | 0.674601 | 1 |
| Mapre2 | 0.013001 | 0.253862 | 1 |
| Psma7 | 0.01298 | 0.102715 | 1 |
| Srebf2 | 0.012956 | 0.00922 | 1 |
| Slc38a9 | 0.012933 | 0.830198 | 1 |
| Aars | 0.012931 | 0.697994 | 1 |
| Rrm2b | 0.012852 | 0.750071 | 1 |
| 4930556J24Rik | 0.012847 | 0.219373 | 1 |
| Myc | 0.012814 | 0.79871 | 1 |
| Leo1 | 0.012786 | 0.098425 | 1 |
| Dcaf10 | 0.01278 | 0.773118 | 1 |

| | | | |
|---|---|---|---|
| Trappc1 | 0.01278 | 0.337872 | 1 |
| Rbmx | 0.012762 | 0.492702 | 1 |
| Atpaf1 | 0.01274 | 0.18169 | 1 |
| Pdcd10 | 0.012738 | 0.272496 | 1 |
| Pigc | 0.012735 | 0.482042 | 1 |
| Snn | 0.012732 | 0.733916 | 1 |
| Trmt6 | 0.012715 | 0.724973 | 1 |
| Yju2 | 0.012703 | 0.241599 | 1 |
| Mcm6 | 0.012698 | 0.262946 | 1 |
| Ercc1 | 0.012674 | 0.204959 | 1 |
| 4931406C07Rik | 0.012672 | 0.72228 | 1 |
| Rab10os | 0.012657 | 0.572662 | 1 |
| Tnfaip3 | 0.012656 | 0.505254 | 1 |
| Tmem106c | 0.012629 | 0.197518 | 1 |
| Gtpbp10 | 0.012629 | 0.868279 | 1 |
| Msn | 0.012607 | 0.300207 | 1 |
| Cox20 | 0.012606 | 0.63087 | 1 |
| Snx12 | 0.012598 | 0.895038 | 1 |
| Utp14a | 0.012594 | 0.570227 | 1 |
| Git2 | 0.012588 | 0.46109 | 1 |
| Gm16541 | 0.012586 | 0.722306 | 1 |
| Dmac2 | 0.012565 | 0.331283 | 1 |
| Nfic | 0.012555 | 0.996987 | 1 |
| Czib | 0.012551 | 0.658303 | 1 |
| Polr2i | 0.012513 | 0.125122 | 1 |
| Arf5 | 0.012487 | 0.63065 | 1 |
| Rab33b | 0.012473 | 0.24815 | 1 |
| Smim19 | 0.012473 | 0.118326 | 1 |
| Smim15 | 0.012453 | 0.757448 | 1 |
| Arf6 | 0.012447 | 0.593058 | 1 |
| Psmb10 | 0.012445 | 0.117447 | 1 |
| Cdk10 | 0.012445 | 0.551736 | 1 |
| Rpap1 | 0.012365 | 0.873732 | 1 |
| Maip1 | 0.01236 | 0.616791 | 1 |
| Arap2 | 0.012355 | 0.244085 | 1 |
| Psmb1 | 0.012352 | 0.204812 | 1 |
| Ssbp4 | 0.01235 | 0.245452 | 1 |
| Mlh1 | 0.01235 | 0.567858 | 1 |
| AW011738 | 0.012343 | 0.818559 | 1 |
| Gtpbp3 | 0.012341 | 0.856041 | 1 |
| Dhrs7b | 0.01234 | 0.964351 | 1 |
| C1d | 0.012336 | 0.540293 | 1 |
| Rab2a | 0.012335 | 0.47775 | 1 |
| Esd | 0.012329 | 0.419105 | 1 |
| Elp5 | 0.012309 | 0.348892 | 1 |
| Tial1 | 0.012287 | 0.423615 | 1 |
| Pnpla2 | 0.012284 | 0.404106 | 1 |
| Nfx1 | 0.012282 | 0.815964 | 1 |

| | | | |
|---|---|---|---|
| Ccdc51 | 0.01226 | 0.636553 | 1 |
| 2610001J05Rik | 0.012255 | 0.200763 | 1 |
| Herc2 | 0.012255 | 0.698671 | 1 |
| Gtf3c4 | 0.012252 | 0.139398 | 1 |
| Mrpl37 | 0.012248 | 0.586096 | 1 |
| Alg1 | 0.012244 | 0.915764 | 1 |
| Dolpp1 | 0.012235 | 0.119126 | 1 |
| Dbi | 0.012235 | 0.044578 | 1 |
| Slc7a1 | 0.012227 | 0.126588 | 1 |
| Gins1 | 0.012224 | 0.202553 | 1 |
| Hsbp1 | 0.012205 | 0.310238 | 1 |
| Zscan25 | 0.012191 | 0.099595 | 1 |
| Trpv4 | 0.012173 | 0.201238 | 1 |
| Psme2 | 0.012161 | 0.392994 | 1 |
| Ormdl1 | 0.012155 | 0.017693 | 1 |
| Ube2e3 | 0.012152 | 0.738816 | 1 |
| Bet1l | 0.012138 | 0.325114 | 1 |
| B230307C23Rik | 0.012128 | 0.427384 | 1 |
| Entr1 | 0.012114 | 0.702028 | 1 |
| Rgl3 | 0.012107 | 0.167353 | 1 |
| Chm | 0.012103 | 0.517321 | 1 |
| Batf3 | 0.012087 | 0.720621 | 1 |
| Gm39556 | 0.012086 | 0.661367 | 1 |
| Abhd6 | 0.012081 | 0.174496 | 1 |
| Casp3 | 0.012056 | 0.963082 | 1 |
| Mrpl20 | 0.012045 | 0.064795 | 1 |
| Hspbp1 | 0.012043 | 0.727109 | 1 |
| Sema4a | 0.012026 | 0.564098 | 1 |
| Inka1 | 0.012023 | 0.108647 | 1 |
| Lsm8 | 0.012002 | 0.281782 | 1 |
| Pdcl3 | 0.011991 | 0.197522 | 1 |
| Dnajc21 | 0.011977 | 0.234044 | 1 |
| Mtf2 | 0.011949 | 0.29523 | 1 |
| Rab24 | 0.011944 | 0.76883 | 1 |
| Sun2 | 0.011931 | 0.379804 | 1 |
| Cln8 | 0.011927 | 0.635448 | 1 |
| Fem1b | 0.011919 | 0.696889 | 1 |
| Tiparp | 0.011915 | 0.723506 | 1 |
| Haus6 | 0.011892 | 0.157114 | 1 |
| Pdcd6ip | 0.011885 | 0.318302 | 1 |
| Acvr2a | 0.011869 | 0.688707 | 1 |
| Axin1 | 0.011825 | 0.538375 | 1 |
| Xylt2 | 0.011812 | 0.32971 | 1 |
| Atmin | 0.01181 | 0.276019 | 1 |
| Scaper | 0.011805 | 0.514284 | 1 |
| Slc35a1 | 0.011801 | 0.175873 | 1 |
| Erlin2 | 0.011795 | 0.161671 | 1 |
| Rnf219 | 0.011787 | 0.681994 | 1 |

| | | | |
|---|---|---|---|
| Zfp641 | 0.011787 | 0.529917 | 1 |
| Uba2 | 0.011772 | 0.268325 | 1 |
| Cox17 | 0.011769 | 0.053287 | 1 |
| Zfp955a | 0.011764 | 0.704169 | 1 |
| Gm30329 | 0.011743 | 0.014777 | 1 |
| Ddx52 | 0.01174 | 0.35113 | 1 |
| Mrps14 | 0.011735 | 0.313841 | 1 |
| Mmgt2 | 0.011728 | 0.42395 | 1 |
| Bola2 | 0.011709 | 0.062515 | 1 |
| Vegfb | 0.011695 | 0.217284 | 1 |
| Elp3 | 0.011684 | 0.365869 | 1 |
| Zdhhc9 | 0.01168 | 0.295606 | 1 |
| Zfp593 | 0.011677 | 0.126413 | 1 |
| Rnpc3 | 0.011663 | 0.500527 | 1 |
| Cdca7 | 0.011654 | 0.266044 | 1 |
| Zfp52 | 0.011636 | 0.742837 | 1 |
| Itgb2 | 0.011623 | 0.318702 | 1 |
| Zfp993 | 0.011619 | 0.190142 | 1 |
| Rnh1 | 0.011619 | 0.565036 | 1 |
| Nsd2 | 0.011557 | 0.352117 | 1 |
| Ephx1 | 0.011555 | 0.651518 | 1 |
| Tsc22d2 | 0.011553 | 0.404909 | 1 |
| Ube2n | 0.01154 | 0.14878 | 1 |
| Sema4c | 0.011525 | 0.903262 | 1 |
| Mrpl55 | 0.011519 | 0.413866 | 1 |
| Dvl3 | 0.011518 | 0.334679 | 1 |
| Srsf10 | 0.011517 | 0.507083 | 1 |
| Ergic2 | 0.011511 | 0.151482 | 1 |
| Wdsub1 | 0.01151 | 0.977146 | 1 |
| Tradd | 0.011509 | 0.77239 | 1 |
| Pdlim5 | 0.011497 | 0.439461 | 1 |
| Hnrnph1 | 0.011438 | 0.116468 | 1 |
| Hint1 | 0.011437 | 0.048723 | 1 |
| Rab18 | 0.011431 | 0.116767 | 1 |
| Suv39h1 | 0.01142 | 0.503804 | 1 |
| Arpc5l | 0.011407 | 0.122006 | 1 |
| Tarbp2 | 0.011399 | 0.539911 | 1 |
| P2ry6 | 0.011397 | 0.288584 | 1 |
| U2af2 | 0.011392 | 0.646493 | 1 |
| Poglut3 | 0.011383 | 0.544051 | 1 |
| Smap1 | 0.011376 | 0.722844 | 1 |
| Alg9 | 0.011348 | 0.172717 | 1 |
| Atrip | 0.011332 | 0.411792 | 1 |
| Mafk | 0.011322 | 0.204602 | 1 |
| Tnfaip8 | 0.011313 | 0.590725 | 1 |
| Sbds | 0.011298 | 0.682545 | 1 |
| Trpm7 | 0.011284 | 0.071097 | 1 |
| Cnp | 0.011262 | 0.374127 | 1 |

| | | | |
|---|---|---|---|
| Rai1 | 0.011254 | 0.349404 | 1 |
| Inpp5f | 0.011226 | 0.761468 | 1 |
| Clcc1 | 0.011215 | 0.094856 | 1 |
| C9orf72 | 0.011205 | 0.278773 | 1 |
| Cript | 0.011201 | 0.926221 | 1 |
| Cstf2t | 0.011193 | 0.167616 | 1 |
| Dhrs1 | 0.011187 | 0.685317 | 1 |
| Psmd12 | 0.011187 | 0.235538 | 1 |
| Cd82 | 0.011177 | 0.192361 | 1 |
| Gpatch3 | 0.011176 | 0.351419 | 1 |
| Ccdc59 | 0.011175 | 0.032976 | 1 |
| Enpp1 | 0.011168 | 0.700175 | 1 |
| Mettl9 | 0.011141 | 0.333478 | 1 |
| Afmid | 0.011127 | 0.428878 | 1 |
| Ly6m | 0.011121 | 0.482932 | 1 |
| Ciapin1 | 0.011094 | 0.590461 | 1 |
| Chchd7 | 0.011093 | 0.107071 | 1 |
| Rft1 | 0.011091 | 0.730412 | 1 |
| 1810030O07Rik | 0.011074 | 0.57212 | 1 |
| Heca | 0.011061 | 0.246843 | 1 |
| Sec13 | 0.011059 | 0.237468 | 1 |
| Snhg5 | 0.011056 | 0.670746 | 1 |
| Hook3 | 0.011052 | 0.823525 | 1 |
| Cybb | 0.011035 | 0.095724 | 1 |
| Spns1 | 0.011033 | 0.136573 | 1 |
| Polr3d | 0.011022 | 0.384617 | 1 |
| Nit1 | 0.010996 | 0.243739 | 1 |
| 2300009A05Rik | 0.010986 | 0.115839 | 1 |
| 2610021A01Rik | 0.010966 | 0.686874 | 1 |
| Bptf | 0.010963 | 0.586753 | 1 |
| Sdhc | 0.01096 | 0.305234 | 1 |
| Hagh | 0.010958 | 0.903459 | 1 |
| Vbp1 | 0.010954 | 0.29742 | 1 |
| Itgb1bp1 | 0.010911 | 0.146222 | 1 |
| Zfp868 | 0.010908 | 0.519317 | 1 |
| Ttyh3 | 0.010907 | 0.203126 | 1 |
| 1810037I17Rik | 0.010894 | 0.037984 | 1 |
| Xpnpep3 | 0.010881 | 0.837873 | 1 |
| N4bp2 | 0.010878 | 0.461917 | 1 |
| Ghdc | 0.010877 | 0.603129 | 1 |
| Alkbh5 | 0.010863 | 0.993029 | 1 |
| Tmem38b | 0.010854 | 0.44024 | 1 |
| Unc50 | 0.01085 | 0.082877 | 1 |
| Ttc4 | 0.010848 | 0.339457 | 1 |
| Mfsd8 | 0.010834 | 0.49569 | 1 |
| Ptrhd1 | 0.010834 | 0.592612 | 1 |
| Scnm1 | 0.010825 | 0.319273 | 1 |
| Dynlt3 | 0.010822 | 0.125199 | 1 |

| | | | |
|---|---|---|---|
| Arvcf | 0.01081 | 0.822662 | 1 |
| Ormdl2 | 0.01081 | 0.028505 | 1 |
| Oat | 0.010804 | 0.31337 | 1 |
| Arhgap29 | 0.010797 | 0.792 | 1 |
| 2310009B15Rik | 0.010793 | 0.414875 | 1 |
| Mrpl11 | 0.010791 | 0.120476 | 1 |
| Jrkl | 0.010785 | 0.562062 | 1 |
| Cdk4 | 0.010781 | 0.798217 | 1 |
| Traf2 | 0.010767 | 0.7892 | 1 |
| Prpf31 | 0.010762 | 0.696883 | 1 |
| Hspa8 | 0.010755 | 0.430442 | 1 |
| Phpt1 | 0.010748 | 0.039892 | 1 |
| Csgalnact2 | 0.01074 | 0.697194 | 1 |
| Man2c1 | 0.010736 | 0.804628 | 1 |
| Polr1d | 0.010724 | 0.359502 | 1 |
| Arpp19 | 0.010713 | 0.103067 | 1 |
| Ghitm | 0.01068 | 0.159349 | 1 |
| Ube2m | 0.010671 | 0.19589 | 1 |
| Gm15964 | 0.010669 | 0.300924 | 1 |
| Dohh | 0.010662 | 0.432863 | 1 |
| Zfp770 | 0.010636 | 0.13752 | 1 |
| Dkc1 | 0.010633 | 0.785308 | 1 |
| Eif2b2 | 0.01062 | 0.425261 | 1 |
| Rps19bp1 | 0.010615 | 0.156153 | 1 |
| Map1lc3a | 0.010591 | 0.657947 | 1 |
| Casc4 | 0.010583 | 0.320771 | 1 |
| Psme3 | 0.010578 | 0.086462 | 1 |
| Mcur1 | 0.010565 | 0.634869 | 1 |
| Pfdn6 | 0.010563 | 0.587219 | 1 |
| Zscan22 | 0.010559 | 0.712683 | 1 |
| Ndufa3 | 0.010533 | 0.21732 | 1 |
| Ctnnb1 | 0.010524 | 0.06152 | 1 |
| Nap1l4 | 0.010521 | 0.903743 | 1 |
| Lrig2 | 0.010514 | 0.904171 | 1 |
| Psme1 | 0.010501 | 0.3248 | 1 |
| Rhbdd2 | 0.010496 | 0.081155 | 1 |
| Kri1 | 0.010495 | 0.493692 | 1 |
| Zfp414 | 0.01049 | 0.615246 | 1 |
| Mllt1 | 0.010489 | 0.874654 | 1 |
| Tex264 | 0.010485 | 0.260854 | 1 |
| Alg3 | 0.010478 | 0.53462 | 1 |
| Rabggtb | 0.010465 | 0.746896 | 1 |
| Cdc5l | 0.010464 | 0.796369 | 1 |
| Tefm | 0.010459 | 0.549195 | 1 |
| Gdpd5 | 0.010458 | 0.580129 | 1 |
| Zfas1 | 0.010449 | 0.150802 | 1 |
| Fam122a | 0.010438 | 0.162372 | 1 |
| 6430590A07Rik | 0.010408 | 0.166473 | 1 |

| | | | |
|---|---|---|---|
| Mrgbp | 0.0104 | 0.958104 | 1 |
| Ilvbl | 0.010396 | 0.315036 | 1 |
| Smug1 | 0.010392 | 0.676547 | 1 |
| Endod1 | 0.010391 | 0.25737 | 1 |
| Cox11 | 0.010368 | 0.348794 | 1 |
| Sec24d | 0.010366 | 0.744235 | 1 |
| Uqcc2 | 0.010353 | 0.427136 | 1 |
| Vps35 | 0.01035 | 0.46644 | 1 |
| Nudt5 | 0.010322 | 0.309133 | 1 |
| Adgra3 | 0.01032 | 0.043835 | 1 |
| Olfml2b | 0.010316 | 0.410169 | 1 |
| Pigq | 0.010295 | 0.103088 | 1 |
| Oxld1 | 0.010293 | 0.327179 | 1 |
| Pfn1 | 0.010271 | 0.297852 | 1 |
| Ammecr1l | 0.01026 | 0.965291 | 1 |
| Esco1 | 0.010244 | 0.229792 | 1 |
| 6230400D17Rik | 0.010236 | 0.221945 | 1 |
| Lzic | 0.010222 | 0.890142 | 1 |
| Csnk2a1 | 0.010219 | 0.377189 | 1 |
| Parp6 | 0.010208 | 0.953549 | 1 |
| Cyp2r1 | 0.010193 | 0.043269 | 1 |
| Umps | 0.010181 | 0.595717 | 1 |
| Mboat7 | 0.010167 | 0.356057 | 1 |
| Dele1 | 0.010167 | 0.851246 | 1 |
| Trabd | 0.010156 | 0.256735 | 1 |
| Rundc1 | 0.010148 | 0.634722 | 1 |
| Pbx1 | 0.010119 | 0.797296 | 1 |
| Pigx | 0.010084 | 0.692564 | 1 |
| Mrps10 | 0.010079 | 0.167837 | 1 |
| Ndufb6 | 0.01006 | 0.277016 | 1 |
| Gm43462 | 0.010057 | 0.035785 | 1 |
| Trmt12 | 0.010044 | 0.708207 | 1 |
| Gm17066 | 0.010037 | 0.446361 | 1 |
| Trappc2 | 0.010032 | 0.200657 | 1 |
| Abhd10 | 0.010018 | 0.850824 | 1 |
| Chst1 | 0.010009 | 0.286123 | 1 |
| Prrc2b | -0.01 | 0.740367 | 1 |
| Acyp1 | -0.01001 | 0.252092 | 1 |
| Cracr2b | -0.01001 | 0.583314 | 1 |
| Pts | -0.01001 | 0.942703 | 1 |
| Ttc38 | -0.01001 | 0.596399 | 1 |
| Nkapd1 | -0.01003 | 0.34888 | 1 |
| Cklf | -0.01003 | 0.655762 | 1 |
| Zswim3 | -0.01003 | 0.160172 | 1 |
| Znhit6 | -0.01003 | 0.21932 | 1 |
| Scyl2 | -0.01004 | 0.782934 | 1 |
| Pafah1b2 | -0.01004 | 0.826649 | 1 |
| Lyar | -0.01006 | 0.741915 | 1 |

| | | | |
|---|---|---|---|
| Rnf17 | -0.01006 | 0.957827 | 1 |
| BC005561 | -0.01006 | 0.277931 | 1 |
| Prdm9 | -0.01007 | 0.19886 | 1 |
| Azi2 | -0.01009 | 0.944237 | 1 |
| Klhl24 | -0.0101 | 0.697379 | 1 |
| Rdh5 | -0.0101 | 0.437845 | 1 |
| Rdh11 | -0.01011 | 0.507284 | 1 |
| Exosc7 | -0.01012 | 0.459489 | 1 |
| Gcsh | -0.01016 | 0.950244 | 1 |
| Arglu1 | -0.01016 | 0.79113 | 1 |
| Mtmr9 | -0.01017 | 0.350588 | 1 |
| Dnase1l1 | -0.01018 | 0.288878 | 1 |
| Akr1e1 | -0.0102 | 0.9266 | 1 |
| Setd4 | -0.01021 | 0.319539 | 1 |
| Mtor | -0.01024 | 0.674742 | 1 |
| Lin52 | -0.01025 | 0.679896 | 1 |
| Trim3 | -0.01026 | 0.164368 | 1 |
| Mcph1 | -0.01027 | 0.4496 | 1 |
| Gtf3c1 | -0.01027 | 0.887041 | 1 |
| Gm32591 | -0.01028 | 0.461853 | 1 |
| Hnrnpr | -0.01028 | 0.975805 | 1 |
| Rbfa | -0.01028 | 0.788909 | 1 |
| Tirap | -0.01028 | 0.491925 | 1 |
| Rbm41 | -0.01029 | 0.28686 | 1 |
| Insig2 | -0.01033 | 0.86037 | 1 |
| Abhd14a | -0.01033 | 0.346371 | 1 |
| Nde1 | -0.01033 | 0.897234 | 1 |
| Usp8 | -0.01035 | 0.505663 | 1 |
| Ints8 | -0.01035 | 0.432768 | 1 |
| Tcp11l2 | -0.01036 | 0.595516 | 1 |
| Gemin2 | -0.01037 | 0.228291 | 1 |
| Bud23 | -0.01037 | 0.362855 | 1 |
| Mrpl24 | -0.01038 | 0.960668 | 1 |
| 4933412E12Rik | -0.0104 | 0.878171 | 1 |
| Btbd8 | -0.0104 | 0.529926 | 1 |
| Lgals9 | -0.01041 | 0.992134 | 1 |
| Usf1 | -0.01044 | 0.740549 | 1 |
| Zdhhc8 | -0.01044 | 0.360426 | 1 |
| Oaz2 | -0.01046 | 0.348044 | 1 |
| Sdhaf1 | -0.01046 | 0.913227 | 1 |
| AU022252 | -0.01047 | 0.757012 | 1 |
| Hpf1 | -0.01048 | 0.315766 | 1 |
| Usp47 | -0.01049 | 0.331098 | 1 |
| Snrpd2 | -0.01049 | 0.60669 | 1 |
| Faap100 | -0.01049 | 0.267079 | 1 |
| 1600020E01Rik | -0.01049 | 0.505737 | 1 |
| Pknox1 | -0.0105 | 0.443464 | 1 |
| Mypopos | -0.0105 | 0.764359 | 1 |

| | | | |
|---|---|---|---|
| Taf6 | -0.0105 | 0.936174 | 1 |
| Mphosph9 | -0.01051 | 0.586262 | 1 |
| Jarid2 | -0.01052 | 0.301883 | 1 |
| Sp140 | -0.01052 | 0.658965 | 1 |
| A630001O12Rik | -0.01053 | 0.23986 | 1 |
| Borcs5 | -0.01054 | 0.150137 | 1 |
| Ppp2ca | -0.01054 | 0.805001 | 1 |
| Gclc | -0.01054 | 0.569407 | 1 |
| Kansl2 | -0.01054 | 0.528795 | 1 |
| Kctd3 | -0.01055 | 0.643127 | 1 |
| Smarca2 | -0.01055 | 0.645955 | 1 |
| Cmya5 | -0.01056 | 0.109645 | 1 |
| Rad21 | -0.01056 | 0.89996 | 1 |
| Ppp1r8 | -0.01056 | 0.723202 | 1 |
| Scyl1 | -0.01057 | 0.480221 | 1 |
| Scarb1 | -0.01058 | 0.854982 | 1 |
| Cdk16 | -0.01058 | 0.812533 | 1 |
| Rrs1 | -0.01059 | 0.370477 | 1 |
| Cgas | -0.0106 | 0.140237 | 1 |
| Gm11713 | -0.01064 | 0.148883 | 1 |
| Sod2 | -0.01065 | 0.930863 | 1 |
| Sarnp | -0.01065 | 0.979539 | 1 |
| Nop58 | -0.01066 | 0.741387 | 1 |
| Galnt11 | -0.01067 | 0.588677 | 1 |
| Yme1l1 | -0.01068 | 0.655695 | 1 |
| Zfp260 | -0.01073 | 0.911464 | 1 |
| Cluap1 | -0.01073 | 0.886835 | 1 |
| Rcn1 | -0.01073 | 0.745098 | 1 |
| Flcn | -0.01074 | 0.487804 | 1 |
| Zfp384 | -0.01075 | 0.930447 | 1 |
| Gna13 | -0.01076 | 0.310594 | 1 |
| Ctsc | -0.01077 | 0.527199 | 1 |
| Ythdf2 | -0.01078 | 0.561717 | 1 |
| Akt1s1 | -0.01078 | 0.387667 | 1 |
| Kxd1 | -0.01079 | 0.979268 | 1 |
| Crkl | -0.01079 | 0.341159 | 1 |
| Trappc3 | -0.0108 | 0.390707 | 1 |
| Fdx2 | -0.01081 | 0.962058 | 1 |
| 2810006K23Rik | -0.01081 | 0.556331 | 1 |
| Ppip5k1 | -0.01081 | 0.128219 | 1 |
| Pctp | -0.01081 | 0.568744 | 1 |
| Aldh3a2 | -0.01082 | 0.922796 | 1 |
| Fubp3 | -0.01083 | 0.701098 | 1 |
| Rab9 | -0.01083 | 0.782424 | 1 |
| Neurl4 | -0.01084 | 0.396168 | 1 |
| Abi2 | -0.01086 | 0.627309 | 1 |
| Bckdha | -0.01087 | 0.302437 | 1 |
| Gm16201 | -0.01088 | 0.059938 | 1 |

| | | | |
|---|---|---|---|
| Rap2c | -0.01088 | 0.682606 | 1 |
| Nono | -0.01088 | 0.686616 | 1 |
| Faf2 | -0.01089 | 0.150045 | 1 |
| Rab31 | -0.01089 | 0.862102 | 1 |
| Sqor | -0.0109 | 0.61849 | 1 |
| Rpl10-ps3 | -0.0109 | 0.326868 | 1 |
| Gbgt1 | -0.01094 | 0.487032 | 1 |
| Narf | -0.01095 | 0.601804 | 1 |
| BC002059 | -0.01095 | 0.429148 | 1 |
| Fbl | -0.01095 | 0.980642 | 1 |
| Apeh | -0.01095 | 0.474443 | 1 |
| Ccdc58 | -0.01095 | 0.740083 | 1 |
| Usp12 | -0.01097 | 0.517732 | 1 |
| Szt2 | -0.01097 | 0.348708 | 1 |
| Mettl7a1 | -0.01098 | 0.328567 | 1 |
| Chpf2 | -0.01098 | 0.34179 | 1 |
| Grpel2 | -0.01099 | 0.930842 | 1 |
| H2-DMa | -0.01099 | 0.811661 | 1 |
| Cyb5r1 | -0.01099 | 0.508653 | 1 |
| Dnajc4 | -0.011 | 0.913278 | 1 |
| Xbp1 | -0.01102 | 0.753141 | 1 |
| Smim20 | -0.01104 | 0.985734 | 1 |
| Slx4 | -0.01104 | 0.305282 | 1 |
| Actr8 | -0.01105 | 0.434538 | 1 |
| Hectd3 | -0.01105 | 0.321265 | 1 |
| Ints4 | -0.01109 | 0.990742 | 1 |
| Hspb11 | -0.0111 | 0.619436 | 1 |
| Prkd2 | -0.01111 | 0.492099 | 1 |
| Rcc2 | -0.01111 | 0.65403 | 1 |
| Ccdc61 | -0.01111 | 0.374154 | 1 |
| Zfp90 | -0.01111 | 0.718318 | 1 |
| Gpatch4 | -0.01112 | 0.225877 | 1 |
| Sec31a | -0.01112 | 0.926934 | 1 |
| B4galnt1 | -0.01113 | 0.888287 | 1 |
| Slc25a36 | -0.01113 | 0.779186 | 1 |
| Ppp2r5d | -0.01114 | 0.19459 | 1 |
| Mtus1 | -0.01114 | 0.928435 | 1 |
| Tifab | -0.01116 | 0.55724 | 1 |
| Sipa1 | -0.01119 | 0.705492 | 1 |
| Zdhhc7 | -0.0112 | 0.679182 | 1 |
| Zfp788 | -0.01122 | 0.14934 | 1 |
| Cant1 | -0.01122 | 0.67362 | 1 |
| Abraxas1 | -0.01122 | 0.411103 | 1 |
| Dnajc10 | -0.01123 | 0.970045 | 1 |
| Hibadh | -0.01131 | 0.899945 | 1 |
| Gsk3b | -0.01132 | 0.775227 | 1 |
| Cd46 | -0.01134 | 0.957455 | 1 |
| Ppp1r16a | -0.01134 | 0.288817 | 1 |

| | | | |
|---|---|---|---|
| Eif2d | -0.01134 | 0.452775 | 1 |
| Tmem184c | -0.01135 | 0.196244 | 1 |
| Rnf44 | -0.01135 | 0.835117 | 1 |
| Pitpna | -0.01136 | 0.794475 | 1 |
| 4933434E20Rik | -0.01136 | 0.932232 | 1 |
| Rab5b | -0.01138 | 0.626697 | 1 |
| Gosr1 | -0.01138 | 0.38715 | 1 |
| Slc27a1 | -0.01139 | 0.273137 | 1 |
| Abl2 | -0.01139 | 0.458928 | 1 |
| 2210016F16Rik | -0.0114 | 0.679501 | 1 |
| Ankrd13d | -0.0114 | 0.278104 | 1 |
| Aar2 | -0.0114 | 0.276076 | 1 |
| Siah2 | -0.0114 | 0.406563 | 1 |
| Srp68 | -0.01142 | 0.47576 | 1 |
| Drosha | -0.01142 | 0.586735 | 1 |
| Itprid2 | -0.01143 | 0.729203 | 1 |
| Hps3 | -0.01145 | 0.555271 | 1 |
| Tmem39a | -0.01146 | 0.70355 | 1 |
| Mapk1ip1 | -0.01148 | 0.975982 | 1 |
| Capn3 | -0.01149 | 0.385849 | 1 |
| Impdh2 | -0.01151 | 0.534511 | 1 |
| Rmi1 | -0.01154 | 0.288486 | 1 |
| Klhl26 | -0.01156 | 0.617299 | 1 |
| Prkar2a | -0.01158 | 0.524353 | 1 |
| Lrrc14 | -0.0116 | 0.339951 | 1 |
| Cdadc1 | -0.01162 | 0.589545 | 1 |
| Ascc2 | -0.01162 | 0.999487 | 1 |
| Med25 | -0.01163 | 0.700977 | 1 |
| Ddx6 | -0.01164 | 0.671556 | 1 |
| Gm26672 | -0.01164 | 0.782997 | 1 |
| Dera | -0.01164 | 0.452724 | 1 |
| Klhl36 | -0.01165 | 0.712351 | 1 |
| Plekhn1 | -0.01174 | 0.61213 | 1 |
| Hs6st1 | -0.01174 | 0.3714 | 1 |
| Sel1l | -0.01175 | 0.898739 | 1 |
| Mfsd4b4 | -0.01175 | 0.559033 | 1 |
| Ulbp1 | -0.01175 | 0.09552 | 1 |
| Pigg | -0.01176 | 0.030876 | 1 |
| Tesk2 | -0.01176 | 0.716872 | 1 |
| Hook2 | -0.01179 | 0.928078 | 1 |
| Bysl | -0.01179 | 0.47357 | 1 |
| Esrra | -0.01181 | 0.303193 | 1 |
| Coasy | -0.01181 | 0.928359 | 1 |
| Lsm10 | -0.01182 | 0.468199 | 1 |
| Zfp607a | -0.01182 | 0.618673 | 1 |
| Rasal3 | -0.01183 | 0.390171 | 1 |
| Tfrc | -0.01183 | 0.584133 | 1 |
| Tmem39b | -0.01185 | 0.539372 | 1 |

| | | | |
|---|---|---|---|
| Hmg20b | -0.01186 | 0.863117 | 1 |
| Triap1 | -0.01187 | 0.363501 | 1 |
| Hes6 | -0.01189 | 0.80104 | 1 |
| Supt5 | -0.01189 | 0.554708 | 1 |
| Mrpl34 | -0.01191 | 0.721122 | 1 |
| C130050O18Rik | -0.01192 | 0.572217 | 1 |
| Lbr | -0.01193 | 0.327693 | 1 |
| Dpm2 | -0.01196 | 0.803132 | 1 |
| Clp1 | -0.01196 | 0.165382 | 1 |
| BC004004 | -0.01197 | 0.467952 | 1 |
| Uvssa | -0.01198 | 0.500699 | 1 |
| Zfyve19 | -0.01198 | 0.673569 | 1 |
| Cep135 | -0.01198 | 0.833625 | 1 |
| Zfp326 | -0.01199 | 0.986262 | 1 |
| Clk3 | -0.012 | 0.540471 | 1 |
| Zfp512b | -0.01201 | 0.447456 | 1 |
| Lamtor1 | -0.01201 | 0.712963 | 1 |
| Panx1 | -0.01202 | 0.165889 | 1 |
| Psmd5 | -0.01202 | 0.393011 | 1 |
| A530017D24Rik | -0.01203 | 0.073334 | 1 |
| Dynlt1f | -0.01205 | 0.785676 | 1 |
| Cox14 | -0.01207 | 0.941967 | 1 |
| Dnaaf5 | -0.01207 | 0.19314 | 1 |
| Mrps26 | -0.01207 | 0.305272 | 1 |
| Tsr3 | -0.01208 | 0.505685 | 1 |
| Bcl2l12 | -0.01209 | 0.106056 | 1 |
| Fggy | -0.01209 | 0.720927 | 1 |
| Rbbp4 | -0.0121 | 0.683512 | 1 |
| 9530062K07Rik | -0.01211 | 0.487583 | 1 |
| Eif3d | -0.01211 | 0.927054 | 1 |
| Ppp2r5a | -0.01212 | 0.573835 | 1 |
| Prrc1 | -0.01214 | 0.368494 | 1 |
| B230219D22Rik | -0.01215 | 0.865782 | 1 |
| Batf | -0.01215 | 0.446381 | 1 |
| Cnksr3 | -0.01215 | 0.576666 | 1 |
| Tmem156 | -0.01216 | 0.248538 | 1 |
| Rfxap | -0.01218 | 0.51062 | 1 |
| AW549877 | -0.01219 | 0.43084 | 1 |
| Lclat1 | -0.01219 | 0.456792 | 1 |
| 4930549G23Rik | -0.0122 | 0.26695 | 1 |
| Cdc42se1 | -0.01222 | 0.255953 | 1 |
| Ubxn2b | -0.01222 | 0.817533 | 1 |
| Inppl1 | -0.01228 | 0.370686 | 1 |
| Aste1 | -0.01229 | 0.175667 | 1 |
| Rcc1 | -0.0123 | 0.404277 | 1 |
| Med8 | -0.01231 | 0.299541 | 1 |
| 1500011B03Rik | -0.01232 | 0.641869 | 1 |
| S100a10 | -0.01234 | 0.923202 | 1 |

| | | | |
|---|---|---|---|
| Slc25a19 | -0.01235 | 0.038739 | 1 |
| Dip2b | -0.01236 | 0.577832 | 1 |
| Flna | -0.01236 | 0.705867 | 1 |
| Nat10 | -0.01237 | 0.69532 | 1 |
| Smox | -0.01237 | 0.536828 | 1 |
| Med28 | -0.01238 | 0.652103 | 1 |
| Zc3hc1 | -0.01239 | 0.043276 | 1 |
| Wdr82 | -0.01239 | 0.598423 | 1 |
| Gm32036 | -0.0124 | 0.283779 | 1 |
| Snx8 | -0.01241 | 0.993261 | 1 |
| Rgp1 | -0.01241 | 0.600621 | 1 |
| Itga5 | -0.01241 | 0.955168 | 1 |
| Il10rb | -0.01243 | 0.8703 | 1 |
| 1700123O20Rik | -0.01243 | 0.670944 | 1 |
| Exosc8 | -0.01244 | 0.587789 | 1 |
| Slc35e3 | -0.01245 | 0.623346 | 1 |
| Zfp87 | -0.0125 | 0.008462 | 1 |
| Klhl23 | -0.0125 | 0.247406 | 1 |
| Cyren | -0.01251 | 0.127113 | 1 |
| Polr1b | -0.01252 | 0.18168 | 1 |
| Prkcq | -0.01253 | 0.775614 | 1 |
| Zfp821 | -0.01254 | 0.545155 | 1 |
| Pitpnm1 | -0.01256 | 0.833881 | 1 |
| Nsmf | -0.01256 | 0.079322 | 1 |
| Yae1d1 | -0.01256 | 0.549916 | 1 |
| Trim35 | -0.01257 | 0.900508 | 1 |
| Tulp3 | -0.01257 | 0.196806 | 1 |
| Ndufaf7 | -0.01257 | 0.652595 | 1 |
| Tmem150a | -0.01259 | 0.649734 | 1 |
| Pus1 | -0.0126 | 0.507891 | 1 |
| Sp3os | -0.01262 | 0.67838 | 1 |
| Zfp956 | -0.01262 | 0.040029 | 1 |
| Slc35f6 | -0.01264 | 0.834382 | 1 |
| Cdk2 | -0.01266 | 0.373581 | 1 |
| Cops5 | -0.01266 | 0.328418 | 1 |
| Ndufa9 | -0.01268 | 0.688853 | 1 |
| Ahdc1 | -0.01269 | 0.474867 | 1 |
| Gm50012 | -0.01271 | 0.977642 | 1 |
| Gprasp1 | -0.01271 | 0.285103 | 1 |
| Cmtr1 | -0.01272 | 0.251659 | 1 |
| Tlr3 | -0.01272 | 0.862162 | 1 |
| Ubxn8 | -0.01272 | 0.624047 | 1 |
| Tmem154 | -0.01274 | 0.306689 | 1 |
| Magohb | -0.01274 | 0.271246 | 1 |
| Tnni2 | -0.01276 | 0.876538 | 1 |
| Arhgap9 | -0.0128 | 0.952445 | 1 |
| Psme4 | -0.0128 | 0.985484 | 1 |
| Agpat5 | -0.01281 | 0.861195 | 1 |

| | | | |
|---|---|---|---|
| Irak1 | -0.01282 | 0.885607 | 1 |
| Gpr137 | -0.01282 | 0.670224 | 1 |
| Bloc1s6 | -0.01282 | 0.407818 | 1 |
| Abracl | -0.01282 | 0.676257 | 1 |
| Cbx1 | -0.01283 | 0.674836 | 1 |
| Klc2 | -0.01283 | 0.693198 | 1 |
| Foxo1 | -0.01285 | 0.836153 | 1 |
| P3h3 | -0.01286 | 0.921329 | 1 |
| Ptger4 | -0.01286 | 0.583959 | 1 |
| Phyh | -0.01287 | 0.924844 | 1 |
| Lmbrd1 | -0.0129 | 0.412532 | 1 |
| Vps9d1 | -0.01291 | 0.718602 | 1 |
| Rab14 | -0.01292 | 0.53433 | 1 |
| Zfp608 | -0.01295 | 0.366057 | 1 |
| Wdr75 | -0.01298 | 0.748187 | 1 |
| Lncppara | -0.01298 | 0.572997 | 1 |
| Ggnbp2 | -0.01298 | 0.798828 | 1 |
| Arap1 | -0.01298 | 0.356848 | 1 |
| Ankrd13a | -0.01299 | 0.885931 | 1 |
| AI480526 | -0.01301 | 0.250904 | 1 |
| Eci1 | -0.01302 | 0.803021 | 1 |
| Isoc1 | -0.01303 | 0.740346 | 1 |
| Ube2v2 | -0.01304 | 0.540693 | 1 |
| Med6 | -0.01306 | 0.58892 | 1 |
| Lrrk1 | -0.01307 | 0.791112 | 1 |
| Gtf2h1 | -0.01309 | 0.130661 | 1 |
| Phactr2 | -0.01309 | 0.673817 | 1 |
| Fam160a2 | -0.0131 | 0.169944 | 1 |
| Dubr | -0.0131 | 0.423743 | 1 |
| Usp42 | -0.0131 | 0.053767 | 1 |
| Vamp8 | -0.01311 | 0.933544 | 1 |
| Zranb2 | -0.01312 | 0.77232 | 1 |
| Mrpl4 | -0.01313 | 0.954596 | 1 |
| Dolk | -0.01315 | 0.148533 | 1 |
| Ubp1 | -0.01316 | 0.949978 | 1 |
| Acrbp | -0.01317 | 0.305486 | 1 |
| Qars | -0.01318 | 0.40009 | 1 |
| Fbxo45 | -0.01319 | 0.383723 | 1 |
| Tipin | -0.01319 | 0.900615 | 1 |
| Plin3 | -0.01321 | 0.764335 | 1 |
| Zfp429 | -0.01322 | 0.280698 | 1 |
| Gm10125 | -0.01323 | 0.960981 | 1 |
| Ndufs4 | -0.01323 | 0.762853 | 1 |
| Grk5 | -0.01324 | 0.06751 | 1 |
| Nr2f6 | -0.01324 | 0.536637 | 1 |
| Cbr4 | -0.01325 | 0.408492 | 1 |
| Nr1h2 | -0.01325 | 0.650072 | 1 |
| Adam10 | -0.01327 | 0.76348 | 1 |

| | | | |
|---|---|---|---|
| Ppm1d | -0.01328 | 0.686555 | 1 |
| E2f3 | -0.01328 | 0.566971 | 1 |
| Thoc3 | -0.0133 | 0.089717 | 1 |
| Cd2bp2 | -0.01331 | 0.636222 | 1 |
| Btbd2 | -0.01331 | 0.491614 | 1 |
| Tmem165 | -0.01332 | 0.519972 | 1 |
| Zfp748 | -0.01332 | 0.75106 | 1 |
| Naf1 | -0.01333 | 0.920566 | 1 |
| Dgka | -0.01335 | 0.511505 | 1 |
| Agl | -0.01335 | 0.47594 | 1 |
| Ipo13 | -0.01335 | 0.356258 | 1 |
| Tdp1 | -0.01336 | 0.926869 | 1 |
| Fbxl17 | -0.01336 | 0.667874 | 1 |
| Atp5l | -0.01336 | 0.550984 | 1 |
| Sirt3 | -0.01337 | 0.949917 | 1 |
| Stoml1 | -0.01338 | 0.493464 | 1 |
| Med20 | -0.01338 | 0.640267 | 1 |
| Cep83os | -0.01339 | 0.386564 | 1 |
| Dnmt1 | -0.0134 | 0.713272 | 1 |
| Atp5c1 | -0.01342 | 0.977003 | 1 |
| Abi1 | -0.01343 | 0.73645 | 1 |
| Rab1b | -0.01344 | 0.81805 | 1 |
| Zscan29 | -0.01344 | 0.197666 | 1 |
| Fra10ac1 | -0.01346 | 0.619829 | 1 |
| Ergic1 | -0.01346 | 0.429911 | 1 |
| Nt5c3 | -0.01347 | 0.446854 | 1 |
| Mrpl53 | -0.01347 | 0.30132 | 1 |
| Gm15635 | -0.01349 | 0.377099 | 1 |
| Rbm19 | -0.01349 | 0.141766 | 1 |
| Pbx3 | -0.0135 | 0.615875 | 1 |
| Rpgrip1 | -0.0135 | 0.287217 | 1 |
| Mvp | -0.01352 | 0.402202 | 1 |
| Dhx35 | -0.01352 | 0.282146 | 1 |
| Rap1a | -0.01352 | 0.356342 | 1 |
| Gm43331 | -0.01353 | 0.081143 | 1 |
| Ino80b | -0.01353 | 0.668858 | 1 |
| Wdr35 | -0.01354 | 0.189181 | 1 |
| Pmpcb | -0.01355 | 0.986079 | 1 |
| Mrpl47 | -0.01355 | 0.171473 | 1 |
| Rnf24 | -0.01356 | 0.446656 | 1 |
| Sfswap | -0.01357 | 0.092229 | 1 |
| Nup50 | -0.01359 | 0.371834 | 1 |
| Kpna1 | -0.01362 | 0.35738 | 1 |
| 2310061I04Rik | -0.01362 | 0.060446 | 1 |
| Hmbox1 | -0.01363 | 0.369804 | 1 |
| Ndor1 | -0.01364 | 0.34105 | 1 |
| Hpn | -0.01364 | 0.797761 | 1 |
| Slc38a1 | -0.01364 | 0.956667 | 1 |

| | | | |
|---|---|---|---|
| Bnip2 | -0.01367 | 0.826642 | 1 |
| Gm27010 | -0.01367 | 0.013124 | 1 |
| Ticam1 | -0.01367 | 0.969578 | 1 |
| Il1rl2 | -0.01367 | 0.193214 | 1 |
| Slc26a2 | -0.0137 | 0.506153 | 1 |
| Jmy | -0.0137 | 0.385151 | 1 |
| Ncoa6 | -0.01372 | 0.36157 | 1 |
| Suz12 | -0.01372 | 0.482075 | 1 |
| Simc1 | -0.01375 | 0.894369 | 1 |
| Pip4p1 | -0.01375 | 0.914559 | 1 |
| Trmt10c | -0.01376 | 0.632064 | 1 |
| Cox10 | -0.01377 | 0.113954 | 1 |
| Gstcd | -0.01378 | 0.705979 | 1 |
| Usp6nl | -0.01378 | 0.990444 | 1 |
| Gigyf2 | -0.01379 | 0.40802 | 1 |
| Kras | -0.01379 | 0.914541 | 1 |
| Mbd3 | -0.0138 | 0.944915 | 1 |
| Abcb10 | -0.01381 | 0.04283 | 1 |
| Hdhd2 | -0.01381 | 0.753147 | 1 |
| Pkdcc | -0.01383 | 0.628955 | 1 |
| Prune1 | -0.01384 | 0.608664 | 1 |
| Zpr1 | -0.01387 | 0.88198 | 1 |
| Atp5g3 | -0.01387 | 0.77821 | 1 |
| Qsox1 | -0.01387 | 0.33949 | 1 |
| Metap1d | -0.01388 | 0.6563 | 1 |
| Siah1a | -0.01388 | 0.453274 | 1 |
| Gm28417 | -0.01389 | 0.454149 | 1 |
| Tcf12 | -0.01389 | 0.728058 | 1 |
| Gm15513 | -0.01389 | 0.21488 | 1 |
| Tor4a | -0.0139 | 0.325972 | 1 |
| Copb2 | -0.01391 | 0.470431 | 1 |
| Gm19605 | -0.01392 | 0.174876 | 1 |
| Kat6a | -0.01393 | 0.978148 | 1 |
| Gm11084 | -0.01395 | 0.492618 | 1 |
| Sv2a | -0.01397 | 0.399822 | 1 |
| Rcc1l | -0.01397 | 0.870624 | 1 |
| Lias | -0.01397 | 0.646586 | 1 |
| Zfp212 | -0.01397 | 0.773845 | 1 |
| Rybp | -0.01399 | 0.641346 | 1 |
| Bcl2a1a | -0.01399 | 0.893552 | 1 |
| Mapre3 | -0.01399 | 0.261896 | 1 |
| Cdk9 | -0.014 | 0.672705 | 1 |
| Ppfia4 | -0.01401 | 0.535558 | 1 |
| Fbxw8 | -0.01402 | 0.165373 | 1 |
| Brpf1 | -0.01403 | 0.654685 | 1 |
| Eri1 | -0.01407 | 0.65099 | 1 |
| Casc3 | -0.01408 | 0.445656 | 1 |
| Ticam2 | -0.01408 | 0.289238 | 1 |

| | | | |
|---|---|---|---|
| Cdkn2aip | -0.01409 | 0.513474 | 1 |
| Prickle1 | -0.01409 | 0.314755 | 1 |
| Tra2b | -0.0141 | 0.489699 | 1 |
| Il1rap | -0.01414 | 0.742416 | 1 |
| Gm11973 | -0.01414 | 0.295733 | 1 |
| Acvr1b | -0.01417 | 0.271849 | 1 |
| Ube2d2a | -0.01417 | 0.818779 | 1 |
| Adgb | -0.01419 | 0.218329 | 1 |
| Slc41a3 | -0.01419 | 0.92859 | 1 |
| Ctps | -0.0142 | 0.76268 | 1 |
| Dhdh | -0.01421 | 0.10132 | 1 |
| Qser1 | -0.01421 | 0.6042 | 1 |
| 2610008E11Rik | -0.01423 | 0.394376 | 1 |
| Zbtb7b | -0.01424 | 0.11674 | 1 |
| Tnfsf8 | -0.01427 | 0.614063 | 1 |
| Gatad1 | -0.0143 | 0.644828 | 1 |
| Degs2 | -0.01431 | 0.112756 | 1 |
| Mms22l | -0.01431 | 0.150745 | 1 |
| Edrf1 | -0.01432 | 0.239643 | 1 |
| Zhx2 | -0.01434 | 0.695602 | 1 |
| Trdmt1 | -0.01434 | 0.296469 | 1 |
| Mettl16 | -0.01435 | 0.770702 | 1 |
| Rhbdd3 | -0.01437 | 0.315234 | 1 |
| Sdad1 | -0.01439 | 0.640912 | 1 |
| Hexim1 | -0.0144 | 0.404355 | 1 |
| Sf3a1 | -0.0144 | 0.507737 | 1 |
| Ubtf | -0.0144 | 0.822681 | 1 |
| Snhg3 | -0.01443 | 0.648103 | 1 |
| Sirt2 | -0.01443 | 0.562027 | 1 |
| Pot1b | -0.01443 | 0.48679 | 1 |
| Gas8 | -0.01444 | 0.254695 | 1 |
| Cmtm4 | -0.01444 | 0.566138 | 1 |
| Slc35c1 | -0.01445 | 0.586855 | 1 |
| 5530601H04Rik | -0.01445 | 0.208427 | 1 |
| Wdr74 | -0.01445 | 0.147294 | 1 |
| Ppa1 | -0.01445 | 0.690893 | 1 |
| Wdr5 | -0.01446 | 0.283038 | 1 |
| Slc8a1 | -0.01446 | 0.394495 | 1 |
| Pdyn | -0.01448 | 0.44876 | 1 |
| Gpam | -0.01452 | 0.592606 | 1 |
| Mbtps1 | -0.01452 | 0.875465 | 1 |
| Card6 | -0.01453 | 0.144815 | 1 |
| Rcbtb1 | -0.01454 | 0.447284 | 1 |
| Crtc2 | -0.01455 | 0.089335 | 1 |
| Naip2 | -0.01456 | 0.257005 | 1 |
| Mras | -0.01458 | 0.770769 | 1 |
| Homer1 | -0.01459 | 0.178593 | 1 |
| Mettl4 | -0.01459 | 0.699371 | 1 |

| | | | |
|---|---|---|---|
| Ube3a | -0.01459 | 0.518662 | 1 |
| Ccnt2 | -0.0146 | 0.958535 | 1 |
| Acad11 | -0.0146 | 0.86414 | 1 |
| Otulin | -0.01463 | 0.980176 | 1 |
| Ube2l6 | -0.01465 | 0.34299 | 1 |
| Wdr4 | -0.01465 | 0.970553 | 1 |
| Osgepl1 | -0.01466 | 0.466351 | 1 |
| Polr1c | -0.01467 | 0.478116 | 1 |
| Mospd1 | -0.01469 | 0.257861 | 1 |
| Sf3b2 | -0.0147 | 0.941596 | 1 |
| Rxrb | -0.01471 | 0.537542 | 1 |
| Rae1 | -0.01471 | 0.385533 | 1 |
| Camta2 | -0.01471 | 0.495371 | 1 |
| Mfsd11 | -0.01472 | 0.291377 | 1 |
| Fam117a | -0.01474 | 0.090798 | 1 |
| Tut1 | -0.01474 | 0.032331 | 1 |
| Rtca | -0.01474 | 0.968292 | 1 |
| Gclm | -0.01474 | 0.174383 | 1 |
| Thap3 | -0.01476 | 0.772676 | 1 |
| Itpr2 | -0.01477 | 0.466307 | 1 |
| Dip2a | -0.01477 | 0.783088 | 1 |
| Nubp2 | -0.01478 | 0.402421 | 1 |
| Htra3 | -0.01478 | 0.232516 | 1 |
| Gpr18 | -0.01478 | 0.486834 | 1 |
| Abcc3 | -0.01479 | 0.679369 | 1 |
| Rapsn | -0.0148 | 0.758578 | 1 |
| 4930517O19Rik | -0.0148 | 0.883626 | 1 |
| Arhgdib | -0.01481 | 0.401042 | 1 |
| Ccdc73 | -0.01482 | 0.395939 | 1 |
| Trp53rka | -0.01485 | 0.291668 | 1 |
| Slc31a2 | -0.01487 | 0.929019 | 1 |
| Klf3 | -0.01487 | 0.817808 | 1 |
| Chchd6 | -0.01487 | 0.427642 | 1 |
| 9430091E24Rik | -0.01488 | 0.581073 | 1 |
| Dgkh | -0.0149 | 0.490421 | 1 |
| Htra2 | -0.0149 | 0.591985 | 1 |
| Ece1 | -0.0149 | 0.235932 | 1 |
| Chst7 | -0.0149 | 0.954449 | 1 |
| Cbx7 | -0.01491 | 0.073294 | 1 |
| Blnk | -0.01491 | 0.404246 | 1 |
| Psmb9 | -0.01491 | 0.682268 | 1 |
| Dcps | -0.01491 | 0.316862 | 1 |
| Cipc | -0.01491 | 0.137389 | 1 |
| Ubxn7 | -0.01492 | 0.465535 | 1 |
| Sbf2 | -0.01493 | 0.728747 | 1 |
| Snx15 | -0.01495 | 0.890242 | 1 |
| 2310058D17Rik | -0.01496 | 0.011313 | 1 |
| Vps29 | -0.01496 | 0.592728 | 1 |

| | | | |
|---|---|---|---|
| Gpn1 | -0.01497 | 0.055342 | 1 |
| Gm13269 | -0.01497 | 0.993892 | 1 |
| Skiv2l | -0.01497 | 0.110637 | 1 |
| Fxr1 | -0.01497 | 0.660813 | 1 |
| Gpatch2l | -0.01498 | 0.496607 | 1 |
| Hdac1 | -0.01499 | 0.901033 | 1 |
| Sgsm3 | -0.01499 | 0.111845 | 1 |
| Ftsj3 | -0.01499 | 0.411766 | 1 |
| Parp16 | -0.01499 | 0.937717 | 1 |
| Pop5 | -0.015 | 0.380316 | 1 |
| C2cd2l | -0.015 | 0.215264 | 1 |
| Clk4 | -0.01501 | 0.333367 | 1 |
| Zfp426 | -0.01501 | 0.260668 | 1 |
| Zfp1 | -0.01501 | 0.185304 | 1 |
| Pced1b | -0.01501 | 0.839643 | 1 |
| Wdr11 | -0.01504 | 0.336981 | 1 |
| Cdan1 | -0.01505 | 0.212323 | 1 |
| Stk25 | -0.01506 | 0.111014 | 1 |
| Wrn | -0.01507 | 0.753552 | 1 |
| Cryzl2 | -0.01509 | 0.624702 | 1 |
| Ly9 | -0.0151 | 0.495563 | 1 |
| Vps72 | -0.01511 | 0.511705 | 1 |
| Ikbip | -0.01512 | 0.084138 | 1 |
| Vps11 | -0.01512 | 0.579564 | 1 |
| Pa2g4 | -0.01512 | 0.838942 | 1 |
| Ms4a6b | -0.01513 | 0.954437 | 1 |
| Setdb2 | -0.01513 | 0.016013 | 1 |
| Mipep | -0.01514 | 0.505914 | 1 |
| Senp7 | -0.01515 | 0.144781 | 1 |
| Gm28198 | -0.01515 | 0.703615 | 1 |
| Adnp2 | -0.01516 | 0.485848 | 1 |
| Gm32856 | -0.01516 | 0.580406 | 1 |
| Rps6kc1 | -0.01516 | 0.762743 | 1 |
| Fam13a | -0.01517 | 0.218059 | 1 |
| Urm1 | -0.01518 | 0.733357 | 1 |
| Acss1 | -0.0152 | 0.460826 | 1 |
| Sec16a | -0.01521 | 0.939268 | 1 |
| Dpp7 | -0.01521 | 0.971998 | 1 |
| Akr1a1 | -0.01522 | 0.682811 | 1 |
| Kpna6 | -0.01524 | 0.584979 | 1 |
| Fmn1 | -0.01524 | 0.867137 | 1 |
| Epha2 | -0.01524 | 0.664658 | 1 |
| Kars | -0.01525 | 0.320882 | 1 |
| Hdac3 | -0.01525 | 0.738174 | 1 |
| Chac2 | -0.01526 | 0.02113 | 1 |
| D6Wsu163e | -0.01526 | 0.424001 | 1 |
| Qsox2 | -0.01527 | 0.161615 | 1 |
| Tm9sf1 | -0.01528 | 0.728989 | 1 |

| | | | |
|---|---|---|---|
| C130026I21Rik | -0.01528 | 0.193285 | 1 |
| Mlh3 | -0.01528 | 0.092058 | 1 |
| Tcf25 | -0.01529 | 0.871778 | 1 |
| Parvg | -0.01531 | 0.852468 | 1 |
| Pex3 | -0.01533 | 0.735774 | 1 |
| Tom1l1 | -0.01534 | 0.938426 | 1 |
| Sgcb | -0.01535 | 0.500302 | 1 |
| Epc2 | -0.01536 | 0.802241 | 1 |
| Pdcd2 | -0.01536 | 0.939641 | 1 |
| Smc2 | -0.01537 | 0.887832 | 1 |
| Sec23a | -0.01538 | 0.551386 | 1 |
| Rac2 | -0.01539 | 0.494925 | 1 |
| Diaph2 | -0.0154 | 0.432002 | 1 |
| H2-Ob | -0.0154 | 0.660855 | 1 |
| Eif4g2 | -0.0154 | 0.84768 | 1 |
| Gtpbp6 | -0.01542 | 0.6046 | 1 |
| Sfxn2 | -0.01543 | 0.296773 | 1 |
| Baz2a | -0.01545 | 0.189053 | 1 |
| Phf10 | -0.01545 | 0.207712 | 1 |
| Rnf8 | -0.01546 | 0.935957 | 1 |
| 1810013L24Rik | -0.01546 | 0.856373 | 1 |
| Furin | -0.01548 | 0.728633 | 1 |
| Glb1l | -0.01548 | 0.370414 | 1 |
| Gm28791 | -0.01548 | 0.134627 | 1 |
| Ppt1 | -0.01549 | 0.799157 | 1 |
| Smyd3 | -0.01549 | 0.169694 | 1 |
| D130062J10Rik | -0.0155 | 0.205342 | 1 |
| Ninl | -0.0155 | 0.398358 | 1 |
| Mta2 | -0.0155 | 0.688895 | 1 |
| Gga1 | -0.01551 | 0.751226 | 1 |
| Rnf168 | -0.01551 | 0.454575 | 1 |
| Tbc1d25 | -0.01551 | 0.041044 | 1 |
| Ctse | -0.01555 | 0.213763 | 1 |
| Ofd1 | -0.01556 | 0.346606 | 1 |
| Dcaf8 | -0.01556 | 0.127335 | 1 |
| Fbxo25 | -0.01556 | 0.755856 | 1 |
| Atg9a | -0.01556 | 0.601991 | 1 |
| Rbm12b2 | -0.01557 | 0.32413 | 1 |
| Mfn2 | -0.01558 | 0.854457 | 1 |
| Gpcpd1 | -0.01558 | 0.902497 | 1 |
| Ube2v1 | -0.01559 | 0.942299 | 1 |
| Xrcc6 | -0.0156 | 0.300901 | 1 |
| Eya4 | -0.0156 | 0.931441 | 1 |
| Brwd1 | -0.0156 | 0.74548 | 1 |
| Lat2 | -0.01561 | 0.992207 | 1 |
| Patl2 | -0.01561 | 0.914844 | 1 |
| Trnt1 | -0.01561 | 0.308672 | 1 |
| Urb1 | -0.01562 | 0.518162 | 1 |

| | | | |
|---|---|---|---|
| Stk16 | -0.01562 | 0.06357 | 1 |
| Usp54 | -0.01563 | 0.326242 | 1 |
| Ensa | -0.01564 | 0.936083 | 1 |
| Top2b | -0.01564 | 0.87165 | 1 |
| Zfp141 | -0.01565 | 0.690718 | 1 |
| Gm7030 | -0.01565 | 0.024114 | 1 |
| Trappc9 | -0.01566 | 0.372077 | 1 |
| Lrrc29 | -0.01568 | 0.31472 | 1 |
| Auh | -0.01571 | 0.701651 | 1 |
| Api5 | -0.01572 | 0.966984 | 1 |
| Trim30d | -0.01572 | 0.158483 | 1 |
| Cds1 | -0.01574 | 0.511045 | 1 |
| Mrpl43 | -0.01574 | 0.75442 | 1 |
| Zfp984 | -0.01576 | 0.199492 | 1 |
| March3 | -0.01576 | 0.206799 | 1 |
| Ints10 | -0.01576 | 0.827759 | 1 |
| Slc38a2 | -0.01577 | 0.750348 | 1 |
| Rpf2 | -0.01578 | 0.651293 | 1 |
| Pnn | -0.01578 | 0.380031 | 1 |
| Mfhas1 | -0.0158 | 0.726703 | 1 |
| Rad54l2 | -0.01581 | 0.169053 | 1 |
| Adprhl2 | -0.01582 | 0.834867 | 1 |
| Mrpl38 | -0.01582 | 0.579246 | 1 |
| Zrsr2 | -0.01583 | 0.803541 | 1 |
| Stard5 | -0.01584 | 0.132557 | 1 |
| Psd4 | -0.01586 | 0.846664 | 1 |
| Hacd4 | -0.01587 | 0.701054 | 1 |
| Rsf1 | -0.01587 | 0.511191 | 1 |
| Snx13 | -0.01587 | 0.578551 | 1 |
| Clasp2 | -0.01589 | 0.812438 | 1 |
| E430024I08Rik | -0.01589 | 0.361951 | 1 |
| Creb3l2 | -0.01589 | 0.538106 | 1 |
| Sco2 | -0.0159 | 0.647644 | 1 |
| Nup85 | -0.0159 | 0.016969 | 1 |
| Pofut2 | -0.01591 | 0.51517 | 1 |
| Cenpj | -0.01592 | 0.135887 | 1 |
| Il17ra | -0.01592 | 0.992266 | 1 |
| 5830432E09Rik | -0.01592 | 0.136613 | 1 |
| Smad2 | -0.01592 | 0.334149 | 1 |
| 0610009B22Rik | -0.01593 | 0.565087 | 1 |
| Ankrd26 | -0.01594 | 0.161705 | 1 |
| Otub1 | -0.01595 | 0.766921 | 1 |
| Sap130 | -0.01597 | 0.27666 | 1 |
| Pyroxd1 | -0.01597 | 0.560451 | 1 |
| Rhno1 | -0.01597 | 0.231889 | 1 |
| Zbtb14 | -0.01597 | 0.302966 | 1 |
| Tmem203 | -0.01598 | 0.615046 | 1 |
| Slc25a17 | -0.01598 | 0.439782 | 1 |

| | | | |
|---|---|---|---|
| 4930473A02Rik | -0.01598 | 0.507792 | 1 |
| Ccdc117 | -0.01598 | 0.115575 | 1 |
| Smcr8 | -0.01601 | 0.420903 | 1 |
| Bms1 | -0.01601 | 0.475776 | 1 |
| Fbxw2 | -0.01601 | 0.548719 | 1 |
| Fbxo6 | -0.01604 | 0.563679 | 1 |
| Crbn | -0.01604 | 0.370242 | 1 |
| Vps41 | -0.01604 | 0.793661 | 1 |
| Ccdc171 | -0.01605 | 0.937591 | 1 |
| Uspl1 | -0.01606 | 0.552397 | 1 |
| Polg | -0.01607 | 0.834264 | 1 |
| Cic | -0.01609 | 0.525087 | 1 |
| Nkiras1 | -0.01612 | 0.410371 | 1 |
| Gatd1 | -0.01612 | 0.401932 | 1 |
| Cfap97 | -0.01613 | 0.588164 | 1 |
| Rad51d | -0.01614 | 0.050078 | 1 |
| Mb21d2 | -0.01615 | 0.056655 | 1 |
| Zbtb8os | -0.01616 | 0.983158 | 1 |
| Tcp11l1 | -0.01616 | 0.561836 | 1 |
| Ddb2 | -0.01616 | 0.007998 | 1 |
| Gnpda1 | -0.01617 | 0.986575 | 1 |
| Svil | -0.01618 | 0.457234 | 1 |
| Fbxw5 | -0.01618 | 0.884299 | 1 |
| Zfp830 | -0.0162 | 0.579425 | 1 |
| Mks1 | -0.01621 | 0.448647 | 1 |
| Zbtb17 | -0.01622 | 0.640636 | 1 |
| Gm13708 | -0.01622 | 0.237365 | 1 |
| Polr2g | -0.01622 | 0.831553 | 1 |
| Gm29488 | -0.01622 | 0.367478 | 1 |
| 2900005J15Rik | -0.01624 | 0.024606 | 1 |
| Tsc2 | -0.01625 | 0.758011 | 1 |
| Pwwp3a | -0.01625 | 0.120001 | 1 |
| Gm17268 | -0.01626 | 0.032092 | 1 |
| Ppp6c | -0.01628 | 0.744351 | 1 |
| Timm44 | -0.01629 | 0.408812 | 1 |
| Brap | -0.0163 | 0.172293 | 1 |
| Gar1 | -0.0163 | 0.538435 | 1 |
| Psmd14 | -0.01631 | 0.636025 | 1 |
| Tmem205 | -0.01631 | 0.46 | 1 |
| Cdc26 | -0.01632 | 0.77222 | 1 |
| Fam129b | -0.01632 | 0.064482 | 1 |
| Eloa | -0.01633 | 0.896917 | 1 |
| 0610030E20Rik | -0.01635 | 0.655464 | 1 |
| Uhrf1bp1 | -0.01635 | 0.827599 | 1 |
| Spsb3 | -0.01635 | 0.363018 | 1 |
| Gm36198 | -0.01636 | 0.842059 | 1 |
| Bsdc1 | -0.01637 | 0.103419 | 1 |
| Mcat | -0.0164 | 0.437843 | 1 |

| | | | |
|---|---|---|---|
| Ankrd50 | -0.01643 | 0.414332 | 1 |
| Pias4 | -0.01643 | 0.093189 | 1 |
| Ulk1 | -0.01643 | 0.25971 | 1 |
| Zfp983 | -0.01645 | 0.340477 | 1 |
| Sh3gl1 | -0.01645 | 0.88001 | 1 |
| Ibtk | -0.01645 | 0.477376 | 1 |
| 1600010M07Rik | -0.01646 | 0.855968 | 1 |
| Psmf1 | -0.01646 | 0.263823 | 1 |
| Pola2 | -0.01646 | 0.690883 | 1 |
| G3bp2 | -0.01646 | 0.641027 | 1 |
| Zfp408 | -0.01646 | 0.04946 | 1 |
| Cul9 | -0.01647 | 0.904904 | 1 |
| Tardbp | -0.01648 | 0.604858 | 1 |
| Rbm48 | -0.01649 | 0.488451 | 1 |
| St7 | -0.01649 | 0.607215 | 1 |
| Ube2q2 | -0.01651 | 0.485049 | 1 |
| Pus10 | -0.01653 | 0.289337 | 1 |
| Zfp740 | -0.01653 | 0.692393 | 1 |
| Usp37 | -0.01654 | 0.60569 | 1 |
| Pak1ip1 | -0.01654 | 0.517295 | 1 |
| Gmfb | -0.01654 | 0.956437 | 1 |
| Wsb2 | -0.01655 | 0.683914 | 1 |
| Ahsa1 | -0.01657 | 0.098131 | 1 |
| Rfx5 | -0.01659 | 0.508833 | 1 |
| Coq8b | -0.0166 | 0.788491 | 1 |
| Oxr1 | -0.01661 | 0.648238 | 1 |
| Cox7a2 | -0.01662 | 0.63411 | 1 |
| Acadsb | -0.01662 | 0.804798 | 1 |
| Pabpc4 | -0.01663 | 0.734891 | 1 |
| Ccdc84 | -0.01663 | 0.540638 | 1 |
| Abhd16a | -0.01666 | 0.890528 | 1 |
| Prps2 | -0.01667 | 0.77467 | 1 |
| Il2rg | -0.01668 | 0.724554 | 1 |
| Mgll | -0.0167 | 0.649793 | 1 |
| Fdx1 | -0.0167 | 0.941284 | 1 |
| Mib2 | -0.01672 | 0.35194 | 1 |
| Mrpl22 | -0.01673 | 0.222745 | 1 |
| Serinc3 | -0.01673 | 0.132479 | 1 |
| Srl | -0.01674 | 0.038729 | 1 |
| Atl3 | -0.01674 | 0.638696 | 1 |
| Sgsm2 | -0.01675 | 0.246783 | 1 |
| Ndufaf3 | -0.01676 | 0.614603 | 1 |
| Jkamp | -0.01676 | 0.476344 | 1 |
| Hmgn1 | -0.01677 | 0.892709 | 1 |
| Oma1 | -0.01677 | 0.85194 | 1 |
| Nox1 | -0.01682 | 0.727246 | 1 |
| Sugt1 | -0.01683 | 0.909301 | 1 |
| Elk3 | -0.01684 | 0.664521 | 1 |

| | | | |
|---|---|---|---|
| Gtf3c3 | -0.01684 | 0.812892 | 1 |
| Gm4356 | -0.01687 | 0.026567 | 1 |
| Polr3b | -0.0169 | 0.875203 | 1 |
| Hps4 | -0.0169 | 0.993251 | 1 |
| Lasp1 | -0.01691 | 0.898736 | 1 |
| Polr3k | -0.01692 | 0.045972 | 1 |
| R74862 | -0.01693 | 0.292466 | 1 |
| Sat1 | -0.01693 | 0.197222 | 1 |
| Rpa3 | -0.01694 | 0.257354 | 1 |
| Dnajc15 | -0.01694 | 0.9236 | 1 |
| Ddx5 | -0.01695 | 0.319304 | 1 |
| Eif1ad | -0.01695 | 0.374182 | 1 |
| Anapc11 | -0.01695 | 0.844345 | 1 |
| Gm6712 | -0.01696 | 0.147912 | 1 |
| Slc25a16 | -0.01696 | 0.429781 | 1 |
| Myd88 | -0.01697 | 0.247805 | 1 |
| Prelid3b | -0.01699 | 0.970132 | 1 |
| Cpt1a | -0.01702 | 0.56166 | 1 |
| Stam | -0.01702 | 0.927524 | 1 |
| Ndufv1 | -0.01703 | 0.602982 | 1 |
| Zfp472 | -0.01703 | 0.274844 | 1 |
| Ap5z1 | -0.01703 | 0.019461 | 1 |
| Uck1 | -0.01704 | 0.286728 | 1 |
| Rdx | -0.01704 | 0.616143 | 1 |
| E230016M11Rik | -0.01704 | 0.744356 | 1 |
| Bcorl1 | -0.01705 | 0.585881 | 1 |
| Fam76a | -0.01705 | 0.854298 | 1 |
| Zfp687 | -0.01708 | 0.444579 | 1 |
| Zfp180 | -0.01708 | 0.226809 | 1 |
| Traf7 | -0.01712 | 0.184031 | 1 |
| Cep250 | -0.01713 | 0.13177 | 1 |
| Txn2 | -0.01714 | 0.317998 | 1 |
| Rpf1 | -0.01715 | 0.811525 | 1 |
| Cenpt | -0.01715 | 0.077109 | 1 |
| Ring1 | -0.01716 | 0.887933 | 1 |
| Cox18 | -0.01716 | 0.457548 | 1 |
| Atp9b | -0.01717 | 0.365994 | 1 |
| Gm42917 | -0.01718 | 0.099802 | 1 |
| Arfgap1 | -0.01719 | 0.552996 | 1 |
| Leng1 | -0.01719 | 0.820925 | 1 |
| Wdfy3 | -0.0172 | 0.55418 | 1 |
| Hcfc2 | -0.01721 | 0.69185 | 1 |
| Papss1 | -0.01721 | 0.266717 | 1 |
| Arl13b | -0.01722 | 0.056124 | 1 |
| Tmem131 | -0.01725 | 0.143402 | 1 |
| Wdr25 | -0.01725 | 0.667363 | 1 |
| Ctr9 | -0.01727 | 0.658708 | 1 |
| Cars | -0.01727 | 0.340879 | 1 |

| | | | |
|---|---|---|---|
| Efcab2 | -0.01728 | 0.595496 | 1 |
| Tmem265 | -0.01729 | 0.188619 | 1 |
| Ypel3 | -0.01731 | 0.362093 | 1 |
| Pitpnb | -0.01731 | 0.76398 | 1 |
| Farsa | -0.01732 | 0.484615 | 1 |
| Tfip11 | -0.01734 | 0.509631 | 1 |
| Mvb12a | -0.01735 | 0.423847 | 1 |
| Trnau1ap | -0.01735 | 0.562018 | 1 |
| Tsnax | -0.01735 | 0.874414 | 1 |
| Luzp1 | -0.01737 | 0.779598 | 1 |
| Alkbh3 | -0.01737 | 0.826417 | 1 |
| Dram2 | -0.01737 | 0.731475 | 1 |
| Ppp1r3b | -0.01738 | 0.705701 | 1 |
| Iqgap1 | -0.01742 | 0.703588 | 1 |
| Adgrl2 | -0.01745 | 0.976202 | 1 |
| Zfp330 | -0.01747 | 0.851614 | 1 |
| Ric8a | -0.01747 | 0.149886 | 1 |
| Cep95 | -0.0175 | 0.328989 | 1 |
| Hivep1 | -0.0175 | 0.204893 | 1 |
| Adnp | -0.01752 | 0.934386 | 1 |
| Vps4a | -0.01752 | 0.25328 | 1 |
| Tagap | -0.01753 | 0.099712 | 1 |
| Ppil4 | -0.01756 | 0.280775 | 1 |
| Dhodh | -0.01756 | 0.218968 | 1 |
| Rab12 | -0.01758 | 0.524172 | 1 |
| Rab28 | -0.01758 | 0.854602 | 1 |
| 0610010F05Rik | -0.0176 | 0.28997 | 1 |
| Huwe1 | -0.01761 | 0.706635 | 1 |
| Acsf2 | -0.01762 | 0.277861 | 1 |
| Ip6k2 | -0.01763 | 0.474424 | 1 |
| Tcerg1 | -0.01766 | 0.626352 | 1 |
| Ccdc91 | -0.01767 | 0.121865 | 1 |
| Tbc1d16 | -0.01768 | 0.376668 | 1 |
| Acad10 | -0.01768 | 0.38628 | 1 |
| Gpd2 | -0.01769 | 0.398188 | 1 |
| Ubtd2 | -0.01769 | 0.325207 | 1 |
| Git1 | -0.01769 | 0.146149 | 1 |
| Mbtps2 | -0.0177 | 0.242744 | 1 |
| Ccdc90b | -0.0177 | 0.478232 | 1 |
| Fam214b | -0.0177 | 0.075817 | 1 |
| Inpp5a | -0.01771 | 0.579527 | 1 |
| 1110038B12Rik | -0.01771 | 0.703573 | 1 |
| Zfp263 | -0.01772 | 0.786451 | 1 |
| Zfp410 | -0.01772 | 0.644986 | 1 |
| Med19 | -0.01774 | 0.878434 | 1 |
| Nedd1 | -0.01777 | 0.597931 | 1 |
| Usf3 | -0.01778 | 0.833165 | 1 |
| 4833438C02Rik | -0.01778 | 0.27678 | 1 |

| | | | |
|---|---|---|---|
| D030028A08Rik | -0.0178 | 0.129982 | 1 |
| Phip | -0.01782 | 0.694276 | 1 |
| Fam151b | -0.01783 | 0.048897 | 1 |
| Mphosph10 | -0.01784 | 0.754989 | 1 |
| Ptdss2 | -0.01784 | 0.42484 | 1 |
| Alg11 | -0.01786 | 0.337669 | 1 |
| Zfp60 | -0.01787 | 0.136206 | 1 |
| Impa1 | -0.01787 | 0.275995 | 1 |
| Cyb5b | -0.01789 | 0.476761 | 1 |
| Ppt2 | -0.0179 | 0.652229 | 1 |
| H2-DMb2 | -0.01791 | 0.500019 | 1 |
| Mitd1 | -0.01792 | 0.199808 | 1 |
| D2hgdh | -0.01793 | 0.180754 | 1 |
| Frs2 | -0.01794 | 0.696918 | 1 |
| Pgpep1 | -0.01797 | 0.392514 | 1 |
| Mrps31 | -0.01798 | 0.454895 | 1 |
| Prr3 | -0.01799 | 0.257968 | 1 |
| Arcn1 | -0.01799 | 0.720667 | 1 |
| Tmem86b | -0.01801 | 0.10086 | 1 |
| C1galt1c1 | -0.01802 | 0.957644 | 1 |
| Rp9 | -0.01802 | 0.75432 | 1 |
| Wtap | -0.01804 | 0.961783 | 1 |
| Fam98a | -0.01804 | 0.272959 | 1 |
| Supt7l | -0.01804 | 0.039272 | 1 |
| Pop1 | -0.01806 | 0.535318 | 1 |
| Pcyox1 | -0.01806 | 0.607912 | 1 |
| Minpp1 | -0.01809 | 0.927487 | 1 |
| Mrpl10 | -0.01812 | 0.283725 | 1 |
| Rnf146 | -0.01812 | 0.431921 | 1 |
| Nr3c2 | -0.01812 | 0.669287 | 1 |
| Sfxn1 | -0.01817 | 0.873892 | 1 |
| Zfp654 | -0.01818 | 0.439774 | 1 |
| Tasp1 | -0.01818 | 0.103774 | 1 |
| Unc93b1 | -0.01821 | 0.386571 | 1 |
| Fmo5 | -0.01823 | 0.357125 | 1 |
| Pkn1 | -0.01824 | 0.598329 | 1 |
| BC005537 | -0.01825 | 0.935372 | 1 |
| Gtf2e2 | -0.01825 | 0.480142 | 1 |
| Zdhhc1 | -0.01826 | 0.094418 | 1 |
| Ccnt1 | -0.01828 | 0.366337 | 1 |
| 6030458C11Rik | -0.01829 | 0.203711 | 1 |
| Gm4117 | -0.01829 | 0.478378 | 1 |
| Prkaca | -0.01831 | 0.521118 | 1 |
| Adam9 | -0.01831 | 0.629486 | 1 |
| Ptar1 | -0.01831 | 0.383353 | 1 |
| Abhd5 | -0.01832 | 0.094082 | 1 |
| Ap1s1 | -0.01832 | 0.735326 | 1 |
| Ipo4 | -0.01833 | 0.35756 | 1 |

| | | | |
|---|---|---|---|
| Nt5c | -0.01834 | 0.544014 | 1 |
| Arhgap5 | -0.01834 | 0.853117 | 1 |
| Kat2a | -0.01834 | 0.071672 | 1 |
| Zer1 | -0.01834 | 0.71854 | 1 |
| Slc30a1 | -0.01835 | 0.236612 | 1 |
| Gm26749 | -0.01838 | 0.436046 | 1 |
| Edc4 | -0.01838 | 0.131856 | 1 |
| Ppfibp1 | -0.01842 | 0.207909 | 1 |
| Cdc42 | -0.01844 | 0.692412 | 1 |
| Otud5 | -0.01845 | 0.765641 | 1 |
| Creb5 | -0.01845 | 0.739327 | 1 |
| Itpripl1 | -0.01846 | 0.766888 | 1 |
| Nf2 | -0.01846 | 0.101681 | 1 |
| Josd2 | -0.01847 | 0.415863 | 1 |
| Prelid1 | -0.01848 | 0.545553 | 1 |
| Prox1 | -0.01849 | 0.225271 | 1 |
| Ppm1g | -0.01849 | 0.282944 | 1 |
| 4833420G17Rik | -0.01854 | 0.88025 | 1 |
| Rbm5 | -0.01856 | 0.677383 | 1 |
| Gm15952 | -0.01858 | 0.339578 | 1 |
| Mapk7 | -0.01858 | 0.08518 | 1 |
| Hebp1 | -0.0186 | 0.575706 | 1 |
| Map3k2 | -0.01862 | 0.61496 | 1 |
| Ccl2 | -0.01862 | 0.194686 | 1 |
| Relt | -0.01863 | 0.163823 | 1 |
| Txlna | -0.01863 | 0.449033 | 1 |
| Msh2 | -0.01864 | 0.134348 | 1 |
| Sdhaf3 | -0.01865 | 0.188653 | 1 |
| Gm44899 | -0.01867 | 0.287266 | 1 |
| Ddx19b | -0.01868 | 0.438278 | 1 |
| Zfp626 | -0.01869 | 0.802977 | 1 |
| Man2a2 | -0.01869 | 0.919752 | 1 |
| Rnf103 | -0.0187 | 0.990203 | 1 |
| Opa1 | -0.0187 | 0.537477 | 1 |
| Nupr1 | -0.0187 | 0.571304 | 1 |
| Bend6 | -0.01872 | 0.970593 | 1 |
| Cars2 | -0.01872 | 0.775058 | 1 |
| Med27 | -0.01872 | 0.153443 | 1 |
| Card9 | -0.01873 | 0.682383 | 1 |
| Usp32 | -0.01873 | 0.715372 | 1 |
| Ercc6 | -0.01873 | 0.385238 | 1 |
| Surf6 | -0.01874 | 0.166452 | 1 |
| Rprd1b | -0.01875 | 0.416425 | 1 |
| 4921524J17Rik | -0.01876 | 0.243943 | 1 |
| Gys1 | -0.01877 | 0.143624 | 1 |
| Zfp672 | -0.01879 | 0.588038 | 1 |
| E130102H24Rik | -0.01879 | 0.523227 | 1 |
| Ints14 | -0.01881 | 0.693224 | 1 |

| | | | |
|---|---|---|---|
| Nln | -0.01882 | 0.993224 | 1 |
| Ddx39 | -0.01883 | 0.490186 | 1 |
| Scpep1 | -0.01884 | 0.403864 | 1 |
| Ccdc32 | -0.01887 | 0.986248 | 1 |
| Catspere2 | -0.01888 | 0.815488 | 1 |
| Gm20559 | -0.01889 | 0.14019 | 1 |
| Slc35d1 | -0.01891 | 0.040836 | 1 |
| Vps50 | -0.01892 | 0.438482 | 1 |
| Pum1 | -0.01892 | 0.932322 | 1 |
| Tbc1d10b | -0.01894 | 0.599328 | 1 |
| Bcl3 | -0.01894 | 0.081587 | 1 |
| E4f1 | -0.01896 | 0.98717 | 1 |
| Wdr3 | -0.01896 | 0.105666 | 1 |
| Agap1 | -0.01896 | 0.709662 | 1 |
| Senp3 | -0.01898 | 0.108116 | 1 |
| Ube2i | -0.01898 | 0.306939 | 1 |
| Rnf19a | -0.019 | 0.780065 | 1 |
| Slc25a40 | -0.019 | 0.862787 | 1 |
| Gab1 | -0.01903 | 0.70306 | 1 |
| Galnt7 | -0.01905 | 0.914121 | 1 |
| Taf11 | -0.01906 | 0.128552 | 1 |
| Ap1b1 | -0.01906 | 0.58313 | 1 |
| Bscl2 | -0.01907 | 0.916919 | 1 |
| Evi2 | -0.0191 | 0.030834 | 1 |
| Snhg9 | -0.0191 | 0.681603 | 1 |
| Slc25a43 | -0.01911 | 0.074139 | 1 |
| Vps13b | -0.01911 | 0.450826 | 1 |
| Slc25a46 | -0.01912 | 0.322678 | 1 |
| Alg13 | -0.01912 | 0.085421 | 1 |
| Mettl25 | -0.01912 | 0.518118 | 1 |
| Pphln1 | -0.01912 | 0.519074 | 1 |
| Hmgxb3 | -0.01913 | 0.731925 | 1 |
| Bcl10 | -0.01915 | 0.198708 | 1 |
| Snapin | -0.01916 | 0.936628 | 1 |
| Ocrl | -0.01917 | 0.403093 | 1 |
| Fam160b1 | -0.01917 | 0.40744 | 1 |
| Zbtb4 | -0.01917 | 0.532816 | 1 |
| Slc25a53 | -0.01919 | 0.772326 | 1 |
| Slc25a13 | -0.01921 | 0.11479 | 1 |
| Cbfa2t2 | -0.01921 | 0.699621 | 1 |
| Calcrl | -0.01923 | 0.184963 | 1 |
| Aasdh | -0.01924 | 0.069608 | 1 |
| Mcm7 | -0.01924 | 0.384344 | 1 |
| Usp10 | -0.01925 | 0.288843 | 1 |
| Afg3l2 | -0.01926 | 0.475124 | 1 |
| Eef1akmt4 | -0.01927 | 0.094835 | 1 |
| Aarsd1 | -0.01928 | 0.800449 | 1 |
| Cr1l | -0.01928 | 0.586355 | 1 |

| | | | |
|---|---|---|---|
| Lsm12 | -0.01928 | 0.650768 | 1 |
| Hsd17b4 | -0.01929 | 0.980188 | 1 |
| Cope | -0.01931 | 0.747504 | 1 |
| Dpy19l1 | -0.01931 | 0.306496 | 1 |
| Mlkl | -0.01932 | 0.192321 | 1 |
| Samd8 | -0.01933 | 0.688299 | 1 |
| Rc3h2 | -0.01933 | 0.090545 | 1 |
| Smarcd2 | -0.01933 | 0.149282 | 1 |
| Ap3d1 | -0.01934 | 0.818805 | 1 |
| Map4k2 | -0.01935 | 0.658269 | 1 |
| Ftx | -0.01935 | 0.473577 | 1 |
| Cluh | -0.01938 | 0.7488 | 1 |
| Col15a1 | -0.01939 | 0.100419 | 1 |
| Smndc1 | -0.01939 | 0.685365 | 1 |
| Pprc1 | -0.01939 | 0.789784 | 1 |
| Tmem87a | -0.0194 | 0.303898 | 1 |
| Taf12 | -0.01941 | 0.271639 | 1 |
| Gm3448 | -0.01941 | 0.027263 | 1 |
| Galc | -0.01943 | 0.729439 | 1 |
| Ice2 | -0.01944 | 0.597108 | 1 |
| Nt5c2 | -0.01944 | 0.856884 | 1 |
| Atp6v1g2 | -0.01945 | 0.056774 | 1 |
| Rtcb | -0.01948 | 0.548198 | 1 |
| Mief1 | -0.01949 | 0.444646 | 1 |
| Psmd10 | -0.0195 | 0.86545 | 1 |
| Sun1 | -0.0195 | 0.33731 | 1 |
| Zfp729a | -0.01955 | 0.146823 | 1 |
| Cnot6l | -0.01956 | 0.537821 | 1 |
| Vps37b | -0.01956 | 0.429555 | 1 |
| Zfp182 | -0.01957 | 0.601059 | 1 |
| Hdhd5 | -0.01957 | 0.166473 | 1 |
| C1galt1 | -0.01957 | 0.771374 | 1 |
| Gm37768 | -0.01958 | 0.345468 | 1 |
| Golga7 | -0.01959 | 0.758462 | 1 |
| Gm41764 | -0.01961 | 0.560191 | 1 |
| Sept2 | -0.01963 | 0.501017 | 1 |
| Ganab | -0.01964 | 0.348098 | 1 |
| Gm21860 | -0.01964 | 0.346568 | 1 |
| Ing4 | -0.01964 | 0.4856 | 1 |
| Wasl | -0.01966 | 0.7271 | 1 |
| Plcl1 | -0.01968 | 0.358744 | 1 |
| Ppp4r2 | -0.01969 | 0.406349 | 1 |
| Cxxc1 | -0.01969 | 0.341764 | 1 |
| Zbtb44 | -0.0197 | 0.202478 | 1 |
| Zbtb41 | -0.0197 | 0.815512 | 1 |
| Paxip1 | -0.01971 | 0.593522 | 1 |
| Nlrx1 | -0.01971 | 0.041726 | 1 |
| Zfp74 | -0.01973 | 0.104416 | 1 |

| | | | |
|---|---|---|---|
| Acaa1a | -0.01975 | 0.632722 | 1 |
| Fam172a | -0.01975 | 0.368019 | 1 |
| Dlg3 | -0.01975 | 0.248444 | 1 |
| Rad9b | -0.01975 | 0.012316 | 1 |
| St14 | -0.01976 | 0.829655 | 1 |
| Tigd2 | -0.01979 | 0.633874 | 1 |
| 6330418K02Rik | -0.01981 | 0.27768 | 1 |
| Cnep1r1 | -0.01981 | 0.56544 | 1 |
| Cd101 | -0.01982 | 0.365953 | 1 |
| Pkd1 | -0.01983 | 0.980153 | 1 |
| Adsl | -0.01983 | 0.542054 | 1 |
| Shfl | -0.01986 | 0.406563 | 1 |
| Ipo11 | -0.01987 | 0.804581 | 1 |
| Kmt5a | -0.01988 | 0.629888 | 1 |
| Tmem120a | -0.0199 | 0.368616 | 1 |
| Nol9 | -0.01991 | 0.317328 | 1 |
| Tfg | -0.01992 | 0.502863 | 1 |
| Aldh16a1 | -0.01994 | 0.197834 | 1 |
| Tspan5 | -0.01996 | 0.482744 | 1 |
| Trappc6b | -0.01997 | 0.065993 | 1 |
| Nipal3 | -0.01997 | 0.00278 | 1 |
| Nsun4 | -0.01997 | 0.173487 | 1 |
| Sh2b3 | -0.01998 | 0.516556 | 1 |
| Sirt7 | -0.01999 | 0.876832 | 1 |
| Pdcd7 | -0.01999 | 0.296393 | 1 |
| Cyhr1 | -0.02 | 0.18834 | 1 |
| Krr1 | -0.02 | 0.509488 | 1 |
| Hdac11 | -0.02001 | 0.573563 | 1 |
| Spg7 | -0.02001 | 0.909021 | 1 |
| Nmt1 | -0.02001 | 0.236274 | 1 |
| Xndc1 | -0.02005 | 0.985754 | 1 |
| Ubl4a | -0.02005 | 0.539767 | 1 |
| Snapc4 | -0.02006 | 0.108499 | 1 |
| Phrf1 | -0.02007 | 0.04496 | 1 |
| Rad52 | -0.02007 | 0.132994 | 1 |
| Tmem216 | -0.02007 | 0.575281 | 1 |
| Pwp1 | -0.02008 | 0.304956 | 1 |
| Snx20 | -0.02008 | 0.271989 | 1 |
| Mfsd13a | -0.02008 | 0.161969 | 1 |
| Psen1 | -0.02009 | 0.803124 | 1 |
| 4930469K13Rik | -0.02009 | 0.480504 | 1 |
| Bnip3l | -0.0201 | 0.758386 | 1 |
| Glud1 | -0.0201 | 0.976333 | 1 |
| Fars2 | -0.02011 | 0.503382 | 1 |
| Usp48 | -0.02011 | 0.377281 | 1 |
| Paip2b | -0.02011 | 0.330924 | 1 |
| Pfkfb2 | -0.02012 | 0.396198 | 1 |
| Glcci1 | -0.02012 | 0.613762 | 1 |

| | | | |
|---|---|---|---|
| Snap47 | -0.02013 | 0.045986 | 1 |
| Aebp2 | -0.02014 | 0.272883 | 1 |
| Ddx46 | -0.02015 | 0.678858 | 1 |
| Cep104 | -0.02016 | 0.116018 | 1 |
| Slc25a26 | -0.02016 | 0.922282 | 1 |
| Zc3h3 | -0.02016 | 0.133505 | 1 |
| Fastkd1 | -0.02017 | 0.016964 | 1 |
| Cdk2ap1 | -0.02017 | 0.714926 | 1 |
| Mta1 | -0.02018 | 0.268631 | 1 |
| Rpp25l | -0.02018 | 0.541394 | 1 |
| Strn4 | -0.02018 | 0.808731 | 1 |
| Rbms2 | -0.02019 | 0.109217 | 1 |
| Rpa1 | -0.02019 | 0.503755 | 1 |
| Mob3c | -0.02019 | 0.909665 | 1 |
| Igip | -0.0202 | 0.159532 | 1 |
| Usp31 | -0.02021 | 0.09609 | 1 |
| Gnptab | -0.02022 | 0.286287 | 1 |
| Ipo7 | -0.02022 | 0.553152 | 1 |
| Nos1ap | -0.02024 | 0.673551 | 1 |
| Trim47 | -0.02026 | 0.356342 | 1 |
| Rpp14 | -0.02026 | 0.117422 | 1 |
| Chmp3 | -0.02027 | 0.284951 | 1 |
| Gas7 | -0.02027 | 0.958472 | 1 |
| Pgm1 | -0.02027 | 0.340481 | 1 |
| Stambp | -0.02028 | 0.77202 | 1 |
| Ctdp1 | -0.02028 | 0.481699 | 1 |
| Abce1 | -0.02029 | 0.99089 | 1 |
| Mrps18b | -0.02031 | 0.489196 | 1 |
| Sharpin | -0.02031 | 0.709851 | 1 |
| Adap1 | -0.02031 | 0.738698 | 1 |
| Ccnh | -0.02032 | 0.310142 | 1 |
| Sod1 | -0.02032 | 0.965617 | 1 |
| Gdi1 | -0.02034 | 0.425895 | 1 |
| Slc35e2 | -0.02034 | 0.722356 | 1 |
| Wwp1 | -0.02036 | 0.754973 | 1 |
| Dph5 | -0.02037 | 0.873508 | 1 |
| Usp45 | -0.02038 | 0.547571 | 1 |
| Ddx42 | -0.02038 | 0.64459 | 1 |
| Lsg1 | -0.02038 | 0.782631 | 1 |
| Gpatch2 | -0.02038 | 0.319832 | 1 |
| Mfn1 | -0.02039 | 0.53022 | 1 |
| 2900097C17Rik | -0.02039 | 0.893594 | 1 |
| Tulp4 | -0.02041 | 0.257418 | 1 |
| Zbtb37 | -0.02045 | 0.263649 | 1 |
| Cab39l | -0.02045 | 0.254823 | 1 |
| Brms1l | -0.02047 | 0.831684 | 1 |
| Camkk2 | -0.02048 | 0.416833 | 1 |
| Cyb561d1 | -0.02048 | 0.024303 | 1 |

| | | | |
|---|---|---|---|
| Ifi27 | -0.02052 | 0.592402 | 1 |
| Pag1 | -0.02057 | 0.494214 | 1 |
| Crebl2 | -0.02059 | 0.257117 | 1 |
| Inpp5d | -0.02059 | 0.100647 | 1 |
| Psip1 | -0.02059 | 0.442321 | 1 |
| Imp4 | -0.0206 | 0.939305 | 1 |
| Dnajc14 | -0.0206 | 0.250056 | 1 |
| Kdm4b | -0.02061 | 0.503752 | 1 |
| Pald1 | -0.02061 | 0.268051 | 1 |
| Prrc2c | -0.02061 | 0.439549 | 1 |
| Bahd1 | -0.02061 | 0.104863 | 1 |
| Fam234a | -0.02061 | 0.871121 | 1 |
| Dock9 | -0.02062 | 0.387615 | 1 |
| Tent4a | -0.02062 | 0.167723 | 1 |
| Rhbdd1 | -0.02065 | 0.847738 | 1 |
| Purg | -0.02067 | 0.533232 | 1 |
| Nek4 | -0.02068 | 0.414033 | 1 |
| Rpap2 | -0.02068 | 0.411779 | 1 |
| Tmem186 | -0.02071 | 0.211649 | 1 |
| Atad2b | -0.02071 | 0.168913 | 1 |
| Tmem68 | -0.02071 | 0.542531 | 1 |
| Clcn5 | -0.02073 | 0.824989 | 1 |
| Stx6 | -0.02075 | 0.77027 | 1 |
| Lyrm1 | -0.02075 | 0.633421 | 1 |
| Tbcel | -0.02076 | 0.136089 | 1 |
| Rpe | -0.02077 | 0.949438 | 1 |
| Tgs1 | -0.02078 | 0.768501 | 1 |
| Mrps25 | -0.02079 | 0.36474 | 1 |
| Polr3h | -0.0208 | 0.038823 | 1 |
| Taf1a | -0.02081 | 0.006006 | 1 |
| Fastk | -0.02084 | 0.125251 | 1 |
| 1700030K09Rik | -0.02085 | 0.080749 | 1 |
| Pnpla6 | -0.02086 | 0.12581 | 1 |
| Rrp15 | -0.02086 | 0.630692 | 1 |
| Ms4a6c | -0.02088 | 0.421828 | 1 |
| Pld2 | -0.0209 | 0.200832 | 1 |
| Mrpl13 | -0.0209 | 0.611418 | 1 |
| Gpn2 | -0.02095 | 0.223767 | 1 |
| Atp13a2 | -0.02096 | 0.598358 | 1 |
| Dbnl | -0.02097 | 0.142771 | 1 |
| Pcid2 | -0.02098 | 0.138053 | 1 |
| Neu1 | -0.02099 | 0.300892 | 1 |
| Dhx33 | -0.02101 | 0.192215 | 1 |
| Ap4b1 | -0.02102 | 0.042333 | 1 |
| BC051142 | -0.02102 | 0.093028 | 1 |
| Zbtb46 | -0.02103 | 0.398032 | 1 |
| Ildr2 | -0.02103 | 0.490606 | 1 |
| Baz1b | -0.02103 | 0.894151 | 1 |

| | | | |
|---|---|---|---|
| Pef1 | -0.02107 | 0.273076 | 1 |
| Ecpas | -0.02109 | 0.554727 | 1 |
| Nsmce4a | -0.0211 | 0.866987 | 1 |
| Zmym4 | -0.02111 | 0.671993 | 1 |
| Oip5os1 | -0.02113 | 0.922453 | 1 |
| Immt | -0.02114 | 0.445178 | 1 |
| Lcp1 | -0.02116 | 0.501703 | 1 |
| Gnpat | -0.02117 | 0.955782 | 1 |
| Mpi | -0.02117 | 0.69296 | 1 |
| Spsb2 | -0.02118 | 0.086366 | 1 |
| Slc9a1 | -0.02118 | 0.437006 | 1 |
| Helz2 | -0.02119 | 0.037592 | 1 |
| Nbn | -0.0212 | 0.265121 | 1 |
| Fbxo36 | -0.0212 | 0.109859 | 1 |
| Snx9 | -0.02122 | 0.367878 | 1 |
| Snap29 | -0.02123 | 0.124257 | 1 |
| Dnal1 | -0.02124 | 0.087079 | 1 |
| Lysmd3 | -0.02124 | 0.063484 | 1 |
| Mosmo | -0.02124 | 0.213695 | 1 |
| Tk1 | -0.02125 | 0.27127 | 1 |
| Tubb5 | -0.02125 | 0.542344 | 1 |
| Mxd4 | -0.02125 | 0.860999 | 1 |
| Ehd1 | -0.02129 | 0.808747 | 1 |
| Sik3 | -0.0213 | 0.953543 | 1 |
| Ebf3 | -0.02132 | 0.400167 | 1 |
| Slf1 | -0.02133 | 0.20432 | 1 |
| A530088E08Rik | -0.02133 | 0.042411 | 1 |
| Tlnrd1 | -0.02133 | 0.014702 | 1 |
| Znrf3 | -0.02133 | 0.55478 | 1 |
| Usp38 | -0.02136 | 0.412691 | 1 |
| Trappc13 | -0.02137 | 0.385439 | 1 |
| Alg8 | -0.02138 | 0.475626 | 1 |
| Asph | -0.02138 | 0.825445 | 1 |
| Sgms1 | -0.02139 | 0.431237 | 1 |
| Isg20l2 | -0.0214 | 0.234644 | 1 |
| Xpa | -0.0214 | 0.228273 | 1 |
| Spindoc | -0.02142 | 0.1588 | 1 |
| Pdik1l | -0.02142 | 0.335526 | 1 |
| H2afy | -0.02144 | 0.812262 | 1 |
| Wdr73 | -0.02145 | 0.608245 | 1 |
| Ifit3 | -0.02146 | 0.000767 | 1 |
| Man1b1 | -0.02146 | 0.567871 | 1 |
| Gm47601 | -0.02147 | 0.648213 | 1 |
| Cyb561a3 | -0.02151 | 0.916528 | 1 |
| Mrps11 | -0.02152 | 0.426459 | 1 |
| Oxsm | -0.02153 | 0.087339 | 1 |
| Atat1 | -0.02158 | 0.017425 | 1 |
| Slc20a2 | -0.02158 | 0.680248 | 1 |

| | | | |
|---|---|---|---|
| AI413582 | -0.02158 | 0.392567 | 1 |
| Etfa | -0.0216 | 0.439935 | 1 |
| Flywch1 | -0.02161 | 0.04084 | 1 |
| Ern1 | -0.02162 | 0.188266 | 1 |
| Zfp655 | -0.02163 | 0.975841 | 1 |
| Scrib | -0.02163 | 0.065584 | 1 |
| Trub2 | -0.02164 | 0.233785 | 1 |
| Pnpt1 | -0.02164 | 0.089244 | 1 |
| Rassf2 | -0.02166 | 0.582855 | 1 |
| Gm47167 | -0.02167 | 0.636041 | 1 |
| Dnajb12 | -0.02168 | 0.172389 | 1 |
| Kdm5b | -0.0217 | 0.132025 | 1 |
| Slc9a3r1 | -0.02171 | 0.257307 | 1 |
| 2410004B18Rik | -0.02171 | 0.274044 | 1 |
| Nufip1 | -0.02172 | 0.158584 | 1 |
| Lhpp | -0.02172 | 0.109921 | 1 |
| Zc3h8 | -0.02174 | 0.267911 | 1 |
| Vamp1 | -0.02174 | 0.191506 | 1 |
| Sbf1 | -0.02175 | 0.434282 | 1 |
| Gatb | -0.02175 | 0.518621 | 1 |
| Atp1a1 | -0.02176 | 0.749455 | 1 |
| Creb3 | -0.02177 | 0.590184 | 1 |
| Ccr1 | -0.02177 | 0.91292 | 1 |
| Rbm28 | -0.02178 | 0.684556 | 1 |
| Nudt7 | -0.02179 | 0.341095 | 1 |
| Erp44 | -0.02181 | 0.33654 | 1 |
| Nmi | -0.02182 | 0.474402 | 1 |
| Rgmb | -0.02183 | 0.371943 | 1 |
| Map1s | -0.02183 | 0.800345 | 1 |
| Tspan32 | -0.02185 | 0.58619 | 1 |
| Pde7a | -0.02187 | 0.323921 | 1 |
| Shld2 | -0.02187 | 0.362066 | 1 |
| Ptpn12 | -0.02188 | 0.248462 | 1 |
| Helq | -0.02189 | 0.082398 | 1 |
| Nol10 | -0.0219 | 0.130171 | 1 |
| Zbtb25 | -0.02191 | 0.051075 | 1 |
| Ndufs1 | -0.02191 | 0.034655 | 1 |
| Amotl1 | -0.02193 | 0.63731 | 1 |
| Nudt13 | -0.02193 | 0.059683 | 1 |
| Dgkq | -0.02193 | 0.107834 | 1 |
| Wdr91 | -0.02195 | 0.140075 | 1 |
| Uhrf1bp1l | -0.02195 | 0.464375 | 1 |
| Trim2 | -0.02196 | 0.14338 | 1 |
| Bad | -0.02196 | 0.280609 | 1 |
| Scarf2 | -0.02197 | 0.233176 | 1 |
| Fbxw11 | -0.02198 | 0.395059 | 1 |
| Dazap1 | -0.022 | 0.753546 | 1 |
| Snrnp70 | -0.02201 | 0.316185 | 1 |

| | | | |
|---|---|---|---|
| Slamf1 | -0.02202 | 0.055309 | 1 |
| Nab1 | -0.02203 | 0.312618 | 1 |
| Rabif | -0.02204 | 0.04453 | 1 |
| Rbm39 | -0.02205 | 0.352558 | 1 |
| Abcf3 | -0.02208 | 0.544959 | 1 |
| Morc2a | -0.02209 | 0.187482 | 1 |
| Deptor | -0.0221 | 0.194734 | 1 |
| Prr13 | -0.02212 | 0.350854 | 1 |
| Rrp7a | -0.02213 | 0.101881 | 1 |
| Aatk | -0.02214 | 0.017782 | 1 |
| Lemd3 | -0.02215 | 0.541704 | 1 |
| Gm43378 | -0.02217 | 0.012761 | 1 |
| Eif4ebp2 | -0.02217 | 0.039959 | 1 |
| Zfp276 | -0.0222 | 0.391336 | 1 |
| Cenpp | -0.02221 | 0.79852 | 1 |
| Gm16118 | -0.02223 | 0.124961 | 1 |
| Tes | -0.02224 | 0.588789 | 1 |
| Lilrb4a | -0.02224 | 0.532261 | 1 |
| Hgs | -0.02225 | 0.316391 | 1 |
| Kif1c | -0.02225 | 0.023462 | 1 |
| Camsap2 | -0.02225 | 0.761506 | 1 |
| Hs1bp3 | -0.02227 | 0.221499 | 1 |
| Cab39 | -0.02227 | 0.969153 | 1 |
| Helz | -0.02228 | 0.034309 | 1 |
| Fam13b | -0.02228 | 0.311155 | 1 |
| Fech | -0.02231 | 0.641972 | 1 |
| Unk | -0.02233 | 0.522013 | 1 |
| Micos10 | -0.02234 | 0.361237 | 1 |
| Aig1 | -0.02235 | 0.588306 | 1 |
| Polr3f | -0.02235 | 0.278124 | 1 |
| Unc119b | -0.02237 | 0.189877 | 1 |
| 8030462N17Rik | -0.02239 | 0.449994 | 1 |
| Stx17 | -0.0224 | 0.688711 | 1 |
| Hlcs | -0.0224 | 0.493253 | 1 |
| Arfrp1 | -0.02246 | 0.667113 | 1 |
| Zfp664 | -0.02247 | 0.035593 | 1 |
| Cwf19l2 | -0.02247 | 0.990759 | 1 |
| Pxmp4 | -0.02247 | 0.060603 | 1 |
| Ehbp1l1 | -0.02249 | 0.067416 | 1 |
| Pros1 | -0.0225 | 0.374416 | 1 |
| Fnip1 | -0.0225 | 0.194986 | 1 |
| Spg11 | -0.02253 | 0.207817 | 1 |
| Fam162a | -0.02253 | 0.271518 | 1 |
| Braf | -0.02255 | 0.786403 | 1 |
| Emc2 | -0.02255 | 0.429064 | 1 |
| Pogk | -0.02255 | 0.321435 | 1 |
| Stag1 | -0.02256 | 0.28666 | 1 |
| Farp2 | -0.02257 | 0.135713 | 1 |

| | | | |
|---|---|---|---|
| Aimp1 | -0.02258 | 0.820857 | 1 |
| Clk1 | -0.02258 | 0.618755 | 1 |
| Appbp2 | -0.0226 | 0.602434 | 1 |
| Ezh1 | -0.02261 | 0.370569 | 1 |
| Ppp1r12c | -0.02263 | 0.269553 | 1 |
| Selenoo | -0.02263 | 0.297932 | 1 |
| Rubcn | -0.02264 | 0.265839 | 1 |
| Copg1 | -0.02265 | 0.165038 | 1 |
| Ctps2 | -0.02266 | 0.155156 | 1 |
| Gm16552 | -0.02267 | 0.292243 | 1 |
| Fktn | -0.02268 | 0.109973 | 1 |
| Dnhd1 | -0.0227 | 0.029614 | 1 |
| Ethe1 | -0.0227 | 0.904338 | 1 |
| Mrpl44 | -0.02271 | 0.066505 | 1 |
| Hdac2 | -0.02271 | 0.697462 | 1 |
| Tmem229b | -0.02273 | 0.225686 | 1 |
| Sac3d1 | -0.02274 | 0.214619 | 1 |
| Acadm | -0.02274 | 0.27846 | 1 |
| Nipsnap2 | -0.02275 | 0.377647 | 1 |
| Wipi2 | -0.02277 | 0.13787 | 1 |
| Gm5431 | -0.02278 | 0.075088 | 1 |
| Stip1 | -0.0228 | 0.310935 | 1 |
| Zc3h14 | -0.02281 | 0.332828 | 1 |
| Ap3s2 | -0.02282 | 0.407916 | 1 |
| Rsf1os1 | -0.02283 | 0.363856 | 1 |
| Smu1 | -0.02283 | 0.589214 | 1 |
| Gm36279 | -0.02283 | 0.023622 | 1 |
| Macf1 | -0.02283 | 0.078859 | 1 |
| Rmdn2 | -0.02286 | 0.005229 | 1 |
| Kcnq1ot1 | -0.02287 | 0.629487 | 1 |
| Zfp346 | -0.02288 | 0.754748 | 1 |
| Cnnm4 | -0.02288 | 0.084972 | 1 |
| Tlr13 | -0.02289 | 0.956594 | 1 |
| Dnajc19 | -0.02289 | 0.795697 | 1 |
| Dnttip2 | -0.02291 | 0.109343 | 1 |
| Tfcp2 | -0.02291 | 0.135689 | 1 |
| Xkr6 | -0.02292 | 0.506859 | 1 |
| Rptor | -0.02293 | 0.418101 | 1 |
| Fes | -0.02293 | 0.803893 | 1 |
| Acad12 | -0.02298 | 0.334342 | 1 |
| Xpc | -0.02298 | 0.038813 | 1 |
| Rbck1 | -0.02299 | 0.060682 | 1 |
| Hbp1 | -0.023 | 0.494718 | 1 |
| Clic4 | -0.023 | 0.104465 | 1 |
| Atp6v1h | -0.02301 | 0.736518 | 1 |
| Ptbp3 | -0.02301 | 0.524949 | 1 |
| Dusp12 | -0.02305 | 0.518741 | 1 |
| Gm32849 | -0.02305 | 0.153376 | 1 |

| | | | |
|---|---|---|---|
| Polb | -0.02305 | 0.118888 | 1 |
| Zfp933 | -0.02305 | 0.411011 | 1 |
| Ube2w | -0.02305 | 0.50387 | 1 |
| Rtf2 | -0.02306 | 0.40355 | 1 |
| Stk11ip | -0.02306 | 0.15695 | 1 |
| Sdhd | -0.02312 | 0.945625 | 1 |
| Parn | -0.02312 | 0.901629 | 1 |
| Lhfpl2 | -0.02313 | 0.512458 | 1 |
| Mlycd | -0.02315 | 0.424818 | 1 |
| 6430548M08Rik | -0.02317 | 0.10241 | 1 |
| Adprm | -0.02318 | 0.069206 | 1 |
| Gcnt1 | -0.0232 | 0.516458 | 1 |
| Pcnx | -0.0232 | 0.151118 | 1 |
| Ptgfrn | -0.02321 | 0.114839 | 1 |
| Rnf149 | -0.02325 | 0.682955 | 1 |
| Abcf1 | -0.02326 | 0.799927 | 1 |
| Rfesd | -0.02327 | 0.172911 | 1 |
| Zcwpw2 | -0.02327 | 0.060159 | 1 |
| Stxbp5 | -0.02328 | 0.415046 | 1 |
| Pgs1 | -0.02331 | 0.683614 | 1 |
| Gtf2f1 | -0.02332 | 0.816897 | 1 |
| Amd1 | -0.02334 | 0.177577 | 1 |
| Dmxl2 | -0.02337 | 0.189937 | 1 |
| Zfp397 | -0.02338 | 0.157497 | 1 |
| Rnf40 | -0.0234 | 0.080451 | 1 |
| Dcaf13 | -0.0234 | 0.620917 | 1 |
| Taf4b | -0.02342 | 0.700949 | 1 |
| Xxylt1 | -0.02345 | 0.48728 | 1 |
| Ncaph2 | -0.02345 | 0.168815 | 1 |
| Paip2 | -0.02348 | 0.635671 | 1 |
| Acbd5 | -0.02351 | 0.637726 | 1 |
| Fbf1 | -0.02351 | 0.003666 | 1 |
| Rsrc1 | -0.02351 | 0.428647 | 1 |
| Cdkal1 | -0.02351 | 0.245369 | 1 |
| Ttc8 | -0.02352 | 0.689441 | 1 |
| Tomm40 | -0.02352 | 0.062708 | 1 |
| Cd84 | -0.02355 | 0.681446 | 1 |
| Tpm3 | -0.02356 | 0.351937 | 1 |
| Gm49359 | -0.02359 | 0.863269 | 1 |
| Tsen2 | -0.02361 | 0.113454 | 1 |
| Pcgf3 | -0.02361 | 0.125762 | 1 |
| Gan | -0.02361 | 0.062765 | 1 |
| Slc10a7 | -0.02365 | 0.719319 | 1 |
| Casd1 | -0.02365 | 0.238835 | 1 |
| Ammecr1 | -0.02366 | 0.137379 | 1 |
| Ints7 | -0.02369 | 0.788698 | 1 |
| Gm27003 | -0.0237 | 0.061068 | 1 |
| Erich1 | -0.0237 | 0.130478 | 1 |

| | | | |
|---|---|---|---|
| Abraxas2 | -0.02371 | 0.685912 | 1 |
| Chuk | -0.02371 | 0.07505 | 1 |
| Sh3kbp1 | -0.02373 | 0.395914 | 1 |
| Tmem126b | -0.02374 | 0.041055 | 1 |
| Scfd1 | -0.02375 | 0.987044 | 1 |
| Fbxo30 | -0.02378 | 0.825531 | 1 |
| Wdr13 | -0.02382 | 0.207685 | 1 |
| AU041133 | -0.02384 | 0.148786 | 1 |
| Tlk1 | -0.02385 | 0.33157 | 1 |
| Myo18a | -0.02386 | 0.124789 | 1 |
| Cep295 | -0.02386 | 0.235126 | 1 |
| Eif2b3 | -0.02386 | 0.465907 | 1 |
| Tle4 | -0.02386 | 0.572706 | 1 |
| Castor2 | -0.0239 | 0.137178 | 1 |
| Fam214a | -0.0239 | 0.347784 | 1 |
| Kyat3 | -0.0239 | 0.000277 | 1 |
| Smap2 | -0.0239 | 0.187853 | 1 |
| Arhgef10l | -0.02392 | 0.037646 | 1 |
| Haus2 | -0.02395 | 0.187274 | 1 |
| Zfp943 | -0.02395 | 0.353581 | 1 |
| Osbp | -0.02396 | 0.365491 | 1 |
| Ranbp2 | -0.02396 | 0.651397 | 1 |
| Nop56 | -0.02397 | 0.463178 | 1 |
| Zmat3 | -0.02397 | 0.134074 | 1 |
| Cmc1 | -0.024 | 0.431662 | 1 |
| Eif2ak1 | -0.024 | 0.666735 | 1 |
| Gm13091 | -0.02402 | 0.005647 | 1 |
| Ciao2b | -0.02403 | 0.076508 | 1 |
| Tal1 | -0.02405 | 0.045971 | 1 |
| Nkiras2 | -0.02405 | 0.096751 | 1 |
| Layn | -0.02407 | 0.463129 | 1 |
| Zfp958 | -0.02408 | 0.391219 | 1 |
| Gm14798 | -0.02409 | 0.006191 | 1 |
| Slc33a1 | -0.0241 | 0.749689 | 1 |
| Bri3bp | -0.02412 | 0.065171 | 1 |
| Vamp2 | -0.02412 | 0.048915 | 1 |
| Vamp4 | -0.02412 | 0.161711 | 1 |
| Armc7 | -0.02412 | 0.334844 | 1 |
| Hk1 | -0.02413 | 0.005959 | 1 |
| Ganc | -0.02413 | 0.571311 | 1 |
| F3 | -0.02414 | 0.02448 | 1 |
| Sap30l | -0.02415 | 0.109304 | 1 |
| Ube2g2 | -0.02416 | 0.134045 | 1 |
| Mcee | -0.02418 | 0.390208 | 1 |
| Shq1 | -0.02419 | 0.29991 | 1 |
| Ssb | -0.02421 | 0.811059 | 1 |
| Mpp5 | -0.02422 | 0.51948 | 1 |
| Myg1 | -0.02422 | 0.550293 | 1 |

| | | | |
|---|---|---|---|
| Gm15886 | -0.02422 | 0.03071 | 1 |
| Pitpnc1 | -0.02423 | 0.388177 | 1 |
| Slirp | -0.02428 | 0.412704 | 1 |
| Mmaa | -0.02428 | 0.375138 | 1 |
| Mapkbp1 | -0.02428 | 0.161415 | 1 |
| Smarce1 | -0.02431 | 0.890998 | 1 |
| Dpf2 | -0.02432 | 0.33152 | 1 |
| Foxp1 | -0.02435 | 0.317726 | 1 |
| Rac1 | -0.02435 | 0.34906 | 1 |
| Slc35a5 | -0.02436 | 0.096927 | 1 |
| Lrp1 | -0.02438 | 0.538879 | 1 |
| Cul5 | -0.02439 | 0.981336 | 1 |
| Zfp444 | -0.02441 | 0.033895 | 1 |
| Khnyn | -0.02442 | 0.225539 | 1 |
| Tmlhe | -0.02443 | 0.692334 | 1 |
| Ube2o | -0.02443 | 0.294704 | 1 |
| Ubald2 | -0.02445 | 0.570794 | 1 |
| Pacsin1 | -0.02448 | 0.080731 | 1 |
| Tinf2 | -0.02451 | 0.251711 | 1 |
| Cd200r1 | -0.02453 | 0.026418 | 1 |
| Nudt19 | -0.02453 | 0.145431 | 1 |
| Ap4m1 | -0.02454 | 0.499097 | 1 |
| Dpp3 | -0.02455 | 0.242216 | 1 |
| Emc1 | -0.02456 | 0.126295 | 1 |
| Fmr1 | -0.02456 | 0.996079 | 1 |
| Galns | -0.02457 | 0.035536 | 1 |
| Dab2 | -0.02458 | 0.711261 | 1 |
| 2700062C07Rik | -0.02459 | 0.10855 | 1 |
| Tpk1 | -0.02459 | 0.977095 | 1 |
| Nsfl1c | -0.0246 | 0.293775 | 1 |
| Dzip3 | -0.0246 | 0.049909 | 1 |
| Preb | -0.02462 | 0.64505 | 1 |
| Glod4 | -0.02465 | 0.288083 | 1 |
| Fry | -0.02466 | 0.743022 | 1 |
| Ralbp1 | -0.02466 | 0.17149 | 1 |
| Gm20342 | -0.02468 | 0.111502 | 1 |
| Gm27201 | -0.02468 | 0.178281 | 1 |
| Nckipsd | -0.02469 | 0.17987 | 1 |
| Asb7 | -0.0247 | 0.245747 | 1 |
| Brf1 | -0.0247 | 0.337971 | 1 |
| Gch1 | -0.0247 | 0.198624 | 1 |
| Plcb3 | -0.0247 | 0.735089 | 1 |
| Carf | -0.02471 | 0.074866 | 1 |
| Lrmda | -0.02471 | 0.099114 | 1 |
| Atp11b | -0.02472 | 0.318093 | 1 |
| Tubgcp4 | -0.02474 | 0.391723 | 1 |
| Napg | -0.02474 | 0.243321 | 1 |
| Kntc1 | -0.02477 | 0.960628 | 1 |

| | | | |
|---|---|---|---|
| Ttc27 | -0.02477 | 0.032766 | 1 |
| Zfp668 | -0.02477 | 0.034265 | 1 |
| 1700109H08Rik | -0.02478 | 0.180511 | 1 |
| Foxk2 | -0.02478 | 0.600589 | 1 |
| Mbd5 | -0.02479 | 0.354317 | 1 |
| Aen | -0.02481 | 0.091309 | 1 |
| Phf11a | -0.02481 | 0.067996 | 1 |
| Sympk | -0.02482 | 0.287518 | 1 |
| Dgcr8 | -0.02482 | 0.048783 | 1 |
| Arl8a | -0.02482 | 0.637796 | 1 |
| Stat5a | -0.02484 | 0.101352 | 1 |
| Malt1 | -0.02485 | 0.992848 | 1 |
| Commd6 | -0.02486 | 0.549444 | 1 |
| 8030453O22Rik | -0.02487 | 0.159598 | 1 |
| Gcn1 | -0.02488 | 0.292014 | 1 |
| Nip7 | -0.02489 | 0.098872 | 1 |
| Sh2b2 | -0.0249 | 0.427481 | 1 |
| Map3k12 | -0.02494 | 0.001358 | 1 |
| Slco4a1 | -0.02495 | 0.968277 | 1 |
| Ppp1r13b | -0.02496 | 0.215393 | 1 |
| Qtrt2 | -0.02497 | 0.069889 | 1 |
| Epg5 | -0.02497 | 0.36966 | 1 |
| 4833411C07Rik | -0.02498 | 0.04346 | 1 |
| Ddx56 | -0.02499 | 0.116176 | 1 |
| Samd4b | -0.02499 | 0.924045 | 1 |
| Zfp329 | -0.02502 | 0.960355 | 1 |
| Zfp942 | -0.02504 | 0.144753 | 1 |
| Eif2b5 | -0.02506 | 0.210859 | 1 |
| Il20rb | -0.02507 | 0.017997 | 1 |
| Mrpl57 | -0.02512 | 0.171385 | 1 |
| Zfp169 | -0.02516 | 0.456011 | 1 |
| Gorasp2 | -0.02517 | 0.105714 | 1 |
| Shoc2 | -0.02518 | 0.47604 | 1 |
| Ncoa5 | -0.02519 | 0.014448 | 1 |
| Rap1gap2 | -0.0252 | 0.236916 | 1 |
| Ncor1 | -0.02521 | 0.071962 | 1 |
| Rps6kb2 | -0.02522 | 0.044818 | 1 |
| Vezt | -0.02523 | 0.164339 | 1 |
| Dctn6 | -0.02525 | 0.202929 | 1 |
| Fblim1 | -0.02525 | 0.812075 | 1 |
| Mllt11 | -0.02528 | 0.646852 | 1 |
| Lrsam1 | -0.0253 | 0.139743 | 1 |
| Cdc123 | -0.02533 | 0.307273 | 1 |
| Specc1 | -0.02534 | 0.370841 | 1 |
| Tle1 | -0.02534 | 0.004324 | 1 |
| Pex6 | -0.02535 | 0.229745 | 1 |
| Pygo2 | -0.02537 | 0.054069 | 1 |
| Dyrk4 | -0.02538 | 0.003054 | 1 |

| | | | |
|---|---|---|---|
| Bet1 | -0.02538 | 0.343963 | 1 |
| Zfp771 | -0.02538 | 0.115036 | 1 |
| Armt1 | -0.0254 | 0.05856 | 1 |
| Pik3ca | -0.02542 | 0.576893 | 1 |
| Mtx2 | -0.02542 | 0.16488 | 1 |
| Senp5 | -0.02544 | 0.149568 | 1 |
| Eif4e3 | -0.02544 | 0.829253 | 1 |
| Alms1 | -0.02546 | 0.578746 | 1 |
| Zfp800 | -0.02546 | 0.416772 | 1 |
| Sec22b | -0.02548 | 0.097922 | 1 |
| Tnfsf13 | -0.0255 | 0.052612 | 1 |
| Hrh2 | -0.02554 | 0.067547 | 1 |
| Vps26a | -0.02555 | 0.396757 | 1 |
| Slc25a51 | -0.02556 | 0.129842 | 1 |
| Arid3a | -0.02557 | 0.735893 | 1 |
| Itgax | -0.02558 | 0.466473 | 1 |
| Tbc1d19 | -0.02561 | 0.458625 | 1 |
| Mocs1 | -0.02561 | 0.017702 | 1 |
| Atp13a1 | -0.02564 | 0.188821 | 1 |
| Champ1 | -0.02565 | 0.167729 | 1 |
| Prkce | -0.02566 | 0.444169 | 1 |
| Csf2ra | -0.02566 | 0.273048 | 1 |
| Fntb | -0.02568 | 0.019155 | 1 |
| Kmt2b | -0.02568 | 0.215477 | 1 |
| Zfp715 | -0.02568 | 0.724431 | 1 |
| Lmf2 | -0.02569 | 0.014988 | 1 |
| Cwc22 | -0.02571 | 0.59802 | 1 |
| Rtn4ip1 | -0.0258 | 0.291498 | 1 |
| Fubp1 | -0.0258 | 0.452952 | 1 |
| Megf11 | -0.02582 | 0.090968 | 1 |
| Setd7 | -0.02582 | 0.697187 | 1 |
| Zswim6 | -0.02583 | 0.530953 | 1 |
| Tlk2 | -0.02585 | 0.271054 | 1 |
| Sf3b3 | -0.02585 | 0.643442 | 1 |
| Tollip | -0.02586 | 0.59783 | 1 |
| Rbm43 | -0.02587 | 0.074208 | 1 |
| Akt2 | -0.02589 | 0.083422 | 1 |
| Ogdh | -0.02589 | 0.530003 | 1 |
| Ermp1 | -0.0259 | 0.136928 | 1 |
| Mtbp | -0.02591 | 0.062558 | 1 |
| Lsm14a | -0.02592 | 0.383594 | 1 |
| Mindy3 | -0.02597 | 0.201411 | 1 |
| Rcor3 | -0.02598 | 0.001909 | 1 |
| Vps33b | -0.02599 | 0.019698 | 1 |
| Katnbl1 | -0.02601 | 0.321706 | 1 |
| Zfp780b | -0.02601 | 0.286037 | 1 |
| Trim11 | -0.02604 | 0.737279 | 1 |
| Akap7 | -0.02604 | 0.603569 | 1 |

| | | | |
|---|---|---|---|
| Gm13684 | -0.02605 | 0.088384 | 1 |
| Rnf144b | -0.02606 | 0.91151 | 1 |
| Arrb1 | -0.02609 | 0.434056 | 1 |
| Parg | -0.02609 | 0.164605 | 1 |
| Mgat1 | -0.02614 | 0.506008 | 1 |
| Csad | -0.02614 | 0.461458 | 1 |
| Eif4e | -0.02614 | 0.302611 | 1 |
| Wwp2 | -0.02616 | 0.270958 | 1 |
| Pex11b | -0.02616 | 0.330062 | 1 |
| Zfp395 | -0.02619 | 0.021562 | 1 |
| Nop9 | -0.0262 | 0.20147 | 1 |
| Polk | -0.02621 | 0.804145 | 1 |
| Zfhx2 | -0.02622 | 0.013959 | 1 |
| Map2k4 | -0.02624 | 0.274361 | 1 |
| Ppp4r3b | -0.02624 | 0.143345 | 1 |
| Cemip2 | -0.02627 | 0.021733 | 1 |
| Bola3 | -0.02627 | 0.125227 | 1 |
| Kdm2b | -0.02627 | 0.902432 | 1 |
| 2900076A07Rik | -0.0263 | 0.133485 | 1 |
| Fam89a | -0.02632 | 0.114051 | 1 |
| Spock1 | -0.02632 | 0.123184 | 1 |
| Gm17018 | -0.02636 | 0.029491 | 1 |
| Akt1 | -0.02636 | 0.156492 | 1 |
| Atrn | -0.02638 | 0.153469 | 1 |
| Fam185a | -0.02642 | 0.081771 | 1 |
| Fam126a | -0.02642 | 0.329173 | 1 |
| Ube2b | -0.02642 | 0.609075 | 1 |
| Cnot10 | -0.02643 | 0.081808 | 1 |
| 3300002I08Rik | -0.02643 | 0.153094 | 1 |
| Psma5 | -0.02644 | 0.2977 | 1 |
| Zbtb18 | -0.02644 | 0.658366 | 1 |
| Bcl6 | -0.02645 | 0.375747 | 1 |
| Yipf1 | -0.02646 | 0.320002 | 1 |
| Lztr1 | -0.02647 | 0.044343 | 1 |
| Cdkn2aipnl | -0.02648 | 0.132209 | 1 |
| Zfp113 | -0.02648 | 0.097494 | 1 |
| Tsg101 | -0.02649 | 0.448646 | 1 |
| Siglec1 | -0.02649 | 0.376329 | 1 |
| Ruvbl2 | -0.0265 | 0.46309 | 1 |
| Tsacc | -0.02653 | 0.020421 | 1 |
| Zfp563 | -0.02653 | 0.056848 | 1 |
| Dnajc13 | -0.02653 | 0.272915 | 1 |
| Pelp1 | -0.02654 | 0.350171 | 1 |
| AI987944 | -0.02655 | 0.079512 | 1 |
| Kif13a | -0.02655 | 0.168397 | 1 |
| Mthfd2l | -0.02656 | 0.076279 | 1 |
| Nfkbil1 | -0.02657 | 0.334159 | 1 |
| Ppp1r9b | -0.02658 | 0.363309 | 1 |

| | | | |
|---|---|---|---|
| Dop1a | -0.0266 | 0.269803 | 1 |
| Hdgfl2 | -0.02664 | 0.064839 | 1 |
| Kdm3b | -0.02665 | 0.066279 | 1 |
| Baz1a | -0.02668 | 0.324527 | 1 |
| Uba1 | -0.0267 | 0.144157 | 1 |
| Uros | -0.0267 | 0.040071 | 1 |
| Rbm4b | -0.0267 | 0.791775 | 1 |
| Mbnl2 | -0.02671 | 0.186193 | 1 |
| Zkscan17 | -0.02673 | 0.568567 | 1 |
| Tmem127 | -0.02673 | 0.533104 | 1 |
| Cdk19 | -0.02675 | 0.350545 | 1 |
| Snx7 | -0.02679 | 0.087587 | 1 |
| D930016D06Rik | -0.0268 | 0.026435 | 1 |
| Arl6ip6 | -0.0268 | 0.98001 | 1 |
| Gtf2a1 | -0.02683 | 0.382604 | 1 |
| Ppip5k2 | -0.02684 | 0.057466 | 1 |
| Icosl | -0.02684 | 0.378836 | 1 |
| Myo6 | -0.02685 | 0.767804 | 1 |
| Cryl1 | -0.02686 | 0.083945 | 1 |
| Ywhag | -0.02686 | 0.434522 | 1 |
| Gm26936 | -0.02687 | 0.11392 | 1 |
| Washc4 | -0.02689 | 0.324775 | 1 |
| Fam207a | -0.02689 | 0.301336 | 1 |
| Pacs2 | -0.02693 | 0.255321 | 1 |
| Tpp2 | -0.02695 | 0.690758 | 1 |
| Ccdc122 | -0.02695 | 0.451863 | 1 |
| Zfp777 | -0.02696 | 0.0521 | 1 |
| Rala | -0.02696 | 0.587355 | 1 |
| Ap3s1 | -0.02696 | 0.330956 | 1 |
| Slc4a7 | -0.02699 | 0.451073 | 1 |
| Masp1 | -0.027 | 0.341831 | 1 |
| Wdr34 | -0.027 | 0.009753 | 1 |
| Gpr146 | -0.02701 | 0.542266 | 1 |
| Napb | -0.02702 | 0.007086 | 1 |
| Gab3 | -0.02709 | 0.214641 | 1 |
| 2610044O15Rik8 | -0.02709 | 0.095246 | 1 |
| Mxi1 | -0.0271 | 0.836001 | 1 |
| Smyd2 | -0.0271 | 0.121082 | 1 |
| Tbl1x | -0.0271 | 0.476966 | 1 |
| Rsad2 | -0.0271 | 0.101627 | 1 |
| Taf1 | -0.02711 | 0.026445 | 1 |
| Stk24 | -0.02711 | 0.364381 | 1 |
| Cd274 | -0.02714 | 0.102794 | 1 |
| Rnft1 | -0.02715 | 0.858719 | 1 |
| Glrx | -0.02716 | 0.111071 | 1 |
| Chd9 | -0.02716 | 0.233108 | 1 |
| Prkaa1 | -0.02719 | 0.3245 | 1 |
| Pms1 | -0.0272 | 0.07144 | 1 |

| | | | |
|---|---|---|---|
| Wdr77 | -0.0272 | 0.263413 | 1 |
| Zfp658 | -0.02721 | 0.786953 | 1 |
| Polh | -0.02723 | 0.243844 | 1 |
| Gemin7 | -0.02724 | 0.876891 | 1 |
| Slc39a11 | -0.02729 | 0.135964 | 1 |
| Traf6 | -0.0273 | 0.743289 | 1 |
| Dagla | -0.0273 | 0.129843 | 1 |
| Mapk8ip3 | -0.0273 | 0.021095 | 1 |
| Fam135a | -0.02732 | 0.119294 | 1 |
| B430306N03Rik | -0.02734 | 0.067676 | 1 |
| Nup188 | -0.02735 | 0.087457 | 1 |
| Mxd1 | -0.02736 | 0.121934 | 1 |
| Vipas39 | -0.02739 | 0.057305 | 1 |
| Mink1 | -0.0274 | 0.066561 | 1 |
| Gm26724 | -0.0274 | 0.04089 | 1 |
| Hpgds | -0.0274 | 0.351857 | 1 |
| Clpx | -0.02741 | 0.420096 | 1 |
| Gm7480 | -0.02744 | 0.016783 | 1 |
| Gid4 | -0.02745 | 0.270569 | 1 |
| Slc45a4 | -0.02746 | 0.249169 | 1 |
| Letm1 | -0.02747 | 0.180298 | 1 |
| Mical3 | -0.02749 | 0.524664 | 1 |
| Sesn2 | -0.02749 | 0.088683 | 1 |
| Mss51 | -0.0275 | 0.011928 | 1 |
| Recql5 | -0.0275 | 0.096072 | 1 |
| Ipo8 | -0.02753 | 0.419753 | 1 |
| Tbc1d31 | -0.02754 | 0.018389 | 1 |
| Mrps6 | -0.02756 | 0.164268 | 1 |
| Rbm10 | -0.02758 | 0.081921 | 1 |
| Tmem120b | -0.02758 | 0.516284 | 1 |
| Eif4g1 | -0.02759 | 0.406863 | 1 |
| Pms2 | -0.0276 | 0.183771 | 1 |
| Sacm1l | -0.02761 | 0.265049 | 1 |
| Arid4b | -0.02761 | 0.724899 | 1 |
| Ttc17 | -0.02763 | 0.116132 | 1 |
| Fam210a | -0.02764 | 0.472838 | 1 |
| Nae1 | -0.02764 | 0.403628 | 1 |
| Snrk | -0.02766 | 0.184878 | 1 |
| Sord | -0.02767 | 0.135185 | 1 |
| Calhm2 | -0.0277 | 0.389185 | 1 |
| Irf5 | -0.02772 | 0.542058 | 1 |
| Uhmk1 | -0.02773 | 0.884052 | 1 |
| Sin3a | -0.02773 | 0.774927 | 1 |
| Sltm | -0.02774 | 0.375063 | 1 |
| Cacul1 | -0.02775 | 0.281102 | 1 |
| Sprtn | -0.02776 | 0.360384 | 1 |
| Atg5 | -0.02778 | 0.530814 | 1 |
| Cdk5rap1 | -0.02779 | 0.342612 | 1 |

| | | | |
|---|---|---|---|
| Pter | -0.02781 | 0.416339 | 1 |
| Gsap | -0.02784 | 0.117985 | 1 |
| Atg16l2 | -0.02785 | 0.514502 | 1 |
| Scaf1 | -0.02786 | 0.203913 | 1 |
| Igf2bp3 | -0.02788 | 0.36481 | 1 |
| Pik3c3 | -0.02788 | 0.218734 | 1 |
| Brd1 | -0.02788 | 0.397463 | 1 |
| Dnajc16 | -0.02789 | 0.556043 | 1 |
| Ppp1r12a | -0.02791 | 0.14627 | 1 |
| Slc6a6 | -0.02792 | 0.57732 | 1 |
| Tlr1 | -0.02794 | 0.344648 | 1 |
| Fam129a | -0.02798 | 0.465243 | 1 |
| Tubb4b | -0.02803 | 0.227177 | 1 |
| Angel2 | -0.02803 | 0.620622 | 1 |
| Vps53 | -0.02804 | 0.717252 | 1 |
| Gm15764 | -0.02805 | 0.251716 | 1 |
| Zfp385a | -0.02805 | 0.404992 | 1 |
| Bpnt1 | -0.02806 | 0.117166 | 1 |
| Txlng | -0.02806 | 0.038405 | 1 |
| Pgm3 | -0.02807 | 0.028187 | 1 |
| Ulk2 | -0.02808 | 0.22393 | 1 |
| Ptpra | -0.02808 | 0.189507 | 1 |
| Llph | -0.02808 | 0.9495 | 1 |
| B3glct | -0.02808 | 0.789673 | 1 |
| Zfp746 | -0.0281 | 0.052809 | 1 |
| G2e3 | -0.02812 | 0.089221 | 1 |
| Lpcat1 | -0.02812 | 0.350454 | 1 |
| Abca7 | -0.02812 | 0.47407 | 1 |
| Pccb | -0.02812 | 0.147339 | 1 |
| Tbc1d22a | -0.02814 | 0.088864 | 1 |
| Gm34408 | -0.02816 | 0.004615 | 1 |
| Rin3 | -0.02816 | 0.172802 | 1 |
| Pskh1 | -0.02818 | 0.051731 | 1 |
| Casp12 | -0.02818 | 0.052809 | 1 |
| 2510009E07Rik | -0.02819 | 0.467434 | 1 |
| Rmdn3 | -0.0282 | 0.103956 | 1 |
| Zfp810 | -0.02824 | 0.706494 | 1 |
| Ddx39b | -0.02825 | 0.350536 | 1 |
| Dock7 | -0.02825 | 0.960192 | 1 |
| Cc2d1b | -0.02828 | 0.494421 | 1 |
| Orai1 | -0.02828 | 0.626602 | 1 |
| Fbxl18 | -0.02828 | 0.002127 | 1 |
| Cnot1 | -0.02831 | 0.209028 | 1 |
| Recql | -0.02834 | 0.116249 | 1 |
| Ubfd1 | -0.02837 | 0.45907 | 1 |
| Ssbp1 | -0.02837 | 0.22994 | 1 |
| Rab8a | -0.02838 | 0.428491 | 1 |
| Tmf1 | -0.02838 | 0.945366 | 1 |

| | | | |
|---|---|---|---|
| Poli | -0.02839 | 0.001398 | 1 |
| Pias2 | -0.02839 | 0.728432 | 1 |
| Gm10130 | -0.0284 | 0.029951 | 1 |
| Wdr37 | -0.02841 | 0.546525 | 1 |
| Tab3 | -0.02846 | 0.296404 | 1 |
| Scamp1 | -0.02848 | 0.592571 | 1 |
| Gnl1 | -0.02848 | 0.012506 | 1 |
| Piezo1 | -0.02849 | 0.342703 | 1 |
| Atp10a | -0.0285 | 0.169006 | 1 |
| Szrd1 | -0.0285 | 0.437166 | 1 |
| Psmg1 | -0.02852 | 0.520705 | 1 |
| Otulinl | -0.02854 | 0.090675 | 1 |
| Ugcg | -0.02855 | 0.172518 | 1 |
| Gatad2a | -0.02857 | 0.74525 | 1 |
| Dcaf1 | -0.0286 | 0.735298 | 1 |
| Prmt5 | -0.02862 | 0.04667 | 1 |
| Osbpl1a | -0.02863 | 0.706876 | 1 |
| Cdk5rap2 | -0.02865 | 0.275737 | 1 |
| Enoph1 | -0.02865 | 0.033879 | 1 |
| Epn1 | -0.02866 | 0.112726 | 1 |
| Capg | -0.02867 | 0.49638 | 1 |
| Trip12 | -0.02868 | 0.521044 | 1 |
| Gtf3c2 | -0.02873 | 0.762156 | 1 |
| Pptc7 | -0.02875 | 0.102051 | 1 |
| Ubl7 | -0.02876 | 0.010796 | 1 |
| Tshz1 | -0.02876 | 0.349435 | 1 |
| Lrrc8a | -0.02877 | 0.263323 | 1 |
| Erap1 | -0.02877 | 0.415973 | 1 |
| Cdk13 | -0.02877 | 0.161135 | 1 |
| Slc25a45 | -0.02879 | 0.109095 | 1 |
| Dync2h1 | -0.0288 | 0.380442 | 1 |
| Carmil1 | -0.0288 | 0.652174 | 1 |
| Eif4h | -0.02881 | 0.247476 | 1 |
| Ugp2 | -0.02883 | 0.164322 | 1 |
| Akap10 | -0.02883 | 0.412846 | 1 |
| Gpr35 | -0.02885 | 0.876927 | 1 |
| Prdm2 | -0.02885 | 0.667999 | 1 |
| Zfp142 | -0.02887 | 0.251705 | 1 |
| Odf2l | -0.02887 | 0.036394 | 1 |
| Ttc13 | -0.02887 | 0.635844 | 1 |
| Agps | -0.02888 | 0.326295 | 1 |
| Med14 | -0.02889 | 0.222879 | 1 |
| Fnbp4 | -0.02893 | 0.434309 | 1 |
| Tdrd3 | -0.02893 | 0.369287 | 1 |
| Tab1 | -0.02893 | 0.302153 | 1 |
| Glmn | -0.02896 | 0.098268 | 1 |
| Vps16 | -0.02896 | 0.238153 | 1 |
| Lpar6 | -0.02898 | 0.321137 | 1 |

| | | | |
|---|---|---|---|
| Os9 | -0.02899 | 0.873664 | 1 |
| Coro1a | -0.029 | 0.512932 | 1 |
| Dnaja2 | -0.02901 | 0.642823 | 1 |
| Pank2 | -0.02901 | 0.18022 | 1 |
| Gm21057 | -0.02902 | 0.073668 | 1 |
| Gm15965 | -0.02904 | 0.019889 | 1 |
| Zfp280d | -0.02907 | 0.223379 | 1 |
| Dpysl2 | -0.02907 | 0.083565 | 1 |
| Ist1 | -0.02908 | 0.087604 | 1 |
| Shb | -0.0291 | 0.30908 | 1 |
| Gm13212 | -0.02911 | 0.032922 | 1 |
| Cish | -0.02912 | 4.87E-05 | 1 |
| Ms4a4c | -0.02912 | 0.489199 | 1 |
| Gm16091 | -0.02913 | 0.023623 | 1 |
| Mtmr12 | -0.02914 | 0.476494 | 1 |
| Rtn1 | -0.02916 | 0.57127 | 1 |
| Cryba4 | -0.02916 | 0.560682 | 1 |
| Smim13 | -0.02917 | 0.111529 | 1 |
| Arl6ip5 | -0.02917 | 0.802717 | 1 |
| Bahcc1 | -0.02918 | 0.309826 | 1 |
| Prdm15 | -0.02918 | 0.002923 | 1 |
| Zc3h7a | -0.02918 | 0.125179 | 1 |
| Cebpg | -0.02919 | 0.417111 | 1 |
| Uckl1 | -0.02922 | 0.008362 | 1 |
| Ramac | -0.02923 | 0.584999 | 1 |
| Trappc10 | -0.02925 | 0.130558 | 1 |
| Relch | -0.02928 | 0.418782 | 1 |
| Sf3b6 | -0.02929 | 0.304683 | 1 |
| Acp6 | -0.02929 | 0.536414 | 1 |
| Blm | -0.02929 | 0.331177 | 1 |
| Il7r | -0.02933 | 0.283338 | 1 |
| Rlf | -0.02933 | 0.377741 | 1 |
| Tgfbrap1 | -0.02934 | 0.343197 | 1 |
| Ampd3 | -0.02934 | 0.437844 | 1 |
| Uchl3 | -0.02934 | 0.170337 | 1 |
| Nfrkb | -0.02934 | 0.208919 | 1 |
| Ccl7 | -0.02936 | 0.002694 | 1 |
| Klc1 | -0.02939 | 0.089503 | 1 |
| 4930402H24Rik | -0.0294 | 0.226716 | 1 |
| Hip1 | -0.0294 | 0.504453 | 1 |
| Agpat1 | -0.02941 | 0.075888 | 1 |
| P2rx1 | -0.02942 | 0.006308 | 1 |
| Agtpbp1 | -0.02944 | 0.190217 | 1 |
| Mcu | -0.02944 | 0.09311 | 1 |
| Ap3m2 | -0.02944 | 0.142275 | 1 |
| Hacl1 | -0.02947 | 0.050329 | 1 |
| Sik2 | -0.02948 | 0.412748 | 1 |
| Ypel5 | -0.02948 | 0.373444 | 1 |

| | | | |
|---|---|---|---|
| Ralgapb | -0.02949 | 0.328125 | 1 |
| Aldh2 | -0.02951 | 0.404875 | 1 |
| Trip6 | -0.02951 | 0.026826 | 1 |
| Rnf181 | -0.02953 | 0.207643 | 1 |
| Daxx | -0.02953 | 0.063322 | 1 |
| Gbp9 | -0.02953 | 0.328805 | 1 |
| Nfe2l2 | -0.02956 | 0.176511 | 1 |
| Eps8 | -0.02956 | 0.025988 | 1 |
| Msi2 | -0.02956 | 0.6157 | 1 |
| Ppp1cc | -0.02956 | 0.488146 | 1 |
| Bmpr1a | -0.02957 | 0.41096 | 1 |
| Dcaf17 | -0.02957 | 0.514079 | 1 |
| Eef2k | -0.0296 | 0.27218 | 1 |
| Nek9 | -0.0296 | 0.437941 | 1 |
| Gm6034 | -0.02961 | 0.122925 | 1 |
| Ly6a | -0.02963 | 0.460234 | 1 |
| Srp72 | -0.02963 | 0.327872 | 1 |
| Ncoa3 | -0.02963 | 0.318656 | 1 |
| Tsga10 | -0.02963 | 0.98897 | 1 |
| E130307A14Rik | -0.02964 | 0.922105 | 1 |
| Ptrh2 | -0.02967 | 0.134784 | 1 |
| Ascc1 | -0.02968 | 0.226233 | 1 |
| Tbc1d24 | -0.0297 | 0.186636 | 1 |
| Ythdf1 | -0.0297 | 0.116524 | 1 |
| Znrf1 | -0.02971 | 0.49088 | 1 |
| Rbm14 | -0.02972 | 0.044287 | 1 |
| Riok2 | -0.02974 | 0.898067 | 1 |
| Rilpl1 | -0.02976 | 0.816886 | 1 |
| 6330562C20Rik | -0.02979 | 0.088683 | 1 |
| Tmem242 | -0.02979 | 0.128395 | 1 |
| Tgds | -0.02981 | 0.118438 | 1 |
| Tjp1 | -0.02983 | 0.118484 | 1 |
| Gm49662 | -0.02985 | 0.02261 | 1 |
| Ccdc85c | -0.02985 | 0.00849 | 1 |
| Pram1 | -0.02986 | 0.002201 | 1 |
| Elf1 | -0.02987 | 0.073633 | 1 |
| Acsl5 | -0.02989 | 0.355146 | 1 |
| Slc39a9 | -0.02989 | 0.236453 | 1 |
| Plekha2 | -0.02992 | 0.267261 | 1 |
| Trip11 | -0.02992 | 0.727047 | 1 |
| Sec22a | -0.02992 | 0.805758 | 1 |
| Fam20b | -0.02995 | 0.069948 | 1 |
| Enox2 | -0.02995 | 0.009901 | 1 |
| Elac1 | -0.02996 | 0.14289 | 1 |
| Tnfsf10 | -0.02997 | 0.023884 | 1 |
| Skp2 | -0.02998 | 0.238838 | 1 |
| Gm14966 | -0.02999 | 0.023959 | 1 |
| Arl2 | -0.03 | 0.659754 | 1 |

| | | | |
|---|---|---|---|
| Reep4 | -0.03002 | 0.19777 | 1 |
| Plekhb2 | -0.03003 | 0.165357 | 1 |
| C2cd5 | -0.03009 | 0.54136 | 1 |
| Gm26827 | -0.0301 | 0.840588 | 1 |
| Cpne1 | -0.0301 | 0.166217 | 1 |
| Ppp1r9a | -0.03012 | 0.630391 | 1 |
| Lrrfip1 | -0.03013 | 0.172808 | 1 |
| Psme2b | -0.03014 | 0.289194 | 1 |
| Abcd1 | -0.03015 | 0.86034 | 1 |
| Dcun1d3 | -0.03015 | 0.164517 | 1 |
| Pold3 | -0.03018 | 0.341892 | 1 |
| Asb8 | -0.03019 | 0.264078 | 1 |
| Ireb2 | -0.03022 | 0.347908 | 1 |
| Cln3 | -0.03023 | 0.210383 | 1 |
| Nmrk1 | -0.03023 | 0.014399 | 1 |
| Orc3 | -0.03028 | 0.085413 | 1 |
| Stau1 | -0.03029 | 0.198048 | 1 |
| Phf1 | -0.0303 | 0.014804 | 1 |
| Bcl2l11 | -0.03031 | 0.895841 | 1 |
| Glt8d1 | -0.03031 | 0.275548 | 1 |
| Raet1e | -0.03032 | 0.15918 | 1 |
| Arl14ep | -0.03032 | 0.6252 | 1 |
| Nfkb2 | -0.03032 | 0.600976 | 1 |
| Ogfr | -0.03033 | 0.957615 | 1 |
| Wdr12 | -0.03033 | 0.046353 | 1 |
| Zbtb11 | -0.03035 | 0.097088 | 1 |
| Trap1 | -0.03036 | 0.065338 | 1 |
| Sos2 | -0.03038 | 0.727631 | 1 |
| Lmbr1 | -0.03042 | 0.115292 | 1 |
| Rab6a | -0.03043 | 0.780984 | 1 |
| Arfip2 | -0.03045 | 0.037505 | 1 |
| Hars | -0.03052 | 0.018924 | 1 |
| Ccl25 | -0.03054 | 0.0493 | 1 |
| Tm9sf3 | -0.03057 | 0.310703 | 1 |
| Zfp644 | -0.03057 | 0.725347 | 1 |
| Rcan1 | -0.0306 | 0.172088 | 1 |
| Relb | -0.03063 | 0.142434 | 1 |
| Brd9 | -0.03064 | 0.507646 | 1 |
| Ccdc186 | -0.03066 | 0.560857 | 1 |
| Eif2ak2 | -0.0307 | 0.181342 | 1 |
| Gm11944 | -0.03072 | 0.017871 | 1 |
| Rassf5 | -0.03072 | 0.206933 | 1 |
| Ints9 | -0.03073 | 0.212737 | 1 |
| Atg4b | -0.03074 | 0.525207 | 1 |
| Itfg1 | -0.03074 | 0.686292 | 1 |
| Kantr | -0.03074 | 0.136787 | 1 |
| Prpf38b | -0.03076 | 0.346161 | 1 |
| Det1 | -0.03076 | 0.101903 | 1 |

| | | | |
|---|---|---|---|
| Rnf220 | -0.03079 | 0.13434 | 1 |
| Mtm1 | -0.0308 | 0.502784 | 1 |
| Dlg4 | -0.03081 | 0.041698 | 1 |
| Map3k8 | -0.03081 | 0.269695 | 1 |
| Utp6 | -0.03082 | 0.693606 | 1 |
| Ext1 | -0.03084 | 0.295513 | 1 |
| AW554918 | -0.03085 | 0.578711 | 1 |
| Zcchc14 | -0.03086 | 0.057435 | 1 |
| Mex3c | -0.03087 | 0.179935 | 1 |
| Pmm2 | -0.03088 | 0.118182 | 1 |
| Ppp1r18 | -0.03088 | 0.990597 | 1 |
| Fam118b | -0.03088 | 0.126589 | 1 |
| Zfp976 | -0.03089 | 0.034069 | 1 |
| Tigar | -0.03089 | 0.046058 | 1 |
| Cpne3 | -0.03091 | 0.208799 | 1 |
| Map3k20 | -0.03091 | 0.420114 | 1 |
| Rnf6 | -0.03092 | 0.166951 | 1 |
| 1700037C18Rik | -0.03092 | 0.382132 | 1 |
| Adck1 | -0.03096 | 0.052304 | 1 |
| Tti1 | -0.03099 | 0.10805 | 1 |
| Pcnx4 | -0.03101 | 0.031283 | 1 |
| Rbm4 | -0.03102 | 0.516708 | 1 |
| Foxj2 | -0.03103 | 0.164538 | 1 |
| Akna | -0.03107 | 0.412779 | 1 |
| Rars2 | -0.03109 | 0.176605 | 1 |
| Fam98c | -0.03112 | 0.043775 | 1 |
| Nlrp1b | -0.03113 | 0.05767 | 1 |
| Ptpre | -0.03113 | 0.297361 | 1 |
| Ttc9c | -0.03114 | 0.100551 | 1 |
| Btbd1 | -0.03114 | 0.488378 | 1 |
| Ttc3 | -0.03115 | 0.966768 | 1 |
| Gemin8 | -0.03117 | 0.266719 | 1 |
| Taok2 | -0.03118 | 0.127374 | 1 |
| Taf2 | -0.03119 | 0.202985 | 1 |
| Nme7 | -0.0312 | 0.009703 | 1 |
| Gm49980 | -0.0312 | 0.001504 | 1 |
| Adat1 | -0.03122 | 0.013762 | 1 |
| Nt5dc1 | -0.03125 | 0.571503 | 1 |
| Utp25 | -0.03128 | 0.0726 | 1 |
| Gm50237 | -0.03131 | 0.031838 | 1 |
| Pmvk | -0.03133 | 0.064477 | 1 |
| Gpatch8 | -0.03135 | 0.520206 | 1 |
| Sh2b1 | -0.03137 | 0.230776 | 1 |
| Adck2 | -0.03139 | 0.104347 | 1 |
| Stxbp2 | -0.03145 | 0.187189 | 1 |
| Pi4k2a | -0.03145 | 0.884103 | 1 |
| Plekhm1 | -0.03146 | 0.034077 | 1 |
| Pde6d | -0.03148 | 0.145483 | 1 |

| | | | |
|---|---|---|---|
| Nsun3 | -0.03148 | 0.290616 | 1 |
| Ncf1 | -0.03153 | 0.412244 | 1 |
| Ano6 | -0.03154 | 0.849887 | 1 |
| Tfdp2 | -0.03155 | 0.140484 | 1 |
| Cbx5 | -0.03156 | 0.405037 | 1 |
| Cstf3 | -0.03159 | 0.20213 | 1 |
| Snx4 | -0.03161 | 0.404087 | 1 |
| Tmem181a | -0.03163 | 0.83768 | 1 |
| Prpf4 | -0.03164 | 0.063068 | 1 |
| Ube2k | -0.03166 | 0.152771 | 1 |
| Oga | -0.03167 | 0.584959 | 1 |
| Zbtb26 | -0.03167 | 0.007788 | 1 |
| D1Ertd622e | -0.03168 | 0.209971 | 1 |
| Hmg20a | -0.03171 | 0.079396 | 1 |
| Mecr | -0.03171 | 0.776217 | 1 |
| Gpalpp1 | -0.03174 | 0.327404 | 1 |
| Cog4 | -0.03174 | 0.183318 | 1 |
| Xrcc1 | -0.03174 | 0.017489 | 1 |
| Speg | -0.03175 | 0.477033 | 1 |
| Coa5 | -0.03177 | 0.046624 | 1 |
| Sirt6 | -0.03177 | 0.005615 | 1 |
| Prdm11 | -0.03178 | 0.155031 | 1 |
| Naa15 | -0.03179 | 0.664087 | 1 |
| Irf7 | -0.0318 | 0.007799 | 1 |
| Mettl1 | -0.03181 | 0.290734 | 1 |
| Ythdf3 | -0.03183 | 0.312781 | 1 |
| Prmt1 | -0.03185 | 0.017565 | 1 |
| Zzz3 | -0.03185 | 0.133571 | 1 |
| Rbbp8 | -0.03187 | 0.499032 | 1 |
| Pfkl | -0.03191 | 0.077007 | 1 |
| Tsen34 | -0.03192 | 0.23639 | 1 |
| Sntb2 | -0.03192 | 0.370925 | 1 |
| Yars | -0.03194 | 0.293937 | 1 |
| Ulk4 | -0.03195 | 0.763167 | 1 |
| Depdc5 | -0.03196 | 0.259679 | 1 |
| Scarb2 | -0.03197 | 0.352525 | 1 |
| Aopep | -0.03198 | 0.080091 | 1 |
| Mdm2 | -0.03199 | 0.381162 | 1 |
| Usp33 | -0.032 | 0.186259 | 1 |
| Ikzf2 | -0.032 | 0.167814 | 1 |
| Gm10848 | -0.032 | 0.984002 | 1 |
| Polr3a | -0.03201 | 0.380788 | 1 |
| Rint1 | -0.03201 | 0.116189 | 1 |
| Abhd17a | -0.03204 | 0.181633 | 1 |
| Slc35b3 | -0.03205 | 0.373511 | 1 |
| Apobr | -0.03207 | 0.363065 | 1 |
| Siglecf | -0.03207 | 0.191464 | 1 |
| Vrk3 | -0.03208 | 0.083986 | 1 |

| | | | |
|---|---|---|---|
| Rsph3b | -0.03212 | 0.012688 | 1 |
| Arfgef2 | -0.03212 | 0.054513 | 1 |
| Styx | -0.03213 | 0.129476 | 1 |
| Msra | -0.03214 | 0.92434 | 1 |
| Gm46218 | -0.03214 | 0.330823 | 1 |
| Klhdc10 | -0.03216 | 0.089875 | 1 |
| Parp2 | -0.03216 | 0.259005 | 1 |
| Uhrf2 | -0.03217 | 0.068972 | 1 |
| Zfp938 | -0.03219 | 0.001092 | 1 |
| Fan1 | -0.03222 | 0.354557 | 1 |
| Rabep1 | -0.03222 | 0.114705 | 1 |
| Stard7 | -0.03223 | 0.283463 | 1 |
| Tstd2 | -0.03223 | 0.054444 | 1 |
| Ikzf5 | -0.03224 | 0.04938 | 1 |
| Pstpip1 | -0.03226 | 0.045791 | 1 |
| Zfp518a | -0.03229 | 0.180786 | 1 |
| Mob2 | -0.03229 | 0.351682 | 1 |
| Usp30 | -0.03231 | 0.012268 | 1 |
| Extl2 | -0.03231 | 0.108852 | 1 |
| Sptbn1 | -0.03232 | 0.313884 | 1 |
| Usp14 | -0.03233 | 0.50621 | 1 |
| Eif1b | -0.03234 | 0.481786 | 1 |
| Ptpn6 | -0.03236 | 0.37033 | 1 |
| Pdxk | -0.03236 | 0.010007 | 1 |
| Nr6a1 | -0.03237 | 0.7993 | 1 |
| Rnf135 | -0.03238 | 0.106413 | 1 |
| Capn1 | -0.03239 | 0.012001 | 1 |
| Atp6v1b2 | -0.03239 | 0.548949 | 1 |
| Wdr20 | -0.0324 | 0.316315 | 1 |
| Rbm33 | -0.03241 | 0.830877 | 1 |
| Ino80 | -0.03241 | 0.268136 | 1 |
| Rufy1 | -0.03242 | 0.837531 | 1 |
| Cyp4f18 | -0.03242 | 0.307058 | 1 |
| Map3k1 | -0.03244 | 0.291481 | 1 |
| Cul3 | -0.03244 | 0.579422 | 1 |
| Supt20 | -0.03246 | 0.953118 | 1 |
| Mpv17 | -0.03246 | 0.023877 | 1 |
| Ncoa7 | -0.03247 | 0.02719 | 1 |
| Trip4 | -0.03247 | 0.202499 | 1 |
| Araf | -0.03247 | 0.474642 | 1 |
| Tor1aip2 | -0.03248 | 0.341348 | 1 |
| Mia2 | -0.03248 | 0.381844 | 1 |
| Gmip | -0.03249 | 0.192176 | 1 |
| Zfp809 | -0.0325 | 0.11719 | 1 |
| Malat1 | -0.0325 | 0.09112 | 1 |
| Dlat | -0.03251 | 0.261498 | 1 |
| Lsm2 | -0.03253 | 0.040448 | 1 |
| Tmem87b | -0.03255 | 0.261373 | 1 |

| | | | |
|---|---|---|---|
| Slc25a22 | -0.0326 | 0.007501 | 1 |
| Gm4070 | -0.03262 | 0.128721 | 1 |
| Mpv17l | -0.03262 | 0.395673 | 1 |
| Slx4ip | -0.03264 | 0.345114 | 1 |
| Poldip3 | -0.03267 | 0.13303 | 1 |
| Cttnbp2nl | -0.0327 | 0.285016 | 1 |
| Uap1 | -0.03271 | 0.11979 | 1 |
| Knop1 | -0.03273 | 0.212279 | 1 |
| R3hdm1 | -0.03273 | 0.113379 | 1 |
| Tmem71 | -0.03273 | 0.037227 | 1 |
| Eri3 | -0.03274 | 0.066236 | 1 |
| Mtmr6 | -0.03274 | 0.038387 | 1 |
| Ten1 | -0.03276 | 0.161486 | 1 |
| Prmt9 | -0.03277 | 0.090606 | 1 |
| Chil1 | -0.03278 | 0.001908 | 1 |
| Zdhhc14 | -0.03278 | 0.913353 | 1 |
| Iqce | -0.03279 | 0.001744 | 1 |
| Kctd21 | -0.03279 | 0.012264 | 1 |
| Tecpr1 | -0.03281 | 0.240579 | 1 |
| E530011L22Rik | -0.03281 | 0.031428 | 1 |
| Snrnp48 | -0.03281 | 0.366391 | 1 |
| Runx1 | -0.03284 | 0.08253 | 1 |
| Babam2 | -0.03285 | 0.06902 | 1 |
| Ccdc66 | -0.03286 | 0.013081 | 1 |
| Fhod1 | -0.03287 | 0.528927 | 1 |
| Rasa4 | -0.03287 | 0.320567 | 1 |
| Kdm1a | -0.03289 | 0.483159 | 1 |
| Zc3h18 | -0.03289 | 0.208727 | 1 |
| Oas1a | -0.03293 | 0.08498 | 1 |
| Irf3 | -0.03293 | 0.121935 | 1 |
| Acox1 | -0.03293 | 0.066643 | 1 |
| Brwd3 | -0.03293 | 0.326445 | 1 |
| Brd7 | -0.03295 | 0.362821 | 1 |
| Ythdc2 | -0.03296 | 0.205895 | 1 |
| Papolg | -0.03299 | 0.114022 | 1 |
| Npepl1 | -0.03302 | 0.091184 | 1 |
| Zfp277 | -0.03303 | 0.318152 | 1 |
| Pex14 | -0.03304 | 0.092996 | 1 |
| Nmt2 | -0.03307 | 0.916359 | 1 |
| Incenp | -0.03308 | 0.069593 | 1 |
| Ngrn | -0.03309 | 0.020666 | 1 |
| Larp4 | -0.03309 | 0.264883 | 1 |
| Usp20 | -0.03309 | 0.022368 | 1 |
| Spata13 | -0.0331 | 0.161201 | 1 |
| Pcgf5 | -0.03311 | 0.655703 | 1 |
| Gm26520 | -0.03313 | 0.279334 | 1 |
| Heatr6 | -0.03314 | 0.103673 | 1 |
| Gm12353 | -0.03314 | 0.014142 | 1 |

| | | | |
|---|---|---|---|
| R3hdm2 | -0.03314 | 0.181324 | 1 |
| Apba1 | -0.03319 | 0.248415 | 1 |
| Dtx2 | -0.0332 | 0.170403 | 1 |
| Tor1b | -0.03321 | 0.127531 | 1 |
| Slc30a6 | -0.03323 | 0.064628 | 1 |
| Gnl3l | -0.03324 | 0.008294 | 1 |
| Gm45894 | -0.03325 | 0.001401 | 1 |
| Cyb5r4 | -0.03326 | 0.142004 | 1 |
| Vars | -0.03326 | 0.550819 | 1 |
| Ift74 | -0.0333 | 0.098216 | 1 |
| Tbk1 | -0.03332 | 0.468257 | 1 |
| Agap3 | -0.03332 | 0.075013 | 1 |
| Clpb | -0.03335 | 0.339265 | 1 |
| Ctdspl2 | -0.03337 | 0.514025 | 1 |
| Tmem135 | -0.0334 | 0.35125 | 1 |
| C330018D20Rik | -0.0334 | 0.0402 | 1 |
| Cnnm2 | -0.03342 | 0.941269 | 1 |
| Gpc2 | -0.03343 | 0.037958 | 1 |
| Rab3gap1 | -0.03346 | 0.334847 | 1 |
| Timmdc1 | -0.03348 | 0.063155 | 1 |
| Arl5c | -0.03349 | 0.168028 | 1 |
| Klhdc1 | -0.0335 | 0.008672 | 1 |
| Parp10 | -0.03353 | 0.016797 | 1 |
| Frmd8 | -0.03354 | 0.133188 | 1 |
| Tjp2 | -0.03354 | 0.058652 | 1 |
| Fbxl4 | -0.03355 | 0.460057 | 1 |
| Rbm6 | -0.03357 | 0.098507 | 1 |
| Zfp120 | -0.0336 | 0.348373 | 1 |
| Rnf34 | -0.03363 | 0.546087 | 1 |
| Ptpn11 | -0.03364 | 0.281108 | 1 |
| Nfat5 | -0.03365 | 0.882566 | 1 |
| Zfp579 | -0.03367 | 0.139939 | 1 |
| Coro7 | -0.03367 | 0.243697 | 1 |
| Rpgr | -0.03367 | 0.004026 | 1 |
| Slc7a7 | -0.03367 | 0.163083 | 1 |
| Etv1 | -0.03368 | 0.360726 | 1 |
| Gm4924 | -0.03369 | 0.365277 | 1 |
| 2610037D02Rik | -0.03371 | 0.244761 | 1 |
| Hdac4 | -0.03374 | 0.083193 | 1 |
| Chfr | -0.03376 | 0.116615 | 1 |
| Fbxl19 | -0.03376 | 0.003893 | 1 |
| Apopt1 | -0.03377 | 0.075195 | 1 |
| Dnajc24 | -0.03379 | 0.039618 | 1 |
| Cep192 | -0.0338 | 0.505383 | 1 |
| Anapc1 | -0.03381 | 0.124177 | 1 |
| Dis3 | -0.03386 | 0.358549 | 1 |
| Nup160 | -0.03388 | 0.039495 | 1 |
| Fam45a | -0.03388 | 0.445451 | 1 |

| | | | |
|---|---|---|---|
| Flad1 | -0.0339 | 0.274686 | 1 |
| 4930444A19Rik | -0.0339 | 0.024608 | 1 |
| Rfx7 | -0.03392 | 0.375933 | 1 |
| Uso1 | -0.03392 | 0.256075 | 1 |
| Uba3 | -0.03394 | 0.049061 | 1 |
| Qtrt1 | -0.03395 | 0.790164 | 1 |
| D10Wsu102e | -0.03396 | 0.100695 | 1 |
| Rnf10 | -0.03396 | 0.271184 | 1 |
| Psma2 | -0.03399 | 0.22834 | 1 |
| Anxa7 | -0.03402 | 0.235875 | 1 |
| Fbxl3 | -0.03402 | 0.98244 | 1 |
| Ccnk | -0.03403 | 0.072632 | 1 |
| Agpat3 | -0.03403 | 0.786245 | 1 |
| Casp7 | -0.03404 | 0.633125 | 1 |
| Slc11a2 | -0.03406 | 0.05772 | 1 |
| Sec23ip | -0.03407 | 0.024295 | 1 |
| Nbr1 | -0.03411 | 0.088676 | 1 |
| Cmklr1 | -0.03413 | 0.198919 | 1 |
| Armc8 | -0.03416 | 0.036081 | 1 |
| Phf3 | -0.03416 | 0.932244 | 1 |
| Fam114a1 | -0.03417 | 0.060949 | 1 |
| Pspc1 | -0.03418 | 0.717075 | 1 |
| Anxa11 | -0.03419 | 0.008356 | 1 |
| Capn7 | -0.03421 | 0.242927 | 1 |
| Pan3 | -0.03421 | 0.301726 | 1 |
| Zfp609 | -0.03423 | 0.374774 | 1 |
| Dennd4c | -0.03423 | 0.285852 | 1 |
| Snx25 | -0.03424 | 0.676184 | 1 |
| Ndufa10 | -0.03426 | 0.828159 | 1 |
| Kbtbd2 | -0.03426 | 0.704523 | 1 |
| Itgb3bp | -0.03428 | 0.066837 | 1 |
| Tmem106b | -0.03428 | 0.269695 | 1 |
| Tfe3 | -0.03428 | 0.430877 | 1 |
| Rfwd3 | -0.0343 | 0.167726 | 1 |
| Lats1 | -0.03431 | 0.277129 | 1 |
| Megf9 | -0.03433 | 0.020657 | 1 |
| Sdhb | -0.03441 | 0.671408 | 1 |
| I830077J02Rik | -0.03442 | 0.046728 | 1 |
| Lima1 | -0.03444 | 0.206304 | 1 |
| Gpr180 | -0.03444 | 0.245123 | 1 |
| Kdm4c | -0.03445 | 0.349644 | 1 |
| Zfp281 | -0.03447 | 0.006374 | 1 |
| Camkmt | -0.03448 | 0.349603 | 1 |
| Map4k3 | -0.03449 | 0.247776 | 1 |
| Ppp2cb | -0.0345 | 0.039269 | 1 |
| Ap1ar | -0.03452 | 0.310383 | 1 |
| Ice1 | -0.03455 | 0.367855 | 1 |
| Fxr2 | -0.03456 | 0.301426 | 1 |

| | | | |
|---|---|---|---|
| Crk | -0.03458 | 0.117059 | 1 |
| Cog3 | -0.03462 | 0.063241 | 1 |
| Wdr70 | -0.03463 | 0.221335 | 1 |
| Utp11 | -0.03463 | 0.098271 | 1 |
| Snhg20 | -0.03465 | 0.018969 | 1 |
| Paqr3 | -0.03466 | 0.208802 | 1 |
| Thap4 | -0.03466 | 0.020772 | 1 |
| Tcf20 | -0.03468 | 0.58871 | 1 |
| Sfxn5 | -0.03469 | 0.077095 | 1 |
| Nfxl1 | -0.0347 | 0.071732 | 1 |
| Xpo5 | -0.03471 | 0.018963 | 1 |
| Raph1 | -0.03471 | 0.02046 | 1 |
| Zkscan8 | -0.03474 | 0.109209 | 1 |
| Hspa12a | -0.03476 | 0.098509 | 1 |
| Trio | -0.03476 | 0.338573 | 1 |
| Orai2 | -0.03476 | 0.494744 | 1 |
| Dmtf1 | -0.03478 | 0.176098 | 1 |
| Atp6v0a2 | -0.03479 | 0.245003 | 1 |
| Dpy19l4 | -0.03479 | 0.057536 | 1 |
| Galm | -0.0348 | 0.097707 | 1 |
| Dync1li2 | -0.03481 | 0.383808 | 1 |
| Otud6b | -0.03481 | 0.13241 | 1 |
| Gsdmd | -0.03482 | 0.141702 | 1 |
| Rbm18 | -0.03485 | 0.481374 | 1 |
| Ckap5 | -0.03486 | 0.061407 | 1 |
| Als2 | -0.03487 | 0.470452 | 1 |
| Rnf14 | -0.03494 | 0.041288 | 1 |
| Tpst2 | -0.03495 | 0.118173 | 1 |
| Atp2b1 | -0.03497 | 0.298411 | 1 |
| Atic | -0.03498 | 0.029254 | 1 |
| Rheb | -0.03498 | 0.767117 | 1 |
| Gm17435 | -0.03501 | 0.093022 | 1 |
| Inka2 | -0.03502 | 0.042649 | 1 |
| Ifit3b | -0.03503 | 0.012964 | 1 |
| Crim1 | -0.03505 | 0.443441 | 1 |
| Pigl | -0.03508 | 0.225338 | 1 |
| Gm46411 | -0.03509 | 0.006257 | 1 |
| Mrtfb | -0.03512 | 0.073057 | 1 |
| Kcnk13 | -0.03513 | 0.269738 | 1 |
| Smurf2 | -0.03514 | 0.184355 | 1 |
| Glb1 | -0.03514 | 0.121649 | 1 |
| Plekha1 | -0.03515 | 0.299384 | 1 |
| Sdha | -0.03515 | 0.365812 | 1 |
| Lars | -0.03517 | 0.455445 | 1 |
| Naa60 | -0.03517 | 0.523984 | 1 |
| Nupl2 | -0.03519 | 0.297028 | 1 |
| Gemin5 | -0.03519 | 0.112065 | 1 |
| Vps13d | -0.03519 | 0.276177 | 1 |

| | | | |
|---|---|---|---|
| Pou2f1 | -0.03521 | 0.516231 | 1 |
| Polr3gl | -0.03522 | 0.31004 | 1 |
| Phactr4 | -0.03528 | 0.21097 | 1 |
| Tmx3 | -0.03529 | 0.166121 | 1 |
| Ift80 | -0.03534 | 0.001541 | 1 |
| Ndufs2 | -0.03535 | 0.097252 | 1 |
| Scap | -0.03537 | 0.317965 | 1 |
| Tmem63b | -0.0354 | 0.036085 | 1 |
| Otud7b | -0.03543 | 0.23122 | 1 |
| Wdfy1 | -0.03544 | 0.513423 | 1 |
| 2310001H17Rik | -0.03544 | 0.058514 | 1 |
| Nifk | -0.03545 | 0.081013 | 1 |
| Gng10 | -0.03547 | 0.071632 | 1 |
| Dync1i2 | -0.03548 | 0.521164 | 1 |
| Sap30 | -0.03548 | 0.012328 | 1 |
| Irf2bp1 | -0.03549 | 0.000312 | 1 |
| Rock2 | -0.0355 | 0.051686 | 1 |
| Gtf2e1 | -0.0355 | 0.019104 | 1 |
| Limk2 | -0.03551 | 0.080665 | 1 |
| Chmp4b | -0.03554 | 0.163106 | 1 |
| Baiap2l1 | -0.03557 | 0.000586 | 1 |
| Sinhcaf | -0.03558 | 0.031639 | 1 |
| H13 | -0.03559 | 0.230074 | 1 |
| Nckap1l | -0.0356 | 0.129973 | 1 |
| Zcchc2 | -0.03561 | 0.111598 | 1 |
| 4933423P22Rik | -0.03562 | 0.262384 | 1 |
| Spout1 | -0.03562 | 0.0332 | 1 |
| Mfsd4a | -0.03562 | 0.008368 | 1 |
| Zfp974 | -0.03563 | 0.529921 | 1 |
| Spata2 | -0.03564 | 0.031694 | 1 |
| Cebpd | -0.03565 | 0.166854 | 1 |
| Maco1 | -0.03566 | 0.149977 | 1 |
| Cep78 | -0.03566 | 0.027154 | 1 |
| Dars2 | -0.03567 | 0.00772 | 1 |
| St3gal2 | -0.03571 | 0.019804 | 1 |
| Katna1 | -0.03571 | 0.005176 | 1 |
| Caml | -0.03572 | 0.093089 | 1 |
| Setd5 | -0.03572 | 0.055162 | 1 |
| C030034L19Rik | -0.03572 | 0.103631 | 1 |
| Creb1 | -0.03577 | 0.185359 | 1 |
| Arih2 | -0.03581 | 0.118277 | 1 |
| Arntl | -0.03581 | 0.183235 | 1 |
| Atl2 | -0.03582 | 0.079882 | 1 |
| Prpf40b | -0.03582 | 0.006479 | 1 |
| Stx4a | -0.03584 | 0.090272 | 1 |
| Tbc1d32 | -0.03584 | 0.058553 | 1 |
| Gm17227 | -0.03585 | 0.420616 | 1 |
| Nfkb1 | -0.03586 | 0.093608 | 1 |

| | | | |
|---|---|---|---|
| Ttf1 | -0.03587 | 0.197705 | 1 |
| Sil1 | -0.03589 | 0.549177 | 1 |
| Sumf2 | -0.03591 | 0.094229 | 1 |
| Cacna1a | -0.03595 | 0.367079 | 1 |
| Mtmr10 | -0.03598 | 0.284116 | 1 |
| Cs | -0.03599 | 0.067671 | 1 |
| Atxn2l | -0.036 | 0.058948 | 1 |
| Pak2 | -0.03601 | 0.117646 | 1 |
| Rhobtb2 | -0.03601 | 0.090944 | 1 |
| Mga | -0.03603 | 0.38256 | 1 |
| Gm28375 | -0.03603 | 0.114123 | 1 |
| Slc30a9 | -0.03605 | 0.22218 | 1 |
| Rab22a | -0.03606 | 0.216551 | 1 |
| Shprh | -0.03615 | 0.167677 | 1 |
| Slc12a9 | -0.03616 | 0.033329 | 1 |
| 6030442K20Rik | -0.03617 | 0.049003 | 1 |
| Rab39 | -0.03619 | 0.287051 | 1 |
| Zfp217 | -0.03619 | 0.116157 | 1 |
| Safb | -0.0362 | 0.021543 | 1 |
| Gtf2f2 | -0.0362 | 0.071327 | 1 |
| Trappc12 | -0.0362 | 0.353918 | 1 |
| Tgtp2 | -0.03622 | 0.007986 | 1 |
| Abcb1b | -0.03626 | 0.910871 | 1 |
| Tomm70a | -0.03627 | 0.072596 | 1 |
| Exoc2 | -0.03627 | 0.388198 | 1 |
| Opa3 | -0.03629 | 0.043347 | 1 |
| Myo9a | -0.03629 | 0.097472 | 1 |
| Slc22a5 | -0.0363 | 0.205441 | 1 |
| Aven | -0.0363 | 0.004695 | 1 |
| Got2 | -0.03631 | 0.274034 | 1 |
| Ube3b | -0.03636 | 0.337658 | 1 |
| Aagab | -0.03636 | 0.162879 | 1 |
| Ube2r2 | -0.03636 | 0.303757 | 1 |
| Ep400 | -0.03639 | 0.372332 | 1 |
| Tlr12 | -0.03641 | 0.083928 | 1 |
| Btbd10 | -0.03642 | 0.039239 | 1 |
| Nsun5 | -0.03642 | 0.011717 | 1 |
| Ppm1b | -0.03643 | 0.023011 | 1 |
| Pgam5 | -0.03643 | 0.063392 | 1 |
| Apobec1 | -0.03644 | 0.009222 | 1 |
| Dtnbp1 | -0.03645 | 0.569902 | 1 |
| Tex2 | -0.03645 | 0.218621 | 1 |
| Gtpbp4 | -0.03651 | 0.392098 | 1 |
| Tet2 | -0.03655 | 0.120246 | 1 |
| Zfp131 | -0.03655 | 0.461749 | 1 |
| Pcgf6 | -0.03656 | 0.02512 | 1 |
| Fig4 | -0.03658 | 0.333078 | 1 |
| Uty | -0.03661 | 0.13209 | 1 |

| | | | |
|---|---|---|---|
| Ubr1 | -0.03665 | 0.112309 | 1 |
| 2810403D21Rik | -0.03666 | 0.023033 | 1 |
| Rprd1a | -0.03666 | 0.055898 | 1 |
| Tmc6 | -0.0367 | 0.094029 | 1 |
| Ptpn4 | -0.0367 | 0.326065 | 1 |
| Kitl | -0.03671 | 0.01891 | 1 |
| Abcc1 | -0.03674 | 0.743714 | 1 |
| Erlin1 | -0.03676 | 0.042308 | 1 |
| Prcp | -0.03677 | 0.08226 | 1 |
| Zcchc4 | -0.03678 | 0.031962 | 1 |
| Actr2 | -0.03679 | 0.036942 | 1 |
| Gm20404 | -0.03679 | 0.031698 | 1 |
| Kat6b | -0.03681 | 0.326286 | 1 |
| Sh3bp4 | -0.03682 | 0.005764 | 1 |
| Clec2d | -0.03682 | 0.083396 | 1 |
| Pnisr | -0.03683 | 0.083525 | 1 |
| Mast3 | -0.03683 | 0.563363 | 1 |
| Rassf3 | -0.03684 | 0.424749 | 1 |
| Tm6sf1 | -0.03686 | 0.632844 | 1 |
| Capzb | -0.03686 | 0.127603 | 1 |
| Clasrp | -0.03686 | 0.01812 | 1 |
| Usp19 | -0.03689 | 0.45329 | 1 |
| Gapdh | -0.0369 | 0.997355 | 1 |
| Brpf3 | -0.03691 | 0.305774 | 1 |
| Prkd3 | -0.03691 | 0.354523 | 1 |
| Aspscr1 | -0.03692 | 0.019796 | 1 |
| Bak1 | -0.03693 | 0.026366 | 1 |
| Zfp606 | -0.03693 | 0.008137 | 1 |
| Srgap2 | -0.03694 | 0.030827 | 1 |
| Nin | -0.03694 | 0.368525 | 1 |
| Gpd1l | -0.03697 | 0.145599 | 1 |
| Ext2 | -0.03701 | 0.254082 | 1 |
| Med26 | -0.03702 | 0.134643 | 1 |
| Cep85l | -0.03704 | 0.323453 | 1 |
| Cdv3 | -0.03705 | 0.280967 | 1 |
| Zfp369 | -0.03708 | 0.033768 | 1 |
| Trak1 | -0.03708 | 0.696153 | 1 |
| Vps8 | -0.03709 | 0.383459 | 1 |
| Rnf141 | -0.03711 | 0.014895 | 1 |
| Rnf185 | -0.03711 | 0.270461 | 1 |
| Cdc37l1 | -0.03713 | 0.060204 | 1 |
| Nacc2 | -0.03714 | 0.073505 | 1 |
| Mipol1 | -0.03717 | 0.573072 | 1 |
| Nabp2 | -0.03718 | 0.095165 | 1 |
| Wdr43 | -0.0372 | 0.215362 | 1 |
| Usp21 | -0.0372 | 0.007527 | 1 |
| Crybg1 | -0.03725 | 0.01762 | 1 |
| Ccdc77 | -0.03726 | 0.014315 | 1 |

| | | | |
|---|---|---|---|
| Gm48765 | -0.03729 | 0.039813 | 1 |
| Tfdp1 | -0.03729 | 0.036125 | 1 |
| Ndst1 | -0.0373 | 0.100191 | 1 |
| Gm16759 | -0.03731 | 0.027579 | 1 |
| 2010315B03Rik | -0.03737 | 0.046395 | 1 |
| Cdk14 | -0.03739 | 0.091038 | 1 |
| Arpc1b | -0.03743 | 0.164169 | 1 |
| Zcchc7 | -0.03743 | 0.067085 | 1 |
| Ptpn2 | -0.03745 | 0.03047 | 1 |
| Kat5 | -0.03745 | 0.013215 | 1 |
| Cog8 | -0.0375 | 0.223171 | 1 |
| Tspan7 | -0.0375 | 0.439172 | 1 |
| Kmt5c | -0.03752 | 0.000599 | 1 |
| Ing2 | -0.03753 | 0.035591 | 1 |
| Zdhhc17 | -0.03757 | 0.350226 | 1 |
| Ccdc130 | -0.03758 | 0.064068 | 1 |
| Arid2 | -0.03759 | 0.264204 | 1 |
| Exoc3 | -0.03764 | 0.383007 | 1 |
| Impa2 | -0.03764 | 0.127928 | 1 |
| Mtx1 | -0.03766 | 0.129392 | 1 |
| Notch1 | -0.03771 | 0.062089 | 1 |
| Tcf3 | -0.03772 | 0.129857 | 1 |
| Pkd2 | -0.03773 | 0.081256 | 1 |
| Slc49a4 | -0.03774 | 0.750209 | 1 |
| Dap3 | -0.03775 | 0.14453 | 1 |
| Nfs1 | -0.03778 | 0.26684 | 1 |
| Krit1 | -0.03783 | 0.142855 | 1 |
| Myo9b | -0.03783 | 0.071928 | 1 |
| Lrpprc | -0.03784 | 0.067656 | 1 |
| Bmpr2 | -0.03786 | 0.543738 | 1 |
| Rilpl2 | -0.03787 | 0.286863 | 1 |
| Mir99ahg | -0.03788 | 0.244986 | 1 |
| Spata5 | -0.03789 | 0.3294 | 1 |
| Gna11 | -0.03791 | 0.01483 | 1 |
| Gramd3 | -0.03793 | 0.071864 | 1 |
| Naa16 | -0.03794 | 0.11699 | 1 |
| Rsrc2 | -0.03794 | 0.454158 | 1 |
| Ttpal | -0.03799 | 0.052173 | 1 |
| Trit1 | -0.03799 | 0.333288 | 1 |
| Tbp | -0.038 | 0.116102 | 1 |
| H2-Q4 | -0.03801 | 0.420864 | 1 |
| Camsap1 | -0.03802 | 0.277171 | 1 |
| Nova1 | -0.03808 | 0.187693 | 1 |
| AW112010 | -0.03815 | 0.984616 | 1 |
| Cfap74 | -0.03815 | 0.035799 | 1 |
| Smc6 | -0.03818 | 0.312463 | 1 |
| Mob1a | -0.03825 | 0.23941 | 1 |
| Chst11 | -0.03826 | 0.458831 | 1 |

| | | | |
|---|---|---|---|
| Tmpo | -0.03828 | 0.22904 | 1 |
| Eif5b | -0.03831 | 0.149049 | 1 |
| Sp110 | -0.03832 | 0.042712 | 1 |
| Cat | -0.03834 | 0.023888 | 1 |
| Lncpint | -0.03834 | 0.805498 | 1 |
| Hinfp | -0.03834 | 0.01868 | 1 |
| Rsu1 | -0.03841 | 0.30078 | 1 |
| Cyp4f13 | -0.03843 | 0.170291 | 1 |
| Stard3 | -0.03843 | 0.205909 | 1 |
| Ubr2 | -0.03843 | 0.037262 | 1 |
| Evi5 | -0.03846 | 0.390583 | 1 |
| Zfyve9 | -0.03849 | 0.22903 | 1 |
| Irak2 | -0.0385 | 0.788896 | 1 |
| Cdh23 | -0.03851 | 0.143798 | 1 |
| Stambpl1 | -0.03853 | 0.332165 | 1 |
| Map4k5 | -0.03856 | 0.269642 | 1 |
| Tgfb1 | -0.03859 | 0.050471 | 1 |
| Plekhm2 | -0.03859 | 0.102708 | 1 |
| Letm2 | -0.03866 | 0.000526 | 1 |
| Fam49b | -0.03866 | 0.071496 | 1 |
| Mttp | -0.03867 | 0.282872 | 1 |
| Nipa2 | -0.03867 | 0.098191 | 1 |
| Alox5 | -0.03872 | 0.309696 | 1 |
| Gopc | -0.03873 | 0.00854 | 1 |
| Lfng | -0.03875 | 0.084991 | 1 |
| Lrrc8c | -0.03877 | 0.191974 | 1 |
| Kdm6b | -0.03878 | 0.854475 | 1 |
| Bfar | -0.03879 | 0.324699 | 1 |
| Naa40 | -0.03879 | 0.061909 | 1 |
| Map3k10 | -0.03881 | 0.012905 | 1 |
| Rnf167 | -0.03881 | 0.252554 | 1 |
| Smg1 | -0.03882 | 0.35711 | 1 |
| Slco2b1 | -0.03883 | 0.255888 | 1 |
| Atp8b1 | -0.03887 | 0.086578 | 1 |
| Ralgapa1 | -0.03889 | 0.163537 | 1 |
| Necap1 | -0.03893 | 0.006828 | 1 |
| Abhd18 | -0.03898 | 0.036002 | 1 |
| Smchd1 | -0.03898 | 0.050788 | 1 |
| Ppp2r5c | -0.03902 | 0.088955 | 1 |
| Arsg | -0.03908 | 0.442282 | 1 |
| Rbl2 | -0.03908 | 0.109636 | 1 |
| Nrd1 | -0.03911 | 0.415982 | 1 |
| Slc25a44 | -0.03913 | 0.025129 | 1 |
| Ssh1 | -0.03914 | 0.033398 | 1 |
| Ctnnbl1 | -0.03914 | 0.348254 | 1 |
| Tent2 | -0.03918 | 0.023734 | 1 |
| Spred1 | -0.03919 | 0.178658 | 1 |
| Fbxo11 | -0.0392 | 0.296931 | 1 |

| | | | |
|---|---|---|---|
| Plekho1 | -0.03923 | 0.125833 | 1 |
| Bach1 | -0.03929 | 0.776589 | 1 |
| Pfdn1 | -0.0393 | 0.156536 | 1 |
| Sf3a3 | -0.03933 | 0.069804 | 1 |
| Lmf1 | -0.03935 | 0.366619 | 1 |
| Dedd | -0.03936 | 0.100118 | 1 |
| Ano10 | -0.0394 | 0.11613 | 1 |
| Mthfsl | -0.03942 | 0.374368 | 1 |
| Mphosph8 | -0.03943 | 0.167988 | 1 |
| Ssbp2 | -0.03944 | 0.152216 | 1 |
| Med1 | -0.03946 | 0.094438 | 1 |
| Dusp22 | -0.03947 | 0.027553 | 1 |
| Tdg | -0.03947 | 0.131281 | 1 |
| A630001G21Rik | -0.03948 | 0.355627 | 1 |
| Lrba | -0.03949 | 0.924275 | 1 |
| Tmem131l | -0.03949 | 0.299285 | 1 |
| Gart | -0.0395 | 0.460648 | 1 |
| Med15 | -0.03955 | 0.198663 | 1 |
| Akap9 | -0.03956 | 0.313139 | 1 |
| Tnfrsf1b | -0.0396 | 0.070289 | 1 |
| Ube3c | -0.03965 | 0.232818 | 1 |
| Ccdc192 | -0.03971 | 0.107908 | 1 |
| Echdc1 | -0.03972 | 0.010846 | 1 |
| Map4 | -0.03975 | 0.12917 | 1 |
| Mettl6 | -0.03976 | 0.135414 | 1 |
| Pdxdc1 | -0.03976 | 0.256563 | 1 |
| Ypel2 | -0.0398 | 0.355899 | 1 |
| Ptpn1 | -0.03982 | 0.218896 | 1 |
| Prpsap2 | -0.03982 | 0.213903 | 1 |
| Hnrnph3 | -0.03982 | 0.321766 | 1 |
| Bcl7c | -0.03984 | 0.047882 | 1 |
| Mff | -0.03984 | 0.116796 | 1 |
| Ralgapa2 | -0.03987 | 0.094448 | 1 |
| Dapk3 | -0.03992 | 0.228509 | 1 |
| 9330159M07Rik | -0.03992 | 0.054198 | 1 |
| Maml1 | -0.03993 | 0.386195 | 1 |
| Vps13a | -0.03996 | 0.221139 | 1 |
| Clint1 | -0.03996 | 0.146913 | 1 |
| Wnk1 | -0.03997 | 0.043308 | 1 |
| Zc3h13 | -0.04 | 0.281808 | 1 |
| Odf2 | -0.04 | 0.028358 | 1 |
| Ccdc50 | -0.04002 | 0.059277 | 1 |
| Xkr4 | -0.04002 | 0.012918 | 1 |
| Atf7ip | -0.04004 | 0.209466 | 1 |
| Pisd | -0.0401 | 0.15876 | 1 |
| Selenoi | -0.0401 | 0.088419 | 1 |
| Dhx9 | -0.04011 | 0.824935 | 1 |
| Crebbp | -0.04013 | 0.07628 | 1 |

| | | | |
|---|---|---|---|
| Cds2 | -0.04013 | 0.110043 | 1 |
| Fbrs | -0.04015 | 0.003656 | 1 |
| Lifr | -0.04015 | 0.858247 | 1 |
| Mpp1 | -0.04016 | 0.44586 | 1 |
| Bbx | -0.04016 | 0.228863 | 1 |
| Rasgrf2 | -0.04018 | 0.03108 | 1 |
| Snx32 | -0.04018 | 0.124467 | 1 |
| Trim30b | -0.0402 | 0.002963 | 1 |
| Ttc7 | -0.04022 | 0.419496 | 1 |
| Cdkl3 | -0.04025 | 0.086067 | 1 |
| Pwwp2a | -0.04027 | 0.247445 | 1 |
| Uri1 | -0.04029 | 0.287644 | 1 |
| Gramd4 | -0.04034 | 0.000558 | 1 |
| Scly | -0.04038 | 0.043328 | 1 |
| Fbxo34 | -0.0404 | 0.152508 | 1 |
| 2310022A10Rik | -0.04041 | 0.00321 | 1 |
| Thap12 | -0.04047 | 0.256176 | 1 |
| Ccar1 | -0.04048 | 0.146931 | 1 |
| Phf11b | -0.04052 | 0.097997 | 1 |
| Gm30881 | -0.04054 | 0.052629 | 1 |
| Zbed4 | -0.04055 | 0.298745 | 1 |
| Snx6 | -0.04057 | 0.088883 | 1 |
| Cyth3 | -0.04063 | 0.229101 | 1 |
| Dse | -0.04066 | 0.057732 | 1 |
| Mapkap1 | -0.0407 | 0.177479 | 1 |
| Zranb3 | -0.04072 | 0.085493 | 1 |
| Slc14a1 | -0.04074 | 0.006904 | 1 |
| Filip1l | -0.04074 | 0.429201 | 1 |
| Eefsec | -0.04075 | 0.571967 | 1 |
| Arhgef18 | -0.04078 | 0.02188 | 1 |
| Mgat4b | -0.04078 | 0.148784 | 1 |
| Farsb | -0.04078 | 0.04368 | 1 |
| Slc44a2 | -0.04085 | 0.22154 | 1 |
| Trim30c | -0.04086 | 0.004737 | 1 |
| Senp2 | -0.04086 | 0.204784 | 1 |
| Rhobtb1 | -0.04086 | 0.405877 | 1 |
| Ptpn23 | -0.04088 | 0.046566 | 1 |
| Arnt | -0.04091 | 0.016075 | 1 |
| Rps27rt | -0.04093 | 0.054916 | 1 |
| Emsy | -0.04094 | 0.007362 | 1 |
| Dhx30 | -0.04094 | 0.070813 | 1 |
| Mmp14 | -0.04095 | 0.10135 | 1 |
| Atg3 | -0.04097 | 0.093228 | 1 |
| Washc3 | -0.04099 | 0.048507 | 1 |
| Parp12 | -0.041 | 0.052098 | 1 |
| Cep57l1 | -0.04101 | 0.111702 | 1 |
| Cast | -0.04102 | 0.30961 | 1 |
| Hhex | -0.04102 | 0.153025 | 1 |

| | | | |
|---|---|---|---|
| Npepps | -0.04102 | 0.094685 | 1 |
| Tex10 | -0.04104 | 0.419914 | 1 |
| Mcoln1 | -0.04108 | 0.089672 | 1 |
| Gm28981 | -0.04109 | 0.011183 | 1 |
| Loxl3 | -0.0411 | 0.163909 | 1 |
| Traf5 | -0.04111 | 0.121661 | 1 |
| Mbd6 | -0.04113 | 0.01129 | 1 |
| Dyrk1a | -0.04118 | 0.415952 | 1 |
| 5430427O19Rik | -0.0412 | 0.284106 | 1 |
| Fam120a | -0.04121 | 0.045267 | 1 |
| Gm50035 | -0.04124 | 0.287798 | 1 |
| Yes1 | -0.04128 | 0.108909 | 1 |
| Invs | -0.04131 | 0.163577 | 1 |
| Galnt2 | -0.04131 | 0.027668 | 1 |
| Oprm1 | -0.04136 | 0.004326 | 1 |
| Traf3ip3 | -0.04137 | 0.025495 | 1 |
| Secisbp2l | -0.04138 | 0.269124 | 1 |
| Socs5 | -0.04141 | 0.078958 | 1 |
| Slc16a10 | -0.04143 | 0.316793 | 1 |
| H2-T22 | -0.04146 | 0.107385 | 1 |
| Senp1 | -0.04148 | 0.017552 | 1 |
| Nrf1 | -0.04148 | 0.069077 | 1 |
| Tcf4 | -0.04148 | 0.116899 | 1 |
| Smg5 | -0.04149 | 0.058532 | 1 |
| Caap1 | -0.04152 | 0.003734 | 1 |
| Tmem161b | -0.04153 | 0.083091 | 1 |
| Heatr3 | -0.04156 | 0.132499 | 1 |
| Arhgef2 | -0.04157 | 0.036253 | 1 |
| Zfand6 | -0.04159 | 0.330886 | 1 |
| Etnk1 | -0.04165 | 0.060288 | 1 |
| Lrrc8d | -0.04167 | 0.384123 | 1 |
| Krcc1 | -0.04174 | 0.214649 | 1 |
| Ppp5c | -0.04174 | 0.097356 | 1 |
| Ift52 | -0.04178 | 0.334105 | 1 |
| Pqlc2 | -0.04187 | 0.010226 | 1 |
| Kansl3 | -0.04188 | 0.162486 | 1 |
| Nipbl | -0.04189 | 0.23233 | 1 |
| Prdm10 | -0.04189 | 0.378262 | 1 |
| Fem1c | -0.04189 | 0.627407 | 1 |
| Gm4890 | -0.0419 | 0.001057 | 1 |
| Sfmbt1 | -0.0419 | 0.055567 | 1 |
| Dgkz | -0.04191 | 0.072733 | 1 |
| Cdc14b | -0.04191 | 0.058808 | 1 |
| Tmem168 | -0.04191 | 0.265587 | 1 |
| Idnk | -0.04192 | 0.037969 | 1 |
| Fxyd5 | -0.04195 | 0.513001 | 1 |
| Cep112 | -0.04197 | 0.085042 | 1 |
| Dock4 | -0.04197 | 0.011556 | 1 |

| | | | |
|---|---|---|---|
| Rnf4 | -0.04199 | 0.058641 | 1 |
| Lekr1 | -0.042 | 0.334535 | 1 |
| Zkscan5 | -0.04201 | 0.002604 | 1 |
| Phkg2 | -0.04202 | 0.041992 | 1 |
| Acly | -0.04203 | 0.157632 | 1 |
| Dctn5 | -0.04205 | 0.207872 | 1 |
| Fndc3a | -0.04205 | 0.067218 | 1 |
| Dgat1 | -0.04216 | 0.033064 | 1 |
| Vta1 | -0.04218 | 0.046787 | 1 |
| Thumpd3 | -0.04218 | 0.02189 | 1 |
| Dcp1a | -0.04226 | 0.046747 | 1 |
| Ppm1l | -0.04227 | 0.039214 | 1 |
| Ube4a | -0.04228 | 0.114015 | 1 |
| Rrp36 | -0.04231 | 0.111117 | 1 |
| Stag2 | -0.04234 | 0.320777 | 1 |
| Phf11d | -0.04242 | 0.009983 | 1 |
| Atp6v0d1 | -0.04242 | 0.294749 | 1 |
| Mkrn1 | -0.04243 | 0.107648 | 1 |
| Rictor | -0.04244 | 0.246316 | 1 |
| AC149090.1 | -0.04245 | 0.113369 | 1 |
| Col4a3bp | -0.04246 | 0.095317 | 1 |
| Med13 | -0.04248 | 0.09541 | 1 |
| Bco2 | -0.04248 | 0.047309 | 1 |
| Trim25 | -0.04251 | 0.022145 | 1 |
| Trim33 | -0.04263 | 0.441921 | 1 |
| Rtp4 | -0.04264 | 0.370821 | 1 |
| Arfgap3 | -0.04264 | 0.109635 | 1 |
| Atxn3 | -0.04267 | 0.241343 | 1 |
| Rfc1 | -0.04269 | 0.438959 | 1 |
| Micu1 | -0.0427 | 0.088344 | 1 |
| Klhl28 | -0.04272 | 0.043721 | 1 |
| Iscu | -0.04272 | 0.150631 | 1 |
| Cep164 | -0.04272 | 0.087173 | 1 |
| Mapkapk3 | -0.04273 | 0.013604 | 1 |
| Eml4 | -0.04273 | 0.088573 | 1 |
| Sergef | -0.04273 | 0.100693 | 1 |
| Zfp646 | -0.04273 | 0.005656 | 1 |
| Ssbp3 | -0.04276 | 0.018363 | 1 |
| Ermard | -0.04279 | 0.111649 | 1 |
| Pgap2 | -0.04282 | 0.061751 | 1 |
| Dock6 | -0.04283 | 0.273952 | 1 |
| Hipk3 | -0.04283 | 0.781655 | 1 |
| Fam102a | -0.04288 | 0.093946 | 1 |
| Gipc1 | -0.04289 | 0.005704 | 1 |
| Phtf1 | -0.0429 | 0.07667 | 1 |
| Kdm3a | -0.04293 | 0.640794 | 1 |
| Tor1aip1 | -0.04297 | 0.015561 | 1 |
| Senp8 | -0.04299 | 0.009324 | 1 |

| | | | |
|---|---|---|---|
| Ube2d1 | -0.04299 | 0.174735 | 1 |
| Arhgap4 | -0.04307 | 0.728176 | 1 |
| Mef2d | -0.04309 | 0.032168 | 1 |
| Cmtm7 | -0.04309 | 0.131911 | 1 |
| Cryzl1 | -0.0431 | 0.025173 | 1 |
| Ppp6r2 | -0.0431 | 0.220277 | 1 |
| Ash1l | -0.04319 | 0.193113 | 1 |
| Dcp1b | -0.04323 | 0.369449 | 1 |
| Olfr56 | -0.04324 | 0.00414 | 1 |
| Zcchc24 | -0.04325 | 0.102032 | 1 |
| Mthfd2 | -0.04327 | 0.161456 | 1 |
| Ccdc25 | -0.0433 | 0.206957 | 1 |
| Cep68 | -0.04332 | 0.036896 | 1 |
| Eps15l1 | -0.04338 | 0.072553 | 1 |
| Psen2 | -0.04341 | 0.058965 | 1 |
| Adora3 | -0.04347 | 0.121909 | 1 |
| 1700010I14Rik | -0.04347 | 0.033319 | 1 |
| Serp1 | -0.04352 | 0.468485 | 1 |
| Xrcc4 | -0.04353 | 0.04231 | 1 |
| Rnasel | -0.04355 | 0.081599 | 1 |
| Vps37a | -0.04358 | 0.341602 | 1 |
| 0610040J01Rik | -0.0436 | 0.223468 | 1 |
| Setx | -0.04362 | 0.43442 | 1 |
| Ahr | -0.04362 | 0.033355 | 1 |
| Ifi209 | -0.04364 | 0.088169 | 1 |
| Bcar3 | -0.04365 | 0.025363 | 1 |
| Pfkfb4 | -0.04366 | 0.364866 | 1 |
| Kmt5b | -0.04372 | 0.317396 | 1 |
| Fnbp1 | -0.04373 | 0.017422 | 1 |
| Dok1 | -0.04377 | 0.003449 | 1 |
| Myom1 | -0.0438 | 0.032645 | 1 |
| Tmem260 | -0.04381 | 0.231388 | 1 |
| Wipf1 | -0.04381 | 0.03185 | 1 |
| Cul4a | -0.04383 | 0.49376 | 1 |
| Cnot4 | -0.04384 | 0.168519 | 1 |
| H2-T24 | -0.04385 | 0.072107 | 1 |
| Gm43713 | -0.04388 | 0.103167 | 1 |
| Stard8 | -0.04388 | 0.229733 | 1 |
| Fbxw7 | -0.0439 | 0.02502 | 1 |
| Hmces | -0.04393 | 0.029447 | 1 |
| Sp1 | -0.04394 | 0.093305 | 1 |
| Lrch1 | -0.04396 | 0.114388 | 1 |
| Sec14l1 | -0.04401 | 0.223789 | 1 |
| Clock | -0.04401 | 0.19745 | 1 |
| Nectin4 | -0.04402 | 0.029346 | 1 |
| Trim12a | -0.04403 | 0.415144 | 1 |
| Fndc3b | -0.04408 | 0.272796 | 1 |
| Carnmt1 | -0.04409 | 0.005624 | 1 |

| | | | |
|---|---|---|---|
| Por | -0.04412 | 0.024577 | 1 |
| Itpr3 | -0.04414 | 0.018855 | 1 |
| Card11 | -0.04414 | 0.005983 | 1 |
| Tbc1d22b | -0.04417 | 0.101374 | 1 |
| Gm42031 | -0.04419 | 0.002943 | 1 |
| Washc2 | -0.0442 | 0.215496 | 1 |
| Tut7 | -0.04422 | 0.644132 | 1 |
| Cerk | -0.04429 | 0.036783 | 1 |
| Ncbp3 | -0.04435 | 0.263011 | 1 |
| Arpc1a | -0.04436 | 0.201635 | 1 |
| Zfp26 | -0.04436 | 0.063849 | 1 |
| Lrig3 | -0.0444 | 0.010015 | 1 |
| 1110002L01Rik | -0.04445 | 0.015638 | 1 |
| Cep162 | -0.04446 | 0.06128 | 1 |
| Nup153 | -0.04447 | 0.324847 | 1 |
| Nagk | -0.04451 | 0.055741 | 1 |
| Cox16 | -0.04452 | 0.137971 | 1 |
| Zfyve1 | -0.04456 | 0.190661 | 1 |
| Gm31763 | -0.04456 | 0.000627 | 1 |
| Cramp1l | -0.04461 | 0.049474 | 1 |
| Rab6b | -0.04466 | 0.056256 | 1 |
| Mtrex | -0.04467 | 0.183047 | 1 |
| Ophn1 | -0.04469 | 0.035764 | 1 |
| Bcl9 | -0.04471 | 0.016427 | 1 |
| Tbl1xr1 | -0.04474 | 0.079465 | 1 |
| Taf4 | -0.04475 | 0.141584 | 1 |
| Atm | -0.0448 | 0.323103 | 1 |
| Dhx58 | -0.04486 | 0.008956 | 1 |
| Gm11808 | -0.0449 | 0.076672 | 1 |
| Srrt | -0.04492 | 0.054932 | 1 |
| Mecp2 | -0.04492 | 0.093182 | 1 |
| Plcb2 | -0.04493 | 0.116403 | 1 |
| Hspbap1 | -0.04494 | 0.024257 | 1 |
| Lrrc58 | -0.04495 | 0.303411 | 1 |
| Atp11c | -0.04497 | 0.168522 | 1 |
| Chd8 | -0.045 | 0.071863 | 1 |
| Daam1 | -0.045 | 0.030737 | 1 |
| Zfp568 | -0.04501 | 0.084992 | 1 |
| Lrp12 | -0.04504 | 0.76679 | 1 |
| Strn3 | -0.04506 | 0.11368 | 1 |
| Hmox2 | -0.04509 | 0.071871 | 1 |
| Exog | -0.0451 | 4.65E-05 | 1 |
| Pde4a | -0.04514 | 0.027116 | 1 |
| Gm28501 | -0.04515 | 0.010499 | 1 |
| Rnf31 | -0.04516 | 0.02797 | 1 |
| Cnst | -0.04517 | 0.036269 | 1 |
| Fut8 | -0.0452 | 0.038324 | 1 |
| Limd1 | -0.04522 | 0.267914 | 1 |

| | | | |
|---|---|---|---|
| Nadk2 | -0.04523 | 0.032407 | 1 |
| Ptpa | -0.04525 | 0.064663 | 1 |
| Pdpr | -0.04529 | 0.084237 | 1 |
| Gcnt7 | -0.0453 | 0.001153 | 1 |
| Soga1 | -0.04535 | 0.21348 | 1 |
| Phka1 | -0.04535 | 0.018299 | 1 |
| Sucla2 | -0.04537 | 0.072367 | 1 |
| Crebzf | -0.0454 | 0.07652 | 1 |
| Taf5l | -0.04549 | 0.000767 | 1 |
| Gm11508 | -0.04551 | 0.017411 | 1 |
| Arhgap17 | -0.04551 | 0.039976 | 1 |
| Ddx31 | -0.04552 | 0.614408 | 1 |
| Trpm2 | -0.04553 | 0.088429 | 1 |
| Inpp5b | -0.04554 | 0.047198 | 1 |
| Parp4 | -0.04558 | 0.194188 | 1 |
| Dnm2 | -0.04561 | 0.010393 | 1 |
| Ogfrl1 | -0.04564 | 0.196297 | 1 |
| Thoc1 | -0.04565 | 0.364877 | 1 |
| Kdm5d | -0.04567 | 0.217976 | 1 |
| Mob4 | -0.04568 | 0.495829 | 1 |
| Ttll5 | -0.04568 | 0.151243 | 1 |
| Pcca | -0.04571 | 0.085078 | 1 |
| Heatr1 | -0.04571 | 0.016856 | 1 |
| Vwa8 | -0.04572 | 0.499884 | 1 |
| Ecd | -0.04573 | 0.111918 | 1 |
| Lrrc41 | -0.04574 | 0.111808 | 1 |
| Smim14 | -0.04576 | 0.16429 | 1 |
| Scfd2 | -0.04577 | 0.073772 | 1 |
| Scaf8 | -0.04579 | 0.089726 | 1 |
| Prmt3 | -0.0458 | 0.33954 | 1 |
| Desi2 | -0.04581 | 0.346604 | 1 |
| Zfp738 | -0.04585 | 0.007103 | 1 |
| Ifit1 | -0.04587 | 0.021426 | 1 |
| Gm28286 | -0.04591 | 0.000126 | 1 |
| Snap23 | -0.04593 | 0.117471 | 1 |
| Kdm4a | -0.04593 | 0.003701 | 1 |
| Adcy9 | -0.04593 | 0.027157 | 1 |
| Ylpm1 | -0.04593 | 0.023477 | 1 |
| Cand1 | -0.04594 | 0.057452 | 1 |
| Ggt5 | -0.04596 | 0.119104 | 1 |
| Pcf11 | -0.04598 | 0.116376 | 1 |
| Myo10 | -0.046 | 0.000987 | 1 |
| Fgd2 | -0.04604 | 0.024646 | 1 |
| Carm1 | -0.0461 | 0.120979 | 1 |
| Kctd5 | -0.04611 | 0.071564 | 1 |
| Smarca4 | -0.04611 | 0.032759 | 1 |
| Gm12743 | -0.04615 | 0.003625 | 1 |
| Add1 | -0.04615 | 0.108108 | 1 |

| | | | |
|---|---|---|---|
| Tubgcp3 | -0.04619 | 0.019477 | 1 |
| Zfp638 | -0.04623 | 0.118395 | 1 |
| Mllt6 | -0.04625 | 0.002364 | 1 |
| Itsn2 | -0.04629 | 0.071684 | 1 |
| Golph3l | -0.04634 | 0.064784 | 1 |
| Kmt2c | -0.04635 | 0.246626 | 1 |
| Anxa3 | -0.04636 | 0.011084 | 1 |
| Nploc4 | -0.04637 | 0.003391 | 1 |
| Actr1a | -0.04639 | 0.077771 | 1 |
| Lrrfip2 | -0.04639 | 0.04272 | 1 |
| Trmt61b | -0.04639 | 0.000449 | 1 |
| Bin3 | -0.04645 | 0.036443 | 1 |
| Zmynd8 | -0.04646 | 0.041055 | 1 |
| Zzef1 | -0.04646 | 0.099854 | 1 |
| Rps6ka2 | -0.04646 | 0.014139 | 1 |
| U2surp | -0.04647 | 0.473606 | 1 |
| Coro1c | -0.04648 | 0.106931 | 1 |
| Sdhaf2 | -0.04652 | 0.03954 | 1 |
| Pik3ip1 | -0.04653 | 0.004951 | 1 |
| Prr14 | -0.04655 | 0.044158 | 1 |
| Zyg11b | -0.04656 | 0.133385 | 1 |
| Pacs1 | -0.04659 | 0.24347 | 1 |
| Galk2 | -0.0466 | 0.104892 | 1 |
| Mtif2 | -0.04661 | 0.097815 | 1 |
| Treml2 | -0.04661 | 0.008013 | 1 |
| Ahcyl2 | -0.04665 | 0.0127 | 1 |
| Gm15478 | -0.04666 | 0.044264 | 1 |
| St3gal5 | -0.04667 | 0.1587 | 1 |
| Retreg1 | -0.04667 | 0.101281 | 1 |
| Cdip1 | -0.04674 | 0.056615 | 1 |
| Zbtb40 | -0.04674 | 0.006039 | 1 |
| Pogz | -0.04675 | 0.113027 | 1 |
| Ppp1cb | -0.04683 | 0.090342 | 1 |
| Gpr65 | -0.04684 | 0.185306 | 1 |
| Large1 | -0.04684 | 0.068631 | 1 |
| Lcor | -0.04684 | 0.035423 | 1 |
| Zfhx3 | -0.04686 | 0.016538 | 1 |
| Mta3 | -0.04688 | 0.043369 | 1 |
| Tmem273 | -0.04689 | 0.175791 | 1 |
| Nub1 | -0.04689 | 0.28882 | 1 |
| Nme6 | -0.04691 | 0.13392 | 1 |
| Pibf1 | -0.04692 | 0.05498 | 1 |
| Ttll3 | -0.04693 | 0.012199 | 1 |
| Prkdc | -0.04696 | 0.08874 | 1 |
| Ap2a1 | -0.04698 | 0.025471 | 1 |
| Zbtb2 | -0.04702 | 0.010056 | 1 |
| Prpf40a | -0.04705 | 0.134656 | 1 |
| Gm15523 | -0.04706 | 0.001624 | 1 |

| | | | |
|---|---|---|---|
| Dtd1 | -0.04708 | 0.011907 | 1 |
| Phf8 | -0.04709 | 0.07019 | 1 |
| Entpd7 | -0.04712 | 0.008163 | 1 |
| Mier1 | -0.04715 | 0.225593 | 1 |
| Snx29 | -0.04715 | 0.029352 | 1 |
| Atg16l1 | -0.04721 | 0.145373 | 1 |
| Anapc4 | -0.04723 | 0.133488 | 1 |
| Ccz1 | -0.04726 | 0.142618 | 1 |
| Commd1 | -0.04726 | 0.49184 | 1 |
| Ankrd27 | -0.04726 | 0.05581 | 1 |
| G6pdx | -0.04727 | 0.017637 | 1 |
| Srcap | -0.04729 | 0.800341 | 1 |
| Insyn2b | -0.04733 | 0.143368 | 1 |
| Slc11a1 | -0.04736 | 0.060399 | 1 |
| Ap3m1 | -0.04742 | 0.163826 | 1 |
| Brox | -0.04742 | 0.080635 | 1 |
| Tnfrsf13b | -0.04742 | 0.106676 | 1 |
| Plagl2 | -0.04742 | 0.036704 | 1 |
| Luc7l | -0.04744 | 0.060237 | 1 |
| 2610020C07Rik | -0.04745 | 0.328825 | 1 |
| Ankib1 | -0.04748 | 0.014051 | 1 |
| Fbxl2 | -0.04749 | 0.008682 | 1 |
| Usp46 | -0.04749 | 0.115724 | 1 |
| Khdc4 | -0.04749 | 0.086471 | 1 |
| Fgr | -0.04753 | 0.05118 | 1 |
| Fanca | -0.04754 | 0.000137 | 1 |
| Stam2 | -0.04757 | 0.088927 | 1 |
| Setdb1 | -0.04758 | 0.092581 | 1 |
| Slf2 | -0.04763 | 0.159044 | 1 |
| Sec24a | -0.04764 | 0.250439 | 1 |
| Traf3 | -0.04771 | 0.086797 | 1 |
| Slc16a7 | -0.04773 | 0.102116 | 1 |
| Gak | -0.04775 | 0.076719 | 1 |
| Plpp1 | -0.04776 | 0.023926 | 1 |
| Vcl | -0.04778 | 0.091012 | 1 |
| Nup210l | -0.0478 | 0.000107 | 1 |
| Fam120b | -0.0478 | 0.537294 | 1 |
| Eapp | -0.04782 | 0.004003 | 1 |
| Ap5m1 | -0.04783 | 0.047424 | 1 |
| Tmem189 | -0.04784 | 0.118604 | 1 |
| Tjap1 | -0.04785 | 0.027905 | 1 |
| Batf2 | -0.04787 | 0.001197 | 1 |
| Rcbtb2 | -0.0479 | 0.770537 | 1 |
| Clasp1 | -0.04791 | 0.004387 | 1 |
| Disp1 | -0.04795 | 0.054428 | 1 |
| Prpf6 | -0.04798 | 0.171723 | 1 |
| Mon1a | -0.04799 | 0.00014 | 1 |
| Il27 | -0.048 | 0.010561 | 1 |

| | | | |
|---|---|---|---|
| Cbfb | -0.04801 | 0.224819 | 1 |
| Mark4 | -0.04804 | 0.039034 | 1 |
| Bri3 | -0.04806 | 0.732248 | 1 |
| Cpeb4 | -0.04809 | 0.077868 | 1 |
| Wasf2 | -0.0481 | 0.006053 | 1 |
| Ccdc174 | -0.0481 | 0.018734 | 1 |
| 2810004N23Rik | -0.04816 | 0.044322 | 1 |
| Twf1 | -0.04818 | 0.382678 | 1 |
| Gtf2a2 | -0.04819 | 0.084274 | 1 |
| St3gal3 | -0.04821 | 0.04869 | 1 |
| Uba7 | -0.04821 | 0.061718 | 1 |
| Pde4dip | -0.04822 | 0.109687 | 1 |
| Xiap | -0.04823 | 0.031137 | 1 |
| Gm46367 | -0.04827 | 0.383515 | 1 |
| Stxbp4 | -0.04828 | 0.002214 | 1 |
| Txnrd1 | -0.04828 | 0.628445 | 1 |
| Zfp65 | -0.04836 | 0.000111 | 1 |
| Sult1a1 | -0.04844 | 0.061773 | 1 |
| Far1os | -0.04847 | 0.000898 | 1 |
| Tmcc1 | -0.04853 | 0.047882 | 1 |
| Pcbd2 | -0.04854 | 0.252304 | 1 |
| Notch4 | -0.04854 | 0.003613 | 1 |
| Nfam1 | -0.04857 | 0.035305 | 1 |
| Hectd1 | -0.04858 | 0.137574 | 1 |
| Lipe | -0.04858 | 0.049031 | 1 |
| Abca3 | -0.04867 | 0.014911 | 1 |
| Gm15564 | -0.04869 | 0.006302 | 1 |
| Fam120c | -0.0487 | 0.003815 | 1 |
| B3galt5 | -0.04879 | 0.043272 | 1 |
| Stimate | -0.04882 | 0.3368 | 1 |
| 2510039O18Rik | -0.04887 | 0.068029 | 1 |
| Lypla1 | -0.04892 | 0.059491 | 1 |
| Rnf41 | -0.04894 | 0.011666 | 1 |
| Ubap2l | -0.04897 | 0.014371 | 1 |
| Rab5c | -0.04897 | 0.157226 | 1 |
| Miip | -0.04897 | 0.014203 | 1 |
| Pde1b | -0.049 | 0.009082 | 1 |
| Gigyf1 | -0.04902 | 0.160792 | 1 |
| Ifi35 | -0.04903 | 0.119308 | 1 |
| Tbc1d20 | -0.04905 | 0.033847 | 1 |
| Ago3 | -0.04908 | 0.084609 | 1 |
| Wdfy4 | -0.0491 | 0.046714 | 1 |
| Pxk | -0.04911 | 0.116571 | 1 |
| Tm9sf4 | -0.04913 | 0.087442 | 1 |
| 2610507B11Rik | -0.04917 | 0.338792 | 1 |
| Prmt7 | -0.04919 | 0.068823 | 1 |
| Brip1 | -0.04921 | 0.024507 | 1 |
| Zmym5 | -0.04922 | 0.015318 | 1 |

| | | | |
|---|---|---|---|
| Psd3 | -0.04922 | 0.050466 | 1 |
| Ppp1r15b | -0.04925 | 0.088874 | 1 |
| Rttn | -0.04928 | 0.03286 | 1 |
| Rest | -0.0493 | 0.02508 | 1 |
| 1500004A13Rik | -0.04933 | 0.10502 | 1 |
| Gpbp1l1 | -0.04933 | 0.009942 | 1 |
| Ncoa2 | -0.04934 | 0.146136 | 1 |
| Zfp532 | -0.04935 | 0.090708 | 1 |
| Pdhx | -0.04937 | 0.003788 | 1 |
| Elk4 | -0.04941 | 0.061467 | 1 |
| Lgals8 | -0.04943 | 0.117721 | 1 |
| Hmox1 | -0.04945 | 0.316709 | 1 |
| Slc24a3 | -0.04947 | 0.256626 | 1 |
| Mapkapk2 | -0.04949 | 0.343812 | 1 |
| Rapgef6 | -0.04954 | 0.165457 | 1 |
| Ccdc88b | -0.04955 | 0.03396 | 1 |
| Snta1 | -0.04958 | 0.016878 | 1 |
| Phf14 | -0.0496 | 0.044345 | 1 |
| Gm10974 | -0.0496 | 0.000522 | 1 |
| Sos1 | -0.04962 | 0.197178 | 1 |
| A930015D03Rik | -0.04965 | 0.029153 | 1 |
| Vamp5 | -0.04968 | 0.013942 | 1 |
| Amz1 | -0.0497 | 0.507155 | 1 |
| Anapc15 | -0.04971 | 0.237418 | 1 |
| Tab2 | -0.04972 | 0.076265 | 1 |
| Mad1l1 | -0.04972 | 0.062497 | 1 |
| Znrf2 | -0.04974 | 0.107214 | 1 |
| Tpcn2 | -0.04977 | 0.002945 | 1 |
| Ngly1 | -0.0498 | 0.043401 | 1 |
| Nfyc | -0.04983 | 0.079059 | 1 |
| Pecr | -0.04987 | 0.180404 | 1 |
| Zfp282 | -0.04992 | 0.045484 | 1 |
| Sh3pxd2a | -0.05003 | 0.073947 | 1 |
| Rab3gap2 | -0.05003 | 0.137291 | 1 |
| Trim26 | -0.05004 | 0.092905 | 1 |
| Chordc1 | -0.05004 | 0.063248 | 1 |
| Tec | -0.05005 | 0.123706 | 1 |
| 2610203C22Rik | -0.05005 | 0.181814 | 1 |
| Ltn1 | -0.05005 | 0.019193 | 1 |
| Zfp592 | -0.05005 | 0.120551 | 1 |
| Zfp871 | -0.05006 | 0.062291 | 1 |
| Cep290 | -0.05007 | 0.01825 | 1 |
| Wdr41 | -0.05013 | 0.032616 | 1 |
| Elavl4 | -0.05015 | 0.028801 | 1 |
| Ap1g1 | -0.05025 | 0.133158 | 1 |
| Gkap1 | -0.05026 | 0.023799 | 1 |
| Grk2 | -0.05028 | 0.015776 | 1 |
| Chchd4 | -0.05029 | 0.017825 | 1 |

| | | | |
|---|---|---|---|
| Rai14 | -0.05033 | 0.094185 | 1 |
| Rb1 | -0.05033 | 0.108874 | 1 |
| Rab21 | -0.05038 | 0.076926 | 1 |
| Kdm6a | -0.05049 | 0.196826 | 1 |
| Zfp866 | -0.05049 | 0.091214 | 1 |
| Gvin1 | -0.0505 | 0.036181 | 1 |
| Pbrm1 | -0.05053 | 0.119389 | 1 |
| Klhl20 | -0.05053 | 0.021548 | 1 |
| Ehmt1 | -0.05057 | 0.073225 | 1 |
| Hira | -0.05061 | 0.18322 | 1 |
| Zfp292 | -0.05064 | 0.022556 | 1 |
| Zfp945 | -0.05065 | 0.04149 | 1 |
| Ubap2 | -0.05068 | 0.020915 | 1 |
| Bbs9 | -0.0507 | 0.065038 | 1 |
| Lonp2 | -0.05071 | 0.097028 | 1 |
| N4bp2l2 | -0.05071 | 0.272347 | 1 |
| Csnk1a1 | -0.05077 | 0.074936 | 1 |
| Gpr107 | -0.0508 | 0.247226 | 1 |
| Slc25a20 | -0.05091 | 0.048344 | 1 |
| Arl5a | -0.05097 | 0.021984 | 1 |
| Kdm5c | -0.05097 | 0.173151 | 1 |
| Atf2 | -0.05099 | 0.080555 | 1 |
| Tpr | -0.051 | 0.872594 | 1 |
| Dcaf12 | -0.05101 | 0.043807 | 1 |
| C5ar2 | -0.05103 | 0.03194 | 1 |
| Polr2c | -0.05106 | 0.191396 | 1 |
| Gm42601 | -0.05107 | 0.13207 | 1 |
| Arhgap45 | -0.05116 | 0.048992 | 1 |
| Msrb1 | -0.05117 | 0.003121 | 1 |
| Egln1 | -0.05118 | 0.166936 | 1 |
| Asb3 | -0.0512 | 0.280466 | 1 |
| Jak1 | -0.05121 | 0.108406 | 1 |
| Max | -0.05124 | 0.169643 | 1 |
| Smg6 | -0.05127 | 0.039358 | 1 |
| Dexi | -0.05129 | 0.000928 | 1 |
| Bdp1 | -0.0513 | 0.140032 | 1 |
| Cep70 | -0.05144 | 0.006354 | 1 |
| Gm15563 | -0.05144 | 0.274877 | 1 |
| Tpi1 | -0.05147 | 0.308924 | 1 |
| Gmcl1 | -0.05153 | 0.023787 | 1 |
| Gabpb2 | -0.05157 | 0.067081 | 1 |
| Gm44148 | -0.0516 | 0.001469 | 1 |
| Khdrbs1 | -0.05161 | 0.264301 | 1 |
| Gatad2b | -0.05162 | 0.139841 | 1 |
| Jpx | -0.05167 | 0.001476 | 1 |
| Ppme1 | -0.05168 | 0.019758 | 1 |
| Pum2 | -0.05175 | 0.120979 | 1 |
| Cntrl | -0.05178 | 0.147124 | 1 |

| | | | |
|---|---|---|---|
| Cdc14a | -0.05178 | 0.070427 | 1 |
| Herc3 | -0.0518 | 0.041101 | 1 |
| Avl9 | -0.0518 | 0.157376 | 1 |
| Klhl2 | -0.05181 | 0.014615 | 1 |
| Zmynd11 | -0.05182 | 0.356511 | 1 |
| Entpd1 | -0.05182 | 0.035367 | 1 |
| Abhd15 | -0.0519 | 0.078972 | 1 |
| Zscan21 | -0.05191 | 0.004519 | 1 |
| Dcun1d1 | -0.05194 | 0.077024 | 1 |
| Cog5 | -0.05194 | 0.060011 | 1 |
| Zcchc17 | -0.05196 | 0.05786 | 1 |
| Qrich1 | -0.05198 | 0.018456 | 1 |
| Ube2j2 | -0.05201 | 0.295243 | 1 |
| Atxn7l1 | -0.05207 | 0.153793 | 1 |
| Smyd4 | -0.05208 | 0.042564 | 1 |
| Luc7l3 | -0.05208 | 0.321722 | 1 |
| Wapl | -0.05209 | 0.010816 | 1 |
| Glce | -0.05213 | 0.002669 | 1 |
| Kif13b | -0.05215 | 0.016197 | 1 |
| Unkl | -0.05218 | 0.349211 | 1 |
| Immp1l | -0.05219 | 0.016272 | 1 |
| Med21 | -0.05223 | 0.037978 | 1 |
| Gm10134 | -0.05224 | 0.000817 | 1 |
| Chd2 | -0.05226 | 0.331704 | 1 |
| AW146154 | -0.05227 | 0.006153 | 1 |
| Shld1 | -0.05228 | 0.096306 | 1 |
| Stat3 | -0.05232 | 0.024157 | 1 |
| Rnf216 | -0.05233 | 0.006917 | 1 |
| Mre11a | -0.05235 | 0.024901 | 1 |
| Upk1b | -0.05237 | 0.021387 | 1 |
| Ptgr2 | -0.05237 | 0.00443 | 1 |
| Kif16b | -0.05242 | 0.160168 | 1 |
| Nek7 | -0.05242 | 0.00464 | 1 |
| Chd7 | -0.05244 | 0.014431 | 1 |
| S100pbp | -0.05248 | 0.032782 | 1 |
| Rasa2 | -0.05259 | 0.121924 | 1 |
| Abr | -0.05262 | 0.084365 | 1 |
| Wdr47 | -0.05262 | 0.041817 | 1 |
| Tada3 | -0.05266 | 0.03367 | 1 |
| Gm7072 | -0.05274 | 0.032685 | 1 |
| Cdk12 | -0.05274 | 0.020055 | 1 |
| Cpeb3 | -0.05279 | 0.059474 | 1 |
| Strada | -0.05281 | 0.0249 | 1 |
| Appl1 | -0.05284 | 0.143162 | 1 |
| Gm20045 | -0.05285 | 0.001245 | 1 |
| Asb13 | -0.05286 | 0.001605 | 1 |
| Wls | -0.05287 | 0.055057 | 1 |
| Fam111a | -0.05288 | 0.012581 | 1 |

| | | | |
|---|---|---|---|
| Acap2 | -0.05289 | 0.024054 | 1 |
| Bach2os | -0.0529 | 0.002411 | 1 |
| Ogt | -0.05293 | 0.255875 | 1 |
| Hnrnpa0 | -0.05295 | 0.055337 | 1 |
| Wdr44 | -0.05295 | 0.209207 | 1 |
| Plaat3 | -0.05298 | 0.051488 | 1 |
| Zfc3h1 | -0.05298 | 0.040245 | 1 |
| Ifih1 | -0.05301 | 0.027093 | 1 |
| Snx14 | -0.05302 | 0.008926 | 1 |
| Tmem243 | -0.05307 | 0.073404 | 1 |
| Prim2 | -0.05311 | 0.08589 | 1 |
| Wdr59 | -0.05311 | 5.99E-05 | 1 |
| Nlrp1a | -0.05316 | 0.029222 | 1 |
| Tor3a | -0.05316 | 0.031169 | 1 |
| Crlf3 | -0.05319 | 0.030175 | 1 |
| Gdap2 | -0.05319 | 0.004058 | 1 |
| Man2c1os | -0.05324 | 0.022745 | 1 |
| Nck1 | -0.05325 | 0.120912 | 1 |
| Otud4 | -0.05326 | 0.009912 | 1 |
| Rsbn1l | -0.05329 | 0.094732 | 1 |
| Arfgef1 | -0.05331 | 0.04555 | 1 |
| Nsmaf | -0.05336 | 0.066772 | 1 |
| Trpv2 | -0.05336 | 0.058715 | 1 |
| Baz2b | -0.05346 | 0.110151 | 1 |
| Gtf2ird2 | -0.05348 | 0.014841 | 1 |
| Rab7 | -0.05357 | 0.009748 | 1 |
| Faf1 | -0.05361 | 0.013771 | 1 |
| Pilra | -0.05365 | 0.305755 | 1 |
| Trappc8 | -0.05368 | 0.020683 | 1 |
| Rit1 | -0.0537 | 0.001342 | 1 |
| Taok1 | -0.05374 | 0.052808 | 1 |
| Med13l | -0.05378 | 0.074666 | 1 |
| Gapvd1 | -0.05378 | 0.043497 | 1 |
| Scmh1 | -0.05379 | 0.462334 | 1 |
| Zfp407 | -0.0538 | 0.194177 | 1 |
| Ubac2 | -0.0538 | 0.095955 | 1 |
| Nrp2 | -0.05381 | 0.866666 | 1 |
| Ube4b | -0.05381 | 0.072531 | 1 |
| Tcf7l2 | -0.05382 | 0.018631 | 1 |
| Tnrc6a | -0.05384 | 0.075885 | 1 |
| Ppp2r5e | -0.0539 | 0.107809 | 1 |
| Zc3h7b | -0.05392 | 0.032945 | 1 |
| Psma6 | -0.05392 | 0.098333 | 1 |
| Bcat2 | -0.05403 | 0.011274 | 1 |
| Zmat5 | -0.05411 | 0.00466 | 1 |
| Ap2a2 | -0.05415 | 0.009653 | 1 |
| Dhx8 | -0.05418 | 0.203107 | 1 |
| Osbpl9 | -0.05418 | 0.19458 | 1 |

| | | | |
|---|---|---|---|
| Tmem184b | -0.0542 | 0.037374 | 1 |
| Smarcc1 | -0.0542 | 0.132689 | 1 |
| Soat1 | -0.0542 | 0.316141 | 1 |
| Afap1 | -0.0542 | 0.001592 | 1 |
| Ankrd44 | -0.05434 | 0.022516 | 1 |
| Paox | -0.05434 | 0.003026 | 1 |
| Nxpe3 | -0.05434 | 0.028249 | 1 |
| Rfc2 | -0.05436 | 0.06125 | 1 |
| Slc25a38 | -0.05439 | 0.008222 | 1 |
| Neurl1a | -0.0544 | 0.015421 | 1 |
| Iars | -0.05441 | 0.034639 | 1 |
| Rc3h1 | -0.05442 | 0.088858 | 1 |
| Lap3 | -0.05442 | 0.016607 | 1 |
| Nabp1 | -0.05445 | 0.003231 | 1 |
| Zfp951 | -0.05452 | 0.038486 | 1 |
| Rogdi | -0.05454 | 0.213617 | 1 |
| Gns | -0.05455 | 0.005081 | 1 |
| Usp9x | -0.05456 | 0.068126 | 1 |
| Ube2l3 | -0.05459 | 0.15073 | 1 |
| Rnf170 | -0.0546 | 0.002487 | 1 |
| Btbd9 | -0.05462 | 0.063286 | 1 |
| Gm43821 | -0.05465 | 0.021683 | 1 |
| Srbd1 | -0.05465 | 0.040662 | 1 |
| Spen | -0.05479 | 0.101606 | 1 |
| Acbd3 | -0.05483 | 0.019523 | 1 |
| Pafah1b1 | -0.05485 | 0.014301 | 1 |
| Gpr137b | -0.05491 | 0.0161 | 1 |
| Zfp703 | -0.05491 | 0.01611 | 1 |
| C87436 | -0.05493 | 0.058641 | 1 |
| Dlg1 | -0.05497 | 0.397354 | 1 |
| Virma | -0.05497 | 0.008285 | 1 |
| Nacc1 | -0.05498 | 0.000357 | 1 |
| Htt | -0.055 | 0.014014 | 1 |
| Sec23b | -0.05501 | 0.010868 | 1 |
| Mtf1 | -0.05512 | 0.012277 | 1 |
| Jmjd6 | -0.05513 | 0.012062 | 1 |
| Hipk1 | -0.05517 | 0.031277 | 1 |
| Larp4b | -0.0552 | 0.147274 | 1 |
| Jdp2 | -0.0552 | 0.078775 | 1 |
| Suco | -0.05525 | 0.199429 | 1 |
| Camta1 | -0.05532 | 0.057695 | 1 |
| Picalm | -0.05533 | 0.01417 | 1 |
| Vps13c | -0.05533 | 0.047247 | 1 |
| Arhgef11 | -0.05538 | 0.045331 | 1 |
| Gse1 | -0.05539 | 0.013676 | 1 |
| Sorl1 | -0.0554 | 0.161677 | 1 |
| Thada | -0.05542 | 0.036545 | 1 |
| 2700049A03Rik | -0.05547 | 0.034169 | 1 |

| | | | |
|---|---|---|---|
| Tg | -0.05551 | 0.005584 | 1 |
| Upf2 | -0.05552 | 0.056267 | 1 |
| Acbd6 | -0.05555 | 0.059025 | 1 |
| 1110051M20Rik | -0.05558 | 0.156914 | 1 |
| Mapk1 | -0.05559 | 0.015536 | 1 |
| Kifap3 | -0.05563 | 0.097488 | 1 |
| Tfeb | -0.05566 | 0.033003 | 1 |
| BC035044 | -0.05567 | 0.00438 | 1 |
| Ino80d | -0.05568 | 0.003576 | 1 |
| Ankle2 | -0.05568 | 0.020264 | 1 |
| Fbxl20 | -0.05572 | 0.028807 | 1 |
| Dnaja3 | -0.05577 | 0.013295 | 1 |
| Ankrd12 | -0.0558 | 0.04169 | 1 |
| Hgf | -0.05587 | 0.297933 | 1 |
| Ccdc191 | -0.05588 | 0.002568 | 1 |
| Nup155 | -0.05588 | 0.026681 | 1 |
| Amn1 | -0.05602 | 0.090199 | 1 |
| Meis1 | -0.05604 | 0.049331 | 1 |
| Sepsecs | -0.0561 | 0.098382 | 1 |
| Fam117b | -0.05614 | 0.277033 | 1 |
| Taco1 | -0.05624 | 0.008612 | 1 |
| Ddx3y | -0.05633 | 0.001282 | 1 |
| Phtf2 | -0.05638 | 0.036957 | 1 |
| Patz1 | -0.05642 | 0.001002 | 1 |
| Arf3 | -0.05648 | 0.004305 | 1 |
| Cfdp1 | -0.05651 | 0.490274 | 1 |
| Nufip2 | -0.05652 | 0.208739 | 1 |
| Vps45 | -0.05652 | 0.015426 | 1 |
| Pdss2 | -0.05657 | 0.033979 | 1 |
| Igsf6 | -0.05658 | 0.114262 | 1 |
| Btk | -0.0566 | 0.081129 | 1 |
| Gm40841 | -0.05661 | 0.187566 | 1 |
| Osbpl7 | -0.05662 | 0.012012 | 1 |
| 4932438A13Rik | -0.0567 | 0.04648 | 1 |
| Acot9 | -0.05672 | 0.009195 | 1 |
| Atxn2 | -0.05677 | 0.191605 | 1 |
| Ikzf1 | -0.05682 | 0.004058 | 1 |
| Upp2 | -0.05682 | 0.006483 | 1 |
| Supt3 | -0.05684 | 0.101964 | 1 |
| Zfyve26 | -0.05684 | 0.023145 | 1 |
| Atg4a | -0.05685 | 0.092346 | 1 |
| AY512915 | -0.05686 | 0.046293 | 1 |
| Dnajc7 | -0.05695 | 0.028395 | 1 |
| Rabl6 | -0.05699 | 0.04888 | 1 |
| Ppp4r3a | -0.057 | 0.074827 | 1 |
| Plcl2 | -0.05701 | 0.060537 | 1 |
| Hace1 | -0.05702 | 0.006163 | 1 |
| AB124611 | -0.05705 | 0.076123 | 1 |

| | | | |
|---|---|---|---|
| Tmod3 | -0.05706 | 0.129225 | 1 |
| Zfyve28 | -0.05714 | 0.089271 | 1 |
| Nt5m | -0.05715 | 0.016056 | 1 |
| Kat14 | -0.05719 | 0.006554 | 1 |
| Gm4258 | -0.0572 | 0.209656 | 1 |
| Ikbkg | -0.05725 | 0.006469 | 1 |
| Uimc1 | -0.05726 | 0.122207 | 1 |
| Tmem140 | -0.0573 | 0.078106 | 1 |
| Tmcc3 | -0.05731 | 0.067705 | 1 |
| Vav1 | -0.0574 | 0.019485 | 1 |
| 5031425E22Rik | -0.0574 | 0.111183 | 1 |
| Maea | -0.05742 | 0.135988 | 1 |
| Gm2000 | -0.05742 | 0.017997 | 1 |
| Ubr4 | -0.05743 | 0.059672 | 1 |
| Ppm1k | -0.05744 | 0.001112 | 1 |
| Rbms1 | -0.05745 | 0.091342 | 1 |
| Fchsd2 | -0.05751 | 0.063623 | 1 |
| Slc35a3 | -0.05772 | 0.076175 | 1 |
| Safb2 | -0.05772 | 0.039843 | 1 |
| Tra2a | -0.05776 | 0.018586 | 1 |
| Myo1g | -0.05779 | 0.007071 | 1 |
| Stim2 | -0.0578 | 0.002433 | 1 |
| Dmxl1 | -0.0578 | 0.032326 | 1 |
| Dnajc5 | -0.05788 | 0.034172 | 1 |
| Nfatc1 | -0.0579 | 0.03423 | 1 |
| Pcsk7 | -0.05791 | 0.018374 | 1 |
| Catsper2 | -0.05792 | 0.021646 | 1 |
| Sae1 | -0.05794 | 0.224224 | 1 |
| Tcirg1 | -0.05795 | 0.014716 | 1 |
| Slc30a7 | -0.05796 | 0.001486 | 1 |
| Pi4ka | -0.05798 | 0.010412 | 1 |
| Ppfia1 | -0.058 | 0.035799 | 1 |
| Rnmt | -0.05815 | 0.015428 | 1 |
| Smad3 | -0.05817 | 0.334677 | 1 |
| Lactb2 | -0.05817 | 0.589932 | 1 |
| 2700097O09Rik | -0.05818 | 0.015188 | 1 |
| Atg4c | -0.05819 | 0.002776 | 1 |
| Ints3 | -0.05822 | 0.00366 | 1 |
| Herc1 | -0.05824 | 0.044007 | 1 |
| Ola1 | -0.05825 | 0.039253 | 1 |
| Pik3r5 | -0.05837 | 0.015588 | 1 |
| Rtn3 | -0.05839 | 0.095933 | 1 |
| Dpp8 | -0.05841 | 0.157791 | 1 |
| Gtf2i | -0.05845 | 0.000905 | 1 |
| Wars | -0.05846 | 0.059602 | 1 |
| Tent4b | -0.05852 | 0.064815 | 1 |
| Ift88 | -0.05854 | 0.051678 | 1 |
| Ppp3cc | -0.05854 | 0.015724 | 1 |

| | | | |
|---|---|---|---|
| Xpnpep1 | -0.05854 | 0.073076 | 1 |
| Arid1b | -0.05855 | 0.004429 | 1 |
| Casp4 | -0.05857 | 0.251542 | 1 |
| Galnt10 | -0.05863 | 0.002604 | 1 |
| Pla2g7 | -0.05866 | 0.003695 | 1 |
| Ifi207 | -0.05869 | 0.00469 | 1 |
| Elmsan1 | -0.05874 | 0.007149 | 1 |
| Pggt1b | -0.05878 | 0.073758 | 1 |
| Oas3 | -0.05879 | 0.002755 | 1 |
| Tmem170b | -0.0589 | 0.033805 | 1 |
| Arhgap18 | -0.05901 | 0.096954 | 1 |
| Map3k5 | -0.05907 | 0.012736 | 1 |
| Zfp512 | -0.05913 | 0.016942 | 1 |
| Hltf | -0.05916 | 0.041067 | 1 |
| Pikfyve | -0.05916 | 0.066389 | 1 |
| Slc37a3 | -0.05918 | 1.01E-05 | 0.326331 |
| Itpr1 | -0.05927 | 0.0007 | 1 |
| Xpr1 | -0.05929 | 0.039279 | 1 |
| Zmiz1 | -0.05932 | 0.016451 | 1 |
| Dnajb6 | -0.05934 | 0.009298 | 1 |
| Tbcd | -0.05937 | 0.025122 | 1 |
| Fam91a1 | -0.05938 | 0.053939 | 1 |
| Rcor1 | -0.05939 | 0.008204 | 1 |
| Foxn3 | -0.05944 | 0.000235 | 1 |
| Nisch | -0.05947 | 0.005495 | 1 |
| Gramd1a | -0.05948 | 0.115602 | 1 |
| Rbpms | -0.05951 | 0.057462 | 1 |
| Srpk1 | -0.05953 | 0.018505 | 1 |
| Tmem126a | -0.05954 | 0.004388 | 1 |
| Cyld | -0.05957 | 0.291932 | 1 |
| Sumo1 | -0.05958 | 0.018969 | 1 |
| Slc9a8 | -0.05961 | 0.168452 | 1 |
| Erbin | -0.05961 | 0.022008 | 1 |
| Stx18 | -0.05961 | 0.119038 | 1 |
| Cnot2 | -0.05966 | 0.033289 | 1 |
| E230029C05Rik | -0.05972 | 0.030152 | 1 |
| Mtr | -0.05985 | 0.029814 | 1 |
| Shtn1 | -0.05987 | 0.01097 | 1 |
| Zdhhc6 | -0.05991 | 0.012204 | 1 |
| Dip2c | -0.05998 | 0.101343 | 1 |
| Slc38a7 | -0.06001 | 0.009895 | 1 |
| Chic2 | -0.06001 | 0.001853 | 1 |
| Hk2 | -0.06003 | 0.012529 | 1 |
| Ncoa1 | -0.06009 | 0.009476 | 1 |
| Dstyk | -0.06016 | 0.031988 | 1 |
| Mafg | -0.06018 | 0.198449 | 1 |
| Il18 | -0.06018 | 0.001788 | 1 |
| Scd2 | -0.06019 | 3.28E-05 | 1 |

| | | | |
|---|---|---|---|
| 4931406P16Rik | -0.06037 | 0.309037 | 1 |
| Mdfic | -0.06039 | 0.033491 | 1 |
| Sp3 | -0.06042 | 0.005624 | 1 |
| Pgap1 | -0.06042 | 0.012528 | 1 |
| Sass6 | -0.06046 | 0.00539 | 1 |
| Lrp6 | -0.06047 | 0.032691 | 1 |
| Prrc2a | -0.0605 | 3.13E-05 | 1 |
| Efl1 | -0.06052 | 0.002184 | 1 |
| Ankrd11 | -0.06055 | 0.001057 | 1 |
| Zmat1 | -0.06061 | 0.012417 | 1 |
| Mark3 | -0.06061 | 0.052257 | 1 |
| Slain2 | -0.06064 | 0.033599 | 1 |
| D130040H23Rik | -0.06066 | 0.024267 | 1 |
| Bcl2a1d | -0.06067 | 0.044185 | 1 |
| Cpped1 | -0.06072 | 0.053853 | 1 |
| Apaf1 | -0.06072 | 0.165652 | 1 |
| Cd40 | -0.06073 | 0.068039 | 1 |
| Tbc1d13 | -0.06075 | 0.07359 | 1 |
| D430042O09Rik | -0.06079 | 0.002857 | 1 |
| Map2k5 | -0.0608 | 0.019806 | 1 |
| Tasor2 | -0.06084 | 0.025411 | 1 |
| Inpp4a | -0.06085 | 0.068741 | 1 |
| Gm20470 | -0.06087 | 0.002531 | 1 |
| Tlr4 | -0.06089 | 0.004146 | 1 |
| Dis3l2 | -0.0609 | 0.024554 | 1 |
| Nsd3 | -0.06093 | 0.012384 | 1 |
| Dennd1c | -0.06093 | 0.16067 | 1 |
| Usp35 | -0.061 | 0.124824 | 1 |
| Sesn1 | -0.06108 | 0.001226 | 1 |
| Atf6 | -0.06108 | 0.063631 | 1 |
| Arid5a | -0.0611 | 0.000976 | 1 |
| Zfyve27 | -0.06113 | 0.008288 | 1 |
| Tmem209 | -0.06114 | 0.007585 | 1 |
| Thoc2 | -0.0612 | 0.005477 | 1 |
| Xpo6 | -0.06121 | 0.007666 | 1 |
| Fam193b | -0.06124 | 0.013621 | 1 |
| Lin54 | -0.06128 | 0.004924 | 1 |
| Drg1 | -0.06134 | 0.002073 | 1 |
| Rnf180 | -0.06151 | 0.022698 | 1 |
| Urgcp | -0.06154 | 0.023978 | 1 |
| Mkln1 | -0.06158 | 0.002939 | 1 |
| Mbtd1 | -0.06158 | 0.071098 | 1 |
| Xrn1 | -0.06159 | 0.008251 | 1 |
| Atrnl1 | -0.06159 | 0.032491 | 1 |
| Umad1 | -0.06162 | 0.010142 | 1 |
| Iqsec2 | -0.06168 | 0.002382 | 1 |
| Slc4a2 | -0.0617 | 0.000163 | 1 |
| Mprip | -0.06172 | 0.029514 | 1 |

| | | | |
|---|---|---|---|
| Apbb2 | -0.06184 | 0.054539 | 1 |
| Pdk1 | -0.06184 | 0.009804 | 1 |
| Gm5086 | -0.06186 | 0.482642 | 1 |
| Trmt1l | -0.06187 | 0.001038 | 1 |
| Rps6ka1 | -0.06189 | 0.010457 | 1 |
| Micu2 | -0.0619 | 0.038068 | 1 |
| Ubr5 | -0.06195 | 0.023525 | 1 |
| Stx12 | -0.06196 | 0.086022 | 1 |
| Nceh1 | -0.06201 | 0.001469 | 1 |
| Gna12 | -0.06202 | 0.153725 | 1 |
| Mysm1 | -0.06204 | 0.115486 | 1 |
| Tnpo1 | -0.06211 | 0.014261 | 1 |
| Litaf | -0.06213 | 0.069159 | 1 |
| Msh3 | -0.06213 | 0.1102 | 1 |
| Tsc22d3 | -0.06214 | 0.000818 | 1 |
| Dcaf5 | -0.06214 | 0.087485 | 1 |
| Rhot1 | -0.06218 | 0.010326 | 1 |
| Rapgef1 | -0.06225 | 0.002084 | 1 |
| Lnpep | -0.06226 | 0.021993 | 1 |
| Sgk3 | -0.06226 | 0.087496 | 1 |
| Gtpbp2 | -0.06229 | 0.071328 | 1 |
| Gng12 | -0.06229 | 0.042614 | 1 |
| Trpc4ap | -0.06235 | 0.018651 | 1 |
| Fto | -0.06238 | 0.060016 | 1 |
| Etv3 | -0.0624 | 0.021171 | 1 |
| Adhfe1 | -0.06241 | 9.34E-05 | 1 |
| Snx24 | -0.06242 | 0.17981 | 1 |
| Magi3 | -0.06244 | 0.001265 | 1 |
| Mbp | -0.06245 | 0.096483 | 1 |
| Cdk17 | -0.06257 | 0.066606 | 1 |
| Usp7 | -0.06258 | 0.024622 | 1 |
| Ehbp1 | -0.06264 | 0.052202 | 1 |
| Mtmr3 | -0.06268 | 0.023816 | 1 |
| Ift140 | -0.06269 | 0.00353 | 1 |
| Tacc1 | -0.06271 | 0.00269 | 1 |
| Gm4841 | -0.06276 | 0.001465 | 1 |
| H2-Oa | -0.06277 | 0.022327 | 1 |
| Tk2 | -0.06283 | 0.002831 | 1 |
| Nras | -0.06286 | 0.171769 | 1 |
| Stmn1 | -0.06289 | 0.33757 | 1 |
| Akt3 | -0.063 | 0.327989 | 1 |
| Vrk2 | -0.06302 | 0.00361 | 1 |
| Slk | -0.06305 | 0.004267 | 1 |
| Cdc42bpb | -0.06306 | 0.024943 | 1 |
| Nod2 | -0.06311 | 0.000302 | 1 |
| Rnf115 | -0.06315 | 0.025081 | 1 |
| Gm37240 | -0.06316 | 0.024264 | 1 |
| Srpk2 | -0.06318 | 0.170072 | 1 |

| | | | |
|---|---|---|---|
| Wdr33 | -0.06321 | 0.039685 | 1 |
| Gbp3 | -0.06322 | 0.017783 | 1 |
| Dym | -0.06324 | 0.041514 | 1 |
| Nemf | -0.06328 | 0.065062 | 1 |
| Tep1 | -0.06329 | 0.000417 | 1 |
| Mavs | -0.06331 | 0.000397 | 1 |
| Abcc4 | -0.06335 | 0.006069 | 1 |
| Nrip1 | -0.06335 | 0.004792 | 1 |
| Sbno2 | -0.0634 | 0.01204 | 1 |
| Pten | -0.0634 | 0.03051 | 1 |
| B4galt4 | -0.06343 | 0.037375 | 1 |
| Itgal | -0.06344 | 0.001086 | 1 |
| Il15 | -0.06347 | 0.049024 | 1 |
| Vapb | -0.06352 | 0.015135 | 1 |
| Tyk2 | -0.06352 | 0.010477 | 1 |
| Btrc | -0.06357 | 0.035335 | 1 |
| Cep85 | -0.06369 | 0.002249 | 1 |
| Tceanc2 | -0.06372 | 0.001631 | 1 |
| Gm16124 | -0.06376 | 0.014381 | 1 |
| 3830406C13Rik | -0.06379 | 0.003761 | 1 |
| Utrn | -0.06386 | 0.015478 | 1 |
| Nedd4l | -0.06388 | 0.020086 | 1 |
| Ambra1 | -0.06395 | 0.000666 | 1 |
| Ikbke | -0.06395 | 1.6E-05 | 0.516215 |
| Gm16337 | -0.06396 | 0.002106 | 1 |
| Ppil2 | -0.06405 | 0.003811 | 1 |
| Isg15 | -0.06413 | 0.045841 | 1 |
| Siglecg | -0.06419 | 0.002801 | 1 |
| Dclre1c | -0.0642 | 0.247714 | 1 |
| Gm42962 | -0.06424 | 0.048692 | 1 |
| Fbh1 | -0.06428 | 0.008065 | 1 |
| Cdkn1a | -0.06433 | 0.021328 | 1 |
| Mettl15 | -0.06437 | 0.028465 | 1 |
| Def6 | -0.0644 | 0.006891 | 1 |
| Dync1h1 | -0.06442 | 0.026111 | 1 |
| Zfp652 | -0.06444 | 0.022394 | 1 |
| Hectd4 | -0.06452 | 0.055596 | 1 |
| Akap13 | -0.06458 | 0.003432 | 1 |
| Iars2 | -0.06461 | 0.023593 | 1 |
| Gm40645 | -0.06461 | 0.012004 | 1 |
| Metap1 | -0.06469 | 1.55E-05 | 0.499636 |
| Crlf2 | -0.06473 | 0.036908 | 1 |
| Samd9l | -0.06475 | 0.013648 | 1 |
| Stard9 | -0.06479 | 0.016125 | 1 |
| Map3k3 | -0.06479 | 0.001891 | 1 |
| Pex13 | -0.0648 | 0.001749 | 1 |
| Tlr2 | -0.06489 | 0.006792 | 1 |
| 2210408F21Rik | -0.0649 | 0.003597 | 1 |

| | | | |
|---|---|---|---|
| Atxn1 | -0.06491 | 0.028818 | 1 |
| Hipk2 | -0.06497 | 0.062829 | 1 |
| Dnajb14 | -0.06497 | 0.235619 | 1 |
| Spred2 | -0.06499 | 0.043127 | 1 |
| Strn | -0.06501 | 0.089649 | 1 |
| Cd80 | -0.06501 | 0.003455 | 1 |
| Abcd2 | -0.06501 | 0.238906 | 1 |
| Nfia | -0.06504 | 0.01026 | 1 |
| F830016B08Rik | -0.06504 | 0.041322 | 1 |
| Lrmp | -0.06506 | 0.031849 | 1 |
| 2210408I21Rik | -0.06507 | 0.00561 | 1 |
| Slc9a7 | -0.0651 | 0.008127 | 1 |
| Ube2h | -0.0651 | 0.082108 | 1 |
| Nlk | -0.06512 | 0.010763 | 1 |
| St6galnac6 | -0.06514 | 0.00125 | 1 |
| Rabgap1 | -0.06521 | 0.024899 | 1 |
| Flnb | -0.06524 | 0.046276 | 1 |
| Rab11fip2 | -0.06526 | 0.006957 | 1 |
| Prex1 | -0.06531 | 0.00364 | 1 |
| Csnk2a2 | -0.06532 | 0.011952 | 1 |
| Zfp934 | -0.06534 | 0.013661 | 1 |
| Rffl | -0.06535 | 0.092886 | 1 |
| Zfp451 | -0.06545 | 0.032936 | 1 |
| Gpbp1 | -0.06548 | 0.01176 | 1 |
| Mnat1 | -0.06553 | 0.069509 | 1 |
| Stard3nl | -0.06553 | 0.016158 | 1 |
| Nup214 | -0.06554 | 0.034824 | 1 |
| Ifnar1 | -0.06556 | 0.005743 | 1 |
| B3galnt1 | -0.06558 | 0.01909 | 1 |
| D530033B14Rik | -0.0656 | 0.014695 | 1 |
| Pde4b | -0.06563 | 0.011259 | 1 |
| Btbd7 | -0.06575 | 0.033829 | 1 |
| Tbck | -0.06583 | 0.279738 | 1 |
| Cers5 | -0.066 | 0.034935 | 1 |
| Il10ra | -0.066 | 0.000196 | 1 |
| Rbm27 | -0.06604 | 0.013292 | 1 |
| Itga9 | -0.06606 | 0.461753 | 1 |
| Nup98 | -0.06616 | 0.060862 | 1 |
| Kif1b | -0.06618 | 0.055354 | 1 |
| Fkbp15 | -0.06618 | 0.002595 | 1 |
| 6530409C15Rik | -0.06624 | 8.86E-05 | 1 |
| Ldah | -0.0663 | 0.018748 | 1 |
| Ifi204 | -0.06641 | 0.000322 | 1 |
| Stxbp3 | -0.06656 | 0.003364 | 1 |
| Bckdhb | -0.06657 | 0.013858 | 1 |
| Cspp1 | -0.06658 | 0.043076 | 1 |
| Kdm2a | -0.0666 | 0.020778 | 1 |
| Rhbdf2 | -0.06661 | 0.000356 | 1 |

| | | | |
|---|---|---|---|
| Txndc16 | -0.06663 | 0.154186 | 1 |
| Smurf1 | -0.06664 | 0.020368 | 1 |
| Lsp1 | -0.06665 | 0.014916 | 1 |
| Ptk2b | -0.06668 | 0.005684 | 1 |
| Cp | -0.06668 | 0.01108 | 1 |
| Togaram1 | -0.06676 | 0.091251 | 1 |
| Dcaf6 | -0.0668 | 0.00481 | 1 |
| Trrap | -0.06685 | 0.005348 | 1 |
| Sec24b | -0.06691 | 0.006204 | 1 |
| Rnf111 | -0.06692 | 0.156833 | 1 |
| Tnip1 | -0.06693 | 0.00082 | 1 |
| Gm13710 | -0.06694 | 0.053253 | 1 |
| Evl | -0.06694 | 0.002233 | 1 |
| Tnpo3 | -0.06702 | 0.003565 | 1 |
| Smarcc2 | -0.06705 | 0.019634 | 1 |
| Edem3 | -0.06707 | 0.050349 | 1 |
| Gm20429 | -0.06708 | 0.000252 | 1 |
| Slc12a6 | -0.06713 | 0.015377 | 1 |
| Heatr5a | -0.06721 | 0.001864 | 1 |
| Sort1 | -0.06722 | 0.036701 | 1 |
| P2ry14 | -0.06741 | 0.003844 | 1 |
| Ehmt2 | -0.06742 | 0.000404 | 1 |
| Stxbp1 | -0.06742 | 6.06E-06 | 0.195749 |
| Fam193a | -0.06746 | 0.0028 | 1 |
| Nudcd3 | -0.0675 | 0.002433 | 1 |
| Tasor | -0.06753 | 0.014522 | 1 |
| Chchd3 | -0.06753 | 0.002451 | 1 |
| Aida | -0.06759 | 0.00014 | 1 |
| Zmym6 | -0.06759 | 0.000499 | 1 |
| Ube2g1 | -0.06763 | 0.042934 | 1 |
| Paxbp1 | -0.06769 | 0.003051 | 1 |
| Tom1l2 | -0.06778 | 0.004837 | 1 |
| Ttc39b | -0.06779 | 0.028962 | 1 |
| Ap3b1 | -0.0679 | 0.003131 | 1 |
| Socs2 | -0.06793 | 0.303814 | 1 |
| Ttc28 | -0.06802 | 0.012575 | 1 |
| Tmem175 | -0.06803 | 0.025325 | 1 |
| Cmip | -0.06805 | 0.004632 | 1 |
| Eps15 | -0.06807 | 0.025099 | 1 |
| B430010I23Rik | -0.06813 | 0.503486 | 1 |
| Tmem241 | -0.06824 | 0.1362 | 1 |
| Mgrn1 | -0.06824 | 0.009 | 1 |
| Arhgef7 | -0.06826 | 0.00678 | 1 |
| C2cd3 | -0.06832 | 0.0006 | 1 |
| Irf8 | -0.06839 | 0.008531 | 1 |
| Ilrun | -0.06841 | 0.009526 | 1 |
| Ints6 | -0.06845 | 0.001625 | 1 |
| Zfp148 | -0.06858 | 0.000358 | 1 |

| | | | |
|---|---|---|---|
| Alpk1 | -0.06859 | 0.001119 | 1 |
| Xpo7 | -0.06864 | 0.016654 | 1 |
| Sppl3 | -0.06866 | 0.002814 | 1 |
| Aatf | -0.0687 | 0.053417 | 1 |
| Phf20 | -0.06876 | 0.007625 | 1 |
| Gm20528 | -0.06879 | 0.06439 | 1 |
| Chd4 | -0.06881 | 0.052591 | 1 |
| Mthfr | -0.06883 | 0.000127 | 1 |
| Gm34280 | -0.06885 | 0.000462 | 1 |
| A330023F24Rik | -0.0689 | 0.005963 | 1 |
| Tyw1 | -0.06894 | 0.006974 | 1 |
| Phka2 | -0.06894 | 0.007405 | 1 |
| Ncor2 | -0.06899 | 0.029237 | 1 |
| Themis2 | -0.069 | 0.005218 | 1 |
| Bmf | -0.069 | 0.001883 | 1 |
| Chd1 | -0.06903 | 0.084494 | 1 |
| Brd4 | -0.06903 | 0.002095 | 1 |
| Trim56 | -0.06908 | 0.000853 | 1 |
| Ric8b | -0.0691 | 0.000389 | 1 |
| Sap25 | -0.06913 | 0.002677 | 1 |
| Manba | -0.06915 | 0.031574 | 1 |
| Setd2 | -0.06926 | 0.001756 | 1 |
| Sh3bp2 | -0.06931 | 0.006932 | 1 |
| Cd2ap | -0.06951 | 0.001832 | 1 |
| Kidins220 | -0.06968 | 0.005286 | 1 |
| Gm16599 | -0.06971 | 0.066833 | 1 |
| Ier3 | -0.06977 | 0.345722 | 1 |
| Rsbn1 | -0.06985 | 0.005456 | 1 |
| Fbxo42 | -0.06986 | 0.008812 | 1 |
| Zfp160 | -0.06987 | 0.001744 | 1 |
| 2900089D17Rik | -0.06988 | 0.023561 | 1 |
| Ppard | -0.06988 | 0.035772 | 1 |
| Terf2 | -0.06993 | 0.007613 | 1 |
| Hcar2 | -0.07004 | 0.0325 | 1 |
| Cfb | -0.07004 | 0.027257 | 1 |
| Gm42477 | -0.07019 | 0.001115 | 1 |
| Smad1 | -0.0702 | 0.002406 | 1 |
| Ctif | -0.07029 | 0.000726 | 1 |
| Adcy7 | -0.07032 | 0.246236 | 1 |
| Adar | -0.07036 | 0.002575 | 1 |
| Mettl8 | -0.07044 | 0.001156 | 1 |
| Gm35154 | -0.07044 | 0.081508 | 1 |
| Herc4 | -0.07044 | 0.002467 | 1 |
| Zbtb5 | -0.07045 | 1.62E-05 | 0.522235 |
| Prr5l | -0.07045 | 0.037794 | 1 |
| Plxdc1 | -0.07048 | 0.007125 | 1 |
| Fam53b | -0.07052 | 0.000117 | 1 |
| Vps35l | -0.07052 | 0.017656 | 1 |

| | | | |
|---|---|---|---|
| Asb2 | -0.07054 | 0.007264 | 1 |
| Stk38 | -0.07056 | 0.013164 | 1 |
| Snx10 | -0.07056 | 0.002562 | 1 |
| Elp4 | -0.07058 | 0.017768 | 1 |
| Cd44 | -0.07058 | 0.000746 | 1 |
| Gramd1b | -0.07064 | 0.007633 | 1 |
| Rbm26 | -0.0707 | 0.047063 | 1 |
| Trim12c | -0.07073 | 0.040657 | 1 |
| Marf1 | -0.07083 | 0.013735 | 1 |
| Phf20l1 | -0.07085 | 0.066914 | 1 |
| Sptan1 | -0.07085 | 0.009587 | 1 |
| Zbtb38 | -0.07089 | 0.000141 | 1 |
| Gtpbp1 | -0.07099 | 0.029808 | 1 |
| Dido1 | -0.07109 | 0.005026 | 1 |
| Galnt1 | -0.07117 | 0.004818 | 1 |
| Ptprm | -0.07121 | 0.165925 | 1 |
| R3hcc1l | -0.07122 | 0.002078 | 1 |
| Nup93 | -0.07124 | 0.002621 | 1 |
| Fam49a | -0.0713 | 0.007793 | 1 |
| Dst | -0.07134 | 5.14E-05 | 1 |
| Numa1 | -0.07137 | 0.00377 | 1 |
| Nvl | -0.07137 | 0.016081 | 1 |
| Nubpl | -0.07137 | 0.077958 | 1 |
| Elf2 | -0.07145 | 0.307795 | 1 |
| Fli1 | -0.07149 | 0.006161 | 1 |
| Hps5 | -0.07149 | 0.003951 | 1 |
| Snx2 | -0.0715 | 0.019757 | 1 |
| Rab37 | -0.07161 | 0.008436 | 1 |
| Cep350 | -0.07177 | 0.002773 | 1 |
| Slfn9 | -0.07177 | 0.000346 | 1 |
| Mast4 | -0.07181 | 0.083956 | 1 |
| Mroh1 | -0.07183 | 0.001205 | 1 |
| Zfp438 | -0.07186 | 0.002685 | 1 |
| Sbno1 | -0.07188 | 0.102559 | 1 |
| Gng2 | -0.0719 | 0.003994 | 1 |
| Zc3h4 | -0.07196 | 0.006756 | 1 |
| Lpgat1 | -0.07199 | 0.065385 | 1 |
| Tdrd7 | -0.07207 | 0.001966 | 1 |
| Slc46a3 | -0.07207 | 0.054436 | 1 |
| Prorp | -0.07208 | 0.001289 | 1 |
| Stx8 | -0.07213 | 0.009822 | 1 |
| Pdpk1 | -0.07215 | 0.036674 | 1 |
| Morc3 | -0.07215 | 0.003515 | 1 |
| Dlgap4 | -0.07216 | 0.055694 | 1 |
| Ppm1e | -0.07217 | 0.003569 | 1 |
| Etv5 | -0.07217 | 0.002991 | 1 |
| Far1 | -0.07229 | 0.004649 | 1 |
| Socs7 | -0.07233 | 0.000688 | 1 |

| | | | |
|---|---|---|---|
| Smarcad1 | -0.07234 | 0.017533 | 1 |
| Tspan14 | -0.07244 | 0.00568 | 1 |
| Gm42047 | -0.07248 | 0.007778 | 1 |
| Lyn | -0.07249 | 1.33E-06 | 0.042966 |
| Bcas3 | -0.07264 | 0.00819 | 1 |
| Armc9 | -0.07267 | 0.008673 | 1 |
| Sipa1l2 | -0.0727 | 0.013236 | 1 |
| Copg2 | -0.07274 | 0.015155 | 1 |
| Zmym2 | -0.07278 | 0.001378 | 1 |
| Wdr7 | -0.07282 | 0.005015 | 1 |
| Dock2 | -0.07282 | 4.25E-07 | 0.013706 |
| Slc12a2 | -0.07294 | 0.044314 | 1 |
| Fryl | -0.07296 | 0.005677 | 1 |
| Kat2b | -0.07297 | 0.001658 | 1 |
| Cyp27a1 | -0.07297 | 0.004364 | 1 |
| Tbc1d8 | -0.07298 | 0.109868 | 1 |
| Ppp6r3 | -0.07303 | 0.002987 | 1 |
| Golim4 | -0.07304 | 0.034395 | 1 |
| Atf7 | -0.07315 | 0.005327 | 1 |
| Pla2g4a | -0.07319 | 0.000755 | 1 |
| Rbpj | -0.07329 | 0.033677 | 1 |
| March5 | -0.07341 | 0.007239 | 1 |
| Ercc6l2 | -0.07354 | 0.000288 | 1 |
| Gm27017 | -0.07358 | 3.64E-05 | 1 |
| Banp | -0.07363 | 9.66E-06 | 0.311768 |
| 5330438D12Rik | -0.07366 | 0.012038 | 1 |
| Cited2 | -0.0737 | 0.030258 | 1 |
| Tsc22d4 | -0.07383 | 0.001542 | 1 |
| Dop1b | -0.07384 | 0.005223 | 1 |
| Adgre1 | -0.07397 | 0.006954 | 1 |
| Abcc5 | -0.074 | 0.005712 | 1 |
| Cep128 | -0.07402 | 0.023504 | 1 |
| Gm37494 | -0.07407 | 0.044134 | 1 |
| Agbl3 | -0.07419 | 0.009755 | 1 |
| 1600014C10Rik | -0.07421 | 0.000838 | 1 |
| Brd2 | -0.07424 | 0.013527 | 1 |
| Kmt2e | -0.07425 | 0.064078 | 1 |
| Ipcef1 | -0.07427 | 8.31E-05 | 1 |
| Fyco1 | -0.07428 | 0.005427 | 1 |
| P2rx4 | -0.07429 | 0.040065 | 1 |
| Anxa6 | -0.0743 | 7.42E-05 | 1 |
| Vti1a | -0.07442 | 0.003131 | 1 |
| Usp50 | -0.07449 | 0.071993 | 1 |
| Acsl4 | -0.07452 | 0.001704 | 1 |
| Tuba1b | -0.07463 | 0.025965 | 1 |
| B3galnt2 | -0.07464 | 4.05E-06 | 0.130649 |
| Raf1 | -0.07464 | 0.000901 | 1 |
| Rbbp6 | -0.07465 | 0.009218 | 1 |

| | | | |
|---|---|---|---|
| Lrp5 | -0.07468 | 0.013422 | 1 |
| Zdhhc21 | -0.07469 | 0.001253 | 1 |
| Gm43813 | -0.07483 | 0.00021 | 1 |
| Celf1 | -0.07484 | 0.001081 | 1 |
| Fbxo33 | -0.07504 | 0.000611 | 1 |
| Pxn | -0.07524 | 0.001383 | 1 |
| Ankfy1 | -0.07525 | 0.012682 | 1 |
| Rap2a | -0.07526 | 0.010645 | 1 |
| Osbpl8 | -0.07541 | 0.011047 | 1 |
| Nfatc3 | -0.0755 | 0.003967 | 1 |
| Pip4k2a | -0.07553 | 0.000403 | 1 |
| Zadh2 | -0.07564 | 0.000354 | 1 |
| Vav2 | -0.07565 | 0.009343 | 1 |
| Ndufs3 | -0.07572 | 0.010067 | 1 |
| Nsmce2 | -0.07574 | 0.005609 | 1 |
| Ap1s2 | -0.07575 | 0.00366 | 1 |
| Ranbp9 | -0.07575 | 0.007893 | 1 |
| 1700012D14Rik | -0.07586 | 8.25E-06 | 0.266435 |
| Fmnl1 | -0.07593 | 0.000666 | 1 |
| Tbc1d5 | -0.07596 | 5.08E-05 | 1 |
| Dars | -0.07603 | 0.008107 | 1 |
| Gm31243 | -0.07603 | 0.011947 | 1 |
| Ep300 | -0.07604 | 0.000327 | 1 |
| Eya3 | -0.07614 | 0.0062 | 1 |
| Stim1 | -0.07616 | 0.003102 | 1 |
| Zfp950 | -0.07626 | 4.63E-05 | 1 |
| Sertad2 | -0.0763 | 0.04084 | 1 |
| Ddx17 | -0.07635 | 0.004161 | 1 |
| Tnrc6b | -0.07639 | 0.00124 | 1 |
| Fmnl2 | -0.07647 | 0.085922 | 1 |
| Fbrsl1 | -0.07658 | 0.000149 | 1 |
| Eed | -0.07684 | 0.014274 | 1 |
| 1110019D14Rik | -0.07688 | 0.003297 | 1 |
| Asxl2 | -0.07696 | 0.006355 | 1 |
| Rnf166 | -0.07711 | 0.005277 | 1 |
| 5830444B04Rik | -0.07717 | 3.07E-05 | 0.989833 |
| Tent5a | -0.07719 | 0.008823 | 1 |
| Prkch | -0.07719 | 0.042468 | 1 |
| Rufy2 | -0.07723 | 3.35E-05 | 1 |
| Rab11fip3 | -0.07724 | 4.25E-05 | 1 |
| Resf1 | -0.07736 | 0.00465 | 1 |
| Tln2 | -0.07741 | 0.089376 | 1 |
| Anapc16 | -0.07742 | 0.006531 | 1 |
| Tifa | -0.07748 | 0.00025 | 1 |
| Kansl1 | -0.07751 | 5.98E-05 | 1 |
| Slc43a2 | -0.07753 | 0.001716 | 1 |
| Cers6 | -0.07757 | 0.061846 | 1 |
| Arih1 | -0.07762 | 0.003952 | 1 |

| | | | |
|---|---|---|---|
| Kmt2a | -0.07766 | 0.002464 | 1 |
| Arhgef12 | -0.07766 | 0.016454 | 1 |
| Tango2 | -0.07772 | 0.001443 | 1 |
| Ubn2 | -0.07773 | 0.00055 | 1 |
| Dennd1a | -0.07777 | 0.002142 | 1 |
| Tia1 | -0.07781 | 0.004477 | 1 |
| Tnfaip2 | -0.07783 | 0.256697 | 1 |
| Nuak1 | -0.07794 | 0.017516 | 1 |
| Ubash3b | -0.07795 | 0.000273 | 1 |
| Larp1 | -0.07796 | 0.00518 | 1 |
| Rasa1 | -0.07796 | 6.3E-05 | 1 |
| Zeb1 | -0.07802 | 0.002332 | 1 |
| Zkscan3 | -0.07802 | 0.000369 | 1 |
| Slc36a1 | -0.07831 | 0.00109 | 1 |
| Mtmr14 | -0.07837 | 0.003065 | 1 |
| Ppp3ca | -0.07837 | 1.52E-05 | 0.491312 |
| 1110059E24Rik | -0.07839 | 0.003405 | 1 |
| Ifi47 | -0.07841 | 3.97E-06 | 0.128215 |
| Zbp1 | -0.07846 | 0.001277 | 1 |
| Setd1b | -0.0786 | 0.001436 | 1 |
| Me2 | -0.07882 | 2.72E-05 | 0.877565 |
| Anapc10 | -0.07891 | 0.004179 | 1 |
| Strbp | -0.07892 | 0.00417 | 1 |
| Gm26510 | -0.07898 | 0.005318 | 1 |
| Nsd1 | -0.07898 | 0.001169 | 1 |
| Pias1 | -0.07901 | 0.000138 | 1 |
| Afg1l | -0.07911 | 0.00382 | 1 |
| Ago2 | -0.07916 | 0.000144 | 1 |
| C4b | -0.07921 | 0.019881 | 1 |
| Bcl2 | -0.07926 | 0.12301 | 1 |
| Mitf | -0.07936 | 0.006516 | 1 |
| Ggact | -0.07957 | 8.71E-05 | 1 |
| Naa25 | -0.07977 | 4.39E-05 | 1 |
| Exoc4 | -0.07989 | 0.000545 | 1 |
| Slc2a9 | -0.07991 | 0.003033 | 1 |
| Snd1 | -0.07992 | 0.000843 | 1 |
| Stat5b | -0.07997 | 0.047768 | 1 |
| Ccdc62 | -0.08014 | 0.028456 | 1 |
| Foxn2 | -0.08026 | 0.010367 | 1 |
| Limk1 | -0.0803 | 0.007123 | 1 |
| Fbxw17 | -0.08031 | 5.02E-05 | 1 |
| Fgd3 | -0.08038 | 0.01139 | 1 |
| Cradd | -0.08056 | 0.181006 | 1 |
| Tpst1 | -0.08073 | 0.000241 | 1 |
| Mertk | -0.08075 | 0.000596 | 1 |
| Hs2st1 | -0.08094 | 0.017903 | 1 |
| Zfp992 | -0.08101 | 0.013109 | 1 |
| Man1c1 | -0.08102 | 0.038952 | 1 |

| | | | |
|---|---|---|---|
| Ptafr | -0.08104 | 0.001155 | 1 |
| Pip4k2b | -0.08112 | 0.000604 | 1 |
| Zfp516 | -0.08115 | 0.010882 | 1 |
| Riok3 | -0.08117 | 0.001245 | 1 |
| Xaf1 | -0.08122 | 0.001076 | 1 |
| Maz | -0.08153 | 5.98E-08 | 0.001931 |
| Rad50 | -0.08156 | 0.013001 | 1 |
| Cdyl2 | -0.08157 | 0.012467 | 1 |
| Usp40 | -0.08164 | 0.001682 | 1 |
| Wbp1l | -0.08172 | 6.51E-05 | 1 |
| Sik1 | -0.0818 | 2.45E-07 | 0.007899 |
| Ccdc6 | -0.0821 | 0.000564 | 1 |
| Ppp1r12b | -0.08218 | 0.000103 | 1 |
| Rasa3 | -0.0823 | 0.031984 | 1 |
| Ube2f | -0.08247 | 0.024017 | 1 |
| Gm12216 | -0.0827 | 0.003826 | 1 |
| Glg1 | -0.08286 | 0.00042 | 1 |
| Nbeal1 | -0.08296 | 0.000864 | 1 |
| Phc2 | -0.08299 | 0.005861 | 1 |
| Spire1 | -0.08303 | 0.001948 | 1 |
| Rab40c | -0.08326 | 0.002047 | 1 |
| N4bp1 | -0.08327 | 6.15E-05 | 1 |
| Ddx58 | -0.08338 | 0.005181 | 1 |
| Ptprj | -0.08344 | 0.000161 | 1 |
| Gnb1 | -0.0835 | 0.000492 | 1 |
| Itch | -0.08352 | 0.000236 | 1 |
| St7l | -0.08364 | 0.000563 | 1 |
| Cflar | -0.08375 | 0.000316 | 1 |
| Frmd4a | -0.08376 | 6.79E-07 | 0.021911 |
| Cop1 | -0.08377 | 0.001391 | 1 |
| Ankrd17 | -0.08393 | 0.002806 | 1 |
| Pnpla7 | -0.08422 | 0.010565 | 1 |
| D5Ertd579e | -0.0843 | 0.000524 | 1 |
| Pld1 | -0.08439 | 0.024304 | 1 |
| Zfand3 | -0.08448 | 0.000328 | 1 |
| Hivep2 | -0.0845 | 0.210842 | 1 |
| Brd3 | -0.08461 | 0.000428 | 1 |
| Rreb1 | -0.08469 | 1.41E-05 | 0.456714 |
| Bcl2l1 | -0.08474 | 0.003914 | 1 |
| A830008E24Rik | -0.08475 | 0.017619 | 1 |
| Celf2 | -0.08477 | 0.000162 | 1 |
| Hyal5 | -0.08486 | 0.00021 | 1 |
| Ankhd1 | -0.08493 | 0.002454 | 1 |
| Pacsin2 | -0.08499 | 0.000722 | 1 |
| Klf7 | -0.08503 | 0.000646 | 1 |
| Tap1 | -0.08522 | 0.000102 | 1 |
| Itpkb | -0.08524 | 0.000295 | 1 |
| Bank1 | -0.08528 | 0.0134 | 1 |

| | | | |
|---|---|---|---|
| Birc3 | -0.08551 | 0.000866 | 1 |
| Rab10 | -0.08553 | 0.001243 | 1 |
| Phf12 | -0.08565 | 0.004537 | 1 |
| AU040320 | -0.08571 | 0.001977 | 1 |
| Cd69 | -0.0858 | 0.026024 | 1 |
| Dennd2a | -0.08591 | 3.27E-05 | 1 |
| P4ha1 | -0.086 | 0.049156 | 1 |
| Prkag2 | -0.08604 | 0.000579 | 1 |
| A530072M11Rik | -0.08609 | 3.69E-06 | 0.119199 |
| Il6ra | -0.08612 | 0.003536 | 1 |
| Kif3b | -0.08613 | 0.00017 | 1 |
| Rspry1 | -0.08614 | 0.004966 | 1 |
| Rbks | -0.08629 | 1.35E-05 | 0.434693 |
| Abtb2 | -0.08638 | 3.52E-05 | 1 |
| Map2k1 | -0.08644 | 3.48E-06 | 0.112482 |
| St6gal1 | -0.08647 | 0.114672 | 1 |
| Mcmbp | -0.08648 | 0.009022 | 1 |
| Ascc3 | -0.08662 | 0.00141 | 1 |
| Pid1 | -0.0867 | 0.038372 | 1 |
| Lcorl | -0.08675 | 0.000976 | 1 |
| Tns3 | -0.08679 | 0.000217 | 1 |
| Maf | -0.08681 | 0.000196 | 1 |
| Pkig | -0.08684 | 2.58E-05 | 0.831761 |
| Foxk1 | -0.08686 | 0.002798 | 1 |
| Pla2g15 | -0.08689 | 0.003035 | 1 |
| Rnf38 | -0.08693 | 0.00129 | 1 |
| 5430405H02Rik | -0.08718 | 0.000466 | 1 |
| Ip6k1 | -0.08718 | 6.89E-06 | 0.222544 |
| Calhm6 | -0.08754 | 5.89E-05 | 1 |
| Prkcd | -0.08777 | 5.82E-05 | 1 |
| Pds5a | -0.08794 | 0.005045 | 1 |
| Pip5k1a | -0.08804 | 0.00229 | 1 |
| Ppp4r1 | -0.08816 | 0.001837 | 1 |
| Mllt10 | -0.0882 | 0.001242 | 1 |
| Fgd4 | -0.0882 | 0.003942 | 1 |
| Tank | -0.08821 | 0.008542 | 1 |
| Atg10 | -0.0883 | 0.000739 | 1 |
| Hk3 | -0.08837 | 0.00222 | 1 |
| Rab27a | -0.08841 | 0.000226 | 1 |
| Syk | -0.08861 | 0.004231 | 1 |
| Arel1 | -0.08862 | 0.001336 | 1 |
| Arhgef3 | -0.08863 | 0.002402 | 1 |
| Gnaq | -0.08865 | 9.92E-08 | 0.003204 |
| Mob1b | -0.0891 | 0.000362 | 1 |
| Npnt | -0.08918 | 0.165642 | 1 |
| Abl1 | -0.08923 | 0.002317 | 1 |
| Klhl5 | -0.08932 | 0.002218 | 1 |
| Rgl1 | -0.08933 | 0.006903 | 1 |

| | | | |
|---|---|---|---|
| Znfx1 | -0.08949 | 5.04E-05 | 1 |
| Ewsr1 | -0.08951 | 0.000171 | 1 |
| Susd6 | -0.0897 | 0.00022 | 1 |
| Extl3 | -0.08976 | 0.000275 | 1 |
| Dock8 | -0.08981 | 8.88E-05 | 1 |
| Csnk1g3 | -0.08989 | 0.027222 | 1 |
| Jak2 | -0.08996 | 0.000622 | 1 |
| Slc25a12 | -0.09007 | 0.003266 | 1 |
| Tnfsf13b | -0.09039 | 0.00011 | 1 |
| Dock10 | -0.09041 | 3.77E-06 | 0.121679 |
| Itpk1 | -0.09044 | 9.32E-06 | 0.300821 |
| Aak1 | -0.0905 | 0.000172 | 1 |
| Oasl1 | -0.09058 | 0.001956 | 1 |
| Eepd1 | -0.09078 | 0.000769 | 1 |
| Gm46224 | -0.09079 | 0.000708 | 1 |
| Cnot3 | -0.0908 | 2.22E-05 | 0.716791 |
| Fam241a | -0.09094 | 5.95E-07 | 0.019219 |
| Axl | -0.09094 | 0.004852 | 1 |
| Nadk | -0.09103 | 0.003054 | 1 |
| Zfr | -0.0914 | 0.000983 | 1 |
| Rcsd1 | -0.09174 | 0.000906 | 1 |
| Abcg1 | -0.09179 | 0.00137 | 1 |
| Tbxas1 | -0.0918 | 0.000104 | 1 |
| Ppp1r11 | -0.09181 | 9.59E-07 | 0.030969 |
| Mau2 | -0.09184 | 0.001681 | 1 |
| Pak1 | -0.09195 | 0.000652 | 1 |
| Zfp280c | -0.092 | 0.000539 | 1 |
| Tmem104 | -0.0922 | 0.000208 | 1 |
| Mycbp2 | -0.09225 | 9.32E-06 | 0.301028 |
| Zfx | -0.09228 | 0.000794 | 1 |
| Ranbp10 | -0.09241 | 0.000746 | 1 |
| Atp6v0a1 | -0.09242 | 0.003638 | 1 |
| Epsti1 | -0.09245 | 9.5E-05 | 1 |
| Cul1 | -0.09273 | 0.000162 | 1 |
| Rhoq | -0.09314 | 0.000291 | 1 |
| Dock11 | -0.09319 | 0.003565 | 1 |
| N4bp2l1 | -0.0932 | 0.000209 | 1 |
| Prpf4b | -0.09325 | 0.014282 | 1 |
| 1700084C06Rik | -0.09326 | 0.000419 | 1 |
| Ifi206 | -0.0934 | 0.000396 | 1 |
| Skap2 | -0.09345 | 3.48E-05 | 1 |
| Fcho2 | -0.0935 | 0.000111 | 1 |
| Rbm47 | -0.09358 | 1.5E-05 | 0.484492 |
| Gab2 | -0.09367 | 7.72E-07 | 0.024923 |
| Ppargc1b | -0.09368 | 2.45E-05 | 0.790097 |
| Gon4l | -0.09369 | 0.000124 | 1 |
| Tbc1d4 | -0.09374 | 0.006269 | 1 |
| Smg7 | -0.09382 | 0.009103 | 1 |

| | | | |
|---|---|---|---|
| Grk3 | -0.09393 | 0.001069 | 1 |
| Ap2b1 | -0.09416 | 5.96E-05 | 1 |
| Gm48678 | -0.09434 | 0.001082 | 1 |
| Swt1 | -0.09441 | 0.001858 | 1 |
| Gm26740 | -0.09451 | 0.001478 | 1 |
| Stk10 | -0.09455 | 0.014939 | 1 |
| Fam222b | -0.09456 | 0.00074 | 1 |
| Maml2 | -0.0947 | 8.71E-05 | 1 |
| Dglucy | -0.09483 | 0.005243 | 1 |
| Cept1 | -0.09489 | 0.00049 | 1 |
| Numb | -0.09512 | 4.17E-05 | 1 |
| Phf21a | -0.09519 | 0.000377 | 1 |
| Heatr5b | -0.09557 | 1.91E-05 | 0.617989 |
| Gm20275 | -0.09558 | 0.000685 | 1 |
| Ubr3 | -0.09572 | 0.000294 | 1 |
| Rps6ka3 | -0.09594 | 0.00125 | 1 |
| Vcpip1 | -0.09628 | 0.000225 | 1 |
| Camk2d | -0.09662 | 1.92E-05 | 0.620834 |
| Fbxw4 | -0.09666 | 3.38E-06 | 0.109277 |
| Gbf1 | -0.09682 | 0.003283 | 1 |
| Nfatc2 | -0.09696 | 0.00923 | 1 |
| Klhl18 | -0.09697 | 1.14E-05 | 0.368638 |
| Csf2rb | -0.09702 | 0.001181 | 1 |
| Sufu | -0.09711 | 0.000239 | 1 |
| Arhgef1 | -0.09716 | 2.9E-05 | 0.936577 |
| Abca9 | -0.09716 | 0.002124 | 1 |
| Peli1 | -0.09722 | 1.38E-05 | 0.446004 |
| Mrtfa | -0.09767 | 5.92E-06 | 0.19116 |
| Gpi1 | -0.09792 | 0.000324 | 1 |
| Camk1d | -0.09799 | 5.01E-06 | 0.161859 |
| Etv6 | -0.098 | 1.8E-07 | 0.005807 |
| Epb41 | -0.09812 | 0.000212 | 1 |
| Bicra | -0.09822 | 0.000494 | 1 |
| H2-DMb1 | -0.09825 | 0.011652 | 1 |
| Sncaip | -0.09826 | 0.030999 | 1 |
| Nod1 | -0.09827 | 1.74E-07 | 0.005611 |
| Gm43305 | -0.09832 | 2.69E-05 | 0.868707 |
| 4732471J01Rik | -0.09859 | 0.000523 | 1 |
| Lars2 | -0.09873 | 0.000206 | 1 |
| Nuak2 | -0.09888 | 1.01E-05 | 0.325491 |
| Diaph1 | -0.09892 | 0.000307 | 1 |
| Pml | -0.09939 | 0.000579 | 1 |
| Wwox | -0.0998 | 0.006494 | 1 |
| Rap1gds1 | -0.09998 | 4.37E-07 | 0.014113 |
| Clec16a | -0.10009 | 5.17E-05 | 1 |
| Kcmf1 | -0.10011 | 0.00042 | 1 |
| Usp25 | -0.10032 | 0.000224 | 1 |
| Snhg16 | -0.10037 | 4.42E-13 | 1.43E-08 |

| | | | |
|---|---|---|---|
| Ston1 | -0.10041 | 0.000206 | 1 |
| Tiam1 | -0.10056 | 0.004348 | 1 |
| Ptpn9 | -0.10073 | 0.001529 | 1 |
| Gm12185 | -0.10086 | 0.000381 | 1 |
| Spop | -0.10094 | 0.000136 | 1 |
| Madd | -0.10095 | 0.000185 | 1 |
| Clcn7 | -0.10105 | 9.77E-05 | 1 |
| Ripk1 | -0.10138 | 0.000207 | 1 |
| 9930111J21Rik2 | -0.10147 | 3.73E-06 | 0.120458 |
| Zfp106 | -0.10165 | 0.000897 | 1 |
| Zbtb34 | -0.10172 | 2.26E-05 | 0.728055 |
| Vcam1 | -0.10179 | 0.016371 | 1 |
| Plxdc2 | -0.10179 | 3.07E-08 | 0.000991 |
| Dennd1b | -0.10193 | 0.000366 | 1 |
| Slfn8 | -0.10194 | 0.000242 | 1 |
| Dleu2 | -0.10199 | 1.58E-05 | 0.509715 |
| Prkca | -0.10207 | 8.66E-05 | 1 |
| Asxl1 | -0.10218 | 0.009117 | 1 |
| Phkb | -0.10231 | 0.000305 | 1 |
| Trafd1 | -0.10243 | 2.45E-05 | 0.79019 |
| Dapp1 | -0.10272 | 0.000151 | 1 |
| March7 | -0.10276 | 0.000207 | 1 |
| Pik3r6 | -0.10303 | 0.000264 | 1 |
| Dtx3l | -0.10316 | 1.08E-05 | 0.347081 |
| Rmnd1 | -0.10326 | 8.29E-06 | 0.267675 |
| Fam219a | -0.10338 | 0.002268 | 1 |
| Tbc1d1 | -0.10372 | 0.008068 | 1 |
| Zbtb7a | -0.10398 | 0.000143 | 1 |
| Trim34a | -0.10445 | 5.92E-05 | 1 |
| Kdm7a | -0.10495 | 1.97E-05 | 0.637516 |
| Arhgap12 | -0.10499 | 0.000461 | 1 |
| Igtp | -0.10546 | 8.74E-05 | 1 |
| Icam1 | -0.10549 | 0.023766 | 1 |
| Slc2a3 | -0.10553 | 2.09E-06 | 0.067463 |
| Zfp40 | -0.10579 | 3.19E-05 | 1 |
| Clybl | -0.10586 | 0.013386 | 1 |
| Msr1 | -0.10599 | 3.24E-05 | 1 |
| Elf4 | -0.10644 | 0.00036 | 1 |
| Man2a1 | -0.10667 | 0.000127 | 1 |
| Csf3r | -0.10697 | 3.22E-06 | 0.104118 |
| Gm49797 | -0.10703 | 1.05E-05 | 0.337904 |
| Dennd6a | -0.10724 | 0.000353 | 1 |
| Stx16 | -0.1074 | 3.34E-05 | 1 |
| Akap8l | -0.10742 | 2.5E-05 | 0.805752 |
| Rprd2 | -0.10748 | 0.000155 | 1 |
| Arid4a | -0.10768 | 5.97E-05 | 1 |
| Mark2 | -0.10769 | 0.000107 | 1 |
| Aftph | -0.10786 | 0.000557 | 1 |

| | | | |
|---|---|---|---|
| Gm35188 | -0.10793 | 0.0242 | 1 |
| Fgfr2 | -0.10803 | 0.000109 | 1 |
| Tnrc18 | -0.10822 | 0.001832 | 1 |
| Rnf150 | -0.1083 | 0.005437 | 1 |
| L3mbtl3 | -0.10836 | 7.01E-06 | 0.226159 |
| Asap1 | -0.10843 | 4.13E-08 | 0.001334 |
| Usp18 | -0.10865 | 2.86E-05 | 0.923441 |
| Sla | -0.10886 | 0.01297 | 1 |
| Tnfrsf11a | -0.10894 | 0.0007 | 1 |
| Rab43 | -0.10917 | 5.77E-05 | 1 |
| H2-T23 | -0.10925 | 2.14E-05 | 0.690484 |
| Sgip1 | -0.10944 | 0.002184 | 1 |
| Arhgap39 | -0.10945 | 5.54E-05 | 1 |
| Rufy3 | -0.10987 | 0.000896 | 1 |
| Zc3h12c | -0.10991 | 0.008257 | 1 |
| Nfkbia | -0.10993 | 9.03E-07 | 0.029153 |
| Plcg2 | -0.11001 | 1.54E-06 | 0.049569 |
| Prkcb | -0.11056 | 1.24E-05 | 0.40122 |
| Rnpep | -0.11107 | 4.84E-06 | 0.156237 |
| Pvt1 | -0.11108 | 3.06E-06 | 0.098907 |
| Tet3 | -0.11144 | 3.95E-06 | 0.12758 |
| Gm17106 | -0.11261 | 4.17E-05 | 1 |
| Tgfa | -0.11294 | 6.85E-06 | 0.221245 |
| Crebrf | -0.11343 | 0.000198 | 1 |
| Dna2 | -0.11345 | 0.000289 | 1 |
| Rras2 | -0.11378 | 0.013903 | 1 |
| Wdfy2 | -0.1138 | 7.92E-06 | 0.255696 |
| Plekhm3 | -0.11381 | 8.64E-05 | 1 |
| Fnip2 | -0.11406 | 0.016089 | 1 |
| Pim1 | -0.1141 | 0.055152 | 1 |
| Vgll4 | -0.11417 | 0.000126 | 1 |
| Gphn | -0.11424 | 0.000914 | 1 |
| Sp100 | -0.11451 | 1.8E-05 | 0.579683 |
| Fgl2 | -0.11454 | 1.85E-07 | 0.005973 |
| Socs1 | -0.11454 | 7.44E-09 | 0.00024 |
| Uvrag | -0.11461 | 3.14E-08 | 0.001013 |
| Stk4 | -0.11466 | 3.41E-07 | 0.01101 |
| Rps6kb1 | -0.11471 | 9.62E-06 | 0.310426 |
| Ifi213 | -0.11511 | 7.75E-08 | 0.002503 |
| Mthfs | -0.11536 | 1.16E-05 | 0.373508 |
| Adap2os | -0.1157 | 6.37E-05 | 1 |
| Slc31a1 | -0.1159 | 1.02E-07 | 0.003309 |
| Acer3 | -0.11594 | 1.35E-07 | 0.00437 |
| Myo1f | -0.11617 | 5.54E-08 | 0.001788 |
| Arid1a | -0.11688 | 1.08E-08 | 0.000349 |
| Gbp7 | -0.11698 | 1.2E-07 | 0.003865 |
| Sh2d3c | -0.11719 | 6.12E-07 | 0.019745 |
| Mbnl1 | -0.11747 | 4.11E-08 | 0.001327 |

| | | | |
|---|---|---|---|
| Bicral | -0.11811 | 2.93E-07 | 0.009469 |
| Slc38a6 | -0.11837 | 0.00176 | 1 |
| Lcp2 | -0.11849 | 1.24E-06 | 0.040194 |
| Slc1a3 | -0.11862 | 0.000168 | 1 |
| Pik3ap1 | -0.11885 | 6.54E-07 | 0.021108 |
| Cadm1 | -0.11923 | 0.008487 | 1 |
| Rassf4 | -0.11962 | 6.07E-07 | 0.019586 |
| Gbp6 | -0.11965 | 7.28E-06 | 0.235103 |
| Cbl | -0.11967 | 2.08E-07 | 0.006718 |
| Pik3cd | -0.11967 | 1.88E-05 | 0.607587 |
| Irs2 | -0.11978 | 0.00017 | 1 |
| Hs3st3b1 | -0.12011 | 0.000123 | 1 |
| Oas2 | -0.12034 | 0.000129 | 1 |
| Mllt3 | -0.12049 | 5.34E-05 | 1 |
| Exoc6 | -0.12141 | 5.49E-07 | 0.017712 |
| Tns1 | -0.12154 | 8.77E-05 | 1 |
| Rapgef2 | -0.12177 | 0.000111 | 1 |
| A930037H05Rik | -0.1222 | 0.000173 | 1 |
| Notch2 | -0.12222 | 1.52E-05 | 0.490231 |
| Ubtd1 | -0.12268 | 1.21E-05 | 0.391722 |
| Foxo3 | -0.12336 | 1E-05 | 0.322766 |
| Mgmt | -0.1236 | 0.025921 | 1 |
| Slfn5 | -0.12383 | 0.001011 | 1 |
| Taok3 | -0.12403 | 1.92E-08 | 0.000621 |
| Atp7a | -0.12469 | 4.31E-08 | 0.001393 |
| Synj1 | -0.12479 | 1.54E-08 | 0.000497 |
| Atp8a1 | -0.12507 | 2.02E-10 | 6.54E-06 |
| Tlr7 | -0.1253 | 6.6E-06 | 0.213103 |
| Irf2 | -0.12548 | 6.51E-08 | 0.002101 |
| Irf1 | -0.12553 | 1.53E-05 | 0.493831 |
| Rab8b | -0.12562 | 0.000125 | 1 |
| Arhgap25 | -0.12568 | 1.6E-07 | 0.005156 |
| Hck | -0.12569 | 2.25E-08 | 0.000728 |
| Lyst | -0.12774 | 4.95E-06 | 0.159881 |
| Rp2 | -0.12788 | 3.98E-05 | 1 |
| Atg7 | -0.12789 | 5.95E-07 | 0.019217 |
| Sfi1 | -0.12851 | 1.2E-09 | 3.87E-05 |
| Zc3hav1 | -0.12855 | 5.48E-06 | 0.176806 |
| Vps54 | -0.1289 | 8.81E-09 | 0.000285 |
| A330040F15Rik | -0.12972 | 2.79E-07 | 0.009018 |
| Apobec3 | -0.13007 | 4.29E-07 | 0.013863 |
| Sipa1l3 | -0.13009 | 5.64E-08 | 0.001822 |
| Lrrc4 | -0.13053 | 0.000718 | 1 |
| Rere | -0.13138 | 9.6E-10 | 3.1E-05 |
| Parp8 | -0.13168 | 1.57E-05 | 0.507964 |
| E230001N04Rik | -0.13267 | 7.21E-20 | 2.33E-15 |
| Eif4g3 | -0.13271 | 8.58E-09 | 0.000277 |
| Jade2 | -0.13293 | 9.79E-08 | 0.00316 |

| | | | |
|---|---|---|---|
| Nsf | -0.13328 | 3.23E-05 | 1 |
| Sdccag8 | -0.13443 | 2.53E-08 | 0.000818 |
| Selenow | -0.1347 | 6.69E-09 | 0.000216 |
| Il15ra | -0.13491 | 6.01E-06 | 0.193915 |
| Hpcal1 | -0.13496 | 7.29E-08 | 0.002352 |
| Tbc1d14 | -0.13668 | 4.81E-08 | 0.001553 |
| Lmln | -0.13683 | 0.000493 | 1 |
| Tbc1d9 | -0.13712 | 2.43E-05 | 0.783004 |
| Cacna1d | -0.13831 | 2.3E-05 | 0.743513 |
| Gm2245 | -0.13984 | 1.45E-05 | 0.46826 |
| Anks1 | -0.13986 | 1.34E-07 | 0.004337 |
| Mapk14 | -0.14166 | 1.31E-08 | 0.000424 |
| Parp11 | -0.14356 | 4.01E-07 | 0.012959 |
| Dnmt3a | -0.14432 | 3.66E-11 | 1.18E-06 |
| Hdac9 | -0.14537 | 0.002216 | 1 |
| Aoah | -0.146 | 0.007522 | 1 |
| Parp9 | -0.14794 | 4.48E-08 | 0.001445 |
| Zfp710 | -0.14809 | 9.9E-12 | 3.2E-07 |
| Ddhd1 | -0.14896 | 5.94E-06 | 0.191632 |
| Psmb7 | -0.14919 | 1.79E-13 | 5.78E-09 |
| March1 | -0.15078 | 3.07E-09 | 9.91E-05 |
| Trim30a | -0.1521 | 8.08E-10 | 2.61E-05 |
| Ccnd2 | -0.15213 | 4.34E-09 | 0.00014 |
| Trim14 | -0.15219 | 2.99E-08 | 0.000965 |
| Osbpl11 | -0.1526 | 9.27E-05 | 1 |
| Frmd4b | -0.15263 | 9.19E-13 | 2.97E-08 |
| Rnf169 | -0.15307 | 1.21E-07 | 0.003894 |
| Gm22146 | -0.15345 | 2.34E-14 | 7.56E-10 |
| Pik3r1 | -0.15376 | 7.39E-11 | 2.39E-06 |
| Tmem164 | -0.15558 | 6.65E-07 | 0.021465 |
| St6galnac2 | -0.1557 | 1.21E-69 | 3.9E-65 |
| Cnr2 | -0.15657 | 2.69E-09 | 8.69E-05 |
| Nrros | -0.1567 | 1.07E-11 | 3.44E-07 |
| Abcg3 | -0.15687 | 7.97E-05 | 1 |
| Lrch3 | -0.15699 | 3.73E-10 | 1.2E-05 |
| Ssh2 | -0.15749 | 1.67E-12 | 5.4E-08 |
| Ppm1h | -0.15801 | 1.41E-13 | 4.54E-09 |
| Nrp1 | -0.15859 | 4.53E-07 | 0.014641 |
| Ppp1r21 | -0.15864 | 7.38E-08 | 0.002384 |
| Mlxip | -0.1591 | 6.08E-08 | 0.001962 |
| Lysmd4 | -0.15981 | 1.59E-05 | 0.513587 |
| Arhgap15 | -0.16058 | 0.004207 | 1 |
| 4930599N23Rik | -0.16144 | 9.11E-12 | 2.94E-07 |
| Klf13 | -0.16214 | 3.46E-06 | 0.111632 |
| Ripor2 | -0.16219 | 1.57E-06 | 0.050537 |
| Cdk8 | -0.16317 | 6.18E-14 | 2E-09 |
| Oasl2 | -0.16423 | 5.8E-07 | 0.018728 |
| Ifi211 | -0.16426 | 9.9E-10 | 3.19E-05 |

| | | | |
|---|---|---|---|
| Prr14l | -0.16459 | 3.07E-09 | 9.92E-05 |
| Ccm2 | -0.16463 | 6.23E-09 | 0.000201 |
| Ddx60 | -0.16514 | 0.000532 | 1 |
| Nol4l | -0.16521 | 0.000294 | 1 |
| Slfn2 | -0.16572 | 4.7E-09 | 0.000152 |
| Parp14 | -0.16573 | 2.18E-10 | 7.05E-06 |
| Coro2a | -0.16612 | 1.29E-06 | 0.041505 |
| Rsrp1 | -0.16654 | 1.57E-11 | 5.08E-07 |
| Arsb | -0.16895 | 2.44E-05 | 0.787135 |
| Stat2 | -0.16916 | 1.72E-10 | 5.56E-06 |
| Gm26917 | -0.16959 | 0.862007 | 1 |
| Fyb | -0.17042 | 7.68E-14 | 2.48E-09 |
| Nlrc5 | -0.17247 | 1.12E-11 | 3.61E-07 |
| Nampt | -0.17424 | 4.67E-10 | 1.51E-05 |
| Morrbid | -0.17449 | 1.47E-06 | 0.047412 |
| Zeb2 | -0.17488 | 6.35E-15 | 2.05E-10 |
| Herc6 | -0.17662 | 2.11E-07 | 0.006798 |
| B4galt1 | -0.17808 | 9.39E-13 | 3.03E-08 |
| Irgm2 | -0.17882 | 1.03E-10 | 3.32E-06 |
| Cd300lf | -0.17902 | 7.81E-08 | 0.002521 |
| Xdh | -0.1808 | 4.25E-16 | 1.37E-11 |
| Pstpip2 | -0.18478 | 4.74E-09 | 0.000153 |
| Ddit4 | -0.18488 | 8.5E-27 | 2.74E-22 |
| Gbp2 | -0.18577 | 1.19E-07 | 0.003839 |
| Arid5b | -0.18848 | 1E-07 | 0.003235 |
| Cd36 | -0.18879 | 1.56E-09 | 5.05E-05 |
| Rnf213 | -0.19225 | 9.98E-11 | 3.22E-06 |
| Cxcl10 | -0.1944 | 0.000103 | 1 |
| Irgm1 | -0.19731 | 1.17E-16 | 3.78E-12 |
| Ciita | -0.19979 | 1.43E-20 | 4.61E-16 |
| Rad51b | -0.20335 | 1.03E-05 | 0.331906 |
| Snx30 | -0.20961 | 8.04E-10 | 2.6E-05 |
| Stat1 | -0.21533 | 2.04E-17 | 6.57E-13 |
| Aff1 | -0.23686 | 6.97E-15 | 2.25E-10 |
| Gbp4 | -0.24005 | 5.28E-19 | 1.7E-14 |
| Csmd3 | -0.25138 | 1.16E-10 | 3.75E-06 |
| Samhd1 | -0.25612 | 1.86E-16 | 6.01E-12 |
| Gm4951 | -0.25659 | 9.26E-15 | 2.99E-10 |
| Gbp8 | -0.26499 | 1.33E-19 | 4.31E-15 |
| AY036118 | -0.27094 | 5.9E-47 | 1.9E-42 |
| Cmss1 | -0.34057 | 2.5E-32 | 8.07E-28 |
| Gbp5 | -0.35455 | 8.07E-26 | 2.6E-21 |
| Fkbp5 | -0.36708 | 2.35E-13 | 7.6E-09 |
| Gm42418 | -0.37335 | 1.56E-91 | 5.04E-87 |
| H2-Eb1 | -0.38619 | 6.49E-20 | 2.1E-15 |
| Cd74 | -0.45547 | 2.03E-27 | 6.55E-23 |
| H2-Aa | -0.46031 | 1.4E-25 | 4.51E-21 |
| H2-Ab1 | -0.46463 | 6.43E-25 | 2.08E-20 |

| | | | |
|---|---|---|---|
| Iigp1 | -0.5506 | 4.29E-24 | 1.39E-19 |

#### Supplemental Table 5: Flow Cytometry Antibodies and Reagents

| Target | Clone | Fluorochrome(s) | Vendor |
|---|---|---|---|
| B220 | RA3-6B2 | APC | BD Biosciences |
| CD4 | RM4-5 | APC | eBioscience |
| CD4 | RM4-5 | e450, FITC | BD Biosciences |
| CD4 | GK1.5 | PE | BD Biosciences |
| CD4 | GK1.5 | SB600 | eBioscience |
| CD8 | 53-6.7 | APC, PE-Cy7, BUV737 | BD Biosciences |
| CD11b | M1/70 | e450 | eBioscience |
| CD45.2 | 104 | APC | BD Biosciences |
| CD45.2 | 104 | BV785 | BioLegend |
| CD80 | 16-10A1 | APC | eBioscience |
| CD86 | GL-1 | PE-Cy5.5 | BioLegend |
| F4/80 | BM8 | PE | eBioscience |
| GM-CSF | MP1-22E9 | PE-Cy7 | eBioscience |
| $H2K^b$ | AF6-88.5 | FITC, BUV395 | BD Biosciences |
| $H2K^b$ | AF6-88.5.5.3 | PE | eBioscience |
| $H2K^d$ | SF1-1.1 | FITC, PE | BD Biosciences |
| $H2K^k$ | 36-7-5 | PE | BD Biosciences |
| IFN$\gamma$ | XMG1.2 | APC | eBioscience |
| IL-6 | MQ2-13A5 | e450 | eBioscience |
| MHC Class II | M5/114.15.2 | AF700 | eBioscience |
| NK1.1 | PK136 | PE | BD Biosciences |
| TCRβ | H57-597 | APC | BD Biosciences |
| TCRβ | H57-597 | PE, APC-e780 | eBioscience |
| Tmem119 | V3RT1GOsz | PE-Cy7 | eBioscience |
| TNF$\alpha$ | MP6-XT22 | PerCP-Cy5.5 | BioLegend |
| LIVE/DEAD | N/A | Aqua | Invitrogen |
| FC Block | 2.4G2 | N/A | BD Biosciences |

PE=Phycoerythrin, FITC=Fluorescein isothiocyanate, APC=Allophycocyanin, PE-Cy7= Phycoerythrin-Cyanine7, e450=eFlour 450, PerCP-Cy5.5=Peridinin chlorophyll protein-Cyanine5.5, PE-Cy5.5=Phycorythrin-Cyanine5.5, BV=Brilliant Violet, BUV=Brilliant Ultraviolet, AF700=Alexa Fluor 700, eAPC-780=Allophycocyanin-eFluor 780, SB600=Super Bright 600

### Supplemental Table 6: RT-qPCR primers (listed 5'-3')

| Target | 5' primer sequence | 3' primer sequence |
|---|---|---|
| IFN$\gamma$ | TCAAGTGGCATAGATGTGGAAGAA | TGGCTCTGCAGGATTTTCATG |
| TNF$\alpha$ | CTTCTGTCTACTGAACTTCGGG | CAGGCTTGTCACTCGAATTTTG |
| IL-6 | GTTCTCTGGGAAATCGTGGA | TCCAGTTTGGTAGCATCCATC |
| GM-CSF | TTTACTTTTCCTGGGCAT | TAGCTGGCTGTCATGTTCAA |
